# Deep Red Blood Cell Proteome Defines the Band 3 N-Terminus Interactome as a Regulator of Hypoxic Adaptation via BLVRB-Dependent *S*-Nitroso Transfer

**DOI:** 10.1101/2025.11.29.691178

**Authors:** Aaron V. Issaian, Monika Dzieciatkowska, Shaun Bevers, Safari Zohreh, Ariel Hay, Anthony Saviola, Jasmina S. Redzic, Julie A Reisz, Gregory R. Keele, Francesca I. Cendali, Zachary B. Haiman, Travis Nemkov, Daniel Stephenson, Christina Lisk, Francesca Vallese, Bernhard O. Palsson, S. Bruce King, Grier P Page, Allan Doctor, Krystalyn Hudson, Kirk C. Hansen, David C. Irwin, Narla Mohandas, James C Zimring, Elan Z. Eisenmesser, Angelo D’Alessandro

**Affiliations:** Department of Biochemistry and Molecular Genetics, University of Colorado Anschutz, Aurora, CO, USA; Center for Blood Oxygen Transport and Hemostasis (CBOTH), University of Maryland School of Medicine, Baltimore, MD; University of Virginia, Charlottesville, VA, USA; RTI International, Research Triangle Park, NC, USA; Omix Technologies Inc, Aurora, CO, USA; Structural Biology Initiative, CUNY Advanced Science Research Center, New York, NY, USA; Department of Medicine, Cardiovascular and Pulmonary Research Laboratory, CU Anschutz, Aurora, CO, USA; Shu Chien-Gene Lay Department of Bioengineering, University of California San Diego, La Jolla, CA, USA; Department of Pediatrics, University of California San Diego, La Jolla, CA, USA; Department of Chemistry, Wake Forest University, Winston-Salem, NC, USA; Department of Pediatrics, University of Maryland, School of Medicine, Baltimore, MD, USA; Laboratory of Transfusion Biology, Department of Pathology and Cell Biology, Columbia University Irving Medical Center, New York, NY, USA; New York Blood Center, New York, NY, USA; University of British Columbia, Vancouver, British Columbia, Canada; Canadian Blood Center, Vancouver, CA

**Author notes:** Corresponding author: Angelo D’Alessandro, PhD, Department of Biochemistry and Molecular Genetics, University of Colorado Anschutz Medical Campus, 12801 East 17th Ave., Aurora, CO 80045, www.dalessandrolab.com. These authors contributed equally and share the first authorship (Aaron V. Issaian, Monika Dzieciatkowska).

**Keywords:** SLC4A1, critical speed, exercise, hypoxia, erythrocyte, interactome

## Abstract

Red blood cells (RBCs) have long been regarded as passive oxygen carriers, yet growing evidence reveals a complex, dynamic proteome independent of de novo gene expression. Here, we define the erythrocyte as an oxygen-responsive system organized around a Band 3 (SLC4A1)–centered metabolon. Using deep proteomics of ultra-pure RBCs and cross-linking interactomics, we identify biliverdin reductase B (BLVRB) as a previously unrecognized Band 3 interactor that binds the N-terminal cytosolic domain under normoxia and dissociates under hypoxia, when band 3-deoxyhemoglobin interactions increase threefold. This reversible interaction forms an oxygen-sensitive switch coupling structural, redox, and metabolic remodeling. In humanized mice, truncation of the Band 3 N-terminus disrupted glycolytic activation, reduced 2,3-bisphosphoglycerate synthesis, and impaired exercise tolerance despite preserved cardiopulmonary function, establishing the physiological relevance of this module. Population-scale proteome quantitative trait locus (pQTL) analyses revealed coordinated variation of SLC4A1 and BLVRB abundance but minimal association of biliverdin levels with BLVRB genotype, suggesting alternative functions beyond heme catabolism. Mechanistically, BLVRB Cys109 acts as a nitric oxide (NO) relay, trans-nitrosating glycolytic enzymes such as GAPDH at active site Cys152, transiently inhibiting glycolysis. This S-nitrosation–mediated feedback mirrors conserved mechanisms in plants, where GAPDH-SNO redirects carbon flow toward the Calvin–Benson cycle under nitrosative stress, revealing an evolutionary convergence in gas-responsive metabolic control. Collectively, our findings define a Band 3–BLVRB–hemoglobin axis that links oxygen sensing, NO signaling, and redox homeostasis, providing a unifying model for how an anucleate cell achieves environmental adaptability through reversible protein–protein interactions and post-translational chemistry.

**Graphic abstract:** Issaian et al. define the most comprehensive proteome of ultra-pure human red blood cells (3,775 proteins) and map the O₂-dependent interactome, revealing a Band 3–BLVRB–hemoglobin module that links oxygen sensing to metabolic remodeling via reversible inhibitory S-nitrosation of GAPDH C152. In plants this redirects carbon toward photosynthesis, illustrating a conserved NO-dependent metabolic reprogramming mechanism across oxygen-regulated systems.

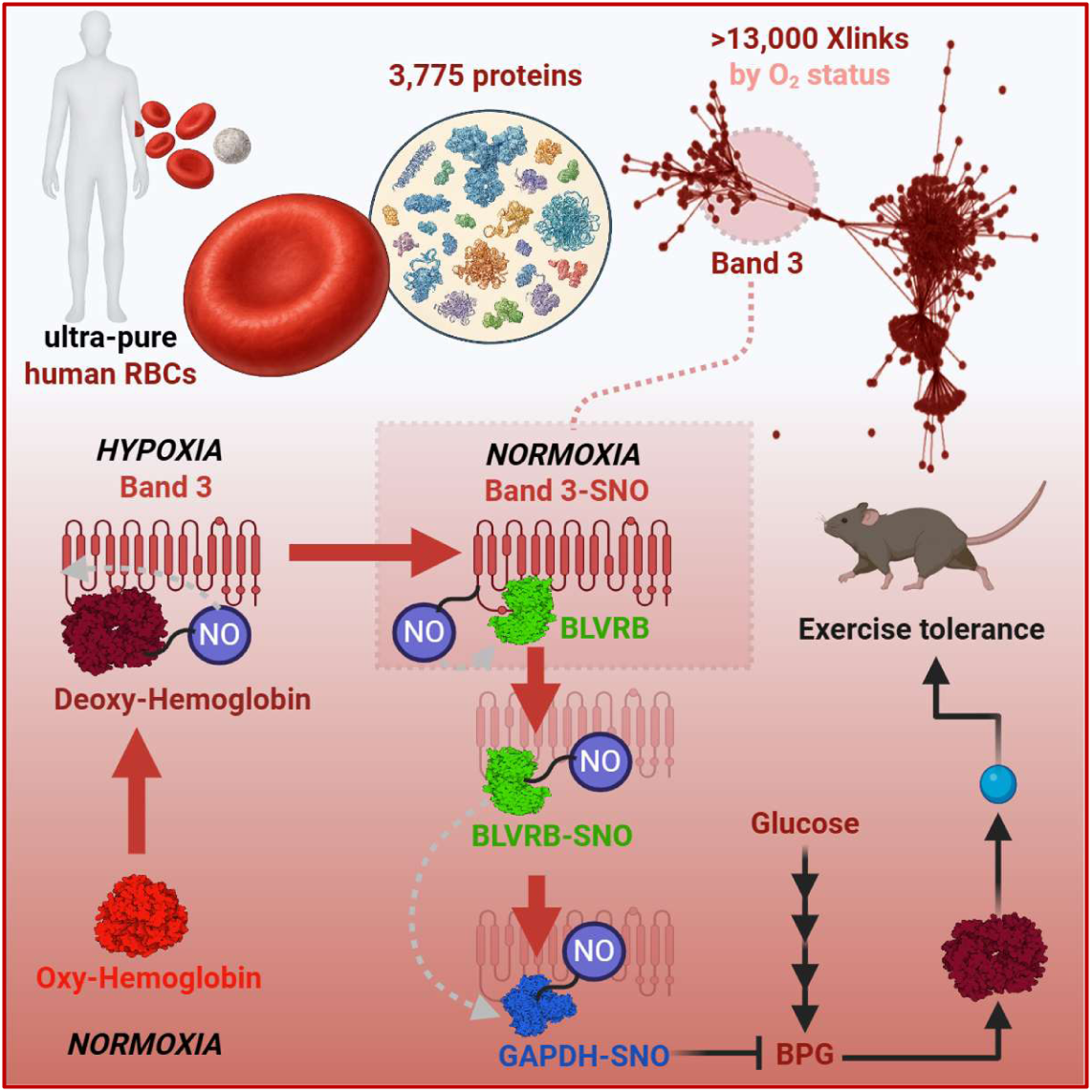

**Highlights:**

- Deep proteomics defines a complete, contamination-free RBC proteome (3,775 proteins)
- Cross-linking proteomics maps an oxygen-sensitive Band 3-centered interactome
- O2-dependent BLVRB–Band 3 binding regulates metabolism via S-nitrosation of GAPDH
- Band 3 N-terminus is required for hypoxic remodeling and exercise tolerance in vivo

## INTRODUCTION

Red blood cells (RBCs) represent approximately 25 of the ∼30 trillion cells in the human body^1,2^, permeate every tissue and account for nearly one-tenth of total body mass^3^. Long considered passive carriers of oxygen and carbon dioxide, a “hapless sack of hemoglobin”^4^, RBCs are now recognized as metabolically active cells with broad roles in redox homeostasis, nitric-oxide (NO) signaling, vascular tone regulation, and immune modulation^5^. Their sheer number and recently unraveled molecular complexity position them as major contributors to systemic physiology and as tractable models for studying cellular adaptation in the absence of protein synthesis.

Comprehensive proteomic and systems-biology analyses have progressively expanded the molecular scope of the mature RBC. Early two-dimensional gel studies detected fewer than two hundred proteins^6^; modern high-resolution datasets now identify several thousand^7–9^, including enzymes classically thought to be restricted to hepatic, renal, or neural metabolism - supporting reactions in amino-acid turnover, lipid remodeling, one-carbon and purine pathways, and even xenobiotic and drug detoxification chemistry^10^. Network reconstructions such as the RBC genome-scale metabolic model (RBC-GEM) integrate 29 RBC proteomics studies to compile more than 4,600 proteins and associated reactions^11^, portraying the erythrocyte as a densely connected metabolic and interactomic network rather than an inert carrier of hemoglobin. These advances have reframed the RBC as a ∼2.5 kg circulating organ^12^ that continuously exchanges metabolites and signaling molecules with every tissue it perfuses.

Despite this progress, uncertainty persists regarding the exact molecular composition of mature erythrocytes. Proteomic depth often trades off against sample purity: datasets reporting thousands of proteins may include contributions from residual leukocytes, platelets, or reticulocytes, whereas the most stringently purified preparations - achieved by fluorescence-activated cell sorting and verified for the absence of reticulocytes - identify roughly 1,400 proteins, upon subtraction of >650 reticulocyte-specific proteins^13^. The resulting disparity has fueled debate over whether the RBC proteome truly approaches the complexity suggested by in-silico reconstructions or whether some of that diversity reflects contamination from nucleated or immature cells. Resolving this issue requires both maximal analytical depth and uncompromising cellular purity to establish a definitive baseline proteome for the human erythrocyte.

The biological imperative for such precision extends beyond cataloguing. Mature RBCs lack nuclei and mitochondria and are incapable of transcription, translation, or canonical signaling through gene regulation. Nevertheless, they circulate for approximately 120 days, completing an estimated 170,000 systemic passages, while preserving membrane integrity, energy homeostasis, and antioxidant capacity under variable oxygen tension and oxidative stress^14^. Adaptation in this context relies entirely on post-translational regulation and dynamic protein–protein interactions. Reversible complex formation, phosphorylation, oxidation, and S-nitrosation modulate enzymatic activity and metabolic flux, allowing the cell to dynamically rewire metabolism in response to environmental cues^15^. The RBC thus provides a natural experiment in post-genomic homeostasis - demonstrating that cells can maintain regulated physiology without new protein synthesis.

A central example of interactome-based regulation is the band 3–deoxyhemoglobin model originally proposed by Low.^16^ Band 3 (SLC4A1), the most abundant membrane protein of the erythrocyte, anchors the spectrin–actin cytoskeleton and serves as a scaffold for metabolic enzymes^17–19^. Under oxygenated conditions, glycolytic enzymes bind to its cytosolic N-terminus, suppressing glycolysis and diverting glucose into the pentose-phosphate pathway to generate NADPH and reinforce antioxidant defenses^17,20^. Deoxygenation promotes hemoglobin binding to the same region, releasing these enzymes to restore glycolytic flux, elevate ATP and 2,3-bisphosphoglycerate, and facilitate oxygen unloading^17,20^. This reversible partitioning of metabolism establishes a direct link between oxygen transport, and redox/energy metabolic balance, and alteration of this system contributes to blunt hypoxic responses in sickle cell disease^21,22^, viral infections (including SARS-CoV-2)^23^ and inherited N-terminal mutations such as band 3 Neapolis (lacking residues 1-11)^24^.

Subsequent studies have broadened this framework. Proteomic analyses and thermal-profiling experiments have revealed additional band 3 interactors spanning antioxidant enzymes, chaperones, and kinases, supporting a model in which metabolic compartmentalization and redox regulation are coupled through large, multiprotein assemblies^25^.

RBCs also harbor nitric oxide synthase^26^ and nitrite-reductase activities^27^, enabling them to generate and exchange reactive nitrogen species with hemoglobin and the vasculature^28^. Enzyme-mediated S-nitrosation and sulfhydryl oxidation provide additional layers of reversible control, linking oxygenation to post-translational signaling. Recent data implicate biliverdin reductase B (BLVRB) - among the ten most abundant RBC cytosolic proteins - as an *S*-Nitroso-CoA-assisted nitrosylase, suggesting a regulated enzymatic mechanism for nitric-oxide transfer within the RBC proteome.

Despite this growing body of evidence, key questions remain unresolved. The absolute proteomic composition of the mature RBC remains uncertain due to contamination artifacts and methodological heterogeneity. In addition, the structural organization and dynamic remodeling of protein-protein interactions that govern oxygen-dependent metabolic control are incompletely mapped, in part because the hemoglobin-dominated cytosol complicates analysis of intact complexes. A major gap also persists in understanding how redox and nitrosative signals are integrated within the band 3/hemoglobin metabolon. In particular, the presence of highly abundant NADPH-dependent reductases in mature RBCs, such as biliverdin reductase B (BLVRB) - whose canonical role in heme catabolism is difficult to reconcile with the minimal heme oxygenase activity in these cells - raises the question of whether these enzymes participate in alternative redox or signaling functions within this oxygen-sensitive scaffold.

In the present study, we address these questions by combining ultrapurified human RBCs with deep quantitative proteomics, interactomics, and structural/functional analyses to define the most comprehensive and molecularly resolved map of the erythrocyte proteome and interactome to date. This unbiased approach revealed BLVRB as a previously unrecognized Band 3-binding partner and redox regulator, prompting mechanistic studies that link oxygen tension, NO transfer, and metabolic remodeling through the Band 3/BLVRB/hemoglobin axis. By resolving the architecture and dynamics of this system, our work refines the conceptual framework of RBC biology and provides a unified model for how a transcriptionally silent cell achieves environmental adaptability through reversible protein assemblies and post-translational chemistry.

## RESULTS

### Isolation of an ultra-pure population of mature human RBCs and construction of the deepest proteome to date

To establish the proteome composition of the mature human RBC in the absence of other cellular contaminants, we first developed a stringent multistage purification workflow. To this end, pooled whole blood samples from 5 donors underwent centrifugation for plasma and buffy coat removal, leukofiltration – a standard procedure in modern transfusion medicine, which removes log4 WBCs and log2.5 PLTs - and fluorescence-activated cell sorting (FACS) to positively isolate CD235⁺CD71⁻ erythrocytes, while simultaneously sorting out residual leukocytes (CD45⁺), platelets (CD41⁺ or CD62P⁺), and reticulocytes (CD235⁺CD71⁺); this workflow yielded an ultrapure RBC population (flow diagrams in **Figure 1A–B**).

**Figure 1.**
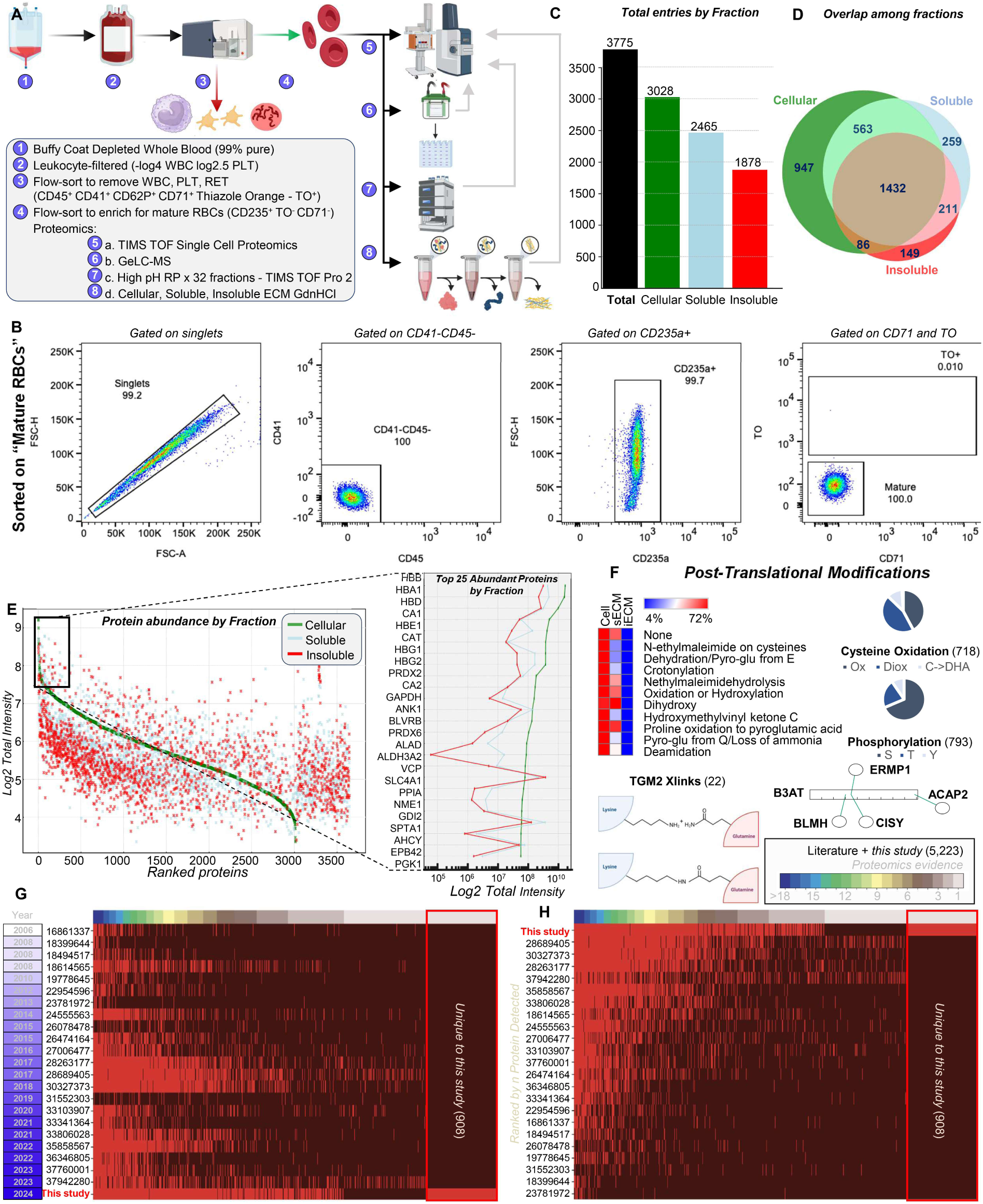
Deep red blood cell proteome identifies 3,775 proteins in ultra-pure RBCs. Comprehensive proteomic workflow yielding the purest human RBC proteome to date. Experimental workflow for RBC purification – including buffy coat removal after centrifugation, leukofiltration (removing log4 WBC and log2.5 PLTs), flow-sorting to remove residual PLTs, WBCs and reticulocytes and positive enrichment for mature RBCs (CD235+ CD71- TO-) (**A**). Proteome fractionation was then performed via both SDS-PAGE and high-pH-RP chromatography coupled to nano-UHPLC-MS/MS of both cellular, soluble and insoluble fractions based on methods optimized for extracellular matrix (ECM) protein recovery **(A)**. Flow charts confirm purity of the mature RBC population (**B**). Quantitative proteome coverage compared with previous RBC proteomics studies; our dataset expands known coverage by ∼3775 proteins (**C**) across fractions – shown as Venn Diagrams (**D**). Rank-ordered abundance plot highlighting major RBC proteins; Band 3 (SLC4A1) quantified as 6–7 × more abundant than prior estimates (**E**). Summary of the post-translational modifications by prevalence in the RBC proteome, including endogenous transglutaminase 2 (TGM2)-catalyzed cross-links of membrane proteins (**F**). Our analysis identifies 908 previously unreported proteins based on a merge meta-analysis of milestone RBC proteomics literature, sorted by year (**G**) or proteome coverage (**I**). Labels for each entry indicate PubMed IDs.

These ultrapure RBCs underwent analytical processing with multiple proteomics approaches, including separation via GeLC-MS/MS of 36 cytosol and membrane bands,^29^ and high-pH reversed-phase fractionation of 96 total fractions. Proteomics workflows have historically struggled to detect low-abundance receptors, transporters, and blood-group proteins (e.g. KCNN4, the Gardos channel, is known from antibody/functional assays but has eluded proteomic detection^30^). To overcome these hurdles, here we leveraged a protocol adapted from extracellular matrix proteomics, which combines strong chaotrope solubilization, sequential ultrasonication, urea/thiourea buffers and high-detergent extraction steps, followed by reduction/alkylation, filter digestion, and peptide cleanup in order to break protein–lipid cross-links and fully solubilize membrane complexes for downstream LC-MS/MS analysis^31^. Deep nano–UHPLC-MS/MS analysis on Orbitrap Fusion Lumos and timsTOF SCP platforms identified 3,775 unique proteins, representing the most comprehensive RBC proteome yet reported (**Supplementary Table 1**). Abundance-ranked mapping of the proteome revealed a continuous distribution spanning more than six orders of magnitude (**Figure 1E-F; Supplementary Figure 1.A-B**), from core cytoskeletal proteins and hemoglobin subunits to low-abundance signaling enzymes. These new identifications included numerous metabolic enzymes typically associated with hepatic, renal, or neural tissues - such as those participating in amino acid turnover, lipid remodeling, one-carbon and purine metabolism, and xenobiotic or drug-detoxification reactions (**Supplementary Figure 2**). This unexpected enzymatic diversity challenges the traditional view of the RBC as metabolically narrow and instead positions it as a multifunctional biochemical hub. Of note, a much more complex network of NADPH-dependent/generating enzymes was observed, among which the most abundant was BLVRB (13 million copies/cell) – ranking number 10 overall, number 4 after excluding all hemoglobin chain subunits. In this view, we confirm results from Roux-Dalvai and colleagues,^32^ reporting up to 32.6 million copies/cell of fetal hemoglobin subunits HBG1 and HBG2 in adult-derived ultrapure mature RBCs, a finding previously questioned as the potential result of contamination from other sources given silencing of fetal hemoglobin expression in adults.

Unsupervised post-translational modification analysis (including enriched PTMs from error tolerant searches) uncovered extensive phosphorylation, oxidation (**Figure 1F–G; Supplementary Table 1**), and transglutaminase-mediated crosslinking (**Supplementary Figure 3**), indicating that the RBC proteome is dynamically regulated despite the absence of new protein synthesis. These observations define a stable yet adaptable proteome capable of sophisticated redox and metabolic control.

Comparative analysis with 29 publicly available RBC proteomic datasets (as reviewed in Haiman et al. 2025^33^) showed a recovery of nearly every protein previously reported by more than 1 study, plus 908 newly identified proteins absent from the combined literature (**Figure 1.G-H**). Among these, KCNN4 was identified with an estimated 286,000 copies/cell. Quantitative label-free analysis further confirmed Band 3 (SLC4A1) as the dominant membrane component, with an estimated ∼6.46 × 10^6^ copies per cell—approximately 6–7 times higher than previous estimates. This increase largely reflects improved recovery of hydrophobic and detergent-insoluble proteins following our combined soluble and insoluble fractionation strategy, which minimizes underrepresentation of membrane-embedded species in traditional proteomics workflows. Endogenous TGM2-derived cross-links were mostly identified around the cytoskeletal protein compartment, especially SLC4A1, spectrin, ankyrin, titin, and metabolic enzymes G6PD and PFKAM consistent with a role of this cross-linking enzyme in regulating RBC deformability and glucose metabolism in response to hypoxia.^15^ These cross-links involved ε-(γ-glutamyl)–lysine isopeptide bonds formed between reactive glutamine and lysine residues, characteristic of TGM2 catalytic activity (**Supplementary Table 1**). Overall, the proteomics and PTM data are now offered as a publicly available, interactive resource – the **Deep Red database** (**Supplementary dataset** https://angelo-dalessandro.github.io/deep-red-supplementary-site/).

### Construction of the red blood cell interactome through orthogonal cross-linking mass spectrometry

To reveal the physical architecture of protein–protein interactions in mature RBCs, and expand upon the observation of endogenous TGM2-dependent cross-links, we performed parallel cross-linking experiments using amine-reactive disuccinimidyl sulfoxide (DSSO) and carboxyl-to-amine 4-(4,6-dimethoxy-1,3,5-triazin-2-yl)-4-methylmorpholinium chloride (DMTMM) chemistries. Each reagent captured complementary classes of interactions between cytosolic and membrane-associated complexes, identifying three domains within the RBC interactome network: the cytosolic hemoglobin interactome centered around hemoglobin chains, the cytoskeletal one centered around titin, and the membrane interactome centered around band 3 (**Figure 2A–B**). The combined datasets yielded over 13,035 high-confidence cross-linked MS^3^ spectral matches with a 1 % false-discovery rate (full list with Uniprot names for protein pairs and linked residues in **Supplementary Table 1**). The top 10 hubs in the RBC interactome are summarized in **Figure 2C** and reveal a previously unrecognized central role for titin (TTN) - the largest known human protein, spanning ∼3.8 MDa and over 34,000 amino acids (∼10,000 copies/cell) - and non-muscle myosin 9 (MYH9), in addition to the more abundant and expected hubs such as hemoglobin and Band 3 (SLC4A1). This finding underscores an expanded structural framework in which traditionally muscle-associated scaffolding proteins participate in maintaining the mechanical and organizational integrity of the mature erythrocyte. These results confirm and expand upon recent Xlinking proteomics studies on the band 3 interactome, which were limited to the very N-terminus peptide (1-56, i.e., the first tryptic cleavable K residue),^25^ revealing a much more complex interactome at different region of the larger N-terminal cytosolic domain of band 3 1-405 (**Figure 2D-E**), one that is compatible with SLC4A1 structural role in the ankyrin 1-complex^34^ (**Supplementary Figure 4**), while suggesting novel, as of yet unexplored roles for the band 3 N-term interactome in RBC physiology. Notably, BLVRB was identified as a novel interactor with the very 1-11 N-terminus residues (**Figure 2F-H**). Mapping of individual cross-link sites onto available crystal structures and AlphaFold models confirmed that the Band 3 N-terminal cytosolic region (residues 1–56) forms multiple discrete contact zones both within and across subunits (**Figure 2H**). This region has long been postulated to mediate reversible interactions with hemoglobin and glycolytic enzymes, but its direct molecular connectivity has not been structurally visualized until now, and has eluded early Xlinking proteomics studies^20,35^. In summary, after quantitatively defining the baseline RBC proteome, we established a baseline interactome in normoxia. Among the newly identified interactors, the Band 3–BLVRB connections suggested an unanticipated link between oxygen sensing, heme catabolism, and nitrosative chemistry - providing a rationale for subsequent detailed structural and functional studies.

**Figure 2.**
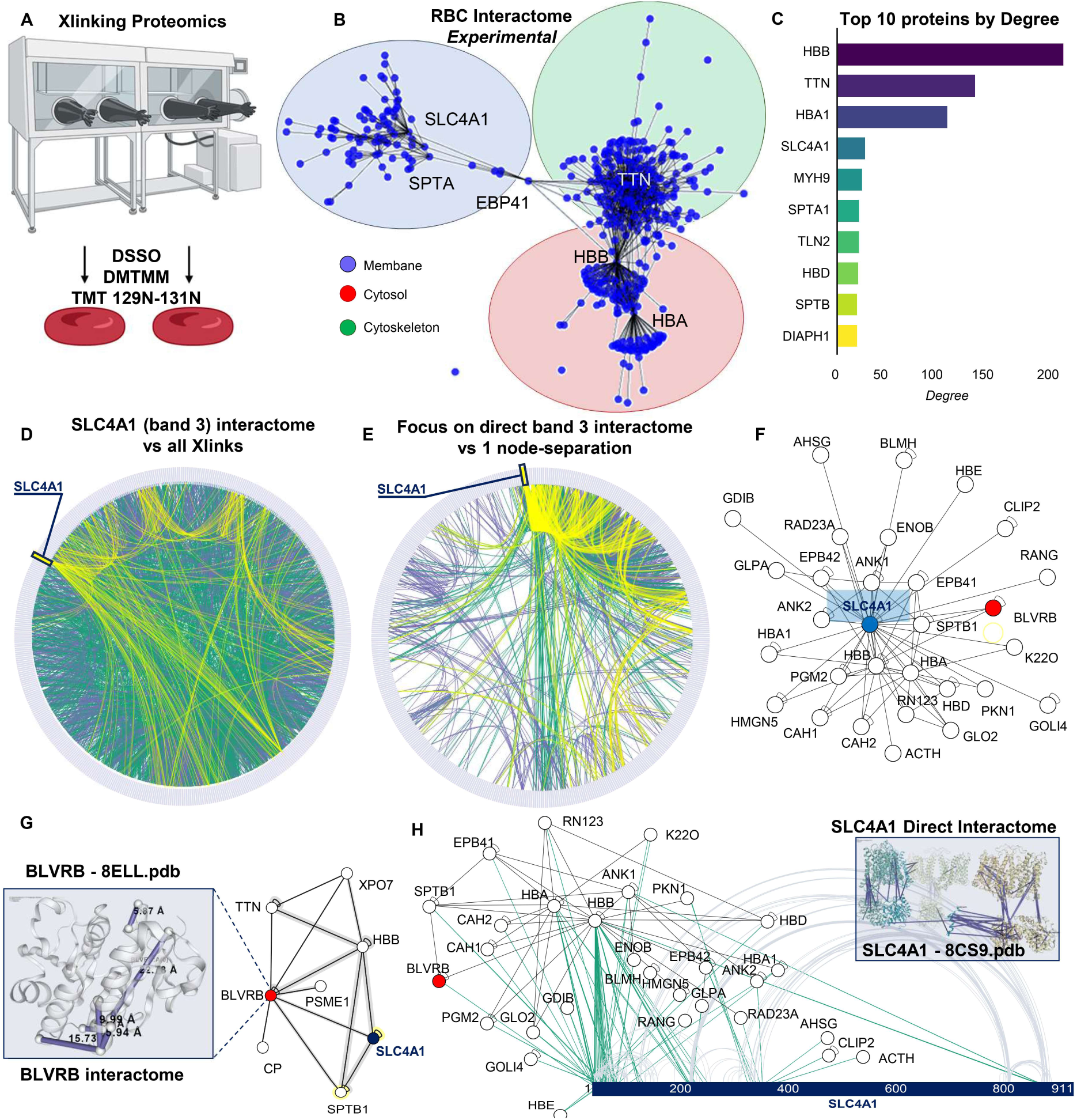
Chemical cross-linking proteomics defines the red blood cell interactome. System-wide mapping of RBC protein–protein interactions by orthogonal cross-linking approaches. Experimental workflow for DSSO (amine-reactive) and DMTMM (carboxyl-to-amine) cross-linking coupled to LC-MS/MS, coupled to TMT labeling for quantitation **(A).** Global RBC interactome network, highlighting high-confidence edges (cross-link spectral counts ≥ 2) and node degree proportional to interaction frequency **(B)**. Overview of top 10 detected intra- and inter-protein cross-links, with confidence scoring and subcellular distribution **(C).** Circos plot of all protein-protein interactions observed **(D)**, with a focus on the direct band 3 (SLC4A1) interactors **(E).** Network view of band 3 **(F)** or BLVRB interactors **(G)**. Focus on the interactome of the cytosolic N-terminus of band 3 (**H**).

### Hypoxia remodels the red cell interactome through dynamic reorganization of Band 3–hemoglobin contacts

We next explored how environmental oxygen tension influences the organization of the RBC interactome. Intact RBCs were equilibrated under either normoxic (21% O₂) or hypoxic (1% O₂) conditions, and subsequently subjected to cross-linking proteomics using identical conditions and relative quantitation by tandem mass tags. The resulting dataset revealed extensive, oxygen-dependent remodeling of protein–protein connectivity, involving nearly one-third of all detected cross-links (**Figure 3A–E; Supplementary Table 1**).

**Figure 3.**
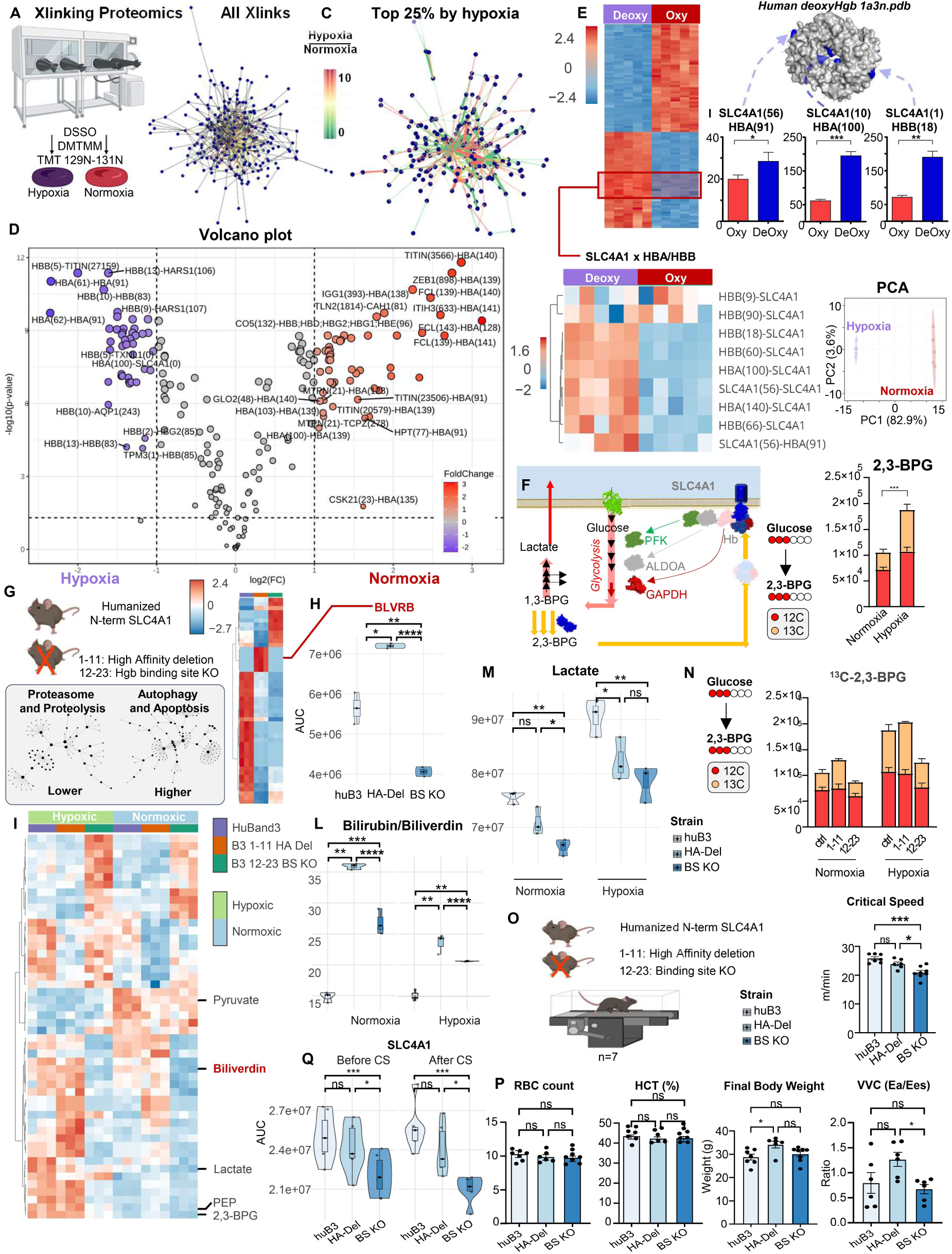
Hypoxia remodels the red blood cell interactome and strengthens the Band 3–deoxyhemoglobin interface, while loss of N-term band 3 blunts hypoxic responses and elicits exercise intolerance. While devoid of mitochondria, the RBC proteome dynamically responds to environmental stimuli like hypoxia. Schematic of normoxia and hypoxia treatment conditions and cross-linking strategy **(A)**. Overview of all Xlinks **(B)** and the top 25% of Xlinks impacted by oxy/deoxy status **(C)**. Volcano plot **(D)** and heat map **(E)** of differential interaction intensities (hypoxia vs normoxia) identifying the Band 3–Hb subnetwork as the most affected. Structural mapping of cross-linked residues on Band 3 N-terminus and deoxyHb surface, revealing multiple contact regions unique to hypoxia (**E**). Quantification of cross-link spectral counts for Band 3–Hb pairs under both conditions (n = 3 independent experiments, *p* < 0.01, paired t-test – heat map and selected bar plots in **E**). These observations are consistent with the model proposed by *Low et al.,* which posits a role for deoxyhemoglobin binding to band 3 in releasing glycolytic enzymes from the bound state to band 3, which inhibits them in normoxia; this dynamic changes in the band 3 interactome favor activation of glycolysis and Rapoport-Luebering shunt for the synthesis of 2,3-bisphosphoglycerate (2,3-BPG) in hypoxia, as confirmed here with 1,2,3-^13^C_3_-glucose tracing **(F)** New C57BL6/J mice were engineered to express human the human canonical band 3 N-terminus, either full length or lacking residues 1-11 (high affinity deletion – HA) or 12-23 (binding site - BS KO). These mice were generated de novo from a clean C57BL6/J background, to independently validate observation in the original strains generated by *Low et al.* on 129 background and further backcrossed to C57BL6 for 7 generation, owing to the preservation of 129-derived elements on chromosome 1 in the previously reported mice **(G).** Proteomics (**H**) and metabolomics differences across strains at baseline (normoxia) and in response to hypoxia **(I**). Band 3 N-term KO mice also showed different levels of BLVRB proteins, bilirubin/biliverdin ratios **(L)**. Elevation of lactate at steady state (**M**) and synthesis of 2,3-BPG **(N)** were observed in control mice, but not in mice lacking residues 12-23. These impaired metabolic responses to hypoxia were accompanied by significantly lower critical speed (higher exercise intolerance **(O)**) with negligible physiological differences and no significant differences in complete blood counts or hemodynamics parameters **(P)**. RBC Band 3 protein levels were not affected by CS test, but were significantly lower in KO mice, especially BS KO **(Q)**.

The most pronounced changes occurred within the Band 3–hemoglobin subnetwork, corroborating the long-standing Band 3–deoxyhemoglobin model^16^ with direct structural evidence of three-fold increases in interactions between SLC4A1 residues 1, 10 and 56 (very N-terminus), HBA (residues 60, 91, 100, 140) and HBB (18, 66, 90), in the surface accessible region of the inner deoxyhemoglobin tetramer (mapped against 1a3n.pdb in **Figure 3E**). These data demonstrate direct, reversible contact formation between Band 3 and deoxyhemoglobin under low oxygen tension. Consistent with the model, under hypoxia 1,2,3-^13^C_3_-glucose consumption and 2,3-bisphosphoglycerate (2,3-BPG) synthesis increased two-fold (**Figure 3F**), in keeping with displacement to cytosol and activation of otherwise bound/inhibited glycolytic enzymes^17^ (e.g., GAPDH^25^). Thus, structural rearrangement of the Band 3–hemoglobin interface is accompanied by functional metabolic reprogramming, linking oxygenation state to cellular energy production.

### Deletion of the Band 3 N-terminus disrupts metabolic remodeling and exercise tolerance in vivo

To probe the physiological relevance of the Band 3 N-terminal region, we generated humanized transgenic mice in which the N-terminal 1-45 amino acids are replaced with the human Band 3 amino acid sequence 1-35, or truncated variants, lacking either residues 1–11 (the High Affinity – HA binding site for glycolytic enzymes, equivalent to human band 3 *Neapolis*^24,36^) or 12–23 (deoxy-hemoglobin Binding Site – BS^20^ - **Figure 3G**). The original colonies had been generated by Low’s lab and reportedly had 10% lower RBC counts and 6% lower hematocrit, with no other significant complete blood count abnormalities at baseline.^20^ For the mice reported here, all lines were re-established on a congenic C57BL/6J background, after observing excess iron-dependent lipid peroxidation in RBCs from the NIH repository-obtained original mice, due to residual contamination from the 129 background on chromosome 1 (*Steap3* coding region^37^), despite 7 generation back-crossing with the C57BL6 mice.^38^ No hematological abnormalities were observed in the new mice (n=7; **Supplementary Figure 5A**), for which ablation of band 3 residues 1-11 and 12-23 were confirmed via mass spectrometry, upon digestion with either trypsin (first cleavage site at residue 56) or AspN in separate runs.

RBCs from these mice were extensively characterized through omics approaches at baseline, showing elevation in proteasome components in HA and BS KO mice compared to humanized controls, and elevation in autophagy and apoptosis in BS KO mice (**Figure 3H**). When exposed to hypoxia, widespread metabolic differences were observed (**Figure 3I**). Of note, KO mice displayed altered BLVRB protein levels (highest in the cytosol of HA KO mice) and bilirubin-to-biliverdin ratios, consistent with lower BLVRB sequestration in HA KO mice and higher biliverdin metabolism in both KOs – as manifested by higher biliverdin/bilirubin ratios in HA KO mice or lower total biliverdin in BS KO ones (**Figure 3H-L**). Above all, wild-type RBCs increased lactate output (**Figure 3M**) and 2,3-BPG synthesis – as gleaned by steady state and 1,2,3-^13^C_3_-glucose tracing (**Figure 3N**). Glycolytic fluxes were significantly blunted in both HA and BS KO mice, with blunt Rapoport-Luebering shunt specifically observed in the BS KO mice.

In vivo, BS KO animals showed markedly reduced exercise endurance on treadmill-based critical-speed testing, defined as the highest work intensity at which a mouse can maintain a physiological steady state^39^ (**Figure 3O**). Of note, the lower exercise tolerance of KO mice was accompanied by normal cardiac and pulmonary performance, as measured via hemodynamics, with the only significant difference noted being higher final body weight in HA KO mice (**Figure 3P; Supplementary Figure 5B**). Deletion of the N-term of band 3 had measurable negative effects on exercise tolerance (a milder defect in HA KO mice, 5%, p>0.05; 15% drop in BS KO, p<0.001). These differences were mirrored by progressively aggravated differences in post-exercise RBC and plasma metabolome and RBC proteomes – with SLC4A1 levels paralleling drops in exercise tolerance (**Figure 3Q**), and markers of hypoxia (lactate, hypoxanthine), oxidant stress (pentose phosphate isomers, methionine sulfoxide) and mitochondrial dysfunction (succinate, fumarate, malate) increasing after exercise the most in BS KO mice, followed by HA KO and humanized band 3 controls (**Supplementary Figure 6**). Altogether, our results confirm that the N-terminus of Band 3 is essential for adaptive metabolic remodeling during hypoxia and exertion. These results establish that the N-terminal cytosolic domain of Band 3 is required for physiological flexibility and energy management in circulating RBCs.

### BLVRB directly binds the Band 3 N-terminus through defined structural interfaces

The recurring detection of Band 3–BLVRB cross-links, and the effects of genetic deletion of band 3 N-term on BLVRB protein levels and biliverdin metabolism, prompted us to assess whether these proteins interact directly and what was the structural specificity. Human BLVRB and progressive N-terminal truncations of Band 3 (full 1-404, 1-379, or just the very N-term residues 1–56, 1–30) were recombinantly expressed and purified for biophysical analysis (**Figure 4; Supplementary Figure 7**).

**Figure 4.**
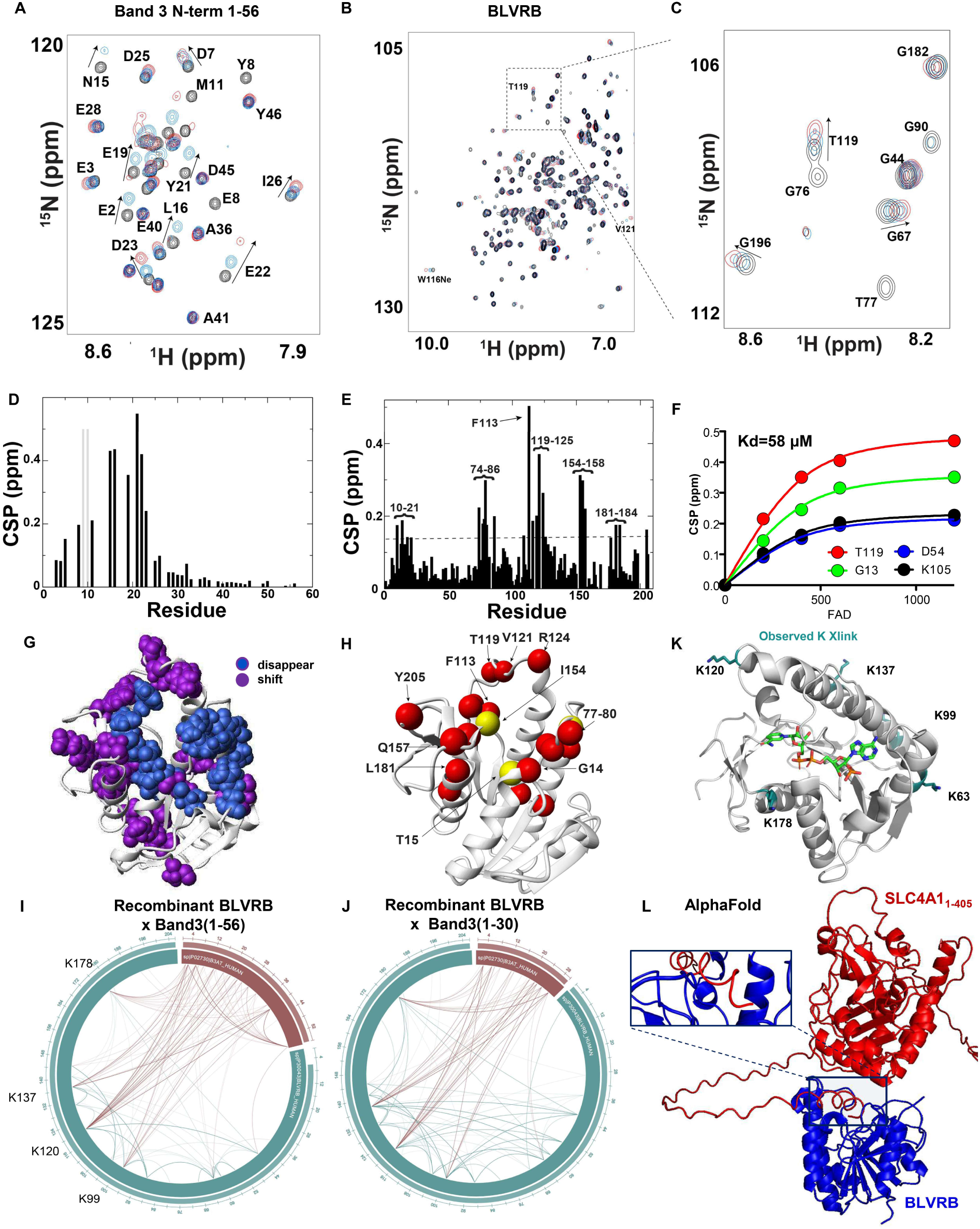
Biophysical validation of a Band 3–BLVRB interaction by NMR and Xlinking proteomics. Unsupervised analysis of cross-linking proteomics date identified BLVRB as a novel Band 3 interactor. Recombinant expression of BLVRB and band 3 N-terminus (either residues 1-379, 1-56 and 1-30) allowed to perform biophysical validations of this observation, including Nuclear Magnetic Resonance (NMR) of ^15^N-labeled band 3 upon dosing of BLVRB **(A)** or vice versa **(B)** (HSQC zoom in **(C)**). Chemical Shift Perturbations (CSP) reveal hotspot of interactions around residues 10-12 and 20-23 of band 3 **(D)** and 110-125 (proximal to the active site S111), 74-86 and 154-158 of BLVRB **(E).** CSP of key residues as a function of BLVRB substrate (FAD) confirm direct interaction with μM affinity **(F)** in proximity to the BLVRB active site, as confirmed via mapping of CSP on BLVRB structure **(G-H).** CSP comprises measured shifts (dark black in **D** and blue in **G**) and those resonances that disappear (light bars in **D** and purple in **G).** These interactions were validated via Xlinking proteomics of BLVRB to Band 3 1-56 or 1-30 (**I-J**), as mapped against BLVRB structure **(K)** and modeled via AlphaFold **(L)**.

Heteronuclear NMR titrations of ¹⁵N-labeled Band 3 peptides with unlabeled BLVRB induced discrete chemical-shift perturbations (CSPs) in Band 3 residues 10–12 and 20–23, while reciprocal titrations revealed BLVRB perturbations in residues 74–86, 110–125, and 154–158 (**Figure 4A–C**). The latter regions encompass the catalytic site and adjacent helical domain, indicating a docking interface near the FAD-binding cleft (**Figure 4F-H**).

Since BLVRB is a flavin reductase, titration of BLVRB with FAD monitored by NMR CSP revealed residue-specific binding responses, with the largest perturbations observed for G13 and T119 near the catalytic cleft (S111), yielding a dissociation constant (K_d_) of ∼58 µM, consistent with a moderate-affinity, reversible interaction characteristic of flavoproteins (**Figure 4F**). Cross-linking mass spectrometry confirmed covalent linkages between Band 3 (1–56) and BLVRB (100, 120, 137) (**Figure 4K-J**), validating the contact sites observed in solution. Of note, Xlinking proteomics studies confirmed the loss of the band 3 – BLVRB interaction when probing recombinant proteoforms of band 3 lacking the vey N-term residues (i.e., 57-379; 350-391 – **Supplementary Figure 7**). Titration of BLVRB with increasing concentrations of Band 3_1–56_ peptides induced residue-specific NMR chemical shift perturbations, with the largest displacements observed for L33 and M98 near the catalytic pocket and smaller changes at distal residue A162, confirming a direct, saturable interaction between the two proteins (**Supplementary Figure 7I**).

Computational docking using AlphaFold Multimer accurately recapitulated this orientation, placing the Band 3 N-terminus along the catalytic groove of BLVRB (**Figure 4L; Supplementary Figure 7G**). These findings demonstrate that BLVRB is a *bona fide* Band 3-binding redox enzyme, poised to integrate heme metabolism and oxygen sensing at the membrane–cytosolic interface.

### Oxygen tension regulates Band 3–BLVRB association in intact RBCs

To determine whether the Band 3–BLVRB complex forms and dissociates dynamically in living cells, we examined primary human RBCs under normoxic and hypoxic conditions using immunofluorescence confocal microscopy, proximity-ligation assays (PLA) and super-resolution microscopy (**Figure 5**). Under normoxia, BLVRB co-localized extensively with Band 3 at the plasma membrane, appearing as discrete puncta distributed along the cytoskeleton (**Figure 5A–B**). Upon hypoxia, this pattern shifted: BLVRB fluorescence redistributed toward the cytosol, and median PLA signal intensity between Band 3 and BLVRB decreased by >70 % (**Figure 5C–D**). Stimulated Emission Depletion (STED) imaging confirmed the spatial separation of the two proteins in hypoxic cells, with the estimated percentage of BLVRB-Band 3 interactions dropping from 55 to 30% under hypoxic conditions (**Figure 5E–F**).

**Figure 5.**
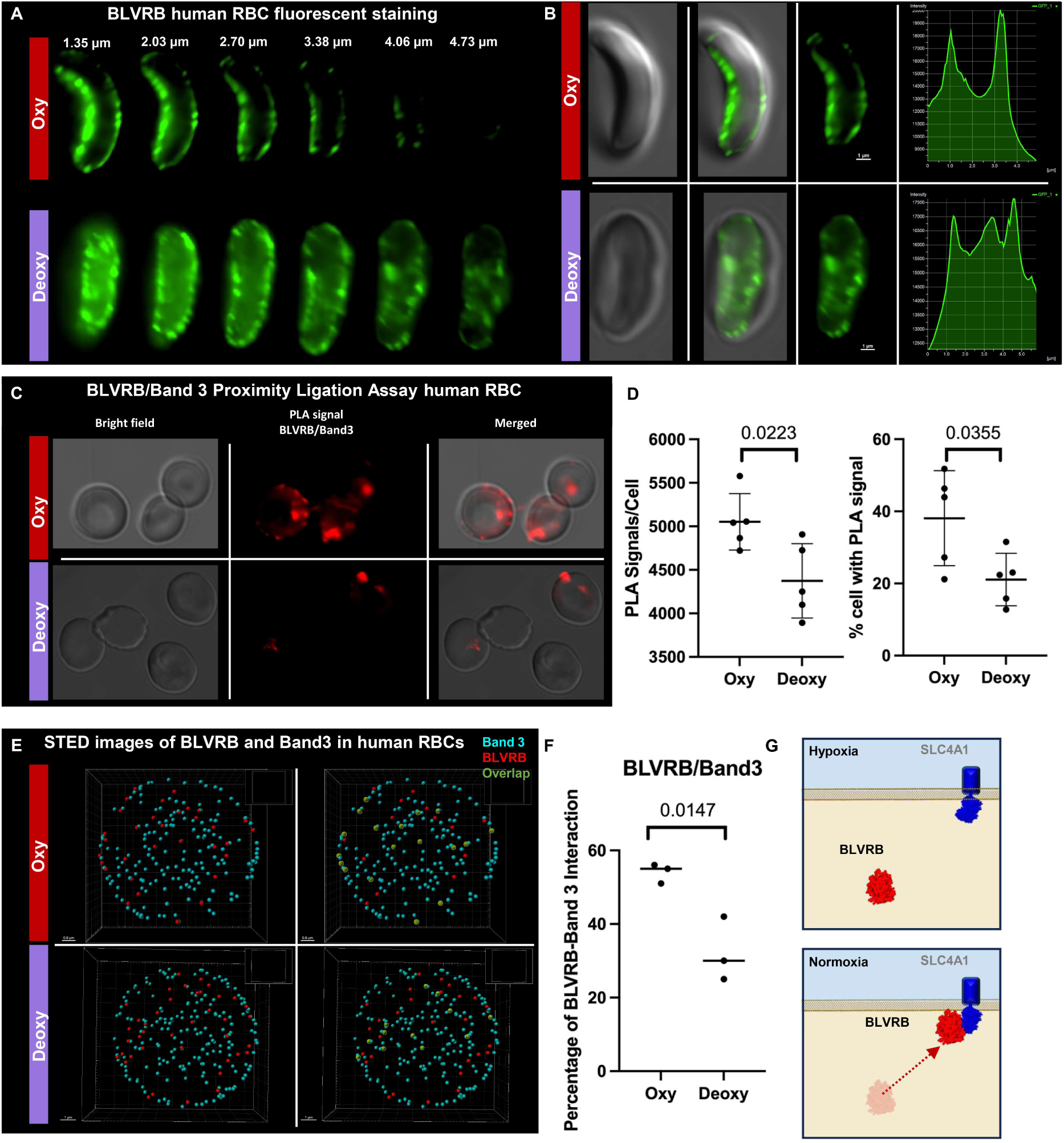
Band 3–BLVRB interactions are regulated by normoxia/hypoxia status in intact RBCs. High-resolution visualization of Band 3–BLVRB complexes and their hypoxia-dependent dynamics. Immunofluorescence microscopy of human RBCs showing BLVRB localization to the membrane in normoxia and cytosol in hypoxia **(A-B)**. Proximity ligation assay (PLA) quantification of Band 3–BLVRB interactions in normoxia, but not hypoxia (n = 5 donors, *p* < 0.001, paired t-test) **(C-D).** Simulated Emission Depletion (STED) super-resolution microscopy demonstrates loss of co-localization of Band 3 and BLVRB under hypoxia (**E-F**). Proposed model **(G**).

These results establish that indeed Band 3-BLVRB interact in vivo, that these interactions are oxygen-sensitive and reversible, supporting the notion that the RBC interactome dynamically remodels in response to physiological oxygen gradients. The dissociation of BLVRB from Band 3 under hypoxia parallels the displacement of glycolytic enzymes by deoxyhemoglobin, suggesting that similar conformational cues orchestrate multiple, concurrent remodeling events.

### Population studies confirm cis-genetic effects on BLVRB and SLC4A1 protein levels but fail to link RBC biliverdin levels to polymorphic *BLVRB,* suggestive of alternative metabolic roles

We have established that BLVRB is a Band 3 interactor under normoxia and that docking near the catalytic pocket modulates BLVRB activity, and we showed that truncating the Band 3 N-terminus in mice blunts hypoxic responses and reduces exercise tolerance without cardiopulmonary defects. We next asked whether common genetic variation - less severe than the rare pathogenic Band 3 Neapolis deletion yet more prevalent in the population - recapitulates aspects of this module in humans. In a cohort of 13,091 healthy blood donors, protein quantitative trait locus analyses revealed robust cis-pQTLs for both SLC4A1 (common missense mutation K56E and D38A in the very N-term region - relevant to the present study, and V862I, linked to ovalocytosis and distal renal tubular acidosis^40^) and BLVRB, confirming that local genetic variation modulates their steady-state protein abundance in RBCs (**Supplementary Figure 8A–E**). Notably, SLC4A1 pQTL signals co-segregated with indices of RBC hemolytic stress, consistent with the structural and metabolic roles defined above (**Supplementary Figure 8D**). By contrast, metabolite QTLs for biliverdin - the canonical BLVRB substrate - did not show a strong genome-wide adjusted signal at BLVRB; instead, associations mapped to other loci, indicating that biliverdin levels in mature RBCs are largely determined by trans-acting pathways and substrate supply rather than BLVRB coding variation (**Supplementary Figure 9A–F**). Specifically, top mQTL hits for biliverdin were on G6PD (biliverdin reductases are NADPH-dependent) and UGT1A1 (UDP-glucoronosyl transferase - primary enzyme responsible for converting unconjugated bilirubin into water-soluble, excretable conjugated bilirubin). BLVRB and BLVRA catalyze the final reduction step of distinct biliverdin isomers: BLVRB preferentially acts on biliverdin IXβ (and to a lesser extent IXγ and IXδ), while BLVRA reduces biliverdin IXα, the principal product of heme oxygenase (HO1/HO2) activity on heme^41^. In mammals, biliverdin IXα originates from enzymatic heme catabolism via HO1 and HO2, generating carbon monoxide, ferrous iron, and biliverdin IXα as products. By contrast, biliverdin IXβ arises largely through non-canonical or oxidative heme cleavage reactions independent of heme oxygenase, particularly in erythroid or hypoxic contexts. Interestingly, BLVRB has also been shown to exhibit ferric reductase activity, with an apparent K(m) of 2.5 μM for the ferric iron, a reaction that requires NAD(P)H and FMN. Heme cleavage in the foetus produces non-alpha isomers of biliverdin and ferric iron, both of which are eligible substrates for BLVRB^42^. Quantitative proteomics from our Deep Red resource indicate that heme oxygenase isoforms are scarce in mature RBCs - only ∼1,000 copies/cell of HO1 and ∼850 copies/cell of HO2 - whereas BLVRB is exceptionally abundant (∼13 million copies/cell vs ∼950k copies/cell for BLVRA – **Supplementary Table 1**). This stark stoichiometric imbalance and the poor GWAS linkage to biliverdin together suggest that BLVRB’s physiological functions in mature erythrocytes may extend well beyond canonical biliverdin IXβ reduction. In line with the recent literature^41^, BLVRB may thus operate as a pleiotropic redox regulator, coupling its FAD-dependent oxidoreductase activity^43^ to other biological functions.

### The Band 3/BLVRB module mediates *S*-Nitroso transfer within RBCs

Recently, elegant work from the Stamler lab has suggested a role for BLVRB as a S-NO-CoA-assisted *S-*nitrosyl transferase^44^. Given the proximity of reactive cysteines in both Band 3 and BLVRB, and the established nitrite reductase activity of hemoglobin,^27^ which generates NO and methemoglobin^45^ – a substrate for BLVRB as an NADPH-dependent methemoglobin reductase - we hypothesized that this module participates in nitric oxide (NO) transfer reactions. Overnight incubation of recombinantly expressed, purified BLVRB with *S*-Nitrosoglutathione (GS-NO) or S-nitroso-CoA (SNO-CoA) resulted in selective nitrosation of BLVRB C109 (up to 36%) and C188 (up to 9.5%), directly verified by mass spectrometry fragmentation and NMR – with highest CSPs observed upon GSNO incubation around residues 72-74, 107-114, 149-154, 178-187 (**Figure 6A**). Please, note that while both *S*-Nitrosylation and *S*-Nitrosation appear in the literature to describe formation of *S*-Nitrosothiols, here we use *S*-Nitrosation (transfer of the nitrosonium ion (NO+)) to a thiol/thiolate rather than S-nitrosylation (coupling of NO to another radical or metal) to describe these reactions. We feel *S*-Nitrosation better mechanistically describes observed protein S-nitrosothiols formation upon treatment with GSNO or *S*-Nitroso CoA in these experiments.

**Figure 6.**
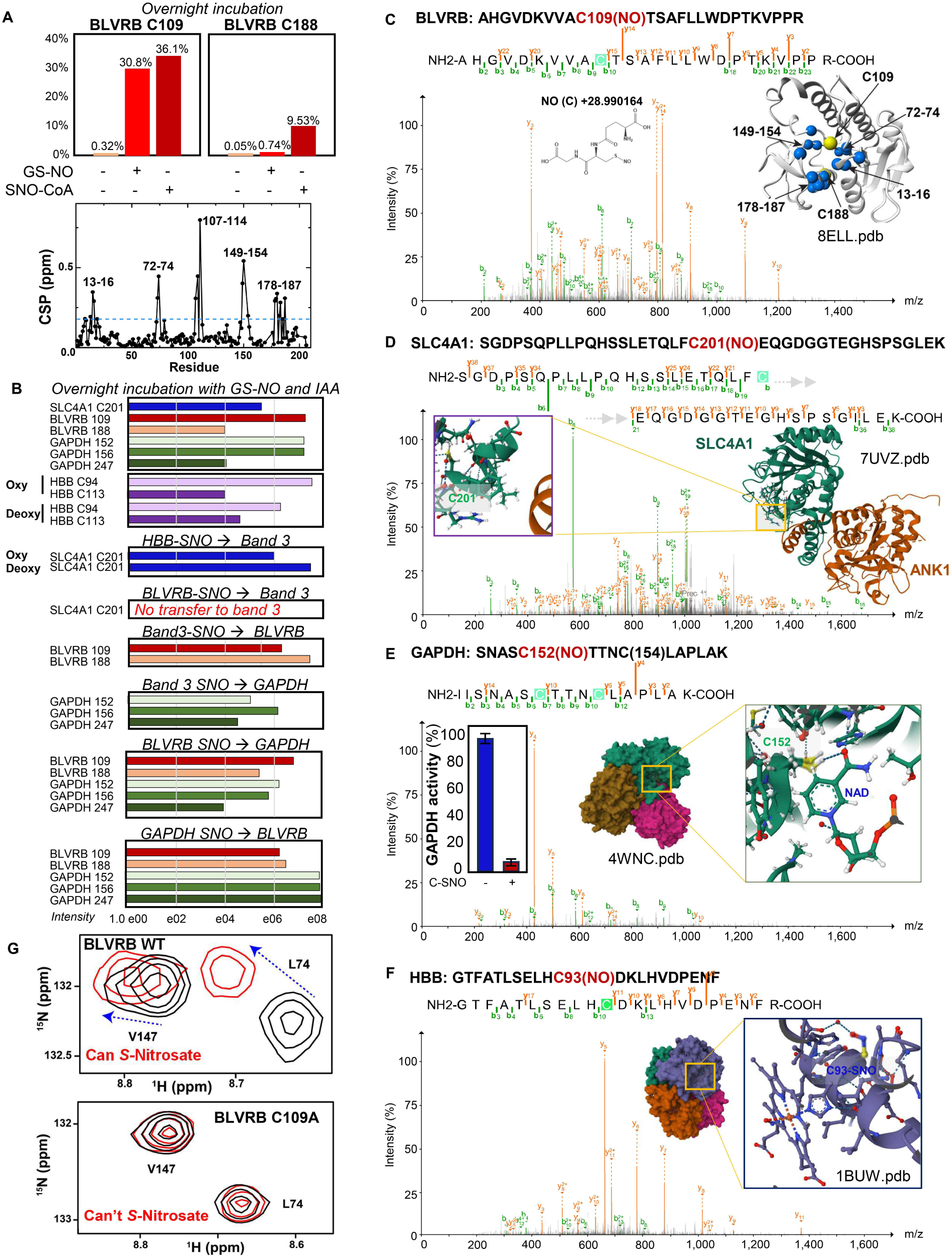
The Band 3/BLVRB (C109) – hemoglobin interactome mediates *S-*Nitroso-transferase activity. Overnight incubation with of BLVRB with *S*-Nitroso-Glutathione (GSNO) or *S*-Nitroso-CoA (SNO-CoA) or both can favor *S-*Nitrosation of BLVRB at cysteine residues 109 and 188 **(A)**, and results in noticeable chemical shift perturbation (CSP) especially in residues 107-114 (around the active site S111). In-vitro assays were performed to determine whether HBB, SLC4A1, BLVRB, GAPDH can be *S*-Nitrosated at key functional cysteine residues in vitro **(B).** Results show that HBB (C94, 113), BLVRB (C109, 188), SLC4A1 (C201) and GAPDH (C152, 156, 247) can be *S*-Nitrosated in vitro, that oxy HBB loads NO better than deoxy, but deoxy-HBB increases five-fold Band 3 S-NO formation only at C201 (not C317). S-NO Band 3 can trans-nitrosate BLVRB at C109 and 188; both S-NO Band 3 and S-NO BLVRB can trans-nitrosate GAPDH cysteines, but BLVRB can also load NO from GAPDH-SNO **(B)**. Representative mass fragmentation spectra are shown for relevant peptides for BLVRB, SLC4A1, GAPDH and HBB (C93) **(C-F**), the latter being inhibited by *S*-nitrosation of C152 in the active site, as shown by a direct activity assay in vitro (indent in **E**). Mutation (to Alanine residues) of BLVRB show that the S-Nitroso transferase activity is dependent upon preservation of C109, but not C188 **(G)**.

In vitro GS-NO incubation of pure recombinant protein, confirmed *S*-Nitrosation adducts on Band 3 (C201), GAPDH (C152, C156, C247) and hemoglobin β (C93 – 94 counting initiator methionine - and 113), on top of validating the SNO-adducts on BLVRB C109 and C188, as determined via mass spectrometry (**Figure 6A–F**). Of note, experiments in normoxia vs hypoxia showed that HBB C93 is up to 5-fold more efficient in trans-*S*-Nitrosating Band 3 (C201, but not 317) under hypoxia (**Figure 6B**). When *S*-Nitrosated BLVRB was mixed with reduced Band 3 (1–405), the nitroso group did not transfer to Band 3 C201. On the other hand, *S*-Nitrosated band 3 (C201) can transfer NO to BLVRB at both C109 and 188, but preferentially to the latter (∼4fold more). Both S-Nitroso BLVRB C109-C188 and S-Nitroso C201 Band 3 – independently or mixed - successfully trans-nitrosated GAPDH C152, 156 and 247, demonstrating a *S*-Nitroso-transferase activity. 3 C201 (S-NO spectrum in **Figure 6D**) is involved in the cytoskeletal interaction of band 3 with ankyrin, but not with the DIDS-sensitive ionophore activity.^46,47^ *S*-Nitrosation of GAPDH at the catalytic cysteine (C152) markedly inhibited enzymatic activity (**Figure 6E**), consistent with reversible metabolic gating. Of note, this observation validates prior reports of S-NO-mediated GAPDH inhibition in vitro^48^ and in plants^49^, where the resulting glycolytic blockade diverts flux toward the pentose phosphate pathway to regenerate ribulose-5-phosphate for RuBisCO-dependent carbon fixation. Co-incubation of in vitro nitrosated GAPDH (at C152, 156 and 247) with BLVRB promoted *S*-Nitrosation of C109 and 188.

To further mechanistically probe the role of BLVRB as a *S*-Nitroso-transferase, BLVRB C109A mutants – but not C188A ones - lose the ability to transfer the NO group (**Figure 6G**), confirming this residue as the catalytic thiol. In summary, these observations define a Band 3–BLVRB–GAPDH–HBB redox relay that integrates oxygenation, NO metabolism, and energy control within the RBC cytosol (**Figure 6D–G**).

### Hypoxia-induced nitrosation remodels RBC redox signaling and metabolism

To assess how oxygen tension affects global *S*-Nitrosation, first we performed enrichment for *S*-Nitrosated proteins with PBZyn^50^ to quantify changes under normoxia and hypoxia in cytosol vs membranes. Both in mouse and human RBCs, hypoxia caused a pronounced increase in cytosolic PBZyn-reactive proteins, particularly from Band 3, BLVRB, GAPDH (**Figure 7A–B**). Membrane localization of *S*-nitrosated proteins decreased, consistent with redistribution of NO equivalents toward soluble protein pools or membrane-releasing effect of GAPDH and BLVRB S-NO modifications (**Figure 7A-C**).

**Figure 7.**
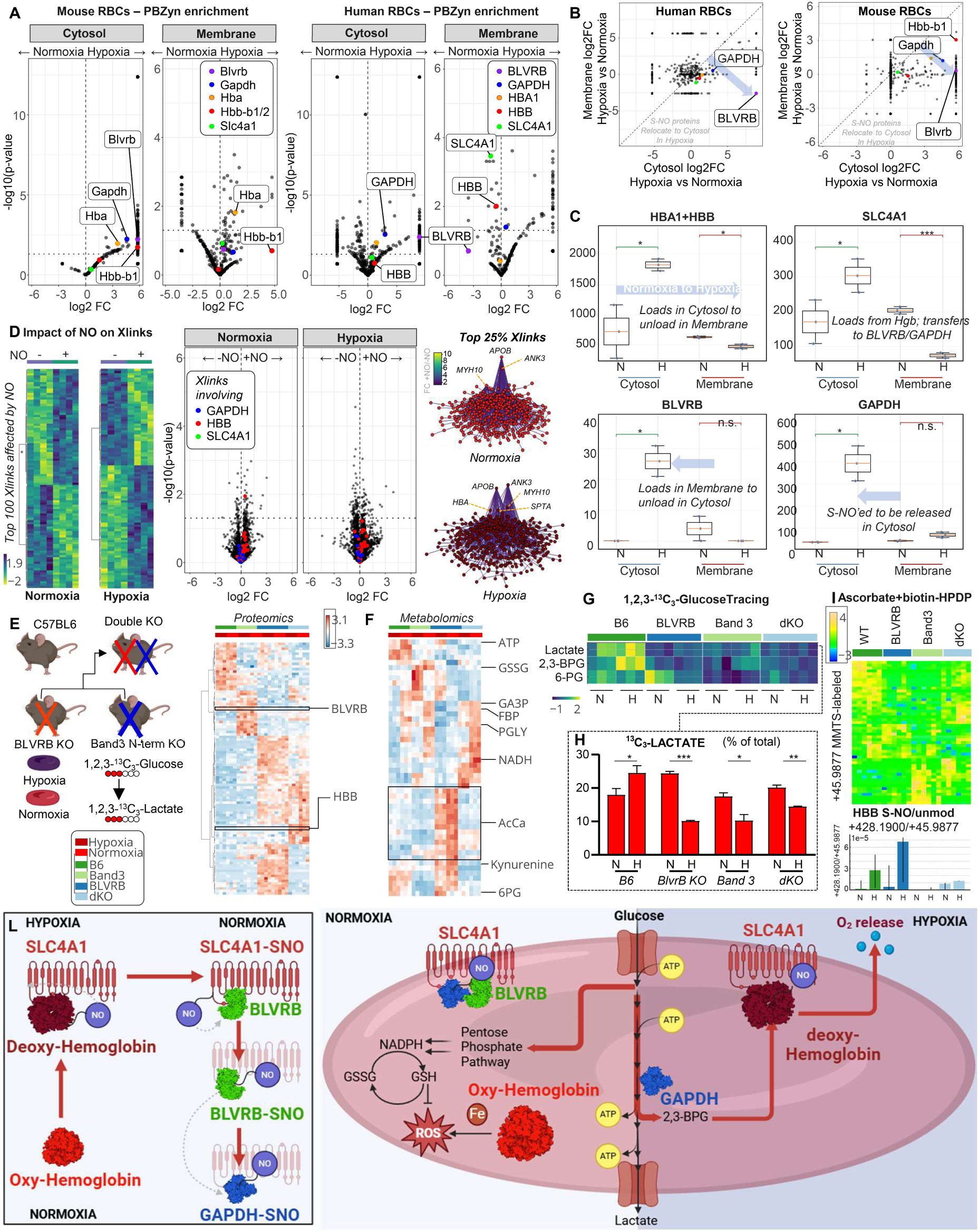
Hypoxia-responsive *S*-Nitrosation network and functional integration of Band 3 and BLVRB. S-nitroso peptide enrichment using PBZyn reagent in normoxic and hypoxic murine and human RBCs **(A)** reveals enrichment in *S*-NO-modified proteins, including BLVRB and GAPDH in the cytosol in hypoxia **(B)**. Comparative analysis of *S-*Nitrosated proteins showing selective compartment-specific induction under hypoxia (higher in cytosol vs membrane) **(C)**. Xlinking proteomics characterization of human RBCs with or without NO supplementation shows widespread alteration of the interactome both in normoxia or hypoxia, including changes to the hemoglobin, GAPDH and SLC4A1 interactomes, with hotspot changes noted in structural membrane proteins (ANK3, SPTA, MYH10, APOB) **(D)**. Phenotypic analysis of mice lacking the Band 3 N-terminal high-affinity domain (ΔHA, residues 1–11) and/or BLVRB (single and double KO) **(C)** significantly decreases BLVRB levels **(E)** and is accompanied by depletion of ATP pools, alterations in glycolysis and redox homeostasis, compatible with lower energetic state, higher oxidant stress and membrane lipid remodeling, including accumulation of hemolytic propensity markers kynurenine and acyl-carnitines. Metabolic tracing experiments with 1,2,3-^13^C_3_-glucose reveal failure to activate glycolysis, synthesize 2,3-BPG and shut down the pentose phosphate pathway in BLVRB KO and double KO mice in response to hypoxia **(F).** Enrichment of S-Nitroso-proteins via mild reduction with ascorbate and desthiobiotin labeling confirms increased *S*-Nitrosation of HBB in hypoxia, altered NO unloading in both BLVRB, SLC4A1 N-term single and double KO mice **(I).** Proposed model summarizing the Hb–Band 3–BLVRB-GAPDH axis as a hypoxia-responsive NO-transfer module that tunes RBC glycolytic enzyme release from the membrane and activity **(L)**.

To understand whether NO treatment is sufficient to impact the RBC interactome, cross-linking proteomics studies of normoxic or hypoxic RBCs were conducted with or without NO supplementation. Results showed that *S*-Nitrosation remodels the RBC interactome, including interactors of HBB, SLC4A1 and GAPDH, with the highest degree nodes in the interactome network impacting structural proteins (ANK3, MYH10, APOB, SPTA - **Figure 7D**).

To test functional relevance of BLVRB in this system, first, we generated constitutive *Blvrb* KO and conditional EpoR*-Cre Blvrb*^fl/fl^, and confirmed ablation of BLVRB protein expression in mature RBCs through omics approaches, which also indicated strain-specific responses to hypoxia (**Supplementary Figure 10**). Next, we examined RBCs from Band 3 HA KO (band 3 *Neapolis*), EpoR*-Cre Blvrb*^fl/fl^, and double-deficient mice (**Figure 7E**). After validating BLVRB ablation in the single and double KOs through proteomics (**Figure 7E**), we observed that BLVRB KO RBCs – like band 3 KO and double KOs - failed to up-regulate glycolysis under hypoxia, as gleaned by both at steady state metabolomics (**Figure 7F**) and tracing of ^13^C-glucose-derived carbons into lactate and ^13^C_3_-BPG (**Figure 7G-H**). Ablation of either BLVRB, Band 3 N-terminus or both was accompanied by depletion of ATP pools, lower glucose consumption, blockade of glycolysis at the fructose bisphosphate and glyceraldehyde 3-phosphate steps. BLVRB KO mice were uniquely characterized by elevation in acyl-carnitines and kynurenine, markers of stress-associated membrane lipid remodeling and increased hemolytic fragility.^51,52^ Interestingly, 6-Phosphogluconate, both at steady state and in tracing experiments, was depleted in WT mice in hypoxia – consistent with glucose oxidation fluxes being redirected towards glycolysis – but not in Band 3 N-term KO mice and even less so in BLVRB KO mice (**Figure 7F**).

Finally, to complement PBZyn studies in the context of genetic ablation of Band 3 N-term and/or BLVRB, normoxic or hypoxic RBC lysates from WT and KO mice were treated with methyl methanethiosulfonate (MMTS) to block free thiols, followed by mild reduction with ascorbate to reduce *S*-Nitrosated peptides and labeling with biotin-HPDP. Results showed a decrease in MMTS-blocked peptides (C(+45.9877 Da)) and an increase in biotin-tagged C(+428.1900 Da) in hypoxia, an effect further exacerbated for S-NO HBB under hypoxia in BLVRB KO mice (**Figure 7I**).

Collectively, our results delineate a hierarchical architecture of RBC regulation, organized around Band 3 as a structural and redox scaffold. Based on the collective data generated, we propose the following model (**Figure 7F**), which is consistent with and expands upon previous ones proposed by Low et al.^20^ and Pawlowski et al.^53^. Under oxygenated conditions, BLVRB is tethered to Band 3, maintaining antioxidant balance through NADPH, methemoglobin reduction and generation of bilirubin. Upon deoxygenation, *S*-NO-deoxyhemoglobin binds the Band 3 N-terminus, transferring NO to band 3 at C201 – triggering remodeling of the ANK1 complex, while also displacing both BLVRB and GAPDH. This process is in part regulated by NO transfer to GAPDH C152-156, an inhibitory PTM that is promoted by transfer of S-NO at C201 of band 3 or C109 of BLVRB onto GAPDH. Our *in vitro* data also suggest that cytosolic BLVRB can load NO from S-nitrosated GAPDH, which would favor its reactivation, promoting 2,3-BPG production, through a sort of regulatory ping-pong mechanism involving NO transfer between BLVRB and GAPDH. At the same time, it should be noted that the canonical NADPH-dependent methemoglobin reductase activity of BLVRB may also further modulate C93 S-nitrosation/unloading dynamics, by favoring a right-shift in the equation of nitrite reduction to NO by deoxyhemoglobin, through recycling of HbFeIII back to the active reactant HbFeII.^45^ These dynamic transitions integrate oxygen sensing, nitric oxide signaling, and metabolic control within a single, protein-level regulatory circuit.

## DISCUSSION

Once regarded as inert vessels for gas transport, red blood cells (RBCs) are now recognized as metabolically flexible, redox-responsive systems. The present study extends this transformation in our understanding by revealing a well-regulated and organized proteome - much more complex than previously reported - and a dynamic O_2_-responsive interactome, centered on Band 3 (SLC4A1) – deoxyhemoglobin interactions. Our analysis identifies biliverdin reductase B (BLVRB) as a new player in what appears to be a dynamic metabolon that integrates oxygen tension, redox chemistry, and nitric oxide (NO) signaling. Through multi-omic, structural, and genetic approaches, we demonstrate that reversible binding between Band 3 and BLVRB constitutes an oxygen-sensitive regulatory switch that orchestrates metabolic remodeling and NO transfer within the mature erythrocyte.

Our findings refine and extend the classical Low model of the Band 3–deoxyhemoglobin axis.^16,17,20,35^ We show that the same N-terminal domain that recruits glycolytic enzymes and hemoglobin also directly engages BLVRB, one of the ten most abundant RBC proteins. Under normoxia, BLVRB docks to Band 3 in proximity to its catalytic pocket, constraining its flavin-dependent activity. Hypoxia promotes deoxyhemoglobin binding to this site, releasing both glycolytic enzymes and BLVRB to the cytosol, where BLVRB gains access to its substrates and nitrosation partners. This reversible complex formation provides a mechanistic basis for how an anucleate cell transduces environmental oxygen fluctuations into biochemical reprogramming.

Population-scale proteome-wide association analyses reinforce this framework. Common SLC4A1 missense variants (D38A, K56E) and BLVRB cis-pQTLs modulated protein abundance, whereas biliverdin levels did not show strong BLVRB genotype association, suggesting broader oxidoreductase functions beyond canonical biliverdin IXβ reduction^41^. The stark stoichiometric disparity between scarce heme oxygenases (HO-1/HO-2; <2,000 copies per cell) and highly abundant BLVRB (∼13 million copies per cell) further supports its pleiotropic redox role. In agreement with recent studies by Stamler and colleagues^44^, we show that BLVRB mediates S-nitroso-transfer reactions, passing NO equivalents to GAPDH and hemoglobin. S-nitrosation of GAPDH at its catalytic cysteine reversibly inhibits glycolysis, thereby diverting glucose toward the pentose-phosphate pathway for NADPH regeneration and antioxidant defense.

Genetic and physiological evidence corroborate this concept. Truncation of the Band 3 N-terminus in humanized mice blunted glycolytic activation under hypoxia, disrupted 2,3-BPG synthesis, and impaired exercise tolerance, despite preserved cardiopulmonary performance - phenotypes reminiscent of the human Band 3 Neapolis variant. These results confirm that the 1–23 amino-acid segment of Band 3 is indispensable for metabolic plasticity and energy homeostasis in circulating RBCs. The concurrent alterations in BLVRB abundance and biliverdin/bilirubin ratios further underscore that this interaction coordinates redox and heme-derived metabolism with oxygen-dependent energy control. Of note, Band 3 N-term KO mice favor glycolysis over the pentose phosphate pathway, owing to the decreased inactivation of glycolytic enzymes with direct binding to Band 3;^25^ however, their phenotype remains different from that of G6PD deficient mice, which unexpectedly appear to have increased exercise tolerance compared to control mice.^54,55^ This observation is in part explained by recent evidence suggesting that exercise-induced oxidant stress limits both glycolysis and NADPH-generation capacity,^56^ and in part are explained by other potential regulatory mechanisms of glycolysis beyond enzyme inhibition by binding to Band 3. One example could be oxidation of redox sensitive thiols of rate-limiting glycolytic enzymes, such as GAPDH.^57^ The present study offers an intriguing, novel explanation, involving a potential role for GAPDH S-nitrosation as a complementary molecular strategy that RBCs have evolved to slow down glycolysis in response to environmental clues. As GAPDH and BLVRB both migrate to Band 3 in normoxia, S-NO transfer may contribute to further inactivating glycolysis beyond direct interactions, a process that would be rate-limited by NO availability rather than full occupancy of available Band 3 N-term domain, addressing in part a common critique to the band 3 glycolytic metabolon model.

Remarkably, this reaction recapitulates a conserved mechanism observed in plants, where *S*-Nitrosation of GAPDH under high-NO conditions or nitrosative stress inhibits glycolysis to shunt carbon toward the Calvin–Benson cycle^49^, fueling RuBisCO-dependent CO₂ fixation. That evolution converged on the same molecular logic - using *S*-Nitroso GAPDH to toggle between energy and redox balance - in photosynthetic cells that fix CO₂ and in erythrocytes that transport O₂ is a striking example of biochemical symmetry across kingdoms. Both systems deploy NO-based post-translational control to couple environmental gases to core carbon metabolism, underscoring the universality of redox signaling as a regulatory language of life. This interpretation - that BLVRB regulates glycolytic and pentose phosphate fluxes through GAPDH S-nitrosation - carries potential clinical implications, particularly for nitrate poisoning, where the standard therapy, methylene blue, serves as a pharmacologic cofactor for BLVRB in methemoglobin reduction.^58^

Importantly, in the present study a deeper characterization of the insoluble membrane proteome allowed us to revise the estimated abundance of Band 3, from 1 to 6-7 million copies/cell, further mitigating stoichiometry concerns against the model.

Mutation studies suggest that while both C188 and C109 can be *S*-Nitrosated, only the latter is essential for the *S*-Nitroso-transferase activity. Of note, C109 resides in proximity to the active site S111. Sequence alignment of BLVRB orthologs from mammals, birds, and reptiles shows that Cys109 is invariant across species, consistent with strong evolutionary constraint and supporting its role as a conserved redox-active residue mediating S-nitroso transfer.

Altogether, the Band 3–BLVRB–hemoglobin axis exemplifies how a transcriptionally silent cell achieves environmental responsiveness through reversible protein–protein interactions and post-translational chemistry. By integrating oxygen sensing, nitrosative signaling, and metabolic gating within a single structural module, this system transforms the erythrocyte into a distributed molecular sensor of tissue hypoxia. Beyond redefining RBC physiology, these findings highlight fundamental principles of convergent evolution linking oxygen transport, carbon fixation, and redox-regulated metabolism across the tree of life.

Despite the extensive work that went into the present study, several limitations ought to be acknowledged. It is unclear whether the deleterious effects on exercise tolerance elicited by the ablation of band 3 N-terminus can be extended to other physiologically relevant context of oxidant stress or hypoxia, such as high-energy demanding conditions (e.g., fat burning, thermoregulation) or pathological conditions leading to oxidant stress or enzyme-dependent fragmentation of the intrinsically disordered N-term domain of Band 3, such as RBC storage in the blood bank^25,59^, or infections^23^. Indeed, ablation or fragmentation of the N-terminus of band 3 leads to a progressive loss of metabolic modulation^60^ in oxidant stress in vitro, leading to the cytosolic relocation of glycolytic enzymes under pseudo-hypoxic conditions (e.g., blood storage^25,59^ or COVID-19^61^). It remains to be tested whether BLVRB ablation phenocopies some of these pathological effects. *S*-Nitrosated proteins were here enriched through S-NO-reactive reagents^50^, but remain hard to directly detect without prior enrichment strategies in physiologically relevant contexts. At the net of the limitations noted above, the present studies shares with the literature the most extensive RBC proteome and interactome mapping to date, while identifying a novel mechanism of oxygen-dependent protein thiol *S*-Nitrosation that involves not just Band 3 and deoxyhemoglobin, but also BLVRB as a previously unrecognized player.

## ONLINE METHODS

### Preparation of pure mature RBCs

Whole blood was obtained from 6 healthy donor volunteers at Columbia University in New York, NY, USA, upon signing of informed consent. Whole blood from 5 healthy volunteers was collected into EDTA tubes. Blood was spun at 800g for 10 minutes, plasma was removed, and 50uL of packed RBCs from each donor was combined prior to leukoreduction via filtration (Pall Medical) to remove log4 (99.99%) and log2.5 residual white blood cells (WBCs) and platelets (PLTs), respectively, as standard workflow in modern blood banks in most US states.^62^. The mixed RBCs were washed in PBS and 200uL of packed RBCs was resuspended in 10mL PBS and stained with antibodies CD45 PE-Cy7 (1:400), CD41 APC-Cy7 (1:400), CD235a BV605 (1:1000), and CD71 PE (1:100) for 20 minutes at room temperature. Samples were washed with PBS and resuspended to a final volume of 1mL. Immediately before sorting, 30uL of RBCs was diluted into 10mL PBS supplemented with 20uL thiazole orange (TO). Purified RBC populations were achieved using BD FACSAria cell sorters. CD45+ white blood cell and CD41+ platelets were excluded from RBC singlets and reticulocytes (CD235a+TO+) and mature RBCs (CD23a+ CD71-TO-) were sorted. Aliquots of sorted cells were re-analyzed on the cell sorter to determine purity, indicating >99.99% mature RBCs.

### GeLC-MS

Proteomic analyses were performed as previously described using a GeLC-MS approach. Briefly, 10 μL of RBCs were lysed in 90 μL of distilled water and incubated with 20 mM N-ethylmaleimide (NEM) for 30 minutes at room temperature in the dark. The samples were then loaded onto a 1.5 mm thick NuPAGE Bis-Tris 4–12% gradient gel (Invitrogen) and visualized using SimplyBlue™ SafeStain (Invitrogen). Excised gel bands were destained with 50% acetonitrile in 50 mM ammonium bicarbonate and subsequently dehydrated with 100% acetonitrile. Modified sequencing-grade trypsin (Promega) was added at a 1:50 (enzyme:substrate, w/w) ratio, and samples were digested overnight at 37 °C. The digestion was quenched by adding 5% formic acid, and organic solvent was removed using a SpeedVac concentrator. Peptides were then desalted and concentrated using Thermo Scientific Pierce C18 Tips prior to LC-MS analysis.

### Fractionation

The mature RBCs were processed through a stepwise extraction protocol using CHAPS and high salt, followed by guanidine hydrochloride and chemical digestion with hydroxylamine hydrochloride (HA) in Gnd-HCl. This procedure generated three distinct fractions: cellular, soluble ECM (sECM), and insoluble ECM (iECM). Each fraction was subjected to proteolytic digestion using a filter-aided sample preparation (FASP) protocol with 10 kDa molecular weight cutoff filters (Sartorius Vivacon 500, Sartorius, Göttingen, Germany; #VN01H02). Samples were alkylated with 20 mM N-ethylmaleimide (NEM) for 30 minutes at room temperature in the dark, then digested overnight with trypsin at an enzyme-to-substrate ratio of 1:100 at 37 °C. Peptides were recovered from the filters using successive washes with 0.2% formic acid.

Digested samples were separated by high-pH reversed-phase chromatography on a Gemini-NH C18 analytical column (4.6 × 250 mm, 3 µm particles) at a flow rate of 0.6 mL/min. The mobile phases consisted of 10 mM ammonium bicarbonate (pH 10) as phase A and 10 mM ammonium bicarbonate with 75% acetonitrile (pH 10) as phase B. Peptide separation was achieved using the following linear gradient: 0–5% B in 10 min, 5–50% B in 80 min, 50–100% B in 10 min, followed by a 10 min hold at 100% B. A total of 96 fractions were collected during LC separation and concatenated into 24 pooled fractions by combining every 24th fraction (e.g., 1, 25, 49, 73, etc.). Samples were then dried in a SpeedVac, desalted, and concentrated using Thermo Scientific Pierce C18 Tips.

### LC–MS Data Acquisition

The samples were analyzed on a nano-UHPLC (nanoElute, Bruker Daltonics) coupled to a timsTOF SCP mass spectrometer (Bruker Daltonics). Samples were separated on an Aurora Ultimate column 25 cm x 75 mm, C18 1.7 mm (IonOptics) which was heated to 50 °C in a column oven. Mobile phases consisted of 0.1% (v/v) formic acid in water (phase A) and acetonitrile (phase B). Samples were separated on a 90 min stepped gradient ranging from 4-35% B at a flow rate of 300 nL/min. The gradient is built of sequential steps ranging from 4 to 25% B in 60 min, 25 to 35% B in 20 min, and finally, 35 to 95% in 10 min. The performance of the mass spectrometer was monitored by standardized measurements of a K562 tryptic digest (Pierce), where the number of peptides and the total ion chromatogram (TIC) shape were considered.

The mass spectrometer was operated in PASEF mode. The accumulation and ramp time was set to 166 ms and 10 PASEF MS/MS scans per topN acquisition cycle were acquired. MS and MS/MS spectra were recorded from *m/z* 100 to 1700. The ion mobility was scanned from 0.6 to 1.40 Vs/cm^2^. Precursors for data-dependent acquisition were isolated within ± 1 Th and fragmented with an ion mobility-dependent collision energy, which was linearly increased from 20 to 65 eV in positive mode. Low-abundance precursor ions with an intensity above a threshold of 1000 counts but below a target value of 12500 counts were repeatedly scheduled and otherwise dynamically excluded for 0.2 min.

### Database Searching and Protein Identification

Raw data files conversion to peak lists in the MGF format, downstream identification, validation, filtering and quantification were managed using FragPipe version 13.0. MSFragger version 3.0 was used for database searches against a mouse database with decoys and common contaminants added. The identification settings were as follows: Trypsin, Specific, with a maximum of 2 missed cleavages, up to 2 isotope errors in precursor selection allowed for, 10.0 ppm as MS1 and 0.04 Da as MS2 tolerances. Oxidation of M (+15.994915 Da), ), Hydroxylation of P (+15.994915 Da), Acetylation of protein N-term (+42.010565 Da), Pyrolidone from peptide N-term Q or C (-17.026549 Da) and Cys conjugation by NEM were set as variable modifications.

The copy number of protein was calculated as described,^8^ using the relationship

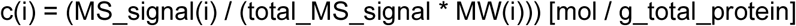

where and *N*_A_ is the Avogadro number. We assumed the mean volume of RBCs of 90 fL and the average total protein content per cell as 20% of RBCs mass.

### Band 3 Mouse Models

Novel humanized mouse models were generated at the University of Virginia carrying either (i) human canonical band3 (HUB3), replacing murine residues 1-45 with human 1-35 given sequence divergencies in the N-terminal region, or two different band3 knock out variations: (ii) high affinity deletion (first 11 amino acids deleted, HA-Del) or (iii) binding site knock out (amino acid residues 12-23, BS-KO). The original AE1 mice were generated by the Low lab on a 129 background, prior to backcrossing for 6-7 generations to C57BL6.^20^ These new mice were generated *de novo* on a clean C57BL6/J background. All animal studies were reviewed and approved by the University of Virginia Institutional Animal Care and Use Committee (protocol n: 4269).^63^

### BLVRB Mouse Models

Novel BLVRB mouse models were generated at the University of Virginia either (i) constitutive KO (BLVRB KO); (ii) conditional KO (EpoR-Cre BLVRB^fl/fl^), both generated on a C57BL6/J background.

### Treadmill exercise and constant speed tests

Prior to the determination of critical speed (CS), mice completed a treadmill familiarization phase, which consisted of four ∼5 min runs on a motor-driven rodent treadmill (Exer 3/6, Columbus Instruments, Columbus, Ohio, USA). For the first several runs, the treadmill speed was maintained at 10-15 m/min (up a 5° grade, which was maintained throughout all treadmill tests). For the last several runs, the speed of the treadmill was increased progressively over the last minute to ∼30-35 m/min to familiarize the mice with high speed running. Animals were encouraged to run with intermittent bursts of compressed room air aimed at the hind limbs from directly above the animal (so as not to push the mouse up the treadmill). All treadmill testing protocols were designed and conducted by experienced staff and strictly followed the guidelines set by the American Physiological Society’s resource book for the design of animal exercise protocols.^64^

The CS was determined using a modified version of the methodology used by Copp et al, 2010^65^ for rats, as well as following the guidelines set forth by Poole et al.^64^ After completion of the treadmill familiarization period, each mouse performed 3-5 runs to exhaustion, in random order, at a constant speed that resulted in fatigue between 1 and 15 minutes (speeds ranging from 30-50 m/min). Each test was performed on separate days with a minimum of 24 hours between tests. For each constant-speed trial, mice were given a 2-minute warm-up period where they ran at 15-20 m/min followed by a 1-minute period of quiet resting. To start the test, the treadmill speed was increased rapidly over a 10 second period to the desired speed at which point a stopwatch was started. Testing was terminated and time to exhaustion was measured to the nearest tenth of a second whenever the mouse could no longer maintain pace with the treadmill despite obvious exertion of effort. A successful constant-speed test was determined if 1) the mouse could quickly adapt to the treadmill speed at the beginning of the test (e.g., did not waste energy), 2) a noticeable change in gait occurred preceding exhaustion (i.e., lowering of the hindlimbs and rising of the snout), and 3) the animal’s righting reflex was markedly attenuated when placed on their back in a supine position (an unexhausted quadruped will typically attempt to right themselves within ∼1 second).

### Data modeling for the determination of CS

Following successful completion of the constant speed treadmill tests, the CS and finite distance capacity (D’) were calculated for each mouse using the linear 1/time model (Speed=D^’×1/time+CS) as described previously.^66,67^ In this model, the treadmill speed used for the constant speed test is plotted as a function of the inverse of time to exhaustion and the y-intercept of the regression line yields the CS and the slope is the D’.

### Hemodynamics

After the CS test, mice underwent terminal open chest right ventricular (RV) and left ventricular (LV) function measurements with a 1.2F, FTE-1212B-4018 pressure volume catheter (Transonic Systems Inc., Ithaca, NY) inserted by direct cardiac puncture. Mice were induced inhaled isoflurane (4-5%), and tracheal incision (∼ 1 cm) was performed. Next, a tracheal tube was inserted and connected to an Anesthesia Workstation or Hallowell EMC Microvent and an anesthetic plain was maintained at 1.0-2.5% isoflurane in 100% oxygen. After which, a thoracotomy was performed exposing the heart, the pericardium was resected, and a small hole made at the base of the RV/LR with a 30g needle for insertion of the pressure-volume catheter. Steady state hemodynamics are collected with short pauses in ventilation (up to 10 seconds) or high frequency oscillatory ventilation to eliminate ventilator artifact from the pressure-volume recordings. Occlusions of the inferior vena cava were performed by applying pressure to the inferior vena cava (up to 10 seconds) through the abdominal opening. After pressure volume and hemodynamic measurements completed mice were humanely euthanized by exsanguination and cervical dislocation. Data was recorded continuously using LabScribe2 and analyzed offline.

### Biotin-Switch Assay and Streptavidin Pull-Down

Lysed RBCs were treated to block free thiols by adding methyl methanethiosulfonate (MMTS) to a final concentration of 20 mM, followed by incubation at 50°C for 20 min with gentle shaking to prevent disulfide formation. Excess MMTS was removed by acetone precipitation (three volumes of cold acetone, −20°C for 1 h, followed by centrifugation at 5,000 × g for 5 min). The protein pellet was resuspended in 100 µL of HEN buffer (250 mM HEPES, 1 mM EDTA, 0.1 mM neocuproine, pH 7.7). S–NO bonds were reduced by adding ascorbate to a final concentration of 20 mM, followed by biotinylation with biotin-HPDP (30 µL of a 4 mM stock in DMSO) for 2 h at room temperature in the dark. Excess reagents were again removed by acetone precipitation. Pellets were resuspended in 25 µL of 10× diluted HEN buffer and 75 µL of neutralization buffer (25 mM HEPES, 100 mM NaCl, 1 mM EDTA, 0.5% Triton X-100, pH 7.5).

For enrichment of biotinylated proteins, 50 µL of pre-washed streptavidin agarose bead slurry was added to each sample, and samples were rotated overnight at 4°C. Beads were washed five times with neutralization buffer containing 500 mM NaCl and once with 50 mM ammonium bicarbonate (ABC). Bound proteins were subjected to on-bead digestion by resuspending the beads in 50 mM ABC containing 5 mM DTT and incubating at 56°C for 30 min to cleave disulfide bonds, including the HPDP linker. Trypsin was added at a 1:50 (enzyme:protein) ratio (∼3 µg per sample), and digestion was carried out overnight at 37°C with agitation at 600 rpm. The resulting peptides were collected for LC–MS/MS analysis.

Positive and negative controls were included in each experiment. The positive control consisted of GSNO-treated samples, while the negative control was processed identically but without the addition of ascorbate during the reduction step to confirm the specificity of S–nitrosation detection.

The samples were analyzed on a nano-UHPLC (nanoElute, Bruker Daltonics) coupled to a timsTOF SCP mass spectrometer (Bruker Daltonics). Samples were separated on an Aurora Ultimate column 25 cm x 75 mm, C18 1.7 mm (IonOptics) which was heated to 50 °C in a column oven. Mobile phases consisted of 0.1% (v/v) formic acid in water (phase A) and acetonitrile (phase B). Samples were separated on a 90 min stepped gradient ranging from 4-35% B at a flow rate of 300 nL/min. The gradient is built of sequential steps ranging from 4 to 25% B in 60 min, 25 to 35% B in 20 min, and finally, 35 to 95% in 10 min. The performance of the mass spectrometer was monitored by standardized measurements of a K562 tryptic digest (Pierce), where the number of peptides and the total ion chromatogram (TIC) shape were considered.

The mass spectrometer was operated in PASEF mode. The accumulation and ramp time was set to 166 ms and 10 PASEF MS/MS scans per topN acquisition cycle were acquired. MS and MS/MS spectra were recorded from *m/z* 100 to 1700. The ion mobility was scanned from 0.6 to 1.40 Vs/cm^2^. Precursors for data-dependent acquisition were isolated within ± 1 Th and fragmented with an ion mobility-dependent collision energy, which was linearly increased from 20 to 65 eV in positive mode. Low-abundance precursor ions with an intensity above a threshold of 1000 counts but below a target value of 12500 counts were repeatedly scheduled and otherwise dynamically excluded for 0.2 min. Raw data files conversion to peak lists in the MGF format, downstream identification, validation, filtering and quantification were managed using FragPipe version 13.0. MSFragger version 3.0 was used for database searches against a human database with decoys and common contaminants added. Search parameters were set for trypsin-specific cleavage allowing up to two missed cleavages, with up to two isotope errors permitted in precursor selection. The precursor and fragment mass tolerances were set to 25 ppm (MS1) and 0.04 Da (MS2), respectively.

Dynamic modifications included oxidation of methionine (+15.9949 Da), MMTS-labeled cysteine (+45.9877 Da), and biotin-tagged cysteine (+428.1900 Da) to account for modifications introduced during the biotin-switch procedure.

### Recipient Epidemiology and Donor evaluation Study (REDS) RBC Omics

A detailed description of the study design, enrollment criteria and main outcomes has been provided in previous publications,^68,69^ and – while redundant with the literature - will be included below with significant overlap with published methods.

### Donor recruitment in the REDS RBC Omics study

#### Index donors

A total of 13,758 donors were enrolled in the Recipient Epidemiology and Donor evaluation Study (REDS) RBC Omics at four different blood centers across the United States (https://biolincc.nhlbi.nih.gov/studies/reds_iii/). Of these, 97% (13,403) provided informed consent and 13,091 were available for metabolomics analyses in this study – referred to as “index donors”. A subset of these donors were evaluable for hemolysis parameters, including spontaneous (n=12,753) and stress (oxidative and osmotic) hemolysis analysis (n=10,476 and 12,799, respectively) in ∼42-day stored leukocyte-filtered packed RBCs derived from whole blood donations from this cohort ^70^. Methods for the determination of FDA-standard spontaneous (storage) hemolysis test, osmotic hemolysis (pink test) and oxidative hemolysis upon challenge with AAPH have been extensively described elsewhere^71^.

### pQTL for SLC4A1 and BLVRB and mQTL for biliverdin

The workflow for the mQTL analysis of biliverdin and pQTL analysis for SLC4A1 and BLVRB protein levels in REDS Index donors is consistent with previously described methods from our pilot mQTL study on 250 recalled donors^72^ and follow up studies on specific metabolic pathways (e.g., glycolysis^73^) other proteins (e.g., GPX4^74^) on the full index cohort. Details of the genotyping and imputation of the RBC Omics study participants have been previously described by Page, et al.^75^ Briefly, genotyping was performed using a Transfusion Medicine microarray^76^ consisting of 879,000 single nucleotide polymorphisms (SNPs); the data are available in dbGAP accession number phs001955.v1.p1. Imputation was performed using 811,782 SNPs that passed quality control. After phasing using Shape-IT ^77^, imputation was performed using Impute2 ^78^ with the 1000 Genomes Project phase 3 ^78^ all-ancestry reference haplotypes. We used the R package SNPRelate ^79^ to calculate principal components (PCs) of ancestry. We performed association analyses using an additive SNP model in the R package ProbABEL^80^ and 13,062 REDS donor study participants who had SLC4A1 and BLVRB protein levels and 13,091 REDS donors who had biliverdin measurements. Datasets were adjusted for sex, age (continuous), frequency of blood donation in the last two years (continuous), blood donor center, and ten ancestry PCs. Statistical (genome-wide adjusted) significance was determined using a p-value threshold of 5x10^-8^. We only considered variants with a minimum minor allele frequency of 1% and a minimum imputation quality score of 0.80. The OASIS: Omics Analysis, Search & Information a TOPMED funded resources^81^, was used to annotate the top SNPs. OASIS annotation includes information on position, chromosome, allele frequencies, closest gene, type of variant, position relative to closest gene model, if predicted to functionally consequential, tissues specific gene expression, and other information.

### High-throughput metabolomics

Metabolomics extraction and analyses in 96 well-plate format were performed as described, with identical protocols for human or murine RBCs^82,83^. For the human and murine studies, RBC samples were transferred on ice on 96 well plate and frozen at -80 °C at Vitalant San Francisco (human RBCs) or University of Virginia (murine RBCs), respectively, prior to shipment in dry ice to the University of Colorado Anschutz Medical Campus. Plates were thawed on ice then a 10 uL aliquot was transferred with a multi-channel pipettor to 96-well extraction plates. A volume of 90 uL of ice cold 5:3:2 MeOH:MeCN:water (*v/v/v*) was added to each well, with an electronically-assisted cycle of sample mixing repeated three times. Extracts were transferred to 0.2 µm filter plates (Biotage) and insoluble material was removed under positive pressure using nitrogen applied via a 96-well plate manifold. Filtered extracts were transferred to an ultra-high-pressure liquid chromatography (UHPLC-MS — Vanquish) equipped with a plate charger. A blank containing a mix of standards detailed before ^84^ and a quality control sample (the same across all plates) were injected 2 or 5 times each per plate, respectively, and used to monitor instrument performance throughout the analysis. Metabolites were resolved on a Phenomenex Kinetex C18 column (2.1 x 30 mm, 1.7 um) at 45 °C using a 1-minute ballistic gradient method in positive and negative ion modes (separate runs) over the scan range 65-975 m/z exactly as previously described.^82^ The UHPLC was coupled online to a Q Exactive mass spectrometer (Thermo Fisher). The Q Exactive MS was operated in negative ion mode, scanning in Full MS mode (2 μscans) from 90 to 900 m/z at 70,000 resolution, with 4 kV spray voltage, 45 sheath gas, 15 auxiliary gas. Following data acquisition, .raw files were converted to .mzXML using RawConverter then metabolites assigned and peaks integrated using ElMaven (Elucidata) in conjunction with an in-house standard library^85^.

### Tracing experiments with labeled 1,2,3-^13^C_3_-glucose

RBCs from all the mouse strains investigated in this study (100 ul) (n=3) were incubated at 37°C for 1h in presence of stable isotope-labeled substrates, as described previously.^86^

### Statistical analysis

Samples were processed and run in randomized order with a technical mixture injected interspersed throughout the run to qualify instrument performance. Data analysis and Statistical analyses – including hierarchical clustering analysis (HCA), linear discriminant analysis (LDA), uniform Manifold Approximation and Projection (uMAP), network plots, heat maps, volcano plots were performed using both MetaboAnalyst 5.0 and RStudio (2024.12.1 Build 563).

### Comparison of Deep Red results vs RBC-GEM

The human RBC-GEM (version 1.2.0)^87^ and the vertebrate homology data from Mouse Genome Informatics database (MGI version 6.24)^88^ were utilized to create a mouse-specific RBC-GEM by mapping human genes to their mouse orthologs, based on a compiled list of 29 published proteomics studies, as detailed in Haiman et al. ^87^

### Recombinant BLVRB and Band 3 protein expression and structural studies

Optimized sequences encoding human BLVRB (full length), Band 3 (SLC4A1) N-terminal residues 1-30; 1-56; 1-379; 1-404; 57-379; or 350-391 were cloned into pET21 or pET28 vector, respectively with an N-terminal 6xHis-TEV-Thrombin sequence (GenScript). Expression and purification of all three constructs were performed similarly. Expression plasmids were transformed into BL21/DE3 or C41 cells for BLVRB and Band 3 N-terminus peptides, respectively (LGC Biosearch Technologies) and starter cultures were grown at 37 °C overnight by inoculating 200 mL of LB with a single colony. The starter culture was used to inoculate 12 L of TB at a 1:25 ratio. The cultures were shaken at 37 °C until an OD of 0.6 was reached. The temperature was reduced to 25 °C and 0.15 mM IPTG was added to induce protein expression once the OD reached 0.8. Cells were harvested by centrifugation after 16 hrs.

All purification steps were performed at 4 °C. Bacterial pellets were resuspended in lysis buffer (50 mM Tris pH 7.5, 300 mM NaCl, 5% glycerol, 10 mM imidazole, 0.1% Triton X-100, 1 mM MgSO_4_). Protease inhibitor cocktail (Xpert Protease Inhibitor Cocktail, GenDEPOT) was added to a final concentration of 0.5X. Lysozyme and Deoxyribonuclease I (Worthington) were added to the lysate. The lysate was subjected to sonication on ice and cell debris was removed by centrifugation for 25 min at 15,000 RPM. Clarified lysate was loaded into a 20 mL Ni-Excel column (Cytiva) and washed with six column volumes of loading buffer (50 mM Tris pH 7.5, 300 mM NaCl, 5% glycerol). G6PD protein was eluted with three column volumes of elution buffer (50 mM Tris pH 7.5, 300 mM NaCl, 400 mM imidazole, 5% glycerol). The eluent was diluted with 20 mM Tris pH 8.0 until a final NaCl concentration of 50 mM was achieved. The diluted protein sample was loaded onto a 12 mL Source 15Q column (Cytiva) and washed with two column volumes of A buffer (20 mM Tris pH 8.0). Protein was eluted with a linear gradient from 0-50% B buffer (20 mM Tris pH 8.0, 1 M NaCl) over eight column volumes with a fraction size of 2 mL. SDS-PAGE was performed on the eluted fractions to identify the peak corresponding to BLVRB or Band 3 – followed by mass spectrometry-based validation of protein sequence coverage.

Fractions containing BLVRB or Band 3 peptides were pooled and concentrated for size exclusion chromatography on a pre-equilibrated (20 mM HEPES pH 7.5, 150 mM NaCl) Superdex 75 pg column (Cytiva). SDS-PAGE was performed on the eluted fractions to identify the peak corresponding to BLVRB or Band 3. The fractions correspond to pure BLVRB or Band 3 peptides were pooled and concentrated by centrifugation using a 10 kDa cutoff concentrator (Sartoris). Protein was aliquoted and kept frozen at -80 °C.

### GAPDH Activity assay

GAPDH activity assays were performed with commercial kits (BioVision, Inc, Milpitas, CA), as previously described.^57,59^

### *S*-Nitrosation Transfer Experiments

Recombinant GAPDH, Band3(1-404) or BLVRB were diluted to a concentration of 100µM in 50mM Bis-Tris/50mM NaCl pH 6.5 buffer. Separately, 5mg DEA NONOate (Enzo Life Sciences, Farmingdale, NY USA) was resolubilized in the same buffer at a concentration of 20mM. This 20mM solution of DEA NONOate was then added to each individual protein solution to a final concentration of 1mM and incubated with shaking at 37°C for 30min to S-nistrosate the target proteins. After incubation, *S*-Nitrosated proteins were mixed at equal molar concentrations with recombinant GAPDH, Band3(1-404) or BLVRB protein to measure transfer of the SNO group to the un-nitrosated protein. These mixed solutions of *S*-Nitrosated/un-nitrosated proteins were incubated at 25°C for 1hr upon which they were sampled for LC-MS/MS analysis. Immediately upon sampling, SNO transfer samples were diluted in 8M Urea to a final concentration of 4M and 200mM N-Ethylmaleimide (stock diluted in LC-MS grade H2O) was added to a final concentration of 20mM to alkylate any unreacted cysteines. This alkylation reaction was allowed to proceed for 20min at 25°C for 20min. After alkylation, proteins samples were subjected to an in-solution trypsin digest (sequencing-grade Trypsin (Promega) added at an enzyme:substrate (w/w) ratio of 1:10) for 3hr at 25°C. Post-digestion, the samples were acidified with 10% (v/v) FA and peptides were cleaned up using Pierce C18 Spin Tips (Thermo Fisher Scientific, Waltham, MA). LC-MS/MS analysis was conducted on a Thermo Fisher Orbitrap Fusion Lumos Mass Spectrometer coupled to an EASY-nLC 1200 nano LC system. An Aurora Ultimate column 25 cm x 75 mm, C18 1.7 mm (IonOpticks, Collingwood VIC, Australia) heated to 50 °C in a column oven was used for peptide separation. A/B buffers for the analysis were 0.1%FA in H2O and 0.1% FA in 80% (v/v) Acetonitrile, respectively, and the gradient used for the experiment was a 5% B to 40% B gradient over 55min followed by a 4min 40% B to 68% step at a flowrate of 450nL/min for the entire analysis. The mass spectrometer was operated in positive ion, data-dependent mode with MS1 scans run in the orbitrap from 375-1600 m/z at 120,000 resolution. Full scan automatic gain control (AGC) and maximum injection time was set to 4E5 ions and 50 ms. Ions above an intensity threshold of 1E4 were selected for MS2 analysis with fragmentation by collision induced dissociation (CID) (30% power). Filtering was performed by the quadrupole with an isolation window of 1.6 m/z. AGC and maximum injection time was set to 1E4 ions and 50 ms.

### Xlinking proteomics with TMT10

To interrogate structural remodeling of the RBC proteome variants in normoxia or hypoxia, cross-linking proteomics with tandem mass tag (TMT2) multiplexing was performed under room air or hypoxic conditions in a glove box. Recombinant proteins (0.5 mg/mL) were incubated with 1 mM di(sulfosuccinimidyl)suberate (DSSO) or 4-(4,6-dimetoxy-1,3,5-triazin-2-yl)-4-metylmorpholinium – DMTMM at 25 °C for 30 min, followed by quenching with 50 mM Tris-HCl (pH 7.5). Cross-linked proteins were digested with trypsin, and peptides were labeled with TMT10 reagents according to the manufacturer’s instructions (Thermo Fisher). Labeled peptides were pooled, fractionated by high-pH reversed-phase chromatography, and analyzed by nanoLC-MS/MS on an Orbitrap Fusion Lumos mass spectrometer. Cross-linked peptides were identified using the XlinkX search node in Proteome Discoverer, and relative abundances across conditions (e.g., normoxia vs hypoxia) were quantified by TMT reporter ion intensities. Similar studies were conducted in presence or absence of DEA NONOate at 20 mM.

### Nuclear Magnetic Resonance and structural models

NMR studies were performed as described.^25,43^ ^15^N-heteronuclear single quantum coherence (HSQC) spectra were collected at 25°C for the recombinantly expressed band 3 peptide 1-30, 1-56 in the presence of 100 and 200 uM BLVRB. Data were collected on a Varian 900 using a standard ^15^N-HSQC sequence.

## Supporting information

Supplementary Table - Raw data and Elaborations

Supplementary Resource - Deep Red

## Author Contribution

Animal studies: AH, FIC, DCI, JCZ. RBC purification: KH; Proteomics: AVI, MD, SB, KCH. RBC biology: MD, KH, NM, ADA. Metabolomics and lipidomics analyses: JAR, DS, FC, TN, ADA. Biostatistics and Bioinformatics: GRK, GPP, ADA. Structural Studies and SNO Biochemistry: AVI, SB, JAR, FV, SBK, EZE, ADA; Systems biology models: ZBH, AMK, BOP; Vein-to-vein database: NHR; mQTL and pQTL analyses: GRK, GPP. Figure preparation: AVI, ZBH, GRK, ADA. Microscopy studies: SZ, ADO. Conceptualization: ADA. Writing and finalization: first draft by AD and all co-authors reviewed and approved the final version.

## Data and Materials availability

The Deep Red database is accessible at https://angelo-dalessandro.github.io/deep-red-supplementary-site/. All raw data and elaborations are included in Supplementary Table 1.xlsx. The BLVRB KO or conditional EpoR-Cre BLVRB^fl/fl^, or the new humanized Band 3 N-term canonical or deficient mice (HA Del and BS KO) – the latter generated on a clean C57BL6/J background and based on prior models by Low’s group - are available upon reasonable request, finalization of material transfer agreement and after institutional ACUC approval through Dr James C Zimring Lab at the University of Virginia. Further information and requests for resources and reagents should be directed to and will be fulfilled by the Lead Contact, Angelo D’Alessandro.

