## Supplementary Figures for "Deep Red Blood Cell Proteome Defines the Band 3 N-Terminus Interactome as a Regulator of Hypoxic Adaptation via BLVRB-Dependent *S*-Nitroso Transfer"

***Corresponding author:**

**SUPPLEMENTARY TABLE 1** .......... XSLX

### SUPPLEMENTARY FIGURES

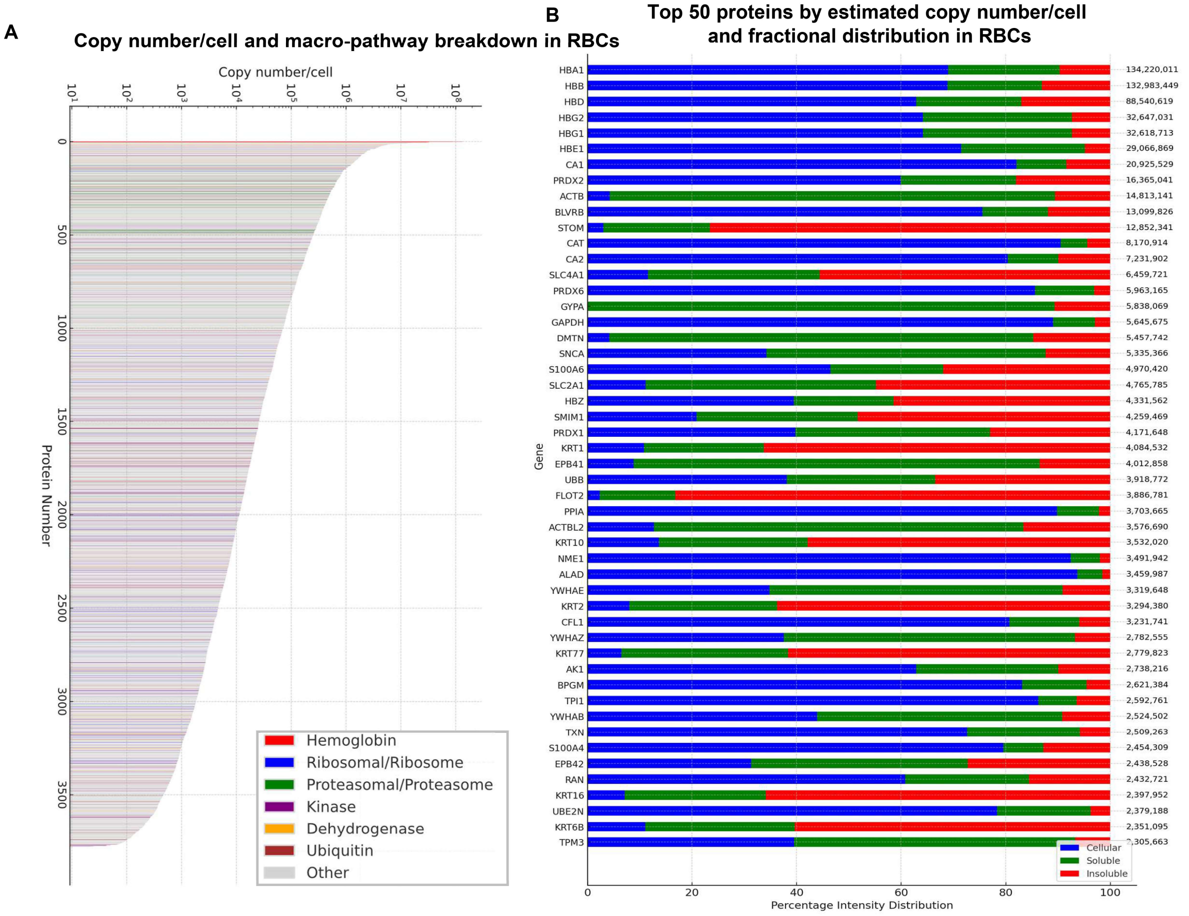

**Supplementary Figure 1. Overview of proteomics results.** Estimated copy number/cell across 3,750 proteins in ultra-pure RBCs, color-coded by class **(A)** and overview of fractional distribution across cytosol (cellular), soluble or insoluble membrane fractions for the top 50 proteins **(B)**.

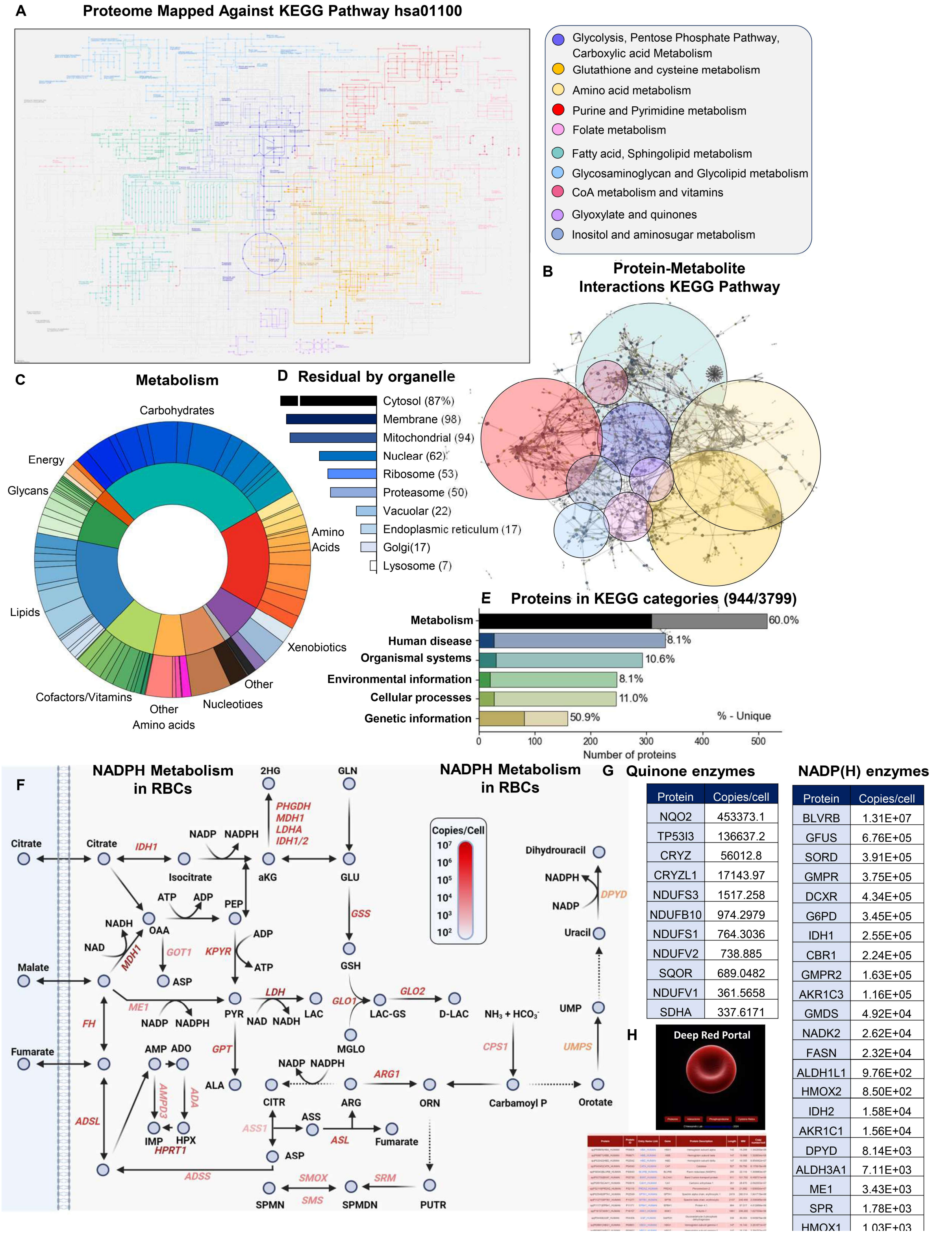

| Protein | Copies/cell |
|---|---|
| NQO2 | 453373.1 |
| TP53I3 | 136637.2 |
| CRYZ | 56012.8 |
| CRYZL1 | 17143.97 |
| NDUFS3 | 1517.258 |
| NDUFB10 | 974.2979 |
| NDUFS1 | 764.3036 |
| NDUFV2 | 738.885 |
| SQOR | 689.0482 |
| NDUFV1 | 361.5658 |
| SDHA | 337.6171 |

| Protein | Copies/cell |
|---|---|
| BLVRB | 1.31E+07 |
| GFUS | 6.76E+05 |
| SORD | 3.91E+05 |
| GMPR | 3.75E+05 |
| DCXR | 4.34E+05 |
| G6PD | 3.45E+05 |
| IDH1 | 2.55E+05 |
| CBR1 | 2.24E+05 |
| GMPR2 | 1.63E+05 |
| AKR1C3 | 1.16E+05 |
| GMDS | 4.92E+04 |
| NADK2 | 2.62E+04 |
| FASN | 2.32E+04 |
| ALDH1L1 | 9.76E+02 |
| HMOX2 | 8.50E+02 |
| IDH2 | 1.58E+04 |
| AKR1C1 | 1.56E+04 |
| DPYD | 8.14E+03 |
| ALDH3A1 | 7.11E+03 |
| ME1 | 3.43E+03 |
| SPR | 1.78E+03 |
| HMOX1 | 1.03E+03 |

**Supplementary Figure 2. Mapping of proteomics data against metabolic pathways and gene ontologies.** Protein entries mapped against Kegg hsa01100 color-coded by pathway **(A)**. Protein-metabolite interaction map in KEGG **(B)** and protein ontology by metabolic pathway **(C),** organelle origin **(D)** and KEGG categories **(E)**. Highlights of NADPH-associated metabolic reactions catalyzed by enzymes detected in ultra-pure RBCs **(F)**, including quinone and **NADPH**-dependent enzymes **(G).** Preview of the Deep Red database **(H)**.

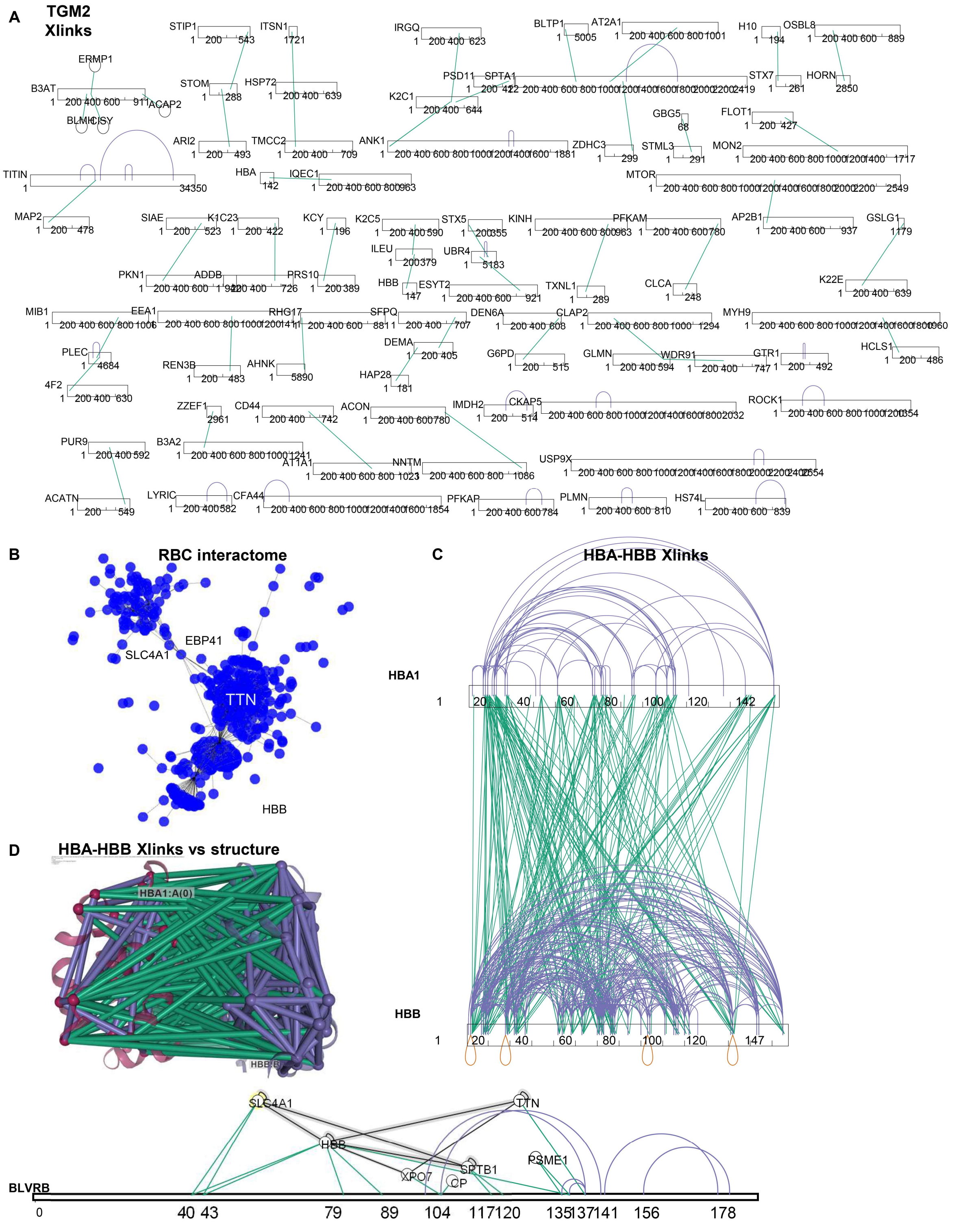

**Supplementary Figure 3. Endogenous and chemical cross-links.** Endogenous TGM2 cross-links (A) and overview of DMTMM and DSSO-generated interactomics networks (B), with highlights on HBA-HBB against the established structure of the oxy-hemoglobin tetramer (C-D).

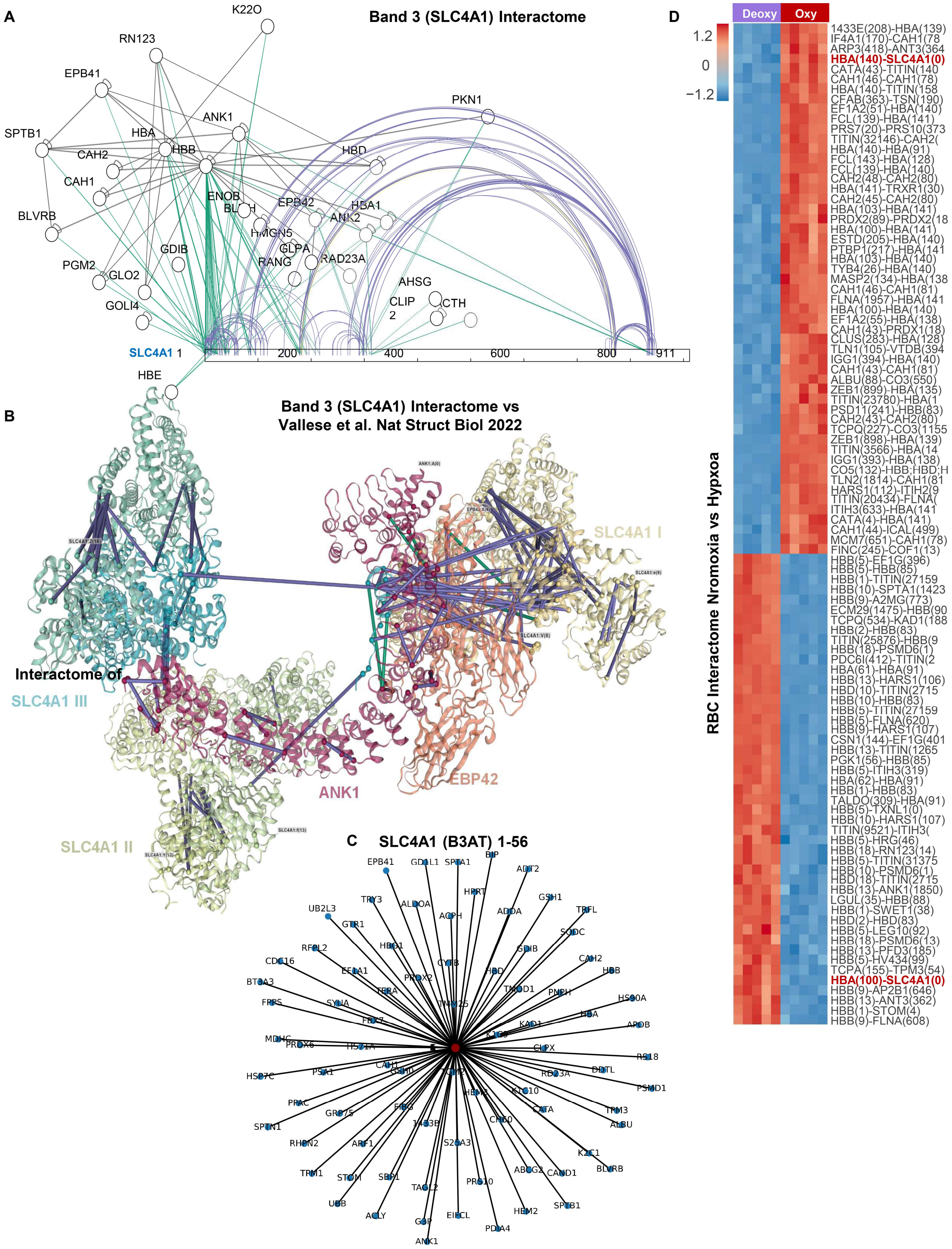

**Supplementary Figure 4. Band 3 cross-links (A), mapped against the structural interactome with ankyrin and spectrin (from Vallese et al. (B).** Focus on the interactome of the N-terminus 1-56 of band 3 **(C)**. Heat map of the top 100 most significant interactome changes in hypoxia vs normoxia **(D)**.

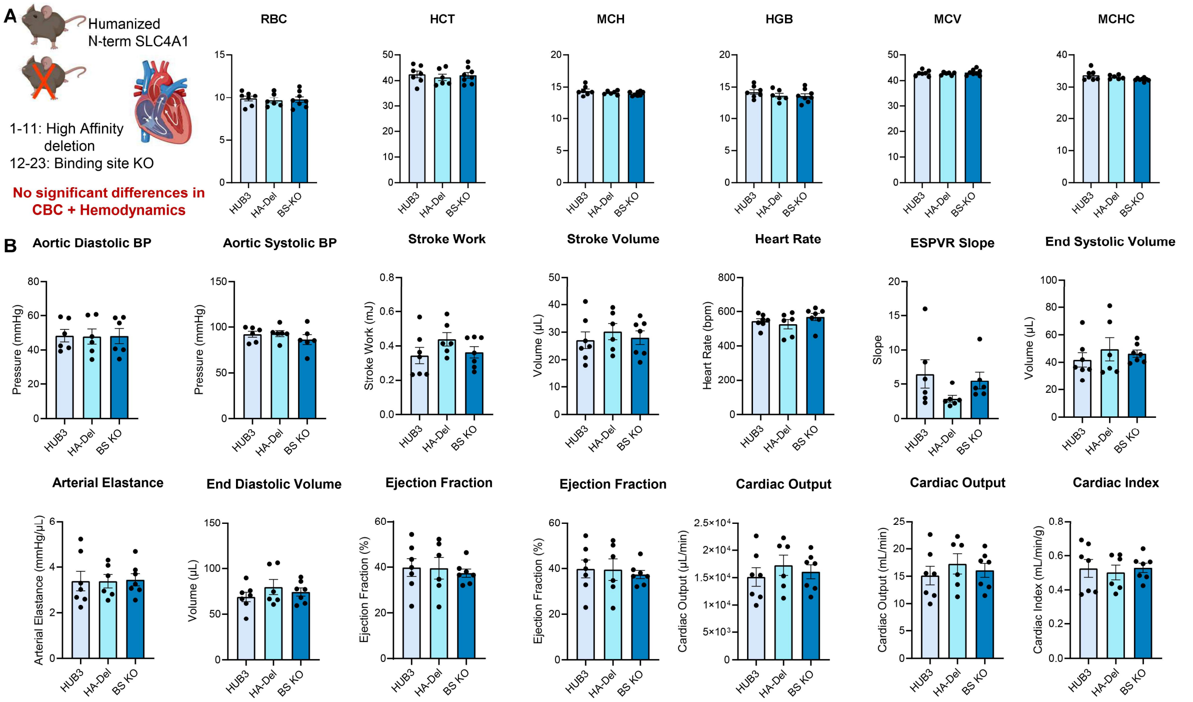

**Supplementary Figure 5. Complete blood counts (A) and hemodynamics results (B) of band 3 N-term KO mice vs humanized control mice.**

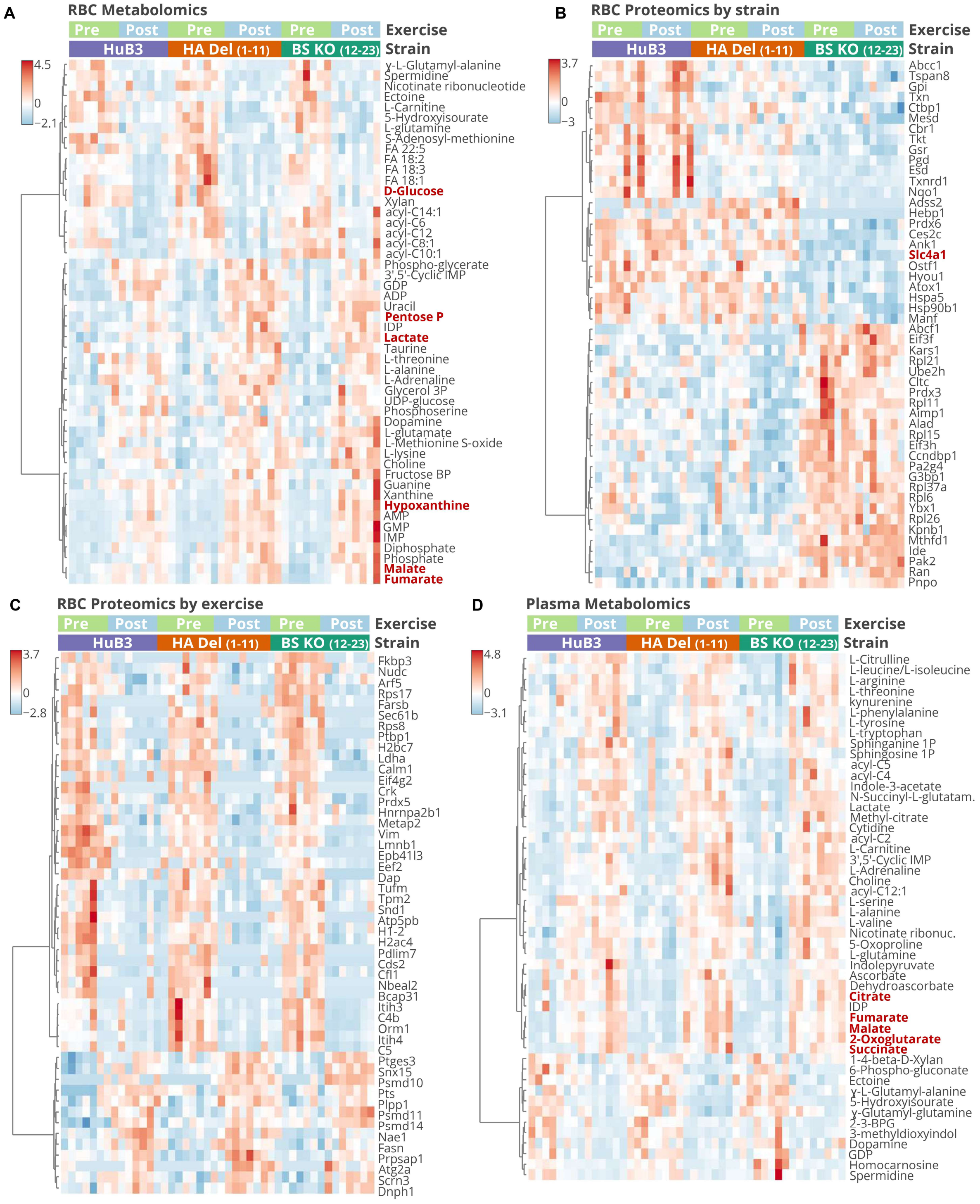

**Supplementary Figure 6. RBC and plasma proteomics and metabolomics after exercise in band 3 KO mice.** Metabolomics **(A)** and proteomics results of RBCs by strain **(B)** or exercise **(C)**, and plasma metabolomics **(D)** as a function of band 3 genotypes (i.e., humanized control, HA Del lacking residues 1-11; BS KO lacking 12-23) before and after critical speed test.

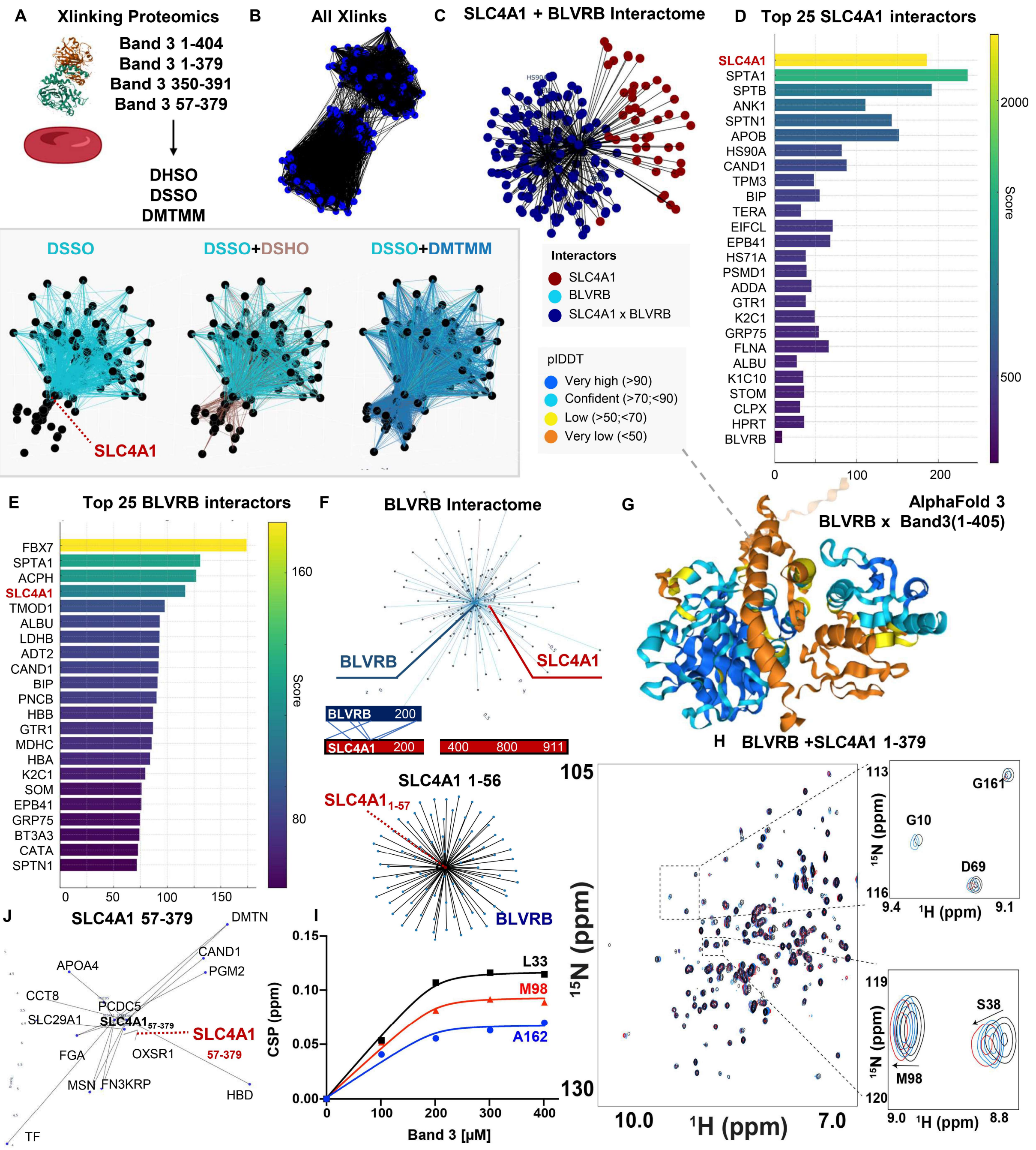

**Supplementary Figure 7. Interactome of the N-terminus of band 3 upon recombinant expression of the cytosolic domain with different lengths.** Recombinant expression of band 3 1-404 (full N-term), 1-379 (before the intermembrane domain), 350-391 (lacking the very N-terminus, but including residues previously reported to bind to glycolytic enzymes via Xlinking proteomics), or 57-379 (lacking the very N-terminus **(A)** allowed in vitro cross-linking proteomics studies with RBC protein extracts. All Xlinks with different agents (DSSO, DSHO, DMTMM) are shown in **(B)**. Focus on the band 3-BLVRB interactome **(C)**. Top 25 band 3 and BLVRB interactors (**D-E**, respectively). Network view of the BLVRB interactome **(F)**. AlphaFold model of the BLVRB x Band 3 interaction **(G).** HSQC from NMR studies of labeled BLVRB incubated with band 3 (1-379 residues **(H)**) and summary CSP for BLVRB residues as a function of band 3 dosage **(I)**. Network view of band 3 57-379 interactors does not have BLVRB **(K)**.

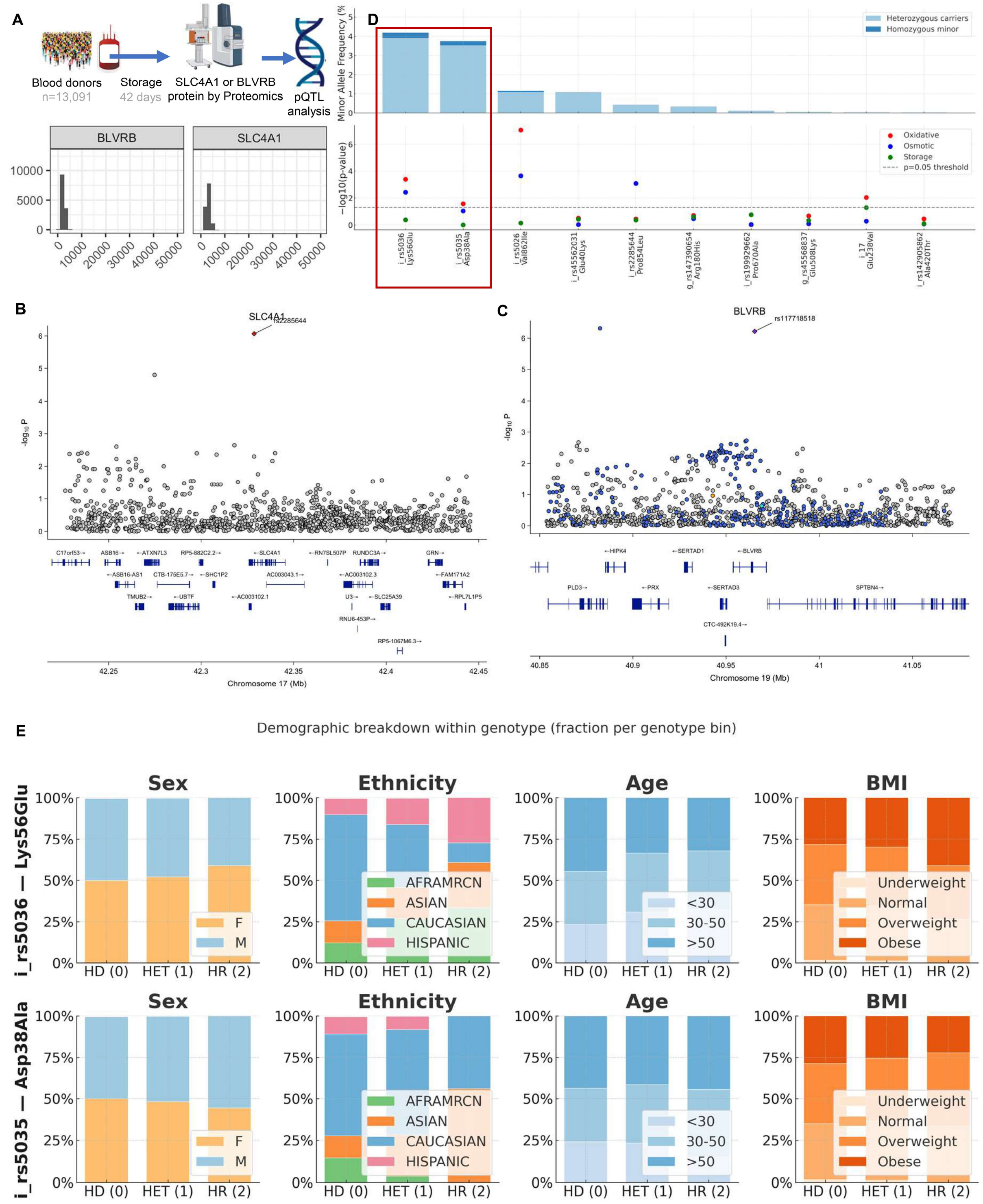

**Supplementary Figure 8. pQTL analysis identifies genetic determinants of SLC4A1 and BLVRB abundance in stored RBCs.** Quantitative proteomics and genetic association analyses identify loci linked to variation in the abundance of Band 3 (SLC4A1) and biliverdin reductase B (BLVRB) in stored red blood cells. A schematic summarizes the workflow for protein quantitative trait locus (pQTL) analysis across 13,091 blood donors, integrating 42-day stored RBC proteomics with 879,000 SNPs **(A)**. Locus zoom plots display genome-wide associations for SLC4A1 and BLVRB protein levels, highlighting significant cis-QTL peaks near the respective genes **(B–C)**. The top panel shows minor allele frequencies and significance levels for leading variants, with oxidative, osmotic, and storage hemolysis correlations indicated by color **(D)**. Demographic breakdowns of donors across genotypes for the two lead variants (rs5036 Lys56Glu and rs5035 Asp38Ala –

both falling in the very N-terminus of band 3 involved in the bulk of protein-protein interactions reported herein) are shown by sex, ethnicity, age, and BMI, illustrating balanced representation across cohorts **(E)**.

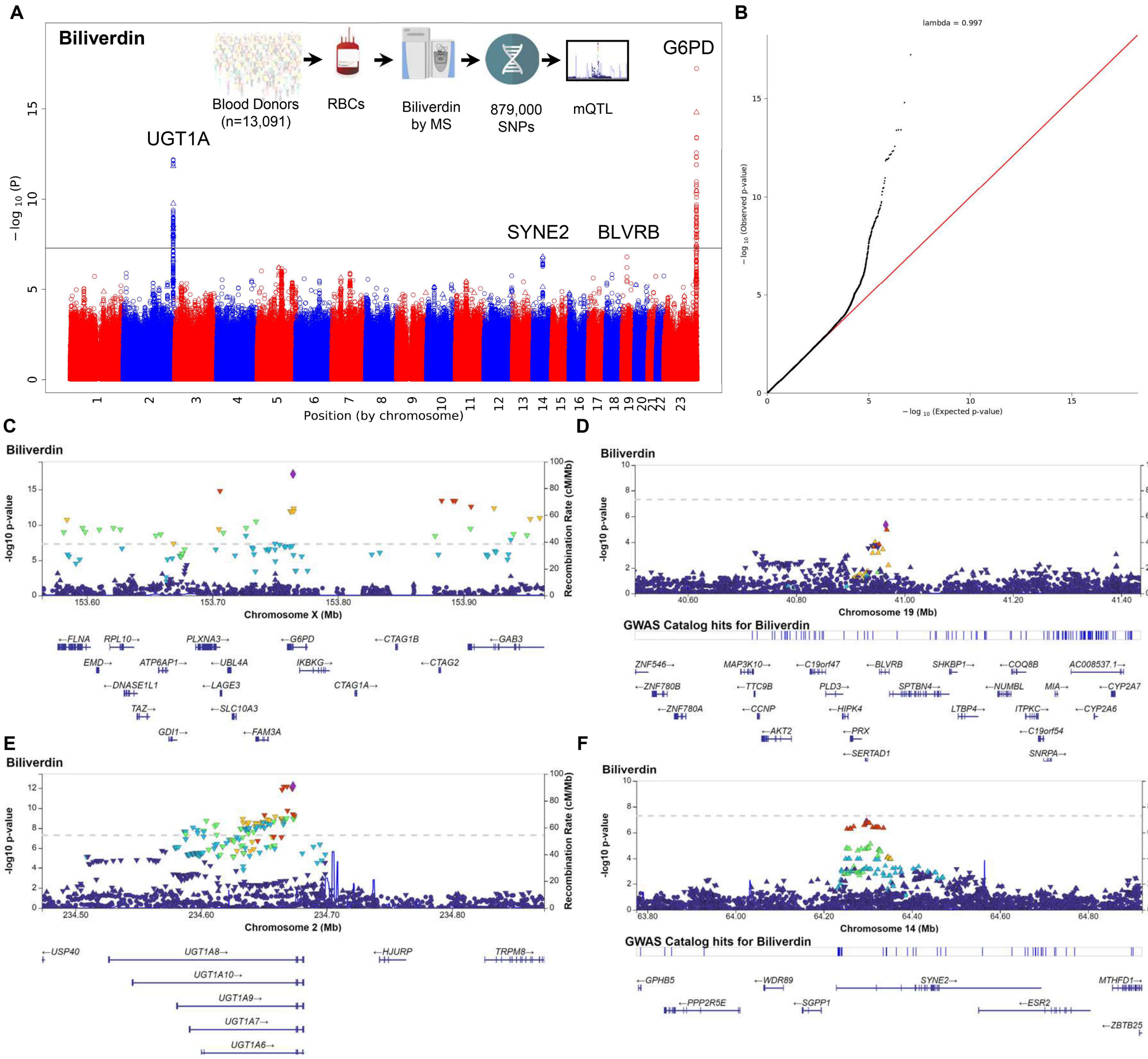

**Supplementary Figure 9. Metabolite Quantitative Trait Loci analysis for BLVRB in 13,091 blood donor volunteers.** mQTL results for BLVRB based on measurements in 13,091 packed RBC units from blood donor volunteers genotyped at 879,000 SNPs **(A)**. Quantile–Quantile (QQ) plot shows the distribution of observed versus expected $-\log_{10}(p)$ values for the mQTL/GWAS analysis of biliverdin levels **(B)**. Locus zoom plots for key genome-wide adjusted significant hits **(C-F).**

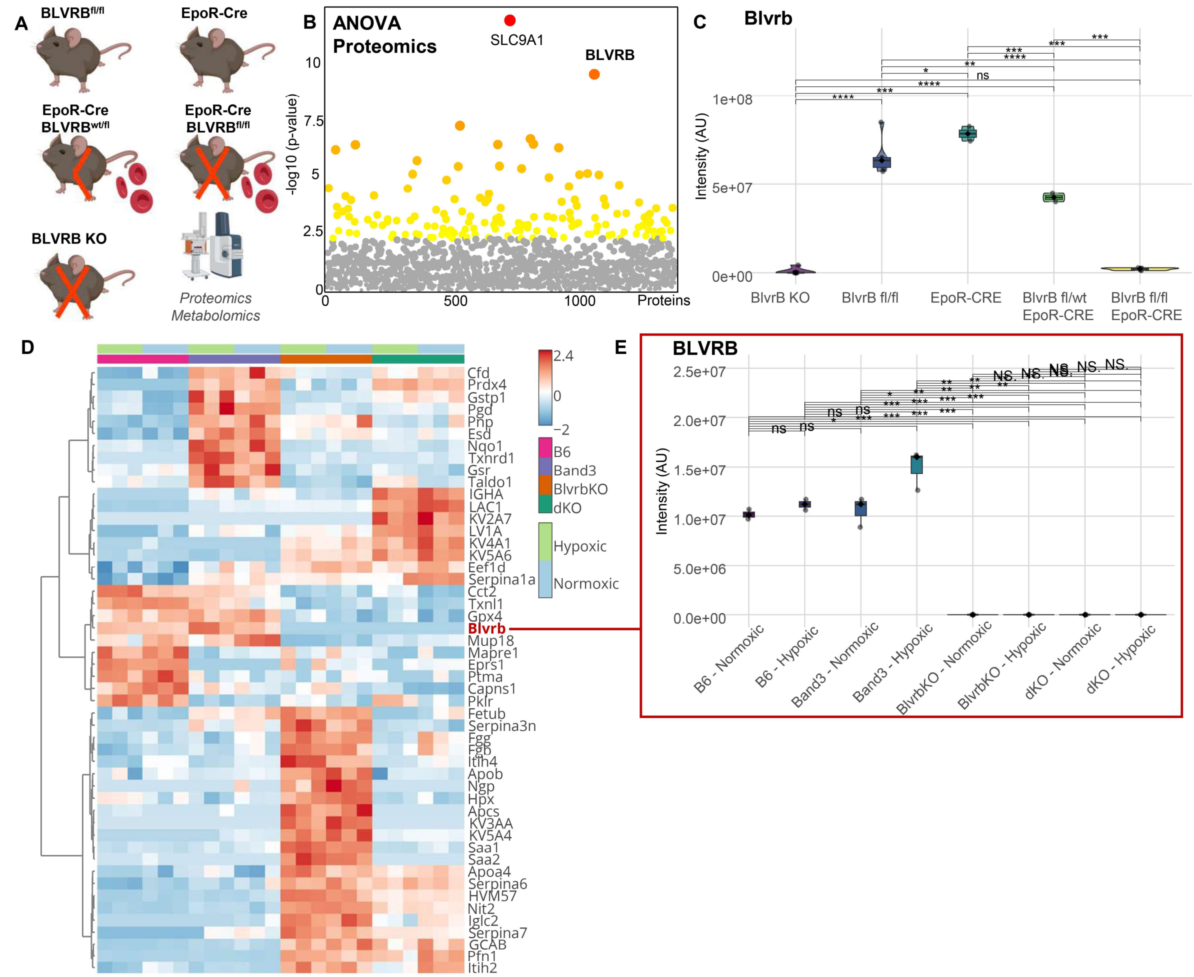

**Supplementary Figure 10. BLVRB deletion and hypoxia modulate the red blood cell proteome.** Genetic ablation of *Blvrb* and hypoxic exposure reveal its role in RBC protein regulation. Schematic of mouse models used for conditional *Blvrb* deletion (EpoR-Cre; Blvrb$^{fl/fl}$) and the global knockout (*Blvrb* KO), followed by proteomic and metabolomic profiling **(A)**. ANOVA of proteomics data highlights *BLVRB* and *SLC9A1* among the most significantly altered proteins **(B)**. Quantification of BLVRB protein intensity confirms complete loss in *Blvrb* KO mice and intermediate expression in erythroid conditional models **(C)**. Heatmap of significantly changing proteins across genotypes and oxygen conditions shows distinct clusters enriched for redox-regulatory enzymes such as GPX4 and PRDX4, as well as hypoxia-responsive factors **(D)**. Boxplots of BLVRB intensity across wild-type, *Band3* KO, *Blvrb* KO, and double-knockout (dKO) RBCs under normoxia and hypoxia demonstrate oxygen-dependent regulation and complete ablation in knockout lines **(E)**.
