## Supplementary Resource - Deep Red for "Deep Red Blood Cell Proteome Defines the Band 3 N-Terminus Interactome as a Regulator of Hypoxic Adaptation via BLVRB-Dependent *S*-Nitroso Transfer": Cysteine Oxidation 2025.html

Cysteine Oxidation 2024


| Official Gene Symbol | Uniprot ID | Protein Description | Position | Peptide start | Official Gene Symbol | Uniprot ID | Protein Description | Assigned Modifications | C (IAA) | C (NEM) | C Oxid | C Dioxid | C-->DHA | C-GS | Total | Cell | sECM | iECM | Peptide |
| --- | --- | --- | --- | --- | --- | --- | --- | --- | --- | --- | --- | --- | --- | --- | --- | --- | --- | --- | --- |
| HBB | HBB\_HUMAN | Hemoglobin subunit beta | 113, 113, 113, 113, 113 | 106 | HBB | HBB\_HUMAN | Hemoglobin subunit beta | 8C(-34.0809) 8C(125.0477) 8C(15.9949) 8C(31.9988) 8C(57.0215) | X | X | X | X | X | NaN | 6.850000e+08 | 6.850000e+08 | NaN | NaN | LLGNVLVCVLAHHFGK |
| HBB | HBB\_HUMAN | Hemoglobin subunit beta | 94, 94, 94, 94 | 84 | HBB | HBB\_HUMAN | Hemoglobin subunit beta | 11C(125.0477) 11C(15.9949) 11C(31.9988) 11C(57.0215) | X | X | X | X | NaN | NaN | 4.039000e+08 | 3.760000e+08 | 2.790000e+07 | NaN | GTFATLSELHCDK |
| HBA1 | HBA\_HUMAN | Hemoglobin subunit alpha | 105, 105, 105, 105, 105, 105 | 101 | HBA1 | HBA\_HUMAN | Hemoglobin subunit alpha | 5C(-34.0809) 5C(125.0477) 5C(15.9949) 5C(305.3076) 5C(31.9988) 5C(57.0215) | X | X | X | X | X | X | 1.310000e+08 | 1.310000e+08 | NaN | NaN | LLSHCLLVTLAAHLPAEFTPAVHASLDK |
| HBE1 | HBE\_HUMAN | Hemoglobin subunit epsilon | 94, 94 | 89 | HBE1 | HBE\_HUMAN | Hemoglobin subunit epsilon | 6C(125.0477) 6C(31.9988) | NaN | X | NaN | X | NaN | NaN | 1.200000e+08 | 1.200000e+08 | NaN | NaN | LSELHCDK |
| GAPDH | G3P\_HUMAN | Glyceraldehyde-3-phosphate dehydrogenase | 247, 247 | 235 | GAPDH | G3P\_HUMAN | Glyceraldehyde-3-phosphate dehydrogenase | 13C(125.0477) 13C(31.9988) | NaN | X | NaN | X | NaN | NaN | 7.990000e+07 | 7.990000e+07 | NaN | NaN | VPTANVSVVDLTCR |
| HBB | HBB\_HUMAN | Hemoglobin subunit beta | 94, 94, 94, 94, 94 | 84 | HBB | HBB\_HUMAN | Hemoglobin subunit beta | 11C(-34.0809) 11C(125.0477) 11C(15.9949) 11C(31.9988) 11C(57.0215) | X | X | X | X | X | NaN | 7.680000e+07 | 7.680000e+07 | NaN | NaN | GTFATLSELHCDKLHVDPENFR |
| SPTB | SPTB1\_HUMAN | Spectrin beta chain, erythrocytic | 735 | 730 | SPTB | SPTB1\_HUMAN | Spectrin beta chain, erythrocytic | 6C(125.0477) | NaN | X | NaN | NaN | NaN | NaN | 6.997951e+07 | NaN | 6.980000e+07 | 179512.3000 | DLAAFCK |
| HBB | HBB\_HUMAN | Hemoglobin subunit beta | 94, 94, 94, 94, 94, 113, 113, 113, 113, 113 | 84 | HBB | HBB\_HUMAN | Hemoglobin subunit beta | 11C(-34.0809) 11C(125.0477) 11C(15.9949) 11C(31.9988) 11C(57.0215) 30C(-34.0809) 30C(125.0477) 30C(15.9949) 30C(31.9988) 30C(57.0215) | X | X | X | X | X | NaN | 6.840000e+07 | 6.840000e+07 | NaN | NaN | GTFATLSELHCDKLHVDPENFRLLGNVLVCVLAHHFGK |
| SLC4A1 | B3AT\_HUMAN | Band 3 anion transport protein | 885, 885, 885, 885 | 880 | SLC4A1 | B3AT\_HUMAN | Band 3 anion transport protein | 6C(125.0477) 6C(15.9949) 6C(31.9988) 6C(57.0215) | X | X | X | X | NaN | NaN | 6.267465e+07 | NaN | 5.370000e+07 | 8974651.0000 | NVELQCLDADDAK |
| HBB | HBB\_HUMAN | Hemoglobin subunit beta | 113, 113, 113, 113 | 106 | HBB | HBB\_HUMAN | Hemoglobin subunit beta | 8C(125.0477) 8C(15.9949) 8C(31.9988) 8C(57.0215) | X | X | X | X | NaN | NaN | 4.580000e+07 | NaN | 4.580000e+07 | NaN | LLGNVLVCVLAHHFGK |
| CAT | CATA\_HUMAN | Catalase | 460, 460, 460, 460 | 459 | CAT | CATA\_HUMAN | Catalase | 2C(-34.0809) 2C(125.0477) 2C(31.9988) 2C(57.0215) | X | X | NaN | X | X | NaN | 4.540000e+07 | 4.540000e+07 | NaN | NaN | LCENIAGHLK |
| BRAF | BRAF\_HUMAN | Serine/threonine-protein kinase B-raf | 173 | 167 | BRAF | BRAF\_HUMAN | Serine/threonine-protein kinase B-raf | 7C(57.0215) | X | NaN | NaN | NaN | NaN | NaN | 4.050000e+07 | 4.050000e+07 | NaN | NaN | TVVPARCGVTVR |
| CAT | CATA\_HUMAN | Catalase | 377, 377, 377, 377 | 366 | CAT | CATA\_HUMAN | Catalase | 12C(125.0477) 12C(15.9949) 12C(31.9988) 12C(57.0215) | X | X | X | X | NaN | NaN | 4.010000e+07 | 4.010000e+07 | NaN | NaN | LGPNYLHIPVNCPYR |
| HBD | HBD\_HUMAN | Hemoglobin subunit delta | 113, 113, 113, 113 | 106 | HBD | HBD\_HUMAN | Hemoglobin subunit delta | 8C(-34.0809) 8C(125.0477) 8C(15.9949) 8C(31.9988) | NaN | X | X | X | X | NaN | 3.180000e+07 | 3.180000e+07 | NaN | NaN | LLGNVLVCVLARNFGKEFTPQMQAAYQK |
| CFL1 | COF1\_HUMAN | Cofilin-1 | 139, 139, 139, 139 | 133 | CFL1 | COF1\_HUMAN | Cofilin-1 | 7C(125.0477) 7C(15.9949) 7C(31.9988) 7C(57.0215) | X | X | X | X | NaN | NaN | 3.150000e+07 | 3.150000e+07 | NaN | NaN | HELQANCYEEVKDR |
| SPTB | SPTB1\_HUMAN | Spectrin beta chain, erythrocytic | 603, 603, 603, 603 | 599 | SPTB | SPTB1\_HUMAN | Spectrin beta chain, erythrocytic | 5C(125.0477) 5C(15.9949) 5C(31.9988) 5C(57.0215) | X | X | X | X | NaN | NaN | 2.864934e+07 | NaN | 2.700000e+07 | 1649340.4000 | GYQPCDPQVIQDR |
| SPTA1 | SPTA1\_HUMAN | Spectrin alpha chain, erythrocytic 1 | 2059, 2059, 2059 | 2052 | SPTA1 | SPTA1\_HUMAN | Spectrin alpha chain, erythrocytic 1 | 8C(125.0477) 8C(15.9949) 8C(31.9988) | NaN | X | X | X | NaN | NaN | 2.740000e+07 | NaN | 2.740000e+07 | NaN | ASALNNWCEK |
| PPIA | PPIA\_HUMAN | Peptidyl-prolyl cis-trans isomerase A | 62, 62, 62 | 56 | PPIA | PPIA\_HUMAN | Peptidyl-prolyl cis-trans isomerase A | 7C(125.0477) 7C(15.9949) 7C(31.9988) | NaN | X | X | X | NaN | NaN | 2.700000e+07 | 2.700000e+07 | NaN | NaN | IIPGFMCQGGDFTR |
| HBD | HBD\_HUMAN | Hemoglobin subunit delta | 94, 94, 94 | 84 | HBD | HBD\_HUMAN | Hemoglobin subunit delta | 11C(125.0477) 11C(31.9988) 11C(57.0215) | X | X | NaN | X | NaN | NaN | 2.490000e+07 | 2.490000e+07 | NaN | NaN | GTFSQLSELHCDK |
| BLVRB | BLVRB\_HUMAN | Flavin reductase (NADPH) | 109, 109, 109 | 106 | BLVRB | BLVRB\_HUMAN | Flavin reductase (NADPH) | 4C(-34.0809) 4C(125.0477) 4C(31.9988) | NaN | X | NaN | X | X | NaN | 2.380000e+07 | 2.380000e+07 | NaN | NaN | VVACTSAFLLWDPTK |
| QDPR | DHPR\_HUMAN | Dihydropteridine reductase | 161, 161 | 155 | QDPR | DHPR\_HUMAN | Dihydropteridine reductase | 7C(125.0477) 7C(31.9988) | NaN | X | NaN | X | NaN | NaN | 2.340000e+07 | 2.340000e+07 | NaN | NaN | GAVHQLCQSLAGK |
| HBB | HBB\_HUMAN | Hemoglobin subunit beta | 113, 113, 113, 113 | 106 | HBB | HBB\_HUMAN | Hemoglobin subunit beta | 8C(125.0477) 8C(15.9949) 8C(31.9988) 8C(57.0215) | X | X | X | X | NaN | NaN | 2.260000e+07 | 2.260000e+07 | NaN | NaN | LLGNVLVCVLAHHFGKEFTPPVQAAYQK |
| SPTA1 | SPTA1\_HUMAN | Spectrin alpha chain, erythrocytic 1 | 965, 965, 965, 965, 965 | 960 | SPTA1 | SPTA1\_HUMAN | Spectrin alpha chain, erythrocytic 1 | 6C(-34.0809) 6C(125.0477) 6C(15.9949) 6C(31.9988) 6C(57.0215) | X | X | X | X | X | NaN | 2.040000e+07 | NaN | 2.040000e+07 | NaN | NQANACQQQQAAPVEGVAGEQR |
| SPTA1 | SPTA1\_HUMAN | Spectrin alpha chain, erythrocytic 1 | 1879, 1879, 1879, 1879, 1879 | 1875 | SPTA1 | SPTA1\_HUMAN | Spectrin alpha chain, erythrocytic 1 | 5C(-34.0809) 5C(125.0477) 5C(15.9949) 5C(31.9988) 5C(57.0215) | X | X | X | X | X | NaN | 2.031588e+07 | NaN | 1.980000e+07 | 515878.0600 | VQNVCAQGEDILNK |
| HYI | HYI\_HUMAN | Putative hydroxypyruvate isomerase | 102 | 99 | HYI | HYI\_HUMAN | Putative hydroxypyruvate isomerase | 4C(15.9949) | NaN | NaN | X | NaN | NaN | NaN | 2.000000e+07 | 2.000000e+07 | NaN | NaN | ALGCPRIHLMAGRVPQGADR |
| ANK1 | ANK1\_HUMAN | Ankyrin-1 | 379, 379 | 368 | ANK1 | ANK1\_HUMAN | Ankyrin-1 | 12C(125.0477) 12C(31.9988) | NaN | X | NaN | X | NaN | NaN | 1.833553e+07 | 1.740000e+07 | 9.355318e+05 | NaN | ALNGFTPLHIACK |
| PGK1 | PGK1\_HUMAN | Phosphoglycerate kinase 1 | 108, 108, 108, 108 | 107 | PGK1 | PGK1\_HUMAN | Phosphoglycerate kinase 1 | 2C(125.0477) 2C(15.9949) 2C(31.9988) 2C(57.0215) | X | X | X | X | NaN | NaN | 1.830000e+07 | 1.830000e+07 | NaN | NaN | ACANPAAGSVILLENLR |
| BROX | BROX\_HUMAN | BRO1 domain-containing protein BROX | 187 | 179 | BROX | BROX\_HUMAN | BRO1 domain-containing protein BROX | 9C(125.0477) | NaN | X | NaN | NaN | NaN | NaN | 1.830000e+07 | 1.830000e+07 | NaN | NaN | LIEAYVIQCQAEAQEVTIAR |
| ALAD | HEM2\_HUMAN | Delta-aminolevulinic acid dehydratase | 75 | 75 | ALAD | HEM2\_HUMAN | Delta-aminolevulinic acid dehydratase | 1C(125.0477) | NaN | X | NaN | NaN | NaN | NaN | 1.770000e+07 | 1.770000e+07 | NaN | NaN | CVLIFGVPSR |
| RNH1 | RINI\_HUMAN | Ribonuclease inhibitor | 30, 30 | 20 | RNH1 | RINI\_HUMAN | Ribonuclease inhibitor | 11C(125.0477) 11C(31.9988) | NaN | X | NaN | X | NaN | NaN | 1.630000e+07 | 1.630000e+07 | NaN | NaN | WAELLPLLQQCQVVR |
| TALDO1 | TALDO\_HUMAN | Transaldolase | 250, 250, 250 | 246 | TALDO1 | TALDO\_HUMAN | Transaldolase | 5C(125.0477) 5C(15.9949) 5C(31.9988) | NaN | X | X | X | NaN | NaN | 1.490000e+07 | 1.490000e+07 | NaN | NaN | ALAGCDFLTISPK |
| ANK1 | ANK1\_HUMAN | Ankyrin-1 | 476, 476, 476, 476 | 467 | ANK1 | ANK1\_HUMAN | Ankyrin-1 | 10C(125.0477) 10C(15.9949) 10C(31.9988) 10C(57.0215) | X | X | X | X | NaN | NaN | 1.420000e+07 | 1.420000e+07 | NaN | NaN | AKDDQTPLHCAAR |
| SPTB | SPTB1\_HUMAN | Spectrin beta chain, erythrocytic | 112, 112, 112, 112 | 110 | SPTB | SPTB1\_HUMAN | Spectrin beta chain, erythrocytic | 3C(125.0477) 3C(15.9949) 3C(31.9988) 3C(57.0215) | X | X | X | X | NaN | NaN | 1.390000e+07 | NaN | 1.390000e+07 | NaN | IHCLENVDK |
| EPB41 | EPB41\_HUMAN | Protein 4.1 | 224, 224, 224, 224 | 214 | EPB41 | EPB41\_HUMAN | Protein 4.1 | 11C(125.0477) 11C(15.9949) 11C(31.9988) 11C(57.0215) | X | X | X | X | NaN | NaN | 1.340000e+07 | NaN | 1.340000e+07 | NaN | VSLLDDTVYECVVEK |
| ANK1 | ANK1\_HUMAN | Ankyrin-1 | 1070, 1070, 1070, 1070 | 1069 | ANK1 | ANK1\_HUMAN | Ankyrin-1 | 2C(125.0477) 2C(15.9949) 2C(31.9988) 2C(57.0215) | X | X | X | X | NaN | NaN | 1.330000e+07 | NaN | 1.330000e+07 | NaN | LCQDYDTIGPEGGSLK |
| PRDX1 | PRDX1\_HUMAN | Peroxiredoxin-1 | 173, 173, 173, 173 | 169 | PRDX1 | PRDX1\_HUMAN | Peroxiredoxin-1 | 5C(125.0477) 5C(15.9949) 5C(31.9988) 5C(57.0215) | X | X | X | X | NaN | NaN | 1.300000e+07 | 1.300000e+07 | NaN | NaN | HGEVCPAGWKPGSDTIKPDVQK |
| HBD | HBD\_HUMAN | Hemoglobin subunit delta | 113, 113, 113 | 106 | HBD | HBD\_HUMAN | Hemoglobin subunit delta | 8C(125.0477) 8C(31.9988) 8C(57.0215) | X | X | NaN | X | NaN | NaN | 1.280000e+07 | 1.280000e+07 | NaN | NaN | LLGNVLVCVLAR |
| GCLM | GSH0\_HUMAN | Glutamate--cysteine ligase regulatory subunit | 72, 72 | 65 | GCLM | GSH0\_HUMAN | Glutamate--cysteine ligase regulatory subunit | 8C(125.0477) 8C(31.9988) | NaN | X | NaN | X | NaN | NaN | 1.250000e+07 | 1.250000e+07 | NaN | NaN | EFPDVLECTVSHAVEK |
| TPI1 | TPIS\_HUMAN | Triosephosphate isomerase | 218, 218 | 207 | TPI1 | TPIS\_HUMAN | Triosephosphate isomerase | 12C(125.0477) 12C(57.0215) | X | X | NaN | NaN | NaN | NaN | 1.200000e+07 | 1.200000e+07 | NaN | NaN | IIYGGSVTGATCK |
| ANK1 | ANK1\_HUMAN | Ankyrin-1 | 1216 | 1211 | ANK1 | ANK1\_HUMAN | Ankyrin-1 | 6C(125.0477) | NaN | X | NaN | NaN | NaN | NaN | 1.162667e+07 | 1.010000e+07 | 1.526673e+06 | NaN | FWLSDCPR |
| HBA1 | HBA\_HUMAN | Hemoglobin subunit alpha | 105, 105 | 101 | HBA1 | HBA\_HUMAN | Hemoglobin subunit alpha | 5C(125.0477) 5C(31.9988) | NaN | X | NaN | X | NaN | NaN | 1.128336e+07 | NaN | 1.050000e+07 | 783356.4400 | LLSHCLLVTLAAHLPAEFTPAVHASLDK |
| TCP1 | TCPA\_HUMAN | T-complex protein 1 subunit alpha | 357 | 356 | TCP1 | TCPA\_HUMAN | T-complex protein 1 subunit alpha | 2C(125.0477) | NaN | X | NaN | NaN | NaN | NaN | 1.123558e+07 | 1.120000e+07 | 3.557813e+04 | NaN | ICDDELILIK |
| HBA1 | HBA\_HUMAN | Hemoglobin subunit alpha | 105, 105, 105, 105, 105, 105 | 101 | HBA1 | HBA\_HUMAN | Hemoglobin subunit alpha | 5C(-34.0809) 5C(125.0477) 5C(15.9949) 5C(305.3076) 5C(31.9988) 5C(57.0215) | X | X | X | X | X | X | 1.060000e+07 | 1.060000e+07 | NaN | NaN | LLSHCLLVTLAAHLPAEFTPAVHASLDKFLASVSTVLTSK |
| SPTA1 | SPTA1\_HUMAN | Spectrin alpha chain, erythrocytic 1 | 773, 773 | 766 | SPTA1 | SPTA1\_HUMAN | Spectrin alpha chain, erythrocytic 1 | 8C(125.0477) 8C(15.9949) | NaN | X | X | NaN | NaN | NaN | 1.050000e+07 | NaN | 1.050000e+07 | NaN | ARQESLVCR |
| EPB42 | EPB42\_HUMAN | Protein 4.2 | 10 | 9 | EPB42 | EPB42\_HUMAN | Protein 4.2 | 2C(125.0477) | NaN | X | NaN | NaN | NaN | NaN | 1.037110e+07 | 9.694284e+06 | 3.252459e+05 | 351565.9000 | SCDFQAAR |
| GSTO1 | GSTO1\_HUMAN | Glutathione S-transferase omega-1 | 192, 192, 192 | 189 | GSTO1 | GSTO1\_HUMAN | Glutathione S-transferase omega-1 | 4C(125.0477) 4C(15.9949) 4C(31.9988) | NaN | X | X | X | NaN | NaN | 9.875288e+06 | 9.875288e+06 | NaN | NaN | LNECVDHTPK |
| HBB | HBB\_HUMAN | Hemoglobin subunit beta | 113, 113, 113, 113, 113 | 97 | HBB | HBB\_HUMAN | Hemoglobin subunit beta | 17C(-34.0809) 17C(125.0477) 17C(15.9949) 17C(31.9988) 17C(57.0215) | X | X | X | X | X | NaN | 9.497031e+06 | 9.497031e+06 | NaN | NaN | LHVDPENFRLLGNVLVCVLAHHFGK |
| ANK1 | ANK1\_HUMAN | Ankyrin-1 | 278, 278, 278 | 269 | ANK1 | ANK1\_HUMAN | Ankyrin-1 | 10C(125.0477) 10C(15.9949) 10C(31.9988) | NaN | X | X | X | NaN | NaN | 9.478578e+06 | NaN | 9.478578e+06 | NaN | TKDELTPLHCAAR |
| CFL1 | COF1\_HUMAN | Cofilin-1 | 147, 147, 139, 139, 139 | 133 | CFL1 | COF1\_HUMAN | Cofilin-1 | 15C(-34.0809) 15C(15.9949) 7C(125.0477) 7C(15.9949) 7C(31.9988) | NaN | X | X | X | X | NaN | 9.320681e+06 | 9.320681e+06 | NaN | NaN | HELQANCYEEVKDRCTLAEK |
| BLVRB | BLVRB\_HUMAN | Flavin reductase (NADPH) | 188, 188, 188, 188 | 188 | BLVRB | BLVRB\_HUMAN | Flavin reductase (NADPH) | 1C(125.0477) 1C(15.9949) 1C(31.9988) 1C(57.0215) | X | X | X | X | NaN | NaN | 9.271423e+06 | 9.271423e+06 | NaN | NaN | CLTTDEYDGHSTYPSHQYQ |
| GAPDH | G3P\_HUMAN | Glyceraldehyde-3-phosphate dehydrogenase | 156, 156, 156, 156, 152, 152, 152, 152 | 146 | GAPDH | G3P\_HUMAN | Glyceraldehyde-3-phosphate dehydrogenase | 11C(-34.0809) 11C(125.0477) 11C(15.9949) 11C(31.9988) 7C(125.0477) 7C(15.9949) 7C(31.9988) 7C(57.0215) | X | X | X | X | X | NaN | 9.153526e+06 | 9.153526e+06 | NaN | NaN | IISNASCTTNCLAPLAK |
| CCT5 | TCPE\_HUMAN | T-complex protein 1 subunit epsilon | 253 | 248 | CCT5 | TCPE\_HUMAN | T-complex protein 1 subunit epsilon | 6C(125.0477) | NaN | X | NaN | NaN | NaN | NaN | 8.857824e+06 | 8.857824e+06 | NaN | NaN | IAILTCPFEPPKPK |
| HBD | HBD\_HUMAN | Hemoglobin subunit delta | 94, 94, 94, 94, 94 | 84 | HBD | HBD\_HUMAN | Hemoglobin subunit delta | 11C(-34.0809) 11C(125.0477) 11C(15.9949) 11C(31.9988) 11C(57.0215) | X | X | X | X | X | NaN | 8.618547e+06 | 8.618547e+06 | NaN | NaN | GTFSQLSELHCDKLHVDPENFR |
| ATIC | PUR9\_HUMAN | Bifunctional purine biosynthesis protein ATIC | 325 | 312 | ATIC | PUR9\_HUMAN | Bifunctional purine biosynthesis protein ATIC | 14C(125.0477) | NaN | X | NaN | NaN | NaN | NaN | 8.414474e+06 | 8.414474e+06 | NaN | NaN | MSSFGDFVALSDVCDVPTAK |
| ATIC | PUR9\_HUMAN | Bifunctional purine biosynthesis protein ATIC | 101 | 98 | ATIC | PUR9\_HUMAN | Bifunctional purine biosynthesis protein ATIC | 4C(125.0477) | NaN | X | NaN | NaN | NaN | NaN | 8.409150e+06 | 8.409150e+06 | NaN | NaN | VVACNLYPFVK |
| PRDX2 | PRDX2\_HUMAN | Peroxiredoxin-2 | 172, 172, 172, 172 | 158 | PRDX2 | PRDX2\_HUMAN | Peroxiredoxin-2 | 15C(125.0477) 15C(15.9949) 15C(31.9988) 15C(57.0215) | X | X | X | X | NaN | NaN | 7.925279e+06 | 7.925279e+06 | NaN | NaN | LVQAFQYTDEHGEVCPAGWKPGSDTIKPNVDDSKEYFSK |
| ANK1 | ANK1\_HUMAN | Ankyrin-1 | 476, 476, 476 | 469 | ANK1 | ANK1\_HUMAN | Ankyrin-1 | 8C(125.0477) 8C(15.9949) 8C(31.9988) | NaN | X | X | X | NaN | NaN | 7.437814e+06 | NaN | 7.437814e+06 | NaN | DDQTPLHCAAR |
| RNH1 | RINI\_HUMAN | Ribonuclease inhibitor | 102, 102, 102, 102, 102, 95, 95, 95, 95, 95, 96, 96, 96 | 90 | RNH1 | RINI\_HUMAN | Ribonuclease inhibitor | 13C(-34.0809) 13C(125.0477) 13C(15.9949) 13C(31.9988) 13C(57.0215) 6C(-34.0809) 6C(125.0477) 6C(15.9949) 6C(31.9988) 6C(57.0215) 7C(-34.0809) 7C(15.9949) 7C(31.9988) | X | X | X | X | X | NaN | 7.217412e+06 | 7.217412e+06 | NaN | NaN | LSLQNCCLTGAGCGVLSSTLR |
| STIP1 | STIP1\_HUMAN | Stress-induced-phosphoprotein 1 | 62 | 57 | STIP1 | STIP1\_HUMAN | Stress-induced-phosphoprotein 1 | 6C(125.0477) | NaN | X | NaN | NaN | NaN | NaN | 6.910304e+06 | 6.845378e+06 | 6.492647e+04 | NaN | AYEDGCK |
| SPTA1 | SPTA1\_HUMAN | Spectrin alpha chain, erythrocytic 1 | 773 | 768 | SPTA1 | SPTA1\_HUMAN | Spectrin alpha chain, erythrocytic 1 | 6C(125.0477) | NaN | X | NaN | NaN | NaN | NaN | 6.775530e+06 | NaN | 6.234012e+06 | 541518.7500 | QESLVCR |
| SPTB | SPTB1\_HUMAN | Spectrin beta chain, erythrocytic | 960, 960, 960, 960, 963, 963, 963 | 956 | SPTB | SPTB1\_HUMAN | Spectrin beta chain, erythrocytic | 5C(125.0477) 5C(15.9949) 5C(31.9988) 5C(57.0215) 8C(125.0477) 8C(15.9949) 8C(31.9988) | X | X | X | X | NaN | NaN | 6.693554e+06 | NaN | 6.693554e+06 | NaN | VHNYCVDCEETSK |
| FBXO7 | FBX7\_HUMAN | F-box only protein 7 | 286, 286 | 277 | FBXO7 | FBX7\_HUMAN | F-box only protein 7 | 10C(125.0477) 10C(57.0215) | X | X | NaN | NaN | NaN | NaN | 6.626614e+06 | 6.626614e+06 | NaN | NaN | LQLLPESFICK |
| TXN | THIO\_HUMAN | Thioredoxin | 73, 73 | 73 | TXN | THIO\_HUMAN | Thioredoxin | 1C(125.0477) 1C(31.9988) | NaN | X | NaN | X | NaN | NaN | 6.615657e+06 | 6.615657e+06 | NaN | NaN | CMPTFQFFK |
| PRDX2 | PRDX2\_HUMAN | Peroxiredoxin-2 | 172, 172 | 158 | PRDX2 | PRDX2\_HUMAN | Peroxiredoxin-2 | 15C(125.0477) 15C(31.9988) | NaN | X | NaN | X | NaN | NaN | 6.379848e+06 | 6.379848e+06 | NaN | NaN | LVQAFQYTDEHGEVCPAGWKPGSDTIKPNVDDSKEYFSKHN |
| HBG2 | HBG2\_HUMAN | Hemoglobin subunit gamma-2 | 94, 94, 94, 94 | 84 | HBG2 | HBG2\_HUMAN | Hemoglobin subunit gamma-2 | 11C(125.0477) 11C(15.9949) 11C(31.9988) 11C(57.0215) | X | X | X | X | NaN | NaN | 6.344998e+06 | 6.344998e+06 | NaN | NaN | GTFAQLSELHCDK |
| PPME1 | PPME1\_HUMAN | Protein phosphatase methylesterase 1 | 312 | 304 | PPME1 | PPME1\_HUMAN | Protein phosphatase methylesterase 1 | 9C(125.0477) | NaN | X | NaN | NaN | NaN | NaN | 6.263162e+06 | 6.263162e+06 | NaN | NaN | GLSNLFLSCPIPK |
| HBA1 | HBA\_HUMAN | Hemoglobin subunit alpha | 105, 105, 105, 105 | 92 | HBA1 | HBA\_HUMAN | Hemoglobin subunit alpha | 14C(-34.0809) 14C(125.0477) 14C(15.9949) 14C(31.9988) | NaN | X | X | X | X | NaN | 6.257328e+06 | 6.257328e+06 | NaN | NaN | LRVDPVNFKLLSHCLLVTLAAHLPAEFTPAVHASLDK |
| DMTN | DEMA\_HUMAN | Dematin | 336 | 331 | DMTN | DEMA\_HUMAN | Dematin | 6C(125.0477) | NaN | X | NaN | NaN | NaN | NaN | 6.237213e+06 | NaN | 6.237213e+06 | NaN | GNSLPCVLEQK |
| PRPS1 | PRPS1\_HUMAN | Ribose-phosphate pyrophosphokinase 1 | 91, 91 | 85 | PRPS1 | PRPS1\_HUMAN | Ribose-phosphate pyrophosphokinase 1 | 7C(125.0477) 7C(31.9988) | NaN | X | NaN | X | NaN | NaN | 6.210483e+06 | 6.210483e+06 | NaN | NaN | VTAVIPCFPYAR |
| PGK1 | PGK1\_HUMAN | Phosphoglycerate kinase 1 | 108, 99 | 98 | PGK1 | PGK1\_HUMAN | Phosphoglycerate kinase 1 | 11C(15.9949) 2C(15.9949) | NaN | NaN | X | NaN | NaN | NaN | 6.107906e+06 | 6.107906e+06 | NaN | NaN | DCVGPEVEKACANPAAGSVILLENLR |
| PRDX6 | PRDX6\_HUMAN | Peroxiredoxin-6 | 47 | 42 | PRDX6 | PRDX6\_HUMAN | Peroxiredoxin-6 | 6C(125.0477) | NaN | X | NaN | NaN | NaN | NaN | 6.068051e+06 | 6.068051e+06 | NaN | NaN | DFTPVCTTELGR |
| SPTA1 | SPTA1\_HUMAN | Spectrin alpha chain, erythrocytic 1 | 2101, 2101, 2101, 2101 | 2101 | SPTA1 | SPTA1\_HUMAN | Spectrin alpha chain, erythrocytic 1 | 1C(125.0477) 1C(15.9949) 1C(31.9988) 1C(57.0215) | X | X | X | X | NaN | NaN | 5.995584e+06 | NaN | 5.995584e+06 | NaN | CLLELDQQIK |
| ADD2 | ADDB\_HUMAN | Beta-adducin | 177, 177 | 174 | ADD2 | ADDB\_HUMAN | Beta-adducin | 4C(125.0477) 4C(31.9988) | NaN | X | NaN | X | NaN | NaN | 5.792580e+06 | NaN | 5.792580e+06 | NaN | GVSCSEVTASSLIK |
| CAPN1 | CAN1\_HUMAN | Calpain-1 catalytic subunit | 49 | 49 | CAPN1 | CAN1\_HUMAN | Calpain-1 catalytic subunit | 1C(125.0477) | NaN | X | NaN | NaN | NaN | NaN | 5.761893e+06 | 5.282242e+06 | 4.652801e+05 | 14371.6550 | CLQSGTLFR |
| HPS1 | HPS1\_HUMAN | Hermansky-Pudlak syndrome 1 protein | 165, 165, 165 | 161 | HPS1 | HPS1\_HUMAN | Hermansky-Pudlak syndrome 1 protein | 5C(15.9949) 5C(31.9988) 5C(57.0215) | X | NaN | X | X | NaN | NaN | 5.592054e+06 | 5.592054e+06 | NaN | NaN | EQEQCFAVEALER |
| GAPDH | G3P\_HUMAN | Glyceraldehyde-3-phosphate dehydrogenase | 247, 247 | 235 | GAPDH | G3P\_HUMAN | Glyceraldehyde-3-phosphate dehydrogenase | 13C(125.0477) 13C(31.9988) | NaN | X | NaN | X | NaN | NaN | 5.538490e+06 | 5.538490e+06 | NaN | NaN | VPTANVSVVDLTCRLEKPAK |
| CARM1 | CARM1\_HUMAN | Histone-arginine methyltransferase CARM1 | 420 | 420 | CARM1 | CARM1\_HUMAN | Histone-arginine methyltransferase CARM1 | 1C(125.0477) | NaN | X | NaN | NaN | NaN | NaN | 5.433890e+06 | 5.433890e+06 | NaN | NaN | CLFQSPLFAK |
| SLC4A1 | B3AT\_HUMAN | Band 3 anion transport protein | 201 | 181 | SLC4A1 | B3AT\_HUMAN | Band 3 anion transport protein | 21C(125.0477) | NaN | X | NaN | NaN | NaN | NaN | 5.428244e+06 | 5.411606e+06 | 1.663725e+04 | NaN | SGDPSQPLLPQHSSLETQLFCEQGDGGTEGHSPSGILEK |
| DDI2 | DDI2\_HUMAN | Protein DDI1 homolog 2 | 361 | 339 | DDI2 | DDI2\_HUMAN | Protein DDI1 homolog 2 | 23C(125.0477) | NaN | X | NaN | NaN | NaN | NaN | 5.410214e+06 | 5.410214e+06 | 0.000000e+00 | NaN | NVLVIGTTGSQTTFLPEGELPECAR |
| UBE2L3 | UB2L3\_HUMAN | Ubiquitin-conjugating enzyme E2 L3 | 86 | 83 | UBE2L3 | UB2L3\_HUMAN | Ubiquitin-conjugating enzyme E2 L3 | 4C(125.0477) | NaN | X | NaN | NaN | NaN | NaN | 5.278530e+06 | 5.278530e+06 | NaN | NaN | GQVCLPVISAENWKPATK |
| ANK1 | ANK1\_HUMAN | Ankyrin-1 | 864, 864, 864 | 862 | ANK1 | ANK1\_HUMAN | Ankyrin-1 | 3C(125.0477) 3C(15.9949) 3C(31.9988) | NaN | X | X | X | NaN | NaN | 5.269361e+06 | NaN | 5.269361e+06 | NaN | IPCAMPETVVIR |
| CLIC1 | CLIC1\_HUMAN | Chloride intracellular channel protein 1 | 178 | 167 | CLIC1 | CLIC1\_HUMAN | Chloride intracellular channel protein 1 | 12C(125.0477) | NaN | X | NaN | NaN | NaN | NaN | 5.229126e+06 | 5.229126e+06 | NaN | NaN | FLDGNELTLADCNLLPK |
| PSMD9 | PSMD9\_HUMAN | 26S proteasome non-ATPase regulatory subunit 9 | 216 | 212 | PSMD9 | PSMD9\_HUMAN | 26S proteasome non-ATPase regulatory subunit 9 | 5C(125.0477) | NaN | X | NaN | NaN | NaN | NaN | 5.064656e+06 | 5.064656e+06 | NaN | NaN | GLLGCNIIPLQR |
| ESD | ESTD\_HUMAN | S-formylglutathione hydrolase | 28, 28 | 18 | ESD | ESTD\_HUMAN | S-formylglutathione hydrolase | 11C(125.0477) 11C(31.9988) | NaN | X | NaN | X | NaN | NaN | 5.005562e+06 | 5.005562e+06 | NaN | NaN | VFEHDSVELNCK |
| PNPLA2 | PLPL2\_HUMAN | Patatin-like phospholipase domain-containing protein 2 | 28, 15 | 7 | PNPLA2 | PLPL2\_HUMAN | Patatin-like phospholipase domain-containing protein 2 | 22C(31.9988) 9C(57.0215) | X | NaN | NaN | X | NaN | NaN | 4.956949e+06 | 4.956949e+06 | NaN | NaN | TWNISFAGCGFLGVYYVGVASCLR |
| ANK1 | ANK1\_HUMAN | Ankyrin-1 | 476, 476, 476 | 467 | ANK1 | ANK1\_HUMAN | Ankyrin-1 | 10C(125.0477) 10C(15.9949) 10C(31.9988) | NaN | X | X | X | NaN | NaN | 4.945334e+06 | NaN | 4.945334e+06 | NaN | AKDDQTPLHCAAR |
| HBQ1 | HBAT\_HUMAN | Hemoglobin subunit theta-1 | 90, 90 | 74 | HBQ1 | HBAT\_HUMAN | Hemoglobin subunit theta-1 | 17C(125.0477) 17C(15.9949) | NaN | X | X | NaN | NaN | NaN | 4.812272e+06 | 4.812272e+06 | NaN | NaN | LDDLPHALSALSHLHACQLR |
| GPS1 | CSN1\_HUMAN | COP9 signalosome complex subunit 1 | 67 | 60 | GPS1 | CSN1\_HUMAN | COP9 signalosome complex subunit 1 | 8C(125.0477) | NaN | X | NaN | NaN | NaN | NaN | 4.554084e+06 | 4.554084e+06 | NaN | NaN | LQFIADHCPTLR |
| TPI1 | TPIS\_HUMAN | Triosephosphate isomerase | 127 | 114 | TPI1 | TPIS\_HUMAN | Triosephosphate isomerase | 14C(125.0477) | NaN | X | NaN | NaN | NaN | NaN | 4.499599e+06 | 4.499599e+06 | NaN | NaN | VAHALAEGLGVIACIGEK |
| EPB42 | EPB42\_HUMAN | Protein 4.2 | 276 | 260 | EPB42 | EPB42\_HUMAN | Protein 4.2 | 17C(125.0477) | NaN | X | NaN | NaN | NaN | NaN | 4.429767e+06 | 4.426536e+06 | 3.230989e+03 | NaN | GRPVYDGQAWVLAAVACTVLR |
| SPTA1 | SPTA1\_HUMAN | Spectrin alpha chain, erythrocytic 1 | 224 | 214 | SPTA1 | SPTA1\_HUMAN | Spectrin alpha chain, erythrocytic 1 | 11C(125.0477) | NaN | X | NaN | NaN | NaN | NaN | 4.404008e+06 | 4.404008e+06 | NaN | NaN | VVEVNQYANECAEENHPDLPLIQSK |
| ANK1 | ANK1\_HUMAN | Ankyrin-1 | 943 | 942 | ANK1 | ANK1\_HUMAN | Ankyrin-1 | 2C(125.0477) | NaN | X | NaN | NaN | NaN | NaN | 4.355486e+06 | 1.417739e+05 | 4.213712e+06 | NaN | TCAAPTR |
| HBA1 | HBA\_HUMAN | Hemoglobin subunit alpha | 105, 105, 105, 105, 105 | 94 | HBA1 | HBA\_HUMAN | Hemoglobin subunit alpha | 12C(-34.0809) 12C(125.0477) 12C(15.9949) 12C(31.9988) 12C(57.0215) | X | X | X | X | X | NaN | 4.351577e+06 | 4.351577e+06 | NaN | NaN | VDPVNFKLLSHCLLVTLAAHLPAEFTPAVHASLDK |
| PRDX1 | PRDX1\_HUMAN | Peroxiredoxin-1 | 83, 83, 71, 71 | 69 | PRDX1 | PRDX1\_HUMAN | Peroxiredoxin-1 | 15C(125.0477) 15C(15.9949) 3C(125.0477) 3C(15.9949) | NaN | X | X | NaN | NaN | NaN | 4.331699e+06 | 4.331699e+06 | NaN | NaN | LNCQVIGASVDSHFCHLAWVNTPK |
| PGD | 6PGD\_HUMAN | 6-phosphogluconate dehydrogenase, decarboxylating | 289 | 289 | PGD | 6PGD\_HUMAN | 6-phosphogluconate dehydrogenase, decarboxylating | 1C(125.0477) | NaN | X | NaN | NaN | NaN | NaN | 4.210028e+06 | 4.092720e+06 | 1.173081e+05 | NaN | CLSSLKDER |
| HINT1 | HINT1\_HUMAN | Adenosine 5'-monophosphoramidase HINT1 | 38 | 38 | HINT1 | HINT1\_HUMAN | Adenosine 5'-monophosphoramidase HINT1 | 1C(125.0477) | NaN | X | NaN | NaN | NaN | NaN | 4.204200e+06 | 4.204200e+06 | NaN | NaN | CLAFHDISPQAPTHFLVIPK |
| RPIA | RPIA\_HUMAN | Ribose-5-phosphate isomerase | 128 | 121 | RPIA | RPIA\_HUMAN | Ribose-5-phosphate isomerase | 8C(125.0477) | NaN | X | NaN | NaN | NaN | NaN | 4.166238e+06 | 4.166238e+06 | NaN | NaN | QENLNLVCIPTSFQAR |
| HBG2 | HBG2\_HUMAN | Hemoglobin subunit gamma-2 | 94, 94 | 78 | HBG2 | HBG2\_HUMAN | Hemoglobin subunit gamma-2 | 17C(125.0477) 17C(15.9949) | NaN | X | X | NaN | NaN | NaN | 4.153878e+06 | 4.153878e+06 | NaN | NaN | HLDDLKGTFAQLSELHCDK |
| PNP | PNPH\_HUMAN | Purine nucleoside phosphorylase | 31, 31 | 23 | PNP | PNPH\_HUMAN | Purine nucleoside phosphorylase | 9C(125.0477) 9C(31.9988) | NaN | X | NaN | X | NaN | NaN | 4.140222e+06 | 4.140222e+06 | NaN | NaN | HRPQVAIICGSGLGGLTDK |
| CCT2 | TCPB\_HUMAN | T-complex protein 1 subunit beta | 395, 395 | 389 | CCT2 | TCPB\_HUMAN | T-complex protein 1 subunit beta | 7C(125.0477) 7C(31.9988) | NaN | X | NaN | X | NaN | NaN | 4.073701e+06 | 4.073701e+06 | NaN | NaN | SLHDALCVLAQTVK |
| GCLC | GSH1\_HUMAN | Glutamate--cysteine ligase catalytic subunit | 152, 152 | 150 | GCLC | GSH1\_HUMAN | Glutamate--cysteine ligase catalytic subunit | 3C(125.0477) 3C(57.0215) | X | X | NaN | NaN | NaN | NaN | 4.029920e+06 | 4.029920e+06 | NaN | NaN | LGCPGFTLPEVKPNPVEGGASK |
| TCP1 | TCPA\_HUMAN | T-complex protein 1 subunit alpha | 236 | 234 | TCP1 | TCPA\_HUMAN | T-complex protein 1 subunit alpha | 3C(15.9949) | NaN | NaN | X | NaN | NaN | NaN | 4.025308e+06 | NaN | 1.674457e+06 | 2350851.0000 | IACLDFSLQK |
| FBXO7 | FBX7\_HUMAN | F-box only protein 7 | 286 | 276 | FBXO7 | FBX7\_HUMAN | F-box only protein 7 | 11C(125.0477) | NaN | X | NaN | NaN | NaN | NaN | 4.020314e+06 | 4.020314e+06 | NaN | NaN | RLQLLPESFICK |
| HBB | HBB\_HUMAN | Hemoglobin subunit beta | 94, 94, 94 | 84 | HBB | HBB\_HUMAN | Hemoglobin subunit beta | 11C(125.0477) 11C(31.9988) 11C(57.0215) | X | X | NaN | X | NaN | NaN | 3.978702e+06 | NaN | NaN | 3978701.5000 | GTFATLSELHCDK |
| GOT1 | AATC\_HUMAN | Aspartate aminotransferase, cytoplasmic | 46 | 43 | GOT1 | AATC\_HUMAN | Aspartate aminotransferase, cytoplasmic | 4C(125.0477) | NaN | X | NaN | NaN | NaN | NaN | 3.966680e+06 | 3.966680e+06 | NaN | NaN | TDDCHPWVLPVVK |
| CAT | CATA\_HUMAN | Catalase | 232, 232 | 222 | CAT | CATA\_HUMAN | Catalase | 11C(125.0477) 11C(31.9988) | NaN | X | NaN | X | NaN | NaN | 3.926064e+06 | 3.926064e+06 | NaN | NaN | LVNANGEAVYCK |
| PRPS1 | PRPS1\_HUMAN | Ribose-phosphate pyrophosphokinase 1 | 41, 41 | 35 | PRPS1 | PRPS1\_HUMAN | Ribose-phosphate pyrophosphokinase 1 | 7C(125.0477) 7C(31.9988) | NaN | X | NaN | X | NaN | NaN | 3.888244e+06 | 3.888244e+06 | NaN | NaN | FSNQETCVEIGESVR |
| ANXA7 | ANXA7\_HUMAN | Annexin A7 | 285 | 279 | ANXA7 | ANXA7\_HUMAN | Annexin A7 | 7C(125.0477) | NaN | X | NaN | NaN | NaN | NaN | 3.787050e+06 | 3.787050e+06 | NaN | NaN | VLIEILCTR |
| GDI2 | GDIB\_HUMAN | Rab GDP dissociation inhibitor beta | 335, 335 | 330 | GDI2 | GDIB\_HUMAN | Rab GDP dissociation inhibitor beta | 6C(125.0477) 6C(31.9988) | NaN | X | NaN | X | NaN | NaN | 3.757884e+06 | 3.757884e+06 | NaN | NaN | SDIYVCMISFAHNVAAQGK |
| EPB42 | EPB42\_HUMAN | Protein 4.2 | 428, 428 | 422 | EPB42 | EPB42\_HUMAN | Protein 4.2 | 7C(125.0477) 7C(31.9988) | NaN | X | NaN | X | NaN | NaN | 3.660909e+06 | 3.660909e+06 | NaN | NaN | GVGSDRCEDITQNYK |
| PTPA | PTPA\_HUMAN | Serine/threonine-protein phosphatase 2A activator | 334 | 332 | PTPA | PTPA\_HUMAN | Serine/threonine-protein phosphatase 2A activator | 3C(125.0477) | NaN | X | NaN | NaN | NaN | NaN | 3.621177e+06 | 3.621177e+06 | NaN | NaN | AECLEKFPVIQHFK |
| KPNA4 | IMA3\_HUMAN | Importin subunit alpha-3 | 122, 122 | 113 | KPNA4 | IMA3\_HUMAN | Importin subunit alpha-3 | 10C(125.0477) 10C(31.9988) | NaN | X | NaN | X | NaN | NaN | 3.612845e+06 | 3.612845e+06 | NaN | NaN | SGILPILVHCLER |
| HBB | HBB\_HUMAN | Hemoglobin subunit beta | 113, 113 | 106 | HBB | HBB\_HUMAN | Hemoglobin subunit beta | 8C(125.0477) 8C(31.9988) | NaN | X | NaN | X | NaN | NaN | 3.612563e+06 | NaN | NaN | 3612562.8000 | LLGNVLVCVLAHHFGK |
| TGM2 | TGM2\_HUMAN | Protein-glutamine gamma-glutamyltransferase 2 | 27, 27 | 20 | TGM2 | TGM2\_HUMAN | Protein-glutamine gamma-glutamyltransferase 2 | 8C(125.0477) 8C(15.9949) | NaN | X | X | NaN | NaN | NaN | 3.605832e+06 | 3.605832e+06 | NaN | NaN | DHHTADLCR |
| ADD2 | ADDB\_HUMAN | Beta-adducin | 224 | 224 | ADD2 | ADDB\_HUMAN | Beta-adducin | 1C(125.0477) | NaN | X | NaN | NaN | NaN | NaN | 3.593341e+06 | 3.586012e+06 | 7.329385e+03 | NaN | CIIHLHTPATAAVSAMK |
| HBE1 | HBE\_HUMAN | Hemoglobin subunit epsilon | 94 | 89 | HBE1 | HBE\_HUMAN | Hemoglobin subunit epsilon | 6C(125.0477) | NaN | X | NaN | NaN | NaN | NaN | 3.574960e+06 | NaN | 3.524486e+06 | 50473.7700 | LSELHCDK |
| CFL1 | COF1\_HUMAN | Cofilin-1 | 39 | 34 | CFL1 | COF1\_HUMAN | Cofilin-1 | 6C(125.0477) | NaN | X | NaN | NaN | NaN | NaN | 3.554185e+06 | 3.554185e+06 | NaN | NaN | KAVLFCLSEDKK |
| RNF123 | RN123\_HUMAN | E3 ubiquitin-protein ligase RNF123 | 1180 | 1169 | RNF123 | RN123\_HUMAN | E3 ubiquitin-protein ligase RNF123 | 12C(125.0477) | NaN | X | NaN | NaN | NaN | NaN | 3.496444e+06 | 3.496444e+06 | NaN | NaN | EQATSVLLADPCFQLR |
| ALDOA | ALDOA\_HUMAN | Fructose-bisphosphate aldolase A | 178, 178 | 174 | ALDOA | ALDOA\_HUMAN | Fructose-bisphosphate aldolase A | 5C(125.0477) 5C(31.9988) | NaN | X | NaN | X | NaN | NaN | 3.467094e+06 | 3.467094e+06 | NaN | NaN | YASICQQNGIVPIVEPEILPDGDHDLK |
| HBD | HBD\_HUMAN | Hemoglobin subunit delta | 94, 94 | 84 | HBD | HBD\_HUMAN | Hemoglobin subunit delta | 11C(125.0477) 11C(31.9988) | NaN | X | NaN | X | NaN | NaN | 3.454615e+06 | NaN | 3.454615e+06 | NaN | GTFSQLSELHCDK |
| HBB | HBB\_HUMAN | Hemoglobin subunit beta | 94, 94 | 84 | HBB | HBB\_HUMAN | Hemoglobin subunit beta | 11C(125.0477) 11C(31.9988) | NaN | X | NaN | X | NaN | NaN | 3.423444e+06 | NaN | 3.423444e+06 | NaN | GTFATLSELHCDKLHVDPENFR |
| SEC14L4 | S14L4\_HUMAN | SEC14-like protein 4 | 301 | 287 | SEC14L4 | S14L4\_HUMAN | SEC14-like protein 4 | 15C(125.0477) | NaN | X | NaN | NaN | NaN | NaN | 3.408805e+06 | 3.408805e+06 | NaN | NaN | GSSLQVENEILFPGCVLR |
| COPS4 | CSN4\_HUMAN | COP9 signalosome complex subunit 4 | 70 | 64 | COPS4 | CSN4\_HUMAN | COP9 signalosome complex subunit 4 | 7C(125.0477) | NaN | X | NaN | NaN | NaN | NaN | 3.389796e+06 | 3.389796e+06 | NaN | NaN | QLLTDFCTHLPNLPDSTAK |
| GPX1 | GPX1\_HUMAN | Glutathione peroxidase 1 | 78 | 70 | GPX1 | GPX1\_HUMAN | Glutathione peroxidase 1 | 9C(125.0477) | NaN | X | NaN | NaN | NaN | NaN | 3.383416e+06 | 3.383416e+06 | NaN | NaN | GLVVLGFPCNQFGHQENAK |
| TFPI | TFPI1\_HUMAN | Tissue factor pathway inhibitor | 63 | 58 | TFPI | TFPI1\_HUMAN | Tissue factor pathway inhibitor | 6C(125.0477) | NaN | X | NaN | NaN | NaN | NaN | 3.381008e+06 | NaN | 3.381008e+06 | NaN | ADDGPCK |
| ATP1A1 | AT1A1\_HUMAN | Sodium/potassium-transporting ATPase subunit alpha-1 | 606 | 597 | ATP1A1 | AT1A1\_HUMAN | Sodium/potassium-transporting ATPase subunit alpha-1 | 10C(57.0215) | X | NaN | NaN | NaN | NaN | NaN | 3.369020e+06 | NaN | 3.369020e+06 | NaN | AAVPDAVGKCRSAGIK |
| CTPS2 | PYRG2\_HUMAN | CTP synthase 2 | 216 | 206 | CTPS2 | PYRG2\_HUMAN | CTP synthase 2 | 11C(125.0477) | NaN | X | NaN | NaN | NaN | NaN | 3.336743e+06 | 3.313311e+06 | 2.343130e+04 | NaN | GLGLSPDLIVCR |
| APEH | ACPH\_HUMAN | Acylamino-acid-releasing enzyme | 292 | 292 | APEH | ACPH\_HUMAN | Acylamino-acid-releasing enzyme | 1C(125.0477) | NaN | X | NaN | NaN | NaN | NaN | 3.336100e+06 | 3.316907e+06 | 1.919340e+04 | NaN | CELLSDDSLAVSSPR |
| GPX1 | GPX1\_HUMAN | Glutathione peroxidase 1 | 115 | 115 | GPX1 | GPX1\_HUMAN | Glutathione peroxidase 1 | 1C(125.0477) | NaN | X | NaN | NaN | NaN | NaN | 3.322036e+06 | 3.322036e+06 | NaN | NaN | CEVNGAGAHPLFAFLR |
| TKTL2 | TKTL2\_HUMAN | Transketolase-like protein 2 | 577, 577 | 569 | TKTL2 | TKTL2\_HUMAN | Transketolase-like protein 2 | 9C(15.9949) 9C(31.9988) | NaN | NaN | X | X | NaN | NaN | 3.301927e+06 | NaN | NaN | 3301927.0000 | EGGIGEAVCAAVSR |
| PEPD | PEPD\_HUMAN | Xaa-Pro dipeptidase | 58 | 37 | PEPD | PEPD\_HUMAN | Xaa-Pro dipeptidase | 22C(57.0215) | X | NaN | NaN | NaN | NaN | NaN | 3.287179e+06 | 3.287179e+06 | NaN | NaN | NPAVQAGSIVVLQGGEETQRYCTDTGVLFR |
| TPI1 | TPIS\_HUMAN | Triosephosphate isomerase | 87, 87, 87 | 86 | TPI1 | TPIS\_HUMAN | Triosephosphate isomerase | 2C(125.0477) 2C(31.9988) 2C(57.0215) | X | X | NaN | X | NaN | NaN | 3.269196e+06 | 3.269196e+06 | NaN | NaN | DCGATWVVLGHSER |
| EPB42 | EPB42\_HUMAN | Protein 4.2 | 281 | 281 | EPB42 | EPB42\_HUMAN | Protein 4.2 | 1C(125.0477) | NaN | X | NaN | NaN | NaN | NaN | 3.261232e+06 | NaN | 3.250046e+06 | 11186.8020 | CLGIPAR |
| GPI | G6PI\_HUMAN | Glucose-6-phosphate isomerase | 404 | 401 | GPI | G6PI\_HUMAN | Glucose-6-phosphate isomerase | 4C(125.0477) | NaN | X | NaN | NaN | NaN | NaN | 3.227065e+06 | 3.227065e+06 | NaN | NaN | MIPCDFLIPVQTQHPIR |
| FERMT3 | URP2\_HUMAN | Fermitin family homolog 3 | 439 | 439 | FERMT3 | URP2\_HUMAN | Fermitin family homolog 3 | 1C(125.0477) | NaN | X | NaN | NaN | NaN | NaN | 3.197567e+06 | 3.119353e+06 | 7.821466e+04 | NaN | CQDEQQYAR |
| MDH1 | MDHC\_HUMAN | Malate dehydrogenase, cytoplasmic | 137, 137, 137 | 126 | MDH1 | MDHC\_HUMAN | Malate dehydrogenase, cytoplasmic | 12C(125.0477) 12C(31.9988) 12C(57.0215) | X | X | NaN | X | NaN | NaN | 3.197152e+06 | 3.197152e+06 | NaN | NaN | VIVVGNPANTNCLTASK |
| RNH1 | RINI\_HUMAN | Ribonuclease inhibitor | 38 | 35 | RNH1 | RINI\_HUMAN | Ribonuclease inhibitor | 4C(125.0477) | NaN | X | NaN | NaN | NaN | NaN | 3.189470e+06 | 3.189470e+06 | NaN | NaN | LDDCGLTEAR |
| HBB | HBB\_HUMAN | Hemoglobin subunit beta | 113, 113 | 97 | HBB | HBB\_HUMAN | Hemoglobin subunit beta | 17C(125.0477) 17C(31.9988) | NaN | X | NaN | X | NaN | NaN | 3.187362e+06 | 3.187362e+06 | NaN | NaN | LHVDPENFRLLGNVLVCVLAHHFGKEFTPPVQAAYQK |
| CFH | CFAH\_HUMAN | Complement factor H | 901 | 897 | CFH | CFAH\_HUMAN | Complement factor H | 5C(31.9988) | NaN | NaN | NaN | X | NaN | NaN | 3.088210e+06 | 8.951769e+03 | 9.876228e+05 | 2091635.4000 | LSYTCEGGFR |
| ANK1 | ANK1\_HUMAN | Ankyrin-1 | 1200 | 1194 | ANK1 | ANK1\_HUMAN | Ankyrin-1 | 7C(125.0477) | NaN | X | NaN | NaN | NaN | NaN | 3.079511e+06 | 2.231199e+06 | 8.483122e+05 | NaN | LVYANECANFTTNVSAR |
| C20orf27 | CT027\_HUMAN | UPF0687 protein C20orf27 | 106 | 106 | C20orf27 | CT027\_HUMAN | UPF0687 protein C20orf27 | 1C(125.0477) | NaN | X | NaN | NaN | NaN | NaN | 3.045702e+06 | 3.045702e+06 | NaN | NaN | CEYSAHK |
| EPB41 | EPB41\_HUMAN | Protein 4.1 | 179, 179 | 172 | EPB41 | EPB41\_HUMAN | Protein 4.1 | 8C(125.0477) 8C(31.9988) | NaN | X | NaN | X | NaN | NaN | 3.016092e+06 | NaN | 3.016092e+06 | NaN | EGEGLEECSK |
| RAB8A | RAB8A\_HUMAN | Ras-related protein Rab-8A | 123 | 123 | RAB8A | RAB8A\_HUMAN | Ras-related protein Rab-8A | 1C(125.0477) | NaN | X | NaN | NaN | NaN | NaN | 3.010725e+06 | 3.010725e+06 | NaN | NaN | CDVNDKR |
| TKFC | TKFC\_HUMAN | Triokinase/FMN cyclase | 404 | 398 | TKFC | TKFC\_HUMAN | Triokinase/FMN cyclase | 7C(125.0477) | NaN | X | NaN | NaN | NaN | NaN | 2.975428e+06 | 2.945338e+06 | 3.009030e+04 | NaN | AAGDGDCGTTHSR |
| CCT7 | TCPH\_HUMAN | T-complex protein 1 subunit eta | 370 | 369 | CCT7 | TCPH\_HUMAN | T-complex protein 1 subunit eta | 2C(125.0477) | NaN | X | NaN | NaN | NaN | NaN | 2.962896e+06 | 2.906050e+06 | 5.684618e+04 | NaN | TCTFILR |
| G6PD | G6PD\_HUMAN | Glucose-6-phosphate 1-dehydrogenase | 385 | 371 | G6PD | G6PD\_HUMAN | Glucose-6-phosphate 1-dehydrogenase | 15C(125.0477) | NaN | X | NaN | NaN | NaN | NaN | 2.958901e+06 | 2.958901e+06 | NaN | NaN | LQFHDVAGDIFHQQCK |
| ATIC | PUR9\_HUMAN | Bifunctional purine biosynthesis protein ATIC | 434 | 427 | ATIC | PUR9\_HUMAN | Bifunctional purine biosynthesis protein ATIC | 8C(125.0477) | NaN | X | NaN | NaN | NaN | NaN | 2.946306e+06 | 2.922511e+06 | 2.379431e+04 | NaN | YTQSNSVCYAK |
| HBG2 | HBG2\_HUMAN | Hemoglobin subunit gamma-2 | 94, 94 | 78 | HBG2 | HBG2\_HUMAN | Hemoglobin subunit gamma-2 | 17C(125.0477) 17C(31.9988) | NaN | X | NaN | X | NaN | NaN | 2.945094e+06 | 2.945094e+06 | NaN | NaN | HLDDLKGTFAQLSELHCDKLHVDPENFK |
| EPB42 | EPB42\_HUMAN | Protein 4.2 | 398, 399 | 398 | EPB42 | EPB42\_HUMAN | Protein 4.2 | 1C(31.9988) 2C(15.9949) | NaN | NaN | X | X | NaN | NaN | 2.917238e+06 | NaN | 2.917238e+06 | NaN | CCEDGTLELTDSNTK |
| ALDH1A1 | AL1A1\_HUMAN | Aldehyde dehydrogenase 1A1 | 186, 186 | 180 | ALDH1A1 | AL1A1\_HUMAN | Aldehyde dehydrogenase 1A1 | 7C(125.0477) 7C(31.9988) | NaN | X | NaN | X | NaN | NaN | 2.882186e+06 | 2.882186e+06 | NaN | NaN | IGPALSCGNTVVVKPAEQTPLTALHVASLIK |
| SEPTIN6 | SEPT6\_HUMAN | Septin-6 | 151, 151 | 148 | SEPTIN6 | SEPT6\_HUMAN | Septin-6 | 4C(125.0477) 4C(57.0215) | X | X | NaN | NaN | NaN | NaN | 2.819672e+06 | 2.819672e+06 | NaN | NaN | IHVCLYFIAPTGHSLK |
| STIP1 | STIP1\_HUMAN | Stress-induced-phosphoprotein 1 | 26, 26 | 14 | STIP1 | STIP1\_HUMAN | Stress-induced-phosphoprotein 1 | 13C(125.0477) 13C(31.9988) | NaN | X | NaN | X | NaN | NaN | 2.774858e+06 | 2.774858e+06 | NaN | NaN | ALSVGNIDDALQCYSEAIK |
| PARK7 | PARK7\_HUMAN | Parkinson disease protein 7 | 46, 46, 46, 46 | 33 | PARK7 | PARK7\_HUMAN | Parkinson disease protein 7 | 14C(125.0477) 14C(15.9949) 14C(31.9988) 14C(57.0215) | X | X | X | X | NaN | NaN | 2.760900e+06 | 2.760900e+06 | NaN | NaN | VTVAGLAGKDPVQCSR |
| USP14 | UBP14\_HUMAN | Ubiquitin carboxyl-terminal hydrolase 14 | 359 | 349 | USP14 | UBP14\_HUMAN | Ubiquitin carboxyl-terminal hydrolase 14 | 11C(125.0477) | NaN | X | NaN | NaN | NaN | NaN | 2.734494e+06 | 2.734494e+06 | NaN | NaN | FPLMLDMYELCTPELQEK |
| SLC4A1 | B3AT\_HUMAN | Band 3 anion transport protein | 885 | 880 | SLC4A1 | B3AT\_HUMAN | Band 3 anion transport protein | 6C(125.0477) | NaN | X | NaN | NaN | NaN | NaN | 2.722529e+06 | 2.722529e+06 | NaN | NaN | NVELQCLDADDAK |
| USP15 | UBP15\_HUMAN | Ubiquitin carboxyl-terminal hydrolase 15 | 570 | 559 | USP15 | UBP15\_HUMAN | Ubiquitin carboxyl-terminal hydrolase 15 | 12C(125.0477) | NaN | X | NaN | NaN | NaN | NaN | 2.713980e+06 | 2.713980e+06 | NaN | NaN | TEDTEHVIIPVCLR |
| ANK1 | ANK1\_HUMAN | Ankyrin-1 | 476, 476 | 469 | ANK1 | ANK1\_HUMAN | Ankyrin-1 | 8C(125.0477) 8C(31.9988) | NaN | X | NaN | X | NaN | NaN | 2.710083e+06 | 2.552598e+06 | NaN | 157484.4200 | DDQTPLHCAAR |
| SPTA1 | SPTA1\_HUMAN | Spectrin alpha chain, erythrocytic 1 | 224, 224, 224 | 214 | SPTA1 | SPTA1\_HUMAN | Spectrin alpha chain, erythrocytic 1 | 11C(125.0477) 11C(31.9988) 11C(57.0215) | X | X | NaN | X | NaN | NaN | 2.706762e+06 | NaN | 2.706762e+06 | NaN | VVEVNQYANECAEENHPDLPLIQSK |
| RNF123 | RN123\_HUMAN | E3 ubiquitin-protein ligase RNF123 | 1187 | 1185 | RNF123 | RN123\_HUMAN | E3 ubiquitin-protein ligase RNF123 | 3C(125.0477) | NaN | X | NaN | NaN | NaN | NaN | 2.665424e+06 | 2.665424e+06 | NaN | NaN | SICYLLGQPEPPAPGTALPAPDR |
| BLVRB | BLVRB\_HUMAN | Flavin reductase (NADPH) | 109 | 100 | BLVRB | BLVRB\_HUMAN | Flavin reductase (NADPH) | 10C(125.0477) | NaN | X | NaN | NaN | NaN | NaN | 2.663937e+06 | 2.663937e+06 | NaN | NaN | AHGVDKVVACTSAFLLWDPTKVPPR |
| LCP1 | PLSL\_HUMAN | Plastin-2 | 140 | 133 | LCP1 | PLSL\_HUMAN | Plastin-2 | 8C(125.0477) | NaN | X | NaN | NaN | NaN | NaN | 2.662619e+06 | 2.598224e+06 | 6.439475e+04 | NaN | ALENDPDCR |
| ANK1 | ANK1\_HUMAN | Ankyrin-1 | 1258, 1258, 1260, 1260 | 1258 | ANK1 | ANK1\_HUMAN | Ankyrin-1 | 1C(125.0477) 1C(31.9988) 3C(15.9949) 3C(31.9988) | NaN | X | X | X | NaN | NaN | 2.658018e+06 | NaN | 2.658018e+06 | NaN | CYCMTDDKVDK |
| LGALS3 | LEG3\_HUMAN | Galectin-3 | 173 | 170 | LGALS3 | LEG3\_HUMAN | Galectin-3 | 4C(125.0477) | NaN | X | NaN | NaN | NaN | NaN | 2.645979e+06 | 2.712267e+05 | 2.106631e+06 | 268120.9400 | VIVCNTK |
| RPIA | RPIA\_HUMAN | Ribose-5-phosphate isomerase | 128 | 119 | RPIA | RPIA\_HUMAN | Ribose-5-phosphate isomerase | 10C(125.0477) | NaN | X | NaN | NaN | NaN | NaN | 2.641106e+06 | 2.641106e+06 | NaN | NaN | VKQENLNLVCIPTSFQAR |
| PHPT1 | PHP14\_HUMAN | 14 kDa phosphohistidine phosphatase | 69, 71, 71, 73, 73 | 67 | PHPT1 | PHP14\_HUMAN | 14 kDa phosphohistidine phosphatase | 3C(125.0477) 5C(125.0477) 5C(15.9949) 7C(125.0477) 7C(15.9949) | NaN | X | X | NaN | NaN | NaN | 2.638944e+06 | 2.638944e+06 | NaN | NaN | QGCDCECLGGGR |
| ADD1 | ADDA\_HUMAN | Alpha-adducin | 236 | 236 | ADD1 | ADDA\_HUMAN | Alpha-adducin | 1C(125.0477) | NaN | X | NaN | NaN | NaN | NaN | 2.619815e+06 | 2.619815e+06 | NaN | NaN | CVVHIHTPAGAAVSAMK |
| CAND1 | CAND1\_HUMAN | Cullin-associated NEDD8-dissociated protein 1 | 1134 | 1130 | CAND1 | CAND1\_HUMAN | Cullin-associated NEDD8-dissociated protein 1 | 5C(125.0477) | NaN | X | NaN | NaN | NaN | NaN | 2.616014e+06 | 2.616014e+06 | NaN | NaN | LSTLCPSAVLQR |
| PRDX2 | PRDX2\_HUMAN | Peroxiredoxin-2 | 172, 172, 172, 172 | 158 | PRDX2 | PRDX2\_HUMAN | Peroxiredoxin-2 | 15C(125.0477) 15C(15.9949) 15C(31.9988) 15C(57.0215) | X | X | X | X | NaN | NaN | 2.608165e+06 | 2.608165e+06 | NaN | NaN | LVQAFQYTDEHGEVCPAGWKPGSDTIKPNVDDSK |
| SPTB | SPTB1\_HUMAN | Spectrin beta chain, erythrocytic | 2012 | 2007 | SPTB | SPTB1\_HUMAN | Spectrin beta chain, erythrocytic | 6C(125.0477) | NaN | X | NaN | NaN | NaN | NaN | 2.603378e+06 | 1.456321e+06 | 1.147057e+06 | NaN | MLLEVCQFSR |
| CAPNS1 | CPNS1\_HUMAN | Calpain small subunit 1 | 144 | 136 | CAPNS1 | CPNS1\_HUMAN | Calpain small subunit 1 | 9C(125.0477) | NaN | X | NaN | NaN | NaN | NaN | 2.597390e+06 | 5.102361e+04 | 2.546367e+06 | NaN | TDGFGIDTCR |
| ANKFY1 | ANFY1\_HUMAN | Rabankyrin-5 | 675 | 665 | ANKFY1 | ANFY1\_HUMAN | Rabankyrin-5 | 11C(125.0477) | NaN | X | NaN | NaN | NaN | NaN | 2.572388e+06 | 2.549778e+06 | 2.260984e+04 | NaN | NQLPLVVDAICTR |
| UBA1 | UBA1\_HUMAN | Ubiquitin-like modifier-activating enzyme 1 | 179 | 175 | UBA1 | UBA1\_HUMAN | Ubiquitin-like modifier-activating enzyme 1 | 5C(125.0477) | NaN | X | NaN | NaN | NaN | NaN | 2.519944e+06 | 2.513194e+06 | 6.749064e+03 | NaN | VGEFCHNR |
| UROS | HEM4\_HUMAN | Uroporphyrinogen-III synthase | 143 | 133 | UROS | HEM4\_HUMAN | Uroporphyrinogen-III synthase | 11C(125.0477) | NaN | X | NaN | NaN | NaN | NaN | 2.487088e+06 | 2.487088e+06 | NaN | NaN | ESSALPLLFPCGNLK |
| TGM2 | TGM2\_HUMAN | Protein-glutamine gamma-glutamyltransferase 2 | 10 | 10 | TGM2 | TGM2\_HUMAN | Protein-glutamine gamma-glutamyltransferase 2 | 1C(125.0477) | NaN | X | NaN | NaN | NaN | NaN | 2.482066e+06 | 2.449487e+06 | 3.257888e+04 | NaN | CDLELETNGR |
| SELENBP1 | SBP1\_HUMAN | Methanethiol oxidase | 268, 268 | 255 | SELENBP1 | SBP1\_HUMAN | Methanethiol oxidase | 14C(125.0477) 14C(31.9988) | NaN | X | NaN | X | NaN | NaN | 2.449449e+06 | 2.449449e+06 | NaN | NaN | FLHNPDAAQGFVGCALSSTIQR |
| AGO2 | AGO2\_HUMAN | Protein argonaute-2 | 564 | 555 | AGO2 | AGO2\_HUMAN | Protein argonaute-2 | 10C(125.0477) | NaN | X | NaN | NaN | NaN | NaN | 2.433759e+06 | 2.367956e+06 | 6.580288e+04 | NaN | TTPQTLSNLCLK |
| GNPDA1 | GNPI1\_HUMAN | Glucosamine-6-phosphate isomerase 1 | 48 | 34 | GNPDA1 | GNPI1\_HUMAN | Glucosamine-6-phosphate isomerase 1 | 15C(125.0477) | NaN | X | NaN | NaN | NaN | NaN | 2.432794e+06 | 2.432794e+06 | NaN | NaN | YFTLGLPTGSTPLGCYK |
| CCT2 | TCPB\_HUMAN | T-complex protein 1 subunit beta | 535 | 529 | CCT2 | TCPB\_HUMAN | T-complex protein 1 subunit beta | 7C(125.0477) | NaN | X | NaN | NaN | NaN | NaN | 2.430085e+06 | 2.205060e+06 | 2.250255e+05 | NaN | VPDHHPC |
| BPGM | PMGE\_HUMAN | Bisphosphoglycerate mutase | 145 | 144 | BPGM | PMGE\_HUMAN | Bisphosphoglycerate mutase | 2C(125.0477) | NaN | X | NaN | NaN | NaN | NaN | 2.425568e+06 | 2.403712e+06 | 2.185588e+04 | NaN | VCDVPLDQLPR |
| ALDOA | ALDOA\_HUMAN | Fructose-bisphosphate aldolase A | 178 | 174 | ALDOA | ALDOA\_HUMAN | Fructose-bisphosphate aldolase A | 5C(125.0477) | NaN | X | NaN | NaN | NaN | NaN | 2.405666e+06 | 2.405666e+06 | 0.000000e+00 | NaN | YASICQQNGIVPIVEPEILPDGDHDLKR |
| RNH1 | RINI\_HUMAN | Ribonuclease inhibitor | 248 | 239 | RNH1 | RINI\_HUMAN | Ribonuclease inhibitor | 10C(125.0477) | NaN | X | NaN | NaN | NaN | NaN | 2.397881e+06 | 2.397881e+06 | NaN | NaN | LGDVGMAELCPGLLHPSSR |
| CLTC | CLH1\_HUMAN | Clathrin heavy chain 1 | 491 | 488 | CLTC | CLH1\_HUMAN | Clathrin heavy chain 1 | 4C(125.0477) | NaN | X | NaN | NaN | NaN | NaN | 2.396202e+06 | 2.361236e+06 | 3.496700e+04 | NaN | VIQCFAETGQVQK |
| CAND1 | CAND1\_HUMAN | Cullin-associated NEDD8-dissociated protein 1 | 237 | 233 | CAND1 | CAND1\_HUMAN | Cullin-associated NEDD8-dissociated protein 1 | 5C(125.0477) | NaN | X | NaN | NaN | NaN | NaN | 2.368531e+06 | 2.368531e+06 | NaN | NaN | TYIQCIAAISR |
| GCLC | GSH1\_HUMAN | Glutamate--cysteine ligase catalytic subunit | 553 | 553 | GCLC | GSH1\_HUMAN | Glutamate--cysteine ligase catalytic subunit | 1C(125.0477) | NaN | X | NaN | NaN | NaN | NaN | 2.343018e+06 | 2.343018e+06 | NaN | NaN | CSILNYLK |
| HGS | HGS\_HUMAN | Hepatocyte growth factor-regulated tyrosine kinase substrate | 166 | 157 | HGS | HGS\_HUMAN | Hepatocyte growth factor-regulated tyrosine kinase substrate | 10C(125.0477) | NaN | X | NaN | NaN | NaN | NaN | 2.326604e+06 | 2.326604e+06 | NaN | NaN | APDWVDAEECHR |
| NAPRT | PNCB\_HUMAN | Nicotinate phosphoribosyltransferase | 385 | 366 | NAPRT | PNCB\_HUMAN | Nicotinate phosphoribosyltransferase | 20C(125.0477) | NaN | X | NaN | NaN | NaN | NaN | 2.281662e+06 | 2.281662e+06 | NaN | NaN | LAQEGSEVNVIGIGTSVVTCPQQPSLGGVYK |
| HBB | HBB\_HUMAN | Hemoglobin subunit beta | 94 | 67 | HBB | HBB\_HUMAN | Hemoglobin subunit beta | 28C(125.0477) | NaN | X | NaN | NaN | NaN | NaN | 2.278448e+06 | 2.278448e+06 | 0.000000e+00 | NaN | KVLGAFSDGLAHLDNLKGTFATLSELHCDK |
| PFKM | PFKAM\_HUMAN | ATP-dependent 6-phosphofructokinase, muscle type | 351, 351 | 346 | PFKM | PFKAM\_HUMAN | ATP-dependent 6-phosphofructokinase, muscle type | 6C(125.0477) 6C(15.9949) | NaN | X | X | NaN | NaN | NaN | 2.254225e+06 | 2.254225e+06 | NaN | NaN | LPLMECVQVTK |
| ANK1 | ANK1\_HUMAN | Ankyrin-1 | 320, 320 | 304 | ANK1 | ANK1\_HUMAN | Ankyrin-1 | 17C(125.0477) 17C(15.9949) | NaN | X | X | NaN | NaN | NaN | 2.250224e+06 | 2.250224e+06 | NaN | NaN | NGLSPIHMAAQGDHLDCVR |
| PREP | PPCE\_HUMAN | Prolyl endopeptidase | 57 | 49 | PREP | PPCE\_HUMAN | Prolyl endopeptidase | 9C(125.0477) | NaN | X | NaN | NaN | NaN | NaN | 2.240058e+06 | 2.240058e+06 | NaN | NaN | ITVPFLEQCPIR |
| HPS1 | HPS1\_HUMAN | Hermansky-Pudlak syndrome 1 protein | 165, 165 | 161 | HPS1 | HPS1\_HUMAN | Hermansky-Pudlak syndrome 1 protein | 5C(15.9949) 5C(57.0215) | X | NaN | X | NaN | NaN | NaN | 2.193722e+06 | NaN | 2.193722e+06 | NaN | EQEQCFAVEALER |
| GDI1 | GDIA\_HUMAN | Rab GDP dissociation inhibitor alpha | 282 | 279 | GDI1 | GDIA\_HUMAN | Rab GDP dissociation inhibitor alpha | 4C(125.0477) | NaN | X | NaN | NaN | NaN | NaN | 2.191396e+06 | 2.191396e+06 | NaN | NaN | QLICDPSYIPDR |
| SLC14A1 | UT1\_HUMAN | Urea transporter 1 | 25, 25 | 18 | SLC14A1 | UT1\_HUMAN | Urea transporter 1 | 8C(125.0477) 8C(31.9988) | NaN | X | NaN | X | NaN | NaN | 2.180530e+06 | NaN | NaN | 2180529.5000 | GENQVSPCQGR |
| HSPA8 | HSP7C\_HUMAN | Heat shock cognate 71 kDa protein | 603, 603 | 602 | HSPA8 | HSP7C\_HUMAN | Heat shock cognate 71 kDa protein | 2C(125.0477) 2C(31.9988) | NaN | X | NaN | X | NaN | NaN | 2.175358e+06 | NaN | 2.175358e+06 | NaN | VCNPIITK |
| TGM2 | TGM2\_HUMAN | Protein-glutamine gamma-glutamyltransferase 2 | 545 | 541 | TGM2 | TGM2\_HUMAN | Protein-glutamine gamma-glutamyltransferase 2 | 5C(125.0477) | NaN | X | NaN | NaN | NaN | NaN | 2.152230e+06 | 2.126881e+06 | 2.534882e+04 | NaN | SVPLCILYEK |
| DDB1 | DDB1\_HUMAN | DNA damage-binding protein 1 | 652 | 643 | DDB1 | DDB1\_HUMAN | DNA damage-binding protein 1 | 10C(125.0477) | NaN | X | NaN | NaN | NaN | NaN | 2.133230e+06 | 2.133230e+06 | NaN | NaN | SLSTTNVFACSDRPTVIYSSNHK |
| CFL1 | COF1\_HUMAN | Cofilin-1 | 139 | 128 | CFL1 | COF1\_HUMAN | Cofilin-1 | 12C(125.0477) | NaN | X | NaN | NaN | NaN | NaN | 2.126890e+06 | 2.126890e+06 | NaN | NaN | LTGIKHELQANCYEEVKDR |
| HINT1 | HINT1\_HUMAN | Adenosine 5'-monophosphoramidase HINT1 | 38 | 31 | HINT1 | HINT1\_HUMAN | Adenosine 5'-monophosphoramidase HINT1 | 8C(125.0477) | NaN | X | NaN | NaN | NaN | NaN | 2.126516e+06 | 2.126516e+06 | NaN | NaN | IIFEDDRCLAFHDISPQAPTHFLVIPK |
| MIF | MIF\_HUMAN | Macrophage migration inhibitory factor | 81 | 79 | MIF | MIF\_HUMAN | Macrophage migration inhibitory factor | 3C(125.0477) | NaN | X | NaN | NaN | NaN | NaN | 2.118260e+06 | 2.118260e+06 | NaN | NaN | LLCGLLAER |
| COPS3 | CSN3\_HUMAN | COP9 signalosome complex subunit 3 | 383 | 383 | COPS3 | CSN3\_HUMAN | COP9 signalosome complex subunit 3 | 1C(125.0477) | NaN | X | NaN | NaN | NaN | NaN | 2.113180e+06 | 1.847196e+06 | 2.659832e+05 | NaN | CIELDER |
| RIC8A | RIC8A\_HUMAN | Synembryn-A | 341 | 329 | RIC8A | RIC8A\_HUMAN | Synembryn-A | 13C(125.0477) | NaN | X | NaN | NaN | NaN | NaN | 2.095966e+06 | 2.095966e+06 | NaN | NaN | ESVAPVLSVLTECAR |
| CUL2 | CUL2\_HUMAN | Cullin-2 | 46 | 39 | CUL2 | CUL2\_HUMAN | Cullin-2 | 8C(125.0477) | NaN | X | NaN | NaN | NaN | NaN | 2.083733e+06 | 2.083733e+06 | NaN | NaN | FSDIYALCVAYPEPLGER |
| UBE2O | UBE2O\_HUMAN | (E3-independent) E2 ubiquitin-conjugating enzyme | 182 | 177 | UBE2O | UBE2O\_HUMAN | (E3-independent) E2 ubiquitin-conjugating enzyme | 6C(125.0477) | NaN | X | NaN | NaN | NaN | NaN | 2.078366e+06 | 2.071187e+06 | 7.179520e+03 | NaN | LIGTNCIIYPVNSK |
| PITPNB | PIPNB\_HUMAN | Phosphatidylinositol transfer protein beta isoform | 230 | 227 | PITPNB | PIPNB\_HUMAN | Phosphatidylinositol transfer protein beta isoform | 4C(125.0477) | NaN | X | NaN | NaN | NaN | NaN | 2.077901e+06 | 2.077901e+06 | NaN | NaN | QLFCWIDK |
| PGM2L1 | PGM2L\_HUMAN | Glucose 1,6-bisphosphate synthase | 125 | 119 | PGM2L1 | PGM2L\_HUMAN | Glucose 1,6-bisphosphate synthase | 7C(125.0477) | NaN | X | NaN | NaN | NaN | NaN | 2.074804e+06 | 2.043180e+06 | 3.162390e+04 | NaN | GQVTSSCSSQR |
| USP14 | UBP14\_HUMAN | Ubiquitin carboxyl-terminal hydrolase 14 | 257, 257 | 257 | USP14 | UBP14\_HUMAN | Ubiquitin carboxyl-terminal hydrolase 14 | 1C(125.0477) 1C(57.0215) | X | X | NaN | NaN | NaN | NaN | 2.063668e+06 | 2.063668e+06 | NaN | NaN | CTESEEEEVTK |
| PARK7 | PARK7\_HUMAN | Parkinson disease protein 7 | 53, 46 | 42 | PARK7 | PARK7\_HUMAN | Parkinson disease protein 7 | 12C(15.9949) 5C(15.9949) | NaN | NaN | X | NaN | NaN | NaN | 2.024963e+06 | 2.024963e+06 | NaN | NaN | DPVQCSRDVVICPDASLEDAK |
| PSME1 | PSME1\_HUMAN | Proteasome activator complex subunit 1 | 101, 101, 106 | 91 | PSME1 | PSME1\_HUMAN | Proteasome activator complex subunit 1 | 11C(125.0477) 11C(31.9988) 16C(15.9949) | NaN | X | X | X | NaN | NaN | 2.013688e+06 | 2.013688e+06 | NaN | NaN | KGEDEDKGPPCGPVNCNEK |
| SKP1 | SKP1\_HUMAN | S-phase kinase-associated protein 1 | 160 | 155 | SKP1 | SKP1\_HUMAN | S-phase kinase-associated protein 1 | 6C(125.0477) | NaN | X | NaN | NaN | NaN | NaN | 2.013526e+06 | 1.911398e+06 | 1.021283e+05 | NaN | KENQWCEEK |
| SNAP91 | AP180\_HUMAN | Clathrin coat assembly protein AP180 | 27 | 10 | SNAP91 | AP180\_HUMAN | Clathrin coat assembly protein AP180 | 18C(31.9988) | NaN | NaN | NaN | X | NaN | NaN | 2.003090e+06 | 2.003090e+06 | NaN | NaN | IAAAQYSVTGSAVARAVCK |
| GDI2 | GDIB\_HUMAN | Rab GDP dissociation inhibitor beta | 414 | 403 | GDI2 | GDIB\_HUMAN | Rab GDP dissociation inhibitor beta | 12C(125.0477) | NaN | X | NaN | NaN | NaN | NaN | 2.000442e+06 | 1.975255e+06 | 2.518753e+04 | NaN | TYDATTHFETTCDDIK |
| USP7 | UBP7\_HUMAN | Ubiquitin carboxyl-terminal hydrolase 7 | 90 | 81 | USP7 | UBP7\_HUMAN | Ubiquitin carboxyl-terminal hydrolase 7 | 10C(125.0477) | NaN | X | NaN | NaN | NaN | NaN | 1.993325e+06 | 1.993325e+06 | NaN | NaN | LSESVLSPPCFVR |
| FKBP1A | FKB1A\_HUMAN | Peptidyl-prolyl cis-trans isomerase FKBP1A | 23, 23, 23 | 19 | FKBP1A | FKB1A\_HUMAN | Peptidyl-prolyl cis-trans isomerase FKBP1A | 5C(125.0477) 5C(15.9949) 5C(31.9988) | NaN | X | X | X | NaN | NaN | 1.982312e+06 | 1.982312e+06 | NaN | NaN | RGQTCVVHYTGMLEDGKK |
| SORD | DHSO\_HUMAN | Sorbitol dehydrogenase | 301 | 300 | SORD | DHSO\_HUMAN | Sorbitol dehydrogenase | 2C(125.0477) | NaN | X | NaN | NaN | NaN | NaN | 1.977397e+06 | 1.977397e+06 | NaN | NaN | YCNTWPVAISMLASK |
| CAT | CATA\_HUMAN | Catalase | 460, 460 | 459 | CAT | CATA\_HUMAN | Catalase | 2C(125.0477) 2C(57.0215) | X | X | NaN | NaN | NaN | NaN | 1.977386e+06 | 1.977386e+06 | NaN | NaN | LCENIAGHLKDAQIFIQK |
| FERMT3 | URP2\_HUMAN | Fermitin family homolog 3 | 128 | 114 | FERMT3 | URP2\_HUMAN | Fermitin family homolog 3 | 15C(125.0477) | NaN | X | NaN | NaN | NaN | NaN | 1.966700e+06 | 1.966700e+06 | NaN | NaN | ASFSQPLFQAVAAICR |
| ECPAS | ECM29\_HUMAN | Proteasome adapter and scaffold protein ECM29 | 1257 | 1250 | ECPAS | ECM29\_HUMAN | Proteasome adapter and scaffold protein ECM29 | 8C(125.0477) | NaN | X | NaN | NaN | NaN | NaN | 1.950210e+06 | 1.950210e+06 | NaN | NaN | TIAALLPCLLDK |
| BLVRA | BIEA\_HUMAN | Biliverdin reductase A | 281 | 278 | BLVRA | BIEA\_HUMAN | Biliverdin reductase A | 4C(125.0477) | NaN | X | NaN | NaN | NaN | NaN | 1.949254e+06 | 1.949254e+06 | NaN | NaN | ILHCLGLAEEIQK |
| SKP1 | SKP1\_HUMAN | S-phase kinase-associated protein 1 | 160 | 156 | SKP1 | SKP1\_HUMAN | S-phase kinase-associated protein 1 | 5C(125.0477) | NaN | X | NaN | NaN | NaN | NaN | 1.946963e+06 | 1.837869e+06 | 1.090945e+05 | NaN | ENQWCEEK |
| PSMD2 | PSMD2\_HUMAN | 26S proteasome non-ATPase regulatory subunit 2 | 448 | 442 | PSMD2 | PSMD2\_HUMAN | 26S proteasome non-ATPase regulatory subunit 2 | 7C(125.0477) | NaN | X | NaN | NaN | NaN | NaN | 1.933204e+06 | 1.933204e+06 | NaN | NaN | SGALLACGIVNSGVR |
| CARM1 | CARM1\_HUMAN | Histone-arginine methyltransferase CARM1 | 438, 438 | 430 | CARM1 | CARM1\_HUMAN | Histone-arginine methyltransferase CARM1 | 9C(125.0477) 9C(31.9988) | NaN | X | NaN | X | NaN | NaN | 1.932853e+06 | 1.932853e+06 | NaN | NaN | AGDTLSGTCLLIANKR |
| AP2A1 | AP2A1\_HUMAN | AP-2 complex subunit alpha-1 | 331 | 330 | AP2A1 | AP2A1\_HUMAN | AP-2 complex subunit alpha-1 | 2C(125.0477) | NaN | X | NaN | NaN | NaN | NaN | 1.920691e+06 | 1.920691e+06 | NaN | NaN | ACNQLGQFLQHR |
| PFKL | PFKAL\_HUMAN | ATP-dependent 6-phosphofructokinase, liver type | 351 | 346 | PFKL | PFKAL\_HUMAN | ATP-dependent 6-phosphofructokinase, liver type | 6C(125.0477) | NaN | X | NaN | NaN | NaN | NaN | 1.914914e+06 | 1.914914e+06 | NaN | NaN | LPLMECVQMTK |
| COPS2 | CSN2\_HUMAN | COP9 signalosome complex subunit 2 | 179, 179 | 174 | COPS2 | CSN2\_HUMAN | COP9 signalosome complex subunit 2 | 6C(125.0477) 6C(31.9988) | NaN | X | NaN | X | NaN | NaN | 1.904420e+06 | 1.904420e+06 | NaN | NaN | QLHQSCQTDDGEDDLKK |
| EPB41 | EPB41\_HUMAN | Protein 4.1 | 224 | 214 | EPB41 | EPB41\_HUMAN | Protein 4.1 | 11C(125.0477) | NaN | X | NaN | NaN | NaN | NaN | 1.901268e+06 | 1.901268e+06 | NaN | NaN | VSLLDDTVYECVVEK |
| SPTA1 | SPTA1\_HUMAN | Spectrin alpha chain, erythrocytic 1 | 1203 | 1203 | SPTA1 | SPTA1\_HUMAN | Spectrin alpha chain, erythrocytic 1 | 1C(125.0477) | NaN | X | NaN | NaN | NaN | NaN | 1.887289e+06 | 1.887289e+06 | NaN | NaN | CQALSAADPGSDLFSVQALQR |
| PRDX6 | PRDX6\_HUMAN | Peroxiredoxin-6 | 91, 91, 91, 91 | 85 | PRDX6 | PRDX6\_HUMAN | Peroxiredoxin-6 | 7C(125.0477) 7C(15.9949) 7C(31.9988) 7C(57.0215) | X | X | X | X | NaN | NaN | 1.866649e+06 | 1.866649e+06 | NaN | NaN | DINAYNCEEPTEK |
| HBD | HBD\_HUMAN | Hemoglobin subunit delta | 113, 113 | 106 | HBD | HBD\_HUMAN | Hemoglobin subunit delta | 8C(125.0477) 8C(31.9988) | NaN | X | NaN | X | NaN | NaN | 1.864962e+06 | NaN | 1.864962e+06 | NaN | LLGNVLVCVLAR |
| ANK1 | ANK1\_HUMAN | Ankyrin-1 | 278, 278 | 271 | ANK1 | ANK1\_HUMAN | Ankyrin-1 | 8C(125.0477) 8C(31.9988) | NaN | X | NaN | X | NaN | NaN | 1.845761e+06 | NaN | 1.845761e+06 | NaN | DELTPLHCAAR |
| PGAM1 | PGAM1\_HUMAN | Phosphoglycerate mutase 1 | 153 | 142 | PGAM1 | PGAM1\_HUMAN | Phosphoglycerate mutase 1 | 12C(125.0477) | NaN | X | NaN | NaN | NaN | NaN | 1.837559e+06 | 1.812391e+06 | 2.516822e+04 | NaN | YADLTEDQLPSCESLK |
| KLC3 | KLC3\_HUMAN | Kinesin light chain 3 | 343 | 324 | KLC3 | KLC3\_HUMAN | Kinesin light chain 3 | 20C(15.9949) | NaN | NaN | X | NaN | NaN | NaN | 1.834073e+06 | 1.834073e+06 | NaN | NaN | VLGADHPDVAKQLNNLALLCQNQGKFEDVER |
| SELENBP1 | SBP1\_HUMAN | Methanethiol oxidase | 31 | 24 | SELENBP1 | SBP1\_HUMAN | Methanethiol oxidase | 8C(125.0477) | NaN | X | NaN | NaN | NaN | NaN | 1.819351e+06 | 1.819351e+06 | NaN | NaN | EEIVYLPCIYR |
| YWHAG | 1433G\_HUMAN | 14-3-3 protein gamma | 112, 112 | 111 | YWHAG | 1433G\_HUMAN | 14-3-3 protein gamma | 2C(125.0477) 2C(31.9988) | NaN | X | NaN | X | NaN | NaN | 1.812438e+06 | 1.812438e+06 | NaN | NaN | NCSETQYESK |
| AGO2 | AGO2\_HUMAN | Protein argonaute-2 | 717 | 715 | AGO2 | AGO2\_HUMAN | Protein argonaute-2 | 3C(125.0477) | NaN | X | NaN | NaN | NaN | NaN | 1.810396e+06 | 1.758749e+06 | 5.164723e+04 | NaN | LFCTDKNER |
| SPTB | SPTB1\_HUMAN | Spectrin beta chain, erythrocytic | 1892 | 1878 | SPTB | SPTB1\_HUMAN | Spectrin beta chain, erythrocytic | 15C(125.0477) | NaN | X | NaN | NaN | NaN | NaN | 1.809736e+06 | 1.809736e+06 | NaN | NaN | EQEVSAAWQALLDACAGR |
| AP2A1 | AP2A1\_HUMAN | AP-2 complex subunit alpha-1 | 171 | 166 | AP2A1 | AP2A1\_HUMAN | AP-2 complex subunit alpha-1 | 6C(125.0477) | NaN | X | NaN | NaN | NaN | NaN | 1.794181e+06 | 1.732064e+06 | 6.211789e+04 | NaN | QSAALCLLR |
| GDI2 | GDIB\_HUMAN | Rab GDP dissociation inhibitor beta | 335 | 329 | GDI2 | GDIB\_HUMAN | Rab GDP dissociation inhibitor beta | 7C(125.0477) | NaN | X | NaN | NaN | NaN | NaN | 1.784228e+06 | 1.784228e+06 | NaN | NaN | KSDIYVCMISFAHNVAAQGK |
| AK1 | KAD1\_HUMAN | Adenylate kinase isoenzyme 1 | 187 | 172 | AK1 | KAD1\_HUMAN | Adenylate kinase isoenzyme 1 | 16C(125.0477) | NaN | X | NaN | NaN | NaN | NaN | 1.764915e+06 | 1.764915e+06 | NaN | NaN | KVNAEGSVDSVFSQVCTHLDALK |
| SPTA1 | SPTA1\_HUMAN | Spectrin alpha chain, erythrocytic 1 | 2101 | 2101 | SPTA1 | SPTA1\_HUMAN | Spectrin alpha chain, erythrocytic 1 | 1C(125.0477) | NaN | X | NaN | NaN | NaN | NaN | 1.758428e+06 | 1.758428e+06 | NaN | NaN | CLLELDQQIK |
| PRDX2 | PRDX2\_HUMAN | Peroxiredoxin-2 | 70, 70 | 68 | PRDX2 | PRDX2\_HUMAN | Peroxiredoxin-2 | 3C(125.0477) 3C(31.9988) | NaN | X | NaN | X | NaN | NaN | 1.737774e+06 | 1.737774e+06 | NaN | NaN | LGCEVLGVSVDSQFTHLAWINTPR |
| C9orf64 | QSPP\_HUMAN | Queuosine salvage protein | 292 | 291 | C9orf64 | QSPP\_HUMAN | Queuosine salvage protein | 2C(125.0477) | NaN | X | NaN | NaN | NaN | NaN | 1.733836e+06 | 1.733836e+06 | NaN | NaN | DCLLELIEQK |
| WDR1 | WDR1\_HUMAN | WD repeat-containing protein 1 | 507 | 490 | WDR1 | WDR1\_HUMAN | WD repeat-containing protein 1 | 18C(125.0477) | NaN | X | NaN | NaN | NaN | NaN | 1.730175e+06 | 1.730175e+06 | NaN | NaN | GPVTDVAYSHDGAFLAVCDASK |
| MSN | MOES\_HUMAN | Moesin | 284 | 280 | MSN | MOES\_HUMAN | Moesin | 5C(125.0477) | NaN | X | NaN | NaN | NaN | NaN | 1.721431e+06 | 1.721431e+06 | NaN | NaN | ILALCMGNHELYMR |
| UBE2O | UBE2O\_HUMAN | (E3-independent) E2 ubiquitin-conjugating enzyme | 230 | 230 | UBE2O | UBE2O\_HUMAN | (E3-independent) E2 ubiquitin-conjugating enzyme | 1C(125.0477) | NaN | X | NaN | NaN | NaN | NaN | 1.715199e+06 | 1.703208e+06 | 1.199141e+04 | NaN | CSMNTEDGAK |
| CAPZA1 | CAZA1\_HUMAN | F-actin-capping protein subunit alpha-1 | 157 | 147 | CAPZA1 | CAZA1\_HUMAN | F-actin-capping protein subunit alpha-1 | 11C(125.0477) | NaN | X | NaN | NaN | NaN | NaN | 1.702753e+06 | 1.702753e+06 | NaN | NaN | TIDGQQTIIACIESHQFQPK |
| G6PD | G6PD\_HUMAN | Glucose-6-phosphate 1-dehydrogenase | 446 | 440 | G6PD | G6PD\_HUMAN | Glucose-6-phosphate 1-dehydrogenase | 7C(125.0477) | NaN | X | NaN | NaN | NaN | NaN | 1.697902e+06 | 1.697902e+06 | NaN | NaN | LILDVFCGSQMHFVR |
| RANBP1 | RANG\_HUMAN | Ran-specific GTPase-activating protein | 158 | 153 | RANBP1 | RANG\_HUMAN | Ran-specific GTPase-activating protein | 6C(125.0477) | NaN | X | NaN | NaN | NaN | NaN | 1.685162e+06 | 1.685162e+06 | NaN | NaN | TKFEECRK |
| UCHL3 | UCHL3\_HUMAN | Ubiquitin carboxyl-terminal hydrolase isozyme L3 | 95, 95 | 89 | UCHL3 | UCHL3\_HUMAN | Ubiquitin carboxyl-terminal hydrolase isozyme L3 | 7C(125.0477) 7C(31.9988) | NaN | X | NaN | X | NaN | NaN | 1.684115e+06 | 1.684115e+06 | NaN | NaN | QTISNACGTIGLIHAIANNK |
| CCT7 | TCPH\_HUMAN | T-complex protein 1 subunit eta | 29 | 11 | CCT7 | TCPH\_HUMAN | T-complex protein 1 subunit eta | 19C(125.0477) | NaN | X | NaN | NaN | NaN | NaN | 1.681183e+06 | 1.681183e+06 | NaN | NaN | EGTDSSQGIPQLVSNISACQVIAEAVR |
| GMPR | GMPR1\_HUMAN | GMP reductase 1 | 205 | 192 | GMPR | GMPR1\_HUMAN | GMP reductase 1 | 14C(125.0477) | NaN | X | NaN | NaN | NaN | NaN | 1.674544e+06 | 1.674544e+06 | NaN | NaN | TGVGYPQLSAVIECADSAHGLK |
| GCLC | GSH1\_HUMAN | Glutamate--cysteine ligase catalytic subunit | 142, 142 | 130 | GCLC | GSH1\_HUMAN | Glutamate--cysteine ligase catalytic subunit | 13C(125.0477) 13C(31.9988) | NaN | X | NaN | X | NaN | NaN | 1.668310e+06 | 1.668310e+06 | NaN | NaN | EATSILEENQALCTITSFPR |
| ARG1 | ARGI1\_HUMAN | Arginase-1 | 168 | 154 | ARG1 | ARGI1\_HUMAN | Arginase-1 | 15C(125.0477) | NaN | X | NaN | NaN | NaN | NaN | 1.664531e+06 | 1.664531e+06 | NaN | NaN | GKIPDVPGFSWVTPCISAK |
| HAGH | GLO2\_HUMAN | Hydroxyacylglutathione hydrolase, mitochondrial | 219, 219 | 217 | HAGH | GLO2\_HUMAN | Hydroxyacylglutathione hydrolase, mitochondrial | 3C(125.0477) 3C(57.0215) | X | X | NaN | NaN | NaN | NaN | 1.660123e+06 | 1.660123e+06 | NaN | NaN | VYCGHEYTINNLK |
| PFKL | PFKAL\_HUMAN | ATP-dependent 6-phosphofructokinase, liver type | 114, 114 | 99 | PFKL | PFKAL\_HUMAN | ATP-dependent 6-phosphofructokinase, liver type | 16C(-34.0809) 16C(125.0477) | NaN | X | NaN | NaN | X | NaN | 1.655631e+06 | 1.655631e+06 | NaN | NaN | RAAAYNLVQHGITNLCVIGGDGSLTGANIFR |
| ALDOA | ALDOA\_HUMAN | Fructose-bisphosphate aldolase A | 202 | 202 | ALDOA | ALDOA\_HUMAN | Fructose-bisphosphate aldolase A | 1C(125.0477) | NaN | X | NaN | NaN | NaN | NaN | 1.654957e+06 | 1.256402e+06 | 2.120695e+05 | 186484.7700 | CQYVTEK |
| BLVRB | BLVRB\_HUMAN | Flavin reductase (NADPH) | 109, 109, 109 | 106 | BLVRB | BLVRB\_HUMAN | Flavin reductase (NADPH) | 4C(125.0477) 4C(31.9988) 4C(57.0215) | X | X | NaN | X | NaN | NaN | 1.649774e+06 | 1.649774e+06 | NaN | NaN | VVACTSAFLLWDPTKVPPR |
| WDR81 | WDR81\_HUMAN | WD repeat-containing protein 81 | 1894 | 1891 | WDR81 | WDR81\_HUMAN | WD repeat-containing protein 81 | 4C(125.0477) | NaN | X | NaN | NaN | NaN | NaN | 1.642687e+06 | 1.642687e+06 | NaN | NaN | IGVCSLLEPPSQATTK |
| SPTB | SPTB1\_HUMAN | Spectrin beta chain, erythrocytic | 1167, 1167 | 1161 | SPTB | SPTB1\_HUMAN | Spectrin beta chain, erythrocytic | 7C(125.0477) 7C(31.9988) | NaN | X | NaN | X | NaN | NaN | 1.635522e+06 | 1.159353e+06 | 4.761692e+05 | NaN | SHTLAQCLGFQEFQK |
| EIF5A | IF5A1\_HUMAN | Eukaryotic translation initiation factor 5A-1 | 73 | 69 | EIF5A | IF5A1\_HUMAN | Eukaryotic translation initiation factor 5A-1 | 5C(125.0477) | NaN | X | NaN | NaN | NaN | NaN | 1.632092e+06 | 1.632092e+06 | NaN | NaN | YEDICPSTHNMDVPNIK |
| CAPN7 | CAN7\_HUMAN | Calpain-7 | 361 | 357 | CAPN7 | CAN7\_HUMAN | Calpain-7 | 5C(57.0215) | X | NaN | NaN | NaN | NaN | NaN | 1.627092e+06 | NaN | 1.547517e+06 | 79574.8900 | GELLCSYSNNK |
| PTGES3 | TEBP\_HUMAN | Prostaglandin E synthase 3 | 58 | 49 | PTGES3 | TEBP\_HUMAN | Prostaglandin E synthase 3 | 10C(125.0477) | NaN | X | NaN | NaN | NaN | NaN | 1.616796e+06 | 1.616796e+06 | NaN | NaN | HLNEIDLFHCIDPNDSK |
| ST13 | F10A1\_HUMAN | Hsc70-interacting protein | 209 | 194 | ST13 | F10A1\_HUMAN | Hsc70-interacting protein | 16C(125.0477) | NaN | X | NaN | NaN | NaN | NaN | 1.616260e+06 | 1.616260e+06 | NaN | NaN | LLGHWEEAAHDLALACK |
| PSMA6 | PSA6\_HUMAN | Proteasome subunit alpha type-6 | 78 | 72 | PSMA6 | PSA6\_HUMAN | Proteasome subunit alpha type-6 | 7C(125.0477) | NaN | X | NaN | NaN | NaN | NaN | 1.611452e+06 | 1.611452e+06 | NaN | NaN | ITENIGCVMTGMTADSR |
| BOLA2 | BOLA2\_HUMAN | BolA-like protein 2 | 59 | 55 | BOLA2 | BOLA2\_HUMAN | BolA-like protein 2 | 5C(125.0477) | NaN | X | NaN | NaN | NaN | NaN | 1.608592e+06 | 1.608592e+06 | NaN | NaN | LVNACLAEELPHIHAFEQK |
| PSME1 | PSME1\_HUMAN | Proteasome activator complex subunit 1 | 22 | 14 | PSME1 | PSME1\_HUMAN | Proteasome activator complex subunit 1 | 9C(125.0477) | NaN | X | NaN | NaN | NaN | NaN | 1.604737e+06 | 1.579848e+06 | 2.488834e+04 | NaN | VDVFREDLCTK |
| PAICS | PUR6\_HUMAN | Multifunctional protein ADE2 | 281 | 268 | PAICS | PUR6\_HUMAN | Multifunctional protein ADE2 | 14C(125.0477) | NaN | X | NaN | NaN | NaN | NaN | 1.588272e+06 | 1.588272e+06 | NaN | NaN | VVVLMGSTSDLGHCEK |
| NAPRT | PNCB\_HUMAN | Nicotinate phosphoribosyltransferase | 484 | 476 | NAPRT | PNCB\_HUMAN | Nicotinate phosphoribosyltransferase | 9C(125.0477) | NaN | X | NaN | NaN | NaN | NaN | 1.582198e+06 | 1.582198e+06 | NaN | NaN | LCLQQGQLCEPLPSLAESR |
| HSPA1A | HS71A\_HUMAN | Heat shock 70 kDa protein 1A | 306 | 300 | HSPA1A | HS71A\_HUMAN | Heat shock 70 kDa protein 1A | 7C(125.0477) | NaN | X | NaN | NaN | NaN | NaN | 1.580585e+06 | 1.580585e+06 | NaN | NaN | ARFEELCSDLFR |
| ANK1 | ANK1\_HUMAN | Ankyrin-1 | 278, 278 | 269 | ANK1 | ANK1\_HUMAN | Ankyrin-1 | 10C(125.0477) 10C(31.9988) | NaN | X | NaN | X | NaN | NaN | 1.572406e+06 | 1.572406e+06 | NaN | NaN | TKDELTPLHCAAR |
| PIP4K2A | PI42A\_HUMAN | Phosphatidylinositol 5-phosphate 4-kinase type-2 alpha | 343 | 343 | PIP4K2A | PI42A\_HUMAN | Phosphatidylinositol 5-phosphate 4-kinase type-2 alpha | 1C(125.0477) | NaN | X | NaN | NaN | NaN | NaN | 1.530135e+06 | 1.520186e+06 | 9.949325e+03 | NaN | CHENSPR |
| HBA1 | HBA\_HUMAN | Hemoglobin subunit alpha | 105, 105, 105, 105, 105 | 101 | HBA1 | HBA\_HUMAN | Hemoglobin subunit alpha | 5C(-34.0809) 5C(125.0477) 5C(15.9949) 5C(31.9988) 5C(57.0215) | X | X | X | X | X | NaN | 1.509443e+06 | NaN | 1.509443e+06 | NaN | LLSHCLLVTLAAHLPAEFTPAVHASLDKFLASVSTVLTSK |
| LAP3 | AMPL\_HUMAN | Cytosol aminopeptidase | 376 | 369 | LAP3 | AMPL\_HUMAN | Cytosol aminopeptidase | 8C(125.0477) | NaN | X | NaN | NaN | NaN | NaN | 1.504213e+06 | 1.504213e+06 | NaN | NaN | LILADALCYAHTFNPK |
| ELP1 | ELP1\_HUMAN | Elongator complex protein 1 | 319 | 318 | ELP1 | ELP1\_HUMAN | Elongator complex protein 1 | 2C(15.9949) | NaN | NaN | X | NaN | NaN | NaN | 1.500676e+06 | 1.500676e+06 | NaN | NaN | TCVQLWTVGNYHWYLKQSLSFSTCGK |
| PPP2CA | PP2AA\_HUMAN | Serine/threonine-protein phosphatase 2A catalytic subunit alpha isoform | 266 | 255 | PPP2CA | PP2AA\_HUMAN | Serine/threonine-protein phosphatase 2A catalytic subunit alpha isoform | 12C(125.0477) | NaN | X | NaN | NaN | NaN | NaN | 1.481928e+06 | 1.481928e+06 | NaN | NaN | NVVTIFSAPNYCYR |
| HPRT1 | HPRT\_HUMAN | Hypoxanthine-guanine phosphoribosyltransferase | 66, 66 | 56 | HPRT1 | HPRT\_HUMAN | Hypoxanthine-guanine phosphoribosyltransferase | 11C(125.0477) 11C(31.9988) | NaN | X | NaN | X | NaN | NaN | 1.467822e+06 | 1.467822e+06 | NaN | NaN | EMGGHHIVALCVLK |
| YWHAZ | 1433Z\_HUMAN | 14-3-3 protein zeta/delta | 94 | 92 | YWHAZ | 1433Z\_HUMAN | 14-3-3 protein zeta/delta | 3C(125.0477) | NaN | X | NaN | NaN | NaN | NaN | 1.459450e+06 | 1.459450e+06 | 0.000000e+00 | NaN | DICNDVLSLLEK |
| XPO1 | XPO1\_HUMAN | Exportin-1 | 119, 119 | 113 | XPO1 | XPO1\_HUMAN | Exportin-1 | 7C(125.0477) 7C(31.9988) | NaN | X | NaN | X | NaN | NaN | 1.450867e+06 | 1.450867e+06 | NaN | NaN | TSSDPTCVEK |
| CBR1 | CBR1\_HUMAN | Carbonyl reductase [NADPH] 1 | 122 | 120 | CBR1 | CBR1\_HUMAN | Carbonyl reductase [NADPH] 1 | 3C(125.0477) | NaN | X | NaN | NaN | NaN | NaN | 1.450001e+06 | 1.450001e+06 | NaN | NaN | DVCTELLPLIKPQGR |
| ALB | ALBU\_HUMAN | Albumin | 289, 289, 289 | 287 | ALB | ALBU\_HUMAN | Albumin | 3C(125.0477) 3C(31.9988) 3C(57.0215) | X | X | NaN | X | NaN | NaN | 1.445863e+06 | NaN | 1.445863e+06 | NaN | YICENQDSISSK |
| TCP1 | TCPA\_HUMAN | T-complex protein 1 subunit alpha | 397 | 391 | TCP1 | TCPA\_HUMAN | T-complex protein 1 subunit alpha | 7C(125.0477) | NaN | X | NaN | NaN | NaN | NaN | 1.444658e+06 | 1.444658e+06 | NaN | NaN | SLHDALCVVKR |
| MDH1 | MDHC\_HUMAN | Malate dehydrogenase, cytoplasmic | 251 | 249 | MDH1 | MDHC\_HUMAN | Malate dehydrogenase, cytoplasmic | 3C(125.0477) | NaN | X | NaN | NaN | NaN | NaN | 1.441823e+06 | 1.383634e+06 | 5.818930e+04 | NaN | AICDHVR |
| SPTA1 | SPTA1\_HUMAN | Spectrin alpha chain, erythrocytic 1 | 773 | 766 | SPTA1 | SPTA1\_HUMAN | Spectrin alpha chain, erythrocytic 1 | 8C(125.0477) | NaN | X | NaN | NaN | NaN | NaN | 1.428837e+06 | 1.428837e+06 | NaN | NaN | ARQESLVCR |
| ALDOC | ALDOC\_HUMAN | Fructose-bisphosphate aldolase C | 240 | 216 | ALDOC | ALDOC\_HUMAN | Fructose-bisphosphate aldolase C | 25C(125.0477) | NaN | X | NaN | NaN | NaN | NaN | 1.425910e+06 | 1.425910e+06 | NaN | NaN | ALSDHHVYLEGTLLKPNMVTPGHACPIK |
| CAT | CATA\_HUMAN | Catalase | 460, 460 | 459 | CAT | CATA\_HUMAN | Catalase | 2C(125.0477) 2C(31.9988) | NaN | X | NaN | X | NaN | NaN | 1.424396e+06 | NaN | 1.424396e+06 | NaN | LCENIAGHLK |
| YWHAQ | 1433T\_HUMAN | 14-3-3 protein theta | 134 | 128 | YWHAQ | 1433T\_HUMAN | 14-3-3 protein theta | 7C(125.0477) | NaN | X | NaN | NaN | NaN | NaN | 1.422949e+06 | 1.051343e+06 | 3.716056e+05 | NaN | YLAEVACGDDRK |
| USP5 | UBP5\_HUMAN | Ubiquitin carboxyl-terminal hydrolase 5 | 838, 838 | 837 | USP5 | UBP5\_HUMAN | Ubiquitin carboxyl-terminal hydrolase 5 | 2C(125.0477) 2C(15.9949) | NaN | X | X | NaN | NaN | NaN | 1.421042e+06 | 1.421042e+06 | NaN | NaN | VCASEKPPK |
| CD59 | CD59\_HUMAN | CD59 glycoprotein | 70 | 67 | CD59 | CD59\_HUMAN | CD59 glycoprotein | 4C(125.0477) | NaN | X | NaN | NaN | NaN | NaN | 1.419813e+06 | NaN | 1.419813e+06 | NaN | FEHCNFNDVTTR |
| KRT2 | K22E\_HUMAN | Keratin, type II cytoskeletal 2 epidermal | 489 | 483 | KRT2 | K22E\_HUMAN | Keratin, type II cytoskeletal 2 epidermal | 7C(125.0477) | NaN | X | NaN | NaN | NaN | NaN | 1.417409e+06 | 1.449719e+05 | 9.396749e+05 | 332761.7200 | LLEGEECR |
| MYH9 | MYH9\_HUMAN | Myosin-9 | 988 | 975 | MYH9 | MYH9\_HUMAN | Myosin-9 | 14C(125.0477) | NaN | X | NaN | NaN | NaN | NaN | 1.416884e+06 | 1.416884e+06 | NaN | NaN | KLEEEQIILEDQNCK |
| ANK1 | ANK1\_HUMAN | Ankyrin-1 | 1258, 1258, 1260 | 1258 | ANK1 | ANK1\_HUMAN | Ankyrin-1 | 1C(125.0477) 1C(31.9988) 3C(15.9949) | NaN | X | X | X | NaN | NaN | 1.415141e+06 | 1.415141e+06 | NaN | NaN | CYCMTDDKVDK |
| GFUS | FCL\_HUMAN | GDP-L-fucose synthase | 309 | 307 | GFUS | FCL\_HUMAN | GDP-L-fucose synthase | 3C(125.0477) | NaN | X | NaN | NaN | NaN | NaN | 1.414866e+06 | 1.414866e+06 | NaN | NaN | ETCAWFTDNYEQAR |
| VCP | TERA\_HUMAN | Transitional endoplasmic reticulum ATPase | 105 | 94 | VCP | TERA\_HUMAN | Transitional endoplasmic reticulum ATPase | 12C(125.0477) | NaN | X | NaN | NaN | NaN | NaN | 1.401102e+06 | 1.401102e+06 | NaN | NaN | VRLGDVISIQPCPDVK |
| DCXR | DCXR\_HUMAN | L-xylulose reductase | 138, 150 | 126 | DCXR | DCXR\_HUMAN | L-xylulose reductase | 13C(15.9949) 25C(15.9949) | NaN | NaN | X | NaN | NaN | NaN | 1.400885e+06 | 1.400885e+06 | NaN | NaN | GVPGAIVNVSSQCSQRAVTNHSVYCSTK |
| SPTB | SPTB1\_HUMAN | Spectrin beta chain, erythrocytic | 603 | 599 | SPTB | SPTB1\_HUMAN | Spectrin beta chain, erythrocytic | 5C(125.0477) | NaN | X | NaN | NaN | NaN | NaN | 1.399352e+06 | 1.399352e+06 | NaN | NaN | GYQPCDPQVIQDR |
| PEBP1 | PEBP1\_HUMAN | Phosphatidylethanolamine-binding protein 1 | 168, 168, 168 | 162 | PEBP1 | PEBP1\_HUMAN | Phosphatidylethanolamine-binding protein 1 | 7C(125.0477) 7C(31.9988) 7C(57.0215) | X | X | NaN | X | NaN | NaN | 1.388938e+06 | 1.388938e+06 | NaN | NaN | APVAGTCYQAEWDDYVPK |
| LDHB | LDHB\_HUMAN | L-lactate dehydrogenase B chain | 36, 36, 36 | 24 | LDHB | LDHB\_HUMAN | L-lactate dehydrogenase B chain | 13C(-34.0809) 13C(125.0477) 13C(57.0215) | X | X | NaN | NaN | X | NaN | 1.385228e+06 | 1.385228e+06 | NaN | NaN | ITVVGVGQVGMACAISILGK |
| PAICS | PUR6\_HUMAN | Multifunctional protein ADE2 | 63, 63 | 60 | PAICS | PUR6\_HUMAN | Multifunctional protein ADE2 | 4C(125.0477) 4C(31.9988) | NaN | X | NaN | X | NaN | NaN | 1.382894e+06 | 1.382894e+06 | NaN | NaN | ITSCIFQLLQEAGIK |
| ALDH1A1 | AL1A1\_HUMAN | Aldehyde dehydrogenase 1A1 | 50 | 37 | ALDH1A1 | AL1A1\_HUMAN | Aldehyde dehydrogenase 1A1 | 14C(125.0477) | NaN | X | NaN | NaN | NaN | NaN | 1.379411e+06 | 1.372153e+06 | 7.258041e+03 | NaN | KFPVFNPATEEELCQVEEGDKEDVDK |
| ANK1 | ANK1\_HUMAN | Ankyrin-1 | 1070, 1070, 1070 | 1069 | ANK1 | ANK1\_HUMAN | Ankyrin-1 | 2C(125.0477) 2C(31.9988) 2C(57.0215) | X | X | NaN | X | NaN | NaN | 1.367667e+06 | 1.367667e+06 | NaN | NaN | LCQDYDTIGPEGGSLK |
| ALDOA | ALDOA\_HUMAN | Fructose-bisphosphate aldolase A | 73, 73 | 70 | ALDOA | ALDOA\_HUMAN | Fructose-bisphosphate aldolase A | 4C(125.0477) 4C(15.9949) | NaN | X | X | NaN | NaN | NaN | 1.353571e+06 | 1.353571e+06 | NaN | NaN | VNPCIGGVILFHETLYQK |
| DERA | DEOC\_HUMAN | Deoxyribose-phosphate aldolase | 100 | 93 | DERA | DEOC\_HUMAN | Deoxyribose-phosphate aldolase | 8C(125.0477) | NaN | X | NaN | NaN | NaN | NaN | 1.352202e+06 | 1.337181e+06 | 1.502161e+04 | NaN | GITTAAVCVYPAR |
| HSPA8 | HSP7C\_HUMAN | Heat shock cognate 71 kDa protein | 17 | 4 | HSPA8 | HSP7C\_HUMAN | Heat shock cognate 71 kDa protein | 14C(125.0477) | NaN | X | NaN | NaN | NaN | NaN | 1.349863e+06 | 1.341631e+06 | 8.231568e+03 | NaN | GPAVGIDLGTTYSCVGVFQHGK |
| MAN2C1 | MA2C1\_HUMAN | Alpha-mannosidase 2C1 | 29 | 20 | MAN2C1 | MA2C1\_HUMAN | Alpha-mannosidase 2C1 | 10C(125.0477) | NaN | X | NaN | NaN | NaN | NaN | 1.347478e+06 | 1.347478e+06 | NaN | NaN | FVSPLYFTDCNLR |
| AKR1A1 | AK1A1\_HUMAN | Aldo-keto reductase family 1 member A1 | 134 | 128 | AKR1A1 | AK1A1\_HUMAN | Aldo-keto reductase family 1 member A1 | 7C(125.0477) | NaN | X | NaN | NaN | NaN | NaN | 1.346429e+06 | 1.320394e+06 | 2.603542e+04 | NaN | NADGTICYDSTHYK |
| OLA1 | OLA1\_HUMAN | Obg-like ATPase 1 | 75 | 64 | OLA1 | OLA1\_HUMAN | Obg-like ATPase 1 | 12C(125.0477) | NaN | X | NaN | NaN | NaN | NaN | 1.343442e+06 | 1.343442e+06 | NaN | NaN | VPVPDERFDFLCQYHKPASK |
| PEBP1 | PEBP1\_HUMAN | Phosphatidylethanolamine-binding protein 1 | 133 | 133 | PEBP1 | PEBP1\_HUMAN | Phosphatidylethanolamine-binding protein 1 | 1C(125.0477) | NaN | X | NaN | NaN | NaN | NaN | 1.341392e+06 | 7.879607e+05 | 5.534312e+05 | NaN | CDEPILSNR |
| CCT3 | TCPG\_HUMAN | T-complex protein 1 subunit gamma | 455 | 450 | CCT3 | TCPG\_HUMAN | T-complex protein 1 subunit gamma | 6C(125.0477) | NaN | X | NaN | NaN | NaN | NaN | 1.337561e+06 | 1.221037e+06 | 1.165243e+05 | NaN | TLIQNCGASTIR |
| CSE1L | XPO2\_HUMAN | Exportin-2 | 344 | 332 | CSE1L | XPO2\_HUMAN | Exportin-2 | 13C(125.0477) | NaN | X | NaN | NaN | NaN | NaN | 1.334775e+06 | 1.334775e+06 | NaN | NaN | NLFEDQNTLTSICEK |
| ALDH9A1 | AL9A1\_HUMAN | 4-trimethylaminobutyraldehyde dehydrogenase | 355 | 353 | ALDH9A1 | AL9A1\_HUMAN | 4-trimethylaminobutyraldehyde dehydrogenase | 3C(125.0477) | NaN | X | NaN | NaN | NaN | NaN | 1.334094e+06 | 1.328865e+06 | 5.228657e+03 | NaN | VLCGGDIYVPEDPK |
| ALDOA | ALDOA\_HUMAN | Fructose-bisphosphate aldolase A | 290, 290 | 290 | ALDOA | ALDOA\_HUMAN | Fructose-bisphosphate aldolase A | 1C(125.0477) 1C(31.9988) | NaN | X | NaN | X | NaN | NaN | 1.331675e+06 | 1.331675e+06 | NaN | NaN | CPLLKPWALTFSYGR |
| GBF1 | GBF1\_HUMAN | Golgi-specific brefeldin A-resistance guanine nucleotide exchange factor 1 | 1110 | 1106 | GBF1 | GBF1\_HUMAN | Golgi-specific brefeldin A-resistance guanine nucleotide exchange factor 1 | 5C(31.9988) | NaN | NaN | NaN | X | NaN | NaN | 1.331454e+06 | NaN | 1.331454e+06 | NaN | VALECIK |
| SYNCRIP | HNRPQ\_HUMAN | Heterogeneous nuclear ribonucleoprotein Q | 289 | 287 | SYNCRIP | HNRPQ\_HUMAN | Heterogeneous nuclear ribonucleoprotein Q | 3C(125.0477) | NaN | X | NaN | NaN | NaN | NaN | 1.329396e+06 | 1.329396e+06 | NaN | NaN | GFCFLEYEDHK |
| IPO5 | IPO5\_HUMAN | Importin-5 | 560, 560 | 557 | IPO5 | IPO5\_HUMAN | Importin-5 | 4C(125.0477) 4C(31.9988) | NaN | X | NaN | X | NaN | NaN | 1.328726e+06 | 1.328726e+06 | NaN | NaN | TIECISLIGLAVGK |
| AP2B1 | AP2B1\_HUMAN | AP-2 complex subunit beta | 95 | 94 | AP2B1 | AP2B1\_HUMAN | AP-2 complex subunit beta | 2C(125.0477) | NaN | X | NaN | NaN | NaN | NaN | 1.327373e+06 | NaN | 1.327373e+06 | NaN | DCEDPNPLIR |
| AK1 | KAD1\_HUMAN | Adenylate kinase isoenzyme 1 | 187 | 173 | AK1 | KAD1\_HUMAN | Adenylate kinase isoenzyme 1 | 15C(125.0477) | NaN | X | NaN | NaN | NaN | NaN | 1.325691e+06 | 1.325691e+06 | NaN | NaN | VNAEGSVDSVFSQVCTHLDALK |
| CMBL | CMBL\_HUMAN | Carboxymethylenebutenolidase homolog | 222 | 220 | CMBL | CMBL\_HUMAN | Carboxymethylenebutenolidase homolog | 3C(125.0477) | NaN | X | NaN | NaN | NaN | NaN | 1.324383e+06 | 1.324383e+06 | NaN | NaN | EDCSPADKPYIDEAR |
| PFKL | PFKAL\_HUMAN | ATP-dependent 6-phosphofructokinase, liver type | 708 | 699 | PFKL | PFKAL\_HUMAN | ATP-dependent 6-phosphofructokinase, liver type | 10C(125.0477) | NaN | X | NaN | NaN | NaN | NaN | 1.322519e+06 | 1.230876e+06 | 9.164373e+04 | NaN | VFANAPDSACVIGLK |
| CUL1 | CUL1\_HUMAN | Cullin-1 | 355 | 355 | CUL1 | CUL1\_HUMAN | Cullin-1 | 1C(125.0477) | NaN | X | NaN | NaN | NaN | NaN | 1.307910e+06 | 1.223565e+06 | 8.434495e+04 | NaN | CGEAALNDPK |
| GAPVD1 | GAPD1\_HUMAN | GTPase-activating protein and VPS9 domain-containing protein 1 | 1129 | 1126 | GAPVD1 | GAPD1\_HUMAN | GTPase-activating protein and VPS9 domain-containing protein 1 | 4C(125.0477) | NaN | X | NaN | NaN | NaN | NaN | 1.290799e+06 | 1.290799e+06 | NaN | NaN | LALCSADSVAFPVLTHSTR |
| AGO2 | AGO2\_HUMAN | Protein argonaute-2 | 344 | 336 | AGO2 | AGO2\_HUMAN | Protein argonaute-2 | 9C(125.0477) | NaN | X | NaN | NaN | NaN | NaN | 1.290155e+06 | 1.290155e+06 | NaN | NaN | HTYLPLEVCNIVAGQR |
| CUL2 | CUL2\_HUMAN | Cullin-2 | 266 | 262 | CUL2 | CUL2\_HUMAN | Cullin-2 | 5C(125.0477) | NaN | X | NaN | NaN | NaN | NaN | 1.290151e+06 | 1.290151e+06 | NaN | NaN | VIHECQQR |
| PSMD2 | PSMD2\_HUMAN | 26S proteasome non-ATPase regulatory subunit 2 | 251 | 251 | PSMD2 | PSMD2\_HUMAN | 26S proteasome non-ATPase regulatory subunit 2 | 1C(125.0477) | NaN | X | NaN | NaN | NaN | NaN | 1.286338e+06 | 1.255130e+06 | 3.120726e+04 | NaN | CALGVFR |
| BSG | BASI\_HUMAN | Basigin | 242 | 228 | BSG | BASI\_HUMAN | Basigin | 15C(125.0477) | NaN | X | NaN | NaN | NaN | NaN | 1.277647e+06 | NaN | 1.277647e+06 | NaN | SSEHINEGETAMLVCK |
| VAC14 | VAC14\_HUMAN | Protein VAC14 homolog | 719 | 717 | VAC14 | VAC14\_HUMAN | Protein VAC14 homolog | 3C(125.0477) | NaN | X | NaN | NaN | NaN | NaN | 1.272202e+06 | 1.272202e+06 | NaN | NaN | LQCVPNPELLQTEDSLK |
| SPTA1 | SPTA1\_HUMAN | Spectrin alpha chain, erythrocytic 1 | 1568 | 1552 | SPTA1 | SPTA1\_HUMAN | Spectrin alpha chain, erythrocytic 1 | 17C(15.9949) | NaN | NaN | X | NaN | NaN | NaN | 1.269705e+06 | NaN | 1.269705e+06 | NaN | SEQVHGVINLGNSLIECSACDGNEEAMK |
| ACAT2 | THIC\_HUMAN | Acetyl-CoA acetyltransferase, cytosolic | 65 | 45 | ACAT2 | THIC\_HUMAN | Acetyl-CoA acetyltransferase, cytosolic | 21C(125.0477) | NaN | X | NaN | NaN | NaN | NaN | 1.266145e+06 | 1.266145e+06 | NaN | NaN | ATVAPEDVSEVIFGHVLAAGCGQNPVR |
| CALR | CALR\_HUMAN | Calreticulin | 105 | 99 | CALR | CALR\_HUMAN | Calreticulin | 7C(125.0477) | NaN | X | NaN | NaN | NaN | NaN | 1.258767e+06 | 1.023486e+05 | 1.156418e+06 | NaN | HEQNIDCGGGYVK |
| PSMD1 | PSMD1\_HUMAN | 26S proteasome non-ATPase regulatory subunit 1 | 703 | 685 | PSMD1 | PSMD1\_HUMAN | 26S proteasome non-ATPase regulatory subunit 1 | 19C(125.0477) | NaN | X | NaN | NaN | NaN | NaN | 1.258679e+06 | 1.258679e+06 | NaN | NaN | QGALIASALIMIQQTEITCPK |
| GGCT | GGCT\_HUMAN | Gamma-glutamylcyclotransferase | 42 | 35 | GGCT | GGCT\_HUMAN | Gamma-glutamylcyclotransferase | 8C(125.0477) | NaN | X | NaN | NaN | NaN | NaN | 1.256503e+06 | 1.256503e+06 | NaN | NaN | NPSAAFFCVAR |
| PRKAR1A | KAP0\_HUMAN | cAMP-dependent protein kinase type I-alpha regulatory subunit | 362 | 358 | PRKAR1A | KAP0\_HUMAN | cAMP-dependent protein kinase type I-alpha regulatory subunit | 5C(125.0477) | NaN | X | NaN | NaN | NaN | NaN | 1.254674e+06 | 1.640946e+04 | 1.238265e+06 | NaN | VLGPCSDILK |
| EPB42 | EPB42\_HUMAN | Protein 4.2 | 428 | 428 | EPB42 | EPB42\_HUMAN | Protein 4.2 | 1C(125.0477) | NaN | X | NaN | NaN | NaN | NaN | 1.247462e+06 | 1.212159e+05 | 1.126246e+06 | NaN | CEDITQNYK |
| PDCD6IP | PDC6I\_HUMAN | Programmed cell death 6-interacting protein | 40 | 24 | PDCD6IP | PDC6I\_HUMAN | Programmed cell death 6-interacting protein | 17C(125.0477) | NaN | X | NaN | NaN | NaN | NaN | 1.246565e+06 | 1.246565e+06 | NaN | NaN | FIQQTYPSGGEEQAQYCR |
| ARG1 | ARGI1\_HUMAN | Arginase-1 | 303 | 292 | ARG1 | ARGI1\_HUMAN | Arginase-1 | 12C(125.0477) | NaN | X | NaN | NaN | NaN | NaN | 1.246056e+06 | 1.246056e+06 | NaN | NaN | TVNTAVAITLACFGLAR |
| RAB8B | RAB8B\_HUMAN | Ras-related protein Rab-8B | 123 | 123 | RAB8B | RAB8B\_HUMAN | Ras-related protein Rab-8B | 1C(125.0477) | NaN | X | NaN | NaN | NaN | NaN | 1.234568e+06 | 1.234568e+06 | NaN | NaN | CDMNDKR |
| ARG1 | ARGI1\_HUMAN | Arginase-1 | 45 | 40 | ARG1 | ARGI1\_HUMAN | Arginase-1 | 6C(125.0477) | NaN | X | NaN | NaN | NaN | NaN | 1.231662e+06 | 1.042652e+06 | 1.890101e+05 | NaN | LKEQECDVK |
| METAP2 | MAP2\_HUMAN | Methionine aminopeptidase 2 | 121 | 111 | METAP2 | MAP2\_HUMAN | Methionine aminopeptidase 2 | 11C(125.0477) | NaN | X | NaN | NaN | NaN | NaN | 1.226386e+06 | 1.226386e+06 | NaN | NaN | VQTDPPSVPICDLYPNGVFPK |
| PRPS1 | PRPS1\_HUMAN | Ribose-phosphate pyrophosphokinase 1 | 265 | 261 | PRPS1 | PRPS1\_HUMAN | Ribose-phosphate pyrophosphokinase 1 | 5C(125.0477) | NaN | X | NaN | NaN | NaN | NaN | 1.218348e+06 | 1.218348e+06 | NaN | NaN | INNACFEAVVVTNTIPQEDK |
| ALB | ALBU\_HUMAN | Albumin | 58, 58 | 45 | ALB | ALBU\_HUMAN | Albumin | 14C(125.0477) 14C(57.0215) | X | X | NaN | NaN | NaN | NaN | 1.215888e+06 | 1.215888e+06 | NaN | NaN | ALVLIAFAQYLQQCPFEDHVK |
| PSMD13 | PSD13\_HUMAN | 26S proteasome non-ATPase regulatory subunit 13 | 182 | 179 | PSMD13 | PSD13\_HUMAN | 26S proteasome non-ATPase regulatory subunit 13 | 4C(125.0477) | NaN | X | NaN | NaN | NaN | NaN | 1.211327e+06 | 1.175960e+06 | 3.536706e+04 | NaN | FLGCVDIK |
| HUWE1 | HUWE1\_HUMAN | E3 ubiquitin-protein ligase HUWE1 | 29 | 28 | HUWE1 | HUWE1\_HUMAN | E3 ubiquitin-protein ligase HUWE1 | 2C(125.0477) | NaN | X | NaN | NaN | NaN | NaN | 1.210579e+06 | 1.210579e+06 | NaN | NaN | VCNDEQLLLELQQIK |
| KPNA6 | IMA7\_HUMAN | Importin subunit alpha-7 | 238 | 229 | KPNA6 | IMA7\_HUMAN | Importin subunit alpha-7 | 10C(125.0477) | NaN | X | NaN | NaN | NaN | NaN | 1.202636e+06 | 1.202636e+06 | NaN | NaN | NAVWALSNLCR |
| CUL1 | CUL1\_HUMAN | Cullin-1 | 170 | 169 | CUL1 | CUL1\_HUMAN | Cullin-1 | 2C(125.0477) | NaN | X | NaN | NaN | NaN | NaN | 1.194034e+06 | 1.194034e+06 | NaN | NaN | DCLFRPLNK |
| MTOR | MTOR\_HUMAN | Serine/threonine-protein kinase mTOR | 374 | 370 | MTOR | MTOR\_HUMAN | Serine/threonine-protein kinase mTOR | 5C(125.0477) | NaN | X | NaN | NaN | NaN | NaN | 1.193361e+06 | 1.193361e+06 | NaN | NaN | FDQVCQWVLK |
| ALDOA | ALDOA\_HUMAN | Fructose-bisphosphate aldolase A | 339 | 332 | ALDOA | ALDOA\_HUMAN | Fructose-bisphosphate aldolase A | 8C(125.0477) | NaN | X | NaN | NaN | NaN | NaN | 1.188453e+06 | 1.813552e+05 | 7.709562e+05 | 236142.0000 | ALANSLACQGK |
| CLTC | CLH1\_HUMAN | Clathrin heavy chain 1 | 617 | 611 | CLTC | CLH1\_HUMAN | Clathrin heavy chain 1 | 7C(125.0477) | NaN | X | NaN | NaN | NaN | NaN | 1.185552e+06 | 1.158764e+06 | 2.678795e+04 | NaN | AHIAQLCEK |
| GSTM3 | GSTM3\_HUMAN | Glutathione S-transferase Mu 3 | 178 | 178 | GSTM3 | GSTM3\_HUMAN | Glutathione S-transferase Mu 3 | 1C(125.0477) | NaN | X | NaN | NaN | NaN | NaN | 1.173672e+06 | 1.173672e+06 | NaN | NaN | CLDEFPNLK |
| UBE2O | UBE2O\_HUMAN | (E3-independent) E2 ubiquitin-conjugating enzyme | 598 | 595 | UBE2O | UBE2O\_HUMAN | (E3-independent) E2 ubiquitin-conjugating enzyme | 4C(125.0477) | NaN | X | NaN | NaN | NaN | NaN | 1.166277e+06 | 1.166277e+06 | NaN | NaN | VQSCPDPAVYGVVQSGDHIGR |
| GDI2 | GDIB\_HUMAN | Rab GDP dissociation inhibitor beta | 317, 317 | 310 | GDI2 | GDIB\_HUMAN | Rab GDP dissociation inhibitor beta | 8C(125.0477) 8C(31.9988) | NaN | X | NaN | X | NaN | NaN | 1.166052e+06 | 1.166052e+06 | NaN | NaN | NTNDANSCQIIIPQNQVNR |
| UBA1 | UBA1\_HUMAN | Ubiquitin-like modifier-activating enzyme 1 | 481, 481 | 471 | UBA1 | UBA1\_HUMAN | Ubiquitin-like modifier-activating enzyme 1 | 11C(125.0477) 11C(31.9988) | NaN | X | NaN | X | NaN | NaN | 1.163748e+06 | 1.163748e+06 | NaN | NaN | YFLVGAGAIGCELLK |
| ATP6V1B2 | VATB2\_HUMAN | V-type proton ATPase subunit B, brain isoform | 207 | 189 | ATP6V1B2 | VATB2\_HUMAN | V-type proton ATPase subunit B, brain isoform | 19C(125.0477) | NaN | X | NaN | NaN | NaN | NaN | 1.161630e+06 | 1.161630e+06 | NaN | NaN | IPIFSAAGLPHNEIAAQICR |
| USP9X | USP9X\_HUMAN | Probable ubiquitin carboxyl-terminal hydrolase FAF-X | 2293 | 2289 | USP9X | USP9X\_HUMAN | Probable ubiquitin carboxyl-terminal hydrolase FAF-X | 5C(125.0477) | NaN | X | NaN | NaN | NaN | NaN | 1.156323e+06 | 1.156323e+06 | NaN | NaN | IIEDCSNSEETVK |
| PTGES3 | TEBP\_HUMAN | Prostaglandin E synthase 3 | 40 | 36 | PTGES3 | TEBP\_HUMAN | Prostaglandin E synthase 3 | 5C(125.0477) | NaN | X | NaN | NaN | NaN | NaN | 1.153688e+06 | 1.109589e+06 | 4.409939e+04 | NaN | LTFSCLGGSDNFK |
| CCT6A | TCPZ\_HUMAN | T-complex protein 1 subunit zeta | 406 | 400 | CCT6A | TCPZ\_HUMAN | T-complex protein 1 subunit zeta | 7C(125.0477) | NaN | X | NaN | NaN | NaN | NaN | 1.152738e+06 | 1.152738e+06 | NaN | NaN | NAIDDGCVVPGAGAVEVAMAEALIK |
| EIF5A | IF5A1\_HUMAN | Eukaryotic translation initiation factor 5A-1 | 73, 73, 73 | 68 | EIF5A | IF5A1\_HUMAN | Eukaryotic translation initiation factor 5A-1 | 6C(125.0477) 6C(15.9949) 6C(31.9988) | NaN | X | X | X | NaN | NaN | 1.148165e+06 | 1.148165e+06 | NaN | NaN | KYEDICPSTHNMDVPNIK |
| METTL7A | MET7A\_HUMAN | Putative methyltransferase-like protein 7A | 79 | 72 | METTL7A | MET7A\_HUMAN | Putative methyltransferase-like protein 7A | 8C(125.0477) | NaN | X | NaN | NaN | NaN | NaN | 1.146196e+06 | 1.146196e+06 | NaN | NaN | LSLLEVGCGTGANFK |
| CCT2 | TCPB\_HUMAN | T-complex protein 1 subunit beta | 535 | 528 | CCT2 | TCPB\_HUMAN | T-complex protein 1 subunit beta | 8C(125.0477) | NaN | X | NaN | NaN | NaN | NaN | 1.144002e+06 | 1.144002e+06 | NaN | NaN | RVPDHHPC |
| LACTB2 | LACB2\_HUMAN | Endoribonuclease LACTB2 | 58 | 43 | LACTB2 | LACB2\_HUMAN | Endoribonuclease LACTB2 | 16C(125.0477) | NaN | X | NaN | NaN | NaN | NaN | 1.122047e+06 | 1.122047e+06 | NaN | NaN | ILIDTGEPAIPEYISCLK |
| TCP1 | TCPA\_HUMAN | T-complex protein 1 subunit alpha | 236 | 234 | TCP1 | TCPA\_HUMAN | T-complex protein 1 subunit alpha | 3C(125.0477) | NaN | X | NaN | NaN | NaN | NaN | 1.121096e+06 | 1.121096e+06 | NaN | NaN | IACLDFSLQK |
| ENO1 | ENOA\_HUMAN | Alpha-enolase | 119 | 106 | ENO1 | ENOA\_HUMAN | Alpha-enolase | 14C(125.0477) | NaN | X | NaN | NaN | NaN | NaN | 1.117795e+06 | 1.111726e+06 | 6.068225e+03 | NaN | FGANAILGVSLAVCK |
| ATP2A2 | AT2A2\_HUMAN | Sarcoplasmic/endoplasmic reticulum calcium ATPase 2 | 447 | 437 | ATP2A2 | AT2A2\_HUMAN | Sarcoplasmic/endoplasmic reticulum calcium ATPase 2 | 11C(125.0477) | NaN | X | NaN | NaN | NaN | NaN | 1.113806e+06 | 1.113806e+06 | NaN | NaN | VGEATETALTCLVEK |
| ANK1 | ANK1\_HUMAN | Ankyrin-1 | 41 | 34 | ANK1 | ANK1\_HUMAN | Ankyrin-1 | 8C(125.0477) | NaN | X | NaN | NaN | NaN | NaN | 1.111613e+06 | 1.111613e+06 | 0.000000e+00 | NaN | NGVDINTCNQNGLNGLHLASK |
| CFL1 | COF1\_HUMAN | Cofilin-1 | 139, 139, 139 | 133 | CFL1 | COF1\_HUMAN | Cofilin-1 | 7C(125.0477) 7C(15.9949) 7C(31.9988) | NaN | X | X | X | NaN | NaN | 1.107436e+06 | 1.107436e+06 | NaN | NaN | HELQANCYEEVK |
| SELENBP1 | SBP1\_HUMAN | Methanethiol oxidase | 57, 57, 57 | 53 | SELENBP1 | SBP1\_HUMAN | Methanethiol oxidase | 5C(125.0477) 5C(31.9988) 5C(57.0215) | X | X | NaN | X | NaN | NaN | 1.105209e+06 | 1.105209e+06 | NaN | NaN | SPQYCQVIHR |
| SPTB | SPTB1\_HUMAN | Spectrin beta chain, erythrocytic | 960, 960, 960, 963, 963 | 956 | SPTB | SPTB1\_HUMAN | Spectrin beta chain, erythrocytic | 5C(125.0477) 5C(15.9949) 5C(31.9988) 8C(15.9949) 8C(31.9988) | NaN | X | X | X | NaN | NaN | 1.103155e+06 | NaN | NaN | 1103155.0000 | VHNYCVDCEETSK |
| HBA1 | HBA\_HUMAN | Hemoglobin subunit alpha | 105, 105, 105 | 101 | HBA1 | HBA\_HUMAN | Hemoglobin subunit alpha | 5C(-34.0809) 5C(125.0477) 5C(31.9988) | NaN | X | NaN | X | X | NaN | 1.098754e+06 | NaN | NaN | 1098754.0000 | LLSHCLLVTLAAHLPAEFTPAVHASLDKFLASVSTVLTSK |
| XPO7 | XPO7\_HUMAN | Exportin-7 | 767 | 757 | XPO7 | XPO7\_HUMAN | Exportin-7 | 11C(125.0477) | NaN | X | NaN | NaN | NaN | NaN | 1.092488e+06 | 1.092488e+06 | NaN | NaN | AIELWYHDPACTTPVLK |
| CCT2 | TCPB\_HUMAN | T-complex protein 1 subunit beta | 289 | 285 | CCT2 | TCPB\_HUMAN | T-complex protein 1 subunit beta | 5C(125.0477) | NaN | X | NaN | NaN | NaN | NaN | 1.086339e+06 | 1.053166e+06 | 3.317341e+04 | NaN | HGINCFINR |
| HERC2 | HERC2\_HUMAN | E3 ubiquitin-protein ligase HERC2 | 3864, 3865, 3855 | 3849 | HERC2 | HERC2\_HUMAN | E3 ubiquitin-protein ligase HERC2 | 16C(-34.0809) 17C(31.9988) 7C(57.0215) | X | NaN | NaN | X | X | NaN | 1.079756e+06 | 1.079756e+06 | NaN | NaN | VLVALACDLELDTLPCCAETHK |
| PSMD1 | PSMD1\_HUMAN | 26S proteasome non-ATPase regulatory subunit 1 | 112 | 111 | PSMD1 | PSMD1\_HUMAN | 26S proteasome non-ATPase regulatory subunit 1 | 2C(125.0477) | NaN | X | NaN | NaN | NaN | NaN | 1.075546e+06 | 1.075546e+06 | NaN | NaN | QCVENADLPEGEKKPIDQR |
| HBQ1 | HBAT\_HUMAN | Hemoglobin subunit theta-1 | 105 | 94 | HBQ1 | HBAT\_HUMAN | Hemoglobin subunit theta-1 | 12C(125.0477) | NaN | X | NaN | NaN | NaN | NaN | 1.070781e+06 | 1.070781e+06 | NaN | NaN | VDPASFQLLGHCLLVTLAR |
| EPB41 | EPB41\_HUMAN | Protein 4.1 | 480, 480, 480 | 479 | EPB41 | EPB41\_HUMAN | Protein 4.1 | 2C(125.0477) 2C(15.9949) 2C(31.9988) | NaN | X | X | X | NaN | NaN | 1.067440e+06 | NaN | 1.067440e+06 | NaN | VCVEHHTFFR |
| CCT3 | TCPG\_HUMAN | T-complex protein 1 subunit gamma | 372 | 371 | CCT3 | TCPG\_HUMAN | T-complex protein 1 subunit gamma | 2C(125.0477) | NaN | X | NaN | NaN | NaN | NaN | 1.067177e+06 | 1.039610e+06 | 2.756691e+04 | NaN | ACTILLR |
| DNAJA4 | DNJA4\_HUMAN | DnaJ homolog subfamily A member 4 | 272 | 265 | DNAJA4 | DNJA4\_HUMAN | DnaJ homolog subfamily A member 4 | 8C(125.0477) | NaN | X | NaN | NaN | NaN | NaN | 1.062646e+06 | 1.062646e+06 | NaN | NaN | IQLSEALCGFK |
| DDT | DOPD\_HUMAN | D-dopachrome decarboxylase | 24, 24, 24, 24, 24 | 23 | DDT | DOPD\_HUMAN | D-dopachrome decarboxylase | 2C(-34.0809) 2C(125.0477) 2C(15.9949) 2C(31.9988) 2C(57.0215) | X | X | X | X | X | NaN | 1.052797e+06 | 1.052797e+06 | NaN | NaN | LCAAAASILGKPADR |
| MYH9 | MYH9\_HUMAN | Myosin-9 | 172 | 166 | MYH9 | MYH9\_HUMAN | Myosin-9 | 7C(125.0477) | NaN | X | NaN | NaN | NaN | NaN | 1.050949e+06 | 1.001542e+06 | 4.940786e+04 | NaN | EDQSILCTGESGAGK |
| SPTB | SPTB1\_HUMAN | Spectrin beta chain, erythrocytic | 112, 112, 112 | 110 | SPTB | SPTB1\_HUMAN | Spectrin beta chain, erythrocytic | 3C(125.0477) 3C(15.9949) 3C(31.9988) | NaN | X | X | X | NaN | NaN | 1.042777e+06 | NaN | NaN | 1042777.3000 | IHCLENVDK |
| GDI2 | GDIB\_HUMAN | Rab GDP dissociation inhibitor beta | 277 | 277 | GDI2 | GDIB\_HUMAN | Rab GDP dissociation inhibitor beta | 1C(15.9949) | NaN | NaN | X | NaN | NaN | NaN | 1.040880e+06 | 7.795582e+05 | 2.613213e+05 | NaN | CKQLICDPSYVK |
| IGKC | IGKC\_HUMAN | Immunoglobulin kappa constant | 27, 27 | 20 | IGKC | IGKC\_HUMAN | Immunoglobulin kappa constant | 8C(125.0477) 8C(57.0215) | X | X | NaN | NaN | NaN | NaN | 1.038623e+06 | 1.038623e+06 | NaN | NaN | SGTASVVCLLNNFYPR |
| FSCN1 | FSCN1\_HUMAN | Fascin | 260 | 248 | FSCN1 | FSCN1\_HUMAN | Fascin | 13C(125.0477) | NaN | X | NaN | NaN | NaN | NaN | 1.034677e+06 | 1.034677e+06 | NaN | NaN | VGKDELFALEQSCAQVVLQAANER |
| SPTB | SPTB1\_HUMAN | Spectrin beta chain, erythrocytic | 183, 183, 183 | 177 | SPTB | SPTB1\_HUMAN | Spectrin beta chain, erythrocytic | 7C(125.0477) 7C(15.9949) 7C(31.9988) | NaN | X | X | X | NaN | NaN | 1.031893e+06 | NaN | 1.031893e+06 | NaN | DALLLWCQMK |
| PSMD9 | PSMD9\_HUMAN | 26S proteasome non-ATPase regulatory subunit 9 | 81, 81, 81 | 77 | PSMD9 | PSMD9\_HUMAN | 26S proteasome non-ATPase regulatory subunit 9 | 5C(125.0477) 5C(15.9949) 5C(57.0215) | X | X | X | NaN | NaN | NaN | 1.031283e+06 | 1.031283e+06 | NaN | NaN | HNIICLQNDHK |
| KRT14 | K1C14\_HUMAN | Keratin, type I cytoskeletal 14 | 40 | 31 | KRT14 | K1C14\_HUMAN | Keratin, type I cytoskeletal 14 | 10C(125.0477) | NaN | X | NaN | NaN | NaN | NaN | 1.029460e+06 | NaN | 5.557041e+05 | 473755.8800 | ISSVLAGGSCR |
| HBB | HBB\_HUMAN | Hemoglobin subunit beta | 94 | 68 | HBB | HBB\_HUMAN | Hemoglobin subunit beta | 27C(125.0477) | NaN | X | NaN | NaN | NaN | NaN | 1.029168e+06 | 1.025383e+06 | 3.784865e+03 | NaN | VLGAFSDGLAHLDNLKGTFATLSELHCDK |
| EPB42 | EPB42\_HUMAN | Protein 4.2 | 398, 398, 398, 399 | 398 | EPB42 | EPB42\_HUMAN | Protein 4.2 | 1C(15.9949) 1C(31.9988) 1C(57.0215) 2C(15.9949) | X | NaN | X | X | NaN | NaN | 1.025293e+06 | NaN | NaN | 1025292.7500 | CCEDGTLELTDSNTK |
| EPB41 | EPB41\_HUMAN | Protein 4.1 | 480 | 479 | EPB41 | EPB41\_HUMAN | Protein 4.1 | 2C(125.0477) | NaN | X | NaN | NaN | NaN | NaN | 1.016391e+06 | 1.016391e+06 | NaN | NaN | VCVEHHTFFR |
| UBA6 | UBA6\_HUMAN | Ubiquitin-like modifier-activating enzyme 6 | 721 | 715 | UBA6 | UBA6\_HUMAN | Ubiquitin-like modifier-activating enzyme 6 | 7C(125.0477) | NaN | X | NaN | NaN | NaN | NaN | 1.009979e+06 | 1.009979e+06 | NaN | NaN | ALQLLHCFPLDIR |
| ATP2B4 | AT2B4\_HUMAN | Plasma membrane calcium-transporting ATPase 4 | 537 | 535 | ATP2B4 | AT2B4\_HUMAN | Plasma membrane calcium-transporting ATPase 4 | 3C(125.0477) | NaN | X | NaN | NaN | NaN | NaN | 1.006683e+06 | 1.002599e+06 | 4.084283e+03 | NaN | TECALLGFVTDLK |
| PRDX6 | PRDX6\_HUMAN | Peroxiredoxin-6 | 47, 47 | 42 | PRDX6 | PRDX6\_HUMAN | Peroxiredoxin-6 | 6C(125.0477) 6C(31.9988) | NaN | X | NaN | X | NaN | NaN | 1.005278e+06 | NaN | 1.005278e+06 | NaN | DFTPVCTTELGR |
| UBE2O | UBE2O\_HUMAN | (E3-independent) E2 ubiquitin-conjugating enzyme | 353 | 353 | UBE2O | UBE2O\_HUMAN | (E3-independent) E2 ubiquitin-conjugating enzyme | 1C(125.0477) | NaN | X | NaN | NaN | NaN | NaN | 1.004726e+06 | 9.898796e+05 | 1.484672e+04 | NaN | CLYVFPAK |
| YWHAG | 1433G\_HUMAN | 14-3-3 protein gamma | 112 | 111 | YWHAG | 1433G\_HUMAN | 14-3-3 protein gamma | 2C(125.0477) | NaN | X | NaN | NaN | NaN | NaN | 1.001415e+06 | NaN | 1.001415e+06 | NaN | NCSETQYESK |
| VCP | TERA\_HUMAN | Transitional endoplasmic reticulum ATPase | 522 | 513 | VCP | TERA\_HUMAN | Transitional endoplasmic reticulum ATPase | 10C(125.0477) | NaN | X | NaN | NaN | NaN | NaN | 1.000942e+06 | 9.003844e+05 | 1.005574e+05 | NaN | GVLFYGPPGCGK |
| COPS4 | CSN4\_HUMAN | COP9 signalosome complex subunit 4 | 378 | 373 | COPS4 | CSN4\_HUMAN | COP9 signalosome complex subunit 4 | 6C(125.0477) | NaN | X | NaN | NaN | NaN | NaN | 9.981352e+05 | 9.981352e+05 | NaN | NaN | QIQSLCFQVNNLLEK |
| CCT3 | TCPG\_HUMAN | T-complex protein 1 subunit gamma | 213 | 201 | CCT3 | TCPG\_HUMAN | T-complex protein 1 subunit gamma | 13C(125.0477) | NaN | X | NaN | NaN | NaN | NaN | 9.936980e+05 | 9.936980e+05 | NaN | NaN | VEKIPGGIIEDSCVLR |
| YWHAQ | 1433T\_HUMAN | 14-3-3 protein theta | 134 | 128 | YWHAQ | 1433T\_HUMAN | 14-3-3 protein theta | 7C(125.0477) | NaN | X | NaN | NaN | NaN | NaN | 9.930912e+05 | NaN | 9.930912e+05 | NaN | YLAEVACGDDR |
| GMPPB | GMPPB\_HUMAN | Mannose-1-phosphate guanyltransferase beta | 245 | 244 | GMPPB | GMPPB\_HUMAN | Mannose-1-phosphate guanyltransferase beta | 2C(125.0477) | NaN | X | NaN | NaN | NaN | NaN | 9.904179e+05 | 9.904179e+05 | NaN | NaN | LCSGPGIVGNVLVDPSAR |
| MPP1 | EM55\_HUMAN | 55 kDa erythrocyte membrane protein | 179 | 174 | MPP1 | EM55\_HUMAN | 55 kDa erythrocyte membrane protein | 6C(125.0477) | NaN | X | NaN | NaN | NaN | NaN | 9.873048e+05 | NaN | 9.873048e+05 | NaN | DNLIPCK |
| DNPH1 | DNPH1\_HUMAN | 2'-deoxynucleoside 5'-phosphate N-hydrolase 1 | 26 | 16 | DNPH1 | DNPH1\_HUMAN | 2'-deoxynucleoside 5'-phosphate N-hydrolase 1 | 11C(125.0477) | NaN | X | NaN | NaN | NaN | NaN | 9.851390e+05 | 9.851390e+05 | NaN | NaN | GEPGRPALYFCGSIR |
| PSMC2 | PRS7\_HUMAN | 26S proteasome regulatory subunit 7 | 270 | 269 | PSMC2 | PRS7\_HUMAN | 26S proteasome regulatory subunit 7 | 2C(125.0477) | NaN | X | NaN | NaN | NaN | NaN | 9.839653e+05 | 9.839653e+05 | NaN | NaN | ACLIFFDEIDAIGGAR |
| CUL4A | CUL4A\_HUMAN | Cullin-4A | 289 | 274 | CUL4A | CUL4A\_HUMAN | Cullin-4A | 16C(125.0477) | NaN | X | NaN | NaN | NaN | NaN | 9.833615e+05 | 9.833615e+05 | NaN | NaN | VITYLDHSTQKPLIACVEK |
| CLIC6 | CLIC6\_HUMAN | Chloride intracellular channel protein 6 | 641 | 630 | CLIC6 | CLIC6\_HUMAN | Chloride intracellular channel protein 6 | 12C(125.0477) | NaN | X | NaN | NaN | NaN | NaN | 9.781798e+05 | 9.781798e+05 | NaN | NaN | FLDGDELTLADCNLLPK |
| AKR7A2 | ARK72\_HUMAN | Aflatoxin B1 aldehyde reductase member 2 | 214 | 206 | AKR7A2 | ARK72\_HUMAN | Aflatoxin B1 aldehyde reductase member 2 | 9C(125.0477) | NaN | X | NaN | NaN | NaN | NaN | 9.719171e+05 | 9.719171e+05 | NaN | NaN | QVETELFPCLR |
| MPP1 | EM55\_HUMAN | 55 kDa erythrocyte membrane protein | 179 | 173 | MPP1 | EM55\_HUMAN | 55 kDa erythrocyte membrane protein | 7C(125.0477) | NaN | X | NaN | NaN | NaN | NaN | 9.705718e+05 | 1.150614e+05 | 8.555104e+05 | NaN | KDNLIPCK |
| UBA3 | UBA3\_HUMAN | NEDD8-activating enzyme E1 catalytic subunit | 82 | 72 | UBA3 | UBA3\_HUMAN | NEDD8-activating enzyme E1 catalytic subunit | 11C(125.0477) | NaN | X | NaN | NaN | NaN | NaN | 9.676627e+05 | 9.676627e+05 | NaN | NaN | VLVIGAGGLGCELLK |
| PSMB3 | PSB3\_HUMAN | Proteasome subunit beta type-3 | 19, 19 | 18 | PSMB3 | PSB3\_HUMAN | Proteasome subunit beta type-3 | 2C(125.0477) 2C(15.9949) | NaN | X | X | NaN | NaN | NaN | 9.647519e+05 | 9.647519e+05 | NaN | NaN | NCVAIAADRR |
| VCP | TERA\_HUMAN | Transitional endoplasmic reticulum ATPase | 572, 572 | 568 | VCP | TERA\_HUMAN | Transitional endoplasmic reticulum ATPase | 5C(125.0477) 5C(31.9988) | NaN | X | NaN | X | NaN | NaN | 9.642588e+05 | 9.642588e+05 | NaN | NaN | QAAPCVLFFDELDSIAK |
| ADRM1 | ADRM1\_HUMAN | Proteasomal ubiquitin receptor ADRM1 | 121 | 114 | ADRM1 | ADRM1\_HUMAN | Proteasomal ubiquitin receptor ADRM1 | 8C(125.0477) | NaN | X | NaN | NaN | NaN | NaN | 9.612306e+05 | 9.612306e+05 | NaN | NaN | TDQDEEHCR |
| PPP1R7 | PP1R7\_HUMAN | Protein phosphatase 1 regulatory subunit 7 | 112 | 112 | PPP1R7 | PP1R7\_HUMAN | Protein phosphatase 1 regulatory subunit 7 | 1C(125.0477) | NaN | X | NaN | NaN | NaN | NaN | 9.585309e+05 | 9.585309e+05 | NaN | NaN | CIENLEELQSLR |
| COPS6 | CSN6\_HUMAN | COP9 signalosome complex subunit 6 | 143 | 141 | COPS6 | CSN6\_HUMAN | COP9 signalosome complex subunit 6 | 3C(125.0477) | NaN | X | NaN | NaN | NaN | NaN | 9.559871e+05 | 9.441887e+05 | 1.179837e+04 | NaN | QVCEIIESPLFLK |
| ESD | ESTD\_HUMAN | S-formylglutathione hydrolase | 11 | 11 | ESD | ESTD\_HUMAN | S-formylglutathione hydrolase | 1C(15.9949) | NaN | NaN | X | NaN | NaN | NaN | 9.480232e+05 | 9.480232e+05 | NaN | NaN | CFGGLQKVFEHDSVELNCK |
| EIF2B1 | EI2BA\_HUMAN | Translation initiation factor eIF-2B subunit alpha | 115 | 111 | EIF2B1 | EI2BA\_HUMAN | Translation initiation factor eIF-2B subunit alpha | 5C(125.0477) | NaN | X | NaN | NaN | NaN | NaN | 9.333261e+05 | 9.333261e+05 | NaN | NaN | IADLCHTFIK |
| HBB | HBB\_HUMAN | Hemoglobin subunit beta | 94, 94, 94, 94 | 68 | HBB | HBB\_HUMAN | Hemoglobin subunit beta | 27C(125.0477) 27C(15.9949) 27C(31.9988) 27C(57.0215) | X | X | X | X | NaN | NaN | 9.295886e+05 | 9.295886e+05 | NaN | NaN | VLGAFSDGLAHLDNLKGTFATLSELHCDKLHVDPENFR |
| SH3GLB2 | SHLB2\_HUMAN | Endophilin-B2 | 252 | 252 | SH3GLB2 | SHLB2\_HUMAN | Endophilin-B2 | 1C(125.0477) | NaN | X | NaN | NaN | NaN | NaN | 9.293829e+05 | 8.157386e+05 | 1.136443e+05 | NaN | CLHEFVK |
| CIAO1 | CIAO1\_HUMAN | Probable cytosolic iron-sulfur protein assembly protein CIAO1 | 212 | 209 | CIAO1 | CIAO1\_HUMAN | Probable cytosolic iron-sulfur protein assembly protein CIAO1 | 4C(125.0477) | NaN | X | NaN | NaN | NaN | NaN | 9.101980e+05 | 9.101980e+05 | NaN | NaN | LASCSDDR |
| DERA | DEOC\_HUMAN | Deoxyribose-phosphate aldolase | 118 | 115 | DERA | DEOC\_HUMAN | Deoxyribose-phosphate aldolase | 4C(125.0477) | NaN | X | NaN | NaN | NaN | NaN | 9.095850e+05 | 9.095850e+05 | NaN | NaN | AAGCNIPVASVAAGFPAGQTHLK |
| ACTB | ACTB\_HUMAN | Actin, cytoplasmic 1 | 285, 285 | 285 | ACTB | ACTB\_HUMAN | Actin, cytoplasmic 1 | 1C(125.0477) 1C(15.9949) | NaN | X | X | NaN | NaN | NaN | 9.053709e+05 | NaN | 9.053709e+05 | NaN | CDVDIRK |
| RNH1 | RINI\_HUMAN | Ribonuclease inhibitor | 45 | 45 | RNH1 | RINI\_HUMAN | Ribonuclease inhibitor | 1C(125.0477) | NaN | X | NaN | NaN | NaN | NaN | 9.047137e+05 | 8.482297e+05 | 5.648399e+04 | NaN | CKDISSALR |
| MTHFD1 | C1TC\_HUMAN | C-1-tetrahydrofolate synthase, cytoplasmic | 863 | 855 | MTHFD1 | C1TC\_HUMAN | C-1-tetrahydrofolate synthase, cytoplasmic | 9C(125.0477) | NaN | X | NaN | NaN | NaN | NaN | 9.040198e+05 | 9.040198e+05 | NaN | NaN | QGFGNLPICMAK |
| PRKAR1A | KAP0\_HUMAN | cAMP-dependent protein kinase type I-alpha regulatory subunit | 18 | 17 | PRKAR1A | KAP0\_HUMAN | cAMP-dependent protein kinase type I-alpha regulatory subunit | 2C(125.0477) | NaN | X | NaN | NaN | NaN | NaN | 9.015268e+05 | NaN | 9.015268e+05 | NaN | ECELYVQK |
| FN3KRP | KT3K\_HUMAN | Ketosamine-3-kinase | 11 | 8 | FN3KRP | KT3K\_HUMAN | Ketosamine-3-kinase | 4C(15.9949) | NaN | NaN | X | NaN | NaN | NaN | 9.006774e+05 | 9.006774e+05 | NaN | NaN | ELGCSSVRATGHSGGGCISQGR |
| PTGES3 | TEBP\_HUMAN | Prostaglandin E synthase 3 | 40 | 34 | PTGES3 | TEBP\_HUMAN | Prostaglandin E synthase 3 | 7C(125.0477) | NaN | X | NaN | NaN | NaN | NaN | 8.982466e+05 | 8.982466e+05 | NaN | NaN | SKLTFSCLGGSDNFK |
| RNPEP | AMPB\_HUMAN | Aminopeptidase B | 181 | 177 | RNPEP | AMPB\_HUMAN | Aminopeptidase B | 5C(125.0477) | NaN | X | NaN | NaN | NaN | NaN | 8.968205e+05 | 8.968205e+05 | NaN | NaN | AFFPCFDTPAVK |
| ERP44 | ERP44\_HUMAN | Endoplasmic reticulum resident protein 44 | 92 | 90 | ERP44 | ERP44\_HUMAN | Endoplasmic reticulum resident protein 44 | 3C(125.0477) | NaN | X | NaN | NaN | NaN | NaN | 8.938863e+05 | 4.292014e+05 | 4.646849e+05 | NaN | VDCDQHSDIAQR |
| PRDX1 | PRDX1\_HUMAN | Peroxiredoxin-1 | 71 | 68 | PRDX1 | PRDX1\_HUMAN | Peroxiredoxin-1 | 4C(15.9949) | NaN | NaN | X | NaN | NaN | NaN | 8.812855e+05 | 8.812855e+05 | NaN | NaN | KLNCQVIGASVDSHFCHLAWVNTPK |
| BPNT1 | BPNT1\_HUMAN | 3'(2'),5'-bisphosphate nucleotidase 1 | 249 | 246 | BPNT1 | BPNT1\_HUMAN | 3'(2'),5'-bisphosphate nucleotidase 1 | 4C(125.0477) | NaN | X | NaN | NaN | NaN | NaN | 8.784815e+05 | 8.784815e+05 | NaN | NaN | WDTCAPEVILHAVGGK |
| PA2G4 | PA2G4\_HUMAN | Proliferation-associated protein 2G4 | 179 | 173 | PA2G4 | PA2G4\_HUMAN | Proliferation-associated protein 2G4 | 7C(125.0477) | NaN | X | NaN | NaN | NaN | NaN | 8.778541e+05 | 8.778541e+05 | NaN | NaN | VAHSFNCTPIEGMLSHQLK |
| SKP1 | SKP1\_HUMAN | S-phase kinase-associated protein 1 | 120 | 114 | SKP1 | SKP1\_HUMAN | S-phase kinase-associated protein 1 | 7C(125.0477) | NaN | X | NaN | NaN | NaN | NaN | 8.764409e+05 | NaN | 8.764409e+05 | NaN | GLLDVTCK |
| DCXR | DCXR\_HUMAN | L-xylulose reductase | 244 | 227 | DCXR | DCXR\_HUMAN | L-xylulose reductase | 18C(125.0477) | NaN | X | NaN | NaN | NaN | NaN | 8.759054e+05 | 8.759054e+05 | NaN | NaN | SGMTTGSTLPVEGGFWAC |
| CCT4 | TCPD\_HUMAN | T-complex protein 1 subunit delta | 379 | 376 | CCT4 | TCPD\_HUMAN | T-complex protein 1 subunit delta | 4C(125.0477) | NaN | X | NaN | NaN | NaN | NaN | 8.724588e+05 | 8.272948e+05 | 4.516398e+04 | NaN | ITGCASPGK |
| ALAD | HEM2\_HUMAN | Delta-aminolevulinic acid dehydratase | 203 | 200 | ALAD | HEM2\_HUMAN | Delta-aminolevulinic acid dehydratase | 4C(125.0477) | NaN | X | NaN | NaN | NaN | NaN | 8.699453e+05 | 8.589556e+05 | 1.098977e+04 | NaN | FASCFYGPFR |
| ALDH1A1 | AL1A1\_HUMAN | Aldehyde dehydrogenase 1A1 | 302, 276, 276 | 274 | ALDH1A1 | AL1A1\_HUMAN | Aldehyde dehydrogenase 1A1 | 29C(15.9949) 3C(125.0477) 3C(31.9988) | NaN | X | X | X | NaN | NaN | 8.691191e+05 | 8.691191e+05 | NaN | NaN | SPCIVLADADLDNAVEFAHHGVFYHQGQCCIAASR |
| PRDX6 | PRDX6\_HUMAN | Peroxiredoxin-6 | 91 | 85 | PRDX6 | PRDX6\_HUMAN | Peroxiredoxin-6 | 7C(125.0477) | NaN | X | NaN | NaN | NaN | NaN | 8.609888e+05 | NaN | 8.609888e+05 | NaN | DINAYNCEEPTEK |
| ALB | ALBU\_HUMAN | Albumin | 99, 99 | 98 | ALB | ALBU\_HUMAN | Albumin | 2C(125.0477) 2C(57.0215) | X | X | NaN | NaN | NaN | NaN | 8.596279e+05 | NaN | 8.596279e+05 | NaN | LCTVATLR |
| PSMA4 | PSA4\_HUMAN | Proteasome subunit alpha type-4 | 107, 115 | 97 | PSMA4 | PSA4\_HUMAN | Proteasome subunit alpha type-4 | 11C(-34.0809) 19C(31.9988) | NaN | NaN | NaN | X | X | NaN | 8.561241e+05 | 8.561241e+05 | NaN | NaN | YLLQYQEPIPCEQLVTALCDIK |
| C11orf54 | CK054\_HUMAN | Ester hydrolase C11orf54 | 187 | 179 | C11orf54 | CK054\_HUMAN | Ester hydrolase C11orf54 | 9C(125.0477) | NaN | X | NaN | NaN | NaN | NaN | 8.540119e+05 | 8.540119e+05 | NaN | NaN | TGPLNFVTCMR |
| PARK7 | PARK7\_HUMAN | Parkinson disease protein 7 | 106, 106 | 100 | PARK7 | PARK7\_HUMAN | Parkinson disease protein 7 | 7C(125.0477) 7C(31.9988) | NaN | X | NaN | X | NaN | NaN | 8.525694e+05 | 8.525694e+05 | NaN | NaN | GLIAAICAGPTALLAHEIGFGSK |
| RHOA | RHOA\_HUMAN | Transforming protein RhoA | 16 | 7 | RHOA | RHOA\_HUMAN | Transforming protein RhoA | 10C(125.0477) | NaN | X | NaN | NaN | NaN | NaN | 8.442062e+05 | 8.442062e+05 | NaN | NaN | KLVIVGDGACGK |
| ANK1 | ANK1\_HUMAN | Ankyrin-1 | 278 | 271 | ANK1 | ANK1\_HUMAN | Ankyrin-1 | 8C(125.0477) | NaN | X | NaN | NaN | NaN | NaN | 8.422075e+05 | 8.422075e+05 | NaN | NaN | DELTPLHCAAR |
| UBAC1 | UBAC1\_HUMAN | Ubiquitin-associated domain-containing protein 1 | 47 | 46 | UBAC1 | UBAC1\_HUMAN | Ubiquitin-associated domain-containing protein 1 | 2C(125.0477) | NaN | X | NaN | NaN | NaN | NaN | 8.404146e+05 | 8.250878e+05 | 1.532688e+04 | NaN | HCAHGSLEDPK |
| WDR1 | WDR1\_HUMAN | WD repeat-containing protein 1 | 325 | 322 | WDR1 | WDR1\_HUMAN | WD repeat-containing protein 1 | 4C(125.0477) | NaN | X | NaN | NaN | NaN | NaN | 8.390001e+05 | 8.331057e+05 | 5.894365e+03 | NaN | SIQCLTVHK |
| PARK7 | PARK7\_HUMAN | Parkinson disease protein 7 | 53, 53, 53 | 49 | PARK7 | PARK7\_HUMAN | Parkinson disease protein 7 | 5C(125.0477) 5C(15.9949) 5C(31.9988) | NaN | X | X | X | NaN | NaN | 8.373536e+05 | 8.373536e+05 | NaN | NaN | DVVICPDASLEDAKK |
| KPNA4 | IMA3\_HUMAN | Importin subunit alpha-3 | 57 | 50 | KPNA4 | IMA3\_HUMAN | Importin subunit alpha-3 | 8C(125.0477) | NaN | X | NaN | NaN | NaN | NaN | 8.366881e+05 | 8.366881e+05 | NaN | NaN | NVPHEDICEDSDIDGDYR |
| BLMH | BLMH\_HUMAN | Bleomycin hydrolase | 269 | 268 | BLMH | BLMH\_HUMAN | Bleomycin hydrolase | 2C(125.0477) | NaN | X | NaN | NaN | NaN | NaN | 8.346857e+05 | 8.346857e+05 | NaN | NaN | ICLVNDPRPQHK |
| KRT2 | K22E\_HUMAN | Keratin, type II cytoskeletal 2 epidermal | 42, 42 | 36 | KRT2 | K22E\_HUMAN | Keratin, type II cytoskeletal 2 epidermal | 7C(125.0477) 7C(31.9988) | NaN | X | NaN | X | NaN | NaN | 8.333623e+05 | NaN | NaN | 833362.3000 | STSSFSCLSR |
| CAND1 | CAND1\_HUMAN | Cullin-associated NEDD8-dissociated protein 1 | 301 | 289 | CAND1 | CAND1\_HUMAN | Cullin-associated NEDD8-dissociated protein 1 | 13C(125.0477) | NaN | X | NaN | NaN | NaN | NaN | 8.328028e+05 | 8.328028e+05 | NaN | NaN | EVYPHVSTIINICLK |
| PITHD1 | PITH1\_HUMAN | PITH domain-containing protein 1 | 14 | 14 | PITHD1 | PITH1\_HUMAN | PITH domain-containing protein 1 | 1C(125.0477) | NaN | X | NaN | NaN | NaN | NaN | 8.326974e+05 | 8.326974e+05 | NaN | NaN | CAAEREEPPEQR |
| CAP1 | CAP1\_HUMAN | Adenylyl cyclase-associated protein 1 | 375 | 367 | CAP1 | CAP1\_HUMAN | Adenylyl cyclase-associated protein 1 | 9C(125.0477) | NaN | X | NaN | NaN | NaN | NaN | 8.318001e+05 | 8.228320e+05 | 8.968112e+03 | NaN | INSITVDNCKK |
| ANK1 | ANK1\_HUMAN | Ankyrin-1 | 531, 531 | 526 | ANK1 | ANK1\_HUMAN | Ankyrin-1 | 6C(125.0477) 6C(15.9949) | NaN | X | X | NaN | NaN | NaN | 8.310566e+05 | NaN | 8.310566e+05 | NaN | EASQACMTK |
| TCP1 | TCPA\_HUMAN | T-complex protein 1 subunit alpha | 397 | 391 | TCP1 | TCPA\_HUMAN | T-complex protein 1 subunit alpha | 7C(125.0477) | NaN | X | NaN | NaN | NaN | NaN | 8.309163e+05 | 7.872556e+05 | 4.366071e+04 | NaN | SLHDALCVVK |
| RAP1B | RAP1B\_HUMAN | Ras-related protein Rap-1b | 118 | 118 | RAP1B | RAP1B\_HUMAN | Ras-related protein Rap-1b | 1C(125.0477) | NaN | X | NaN | NaN | NaN | NaN | 8.305290e+05 | 8.305290e+05 | NaN | NaN | CDLEDERVVGK |
| HGS | HGS\_HUMAN | Hepatocyte growth factor-regulated tyrosine kinase substrate | 75, 75, 75 | 74 | HGS | HGS\_HUMAN | Hepatocyte growth factor-regulated tyrosine kinase substrate | 2C(125.0477) 2C(15.9949) 2C(31.9988) | NaN | X | X | X | NaN | NaN | 8.276119e+05 | 8.276119e+05 | NaN | NaN | NCGQTVHDEVANK |
| TPI1 | TPIS\_HUMAN | Triosephosphate isomerase | 42, 42, 42 | 34 | TPI1 | TPIS\_HUMAN | Triosephosphate isomerase | 9C(125.0477) 9C(31.9988) 9C(57.0215) | X | X | NaN | X | NaN | NaN | 8.251054e+05 | 8.251054e+05 | NaN | NaN | VPADTEVVCAPPTAYIDFAR |
| RAP2B | RAP2B\_HUMAN | Ras-related protein Rap-2b | 140 | 133 | RAP2B | RAP2B\_HUMAN | Ras-related protein Rap-2b | 8C(125.0477) | NaN | X | NaN | NaN | NaN | NaN | 8.210384e+05 | 8.210384e+05 | NaN | NaN | ALAEEWSCPFMETSAK |
| DIAPH1 | DIAP1\_HUMAN | Protein diaphanous homolog 1 | 796 | 787 | DIAPH1 | DIAP1\_HUMAN | Protein diaphanous homolog 1 | 10C(125.0477) | NaN | X | NaN | NaN | NaN | NaN | 8.190163e+05 | 8.190163e+05 | NaN | NaN | LVAEDLSQDCFWTK |
| ALDH1A1 | AL1A1\_HUMAN | Aldehyde dehydrogenase 1A1 | 133 | 132 | ALDH1A1 | AL1A1\_HUMAN | Aldehyde dehydrogenase 1A1 | 2C(125.0477) | NaN | X | NaN | NaN | NaN | NaN | 8.145566e+05 | 8.145566e+05 | NaN | NaN | YCAGWADKIQGR |
| VBP1 | PFD3\_HUMAN | Prefoldin subunit 3 | 113 | 105 | VBP1 | PFD3\_HUMAN | Prefoldin subunit 3 | 9C(125.0477) | NaN | X | NaN | NaN | NaN | NaN | 8.116281e+05 | 8.116281e+05 | NaN | NaN | FLLADNLYCK |
| MTPN | MTPN\_HUMAN | Myotrophin | 83 | 58 | MTPN | MTPN\_HUMAN | Myotrophin | 26C(125.0477) | NaN | X | NaN | NaN | NaN | NaN | 8.105118e+05 | 8.105118e+05 | NaN | NaN | GADINAPDKHHITPLLSAVYEGHVSCVK |
| GBE1 | GLGB\_HUMAN | 1,4-alpha-glucan-branching enzyme | 221 | 218 | GBE1 | GLGB\_HUMAN | 1,4-alpha-glucan-branching enzyme | 4C(125.0477) | NaN | X | NaN | NaN | NaN | NaN | 8.057025e+05 | 8.057025e+05 | NaN | NaN | HFTCNVLPR |
| AP2A1 | AP2A1\_HUMAN | AP-2 complex subunit alpha-1 | 283 | 280 | AP2A1 | AP2A1\_HUMAN | AP-2 complex subunit alpha-1 | 4C(125.0477) | NaN | X | NaN | NaN | NaN | NaN | 8.054358e+05 | 2.150166e+05 | 5.904192e+05 | NaN | LVECLETVLNK |
| CDC42 | CDC42\_HUMAN | Cell division control protein 42 homolog | 105 | 97 | CDC42 | CDC42\_HUMAN | Cell division control protein 42 homolog | 9C(125.0477) | NaN | X | NaN | NaN | NaN | NaN | 8.047653e+05 | 8.047653e+05 | NaN | NaN | WVPEITHHCPK |
| PSMA1 | PSA1\_HUMAN | Proteasome subunit alpha type-1 | 92 | 90 | PSMA1 | PSA1\_HUMAN | Proteasome subunit alpha type-1 | 3C(125.0477) | NaN | X | NaN | NaN | NaN | NaN | 8.027023e+05 | 7.378029e+05 | 6.489938e+04 | NaN | QECLDSR |
| PGAM1 | PGAM1\_HUMAN | Phosphoglycerate mutase 1 | 153 | 142 | PGAM1 | PGAM1\_HUMAN | Phosphoglycerate mutase 1 | 12C(125.0477) | NaN | X | NaN | NaN | NaN | NaN | 8.000048e+05 | 8.000048e+05 | NaN | NaN | YADLTEDQLPSCESLKDTIAR |
| CRLF3 | CRLF3\_HUMAN | Cytokine receptor-like factor 3 | 313 | 308 | CRLF3 | CRLF3\_HUMAN | Cytokine receptor-like factor 3 | 6C(125.0477) | NaN | X | NaN | NaN | NaN | NaN | 7.990718e+05 | 7.990718e+05 | NaN | NaN | APTYFCGQTLTFR |
| RNH1 | RINI\_HUMAN | Ribonuclease inhibitor | 199 | 196 | RNH1 | RINI\_HUMAN | Ribonuclease inhibitor | 4C(125.0477) | NaN | X | NaN | NaN | NaN | NaN | 7.972680e+05 | 1.452383e+05 | 6.163588e+05 | 35670.8750 | DSPCQLEALK |
| PFAS | PUR4\_HUMAN | Phosphoribosylformylglycinamidine synthase | 108 | 94 | PFAS | PUR4\_HUMAN | Phosphoribosylformylglycinamidine synthase | 15C(125.0477) | NaN | X | NaN | NaN | NaN | NaN | 7.963558e+05 | 7.963558e+05 | NaN | NaN | LNFSTPTSTNIVSVCR |
| SEC13 | SEC13\_HUMAN | Protein SEC13 homolog | 187 | 182 | SEC13 | SEC13\_HUMAN | Protein SEC13 homolog | 6C(57.0215) | X | NaN | NaN | NaN | NaN | NaN | 7.960618e+05 | NaN | 7.960618e+05 | NaN | FASGGCDNLIK |
| HSP90AA1 | HS90A\_HUMAN | Heat shock protein HSP 90-alpha | 420 | 420 | HSP90AA1 | HS90A\_HUMAN | Heat shock protein HSP 90-alpha | 1C(125.0477) | NaN | X | NaN | NaN | NaN | NaN | 7.931248e+05 | 7.931248e+05 | NaN | NaN | CLELFTELAEDKENYKK |
| PPP6C | PPP6\_HUMAN | Serine/threonine-protein phosphatase 6 catalytic subunit | 262 | 252 | PPP6C | PPP6\_HUMAN | Serine/threonine-protein phosphatase 6 catalytic subunit | 11C(125.0477) | NaN | X | NaN | NaN | NaN | NaN | 7.924192e+05 | 7.924192e+05 | NaN | NaN | LVTVWSAPNYCYR |
| PDCD6IP | PDC6I\_HUMAN | Programmed cell death 6-interacting protein | 250, 250 | 249 | PDCD6IP | PDC6I\_HUMAN | Programmed cell death 6-interacting protein | 2C(125.0477) 2C(15.9949) | NaN | X | X | NaN | NaN | NaN | 7.815831e+05 | 7.815831e+05 | NaN | NaN | HCIMQANAEYHQSILAK |
| CAT | CATA\_HUMAN | Catalase | 232 | 222 | CAT | CATA\_HUMAN | Catalase | 11C(125.0477) | NaN | X | NaN | NaN | NaN | NaN | 7.764642e+05 | NaN | 7.764642e+05 | NaN | LVNANGEAVYCK |
| PPP5C | PPP5\_HUMAN | Serine/threonine-protein phosphatase 5 | 11 | 9 | PPP5C | PPP5\_HUMAN | Serine/threonine-protein phosphatase 5 | 3C(125.0477) | NaN | X | NaN | NaN | NaN | NaN | 7.740966e+05 | 7.740966e+05 | NaN | NaN | TECAEPPRDEPPADGALK |
| AKR1C3 | AK1C3\_HUMAN | Aldo-keto reductase family 1 member C3 | 242 | 227 | AKR1C3 | AK1C3\_HUMAN | Aldo-keto reductase family 1 member C3 | 16C(125.0477) | NaN | X | NaN | NaN | NaN | NaN | 7.712138e+05 | 7.712138e+05 | NaN | NaN | WVDPNSPVLLEDPVLCALAK |
| BAG5 | BAG5\_HUMAN | BAG family molecular chaperone regulator 5 | 360 | 357 | BAG5 | BAG5\_HUMAN | BAG family molecular chaperone regulator 5 | 4C(125.0477) | NaN | X | NaN | NaN | NaN | NaN | 7.710538e+05 | 7.710538e+05 | NaN | NaN | LFACEEHPSHK |
| CLTC | CLH1\_HUMAN | Clathrin heavy chain 1 | 870, 870 | 866 | CLTC | CLH1\_HUMAN | Clathrin heavy chain 1 | 5C(125.0477) 5C(31.9988) | NaN | X | NaN | X | NaN | NaN | 7.708173e+05 | 7.708173e+05 | NaN | NaN | IHEGCEEPATHNALAK |
| PSMA3 | PSA3\_HUMAN | Proteasome subunit alpha type-3 | 42 | 42 | PSMA3 | PSA3\_HUMAN | Proteasome subunit alpha type-3 | 1C(125.0477) | NaN | X | NaN | NaN | NaN | NaN | 7.672790e+05 | 6.644688e+05 | 1.028102e+05 | NaN | CKDGVVFGVEK |
| UBA3 | UBA3\_HUMAN | NEDD8-activating enzyme E1 catalytic subunit | 139 | 136 | UBA3 | UBA3\_HUMAN | NEDD8-activating enzyme E1 catalytic subunit | 4C(125.0477) | NaN | X | NaN | NaN | NaN | NaN | 7.645775e+05 | 7.645775e+05 | NaN | NaN | VPNCNVVPHFNK |
| PCBP1 | PCBP1\_HUMAN | Poly(rC)-binding protein 1 | 109 | 102 | PCBP1 | PCBP1\_HUMAN | Poly(rC)-binding protein 1 | 8C(125.0477) | NaN | X | NaN | NaN | NaN | NaN | 7.617364e+05 | 7.528144e+05 | 8.921997e+03 | NaN | LVVPATQCGSLIGK |
| DARS1 | SYDC\_HUMAN | Aspartate--tRNA ligase, cytoplasmic | 130 | 127 | DARS1 | SYDC\_HUMAN | Aspartate--tRNA ligase, cytoplasmic | 4C(125.0477) | NaN | X | NaN | NaN | NaN | NaN | 7.587018e+05 | 7.587018e+05 | NaN | NaN | IGSCTQQDVELHVQK |
| XPO7 | XPO7\_HUMAN | Exportin-7 | 66 | 54 | XPO7 | XPO7\_HUMAN | Exportin-7 | 13C(125.0477) | NaN | X | NaN | NaN | NaN | NaN | 7.577742e+05 | 7.577742e+05 | NaN | NaN | GSSSYSQLLAATCLTK |
| IPO13 | IPO13\_HUMAN | Importin-13 | 124 | 118 | IPO13 | IPO13\_HUMAN | Importin-13 | 7C(-34.0809) | NaN | NaN | NaN | NaN | X | NaN | 7.569696e+05 | 7.569696e+05 | NaN | NaN | IVLTRLCVALASLALSMMPDAWPCAVADMVR |
| AGO2 | AGO2\_HUMAN | Protein argonaute-2 | 352, 344 | 336 | AGO2 | AGO2\_HUMAN | Protein argonaute-2 | 17C(15.9949) 9C(15.9949) | NaN | NaN | X | NaN | NaN | NaN | 7.566115e+05 | 7.566115e+05 | NaN | NaN | HTYLPLEVCNIVAGQRCIK |
| ALAD | HEM2\_HUMAN | Delta-aminolevulinic acid dehydratase | 203 | 200 | ALAD | HEM2\_HUMAN | Delta-aminolevulinic acid dehydratase | 4C(125.0477) | NaN | X | NaN | NaN | NaN | NaN | 7.530574e+05 | 7.530574e+05 | NaN | NaN | FASCFYGPFRDAAK |
| USP15 | UBP15\_HUMAN | Ubiquitin carboxyl-terminal hydrolase 15 | 139, 139 | 138 | USP15 | UBP15\_HUMAN | Ubiquitin carboxyl-terminal hydrolase 15 | 2C(125.0477) 2C(15.9949) | NaN | X | X | NaN | NaN | NaN | 7.489156e+05 | 7.489156e+05 | NaN | NaN | LCENGNMNNVVTR |
| GDI2 | GDIB\_HUMAN | Rab GDP dissociation inhibitor beta | 202, 202 | 194 | GDI2 | GDIB\_HUMAN | Rab GDP dissociation inhibitor beta | 9C(125.0477) 9C(57.0215) | X | X | NaN | NaN | NaN | NaN | 7.483340e+05 | 7.483340e+05 | NaN | NaN | TDDYLDQPCYETINR |
| SERPINB6 | SPB6\_HUMAN | Serpin B6 | 350 | 349 | SERPINB6 | SPB6\_HUMAN | Serpin B6 | 2C(125.0477) | NaN | X | NaN | NaN | NaN | NaN | 7.420252e+05 | 7.420252e+05 | NaN | NaN | FCADHPFLFFIQHSK |
| USP47 | UBP47\_HUMAN | Ubiquitin carboxyl-terminal hydrolase 47 | 45 | 43 | USP47 | UBP47\_HUMAN | Ubiquitin carboxyl-terminal hydrolase 47 | 3C(125.0477) | NaN | X | NaN | NaN | NaN | NaN | 7.413118e+05 | 7.413118e+05 | NaN | NaN | VLCIIQDTTNSK |
| CCT4 | TCPD\_HUMAN | T-complex protein 1 subunit delta | 120 | 97 | CCT4 | TCPD\_HUMAN | T-complex protein 1 subunit delta | 24C(125.0477) | NaN | X | NaN | NaN | NaN | NaN | 7.412919e+05 | 7.412919e+05 | NaN | NaN | AQDIEAGDGTTSVVIIAGSLLDSCTK |
| KCMF1 | KCMF1\_HUMAN | E3 ubiquitin-protein ligase KCMF1 | 54 | 46 | KCMF1 | KCMF1\_HUMAN | E3 ubiquitin-protein ligase KCMF1 | 9C(125.0477) | NaN | X | NaN | NaN | NaN | NaN | 7.397755e+05 | 7.397755e+05 | NaN | NaN | HTTDHPMQCILTR |
| GSTZ1 | MAAI\_HUMAN | Maleylacetoacetate isomerase | 205 | 193 | GSTZ1 | MAAI\_HUMAN | Maleylacetoacetate isomerase | 13C(125.0477) | NaN | X | NaN | NaN | NaN | NaN | 7.390376e+05 | 7.390376e+05 | NaN | NaN | LLVLEAFQVSHPCR |
| SORD | DHSO\_HUMAN | Sorbitol dehydrogenase | 165 | 143 | SORD | DHSO\_HUMAN | Sorbitol dehydrogenase | 23C(125.0477) | NaN | X | NaN | NaN | NaN | NaN | 7.386690e+05 | 7.386690e+05 | NaN | NaN | LPDNVTFEEGALIEPLSVGIHACR |
| FAH | FAAA\_HUMAN | Fumarylacetoacetase | 315 | 305 | FAH | FAAA\_HUMAN | Fumarylacetoacetase | 11C(125.0477) | NaN | X | NaN | NaN | NaN | NaN | 7.372758e+05 | 7.372758e+05 | NaN | NaN | GEGMSQAATICK |
| NAE1 | ULA1\_HUMAN | NEDD8-activating enzyme E1 regulatory subunit | 153 | 139 | NAE1 | ULA1\_HUMAN | NEDD8-activating enzyme E1 regulatory subunit | 15C(125.0477) | NaN | X | NaN | NaN | NaN | NaN | 7.356629e+05 | 7.356629e+05 | NaN | NaN | LADVLWNSQIPLLICR |
| RPE | RPE\_HUMAN | Ribulose-phosphate 3-epimerase | 217 | 215 | RPE | RPE\_HUMAN | Ribulose-phosphate 3-epimerase | 3C(125.0477) | NaN | X | NaN | NaN | NaN | NaN | 7.335864e+05 | 6.899978e+05 | 4.358862e+04 | NaN | NVCSEAAQK |
| TPP2 | TPP2\_HUMAN | Tripeptidyl-peptidase 2 | 614 | 606 | TPP2 | TPP2\_HUMAN | Tripeptidyl-peptidase 2 | 9C(125.0477) | NaN | X | NaN | NaN | NaN | NaN | 7.287614e+05 | 7.287614e+05 | NaN | NaN | EGLHYTEVCGYDIASPNAGPLFR |
| VTA1 | VTA1\_HUMAN | Vacuolar protein sorting-associated protein VTA1 homolog | 155 | 149 | VTA1 | VTA1\_HUMAN | Vacuolar protein sorting-associated protein VTA1 homolog | 7C(125.0477) | NaN | X | NaN | NaN | NaN | NaN | 7.281120e+05 | 7.281120e+05 | NaN | NaN | ATYIHNCLK |
| PPP2R5D | 2A5D\_HUMAN | Serine/threonine-protein phosphatase 2A 56 kDa regulatory subunit delta isoform | 410 | 410 | PPP2R5D | 2A5D\_HUMAN | Serine/threonine-protein phosphatase 2A 56 kDa regulatory subunit delta isoform | 1C(125.0477) | NaN | X | NaN | NaN | NaN | NaN | 7.275353e+05 | 7.104646e+05 | 1.707078e+04 | NaN | CVSSPHFQVAER |
| HSPA1A | HS71A\_HUMAN | Heat shock 70 kDa protein 1A | 17 | 4 | HSPA1A | HS71A\_HUMAN | Heat shock 70 kDa protein 1A | 14C(125.0477) | NaN | X | NaN | NaN | NaN | NaN | 7.273011e+05 | 7.273011e+05 | NaN | NaN | AAAIGIDLGTTYSCVGVFQHGK |
| PSMD6 | PSMD6\_HUMAN | 26S proteasome non-ATPase regulatory subunit 6 | 266 | 258 | PSMD6 | PSMD6\_HUMAN | 26S proteasome non-ATPase regulatory subunit 6 | 9C(125.0477) | NaN | X | NaN | NaN | NaN | NaN | 7.194894e+05 | 7.194894e+05 | NaN | NaN | QYLFSLYECR |
| ALDH1A1 | AL1A1\_HUMAN | Aldehyde dehydrogenase 1A1 | 126 | 114 | ALDH1A1 | AL1A1\_HUMAN | Aldehyde dehydrogenase 1A1 | 13C(125.0477) | NaN | X | NaN | NaN | NaN | NaN | 7.167863e+05 | 7.167863e+05 | NaN | NaN | LYSNAYLNDLAGCIK |
| PAK2 | PAK2\_HUMAN | Serine/threonine-protein kinase PAK 2 | 352 | 351 | PAK2 | PAK2\_HUMAN | Serine/threonine-protein kinase PAK 2 | 2C(125.0477) | NaN | X | NaN | NaN | NaN | NaN | 7.162039e+05 | 7.162039e+05 | NaN | NaN | ECLQALEFLHANQVIHR |
| PSMD12 | PSD12\_HUMAN | 26S proteasome non-ATPase regulatory subunit 12 | 108 | 104 | PSMD12 | PSD12\_HUMAN | 26S proteasome non-ATPase regulatory subunit 12 | 5C(15.9949) | NaN | NaN | X | NaN | NaN | NaN | 7.115743e+05 | 7.115743e+05 | NaN | NaN | MVQQCCTYVEEITDLPIK |
| HBG2 | HBG2\_HUMAN | Hemoglobin subunit gamma-2 | 94, 94 | 84 | HBG2 | HBG2\_HUMAN | Hemoglobin subunit gamma-2 | 11C(125.0477) 11C(31.9988) | NaN | X | NaN | X | NaN | NaN | 7.095636e+05 | 7.095636e+05 | NaN | NaN | GTFAQLSELHCDKLHVDPENFK |
| PLSCR1 | PLS1\_HUMAN | Phospholipid scramblase 1 | 254 | 249 | PLSCR1 | PLS1\_HUMAN | Phospholipid scramblase 1 | 6C(125.0477) | NaN | X | NaN | NaN | NaN | NaN | 7.089028e+05 | NaN | 6.257596e+05 | 83143.2000 | SLDEQCVVGK |
| ESD | ESTD\_HUMAN | S-formylglutathione hydrolase | 181, 176 | 168 | ESD | ESTD\_HUMAN | S-formylglutathione hydrolase | 14C(15.9949) 9C(125.0477) | NaN | X | X | NaN | NaN | NaN | 7.039958e+05 | 7.039958e+05 | NaN | NaN | SVSAFAPICNPVLCPWGK |
| PDXP | PLPP\_HUMAN | Chronophin | 250 | 250 | PDXP | PLPP\_HUMAN | Chronophin | 1C(125.0477) | NaN | X | NaN | NaN | NaN | NaN | 7.031662e+05 | 7.031662e+05 | NaN | NaN | CGMTTVLTLTGVSR |
| PPCDC | COAC\_HUMAN | Phosphopantothenoylcysteine decarboxylase | 166 | 158 | PPCDC | COAC\_HUMAN | Phosphopantothenoylcysteine decarboxylase | 9C(125.0477) | NaN | X | NaN | NaN | NaN | NaN | 6.937412e+05 | 6.937412e+05 | NaN | NaN | AFGYVEIPCVAK |
| HEATR5B | HTR5B\_HUMAN | HEAT repeat-containing protein 5B | 643 | 637 | HEATR5B | HTR5B\_HUMAN | HEAT repeat-containing protein 5B | 7C(57.0215) | X | NaN | NaN | NaN | NaN | NaN | 6.933008e+05 | 6.933008e+05 | NaN | NaN | LMTPIECAMTMMSHIPSVMKAHGAHLK |
| CRLF3 | CRLF3\_HUMAN | Cytokine receptor-like factor 3 | 95 | 74 | CRLF3 | CRLF3\_HUMAN | Cytokine receptor-like factor 3 | 22C(125.0477) | NaN | X | NaN | NaN | NaN | NaN | 6.920386e+05 | 6.920386e+05 | NaN | NaN | LVTLLQEVDTIEQETIKPLDDCQK |
| HUWE1 | HUWE1\_HUMAN | E3 ubiquitin-protein ligase HUWE1 | 612 | 599 | HUWE1 | HUWE1\_HUMAN | E3 ubiquitin-protein ligase HUWE1 | 14C(125.0477) | NaN | X | NaN | NaN | NaN | NaN | 6.890594e+05 | 6.890594e+05 | NaN | NaN | EVLGSLPNVFSALCLNAR |
| XPO7 | XPO7\_HUMAN | Exportin-7 | 331 | 314 | XPO7 | XPO7\_HUMAN | Exportin-7 | 18C(125.0477) | NaN | X | NaN | NaN | NaN | NaN | 6.876903e+05 | 6.876903e+05 | NaN | NaN | ILENPQSLSDPNNYHEFCR |
| DDI2 | DDI2\_HUMAN | Protein DDI1 homolog 2 | 34 | 32 | DDI2 | DDI2\_HUMAN | Protein DDI1 homolog 2 | 3C(125.0477) | NaN | X | NaN | NaN | NaN | NaN | 6.856810e+05 | 6.856810e+05 | NaN | NaN | ALCELESGIPAAESQIVYAERPLTDNHR |
| PITHD1 | PITH1\_HUMAN | PITH domain-containing protein 1 | 187, 187, 187 | 182 | PITHD1 | PITH1\_HUMAN | PITH domain-containing protein 1 | 6C(125.0477) 6C(31.9988) 6C(57.0215) | X | X | NaN | X | NaN | NaN | 6.841246e+05 | 6.841246e+05 | NaN | NaN | HEVTICNYEASANPADHR |
| SKP1 | SKP1\_HUMAN | S-phase kinase-associated protein 1 | 62 | 58 | SKP1 | SKP1\_HUMAN | S-phase kinase-associated protein 1 | 5C(125.0477) | NaN | X | NaN | NaN | NaN | NaN | 6.829009e+05 | 6.829009e+05 | NaN | NaN | VIQWCTHHKDDPPPPEDDENKEK |
| FSCN1 | FSCN1\_HUMAN | Fascin | 80 | 69 | FSCN1 | FSCN1\_HUMAN | Fascin | 12C(125.0477) | NaN | X | NaN | NaN | NaN | NaN | 6.813294e+05 | 6.813294e+05 | NaN | NaN | YLAADKDGNVTCER |
| GCLC | GSH1\_HUMAN | Glutamate--cysteine ligase catalytic subunit | 142, 142 | 129 | GCLC | GSH1\_HUMAN | Glutamate--cysteine ligase catalytic subunit | 14C(125.0477) 14C(15.9949) | NaN | X | X | NaN | NaN | NaN | 6.788132e+05 | 6.788132e+05 | NaN | NaN | KEATSILEENQALCTITSFPR |
| HUWE1 | HUWE1\_HUMAN | E3 ubiquitin-protein ligase HUWE1 | 19 | 11 | HUWE1 | HUWE1\_HUMAN | E3 ubiquitin-protein ligase HUWE1 | 9C(125.0477) | NaN | X | NaN | NaN | NaN | NaN | 6.728354e+05 | 6.545083e+05 | 1.832710e+04 | NaN | TPTEAPADCR |
| ANKFY1 | ANFY1\_HUMAN | Rabankyrin-5 | 995 | 990 | ANKFY1 | ANFY1\_HUMAN | Rabankyrin-5 | 6C(125.0477) | NaN | X | NaN | NaN | NaN | NaN | 6.722378e+05 | 6.722378e+05 | NaN | NaN | VLLTECTVDAEAFNLR |
| PSMD2 | PSMD2\_HUMAN | 26S proteasome non-ATPase regulatory subunit 2 | 459 | 457 | PSMD2 | PSMD2\_HUMAN | 26S proteasome non-ATPase regulatory subunit 2 | 3C(125.0477) | NaN | X | NaN | NaN | NaN | NaN | 6.709584e+05 | 6.709584e+05 | NaN | NaN | NECDPALALLSDYVLHNSNTMR |
| RANGAP1 | RAGP1\_HUMAN | Ran GTPase-activating protein 1 | 169 | 161 | RANGAP1 | RAGP1\_HUMAN | Ran GTPase-activating protein 1 | 9C(125.0477) | NaN | X | NaN | NaN | NaN | NaN | 6.662827e+05 | 6.467504e+05 | 1.953227e+04 | NaN | ILAAALTECHR |
| PSMD8 | PSMD8\_HUMAN | 26S proteasome non-ATPase regulatory subunit 8 | 112 | 112 | PSMD8 | PSMD8\_HUMAN | 26S proteasome non-ATPase regulatory subunit 8 | 1C(125.0477) | NaN | X | NaN | NaN | NaN | NaN | 6.645722e+05 | 6.645722e+05 | NaN | NaN | CGEELGR |
| RAN | RAN\_HUMAN | GTP-binding nuclear protein Ran | 120, 120, 112, 112, 112 | 111 | RAN | RAN\_HUMAN | GTP-binding nuclear protein Ran | 10C(125.0477) 10C(31.9988) 2C(125.0477) 2C(31.9988) 2C(57.0215) | X | X | NaN | X | NaN | NaN | 6.621466e+05 | 6.621466e+05 | NaN | NaN | VCENIPIVLCGNK |
| TGM2 | TGM2\_HUMAN | Protein-glutamine gamma-glutamyltransferase 2 | 524, 524 | 513 | TGM2 | TGM2\_HUMAN | Protein-glutamine gamma-glutamyltransferase 2 | 12C(125.0477) 12C(31.9988) | NaN | X | NaN | X | NaN | NaN | 6.603954e+05 | 6.603954e+05 | NaN | NaN | TVSYNGILGPECGTK |
| METAP2 | MAP2\_HUMAN | Methionine aminopeptidase 2 | 135 | 132 | METAP2 | MAP2\_HUMAN | Methionine aminopeptidase 2 | 4C(125.0477) | NaN | X | NaN | NaN | NaN | NaN | 6.572107e+05 | 6.572107e+05 | NaN | NaN | GQECEYPPTQDGR |
| CCT6A | TCPZ\_HUMAN | T-complex protein 1 subunit zeta | 282 | 281 | CCT6A | TCPZ\_HUMAN | T-complex protein 1 subunit zeta | 2C(125.0477) | NaN | X | NaN | NaN | NaN | NaN | 6.569449e+05 | 6.569449e+05 | NaN | NaN | VCGDSDKGFVVINQK |
| PARK7 | PARK7\_HUMAN | Parkinson disease protein 7 | 106, 106 | 99 | PARK7 | PARK7\_HUMAN | Parkinson disease protein 7 | 8C(125.0477) 8C(15.9949) | NaN | X | X | NaN | NaN | NaN | 6.563154e+05 | 6.563154e+05 | NaN | NaN | KGLIAAICAGPTALLAHEIGFGSK |
| EPB42 | EPB42\_HUMAN | Protein 4.2 | 428 | 428 | EPB42 | EPB42\_HUMAN | Protein 4.2 | 1C(125.0477) | NaN | X | NaN | NaN | NaN | NaN | 6.559111e+05 | 6.559111e+05 | NaN | NaN | CEDITQNYKYPEGSLQEK |
| DCXR | DCXR\_HUMAN | L-xylulose reductase | 150 | 142 | DCXR | DCXR\_HUMAN | L-xylulose reductase | 9C(125.0477) | NaN | X | NaN | NaN | NaN | NaN | 6.548995e+05 | 6.390374e+05 | 1.586204e+04 | NaN | AVTNHSVYCSTK |
| PFAS | PUR4\_HUMAN | Phosphoribosylformylglycinamidine synthase | 270 | 269 | PFAS | PUR4\_HUMAN | Phosphoribosylformylglycinamidine synthase | 2C(125.0477) | NaN | X | NaN | NaN | NaN | NaN | 6.521448e+05 | 5.983608e+05 | 5.378402e+04 | NaN | FCDNSSAIQGK |
| DDI2 | DDI2\_HUMAN | Protein DDI1 homolog 2 | 332 | 330 | DDI2 | DDI2\_HUMAN | Protein DDI1 homolog 2 | 3C(125.0477) | NaN | X | NaN | NaN | NaN | NaN | 6.504224e+05 | 6.504224e+05 | NaN | NaN | HQCSIDLKK |
| ALB | ALBU\_HUMAN | Albumin | 201, 201 | 199 | ALB | ALBU\_HUMAN | Albumin | 3C(125.0477) 3C(57.0215) | X | X | NaN | NaN | NaN | NaN | 6.483430e+05 | NaN | 6.097859e+05 | 38557.1330 | AACLLPK |
| RHOA | RHOA\_HUMAN | Transforming protein RhoA | 107 | 105 | RHOA | RHOA\_HUMAN | Transforming protein RhoA | 3C(125.0477) | NaN | X | NaN | NaN | NaN | NaN | 6.482161e+05 | 6.482161e+05 | NaN | NaN | HFCPNVPIILVGNKK |
| SGTA | SGTA\_HUMAN | Small glutamine-rich tetratricopeptide repeat-containing protein alpha | 129 | 117 | SGTA | SGTA\_HUMAN | Small glutamine-rich tetratricopeptide repeat-containing protein alpha | 13C(125.0477) | NaN | X | NaN | NaN | NaN | NaN | 6.479501e+05 | 6.479501e+05 | NaN | NaN | AIELNPANAVYFCNR |
| NUTF2 | NTF2\_HUMAN | Nuclear transport factor 2 | 114 | 107 | NUTF2 | NTF2\_HUMAN | Nuclear transport factor 2 | 8C(125.0477) | NaN | X | NaN | NaN | NaN | NaN | 6.478561e+05 | 6.478561e+05 | NaN | NaN | NINDAWVCTNDMFR |
| PARK7 | PARK7\_HUMAN | Parkinson disease protein 7 | 53 | 49 | PARK7 | PARK7\_HUMAN | Parkinson disease protein 7 | 5C(125.0477) | NaN | X | NaN | NaN | NaN | NaN | 6.464145e+05 | 5.246319e+05 | 1.217825e+05 | NaN | DVVICPDASLEDAK |
| GNAI2 | GNAI2\_HUMAN | Guanine nucleotide-binding protein G(i) subunit alpha-2 | 66 | 55 | GNAI2 | GNAI2\_HUMAN | Guanine nucleotide-binding protein G(i) subunit alpha-2 | 12C(125.0477) | NaN | X | NaN | NaN | NaN | NaN | 6.459982e+05 | 5.414021e+05 | 1.045961e+05 | NaN | IIHEDGYSEEECR |
| DDAH2 | DDAH2\_HUMAN | N(G),N(G)-dimethylarginine dimethylaminohydrolase 2 | 262 | 252 | DDAH2 | DDAH2\_HUMAN | N(G),N(G)-dimethylarginine dimethylaminohydrolase 2 | 11C(125.0477) | NaN | X | NaN | NaN | NaN | NaN | 6.442129e+05 | 6.442129e+05 | NaN | NaN | LSDVTLVPVSCSELEK |
| KRT13 | K1C13\_HUMAN | Keratin, type I cytoskeletal 13 | 32 | 28 | KRT13 | K1C13\_HUMAN | Keratin, type I cytoskeletal 13 | 5C(125.0477) | NaN | X | NaN | NaN | NaN | NaN | 6.423778e+05 | NaN | 6.423778e+05 | NaN | GVSTCSTR |
| RNH1 | RINI\_HUMAN | Ribonuclease inhibitor | 38, 38 | 35 | RNH1 | RINI\_HUMAN | Ribonuclease inhibitor | 4C(125.0477) 4C(31.9988) | NaN | X | NaN | X | NaN | NaN | 6.421579e+05 | NaN | 6.421579e+05 | NaN | LDDCGLTEAR |
| LHPP | LHPP\_HUMAN | Phospholysine phosphohistidine inorganic pyrophosphate phosphatase | 53 | 52 | LHPP | LHPP\_HUMAN | Phospholysine phosphohistidine inorganic pyrophosphate phosphatase | 2C(125.0477) | NaN | X | NaN | NaN | NaN | NaN | 6.406216e+05 | 6.406216e+05 | NaN | NaN | FCTNESQK |
| ATP8A2 | AT8A2\_HUMAN | Phospholipid-transporting ATPase IB | 663 | 660 | ATP8A2 | AT8A2\_HUMAN | Phospholipid-transporting ATPase IB | 4C(57.0215) | X | NaN | NaN | NaN | NaN | NaN | 6.367916e+05 | NaN | NaN | 636791.5600 | LEECYEIIEK |
| PYCR3 | P5CR3\_HUMAN | Pyrroline-5-carboxylate reductase 3 | 129 | 123 | PYCR3 | P5CR3\_HUMAN | Pyrroline-5-carboxylate reductase 3 | 7C(125.0477) | NaN | X | NaN | NaN | NaN | NaN | 6.348269e+05 | 6.348269e+05 | NaN | NaN | VLPNLPCVVQEGAIVMAR |
| GSR | GSHR\_HUMAN | Glutathione reductase, mitochondrial | 467 | 465 | GSR | GSHR\_HUMAN | Glutathione reductase, mitochondrial | 3C(125.0477) | NaN | X | NaN | NaN | NaN | NaN | 6.336796e+05 | 6.336796e+05 | NaN | NaN | MVCANKEEK |
| CAPN1 | CAN1\_HUMAN | Calpain-1 catalytic subunit | 351 | 344 | CAPN1 | CAN1\_HUMAN | Calpain-1 catalytic subunit | 8C(125.0477) | NaN | X | NaN | NaN | NaN | NaN | 6.295240e+05 | 6.295240e+05 | NaN | NaN | EFTRLEICNLTPDALK |
| AP2A1 | AP2A1\_HUMAN | AP-2 complex subunit alpha-1 | 354 | 347 | AP2A1 | AP2A1\_HUMAN | AP-2 complex subunit alpha-1 | 8C(125.0477) | NaN | X | NaN | NaN | NaN | NaN | 6.292743e+05 | 6.292743e+05 | NaN | NaN | YLALESMCTLASSEFSHEAVK |
| ENOSF1 | ENOF1\_HUMAN | Mitochondrial enolase superfamily member 1 | 330 | 321 | ENOSF1 | ENOF1\_HUMAN | Mitochondrial enolase superfamily member 1 | 10C(125.0477) | NaN | X | NaN | NaN | NaN | NaN | 6.279119e+05 | 6.279119e+05 | NaN | NaN | ALQFLQIDSCR |
| TPI1 | TPIS\_HUMAN | Triosephosphate isomerase | 218 | 207 | TPI1 | TPIS\_HUMAN | Triosephosphate isomerase | 12C(125.0477) | NaN | X | NaN | NaN | NaN | NaN | 6.196765e+05 | NaN | 6.196765e+05 | NaN | IIYGGSVTGATCK |
| PSMD7 | PSMD7\_HUMAN | 26S proteasome non-ATPase regulatory subunit 7 | 116, 116 | 115 | PSMD7 | PSMD7\_HUMAN | 26S proteasome non-ATPase regulatory subunit 7 | 2C(125.0477) 2C(31.9988) | NaN | X | NaN | X | NaN | NaN | 6.176294e+05 | 6.176294e+05 | NaN | NaN | YCPNSVLVIIDVKPK |
| KRT2 | K22E\_HUMAN | Keratin, type II cytoskeletal 2 epidermal | 42 | 36 | KRT2 | K22E\_HUMAN | Keratin, type II cytoskeletal 2 epidermal | 7C(125.0477) | NaN | X | NaN | NaN | NaN | NaN | 6.174358e+05 | NaN | 6.174358e+05 | NaN | STSSFSCLSR |
| HBZ | HBAZ\_HUMAN | Hemoglobin subunit zeta | 105 | 101 | HBZ | HBAZ\_HUMAN | Hemoglobin subunit zeta | 5C(125.0477) | NaN | X | NaN | NaN | NaN | NaN | 6.155211e+05 | 6.155211e+05 | NaN | NaN | LLSHCLLVTLAAR |
| HUWE1 | HUWE1\_HUMAN | E3 ubiquitin-protein ligase HUWE1 | 1567 | 1560 | HUWE1 | HUWE1\_HUMAN | E3 ubiquitin-protein ligase HUWE1 | 8C(125.0477) | NaN | X | NaN | NaN | NaN | NaN | 6.143020e+05 | 6.143020e+05 | NaN | NaN | LLEVVQPCLQAAK |
| PGM2L1 | PGM2L\_HUMAN | Glucose 1,6-bisphosphate synthase | 520 | 519 | PGM2L1 | PGM2L\_HUMAN | Glucose 1,6-bisphosphate synthase | 2C(125.0477) | NaN | X | NaN | NaN | NaN | NaN | 6.133377e+05 | 6.133377e+05 | NaN | NaN | FCGTFAILHVR |
| HBB | HBB\_HUMAN | Hemoglobin subunit beta | 94, 113 | 84 | HBB | HBB\_HUMAN | Hemoglobin subunit beta | 11C(15.9949) 30C(15.9949) | NaN | NaN | X | NaN | NaN | NaN | 6.120956e+05 | NaN | 6.120956e+05 | NaN | GTFATLSELHCDKLHVDPENFRLLGNVLVCVLAHHFGK |
| WDR81 | WDR81\_HUMAN | WD repeat-containing protein 81 | 1653 | 1653 | WDR81 | WDR81\_HUMAN | WD repeat-containing protein 81 | 1C(125.0477) | NaN | X | NaN | NaN | NaN | NaN | 6.102879e+05 | 6.102879e+05 | NaN | NaN | CVAPLSSEDFFLSGSK |
| ADK | ADK\_HUMAN | Adenosine kinase | 160 | 150 | ADK | ADK\_HUMAN | Adenosine kinase | 11C(125.0477) | NaN | X | NaN | NaN | NaN | NaN | 6.094686e+05 | 6.094686e+05 | NaN | NaN | SLIANLAAANCYK |
| HUWE1 | HUWE1\_HUMAN | E3 ubiquitin-protein ligase HUWE1 | 3635 | 3629 | HUWE1 | HUWE1\_HUMAN | E3 ubiquitin-protein ligase HUWE1 | 7C(125.0477) | NaN | X | NaN | NaN | NaN | NaN | 6.038448e+05 | 6.038448e+05 | NaN | NaN | HLGYTLCK |
| AP2B1 | AP2B1\_HUMAN | AP-2 complex subunit beta | 129 | 129 | AP2B1 | AP2B1\_HUMAN | AP-2 complex subunit beta | 1C(125.0477) | NaN | X | NaN | NaN | NaN | NaN | 6.020581e+05 | 1.324672e+05 | 4.695908e+05 | NaN | CLKDEDPYVR |
| TTLL12 | TTL12\_HUMAN | Tubulin--tyrosine ligase-like protein 12 | 98 | 88 | TTLL12 | TTL12\_HUMAN | Tubulin--tyrosine ligase-like protein 12 | 11C(125.0477) | NaN | X | NaN | NaN | NaN | NaN | 6.011430e+05 | 6.011430e+05 | NaN | NaN | KQQPNPGNELCYK |
| IPO7 | IPO7\_HUMAN | Importin-7 | 736 | 725 | IPO7 | IPO7\_HUMAN | Importin-7 | 12C(125.0477) | NaN | X | NaN | NaN | NaN | NaN | 5.999108e+05 | 5.219553e+05 | 7.795548e+04 | NaN | VLTGVAGEDAECHAAK |
| NIT2 | NIT2\_HUMAN | Omega-amidase NIT2 | 44 | 38 | NIT2 | NIT2\_HUMAN | Omega-amidase NIT2 | 7C(125.0477) | NaN | X | NaN | NaN | NaN | NaN | 5.989419e+05 | 5.989419e+05 | NaN | NaN | IVSLPECFNSPYGAK |
| CAND1 | CAND1\_HUMAN | Cullin-associated NEDD8-dissociated protein 1 | 71 | 71 | CAND1 | CAND1\_HUMAN | Cullin-associated NEDD8-dissociated protein 1 | 1C(125.0477) | NaN | X | NaN | NaN | NaN | NaN | 5.975435e+05 | 4.288896e+05 | 1.686539e+05 | NaN | CLGPLVSK |
| SPTB | SPTB1\_HUMAN | Spectrin beta chain, erythrocytic | 735 | 730 | SPTB | SPTB1\_HUMAN | Spectrin beta chain, erythrocytic | 6C(31.9988) | NaN | NaN | NaN | X | NaN | NaN | 5.968866e+05 | NaN | NaN | 596886.6000 | DLAAFCKK |
| HPRT1 | HPRT\_HUMAN | Hypoxanthine-guanine phosphoribosyltransferase | 206 | 201 | HPRT1 | HPRT\_HUMAN | Hypoxanthine-guanine phosphoribosyltransferase | 6C(125.0477) | NaN | X | NaN | NaN | NaN | NaN | 5.952108e+05 | 4.391042e+05 | 1.561066e+05 | NaN | DLNHVCVISETGK |
| IGHG1 | IGHG1\_HUMAN | Immunoglobulin heavy constant gamma 1 | 27, 27, 27 | 17 | IGHG1 | IGHG1\_HUMAN | Immunoglobulin heavy constant gamma 1 | 11C(125.0477) 11C(31.9988) 11C(57.0215) | X | X | NaN | X | NaN | NaN | 5.943723e+05 | NaN | 5.943723e+05 | NaN | STSGGTAALGCLVK |
| BPGM | PMGE\_HUMAN | Bisphosphoglycerate mutase | 23 | 22 | BPGM | PMGE\_HUMAN | Bisphosphoglycerate mutase | 2C(125.0477) | NaN | X | NaN | NaN | NaN | NaN | 5.933991e+05 | 2.995682e+05 | 2.938310e+05 | NaN | FCSWVDQK |
| PLEKHF2 | PKHF2\_HUMAN | Pleckstrin homology domain-containing family F member 2 | 21 | 15 | PLEKHF2 | PKHF2\_HUMAN | Pleckstrin homology domain-containing family F member 2 | 7C(125.0477) | NaN | X | NaN | NaN | NaN | NaN | 5.900699e+05 | 5.900699e+05 | NaN | NaN | ISIVENCFGAAGQPLTIPGR |
| TRPV2 | TRPV2\_HUMAN | Transient receptor potential cation channel subfamily V member 2 | 281, 281 | 280 | TRPV2 | TRPV2\_HUMAN | Transient receptor potential cation channel subfamily V member 2 | 2C(-34.0809) 2C(15.9949) | NaN | NaN | X | NaN | X | NaN | 5.886456e+05 | NaN | 5.886456e+05 | NaN | LCPTVQLEDIR |
| PARK7 | PARK7\_HUMAN | Parkinson disease protein 7 | 53, 46 | 42 | PARK7 | PARK7\_HUMAN | Parkinson disease protein 7 | 12C(15.9949) 5C(15.9949) | NaN | NaN | X | NaN | NaN | NaN | 5.884914e+05 | 5.884914e+05 | NaN | NaN | DPVQCSRDVVICPDASLEDAKK |
| RAB10 | RAB10\_HUMAN | Ras-related protein Rab-10 | 124 | 124 | RAB10 | RAB10\_HUMAN | Ras-related protein Rab-10 | 1C(125.0477) | NaN | X | NaN | NaN | NaN | NaN | 5.862425e+05 | 5.862425e+05 | NaN | NaN | CDMDDKR |
| CLTC | CLH1\_HUMAN | Clathrin heavy chain 1 | 778 | 768 | CLTC | CLH1\_HUMAN | Clathrin heavy chain 1 | 11C(125.0477) | NaN | X | NaN | NaN | NaN | NaN | 5.853704e+05 | 5.853704e+05 | NaN | NaN | LTDQLPLIIVCDR |
| KRT13 | K1C13\_HUMAN | Keratin, type I cytoskeletal 13 | 354 | 343 | KRT13 | K1C13\_HUMAN | Keratin, type I cytoskeletal 13 | 12C(125.0477) | NaN | X | NaN | NaN | NaN | NaN | 5.849563e+05 | 1.552806e+05 | 4.296758e+05 | NaN | AGLENTVAETECR |
| SLC40A1 | S40A1\_HUMAN | Solute carrier family 40 member 1 | 553 | 550 | SLC40A1 | S40A1\_HUMAN | Solute carrier family 40 member 1 | 4C(125.0477) | NaN | X | NaN | NaN | NaN | NaN | 5.841677e+05 | NaN | 5.556706e+05 | 28497.1640 | LFACGPDAK |
| HSPA8 | HSP7C\_HUMAN | Heat shock cognate 71 kDa protein | 574 | 570 | HSPA8 | HSP7C\_HUMAN | Heat shock cognate 71 kDa protein | 5C(125.0477) | NaN | X | NaN | NaN | NaN | NaN | 5.840007e+05 | 5.840007e+05 | NaN | NaN | ILDKCNEIINWLDK |
| RAB3GAP1 | RB3GP\_HUMAN | Rab3 GTPase-activating protein catalytic subunit | 858 | 858 | RAB3GAP1 | RB3GP\_HUMAN | Rab3 GTPase-activating protein catalytic subunit | 1C(125.0477) | NaN | X | NaN | NaN | NaN | NaN | 5.838465e+05 | 4.488307e+05 | 1.350158e+05 | NaN | CEQEEEKEDLER |
| SPTA1 | SPTA1\_HUMAN | Spectrin alpha chain, erythrocytic 1 | 1847 | 1845 | SPTA1 | SPTA1\_HUMAN | Spectrin alpha chain, erythrocytic 1 | 3C(125.0477) | NaN | X | NaN | NaN | NaN | NaN | 5.783966e+05 | 4.814384e+05 | 9.695820e+04 | NaN | GDCGDTLAATQSLLMK |
| SELENBP1 | SBP1\_HUMAN | Methanethiol oxidase | 31 | 21 | SELENBP1 | SBP1\_HUMAN | Methanethiol oxidase | 11C(125.0477) | NaN | X | NaN | NaN | NaN | NaN | 5.779954e+05 | 5.779954e+05 | NaN | NaN | GPREEIVYLPCIYR |
| EHBP1L1 | EH1L1\_HUMAN | EH domain-binding protein 1-like protein 1 | 346, 346 | 340 | EHBP1L1 | EH1L1\_HUMAN | EH domain-binding protein 1-like protein 1 | 7C(125.0477) 7C(31.9988) | NaN | X | NaN | X | NaN | NaN | 5.769769e+05 | 5.769769e+05 | NaN | NaN | ETQAQACPQEGTEAHGAR |
| HUWE1 | HUWE1\_HUMAN | E3 ubiquitin-protein ligase HUWE1 | 349 | 343 | HUWE1 | HUWE1\_HUMAN | E3 ubiquitin-protein ligase HUWE1 | 7C(125.0477) | NaN | X | NaN | NaN | NaN | NaN | 5.763155e+05 | 5.763155e+05 | NaN | NaN | LSSIIDCTGTASYHGFLPVLVR |
| PPIA | PPIA\_HUMAN | Peptidyl-prolyl cis-trans isomerase A | 115 | 92 | PPIA | PPIA\_HUMAN | Peptidyl-prolyl cis-trans isomerase A | 24C(125.0477) | NaN | X | NaN | NaN | NaN | NaN | 5.712236e+05 | 5.712236e+05 | NaN | NaN | HTGPGILSMANAGPNTNGSQFFICTAK |
| CLTC | CLH1\_HUMAN | Clathrin heavy chain 1 | 918 | 912 | CLTC | CLH1\_HUMAN | Clathrin heavy chain 1 | 7C(125.0477) | NaN | X | NaN | NaN | NaN | NaN | 5.705932e+05 | 5.705932e+05 | NaN | NaN | RDPHLACVAYER |
| WNK1 | WNK1\_HUMAN | Serine/threonine-protein kinase WNK1 | 1135, 1135, 1135 | 1132 | WNK1 | WNK1\_HUMAN | Serine/threonine-protein kinase WNK1 | 4C(125.0477) 4C(15.9949) 4C(31.9988) | NaN | X | X | X | NaN | NaN | 5.703621e+05 | 5.703621e+05 | NaN | NaN | VVECQLETHNR |
| SPTA1 | SPTA1\_HUMAN | Spectrin alpha chain, erythrocytic 1 | 1203, 1203 | 1202 | SPTA1 | SPTA1\_HUMAN | Spectrin alpha chain, erythrocytic 1 | 2C(125.0477) 2C(15.9949) | NaN | X | X | NaN | NaN | NaN | 5.675238e+05 | 5.675238e+05 | NaN | NaN | KCQALSAADPGSDLFSVQALQR |
| SNX9 | SNX9\_HUMAN | Sorting nexin-9 | 502 | 498 | SNX9 | SNX9\_HUMAN | Sorting nexin-9 | 5C(125.0477) | NaN | X | NaN | NaN | NaN | NaN | 5.640462e+05 | 5.640462e+05 | NaN | NaN | GFLGCFPDIIGTHK |
| PSMD4 | PSMD4\_HUMAN | 26S proteasome non-ATPase regulatory subunit 4 | 37 | 26 | PSMD4 | PSMD4\_HUMAN | 26S proteasome non-ATPase regulatory subunit 4 | 12C(125.0477) | NaN | X | NaN | NaN | NaN | NaN | 5.638859e+05 | 5.638859e+05 | NaN | NaN | LQAQQDAVNIVCHSK |
| VCP | TERA\_HUMAN | Transitional endoplasmic reticulum ATPase | 105 | 96 | VCP | TERA\_HUMAN | Transitional endoplasmic reticulum ATPase | 10C(125.0477) | NaN | X | NaN | NaN | NaN | NaN | 5.620664e+05 | 4.056895e+05 | 1.563770e+05 | NaN | LGDVISIQPCPDVK |
| RARS1 | SYRC\_HUMAN | Arginine--tRNA ligase, cytoplasmic | 150 | 144 | RARS1 | SYRC\_HUMAN | Arginine--tRNA ligase, cytoplasmic | 7C(125.0477) | NaN | X | NaN | NaN | NaN | NaN | 5.613124e+05 | 5.613124e+05 | NaN | NaN | HLPDNECIEK |
| TNPO3 | TNPO3\_HUMAN | Transportin-3 | 303 | 298 | TNPO3 | TNPO3\_HUMAN | Transportin-3 | 6C(125.0477) | NaN | X | NaN | NaN | NaN | NaN | 5.601972e+05 | 5.601972e+05 | NaN | NaN | IFTELCETFLEK |
| ARG1 | ARGI1\_HUMAN | Arginase-1 | 45 | 40 | ARG1 | ARGI1\_HUMAN | Arginase-1 | 6C(125.0477) | NaN | X | NaN | NaN | NaN | NaN | 5.585328e+05 | 5.585328e+05 | NaN | NaN | LKEQECDVKDYGDLPFADIPNDSPFQIVK |
| PGM2 | PGM2\_HUMAN | Phosphoglucomutase-2 | 484 | 479 | PGM2 | PGM2\_HUMAN | Phosphoglucomutase-2 | 6C(125.0477) | NaN | X | NaN | NaN | NaN | NaN | 5.550728e+05 | 5.550728e+05 | NaN | NaN | ASYFICHDQETIKK |
| GSTO1 | GSTO1\_HUMAN | Glutathione S-transferase omega-1 | 192 | 189 | GSTO1 | GSTO1\_HUMAN | Glutathione S-transferase omega-1 | 4C(125.0477) | NaN | X | NaN | NaN | NaN | NaN | 5.532506e+05 | NaN | 5.532506e+05 | NaN | LNECVDHTPK |
| HBD | HBD\_HUMAN | Hemoglobin subunit delta | 113, 113, 113, 113 | 106 | HBD | HBD\_HUMAN | Hemoglobin subunit delta | 8C(-34.0809) 8C(125.0477) 8C(15.9949) 8C(31.9988) | NaN | X | X | X | X | NaN | 5.529632e+05 | 5.529632e+05 | NaN | NaN | LLGNVLVCVLARNFGK |
| ADD1 | ADDA\_HUMAN | Alpha-adducin | 253 | 253 | ADD1 | ADDA\_HUMAN | Alpha-adducin | 1C(125.0477) | NaN | X | NaN | NaN | NaN | NaN | 5.517638e+05 | 5.517638e+05 | NaN | NaN | CGLLPISPEALSLGEVAYHDYHGILVDEEEK |
| LASP1 | LASP1\_HUMAN | LIM and SH3 domain protein 1 | 53 | 50 | LASP1 | LASP1\_HUMAN | LIM and SH3 domain protein 1 | 4C(125.0477) | NaN | X | NaN | NaN | NaN | NaN | 5.511444e+05 | 5.511444e+05 | NaN | NaN | KPYCNAHYPK |
| LEPROT | OBRG\_HUMAN | Leptin receptor gene-related protein | 66 | 55 | LEPROT | OBRG\_HUMAN | Leptin receptor gene-related protein | 12C(125.0477) | NaN | X | NaN | NaN | NaN | NaN | 5.503776e+05 | NaN | 5.503776e+05 | NaN | VTYDSDATSSACR |
| PSMD1 | PSMD1\_HUMAN | 26S proteasome non-ATPase regulatory subunit 1 | 104 | 104 | PSMD1 | PSMD1\_HUMAN | 26S proteasome non-ATPase regulatory subunit 1 | 1C(125.0477) | NaN | X | NaN | NaN | NaN | NaN | 5.500456e+05 | 5.500456e+05 | NaN | NaN | CIDHYTK |
| G6PD | G6PD\_HUMAN | Glucose-6-phosphate 1-dehydrogenase | 385 | 371 | G6PD | G6PD\_HUMAN | Glucose-6-phosphate 1-dehydrogenase | 15C(125.0477) | NaN | X | NaN | NaN | NaN | NaN | 5.487831e+05 | 5.487831e+05 | NaN | NaN | LQFHDVAGDIFHQQCKR |
| USP47 | UBP47\_HUMAN | Ubiquitin carboxyl-terminal hydrolase 47 | 856 | 845 | USP47 | UBP47\_HUMAN | Ubiquitin carboxyl-terminal hydrolase 47 | 12C(125.0477) | NaN | X | NaN | NaN | NaN | NaN | 5.484184e+05 | 5.282064e+05 | 2.021196e+04 | NaN | AGGDSGNVDDDCER |
| ACTR3 | ARP3\_HUMAN | Actin-related protein 3 | 189 | 162 | ACTR3 | ARP3\_HUMAN | Actin-related protein 3 | 28C(125.0477) | NaN | X | NaN | NaN | NaN | NaN | 5.483886e+05 | 5.483886e+05 | NaN | NaN | TLTGTVIDSGDGVTHVIPVAEGYVIGSCIK |
| HUWE1 | HUWE1\_HUMAN | E3 ubiquitin-protein ligase HUWE1 | 3333 | 3315 | HUWE1 | HUWE1\_HUMAN | E3 ubiquitin-protein ligase HUWE1 | 19C(125.0477) | NaN | X | NaN | NaN | NaN | NaN | 5.483026e+05 | 5.483026e+05 | NaN | NaN | HASGGSTVHIHPQAAPVVCR |
| SPTB | SPTB1\_HUMAN | Spectrin beta chain, erythrocytic | 1962 | 1958 | SPTB | SPTB1\_HUMAN | Spectrin beta chain, erythrocytic | 5C(125.0477) | NaN | X | NaN | NaN | NaN | NaN | 5.466487e+05 | 5.121920e+05 | 3.445668e+04 | NaN | NFSACLELGESLLQR |
| FBXO7 | FBX7\_HUMAN | F-box only protein 7 | 286 | 277 | FBXO7 | FBX7\_HUMAN | F-box only protein 7 | 10C(125.0477) | NaN | X | NaN | NaN | NaN | NaN | 5.460050e+05 | 5.460050e+05 | NaN | NaN | LQLLPESFICKEK |
| SHPK | SHPK\_HUMAN | Sedoheptulokinase | 132 | 132 | SHPK | SHPK\_HUMAN | Sedoheptulokinase | 1C(125.0477) | NaN | X | NaN | NaN | NaN | NaN | 5.455954e+05 | 5.455954e+05 | NaN | NaN | CSSEFLASLPQPK |
| TLN1 | TLN1\_HUMAN | Talin-1 | 2442 | 2430 | TLN1 | TLN1\_HUMAN | Talin-1 | 13C(125.0477) | NaN | X | NaN | NaN | NaN | NaN | 5.424872e+05 | 5.424872e+05 | NaN | NaN | QVAASTAQLLVACK |
| HSPH1 | HS105\_HUMAN | Heat shock protein 105 kDa | 167 | 155 | HSPH1 | HS105\_HUMAN | Heat shock protein 105 kDa | 13C(125.0477) | NaN | X | NaN | NaN | NaN | NaN | 5.420955e+05 | 5.420955e+05 | NaN | NaN | SVLDAAQIVGLNCLR |
| AGO2 | AGO2\_HUMAN | Protein argonaute-2 | 137 | 134 | AGO2 | AGO2\_HUMAN | Protein argonaute-2 | 4C(125.0477) | NaN | X | NaN | NaN | NaN | NaN | 5.418755e+05 | 5.418755e+05 | NaN | NaN | WVSCVSLQALHDALSGR |
| ASL | ARLY\_HUMAN | Argininosuccinate lyase | 307 | 307 | ASL | ARLY\_HUMAN | Argininosuccinate lyase | 1C(125.0477) | NaN | X | NaN | NaN | NaN | NaN | 5.362942e+05 | 5.362942e+05 | NaN | NaN | CAGLLMTLK |
| GMPS | GUAA\_HUMAN | GMP synthase [glutamine-hydrolyzing] | 489 | 488 | GMPS | GUAA\_HUMAN | GMP synthase [glutamine-hydrolyzing] | 2C(125.0477) | NaN | X | NaN | NaN | NaN | NaN | 5.333464e+05 | 5.094235e+05 | 2.392294e+04 | NaN | ACTTEEDQEK |
| PPP6R1 | PP6R1\_HUMAN | Serine/threonine-protein phosphatase 6 regulatory subunit 1 | 437 | 432 | PPP6R1 | PP6R1\_HUMAN | Serine/threonine-protein phosphatase 6 regulatory subunit 1 | 6C(125.0477) | NaN | X | NaN | NaN | NaN | NaN | 5.325735e+05 | 5.325735e+05 | NaN | NaN | HLLQQCR |
| ACTR1A | ACTZ\_HUMAN | Alpha-centractin | 222 | 221 | ACTR1A | ACTZ\_HUMAN | Alpha-centractin | 2C(125.0477) | NaN | X | NaN | NaN | NaN | NaN | 5.321334e+05 | 5.321334e+05 | NaN | NaN | ACYLSINPQKDETLETEK |
| USP14 | UBP14\_HUMAN | Ubiquitin carboxyl-terminal hydrolase 14 | 257 | 257 | USP14 | UBP14\_HUMAN | Ubiquitin carboxyl-terminal hydrolase 14 | 1C(125.0477) | NaN | X | NaN | NaN | NaN | NaN | 5.311686e+05 | 5.311686e+05 | NaN | NaN | CTESEEEEVTKGK |
| SPTB | SPTB1\_HUMAN | Spectrin beta chain, erythrocytic | 112 | 110 | SPTB | SPTB1\_HUMAN | Spectrin beta chain, erythrocytic | 3C(125.0477) | NaN | X | NaN | NaN | NaN | NaN | 5.299590e+05 | 5.299590e+05 | NaN | NaN | IHCLENVDK |
| UROS | HEM4\_HUMAN | Uroporphyrinogen-III synthase | 119, 119 | 110 | UROS | HEM4\_HUMAN | Uroporphyrinogen-III synthase | 10C(125.0477) 10C(31.9988) | NaN | X | NaN | X | NaN | NaN | 5.239289e+05 | 5.239289e+05 | NaN | NaN | IGLDTEGETCGNAEK |
| CD44 | CD44\_HUMAN | CD44 antigen | 53 | 47 | CD44 | CD44\_HUMAN | CD44 antigen | 7C(125.0477) | NaN | X | NaN | NaN | NaN | NaN | 5.219986e+05 | NaN | 1.173054e+04 | 510268.0300 | TEAADLCK |
| AP2M1 | AP2M1\_HUMAN | AP-2 complex subunit mu | 337 | 324 | AP2M1 | AP2M1\_HUMAN | AP-2 complex subunit mu | 14C(125.0477) | NaN | X | NaN | NaN | NaN | NaN | 5.207316e+05 | 5.207316e+05 | NaN | NaN | IPTPLNTSGVQVICMK |
| ACSL4 | ACSL4\_HUMAN | Long-chain-fatty-acid--CoA ligase 4 | 420 | 414 | ACSL4 | ACSL4\_HUMAN | Long-chain-fatty-acid--CoA ligase 4 | 7C(125.0477) | NaN | X | NaN | NaN | NaN | NaN | 5.203991e+05 | 5.203991e+05 | NaN | NaN | GYDAPLCNLLLFK |
| PRPS1 | PRPS1\_HUMAN | Ribose-phosphate pyrophosphokinase 1 | 165, 165 | 164 | PRPS1 | PRPS1\_HUMAN | Ribose-phosphate pyrophosphokinase 1 | 2C(125.0477) 2C(15.9949) | NaN | X | X | NaN | NaN | NaN | 5.177282e+05 | 5.177282e+05 | NaN | NaN | NCTIVSPDAGGAKR |
| RNF123 | RN123\_HUMAN | E3 ubiquitin-protein ligase RNF123 | 1277 | 1276 | RNF123 | RN123\_HUMAN | E3 ubiquitin-protein ligase RNF123 | 2C(125.0477) | NaN | X | NaN | NaN | NaN | NaN | 5.174601e+05 | 5.174601e+05 | NaN | NaN | ACINQHLMNNK |
| FN3KRP | KT3K\_HUMAN | Ketosamine-3-kinase | 24, 24 | 16 | FN3KRP | KT3K\_HUMAN | Ketosamine-3-kinase | 9C(125.0477) 9C(31.9988) | NaN | X | NaN | X | NaN | NaN | 5.149139e+05 | 5.149139e+05 | NaN | NaN | ATGHSGGGCISQGR |
| GNAI3 | GNAI3\_HUMAN | Guanine nucleotide-binding protein G(i) subunit alpha-3 | 66 | 55 | GNAI3 | GNAI3\_HUMAN | Guanine nucleotide-binding protein G(i) subunit alpha-3 | 12C(125.0477) | NaN | X | NaN | NaN | NaN | NaN | 5.136369e+05 | 3.802598e+05 | 1.333770e+05 | NaN | IIHEDGYSEDECK |
| GMPR2 | GMPR2\_HUMAN | GMP reductase 2 | 348 | 336 | GMPR2 | GMPR2\_HUMAN | GMP reductase 2 | 13C(125.0477) | NaN | X | NaN | NaN | NaN | NaN | 5.116904e+05 | 5.116904e+05 | NaN | NaN | VTQQVNPIFSEAC |
| SCYL1 | SCYL1\_HUMAN | N-terminal kinase-like protein | 512 | 507 | SCYL1 | SCYL1\_HUMAN | N-terminal kinase-like protein | 6C(125.0477) | NaN | X | NaN | NaN | NaN | NaN | 5.113851e+05 | 5.113851e+05 | NaN | NaN | ILPVLCGLTVDPEK |
| FAM114A2 | F1142\_HUMAN | Protein FAM114A2 | 425 | 414 | FAM114A2 | F1142\_HUMAN | Protein FAM114A2 | 12C(125.0477) | NaN | X | NaN | NaN | NaN | NaN | 5.113202e+05 | 5.113202e+05 | NaN | NaN | SQTLSQMTIVLCK |
| ACTBL2 | ACTBL\_HUMAN | Beta-actin-like protein 2 | 53, 53, 53 | 41 | ACTBL2 | ACTBL\_HUMAN | Beta-actin-like protein 2 | 13C(125.0477) 13C(15.9949) 13C(31.9988) | NaN | X | X | X | NaN | NaN | 5.081986e+05 | NaN | 5.081986e+05 | NaN | HQGVMVGMGQKDCYVGDEAQSK |
| PSMC5 | PRS8\_HUMAN | 26S proteasome regulatory subunit 8 | 209 | 202 | PSMC5 | PRS8\_HUMAN | 26S proteasome regulatory subunit 8 | 8C(125.0477) | NaN | X | NaN | NaN | NaN | NaN | 5.081664e+05 | 5.081664e+05 | NaN | NaN | AVAHHTDCTFIR |
| UBR4 | UBR4\_HUMAN | E3 ubiquitin-protein ligase UBR4 | 1962 | 1945 | UBR4 | UBR4\_HUMAN | E3 ubiquitin-protein ligase UBR4 | 18C(125.0477) | NaN | X | NaN | NaN | NaN | NaN | 5.039149e+05 | 5.039149e+05 | NaN | NaN | LASAPVPFTVLSLTGNPCK |
| UBR4 | UBR4\_HUMAN | E3 ubiquitin-protein ligase UBR4 | 3864 | 3858 | UBR4 | UBR4\_HUMAN | E3 ubiquitin-protein ligase UBR4 | 7C(125.0477) | NaN | X | NaN | NaN | NaN | NaN | 5.027718e+05 | 5.027718e+05 | NaN | NaN | ALSVLGCGHTSSTK |
| CLTC | CLH1\_HUMAN | Clathrin heavy chain 1 | 459 | 451 | CLTC | CLH1\_HUMAN | Clathrin heavy chain 1 | 9C(125.0477) | NaN | X | NaN | NaN | NaN | NaN | 5.017341e+05 | 5.017341e+05 | NaN | NaN | WLKEDKLECSEELGDLVK |
| USP14 | UBP14\_HUMAN | Ubiquitin carboxyl-terminal hydrolase 14 | 257 | 257 | USP14 | UBP14\_HUMAN | Ubiquitin carboxyl-terminal hydrolase 14 | 1C(125.0477) | NaN | X | NaN | NaN | NaN | NaN | 5.012366e+05 | NaN | 5.012366e+05 | NaN | CTESEEEEVTK |
| PEPD | PEPD\_HUMAN | Xaa-Pro dipeptidase | 183 | 169 | PEPD | PEPD\_HUMAN | Xaa-Pro dipeptidase | 15C(125.0477) | NaN | X | NaN | NaN | NaN | NaN | 5.005485e+05 | 5.005485e+05 | NaN | NaN | FEVNNTILHPEIVECR |
| IPO9 | IPO9\_HUMAN | Importin-9 | 90 | 83 | IPO9 | IPO9\_HUMAN | Importin-9 | 8C(125.0477) | NaN | X | NaN | NaN | NaN | NaN | 5.004728e+05 | 5.004728e+05 | NaN | NaN | QYVETHWCAQSEK |
| EIF2S2 | IF2B\_HUMAN | Eukaryotic translation initiation factor 2 subunit 2 | 226 | 217 | EIF2S2 | IF2B\_HUMAN | Eukaryotic translation initiation factor 2 subunit 2 | 10C(125.0477) | NaN | X | NaN | NaN | NaN | NaN | 5.000628e+05 | 5.000628e+05 | NaN | NaN | TSFVNFTDICK |
| TPP2 | TPP2\_HUMAN | Tripeptidyl-peptidase 2 | 209, 209 | 208 | TPP2 | TPP2\_HUMAN | Tripeptidyl-peptidase 2 | 2C(125.0477) 2C(31.9988) | NaN | X | NaN | X | NaN | NaN | 4.995008e+05 | 4.995008e+05 | NaN | NaN | ACIDSNEDGDLSK |
| PRDX5 | PRDX5\_HUMAN | Peroxiredoxin-5, mitochondrial | 100 | 87 | PRDX5 | PRDX5\_HUMAN | Peroxiredoxin-5, mitochondrial | 14C(125.0477) | NaN | X | NaN | NaN | NaN | NaN | 4.977483e+05 | 4.977483e+05 | NaN | NaN | GVLFGVPGAFTPGCSK |
| SEMA7A | SEM7A\_HUMAN | Semaphorin-7A | 143 | 137 | SEMA7A | SEM7A\_HUMAN | Semaphorin-7A | 7C(125.0477) | NaN | X | NaN | NaN | NaN | NaN | 4.975884e+05 | NaN | 2.894350e+05 | 208153.4000 | SEGLLACGTNAR |
| PARK7 | PARK7\_HUMAN | Parkinson disease protein 7 | 53 | 49 | PARK7 | PARK7\_HUMAN | Parkinson disease protein 7 | 5C(125.0477) | NaN | X | NaN | NaN | NaN | NaN | 4.968058e+05 | NaN | 4.848471e+05 | 11958.7490 | DVVICPDASLEDAKK |
| HINT1 | HINT1\_HUMAN | Adenosine 5'-monophosphoramidase HINT1 | 84 | 84 | HINT1 | HINT1\_HUMAN | Adenosine 5'-monophosphoramidase HINT1 | 1C(125.0477) | NaN | X | NaN | NaN | NaN | NaN | 4.961642e+05 | 3.637609e+04 | 4.597881e+05 | NaN | CAADLGLNK |
| COPS7B | CSN7B\_HUMAN | COP9 signalosome complex subunit 7b | 110 | 110 | COPS7B | CSN7B\_HUMAN | COP9 signalosome complex subunit 7b | 1C(125.0477) | NaN | X | NaN | NaN | NaN | NaN | 4.957759e+05 | 4.957759e+05 | NaN | NaN | CIPYSVLLK |
| CAND1 | CAND1\_HUMAN | Cullin-associated NEDD8-dissociated protein 1 | 264 | 263 | CAND1 | CAND1\_HUMAN | Cullin-associated NEDD8-dissociated protein 1 | 2C(125.0477) | NaN | X | NaN | NaN | NaN | NaN | 4.918032e+05 | 4.543829e+05 | 3.742027e+04 | NaN | FCNVDDDELR |
| RRM1 | RIR1\_HUMAN | Ribonucleoside-diphosphate reductase large subunit | 662 | 657 | RRM1 | RIR1\_HUMAN | Ribonucleoside-diphosphate reductase large subunit | 6C(125.0477) | NaN | X | NaN | NaN | NaN | NaN | 4.902343e+05 | 4.902343e+05 | NaN | NaN | NQIIACNGSIQSIPEIPDDLK |
| ASPSCR1 | ASPC1\_HUMAN | Tether containing UBX domain for GLUT4 | 325 | 310 | ASPSCR1 | ASPC1\_HUMAN | Tether containing UBX domain for GLUT4 | 16C(125.0477) | NaN | X | NaN | NaN | NaN | NaN | 4.890375e+05 | 4.890375e+05 | NaN | NaN | ERPVDREPVDREPVVCHPDLEER |
| ALDOA | ALDOA\_HUMAN | Fructose-bisphosphate aldolase A | 339, 339 | 331 | ALDOA | ALDOA\_HUMAN | Fructose-bisphosphate aldolase A | 9C(125.0477) 9C(31.9988) | NaN | X | NaN | X | NaN | NaN | 4.886075e+05 | 4.886075e+05 | NaN | NaN | RALANSLACQGK |
| LHPP | LHPP\_HUMAN | Phospholysine phosphohistidine inorganic pyrophosphate phosphatase | 88 | 71 | LHPP | LHPP\_HUMAN | Phospholysine phosphohistidine inorganic pyrophosphate phosphatase | 18C(125.0477) | NaN | X | NaN | NaN | NaN | NaN | 4.885793e+05 | 4.885793e+05 | NaN | NaN | LGFDISEQEVTAPAPAACQILK |
| FASN | FAS\_HUMAN | Fatty acid synthase | 1403 | 1394 | FASN | FAS\_HUMAN | Fatty acid synthase | 10C(125.0477) | NaN | X | NaN | NaN | NaN | NaN | 4.838548e+05 | 4.838548e+05 | NaN | NaN | SFYGSTLFLCR |
| TPP2 | TPP2\_HUMAN | Tripeptidyl-peptidase 2 | 28 | 19 | TPP2 | TPP2\_HUMAN | Tripeptidyl-peptidase 2 | 10C(125.0477) | NaN | X | NaN | NaN | NaN | NaN | 4.825123e+05 | 4.825123e+05 | NaN | NaN | KETGAASFLCR |
| LDHB | LDHB\_HUMAN | L-lactate dehydrogenase B chain | 164 | 159 | LDHB | LDHB\_HUMAN | L-lactate dehydrogenase B chain | 6C(125.0477) | NaN | X | NaN | NaN | NaN | NaN | 4.816278e+05 | NaN | 4.816278e+05 | NaN | VIGSGCNLDSAR |
| SMYD5 | SMYD5\_HUMAN | Histone-lysine N-trimethyltransferase SMYD5 | 101 | 87 | SMYD5 | SMYD5\_HUMAN | Histone-lysine N-trimethyltransferase SMYD5 | 15C(125.0477) | NaN | X | NaN | NaN | NaN | NaN | 4.809648e+05 | 4.809648e+05 | NaN | NaN | LTGKPGQVLPHPELCTVR |
| ACLY | ACLY\_HUMAN | ATP-citrate synthase | 633 | 617 | ACLY | ACLY\_HUMAN | ATP-citrate synthase | 17C(125.0477) | NaN | X | NaN | NaN | NaN | NaN | 4.789638e+05 | 4.789638e+05 | NaN | NaN | GVTIIGPATVGGIKPGCFK |
| WDR1 | WDR1\_HUMAN | WD repeat-containing protein 1 | 170 | 162 | WDR1 | WDR1\_HUMAN | WD repeat-containing protein 1 | 9C(125.0477) | NaN | X | NaN | NaN | NaN | NaN | 4.773339e+05 | 4.773339e+05 | NaN | NaN | LATGSDDNCAAFFEGPPFK |
| IPO7 | IPO7\_HUMAN | Importin-7 | 757 | 753 | IPO7 | IPO7\_HUMAN | Importin-7 | 5C(125.0477) | NaN | X | NaN | NaN | NaN | NaN | 4.764896e+05 | 4.764896e+05 | NaN | NaN | GIDQCIPLFVEAALER |
| DTYMK | KTHY\_HUMAN | Thymidylate kinase | 31 | 26 | DTYMK | KTHY\_HUMAN | Thymidylate kinase | 6C(125.0477) | NaN | X | NaN | NaN | NaN | NaN | 4.742887e+05 | 4.181925e+05 | 5.609615e+04 | NaN | LVEALCAAGHR |
| PSMD13 | PSD13\_HUMAN | 26S proteasome non-ATPase regulatory subunit 13 | 357 | 351 | PSMD13 | PSD13\_HUMAN | 26S proteasome non-ATPase regulatory subunit 13 | 7C(125.0477) | NaN | X | NaN | NaN | NaN | NaN | 4.733045e+05 | 4.733045e+05 | NaN | NaN | DRLEFWCTDVK |
| BCAM | BCAM\_HUMAN | Basal cell adhesion molecule | 291 | 283 | BCAM | BCAM\_HUMAN | Basal cell adhesion molecule | 9C(125.0477) | NaN | X | NaN | NaN | NaN | NaN | 4.730476e+05 | NaN | 3.622682e+05 | 110779.4450 | EGDTVQLLCR |
| STAM | STAM1\_HUMAN | Signal transducing adapter molecule 1 | 85 | 79 | STAM | STAM1\_HUMAN | Signal transducing adapter molecule 1 | 7C(125.0477) | NaN | X | NaN | NaN | NaN | NaN | 4.714622e+05 | 4.714622e+05 | NaN | NaN | IFHLEVCSR |
| SELENBP1 | SBP1\_HUMAN | Methanethiol oxidase | 466 | 461 | SELENBP1 | SBP1\_HUMAN | Methanethiol oxidase | 6C(125.0477) | NaN | X | NaN | NaN | NaN | NaN | 4.694340e+05 | 4.659648e+05 | 3.469257e+03 | NaN | YPGGDCSSDIWI |
| UBE2O | UBE2O\_HUMAN | (E3-independent) E2 ubiquitin-conjugating enzyme | 341 | 338 | UBE2O | UBE2O\_HUMAN | (E3-independent) E2 ubiquitin-conjugating enzyme | 4C(125.0477) | NaN | X | NaN | NaN | NaN | NaN | 4.690183e+05 | 4.690183e+05 | NaN | NaN | RLGCFDHAQR |
| ACTR10 | ARP10\_HUMAN | Actin-related protein 10 | 97 | 89 | ACTR10 | ARP10\_HUMAN | Actin-related protein 10 | 9C(125.0477) | NaN | X | NaN | NaN | NaN | NaN | 4.685171e+05 | 4.685171e+05 | NaN | NaN | VVIIESVLCPSHFR |
| GAPDH | G3P\_HUMAN | Glyceraldehyde-3-phosphate dehydrogenase | 247, 247 | 235 | GAPDH | G3P\_HUMAN | Glyceraldehyde-3-phosphate dehydrogenase | 13C(125.0477) 13C(57.0215) | X | X | NaN | NaN | NaN | NaN | 4.681342e+05 | NaN | 4.681342e+05 | NaN | VPTANVSVVDLTCR |
| PRPS1 | PRPS1\_HUMAN | Ribose-phosphate pyrophosphokinase 1 | 165, 165 | 164 | PRPS1 | PRPS1\_HUMAN | Ribose-phosphate pyrophosphokinase 1 | 2C(125.0477) 2C(31.9988) | NaN | X | NaN | X | NaN | NaN | 4.671544e+05 | 4.671544e+05 | NaN | NaN | NCTIVSPDAGGAK |
| KRT17 | K1C17\_HUMAN | Keratin, type I cytoskeletal 17 | 40 | 31 | KRT17 | K1C17\_HUMAN | Keratin, type I cytoskeletal 17 | 10C(125.0477) | NaN | X | NaN | NaN | NaN | NaN | 4.660518e+05 | NaN | 2.340410e+05 | 232010.7300 | LSGGLGAGSCR |
| PGM2 | PGM2\_HUMAN | Phosphoglucomutase-2 | 484, 484 | 479 | PGM2 | PGM2\_HUMAN | Phosphoglucomutase-2 | 6C(125.0477) 6C(31.9988) | NaN | X | NaN | X | NaN | NaN | 4.652668e+05 | 4.652668e+05 | NaN | NaN | ASYFICHDQETIK |
| ANK1 | ANK1\_HUMAN | Ankyrin-1 | 1258, 1260 | 1256 | ANK1 | ANK1\_HUMAN | Ankyrin-1 | 3C(-34.0809) 5C(31.9988) | NaN | NaN | NaN | X | X | NaN | 4.651677e+05 | NaN | 4.651677e+05 | NaN | LRCYCMTDDKVDK |
| HBD | HBD\_HUMAN | Hemoglobin subunit delta | 113 | 97 | HBD | HBD\_HUMAN | Hemoglobin subunit delta | 17C(125.0477) | NaN | X | NaN | NaN | NaN | NaN | 4.650061e+05 | 4.650061e+05 | NaN | NaN | LHVDPENFRLLGNVLVCVLAR |
| NT5C | NT5C\_HUMAN | 5'(3')-deoxyribonucleotidase, cytosolic type | 98 | 86 | NT5C | NT5C\_HUMAN | 5'(3')-deoxyribonucleotidase, cytosolic type | 13C(125.0477) | NaN | X | NaN | NaN | NaN | NaN | 4.636854e+05 | 4.636854e+05 | NaN | NaN | EMNDLPDTQVFICTSPLLK |
| PPP2CA | PP2AA\_HUMAN | Serine/threonine-protein phosphatase 2A catalytic subunit alpha isoform | 269 | 269 | PPP2CA | PP2AA\_HUMAN | Serine/threonine-protein phosphatase 2A catalytic subunit alpha isoform | 1C(125.0477) | NaN | X | NaN | NaN | NaN | NaN | 4.633301e+05 | 4.633301e+05 | NaN | NaN | CGNQAAIMELDDTLK |
| ADH5 | ADHX\_HUMAN | Alcohol dehydrogenase class-3 | 97, 97, 100, 100, 103 | 85 | ADH5 | ADHX\_HUMAN | Alcohol dehydrogenase class-3 | 13C(125.0477) 13C(15.9949) 16C(15.9949) 16C(31.9988) 19C(125.0477) | NaN | X | X | X | NaN | NaN | 4.607942e+05 | 4.607942e+05 | NaN | NaN | AGDTVIPLYIPQCGECKFCLNPK |
| UBE2N | UBE2N\_HUMAN | Ubiquitin-conjugating enzyme E2 N | 87 | 86 | UBE2N | UBE2N\_HUMAN | Ubiquitin-conjugating enzyme E2 N | 2C(125.0477) | NaN | X | NaN | NaN | NaN | NaN | 4.601496e+05 | 4.601496e+05 | NaN | NaN | ICLDILKDK |
| BLVRB | BLVRB\_HUMAN | Flavin reductase (NADPH) | 188 | 188 | BLVRB | BLVRB\_HUMAN | Flavin reductase (NADPH) | 1C(125.0477) | NaN | X | NaN | NaN | NaN | NaN | 4.600769e+05 | NaN | 4.600769e+05 | NaN | CLTTDEYDGHSTYPSHQYQ |
| ABCB6 | ABCB6\_HUMAN | ATP-binding cassette sub-family B member 6 | 646 | 639 | ABCB6 | ABCB6\_HUMAN | ATP-binding cassette sub-family B member 6 | 8C(125.0477) | NaN | X | NaN | NaN | NaN | NaN | 4.597242e+05 | NaN | 4.597242e+05 | NaN | FYDISSGCIR |
| RUVBL1 | RUVB1\_HUMAN | RuvB-like 1 | 49 | 47 | RUVBL1 | RUVB1\_HUMAN | RuvB-like 1 | 3C(125.0477) | NaN | X | NaN | NaN | NaN | NaN | 4.582877e+05 | 4.582877e+05 | NaN | NaN | EACGVIVELIK |
| ANK1 | ANK1\_HUMAN | Ankyrin-1 | 278 | 269 | ANK1 | ANK1\_HUMAN | Ankyrin-1 | 10C(31.9988) | NaN | NaN | NaN | X | NaN | NaN | 4.582351e+05 | NaN | NaN | 458235.1200 | TKDELTPLHCAAR |
| FASN | FAS\_HUMAN | Fatty acid synthase | 1992 | 1977 | FASN | FAS\_HUMAN | Fatty acid synthase | 16C(125.0477) | NaN | X | NaN | NaN | NaN | NaN | 4.567488e+05 | 4.567488e+05 | NaN | NaN | DGLLENQTPEFFQDVCKPK |
| CAND1 | CAND1\_HUMAN | Cullin-associated NEDD8-dissociated protein 1 | 131 | 112 | CAND1 | CAND1\_HUMAN | Cullin-associated NEDD8-dissociated protein 1 | 20C(125.0477) | NaN | X | NaN | NaN | NaN | NaN | 4.564016e+05 | 4.564016e+05 | NaN | NaN | TVIGELPPASSGSALAANVCK |
| TWF2 | TWF2\_HUMAN | Twinfilin-2 | 67 | 56 | TWF2 | TWF2\_HUMAN | Twinfilin-2 | 12C(125.0477) | NaN | X | NaN | NaN | NaN | NaN | 4.553482e+05 | 4.553482e+05 | NaN | NaN | AVLPLLDAQQPCYLLYR |
| RIC8A | RIC8A\_HUMAN | Synembryn-A | 341 | 327 | RIC8A | RIC8A\_HUMAN | Synembryn-A | 15C(125.0477) | NaN | X | NaN | NaN | NaN | NaN | 4.545608e+05 | 4.545608e+05 | NaN | NaN | LKESVAPVLSVLTECAR |
| GPS1 | CSN1\_HUMAN | COP9 signalosome complex subunit 1 | 175, 184 | 164 | GPS1 | CSN1\_HUMAN | COP9 signalosome complex subunit 1 | 12C(15.9949) 21C(15.9949) | NaN | NaN | X | NaN | NaN | NaN | 4.539281e+05 | 4.539281e+05 | NaN | NaN | GHDDLGDHYLDCGDLSNALKCYSR |
| TCP1 | TCPA\_HUMAN | T-complex protein 1 subunit alpha | 296 | 278 | TCP1 | TCPA\_HUMAN | T-complex protein 1 subunit alpha | 19C(125.0477) | NaN | X | NaN | NaN | NaN | NaN | 4.519055e+05 | 4.519055e+05 | NaN | NaN | ILATGANVILTTGGIDDMCLK |
| COMT | COMT\_HUMAN | Catechol O-methyltransferase | 207 | 197 | COMT | COMT\_HUMAN | Catechol O-methyltransferase | 11C(125.0477) | NaN | X | NaN | NaN | NaN | NaN | 4.512836e+05 | 4.512836e+05 | NaN | NaN | YLPDTLLLEECGLLR |
| PSMB2 | PSB2\_HUMAN | Proteasome subunit beta type-2 | 91 | 86 | PSMB2 | PSB2\_HUMAN | Proteasome subunit beta type-2 | 6C(125.0477) | NaN | X | NaN | NaN | NaN | NaN | 4.500434e+05 | 4.500434e+05 | NaN | NaN | RNLADCLR |
| HBG1 | HBG1\_HUMAN | Hemoglobin subunit gamma-1 | 94 | 84 | HBG1 | HBG1\_HUMAN | Hemoglobin subunit gamma-1 | 11C(125.0477) | NaN | X | NaN | NaN | NaN | NaN | 4.490138e+05 | NaN | 4.490138e+05 | NaN | GTFAQLSELHCDK |
| AP2B1 | AP2B1\_HUMAN | AP-2 complex subunit beta | 857, 857 | 856 | AP2B1 | AP2B1\_HUMAN | AP-2 complex subunit beta | 2C(125.0477) 2C(31.9988) | NaN | X | NaN | X | NaN | NaN | 4.488232e+05 | NaN | 4.488232e+05 | NaN | ECHLNADTVSSK |
| NIF3L1 | NIF3L\_HUMAN | NIF3-like protein 1 | 254 | 253 | NIF3L1 | NIF3L\_HUMAN | NIF3-like protein 1 | 2C(125.0477) | NaN | X | NaN | NaN | NaN | NaN | 4.485466e+05 | 4.485466e+05 | NaN | NaN | LCTLDESVSLATMIDR |
| ANP32A | AN32A\_HUMAN | Acidic leucine-rich nuclear phosphoprotein 32 family member A | 87 | 87 | ANP32A | AN32A\_HUMAN | Acidic leucine-rich nuclear phosphoprotein 32 family member A | 1C(125.0477) | NaN | X | NaN | NaN | NaN | NaN | 4.481124e+05 | 4.335694e+05 | 1.454297e+04 | NaN | CPNLTHLNLSGNK |
| XPO1 | XPO1\_HUMAN | Exportin-1 | 119 | 113 | XPO1 | XPO1\_HUMAN | Exportin-1 | 7C(125.0477) | NaN | X | NaN | NaN | NaN | NaN | 4.479629e+05 | 4.479629e+05 | NaN | NaN | TSSDPTCVEKEK |
| PGK1 | PGK1\_HUMAN | Phosphoglycerate kinase 1 | 99 | 98 | PGK1 | PGK1\_HUMAN | Phosphoglycerate kinase 1 | 2C(125.0477) | NaN | X | NaN | NaN | NaN | NaN | 4.473271e+05 | NaN | 4.288118e+05 | 18515.3000 | DCVGPEVEK |
| HNRNPA1 | ROA1\_HUMAN | Heterogeneous nuclear ribonucleoprotein A1 | 175 | 167 | HNRNPA1 | ROA1\_HUMAN | Heterogeneous nuclear ribonucleoprotein A1 | 9C(125.0477) | NaN | X | NaN | NaN | NaN | NaN | 4.470828e+05 | 4.470828e+05 | NaN | NaN | YHTVNGHNCEVR |
| HACE1 | HACE1\_HUMAN | E3 ubiquitin-protein ligase HACE1 | 99 | 87 | HACE1 | HACE1\_HUMAN | E3 ubiquitin-protein ligase HACE1 | 13C(57.0215) | X | NaN | NaN | NaN | NaN | NaN | 4.468767e+05 | 4.468767e+05 | NaN | NaN | KGANPNYQDISGCTPLHLAAR |
| NADSYN1 | NADE\_HUMAN | Glutamine-dependent NAD(+) synthetase | 686 | 686 | NADSYN1 | NADE\_HUMAN | Glutamine-dependent NAD(+) synthetase | 1C(125.0477) | NaN | X | NaN | NaN | NaN | NaN | 4.460373e+05 | 4.460373e+05 | NaN | NaN | CIENQVLQLER |
| NIT2 | NIT2\_HUMAN | Omega-amidase NIT2 | 97 | 93 | NIT2 | NIT2\_HUMAN | Omega-amidase NIT2 | 5C(125.0477) | NaN | X | NaN | NaN | NaN | NaN | 4.452186e+05 | 4.452186e+05 | NaN | NaN | LYNTCAVFGPDGTLLAK |
| CFL1 | COF1\_HUMAN | Cofilin-1 | 39 | 35 | CFL1 | COF1\_HUMAN | Cofilin-1 | 5C(125.0477) | NaN | X | NaN | NaN | NaN | NaN | 4.434123e+05 | 2.860607e+05 | 1.573516e+05 | NaN | AVLFCLSEDKK |
| SERPINB9 | SPB9\_HUMAN | Serpin B9 | 350 | 349 | SERPINB9 | SPB9\_HUMAN | Serpin B9 | 2C(125.0477) | NaN | X | NaN | NaN | NaN | NaN | 4.432887e+05 | 4.432887e+05 | NaN | NaN | FCADHPFLFFIR |
| ECPAS | ECM29\_HUMAN | Proteasome adapter and scaffold protein ECM29 | 1638 | 1633 | ECPAS | ECM29\_HUMAN | Proteasome adapter and scaffold protein ECM29 | 6C(125.0477) | NaN | X | NaN | NaN | NaN | NaN | 4.419820e+05 | 4.419820e+05 | NaN | NaN | IVAISCAADILK |
| ACTB | ACTB\_HUMAN | Actin, cytoplasmic 1 | 285, 285 | 285 | ACTB | ACTB\_HUMAN | Actin, cytoplasmic 1 | 1C(125.0477) 1C(31.9988) | NaN | X | NaN | X | NaN | NaN | 4.418818e+05 | 4.418818e+05 | NaN | NaN | CDVDIRK |
| PIN1 | PIN1\_HUMAN | Peptidyl-prolyl cis-trans isomerase NIMA-interacting 1 | 57 | 57 | PIN1 | PIN1\_HUMAN | Peptidyl-prolyl cis-trans isomerase NIMA-interacting 1 | 1C(125.0477) | NaN | X | NaN | NaN | NaN | NaN | 4.415781e+05 | 4.415781e+05 | NaN | NaN | CSHLLVK |
| DPP3 | DPP3\_HUMAN | Dipeptidyl peptidase 3 | 654 | 639 | DPP3 | DPP3\_HUMAN | Dipeptidyl peptidase 3 | 16C(125.0477) | NaN | X | NaN | NaN | NaN | NaN | 4.401925e+05 | 4.401925e+05 | NaN | NaN | ALYEGYATVTDAPPECFLTLR |
| KRT10 | K1C10\_HUMAN | Keratin, type I cytoskeletal 10 | 25 | 16 | KRT10 | K1C10\_HUMAN | Keratin, type I cytoskeletal 10 | 10C(125.0477) | NaN | X | NaN | NaN | NaN | NaN | 4.401623e+05 | 6.099834e+04 | 2.238113e+05 | 155352.6900 | SGGGGGGGGCGGGGGVSSLR |
| ARIH2 | ARI2\_HUMAN | E3 ubiquitin-protein ligase ARIH2 | 280 | 280 | ARIH2 | ARI2\_HUMAN | E3 ubiquitin-protein ligase ARIH2 | 1C(125.0477) | NaN | X | NaN | NaN | NaN | NaN | 4.396367e+05 | 4.396367e+05 | NaN | NaN | CADDSETANYISAHTK |
| CLTC | CLH1\_HUMAN | Clathrin heavy chain 1 | 824 | 812 | CLTC | CLH1\_HUMAN | Clathrin heavy chain 1 | 13C(125.0477) | NaN | X | NaN | NaN | NaN | NaN | 4.377019e+05 | 4.377019e+05 | NaN | NaN | LPVVIGGLLDVDCSEDVIK |
| IGLC2 | IGLC2\_HUMAN | Immunoglobulin lambda constant 2 | 87, 87 | 84 | IGLC2 | IGLC2\_HUMAN | Immunoglobulin lambda constant 2 | 4C(125.0477) 4C(57.0215) | X | X | NaN | NaN | NaN | NaN | 4.346589e+05 | 1.030877e+05 | 3.315712e+05 | NaN | SYSCQVTHEGSTVEK |
| PCMT1 | PIMT\_HUMAN | Protein-L-isoaspartate(D-aspartate) O-methyltransferase | 95, 102 | 82 | PCMT1 | PIMT\_HUMAN | Protein-L-isoaspartate(D-aspartate) O-methyltransferase | 14C(31.9988) 21C(15.9949) | NaN | NaN | X | X | NaN | NaN | 4.338890e+05 | 4.338890e+05 | NaN | NaN | ALDVGSGSGILTACFARMVGCTGK |
| FAM114A2 | F1142\_HUMAN | Protein FAM114A2 | 388 | 381 | FAM114A2 | F1142\_HUMAN | Protein FAM114A2 | 8C(125.0477) | NaN | X | NaN | NaN | NaN | NaN | 4.324108e+05 | 4.324108e+05 | NaN | NaN | SLAELTACSIELFHK |
| PRDX6 | PRDX6\_HUMAN | Peroxiredoxin-6 | 91 | 85 | PRDX6 | PRDX6\_HUMAN | Peroxiredoxin-6 | 7C(125.0477) | NaN | X | NaN | NaN | NaN | NaN | 4.312435e+05 | 4.312435e+05 | NaN | NaN | DINAYNCEEPTEKLPFPIIDDR |
| UROS | HEM4\_HUMAN | Uroporphyrinogen-III synthase | 241 | 231 | UROS | HEM4\_HUMAN | Uroporphyrinogen-III synthase | 11C(125.0477) | NaN | X | NaN | NaN | NaN | NaN | 4.308017e+05 | 4.308017e+05 | NaN | NaN | ALAAQGLPVSCTAESPTPQALATGIR |
| COPS7B | CSN7B\_HUMAN | COP9 signalosome complex subunit 7b | 240 | 239 | COPS7B | CSN7B\_HUMAN | COP9 signalosome complex subunit 7b | 2C(125.0477) | NaN | X | NaN | NaN | NaN | NaN | 4.304539e+05 | 4.304539e+05 | NaN | NaN | ECPPHAEQR |
| IGHG1 | IGHG1\_HUMAN | Immunoglobulin heavy constant gamma 1 | 250, 250, 250 | 244 | IGHG1 | IGHG1\_HUMAN | Immunoglobulin heavy constant gamma 1 | 7C(125.0477) 7C(31.9988) 7C(57.0215) | X | X | NaN | X | NaN | NaN | 4.303706e+05 | NaN | 4.303706e+05 | NaN | NQVSLTCLVK |
| YWHAH | 1433F\_HUMAN | 14-3-3 protein eta | 112 | 111 | YWHAH | 1433F\_HUMAN | 14-3-3 protein eta | 2C(125.0477) | NaN | X | NaN | NaN | NaN | NaN | 4.298067e+05 | 1.905696e+05 | 2.392371e+05 | NaN | NCNDFQYESK |
| HMBS | HEM3\_HUMAN | Porphobilinogen deaminase | 114 | 99 | HMBS | HEM3\_HUMAN | Porphobilinogen deaminase | 16C(125.0477) | NaN | X | NaN | NaN | NaN | NaN | 4.295836e+05 | 4.295836e+05 | NaN | NaN | DLPTVLPPGFTIGAICK |
| HINT1 | HINT1\_HUMAN | Adenosine 5'-monophosphoramidase HINT1 | 84 | 83 | HINT1 | HINT1\_HUMAN | Adenosine 5'-monophosphoramidase HINT1 | 2C(125.0477) | NaN | X | NaN | NaN | NaN | NaN | 4.292864e+05 | 4.130020e+05 | 1.628438e+04 | NaN | KCAADLGLNK |
| PPP2R1A | 2AAA\_HUMAN | Serine/threonine-protein phosphatase 2A 65 kDa regulatory subunit A alpha isoform | 377 | 375 | PPP2R1A | 2AAA\_HUMAN | Serine/threonine-protein phosphatase 2A 65 kDa regulatory subunit A alpha isoform | 3C(125.0477) | NaN | X | NaN | NaN | NaN | NaN | 4.289219e+05 | 3.013558e+05 | 1.275661e+05 | NaN | DECPEVR |
| ACP1 | PPAC\_HUMAN | Low molecular weight phosphotyrosine protein phosphatase | 146 | 125 | ACP1 | PPAC\_HUMAN | Low molecular weight phosphotyrosine protein phosphatase | 22C(125.0477) | NaN | X | NaN | NaN | NaN | NaN | 4.273551e+05 | 4.273551e+05 | NaN | NaN | QLIIEDPYYGNDSDFETVYQQCVR |
| TMCO1 | TMCO1\_HUMAN | Calcium load-activated calcium channel | 68, 50 | 49 | TMCO1 | TMCO1\_HUMAN | Calcium load-activated calcium channel | 20C(57.0215) 2C(305.3076) | X | NaN | NaN | NaN | NaN | X | 4.271722e+05 | NaN | 4.271722e+05 | NaN | RCEMSTMFADTLLIVFISVCTALLAEGITWVLVYR |
| RNH1 | RINI\_HUMAN | Ribonuclease inhibitor | 45, 38 | 35 | RNH1 | RINI\_HUMAN | Ribonuclease inhibitor | 11C(15.9949) 4C(15.9949) | NaN | NaN | X | NaN | NaN | NaN | 4.269310e+05 | 3.595118e+05 | 6.741923e+04 | NaN | LDDCGLTEARCKDISSALR |
| CIAO1 | CIAO1\_HUMAN | Probable cytosolic iron-sulfur protein assembly protein CIAO1 | 123 | 110 | CIAO1 | CIAO1\_HUMAN | Probable cytosolic iron-sulfur protein assembly protein CIAO1 | 14C(125.0477) | NaN | X | NaN | NaN | NaN | NaN | 4.255288e+05 | 4.255288e+05 | NaN | NaN | SVAWAPSGNLLATCSR |
| DNPEP | DNPEP\_HUMAN | Aspartyl aminopeptidase | 457, 457 | 454 | DNPEP | DNPEP\_HUMAN | Aspartyl aminopeptidase | 4C(-34.0809) 4C(125.0477) | NaN | X | NaN | NaN | X | NaN | 4.243089e+05 | 4.243089e+05 | NaN | NaN | EMACTTGVLQTLTLFK |
| DNAJA4 | DNJA4\_HUMAN | DnaJ homolog subfamily A member 4 | 138 | 138 | DNAJA4 | DNJA4\_HUMAN | DnaJ homolog subfamily A member 4 | 1C(125.0477) | NaN | X | NaN | NaN | NaN | NaN | 4.242224e+05 | 4.242224e+05 | NaN | NaN | CEGVGGK |
| ALDH16A1 | A16A1\_HUMAN | Aldehyde dehydrogenase family 16 member A1 | 350 | 346 | ALDH16A1 | A16A1\_HUMAN | Aldehyde dehydrogenase family 16 member A1 | 5C(125.0477) | NaN | X | NaN | NaN | NaN | NaN | 4.229342e+05 | 2.436679e+05 | 1.792663e+05 | NaN | GAAACDLVQR |
| CORO7 | CORO7\_HUMAN | Coronin-7 | 505 | 492 | CORO7 | CORO7\_HUMAN | Coronin-7 | 14C(125.0477) | NaN | X | NaN | NaN | NaN | NaN | 4.199784e+05 | 4.199784e+05 | NaN | NaN | GLNLTTPGESDGFCANK |
| PPP2R1A | 2AAA\_HUMAN | Serine/threonine-protein phosphatase 2A 65 kDa regulatory subunit A alpha isoform | 447 | 443 | PPP2R1A | 2AAA\_HUMAN | Serine/threonine-protein phosphatase 2A 65 kDa regulatory subunit A alpha isoform | 5C(125.0477) | NaN | X | NaN | NaN | NaN | NaN | 4.194500e+05 | 4.194500e+05 | NaN | NaN | LNSLCMAWLVDHVYAIR |
| ANK1 | ANK1\_HUMAN | Ankyrin-1 | 943, 943 | 942 | ANK1 | ANK1\_HUMAN | Ankyrin-1 | 2C(125.0477) 2C(31.9988) | NaN | X | NaN | X | NaN | NaN | 4.186504e+05 | NaN | NaN | 418650.4000 | TCAAPTR |
| TAOK3 | TAOK3\_HUMAN | Serine/threonine-protein kinase TAO3 | 795 | 787 | TAOK3 | TAOK3\_HUMAN | Serine/threonine-protein kinase TAO3 | 9C(125.0477) | NaN | X | NaN | NaN | NaN | NaN | 4.177504e+05 | 4.177504e+05 | NaN | NaN | LDEAQEAECQALR |
| THOP1 | THOP1\_HUMAN | Thimet oligopeptidase | 246 | 238 | THOP1 | THOP1\_HUMAN | Thimet oligopeptidase | 9C(125.0477) | NaN | X | NaN | NaN | NaN | NaN | 4.169989e+05 | 4.169989e+05 | NaN | NaN | RKVEEAFNCR |
| CPPED1 | CPPED\_HUMAN | Serine/threonine-protein phosphatase CPPED1 | 143 | 118 | CPPED1 | CPPED\_HUMAN | Serine/threonine-protein phosphatase CPPED1 | 26C(125.0477) | NaN | X | NaN | NaN | NaN | NaN | 4.169803e+05 | 4.169803e+05 | NaN | NaN | AIPLVLVSGNHDIGNTPTAETVEEFCR |
| PSMB1 | PSB1\_HUMAN | Proteasome subunit beta type-1 | 82 | 78 | PSMB1 | PSB1\_HUMAN | Proteasome subunit beta type-1 | 5C(31.9988) | NaN | NaN | NaN | X | NaN | NaN | 4.166528e+05 | 4.166528e+05 | NaN | NaN | TVIGCSGFHGDCLTLTK |
| GNL1 | GNL1\_HUMAN | Guanine nucleotide-binding protein-like 1 | 517 | 514 | GNL1 | GNL1\_HUMAN | Guanine nucleotide-binding protein-like 1 | 4C(125.0477) | NaN | X | NaN | NaN | NaN | NaN | 4.148658e+05 | 4.148658e+05 | NaN | NaN | LSLCFHPPGYSEQK |
| LGALS9 | LEG9\_HUMAN | Galectin-9 | 74 | 66 | LGALS9 | LEG9\_HUMAN | Galectin-9 | 9C(125.0477) | NaN | X | NaN | NaN | NaN | NaN | 4.136602e+05 | 9.022941e+04 | 3.234308e+05 | NaN | FEDGGYVVCNTR |
| SERPINB6 | SPB6\_HUMAN | Serpin B6 | 370 | 364 | SERPINB6 | SPB6\_HUMAN | Serpin B6 | 7C(125.0477) | NaN | X | NaN | NaN | NaN | NaN | 4.120434e+05 | 4.120434e+05 | NaN | NaN | TNGILFCGR |
| HNRNPK | HNRPK\_HUMAN | Heterogeneous nuclear ribonucleoprotein K | 145 | 140 | HNRNPK | HNRPK\_HUMAN | Heterogeneous nuclear ribonucleoprotein K | 6C(125.0477) | NaN | X | NaN | NaN | NaN | NaN | 4.104237e+05 | 2.950088e+05 | 1.154148e+05 | NaN | GSDFDCELR |
| ATP6V1A | VATA\_HUMAN | V-type proton ATPase catalytic subunit A | 277 | 266 | ATP6V1A | VATA\_HUMAN | V-type proton ATPase catalytic subunit A | 12C(125.0477) | NaN | X | NaN | NaN | NaN | NaN | 4.103731e+05 | 4.103731e+05 | NaN | NaN | YSNSDVIIYVGCGER |
| PPP6C | PPP6\_HUMAN | Serine/threonine-protein phosphatase 6 catalytic subunit | 129 | 118 | PPP6C | PPP6\_HUMAN | Serine/threonine-protein phosphatase 6 catalytic subunit | 12C(125.0477) | NaN | X | NaN | NaN | NaN | NaN | 4.098764e+05 | 4.098764e+05 | NaN | NaN | QITQVYGFYDECQTK |
| ADK | ADK\_HUMAN | Adenosine kinase | 353 | 351 | ADK | ADK\_HUMAN | Adenosine kinase | 3C(125.0477) | NaN | X | NaN | NaN | NaN | NaN | 4.087456e+05 | 4.087456e+05 | NaN | NaN | TGCTFPEKPDFH |
| TTLL12 | TTL12\_HUMAN | Tubulin--tyrosine ligase-like protein 12 | 126 | 106 | TTLL12 | TTL12\_HUMAN | Tubulin--tyrosine ligase-like protein 12 | 21C(125.0477) | NaN | X | NaN | NaN | NaN | NaN | 4.073232e+05 | 4.073232e+05 | NaN | NaN | ESGLQAAHPNSIFLIDHAWTCR |
| FSCN1 | FSCN1\_HUMAN | Fascin | 121 | 119 | FSCN1 | FSCN1\_HUMAN | Fascin | 3C(125.0477) | NaN | X | NaN | NaN | NaN | NaN | 4.061470e+05 | 3.799839e+05 | 2.616311e+04 | NaN | LSCFAQTVSPAEK |
| UROS | HEM4\_HUMAN | Uroporphyrinogen-III synthase | 14 | 8 | UROS | HEM4\_HUMAN | Uroporphyrinogen-III synthase | 7C(125.0477) | NaN | X | NaN | NaN | NaN | NaN | 4.056265e+05 | 4.056265e+05 | NaN | NaN | DAKEDDCGQDPYIR |
| BAG1 | BAG1\_HUMAN | BAG family molecular chaperone regulator 1 | 213 | 199 | BAG1 | BAG1\_HUMAN | BAG family molecular chaperone regulator 1 | 15C(125.0477) | NaN | X | NaN | NaN | NaN | NaN | 4.048747e+05 | 4.048747e+05 | NaN | NaN | EMETPLSALGIQDGCR |
| MCTS1 | MCTS1\_HUMAN | Malignant T-cell-amplified sequence 1 | 113 | 104 | MCTS1 | MCTS1\_HUMAN | Malignant T-cell-amplified sequence 1 | 10C(125.0477) | NaN | X | NaN | NaN | NaN | NaN | 4.023502e+05 | 4.023502e+05 | NaN | NaN | FVLSGANIMCPGLTSPGAK |
| PSMB5 | PSB5\_HUMAN | Proteasome subunit beta type-5 | 111 | 93 | PSMB5 | PSB5\_HUMAN | Proteasome subunit beta type-5 | 19C(125.0477) | NaN | X | NaN | NaN | NaN | NaN | 4.005766e+05 | 4.005766e+05 | NaN | NaN | VIEINPYLLGTMAGGAADCSFWER |
| IPO5 | IPO5\_HUMAN | Importin-5 | 972, 972 | 962 | IPO5 | IPO5\_HUMAN | Importin-5 | 11C(125.0477) 11C(31.9988) | NaN | X | NaN | X | NaN | NaN | 3.995830e+05 | 3.995830e+05 | NaN | NaN | TKENVNATENCISAVGK |
| LIN7C | LIN7C\_HUMAN | Protein lin-7 homolog C | 47 | 41 | LIN7C | LIN7C\_HUMAN | Protein lin-7 homolog C | 7C(125.0477) | NaN | X | NaN | NaN | NaN | NaN | 3.987220e+05 | 1.784292e+05 | 2.202928e+05 | NaN | VLQSEFCNAVR |
| TBC1D24 | TBC24\_HUMAN | TBC1 domain family member 24 | 386 | 382 | TBC1D24 | TBC24\_HUMAN | TBC1 domain family member 24 | 5C(125.0477) | NaN | X | NaN | NaN | NaN | NaN | 3.980328e+05 | 3.980328e+05 | NaN | NaN | FYFQCEGHEPTLLLIK |
| AP2B1 | AP2B1\_HUMAN | AP-2 complex subunit beta | 241 | 233 | AP2B1 | AP2B1\_HUMAN | AP-2 complex subunit beta | 9C(125.0477) | NaN | X | NaN | NaN | NaN | NaN | 3.976277e+05 | 3.976277e+05 | NaN | NaN | DDREAQSICER |
| COPS3 | CSN3\_HUMAN | COP9 signalosome complex subunit 3 | 104 | 95 | COPS3 | CSN3\_HUMAN | COP9 signalosome complex subunit 3 | 10C(125.0477) | NaN | X | NaN | NaN | NaN | NaN | 3.975019e+05 | 3.975019e+05 | NaN | NaN | YATDTFAGLCHQLTNALVER |
| ESD | ESTD\_HUMAN | S-formylglutathione hydrolase | 158 | 143 | ESD | ESTD\_HUMAN | S-formylglutathione hydrolase | 16C(125.0477) | NaN | X | NaN | NaN | NaN | NaN | 3.970651e+05 | 3.970651e+05 | NaN | NaN | MSIFGHSMGGHGALICALK |
| RNH1 | RINI\_HUMAN | Ribonuclease inhibitor | 220 | 218 | RNH1 | RINI\_HUMAN | Ribonuclease inhibitor | 3C(125.0477) | NaN | X | NaN | NaN | NaN | NaN | 3.944700e+05 | NaN | 3.944700e+05 | NaN | DLCGIVASK |
| USP9X | USP9X\_HUMAN | Probable ubiquitin carboxyl-terminal hydrolase FAF-X | 1771 | 1760 | USP9X | USP9X\_HUMAN | Probable ubiquitin carboxyl-terminal hydrolase FAF-X | 12C(125.0477) | NaN | X | NaN | NaN | NaN | NaN | 3.943468e+05 | 3.943468e+05 | NaN | NaN | GDLLEGANAYHCEK |
| PRDX6 | PRDX6\_HUMAN | Peroxiredoxin-6 | 91 | 85 | PRDX6 | PRDX6\_HUMAN | Peroxiredoxin-6 | 7C(125.0477) | NaN | X | NaN | NaN | NaN | NaN | 3.941710e+05 | 3.941710e+05 | NaN | NaN | DINAYNCEEPTEKLPFPIIDDRNR |
| CARS1 | SYCC\_HUMAN | Cysteine--tRNA ligase, cytoplasmic | 335 | 325 | CARS1 | SYCC\_HUMAN | Cysteine--tRNA ligase, cytoplasmic | 11C(125.0477) | NaN | X | NaN | NaN | NaN | NaN | 3.941316e+05 | 3.941316e+05 | NaN | NaN | ASKPGEPSWPCPWGK |
| SH3BGRL2 | SH3L2\_HUMAN | SH3 domain-binding glutamic acid-rich-like protein 2 | 76 | 75 | SH3BGRL2 | SH3L2\_HUMAN | SH3 domain-binding glutamic acid-rich-like protein 2 | 2C(125.0477) | NaN | X | NaN | NaN | NaN | NaN | 3.938792e+05 | 3.938792e+05 | NaN | NaN | YCGDYDSFFESK |
| A2M | A2MG\_HUMAN | Alpha-2-macroglobulin | 299 | 290 | A2M | A2MG\_HUMAN | Alpha-2-macroglobulin | 10C(57.0215) | X | NaN | NaN | NaN | NaN | NaN | 3.938610e+05 | 3.938610e+05 | NaN | NaN | FSGQLNSHGCFYQQVK |
| ALDH9A1 | AL9A1\_HUMAN | 4-trimethylaminobutyraldehyde dehydrogenase | 45 | 39 | ALDH9A1 | AL9A1\_HUMAN | 4-trimethylaminobutyraldehyde dehydrogenase | 7C(125.0477) | NaN | X | NaN | NaN | NaN | NaN | 3.932748e+05 | 3.932748e+05 | NaN | NaN | VIATFTCSGEK |
| ACTR10 | ARP10\_HUMAN | Actin-related protein 10 | 37 | 37 | ACTR10 | ARP10\_HUMAN | Actin-related protein 10 | 1C(125.0477) | NaN | X | NaN | NaN | NaN | NaN | 3.924222e+05 | 3.826901e+05 | 9.732076e+03 | NaN | CIIPSVIK |
| APIP | MTNB\_HUMAN | Methylthioribulose-1-phosphate dehydratase | 16 | 16 | APIP | MTNB\_HUMAN | Methylthioribulose-1-phosphate dehydratase | 1C(125.0477) | NaN | X | NaN | NaN | NaN | NaN | 3.923918e+05 | 3.923918e+05 | NaN | NaN | CGAQDKEHPR |
| USP9X | USP9X\_HUMAN | Probable ubiquitin carboxyl-terminal hydrolase FAF-X | 1061 | 1059 | USP9X | USP9X\_HUMAN | Probable ubiquitin carboxyl-terminal hydrolase FAF-X | 3C(125.0477) | NaN | X | NaN | NaN | NaN | NaN | 3.913062e+05 | 3.913062e+05 | NaN | NaN | AICLDHAK |
| UBAC1 | UBAC1\_HUMAN | Ubiquitin-associated domain-containing protein 1 | 322 | 313 | UBAC1 | UBAC1\_HUMAN | Ubiquitin-associated domain-containing protein 1 | 10C(125.0477) | NaN | X | NaN | NaN | NaN | NaN | 3.909881e+05 | 3.909881e+05 | NaN | NaN | VNNNQQNAACEWLLGDR |
| PSMA6 | PSA6\_HUMAN | Proteasome subunit alpha type-6 | 115 | 105 | PSMA6 | PSA6\_HUMAN | Proteasome subunit alpha type-6 | 11C(125.0477) | NaN | X | NaN | NaN | NaN | NaN | 3.909834e+05 | 3.909834e+05 | NaN | NaN | YGYEIPVDMLCK |
| MYH9 | MYH9\_HUMAN | Myosin-9 | 91 | 83 | MYH9 | MYH9\_HUMAN | Myosin-9 | 9C(125.0477) | NaN | X | NaN | NaN | NaN | NaN | 3.886637e+05 | 3.886637e+05 | NaN | NaN | VEDMAELTCLNEASVLHNLK |
| GDI2 | GDIB\_HUMAN | Rab GDP dissociation inhibitor beta | 414 | 403 | GDI2 | GDIB\_HUMAN | Rab GDP dissociation inhibitor beta | 12C(125.0477) | NaN | X | NaN | NaN | NaN | NaN | 3.884711e+05 | 3.884711e+05 | NaN | NaN | TYDATTHFETTCDDIKNIYKR |
| DCAF6 | DCAF6\_HUMAN | DDB1- and CUL4-associated factor 6 | 593 | 592 | DCAF6 | DCAF6\_HUMAN | DDB1- and CUL4-associated factor 6 | 2C(125.0477) | NaN | X | NaN | NaN | NaN | NaN | 3.881007e+05 | 3.690660e+05 | 1.903467e+04 | NaN | SCGVPEESASSEK |
| BIRC6 | BIRC6\_HUMAN | Baculoviral IAP repeat-containing protein 6 | 777 | 772 | BIRC6 | BIRC6\_HUMAN | Baculoviral IAP repeat-containing protein 6 | 6C(-34.0809) | NaN | NaN | NaN | NaN | X | NaN | 3.880793e+05 | 3.880793e+05 | NaN | NaN | LNSALCNR |
| CCT3 | TCPG\_HUMAN | T-complex protein 1 subunit gamma | 366 | 355 | CCT3 | TCPG\_HUMAN | T-complex protein 1 subunit gamma | 12C(125.0477) | NaN | X | NaN | NaN | NaN | NaN | 3.873568e+05 | 3.873568e+05 | NaN | NaN | IGDEYFTFITDCKDPK |
| IST1 | IST1\_HUMAN | IST1 homolog | 125 | 119 | IST1 | IST1\_HUMAN | IST1 homolog | 7C(125.0477) | NaN | X | NaN | NaN | NaN | NaN | 3.872101e+05 | NaN | 3.872101e+05 | NaN | IVADQLCAK |
| CLTC | CLH1\_HUMAN | Clathrin heavy chain 1 | 748, 753 | 747 | CLTC | CLH1\_HUMAN | Clathrin heavy chain 1 | 2C(15.9949) 7C(15.9949) | NaN | NaN | X | NaN | NaN | NaN | 3.856515e+05 | 3.856515e+05 | NaN | NaN | ICRESNCYDPER |
| CTPS1 | PYRG1\_HUMAN | CTP synthase 1 | 299 | 295 | CTPS1 | PYRG1\_HUMAN | CTP synthase 1 | 5C(125.0477) | NaN | X | NaN | NaN | NaN | NaN | 3.849993e+05 | 3.849993e+05 | NaN | NaN | LLETCSIALVGK |
| RAN | RAN\_HUMAN | GTP-binding nuclear protein Ran | 120, 112 | 111 | RAN | RAN\_HUMAN | GTP-binding nuclear protein Ran | 10C(57.0215) 2C(125.0477) | X | X | NaN | NaN | NaN | NaN | 3.848575e+05 | 3.848575e+05 | NaN | NaN | VCENIPIVLCGNKVDIKDR |
| FASN | FAS\_HUMAN | Fatty acid synthase | 2273 | 2264 | FASN | FAS\_HUMAN | Fatty acid synthase | 10C(125.0477) | NaN | X | NaN | NaN | NaN | NaN | 3.846531e+05 | 3.846531e+05 | NaN | NaN | LSIPTYGLQCTR |
| ATP6V1G1 | VATG1\_HUMAN | V-type proton ATPase subunit G 1 | 69, 69 | 67 | ATP6V1G1 | VATG1\_HUMAN | V-type proton ATPase subunit G 1 | 3C(125.0477) 3C(31.9988) | NaN | X | NaN | X | NaN | NaN | 3.835183e+05 | 3.835183e+05 | NaN | NaN | GSCSTEVEKETQEK |
| PNPLA6 | PLPL6\_HUMAN | Patatin-like phospholipase domain-containing protein 6 | 208 | 195 | PNPLA6 | PLPL6\_HUMAN | Patatin-like phospholipase domain-containing protein 6 | 14C(125.0477) | NaN | X | NaN | NaN | NaN | NaN | 3.819528e+05 | 3.819528e+05 | NaN | NaN | VLGHFEKPLFLELCR |
| DDB1 | DDB1\_HUMAN | DNA damage-binding protein 1 | 260 | 255 | DDB1 | DDB1\_HUMAN | DNA damage-binding protein 1 | 6C(125.0477) | NaN | X | NaN | NaN | NaN | NaN | 3.816313e+05 | 3.816313e+05 | 0.000000e+00 | NaN | QSTIVCHNR |
| UROS | HEM4\_HUMAN | Uroporphyrinogen-III synthase | 73 | 66 | UROS | HEM4\_HUMAN | Uroporphyrinogen-III synthase | 8C(125.0477) | NaN | X | NaN | NaN | NaN | NaN | 3.789275e+05 | 3.428086e+05 | 3.611887e+04 | NaN | AVEAAELCLEQNNK |
| GDI2 | GDIB\_HUMAN | Rab GDP dissociation inhibitor beta | 282, 282 | 279 | GDI2 | GDIB\_HUMAN | Rab GDP dissociation inhibitor beta | 4C(125.0477) 4C(31.9988) | NaN | X | NaN | X | NaN | NaN | 3.784321e+05 | 3.784321e+05 | NaN | NaN | QLICDPSYVK |
| PAFAH1B1 | LIS1\_HUMAN | Platelet-activating factor acetylhydrolase IB subunit beta | 356 | 352 | PAFAH1B1 | LIS1\_HUMAN | Platelet-activating factor acetylhydrolase IB subunit beta | 5C(125.0477) | NaN | X | NaN | NaN | NaN | NaN | 3.783481e+05 | 3.429909e+05 | 3.535721e+04 | NaN | FILSCADDK |
| GDI2 | GDIB\_HUMAN | Rab GDP dissociation inhibitor beta | 317, 302 | 300 | GDI2 | GDIB\_HUMAN | Rab GDP dissociation inhibitor beta | 18C(15.9949) 3C(15.9949) | NaN | NaN | X | NaN | NaN | NaN | 3.775260e+05 | 3.775260e+05 | NaN | NaN | VICILSHPIKNTNDANSCQIIIPQNQVNR |
| CLU | CLUS\_HUMAN | Clusterin | 313, 313 | 307 | CLU | CLUS\_HUMAN | Clusterin | 7C(125.0477) 7C(57.0215) | X | X | NaN | NaN | NaN | NaN | 3.771650e+05 | NaN | 3.771650e+05 | NaN | EILSVDCSTNNPSQAK |
| ADK | ADK\_HUMAN | Adenosine kinase | 106 | 100 | ADK | ADK\_HUMAN | Adenosine kinase | 7C(125.0477) | NaN | X | NaN | NaN | NaN | NaN | 3.761231e+05 | 3.761231e+05 | NaN | NaN | AATFFGCIGIDK |
| CMBL | CMBL\_HUMAN | Carboxymethylenebutenolidase homolog | 222 | 219 | CMBL | CMBL\_HUMAN | Carboxymethylenebutenolidase homolog | 4C(125.0477) | NaN | X | NaN | NaN | NaN | NaN | 3.756657e+05 | 3.756657e+05 | NaN | NaN | REDCSPADKPYIDEAR |
| PFAS | PUR4\_HUMAN | Phosphoribosylformylglycinamidine synthase | 512 | 508 | PFAS | PUR4\_HUMAN | Phosphoribosylformylglycinamidine synthase | 5C(125.0477) | NaN | X | NaN | NaN | NaN | NaN | 3.751070e+05 | 3.751070e+05 | NaN | NaN | GNPICSLHDQGAGGNGNVLK |
| USP15 | UBP15\_HUMAN | Ubiquitin carboxyl-terminal hydrolase 15 | 127 | 126 | USP15 | UBP15\_HUMAN | Ubiquitin carboxyl-terminal hydrolase 15 | 2C(125.0477) | NaN | X | NaN | NaN | NaN | NaN | 3.746077e+05 | 3.746077e+05 | NaN | NaN | HCKVEVYLTELK |
| PLIN3 | PLIN3\_HUMAN | Perilipin-3 | 60 | 58 | PLIN3 | PLIN3\_HUMAN | Perilipin-3 | 3C(125.0477) | NaN | X | NaN | NaN | NaN | NaN | 3.742936e+05 | 2.989612e+05 | 7.533245e+04 | NaN | TVCDAAEK |
| RRM1 | RIR1\_HUMAN | Ribonucleoside-diphosphate reductase large subunit | 492 | 480 | RRM1 | RIR1\_HUMAN | Ribonucleoside-diphosphate reductase large subunit | 13C(125.0477) | NaN | X | NaN | NaN | NaN | NaN | 3.733801e+05 | 3.733801e+05 | NaN | NaN | IIDINYYPVPEACLSNK |
| ANKRD28 | ANR28\_HUMAN | Serine/threonine-protein phosphatase 6 regulatory ankyrin repeat subunit A | 85 | 81 | ANKRD28 | ANR28\_HUMAN | Serine/threonine-protein phosphatase 6 regulatory ankyrin repeat subunit A | 5C(125.0477) | NaN | X | NaN | NaN | NaN | NaN | 3.718566e+05 | 3.718566e+05 | NaN | NaN | AVASCSEEAVQVLLK |
| GRHPR | GRHPR\_HUMAN | Glyoxylate reductase/hydroxypyruvate reductase | 57 | 48 | GRHPR | GRHPR\_HUMAN | Glyoxylate reductase/hydroxypyruvate reductase | 10C(125.0477) | NaN | X | NaN | NaN | NaN | NaN | 3.707293e+05 | 3.707293e+05 | NaN | NaN | GVAGAHGLLCLLSDHVDK |
| HSP90AA1 | HS90A\_HUMAN | Heat shock protein HSP 90-alpha | 572 | 566 | HSP90AA1 | HS90A\_HUMAN | Heat shock protein HSP 90-alpha | 7C(125.0477) | NaN | X | NaN | NaN | NaN | NaN | 3.700401e+05 | 3.700401e+05 | NaN | NaN | TKFENLCK |
| PPP5C | PPP5\_HUMAN | Serine/threonine-protein phosphatase 5 | 11 | 9 | PPP5C | PPP5\_HUMAN | Serine/threonine-protein phosphatase 5 | 3C(125.0477) | NaN | X | NaN | NaN | NaN | NaN | 3.695160e+05 | 3.695160e+05 | NaN | NaN | TECAEPPRDEPPADGALKR |
| THOP1 | THOP1\_HUMAN | Thimet oligopeptidase | 175 | 170 | THOP1 | THOP1\_HUMAN | Thimet oligopeptidase | 6C(125.0477) | NaN | X | NaN | NaN | NaN | NaN | 3.694347e+05 | 3.694347e+05 | NaN | NaN | KLSLLCIDFNK |
| SPTA1 | SPTA1\_HUMAN | Spectrin alpha chain, erythrocytic 1 | 799 | 789 | SPTA1 | SPTA1\_HUMAN | Spectrin alpha chain, erythrocytic 1 | 11C(125.0477) | NaN | X | NaN | NaN | NaN | NaN | 3.689150e+05 | 2.884871e+05 | 8.042791e+04 | NaN | LLDLLHLQLICR |
| COPS2 | CSN2\_HUMAN | COP9 signalosome complex subunit 2 | 179 | 174 | COPS2 | CSN2\_HUMAN | COP9 signalosome complex subunit 2 | 6C(125.0477) | NaN | X | NaN | NaN | NaN | NaN | 3.683375e+05 | 3.683375e+05 | NaN | NaN | QLHQSCQTDDGEDDLK |
| GBE1 | GLGB\_HUMAN | 1,4-alpha-glucan-branching enzyme | 473 | 473 | GBE1 | GLGB\_HUMAN | 1,4-alpha-glucan-branching enzyme | 1C(125.0477) | NaN | X | NaN | NaN | NaN | NaN | 3.671348e+05 | 3.671348e+05 | NaN | NaN | CIAYAESHDQALVGDK |
| NPL | NPL\_HUMAN | N-acetylneuraminate lyase | 185 | 174 | NPL | NPL\_HUMAN | N-acetylneuraminate lyase | 12C(125.0477) | NaN | X | NaN | NaN | NaN | NaN | 3.665931e+05 | 3.665931e+05 | NaN | NaN | FSDTDLLDFGQCVDQNR |
| UBE2V1 | UB2V1\_HUMAN | Ubiquitin-conjugating enzyme E2 variant 1 | 144 | 136 | UBE2V1 | UB2V1\_HUMAN | Ubiquitin-conjugating enzyme E2 variant 1 | 9C(125.0477) | NaN | X | NaN | NaN | NaN | NaN | 3.649850e+05 | NaN | 3.649850e+05 | NaN | LPQPPEGQCYSN |
| AARSD1 | AASD1\_HUMAN | Alanyl-tRNA editing protein Aarsd1 | 266 | 266 | AARSD1 | AASD1\_HUMAN | Alanyl-tRNA editing protein Aarsd1 | 1C(125.0477) | NaN | X | NaN | NaN | NaN | NaN | 3.638991e+05 | 3.638991e+05 | NaN | NaN | CGAEDHVEAVK |
| CD59 | CD59\_HUMAN | CD59 glycoprotein | 64, 64, 70, 70 | 64 | CD59 | CD59\_HUMAN | CD59 glycoprotein | 1C(15.9949) 1C(31.9988) 7C(15.9949) 7C(31.9988) | NaN | NaN | X | X | NaN | NaN | 3.638516e+05 | 3.638516e+05 | NaN | NaN | CWKFEHCNFNDVTTR |
| LANCL2 | LANC2\_HUMAN | LanC-like protein 2 | 248 | 248 | LANCL2 | LANC2\_HUMAN | LanC-like protein 2 | 1C(125.0477) | NaN | X | NaN | NaN | NaN | NaN | 3.618478e+05 | 3.618478e+05 | NaN | NaN | CPLLYQWHR |
| WDR44 | WDR44\_HUMAN | WD repeat-containing protein 44 | 716 | 716 | WDR44 | WDR44\_HUMAN | WD repeat-containing protein 44 | 1C(125.0477) | NaN | X | NaN | NaN | NaN | NaN | 3.608271e+05 | 3.608271e+05 | NaN | NaN | CIFYDTEHLK |
| PFAS | PUR4\_HUMAN | Phosphoribosylformylglycinamidine synthase | 1044 | 1039 | PFAS | PUR4\_HUMAN | Phosphoribosylformylglycinamidine synthase | 6C(125.0477) | NaN | X | NaN | NaN | NaN | NaN | 3.591078e+05 | 3.591078e+05 | NaN | NaN | MGPSYCLPPTFPK |
| USP24 | UBP24\_HUMAN | Ubiquitin carboxyl-terminal hydrolase 24 | 153 | 153 | USP24 | UBP24\_HUMAN | Ubiquitin carboxyl-terminal hydrolase 24 | 1C(125.0477) | NaN | X | NaN | NaN | NaN | NaN | 3.587536e+05 | 3.587536e+05 | NaN | NaN | CLLASTYLAR |
| AP2B1 | AP2B1\_HUMAN | AP-2 complex subunit beta | 857, 857 | 856 | AP2B1 | AP2B1\_HUMAN | AP-2 complex subunit beta | 2C(125.0477) 2C(57.0215) | X | X | NaN | NaN | NaN | NaN | 3.585911e+05 | 3.585911e+05 | NaN | NaN | ECHLNADTVSSK |
| PSMG1 | PSMG1\_HUMAN | Proteasome assembly chaperone 1 | 235 | 228 | PSMG1 | PSMG1\_HUMAN | Proteasome assembly chaperone 1 | 8C(125.0477) | NaN | X | NaN | NaN | NaN | NaN | 3.581265e+05 | 3.581265e+05 | NaN | NaN | IPAILYLCYTDVMK |
| KRT10 | K1C10\_HUMAN | Keratin, type I cytoskeletal 10 | 427 | 423 | KRT10 | K1C10\_HUMAN | Keratin, type I cytoskeletal 10 | 5C(125.0477) | NaN | X | NaN | NaN | NaN | NaN | 3.564201e+05 | NaN | 3.417107e+05 | 14709.4190 | AETECQNTEYQQLLDIK |
| DDI2 | DDI2\_HUMAN | Protein DDI1 homolog 2 | 332 | 330 | DDI2 | DDI2\_HUMAN | Protein DDI1 homolog 2 | 3C(125.0477) | NaN | X | NaN | NaN | NaN | NaN | 3.545602e+05 | 2.547891e+05 | 9.977116e+04 | NaN | HQCSIDLK |
| TKFC | TKFC\_HUMAN | Triokinase/FMN cyclase | 155 | 153 | TKFC | TKFC\_HUMAN | Triokinase/FMN cyclase | 3C(125.0477) | NaN | X | NaN | NaN | NaN | NaN | 3.530483e+05 | 3.530483e+05 | NaN | NaN | GLCGTVLIHK |
| ALB | ALBU\_HUMAN | Albumin | 289 | 287 | ALB | ALBU\_HUMAN | Albumin | 3C(57.0215) | X | NaN | NaN | NaN | NaN | NaN | 3.526441e+05 | 3.526441e+05 | NaN | NaN | YICENQDSISSK |
| HSPA4 | HSP74\_HUMAN | Heat shock 70 kDa protein 4 | 167 | 155 | HSPA4 | HSP74\_HUMAN | Heat shock 70 kDa protein 4 | 13C(125.0477) | NaN | X | NaN | NaN | NaN | NaN | 3.522518e+05 | 3.522518e+05 | NaN | NaN | SVMDATQIAGLNCLR |
| ANK1 | ANK1\_HUMAN | Ankyrin-1 | 1174, 1174 | 1172 | ANK1 | ANK1\_HUMAN | Ankyrin-1 | 3C(125.0477) 3C(31.9988) | NaN | X | NaN | X | NaN | NaN | 3.518070e+05 | 3.518070e+05 | NaN | NaN | LLCSVIGGTDQAQWEDITGTTK |
| SPTA1 | SPTA1\_HUMAN | Spectrin alpha chain, erythrocytic 1 | 324 | 322 | SPTA1 | SPTA1\_HUMAN | Spectrin alpha chain, erythrocytic 1 | 3C(125.0477) | NaN | X | NaN | NaN | NaN | NaN | 3.506982e+05 | NaN | 3.506982e+05 | NaN | ELCAKAEK |
| CD55 | DAF\_HUMAN | Complement decay-accelerating factor | 98 | 97 | CD55 | DAF\_HUMAN | Complement decay-accelerating factor | 2C(125.0477) | NaN | X | NaN | NaN | NaN | NaN | 3.506982e+05 | NaN | 3.506982e+05 | NaN | SCEVPTR |
| AHCY | SAHH\_HUMAN | Adenosylhomocysteinase | 421 | 413 | AHCY | SAHH\_HUMAN | Adenosylhomocysteinase | 9C(125.0477) | NaN | X | NaN | NaN | NaN | NaN | 3.504502e+05 | 3.504502e+05 | NaN | NaN | QAQYLGMSCDGPFKPDHYR |
| CUL2 | CUL2\_HUMAN | Cullin-2 | 465 | 463 | CUL2 | CUL2\_HUMAN | Cullin-2 | 3C(125.0477) | NaN | X | NaN | NaN | NaN | NaN | 3.502755e+05 | 3.125467e+05 | 3.772883e+04 | NaN | QACGYEFTSK |
| FMO5 | FMO5\_HUMAN | Flavin-containing monooxygenase 5 | 468 | 459 | FMO5 | FMO5\_HUMAN | Flavin-containing monooxygenase 5 | 10C(31.9988) | NaN | NaN | NaN | X | NaN | NaN | 3.494731e+05 | 3.494731e+05 | NaN | NaN | LALHLLLGPCTPIHYR |
| NUDT14 | NUD14\_HUMAN | Uridine diphosphate glucose pyrophosphatase NUDT14 | 104 | 88 | NUDT14 | NUD14\_HUMAN | Uridine diphosphate glucose pyrophosphatase NUDT14 | 17C(15.9949) | NaN | NaN | X | NaN | NaN | NaN | 3.494380e+05 | 3.494380e+05 | NaN | NaN | ELQPALPGSAGVTVELCAGLVDQPGLSLEEVACK |
| TNPO1 | TNPO1\_HUMAN | Transportin-1 | 103 | 101 | TNPO1 | TNPO1\_HUMAN | Transportin-1 | 3C(125.0477) | NaN | X | NaN | NaN | NaN | NaN | 3.493708e+05 | 3.493708e+05 | NaN | NaN | SECLNNIGDSSPLIR |
| DCTN4 | DCTN4\_HUMAN | Dynactin subunit 4 | 30 | 22 | DCTN4 | DCTN4\_HUMAN | Dynactin subunit 4 | 9C(125.0477) | NaN | X | NaN | NaN | NaN | NaN | 3.490519e+05 | 3.490519e+05 | NaN | NaN | APLSQLYFCR |
| PPME1 | PPME1\_HUMAN | Protein phosphatase methylesterase 1 | 238, 238 | 237 | PPME1 | PPME1\_HUMAN | Protein phosphatase methylesterase 1 | 2C(125.0477) 2C(31.9988) | NaN | X | NaN | X | NaN | NaN | 3.482316e+05 | 3.482316e+05 | NaN | NaN | QCEGITSPEGSK |
| GLO1 | LGUL\_HUMAN | Lactoylglutathione lyase | 139 | 124 | GLO1 | LGUL\_HUMAN | Lactoylglutathione lyase | 16C(125.0477) | NaN | X | NaN | NaN | NaN | NaN | 3.481501e+05 | 3.481501e+05 | NaN | NaN | GFGHIGIAVPDVYSACK |
| MCTS1 | MCTS1\_HUMAN | Malignant T-cell-amplified sequence 1 | 14 | 5 | MCTS1 | MCTS1\_HUMAN | Malignant T-cell-amplified sequence 1 | 10C(125.0477) | NaN | X | NaN | NaN | NaN | NaN | 3.481024e+05 | 3.481024e+05 | NaN | NaN | FDEKENVSNCIQLK |
| MTHFD1 | C1TC\_HUMAN | C-1-tetrahydrofolate synthase, cytoplasmic | 785 | 785 | MTHFD1 | C1TC\_HUMAN | C-1-tetrahydrofolate synthase, cytoplasmic | 1C(125.0477) | NaN | X | NaN | NaN | NaN | NaN | 3.478436e+05 | 3.478436e+05 | NaN | NaN | CTHWAEGGK |
| STIP1 | STIP1\_HUMAN | Stress-induced-phosphoprotein 1 | 370 | 365 | STIP1 | STIP1\_HUMAN | Stress-induced-phosphoprotein 1 | 6C(125.0477) | NaN | X | NaN | NaN | NaN | NaN | 3.475928e+05 | 3.475928e+05 | NaN | NaN | NKGNECFQK |
| MARK3 | MARK3\_HUMAN | MAP/microtubule affinity-regulating kinase 3 | 169 | 161 | MARK3 | MARK3\_HUMAN | MAP/microtubule affinity-regulating kinase 3 | 9C(125.0477) | NaN | X | NaN | NaN | NaN | NaN | 3.471313e+05 | 3.361475e+05 | 1.098378e+04 | NaN | QIVSAVQYCHQK |
| PSMA4 | PSA4\_HUMAN | Proteasome subunit alpha type-4 | 74 | 68 | PSMA4 | PSA4\_HUMAN | Proteasome subunit alpha type-4 | 7C(125.0477) | NaN | X | NaN | NaN | NaN | NaN | 3.467701e+05 | 3.467701e+05 | NaN | NaN | LNEDMACSVAGITSDANVLTNELR |
| CCT3 | TCPG\_HUMAN | T-complex protein 1 subunit gamma | 213 | 204 | CCT3 | TCPG\_HUMAN | T-complex protein 1 subunit gamma | 10C(125.0477) | NaN | X | NaN | NaN | NaN | NaN | 3.467532e+05 | 3.467532e+05 | NaN | NaN | IPGGIIEDSCVLR |
| ADK | ADK\_HUMAN | Adenosine kinase | 353 | 350 | ADK | ADK\_HUMAN | Adenosine kinase | 4C(125.0477) | NaN | X | NaN | NaN | NaN | NaN | 3.466149e+05 | 3.466149e+05 | NaN | NaN | RTGCTFPEKPDFH |
| DNM2 | DYN2\_HUMAN | Dynamin-2 | 27 | 16 | DNM2 | DYN2\_HUMAN | Dynamin-2 | 12C(125.0477) | NaN | X | NaN | NaN | NaN | NaN | 3.452513e+05 | 3.452513e+05 | NaN | NaN | LQDAFSSIGQSCHLDLPQIAVVGGQSAGK |
| PEPD | PEPD\_HUMAN | Xaa-Pro dipeptidase | 158 | 149 | PEPD | PEPD\_HUMAN | Xaa-Pro dipeptidase | 10C(125.0477) | NaN | X | NaN | NaN | NaN | NaN | 3.446605e+05 | 3.114841e+05 | 3.317648e+04 | NaN | GVNTDSGSVCR |
| NAPA | SNAA\_HUMAN | Alpha-soluble NSF attachment protein | 42 | 38 | NAPA | SNAA\_HUMAN | Alpha-soluble NSF attachment protein | 5C(125.0477) | NaN | X | NaN | NaN | NaN | NaN | 3.444495e+05 | 4.826289e+04 | 2.961866e+05 | NaN | IEEACEIYAR |
| DAZAP1 | DAZP1\_HUMAN | DAZ-associated protein 1 | 63 | 58 | DAZAP1 | DAZP1\_HUMAN | DAZ-associated protein 1 | 6C(125.0477) | NaN | X | NaN | NaN | NaN | NaN | 3.440748e+05 | 3.440748e+05 | NaN | NaN | FKDPNCVGTVLASRPHTLDGR |
| UBE2O | UBE2O\_HUMAN | (E3-independent) E2 ubiquitin-conjugating enzyme | 101 | 97 | UBE2O | UBE2O\_HUMAN | (E3-independent) E2 ubiquitin-conjugating enzyme | 5C(125.0477) | NaN | X | NaN | NaN | NaN | NaN | 3.423745e+05 | 3.239969e+05 | 1.837760e+04 | NaN | GSSGCSEAGGAGHEEGR |
| PGLS | 6PGL\_HUMAN | 6-phosphogluconolactonase | 237 | 236 | PGLS | 6PGL\_HUMAN | 6-phosphogluconolactonase | 2C(125.0477) | NaN | X | NaN | NaN | NaN | NaN | 3.422119e+05 | 3.422119e+05 | NaN | NaN | LCWFLDEAAAR |
| MCTS1 | MCTS1\_HUMAN | Malignant T-cell-amplified sequence 1 | 14 | 4 | MCTS1 | MCTS1\_HUMAN | Malignant T-cell-amplified sequence 1 | 11C(125.0477) | NaN | X | NaN | NaN | NaN | NaN | 3.417604e+05 | 3.417604e+05 | NaN | NaN | KFDEKENVSNCIQLK |
| SHPK | SHPK\_HUMAN | Sedoheptulokinase | 105 | 101 | SHPK | SHPK\_HUMAN | Sedoheptulokinase | 5C(125.0477) | NaN | X | NaN | NaN | NaN | NaN | 3.417338e+05 | 3.417338e+05 | NaN | NaN | TGQGCEWTEGGITPVFEPR |
| GCLM | GSH0\_HUMAN | Glutamate--cysteine ligase regulatory subunit | 46, 35 | 35 | GCLM | GSH0\_HUMAN | Glutamate--cysteine ligase regulatory subunit | 12C(125.0477) 1C(125.0477) | NaN | X | NaN | NaN | NaN | NaN | 3.405789e+05 | 3.405789e+05 | NaN | NaN | CPSTHSEELHDCIQK |
| APRT | APT\_HUMAN | Adenine phosphoribosyltransferase | 153 | 146 | APRT | APT\_HUMAN | Adenine phosphoribosyltransferase | 8C(125.0477) | NaN | X | NaN | NaN | NaN | NaN | 3.401078e+05 | 3.401078e+05 | NaN | NaN | LQAEVLECVSLVELTSLK |
| EIF5A | IF5A1\_HUMAN | Eukaryotic translation initiation factor 5A-1 | 73 | 69 | EIF5A | IF5A1\_HUMAN | Eukaryotic translation initiation factor 5A-1 | 5C(125.0477) | NaN | X | NaN | NaN | NaN | NaN | 3.398225e+05 | 3.398225e+05 | NaN | NaN | YEDICPSTHNMDVPNIKR |
| PGK1 | PGK1\_HUMAN | Phosphoglycerate kinase 1 | 99, 99 | 98 | PGK1 | PGK1\_HUMAN | Phosphoglycerate kinase 1 | 2C(125.0477) 2C(57.0215) | X | X | NaN | NaN | NaN | NaN | 3.396744e+05 | 3.396744e+05 | NaN | NaN | DCVGPEVEK |
| DCTN1 | DCTN1\_HUMAN | Dynactin subunit 1 | 791 | 786 | DCTN1 | DCTN1\_HUMAN | Dynactin subunit 1 | 6C(125.0477) | NaN | X | NaN | NaN | NaN | NaN | 3.391634e+05 | 3.391634e+05 | NaN | NaN | DLETSCSDIR |
| TBCB | TBCB\_HUMAN | Tubulin-folding cofactor B | 83 | 67 | TBCB | TBCB\_HUMAN | Tubulin-folding cofactor B | 17C(125.0477) | NaN | X | NaN | NaN | NaN | NaN | 3.387953e+05 | 3.387953e+05 | NaN | NaN | LDQEDALLGSYPVDDGCR |
| TBC1D15 | TBC15\_HUMAN | TBC1 domain family member 15 | 605 | 598 | TBC1D15 | TBC15\_HUMAN | TBC1 domain family member 15 | 8C(125.0477) | NaN | X | NaN | NaN | NaN | NaN | 3.384790e+05 | 3.384790e+05 | NaN | NaN | IDVEDILCK |
| CAND1 | CAND1\_HUMAN | Cullin-associated NEDD8-dissociated protein 1 | 92 | 81 | CAND1 | CAND1\_HUMAN | Cullin-associated NEDD8-dissociated protein 1 | 12C(125.0477) | NaN | X | NaN | NaN | NaN | NaN | 3.364719e+05 | 3.364719e+05 | NaN | NaN | EYQVETIVDTLCTNMLSDKEQLR |
| NIT2 | NIT2\_HUMAN | Omega-amidase NIT2 | 76 | 75 | NIT2 | NIT2\_HUMAN | Omega-amidase NIT2 | 2C(125.0477) | NaN | X | NaN | NaN | NaN | NaN | 3.360435e+05 | 3.360435e+05 | NaN | NaN | ECSIYLIGGSIPEEDAGK |
| CYB5R3 | NB5R3\_HUMAN | NADH-cytochrome b5 reductase 3 | 204 | 197 | CYB5R3 | NB5R3\_HUMAN | NADH-cytochrome b5 reductase 3 | 8C(125.0477) | NaN | X | NaN | NaN | NaN | NaN | 3.348532e+05 | 3.348532e+05 | NaN | NaN | DPDDHTVCHLLFANQTEK |
| CFL1 | COF1\_HUMAN | Cofilin-1 | 139 | 128 | CFL1 | COF1\_HUMAN | Cofilin-1 | 12C(125.0477) | NaN | X | NaN | NaN | NaN | NaN | 3.335802e+05 | 3.335802e+05 | NaN | NaN | LTGIKHELQANCYEEVK |
| CAT | CATA\_HUMAN | Catalase | 460, 460 | 458 | CAT | CATA\_HUMAN | Catalase | 3C(-34.0809) 3C(125.0477) | NaN | X | NaN | NaN | X | NaN | 3.323795e+05 | 3.323795e+05 | NaN | NaN | RLCENIAGHLK |
| PCBP1 | PCBP1\_HUMAN | Poly(rC)-binding protein 1 | 54 | 47 | PCBP1 | PCBP1\_HUMAN | Poly(rC)-binding protein 1 | 8C(125.0477) | NaN | X | NaN | NaN | NaN | NaN | 3.317765e+05 | 1.304693e+05 | 2.013072e+05 | NaN | INISEGNCPER |
| RBX1 | RBX1\_HUMAN | E3 ubiquitin-protein ligase RBX1 | 94 | 92 | RBX1 | RBX1\_HUMAN | E3 ubiquitin-protein ligase RBX1 | 3C(125.0477) | NaN | X | NaN | NaN | NaN | NaN | 3.298382e+05 | NaN | 3.298382e+05 | NaN | QVCPLDNR |
| NUDT2 | AP4A\_HUMAN | Bis(5'-nucleosyl)-tetraphosphatase [asymmetrical] | 143 | 133 | NUDT2 | AP4A\_HUMAN | Bis(5'-nucleosyl)-tetraphosphatase [asymmetrical] | 11C(125.0477) | NaN | X | NaN | NaN | NaN | NaN | 3.297497e+05 | 3.297497e+05 | NaN | NaN | AALQEGHQFLCSIEA |
| MINPP1 | MINP1\_HUMAN | Multiple inositol polyphosphate phosphatase 1 | 232 | 228 | MINPP1 | MINP1\_HUMAN | Multiple inositol polyphosphate phosphatase 1 | 5C(125.0477) | NaN | X | NaN | NaN | NaN | NaN | 3.296546e+05 | NaN | 3.296546e+05 | NaN | FFDHCEK |
| C2CD2L | C2C2L\_HUMAN | Phospholipid transfer protein C2CD2L | 154 | 154 | C2CD2L | C2C2L\_HUMAN | Phospholipid transfer protein C2CD2L | 1C(125.0477) | NaN | X | NaN | NaN | NaN | NaN | 3.292620e+05 | NaN | 2.957258e+05 | 33536.2770 | CQLSAEEVR |
| CCT8 | TCPQ\_HUMAN | T-complex protein 1 subunit theta | 430 | 422 | CCT8 | TCPQ\_HUMAN | T-complex protein 1 subunit theta | 9C(125.0477) | NaN | X | NaN | NaN | NaN | NaN | 3.289579e+05 | 3.289579e+05 | NaN | NaN | QITSYGETCPGLEQYAIK |
| OXSR1 | OXSR1\_HUMAN | Serine/threonine-protein kinase | 53 | 53 | OXSR1 | OXSR1\_HUMAN | Serine/threonine-protein kinase | 1C(125.0477) | NaN | X | NaN | NaN | NaN | NaN | 3.279309e+05 | 3.224921e+05 | 5.438814e+03 | NaN | CQTSMDELLK |
| ADD2 | ADDB\_HUMAN | Beta-adducin | 277, 277, 282, 282 | 277 | ADD2 | ADDB\_HUMAN | Beta-adducin | 1C(125.0477) 1C(15.9949) 6C(125.0477) 6C(15.9949) | NaN | X | X | NaN | NaN | NaN | 3.277623e+05 | NaN | 3.277623e+05 | NaN | CLGPTCKILVLR |
| DGKA | DGKA\_HUMAN | Diacylglycerol kinase alpha | 682 | 682 | DGKA | DGKA\_HUMAN | Diacylglycerol kinase alpha | 1C(125.0477) | NaN | X | NaN | NaN | NaN | NaN | 3.273865e+05 | 2.867421e+05 | 4.064449e+04 | NaN | CSEITFHTTK |
| ALDOA | ALDOA\_HUMAN | Fructose-bisphosphate aldolase A | 202 | 201 | ALDOA | ALDOA\_HUMAN | Fructose-bisphosphate aldolase A | 2C(125.0477) | NaN | X | NaN | NaN | NaN | NaN | 3.273077e+05 | 3.273077e+05 | NaN | NaN | RCQYVTEK |
| SUGT1 | SGT1\_HUMAN | Protein SGT1 homolog | 88 | 85 | SUGT1 | SGT1\_HUMAN | Protein SGT1 homolog | 4C(125.0477) | NaN | X | NaN | NaN | NaN | NaN | 3.267013e+05 | 3.267013e+05 | NaN | NaN | KGICEYHEK |
| LCP1 | PLSL\_HUMAN | Plastin-2 | 460 | 456 | LCP1 | PLSL\_HUMAN | Plastin-2 | 5C(125.0477) | NaN | X | NaN | NaN | NaN | NaN | 3.256291e+05 | 3.256291e+05 | NaN | NaN | KLENCNYAVELGK |
| HSP90AB1 | HS90B\_HUMAN | Heat shock protein HSP 90-beta | 412 | 412 | HSP90AB1 | HS90B\_HUMAN | Heat shock protein HSP 90-beta | 1C(125.0477) | NaN | X | NaN | NaN | NaN | NaN | 3.248503e+05 | 3.248503e+05 | NaN | NaN | CLELFSELAEDKENYK |
| ACAT2 | THIC\_HUMAN | Acetyl-CoA acetyltransferase, cytosolic | 360 | 342 | ACAT2 | THIC\_HUMAN | Acetyl-CoA acetyltransferase, cytosolic | 19C(125.0477) | NaN | X | NaN | NaN | NaN | NaN | 3.246380e+05 | 3.246380e+05 | NaN | NaN | VNIEGGAIALGHPLGASGCR |
| ABCB6 | ABCB6\_HUMAN | ATP-binding cassette sub-family B member 6 | 801 | 799 | ABCB6 | ABCB6\_HUMAN | ATP-binding cassette sub-family B member 6 | 3C(125.0477) | NaN | X | NaN | NaN | NaN | NaN | 3.245162e+05 | NaN | 1.312500e+05 | 193266.2000 | DGCIVER |
| RUVBL2 | RUVB2\_HUMAN | RuvB-like 2 | 227 | 224 | RUVBL2 | RUVB2\_HUMAN | RuvB-like 2 | 4C(125.0477) | NaN | X | NaN | NaN | NaN | NaN | 3.238820e+05 | 2.055690e+05 | 1.183130e+05 | NaN | FVQCPDGELQK |
| SCRN3 | SCRN3\_HUMAN | Secernin-3 | 143 | 137 | SCRN3 | SCRN3\_HUMAN | Secernin-3 | 7C(125.0477) | NaN | X | NaN | NaN | NaN | NaN | 3.237192e+05 | 3.237192e+05 | NaN | NaN | YGQGGNCTEGR |
| C2CD2L | C2C2L\_HUMAN | Phospholipid transfer protein C2CD2L | 115 | 109 | C2CD2L | C2C2L\_HUMAN | Phospholipid transfer protein C2CD2L | 7C(125.0477) | NaN | X | NaN | NaN | NaN | NaN | 3.227328e+05 | NaN | 3.227328e+05 | NaN | ALNEQACR |
| HBB | HBB\_HUMAN | Hemoglobin subunit beta | 94 | 84 | HBB | HBB\_HUMAN | Hemoglobin subunit beta | 11C(125.0477) | NaN | X | NaN | NaN | NaN | NaN | 3.219623e+05 | NaN | NaN | 321962.3400 | GTFATLSELHCDKLHVDPENFR |
| GDI1 | GDIA\_HUMAN | Rab GDP dissociation inhibitor alpha | 202 | 194 | GDI1 | GDIA\_HUMAN | Rab GDP dissociation inhibitor alpha | 9C(125.0477) | NaN | X | NaN | NaN | NaN | NaN | 3.217663e+05 | 3.170719e+05 | 4.694448e+03 | NaN | TDDYLDQPCLETVNR |
| DHX9 | DHX9\_HUMAN | ATP-dependent RNA helicase A | 1099 | 1090 | DHX9 | DHX9\_HUMAN | ATP-dependent RNA helicase A | 10C(125.0477) | NaN | X | NaN | NaN | NaN | NaN | 3.214159e+05 | 3.214159e+05 | NaN | NaN | LQISHEAAACITGLR |
| ATP2B4 | AT2B4\_HUMAN | Plasma membrane calcium-transporting ATPase 4 | 24 | 19 | ATP2B4 | AT2B4\_HUMAN | Plasma membrane calcium-transporting ATPase 4 | 6C(125.0477) | NaN | X | NaN | NaN | NaN | NaN | 3.209047e+05 | 3.209047e+05 | NaN | NaN | EGDFGCTVMELR |
| AP2B1 | AP2B1\_HUMAN | AP-2 complex subunit beta | 123 | 115 | AP2B1 | AP2B1\_HUMAN | AP-2 complex subunit beta | 9C(125.0477) | NaN | X | NaN | NaN | NaN | NaN | 3.204988e+05 | 3.204988e+05 | NaN | NaN | VDKITEYLCEPLRK |
| CAND1 | CAND1\_HUMAN | Cullin-associated NEDD8-dissociated protein 1 | 179 | 165 | CAND1 | CAND1\_HUMAN | Cullin-associated NEDD8-dissociated protein 1 | 15C(125.0477) | NaN | X | NaN | NaN | NaN | NaN | 3.200919e+05 | 3.200919e+05 | NaN | NaN | QGGLLVNFHPSILTCLLPQLTSPR |
| LDHB | LDHB\_HUMAN | L-lactate dehydrogenase B chain | 164, 164, 164 | 159 | LDHB | LDHB\_HUMAN | L-lactate dehydrogenase B chain | 6C(125.0477) 6C(31.9988) 6C(57.0215) | X | X | NaN | X | NaN | NaN | 3.199103e+05 | 3.199103e+05 | NaN | NaN | VIGSGCNLDSAR |
| PRPS1 | PRPS1\_HUMAN | Ribose-phosphate pyrophosphokinase 1 | 41 | 34 | PRPS1 | PRPS1\_HUMAN | Ribose-phosphate pyrophosphokinase 1 | 8C(125.0477) | NaN | X | NaN | NaN | NaN | NaN | 3.192648e+05 | 3.192648e+05 | NaN | NaN | KFSNQETCVEIGESVR |
| FN3KRP | KT3K\_HUMAN | Ketosamine-3-kinase | 11, 11 | 7 | FN3KRP | KT3K\_HUMAN | Ketosamine-3-kinase | 5C(125.0477) 5C(15.9949) | NaN | X | X | NaN | NaN | NaN | 3.190112e+05 | 3.190112e+05 | NaN | NaN | RELGCSSVR |
| CMPK2 | CMPK2\_HUMAN | UMP-CMP kinase 2, mitochondrial | 226 | 221 | CMPK2 | CMPK2\_HUMAN | UMP-CMP kinase 2, mitochondrial | 6C(125.0477) | NaN | X | NaN | NaN | NaN | NaN | 3.185568e+05 | 3.185568e+05 | NaN | NaN | AVLEECTSFIPEAR |
| RAB1A | RAB1A\_HUMAN | Ras-related protein Rab-1A | 126 | 126 | RAB1A | RAB1A\_HUMAN | Ras-related protein Rab-1A | 1C(125.0477) | NaN | X | NaN | NaN | NaN | NaN | 3.174999e+05 | 3.174999e+05 | NaN | NaN | CDLTTKK |
| UBA3 | UBA3\_HUMAN | NEDD8-activating enzyme E1 catalytic subunit | 324 | 309 | UBA3 | UBA3\_HUMAN | NEDD8-activating enzyme E1 catalytic subunit | 16C(125.0477) | NaN | X | NaN | NaN | NaN | NaN | 3.171929e+05 | 3.171929e+05 | NaN | NaN | IIPAVASTNAVIAAVCATEVFK |
| EIF4B | IF4B\_HUMAN | Eukaryotic translation initiation factor 4B | 457 | 445 | EIF4B | IF4B\_HUMAN | Eukaryotic translation initiation factor 4B | 13C(125.0477) | NaN | X | NaN | NaN | NaN | NaN | 3.165176e+05 | 3.165176e+05 | NaN | NaN | SLENETLNKEEDCHSPTSKPPKPDQPLK |
| CCT8 | TCPQ\_HUMAN | T-complex protein 1 subunit theta | 136 | 121 | CCT8 | TCPQ\_HUMAN | T-complex protein 1 subunit theta | 16C(125.0477) | NaN | X | NaN | NaN | NaN | NaN | 3.164458e+05 | 3.164458e+05 | NaN | NaN | IGLSVSEVIEGYEIACR |
| TANGO2 | TNG2\_HUMAN | Transport and Golgi organization protein 2 homolog | 231 | 231 | TANGO2 | TNG2\_HUMAN | Transport and Golgi organization protein 2 homolog | 1C(125.0477) | NaN | X | NaN | NaN | NaN | NaN | 3.158650e+05 | 3.158650e+05 | NaN | NaN | CPGYGTR |
| CAPN5 | CAN5\_HUMAN | Calpain-5 | 544 | 544 | CAPN5 | CAN5\_HUMAN | Calpain-5 | 1C(125.0477) | NaN | X | NaN | NaN | NaN | NaN | 3.156976e+05 | 3.156976e+05 | NaN | NaN | CEGDKVR |
| VTA1 | VTA1\_HUMAN | Vacuolar protein sorting-associated protein VTA1 homolog | 155, 155 | 149 | VTA1 | VTA1\_HUMAN | Vacuolar protein sorting-associated protein VTA1 homolog | 7C(-34.0809) 7C(125.0477) | NaN | X | NaN | NaN | X | NaN | 3.155909e+05 | NaN | 3.155909e+05 | NaN | ATYIHNCLK |
| CNP | CN37\_HUMAN | 2',3'-cyclic-nucleotide 3'-phosphodiesterase | 252 | 245 | CNP | CN37\_HUMAN | 2',3'-cyclic-nucleotide 3'-phosphodiesterase | 8C(125.0477) | NaN | X | NaN | NaN | NaN | NaN | 3.155423e+05 | 3.155423e+05 | NaN | NaN | RPPGVLHCTTK |
| HBA1 | HBA\_HUMAN | Hemoglobin subunit alpha | 105, 105 | 101 | HBA1 | HBA\_HUMAN | Hemoglobin subunit alpha | 5C(125.0477) 5C(31.9988) | NaN | X | NaN | X | NaN | NaN | 3.154461e+05 | 3.154461e+05 | NaN | NaN | LLSHCLLVTLAAHLPAEFTPAVHASLDKFLASVSTVLTSKYR |
| ADD3 | ADDG\_HUMAN | Gamma-adducin | 286 | 281 | ADD3 | ADDG\_HUMAN | Gamma-adducin | 6C(125.0477) | NaN | X | NaN | NaN | NaN | NaN | 3.148960e+05 | NaN | 3.148960e+05 | NaN | VLGPSCK |
| CAND1 | CAND1\_HUMAN | Cullin-associated NEDD8-dissociated protein 1 | 92 | 79 | CAND1 | CAND1\_HUMAN | Cullin-associated NEDD8-dissociated protein 1 | 14C(125.0477) | NaN | X | NaN | NaN | NaN | NaN | 3.144146e+05 | 3.144146e+05 | NaN | NaN | VKEYQVETIVDTLCTNMLSDKEQLR |
| PRDX2 | PRDX2\_HUMAN | Peroxiredoxin-2 | 70, 70, 70 | 67 | PRDX2 | PRDX2\_HUMAN | Peroxiredoxin-2 | 4C(-34.0809) 4C(125.0477) 4C(31.9988) | NaN | X | NaN | X | X | NaN | 3.141029e+05 | 3.141029e+05 | NaN | NaN | KLGCEVLGVSVDSQFTHLAWINTPR |
| USP9X | USP9X\_HUMAN | Probable ubiquitin carboxyl-terminal hydrolase FAF-X | 1908 | 1900 | USP9X | USP9X\_HUMAN | Probable ubiquitin carboxyl-terminal hydrolase FAF-X | 9C(125.0477) | NaN | X | NaN | NaN | NaN | NaN | 3.139988e+05 | 3.012311e+05 | 1.276771e+04 | NaN | FDDGDVTECK |
| ACTBL2 | ACTBL\_HUMAN | Beta-actin-like protein 2 | 53, 53, 53 | 52 | ACTBL2 | ACTBL\_HUMAN | Beta-actin-like protein 2 | 2C(125.0477) 2C(15.9949) 2C(31.9988) | NaN | X | X | X | NaN | NaN | 3.137736e+05 | NaN | 3.137736e+05 | NaN | DCYVGDEAQSK |
| STIP1 | STIP1\_HUMAN | Stress-induced-phosphoprotein 1 | 461, 461 | 454 | STIP1 | STIP1\_HUMAN | Stress-induced-phosphoprotein 1 | 8C(125.0477) 8C(31.9988) | NaN | X | NaN | X | NaN | NaN | 3.136199e+05 | 3.136199e+05 | NaN | NaN | ALDLDSSCKEAADGYQR |
| SHMT1 | GLYC\_HUMAN | Serine hydroxymethyltransferase, cytosolic | 68 | 60 | SHMT1 | GLYC\_HUMAN | Serine hydroxymethyltransferase, cytosolic | 9C(125.0477) | NaN | X | NaN | NaN | NaN | NaN | 3.127118e+05 | 3.127118e+05 | NaN | NaN | AVLEALGSCLNNK |
| DHRS7B | DRS7B\_HUMAN | Dehydrogenase/reductase SDR family member 7B | 68 | 52 | DHRS7B | DRS7B\_HUMAN | Dehydrogenase/reductase SDR family member 7B | 17C(57.0215) | X | NaN | NaN | NaN | NaN | NaN | 3.124947e+05 | 3.124947e+05 | NaN | NaN | NAVVVITGATSGLGKECAK |
| USP9X | USP9X\_HUMAN | Probable ubiquitin carboxyl-terminal hydrolase FAF-X | 864 | 859 | USP9X | USP9X\_HUMAN | Probable ubiquitin carboxyl-terminal hydrolase FAF-X | 6C(125.0477) | NaN | X | NaN | NaN | NaN | NaN | 3.124802e+05 | 3.124802e+05 | NaN | NaN | EYINECDSDYHEER |
| TPP2 | TPP2\_HUMAN | Tripeptidyl-peptidase 2 | 322 | 322 | TPP2 | TPP2\_HUMAN | Tripeptidyl-peptidase 2 | 1C(125.0477) | NaN | X | NaN | NaN | NaN | NaN | 3.122271e+05 | 3.122271e+05 | NaN | NaN | CDLVNYSYGEATHWPNSGR |
| EPB41 | EPB41\_HUMAN | Protein 4.1 | 306 | 303 | EPB41 | EPB41\_HUMAN | Protein 4.1 | 4C(125.0477) | NaN | X | NaN | NaN | NaN | NaN | 3.102718e+05 | 3.102718e+05 | NaN | NaN | YYLCLQLR |
| UBE2O | UBE2O\_HUMAN | (E3-independent) E2 ubiquitin-conjugating enzyme | 585, 585 | 570 | UBE2O | UBE2O\_HUMAN | (E3-independent) E2 ubiquitin-conjugating enzyme | 16C(125.0477) 16C(57.0215) | X | X | NaN | NaN | NaN | NaN | 3.100262e+05 | 3.100262e+05 | NaN | NaN | SNDLFPVHHLDNNEFCPGDFVVDKR |
| PSMD2 | PSMD2\_HUMAN | 26S proteasome non-ATPase regulatory subunit 2 | 616 | 609 | PSMD2 | PSMD2\_HUMAN | 26S proteasome non-ATPase regulatory subunit 2 | 8C(125.0477) | NaN | X | NaN | NaN | NaN | NaN | 3.095884e+05 | 3.095884e+05 | NaN | NaN | VQQLLHICSEHFDSK |
| NAPRT | PNCB\_HUMAN | Nicotinate phosphoribosyltransferase | 533 | 525 | NAPRT | PNCB\_HUMAN | Nicotinate phosphoribosyltransferase | 9C(125.0477) | NaN | X | NaN | NaN | NaN | NaN | 3.092812e+05 | 3.092812e+05 | NaN | NaN | LQALVNSLCAGQSP |
| HNRNPAB | ROAA\_HUMAN | Heterogeneous nuclear ribonucleoprotein A/B | 98 | 92 | HNRNPAB | ROAA\_HUMAN | Heterogeneous nuclear ribonucleoprotein A/B | 7C(125.0477) | NaN | X | NaN | NaN | NaN | NaN | 3.085700e+05 | 2.949550e+05 | 1.361492e+04 | NaN | FGEVVDCTIK |
| TTC7A | TTC7A\_HUMAN | Tetratricopeptide repeat protein 7A | 20 | 13 | TTC7A | TTC7A\_HUMAN | Tetratricopeptide repeat protein 7A | 8C(31.9988) | NaN | NaN | NaN | X | NaN | NaN | 3.080269e+05 | NaN | 3.080269e+05 | NaN | VESELERCRAEGHWDR |
| CCT5 | TCPE\_HUMAN | T-complex protein 1 subunit epsilon | 407 | 401 | CCT5 | TCPE\_HUMAN | T-complex protein 1 subunit epsilon | 7C(125.0477) | NaN | X | NaN | NaN | NaN | NaN | 3.070372e+05 | 1.769140e+05 | 1.301233e+05 | NaN | SLHDALCVIR |
| VAT1 | VAT1\_HUMAN | Synaptic vesicle membrane protein VAT-1 homolog | 50 | 50 | VAT1 | VAT1\_HUMAN | Synaptic vesicle membrane protein VAT-1 homolog | 1C(125.0477) | NaN | X | NaN | NaN | NaN | NaN | 3.064304e+05 | 3.064304e+05 | NaN | NaN | CLVLTGFGGYDK |
| PRDX1 | PRDX1\_HUMAN | Peroxiredoxin-1 | 173 | 159 | PRDX1 | PRDX1\_HUMAN | Peroxiredoxin-1 | 15C(125.0477) | NaN | X | NaN | NaN | NaN | NaN | 3.061046e+05 | 3.061046e+05 | NaN | NaN | LVQAFQFTDKHGEVCPAGWKPGSDTIKPDVQK |
| PDXP | PLPP\_HUMAN | Chronophin | 171 | 168 | PDXP | PLPP\_HUMAN | Chronophin | 4C(125.0477) | NaN | X | NaN | NaN | NaN | NaN | 3.053920e+05 | 3.053920e+05 | NaN | NaN | DPECLLVATDRDPWHPLSDGSR |
| HPS1 | HPS1\_HUMAN | Hermansky-Pudlak syndrome 1 protein | 165 | 161 | HPS1 | HPS1\_HUMAN | Hermansky-Pudlak syndrome 1 protein | 5C(57.0215) | X | NaN | NaN | NaN | NaN | NaN | 3.050097e+05 | NaN | NaN | 305009.7000 | EQEQCFAVEALER |
| ALAD | HEM2\_HUMAN | Delta-aminolevulinic acid dehydratase | 223 | 222 | ALAD | HEM2\_HUMAN | Delta-aminolevulinic acid dehydratase | 2C(125.0477) | NaN | X | NaN | NaN | NaN | NaN | 3.049599e+05 | 3.049599e+05 | NaN | NaN | RCYQLPPGAR |
| DCAF11 | DCA11\_HUMAN | DDB1- and CUL4-associated factor 11 | 288 | 279 | DCAF11 | DCA11\_HUMAN | DDB1- and CUL4-associated factor 11 | 10C(125.0477) | NaN | X | NaN | NaN | NaN | NaN | 3.047762e+05 | 3.047762e+05 | NaN | NaN | EVLGGANDGCLYVFDR |
| FAH | FAAA\_HUMAN | Fumarylacetoacetase | 193 | 175 | FAH | FAAA\_HUMAN | Fumarylacetoacetase | 19C(125.0477) | NaN | X | NaN | NaN | NaN | NaN | 3.047616e+05 | 3.047616e+05 | NaN | NaN | RPMGQMKPDDSKPPVYGACK |
| CUL2 | CUL2\_HUMAN | Cullin-2 | 560 | 553 | CUL2 | CUL2\_HUMAN | Cullin-2 | 8C(125.0477) | NaN | X | NaN | NaN | NaN | NaN | 3.046898e+05 | 3.046898e+05 | NaN | NaN | LTWLHYLCTGEVK |
| TGM2 | TGM2\_HUMAN | Protein-glutamine gamma-glutamyltransferase 2 | 554 | 553 | TGM2 | TGM2\_HUMAN | Protein-glutamine gamma-glutamyltransferase 2 | 2C(125.0477) | NaN | X | NaN | NaN | NaN | NaN | 3.045953e+05 | 2.346217e+05 | 6.997355e+04 | NaN | DCLTESNLIK |
| ADH5 | ADHX\_HUMAN | Alcohol dehydrogenase class-3 | 282 | 277 | ADH5 | ADHX\_HUMAN | Alcohol dehydrogenase class-3 | 6C(125.0477) | NaN | X | NaN | NaN | NaN | NaN | 3.025958e+05 | 3.025958e+05 | NaN | NaN | AALEACHK |
| RANBP10 | RBP10\_HUMAN | Ran-binding protein 10 | 243 | 237 | RANBP10 | RBP10\_HUMAN | Ran-binding protein 10 | 7C(125.0477) | NaN | X | NaN | NaN | NaN | NaN | 3.017865e+05 | 3.017865e+05 | NaN | NaN | VQGTVHCFPISAR |
| ACLY | ACLY\_HUMAN | ATP-citrate synthase | 20, 20 | 18 | ACLY | ACLY\_HUMAN | ATP-citrate synthase | 3C(125.0477) 3C(57.0215) | X | X | NaN | NaN | NaN | NaN | 3.014004e+05 | 3.014004e+05 | NaN | NaN | FICTTSAIQNR |
| GLOD4 | GLOD4\_HUMAN | Glyoxalase domain-containing protein 4 | 221 | 217 | GLOD4 | GLOD4\_HUMAN | Glyoxalase domain-containing protein 4 | 5C(125.0477) | NaN | X | NaN | NaN | NaN | NaN | 3.008407e+05 | 2.650827e+05 | 3.575807e+04 | NaN | IAFSCPQK |
| CHORDC1 | CHRD1\_HUMAN | Cysteine and histidine-rich domain-containing protein 1 | 211 | 199 | CHORDC1 | CHRD1\_HUMAN | Cysteine and histidine-rich domain-containing protein 1 | 13C(125.0477) | NaN | X | NaN | NaN | NaN | NaN | 3.003812e+05 | 3.003812e+05 | NaN | NaN | TSDFNTFLAQEGCTK |
| ANK1 | ANK1\_HUMAN | Ankyrin-1 | 379 | 368 | ANK1 | ANK1\_HUMAN | Ankyrin-1 | 12C(125.0477) | NaN | X | NaN | NaN | NaN | NaN | 2.999141e+05 | 2.999141e+05 | NaN | NaN | ALNGFTPLHIACKK |
| AP2B1 | AP2B1\_HUMAN | AP-2 complex subunit beta | 241 | 236 | AP2B1 | AP2B1\_HUMAN | AP-2 complex subunit beta | 6C(125.0477) | NaN | X | NaN | NaN | NaN | NaN | 2.989719e+05 | 2.989719e+05 | NaN | NaN | EAQSICER |
| CLIC1 | CLIC1\_HUMAN | Chloride intracellular channel protein 1 | 24 | 21 | CLIC1 | CLIC1\_HUMAN | Chloride intracellular channel protein 1 | 4C(125.0477) | NaN | X | NaN | NaN | NaN | NaN | 2.988319e+05 | 2.988319e+05 | NaN | NaN | IGNCPFSQR |
| SPTB | SPTB1\_HUMAN | Spectrin beta chain, erythrocytic | 1552, 1552 | 1541 | SPTB | SPTB1\_HUMAN | Spectrin beta chain, erythrocytic | 12C(125.0477) 12C(31.9988) | NaN | X | NaN | X | NaN | NaN | 2.986944e+05 | NaN | 2.986944e+05 | NaN | GQQLVEAAEIDCQDLEER |
| GSTM3 | GSTM3\_HUMAN | Glutathione S-transferase Mu 3 | 39 | 37 | GSTM3 | GSTM3\_HUMAN | Glutathione S-transferase Mu 3 | 3C(125.0477) | NaN | X | NaN | NaN | NaN | NaN | 2.984540e+05 | 2.984540e+05 | NaN | NaN | YTCGEAPDYDR |
| PSMA5 | PSA5\_HUMAN | Proteasome subunit alpha type-5 | 47, 47 | 33 | PSMA5 | PSA5\_HUMAN | Proteasome subunit alpha type-5 | 15C(125.0477) 15C(57.0215) | X | X | NaN | NaN | NaN | NaN | 2.978026e+05 | 2.978026e+05 | NaN | NaN | LGSTAIGIQTSEGVCLAVEK |
| NUDT2 | AP4A\_HUMAN | Bis(5'-nucleosyl)-tetraphosphatase [asymmetrical] | 6 | 5 | NUDT2 | AP4A\_HUMAN | Bis(5'-nucleosyl)-tetraphosphatase [asymmetrical] | 2C(125.0477) | NaN | X | NaN | NaN | NaN | NaN | 2.976317e+05 | 2.976317e+05 | NaN | NaN | ACGLIIFR |
| WDR81 | WDR81\_HUMAN | WD repeat-containing protein 81 | 130 | 115 | WDR81 | WDR81\_HUMAN | WD repeat-containing protein 81 | 16C(125.0477) | NaN | X | NaN | NaN | NaN | NaN | 2.964786e+05 | 2.964786e+05 | NaN | NaN | LSYPLGGGLPFEDGSCGPETLTR |
| PSMD4 | PSMD4\_HUMAN | 26S proteasome non-ATPase regulatory subunit 4 | 87 | 84 | PSMD4 | PSMD4\_HUMAN | 26S proteasome non-ATPase regulatory subunit 4 | 4C(15.9949) | NaN | NaN | X | NaN | NaN | NaN | 2.952397e+05 | NaN | 2.952397e+05 | NaN | ITFCTGIR |
| RANBP1 | RANG\_HUMAN | Ran-specific GTPase-activating protein | 99, 99, 99 | 98 | RANBP1 | RANG\_HUMAN | Ran-specific GTPase-activating protein | 2C(125.0477) 2C(31.9988) 2C(57.0215) | X | X | NaN | X | NaN | NaN | 2.948939e+05 | 2.948939e+05 | NaN | NaN | ICANHYITPMMELKPNAGSDR |
| PPP2CB | PP2AB\_HUMAN | Serine/threonine-protein phosphatase 2A catalytic subunit beta isoform | 133 | 122 | PPP2CB | PP2AB\_HUMAN | Serine/threonine-protein phosphatase 2A catalytic subunit beta isoform | 12C(125.0477) | NaN | X | NaN | NaN | NaN | NaN | 2.948432e+05 | 2.948432e+05 | NaN | NaN | QITQVYGFYDECLR |
| HSPA1A | HS71A\_HUMAN | Heat shock 70 kDa protein 1A | 306 | 302 | HSPA1A | HS71A\_HUMAN | Heat shock 70 kDa protein 1A | 5C(125.0477) | NaN | X | NaN | NaN | NaN | NaN | 2.946600e+05 | 2.486055e+05 | 4.605448e+04 | NaN | FEELCSDLFR |
| SPTA1 | SPTA1\_HUMAN | Spectrin alpha chain, erythrocytic 1 | 1879, 1879 | 1875 | SPTA1 | SPTA1\_HUMAN | Spectrin alpha chain, erythrocytic 1 | 5C(125.0477) 5C(31.9988) | NaN | X | NaN | X | NaN | NaN | 2.945901e+05 | 2.945901e+05 | NaN | NaN | VQNVCAQGEDILNK |
| ADRM1 | ADRM1\_HUMAN | Proteasomal ubiquitin receptor ADRM1 | 88 | 85 | ADRM1 | ADRM1\_HUMAN | Proteasomal ubiquitin receptor ADRM1 | 4C(125.0477) | NaN | X | NaN | NaN | NaN | NaN | 2.944984e+05 | 2.944984e+05 | NaN | NaN | VPQCPSGR |
| PRPS1 | PRPS1\_HUMAN | Ribose-phosphate pyrophosphokinase 1 | 165 | 164 | PRPS1 | PRPS1\_HUMAN | Ribose-phosphate pyrophosphokinase 1 | 2C(125.0477) | NaN | X | NaN | NaN | NaN | NaN | 2.937812e+05 | NaN | 2.937812e+05 | NaN | NCTIVSPDAGGAK |
| DPYSL2 | DPYL2\_HUMAN | Dihydropyrimidinase-related protein 2 | 504 | 497 | DPYSL2 | DPYL2\_HUMAN | Dihydropyrimidinase-related protein 2 | 8C(125.0477) | NaN | X | NaN | NaN | NaN | NaN | 2.929775e+05 | 2.929775e+05 | NaN | NaN | GLYDGPVCEVSVTPK |
| TPI1 | TPIS\_HUMAN | Triosephosphate isomerase | 67, 67 | 60 | TPI1 | TPIS\_HUMAN | Triosephosphate isomerase | 8C(125.0477) 8C(57.0215) | X | X | NaN | NaN | NaN | NaN | 2.928145e+05 | 2.928145e+05 | NaN | NaN | IAVAAQNCYK |
| KRT31 | K1H1\_HUMAN | Keratin, type I cuticular Ha1 | 351 | 349 | KRT31 | K1H1\_HUMAN | Keratin, type I cuticular Ha1 | 3C(125.0477) | NaN | X | NaN | NaN | NaN | NaN | 2.925057e+05 | NaN | 2.925057e+05 | NaN | LECEINTYR |
| SUGT1 | SGT1\_HUMAN | Protein SGT1 homolog | 49 | 37 | SUGT1 | SGT1\_HUMAN | Protein SGT1 homolog | 13C(125.0477) | NaN | X | NaN | NaN | NaN | NaN | 2.924613e+05 | 2.924613e+05 | NaN | NaN | ALEQKPDDAQYYCQR |
| SPTA1 | SPTA1\_HUMAN | Spectrin alpha chain, erythrocytic 1 | 965, 965, 965, 965 | 960 | SPTA1 | SPTA1\_HUMAN | Spectrin alpha chain, erythrocytic 1 | 6C(125.0477) 6C(15.9949) 6C(31.9988) 6C(57.0215) | X | X | X | X | NaN | NaN | 2.923728e+05 | 2.923728e+05 | NaN | NaN | NQANACQQQQAAPVEGVAGEQR |
| BPGM | PMGE\_HUMAN | Bisphosphoglycerate mutase | 145 | 142 | BPGM | PMGE\_HUMAN | Bisphosphoglycerate mutase | 4C(125.0477) | NaN | X | NaN | NaN | NaN | NaN | 2.919974e+05 | 2.919974e+05 | NaN | NaN | YKVCDVPLDQLPR |
| EPB42 | EPB42\_HUMAN | Protein 4.2 | 428, 428 | 428 | EPB42 | EPB42\_HUMAN | Protein 4.2 | 1C(125.0477) 1C(31.9988) | NaN | X | NaN | X | NaN | NaN | 2.914512e+05 | NaN | NaN | 291451.2000 | CEDITQNYK |
| P4HB | PDIA1\_HUMAN | Protein disulfide-isomerase | 312 | 309 | P4HB | PDIA1\_HUMAN | Protein disulfide-isomerase | 4C(125.0477) | NaN | X | NaN | NaN | NaN | NaN | 2.891620e+05 | 2.196738e+05 | 6.948826e+04 | NaN | KEECPAVR |
| WDFY1 | WDFY1\_HUMAN | WD repeat and FYVE domain-containing protein 1 | 287 | 276 | WDFY1 | WDFY1\_HUMAN | WD repeat and FYVE domain-containing protein 1 | 12C(125.0477) | NaN | X | NaN | NaN | NaN | NaN | 2.882862e+05 | 2.882862e+05 | NaN | NaN | EEAPQWLESDSCQK |
| FCSK | FCSK\_HUMAN | L-fucose kinase | 970 | 966 | FCSK | FCSK\_HUMAN | L-fucose kinase | 5C(125.0477) | NaN | X | NaN | NaN | NaN | NaN | 2.876918e+05 | 2.876918e+05 | NaN | NaN | QTEECAEGFR |
| TNPO2 | TNPO2\_HUMAN | Transportin-2 | 93 | 91 | TNPO2 | TNPO2\_HUMAN | Transportin-2 | 3C(125.0477) | NaN | X | NaN | NaN | NaN | NaN | 2.870830e+05 | 2.870830e+05 | NaN | NaN | QECLNNIGDASSLIR |
| STIP1 | STIP1\_HUMAN | Stress-induced-phosphoprotein 1 | 26 | 14 | STIP1 | STIP1\_HUMAN | Stress-induced-phosphoprotein 1 | 13C(125.0477) | NaN | X | NaN | NaN | NaN | NaN | 2.868122e+05 | 2.868122e+05 | NaN | NaN | ALSVGNIDDALQCYSEAIKLDPHNHVLYSNR |
| RPS17 | RS17\_HUMAN | 40S ribosomal protein S17 | 35 | 34 | RPS17 | RS17\_HUMAN | 40S ribosomal protein S17 | 2C(125.0477) | NaN | X | NaN | NaN | NaN | NaN | 2.866921e+05 | 2.866921e+05 | NaN | NaN | VCEEIAIIPSKK |
| DNPEP | DNPEP\_HUMAN | Aspartyl aminopeptidase | 48 | 38 | DNPEP | DNPEP\_HUMAN | Aspartyl aminopeptidase | 11C(125.0477) | NaN | X | NaN | NaN | NaN | NaN | 2.865370e+05 | 2.865370e+05 | NaN | NaN | SPSPFHAVAECR |
| RNPEP | AMPB\_HUMAN | Aminopeptidase B | 369 | 358 | RNPEP | AMPB\_HUMAN | Aminopeptidase B | 12C(125.0477) | NaN | X | NaN | NaN | NaN | NaN | 2.860170e+05 | 2.860170e+05 | NaN | NaN | ISTILFGAAYTCLEAATGR |
| TBC1D24 | TBC24\_HUMAN | TBC1 domain family member 24 | 430 | 422 | TBC1D24 | TBC24\_HUMAN | TBC1 domain family member 24 | 9C(125.0477) | NaN | X | NaN | NaN | NaN | NaN | 2.857180e+05 | 2.857180e+05 | NaN | NaN | LGFFGTGECFVFR |
| PPP6R3 | PP6R3\_HUMAN | Serine/threonine-protein phosphatase 6 regulatory subunit 3 | 467 | 464 | PPP6R3 | PP6R3\_HUMAN | Serine/threonine-protein phosphatase 6 regulatory subunit 3 | 4C(125.0477) | NaN | X | NaN | NaN | NaN | NaN | 2.857129e+05 | 2.857129e+05 | NaN | NaN | IANCIVHSTDK |
| ARHGDIB | GDIR2\_HUMAN | Rho GDP-dissociation inhibitor 2 | 76 | 72 | ARHGDIB | GDIR2\_HUMAN | Rho GDP-dissociation inhibitor 2 | 5C(125.0477) | NaN | X | NaN | NaN | NaN | NaN | 2.853691e+05 | 2.853691e+05 | NaN | NaN | LTLVCESAPGPITMDLTGDLEALKK |
| ANK1 | ANK1\_HUMAN | Ankyrin-1 | 864 | 862 | ANK1 | ANK1\_HUMAN | Ankyrin-1 | 3C(15.9949) | NaN | NaN | X | NaN | NaN | NaN | 2.852746e+05 | NaN | NaN | 285274.5600 | IPCAMPETVVIR |
| ATIC | PUR9\_HUMAN | Bifunctional purine biosynthesis protein ATIC | 363 | 359 | ATIC | PUR9\_HUMAN | Bifunctional purine biosynthesis protein ATIC | 5C(125.0477) | NaN | X | NaN | NaN | NaN | NaN | 2.852665e+05 | 2.852665e+05 | NaN | NaN | NGNYCVLQMDQSYKPDENEVR |
| ANKRD54 | ANR54\_HUMAN | Ankyrin repeat domain-containing protein 54 | 185 | 173 | ANKRD54 | ANR54\_HUMAN | Ankyrin repeat domain-containing protein 54 | 13C(125.0477) | NaN | X | NaN | NaN | NaN | NaN | 2.850742e+05 | 2.850742e+05 | NaN | NaN | DGLGNTPLHLAACTNHVPVITTLLR |
| XK | XK\_HUMAN | Membrane transport protein XK | 442 | 428 | XK | XK\_HUMAN | Membrane transport protein XK | 15C(125.0477) | NaN | X | NaN | NaN | NaN | NaN | 2.850352e+05 | 2.298924e+05 | 5.514275e+04 | NaN | TSPEPGQFLNAEDLCSA |
| USP9X | USP9X\_HUMAN | Probable ubiquitin carboxyl-terminal hydrolase FAF-X | 2107 | 2100 | USP9X | USP9X\_HUMAN | Probable ubiquitin carboxyl-terminal hydrolase FAF-X | 8C(125.0477) | NaN | X | NaN | NaN | NaN | NaN | 2.847753e+05 | 2.847753e+05 | NaN | NaN | FSEYLLECPSAEVR |
| IMPACT | IMPCT\_HUMAN | Protein IMPACT | 195 | 184 | IMPACT | IMPCT\_HUMAN | Protein IMPACT | 12C(125.0477) | NaN | X | NaN | NaN | NaN | NaN | 2.847170e+05 | 2.847170e+05 | NaN | NaN | STFQAHLAPVVCPK |
| PCMT1 | PIMT\_HUMAN | Protein-L-isoaspartate(D-aspartate) O-methyltransferase | 95, 95 | 82 | PCMT1 | PIMT\_HUMAN | Protein-L-isoaspartate(D-aspartate) O-methyltransferase | 14C(125.0477) 14C(31.9988) | NaN | X | NaN | X | NaN | NaN | 2.843084e+05 | 2.843084e+05 | NaN | NaN | ALDVGSGSGILTACFAR |
| GDI1 | GDIA\_HUMAN | Rab GDP dissociation inhibitor alpha | 335, 335 | 330 | GDI1 | GDIA\_HUMAN | Rab GDP dissociation inhibitor alpha | 6C(125.0477) 6C(31.9988) | NaN | X | NaN | X | NaN | NaN | 2.837826e+05 | 2.837826e+05 | NaN | NaN | SDIYVCMISYAHNVAAQGK |
| PSMD10 | PSD10\_HUMAN | 26S proteasome non-ATPase regulatory subunit 10 | 107 | 96 | PSMD10 | PSD10\_HUMAN | 26S proteasome non-ATPase regulatory subunit 10 | 12C(125.0477) | NaN | X | NaN | NaN | NaN | NaN | 2.832028e+05 | 2.832028e+05 | NaN | NaN | GAQVNAVNQNGCTPLHYAASK |
| GSTM2 | GSTM2\_HUMAN | Glutathione S-transferase Mu 2 | 174 | 167 | GSTM2 | GSTM2\_HUMAN | Glutathione S-transferase Mu 2 | 8C(125.0477) | NaN | X | NaN | NaN | NaN | NaN | 2.826828e+05 | 2.826828e+05 | NaN | NaN | NQVFEPSCLDAFPNLK |
| PSME2 | PSME2\_HUMAN | Proteasome activator complex subunit 2 | 91 | 91 | PSME2 | PSME2\_HUMAN | Proteasome activator complex subunit 2 | 1C(125.0477) | NaN | X | NaN | NaN | NaN | NaN | 2.822808e+05 | NaN | 2.822808e+05 | NaN | CGFLPGNEK |
| ECPAS | ECM29\_HUMAN | Proteasome adapter and scaffold protein ECM29 | 1070 | 1068 | ECPAS | ECM29\_HUMAN | Proteasome adapter and scaffold protein ECM29 | 3C(125.0477) | NaN | X | NaN | NaN | NaN | NaN | 2.819898e+05 | 2.819898e+05 | NaN | NaN | ELCSLASDLSQPDLVYK |
| C9orf64 | QSPP\_HUMAN | Queuosine salvage protein | 170 | 163 | C9orf64 | QSPP\_HUMAN | Queuosine salvage protein | 8C(125.0477) | NaN | X | NaN | NaN | NaN | NaN | 2.819482e+05 | 2.819482e+05 | NaN | NaN | FGGSFLNCVR |
| XK | XK\_HUMAN | Membrane transport protein XK | 187 | 187 | XK | XK\_HUMAN | Membrane transport protein XK | 1C(125.0477) | NaN | X | NaN | NaN | NaN | NaN | 2.817254e+05 | NaN | 2.425945e+05 | 39130.9020 | CNILAIK |
| SKP1 | SKP1\_HUMAN | S-phase kinase-associated protein 1 | 62 | 58 | SKP1 | SKP1\_HUMAN | S-phase kinase-associated protein 1 | 5C(125.0477) | NaN | X | NaN | NaN | NaN | NaN | 2.814762e+05 | 2.814762e+05 | NaN | NaN | VIQWCTHHK |
| CD55 | DAF\_HUMAN | Complement decay-accelerating factor | 145 | 143 | CD55 | DAF\_HUMAN | Complement decay-accelerating factor | 3C(125.0477) | NaN | X | NaN | NaN | NaN | NaN | 2.804932e+05 | 1.361976e+05 | 1.442956e+05 | NaN | LTCLQNLK |
| SPRR3 | SPRR3\_HUMAN | Small proline-rich protein 3 | 66 | 61 | SPRR3 | SPRR3\_HUMAN | Small proline-rich protein 3 | 6C(125.0477) | NaN | X | NaN | NaN | NaN | NaN | 2.786980e+05 | NaN | 1.840003e+05 | 94697.7300 | VPEPGCTK |
| PDCD6IP | PDC6I\_HUMAN | Programmed cell death 6-interacting protein | 512 | 511 | PDCD6IP | PDC6I\_HUMAN | Programmed cell death 6-interacting protein | 2C(125.0477) | NaN | X | NaN | NaN | NaN | NaN | 2.784384e+05 | 2.646812e+05 | 1.375715e+04 | NaN | ECYQSHR |
| BLVRB | BLVRB\_HUMAN | Flavin reductase (NADPH) | 109 | 100 | BLVRB | BLVRB\_HUMAN | Flavin reductase (NADPH) | 10C(125.0477) | NaN | X | NaN | NaN | NaN | NaN | 2.769793e+05 | 2.769793e+05 | NaN | NaN | AHGVDKVVACTSAFLLWDPTK |
| HMBS | HEM3\_HUMAN | Porphobilinogen deaminase | 261 | 256 | HMBS | HEM3\_HUMAN | Porphobilinogen deaminase | 6C(125.0477) | NaN | X | NaN | NaN | NaN | NaN | 2.769758e+05 | 2.769758e+05 | NaN | NaN | HLEGGCSVPVAVHTAMK |
| PSMD1 | PSMD1\_HUMAN | 26S proteasome non-ATPase regulatory subunit 1 | 571 | 560 | PSMD1 | PSMD1\_HUMAN | 26S proteasome non-ATPase regulatory subunit 1 | 12C(125.0477) | NaN | X | NaN | NaN | NaN | NaN | 2.766781e+05 | 2.766781e+05 | NaN | NaN | MEEADALIESLCR |
| TNPO3 | TNPO3\_HUMAN | Transportin-3 | 908 | 901 | TNPO3 | TNPO3\_HUMAN | Transportin-3 | 8C(125.0477) | NaN | X | NaN | NaN | NaN | NaN | 2.754305e+05 | 2.754305e+05 | NaN | NaN | QVTSAEECK |
| OVCA2 | OVCA2\_HUMAN | Esterase OVCA2 | 41, 41 | 37 | OVCA2 | OVCA2\_HUMAN | Esterase OVCA2 | 5C(31.9988) 5C(57.0215) | X | NaN | NaN | X | NaN | NaN | 2.754048e+05 | 2.754048e+05 | NaN | NaN | AELVCLSGPHPVPDPPGPEGAR |
| CAT | CATA\_HUMAN | Catalase | 232, 232 | 222 | CAT | CATA\_HUMAN | Catalase | 11C(125.0477) 11C(15.9949) | NaN | X | X | NaN | NaN | NaN | 2.753196e+05 | 2.753196e+05 | NaN | NaN | LVNANGEAVYCKFHYK |
| TLN1 | TLN1\_HUMAN | Talin-1 | 709, 719 | 693 | TLN1 | TLN1\_HUMAN | Talin-1 | 17C(31.9988) 27C(57.0215) | X | NaN | NaN | X | NaN | NaN | 2.737605e+05 | 2.737605e+05 | NaN | NaN | TEDSGLQTQVIAAATQCALSTSQLVACTK |
| TTC38 | TTC38\_HUMAN | Tetratricopeptide repeat protein 38 | 429 | 416 | TTC38 | TTC38\_HUMAN | Tetratricopeptide repeat protein 38 | 14C(125.0477) | NaN | X | NaN | NaN | NaN | NaN | 2.736105e+05 | 2.736105e+05 | NaN | NaN | DVFNQLLIHAALNCTSSVHK |
| PFAS | PUR4\_HUMAN | Phosphoribosylformylglycinamidine synthase | 871 | 863 | PFAS | PUR4\_HUMAN | Phosphoribosylformylglycinamidine synthase | 9C(125.0477) | NaN | X | NaN | NaN | NaN | NaN | 2.734048e+05 | 2.734048e+05 | NaN | NaN | LGGTALAQCFSQLGEHPPDLDLPENLVR |
| CAPZA2 | CAZA2\_HUMAN | F-actin-capping protein subunit alpha-2 | 157 | 148 | CAPZA2 | CAZA2\_HUMAN | F-actin-capping protein subunit alpha-2 | 10C(125.0477) | NaN | X | NaN | NaN | NaN | NaN | 2.728005e+05 | 2.728005e+05 | NaN | NaN | IDGQQTIIACIESHQFQAK |
| CD55 | DAF\_HUMAN | Complement decay-accelerating factor | 81 | 77 | CD55 | DAF\_HUMAN | Complement decay-accelerating factor | 5C(125.0477) | NaN | X | NaN | NaN | NaN | NaN | 2.726054e+05 | NaN | 2.661254e+05 | 6479.9780 | DSVICLK |
| CAND1 | CAND1\_HUMAN | Cullin-associated NEDD8-dissociated protein 1 | 131, 131 | 112 | CAND1 | CAND1\_HUMAN | Cullin-associated NEDD8-dissociated protein 1 | 20C(125.0477) 20C(15.9949) | NaN | X | X | NaN | NaN | NaN | 2.717202e+05 | 2.717202e+05 | NaN | NaN | TVIGELPPASSGSALAANVCKK |
| MYH9 | MYH9\_HUMAN | Myosin-9 | 931 | 924 | MYH9 | MYH9\_HUMAN | Myosin-9 | 8C(125.0477) | NaN | X | NaN | NaN | NaN | NaN | 2.709890e+05 | 2.709890e+05 | NaN | NaN | VEEEEERCQHLQAEK |
| PPP2CA | PP2AA\_HUMAN | Serine/threonine-protein phosphatase 2A catalytic subunit alpha isoform | 251 | 240 | PPP2CA | PP2AA\_HUMAN | Serine/threonine-protein phosphatase 2A catalytic subunit alpha isoform | 12C(125.0477) | NaN | X | NaN | NaN | NaN | NaN | 2.703295e+05 | 2.703295e+05 | NaN | NaN | AHQLVMEGYNWCHDR |
| COPS4 | CSN4\_HUMAN | COP9 signalosome complex subunit 4 | 232, 232 | 228 | COPS4 | CSN4\_HUMAN | COP9 signalosome complex subunit 4 | 5C(125.0477) 5C(31.9988) | NaN | X | NaN | X | NaN | NaN | 2.701863e+05 | 2.701863e+05 | NaN | NaN | HALHCTILASAGQQR |
| ALDH1A1 | AL1A1\_HUMAN | Aldehyde dehydrogenase 1A1 | 370 | 368 | ALDH1A1 | AL1A1\_HUMAN | Aldehyde dehydrogenase 1A1 | 3C(125.0477) | NaN | X | NaN | NaN | NaN | NaN | 2.698912e+05 | 2.056181e+05 | 6.427303e+04 | NaN | LECGGGPWGNK |
| SNAP23 | SNP23\_HUMAN | Synaptosomal-associated protein 23 | 112 | 101 | SNAP23 | SNP23\_HUMAN | Synaptosomal-associated protein 23 | 12C(125.0477) | NaN | X | NaN | NaN | NaN | NaN | 2.691475e+05 | NaN | 2.691475e+05 | NaN | TTWGDGGENSPCNVVSK |
| VPS13A | VP13A\_HUMAN | Vacuolar protein sorting-associated protein 13A | 1626 | 1619 | VPS13A | VP13A\_HUMAN | Vacuolar protein sorting-associated protein 13A | 8C(125.0477) | NaN | X | NaN | NaN | NaN | NaN | 2.688221e+05 | 2.688221e+05 | NaN | NaN | ITTVLQPCDLFYQTTQK |
| CAPN1 | CAN1\_HUMAN | Calpain-1 catalytic subunit | 351 | 348 | CAPN1 | CAN1\_HUMAN | Calpain-1 catalytic subunit | 4C(125.0477) | NaN | X | NaN | NaN | NaN | NaN | 2.687365e+05 | 2.053600e+05 | 6.337652e+04 | NaN | LEICNLTPDALK |
| PGM2L1 | PGM2L\_HUMAN | Glucose 1,6-bisphosphate synthase | 303 | 303 | PGM2L1 | PGM2L\_HUMAN | Glucose 1,6-bisphosphate synthase | 1C(125.0477) | NaN | X | NaN | NaN | NaN | NaN | 2.675692e+05 | 2.675692e+05 | NaN | NaN | CPNPEEGESVLELSLR |
| PGM2 | PGM2\_HUMAN | Phosphoglucomutase-2 | 573 | 568 | PGM2 | PGM2\_HUMAN | Phosphoglucomutase-2 | 6C(125.0477) | NaN | X | NaN | NaN | NaN | NaN | 2.667014e+05 | 2.667014e+05 | NaN | NaN | YYAELCAPPGNSDPEQLKK |
| RPS6KB1 | KS6B1\_HUMAN | Ribosomal protein S6 kinase beta-1 | 90 | 86 | RPS6KB1 | KS6B1\_HUMAN | Ribosomal protein S6 kinase beta-1 | 5C(125.0477) | NaN | X | NaN | NaN | NaN | NaN | 2.664384e+05 | 2.664384e+05 | NaN | NaN | IRPECFELLR |
| CAPZA1 | CAZA1\_HUMAN | F-actin-capping protein subunit alpha-1 | 124 | 122 | CAPZA1 | CAZA1\_HUMAN | F-actin-capping protein subunit alpha-1 | 3C(125.0477) | NaN | X | NaN | NaN | NaN | NaN | 2.659525e+05 | 1.209613e+05 | 1.449912e+05 | NaN | ESCDSALR |
| PLAA | PLAP\_HUMAN | Phospholipase A-2-activating protein | 208 | 196 | PLAA | PLAP\_HUMAN | Phospholipase A-2-activating protein | 13C(125.0477) | NaN | X | NaN | NaN | NaN | NaN | 2.656970e+05 | 2.656970e+05 | NaN | NaN | GLAILSETEFLSCANDASIR |
| AP2B1 | AP2B1\_HUMAN | AP-2 complex subunit beta | 144 | 140 | AP2B1 | AP2B1\_HUMAN | AP-2 complex subunit beta | 5C(125.0477) | NaN | X | NaN | NaN | NaN | NaN | 2.656784e+05 | NaN | 2.575047e+05 | 8173.6840 | TAAVCVAK |
| BRCC3 | BRCC3\_HUMAN | Lys-63-specific deubiquitinase BRCC36 | 172 | 168 | BRCC3 | BRCC3\_HUMAN | Lys-63-specific deubiquitinase BRCC36 | 5C(125.0477) | NaN | X | NaN | NaN | NaN | NaN | 2.653882e+05 | 2.653882e+05 | NaN | NaN | VLYTCFQSIQAQK |
| SPTA1 | SPTA1\_HUMAN | Spectrin alpha chain, erythrocytic 1 | 2072 | 2062 | SPTA1 | SPTA1\_HUMAN | Spectrin alpha chain, erythrocytic 1 | 11C(125.0477) | NaN | X | NaN | NaN | NaN | NaN | 2.643100e+05 | 2.643100e+05 | NaN | NaN | MEENLSEPVHCVSLNEIR |
| SPTA1 | SPTA1\_HUMAN | Spectrin alpha chain, erythrocytic 1 | 965, 965, 965 | 960 | SPTA1 | SPTA1\_HUMAN | Spectrin alpha chain, erythrocytic 1 | 6C(125.0477) 6C(31.9988) 6C(57.0215) | X | X | NaN | X | NaN | NaN | 2.642781e+05 | NaN | NaN | 264278.0600 | NQANACQQQQAAPVEGVAGEQR |
| ACTN4 | ACTN4\_HUMAN | Alpha-actinin-4 | 173 | 167 | ACTN4 | ACTN4\_HUMAN | Alpha-actinin-4 | 7C(125.0477) | NaN | X | NaN | NaN | NaN | NaN | 2.641105e+05 | 2.641105e+05 | NaN | NaN | EGLLLWCQR |
| CARM1 | CARM1\_HUMAN | Histone-arginine methyltransferase CARM1 | 438 | 430 | CARM1 | CARM1\_HUMAN | Histone-arginine methyltransferase CARM1 | 9C(125.0477) | NaN | X | NaN | NaN | NaN | NaN | 2.641065e+05 | 2.290385e+05 | 3.506802e+04 | NaN | AGDTLSGTCLLIANK |
| ALB | ALBU\_HUMAN | Albumin | 224 | 222 | ALB | ALBU\_HUMAN | Albumin | 3C(57.0215) | X | NaN | NaN | NaN | NaN | NaN | 2.639845e+05 | 2.340012e+05 | 2.998329e+04 | NaN | LKCASLQKFGER |
| TPI1 | TPIS\_HUMAN | Triosephosphate isomerase | 67 | 60 | TPI1 | TPIS\_HUMAN | Triosephosphate isomerase | 8C(125.0477) | NaN | X | NaN | NaN | NaN | NaN | 2.637692e+05 | NaN | 2.637692e+05 | NaN | IAVAAQNCYK |
| STK11 | STK11\_HUMAN | Serine/threonine-protein kinase STK11 | 210 | 192 | STK11 | STK11\_HUMAN | Serine/threonine-protein kinase STK11 | 19C(125.0477) | NaN | X | NaN | NaN | NaN | NaN | 2.634369e+05 | 2.634369e+05 | NaN | NaN | ISDLGVAEALHPFAADDTCR |
| GSS | GSHB\_HUMAN | Glutathione synthetase | 409 | 401 | GSS | GSHB\_HUMAN | Glutathione synthetase | 9C(125.0477) | NaN | X | NaN | NaN | NaN | NaN | 2.626740e+05 | 2.626740e+05 | NaN | NaN | IEPEPFENCLLRPGSPAR |
| CUL5 | CUL5\_HUMAN | Cullin-5 | 281 | 275 | CUL5 | CUL5\_HUMAN | Cullin-5 | 7C(125.0477) | NaN | X | NaN | NaN | NaN | NaN | 2.617556e+05 | 2.617556e+05 | NaN | NaN | ETILAECQGMIK |
| HSPBP1 | HPBP1\_HUMAN | Hsp70-binding protein 1 | 310 | 309 | HSPBP1 | HPBP1\_HUMAN | Hsp70-binding protein 1 | 2C(125.0477) | NaN | X | NaN | NaN | NaN | NaN | 2.615398e+05 | 2.615398e+05 | NaN | NaN | ECREPELGLEELLR |
| GOLPH3 | GOLP3\_HUMAN | Golgi phosphoprotein 3 | 280 | 270 | GOLPH3 | GOLP3\_HUMAN | Golgi phosphoprotein 3 | 11C(125.0477) | NaN | X | NaN | NaN | NaN | NaN | 2.607089e+05 | 2.607089e+05 | NaN | NaN | QLLDLDPEVECLK |
| ATG7 | ATG7\_HUMAN | Ubiquitin-like modifier-activating enzyme ATG7 | 298 | 289 | ATG7 | ATG7\_HUMAN | Ubiquitin-like modifier-activating enzyme ATG7 | 10C(125.0477) | NaN | X | NaN | NaN | NaN | NaN | 2.601388e+05 | 2.601388e+05 | NaN | NaN | LPEMAFSPDCPK |
| ANP32A | AN32A\_HUMAN | Acidic leucine-rich nuclear phosphoprotein 32 family member A | 123 | 117 | ANP32A | AN32A\_HUMAN | Acidic leucine-rich nuclear phosphoprotein 32 family member A | 7C(125.0477) | NaN | X | NaN | NaN | NaN | NaN | 2.601242e+05 | 2.601242e+05 | NaN | NaN | SLDLFNCEVTNLNDYR |
| ANKRD28 | ANR28\_HUMAN | Serine/threonine-protein phosphatase 6 regulatory ankyrin repeat subunit A | 898 | 891 | ANKRD28 | ANR28\_HUMAN | Serine/threonine-protein phosphatase 6 regulatory ankyrin repeat subunit A | 8C(125.0477) | NaN | X | NaN | NaN | NaN | NaN | 2.597833e+05 | 2.597833e+05 | NaN | NaN | NTALHLACSK |
| HPRT1 | HPRT\_HUMAN | Hypoxanthine-guanine phosphoribosyltransferase | 106, 106, 106 | 104 | HPRT1 | HPRT\_HUMAN | Hypoxanthine-guanine phosphoribosyltransferase | 3C(125.0477) 3C(15.9949) 3C(31.9988) | NaN | X | X | X | NaN | NaN | 2.591594e+05 | 2.591594e+05 | NaN | NaN | SYCNDQSTGDIK |
| PIK3CB | PK3CB\_HUMAN | Phosphatidylinositol 4,5-bisphosphate 3-kinase catalytic subunit beta isoform | 699 | 680 | PIK3CB | PK3CB\_HUMAN | Phosphatidylinositol 4,5-bisphosphate 3-kinase catalytic subunit beta isoform | 20C(-34.0809) | NaN | NaN | NaN | NaN | X | NaN | 2.586601e+05 | 2.586601e+05 | NaN | NaN | SEVHIPAVSVQFGVILEAYCRGSVGHMK |
| EEF2 | EF2\_HUMAN | Elongation factor 2 | 290, 290 | 288 | EEF2 | EF2\_HUMAN | Elongation factor 2 | 3C(125.0477) 3C(57.0215) | X | X | NaN | NaN | NaN | NaN | 2.581143e+05 | 2.581143e+05 | NaN | NaN | TFCQLILDPIFK |
| SPTA1 | SPTA1\_HUMAN | Spectrin alpha chain, erythrocytic 1 | 324 | 320 | SPTA1 | SPTA1\_HUMAN | Spectrin alpha chain, erythrocytic 1 | 5C(125.0477) | NaN | X | NaN | NaN | NaN | NaN | 2.580087e+05 | NaN | 2.580087e+05 | NaN | VKELCAK |
| AP2B1 | AP2B1\_HUMAN | AP-2 complex subunit beta | 391 | 391 | AP2B1 | AP2B1\_HUMAN | AP-2 complex subunit beta | 1C(125.0477) | NaN | X | NaN | NaN | NaN | NaN | 2.558107e+05 | 2.398717e+05 | 1.593899e+04 | NaN | CVSTLLDLIQTK |
| ARHGAP17 | RHG17\_HUMAN | Rho GTPase-activating protein 17 | 305 | 301 | ARHGAP17 | RHG17\_HUMAN | Rho GTPase-activating protein 17 | 5C(125.0477) | NaN | X | NaN | NaN | NaN | NaN | 2.552968e+05 | 2.552968e+05 | NaN | NaN | AALDCSTSHLDEFYSDPHAVAGALK |
| FLG2 | FILA2\_HUMAN | Filaggrin-2 | 25 | 21 | FLG2 | FILA2\_HUMAN | Filaggrin-2 | 5C(125.0477) | NaN | X | NaN | NaN | NaN | NaN | 2.547390e+05 | 3.794421e+04 | 1.262924e+05 | 90502.4400 | QDGECGTLSK |
| DCUN1D1 | DCNL1\_HUMAN | DCN1-like protein 1 | 115 | 111 | DCUN1D1 | DCNL1\_HUMAN | DCN1-like protein 1 | 5C(125.0477) | NaN | X | NaN | NaN | NaN | NaN | 2.546912e+05 | 2.546912e+05 | NaN | NaN | AATQCEFSK |
| NAPRT | PNCB\_HUMAN | Nicotinate phosphoribosyltransferase | 463 | 454 | NAPRT | PNCB\_HUMAN | Nicotinate phosphoribosyltransferase | 10C(125.0477) | NaN | X | NaN | NaN | NaN | NaN | 2.533825e+05 | 2.533825e+05 | NaN | NaN | VWPPGAQEPCTVRPAQVEPLLR |
| ERMAP | ERMAP\_HUMAN | Erythroid membrane-associated protein | 412 | 405 | ERMAP | ERMAP\_HUMAN | Erythroid membrane-associated protein | 8C(125.0477) | NaN | X | NaN | NaN | NaN | NaN | 2.529282e+05 | NaN | 2.529282e+05 | NaN | NTAPLVICSELHK |
| PPP2R5D | 2A5D\_HUMAN | Serine/threonine-protein phosphatase 2A 56 kDa regulatory subunit delta isoform | 346 | 335 | PPP2R5D | 2A5D\_HUMAN | Serine/threonine-protein phosphatase 2A 56 kDa regulatory subunit delta isoform | 12C(125.0477) | NaN | X | NaN | NaN | NaN | NaN | 2.514284e+05 | 2.514284e+05 | NaN | NaN | SLSVYHPQLAYCVVQFLEK |
| PSMC2 | PRS7\_HUMAN | 26S proteasome regulatory subunit 7 | 377 | 376 | PSMC2 | PRS7\_HUMAN | 26S proteasome regulatory subunit 7 | 2C(125.0477) | NaN | X | NaN | NaN | NaN | NaN | 2.508344e+05 | 2.017008e+05 | 4.913356e+04 | NaN | LCPNSTGAEIR |
| FSCN1 | FSCN1\_HUMAN | Fascin | 80 | 75 | FSCN1 | FSCN1\_HUMAN | Fascin | 6C(125.0477) | NaN | X | NaN | NaN | NaN | NaN | 2.501828e+05 | 2.501828e+05 | NaN | NaN | DGNVTCER |
| ASAP1 | ASAP1\_HUMAN | Arf-GAP with SH3 domain, ANK repeat and PH domain-containing protein 1 | 373 | 368 | ASAP1 | ASAP1\_HUMAN | Arf-GAP with SH3 domain, ANK repeat and PH domain-containing protein 1 | 6C(125.0477) | NaN | X | NaN | NaN | NaN | NaN | 2.500441e+05 | 2.500441e+05 | NaN | NaN | LNLLTCQVKPNAEDKK |
| ALDOA | ALDOA\_HUMAN | Fructose-bisphosphate aldolase A | 240, 240, 240 | 216 | ALDOA | ALDOA\_HUMAN | Fructose-bisphosphate aldolase A | 25C(125.0477) 25C(15.9949) 25C(31.9988) | NaN | X | X | X | NaN | NaN | 2.498847e+05 | 2.498847e+05 | NaN | NaN | ALSDHHIYLEGTLLKPNMVTPGHACTQK |
| ANKRD28 | ANR28\_HUMAN | Serine/threonine-protein phosphatase 6 regulatory ankyrin repeat subunit A | 632 | 619 | ANKRD28 | ANR28\_HUMAN | Serine/threonine-protein phosphatase 6 regulatory ankyrin repeat subunit A | 14C(125.0477) | NaN | X | NaN | NaN | NaN | NaN | 2.496162e+05 | 2.496162e+05 | NaN | NaN | TPIHAAATNGHSECLR |
| DIAPH1 | DIAP1\_HUMAN | Protein diaphanous homolog 1 | 1227 | 1225 | DIAPH1 | DIAP1\_HUMAN | Protein diaphanous homolog 1 | 3C(125.0477) | NaN | X | NaN | NaN | NaN | NaN | 2.495758e+05 | 1.688778e+05 | 8.069795e+04 | NaN | AGCAVTSLLASELTK |
| UFC1 | UFC1\_HUMAN | Ubiquitin-fold modifier-conjugating enzyme 1 | 116 | 115 | UFC1 | UFC1\_HUMAN | Ubiquitin-fold modifier-conjugating enzyme 1 | 2C(125.0477) | NaN | X | NaN | NaN | NaN | NaN | 2.490090e+05 | 2.490090e+05 | NaN | NaN | ICLTDHFKPLWAR |
| KRT6A | K2C6A\_HUMAN | Keratin, type II cytoskeletal 6A | 51 | 43 | KRT6A | K2C6A\_HUMAN | Keratin, type II cytoskeletal 6A | 9C(125.0477) | NaN | X | NaN | NaN | NaN | NaN | 2.486918e+05 | NaN | 2.486918e+05 | NaN | GSGGLGGACGGAGFGSR |
| PPP2R5D | 2A5D\_HUMAN | Serine/threonine-protein phosphatase 2A 56 kDa regulatory subunit delta isoform | 17 | 17 | PPP2R5D | 2A5D\_HUMAN | Serine/threonine-protein phosphatase 2A 56 kDa regulatory subunit delta isoform | 1C(125.0477) | NaN | X | NaN | NaN | NaN | NaN | 2.474867e+05 | 2.273239e+05 | 2.016284e+04 | NaN | CTAKPSSSGK |
| ANK1 | ANK1\_HUMAN | Ankyrin-1 | 348 | 323 | ANK1 | ANK1\_HUMAN | Ankyrin-1 | 26C(125.0477) | NaN | X | NaN | NaN | NaN | NaN | 2.471953e+05 | 2.471953e+05 | NaN | NaN | LLLQYDAEIDDITLDHLTPLHVAAHCGHHR |
| KRT4 | K2C4\_HUMAN | Keratin, type II cytoskeletal 4 | 17 | 14 | KRT4 | K2C4\_HUMAN | Keratin, type II cytoskeletal 4 | 4C(125.0477) | NaN | X | NaN | NaN | NaN | NaN | 2.471834e+05 | NaN | 2.471834e+05 | NaN | GFSCGSAIVGGGK |
| CLIC1 | CLIC1\_HUMAN | Chloride intracellular channel protein 1 | 178 | 166 | CLIC1 | CLIC1\_HUMAN | Chloride intracellular channel protein 1 | 13C(125.0477) | NaN | X | NaN | NaN | NaN | NaN | 2.468241e+05 | 2.468241e+05 | NaN | NaN | KFLDGNELTLADCNLLPK |
| CCT4 | TCPD\_HUMAN | T-complex protein 1 subunit delta | 337 | 327 | CCT4 | TCPD\_HUMAN | T-complex protein 1 subunit delta | 11C(125.0477) | NaN | X | NaN | NaN | NaN | NaN | 2.465931e+05 | 2.465931e+05 | NaN | NaN | DIEREDIEFICK |
| CLIC4 | CLIC4\_HUMAN | Chloride intracellular channel protein 4 | 189 | 178 | CLIC4 | CLIC4\_HUMAN | Chloride intracellular channel protein 4 | 12C(125.0477) | NaN | X | NaN | NaN | NaN | NaN | 2.463776e+05 | 2.463776e+05 | NaN | NaN | FLDGNEMTLADCNLLPK |
| ALB | ALBU\_HUMAN | Albumin | 125 | 123 | ALB | ALBU\_HUMAN | Albumin | 3C(57.0215) | X | NaN | NaN | NaN | NaN | NaN | 2.462995e+05 | 4.578354e+04 | 2.005159e+05 | NaN | NECFLQHKDDNPNLPR |
| PSMD12 | PSD12\_HUMAN | 26S proteasome non-ATPase regulatory subunit 12 | 193 | 192 | PSMD12 | PSD12\_HUMAN | 26S proteasome non-ATPase regulatory subunit 12 | 2C(125.0477) | NaN | X | NaN | NaN | NaN | NaN | 2.460983e+05 | 2.460983e+05 | NaN | NaN | LCLAVKDYIR |
| RNH1 | RINI\_HUMAN | Ribonuclease inhibitor | 277, 266 | 260 | RNH1 | RINI\_HUMAN | Ribonuclease inhibitor | 18C(125.0477) 7C(15.9949) | NaN | X | X | NaN | NaN | NaN | 2.451306e+05 | 2.451306e+05 | NaN | NaN | TLWIWECGITAKGCGDLCR |
| PSME1 | PSME1\_HUMAN | Proteasome activator complex subunit 1 | 101, 101, 106 | 92 | PSME1 | PSME1\_HUMAN | Proteasome activator complex subunit 1 | 10C(125.0477) 10C(31.9988) 15C(15.9949) | NaN | X | X | X | NaN | NaN | 2.446075e+05 | 2.446075e+05 | NaN | NaN | GEDEDKGPPCGPVNCNEK |
| OSTF1 | OSTF1\_HUMAN | Osteoclast-stimulating factor 1 | 114 | 105 | OSTF1 | OSTF1\_HUMAN | Osteoclast-stimulating factor 1 | 10C(125.0477) | NaN | X | NaN | NaN | NaN | NaN | 2.444686e+05 | 2.444686e+05 | NaN | NaN | AGSTALYWACHGGHK |
| TATDN1 | TATD1\_HUMAN | Deoxyribonuclease TATDN1 | 168 | 168 | TATDN1 | TATD1\_HUMAN | Deoxyribonuclease TATDN1 | 1C(125.0477) | NaN | X | NaN | NaN | NaN | NaN | 2.443297e+05 | 2.443297e+05 | NaN | NaN | CVGGVVHSFDGTK |
| NADSYN1 | NADE\_HUMAN | Glutamine-dependent NAD(+) synthetase | 531 | 529 | NADSYN1 | NADE\_HUMAN | Glutamine-dependent NAD(+) synthetase | 3C(125.0477) | NaN | X | NaN | NaN | NaN | NaN | 2.441511e+05 | 2.441511e+05 | NaN | NaN | YDCSSADINPIGGISK |
| UBFD1 | UBFD1\_HUMAN | Ubiquitin domain-containing protein UBFD1 | 185 | 176 | UBFD1 | UBFD1\_HUMAN | Ubiquitin domain-containing protein UBFD1 | 10C(125.0477) | NaN | X | NaN | NaN | NaN | NaN | 2.440585e+05 | 2.440585e+05 | NaN | NaN | AEENKKEPLCR |
| PFKM | PFKAM\_HUMAN | ATP-dependent 6-phosphofructokinase, muscle type | 114 | 109 | PFKM | PFKAM\_HUMAN | ATP-dependent 6-phosphofructokinase, muscle type | 6C(125.0477) | NaN | X | NaN | NaN | NaN | NaN | 2.436535e+05 | 2.436535e+05 | NaN | NaN | GITNLCVIGGDGSLTGADTFR |
| NUDT3 | NUDT3\_HUMAN | Diphosphoinositol polyphosphate phosphohydrolase 1 | 68 | 66 | NUDT3 | NUDT3\_HUMAN | Diphosphoinositol polyphosphate phosphohydrolase 1 | 3C(125.0477) | NaN | X | NaN | NaN | NaN | NaN | 2.428754e+05 | 2.428754e+05 | NaN | NaN | EVCEEAGVK |
| PREP | PPCE\_HUMAN | Prolyl endopeptidase | 352 | 346 | PREP | PPCE\_HUMAN | Prolyl endopeptidase | 7C(125.0477) | NaN | X | NaN | NaN | NaN | NaN | 2.426577e+05 | 2.426577e+05 | NaN | NaN | SNFLVLCYLHDVK |
| FLOT2 | FLOT2\_HUMAN | Flotillin-2 | 88 | 82 | FLOT2 | FLOT2\_HUMAN | Flotillin-2 | 7C(125.0477) | NaN | X | NaN | NaN | NaN | NaN | 2.423887e+05 | 1.356092e+05 | 1.067795e+05 | NaN | ELLAVACEQFLGK |
| TIPRL | TIPRL\_HUMAN | TIP41-like protein | 14 | 12 | TIPRL | TIPRL\_HUMAN | TIP41-like protein | 3C(125.0477) | NaN | X | NaN | NaN | NaN | NaN | 2.417490e+05 | 2.417490e+05 | NaN | NaN | DFCFGPWK |
| PAFAH1B1 | LIS1\_HUMAN | Platelet-activating factor acetylhydrolase IB subunit beta | 252 | 239 | PAFAH1B1 | LIS1\_HUMAN | Platelet-activating factor acetylhydrolase IB subunit beta | 14C(125.0477) | NaN | X | NaN | NaN | NaN | NaN | 2.414547e+05 | 2.414547e+05 | NaN | NaN | MVRPNQDGTLIASCSNDQTVR |
| SPTB | SPTB1\_HUMAN | Spectrin beta chain, erythrocytic | 183 | 177 | SPTB | SPTB1\_HUMAN | Spectrin beta chain, erythrocytic | 7C(125.0477) | NaN | X | NaN | NaN | NaN | NaN | 2.413959e+05 | 2.413959e+05 | NaN | NaN | DALLLWCQMK |
| TTC38 | TTC38\_HUMAN | Tetratricopeptide repeat protein 38 | 28 | 18 | TTC38 | TTC38\_HUMAN | Tetratricopeptide repeat protein 38 | 11C(125.0477) | NaN | X | NaN | NaN | NaN | NaN | 2.404557e+05 | 2.404557e+05 | NaN | NaN | LPLSTTSNEACK |
| RANBP1 | RANG\_HUMAN | Ran-specific GTPase-activating protein | 132 | 119 | RANBP1 | RANG\_HUMAN | Ran-specific GTPase-activating protein | 14C(125.0477) | NaN | X | NaN | NaN | NaN | NaN | 2.401388e+05 | 2.401388e+05 | NaN | NaN | AWVWNTHADFADECPKPELLAIR |
| MTAP | MTAP\_HUMAN | S-methyl-5'-thioadenosine phosphorylase | 55 | 50 | MTAP | MTAP\_HUMAN | S-methyl-5'-thioadenosine phosphorylase | 6C(125.0477) | NaN | X | NaN | NaN | NaN | NaN | 2.393634e+05 | 2.393634e+05 | NaN | NaN | IKNVDCVLLAR |
| GET4 | GET4\_HUMAN | Golgi to ER traffic protein 4 homolog | 160 | 156 | GET4 | GET4\_HUMAN | Golgi to ER traffic protein 4 homolog | 5C(125.0477) | NaN | X | NaN | NaN | NaN | NaN | 2.393009e+05 | 2.393009e+05 | NaN | NaN | EQNYCESR |
| XPOT | XPOT\_HUMAN | Exportin-T | 56 | 54 | XPOT | XPOT\_HUMAN | Exportin-T | 3C(125.0477) | NaN | X | NaN | NaN | NaN | NaN | 2.388130e+05 | 2.388130e+05 | NaN | NaN | FFCFQVLEHQVK |
| PGM2L1 | PGM2L\_HUMAN | Glucose 1,6-bisphosphate synthase | 494 | 489 | PGM2L1 | PGM2L\_HUMAN | Glucose 1,6-bisphosphate synthase | 6C(125.0477) | NaN | X | NaN | NaN | NaN | NaN | 2.386756e+05 | 2.386756e+05 | NaN | NaN | TSYFLCYEPPTIK |
| ACLY | ACLY\_HUMAN | ATP-citrate synthase | 229 | 221 | ACLY | ACLY\_HUMAN | ATP-citrate synthase | 9C(125.0477) | NaN | X | NaN | NaN | NaN | NaN | 2.385315e+05 | 1.080454e+05 | 1.304861e+05 | NaN | VDATADYICK |
| UBR4 | UBR4\_HUMAN | E3 ubiquitin-protein ligase UBR4 | 4146 | 4143 | UBR4 | UBR4\_HUMAN | E3 ubiquitin-protein ligase UBR4 | 4C(125.0477) | NaN | X | NaN | NaN | NaN | NaN | 2.383980e+05 | 2.383980e+05 | NaN | NaN | QAACTIVEALATIPSRK |
| FAHD2A | FAH2A\_HUMAN | Fumarylacetoacetate hydrolase domain-containing protein 2A | 301 | 295 | FAHD2A | FAH2A\_HUMAN | Fumarylacetoacetate hydrolase domain-containing protein 2A | 7C(125.0477) | NaN | X | NaN | NaN | NaN | NaN | 2.383673e+05 | 2.383673e+05 | NaN | NaN | KGDEVQCEIEELGVIINK |
| KPNB1 | IMB1\_HUMAN | Importin subunit beta-1 | 765 | 763 | KPNB1 | IMB1\_HUMAN | Importin subunit beta-1 | 3C(125.0477) | NaN | X | NaN | NaN | NaN | NaN | 2.380054e+05 | 2.380054e+05 | NaN | NaN | ESCLEAYTGIVQGLK |
| NUDCD1 | NUDC1\_HUMAN | NudC domain-containing protein 1 | 376 | 372 | NUDCD1 | NUDC1\_HUMAN | NudC domain-containing protein 1 | 5C(125.0477) | NaN | X | NaN | NaN | NaN | NaN | 2.377063e+05 | 2.377063e+05 | NaN | NaN | DSAQCAAIAER |
| USP14 | UBP14\_HUMAN | Ubiquitin carboxyl-terminal hydrolase 14 | 277, 277, 277 | 270 | USP14 | UBP14\_HUMAN | Ubiquitin carboxyl-terminal hydrolase 14 | 8C(-34.0809) 8C(125.0477) 8C(31.9988) | NaN | X | NaN | X | X | NaN | 2.374827e+05 | 2.374827e+05 | NaN | NaN | ENQLQLSCFINQEVK |
| TXNL1 | TXNL1\_HUMAN | Thioredoxin-like protein 1 | 161 | 153 | TXNL1 | TXNL1\_HUMAN | Thioredoxin-like protein 1 | 9C(125.0477) | NaN | X | NaN | NaN | NaN | NaN | 2.370923e+05 | 2.370923e+05 | NaN | NaN | DTTFLESDCDEQLLITVAFNQPVK |
| PGP | PGP\_HUMAN | Glycerol-3-phosphate phosphatase | 297 | 291 | PGP | PGP\_HUMAN | Glycerol-3-phosphate phosphatase | 7C(125.0477) | NaN | X | NaN | NaN | NaN | NaN | 2.369100e+05 | 2.121901e+05 | 2.471989e+04 | NaN | NNQESDCVSK |
| CARS1 | SYCC\_HUMAN | Cysteine--tRNA ligase, cytoplasmic | 27 | 14 | CARS1 | SYCC\_HUMAN | Cysteine--tRNA ligase, cytoplasmic | 14C(125.0477) | NaN | X | NaN | NaN | NaN | NaN | 2.367135e+05 | 2.367135e+05 | NaN | NaN | VQPQWSPPAGTQPCR |
| HPRT1 | HPRT\_HUMAN | Hypoxanthine-guanine phosphoribosyltransferase | 23 | 5 | HPRT1 | HPRT\_HUMAN | Hypoxanthine-guanine phosphoribosyltransferase | 19C(125.0477) | NaN | X | NaN | NaN | NaN | NaN | 2.357963e+05 | 2.357963e+05 | NaN | NaN | SPGVVISDDEPGYDLDLFCIPNHYAEDLER |
| MYH9 | MYH9\_HUMAN | Myosin-9 | 671 | 671 | MYH9 | MYH9\_HUMAN | Myosin-9 | 1C(125.0477) | NaN | X | NaN | NaN | NaN | NaN | 2.356295e+05 | 1.769271e+05 | 5.870242e+04 | NaN | CIIPNHEK |
| RELL1 | RELL1\_HUMAN | RELT-like protein 1 | 88 | 88 | RELL1 | RELL1\_HUMAN | RELT-like protein 1 | 1C(125.0477) | NaN | X | NaN | NaN | NaN | NaN | 2.355848e+05 | 2.104242e+04 | 2.145424e+05 | NaN | CTTEAEQDIEEEKVEK |
| MPP1 | EM55\_HUMAN | 55 kDa erythrocyte membrane protein | 94 | 92 | MPP1 | EM55\_HUMAN | 55 kDa erythrocyte membrane protein | 3C(125.0477) | NaN | X | NaN | NaN | NaN | NaN | 2.355271e+05 | NaN | 1.897843e+05 | 45742.7800 | QSCTVAR |
| PDE12 | PDE12\_HUMAN | 2',5'-phosphodiesterase 12 | 530 | 530 | PDE12 | PDE12\_HUMAN | 2',5'-phosphodiesterase 12 | 1C(125.0477) | NaN | X | NaN | NaN | NaN | NaN | 2.352803e+05 | 2.352803e+05 | NaN | NaN | CNMSLTHFFK |
| PFKL | PFKAL\_HUMAN | ATP-dependent 6-phosphofructokinase, liver type | 114 | 100 | PFKL | PFKAL\_HUMAN | ATP-dependent 6-phosphofructokinase, liver type | 15C(125.0477) | NaN | X | NaN | NaN | NaN | NaN | 2.351263e+05 | 2.351263e+05 | NaN | NaN | AAAYNLVQHGITNLCVIGGDGSLTGANIFR |
| SOD1 | SODC\_HUMAN | Superoxide dismutase [Cu-Zn] | 112 | 81 | SOD1 | SODC\_HUMAN | Superoxide dismutase [Cu-Zn] | 32C(125.0477) | NaN | X | NaN | NaN | NaN | NaN | 2.349673e+05 | 2.349673e+05 | NaN | NaN | HVGDLGNVTADKDGVADVSIEDSVISLSGDHCIIGR |
| KRT2 | K22E\_HUMAN | Keratin, type II cytoskeletal 2 epidermal | 489, 489 | 482 | KRT2 | K22E\_HUMAN | Keratin, type II cytoskeletal 2 epidermal | 8C(125.0477) 8C(15.9949) | NaN | X | X | NaN | NaN | NaN | 2.349667e+05 | NaN | 2.349667e+05 | NaN | KLLEGEECR |
| HBE1 | HBE\_HUMAN | Hemoglobin subunit epsilon | 94 | 89 | HBE1 | HBE\_HUMAN | Hemoglobin subunit epsilon | 6C(125.0477) | NaN | X | NaN | NaN | NaN | NaN | 2.347792e+05 | 2.347792e+05 | NaN | NaN | LSELHCDKLHVDPENFK |
| PIEZO1 | PIEZ1\_HUMAN | Piezo-type mechanosensitive ion channel component 1 | 2481 | 2480 | PIEZO1 | PIEZ1\_HUMAN | Piezo-type mechanosensitive ion channel component 1 | 2C(125.0477) | NaN | X | NaN | NaN | NaN | NaN | 2.344269e+05 | 2.344269e+05 | 0.000000e+00 | NaN | LCQDIFLVR |
| ALB | ALBU\_HUMAN | Albumin | 125, 125 | 123 | ALB | ALBU\_HUMAN | Albumin | 3C(125.0477) 3C(57.0215) | X | X | NaN | NaN | NaN | NaN | 2.341481e+05 | NaN | 2.341481e+05 | NaN | NECFLQHK |
| HNRNPD | HNRPD\_HUMAN | Heterogeneous nuclear ribonucleoprotein D0 | 226 | 224 | HNRNPD | HNRPD\_HUMAN | Heterogeneous nuclear ribonucleoprotein D0 | 3C(125.0477) | NaN | X | NaN | NaN | NaN | NaN | 2.339870e+05 | 2.339870e+05 | NaN | NaN | GFCFITFKEEEPVKK |
| TLN2 | TLN2\_HUMAN | Talin-2 | 1928 | 1921 | TLN2 | TLN2\_HUMAN | Talin-2 | 8C(125.0477) | NaN | X | NaN | NaN | NaN | NaN | 2.332973e+05 | 2.332973e+05 | NaN | NaN | VQDLGHGCIFLVQK |
| SERPINB9 | SPB9\_HUMAN | Serpin B9 | 370 | 364 | SERPINB9 | SPB9\_HUMAN | Serpin B9 | 7C(125.0477) | NaN | X | NaN | NaN | NaN | NaN | 2.332445e+05 | 2.332445e+05 | NaN | NaN | ANSILFCGR |
| UBR4 | UBR4\_HUMAN | E3 ubiquitin-protein ligase UBR4 | 122 | 115 | UBR4 | UBR4\_HUMAN | E3 ubiquitin-protein ligase UBR4 | 8C(125.0477) | NaN | X | NaN | NaN | NaN | NaN | 2.329799e+05 | 2.329799e+05 | NaN | NaN | LENPDEACAVSQK |
| HUWE1 | HUWE1\_HUMAN | E3 ubiquitin-protein ligase HUWE1 | 3213 | 3211 | HUWE1 | HUWE1\_HUMAN | E3 ubiquitin-protein ligase HUWE1 | 3C(125.0477) | NaN | X | NaN | NaN | NaN | NaN | 2.327102e+05 | 2.327102e+05 | NaN | NaN | NLCYHAQTR |
| HSP90AA1 | HS90A\_HUMAN | Heat shock protein HSP 90-alpha | 420 | 420 | HSP90AA1 | HS90A\_HUMAN | Heat shock protein HSP 90-alpha | 1C(125.0477) | NaN | X | NaN | NaN | NaN | NaN | 2.323207e+05 | 2.323207e+05 | NaN | NaN | CLELFTELAEDKENYK |
| GDI2 | GDIB\_HUMAN | Rab GDP dissociation inhibitor beta | 302 | 300 | GDI2 | GDIB\_HUMAN | Rab GDP dissociation inhibitor beta | 3C(125.0477) | NaN | X | NaN | NaN | NaN | NaN | 2.322895e+05 | 2.322895e+05 | NaN | NaN | VICILSHPIK |
| USP9X | USP9X\_HUMAN | Probable ubiquitin carboxyl-terminal hydrolase FAF-X | 1323 | 1311 | USP9X | USP9X\_HUMAN | Probable ubiquitin carboxyl-terminal hydrolase FAF-X | 13C(125.0477) | NaN | X | NaN | NaN | NaN | NaN | 2.318946e+05 | 2.318946e+05 | NaN | NaN | AWQTFIIDLLLHCHSK |
| CCT4 | TCPD\_HUMAN | T-complex protein 1 subunit delta | 252 | 246 | CCT4 | TCPD\_HUMAN | T-complex protein 1 subunit delta | 7C(125.0477) | NaN | X | NaN | NaN | NaN | NaN | 2.318415e+05 | 2.318415e+05 | NaN | NaN | IGLIQFCLSAPK |
| KPNA6 | IMA7\_HUMAN | Importin subunit alpha-7 | 467 | 459 | KPNA6 | IMA7\_HUMAN | Importin subunit alpha-7 | 9C(125.0477) | NaN | X | NaN | NaN | NaN | NaN | 2.314761e+05 | 2.314761e+05 | NaN | NaN | SGSGVNPYCGLIEEAYGLDK |
| GNAI3 | GNAI3\_HUMAN | Guanine nucleotide-binding protein G(i) subunit alpha-3 | 325 | 318 | GNAI3 | GNAI3\_HUMAN | Guanine nucleotide-binding protein G(i) subunit alpha-3 | 8C(125.0477) | NaN | X | NaN | NaN | NaN | NaN | 2.308905e+05 | 2.217481e+05 | 9.142418e+03 | NaN | EIYTHFTCATDTK |
| MAPK1 | MK01\_HUMAN | Mitogen-activated protein kinase 1 | 65 | 56 | MAPK1 | MK01\_HUMAN | Mitogen-activated protein kinase 1 | 10C(125.0477) | NaN | X | NaN | NaN | NaN | NaN | 2.306741e+05 | 2.306741e+05 | NaN | NaN | ISPFEHQTYCQR |
| CAP1 | CAP1\_HUMAN | Adenylyl cyclase-associated protein 1 | 427 | 423 | CAP1 | CAP1\_HUMAN | Adenylyl cyclase-associated protein 1 | 5C(125.0477) | NaN | X | NaN | NaN | NaN | NaN | 2.302311e+05 | 7.384802e+04 | 1.563831e+05 | NaN | NSLDCEIVSAK |
| CAPN5 | CAN5\_HUMAN | Calpain-5 | 150 | 140 | CAPN5 | CAN5\_HUMAN | Calpain-5 | 11C(125.0477) | NaN | X | NaN | NaN | NaN | NaN | 2.299310e+05 | 2.299310e+05 | NaN | NaN | LPTVNNQLIYCHSNSR |
| ENDOD1 | ENDD1\_HUMAN | Endonuclease domain-containing 1 protein | 269 | 257 | ENDOD1 | ENDD1\_HUMAN | Endonuclease domain-containing 1 protein | 13C(125.0477) | NaN | X | NaN | NaN | NaN | NaN | 2.299186e+05 | 2.299186e+05 | NaN | NaN | LLPFNPQLFQNNCGETEQDTEK |
| NAP1L4 | NP1L4\_HUMAN | Nucleosome assembly protein 1-like 4 | 77 | 77 | NAP1L4 | NP1L4\_HUMAN | Nucleosome assembly protein 1-like 4 | 1C(125.0477) | NaN | X | NaN | NaN | NaN | NaN | 2.299077e+05 | 1.128830e+05 | 1.170247e+05 | NaN | CAHIEAK |
| PSMD9 | PSMD9\_HUMAN | 26S proteasome non-ATPase regulatory subunit 9 | 59 | 49 | PSMD9 | PSMD9\_HUMAN | 26S proteasome non-ATPase regulatory subunit 9 | 11C(125.0477) | NaN | X | NaN | NaN | NaN | NaN | 2.297440e+05 | 2.297440e+05 | NaN | NaN | GIGMNEPLVDCEGYPR |
| CDC42 | CDC42\_HUMAN | Cell division control protein 42 homolog | 6 | 6 | CDC42 | CDC42\_HUMAN | Cell division control protein 42 homolog | 1C(125.0477) | NaN | X | NaN | NaN | NaN | NaN | 2.295259e+05 | NaN | 1.524284e+05 | 77097.5300 | CVVVGDGAVGK |
| TAGLN2 | TAGL2\_HUMAN | Transgelin-2 | 63 | 58 | TAGLN2 | TAGL2\_HUMAN | Transgelin-2 | 6C(125.0477) | NaN | X | NaN | NaN | NaN | NaN | 2.293831e+05 | 2.293831e+05 | NaN | NaN | DGTVLCELINALYPEGQAPVKK |
| PFAS | PUR4\_HUMAN | Phosphoribosylformylglycinamidine synthase | 975 | 968 | PFAS | PUR4\_HUMAN | Phosphoribosylformylglycinamidine synthase | 8C(125.0477) | NaN | X | NaN | NaN | NaN | NaN | 2.289815e+05 | 2.289815e+05 | NaN | NaN | YRDAGLHCLELGHTGEAGPHAMVR |
| PPP6R3 | PP6R3\_HUMAN | Serine/threonine-protein phosphatase 6 regulatory subunit 3 | 500 | 496 | PPP6R3 | PP6R3\_HUMAN | Serine/threonine-protein phosphatase 6 regulatory subunit 3 | 5C(125.0477) | NaN | X | NaN | NaN | NaN | NaN | 2.289053e+05 | 2.289053e+05 | NaN | NaN | WETFCTSSLGETNKR |
| GSS | GSHB\_HUMAN | Glutathione synthetase | 294 | 294 | GSS | GSHB\_HUMAN | Glutathione synthetase | 1C(125.0477) | NaN | X | NaN | NaN | NaN | NaN | 2.284896e+05 | 2.284896e+05 | NaN | NaN | CPDIATQLAGTKK |
| OLA1 | OLA1\_HUMAN | Obg-like ATPase 1 | 75 | 71 | OLA1 | OLA1\_HUMAN | Obg-like ATPase 1 | 5C(125.0477) | NaN | X | NaN | NaN | NaN | NaN | 2.284774e+05 | 2.284774e+05 | NaN | NaN | FDFLCQYHKPASK |
| IGHG3 | IGHG3\_HUMAN | Immunoglobulin heavy constant gamma 3 | 14, 14 | 5 | IGHG3 | IGHG3\_HUMAN | Immunoglobulin heavy constant gamma 3 | 10C(-34.0809) 10C(57.0215) | X | NaN | NaN | NaN | X | NaN | 2.277600e+05 | NaN | 2.277600e+05 | NaN | GPSVFPLAPCSR |
| AP2A2 | AP2A2\_HUMAN | AP-2 complex subunit alpha-2 | 282 | 279 | AP2A2 | AP2A2\_HUMAN | AP-2 complex subunit alpha-2 | 4C(125.0477) | NaN | X | NaN | NaN | NaN | NaN | 2.274787e+05 | 6.209615e+04 | 1.653825e+05 | NaN | LTECLETILNK |
| CAND1 | CAND1\_HUMAN | Cullin-associated NEDD8-dissociated protein 1 | 940, 940, 940, 942, 942 | 939 | CAND1 | CAND1\_HUMAN | Cullin-associated NEDD8-dissociated protein 1 | 2C(125.0477) 2C(15.9949) 2C(31.9988) 4C(125.0477) 4C(31.9988) | NaN | X | X | X | NaN | NaN | 2.274098e+05 | 2.274098e+05 | NaN | NaN | HCECAEEGTR |
| G6PD | G6PD\_HUMAN | Glucose-6-phosphate 1-dehydrogenase | 232 | 228 | G6PD | G6PD\_HUMAN | Glucose-6-phosphate 1-dehydrogenase | 5C(125.0477) | NaN | X | NaN | NaN | NaN | NaN | 2.273674e+05 | 2.273674e+05 | NaN | NaN | DNIACVILTFK |
| DLG1 | DLG1\_HUMAN | Disks large homolog 1 | 378 | 371 | DLG1 | DLG1\_HUMAN | Disks large homolog 1 | 8C(125.0477) | NaN | X | NaN | NaN | NaN | NaN | 2.264985e+05 | 2.264985e+05 | NaN | NaN | LLAVNNVCLEEVTHEEAVTALK |
| RIPOR1 | RIPR1\_HUMAN | Rho family-interacting cell polarization regulator 1 | 1154, 1159 | 1151 | RIPOR1 | RIPR1\_HUMAN | Rho family-interacting cell polarization regulator 1 | 4C(15.9949) 9C(125.0477) | NaN | X | X | NaN | NaN | NaN | 2.264288e+05 | NaN | 2.264288e+05 | NaN | VAGCLALGCIK |
| CD55 | DAF\_HUMAN | Complement decay-accelerating factor | 65 | 65 | CD55 | DAF\_HUMAN | Complement decay-accelerating factor | 1C(125.0477) | NaN | X | NaN | NaN | NaN | NaN | 2.259138e+05 | NaN | 2.259138e+05 | NaN | CEESFVK |
| IPO4 | IPO4\_HUMAN | Importin-4 | 400 | 391 | IPO4 | IPO4\_HUMAN | Importin-4 | 10C(125.0477) | NaN | X | NaN | NaN | NaN | NaN | 2.259026e+05 | 2.259026e+05 | NaN | NaN | LLPPLLQIVCK |
| PITPNB | PIPNB\_HUMAN | Phosphatidylinositol transfer protein beta isoform | 13 | 9 | PITPNB | PIPNB\_HUMAN | Phosphatidylinositol transfer protein beta isoform | 5C(125.0477) | NaN | X | NaN | NaN | NaN | NaN | 2.258799e+05 | 2.258799e+05 | NaN | NaN | VVLPCSVQEYQVGQLYSVAEASK |
| PGM2L1 | PGM2L\_HUMAN | Glucose 1,6-bisphosphate synthase | 236 | 229 | PGM2L1 | PGM2L\_HUMAN | Glucose 1,6-bisphosphate synthase | 8C(125.0477) | NaN | X | NaN | NaN | NaN | NaN | 2.258516e+05 | 2.258516e+05 | NaN | NaN | RDPLQDICR |
| RFESD | RFESD\_HUMAN | Rieske domain-containing protein | 19 | 14 | RFESD | RFESD\_HUMAN | Rieske domain-containing protein | 6C(125.0477) | NaN | X | NaN | NaN | NaN | NaN | 2.256660e+05 | 2.256660e+05 | NaN | NaN | EYSSVCVGREDDIK |
| CAP1 | CAP1\_HUMAN | Adenylyl cyclase-associated protein 1 | 416 | 413 | CAP1 | CAP1\_HUMAN | Adenylyl cyclase-associated protein 1 | 4C(125.0477) | NaN | X | NaN | NaN | NaN | NaN | 2.248082e+05 | 1.887670e+05 | 3.604123e+04 | NaN | TDGCHAYLSK |
| UROS | HEM4\_HUMAN | Uroporphyrinogen-III synthase | 143 | 133 | UROS | HEM4\_HUMAN | Uroporphyrinogen-III synthase | 11C(125.0477) | NaN | X | NaN | NaN | NaN | NaN | 2.246253e+05 | 2.246253e+05 | NaN | NaN | ESSALPLLFPCGNLKR |
| HBD | HBD\_HUMAN | Hemoglobin subunit delta | 113 | 106 | HBD | HBD\_HUMAN | Hemoglobin subunit delta | 8C(125.0477) | NaN | X | NaN | NaN | NaN | NaN | 2.233556e+05 | NaN | NaN | 223355.5800 | LLGNVLVCVLAR |
| EPB42 | EPB42\_HUMAN | Protein 4.2 | 428 | 422 | EPB42 | EPB42\_HUMAN | Protein 4.2 | 7C(125.0477) | NaN | X | NaN | NaN | NaN | NaN | 2.229089e+05 | NaN | 2.229089e+05 | NaN | GVGSDRCEDITQNYK |
| USP9X | USP9X\_HUMAN | Probable ubiquitin carboxyl-terminal hydrolase FAF-X | 2293 | 2288 | USP9X | USP9X\_HUMAN | Probable ubiquitin carboxyl-terminal hydrolase FAF-X | 6C(125.0477) | NaN | X | NaN | NaN | NaN | NaN | 2.228952e+05 | 2.228952e+05 | NaN | NaN | KIIEDCSNSEETVK |
| PRDX5 | PRDX5\_HUMAN | Peroxiredoxin-5, mitochondrial | 204 | 192 | PRDX5 | PRDX5\_HUMAN | Peroxiredoxin-5, mitochondrial | 13C(125.0477) | NaN | X | NaN | NaN | NaN | NaN | 2.224759e+05 | 2.224759e+05 | NaN | NaN | ALNVEPDGTGLTCSLAPNIISQL |
| ANK1 | ANK1\_HUMAN | Ankyrin-1 | 864 | 862 | ANK1 | ANK1\_HUMAN | Ankyrin-1 | 3C(125.0477) | NaN | X | NaN | NaN | NaN | NaN | 2.213575e+05 | 2.213575e+05 | NaN | NaN | IPCAMPETVVIR |
| MRE11 | MRE11\_HUMAN | Double-strand break repair protein MRE11 | 75 | 71 | MRE11 | MRE11\_HUMAN | Double-strand break repair protein MRE11 | 5C(125.0477) | NaN | X | NaN | NaN | NaN | NaN | 2.210217e+05 | 2.210217e+05 | NaN | NaN | TLHTCLELLR |
| CUTA | CUTA\_HUMAN | Protein CutA | 96 | 93 | CUTA | CUTA\_HUMAN | Protein CutA | 4C(125.0477) | NaN | X | NaN | NaN | NaN | NaN | 2.210130e+05 | 2.210130e+05 | NaN | NaN | LAACVNLIPQITSIYEWK |
| THOP1 | THOP1\_HUMAN | Thimet oligopeptidase | 175 | 171 | THOP1 | THOP1\_HUMAN | Thimet oligopeptidase | 5C(125.0477) | NaN | X | NaN | NaN | NaN | NaN | 2.208910e+05 | 2.208910e+05 | NaN | NaN | LSLLCIDFNK |
| ALDH1A1 | AL1A1\_HUMAN | Aldehyde dehydrogenase 1A1 | 133 | 132 | ALDH1A1 | AL1A1\_HUMAN | Aldehyde dehydrogenase 1A1 | 2C(125.0477) | NaN | X | NaN | NaN | NaN | NaN | 2.206344e+05 | 4.120973e+04 | 1.794247e+05 | NaN | YCAGWADK |
| CD2AP | CD2AP\_HUMAN | CD2-associated protein | 595 | 588 | CD2AP | CD2AP\_HUMAN | CD2-associated protein | 8C(125.0477) | NaN | X | NaN | NaN | NaN | NaN | 2.206222e+05 | 2.206222e+05 | NaN | NaN | AQIIELLCIVEALKK |
| ALB | ALBU\_HUMAN | Albumin | 511, 511 | 509 | ALB | ALBU\_HUMAN | Albumin | 3C(125.0477) 3C(57.0215) | X | X | NaN | NaN | NaN | NaN | 2.193277e+05 | 2.193277e+05 | NaN | NaN | RPCFSALEVDETYVPK |
| GCHFR | GFRP\_HUMAN | GTP cyclohydrolase 1 feedback regulatory protein | 80 | 67 | GCHFR | GFRP\_HUMAN | GTP cyclohydrolase 1 feedback regulatory protein | 14C(125.0477) | NaN | X | NaN | NaN | NaN | NaN | 2.190758e+05 | 2.190758e+05 | NaN | NaN | VLSMTGVGQTLVWCLHKE |
| PPP2CB | PP2AB\_HUMAN | Serine/threonine-protein phosphatase 2A catalytic subunit beta isoform | 20 | 9 | PPP2CB | PP2AB\_HUMAN | Serine/threonine-protein phosphatase 2A catalytic subunit beta isoform | 12C(125.0477) | NaN | X | NaN | NaN | NaN | NaN | 2.185553e+05 | 2.185553e+05 | NaN | NaN | ELDQWVEQLNECK |
| CAP1 | CAP1\_HUMAN | Adenylyl cyclase-associated protein 1 | 93, 93 | 85 | CAP1 | CAP1\_HUMAN | Adenylyl cyclase-associated protein 1 | 9C(125.0477) 9C(31.9988) | NaN | X | NaN | X | NaN | NaN | 2.184371e+05 | NaN | 2.184371e+05 | NaN | ALLVTASQCQQPAENK |
| ALDH1A1 | AL1A1\_HUMAN | Aldehyde dehydrogenase 1A1 | 50 | 38 | ALDH1A1 | AL1A1\_HUMAN | Aldehyde dehydrogenase 1A1 | 13C(125.0477) | NaN | X | NaN | NaN | NaN | NaN | 2.183476e+05 | 2.183476e+05 | NaN | NaN | FPVFNPATEEELCQVEEGDKEDVDK |
| MINDY3 | MINY3\_HUMAN | Ubiquitin carboxyl-terminal hydrolase MINDY-3 | 77 | 76 | MINDY3 | MINY3\_HUMAN | Ubiquitin carboxyl-terminal hydrolase MINDY-3 | 2C(125.0477) | NaN | X | NaN | NaN | NaN | NaN | 2.176720e+05 | 2.176720e+05 | NaN | NaN | DCSEEEQK |
| CCT2 | TCPB\_HUMAN | T-complex protein 1 subunit beta | 535 | 527 | CCT2 | TCPB\_HUMAN | T-complex protein 1 subunit beta | 9C(125.0477) | NaN | X | NaN | NaN | NaN | NaN | 2.176262e+05 | 2.176262e+05 | NaN | NaN | KRVPDHHPC |
| STIP1 | STIP1\_HUMAN | Stress-induced-phosphoprotein 1 | 461 | 454 | STIP1 | STIP1\_HUMAN | Stress-induced-phosphoprotein 1 | 8C(125.0477) | NaN | X | NaN | NaN | NaN | NaN | 2.166454e+05 | 2.642212e+04 | 1.491271e+05 | 41096.1450 | ALDLDSSCK |
| AP2B1 | AP2B1\_HUMAN | AP-2 complex subunit beta | 123 | 118 | AP2B1 | AP2B1\_HUMAN | AP-2 complex subunit beta | 6C(125.0477) | NaN | X | NaN | NaN | NaN | NaN | 2.163206e+05 | NaN | 2.163206e+05 | NaN | ITEYLCEPLR |
| NUDCD2 | NUDC2\_HUMAN | NudC domain-containing protein 2 | 45 | 40 | NUDCD2 | NUDC2\_HUMAN | NudC domain-containing protein 2 | 6C(125.0477) | NaN | X | NaN | NaN | NaN | NaN | 2.160639e+05 | 1.483395e+05 | 6.772436e+04 | NaN | AQDIQCGLQSR |
| RHOA | RHOA\_HUMAN | Transforming protein RhoA | 20 | 19 | RHOA | RHOA\_HUMAN | Transforming protein RhoA | 2C(125.0477) | NaN | X | NaN | NaN | NaN | NaN | 2.158346e+05 | 2.158346e+05 | NaN | NaN | TCLLIVFSK |
| WDR77 | MEP50\_HUMAN | Methylosome protein 50 | 186, 172 | 165 | WDR77 | MEP50\_HUMAN | Methylosome protein 50 | 22C(15.9949) 8C(15.9949) | NaN | NaN | X | NaN | NaN | NaN | 2.157009e+05 | 2.157009e+05 | NaN | NaN | AHAAQVTCVAASPHKDSVFLSCSEDNR |
| TGM2 | TGM2\_HUMAN | Protein-glutamine gamma-glutamyltransferase 2 | 554 | 551 | TGM2 | TGM2\_HUMAN | Protein-glutamine gamma-glutamyltransferase 2 | 4C(125.0477) | NaN | X | NaN | NaN | NaN | NaN | 2.148414e+05 | 2.148414e+05 | NaN | NaN | YRDCLTESNLIK |
| ALDH16A1 | A16A1\_HUMAN | Aldehyde dehydrogenase family 16 member A1 | 472, 467, 467 | 459 | ALDH16A1 | A16A1\_HUMAN | Aldehyde dehydrogenase family 16 member A1 | 14C(31.9988) 9C(-34.0809) 9C(15.9949) | NaN | NaN | X | X | X | NaN | 2.147129e+05 | 2.147129e+05 | NaN | NaN | DPSVPTGGCKESGCSWHGGPDGLYEYLRPSGTPAR |
| ADPRS | ADPRS\_HUMAN | ADP-ribosylhydrolase ARH3 | 132 | 132 | ADPRS | ADPRS\_HUMAN | ADP-ribosylhydrolase ARH3 | 1C(125.0477) | NaN | X | NaN | NaN | NaN | NaN | 2.141540e+05 | 2.141540e+05 | NaN | NaN | CRDVFEPAR |
| PPP2R1A | 2AAA\_HUMAN | Serine/threonine-protein phosphatase 2A 65 kDa regulatory subunit A alpha isoform | 294 | 293 | PPP2R1A | 2AAA\_HUMAN | Serine/threonine-protein phosphatase 2A 65 kDa regulatory subunit A alpha isoform | 2C(125.0477) | NaN | X | NaN | NaN | NaN | NaN | 2.138430e+05 | NaN | 2.138430e+05 | NaN | DCEAEVR |
| ANKLE1 | ANKL1\_HUMAN | Ankyrin repeat and LEM domain-containing protein 1 | 571 | 568 | ANKLE1 | ANKL1\_HUMAN | Ankyrin repeat and LEM domain-containing protein 1 | 4C(305.3076) | NaN | NaN | NaN | NaN | NaN | X | 2.133734e+05 | 2.133734e+05 | NaN | NaN | QGHCYGVVAGWPPAR |
| NAA25 | NAA25\_HUMAN | N-alpha-acetyltransferase 25, NatB auxiliary subunit | 381 | 371 | NAA25 | NAA25\_HUMAN | N-alpha-acetyltransferase 25, NatB auxiliary subunit | 11C(125.0477) | NaN | X | NaN | NaN | NaN | NaN | 2.133460e+05 | 2.133460e+05 | NaN | NaN | VFVDLLPATQCTK |
| GPS1 | CSN1\_HUMAN | COP9 signalosome complex subunit 1 | 192 | 190 | GPS1 | CSN1\_HUMAN | COP9 signalosome complex subunit 1 | 3C(125.0477) | NaN | X | NaN | NaN | NaN | NaN | 2.130208e+05 | 2.130208e+05 | NaN | NaN | DYCTSAK |
| SCRN1 | SCRN1\_HUMAN | Secernin-1 | 54 | 52 | SCRN1 | SCRN1\_HUMAN | Secernin-1 | 3C(125.0477) | NaN | X | NaN | NaN | NaN | NaN | 2.130152e+05 | 2.130152e+05 | NaN | NaN | VECTYISIDQVPR |
| ANK1 | ANK1\_HUMAN | Ankyrin-1 | 476 | 467 | ANK1 | ANK1\_HUMAN | Ankyrin-1 | 10C(125.0477) | NaN | X | NaN | NaN | NaN | NaN | 2.129303e+05 | NaN | NaN | 212930.2700 | AKDDQTPLHCAAR |
| IGKC | IGKC\_HUMAN | Immunoglobulin kappa constant | 87, 87, 87 | 84 | IGKC | IGKC\_HUMAN | Immunoglobulin kappa constant | 4C(125.0477) 4C(31.9988) 4C(57.0215) | X | X | NaN | X | NaN | NaN | 2.128802e+05 | 2.128802e+05 | NaN | NaN | VYACEVTHQGLSSPVTK |
| CUL4A | CUL4A\_HUMAN | Cullin-4A | 483 | 479 | CUL4A | CUL4A\_HUMAN | Cullin-4A | 5C(125.0477) | NaN | X | NaN | NaN | NaN | NaN | 2.128776e+05 | 2.128776e+05 | NaN | NaN | LKHECGAAFTSK |
| UBA3 | UBA3\_HUMAN | NEDD8-activating enzyme E1 catalytic subunit | 164 | 158 | UBA3 | UBA3\_HUMAN | NEDD8-activating enzyme E1 catalytic subunit | 7C(125.0477) | NaN | X | NaN | NaN | NaN | NaN | 2.124266e+05 | 2.124266e+05 | NaN | NaN | QFHIIVCGLDSIIAR |
| PFAS | PUR4\_HUMAN | Phosphoribosylformylglycinamidine synthase | 336 | 331 | PFAS | PUR4\_HUMAN | Phosphoribosylformylglycinamidine synthase | 6C(125.0477) | NaN | X | NaN | NaN | NaN | NaN | 2.116568e+05 | 2.116568e+05 | NaN | NaN | IRDVQCTGR |
| PARK7 | PARK7\_HUMAN | Parkinson disease protein 7 | 46 | 33 | PARK7 | PARK7\_HUMAN | Parkinson disease protein 7 | 14C(15.9949) | NaN | NaN | X | NaN | NaN | NaN | 2.112789e+05 | 2.112789e+05 | NaN | NaN | VTVAGLAGKDPVQCSRDVVICPDASLEDAK |
| CTTN | SRC8\_HUMAN | Src substrate cortactin | 246 | 246 | CTTN | SRC8\_HUMAN | Src substrate cortactin | 1C(125.0477) | NaN | X | NaN | NaN | NaN | NaN | 2.110175e+05 | 1.968290e+05 | 1.418852e+04 | NaN | CALGWDHQEK |
| THOP1 | THOP1\_HUMAN | Thimet oligopeptidase | 246 | 239 | THOP1 | THOP1\_HUMAN | Thimet oligopeptidase | 8C(125.0477) | NaN | X | NaN | NaN | NaN | NaN | 2.109638e+05 | 2.109638e+05 | NaN | NaN | KVEEAFNCR |
| PITHD1 | PITH1\_HUMAN | PITH domain-containing protein 1 | 42 | 40 | PITHD1 | PITH1\_HUMAN | PITH domain-containing protein 1 | 3C(125.0477) | NaN | X | NaN | NaN | NaN | NaN | 2.100196e+05 | NaN | 2.100196e+05 | NaN | LQCLNESR |
| ACTBL2 | ACTBL\_HUMAN | Beta-actin-like protein 2 | 53, 53 | 41 | ACTBL2 | ACTBL\_HUMAN | Beta-actin-like protein 2 | 13C(125.0477) 13C(15.9949) | NaN | X | X | NaN | NaN | NaN | 2.090159e+05 | 2.090159e+05 | NaN | NaN | HQGVMVGMGQKDCYVGDEAQSK |
| SH3GLB2 | SHLB2\_HUMAN | Endophilin-B2 | 267 | 259 | SH3GLB2 | SHLB2\_HUMAN | Endophilin-B2 | 9C(125.0477) | NaN | X | NaN | NaN | NaN | NaN | 2.082024e+05 | 2.082024e+05 | NaN | NaN | SQTTYYAQCYR |
| SLC43A2 | LAT4\_HUMAN | Large neutral amino acids transporter small subunit 4 | 392 | 391 | SLC43A2 | LAT4\_HUMAN | Large neutral amino acids transporter small subunit 4 | 2C(125.0477) | NaN | X | NaN | NaN | NaN | NaN | 2.079900e+05 | 5.826896e+04 | 1.497210e+05 | NaN | ECEDASEEPEEK |
| GNB1 | GBB1\_HUMAN | Guanine nucleotide-binding protein G(I)/G(S)/G(T) subunit beta-1 | 204 | 198 | GNB1 | GBB1\_HUMAN | Guanine nucleotide-binding protein G(I)/G(S)/G(T) subunit beta-1 | 7C(125.0477) | NaN | X | NaN | NaN | NaN | NaN | 2.075897e+05 | NaN | 2.075897e+05 | NaN | LFVSGACDASAK |
| GNAS | GNAS1\_HUMAN | Guanine nucleotide-binding protein G(s) subunit alpha isoforms XLas | 817 | 809 | GNAS | GNAS1\_HUMAN | Guanine nucleotide-binding protein G(s) subunit alpha isoforms XLas | 9C(125.0477) | NaN | X | NaN | NaN | NaN | NaN | 2.069612e+05 | 2.069612e+05 | NaN | NaN | SNEYQLIDCAQYFLDK |
| RABGGTA | PGTA\_HUMAN | Geranylgeranyl transferase type-2 subunit alpha | 516 | 510 | RABGGTA | PGTA\_HUMAN | Geranylgeranyl transferase type-2 subunit alpha | 7C(125.0477) | NaN | X | NaN | NaN | NaN | NaN | 2.067558e+05 | 2.067558e+05 | NaN | NaN | LQELLLCNNR |
| GSTT1 | GSTT1\_HUMAN | Glutathione S-transferase theta-1 | 14 | 2 | GSTT1 | GSTT1\_HUMAN | Glutathione S-transferase theta-1 | 13C(125.0477) | NaN | X | NaN | NaN | NaN | NaN | 2.065925e+05 | 2.065925e+05 | NaN | NaN | GLELYLDLLSQPCR |
| PITHD1 | PITH1\_HUMAN | PITH domain-containing protein 1 | 187 | 181 | PITHD1 | PITH1\_HUMAN | PITH domain-containing protein 1 | 7C(125.0477) | NaN | X | NaN | NaN | NaN | NaN | 2.065891e+05 | 2.065891e+05 | NaN | NaN | RHEVTICNYEASANPADHR |
| UBE2O | UBE2O\_HUMAN | (E3-independent) E2 ubiquitin-conjugating enzyme | 585 | 570 | UBE2O | UBE2O\_HUMAN | (E3-independent) E2 ubiquitin-conjugating enzyme | 16C(125.0477) | NaN | X | NaN | NaN | NaN | NaN | 2.062203e+05 | 2.062203e+05 | NaN | NaN | SNDLFPVHHLDNNEFCPGDFVVDK |
| AP2A1 | AP2A1\_HUMAN | AP-2 complex subunit alpha-1 | 267 | 264 | AP2A1 | AP2A1\_HUMAN | AP-2 complex subunit alpha-1 | 4C(125.0477) | NaN | X | NaN | NaN | NaN | NaN | 2.057179e+05 | 1.386814e+05 | 6.703645e+04 | NaN | LLQCYPPPEDAAVK |
| ACTR3 | ARP3\_HUMAN | Actin-related protein 3 | 34 | 19 | ACTR3 | ARP3\_HUMAN | Actin-related protein 3 | 16C(125.0477) | NaN | X | NaN | NaN | NaN | NaN | 2.051160e+05 | 2.051160e+05 | NaN | NaN | LGYAGNTEPQFIIPSCIAIK |
| PDCD6IP | PDC6I\_HUMAN | Programmed cell death 6-interacting protein | 76 | 71 | PDCD6IP | PDC6I\_HUMAN | Programmed cell death 6-interacting protein | 6C(125.0477) | NaN | X | NaN | NaN | NaN | NaN | 2.050821e+05 | 1.862840e+05 | 1.879820e+04 | NaN | YYDQICSIEPK |
| PLAA | PLAP\_HUMAN | Phospholipase A-2-activating protein | 263 | 260 | PLAA | PLAP\_HUMAN | Phospholipase A-2-activating protein | 4C(125.0477) | NaN | X | NaN | NaN | NaN | NaN | 2.047881e+05 | 2.047881e+05 | NaN | NaN | HGECAQTIR |
| COLGALT1 | GT251\_HUMAN | Procollagen galactosyltransferase 1 | 369 | 361 | COLGALT1 | GT251\_HUMAN | Procollagen galactosyltransferase 1 | 9C(125.0477) | NaN | X | NaN | NaN | NaN | NaN | 2.044183e+05 | NaN | 2.044183e+05 | NaN | ALQAQEIECR |
| GCLC | GSH1\_HUMAN | Glutamate--cysteine ligase catalytic subunit | 501 | 493 | GCLC | GSH1\_HUMAN | Glutamate--cysteine ligase catalytic subunit | 9C(125.0477) | NaN | X | NaN | NaN | NaN | NaN | 2.041817e+05 | 1.648213e+05 | 3.936034e+04 | NaN | GGNAVVDGCGK |
| TUBB4B | TBB4B\_HUMAN | Tubulin beta-4B chain | 12 | 3 | TUBB4B | TBB4B\_HUMAN | Tubulin beta-4B chain | 10C(125.0477) | NaN | X | NaN | NaN | NaN | NaN | 2.040350e+05 | 2.040350e+05 | NaN | NaN | EIVHLQAGQCGNQIGAK |
| TPMT | TPMT\_HUMAN | Thiopurine S-methyltransferase | 234 | 229 | TPMT | TPMT\_HUMAN | Thiopurine S-methyltransferase | 6C(125.0477) | NaN | X | NaN | NaN | NaN | NaN | 2.037056e+05 | 2.037056e+05 | NaN | NaN | SWGIDCLFEK |
| TPP2 | TPP2\_HUMAN | Tripeptidyl-peptidase 2 | 1125 | 1118 | TPP2 | TPP2\_HUMAN | Tripeptidyl-peptidase 2 | 8C(125.0477) | NaN | X | NaN | NaN | NaN | NaN | 2.031911e+05 | 2.031911e+05 | NaN | NaN | STLVDALCRK |
| AGO2 | AGO2\_HUMAN | Protein argonaute-2 | 188 | 180 | AGO2 | AGO2\_HUMAN | Protein argonaute-2 | 9C(125.0477) | NaN | X | NaN | NaN | NaN | NaN | 2.027057e+05 | 2.027057e+05 | NaN | NaN | SFFTASEGCSNPLGGGR |
| WDR81 | WDR81\_HUMAN | WD repeat-containing protein 81 | 765 | 757 | WDR81 | WDR81\_HUMAN | WD repeat-containing protein 81 | 9C(125.0477) | NaN | X | NaN | NaN | NaN | NaN | 2.027043e+05 | 2.027043e+05 | NaN | NaN | VQPAVPLQCLLHR |
| CA8 | CAH8\_HUMAN | Carbonic anhydrase-related protein | 266 | 259 | CA8 | CAH8\_HUMAN | Carbonic anhydrase-related protein | 8C(125.0477) | NaN | X | NaN | NaN | NaN | NaN | 2.026506e+05 | 2.026506e+05 | NaN | NaN | GAELVEGCDGILGDNFRPTQPLSDR |
| KRT6C | K2C6C\_HUMAN | Keratin, type II cytoskeletal 6C | 51, 51 | 43 | KRT6C | K2C6C\_HUMAN | Keratin, type II cytoskeletal 6C | 9C(125.0477) 9C(31.9988) | NaN | X | NaN | X | NaN | NaN | 2.025824e+05 | NaN | NaN | 202582.4400 | GSGGLGGACGGAGFGSR |
| PARK7 | PARK7\_HUMAN | Parkinson disease protein 7 | 46 | 42 | PARK7 | PARK7\_HUMAN | Parkinson disease protein 7 | 5C(125.0477) | NaN | X | NaN | NaN | NaN | NaN | 2.025062e+05 | 2.025062e+05 | NaN | NaN | DPVQCSR |
| ANXA11 | ANX11\_HUMAN | Annexin A11 | 226 | 215 | ANXA11 | ANX11\_HUMAN | Annexin A11 | 12C(125.0477) | NaN | X | NaN | NaN | NaN | NaN | 2.022232e+05 | 2.022232e+05 | NaN | NaN | GFGTDEQAIIDCLGSR |
| COPS3 | CSN3\_HUMAN | COP9 signalosome complex subunit 3 | 25 | 14 | COPS3 | CSN3\_HUMAN | COP9 signalosome complex subunit 3 | 12C(125.0477) | NaN | X | NaN | NaN | NaN | NaN | 2.022130e+05 | 2.022130e+05 | NaN | NaN | QLSAQGQMTQLCELINK |
| HP | HPT\_HUMAN | Haptoglobin | 381 | 380 | HP | HPT\_HUMAN | Haptoglobin | 2C(57.0215) | X | NaN | NaN | NaN | NaN | NaN | 2.021740e+05 | NaN | 2.021740e+05 | NaN | SCAVAEYGVYVK |
| VPS35 | VPS35\_HUMAN | Vacuolar protein sorting-associated protein 35 | 286 | 285 | VPS35 | VPS35\_HUMAN | Vacuolar protein sorting-associated protein 35 | 2C(125.0477) | NaN | X | NaN | NaN | NaN | NaN | 2.019315e+05 | 2.019315e+05 | NaN | NaN | ACAELHQNVNVK |
| TNPO1 | TNPO1\_HUMAN | Transportin-1 | 683 | 674 | TNPO1 | TNPO1\_HUMAN | Transportin-1 | 10C(125.0477) | NaN | X | NaN | NaN | NaN | NaN | 2.018442e+05 | 2.018442e+05 | NaN | NaN | SNILTLMYQCMQDK |
| PGP | PGP\_HUMAN | Glycerol-3-phosphate phosphatase | 272 | 255 | PGP | PGP\_HUMAN | Glycerol-3-phosphate phosphatase | 18C(125.0477) | NaN | X | NaN | NaN | NaN | NaN | 2.010254e+05 | 2.010254e+05 | NaN | NaN | TVMVGDRLDTDILLGATCGLK |
| GMPR | GMPR1\_HUMAN | GMP reductase 1 | 316 | 314 | GMPR | GMPR1\_HUMAN | GMP reductase 1 | 3C(125.0477) | NaN | X | NaN | NaN | NaN | NaN | 2.008266e+05 | 2.008266e+05 | NaN | NaN | STCTYVGAAK |
| MDH1 | MDHC\_HUMAN | Malate dehydrogenase, cytoplasmic | 154 | 143 | MDH1 | MDHC\_HUMAN | Malate dehydrogenase, cytoplasmic | 12C(125.0477) | NaN | X | NaN | NaN | NaN | NaN | 2.006765e+05 | 2.006765e+05 | NaN | NaN | SAPSIPKENFSCLTR |
| HSPA4 | HSP74\_HUMAN | Heat shock 70 kDa protein 4 | 417 | 406 | HSPA4 | HSP74\_HUMAN | Heat shock 70 kDa protein 4 | 12C(125.0477) | NaN | X | NaN | NaN | NaN | NaN | 2.006380e+05 | 1.871540e+05 | 1.348403e+04 | NaN | WNSPAEEGSSDCEVFSK |
| PREP | PPCE\_HUMAN | Prolyl endopeptidase | 255 | 253 | PREP | PPCE\_HUMAN | Prolyl endopeptidase | 3C(125.0477) | NaN | X | NaN | NaN | NaN | NaN | 2.006343e+05 | 2.006343e+05 | NaN | NaN | EGCDPVNR |
| MTHFD1 | C1TC\_HUMAN | C-1-tetrahydrofolate synthase, cytoplasmic | 691 | 687 | MTHFD1 | C1TC\_HUMAN | C-1-tetrahydrofolate synthase, cytoplasmic | 5C(125.0477) | NaN | X | NaN | NaN | NaN | NaN | 1.997342e+05 | 1.997342e+05 | NaN | NaN | YSGLCPHVVVLVATVR |
| LGALS3 | LEG3\_HUMAN | Galectin-3 | 173 | 169 | LGALS3 | LEG3\_HUMAN | Galectin-3 | 5C(125.0477) | NaN | X | NaN | NaN | NaN | NaN | 1.996389e+05 | 9.519352e+04 | 1.044454e+05 | NaN | RVIVCNTK |
| ALB | ALBU\_HUMAN | Albumin | 302, 303 | 299 | ALB | ALBU\_HUMAN | Albumin | 4C(57.0215) 5C(57.0215) | X | NaN | NaN | NaN | NaN | NaN | 1.995693e+05 | NaN | 1.995693e+05 | NaN | LKECCEKPLLEK |
| ACTBL2 | ACTBL\_HUMAN | Beta-actin-like protein 2 | 53, 53, 53 | 52 | ACTBL2 | ACTBL\_HUMAN | Beta-actin-like protein 2 | 2C(125.0477) 2C(15.9949) 2C(31.9988) | NaN | X | X | X | NaN | NaN | 1.993762e+05 | NaN | 1.993762e+05 | NaN | DCYVGDEAQSKR |
| TPM3 | TPM3\_HUMAN | Tropomyosin alpha-3 chain | 191 | 191 | TPM3 | TPM3\_HUMAN | Tropomyosin alpha-3 chain | 1C(125.0477) | NaN | X | NaN | NaN | NaN | NaN | 1.990293e+05 | 1.990293e+05 | NaN | NaN | CSELEEELKNVTNNLK |
| HNRNPD | HNRPD\_HUMAN | Heterogeneous nuclear ribonucleoprotein D0 | 226 | 224 | HNRNPD | HNRPD\_HUMAN | Heterogeneous nuclear ribonucleoprotein D0 | 3C(125.0477) | NaN | X | NaN | NaN | NaN | NaN | 1.986760e+05 | 1.986760e+05 | NaN | NaN | GFCFITFK |
| PLG | PLMN\_HUMAN | Plasminogen | 296 | 285 | PLG | PLMN\_HUMAN | Plasminogen | 12C(57.0215) | X | NaN | NaN | NaN | NaN | NaN | 1.984145e+05 | 1.984145e+05 | NaN | NaN | GNVAVTVSGHTCQHWSAQTPHTHNR |
| PRMT5 | ANM5\_HUMAN | Protein arginine N-methyltransferase 5 | 42 | 36 | PRMT5 | ANM5\_HUMAN | Protein arginine N-methyltransferase 5 | 7C(125.0477) | NaN | X | NaN | NaN | NaN | NaN | 1.983863e+05 | 1.983863e+05 | NaN | NaN | QGFDFLCMPVFHPR |
| MTAP | MTAP\_HUMAN | S-methyl-5'-thioadenosine phosphorylase | 131 | 117 | MTAP | MTAP\_HUMAN | S-methyl-5'-thioadenosine phosphorylase | 15C(125.0477) | NaN | X | NaN | NaN | NaN | NaN | 1.983683e+05 | 1.983683e+05 | NaN | NaN | TTMRPQSFYDGSHSCAR |
| LANCL1 | LANC1\_HUMAN | Glutathione S-transferase LANCL1 | 264 | 256 | LANCL1 | LANC1\_HUMAN | Glutathione S-transferase LANCL1 | 9C(125.0477) | NaN | X | NaN | NaN | NaN | NaN | 1.981001e+05 | 1.981001e+05 | NaN | NaN | FPSGNYPPCIGDNR |
| PLPBP | PLPHP\_HUMAN | Pyridoxal phosphate homeostasis protein | 86 | 82 | PLPBP | PLPHP\_HUMAN | Pyridoxal phosphate homeostasis protein | 5C(125.0477) | NaN | X | NaN | NaN | NaN | NaN | 1.978998e+05 | 1.978998e+05 | NaN | NaN | ILSLCPEIK |
| OLA1 | OLA1\_HUMAN | Obg-like ATPase 1 | 55 | 36 | OLA1 | OLA1\_HUMAN | Obg-like ATPase 1 | 20C(125.0477) | NaN | X | NaN | NaN | NaN | NaN | 1.978891e+05 | 1.978891e+05 | NaN | NaN | STFFNVLTNSQASAENFPFCTIDPNESR |
| AARSD1 | AASD1\_HUMAN | Alanyl-tRNA editing protein Aarsd1 | 266 | 266 | AARSD1 | AASD1\_HUMAN | Alanyl-tRNA editing protein Aarsd1 | 1C(125.0477) | NaN | X | NaN | NaN | NaN | NaN | 1.978322e+05 | 1.978322e+05 | NaN | NaN | CGAEDHVEAVKK |
| CAST | ICAL\_HUMAN | Calpastatin | 328 | 328 | CAST | ICAL\_HUMAN | Calpastatin | 1C(125.0477) | NaN | X | NaN | NaN | NaN | NaN | 1.975036e+05 | 8.326228e+04 | 1.142414e+05 | NaN | CGEDDETIPSEYR |
| PAFAH1B3 | PA1B3\_HUMAN | Platelet-activating factor acetylhydrolase IB subunit alpha1 | 205 | 199 | PAFAH1B3 | PA1B3\_HUMAN | Platelet-activating factor acetylhydrolase IB subunit alpha1 | 7C(125.0477) | NaN | X | NaN | NaN | NaN | NaN | 1.973384e+05 | NaN | 1.973384e+05 | NaN | LGYTPVCR |
| TPP2 | TPP2\_HUMAN | Tripeptidyl-peptidase 2 | 197 | 188 | TPP2 | TPP2\_HUMAN | Tripeptidyl-peptidase 2 | 10C(125.0477) | NaN | X | NaN | NaN | NaN | NaN | 1.966742e+05 | 1.966742e+05 | NaN | NaN | YSDPGPVYDCLVWHDGEVWR |
| HDDC3 | MESH1\_HUMAN | Guanosine-3',5'-bis(diphosphate) 3'-pyrophosphohydrolase MESH1 | 133 | 133 | HDDC3 | MESH1\_HUMAN | Guanosine-3',5'-bis(diphosphate) 3'-pyrophosphohydrolase MESH1 | 1C(125.0477) | NaN | X | NaN | NaN | NaN | NaN | 1.963281e+05 | 1.963281e+05 | NaN | NaN | CTPEGWSEHR |
| VAC14 | VAC14\_HUMAN | Protein VAC14 homolog | 101 | 93 | VAC14 | VAC14\_HUMAN | Protein VAC14 homolog | 9C(125.0477) | NaN | X | NaN | NaN | NaN | NaN | 1.963174e+05 | 1.963174e+05 | NaN | NaN | ELIEPVLTCFNDADSR |
| NRBP1 | NRBP\_HUMAN | Nuclear receptor-binding protein | 449 | 444 | NRBP1 | NRBP\_HUMAN | Nuclear receptor-binding protein | 6C(125.0477) | NaN | X | NaN | NaN | NaN | NaN | 1.960948e+05 | 1.960948e+05 | NaN | NaN | VVLMQCNIESVEEGVK |
| PDE12 | PDE12\_HUMAN | 2',5'-phosphodiesterase 12 | 180 | 172 | PDE12 | PDE12\_HUMAN | 2',5'-phosphodiesterase 12 | 9C(125.0477) | NaN | X | NaN | NaN | NaN | NaN | 1.960326e+05 | 1.960326e+05 | NaN | NaN | YIMAGFPVCPK |
| HBA1 | HBA\_HUMAN | Hemoglobin subunit alpha | 105, 105 | 94 | HBA1 | HBA\_HUMAN | Hemoglobin subunit alpha | 12C(-34.0809) 12C(125.0477) | NaN | X | NaN | NaN | X | NaN | 1.958413e+05 | 1.958413e+05 | NaN | NaN | VDPVNFKLLSHCLLVTLAAHLPAEFTPAVHASLDKFLASVSTVLTSK |
| PPIA | PPIA\_HUMAN | Peptidyl-prolyl cis-trans isomerase A | 161 | 156 | PPIA | PPIA\_HUMAN | Peptidyl-prolyl cis-trans isomerase A | 6C(125.0477) | NaN | X | NaN | NaN | NaN | NaN | 1.955293e+05 | 1.570950e+05 | 3.843434e+04 | NaN | ITIADCGQLE |
| RAB18 | RAB18\_HUMAN | Ras-related protein Rab-18 | 110 | 99 | RAB18 | RAB18\_HUMAN | Ras-related protein Rab-18 | 12C(125.0477) | NaN | X | NaN | NaN | NaN | NaN | 1.948789e+05 | 1.948789e+05 | NaN | NaN | LDNWLNELETYCTR |
| ANKFY1 | ANFY1\_HUMAN | Rabankyrin-5 | 499 | 490 | ANKFY1 | ANFY1\_HUMAN | Rabankyrin-5 | 10C(125.0477) | NaN | X | NaN | NaN | NaN | NaN | 1.944266e+05 | 1.944266e+05 | NaN | NaN | WGETPLHTACR |
| ACLY | ACLY\_HUMAN | ATP-citrate synthase | 519 | 508 | ACLY | ACLY\_HUMAN | ATP-citrate synthase | 12C(125.0477) | NaN | X | NaN | NaN | NaN | NaN | 1.943134e+05 | 1.943134e+05 | NaN | NaN | AVQGMLDFDYVCSR |
| ACSL4 | ACSL4\_HUMAN | Long-chain-fatty-acid--CoA ligase 4 | 363 | 361 | ACSL4 | ACSL4\_HUMAN | Long-chain-fatty-acid--CoA ligase 4 | 3C(125.0477) | NaN | X | NaN | NaN | NaN | NaN | 1.942187e+05 | 1.942187e+05 | NaN | NaN | GDCTVLKPTLMAAVPEIMDR |
| PSMG1 | PSMG1\_HUMAN | Proteasome assembly chaperone 1 | 121 | 108 | PSMG1 | PSMG1\_HUMAN | Proteasome assembly chaperone 1 | 14C(125.0477) | NaN | X | NaN | NaN | NaN | NaN | 1.935102e+05 | 1.935102e+05 | NaN | NaN | TTDTTHLSSTEAFCVFYHLK |
| GAPVD1 | GAPD1\_HUMAN | GTPase-activating protein and VPS9 domain-containing protein 1 | 275 | 271 | GAPVD1 | GAPD1\_HUMAN | GTPase-activating protein and VPS9 domain-containing protein 1 | 5C(125.0477) | NaN | X | NaN | NaN | NaN | NaN | 1.931311e+05 | 1.931311e+05 | NaN | NaN | QNTYCFPHSLR |
| RAB3IL1 | R3GEF\_HUMAN | Guanine nucleotide exchange factor for Rab-3A | 97 | 92 | RAB3IL1 | R3GEF\_HUMAN | Guanine nucleotide exchange factor for Rab-3A | 6C(125.0477) | NaN | X | NaN | NaN | NaN | NaN | 1.931251e+05 | 1.931251e+05 | NaN | NaN | LKDEECER |
| GMPR | GMPR1\_HUMAN | GMP reductase 1 | 205, 222, 224 | 192 | GMPR | GMPR1\_HUMAN | GMP reductase 1 | 14C(125.0477) 31C(15.9949) 33C(15.9949) | NaN | X | X | NaN | NaN | NaN | 1.929160e+05 | 1.929160e+05 | NaN | NaN | TGVGYPQLSAVIECADSAHGLKGHIISDGGCTCPGDVAK |
| DIAPH1 | DIAP1\_HUMAN | Protein diaphanous homolog 1 | 437 | 433 | DIAPH1 | DIAP1\_HUMAN | Protein diaphanous homolog 1 | 5C(125.0477) | NaN | X | NaN | NaN | NaN | NaN | 1.925786e+05 | 1.925786e+05 | NaN | NaN | LIEECISQIVLHK |
| CFH | CFAH\_HUMAN | Complement factor H | 494 | 479 | CFH | CFAH\_HUMAN | Complement factor H | 16C(57.0215) | X | NaN | NaN | NaN | NaN | NaN | 1.925748e+05 | 3.368189e+04 | 1.381518e+05 | 20741.1100 | LGYVTADGETSGSITCGK |
| AGO2 | AGO2\_HUMAN | Protein argonaute-2 | 462 | 461 | AGO2 | AGO2\_HUMAN | Protein argonaute-2 | 2C(125.0477) | NaN | X | NaN | NaN | NaN | NaN | 1.922832e+05 | 1.593355e+05 | 3.294773e+04 | NaN | QCTEVHLK |
| ANKRD28 | ANR28\_HUMAN | Serine/threonine-protein phosphatase 6 regulatory ankyrin repeat subunit A | 701 | 693 | ANKRD28 | ANR28\_HUMAN | Serine/threonine-protein phosphatase 6 regulatory ankyrin repeat subunit A | 9C(125.0477) | NaN | X | NaN | NaN | NaN | NaN | 1.922662e+05 | 1.922662e+05 | NaN | NaN | GAVTGHEECVDALLQHGAK |
| INPP5D | SHIP1\_HUMAN | Phosphatidylinositol 3,4,5-trisphosphate 5-phosphatase 1 | 506 | 502 | INPP5D | SHIP1\_HUMAN | Phosphatidylinositol 3,4,5-trisphosphate 5-phosphatase 1 | 5C(125.0477) | NaN | X | NaN | NaN | NaN | NaN | 1.922129e+05 | 1.922129e+05 | NaN | NaN | ISHICTDNVK |
| IDH1 | IDHC\_HUMAN | Isocitrate dehydrogenase [NADP] cytoplasmic | 73 | 73 | IDH1 | IDHC\_HUMAN | Isocitrate dehydrogenase [NADP] cytoplasmic | 1C(125.0477) | NaN | X | NaN | NaN | NaN | NaN | 1.921174e+05 | 1.921174e+05 | NaN | NaN | CATITPDEKR |
| HSPA8 | HSP7C\_HUMAN | Heat shock cognate 71 kDa protein | 603 | 602 | HSPA8 | HSP7C\_HUMAN | Heat shock cognate 71 kDa protein | 2C(125.0477) | NaN | X | NaN | NaN | NaN | NaN | 1.921104e+05 | 6.696784e+04 | NaN | 125142.6000 | VCNPIITK |
| MAPK1 | MK01\_HUMAN | Mitogen-activated protein kinase 1 | 65 | 55 | MAPK1 | MK01\_HUMAN | Mitogen-activated protein kinase 1 | 11C(125.0477) | NaN | X | NaN | NaN | NaN | NaN | 1.920310e+05 | 1.920310e+05 | NaN | NaN | KISPFEHQTYCQR |
| ITPA | ITPA\_HUMAN | Inosine triphosphate pyrophosphatase | 33 | 19 | ITPA | ITPA\_HUMAN | Inosine triphosphate pyrophosphatase | 15C(125.0477) | NaN | X | NaN | NaN | NaN | NaN | 1.920023e+05 | 1.920023e+05 | NaN | NaN | KLEEVVQILGDKFPCTLVAQK |
| PSMD1 | PSMD1\_HUMAN | 26S proteasome non-ATPase regulatory subunit 1 | 104, 104, 112 | 104 | PSMD1 | PSMD1\_HUMAN | 26S proteasome non-ATPase regulatory subunit 1 | 1C(15.9949) 1C(57.0215) 9C(15.9949) | X | NaN | X | NaN | NaN | NaN | 1.919847e+05 | 1.919847e+05 | NaN | NaN | CIDHYTKQCVENADLPEGEK |
| SPTB | SPTB1\_HUMAN | Spectrin beta chain, erythrocytic | 735 | 730 | SPTB | SPTB1\_HUMAN | Spectrin beta chain, erythrocytic | 6C(125.0477) | NaN | X | NaN | NaN | NaN | NaN | 1.916607e+05 | NaN | 1.916607e+05 | NaN | DLAAFCKK |
| CAPNS1 | CPNS1\_HUMAN | Calpain small subunit 1 | 144 | 131 | CAPNS1 | CPNS1\_HUMAN | Calpain small subunit 1 | 14C(125.0477) | NaN | X | NaN | NaN | NaN | NaN | 1.911805e+05 | 1.911805e+05 | NaN | NaN | HPDLKTDGFGIDTCR |
| CLTC | CLH1\_HUMAN | Clathrin heavy chain 1 | 436 | 430 | CLTC | CLH1\_HUMAN | Clathrin heavy chain 1 | 7C(125.0477) | NaN | X | NaN | NaN | NaN | NaN | 1.911742e+05 | 1.911742e+05 | NaN | NaN | YESLELCRPVLQQGR |
| SCFD1 | SCFD1\_HUMAN | Sec1 family domain-containing protein 1 | 485 | 479 | SCFD1 | SCFD1\_HUMAN | Sec1 family domain-containing protein 1 | 7C(125.0477) | NaN | X | NaN | NaN | NaN | NaN | 1.905547e+05 | 1.698272e+05 | 2.072749e+04 | NaN | ALTDAGCNLNPLQYIK |
| DCTN1 | DCTN1\_HUMAN | Dynactin subunit 1 | 677 | 670 | DCTN1 | DCTN1\_HUMAN | Dynactin subunit 1 | 8C(125.0477) | NaN | X | NaN | NaN | NaN | NaN | 1.903028e+05 | 1.903028e+05 | NaN | NaN | YEHALSQCSVDVYK |
| HSPA1A | HS71A\_HUMAN | Heat shock 70 kDa protein 1A | 574 | 574 | HSPA1A | HS71A\_HUMAN | Heat shock 70 kDa protein 1A | 1C(125.0477) | NaN | X | NaN | NaN | NaN | NaN | 1.902319e+05 | 1.902319e+05 | NaN | NaN | CQEVISWLDANTLAEKDEFEHK |
| ASPSCR1 | ASPC1\_HUMAN | Tether containing UBX domain for GLUT4 | 40 | 26 | ASPSCR1 | ASPC1\_HUMAN | Tether containing UBX domain for GLUT4 | 15C(125.0477) | NaN | X | NaN | NaN | NaN | NaN | 1.901267e+05 | 1.901267e+05 | NaN | NaN | VTPSTVLLQVLEDTCR |
| FCSK | FCSK\_HUMAN | L-fucose kinase | 536 | 528 | FCSK | FCSK\_HUMAN | L-fucose kinase | 9C(125.0477) | NaN | X | NaN | NaN | NaN | NaN | 1.898148e+05 | 1.898148e+05 | NaN | NaN | LSWEQLQPCLDR |
| PSMC2 | PRS7\_HUMAN | 26S proteasome regulatory subunit 7 | 180 | 157 | PSMC2 | PRS7\_HUMAN | 26S proteasome regulatory subunit 7 | 24C(125.0477) | NaN | X | NaN | NaN | NaN | NaN | 1.898132e+05 | 1.898132e+05 | NaN | NaN | IDPTVTMMQVEEKPDVTYSDVGGCK |
| UBE2O | UBE2O\_HUMAN | (E3-independent) E2 ubiquitin-conjugating enzyme | 375 | 374 | UBE2O | UBE2O\_HUMAN | (E3-independent) E2 ubiquitin-conjugating enzyme | 2C(125.0477) | NaN | X | NaN | NaN | NaN | NaN | 1.895612e+05 | 1.895612e+05 | NaN | NaN | NCAQGEGSMAK |
| LAP3 | AMPL\_HUMAN | Cytosol aminopeptidase | 335 | 322 | LAP3 | AMPL\_HUMAN | Cytosol aminopeptidase | 14C(125.0477) | NaN | X | NaN | NaN | NaN | NaN | 1.893567e+05 | 1.893567e+05 | NaN | NaN | LNLPINIIGLAPLCENMPSGK |
| FAH | FAAA\_HUMAN | Fumarylacetoacetase | 408 | 403 | FAH | FAAA\_HUMAN | Fumarylacetoacetase | 6C(125.0477) | NaN | X | NaN | NaN | NaN | NaN | 1.891395e+05 | 1.891395e+05 | NaN | NaN | IGFGQCAGK |
| FAHD2A | FAH2A\_HUMAN | Fumarylacetoacetate hydrolase domain-containing protein 2A | 215 | 210 | FAHD2A | FAH2A\_HUMAN | Fumarylacetoacetate hydrolase domain-containing protein 2A | 6C(125.0477) | NaN | X | NaN | NaN | NaN | NaN | 1.890801e+05 | 1.890801e+05 | NaN | NaN | TFDTFCPLGPALVTK |
| CRLF3 | CRLF3\_HUMAN | Cytokine receptor-like factor 3 | 336 | 331 | CRLF3 | CRLF3\_HUMAN | Cytokine receptor-like factor 3 | 6C(125.0477) | NaN | X | NaN | NaN | NaN | NaN | 1.887301e+05 | 1.887301e+05 | NaN | NaN | DSIGVCAEKQDGYDSLQR |
| FAS | TNR6\_HUMAN | Tumor necrosis factor receptor superfamily member 6 | 149 | 149 | FAS | TNR6\_HUMAN | Tumor necrosis factor receptor superfamily member 6 | 1C(125.0477) | NaN | X | NaN | NaN | NaN | NaN | 1.884763e+05 | NaN | 7.970861e+04 | 108767.6900 | CEHGIIK |
| RNPEP | AMPB\_HUMAN | Aminopeptidase B | 69, 69 | 69 | RNPEP | AMPB\_HUMAN | Aminopeptidase B | 1C(125.0477) 1C(31.9988) | NaN | X | NaN | X | NaN | NaN | 1.884158e+05 | NaN | 1.884158e+05 | NaN | CLEPEGAAELR |
| FKBP4 | FKBP4\_HUMAN | Peptidyl-prolyl cis-trans isomerase FKBP4 | 342 | 332 | FKBP4 | FKBP4\_HUMAN | Peptidyl-prolyl cis-trans isomerase FKBP4 | 11C(125.0477) | NaN | X | NaN | NaN | NaN | NaN | 1.883557e+05 | 1.883557e+05 | NaN | NaN | LQAFSAAIESCNK |
| ACLY | ACLY\_HUMAN | ATP-citrate synthase | 845 | 836 | ACLY | ACLY\_HUMAN | ATP-citrate synthase | 10C(125.0477) | NaN | X | NaN | NaN | NaN | NaN | 1.882973e+05 | 1.882973e+05 | NaN | NaN | KPASFMTSICDER |
| IRAK4 | IRAK4\_HUMAN | Interleukin-1 receptor-associated kinase 4 | 13 | 13 | IRAK4 | IRAK4\_HUMAN | Interleukin-1 receptor-associated kinase 4 | 1C(125.0477) | NaN | X | NaN | NaN | NaN | NaN | 1.881419e+05 | 1.881419e+05 | NaN | NaN | CLNVGLIR |
| EIF5 | IF5\_HUMAN | Eukaryotic translation initiation factor 5 | 138 | 137 | EIF5 | IF5\_HUMAN | Eukaryotic translation initiation factor 5 | 2C(125.0477) | NaN | X | NaN | NaN | NaN | NaN | 1.870290e+05 | 1.870290e+05 | NaN | NaN | LCTFILK |
| APEH | ACPH\_HUMAN | Acylamino-acid-releasing enzyme | 152, 149 | 141 | APEH | ACPH\_HUMAN | Acylamino-acid-releasing enzyme | 12C(31.9988) 9C(-34.0809) | NaN | NaN | NaN | X | X | NaN | 1.868558e+05 | 1.868558e+05 | NaN | NaN | HGPVYEDDCFGCLSWSHSETHLLYVAEK |
| MAN2C1 | MA2C1\_HUMAN | Alpha-mannosidase 2C1 | 322 | 317 | MAN2C1 | MA2C1\_HUMAN | Alpha-mannosidase 2C1 | 6C(125.0477) | NaN | X | NaN | NaN | NaN | NaN | 1.864641e+05 | 1.864641e+05 | NaN | NaN | IQEFACR |
| SORD | DHSO\_HUMAN | Sorbitol dehydrogenase | 179 | 176 | SORD | DHSO\_HUMAN | Sorbitol dehydrogenase | 4C(125.0477) | NaN | X | NaN | NaN | NaN | NaN | 1.863628e+05 | 1.863628e+05 | NaN | NaN | VLVCGAGPIGMVTLLVAK |
| CHMP5 | CHMP5\_HUMAN | Charged multivesicular body protein 5 | 20 | 12 | CHMP5 | CHMP5\_HUMAN | Charged multivesicular body protein 5 | 9C(125.0477) | NaN | X | NaN | NaN | NaN | NaN | 1.862227e+05 | 1.862227e+05 | NaN | NaN | APPPSLTDCIGTVDSR |
| FN3K | FN3K\_HUMAN | Fructosamine-3-kinase | 24 | 16 | FN3K | FN3K\_HUMAN | Fructosamine-3-kinase | 9C(125.0477) | NaN | X | NaN | NaN | NaN | NaN | 1.862052e+05 | 1.862052e+05 | NaN | NaN | AFGGPGAGCISEGR |
| ARG1 | ARGI1\_HUMAN | Arginase-1 | 168 | 156 | ARG1 | ARGI1\_HUMAN | Arginase-1 | 13C(125.0477) | NaN | X | NaN | NaN | NaN | NaN | 1.861978e+05 | 1.861978e+05 | NaN | NaN | IPDVPGFSWVTPCISAK |
| SHC1 | SHC1\_HUMAN | SHC-transforming protein 1 | 196 | 190 | SHC1 | SHC1\_HUMAN | SHC-transforming protein 1 | 7C(125.0477) | NaN | X | NaN | NaN | NaN | NaN | 1.860966e+05 | NaN | 1.860966e+05 | NaN | EAISLVCEAVPGAK |
| FCSK | FCSK\_HUMAN | L-fucose kinase | 582 | 580 | FCSK | FCSK\_HUMAN | L-fucose kinase | 3C(125.0477) | NaN | X | NaN | NaN | NaN | NaN | 1.859593e+05 | 1.859593e+05 | NaN | NaN | EGCPGPLLATLDQVAAGAGDPGVAAR |
| BLVRA | BIEA\_HUMAN | Biliverdin reductase A | 281 | 277 | BLVRA | BIEA\_HUMAN | Biliverdin reductase A | 5C(125.0477) | NaN | X | NaN | NaN | NaN | NaN | 1.857776e+05 | 1.857776e+05 | NaN | NaN | RILHCLGLAEEIQK |
| AP1G1 | AP1G1\_HUMAN | AP-1 complex subunit gamma-1 | 353 | 348 | AP1G1 | AP1G1\_HUMAN | AP-1 complex subunit gamma-1 | 6C(125.0477) | NaN | X | NaN | NaN | NaN | NaN | 1.857658e+05 | NaN | 1.857658e+05 | NaN | STIVDCLK |
| PSMA7 | PSA7\_HUMAN | Proteasome subunit alpha type-7 | 91 | 87 | PSMA7 | PSA7\_HUMAN | Proteasome subunit alpha type-7 | 5C(125.0477) | NaN | X | NaN | NaN | NaN | NaN | 1.857191e+05 | 1.857191e+05 | NaN | NaN | ARVECQSHR |
| MPP1 | EM55\_HUMAN | 55 kDa erythrocyte membrane protein | 179 | 165 | MPP1 | EM55\_HUMAN | 55 kDa erythrocyte membrane protein | 15C(125.0477) | NaN | X | NaN | NaN | NaN | NaN | 1.854566e+05 | 1.854566e+05 | NaN | NaN | AQFDYDPKKDNLIPCK |
| NIF3L1 | NIF3L\_HUMAN | NIF3-like protein 1 | 148 | 144 | NIF3L1 | NIF3L\_HUMAN | NIF3-like protein 1 | 5C(125.0477) | NaN | X | NaN | NaN | NaN | NaN | 1.852425e+05 | 1.852425e+05 | NaN | NaN | GLGACTSRPIHPSK |
| HUWE1 | HUWE1\_HUMAN | E3 ubiquitin-protein ligase HUWE1 | 3658 | 3656 | HUWE1 | HUWE1\_HUMAN | E3 ubiquitin-protein ligase HUWE1 | 3C(125.0477) | NaN | X | NaN | NaN | NaN | NaN | 1.844385e+05 | 1.844385e+05 | NaN | NaN | AQCETLSPDGLPEEQPQTTK |
| FASN | FAS\_HUMAN | Fatty acid synthase | 2202 | 2194 | FASN | FAS\_HUMAN | Fatty acid synthase | 9C(125.0477) | NaN | X | NaN | NaN | NaN | NaN | 1.843919e+05 | 4.144585e+04 | 1.429460e+05 | NaN | ADEASELACPTPK |
| DIS3L2 | DI3L2\_HUMAN | DIS3-like exonuclease 2 | 396 | 389 | DIS3L2 | DI3L2\_HUMAN | DIS3-like exonuclease 2 | 8C(125.0477) | NaN | X | NaN | NaN | NaN | NaN | 1.843235e+05 | 1.843235e+05 | NaN | NaN | DLDDALSCKPLADGNFK |
| MPST | THTM\_HUMAN | 3-mercaptopyruvate sulfurtransferase | 65 | 53 | MPST | THTM\_HUMAN | 3-mercaptopyruvate sulfurtransferase | 13C(125.0477) | NaN | X | NaN | NaN | NaN | NaN | 1.840499e+05 | 1.840499e+05 | NaN | NaN | HIPGAAFFDIDQCSDR |
| WDR77 | MEP50\_HUMAN | Methylosome protein 50 | 172 | 165 | WDR77 | MEP50\_HUMAN | Methylosome protein 50 | 8C(125.0477) | NaN | X | NaN | NaN | NaN | NaN | 1.838895e+05 | 1.838895e+05 | NaN | NaN | AHAAQVTCVAASPHK |
| CAPZB | CAPZB\_HUMAN | F-actin-capping protein subunit beta | 206, 206 | 196 | CAPZB | CAPZB\_HUMAN | F-actin-capping protein subunit beta | 11C(125.0477) 11C(15.9949) | NaN | X | X | NaN | NaN | NaN | 1.834542e+05 | 1.834542e+05 | NaN | NaN | QMEKDETVSDCSPHIANIGR |
| ILF3 | ILF3\_HUMAN | Interleukin enhancer-binding factor 3 | 203 | 203 | ILF3 | ILF3\_HUMAN | Interleukin enhancer-binding factor 3 | 1C(125.0477) | NaN | X | NaN | NaN | NaN | NaN | 1.833872e+05 | 1.833872e+05 | NaN | NaN | CLAALASLR |
| MESD | MESD\_HUMAN | LRP chaperone MESD | 180 | 180 | MESD | MESD\_HUMAN | LRP chaperone MESD | 1C(125.0477) | NaN | X | NaN | NaN | NaN | NaN | 1.833828e+05 | NaN | 1.833828e+05 | NaN | CADVTLEGQVYPGK |
| TUBA4A | TBA4A\_HUMAN | Tubulin alpha-4A chain | 347, 347 | 340 | TUBA4A | TBA4A\_HUMAN | Tubulin alpha-4A chain | 8C(125.0477) 8C(57.0215) | X | X | NaN | NaN | NaN | NaN | 1.830600e+05 | 1.830600e+05 | NaN | NaN | SIQFVDWCPTGFK |
| RNH1 | RINI\_HUMAN | Ribonuclease inhibitor | 191 | 189 | RNH1 | RINI\_HUMAN | Ribonuclease inhibitor | 3C(15.9949) | NaN | NaN | X | NaN | NaN | NaN | 1.826106e+05 | 1.826106e+05 | NaN | NaN | VLCQGLKDSPCQLEALK |
| RAC1 | RAC1\_HUMAN | Ras-related C3 botulinum toxin substrate 1 | 105 | 103 | RAC1 | RAC1\_HUMAN | Ras-related C3 botulinum toxin substrate 1 | 3C(125.0477) | NaN | X | NaN | NaN | NaN | NaN | 1.825715e+05 | 1.825715e+05 | NaN | NaN | HHCPNTPIILVGTK |
| TPI1 | TPIS\_HUMAN | Triosephosphate isomerase | 127 | 114 | TPI1 | TPIS\_HUMAN | Triosephosphate isomerase | 14C(125.0477) | NaN | X | NaN | NaN | NaN | NaN | 1.824302e+05 | 1.824302e+05 | NaN | NaN | VAHALAEGLGVIACIGEKLDER |
| ATP1A1 | AT1A1\_HUMAN | Sodium/potassium-transporting ATPase subunit alpha-1 | 518 | 518 | ATP1A1 | AT1A1\_HUMAN | Sodium/potassium-transporting ATPase subunit alpha-1 | 1C(125.0477) | NaN | X | NaN | NaN | NaN | NaN | 1.823831e+05 | 3.523899e+04 | 1.322506e+05 | 14893.5110 | CSSILLHGK |
| FCHO2 | FCHO2\_HUMAN | F-BAR domain only protein 2 | 595 | 584 | FCHO2 | FCHO2\_HUMAN | F-BAR domain only protein 2 | 12C(125.0477) | NaN | X | NaN | NaN | NaN | NaN | 1.823070e+05 | 1.823070e+05 | NaN | NaN | VFTSNPTPAVLCFR |
| DARS1 | SYDC\_HUMAN | Aspartate--tRNA ligase, cytoplasmic | 76 | 75 | DARS1 | SYDC\_HUMAN | Aspartate--tRNA ligase, cytoplasmic | 2C(125.0477) | NaN | X | NaN | NaN | NaN | NaN | 1.822409e+05 | 1.822409e+05 | NaN | NaN | QCFLVLR |
| PPP6R3 | PP6R3\_HUMAN | Serine/threonine-protein phosphatase 6 regulatory subunit 3 | 172 | 169 | PPP6R3 | PP6R3\_HUMAN | Serine/threonine-protein phosphatase 6 regulatory subunit 3 | 4C(125.0477) | NaN | X | NaN | NaN | NaN | NaN | 1.820509e+05 | 1.820509e+05 | NaN | NaN | LLTCIEPPQPR |
| UBA1 | UBA1\_HUMAN | Ubiquitin-like modifier-activating enzyme 1 | 278 | 269 | UBA1 | UBA1\_HUMAN | Ubiquitin-like modifier-activating enzyme 1 | 10C(125.0477) | NaN | X | NaN | NaN | NaN | NaN | 1.820303e+05 | 1.820303e+05 | NaN | NaN | VLGPYTFSICDTSNFSDYIR |
| CHORDC1 | CHRD1\_HUMAN | Cysteine and histidine-rich domain-containing protein 1 | 24 | 14 | CHORDC1 | CHRD1\_HUMAN | Cysteine and histidine-rich domain-containing protein 1 | 11C(125.0477) | NaN | X | NaN | NaN | NaN | NaN | 1.820133e+05 | 1.820133e+05 | NaN | NaN | FDPETNSDDACTYHPGVPVFHDALK |
| TBC1D13 | TBC13\_HUMAN | TBC1 domain family member 13 | 131 | 130 | TBC1D13 | TBC13\_HUMAN | TBC1 domain family member 13 | 2C(125.0477) | NaN | X | NaN | NaN | NaN | NaN | 1.807347e+05 | 1.807347e+05 | NaN | NaN | LCPDISFFQR |
| ALDOA | ALDOA\_HUMAN | Fructose-bisphosphate aldolase A | 73 | 61 | ALDOA | ALDOA\_HUMAN | Fructose-bisphosphate aldolase A | 13C(125.0477) | NaN | X | NaN | NaN | NaN | NaN | 1.806440e+05 | 1.806440e+05 | NaN | NaN | QLLLTADDRVNPCIGGVILFHETLYQK |
| UMPS | UMPS\_HUMAN | Uridine 5'-monophosphate synthase | 304 | 304 | UMPS | UMPS\_HUMAN | Uridine 5'-monophosphate synthase | 1C(125.0477) | NaN | X | NaN | NaN | NaN | NaN | 1.802916e+05 | 1.802916e+05 | NaN | NaN | CHEFLIFEDR |
| OXSR1 | OXSR1\_HUMAN | Serine/threonine-protein kinase | 36 | 15 | OXSR1 | OXSR1\_HUMAN | Serine/threonine-protein kinase | 22C(125.0477) | NaN | X | NaN | NaN | NaN | NaN | 1.801727e+05 | 1.801727e+05 | NaN | NaN | DDYELQEVIGSGATAVVQAAYCAPK |
| IPO7 | IPO7\_HUMAN | Importin-7 | 722 | 714 | IPO7 | IPO7\_HUMAN | Importin-7 | 9C(125.0477) | NaN | X | NaN | NaN | NaN | NaN | 1.801690e+05 | 1.801690e+05 | NaN | NaN | YLEMIYSMCK |
| MYH9 | MYH9\_HUMAN | Myosin-9 | 917 | 910 | MYH9 | MYH9\_HUMAN | Myosin-9 | 8C(125.0477) | NaN | X | NaN | NaN | NaN | NaN | 1.801610e+05 | 1.801610e+05 | NaN | NaN | KQELEEICHDLEAR |
| MAT2A | METK2\_HUMAN | S-adenosylmethionine synthase isoform type-2 | 34 | 33 | MAT2A | METK2\_HUMAN | S-adenosylmethionine synthase isoform type-2 | 2C(125.0477) | NaN | X | NaN | NaN | NaN | NaN | 1.798492e+05 | 1.798492e+05 | NaN | NaN | ICDQISDAVLDAHLQQDPDAK |
| ERMAP | ERMAP\_HUMAN | Erythroid membrane-associated protein | 126 | 126 | ERMAP | ERMAP\_HUMAN | Erythroid membrane-associated protein | 1C(125.0477) | NaN | X | NaN | NaN | NaN | NaN | 1.796059e+05 | NaN | 1.729925e+05 | 6613.3496 | CLIQVGNLSK |
| PRKAA1 | AAPK1\_HUMAN | 5'-AMP-activated protein kinase catalytic subunit alpha-1 | 141 | 130 | PRKAA1 | AAPK1\_HUMAN | 5'-AMP-activated protein kinase catalytic subunit alpha-1 | 12C(125.0477) | NaN | X | NaN | NaN | NaN | NaN | 1.795975e+05 | 1.795975e+05 | NaN | NaN | LFQQILSGVDYCHR |
| NAPRT | PNCB\_HUMAN | Nicotinate phosphoribosyltransferase | 48 | 48 | NAPRT | PNCB\_HUMAN | Nicotinate phosphoribosyltransferase | 1C(125.0477) | NaN | X | NaN | NaN | NaN | NaN | 1.795807e+05 | 1.795807e+05 | NaN | NaN | CPFGGAFALAAGLR |
| HK1 | HXK1\_HUMAN | Hexokinase-1 | 813 | 802 | HK1 | HXK1\_HUMAN | Hexokinase-1 | 12C(125.0477) | NaN | X | NaN | NaN | NaN | NaN | 1.790935e+05 | 1.790935e+05 | NaN | NaN | AILQQLGLNSTCDDSILVK |
| FKBP4 | FKBP4\_HUMAN | Peptidyl-prolyl cis-trans isomerase FKBP4 | 328 | 319 | FKBP4 | FKBP4\_HUMAN | Peptidyl-prolyl cis-trans isomerase FKBP4 | 10C(125.0477) | NaN | X | NaN | NaN | NaN | NaN | 1.785448e+05 | 1.785448e+05 | NaN | NaN | LASHLNLAMCHLK |
| NAA15 | NAA15\_HUMAN | N-alpha-acetyltransferase 15, NatA auxiliary subunit | 322 | 321 | NAA15 | NAA15\_HUMAN | N-alpha-acetyltransferase 15, NatA auxiliary subunit | 2C(125.0477) | NaN | X | NaN | NaN | NaN | NaN | 1.780234e+05 | 1.780234e+05 | NaN | NaN | GCPPVFNTLR |
| CASP6 | CASP6\_HUMAN | Caspase-6 | 87 | 87 | CASP6 | CASP6\_HUMAN | Caspase-6 | 1C(125.0477) | NaN | X | NaN | NaN | NaN | NaN | 1.779607e+05 | 1.779607e+05 | NaN | NaN | CFNDLKAEELLLK |
| PCBP1 | PCBP1\_HUMAN | Poly(rC)-binding protein 1 | 109 | 102 | PCBP1 | PCBP1\_HUMAN | Poly(rC)-binding protein 1 | 8C(15.9949) | NaN | NaN | X | NaN | NaN | NaN | 1.779348e+05 | 1.779348e+05 | NaN | NaN | LVVPATQCGSLIGKGGCK |
| DNAJA4 | DNJA4\_HUMAN | DnaJ homolog subfamily A member 4 | 272 | 265 | DNAJA4 | DNJA4\_HUMAN | DnaJ homolog subfamily A member 4 | 8C(125.0477) | NaN | X | NaN | NaN | NaN | NaN | 1.778657e+05 | 1.778657e+05 | NaN | NaN | IQLSEALCGFKK |
| ALB | ALBU\_HUMAN | Albumin | 148 | 139 | ALB | ALBU\_HUMAN | Albumin | 10C(57.0215) | X | NaN | NaN | NaN | NaN | NaN | 1.774218e+05 | 1.774218e+05 | NaN | NaN | LVRPEVDVMCTAFHDNEETFLKK |
| HSPH1 | HS105\_HUMAN | Heat shock protein 105 kDa | 310 | 304 | HSPH1 | HS105\_HUMAN | Heat shock protein 105 kDa | 7C(125.0477) | NaN | X | NaN | NaN | NaN | NaN | 1.773866e+05 | 1.773866e+05 | NaN | NaN | SQFEELCAELLQK |
| STXBP3 | STXB3\_HUMAN | Syntaxin-binding protein 3 | 546 | 546 | STXBP3 | STXB3\_HUMAN | Syntaxin-binding protein 3 | 1C(125.0477) | NaN | X | NaN | NaN | NaN | NaN | 1.771265e+05 | 1.771265e+05 | NaN | NaN | CAYEVSQAHK |
| CUL3 | CUL3\_HUMAN | Cullin-3 | 636 | 630 | CUL3 | CUL3\_HUMAN | Cullin-3 | 7C(125.0477) | NaN | X | NaN | NaN | NaN | NaN | 1.771041e+05 | 1.771041e+05 | NaN | NaN | ALQSLACGKPTQR |
| ECPAS | ECM29\_HUMAN | Proteasome adapter and scaffold protein ECM29 | 1459 | 1457 | ECPAS | ECM29\_HUMAN | Proteasome adapter and scaffold protein ECM29 | 3C(125.0477) | NaN | X | NaN | NaN | NaN | NaN | 1.766588e+05 | 1.766588e+05 | NaN | NaN | TSCALTIHAIGR |
| ACAT2 | THIC\_HUMAN | Acetyl-CoA acetyltransferase, cytosolic | 360 | 334 | ACAT2 | THIC\_HUMAN | Acetyl-CoA acetyltransferase, cytosolic | 27C(125.0477) | NaN | X | NaN | NaN | NaN | NaN | 1.766560e+05 | 1.766560e+05 | NaN | NaN | ELGLNPEKVNIEGGAIALGHPLGASGCR |
| HBA1 | HBA\_HUMAN | Hemoglobin subunit alpha | 105 | 94 | HBA1 | HBA\_HUMAN | Hemoglobin subunit alpha | 12C(125.0477) | NaN | X | NaN | NaN | NaN | NaN | 1.762922e+05 | NaN | 1.303911e+05 | 45901.0820 | VDPVNFKLLSHCLLVTLAAHLPAEFTPAVHASLDK |
| CTPS1 | PYRG1\_HUMAN | CTP synthase 1 | 30 | 29 | CTPS1 | PYRG1\_HUMAN | CTP synthase 1 | 2C(125.0477) | NaN | X | NaN | NaN | NaN | NaN | 1.761963e+05 | 1.761963e+05 | NaN | NaN | SCGLHVTSIK |
| HBD | HBD\_HUMAN | Hemoglobin subunit delta | 94 | 68 | HBD | HBD\_HUMAN | Hemoglobin subunit delta | 27C(125.0477) | NaN | X | NaN | NaN | NaN | NaN | 1.760218e+05 | 1.760218e+05 | NaN | NaN | VLGAFSDGLAHLDNLKGTFSQLSELHCDK |
| BLMH | BLMH\_HUMAN | Bleomycin hydrolase | 164 | 163 | BLMH | BLMH\_HUMAN | Bleomycin hydrolase | 2C(125.0477) | NaN | X | NaN | NaN | NaN | NaN | 1.758460e+05 | 1.758460e+05 | NaN | NaN | KCFPESYTTEATRR |
| ANKFY1 | ANFY1\_HUMAN | Rabankyrin-5 | 742 | 732 | ANKFY1 | ANFY1\_HUMAN | Rabankyrin-5 | 11C(125.0477) | NaN | X | NaN | NaN | NaN | NaN | 1.753106e+05 | 1.753106e+05 | NaN | NaN | AIDENNEPTACFLIR |
| BROX | BROX\_HUMAN | BRO1 domain-containing protein BROX | 288 | 284 | BROX | BROX\_HUMAN | BRO1 domain-containing protein BROX | 5C(31.9988) | NaN | NaN | NaN | X | NaN | NaN | 1.748366e+05 | 1.748366e+05 | NaN | NaN | AEALCKEYGETK |
| MMS19 | MMS19\_HUMAN | MMS19 nucleotide excision repair protein homolog | 760 | 760 | MMS19 | MMS19\_HUMAN | MMS19 nucleotide excision repair protein homolog | 1C(125.0477) | NaN | X | NaN | NaN | NaN | NaN | 1.747571e+05 | 1.747571e+05 | NaN | NaN | CFAGLLNK |
| TBC1D24 | TBC24\_HUMAN | TBC1 domain family member 24 | 161 | 158 | TBC1D24 | TBC24\_HUMAN | TBC1 domain family member 24 | 4C(125.0477) | NaN | X | NaN | NaN | NaN | NaN | 1.746253e+05 | 1.746253e+05 | NaN | NaN | ILACNDPGRR |
| CCDC9 | CCDC9\_HUMAN | Coiled-coil domain-containing protein 9 | 233 | 233 | CCDC9 | CCDC9\_HUMAN | Coiled-coil domain-containing protein 9 | 1C(125.0477) | NaN | X | NaN | NaN | NaN | NaN | 1.741888e+05 | 1.741888e+05 | NaN | NaN | CGLEHER |
| SSB | LA\_HUMAN | Lupus La protein | 245 | 236 | SSB | LA\_HUMAN | Lupus La protein | 10C(125.0477) | NaN | X | NaN | NaN | NaN | NaN | 1.739724e+05 | 1.739724e+05 | NaN | NaN | FSGDLDDQTCR |
| USP9X | USP9X\_HUMAN | Probable ubiquitin carboxyl-terminal hydrolase FAF-X | 147 | 147 | USP9X | USP9X\_HUMAN | Probable ubiquitin carboxyl-terminal hydrolase FAF-X | 1C(125.0477) | NaN | X | NaN | NaN | NaN | NaN | 1.735885e+05 | 1.735885e+05 | NaN | NaN | CIINNTHR |
| PCBP1 | PCBP1\_HUMAN | Poly(rC)-binding protein 1 | 163 | 161 | PCBP1 | PCBP1\_HUMAN | Poly(rC)-binding protein 1 | 3C(125.0477) | NaN | X | NaN | NaN | NaN | NaN | 1.735262e+05 | 1.735262e+05 | NaN | NaN | QICLVMLETLSQSPQGR |
| TXNRD1 | TRXR1\_HUMAN | Thioredoxin reductase 1, cytoplasmic | 577 | 575 | TXNRD1 | TRXR1\_HUMAN | Thioredoxin reductase 1, cytoplasmic | 3C(125.0477) | NaN | X | NaN | NaN | NaN | NaN | 1.734882e+05 | 1.734882e+05 | NaN | NaN | IICNTKDNER |
| HNRNPK | HNRPK\_HUMAN | Heterogeneous nuclear ribonucleoprotein K | 132, 132 | 104 | HNRNPK | HNRPK\_HUMAN | Heterogeneous nuclear ribonucleoprotein K | 29C(125.0477) 29C(57.0215) | X | X | NaN | NaN | NaN | NaN | 1.730268e+05 | 1.730268e+05 | NaN | NaN | IIPTLEEGLQLPSPTATSQLPLESDAVECLNYQHYK |
| PGM3 | AGM1\_HUMAN | Phosphoacetylglucosamine mutase | 200 | 197 | PGM3 | AGM1\_HUMAN | Phosphoacetylglucosamine mutase | 4C(125.0477) | NaN | X | NaN | NaN | NaN | NaN | 1.726200e+05 | 1.726200e+05 | NaN | NaN | QASCSGDEYR |
| FKBP1A | FKB1A\_HUMAN | Peptidyl-prolyl cis-trans isomerase FKBP1A | 23 | 19 | FKBP1A | FKB1A\_HUMAN | Peptidyl-prolyl cis-trans isomerase FKBP1A | 5C(125.0477) | NaN | X | NaN | NaN | NaN | NaN | 1.722954e+05 | 1.722954e+05 | NaN | NaN | RGQTCVVHYTGMLEDGK |
| SPTA1 | SPTA1\_HUMAN | Spectrin alpha chain, erythrocytic 1 | 2059, 2059 | 2052 | SPTA1 | SPTA1\_HUMAN | Spectrin alpha chain, erythrocytic 1 | 8C(125.0477) 8C(31.9988) | NaN | X | NaN | X | NaN | NaN | 1.722823e+05 | 1.722823e+05 | NaN | NaN | ASALNNWCEK |
| MYO1G | MYO1G\_HUMAN | Unconventional myosin-Ig | 97 | 93 | MYO1G | MYO1G\_HUMAN | Unconventional myosin-Ig | 5C(15.9949) | NaN | NaN | X | NaN | NaN | NaN | 1.721194e+05 | 1.721194e+05 | NaN | NaN | SRDTCIVISGESGAGK |
| CAPN1 | CAN1\_HUMAN | Calpain-1 catalytic subunit | 590 | 584 | CAPN1 | CAN1\_HUMAN | Calpain-1 catalytic subunit | 7C(125.0477) | NaN | X | NaN | NaN | NaN | NaN | 1.720163e+05 | NaN | 1.720163e+05 | NaN | GFSLESCR |
| UBA1 | UBA1\_HUMAN | Ubiquitin-like modifier-activating enzyme 1 | 234 | 227 | UBA1 | UBA1\_HUMAN | Ubiquitin-like modifier-activating enzyme 1 | 8C(125.0477) | NaN | X | NaN | NaN | NaN | NaN | 1.719543e+05 | 7.314468e+04 | 9.880965e+04 | NaN | DNPGVVTCLDEAR |
| PAICS | PUR6\_HUMAN | Multifunctional protein ADE2 | 185 | 179 | PAICS | PUR6\_HUMAN | Multifunctional protein ADE2 | 7C(125.0477) | NaN | X | NaN | NaN | NaN | NaN | 1.716422e+05 | 1.716422e+05 | NaN | NaN | SWLPQNCTLVDMK |
| GDI1 | GDIA\_HUMAN | Rab GDP dissociation inhibitor alpha | 335 | 329 | GDI1 | GDIA\_HUMAN | Rab GDP dissociation inhibitor alpha | 7C(125.0477) | NaN | X | NaN | NaN | NaN | NaN | 1.716243e+05 | 1.716243e+05 | NaN | NaN | KSDIYVCMISYAHNVAAQGK |
| GLOD4 | GLOD4\_HUMAN | Glyoxalase domain-containing protein 4 | 197 | 188 | GLOD4 | GLOD4\_HUMAN | Glyoxalase domain-containing protein 4 | 10C(125.0477) | NaN | X | NaN | NaN | NaN | NaN | 1.712960e+05 | 1.518815e+05 | 1.941455e+04 | NaN | ALLGYADNQCK |
| NSF | NSF\_HUMAN | Vesicle-fusing ATPase | 264 | 255 | NSF | NSF\_HUMAN | Vesicle-fusing ATPase | 10C(125.0477) | NaN | X | NaN | NaN | NaN | NaN | 1.704320e+05 | 1.704320e+05 | NaN | NaN | GILLYGPPGCGK |
| EPB41 | EPB41\_HUMAN | Protein 4.1 | 416 | 403 | EPB41 | EPB41\_HUMAN | Protein 4.1 | 14C(125.0477) | NaN | X | NaN | NaN | NaN | NaN | 1.698702e+05 | 1.698702e+05 | NaN | NaN | AKDLEGVDIILGVCSSGLLVYK |
| AIDA | AIDA\_HUMAN | Axin interactor, dorsalization-associated protein | 184 | 180 | AIDA | AIDA\_HUMAN | Axin interactor, dorsalization-associated protein | 5C(125.0477) | NaN | X | NaN | NaN | NaN | NaN | 1.697779e+05 | 1.697779e+05 | NaN | NaN | DAGQCIDPYITVSVK |
| UCHL3 | UCHL3\_HUMAN | Ubiquitin carboxyl-terminal hydrolase isozyme L3 | 209 | 187 | UCHL3 | UCHL3\_HUMAN | Ubiquitin carboxyl-terminal hydrolase isozyme L3 | 23C(125.0477) | NaN | X | NaN | NaN | NaN | NaN | 1.695880e+05 | 1.695880e+05 | NaN | NaN | KPFPINHGETSDETLLEDAIEVCKK |
| MRI1 | MTNA\_HUMAN | Methylthioribose-1-phosphate isomerase | 199 | 194 | MRI1 | MTNA\_HUMAN | Methylthioribose-1-phosphate isomerase | 6C(125.0477) | NaN | X | NaN | NaN | NaN | NaN | 1.695278e+05 | 1.695278e+05 | NaN | NaN | LEHAFCTETRPYNQGAR |
| CDK2 | CDK2\_HUMAN | Cyclin-dependent kinase 2 | 118 | 106 | CDK2 | CDK2\_HUMAN | Cyclin-dependent kinase 2 | 13C(125.0477) | NaN | X | NaN | NaN | NaN | NaN | 1.690585e+05 | 1.690585e+05 | NaN | NaN | SYLFQLLQGLAFCHSHR |
| BLVRA | BIEA\_HUMAN | Biliverdin reductase A | 204 | 201 | BLVRA | BIEA\_HUMAN | Biliverdin reductase A | 4C(125.0477) | NaN | X | NaN | NaN | NaN | NaN | 1.687640e+05 | 1.514858e+05 | 1.727816e+04 | NaN | MTVCLETEKK |
| WDR81 | WDR81\_HUMAN | WD repeat-containing protein 81 | 76 | 71 | WDR81 | WDR81\_HUMAN | WD repeat-containing protein 81 | 6C(125.0477) | NaN | X | NaN | NaN | NaN | NaN | 1.687621e+05 | 1.687621e+05 | NaN | NaN | LPLGPCPR |
| PARP10 | PAR10\_HUMAN | Protein mono-ADP-ribosyltransferase PARP10 | 757 | 757 | PARP10 | PAR10\_HUMAN | Protein mono-ADP-ribosyltransferase PARP10 | 1C(125.0477) | NaN | X | NaN | NaN | NaN | NaN | 1.686924e+05 | 1.686924e+05 | NaN | NaN | CHGVSVALR |
| RAP1B | RAP1B\_HUMAN | Ras-related protein Rap-1b | 118 | 118 | RAP1B | RAP1B\_HUMAN | Ras-related protein Rap-1b | 1C(125.0477) | NaN | X | NaN | NaN | NaN | NaN | 1.685998e+05 | 1.524962e+05 | 1.610358e+04 | NaN | CDLEDER |
| ANXA7 | ANXA7\_HUMAN | Annexin A7 | 413 | 409 | ANXA7 | ANXA7\_HUMAN | Annexin A7 | 5C(125.0477) | NaN | X | NaN | NaN | NaN | NaN | 1.684577e+05 | 1.684577e+05 | NaN | NaN | TILQCALNRPAFFAER |
| IGKC | IGKC\_HUMAN | Immunoglobulin kappa constant | 87, 87 | 82 | IGKC | IGKC\_HUMAN | Immunoglobulin kappa constant | 6C(125.0477) 6C(57.0215) | X | X | NaN | NaN | NaN | NaN | 1.680450e+05 | 1.680450e+05 | NaN | NaN | HKVYACEVTHQGLSSPVTK |
| PAK2 | PAK2\_HUMAN | Serine/threonine-protein kinase PAK 2 | 480 | 480 | PAK2 | PAK2\_HUMAN | Serine/threonine-protein kinase PAK 2 | 1C(125.0477) | NaN | X | NaN | NaN | NaN | NaN | 1.678552e+05 | 1.281604e+05 | 3.969479e+04 | NaN | CLEMDVEKR |
| HDGF | HDGF\_HUMAN | Hepatoma-derived growth factor | 108 | 107 | HDGF | HDGF\_HUMAN | Hepatoma-derived growth factor | 2C(125.0477) | NaN | X | NaN | NaN | NaN | NaN | 1.677658e+05 | 1.677658e+05 | NaN | NaN | SCVEEPEPEPEAAEGDGDKK |
| RHAG | RHAG\_HUMAN | Ammonium transporter Rh type A | 396 | 385 | RHAG | RHAG\_HUMAN | Ammonium transporter Rh type A | 12C(125.0477) | NaN | X | NaN | NaN | NaN | NaN | 1.676280e+05 | 1.676280e+05 | NaN | NaN | LPLWGQPSDQNCYDDSVYWK |
| SCCPDH | SCPDL\_HUMAN | Saccharopine dehydrogenase-like oxidoreductase | 98 | 91 | SCCPDH | SCPDL\_HUMAN | Saccharopine dehydrogenase-like oxidoreductase | 8C(125.0477) | NaN | X | NaN | NaN | NaN | NaN | 1.674789e+05 | 1.674789e+05 | NaN | NaN | QATVVLNCVGPYR |
| NAA10 | NAA10\_HUMAN | N-alpha-acetyltransferase 10 | 194 | 184 | NAA10 | NAA10\_HUMAN | N-alpha-acetyltransferase 10 | 11C(125.0477) | NaN | X | NaN | NaN | NaN | NaN | 1.672895e+05 | 1.672895e+05 | NaN | NaN | GNSPPSSGEACREEK |
| RPIA | RPIA\_HUMAN | Ribose-5-phosphate isomerase | 178 | 174 | RPIA | RPIA\_HUMAN | Ribose-5-phosphate isomerase | 5C(125.0477) | NaN | X | NaN | NaN | NaN | NaN | 1.668907e+05 | 6.975486e+04 | 9.713587e+04 | NaN | GGGGCLTQEK |
| SKP1 | SKP1\_HUMAN | S-phase kinase-associated protein 1 | 160 | 143 | SKP1 | SKP1\_HUMAN | S-phase kinase-associated protein 1 | 18C(125.0477) | NaN | X | NaN | NaN | NaN | NaN | 1.663999e+05 | 1.663999e+05 | NaN | NaN | NDFTEEEEAQVRKENQWCEEK |
| NHLRC2 | NHLC2\_HUMAN | NHL repeat-containing protein 2 | 609 | 603 | NHLRC2 | NHLC2\_HUMAN | NHL repeat-containing protein 2 | 7C(125.0477) | NaN | X | NaN | NaN | NaN | NaN | 1.662305e+05 | 1.662305e+05 | NaN | NaN | LSPVTACAGQTLQFK |
| PIN1 | PIN1\_HUMAN | Peptidyl-prolyl cis-trans isomerase NIMA-interacting 1 | 113 | 96 | PIN1 | PIN1\_HUMAN | Peptidyl-prolyl cis-trans isomerase NIMA-interacting 1 | 18C(125.0477) | NaN | X | NaN | NaN | NaN | NaN | 1.661468e+05 | 1.661468e+05 | NaN | NaN | IKSGEEDFESLASQFSDCSSAK |
| ATG2A | ATG2A\_HUMAN | Autophagy-related protein 2 homolog A | 1870 | 1860 | ATG2A | ATG2A\_HUMAN | Autophagy-related protein 2 homolog A | 11C(125.0477) | NaN | X | NaN | NaN | NaN | NaN | 1.659250e+05 | 1.659250e+05 | NaN | NaN | EGILDTAQTICDVASR |
| ALDH16A1 | A16A1\_HUMAN | Aldehyde dehydrogenase family 16 member A1 | 249 | 246 | ALDH16A1 | A16A1\_HUMAN | Aldehyde dehydrogenase family 16 member A1 | 4C(125.0477) | NaN | X | NaN | NaN | NaN | NaN | 1.659126e+05 | 1.195957e+05 | 4.631689e+04 | NaN | VAFCGAPEEGR |
| PRMT5 | ANM5\_HUMAN | Protein arginine N-methyltransferase 5 | 622 | 610 | PRMT5 | ANM5\_HUMAN | Protein arginine N-methyltransferase 5 | 13C(125.0477) | NaN | X | NaN | NaN | NaN | NaN | 1.657746e+05 | 1.657746e+05 | NaN | NaN | KVWYEWAVTAPVCSAIHNPTGR |
| MYH9 | MYH9\_HUMAN | Myosin-9 | 931 | 931 | MYH9 | MYH9\_HUMAN | Myosin-9 | 1C(125.0477) | NaN | X | NaN | NaN | NaN | NaN | 1.655007e+05 | 1.655007e+05 | NaN | NaN | CQHLQAEK |
| GLO1 | LGUL\_HUMAN | Lactoylglutathione lyase | 139 | 124 | GLO1 | LGUL\_HUMAN | Lactoylglutathione lyase | 16C(125.0477) | NaN | X | NaN | NaN | NaN | NaN | 1.652925e+05 | 1.652925e+05 | NaN | NaN | GFGHIGIAVPDVYSACKR |
| ARMC8 | ARMC8\_HUMAN | Armadillo repeat-containing protein 8 | 578 | 574 | ARMC8 | ARMC8\_HUMAN | Armadillo repeat-containing protein 8 | 5C(125.0477) | NaN | X | NaN | NaN | NaN | NaN | 1.645087e+05 | 1.645087e+05 | NaN | NaN | EQTLCILANIADGTTAK |
| SOD1 | SODC\_HUMAN | Superoxide dismutase [Cu-Zn] | 147 | 145 | SOD1 | SODC\_HUMAN | Superoxide dismutase [Cu-Zn] | 3C(125.0477) | NaN | X | NaN | NaN | NaN | NaN | 1.644910e+05 | NaN | 1.644910e+05 | NaN | LACGVIGIAQ |
| ISYNA1 | INO1\_HUMAN | Inositol-3-phosphate synthase 1 | 235 | 234 | ISYNA1 | INO1\_HUMAN | Inositol-3-phosphate synthase 1 | 2C(125.0477) | NaN | X | NaN | NaN | NaN | NaN | 1.643112e+05 | 1.643112e+05 | NaN | NaN | FCEVIPGLNDTAENLLR |
| NAXD | NNRD\_HUMAN | ATP-dependent (S)-NAD(P)H-hydrate dehydratase | 310 | 296 | NAXD | NNRD\_HUMAN | ATP-dependent (S)-NAD(P)H-hydrate dehydratase | 15C(15.9949) | NaN | NaN | X | NaN | NaN | NaN | 1.642222e+05 | 1.642222e+05 | NaN | NaN | TNGSSPLLVAAFGACSLTRQCNHQAFQK |
| SH3GLB2 | SHLB2\_HUMAN | Endophilin-B2 | 172 | 166 | SH3GLB2 | SHLB2\_HUMAN | Endophilin-B2 | 7C(125.0477) | NaN | X | NaN | NaN | NaN | NaN | 1.640999e+05 | 1.226099e+05 | 4.149000e+04 | NaN | RLDLDACK |
| YWHAZ | 1433Z\_HUMAN | 14-3-3 protein zeta/delta | 25 | 12 | YWHAZ | 1433Z\_HUMAN | 14-3-3 protein zeta/delta | 14C(125.0477) | NaN | X | NaN | NaN | NaN | NaN | 1.640618e+05 | 1.640618e+05 | NaN | NaN | LAEQAERYDDMAACMK |
| ALB | ALBU\_HUMAN | Albumin | 302, 302, 302, 303, 303 | 299 | ALB | ALBU\_HUMAN | Albumin | 4C(125.0477) 4C(31.9988) 4C(57.0215) 5C(125.0477) 5C(57.0215) | X | X | NaN | X | NaN | NaN | 1.634925e+05 | 1.634925e+05 | NaN | NaN | LKECCEKPLLEK |
| NCBP1 | NCBP1\_HUMAN | Nuclear cap-binding protein subunit 1 | 728 | 728 | NCBP1 | NCBP1\_HUMAN | Nuclear cap-binding protein subunit 1 | 1C(125.0477) | NaN | X | NaN | NaN | NaN | NaN | 1.634640e+05 | 1.634640e+05 | NaN | NaN | CETDGTSVLTPWYK |
| AMBP | AMBP\_HUMAN | Protein AMBP | 277, 281 | 276 | AMBP | AMBP\_HUMAN | Protein AMBP | 2C(57.0215) 6C(15.9949) | X | NaN | X | NaN | NaN | NaN | 1.634096e+05 | NaN | 1.634096e+05 | NaN | ECLQTCR |
| SHMT1 | GLYC\_HUMAN | Serine hydroxymethyltransferase, cytosolic | 335 | 326 | SHMT1 | GLYC\_HUMAN | Serine hydroxymethyltransferase, cytosolic | 10C(125.0477) | NaN | X | NaN | NaN | NaN | NaN | 1.633964e+05 | 1.633964e+05 | NaN | NaN | VYQHQVVANCR |
| LRRC40 | LRC40\_HUMAN | Leucine-rich repeat-containing protein 40 | 264 | 257 | LRRC40 | LRC40\_HUMAN | Leucine-rich repeat-containing protein 40 | 8C(125.0477) | NaN | X | NaN | NaN | NaN | NaN | 1.630411e+05 | 1.630411e+05 | NaN | NaN | FLPEFPSCSLLK |
| SGTA | SGTA\_HUMAN | Small glutamine-rich tetratricopeptide repeat-containing protein alpha | 148 | 138 | SGTA | SGTA\_HUMAN | Small glutamine-rich tetratricopeptide repeat-containing protein alpha | 11C(15.9949) | NaN | NaN | X | NaN | NaN | NaN | 1.629100e+05 | 1.629100e+05 | NaN | NaN | LGNYAGAVQDCERAICIDPAYSK |
| FSCN1 | FSCN1\_HUMAN | Fascin | 397 | 390 | FSCN1 | FSCN1\_HUMAN | Fascin | 8C(125.0477) | NaN | X | NaN | NaN | NaN | NaN | 1.627141e+05 | 1.478527e+05 | 1.486144e+04 | NaN | GEHGFIGCR |
| CFH | CFAH\_HUMAN | Complement factor H | 1201 | 1193 | CFH | CFAH\_HUMAN | Complement factor H | 9C(57.0215) | X | NaN | NaN | NaN | NaN | NaN | 1.626118e+05 | 5.905648e+04 | 1.035553e+05 | NaN | TGESVEFVCKR |
| ZBTB17 | ZBT17\_HUMAN | Zinc finger and BTB domain-containing protein 17 | 448 | 445 | ZBTB17 | ZBT17\_HUMAN | Zinc finger and BTB domain-containing protein 17 | 4C(57.0215) | X | NaN | NaN | NaN | NaN | NaN | 1.625192e+05 | 1.625192e+05 | NaN | NaN | EHKCPHCDK |
| TMX1 | TMX1\_HUMAN | Thioredoxin-related transmembrane protein 1 | 106 | 95 | TMX1 | TMX1\_HUMAN | Thioredoxin-related transmembrane protein 1 | 12C(125.0477) | NaN | X | NaN | NaN | NaN | NaN | 1.624647e+05 | 1.624647e+05 | NaN | NaN | FIITALPTIYHCK |
| RO60 | RO60\_HUMAN | RNA-binding protein RO60 | 38 | 36 | RO60 | RO60\_HUMAN | RNA-binding protein RO60 | 3C(125.0477) | NaN | X | NaN | NaN | NaN | NaN | 1.623454e+05 | 1.623454e+05 | NaN | NaN | FLCFGSEGGTYYIK |
| PPIA | PPIA\_HUMAN | Peptidyl-prolyl cis-trans isomerase A | 161 | 155 | PPIA | PPIA\_HUMAN | Peptidyl-prolyl cis-trans isomerase A | 7C(125.0477) | NaN | X | NaN | NaN | NaN | NaN | 1.621700e+05 | 6.884499e+04 | 9.332499e+04 | NaN | KITIADCGQLE |
| RAP1B | RAP1B\_HUMAN | Ras-related protein Rap-1b | 118 | 118 | RAP1B | RAP1B\_HUMAN | Ras-related protein Rap-1b | 1C(125.0477) | NaN | X | NaN | NaN | NaN | NaN | 1.618961e+05 | 1.618961e+05 | NaN | NaN | CDLEDERVVGKEQGQNLAR |
| MDH1 | MDHC\_HUMAN | Malate dehydrogenase, cytoplasmic | 154 | 150 | MDH1 | MDHC\_HUMAN | Malate dehydrogenase, cytoplasmic | 5C(125.0477) | NaN | X | NaN | NaN | NaN | NaN | 1.618462e+05 | NaN | 1.618462e+05 | NaN | ENFSCLTR |
| HSPA4 | HSP74\_HUMAN | Heat shock 70 kDa protein 4 | 310, 310, 310 | 306 | HSPA4 | HSP74\_HUMAN | Heat shock 70 kDa protein 4 | 5C(125.0477) 5C(15.9949) 5C(31.9988) | NaN | X | X | X | NaN | NaN | 1.613139e+05 | 1.613139e+05 | NaN | NaN | FLEMCNDLLAR |
| AP2B1 | AP2B1\_HUMAN | AP-2 complex subunit beta | 123 | 118 | AP2B1 | AP2B1\_HUMAN | AP-2 complex subunit beta | 6C(125.0477) | NaN | X | NaN | NaN | NaN | NaN | 1.613033e+05 | 6.903820e+04 | 9.226511e+04 | NaN | ITEYLCEPLRK |
| BCAM | BCAM\_HUMAN | Basal cell adhesion molecule | 337 | 330 | BCAM | BCAM\_HUMAN | Basal cell adhesion molecule | 8C(125.0477) | NaN | X | NaN | NaN | NaN | NaN | 1.608515e+05 | NaN | 1.185387e+05 | 42312.7460 | GQSGTYGCR |
| SCRN1 | SCRN1\_HUMAN | Secernin-1 | 324 | 320 | SCRN1 | SCRN1\_HUMAN | Secernin-1 | 5C(125.0477) | NaN | X | NaN | NaN | NaN | NaN | 1.606162e+05 | 1.606162e+05 | NaN | NaN | TQSPCFGDDDPAK |
| PSMD1 | PSMD1\_HUMAN | 26S proteasome non-ATPase regulatory subunit 1 | 112 | 111 | PSMD1 | PSMD1\_HUMAN | 26S proteasome non-ATPase regulatory subunit 1 | 2C(125.0477) | NaN | X | NaN | NaN | NaN | NaN | 1.604103e+05 | 1.273853e+05 | 3.302495e+04 | NaN | QCVENADLPEGEK |
| MYH2 | MYH2\_HUMAN | Myosin-2 | 953 | 948 | MYH2 | MYH2\_HUMAN | Myosin-2 | 6C(57.0215) | X | NaN | NaN | NaN | NaN | NaN | 1.597895e+05 | 1.597895e+05 | NaN | NaN | KLEDECSELKK |
| LANCL2 | LANC2\_HUMAN | LanC-like protein 2 | 350 | 346 | LANCL2 | LANC2\_HUMAN | LanC-like protein 2 | 5C(125.0477) | NaN | X | NaN | NaN | NaN | NaN | 1.596779e+05 | 1.596779e+05 | NaN | NaN | EAMECSDVIWQR |
| NRDC | NRDC\_HUMAN | Nardilysin | 269 | 259 | NRDC | NRDC\_HUMAN | Nardilysin | 11C(125.0477) | NaN | X | NaN | NaN | NaN | NaN | 1.596425e+05 | 1.596425e+05 | NaN | NaN | HGGSDNASTDCER |
| CIAPIN1 | CPIN1\_HUMAN | Anamorsin | 92 | 86 | CIAPIN1 | CPIN1\_HUMAN | Anamorsin | 7C(125.0477) | NaN | X | NaN | NaN | NaN | NaN | 1.594725e+05 | 1.594725e+05 | NaN | NaN | ILRPGGCLFLK |
| RNF40 | BRE1B\_HUMAN | E3 ubiquitin-protein ligase BRE1B | 69, 69 | 67 | RNF40 | BRE1B\_HUMAN | E3 ubiquitin-protein ligase BRE1B | 3C(15.9949) 3C(31.9988) | NaN | NaN | X | X | NaN | NaN | 1.594667e+05 | NaN | NaN | 159466.6600 | QACEDELR |
| USP15 | UBP15\_HUMAN | Ubiquitin carboxyl-terminal hydrolase 15 | 791 | 790 | USP15 | UBP15\_HUMAN | Ubiquitin carboxyl-terminal hydrolase 15 | 2C(125.0477) | NaN | X | NaN | NaN | NaN | NaN | 1.593943e+05 | 1.547015e+05 | 4.692791e+03 | NaN | DCIELFTTK |
| CLTCL1 | CLH2\_HUMAN | Clathrin heavy chain 2 | 926 | 924 | CLTCL1 | CLH2\_HUMAN | Clathrin heavy chain 2 | 3C(57.0215) | X | NaN | NaN | NaN | NaN | NaN | 1.592559e+05 | 1.592559e+05 | NaN | NaN | GQCDLELIK |
| EEFSEC | SELB\_HUMAN | Selenocysteine-specific elongation factor | 406 | 398 | EEFSEC | SELB\_HUMAN | Selenocysteine-specific elongation factor | 9C(125.0477) | NaN | X | NaN | NaN | NaN | NaN | 1.589586e+05 | 1.589586e+05 | NaN | NaN | AGQATEGHCPR |
| CRNN | CRNN\_HUMAN | Cornulin | 25 | 21 | CRNN | CRNN\_HUMAN | Cornulin | 5C(125.0477) | NaN | X | NaN | NaN | NaN | NaN | 1.588810e+05 | NaN | 1.588810e+05 | NaN | TEGNCTALTR |
| ROCK1 | ROCK1\_HUMAN | Rho-associated protein kinase 1 | 231 | 231 | ROCK1 | ROCK1\_HUMAN | Rho-associated protein kinase 1 | 1C(125.0477) | NaN | X | NaN | NaN | NaN | NaN | 1.588382e+05 | 1.588382e+05 | NaN | NaN | CDTAVGTPDYISPEVLK |
| GMPR2 | GMPR2\_HUMAN | GMP reductase 2 | 205 | 192 | GMPR2 | GMPR2\_HUMAN | GMP reductase 2 | 14C(125.0477) | NaN | X | NaN | NaN | NaN | NaN | 1.587707e+05 | 1.587707e+05 | NaN | NaN | TGVGYPQLSAVMECADAAHGLK |
| SORD | DHSO\_HUMAN | Sorbitol dehydrogenase | 140 | 135 | SORD | DHSO\_HUMAN | Sorbitol dehydrogenase | 6C(125.0477) | NaN | X | NaN | NaN | NaN | NaN | 1.582703e+05 | 1.582703e+05 | NaN | NaN | HNAAFCYK |
| COPS5 | CSN5\_HUMAN | COP9 signalosome complex subunit 5 | 218 | 211 | COPS5 | CSN5\_HUMAN | COP9 signalosome complex subunit 5 | 8C(125.0477) | NaN | X | NaN | NaN | NaN | NaN | 1.577431e+05 | NaN | 1.577431e+05 | NaN | IEDFGVHCK |
| PSMD3 | PSMD3\_HUMAN | 26S proteasome non-ATPase regulatory subunit 3 | 210 | 210 | PSMD3 | PSMD3\_HUMAN | 26S proteasome non-ATPase regulatory subunit 3 | 1C(125.0477) | NaN | X | NaN | NaN | NaN | NaN | 1.576180e+05 | 1.576180e+05 | NaN | NaN | CYYYHAR |
| GDI2 | GDIB\_HUMAN | Rab GDP dissociation inhibitor beta | 202 | 194 | GDI2 | GDIB\_HUMAN | Rab GDP dissociation inhibitor beta | 9C(125.0477) | NaN | X | NaN | NaN | NaN | NaN | 1.573164e+05 | NaN | 1.573164e+05 | NaN | TDDYLDQPCYETINR |
| ANK1 | ANK1\_HUMAN | Ankyrin-1 | 320 | 302 | ANK1 | ANK1\_HUMAN | Ankyrin-1 | 19C(125.0477) | NaN | X | NaN | NaN | NaN | NaN | 1.572410e+05 | 1.572410e+05 | NaN | NaN | TKNGLSPIHMAAQGDHLDCVR |
| EPB41 | EPB41\_HUMAN | Protein 4.1 | 39, 49 | 20 | EPB41 | EPB41\_HUMAN | Protein 4.1 | 20C(-34.0809) 30C(31.9988) | NaN | NaN | NaN | X | X | NaN | 1.568876e+05 | NaN | 1.568876e+05 | NaN | EEGEEAINSGQQEPQQEESCQTAAEGDNWCEQK |
| BLMH | BLMH\_HUMAN | Bleomycin hydrolase | 164 | 163 | BLMH | BLMH\_HUMAN | Bleomycin hydrolase | 2C(125.0477) | NaN | X | NaN | NaN | NaN | NaN | 1.568425e+05 | 1.568425e+05 | NaN | NaN | KCFPESYTTEATR |
| CUL1 | CUL1\_HUMAN | Cullin-1 | 752 | 752 | CUL1 | CUL1\_HUMAN | Cullin-1 | 1C(125.0477) | NaN | X | NaN | NaN | NaN | NaN | 1.568059e+05 | 1.568059e+05 | NaN | NaN | CIDILIEK |
| IGHG4 | IGHG4\_HUMAN | Immunoglobulin heavy constant gamma 4 | 27, 27 | 17 | IGHG4 | IGHG4\_HUMAN | Immunoglobulin heavy constant gamma 4 | 11C(125.0477) 11C(57.0215) | X | X | NaN | NaN | NaN | NaN | 1.564303e+05 | NaN | 1.564303e+05 | NaN | STSESTAALGCLVK |
| PSMA5 | PSA5\_HUMAN | Proteasome subunit alpha type-5 | 76 | 67 | PSMA5 | PSA5\_HUMAN | Proteasome subunit alpha type-5 | 10C(125.0477) | NaN | X | NaN | NaN | NaN | NaN | 1.559939e+05 | 1.559939e+05 | NaN | NaN | IVEIDAHIGCAMSGLIADAK |
| TLN1 | TLN1\_HUMAN | Talin-1 | 732 | 722 | TLN1 | TLN1\_HUMAN | Talin-1 | 11C(31.9988) | NaN | NaN | NaN | X | NaN | NaN | 1.559151e+05 | 1.559151e+05 | NaN | NaN | VVAPTISSPVCQEQLVEAGR |
| PSMD3 | PSMD3\_HUMAN | 26S proteasome non-ATPase regulatory subunit 3 | 483 | 480 | PSMD3 | PSMD3\_HUMAN | 26S proteasome non-ATPase regulatory subunit 3 | 4C(125.0477) | NaN | X | NaN | NaN | NaN | NaN | 1.555309e+05 | 1.555309e+05 | NaN | NaN | ISFCLDIHNMSVK |
| CLTC | CLH1\_HUMAN | Clathrin heavy chain 1 | 753 | 750 | CLTC | CLH1\_HUMAN | Clathrin heavy chain 1 | 4C(125.0477) | NaN | X | NaN | NaN | NaN | NaN | 1.554628e+05 | 1.326165e+05 | 2.284632e+04 | NaN | ESNCYDPER |
| HBM | HBM\_HUMAN | Hemoglobin subunit mu | 49 | 41 | HBM | HBM\_HUMAN | Hemoglobin subunit mu | 9C(125.0477) | NaN | X | NaN | NaN | NaN | NaN | 1.552146e+05 | 1.552146e+05 | NaN | NaN | VYFPHLSACQDATQLLSHGQR |
| HMBS | HEM3\_HUMAN | Porphobilinogen deaminase | 114 | 99 | HMBS | HEM3\_HUMAN | Porphobilinogen deaminase | 16C(125.0477) | NaN | X | NaN | NaN | NaN | NaN | 1.551270e+05 | 1.551270e+05 | NaN | NaN | DLPTVLPPGFTIGAICKR |
| GLUL | GLNA\_HUMAN | Glutamine synthetase | 99 | 96 | GLUL | GLNA\_HUMAN | Glutamine synthetase | 4C(125.0477) | NaN | X | NaN | NaN | NaN | NaN | 1.549902e+05 | 1.549902e+05 | NaN | NaN | LVLCEVFK |
| ESD | ESTD\_HUMAN | S-formylglutathione hydrolase | 158 | 143 | ESD | ESTD\_HUMAN | S-formylglutathione hydrolase | 16C(125.0477) | NaN | X | NaN | NaN | NaN | NaN | 1.548924e+05 | 1.548924e+05 | NaN | NaN | MSIFGHSMGGHGALICALKNPGK |
| CUL5 | CUL5\_HUMAN | Cullin-5 | 188 | 182 | CUL5 | CUL5\_HUMAN | Cullin-5 | 7C(125.0477) | NaN | X | NaN | NaN | NaN | NaN | 1.548593e+05 | NaN | 1.548593e+05 | NaN | ESYVNLCSNPEDK |
| GCLC | GSH1\_HUMAN | Glutamate--cysteine ligase catalytic subunit | 615, 615, 605, 605, 613, 613 | 605 | GCLC | GSH1\_HUMAN | Glutamate--cysteine ligase catalytic subunit | 11C(125.0477) 11C(15.9949) 1C(125.0477) 1C(31.9988) 9C(125.0477) 9C(15.9949) | NaN | X | X | X | NaN | NaN | 1.547000e+05 | 1.547000e+05 | NaN | NaN | CNQIANELCECPELLGSAFR |
| TCP1 | TCPA\_HUMAN | T-complex protein 1 subunit alpha | 147 | 146 | TCP1 | TCPA\_HUMAN | T-complex protein 1 subunit alpha | 2C(125.0477) | NaN | X | NaN | NaN | NaN | NaN | 1.535462e+05 | NaN | 1.535462e+05 | NaN | DCLINAAK |
| PAFAH1B1 | LIS1\_HUMAN | Platelet-activating factor acetylhydrolase IB subunit beta | 356 | 352 | PAFAH1B1 | LIS1\_HUMAN | Platelet-activating factor acetylhydrolase IB subunit beta | 5C(125.0477) | NaN | X | NaN | NaN | NaN | NaN | 1.534820e+05 | 1.534820e+05 | NaN | NaN | FILSCADDKTLR |
| CTTN | SRC8\_HUMAN | Src substrate cortactin | 112 | 111 | CTTN | SRC8\_HUMAN | Src substrate cortactin | 2C(125.0477) | NaN | X | NaN | NaN | NaN | NaN | 1.532993e+05 | 1.269098e+05 | 2.638943e+04 | NaN | HCSQVDSVR |
| DHX9 | DHX9\_HUMAN | ATP-dependent RNA helicase A | 578 | 567 | DHX9 | DHX9\_HUMAN | ATP-dependent RNA helicase A | 12C(125.0477) | NaN | X | NaN | NaN | NaN | NaN | 1.531149e+05 | 1.531149e+05 | NaN | NaN | TYPVQEYFLEDCIQMTHFVPPPK |
| PSMG1 | PSMG1\_HUMAN | Proteasome assembly chaperone 1 | 106 | 101 | PSMG1 | PSMG1\_HUMAN | Proteasome assembly chaperone 1 | 6C(125.0477) | NaN | X | NaN | NaN | NaN | NaN | 1.529773e+05 | 1.529773e+05 | NaN | NaN | LWNEWCR |
| CUL4A | CUL4A\_HUMAN | Cullin-4A | 483 | 481 | CUL4A | CUL4A\_HUMAN | Cullin-4A | 3C(125.0477) | NaN | X | NaN | NaN | NaN | NaN | 1.527446e+05 | 1.527446e+05 | NaN | NaN | HECGAAFTSK |
| PFN1 | PROF1\_HUMAN | Profilin-1 | 128 | 128 | PFN1 | PROF1\_HUMAN | Profilin-1 | 1C(125.0477) | NaN | X | NaN | NaN | NaN | NaN | 1.526606e+05 | 1.526606e+05 | NaN | NaN | CYEMASHLR |
| GRHPR | GRHPR\_HUMAN | Glyoxylate reductase/hydroxypyruvate reductase | 57 | 48 | GRHPR | GRHPR\_HUMAN | Glyoxylate reductase/hydroxypyruvate reductase | 10C(125.0477) | NaN | X | NaN | NaN | NaN | NaN | 1.525189e+05 | 1.525189e+05 | NaN | NaN | GVAGAHGLLCLLSDHVDKR |
| VPS4A | VPS4A\_HUMAN | Vacuolar protein sorting-associated protein 4A | 403 | 397 | VPS4A | VPS4A\_HUMAN | Vacuolar protein sorting-associated protein 4A | 7C(15.9949) | NaN | NaN | X | NaN | NaN | NaN | 1.524343e+05 | 1.524343e+05 | NaN | NaN | LLEPVVCMSDMLR |
| MTPN | MTPN\_HUMAN | Myotrophin | 83 | 67 | MTPN | MTPN\_HUMAN | Myotrophin | 17C(125.0477) | NaN | X | NaN | NaN | NaN | NaN | 1.516322e+05 | 1.516322e+05 | NaN | NaN | HHITPLLSAVYEGHVSCVK |
| PSMD5 | PSMD5\_HUMAN | 26S proteasome non-ATPase regulatory subunit 5 | 133 | 129 | PSMD5 | PSMD5\_HUMAN | 26S proteasome non-ATPase regulatory subunit 5 | 5C(125.0477) | NaN | X | NaN | NaN | NaN | NaN | 1.514482e+05 | 1.514482e+05 | NaN | NaN | QIVYCIGGENLSVAK |
| ANXA11 | ANX11\_HUMAN | Annexin A11 | 384 | 379 | ANXA11 | ANX11\_HUMAN | Annexin A11 | 6C(125.0477) | NaN | X | NaN | NaN | NaN | NaN | 1.514063e+05 | 7.093203e+04 | 8.047430e+04 | NaN | FNAVLCSR |
| GALK1 | GALK1\_HUMAN | Galactokinase | 243 | 242 | GALK1 | GALK1\_HUMAN | Galactokinase | 2C(125.0477) | NaN | X | NaN | NaN | NaN | NaN | 1.513074e+05 | 1.513074e+05 | NaN | NaN | QCEEVAR |
| PHKA2 | KPB2\_HUMAN | Phosphorylase b kinase regulatory subunit alpha, liver isoform | 24 | 17 | PHKA2 | KPB2\_HUMAN | Phosphorylase b kinase regulatory subunit alpha, liver isoform | 8C(125.0477) | NaN | X | NaN | NaN | NaN | NaN | 1.508543e+05 | 1.508543e+05 | NaN | NaN | LVQQTILCYQNPVTGLLSASHEQK |
| KRT2 | K22E\_HUMAN | Keratin, type II cytoskeletal 2 epidermal | 489 | 482 | KRT2 | K22E\_HUMAN | Keratin, type II cytoskeletal 2 epidermal | 8C(15.9949) | NaN | NaN | X | NaN | NaN | NaN | 1.502927e+05 | 1.502927e+05 | NaN | NaN | KLLEGEECR |
| TP53I3 | QORX\_HUMAN | Quinone oxidoreductase PIG3 | 76 | 48 | TP53I3 | QORX\_HUMAN | Quinone oxidoreductase PIG3 | 29C(125.0477) | NaN | X | NaN | NaN | NaN | NaN | 1.502025e+05 | 1.502025e+05 | NaN | NaN | QGQYDPPPGASNILGLEASGHVAELGPGCQGHWK |
| PFAS | PUR4\_HUMAN | Phosphoribosylformylglycinamidine synthase | 512, 502 | 501 | PFAS | PUR4\_HUMAN | Phosphoribosylformylglycinamidine synthase | 12C(15.9949) 2C(15.9949) | NaN | NaN | X | NaN | NaN | NaN | 1.502009e+05 | 1.502009e+05 | NaN | NaN | ACVEAPKGNPICSLHDQGAGGNGNVLK |
| ACLY | ACLY\_HUMAN | ATP-citrate synthase | 20 | 13 | ACLY | ACLY\_HUMAN | ATP-citrate synthase | 8C(15.9949) | NaN | NaN | X | NaN | NaN | NaN | 1.501272e+05 | 1.501272e+05 | NaN | NaN | ELLYKFICTTSAIQNR |
| ZZEF1 | ZZEF1\_HUMAN | Zinc finger ZZ-type and EF-hand domain-containing protein 1 | 535 | 524 | ZZEF1 | ZZEF1\_HUMAN | Zinc finger ZZ-type and EF-hand domain-containing protein 1 | 12C(125.0477) | NaN | X | NaN | NaN | NaN | NaN | 1.499163e+05 | 1.499163e+05 | NaN | NaN | YGKPLQLTLQACDVK |
| TXNDC5 | TXND5\_HUMAN | Thioredoxin domain-containing protein 5 | 381 | 376 | TXNDC5 | TXND5\_HUMAN | Thioredoxin domain-containing protein 5 | 6C(15.9949) | NaN | NaN | X | NaN | NaN | NaN | 1.498760e+05 | NaN | 1.498760e+05 | NaN | IAEVDCTAERNICSK |
| CLTCL1 | CLH2\_HUMAN | Clathrin heavy chain 2 | 682 | 677 | CLTCL1 | CLH2\_HUMAN | Clathrin heavy chain 2 | 6C(125.0477) | NaN | X | NaN | NaN | NaN | NaN | 1.497877e+05 | 1.497877e+05 | NaN | NaN | QNLQLCVQVASK |
| HCLS1 | HCLS1\_HUMAN | Hematopoietic lineage cell-specific protein | 470 | 470 | HCLS1 | HCLS1\_HUMAN | Hematopoietic lineage cell-specific protein | 1C(125.0477) | NaN | X | NaN | NaN | NaN | NaN | 1.496898e+05 | 1.496898e+05 | NaN | NaN | CHGHFGLFPANYVK |
| HUWE1 | HUWE1\_HUMAN | E3 ubiquitin-protein ligase HUWE1 | 1401 | 1393 | HUWE1 | HUWE1\_HUMAN | E3 ubiquitin-protein ligase HUWE1 | 9C(125.0477) | NaN | X | NaN | NaN | NaN | NaN | 1.494591e+05 | 1.494591e+05 | NaN | NaN | AESPEEVACRK |
| IDH1 | IDHC\_HUMAN | Isocitrate dehydrogenase [NADP] cytoplasmic | 73 | 73 | IDH1 | IDHC\_HUMAN | Isocitrate dehydrogenase [NADP] cytoplasmic | 1C(125.0477) | NaN | X | NaN | NaN | NaN | NaN | 1.493083e+05 | 9.746421e+04 | 5.184407e+04 | NaN | CATITPDEK |
| GGCT | GGCT\_HUMAN | Gamma-glutamylcyclotransferase | 122 | 112 | GGCT | GGCT\_HUMAN | Gamma-glutamylcyclotransferase | 11C(125.0477) | NaN | X | NaN | NaN | NaN | NaN | 1.491377e+05 | 1.491377e+05 | NaN | NaN | VATQEGKEITCR |
| RELL1 | RELL1\_HUMAN | RELT-like protein 1 | 88 | 88 | RELL1 | RELL1\_HUMAN | RELT-like protein 1 | 1C(125.0477) | NaN | X | NaN | NaN | NaN | NaN | 1.490189e+05 | NaN | 1.490189e+05 | NaN | CTTEAEQDIEEEK |
| JMJD6 | JMJD6\_HUMAN | Bifunctional arginine demethylase and lysyl-hydroxylase JMJD6 | 207 | 206 | JMJD6 | JMJD6\_HUMAN | Bifunctional arginine demethylase and lysyl-hydroxylase JMJD6 | 2C(125.0477) | NaN | X | NaN | NaN | NaN | NaN | 1.488811e+05 | 1.488811e+05 | NaN | NaN | WCLFPTSTPR |
| GDI2 | GDIB\_HUMAN | Rab GDP dissociation inhibitor beta | 414 | 403 | GDI2 | GDIB\_HUMAN | Rab GDP dissociation inhibitor beta | 12C(125.0477) | NaN | X | NaN | NaN | NaN | NaN | 1.487655e+05 | 1.487655e+05 | NaN | NaN | TYDATTHFETTCDDIKNIYK |
| SCFD1 | SCFD1\_HUMAN | Sec1 family domain-containing protein 1 | 623 | 618 | SCFD1 | SCFD1\_HUMAN | Sec1 family domain-containing protein 1 | 6C(125.0477) | NaN | X | NaN | NaN | NaN | NaN | 1.482742e+05 | 1.482742e+05 | NaN | NaN | HILYGCSELFNATQFIK |
| CAT | CATA\_HUMAN | Catalase | 232 | 211 | CAT | CATA\_HUMAN | Catalase | 22C(125.0477) | NaN | X | NaN | NaN | NaN | NaN | 1.479079e+05 | 1.479079e+05 | NaN | NaN | HMNGYGSHTFKLVNANGEAVYCK |
| PPP2CA | PP2AA\_HUMAN | Serine/threonine-protein phosphatase 2A catalytic subunit alpha isoform | 20 | 9 | PPP2CA | PP2AA\_HUMAN | Serine/threonine-protein phosphatase 2A catalytic subunit alpha isoform | 12C(125.0477) | NaN | X | NaN | NaN | NaN | NaN | 1.470888e+05 | 1.470888e+05 | NaN | NaN | ELDQWIEQLNECK |
| FAS | TNR6\_HUMAN | Tumor necrosis factor receptor superfamily member 6 | 304 | 301 | FAS | TNR6\_HUMAN | Tumor necrosis factor receptor superfamily member 6 | 4C(125.0477) | NaN | X | NaN | NaN | NaN | NaN | 1.467457e+05 | NaN | 1.467457e+05 | NaN | ANLCTLAEK |
| KRT14 | K1C14\_HUMAN | Keratin, type I cytoskeletal 14 | 18 | 16 | KRT14 | K1C14\_HUMAN | Keratin, type I cytoskeletal 14 | 3C(31.9988) | NaN | NaN | NaN | X | NaN | NaN | 1.465103e+05 | NaN | NaN | 146510.3300 | GSCGIGGGIGGGSSR |
| LANCL2 | LANC2\_HUMAN | LanC-like protein 2 | 321 | 317 | LANCL2 | LANC2\_HUMAN | LanC-like protein 2 | 5C(125.0477) | NaN | X | NaN | NaN | NaN | NaN | 1.464588e+05 | 1.464588e+05 | NaN | NaN | LVHWCHGAPGVIHMLMQAYK |
| CCT4 | TCPD\_HUMAN | T-complex protein 1 subunit delta | 221 | 214 | CCT4 | TCPD\_HUMAN | T-complex protein 1 subunit delta | 8C(125.0477) | NaN | X | NaN | NaN | NaN | NaN | 1.463188e+05 | 1.463188e+05 | NaN | NaN | LGGTIDDCELVEGLVLTQK |
| NAPRT | PNCB\_HUMAN | Nicotinate phosphoribosyltransferase | 101 | 98 | NAPRT | PNCB\_HUMAN | Nicotinate phosphoribosyltransferase | 4C(125.0477) | NaN | X | NaN | NaN | NaN | NaN | 1.461152e+05 | 6.093791e+04 | 8.517726e+04 | NaN | ALDCSEVTVR |
| FAF1 | FAF1\_HUMAN | FAS-associated factor 1 | 120 | 111 | FAF1 | FAF1\_HUMAN | FAS-associated factor 1 | 10C(125.0477) | NaN | X | NaN | NaN | NaN | NaN | 1.460624e+05 | 1.460624e+05 | NaN | NaN | NVDVVLEDTCTVGEIK |
| EPS15 | EPS15\_HUMAN | Epidermal growth factor receptor substrate 15 | 269 | 255 | EPS15 | EPS15\_HUMAN | Epidermal growth factor receptor substrate 15 | 15C(125.0477) | NaN | X | NaN | NaN | NaN | NaN | 1.459812e+05 | 1.459812e+05 | NaN | NaN | TGLPSTLLAHIWSLCDTK |
| ANXA1 | ANXA1\_HUMAN | Annexin A1 | 263 | 263 | ANXA1 | ANXA1\_HUMAN | Annexin A1 | 1C(125.0477) | NaN | X | NaN | NaN | NaN | NaN | 1.459557e+05 | 1.459557e+05 | NaN | NaN | CLTAIVKCATSKPAFFAEK |
| TCP1 | TCPA\_HUMAN | T-complex protein 1 subunit alpha | 385 | 379 | TCP1 | TCPA\_HUMAN | T-complex protein 1 subunit alpha | 7C(125.0477) | NaN | X | NaN | NaN | NaN | NaN | 1.458744e+05 | 1.458744e+05 | NaN | NaN | GANDFMCDEMER |
| PTPN11 | PTN11\_HUMAN | Tyrosine-protein phosphatase non-receptor type 11 | 104 | 100 | PTPN11 | PTN11\_HUMAN | Tyrosine-protein phosphatase non-receptor type 11 | 5C(125.0477) | NaN | X | NaN | NaN | NaN | NaN | 1.457216e+05 | 9.511945e+04 | 5.060216e+04 | NaN | YPLNCADPTSER |
| CAT | CATA\_HUMAN | Catalase | 393 | 389 | CAT | CATA\_HUMAN | Catalase | 5C(125.0477) | NaN | X | NaN | NaN | NaN | NaN | 1.455727e+05 | 1.455727e+05 | NaN | NaN | DGPMCMQDNQGGAPNYYPNSFGAPEQQPSALEHSIQYSGEVR |
| ANK1 | ANK1\_HUMAN | Ankyrin-1 | 1174 | 1172 | ANK1 | ANK1\_HUMAN | Ankyrin-1 | 3C(125.0477) | NaN | X | NaN | NaN | NaN | NaN | 1.454171e+05 | NaN | 1.454171e+05 | NaN | LLCSVIGGTDQAQWEDITGTTK |
| CCT4 | TCPD\_HUMAN | T-complex protein 1 subunit delta | 252 | 244 | CCT4 | TCPD\_HUMAN | T-complex protein 1 subunit delta | 9C(125.0477) | NaN | X | NaN | NaN | NaN | NaN | 1.451082e+05 | 1.451082e+05 | NaN | NaN | AKIGLIQFCLSAPK |
| PPP2R1A | 2AAA\_HUMAN | Serine/threonine-protein phosphatase 2A 65 kDa regulatory subunit A alpha isoform | 329 | 319 | PPP2R1A | 2AAA\_HUMAN | Serine/threonine-protein phosphatase 2A 65 kDa regulatory subunit A alpha isoform | 11C(125.0477) | NaN | X | NaN | NaN | NaN | NaN | 1.451071e+05 | 1.451071e+05 | NaN | NaN | ENVIMSQILPCIK |
| KRT14 | K1C14\_HUMAN | Keratin, type I cytoskeletal 14 | 18 | 16 | KRT14 | K1C14\_HUMAN | Keratin, type I cytoskeletal 14 | 3C(125.0477) | NaN | X | NaN | NaN | NaN | NaN | 1.445248e+05 | NaN | 1.445248e+05 | NaN | GSCGIGGGIGGGSSR |
| GPS1 | CSN1\_HUMAN | COP9 signalosome complex subunit 1 | 175 | 164 | GPS1 | CSN1\_HUMAN | COP9 signalosome complex subunit 1 | 12C(125.0477) | NaN | X | NaN | NaN | NaN | NaN | 1.443434e+05 | 1.443434e+05 | NaN | NaN | GHDDLGDHYLDCGDLSNALK |
| BCAM | BCAM\_HUMAN | Basal cell adhesion molecule | 473 | 465 | BCAM | BCAM\_HUMAN | Basal cell adhesion molecule | 9C(125.0477) | NaN | X | NaN | NaN | NaN | NaN | 1.440132e+05 | NaN | 1.203190e+05 | 23694.1820 | EGDEVTLICSAR |
| PGK1 | PGK1\_HUMAN | Phosphoglycerate kinase 1 | 316 | 298 | PGK1 | PGK1\_HUMAN | Phosphoglycerate kinase 1 | 19C(125.0477) | NaN | X | NaN | NaN | NaN | NaN | 1.433618e+05 | 1.433618e+05 | NaN | NaN | TGQATVASGIPAGWMGLDCGPESSK |
| ENOSF1 | ENOF1\_HUMAN | Mitochondrial enolase superfamily member 1 | 307 | 293 | ENOSF1 | ENOF1\_HUMAN | Mitochondrial enolase superfamily member 1 | 15C(125.0477) | NaN | X | NaN | NaN | NaN | NaN | 1.432343e+05 | 1.432343e+05 | NaN | NaN | ALVPLGIGIATGEQCHNR |
| CD55 | DAF\_HUMAN | Complement decay-accelerating factor | 163 | 162 | CD55 | DAF\_HUMAN | Complement decay-accelerating factor | 2C(125.0477) | NaN | X | NaN | NaN | NaN | NaN | 1.425516e+05 | NaN | 8.850985e+04 | 54041.7340 | SCPNPGEIR |
| IDE | IDE\_HUMAN | Insulin-degrading enzyme | 414 | 403 | IDE | IDE\_HUMAN | Insulin-degrading enzyme | 12C(125.0477) | NaN | X | NaN | NaN | NaN | NaN | 1.425243e+05 | 1.425243e+05 | NaN | NaN | AEGPQEWVFQECK |
| SEMA7A | SEM7A\_HUMAN | Semaphorin-7A | 511 | 508 | SEMA7A | SEM7A\_HUMAN | Semaphorin-7A | 4C(125.0477) | NaN | X | NaN | NaN | NaN | NaN | 1.423813e+05 | NaN | 1.423813e+05 | NaN | DPYCGWDQGR |
| STOM | STOM\_HUMAN | Stomatin | 87 | 78 | STOM | STOM\_HUMAN | Stomatin | 10C(125.0477) | NaN | X | NaN | NaN | NaN | NaN | 1.421565e+05 | 1.421565e+05 | NaN | NaN | GPGLFFILPCTDSFIK |
| PPM1A | PPM1A\_HUMAN | Protein phosphatase 1A | 144 | 125 | PPM1A | PPM1A\_HUMAN | Protein phosphatase 1A | 20C(125.0477) | NaN | X | NaN | NaN | NaN | NaN | 1.421312e+05 | 1.421312e+05 | NaN | NaN | SGSTAVGVLISPQHTYFINCGDSR |
| VPS35 | VPS35\_HUMAN | Vacuolar protein sorting-associated protein 35 | 253 | 240 | VPS35 | VPS35\_HUMAN | Vacuolar protein sorting-associated protein 35 | 14C(125.0477) | NaN | X | NaN | NaN | NaN | NaN | 1.419117e+05 | 1.419117e+05 | NaN | NaN | QIVLTGILEQVVNCR |
| CUL1 | CUL1\_HUMAN | Cullin-1 | 496 | 494 | CUL1 | CUL1\_HUMAN | Cullin-1 | 3C(125.0477) | NaN | X | NaN | NaN | NaN | NaN | 1.414534e+05 | 1.414534e+05 | NaN | NaN | QACGFEYTSK |
| VAPB | VAPB\_HUMAN | Vesicle-associated membrane protein-associated protein B/C | 173 | 168 | VAPB | VAPB\_HUMAN | Vesicle-associated membrane protein-associated protein B/C | 6C(125.0477) | NaN | X | NaN | NaN | NaN | NaN | 1.414343e+05 | 1.414343e+05 | NaN | NaN | KVMEECKR |
| PITHD1 | PITH1\_HUMAN | PITH domain-containing protein 1 | 42, 42 | 40 | PITHD1 | PITH1\_HUMAN | PITH domain-containing protein 1 | 3C(125.0477) 3C(31.9988) | NaN | X | NaN | X | NaN | NaN | 1.413763e+05 | 1.413763e+05 | NaN | NaN | LQCLNESR |
| PCYT2 | PCY2\_HUMAN | Ethanolamine-phosphate cytidylyltransferase | 278 | 272 | PCYT2 | PCY2\_HUMAN | Ethanolamine-phosphate cytidylyltransferase | 7C(125.0477) | NaN | X | NaN | NaN | NaN | NaN | 1.408256e+05 | 1.408256e+05 | NaN | NaN | TLSVLACR |
| FSCN1 | FSCN1\_HUMAN | Fascin | 61 | 44 | FSCN1 | FSCN1\_HUMAN | Fascin | 18C(125.0477) | NaN | X | NaN | NaN | NaN | NaN | 1.406816e+05 | 1.406816e+05 | NaN | NaN | QIWTLEQPPDEAGSAAVCLR |
| NGLY1 | NGLY1\_HUMAN | Peptide-N(4)-(N-acetyl-beta-glucosaminyl)asparagine amidase | 496 | 492 | NGLY1 | NGLY1\_HUMAN | Peptide-N(4)-(N-acetyl-beta-glucosaminyl)asparagine amidase | 5C(125.0477) | NaN | X | NaN | NaN | NaN | NaN | 1.405797e+05 | 1.405797e+05 | NaN | NaN | QLHLCYNIVK |
| UBR4 | UBR4\_HUMAN | E3 ubiquitin-protein ligase UBR4 | 4890 | 4873 | UBR4 | UBR4\_HUMAN | E3 ubiquitin-protein ligase UBR4 | 18C(125.0477) | NaN | X | NaN | NaN | NaN | NaN | 1.404801e+05 | 1.404801e+05 | NaN | NaN | KQQGYSTVSHFNIVHYDCHLAAVR |
| NT5DC1 | NT5D1\_HUMAN | 5'-nucleotidase domain-containing protein 1 | 197 | 193 | NT5DC1 | NT5D1\_HUMAN | 5'-nucleotidase domain-containing protein 1 | 5C(125.0477) | NaN | X | NaN | NaN | NaN | NaN | 1.402465e+05 | 1.402465e+05 | NaN | NaN | YLHSCPESVK |
| PRKAR2B | KAP3\_HUMAN | cAMP-dependent protein kinase type II-beta regulatory subunit | 388 | 384 | PRKAR2B | KAP3\_HUMAN | cAMP-dependent protein kinase type II-beta regulatory subunit | 5C(125.0477) | NaN | X | NaN | NaN | NaN | NaN | 1.401591e+05 | 1.401591e+05 | NaN | NaN | LLGPCMEIMK |
| NAE1 | ULA1\_HUMAN | NEDD8-activating enzyme E1 regulatory subunit | 483 | 476 | NAE1 | ULA1\_HUMAN | NEDD8-activating enzyme E1 regulatory subunit | 8C(125.0477) | NaN | X | NaN | NaN | NaN | NaN | 1.401089e+05 | 1.233874e+05 | 1.672157e+04 | NaN | DDYVHEFCR |
| USP24 | UBP24\_HUMAN | Ubiquitin carboxyl-terminal hydrolase 24 | 802 | 794 | USP24 | UBP24\_HUMAN | Ubiquitin carboxyl-terminal hydrolase 24 | 9C(125.0477) | NaN | X | NaN | NaN | NaN | NaN | 1.401062e+05 | 1.401062e+05 | NaN | NaN | TFFENVNLCDHR |
| UBA1 | UBA1\_HUMAN | Ubiquitin-like modifier-activating enzyme 1 | 632 | 628 | UBA1 | UBA1\_HUMAN | Ubiquitin-like modifier-activating enzyme 1 | 5C(125.0477) | NaN | X | NaN | NaN | NaN | NaN | 1.397269e+05 | 1.031879e+05 | 3.653893e+04 | NaN | SIPICTLK |
| FCSK | FCSK\_HUMAN | L-fucose kinase | 787 | 785 | FCSK | FCSK\_HUMAN | L-fucose kinase | 3C(125.0477) | NaN | X | NaN | NaN | NaN | NaN | 1.396122e+05 | 1.396122e+05 | NaN | NaN | DYCQPHAPGALLK |
| GSTT1 | GSTT1\_HUMAN | Glutathione S-transferase theta-1 | 182 | 156 | GSTT1 | GSTT1\_HUMAN | Glutathione S-transferase theta-1 | 27C(125.0477) | NaN | X | NaN | NaN | NaN | NaN | 1.394541e+05 | 1.394541e+05 | NaN | NaN | AFLTGPHISLADLVAITELMHPVGAGCQVFEGRPK |
| PSMC1 | PRS4\_HUMAN | 26S proteasome regulatory subunit 4 | 58 | 49 | PSMC1 | PRS4\_HUMAN | 26S proteasome regulatory subunit 4 | 10C(125.0477) | NaN | X | NaN | NaN | NaN | NaN | 1.392298e+05 | 1.088347e+05 | 3.039506e+04 | NaN | LPLVTPHTQCR |
| PKM | KPYM\_HUMAN | Pyruvate kinase PKM | 474 | 468 | PKM | KPYM\_HUMAN | Pyruvate kinase PKM | 7C(125.0477) | NaN | X | NaN | NaN | NaN | NaN | 1.388328e+05 | 1.388328e+05 | NaN | NaN | GIFPVLCK |
| ABCC5 | MRP5\_HUMAN | ATP-binding cassette sub-family C member 5 | 146 | 139 | ABCC5 | MRP5\_HUMAN | ATP-binding cassette sub-family C member 5 | 8C(125.0477) | NaN | X | NaN | NaN | NaN | NaN | 1.384560e+05 | 1.212160e+05 | 1.723999e+04 | NaN | HESSDVNCR |
| PGM1 | PGM1\_HUMAN | Phosphoglucomutase-1 | 101 | 86 | PGM1 | PGM1\_HUMAN | Phosphoglucomutase-1 | 16C(125.0477) | NaN | X | NaN | NaN | NaN | NaN | 1.384312e+05 | 1.384312e+05 | NaN | NaN | LVIGQNGILSTPAVSCIIR |
| UBE2O | UBE2O\_HUMAN | (E3-independent) E2 ubiquitin-conjugating enzyme | 353 | 353 | UBE2O | UBE2O\_HUMAN | (E3-independent) E2 ubiquitin-conjugating enzyme | 1C(125.0477) | NaN | X | NaN | NaN | NaN | NaN | 1.383350e+05 | 1.383350e+05 | NaN | NaN | CLYVFPAKVEPAK |
| CCT4 | TCPD\_HUMAN | T-complex protein 1 subunit delta | 221, 221 | 213 | CCT4 | TCPD\_HUMAN | T-complex protein 1 subunit delta | 9C(125.0477) 9C(15.9949) | NaN | X | X | NaN | NaN | NaN | 1.382288e+05 | 1.382288e+05 | NaN | NaN | KLGGTIDDCELVEGLVLTQK |
| CCT4 | TCPD\_HUMAN | T-complex protein 1 subunit delta | 295 | 293 | CCT4 | TCPD\_HUMAN | T-complex protein 1 subunit delta | 3C(125.0477) | NaN | X | NaN | NaN | NaN | NaN | 1.381832e+05 | 7.404194e+04 | 6.414130e+04 | NaN | TGCNVLLIQK |
| FASN | FAS\_HUMAN | Fatty acid synthase | 1828 | 1828 | FASN | FAS\_HUMAN | Fatty acid synthase | 1C(125.0477) | NaN | X | NaN | NaN | NaN | NaN | 1.380041e+05 | 1.380041e+05 | NaN | NaN | CTVFHGAQVEDAFR |
| XPO7 | XPO7\_HUMAN | Exportin-7 | 1038 | 1032 | XPO7 | XPO7\_HUMAN | Exportin-7 | 7C(125.0477) | NaN | X | NaN | NaN | NaN | NaN | 1.379919e+05 | 1.379919e+05 | NaN | NaN | QQAMHLCFENLMEGIER |
| ATG4A | ATG4A\_HUMAN | Cysteine protease ATG4A | 217 | 212 | ATG4A | ATG4A\_HUMAN | Cysteine protease ATG4A | 6C(125.0477) | NaN | X | NaN | NaN | NaN | NaN | 1.375818e+05 | 1.375818e+05 | NaN | NaN | GTSAYCSAWKPLLLIVPLR |
| PCMT1 | PIMT\_HUMAN | Protein-L-isoaspartate(D-aspartate) O-methyltransferase | 102 | 99 | PCMT1 | PIMT\_HUMAN | Protein-L-isoaspartate(D-aspartate) O-methyltransferase | 4C(125.0477) | NaN | X | NaN | NaN | NaN | NaN | 1.375280e+05 | 1.062158e+05 | 3.131219e+04 | NaN | MVGCTGK |
| GSK3A | GSK3A\_HUMAN | Glycogen synthase kinase-3 alpha | 241 | 231 | GSK3A | GSK3A\_HUMAN | Glycogen synthase kinase-3 alpha | 11C(125.0477) | NaN | X | NaN | NaN | NaN | NaN | 1.374067e+05 | 1.374067e+05 | NaN | NaN | SLAYIHSQGVCHR |
| PGM2 | PGM2\_HUMAN | Phosphoglucomutase-2 | 573 | 566 | PGM2 | PGM2\_HUMAN | Phosphoglucomutase-2 | 8C(125.0477) | NaN | X | NaN | NaN | NaN | NaN | 1.372172e+05 | 1.372172e+05 | NaN | NaN | IKYYAELCAPPGNSDPEQLKK |
| XPO4 | XPO4\_HUMAN | Exportin-4 | 977 | 973 | XPO4 | XPO4\_HUMAN | Exportin-4 | 5C(125.0477) | NaN | X | NaN | NaN | NaN | NaN | 1.366422e+05 | 1.366422e+05 | NaN | NaN | FPTLCNQYYK |
| WDR26 | WDR26\_HUMAN | WD repeat-containing protein 26 | 316 | 316 | WDR26 | WDR26\_HUMAN | WD repeat-containing protein 26 | 1C(125.0477) | NaN | X | NaN | NaN | NaN | NaN | 1.364917e+05 | 1.364917e+05 | NaN | NaN | CLYHNTK |
| DHRS11 | DHR11\_HUMAN | Dehydrogenase/reductase SDR family member 11 | 55 | 44 | DHRS11 | DHR11\_HUMAN | Dehydrogenase/reductase SDR family member 11 | 12C(125.0477) | NaN | X | NaN | NaN | NaN | NaN | 1.363703e+05 | 1.212082e+05 | 1.516207e+04 | NaN | TVGNIEELAAECK |
| CAST | ICAL\_HUMAN | Calpastatin | 661, 661, 676, 676 | 643 | CAST | ICAL\_HUMAN | Calpastatin | 19C(125.0477) 19C(15.9949) 34C(125.0477) 34C(15.9949) | NaN | X | X | NaN | NaN | NaN | 1.363692e+05 | 1.363692e+05 | NaN | NaN | KPADDQDPIDALSGDLDSCPSTTETSQNTAKDKCK |
| PGK1 | PGK1\_HUMAN | Phosphoglycerate kinase 1 | 50 | 49 | PGK1 | PGK1\_HUMAN | Phosphoglycerate kinase 1 | 2C(125.0477) | NaN | X | NaN | NaN | NaN | NaN | 1.362377e+05 | NaN | 1.362377e+05 | NaN | FCLDNGAK |
| PURA | PURA\_HUMAN | Transcriptional activator protein Pur-alpha | 272 | 267 | PURA | PURA\_HUMAN | Transcriptional activator protein Pur-alpha | 6C(125.0477) | NaN | X | NaN | NaN | NaN | NaN | 1.361633e+05 | 1.361633e+05 | NaN | NaN | FGHTFCK |
| SORD | DHSO\_HUMAN | Sorbitol dehydrogenase | 106 | 101 | SORD | DHSO\_HUMAN | Sorbitol dehydrogenase | 6C(125.0477) | NaN | X | NaN | NaN | NaN | NaN | 1.360954e+05 | 1.360954e+05 | NaN | NaN | ENDEFCK |
| EIF6 | IF6\_HUMAN | Eukaryotic translation initiation factor 6 | 110 | 101 | EIF6 | IF6\_HUMAN | Eukaryotic translation initiation factor 6 | 10C(125.0477) | NaN | X | NaN | NaN | NaN | NaN | 1.358437e+05 | 1.358437e+05 | NaN | NaN | LSALGNVTTCNDYVALVHPDLDR |
| HK1 | HXK1\_HUMAN | Hexokinase-1 | 628 | 625 | HK1 | HXK1\_HUMAN | Hexokinase-1 | 4C(125.0477) | NaN | X | NaN | NaN | NaN | NaN | 1.357974e+05 | 1.357974e+05 | NaN | NaN | ATDCVGHDVVTLLR |
| ATP2B4 | AT2B4\_HUMAN | Plasma membrane calcium-transporting ATPase 4 | 632 | 625 | ATP2B4 | AT2B4\_HUMAN | Plasma membrane calcium-transporting ATPase 4 | 8C(125.0477) | NaN | X | NaN | NaN | NaN | NaN | 1.355083e+05 | 1.355083e+05 | NaN | NaN | TVIEPMACDGLR |
| CC2D1A | C2D1A\_HUMAN | Coiled-coil and C2 domain-containing protein 1A | 765 | 758 | CC2D1A | C2D1A\_HUMAN | Coiled-coil and C2 domain-containing protein 1A | 8C(125.0477) | NaN | X | NaN | NaN | NaN | NaN | 1.354574e+05 | 1.304123e+05 | 5.045113e+03 | NaN | LDALEIACEVR |
| FHIP2A | FHI2A\_HUMAN | FHF complex subunit HOOK interacting protein 2A | 574 | 568 | FHIP2A | FHI2A\_HUMAN | FHF complex subunit HOOK interacting protein 2A | 7C(125.0477) | NaN | X | NaN | NaN | NaN | NaN | 1.353346e+05 | 1.353346e+05 | NaN | NaN | IVNSFLCLVPDDAK |
| SNX22 | SNX22\_HUMAN | Sorting nexin-22 | 33 | 29 | SNX22 | SNX22\_HUMAN | Sorting nexin-22 | 5C(125.0477) | NaN | X | NaN | NaN | NaN | NaN | 1.353053e+05 | 5.911344e+04 | 7.619188e+04 | NaN | VEVLCSGR |
| ADD1 | ADDA\_HUMAN | Alpha-adducin | 525, 525 | 516 | ADD1 | ADDA\_HUMAN | Alpha-adducin | 10C(15.9949) 10C(31.9988) | NaN | NaN | X | X | NaN | NaN | 1.352515e+05 | NaN | 1.352515e+05 | NaN | TAGPQSQVLCGVVMDR |
| HSP90AA1 | HS90A\_HUMAN | Heat shock protein HSP 90-alpha | 572 | 565 | HSP90AA1 | HS90A\_HUMAN | Heat shock protein HSP 90-alpha | 8C(125.0477) | NaN | X | NaN | NaN | NaN | NaN | 1.351858e+05 | 1.351858e+05 | NaN | NaN | KTKFENLCK |
| AHCY | SAHH\_HUMAN | Adenosylhomocysteinase | 195 | 187 | AHCY | SAHH\_HUMAN | Adenosylhomocysteinase | 9C(125.0477) | NaN | X | NaN | NaN | NaN | NaN | 1.351683e+05 | 1.351683e+05 | NaN | NaN | SKFDNLYGCR |
| SEC14L2 | S14L2\_HUMAN | SEC14-like protein 2 | 355 | 342 | SEC14L2 | S14L2\_HUMAN | SEC14-like protein 2 | 14C(125.0477) | NaN | X | NaN | NaN | NaN | NaN | 1.348803e+05 | 1.348803e+05 | NaN | NaN | YNSHLVPEDGTLTCSDPGIYVLR |
| PPP6C | PPP6\_HUMAN | Serine/threonine-protein phosphatase 6 catalytic subunit | 265 | 265 | PPP6C | PPP6\_HUMAN | Serine/threonine-protein phosphatase 6 catalytic subunit | 1C(125.0477) | NaN | X | NaN | NaN | NaN | NaN | 1.347377e+05 | 1.347377e+05 | NaN | NaN | CGNIASIMVFK |
| MAP2K2 | MP2K2\_HUMAN | Dual specificity mitogen-activated protein kinase kinase 2 | 384 | 372 | MAP2K2 | MP2K2\_HUMAN | Dual specificity mitogen-activated protein kinase kinase 2 | 13C(125.0477) | NaN | X | NaN | NaN | NaN | NaN | 1.346666e+05 | 1.346666e+05 | NaN | NaN | SEVEEVDFAGWLCK |
| UGP2 | UGPA\_HUMAN | UTP--glucose-1-phosphate uridylyltransferase | 123 | 113 | UGP2 | UGPA\_HUMAN | UTP--glucose-1-phosphate uridylyltransferase | 11C(125.0477) | NaN | X | NaN | NaN | NaN | NaN | 1.346282e+05 | 1.346282e+05 | NaN | NaN | LNGGLGTSMGCK |
| GMPR | GMPR1\_HUMAN | GMP reductase 1 | 127 | 125 | GMPR | GMPR1\_HUMAN | GMP reductase 1 | 3C(125.0477) | NaN | X | NaN | NaN | NaN | NaN | 1.342823e+05 | 1.342823e+05 | NaN | NaN | FICLDVANGYSEHFVEFVK |
| PDE12 | PDE12\_HUMAN | 2',5'-phosphodiesterase 12 | 250 | 246 | PDE12 | PDE12\_HUMAN | 2',5'-phosphodiesterase 12 | 5C(125.0477) | NaN | X | NaN | NaN | NaN | NaN | 1.342705e+05 | 1.342705e+05 | NaN | NaN | LKLHCTPGDGQR |
| ANK1 | ANK1\_HUMAN | Ankyrin-1 | 1070 | 1069 | ANK1 | ANK1\_HUMAN | Ankyrin-1 | 2C(125.0477) | NaN | X | NaN | NaN | NaN | NaN | 1.342317e+05 | NaN | NaN | 134231.7300 | LCQDYDTIGPEGGSLK |
| VCP | TERA\_HUMAN | Transitional endoplasmic reticulum ATPase | 535 | 530 | VCP | TERA\_HUMAN | Transitional endoplasmic reticulum ATPase | 6C(125.0477) | NaN | X | NaN | NaN | NaN | NaN | 1.341421e+05 | 4.801547e+04 | 8.612660e+04 | NaN | AIANECQANFISIK |
| PPP2R1A | 2AAA\_HUMAN | Serine/threonine-protein phosphatase 2A 65 kDa regulatory subunit A alpha isoform | 390 | 382 | PPP2R1A | 2AAA\_HUMAN | Serine/threonine-protein phosphatase 2A 65 kDa regulatory subunit A alpha isoform | 9C(125.0477) | NaN | X | NaN | NaN | NaN | NaN | 1.337974e+05 | 1.337974e+05 | NaN | NaN | LNIISNLDCVNEVIGIR |
| S100A6 | S10A6\_HUMAN | Protein S100-A6 | 3 | 1 | S100A6 | S10A6\_HUMAN | Protein S100-A6 | 3C(125.0477) | NaN | X | NaN | NaN | NaN | NaN | 1.335955e+05 | 1.335955e+05 | NaN | NaN | MACPLDQAIGLLVAIFHK |
| KATNAL1 | KATL1\_HUMAN | Katanin p60 ATPase-containing subunit A-like 1 | 265 | 260 | KATNAL1 | KATL1\_HUMAN | Katanin p60 ATPase-containing subunit A-like 1 | 6C(125.0477) | NaN | X | NaN | NaN | NaN | NaN | 1.333623e+05 | 1.333623e+05 | NaN | NaN | AVATECGTTFFNVSSSTLTSK |
| CMBL | CMBL\_HUMAN | Carboxymethylenebutenolidase homolog | 222 | 220 | CMBL | CMBL\_HUMAN | Carboxymethylenebutenolidase homolog | 3C(125.0477) | NaN | X | NaN | NaN | NaN | NaN | 1.332231e+05 | 1.332231e+05 | NaN | NaN | EDCSPADKPYIDEARR |
| KRT1 | K2C1\_HUMAN | Keratin, type II cytoskeletal 1 | 49, 49 | 46 | KRT1 | K2C1\_HUMAN | Keratin, type II cytoskeletal 1 | 4C(125.0477) 4C(31.9988) | NaN | X | NaN | X | NaN | NaN | 1.331325e+05 | 1.331325e+05 | NaN | NaN | FSSCGGGGGSFGAGGGFGSR |
| PARK7 | PARK7\_HUMAN | Parkinson disease protein 7 | 46 | 42 | PARK7 | PARK7\_HUMAN | Parkinson disease protein 7 | 5C(15.9949) | NaN | NaN | X | NaN | NaN | NaN | 1.330085e+05 | NaN | 1.330085e+05 | NaN | DPVQCSRDVVICPDASLEDAKK |
| PFAS | PUR4\_HUMAN | Phosphoribosylformylglycinamidine synthase | 1027 | 1027 | PFAS | PUR4\_HUMAN | Phosphoribosylformylglycinamidine synthase | 1C(125.0477) | NaN | X | NaN | NaN | NaN | NaN | 1.328955e+05 | 1.328955e+05 | NaN | NaN | CVAEEER |
| MARK3 | MARK3\_HUMAN | MAP/microtubule affinity-regulating kinase 3 | 688 | 682 | MARK3 | MARK3\_HUMAN | MAP/microtubule affinity-regulating kinase 3 | 7C(125.0477) | NaN | X | NaN | NaN | NaN | NaN | 1.326664e+05 | 9.476018e+04 | 3.790622e+04 | NaN | VLDANNCDYEQR |
| AP1B1 | AP1B1\_HUMAN | AP-1 complex subunit beta-1 | 866 | 865 | AP1B1 | AP1B1\_HUMAN | AP-1 complex subunit beta-1 | 2C(125.0477) | NaN | X | NaN | NaN | NaN | NaN | 1.326348e+05 | NaN | 1.326348e+05 | NaN | DCPLNAEAASSK |
| PKLR | KPYR\_HUMAN | Pyruvate kinase PKLR | 401 | 386 | PKLR | KPYR\_HUMAN | Pyruvate kinase PKLR | 16C(125.0477) | NaN | X | NaN | NaN | NaN | NaN | 1.326283e+05 | 1.326283e+05 | NaN | NaN | AETSDVANAVLDGADCIMLSGETAK |
| STK10 | STK10\_HUMAN | Serine/threonine-protein kinase 10 | 947 | 941 | STK10 | STK10\_HUMAN | Serine/threonine-protein kinase 10 | 7C(125.0477) | NaN | X | NaN | NaN | NaN | NaN | 1.325656e+05 | 1.157302e+05 | 1.683542e+04 | NaN | LSEEAECPNPSTPSK |
| SRXN1 | SRXN1\_HUMAN | Sulfiredoxin-1 | 99 | 86 | SRXN1 | SRXN1\_HUMAN | Sulfiredoxin-1 | 14C(125.0477) | NaN | X | NaN | NaN | NaN | NaN | 1.318862e+05 | 1.318862e+05 | NaN | NaN | GAQGGDYFYSFGGCHR |
| CORO7 | CORO7\_HUMAN | Coronin-7 | 695 | 695 | CORO7 | CORO7\_HUMAN | Coronin-7 | 1C(125.0477) | NaN | X | NaN | NaN | NaN | NaN | 1.317604e+05 | 1.317604e+05 | NaN | NaN | CLLVSGFDSQSER |
| TAGLN2 | TAGL2\_HUMAN | Transgelin-2 | 124 | 121 | TAGLN2 | TAGL2\_HUMAN | Transgelin-2 | 4C(125.0477) | NaN | X | NaN | NaN | NaN | NaN | 1.315435e+05 | 1.150874e+05 | 1.645604e+04 | NaN | NMACVQR |
| USP24 | UBP24\_HUMAN | Ubiquitin carboxyl-terminal hydrolase 24 | 1859 | 1848 | USP24 | UBP24\_HUMAN | Ubiquitin carboxyl-terminal hydrolase 24 | 12C(125.0477) | NaN | X | NaN | NaN | NaN | NaN | 1.314748e+05 | 1.314748e+05 | NaN | NaN | GEVLEGSNAYYCEK |
| UBA3 | UBA3\_HUMAN | NEDD8-activating enzyme E1 catalytic subunit | 70 | 48 | UBA3 | UBA3\_HUMAN | NEDD8-activating enzyme E1 catalytic subunit | 23C(125.0477) | NaN | X | NaN | NaN | NaN | NaN | 1.314268e+05 | 1.314268e+05 | NaN | NaN | SGPFTHPDFEPSTESLQFLLDTCK |
| XPO1 | XPO1\_HUMAN | Exportin-1 | 34 | 23 | XPO1 | XPO1\_HUMAN | Exportin-1 | 12C(125.0477) | NaN | X | NaN | NaN | NaN | NaN | 1.314220e+05 | 1.314220e+05 | NaN | NaN | LDINLLDNVVNCLYHGEGAQQR |
| XPO1 | XPO1\_HUMAN | Exportin-1 | 1070 | 1053 | XPO1 | XPO1\_HUMAN | Exportin-1 | 18C(125.0477) | NaN | X | NaN | NaN | NaN | NaN | 1.309801e+05 | 1.309801e+05 | NaN | NaN | QMSVPGIFNPHEIPEEMCD |
| GPHN | GEPH\_HUMAN | Gephyrin | 469 | 466 | GPHN | GEPH\_HUMAN | Gephyrin | 4C(125.0477) | NaN | X | NaN | NaN | NaN | NaN | 1.309634e+05 | 1.309634e+05 | NaN | NaN | RGECVLAK |
| FBXO7 | FBX7\_HUMAN | F-box only protein 7 | 359 | 351 | FBXO7 | FBX7\_HUMAN | F-box only protein 7 | 9C(125.0477) | NaN | X | NaN | NaN | NaN | NaN | 1.307312e+05 | NaN | 1.307312e+05 | NaN | SVLSLSAVCR |
| ZZEF1 | ZZEF1\_HUMAN | Zinc finger ZZ-type and EF-hand domain-containing protein 1 | 69 | 58 | ZZEF1 | ZZEF1\_HUMAN | Zinc finger ZZ-type and EF-hand domain-containing protein 1 | 12C(125.0477) | NaN | X | NaN | NaN | NaN | NaN | 1.306960e+05 | 1.306960e+05 | NaN | NaN | EAAAALLPTPPCESLVSR |
| PPM1B | PPM1B\_HUMAN | Protein phosphatase 1B | 71 | 67 | PPM1B | PPM1B\_HUMAN | Protein phosphatase 1B | 5C(125.0477) | NaN | X | NaN | NaN | NaN | NaN | 1.305026e+05 | 1.305026e+05 | NaN | NaN | VANYCSTHLLEHITTNEDFR |
| TLN2 | TLN2\_HUMAN | Talin-2 | 1355 | 1343 | TLN2 | TLN2\_HUMAN | Talin-2 | 13C(125.0477) | NaN | X | NaN | NaN | NaN | NaN | 1.303418e+05 | 1.303418e+05 | NaN | NaN | AVTESINQLITLCTQQAPGQK |
| KRT5 | K2C5\_HUMAN | Keratin, type II cytoskeletal 5 | 55 | 49 | KRT5 | K2C5\_HUMAN | Keratin, type II cytoskeletal 5 | 7C(125.0477) | NaN | X | NaN | NaN | NaN | NaN | 1.301631e+05 | NaN | 1.301631e+05 | NaN | VSLAGACGVGGYGSR |
| CUL4A | CUL4A\_HUMAN | Cullin-4A | 241 | 235 | CUL4A | CUL4A\_HUMAN | Cullin-4A | 7C(125.0477) | NaN | X | NaN | NaN | NaN | NaN | 1.296002e+05 | 1.296002e+05 | NaN | NaN | FLEETNCLYAAEGQR |
| VPS13A | VP13A\_HUMAN | Vacuolar protein sorting-associated protein 13A | 1848 | 1848 | VPS13A | VP13A\_HUMAN | Vacuolar protein sorting-associated protein 13A | 1C(125.0477) | NaN | X | NaN | NaN | NaN | NaN | 1.295485e+05 | 1.295485e+05 | NaN | NaN | CGLVMLNNLVK |
| PIEZO1 | PIEZ1\_HUMAN | Piezo-type mechanosensitive ion channel component 1 | 50, 50 | 50 | PIEZO1 | PIEZ1\_HUMAN | Piezo-type mechanosensitive ion channel component 1 | 1C(125.0477) 1C(31.9988) | NaN | X | NaN | X | NaN | NaN | 1.295219e+05 | NaN | 1.295219e+05 | NaN | CGLQGHTGR |
| CCT3 | TCPG\_HUMAN | T-complex protein 1 subunit gamma | 173 | 168 | CCT3 | TCPG\_HUMAN | T-complex protein 1 subunit gamma | 6C(125.0477) | NaN | X | NaN | NaN | NaN | NaN | 1.295134e+05 | 1.295134e+05 | NaN | NaN | WSSLACNIALDAVK |
| ANK1 | ANK1\_HUMAN | Ankyrin-1 | 531 | 513 | ANK1 | ANK1\_HUMAN | Ankyrin-1 | 19C(125.0477) | NaN | X | NaN | NaN | NaN | NaN | 1.291248e+05 | 1.291248e+05 | NaN | NaN | EGHVETVLALLEKEASQACMTK |
| ANKRD28 | ANR28\_HUMAN | Serine/threonine-protein phosphatase 6 regulatory ankyrin repeat subunit A | 201 | 191 | ANKRD28 | ANR28\_HUMAN | Serine/threonine-protein phosphatase 6 regulatory ankyrin repeat subunit A | 11C(125.0477) | NaN | X | NaN | NaN | NaN | NaN | 1.291159e+05 | 1.291159e+05 | NaN | NaN | LLVSHGAEVTCK |
| CDSN | CDSN\_HUMAN | Corneodesmosin | 391 | 389 | CDSN | CDSN\_HUMAN | Corneodesmosin | 3C(125.0477) | NaN | X | NaN | NaN | NaN | NaN | 1.288917e+05 | NaN | 2.768643e+04 | 101205.2900 | GPCSPSSSR |
| VPS28 | VPS28\_HUMAN | Vacuolar protein sorting-associated protein 28 homolog | 96 | 83 | VPS28 | VPS28\_HUMAN | Vacuolar protein sorting-associated protein 28 homolog | 14C(125.0477) | NaN | X | NaN | NaN | NaN | NaN | 1.288686e+05 | 1.220406e+05 | 6.827991e+03 | NaN | QVQGSEISSIDEFCR |
| ALB | ALBU\_HUMAN | Albumin | 591, 591, 591 | 585 | ALB | ALBU\_HUMAN | Albumin | 7C(125.0477) 7C(31.9988) 7C(57.0215) | X | X | NaN | X | NaN | NaN | 1.288144e+05 | 7.409142e+04 | 5.472302e+04 | NaN | ADDKETCFAEEGKK |
| AMPD3 | AMPD3\_HUMAN | AMP deaminase 3 | 47 | 36 | AMPD3 | AMPD3\_HUMAN | AMP deaminase 3 | 12C(125.0477) | NaN | X | NaN | NaN | NaN | NaN | 1.287580e+05 | 1.287580e+05 | NaN | NaN | DALSLFTVPEDCPIGQK |
| EIF2S1 | IF2A\_HUMAN | Eukaryotic translation initiation factor 2 subunit 1 | 98 | 98 | EIF2S1 | IF2A\_HUMAN | Eukaryotic translation initiation factor 2 subunit 1 | 1C(125.0477) | NaN | X | NaN | NaN | NaN | NaN | 1.283948e+05 | 1.283948e+05 | NaN | NaN | CEDKFTK |
| PPP6R1 | PP6R1\_HUMAN | Serine/threonine-protein phosphatase 6 regulatory subunit 1 | 216 | 195 | PPP6R1 | PP6R1\_HUMAN | Serine/threonine-protein phosphatase 6 regulatory subunit 1 | 22C(125.0477) | NaN | X | NaN | NaN | NaN | NaN | 1.283698e+05 | 1.283698e+05 | NaN | NaN | LIEQIHPSKDENQHSNASQSLCDIIR |
| FAS | TNR6\_HUMAN | Tumor necrosis factor receptor superfamily member 6 | 157 | 156 | FAS | TNR6\_HUMAN | Tumor necrosis factor receptor superfamily member 6 | 2C(125.0477) | NaN | X | NaN | NaN | NaN | NaN | 1.283085e+05 | NaN | 9.262038e+04 | 35688.1300 | ECTLTSNTK |
| ERP44 | ERP44\_HUMAN | Endoplasmic reticulum resident protein 44 | 301 | 292 | ERP44 | ERP44\_HUMAN | Endoplasmic reticulum resident protein 44 | 10C(125.0477) | NaN | X | NaN | NaN | NaN | NaN | 1.278127e+05 | NaN | 1.278127e+05 | NaN | GTINFLHADCDK |
| SOD1 | SODC\_HUMAN | Superoxide dismutase [Cu-Zn] | 58, 58, 58 | 38 | SOD1 | SODC\_HUMAN | Superoxide dismutase [Cu-Zn] | 21C(125.0477) 21C(31.9988) 21C(57.0215) | X | X | NaN | X | NaN | NaN | 1.275841e+05 | 1.275841e+05 | NaN | NaN | GLTEGLHGFHVHEFGDNTAGCTSAGPHFNPLSR |
| CASP6 | CASP6\_HUMAN | Caspase-6 | 148 | 148 | CASP6 | CASP6\_HUMAN | Caspase-6 | 1C(125.0477) | NaN | X | NaN | NaN | NaN | NaN | 1.273542e+05 | 1.273542e+05 | NaN | NaN | CHSLVGKPK |
| PTPN12 | PTN12\_HUMAN | Tyrosine-protein phosphatase non-receptor type 12 | 164, 164 | 162 | PTPN12 | PTN12\_HUMAN | Tyrosine-protein phosphatase non-receptor type 12 | 3C(125.0477) 3C(57.0215) | X | X | NaN | NaN | NaN | NaN | 1.268254e+05 | 1.268254e+05 | NaN | NaN | ISCEDEQAR |
| PDCD6IP | PDC6I\_HUMAN | Programmed cell death 6-interacting protein | 524 | 518 | PDCD6IP | PDC6I\_HUMAN | Programmed cell death 6-interacting protein | 7C(125.0477) | NaN | X | NaN | NaN | NaN | NaN | 1.266534e+05 | 1.266534e+05 | NaN | NaN | DTIVLLCKPEPELNAAIPSANPAK |
| CAT | CATA\_HUMAN | Catalase | 460 | 459 | CAT | CATA\_HUMAN | Catalase | 2C(125.0477) | NaN | X | NaN | NaN | NaN | NaN | 1.266375e+05 | 1.266375e+05 | NaN | NaN | LCENIAGHLKDAQIFIQKK |
| PPP2R1A | 2AAA\_HUMAN | Serine/threonine-protein phosphatase 2A 65 kDa regulatory subunit A alpha isoform | 174 | 172 | PPP2R1A | 2AAA\_HUMAN | Serine/threonine-protein phosphatase 2A 65 kDa regulatory subunit A alpha isoform | 3C(125.0477) | NaN | X | NaN | NaN | NaN | NaN | 1.264646e+05 | 9.945531e+04 | 2.700932e+04 | NaN | NLCSDDTPMVR |
| SERPINA1 | A1AT\_HUMAN | Alpha-1-antitrypsin | 256 | 247 | SERPINA1 | A1AT\_HUMAN | Alpha-1-antitrypsin | 10C(57.0215) | X | NaN | NaN | NaN | NaN | NaN | 1.259771e+05 | 1.259771e+05 | NaN | NaN | RLGMFNIQHCKK |
| PBK | TOPK\_HUMAN | Lymphokine-activated killer T-cell-originated protein kinase | 22 | 19 | PBK | TOPK\_HUMAN | Lymphokine-activated killer T-cell-originated protein kinase | 4C(125.0477) | NaN | X | NaN | NaN | NaN | NaN | 1.256016e+05 | 1.256016e+05 | NaN | NaN | SVLCSTPTINIPASPFMQK |
| SIRT2 | SIR2\_HUMAN | NAD-dependent protein deacetylase sirtuin-2 | 81 | 79 | SIRT2 | SIR2\_HUMAN | NAD-dependent protein deacetylase sirtuin-2 | 3C(125.0477) | NaN | X | NaN | NaN | NaN | NaN | 1.254039e+05 | 1.254039e+05 | NaN | NaN | VICLVGAGISTSAGIPDFR |
| IGKC | IGKC\_HUMAN | Immunoglobulin kappa constant | 87 | 84 | IGKC | IGKC\_HUMAN | Immunoglobulin kappa constant | 4C(57.0215) | X | NaN | NaN | NaN | NaN | NaN | 1.253717e+05 | NaN | 1.253717e+05 | NaN | VYACEVTHQGLSSPVTK |
| LRRFIP2 | LRRF2\_HUMAN | Leucine-rich repeat flightless-interacting protein 2 | 223 | 219 | LRRFIP2 | LRRF2\_HUMAN | Leucine-rich repeat flightless-interacting protein 2 | 5C(-34.0809) | NaN | NaN | NaN | NaN | X | NaN | 1.252789e+05 | 1.252789e+05 | NaN | NaN | TPSECSYYSSR |
| NAPA | SNAA\_HUMAN | Alpha-soluble NSF attachment protein | 84 | 79 | NAPA | SNAA\_HUMAN | Alpha-soluble NSF attachment protein | 6C(125.0477) | NaN | X | NaN | NaN | NaN | NaN | 1.249321e+05 | 1.249321e+05 | NaN | NaN | HDAATCFVDAGNAFKK |
| GMPPB | GMPPB\_HUMAN | Mannose-1-phosphate guanyltransferase beta | 149 | 144 | GMPPB | GMPPB\_HUMAN | Mannose-1-phosphate guanyltransferase beta | 6C(125.0477) | NaN | X | NaN | NaN | NaN | NaN | 1.249063e+05 | NaN | 1.249063e+05 | NaN | YGVVVCEADTGR |
| KRT84 | KRT84\_HUMAN | Keratin, type II cuticular Hb4 | 432 | 427 | KRT84 | KRT84\_HUMAN | Keratin, type II cuticular Hb4 | 6C(57.0215) | X | NaN | NaN | NaN | NaN | NaN | 1.248933e+05 | NaN | 1.248933e+05 | NaN | LADLECALQQAK |
| WDR1 | WDR1\_HUMAN | WD repeat-containing protein 1 | 438 | 437 | WDR1 | WDR1\_HUMAN | WD repeat-containing protein 1 | 2C(125.0477) | NaN | X | NaN | NaN | NaN | NaN | 1.245195e+05 | 1.245195e+05 | NaN | NaN | KCFSIDNPGYEPEVVAVHPGGDTVAIGGVDGNVR |
| TARS1 | SYTC\_HUMAN | Threonine--tRNA ligase 1, cytoplasmic | 630 | 620 | TARS1 | SYTC\_HUMAN | Threonine--tRNA ligase 1, cytoplasmic | 11C(125.0477) | NaN | X | NaN | NaN | NaN | NaN | 1.244280e+05 | 1.244280e+05 | NaN | NaN | QVMVVPVGPTCDEYAQK |
| STK11 | STK11\_HUMAN | Serine/threonine-protein kinase STK11 | 278 | 270 | STK11 | STK11\_HUMAN | Serine/threonine-protein kinase STK11 | 9C(125.0477) | NaN | X | NaN | NaN | NaN | NaN | 1.238316e+05 | 1.238316e+05 | NaN | NaN | GSYAIPGDCGPPLSDLLK |
| SPTB | SPTB1\_HUMAN | Spectrin beta chain, erythrocytic | 183 | 174 | SPTB | SPTB1\_HUMAN | Spectrin beta chain, erythrocytic | 10C(125.0477) | NaN | X | NaN | NaN | NaN | NaN | 1.236796e+05 | 1.236796e+05 | NaN | NaN | SAKDALLLWCQMK |
| SPTA1 | SPTA1\_HUMAN | Spectrin alpha chain, erythrocytic 1 | 2101, 2101 | 2101 | SPTA1 | SPTA1\_HUMAN | Spectrin alpha chain, erythrocytic 1 | 1C(125.0477) 1C(31.9988) | NaN | X | NaN | X | NaN | NaN | 1.236387e+05 | NaN | NaN | 123638.6640 | CLLELDQQIK |
| CAND1 | CAND1\_HUMAN | Cullin-associated NEDD8-dissociated protein 1 | 356 | 356 | CAND1 | CAND1\_HUMAN | Cullin-associated NEDD8-dissociated protein 1 | 1C(125.0477) | NaN | X | NaN | NaN | NaN | NaN | 1.235022e+05 | NaN | 1.235022e+05 | NaN | CLDAVVSTR |
| CBR1 | CBR1\_HUMAN | Carbonyl reductase [NADPH] 1 | 150 | 149 | CBR1 | CBR1\_HUMAN | Carbonyl reductase [NADPH] 1 | 2C(125.0477) | NaN | X | NaN | NaN | NaN | NaN | 1.234815e+05 | 7.030864e+04 | 5.317291e+04 | NaN | SCSPELQQK |
| USP14 | UBP14\_HUMAN | Ubiquitin carboxyl-terminal hydrolase 14 | 203 | 190 | USP14 | UBP14\_HUMAN | Ubiquitin carboxyl-terminal hydrolase 14 | 14C(125.0477) | NaN | X | NaN | NaN | NaN | NaN | 1.231170e+05 | 1.231170e+05 | NaN | NaN | GEQGQYLQQDANECWIQMMR |
| ALB | ALBU\_HUMAN | Albumin | 416 | 414 | ALB | ALBU\_HUMAN | Albumin | 3C(125.0477) | NaN | X | NaN | NaN | NaN | NaN | 1.231162e+05 | 1.231162e+05 | NaN | NaN | QNCELFEQLGEYK |
| LCN2 | NGAL\_HUMAN | Neutrophil gelatinase-associated lipocalin | 195 | 178 | LCN2 | NGAL\_HUMAN | Neutrophil gelatinase-associated lipocalin | 18C(57.0215) | X | NaN | NaN | NaN | NaN | NaN | 1.229112e+05 | 1.189710e+05 | 3.940203e+03 | NaN | SLGLPENHIVFPVPIDQCIDG |
| RANGAP1 | RAGP1\_HUMAN | Ran GTPase-activating protein 1 | 573 | 554 | RANGAP1 | RAGP1\_HUMAN | Ran GTPase-activating protein 1 | 20C(125.0477) | NaN | X | NaN | NaN | NaN | NaN | 1.228293e+05 | 1.228293e+05 | NaN | NaN | ALAPLLLAFVTKPNSALESCSFAR |
| HSPH1 | HS105\_HUMAN | Heat shock protein 105 kDa | 140 | 135 | HSPH1 | HS105\_HUMAN | Heat shock protein 105 kDa | 6C(125.0477) | NaN | X | NaN | NaN | NaN | NaN | 1.227291e+05 | 1.227291e+05 | NaN | NaN | KPVTDCVISVPSFFTDAER |
| GSTO1 | GSTO1\_HUMAN | Glutathione S-transferase omega-1 | 192 | 184 | GSTO1 | GSTO1\_HUMAN | Glutathione S-transferase omega-1 | 9C(125.0477) | NaN | X | NaN | NaN | NaN | NaN | 1.223420e+05 | 1.223420e+05 | NaN | NaN | LEAMKLNECVDHTPK |
| DENND4C | DEN4C\_HUMAN | DENN domain-containing protein 4C | 88 | 82 | DENND4C | DEN4C\_HUMAN | DENN domain-containing protein 4C | 7C(125.0477) | NaN | X | NaN | NaN | NaN | NaN | 1.223102e+05 | 1.223102e+05 | NaN | NaN | SPELFLCYKR |
| ACTBL2 | ACTBL\_HUMAN | Beta-actin-like protein 2 | 53, 53, 53 | 41 | ACTBL2 | ACTBL\_HUMAN | Beta-actin-like protein 2 | 13C(125.0477) 13C(15.9949) 13C(31.9988) | NaN | X | X | X | NaN | NaN | 1.220963e+05 | 1.220963e+05 | NaN | NaN | HQGVMVGMGQKDCYVGDEAQSKR |
| ROCK2 | ROCK2\_HUMAN | Rho-associated protein kinase 2 | 330 | 327 | ROCK2 | ROCK2\_HUMAN | Rho-associated protein kinase 2 | 4C(125.0477) | NaN | X | NaN | NaN | NaN | NaN | 1.220625e+05 | 1.220625e+05 | NaN | NaN | NLICAFLTDR |
| GRHPR | GRHPR\_HUMAN | Glyoxylate reductase/hydroxypyruvate reductase | 29 | 26 | GRHPR | GRHPR\_HUMAN | Glyoxylate reductase/hydroxypyruvate reductase | 4C(125.0477) | NaN | X | NaN | NaN | NaN | NaN | 1.219314e+05 | 1.219314e+05 | NaN | NaN | AADCEVEQWDSDEPIPAK |
| ALB | ALBU\_HUMAN | Albumin | 224 | 224 | ALB | ALBU\_HUMAN | Albumin | 1C(57.0215) | X | NaN | NaN | NaN | NaN | NaN | 1.216613e+05 | 8.257631e+04 | 2.231385e+04 | 16771.1330 | CASLQKFGER |
| GPX4 | GPX4\_HUMAN | Phospholipid hydroperoxide glutathione peroxidase | 134 | 133 | GPX4 | GPX4\_HUMAN | Phospholipid hydroperoxide glutathione peroxidase | 2C(125.0477) | NaN | X | NaN | NaN | NaN | NaN | 1.215595e+05 | 1.215595e+05 | NaN | NaN | ICVNGDDAHPLWK |
| DCTN1 | DCTN1\_HUMAN | Dynactin subunit 1 | 81 | 77 | DCTN1 | DCTN1\_HUMAN | Dynactin subunit 1 | 5C(125.0477) | NaN | X | NaN | NaN | NaN | NaN | 1.215139e+05 | 1.215139e+05 | NaN | NaN | KYFTCDEGHGIFVR |
| NSF | NSF\_HUMAN | Vesicle-fusing ATPase | 599 | 595 | NSF | NSF\_HUMAN | Vesicle-fusing ATPase | 5C(125.0477) | NaN | X | NaN | NaN | NaN | NaN | 1.213955e+05 | 1.034116e+05 | 1.798388e+04 | NaN | SQLSCVVVDDIER |
| GART | PUR2\_HUMAN | Trifunctional purine biosynthetic protein adenosine-3 | 62 | 45 | GART | PUR2\_HUMAN | Trifunctional purine biosynthetic protein adenosine-3 | 18C(125.0477) | NaN | X | NaN | NaN | NaN | NaN | 1.211698e+05 | 1.211698e+05 | NaN | NaN | ISNTAISISDHTALAQFCK |
| UBE4A | UBE4A\_HUMAN | Ubiquitin conjugation factor E4 A | 465 | 457 | UBE4A | UBE4A\_HUMAN | Ubiquitin conjugation factor E4 A | 9C(125.0477) | NaN | X | NaN | NaN | NaN | NaN | 1.211062e+05 | 1.211062e+05 | NaN | NaN | LLTFNPTYCALK |
| DNPEP | DNPEP\_HUMAN | Aspartyl aminopeptidase | 337 | 332 | DNPEP | DNPEP\_HUMAN | Aspartyl aminopeptidase | 6C(125.0477) | NaN | X | NaN | NaN | NaN | NaN | 1.211029e+05 | 1.211029e+05 | NaN | NaN | RISASCQHPTAFEEAIPK |
| GMPR | GMPR1\_HUMAN | GMP reductase 1 | 186 | 178 | GMPR | GMPR1\_HUMAN | GMP reductase 1 | 9C(125.0477) | NaN | X | NaN | NaN | NaN | NaN | 1.210020e+05 | 1.210020e+05 | NaN | NaN | VGVGPGSVCTTR |
| UBA3 | UBA3\_HUMAN | NEDD8-activating enzyme E1 catalytic subunit | 249 | 245 | UBA3 | UBA3\_HUMAN | NEDD8-activating enzyme E1 catalytic subunit | 5C(125.0477) | NaN | X | NaN | NaN | NaN | NaN | 1.209420e+05 | 1.209420e+05 | NaN | NaN | LPEHCIEYVR |
| EPS15 | EPS15\_HUMAN | Epidermal growth factor receptor substrate 15 | 419 | 419 | EPS15 | EPS15\_HUMAN | Epidermal growth factor receptor substrate 15 | 1C(125.0477) | NaN | X | NaN | NaN | NaN | NaN | 1.208515e+05 | 6.704831e+04 | 5.380320e+04 | NaN | CAEEAQLISSLK |
| ALB | ALBU\_HUMAN | Albumin | 99 | 89 | ALB | ALBU\_HUMAN | Albumin | 11C(57.0215) | X | NaN | NaN | NaN | NaN | NaN | 1.203767e+05 | 1.203767e+05 | NaN | NaN | SLHTLFGDKLCTVATLR |
| CA3 | CAH3\_HUMAN | Carbonic anhydrase 3 | 187, 182 | 178 | CA3 | CAH3\_HUMAN | Carbonic anhydrase 3 | 10C(125.0477) 5C(31.9988) | NaN | X | NaN | X | NaN | NaN | 1.202799e+05 | 1.202799e+05 | NaN | NaN | FDPSCLFPACR |
| ANKRD28 | ANR28\_HUMAN | Serine/threonine-protein phosphatase 6 regulatory ankyrin repeat subunit A | 966 | 949 | ANKRD28 | ANR28\_HUMAN | Serine/threonine-protein phosphatase 6 regulatory ankyrin repeat subunit A | 18C(125.0477) | NaN | X | NaN | NaN | NaN | NaN | 1.201957e+05 | 1.201957e+05 | NaN | NaN | GASVLAVDENGYTPALACAPNK |
| CNP | CN37\_HUMAN | 2',3'-cyclic-nucleotide 3'-phosphodiesterase | 111, 111 | 103 | CNP | CN37\_HUMAN | 2',3'-cyclic-nucleotide 3'-phosphodiesterase | 9C(125.0477) 9C(15.9949) | NaN | X | X | NaN | NaN | NaN | 1.199927e+05 | NaN | 1.199927e+05 | NaN | LDEDLAAYCR |
| ADD1 | ADDA\_HUMAN | Alpha-adducin | 430 | 407 | ADD1 | ADDA\_HUMAN | Alpha-adducin | 24C(125.0477) | NaN | X | NaN | NaN | NaN | NaN | 1.198332e+05 | 1.198332e+05 | NaN | NaN | YSDVEVPASVTGYSFASDGDSGTCSPLR |
| PSMD5 | PSMD5\_HUMAN | 26S proteasome non-ATPase regulatory subunit 5 | 227 | 225 | PSMD5 | PSMD5\_HUMAN | 26S proteasome non-ATPase regulatory subunit 5 | 3C(125.0477) | NaN | X | NaN | NaN | NaN | NaN | 1.195887e+05 | 1.195887e+05 | NaN | NaN | ATCIEMVTSLAYTHHGR |
| SCYL1 | SCYL1\_HUMAN | N-terminal kinase-like protein | 88 | 88 | SCYL1 | SCYL1\_HUMAN | N-terminal kinase-like protein | 1C(125.0477) | NaN | X | NaN | NaN | NaN | NaN | 1.195694e+05 | 1.195694e+05 | NaN | NaN | CLHVVTEAVTPLGIYLK |
| DSP | DESP\_HUMAN | Desmoplakin | 1805 | 1803 | DSP | DESP\_HUMAN | Desmoplakin | 3C(125.0477) | NaN | X | NaN | NaN | NaN | NaN | 1.195352e+05 | NaN | 1.006726e+05 | 18862.5500 | NQCTQVVQER |
| CA3 | CAH3\_HUMAN | Carbonic anhydrase 3 | 66 | 58 | CA3 | CAH3\_HUMAN | Carbonic anhydrase 3 | 9C(125.0477) | NaN | X | NaN | NaN | NaN | NaN | 1.195204e+05 | 1.195204e+05 | NaN | NaN | TILNNGKTCR |
| CRYGD | CRGD\_HUMAN | Gamma-crystallin D | 19 | 16 | CRYGD | CRGD\_HUMAN | Gamma-crystallin D | 4C(125.0477) | NaN | X | NaN | NaN | NaN | NaN | 1.193172e+05 | 1.193172e+05 | NaN | NaN | HYECSSDHPNLQPYLSR |
| SPTA1 | SPTA1\_HUMAN | Spectrin alpha chain, erythrocytic 1 | 1203, 1203 | 1203 | SPTA1 | SPTA1\_HUMAN | Spectrin alpha chain, erythrocytic 1 | 1C(125.0477) 1C(31.9988) | NaN | X | NaN | X | NaN | NaN | 1.192470e+05 | NaN | 1.192470e+05 | NaN | CQALSAADPGSDLFSVQALQR |
| RAD23B | RD23B\_HUMAN | UV excision repair protein RAD23 homolog B | 390 | 375 | RAD23B | RD23B\_HUMAN | UV excision repair protein RAD23 homolog B | 16C(125.0477) | NaN | X | NaN | NaN | NaN | NaN | 1.191526e+05 | 1.191526e+05 | NaN | NaN | ALGFPEGLVIQAYFACEK |
| ALAD | HEM2\_HUMAN | Delta-aminolevulinic acid dehydratase | 75 | 75 | ALAD | HEM2\_HUMAN | Delta-aminolevulinic acid dehydratase | 1C(125.0477) | NaN | X | NaN | NaN | NaN | NaN | 1.191212e+05 | 1.191212e+05 | NaN | NaN | CVLIFGVPSRVPKDER |
| PIN1 | PIN1\_HUMAN | Peptidyl-prolyl cis-trans isomerase NIMA-interacting 1 | 113 | 98 | PIN1 | PIN1\_HUMAN | Peptidyl-prolyl cis-trans isomerase NIMA-interacting 1 | 16C(125.0477) | NaN | X | NaN | NaN | NaN | NaN | 1.187583e+05 | 1.098633e+05 | 8.895033e+03 | NaN | SGEEDFESLASQFSDCSSAK |
| DNM2 | DYN2\_HUMAN | Dynamin-2 | 86 | 78 | DNM2 | DYN2\_HUMAN | Dynamin-2 | 9C(125.0477) | NaN | X | NaN | NaN | NaN | NaN | 1.186950e+05 | 1.186950e+05 | NaN | NaN | TEHAEFLHCK |
| ATP6V1B2 | VATB2\_HUMAN | V-type proton ATPase subunit B, brain isoform | 289 | 277 | ATP6V1B2 | VATB2\_HUMAN | V-type proton ATPase subunit B, brain isoform | 13C(125.0477) | NaN | X | NaN | NaN | NaN | NaN | 1.185593e+05 | 1.185593e+05 | NaN | NaN | LALTTAEFLAYQCEK |
| CAD | PYR1\_HUMAN | CAD protein | 736 | 720 | CAD | PYR1\_HUMAN | CAD protein | 17C(125.0477) | NaN | X | NaN | NaN | NaN | NaN | 1.184614e+05 | 1.184614e+05 | NaN | NaN | NSVTGGTAAFEPSVDYCVVK |
| RBBP7 | RBBP7\_HUMAN | Histone-binding protein RBBP7 | 166 | 156 | RBBP7 | RBBP7\_HUMAN | Histone-binding protein RBBP7 | 11C(125.0477) | NaN | X | NaN | NaN | NaN | NaN | 1.182403e+05 | 1.182403e+05 | NaN | NaN | HPAKPDPSGECNPDLR |
| RAB2B | RAB2B\_HUMAN | Ras-related protein Rab-2B | 21 | 20 | RAB2B | RAB2B\_HUMAN | Ras-related protein Rab-2B | 2C(125.0477) | NaN | X | NaN | NaN | NaN | NaN | 1.181985e+05 | 1.181985e+05 | NaN | NaN | SCLLLQFTDKR |
| DNPEP | DNPEP\_HUMAN | Aspartyl aminopeptidase | 337 | 333 | DNPEP | DNPEP\_HUMAN | Aspartyl aminopeptidase | 5C(125.0477) | NaN | X | NaN | NaN | NaN | NaN | 1.181574e+05 | 1.181574e+05 | NaN | NaN | ISASCQHPTAFEEAIPK |
| IPO4 | IPO4\_HUMAN | Importin-4 | 1027 | 1026 | IPO4 | IPO4\_HUMAN | Importin-4 | 2C(125.0477) | NaN | X | NaN | NaN | NaN | NaN | 1.178051e+05 | 1.178051e+05 | NaN | NaN | ICSLILADNK |
| TUBB4B | TBB4B\_HUMAN | Tubulin beta-4B chain | 12 | 1 | TUBB4B | TBB4B\_HUMAN | Tubulin beta-4B chain | 12C(125.0477) | NaN | X | NaN | NaN | NaN | NaN | 1.177380e+05 | 1.177380e+05 | NaN | NaN | MREIVHLQAGQCGNQIGAK |
| RAB21 | RAB21\_HUMAN | Ras-related protein Rab-21 | 177 | 168 | RAB21 | RAB21\_HUMAN | Ras-related protein Rab-21 | 10C(125.0477) | NaN | X | NaN | NaN | NaN | NaN | 1.176756e+05 | 1.176756e+05 | NaN | NaN | GIEELFLDLCK |
| CFL2 | COF2\_HUMAN | Cofilin-2 | 39 | 35 | CFL2 | COF2\_HUMAN | Cofilin-2 | 5C(125.0477) | NaN | X | NaN | NaN | NaN | NaN | 1.174098e+05 | 1.174098e+05 | NaN | NaN | AVLFCLSDDKR |
| SPRR3 | SPRR3\_HUMAN | Small proline-rich protein 3 | 58 | 53 | SPRR3 | SPRR3\_HUMAN | Small proline-rich protein 3 | 6C(125.0477) | NaN | X | NaN | NaN | NaN | NaN | 1.170785e+05 | NaN | 8.009647e+04 | 36982.0400 | IPEPGCTK |
| PPP1CA | PP1A\_HUMAN | Serine/threonine-protein phosphatase PP1-alpha catalytic subunit | 127 | 123 | PPP1CA | PP1A\_HUMAN | Serine/threonine-protein phosphatase PP1-alpha catalytic subunit | 5C(125.0477) | NaN | X | NaN | NaN | NaN | NaN | 1.170247e+05 | 1.170247e+05 | NaN | NaN | GNHECASINR |
| PGK1 | PGK1\_HUMAN | Phosphoglycerate kinase 1 | 316 | 298 | PGK1 | PGK1\_HUMAN | Phosphoglycerate kinase 1 | 19C(15.9949) | NaN | NaN | X | NaN | NaN | NaN | 1.167154e+05 | 1.167154e+05 | NaN | NaN | TGQATVASGIPAGWMGLDCGPESSKK |
| PTPN4 | PTN4\_HUMAN | Tyrosine-protein phosphatase non-receptor type 4 | 301 | 300 | PTPN4 | PTN4\_HUMAN | Tyrosine-protein phosphatase non-receptor type 4 | 2C(125.0477) | NaN | X | NaN | NaN | NaN | NaN | 1.166999e+05 | 1.166999e+05 | NaN | NaN | ACVEHHTFFR |
| SEC23A | SC23A\_HUMAN | Protein transport protein Sec23A | 423 | 416 | SEC23A | SC23A\_HUMAN | Protein transport protein Sec23A | 8C(125.0477) | NaN | X | NaN | NaN | NaN | NaN | 1.166761e+05 | 1.166761e+05 | NaN | NaN | ISGAIGPCVSLNSK |
| CAND1 | CAND1\_HUMAN | Cullin-associated NEDD8-dissociated protein 1 | 571 | 567 | CAND1 | CAND1\_HUMAN | Cullin-associated NEDD8-dissociated protein 1 | 5C(125.0477) | NaN | X | NaN | NaN | NaN | NaN | 1.166679e+05 | 1.166679e+05 | NaN | NaN | DLFTCTIKR |
| METAP2 | MAP2\_HUMAN | Methionine aminopeptidase 2 | 380 | 372 | METAP2 | MAP2\_HUMAN | Methionine aminopeptidase 2 | 9C(125.0477) | NaN | X | NaN | NaN | NaN | NaN | 1.166212e+05 | 1.166212e+05 | NaN | NaN | GVVHDDMECSHYMK |
| XPO7 | XPO7\_HUMAN | Exportin-7 | 835 | 831 | XPO7 | XPO7\_HUMAN | Exportin-7 | 5C(125.0477) | NaN | X | NaN | NaN | NaN | NaN | 1.166005e+05 | 1.166005e+05 | NaN | NaN | GISICFSMLK |
| ANKFY1 | ANFY1\_HUMAN | Rabankyrin-5 | 499 | 488 | ANKFY1 | ANFY1\_HUMAN | Rabankyrin-5 | 12C(125.0477) | NaN | X | NaN | NaN | NaN | NaN | 1.165964e+05 | 1.165964e+05 | NaN | NaN | NKWGETPLHTACR |
| LTA4H | LKHA4\_HUMAN | Leukotriene A-4 hydrolase | 26 | 26 | LTA4H | LKHA4\_HUMAN | Leukotriene A-4 hydrolase | 1C(125.0477) | NaN | X | NaN | NaN | NaN | NaN | 1.164675e+05 | 1.164675e+05 | NaN | NaN | CSVDFTRR |
| PRDX2 | PRDX2\_HUMAN | Peroxiredoxin-2 | 70 | 67 | PRDX2 | PRDX2\_HUMAN | Peroxiredoxin-2 | 4C(125.0477) | NaN | X | NaN | NaN | NaN | NaN | 1.164270e+05 | 1.164270e+05 | NaN | NaN | KLGCEVLGVSVDSQFTHLAWINTPRK |
| HBS1L | HBS1L\_HUMAN | HBS1-like protein | 491 | 490 | HBS1L | HBS1L\_HUMAN | HBS1-like protein | 2C(125.0477) | NaN | X | NaN | NaN | NaN | NaN | 1.164164e+05 | 1.164164e+05 | NaN | NaN | LCVSDVFK |
| CIAO2A | CIA2A\_HUMAN | Cytosolic iron-sulfur assembly component 2A | 155 | 150 | CIAO2A | CIA2A\_HUMAN | Cytosolic iron-sulfur assembly component 2A | 6C(125.0477) | NaN | X | NaN | NaN | NaN | NaN | 1.161109e+05 | 1.161109e+05 | NaN | NaN | EIVEQCVLEPD |
| PPP2CA | PP2AA\_HUMAN | Serine/threonine-protein phosphatase 2A catalytic subunit alpha isoform | 196 | 186 | PPP2CA | PP2AA\_HUMAN | Serine/threonine-protein phosphatase 2A catalytic subunit alpha isoform | 11C(125.0477) | NaN | X | NaN | NaN | NaN | NaN | 1.157841e+05 | 1.157841e+05 | NaN | NaN | LQEVPHEGPMCDLLWSDPDDR |
| ARL15 | ARL15\_HUMAN | ADP-ribosylation factor-like protein 15 | 175 | 170 | ARL15 | ARL15\_HUMAN | ADP-ribosylation factor-like protein 15 | 6C(125.0477) | NaN | X | NaN | NaN | NaN | NaN | 1.157603e+05 | 1.157603e+05 | NaN | NaN | WILQPCSLDDMDALK |
| UROS | HEM4\_HUMAN | Uroporphyrinogen-III synthase | 14 | 11 | UROS | HEM4\_HUMAN | Uroporphyrinogen-III synthase | 4C(125.0477) | NaN | X | NaN | NaN | NaN | NaN | 1.155584e+05 | 9.324254e+04 | 2.231590e+04 | NaN | EDDCGQDPYIR |
| TCP1 | TCPA\_HUMAN | T-complex protein 1 subunit alpha | 357 | 356 | TCP1 | TCPA\_HUMAN | T-complex protein 1 subunit alpha | 2C(125.0477) | NaN | X | NaN | NaN | NaN | NaN | 1.155409e+05 | 1.155409e+05 | NaN | NaN | ICDDELILIKNTK |
| WDR91 | WDR91\_HUMAN | WD repeat-containing protein 91 | 704 | 700 | WDR91 | WDR91\_HUMAN | WD repeat-containing protein 91 | 5C(125.0477) | NaN | X | NaN | NaN | NaN | NaN | 1.155371e+05 | 1.155371e+05 | NaN | NaN | VLESCLSLGGHR |
| EHBP1L1 | EH1L1\_HUMAN | EH domain-binding protein 1-like protein 1 | 422, 422 | 412 | EHBP1L1 | EH1L1\_HUMAN | EH domain-binding protein 1-like protein 1 | 11C(125.0477) 11C(31.9988) | NaN | X | NaN | X | NaN | NaN | 1.154854e+05 | 1.154854e+05 | NaN | NaN | AGEAEESSAVCQVDAEQR |
| SPTA1 | SPTA1\_HUMAN | Spectrin alpha chain, erythrocytic 1 | 475 | 471 | SPTA1 | SPTA1\_HUMAN | Spectrin alpha chain, erythrocytic 1 | 5C(125.0477) | NaN | X | NaN | NaN | NaN | NaN | 1.152376e+05 | 9.385223e+04 | 2.138539e+04 | NaN | QYEQCLDFHLFYR |
| PGD | 6PGD\_HUMAN | 6-phosphogluconate dehydrogenase, decarboxylating | 422 | 409 | PGD | 6PGD\_HUMAN | 6-phosphogluconate dehydrogenase, decarboxylating | 14C(125.0477) | NaN | X | NaN | NaN | NaN | NaN | 1.152260e+05 | 1.152260e+05 | NaN | NaN | AVSTGVQAGIPMPCFTTALSFYDGYR |
| EPB42 | EPB42\_HUMAN | Protein 4.2 | 428 | 422 | EPB42 | EPB42\_HUMAN | Protein 4.2 | 7C(125.0477) | NaN | X | NaN | NaN | NaN | NaN | 1.151456e+05 | 1.151456e+05 | NaN | NaN | GVGSDRCEDITQNYKYPEGSLQEK |
| CARM1 | CARM1\_HUMAN | Histone-arginine methyltransferase CARM1 | 78 | 62 | CARM1 | CARM1\_HUMAN | Histone-arginine methyltransferase CARM1 | 17C(125.0477) | NaN | X | NaN | NaN | NaN | NaN | 1.148343e+05 | 1.148343e+05 | NaN | NaN | AGPDSAGIALYSHEDVCVFK |
| PCBP2 | PCBP2\_HUMAN | Poly(rC)-binding protein 2 | 158 | 145 | PCBP2 | PCBP2\_HUMAN | Poly(rC)-binding protein 2 | 14C(125.0477) | NaN | X | NaN | NaN | NaN | NaN | 1.145995e+05 | 1.145995e+05 | NaN | NaN | AITIAGIPQSIIECVK |
| MTURN | MTURN\_HUMAN | Maturin | 14 | 13 | MTURN | MTURN\_HUMAN | Maturin | 2C(125.0477) | NaN | X | NaN | NaN | NaN | NaN | 1.142984e+05 | 1.142984e+05 | NaN | NaN | WCSNTPFELIATEETER |
| C2CD2L | C2C2L\_HUMAN | Phospholipid transfer protein C2CD2L | 385 | 383 | C2CD2L | C2C2L\_HUMAN | Phospholipid transfer protein C2CD2L | 3C(125.0477) | NaN | X | NaN | NaN | NaN | NaN | 1.142389e+05 | NaN | 1.142389e+05 | NaN | QLCPLTPGPGK |
| RABGGTA | PGTA\_HUMAN | Geranylgeranyl transferase type-2 subunit alpha | 99 | 91 | RABGGTA | PGTA\_HUMAN | Geranylgeranyl transferase type-2 subunit alpha | 9C(125.0477) | NaN | X | NaN | NaN | NaN | NaN | 1.141766e+05 | 1.141766e+05 | NaN | NaN | AELGFLESCLR |
| CCT7 | TCPH\_HUMAN | T-complex protein 1 subunit eta | 511 | 500 | CCT7 | TCPH\_HUMAN | T-complex protein 1 subunit eta | 12C(125.0477) | NaN | X | NaN | NaN | NaN | NaN | 1.140528e+05 | 1.140528e+05 | NaN | NaN | INALTAASEAACLIVSVDETIKNPR |
| TLN1 | TLN1\_HUMAN | Talin-1 | 709, 709, 719 | 693 | TLN1 | TLN1\_HUMAN | Talin-1 | 17C(15.9949) 17C(31.9988) 27C(57.0215) | X | NaN | X | X | NaN | NaN | 1.139633e+05 | NaN | 1.139633e+05 | NaN | TEDSGLQTQVIAAATQCALSTSQLVACTK |
| SEC31A | SC31A\_HUMAN | Protein transport protein Sec31A | 644 | 639 | SEC31A | SC31A\_HUMAN | Protein transport protein Sec31A | 6C(125.0477) | NaN | X | NaN | NaN | NaN | NaN | 1.139516e+05 | 1.139516e+05 | NaN | NaN | EIVESCDLK |
| ROCK2 | ROCK2\_HUMAN | Rho-associated protein kinase 2 | 887 | 882 | ROCK2 | ROCK2\_HUMAN | Rho-associated protein kinase 2 | 6C(125.0477) | NaN | X | NaN | NaN | NaN | NaN | 1.138874e+05 | 9.428946e+04 | 1.959791e+04 | NaN | ELKEECEEK |
| HTT | HD\_HUMAN | Huntingtin | 807 | 796 | HTT | HD\_HUMAN | Huntingtin | 12C(125.0477) | NaN | X | NaN | NaN | NaN | NaN | 1.138836e+05 | 1.138836e+05 | NaN | NaN | TLTGNTFSLADCIPLLR |
| AP2B1 | AP2B1\_HUMAN | AP-2 complex subunit beta | 144 | 139 | AP2B1 | AP2B1\_HUMAN | AP-2 complex subunit beta | 6C(125.0477) | NaN | X | NaN | NaN | NaN | NaN | 1.138507e+05 | 8.411538e+04 | 2.973531e+04 | NaN | KTAAVCVAK |
| NAXE | NNRE\_HUMAN | NAD(P)H-hydrate epimerase | 283 | 271 | NAXE | NNRE\_HUMAN | NAD(P)H-hydrate epimerase | 13C(125.0477) | NaN | X | NaN | NaN | NaN | NaN | 1.137812e+05 | 1.137812e+05 | NaN | NaN | YQLNLPPYPDTECVYR |
| IPO7 | IPO7\_HUMAN | Importin-7 | 736 | 724 | IPO7 | IPO7\_HUMAN | Importin-7 | 13C(125.0477) | NaN | X | NaN | NaN | NaN | NaN | 1.136770e+05 | 1.136770e+05 | NaN | NaN | KVLTGVAGEDAECHAAK |
| THOP1 | THOP1\_HUMAN | Thimet oligopeptidase | 350 | 349 | THOP1 | THOP1\_HUMAN | Thimet oligopeptidase | 2C(125.0477) | NaN | X | NaN | NaN | NaN | NaN | 1.136607e+05 | 1.136607e+05 | NaN | NaN | YCVDQNLLK |
| PLAA | PLAP\_HUMAN | Phospholipase A-2-activating protein | 193 | 186 | PLAA | PLAP\_HUMAN | Phospholipase A-2-activating protein | 8C(125.0477) | NaN | X | NaN | NaN | NaN | NaN | 1.133142e+05 | 1.133142e+05 | NaN | NaN | TFSGHEDCVR |
| COPS4 | CSN4\_HUMAN | COP9 signalosome complex subunit 4 | 255 | 255 | COPS4 | CSN4\_HUMAN | COP9 signalosome complex subunit 4 | 1C(125.0477) | NaN | X | NaN | NaN | NaN | NaN | 1.131197e+05 | 1.054393e+05 | 7.680407e+03 | NaN | CQQLAAYGILEK |
| USP5 | UBP5\_HUMAN | Ubiquitin carboxyl-terminal hydrolase 5 | 532 | 525 | USP5 | UBP5\_HUMAN | Ubiquitin carboxyl-terminal hydrolase 5 | 8C(125.0477) | NaN | X | NaN | NaN | NaN | NaN | 1.127282e+05 | 1.127282e+05 | NaN | NaN | AQVPFSSCLEAYGAPEQVDDFWSTALQAK |
| PSMD12 | PSD12\_HUMAN | 26S proteasome non-ATPase regulatory subunit 12 | 255 | 249 | PSMD12 | PSD12\_HUMAN | 26S proteasome non-ATPase regulatory subunit 12 | 7C(125.0477) | NaN | X | NaN | NaN | NaN | NaN | 1.126854e+05 | 8.591278e+04 | 2.677267e+04 | NaN | AIYDTPCIQAESEK |
| GAS2L1 | GA2L1\_HUMAN | GAS2-like protein 1 | 139 | 135 | GAS2L1 | GA2L1\_HUMAN | GAS2-like protein 1 | 5C(125.0477) | NaN | X | NaN | NaN | NaN | NaN | 1.124765e+05 | 1.124765e+05 | NaN | NaN | SVVLCLLEVAR |
| CUL3 | CUL3\_HUMAN | Cullin-3 | 251 | 248 | CUL3 | CUL3\_HUMAN | Cullin-3 | 4C(125.0477) | NaN | X | NaN | NaN | NaN | NaN | 1.124749e+05 | 1.124749e+05 | NaN | NaN | VMHCLDK |
| KPRP | KPRP\_HUMAN | Keratinocyte proline-rich protein | 391 | 388 | KPRP | KPRP\_HUMAN | Keratinocyte proline-rich protein | 4C(125.0477) | NaN | X | NaN | NaN | NaN | NaN | 1.123601e+05 | NaN | 6.104645e+04 | 51313.6250 | LDQCPESPLQR |
| KRT4 | K2C4\_HUMAN | Keratin, type II cytoskeletal 4 | 42 | 42 | KRT4 | K2C4\_HUMAN | Keratin, type II cytoskeletal 4 | 1C(125.0477) | NaN | X | NaN | NaN | NaN | NaN | 1.122360e+05 | NaN | 6.942313e+04 | 42812.8360 | CSSGGFGSR |
| GIT1 | GIT1\_HUMAN | ARF GTPase-activating protein GIT1 | 222 | 219 | GIT1 | GIT1\_HUMAN | ARF GTPase-activating protein GIT1 | 4C(125.0477) | NaN | X | NaN | NaN | NaN | NaN | 1.122279e+05 | NaN | 1.122279e+05 | NaN | LVECQYELTDR |
| PLG | PLMN\_HUMAN | Plasminogen | 784 | 770 | PLG | PLMN\_HUMAN | Plasminogen | 15C(57.0215) | X | NaN | NaN | NaN | NaN | NaN | 1.122013e+05 | 1.122013e+05 | NaN | NaN | DKYILQGVTSWGLGCARPNKPGVYVR |
| G6PD | G6PD\_HUMAN | Glucose-6-phosphate 1-dehydrogenase | 294 | 294 | G6PD | G6PD\_HUMAN | Glucose-6-phosphate 1-dehydrogenase | 1C(125.0477) | NaN | X | NaN | NaN | NaN | NaN | 1.121881e+05 | 1.121881e+05 | NaN | NaN | CISEVQANNVVLGQYVGNPDGEGEATK |
| BCAM | BCAM\_HUMAN | Basal cell adhesion molecule | 125 | 122 | BCAM | BCAM\_HUMAN | Basal cell adhesion molecule | 4C(125.0477) | NaN | X | NaN | NaN | NaN | NaN | 1.121813e+05 | NaN | 1.039678e+05 | 8213.5500 | DYVCVVR |
| AGO2 | AGO2\_HUMAN | Protein argonaute-2 | 787 | 774 | AGO2 | AGO2\_HUMAN | Protein argonaute-2 | 14C(125.0477) | NaN | X | NaN | NaN | NaN | NaN | 1.120104e+05 | 1.120104e+05 | NaN | NaN | FSSDELQILTYQLCHTYVR |
| ATP1A1 | AT1A1\_HUMAN | Sodium/potassium-transporting ATPase subunit alpha-1 | 663 | 662 | ATP1A1 | AT1A1\_HUMAN | Sodium/potassium-transporting ATPase subunit alpha-1 | 2C(125.0477) | NaN | X | NaN | NaN | NaN | NaN | 1.116584e+05 | NaN | 5.634612e+04 | 55312.3160 | ACVVHGSDLK |
| PPM1A | PPM1A\_HUMAN | Protein phosphatase 1A | 314 | 305 | PPM1A | PPM1A\_HUMAN | Protein phosphatase 1A | 10C(125.0477) | NaN | X | NaN | NaN | NaN | NaN | 1.116506e+05 | 1.116506e+05 | NaN | NaN | EAELDKYLECR |
| RNH1 | RINI\_HUMAN | Ribonuclease inhibitor | 323, 329, 329, 330 | 322 | RNH1 | RINI\_HUMAN | Ribonuclease inhibitor | 2C(125.0477) 8C(125.0477) 8C(15.9949) 9C(15.9949) | NaN | X | X | NaN | NaN | NaN | 1.116451e+05 | 1.116451e+05 | NaN | NaN | SCSFTAACCSHFSSVLAQNR |
| PPP2R1A | 2AAA\_HUMAN | Serine/threonine-protein phosphatase 2A 65 kDa regulatory subunit A alpha isoform | 377 | 359 | PPP2R1A | 2AAA\_HUMAN | Serine/threonine-protein phosphatase 2A 65 kDa regulatory subunit A alpha isoform | 19C(125.0477) | NaN | X | NaN | NaN | NaN | NaN | 1.114295e+05 | 1.114295e+05 | NaN | NaN | DNTIEHLLPLFLAQLKDECPEVR |
| CAP1 | CAP1\_HUMAN | Adenylyl cyclase-associated protein 1 | 356 | 356 | CAP1 | CAP1\_HUMAN | Adenylyl cyclase-associated protein 1 | 1C(125.0477) | NaN | X | NaN | NaN | NaN | NaN | 1.113211e+05 | NaN | 1.113211e+05 | NaN | CVNTTLQIK |
| UCHL3 | UCHL3\_HUMAN | Ubiquitin carboxyl-terminal hydrolase isozyme L3 | 95 | 89 | UCHL3 | UCHL3\_HUMAN | Ubiquitin carboxyl-terminal hydrolase isozyme L3 | 7C(125.0477) | NaN | X | NaN | NaN | NaN | NaN | 1.112656e+05 | 1.112656e+05 | NaN | NaN | QTISNACGTIGLIHAIANNKDK |
| DAAM1 | DAAM1\_HUMAN | Disheveled-associated activator of morphogenesis 1 | 231 | 221 | DAAM1 | DAAM1\_HUMAN | Disheveled-associated activator of morphogenesis 1 | 11C(125.0477) | NaN | X | NaN | NaN | NaN | NaN | 1.112577e+05 | 1.112577e+05 | NaN | NaN | VAVLEILGAVCLVPGGHK |
| NR2C2AP | NR2CA\_HUMAN | Nuclear receptor 2C2-associated protein | 76, 76 | 75 | NR2C2AP | NR2CA\_HUMAN | Nuclear receptor 2C2-associated protein | 2C(125.0477) 2C(31.9988) | NaN | X | NaN | X | NaN | NaN | 1.110552e+05 | 1.110552e+05 | NaN | NaN | GCLEGSQGTQALHK |
| FSCN1 | FSCN1\_HUMAN | Fascin | 397 | 390 | FSCN1 | FSCN1\_HUMAN | Fascin | 8C(125.0477) | NaN | X | NaN | NaN | NaN | NaN | 1.108307e+05 | 1.108307e+05 | NaN | NaN | GEHGFIGCRK |
| GLMN | GLMN\_HUMAN | Glomulin | 375 | 371 | GLMN | GLMN\_HUMAN | Glomulin | 5C(125.0477) | NaN | X | NaN | NaN | NaN | NaN | 1.106011e+05 | 1.106011e+05 | NaN | NaN | VMTLCPIETLR |
| CAPZA1 | CAZA1\_HUMAN | F-actin-capping protein subunit alpha-1 | 141 | 134 | CAPZA1 | CAZA1\_HUMAN | F-actin-capping protein subunit alpha-1 | 8C(125.0477) | NaN | X | NaN | NaN | NaN | NaN | 1.103989e+05 | 1.103989e+05 | NaN | NaN | DHYSNGFCTVYAK |
| CNP | CN37\_HUMAN | 2',3'-cyclic-nucleotide 3'-phosphodiesterase | 257, 252 | 245 | CNP | CN37\_HUMAN | 2',3'-cyclic-nucleotide 3'-phosphodiesterase | 13C(15.9949) 8C(15.9949) | NaN | NaN | X | NaN | NaN | NaN | 1.103953e+05 | 1.103953e+05 | NaN | NaN | RPPGVLHCTTKFCDYGK |
| PPP6R3 | PP6R3\_HUMAN | Serine/threonine-protein phosphatase 6 regulatory subunit 3 | 216 | 195 | PPP6R3 | PP6R3\_HUMAN | Serine/threonine-protein phosphatase 6 regulatory subunit 3 | 22C(125.0477) | NaN | X | NaN | NaN | NaN | NaN | 1.103764e+05 | 1.103764e+05 | NaN | NaN | LVEIVHPSQEEDRHSNASQSLCEIVR |
| ANKFY1 | ANFY1\_HUMAN | Rabankyrin-5 | 34 | 34 | ANKFY1 | ANFY1\_HUMAN | Rabankyrin-5 | 1C(125.0477) | NaN | X | NaN | NaN | NaN | NaN | 1.103243e+05 | 4.405597e+04 | 6.626830e+04 | NaN | CALLAAQANK |
| RNF14 | RNF14\_HUMAN | E3 ubiquitin-protein ligase RNF14 | 350 | 342 | RNF14 | RNF14\_HUMAN | E3 ubiquitin-protein ligase RNF14 | 9C(125.0477) | NaN | X | NaN | NaN | NaN | NaN | 1.103165e+05 | 1.103165e+05 | NaN | NaN | LTYHGVSPCK |
| ANXA7 | ANXA7\_HUMAN | Annexin A7 | 327 | 322 | ANXA7 | ANXA7\_HUMAN | Annexin A7 | 6C(125.0477) | NaN | X | NaN | NaN | NaN | NaN | 1.098314e+05 | NaN | 1.098314e+05 | NaN | LLVSMCQGNR |
| ABCE1 | ABCE1\_HUMAN | ATP-binding cassette sub-family E member 1 | 201 | 196 | ABCE1 | ABCE1\_HUMAN | ATP-binding cassette sub-family E member 1 | 6C(125.0477) | NaN | X | NaN | NaN | NaN | NaN | 1.096501e+05 | 1.096501e+05 | NaN | NaN | TQAIVCQQLDLTHLK |
| PSMD14 | PSDE\_HUMAN | 26S proteasome non-ATPase regulatory subunit 14 | 238 | 224 | PSMD14 | PSDE\_HUMAN | 26S proteasome non-ATPase regulatory subunit 14 | 15C(125.0477) | NaN | X | NaN | NaN | NaN | NaN | 1.096413e+05 | 1.096413e+05 | NaN | NaN | SWMEGLTLQDYSEHCK |
| XPO7 | XPO7\_HUMAN | Exportin-7 | 43 | 33 | XPO7 | XPO7\_HUMAN | Exportin-7 | 11C(125.0477) | NaN | X | NaN | NaN | NaN | NaN | 1.096164e+05 | 1.096164e+05 | NaN | NaN | ALVEFTNSPDCLSK |
| NAXD | NNRD\_HUMAN | ATP-dependent (S)-NAD(P)H-hydrate dehydratase | 108 | 99 | NAXD | NNRD\_HUMAN | ATP-dependent (S)-NAD(P)H-hydrate dehydratase | 10C(125.0477) | NaN | X | NaN | NaN | NaN | NaN | 1.095796e+05 | 1.095796e+05 | NaN | NaN | VGADLSHVFCASAAAPVIK |
| FABP5 | FABP5\_HUMAN | Fatty acid-binding protein 5 | 67 | 62 | FABP5 | FABP5\_HUMAN | Fatty acid-binding protein 5 | 6C(125.0477) | NaN | X | NaN | NaN | NaN | NaN | 1.095478e+05 | 4.484040e+04 | 6.470742e+04 | NaN | TTQFSCTLGEK |
| ALB | ALBU\_HUMAN | Albumin | 99 | 98 | ALB | ALBU\_HUMAN | Albumin | 2C(57.0215) | X | NaN | NaN | NaN | NaN | NaN | 1.094968e+05 | 0.000000e+00 | NaN | 109496.8000 | LCTVATLR |
| USP9X | USP9X\_HUMAN | Probable ubiquitin carboxyl-terminal hydrolase FAF-X | 819 | 804 | USP9X | USP9X\_HUMAN | Probable ubiquitin carboxyl-terminal hydrolase FAF-X | 16C(125.0477) | NaN | X | NaN | NaN | NaN | NaN | 1.094680e+05 | 1.094680e+05 | NaN | NaN | LQVNQVVIHEDFIQSCFDR |
| GPS1 | CSN1\_HUMAN | COP9 signalosome complex subunit 1 | 192 | 188 | GPS1 | CSN1\_HUMAN | COP9 signalosome complex subunit 1 | 5C(125.0477) | NaN | X | NaN | NaN | NaN | NaN | 1.094164e+05 | 1.094164e+05 | NaN | NaN | ARDYCTSAK |
| MYH10 | MYH10\_HUMAN | Myosin-10 | 1011 | 1008 | MYH10 | MYH10\_HUMAN | Myosin-10 | 4C(125.0477) | NaN | X | NaN | NaN | NaN | NaN | 1.093750e+05 | 1.093750e+05 | NaN | NaN | IAECSSQLAEEEEK |
| UBE2O | UBE2O\_HUMAN | (E3-independent) E2 ubiquitin-conjugating enzyme | 1288 | 1278 | UBE2O | UBE2O\_HUMAN | (E3-independent) E2 ubiquitin-conjugating enzyme | 11C(125.0477) | NaN | X | NaN | NaN | NaN | NaN | 1.089260e+05 | NaN | 1.089260e+05 | NaN | AALLEAGMPECTEDK |
| CCT5 | TCPE\_HUMAN | T-complex protein 1 subunit epsilon | 493 | 484 | CCT5 | TCPE\_HUMAN | T-complex protein 1 subunit epsilon | 10C(125.0477) | NaN | X | NaN | NaN | NaN | NaN | 1.088147e+05 | 1.088147e+05 | NaN | NaN | EMNPALGIDCLHK |
| CSE1L | XPO2\_HUMAN | Exportin-2 | 195 | 190 | CSE1L | XPO2\_HUMAN | Exportin-2 | 6C(125.0477) | NaN | X | NaN | NaN | NaN | NaN | 1.087414e+05 | 1.087414e+05 | NaN | NaN | ATIELCSTHANDASALR |
| UFC1 | UFC1\_HUMAN | Ubiquitin-fold modifier-conjugating enzyme 1 | 69 | 69 | UFC1 | UFC1\_HUMAN | Ubiquitin-fold modifier-conjugating enzyme 1 | 1C(125.0477) | NaN | X | NaN | NaN | NaN | NaN | 1.084707e+05 | 1.084707e+05 | NaN | NaN | CWYIHDLLK |
| FAHD1 | FAHD1\_HUMAN | Acylpyruvase FAHD1, mitochondrial | 129 | 124 | FAHD1 | FAHD1\_HUMAN | Acylpyruvase FAHD1, mitochondrial | 6C(125.0477) | NaN | X | NaN | NaN | NaN | NaN | 1.084112e+05 | 1.084112e+05 | NaN | NaN | SFTASCPVSAFVPK |
| HSPA4 | HSP74\_HUMAN | Heat shock 70 kDa protein 4 | 380 | 375 | HSPA4 | HSP74\_HUMAN | Heat shock 70 kDa protein 4 | 6C(125.0477) | NaN | X | NaN | NaN | NaN | NaN | 1.081719e+05 | 1.081719e+05 | NaN | NaN | GCALQCAILSPAFK |
| ATP6V1H | VATH\_HUMAN | V-type proton ATPase subunit H | 85 | 76 | ATP6V1H | VATH\_HUMAN | V-type proton ATPase subunit H | 10C(125.0477) | NaN | X | NaN | NaN | NaN | NaN | 1.080749e+05 | 1.080749e+05 | NaN | NaN | TFINLMTHICK |
| FLNA | FLNA\_HUMAN | Filamin-A | 717 | 709 | FLNA | FLNA\_HUMAN | Filamin-A | 9C(125.0477) | NaN | X | NaN | NaN | NaN | NaN | 1.078985e+05 | 1.078985e+05 | NaN | NaN | VQVQDNEGCPVEALVK |
| NAP1L1 | NP1L1\_HUMAN | Nucleosome assembly protein 1-like 1 | 388 | 381 | NAP1L1 | NP1L1\_HUMAN | Nucleosome assembly protein 1-like 1 | 8C(15.9949) | NaN | NaN | X | NaN | NaN | NaN | 1.077884e+05 | 1.077884e+05 | NaN | NaN | KDQNPAECK |
| CLTA | CLCA\_HUMAN | Clathrin light chain A | 218 | 217 | CLTA | CLCA\_HUMAN | Clathrin light chain A | 2C(125.0477) | NaN | X | NaN | NaN | NaN | NaN | 1.077809e+05 | NaN | 1.077809e+05 | NaN | LCDFNPK |
| ANKFY1 | ANFY1\_HUMAN | Rabankyrin-5 | 749 | 747 | ANKFY1 | ANFY1\_HUMAN | Rabankyrin-5 | 3C(125.0477) | NaN | X | NaN | NaN | NaN | NaN | 1.076442e+05 | 1.076442e+05 | NaN | NaN | SGCDVNSPR |
| EIF2B5 | EI2BE\_HUMAN | Translation initiation factor eIF-2B subunit epsilon | 571 | 564 | EIF2B5 | EI2BE\_HUMAN | Translation initiation factor eIF-2B subunit epsilon | 8C(125.0477) | NaN | X | NaN | NaN | NaN | NaN | 1.074891e+05 | 1.074891e+05 | NaN | NaN | GKEENISCDNLVLEINSLK |
| ABCF1 | ABCF1\_HUMAN | ATP-binding cassette sub-family F member 1 | 741 | 729 | ABCF1 | ABCF1\_HUMAN | ATP-binding cassette sub-family F member 1 | 13C(125.0477) | NaN | X | NaN | NaN | NaN | NaN | 1.074867e+05 | 1.074867e+05 | NaN | NaN | FGLESHAHTIQICK |
| GGA3 | GGA3\_HUMAN | ADP-ribosylation factor-binding protein GGA3 | 255, 255 | 250 | GGA3 | GGA3\_HUMAN | ADP-ribosylation factor-binding protein GGA3 | 6C(125.0477) 6C(15.9949) | NaN | X | X | NaN | NaN | NaN | 1.073575e+05 | 1.073575e+05 | NaN | NaN | ELFDQCENK |
| RABGAP1L | RBG1L\_HUMAN | Rab GTPase-activating protein 1-like | 330 | 330 | RABGAP1L | RBG1L\_HUMAN | Rab GTPase-activating protein 1-like | 1C(125.0477) | NaN | X | NaN | NaN | NaN | NaN | 1.073275e+05 | 1.073275e+05 | NaN | NaN | CFGMLLSPGR |
| MYL12B | ML12B\_HUMAN | Myosin regulatory light chain 12B | 109 | 105 | MYL12B | ML12B\_HUMAN | Myosin regulatory light chain 12B | 5C(125.0477) | NaN | X | NaN | NaN | NaN | NaN | 1.071535e+05 | 1.071535e+05 | NaN | NaN | NAFACFDEEATGTIQEDYLR |
| PLAA | PLAP\_HUMAN | Phospholipase A-2-activating protein | 631 | 623 | PLAA | PLAP\_HUMAN | Phospholipase A-2-activating protein | 9C(125.0477) | NaN | X | NaN | NaN | NaN | NaN | 1.071103e+05 | 1.071103e+05 | NaN | NaN | HPSVNENFCNEK |
| GRHPR | GRHPR\_HUMAN | Glyoxylate reductase/hydroxypyruvate reductase | 288 | 287 | GRHPR | GRHPR\_HUMAN | Glyoxylate reductase/hydroxypyruvate reductase | 2C(125.0477) | NaN | X | NaN | NaN | NaN | NaN | 1.069616e+05 | 1.069616e+05 | NaN | NaN | NCVILPHIGSATHR |
| NADSYN1 | NADE\_HUMAN | Glutamine-dependent NAD(+) synthetase | 428 | 422 | NADSYN1 | NADE\_HUMAN | Glutamine-dependent NAD(+) synthetase | 7C(125.0477) | NaN | X | NaN | NaN | NaN | NaN | 1.069315e+05 | 1.069315e+05 | NaN | NaN | NSSQETCTR |
| CIAO1 | CIAO1\_HUMAN | Probable cytosolic iron-sulfur protein assembly protein CIAO1 | 52 | 42 | CIAO1 | CIAO1\_HUMAN | Probable cytosolic iron-sulfur protein assembly protein CIAO1 | 11C(125.0477) | NaN | X | NaN | NaN | NaN | NaN | 1.067801e+05 | 1.067801e+05 | NaN | NaN | IWGTEGDSWICK |
| CSE1L | XPO2\_HUMAN | Exportin-2 | 85, 85 | 76 | CSE1L | XPO2\_HUMAN | Exportin-2 | 10C(125.0477) 10C(31.9988) | NaN | X | NaN | X | NaN | NaN | 1.067528e+05 | 1.067528e+05 | NaN | NaN | IVEDEPNKICEADR |
| SAAL1 | SAAL1\_HUMAN | Protein SAAL1 | 248 | 245 | SAAL1 | SAAL1\_HUMAN | Protein SAAL1 | 4C(125.0477) | NaN | X | NaN | NaN | NaN | NaN | 1.066983e+05 | 1.066983e+05 | NaN | NaN | LVPCILEAAK |
| FHOD1 | FHOD1\_HUMAN | FH1/FH2 domain-containing protein 1 | 502 | 495 | FHOD1 | FHOD1\_HUMAN | FH1/FH2 domain-containing protein 1 | 8C(125.0477) | NaN | X | NaN | NaN | NaN | NaN | 1.066238e+05 | 1.066238e+05 | NaN | NaN | TPQSPAPCVLLR |
| NPLOC4 | NPL4\_HUMAN | Nuclear protein localization protein 4 homolog | 130 | 124 | NPLOC4 | NPL4\_HUMAN | Nuclear protein localization protein 4 homolog | 7C(125.0477) | NaN | X | NaN | NaN | NaN | NaN | 1.065784e+05 | 1.065784e+05 | NaN | NaN | SRDPQLCR |
| CARM1 | CARM1\_HUMAN | Histone-arginine methyltransferase CARM1 | 193 | 187 | CARM1 | CARM1\_HUMAN | Histone-arginine methyltransferase CARM1 | 7C(125.0477) | NaN | X | NaN | NaN | NaN | NaN | 1.065473e+05 | 1.065473e+05 | NaN | NaN | IVLDVGCGSGILSFFAAQAGAR |
| VPS35 | VPS35\_HUMAN | Vacuolar protein sorting-associated protein 35 | 653 | 651 | VPS35 | VPS35\_HUMAN | Vacuolar protein sorting-associated protein 35 | 3C(125.0477) | NaN | X | NaN | NaN | NaN | NaN | 1.063349e+05 | 1.063349e+05 | NaN | NaN | TQCALAASK |
| DUSP23 | DUS23\_HUMAN | Dual specificity protein phosphatase 23 | 67 | 63 | DUSP23 | DUS23\_HUMAN | Dual specificity protein phosphatase 23 | 5C(125.0477) | NaN | X | NaN | NaN | NaN | NaN | 1.061439e+05 | 1.061439e+05 | NaN | NaN | IPDFCPPAPDQIDR |
| ENOSF1 | ENOF1\_HUMAN | Mitochondrial enolase superfamily member 1 | 385 | 384 | ENOSF1 | ENOF1\_HUMAN | Mitochondrial enolase superfamily member 1 | 2C(125.0477) | NaN | X | NaN | NaN | NaN | NaN | 1.061060e+05 | 1.061060e+05 | NaN | NaN | VCEYVDHLHEHFK |
| NCAPH | CND2\_HUMAN | Condensin complex subunit 2 | 114 | 102 | NCAPH | CND2\_HUMAN | Condensin complex subunit 2 | 13C(125.0477) | NaN | X | NaN | NaN | NaN | NaN | 1.059628e+05 | 1.059628e+05 | NaN | NaN | FTNTQITEHYSTCIK |
| ANXA7 | ANXA7\_HUMAN | Annexin A7 | 298 | 298 | ANXA7 | ANXA7\_HUMAN | Annexin A7 | 1C(125.0477) | NaN | X | NaN | NaN | NaN | NaN | 1.058785e+05 | NaN | 1.058785e+05 | NaN | CYQSEFGR |
| PTRHD1 | PTRD1\_HUMAN | Putative peptidyl-tRNA hydrolase PTRHD1 | 52 | 34 | PTRHD1 | PTRD1\_HUMAN | Putative peptidyl-tRNA hydrolase PTRHD1 | 19C(125.0477) | NaN | X | NaN | NaN | NaN | NaN | 1.056997e+05 | 1.056997e+05 | NaN | NaN | DLSQAPFSWPAGALVAQACHAATAALHTHR |
| UROD | DCUP\_HUMAN | Uroporphyrinogen decarboxylase | 65, 65, 66, 59 | 51 | UROD | DCUP\_HUMAN | Uroporphyrinogen decarboxylase | 15C(125.0477) 15C(15.9949) 16C(15.9949) 9C(125.0477) | NaN | X | X | NaN | NaN | NaN | 1.056350e+05 | 1.056350e+05 | NaN | NaN | AAQDFFSTCRSPEACCELTLQPLR |
| EIF5A | IF5A1\_HUMAN | Eukaryotic translation initiation factor 5A-1 | 73 | 68 | EIF5A | IF5A1\_HUMAN | Eukaryotic translation initiation factor 5A-1 | 6C(125.0477) | NaN | X | NaN | NaN | NaN | NaN | 1.052977e+05 | 1.052977e+05 | NaN | NaN | KYEDICPSTHNMDVPNIKR |
| GPS1 | CSN1\_HUMAN | COP9 signalosome complex subunit 1 | 251 | 251 | GPS1 | CSN1\_HUMAN | COP9 signalosome complex subunit 1 | 1C(125.0477) | NaN | X | NaN | NaN | NaN | NaN | 1.050620e+05 | 7.196445e+04 | 3.309755e+04 | NaN | CAAGLAELAAR |
| PDCD6IP | PDC6I\_HUMAN | Programmed cell death 6-interacting protein | 691 | 691 | PDCD6IP | PDC6I\_HUMAN | Programmed cell death 6-interacting protein | 1C(125.0477) | NaN | X | NaN | NaN | NaN | NaN | 1.050435e+05 | NaN | 1.050435e+05 | NaN | CSDIVFAR |
| UBE2M | UBC12\_HUMAN | NEDD8-conjugating enzyme Ubc12 | 47 | 46 | UBE2M | UBC12\_HUMAN | NEDD8-conjugating enzyme Ubc12 | 2C(125.0477) | NaN | X | NaN | NaN | NaN | NaN | 1.049753e+05 | 1.049753e+05 | NaN | NaN | TCDISFSDPDDLLNFK |
| APIP | MTNB\_HUMAN | Methylthioribulose-1-phosphate dehydratase | 76 | 68 | APIP | MTNB\_HUMAN | Methylthioribulose-1-phosphate dehydratase | 9C(125.0477) | NaN | X | NaN | NaN | NaN | NaN | 1.047039e+05 | 1.047039e+05 | NaN | NaN | IQPEDMFVCDINEK |
| VCP | TERA\_HUMAN | Transitional endoplasmic reticulum ATPase | 415 | 390 | VCP | TERA\_HUMAN | Transitional endoplasmic reticulum ATPase | 26C(125.0477) | NaN | X | NaN | NaN | NaN | NaN | 1.046972e+05 | 1.046972e+05 | NaN | NaN | LADDVDLEQVANETHGHVGADLAALCSEAALQAIR |
| LANCL2 | LANC2\_HUMAN | LanC-like protein 2 | 149 | 145 | LANCL2 | LANC2\_HUMAN | LanC-like protein 2 | 5C(125.0477) | NaN | X | NaN | NaN | NaN | NaN | 1.042988e+05 | 1.042988e+05 | NaN | NaN | VTFLCGDAGPLAVGAVIYHK |
| AGO2 | AGO2\_HUMAN | Protein argonaute-2 | 546 | 535 | AGO2 | AGO2\_HUMAN | Protein argonaute-2 | 12C(125.0477) | NaN | X | NaN | NaN | NaN | NaN | 1.037161e+05 | 1.037161e+05 | NaN | NaN | VGDTVLGMATQCVQMK |
| PRPS1 | PRPS1\_HUMAN | Ribose-phosphate pyrophosphokinase 1 | 41 | 35 | PRPS1 | PRPS1\_HUMAN | Ribose-phosphate pyrophosphokinase 1 | 7C(125.0477) | NaN | X | NaN | NaN | NaN | NaN | 1.035634e+05 | NaN | 9.186696e+04 | 11696.4420 | FSNQETCVEIGESVR |
| SPTA1 | SPTA1\_HUMAN | Spectrin alpha chain, erythrocytic 1 | 2072, 2072, 2072 | 2062 | SPTA1 | SPTA1\_HUMAN | Spectrin alpha chain, erythrocytic 1 | 11C(125.0477) 11C(15.9949) 11C(31.9988) | NaN | X | X | X | NaN | NaN | 1.031221e+05 | NaN | 1.031221e+05 | NaN | MEENLSEPVHCVSLNEIR |
| DERA | DEOC\_HUMAN | Deoxyribose-phosphate aldolase | 107, 100 | 93 | DERA | DEOC\_HUMAN | Deoxyribose-phosphate aldolase | 15C(15.9949) 8C(15.9949) | NaN | NaN | X | NaN | NaN | NaN | 1.030554e+05 | 1.030554e+05 | NaN | NaN | GITTAAVCVYPARVCDAVK |
| YARS1 | SYYC\_HUMAN | Tyrosine--tRNA ligase, cytoplasmic | 250 | 247 | YARS1 | SYYC\_HUMAN | Tyrosine--tRNA ligase, cytoplasmic | 4C(125.0477) | NaN | X | NaN | NaN | NaN | NaN | 1.029860e+05 | 1.029860e+05 | NaN | NaN | KAFCEPGNVENNGVLSFIK |
| GDI2 | GDIB\_HUMAN | Rab GDP dissociation inhibitor beta | 282 | 279 | GDI2 | GDIB\_HUMAN | Rab GDP dissociation inhibitor beta | 4C(125.0477) | NaN | X | NaN | NaN | NaN | NaN | 1.029845e+05 | 1.029845e+05 | NaN | NaN | QLICDPSYVKDR |
| CPT1A | CPT1A\_HUMAN | Carnitine O-palmitoyltransferase 1, liver isoform | 304 | 296 | CPT1A | CPT1A\_HUMAN | Carnitine O-palmitoyltransferase 1, liver isoform | 9C(125.0477) | NaN | X | NaN | NaN | NaN | NaN | 1.028551e+05 | 1.028551e+05 | NaN | NaN | LLGSTIPLCSAQWER |
| USP14 | UBP14\_HUMAN | Ubiquitin carboxyl-terminal hydrolase 14 | 257 | 257 | USP14 | UBP14\_HUMAN | Ubiquitin carboxyl-terminal hydrolase 14 | 1C(15.9949) | NaN | NaN | X | NaN | NaN | NaN | 1.028550e+05 | 1.028550e+05 | NaN | NaN | CTESEEEEVTKGKENQLQLSCFINQEVK |
| GSR | GSHR\_HUMAN | Glutathione reductase, mitochondrial | 377 | 369 | GSR | GSHR\_HUMAN | Glutathione reductase, mitochondrial | 9C(125.0477) | NaN | X | NaN | NaN | NaN | NaN | 1.027558e+05 | NaN | 1.027558e+05 | NaN | GIYAVGDVCGK |
| BLVRB | BLVRB\_HUMAN | Flavin reductase (NADPH) | 109 | 106 | BLVRB | BLVRB\_HUMAN | Flavin reductase (NADPH) | 4C(125.0477) | NaN | X | NaN | NaN | NaN | NaN | 1.027410e+05 | NaN | 8.408321e+04 | 18657.7750 | VVACTSAFLLWDPTK |
| NAPA | SNAA\_HUMAN | Alpha-soluble NSF attachment protein | 103 | 94 | NAPA | SNAA\_HUMAN | Alpha-soluble NSF attachment protein | 10C(125.0477) | NaN | X | NaN | NaN | NaN | NaN | 1.025417e+05 | 1.025417e+05 | NaN | NaN | KADPQEAINCLMR |
| COPS3 | CSN3\_HUMAN | COP9 signalosome complex subunit 3 | 298 | 297 | COPS3 | CSN3\_HUMAN | COP9 signalosome complex subunit 3 | 2C(125.0477) | NaN | X | NaN | NaN | NaN | NaN | 1.023762e+05 | 7.694211e+04 | 2.543410e+04 | NaN | QCLSSLYK |
| PPCDC | COAC\_HUMAN | Phosphopantothenoylcysteine decarboxylase | 173 | 171 | PPCDC | COAC\_HUMAN | Phosphopantothenoylcysteine decarboxylase | 3C(125.0477) | NaN | X | NaN | NaN | NaN | NaN | 1.021584e+05 | 1.021584e+05 | NaN | NaN | LVCGDEGLGAMAEVGTIVDK |
| UCHL3 | UCHL3\_HUMAN | Ubiquitin carboxyl-terminal hydrolase isozyme L3 | 209 | 187 | UCHL3 | UCHL3\_HUMAN | Ubiquitin carboxyl-terminal hydrolase isozyme L3 | 23C(125.0477) | NaN | X | NaN | NaN | NaN | NaN | 1.021131e+05 | 1.021131e+05 | NaN | NaN | KPFPINHGETSDETLLEDAIEVCK |
| KRT86 | KRT86\_HUMAN | Keratin, type II cuticular Hb6 | 168 | 165 | KRT86 | KRT86\_HUMAN | Keratin, type II cuticular Hb6 | 4C(125.0477) | NaN | X | NaN | NaN | NaN | NaN | 1.020963e+05 | NaN | 2.072194e+04 | 81374.3600 | EAECVEADSGR |
| HTT | HD\_HUMAN | Huntingtin | 718 | 713 | HTT | HD\_HUMAN | Huntingtin | 6C(125.0477) | NaN | X | NaN | NaN | NaN | NaN | 1.020165e+05 | 1.020165e+05 | NaN | NaN | ALALSCVGAAVALHPESFFSK |
| PDE12 | PDE12\_HUMAN | 2',5'-phosphodiesterase 12 | 321 | 315 | PDE12 | PDE12\_HUMAN | 2',5'-phosphodiesterase 12 | 7C(125.0477) | NaN | X | NaN | NaN | NaN | NaN | 1.017919e+05 | 1.017919e+05 | NaN | NaN | TVLYPYCAPYALELDYR |
| GNAO1 | GNAO\_HUMAN | Guanine nucleotide-binding protein G(o) subunit alpha | 3 | 2 | GNAO1 | GNAO\_HUMAN | Guanine nucleotide-binding protein G(o) subunit alpha | 2C(57.0215) | X | NaN | NaN | NaN | NaN | NaN | 1.017589e+05 | 1.017589e+05 | NaN | NaN | GCTLSAEERAALER |
| GPX1 | GPX1\_HUMAN | Glutathione peroxidase 1 | 202 | 182 | GPX1 | GPX1\_HUMAN | Glutathione peroxidase 1 | 21C(125.0477) | NaN | X | NaN | NaN | NaN | NaN | 1.017263e+05 | 1.017263e+05 | NaN | NaN | RFQTIDIEPDIEALLSQGPSCA |
| ACAT2 | THIC\_HUMAN | Acetyl-CoA acetyltransferase, cytosolic | 65 | 44 | ACAT2 | THIC\_HUMAN | Acetyl-CoA acetyltransferase, cytosolic | 22C(125.0477) | NaN | X | NaN | NaN | NaN | NaN | 1.016844e+05 | 1.016844e+05 | NaN | NaN | RATVAPEDVSEVIFGHVLAAGCGQNPVR |
| KIF5B | KINH\_HUMAN | Kinesin-1 heavy chain | 65 | 51 | KIF5B | KINH\_HUMAN | Kinesin-1 heavy chain | 15C(125.0477) | NaN | X | NaN | NaN | NaN | NaN | 1.015938e+05 | 1.015938e+05 | NaN | NaN | VFQSSTSQEQVYNDCAK |
| FERMT3 | URP2\_HUMAN | Fermitin family homolog 3 | 407 | 406 | FERMT3 | URP2\_HUMAN | Fermitin family homolog 3 | 2C(125.0477) | NaN | X | NaN | NaN | NaN | NaN | 1.014081e+05 | NaN | 1.014081e+05 | NaN | GCEVVPDVNVSGQK |
| BLMH | BLMH\_HUMAN | Bleomycin hydrolase | 40 | 18 | BLMH | BLMH\_HUMAN | Bleomycin hydrolase | 23C(125.0477) | NaN | X | NaN | NaN | NaN | NaN | 1.013489e+05 | 1.013489e+05 | NaN | NaN | LNSDPQFVLAQNVGTTHDLLDICLKR |
| CHID1 | CHID1\_HUMAN | Chitinase domain-containing protein 1 | 281 | 280 | CHID1 | CHID1\_HUMAN | Chitinase domain-containing protein 1 | 2C(125.0477) | NaN | X | NaN | NaN | NaN | NaN | 1.011153e+05 | NaN | 1.011153e+05 | NaN | ACVQVLDPK |
| GDI2 | GDIB\_HUMAN | Rab GDP dissociation inhibitor beta | 282 | 279 | GDI2 | GDIB\_HUMAN | Rab GDP dissociation inhibitor beta | 4C(125.0477) | NaN | X | NaN | NaN | NaN | NaN | 1.009232e+05 | 1.009232e+05 | NaN | NaN | QLICDPSYVKDRVEK |
| YWHAZ | 1433Z\_HUMAN | 14-3-3 protein zeta/delta | 25 | 19 | YWHAZ | 1433Z\_HUMAN | 14-3-3 protein zeta/delta | 7C(125.0477) | NaN | X | NaN | NaN | NaN | NaN | 1.009147e+05 | NaN | 1.009147e+05 | NaN | YDDMAACMK |
| STX7 | STX7\_HUMAN | Syntaxin-7 | 28 | 25 | STX7 | STX7\_HUMAN | Syntaxin-7 | 4C(125.0477) | NaN | X | NaN | NaN | NaN | NaN | 1.008863e+05 | 6.167711e+04 | 3.920916e+04 | NaN | ITQCSVEIQR |
| UBR4 | UBR4\_HUMAN | E3 ubiquitin-protein ligase UBR4 | 4946 | 4939 | UBR4 | UBR4\_HUMAN | E3 ubiquitin-protein ligase UBR4 | 8C(125.0477) | NaN | X | NaN | NaN | NaN | NaN | 1.008239e+05 | 1.008239e+05 | NaN | NaN | HNTYLQECTGQR |
| UROD | DCUP\_HUMAN | Uroporphyrinogen decarboxylase | 294 | 291 | UROD | DCUP\_HUMAN | Uroporphyrinogen decarboxylase | 4C(15.9949) | NaN | NaN | X | NaN | NaN | NaN | 1.006717e+05 | 1.006717e+05 | NaN | NaN | ARECVGKTVTLQGNLDPCALYASEEEIGQLVK |
| NCAPD2 | CND1\_HUMAN | Condensin complex subunit 1 | 1229 | 1225 | NCAPD2 | CND1\_HUMAN | Condensin complex subunit 1 | 5C(125.0477) | NaN | X | NaN | NaN | NaN | NaN | 1.006124e+05 | 1.006124e+05 | NaN | NaN | DLAYCVSQLPLTER |
| PIP4K2A | PI42A\_HUMAN | Phosphatidylinositol 5-phosphate 4-kinase type-2 alpha | 94 | 90 | PIP4K2A | PI42A\_HUMAN | Phosphatidylinositol 5-phosphate 4-kinase type-2 alpha | 5C(125.0477) | NaN | X | NaN | NaN | NaN | NaN | 1.005833e+05 | 1.005833e+05 | NaN | NaN | FKEYCPMVFR |
| HSP90AA1 | HS90A\_HUMAN | Heat shock protein HSP 90-alpha | 529 | 514 | HSP90AA1 | HS90A\_HUMAN | Heat shock protein HSP 90-alpha | 16C(125.0477) | NaN | X | NaN | NaN | NaN | NaN | 1.005207e+05 | 1.005207e+05 | NaN | NaN | HGLEVIYMIEPIDEYCVQQLK |
| BLMH | BLMH\_HUMAN | Bleomycin hydrolase | 40 | 18 | BLMH | BLMH\_HUMAN | Bleomycin hydrolase | 23C(125.0477) | NaN | X | NaN | NaN | NaN | NaN | 1.004940e+05 | 1.004940e+05 | NaN | NaN | LNSDPQFVLAQNVGTTHDLLDICLK |
| GSTM3 | GSTM3\_HUMAN | Glutathione S-transferase Mu 3 | 91 | 87 | GSTM3 | GSTM3\_HUMAN | Glutathione S-transferase Mu 3 | 5C(125.0477) | NaN | X | NaN | NaN | NaN | NaN | 1.003655e+05 | 1.003655e+05 | NaN | NaN | KHNMCGETEEEK |
| ESD | ESTD\_HUMAN | S-formylglutathione hydrolase | 11 | 11 | ESD | ESTD\_HUMAN | S-formylglutathione hydrolase | 1C(125.0477) | NaN | X | NaN | NaN | NaN | NaN | 1.001066e+05 | NaN | 1.001066e+05 | NaN | CFGGLQK |
| MTHFD1 | C1TC\_HUMAN | C-1-tetrahydrofolate synthase, cytoplasmic | 195 | 176 | MTHFD1 | C1TC\_HUMAN | C-1-tetrahydrofolate synthase, cytoplasmic | 20C(125.0477) | NaN | X | NaN | NaN | NaN | NaN | 1.000570e+05 | 1.000570e+05 | NaN | NaN | IVGAPMHDLLLWNNATVTTCHSK |
| CUL2 | CUL2\_HUMAN | Cullin-2 | 721 | 721 | CUL2 | CUL2\_HUMAN | Cullin-2 | 1C(125.0477) | NaN | X | NaN | NaN | NaN | NaN | 1.000382e+05 | 1.000382e+05 | NaN | NaN | CIEVLIDK |
| PGP | PGP\_HUMAN | Glycerol-3-phosphate phosphatase | 193 | 187 | PGP | PGP\_HUMAN | Glycerol-3-phosphate phosphatase | 7C(125.0477) | NaN | X | NaN | NaN | NaN | NaN | 1.000295e+05 | 1.000295e+05 | NaN | NaN | YLQQPGCLLVGTNMDNR |
| TAOK3 | TAOK3\_HUMAN | Serine/threonine-protein kinase TAO3 | 727 | 721 | TAOK3 | TAOK3\_HUMAN | Serine/threonine-protein kinase TAO3 | 7C(125.0477) | NaN | X | NaN | NaN | NaN | NaN | 9.999370e+04 | 9.999370e+04 | NaN | NaN | KQFQDTCK |
| METTL7A | MET7A\_HUMAN | Putative methyltransferase-like protein 7A | 96 | 94 | METTL7A | MET7A\_HUMAN | Putative methyltransferase-like protein 7A | 3C(125.0477) | NaN | X | NaN | NaN | NaN | NaN | 9.998774e+04 | 4.478036e+04 | 5.520738e+04 | NaN | VTCIDPNPNFEK |
| MARK3 | MARK3\_HUMAN | MAP/microtubule affinity-regulating kinase 3 | 39 | 39 | MARK3 | MARK3\_HUMAN | MAP/microtubule affinity-regulating kinase 3 | 1C(15.9949) | NaN | NaN | X | NaN | NaN | NaN | 9.983025e+04 | 9.983025e+04 | NaN | NaN | CRNSIASCADEQPHIGNYR |
| TCEA1 | TCEA1\_HUMAN | Transcription elongation factor A protein 1 | 271 | 270 | TCEA1 | TCEA1\_HUMAN | Transcription elongation factor A protein 1 | 2C(125.0477) | NaN | X | NaN | NaN | NaN | NaN | 9.972638e+04 | 9.972638e+04 | NaN | NaN | NCTYTQVQTR |
| DHX15 | DHX15\_HUMAN | ATP-dependent RNA helicase DHX15 | 226 | 223 | DHX15 | DHX15\_HUMAN | ATP-dependent RNA helicase DHX15 | 4C(125.0477) | NaN | X | NaN | NaN | NaN | NaN | 9.971031e+04 | 9.971031e+04 | NaN | NaN | FEDCSSAK |
| CAST | ICAL\_HUMAN | Calpastatin | 408 | 397 | CAST | ICAL\_HUMAN | Calpastatin | 12C(125.0477) | NaN | X | NaN | NaN | NaN | NaN | 9.967588e+04 | NaN | 9.967588e+04 | NaN | AAAPAPVSEAVCR |
| TANGO2 | TNG2\_HUMAN | Transport and Golgi organization protein 2 homolog | 135 | 131 | TANGO2 | TNG2\_HUMAN | Transport and Golgi organization protein 2 homolog | 5C(125.0477) | NaN | X | NaN | NaN | NaN | NaN | 9.964550e+04 | NaN | 9.964550e+04 | NaN | GDVICYYGNR |
| ACTR2 | ARP2\_HUMAN | Actin-related protein 2 | 11 | 7 | ACTR2 | ARP2\_HUMAN | Actin-related protein 2 | 5C(125.0477) | NaN | X | NaN | NaN | NaN | NaN | 9.949990e+04 | 9.949990e+04 | NaN | NaN | KVVVCDNGTGFVK |
| DGKA | DGKA\_HUMAN | Diacylglycerol kinase alpha | 246 | 240 | DGKA | DGKA\_HUMAN | Diacylglycerol kinase alpha | 7C(125.0477) | NaN | X | NaN | NaN | NaN | NaN | 9.892489e+04 | 9.892489e+04 | NaN | NaN | YTVHDQCAMK |
| CCT7 | TCPH\_HUMAN | T-complex protein 1 subunit eta | 450 | 448 | CCT7 | TCPH\_HUMAN | T-complex protein 1 subunit eta | 3C(125.0477) | NaN | X | NaN | NaN | NaN | NaN | 9.890115e+04 | 9.890115e+04 | NaN | NaN | QLCDNAGFDATNILNK |
| PNPO | PNPO\_HUMAN | Pyridoxine-5'-phosphate oxidase | 156 | 148 | PNPO | PNPO\_HUMAN | Pyridoxine-5'-phosphate oxidase | 9C(125.0477) | NaN | X | NaN | NaN | NaN | NaN | 9.875855e+04 | 9.875855e+04 | NaN | NaN | KLPEEEAECYFHSRPK |
| HSPA8 | HSP7C\_HUMAN | Heat shock cognate 71 kDa protein | 574, 574 | 574 | HSPA8 | HSP7C\_HUMAN | Heat shock cognate 71 kDa protein | 1C(125.0477) 1C(57.0215) | X | X | NaN | NaN | NaN | NaN | 9.864249e+04 | 9.864249e+04 | NaN | NaN | CNEIINWLDKNQTAEKEEFEHQQK |
| CFH | CFAH\_HUMAN | Complement factor H | 1201 | 1193 | CFH | CFAH\_HUMAN | Complement factor H | 9C(57.0215) | X | NaN | NaN | NaN | NaN | NaN | 9.853694e+04 | 3.693661e+04 | 6.160033e+04 | NaN | TGESVEFVCK |
| EEF2 | EF2\_HUMAN | Elongation factor 2 | 728 | 728 | EEF2 | EF2\_HUMAN | Elongation factor 2 | 1C(125.0477) | NaN | X | NaN | NaN | NaN | NaN | 9.849878e+04 | 9.849878e+04 | NaN | NaN | CLYASVLTAQPR |
| EIF5 | IF5\_HUMAN | Eukaryotic translation initiation factor 5 | 122 | 116 | EIF5 | IF5\_HUMAN | Eukaryotic translation initiation factor 5 | 7C(15.9949) | NaN | NaN | X | NaN | NaN | NaN | 9.826516e+04 | 9.826516e+04 | NaN | NaN | QTIGNSCKACGYR |
| HSP90AB1 | HS90B\_HUMAN | Heat shock protein HSP 90-beta | 412 | 412 | HSP90AB1 | HS90B\_HUMAN | Heat shock protein HSP 90-beta | 1C(125.0477) | NaN | X | NaN | NaN | NaN | NaN | 9.808123e+04 | 9.808123e+04 | NaN | NaN | CLELFSELAEDKENYKK |
| GDI2 | GDIB\_HUMAN | Rab GDP dissociation inhibitor beta | 317 | 310 | GDI2 | GDIB\_HUMAN | Rab GDP dissociation inhibitor beta | 8C(125.0477) | NaN | X | NaN | NaN | NaN | NaN | 9.799731e+04 | 9.799731e+04 | NaN | NaN | NTNDANSCQIIIPQNQVNRK |
| GPHN | GEPH\_HUMAN | Gephyrin | 212 | 209 | GPHN | GEPH\_HUMAN | Gephyrin | 4C(125.0477) | NaN | X | NaN | NaN | NaN | NaN | 9.797997e+04 | 9.797997e+04 | NaN | NaN | GVQCEEEEEEKK |
| CASP6 | CASP6\_HUMAN | Caspase-6 | 264 | 260 | CASP6 | CASP6\_HUMAN | Caspase-6 | 5C(125.0477) | NaN | X | NaN | NaN | NaN | NaN | 9.797288e+04 | 9.797288e+04 | NaN | NaN | RVDFCKDPSAIGK |
| PSMD1 | PSMD1\_HUMAN | 26S proteasome non-ATPase regulatory subunit 1 | 141 | 141 | PSMD1 | PSMD1\_HUMAN | 26S proteasome non-ATPase regulatory subunit 1 | 1C(125.0477) | NaN | X | NaN | NaN | NaN | NaN | 9.794312e+04 | 9.794312e+04 | NaN | NaN | CLDDHKYK |
| EIF1AY | IF1AY\_HUMAN | Eukaryotic translation initiation factor 1A, Y-chromosomal | 51 | 47 | EIF1AY | IF1AY\_HUMAN | Eukaryotic translation initiation factor 1A, Y-chromosomal | 5C(125.0477) | NaN | X | NaN | NaN | NaN | NaN | 9.789248e+04 | 9.789248e+04 | NaN | NaN | LEALCFDGVKR |
| UBR1 | UBR1\_HUMAN | E3 ubiquitin-protein ligase UBR1 | 1204 | 1202 | UBR1 | UBR1\_HUMAN | E3 ubiquitin-protein ligase UBR1 | 3C(125.0477) | NaN | X | NaN | NaN | NaN | NaN | 9.789178e+04 | 9.789178e+04 | NaN | NaN | SLCNTVIPIIPLQPQK |
| TBCEL | TBCEL\_HUMAN | Tubulin-specific chaperone cofactor E-like protein | 121 | 120 | TBCEL | TBCEL\_HUMAN | Tubulin-specific chaperone cofactor E-like protein | 2C(125.0477) | NaN | X | NaN | NaN | NaN | NaN | 9.768202e+04 | 9.768202e+04 | NaN | NaN | TCAGSFSGVR |
| TRAF2 | TRAF2\_HUMAN | TNF receptor-associated factor 2 | 156 | 151 | TRAF2 | TRAF2\_HUMAN | TNF receptor-associated factor 2 | 6C(125.0477) | NaN | X | NaN | NaN | NaN | NaN | 9.767255e+04 | 9.767255e+04 | NaN | NaN | HLEHECPER |
| LEPROTL1 | LERL1\_HUMAN | Leptin receptor overlapping transcript-like 1 | 66 | 55 | LEPROTL1 | LERL1\_HUMAN | Leptin receptor overlapping transcript-like 1 | 12C(125.0477) | NaN | X | NaN | NaN | NaN | NaN | 9.762934e+04 | NaN | 9.762934e+04 | NaN | LVDDTDAMSNACK |
| PDXP | PLPP\_HUMAN | Chronophin | 26 | 19 | PDXP | PLPP\_HUMAN | Chronophin | 8C(125.0477) | NaN | X | NaN | NaN | NaN | NaN | 9.717797e+04 | 9.717797e+04 | NaN | NaN | AQGVLFDCDGVLWNGER |
| DCAF11 | DCA11\_HUMAN | DDB1- and CUL4-associated factor 11 | 460 | 445 | DCAF11 | DCA11\_HUMAN | DDB1- and CUL4-associated factor 11 | 16C(125.0477) | NaN | X | NaN | NaN | NaN | NaN | 9.698891e+04 | 9.698891e+04 | NaN | NaN | FSPIHSTGQQFIYSGCSTGK |
| MBNL1 | MBNL1\_HUMAN | Muscleblind-like protein 1 | 53 | 50 | MBNL1 | MBNL1\_HUMAN | Muscleblind-like protein 1 | 4C(125.0477) | NaN | X | NaN | NaN | NaN | NaN | 9.635991e+04 | 9.635991e+04 | NaN | NaN | VIACFDSLK |
| STAT5B | STA5B\_HUMAN | Signal transducer and activator of transcription 5B | 508 | 502 | STAT5B | STA5B\_HUMAN | Signal transducer and activator of transcription 5B | 7C(125.0477) | NaN | X | NaN | NaN | NaN | NaN | 9.619668e+04 | 9.619668e+04 | NaN | NaN | VLWPQLCEALNMK |
| IFI35 | IN35\_HUMAN | Interferon-induced 35 kDa protein | 193 | 192 | IFI35 | IN35\_HUMAN | Interferon-induced 35 kDa protein | 2C(125.0477) | NaN | X | NaN | NaN | NaN | NaN | 9.582786e+04 | 9.582786e+04 | NaN | NaN | LCQIGQFTVPLGGQQVPLR |
| ALAD | HEM2\_HUMAN | Delta-aminolevulinic acid dehydratase | 162 | 160 | ALAD | HEM2\_HUMAN | Delta-aminolevulinic acid dehydratase | 3C(125.0477) | NaN | X | NaN | NaN | NaN | NaN | 9.576479e+04 | 9.576479e+04 | NaN | NaN | AGCQVVAPSDMMDGR |
| RAP2A | RAP2A\_HUMAN | Ras-related protein Rap-2a | 140 | 133 | RAP2A | RAP2A\_HUMAN | Ras-related protein Rap-2a | 8C(125.0477) | NaN | X | NaN | NaN | NaN | NaN | 9.576296e+04 | 9.576296e+04 | NaN | NaN | ALAEEWGCPFMETSAK |
| ANKRD28 | ANR28\_HUMAN | Serine/threonine-protein phosphatase 6 regulatory ankyrin repeat subunit A | 122 | 122 | ANKRD28 | ANR28\_HUMAN | Serine/threonine-protein phosphatase 6 regulatory ankyrin repeat subunit A | 1C(125.0477) | NaN | X | NaN | NaN | NaN | NaN | 9.571388e+04 | 9.571388e+04 | NaN | NaN | CAEALVPLLSNVNVSDR |
| DDX1 | DDX1\_HUMAN | ATP-dependent RNA helicase DDX1 | 231 | 224 | DDX1 | DDX1\_HUMAN | ATP-dependent RNA helicase DDX1 | 8C(125.0477) | NaN | X | NaN | NaN | NaN | NaN | 9.566859e+04 | 9.566859e+04 | NaN | NaN | NQALFPACVLK |
| ANKFY1 | ANFY1\_HUMAN | Rabankyrin-5 | 460 | 447 | ANKFY1 | ANFY1\_HUMAN | Rabankyrin-5 | 14C(125.0477) | NaN | X | NaN | NaN | NaN | NaN | 9.564288e+04 | 9.564288e+04 | NaN | NaN | GSHTDAPDTATGNCLLQR |
| AMPD2 | AMPD2\_HUMAN | AMP deaminase 2 | 424 | 414 | AMPD2 | AMPD2\_HUMAN | AMP deaminase 2 | 11C(31.9988) | NaN | NaN | NaN | X | NaN | NaN | 9.561241e+04 | 9.561241e+04 | NaN | NaN | KVDTHIHASSCMNQK |
| AQP3 | AQP3\_HUMAN | Aquaporin-3 | 11 | 11 | AQP3 | AQP3\_HUMAN | Aquaporin-3 | 1C(125.0477) | NaN | X | NaN | NaN | NaN | NaN | 9.559985e+04 | 5.404159e+04 | 4.155826e+04 | NaN | CGEMLHIR |
| SRPRB | SRPRB\_HUMAN | Signal recognition particle receptor subunit beta | 246 | 241 | SRPRB | SRPRB\_HUMAN | Signal recognition particle receptor subunit beta | 6C(15.9949) | NaN | NaN | X | NaN | NaN | NaN | 9.550124e+04 | NaN | 9.550124e+04 | NaN | VEFLECSAK |
| ANXA1 | ANXA1\_HUMAN | Annexin A1 | 270 | 270 | ANXA1 | ANXA1\_HUMAN | Annexin A1 | 1C(125.0477) | NaN | X | NaN | NaN | NaN | NaN | 9.546538e+04 | 9.546538e+04 | NaN | NaN | CATSKPAFFAEK |
| ATE1 | ATE1\_HUMAN | Arginyl-tRNA--protein transferase 1 | 79 | 79 | ATE1 | ATE1\_HUMAN | Arginyl-tRNA--protein transferase 1 | 1C(125.0477) | NaN | X | NaN | NaN | NaN | NaN | 9.545184e+04 | 9.545184e+04 | NaN | NaN | CRPLQFQPSK |
| EHBP1L1 | EH1L1\_HUMAN | EH domain-binding protein 1-like protein 1 | 1073 | 1068 | EHBP1L1 | EH1L1\_HUMAN | EH domain-binding protein 1-like protein 1 | 6C(125.0477) | NaN | X | NaN | NaN | NaN | NaN | 9.544934e+04 | 9.544934e+04 | NaN | NaN | NGLAFCAILHR |
| ACTBL2 | ACTBL\_HUMAN | Beta-actin-like protein 2 | 53, 53 | 52 | ACTBL2 | ACTBL\_HUMAN | Beta-actin-like protein 2 | 2C(15.9949) 2C(31.9988) | NaN | NaN | X | X | NaN | NaN | 9.534984e+04 | NaN | NaN | 95349.8360 | DCYVGDEAQSK |
| GMPR2 | GMPR2\_HUMAN | GMP reductase 2 | 186 | 178 | GMPR2 | GMPR2\_HUMAN | GMP reductase 2 | 9C(125.0477) | NaN | X | NaN | NaN | NaN | NaN | 9.527177e+04 | 9.527177e+04 | NaN | NaN | VGIGPGSVCTTR |
| EPB41L2 | E41L2\_HUMAN | Band 4.1-like protein 2 | 488 | 487 | EPB41L2 | E41L2\_HUMAN | Band 4.1-like protein 2 | 2C(125.0477) | NaN | X | NaN | NaN | NaN | NaN | 9.462089e+04 | 9.462089e+04 | NaN | NaN | VCVEHHTFYR |
| FLOT1 | FLOT1\_HUMAN | Flotillin-1 | 34 | 29 | FLOT1 | FLOT1\_HUMAN | Flotillin-1 | 6C(125.0477) | NaN | X | NaN | NaN | NaN | NaN | 9.460320e+04 | 3.634063e+04 | 5.826257e+04 | NaN | VFVLPCIQQIQR |
| PRDX4 | PRDX4\_HUMAN | Peroxiredoxin-4 | 245 | 241 | PRDX4 | PRDX4\_HUMAN | Peroxiredoxin-4 | 5C(125.0477) | NaN | X | NaN | NaN | NaN | NaN | 9.431922e+04 | 9.431922e+04 | NaN | NaN | HGEVCPAGWKPGSETIIPDPAGK |
| GAPVD1 | GAPD1\_HUMAN | GTPase-activating protein and VPS9 domain-containing protein 1 | 1220 | 1220 | GAPVD1 | GAPD1\_HUMAN | GTPase-activating protein and VPS9 domain-containing protein 1 | 1C(125.0477) | NaN | X | NaN | NaN | NaN | NaN | 9.429319e+04 | 9.429319e+04 | NaN | NaN | CRQGLQTTQAHLER |
| SLC43A1 | LAT3\_HUMAN | Large neutral amino acids transporter small subunit 3 | 384 | 383 | SLC43A1 | LAT3\_HUMAN | Large neutral amino acids transporter small subunit 3 | 2C(125.0477) | NaN | X | NaN | NaN | NaN | NaN | 9.421446e+04 | NaN | 9.421446e+04 | NaN | DCVDAPTQGTVLGDAR |
| TCP1 | TCPA\_HUMAN | T-complex protein 1 subunit alpha | 76 | 74 | TCP1 | TCPA\_HUMAN | T-complex protein 1 subunit alpha | 3C(125.0477) | NaN | X | NaN | NaN | NaN | NaN | 9.397504e+04 | 2.748730e+04 | 6.648773e+04 | NaN | VLCELADLQDK |
| GPT | ALAT1\_HUMAN | Alanine aminotransferase 1 | 450 | 443 | GPT | ALAT1\_HUMAN | Alanine aminotransferase 1 | 8C(125.0477) | NaN | X | NaN | NaN | NaN | NaN | 9.392954e+04 | 9.392954e+04 | NaN | NaN | LLEETGICVVPGSGFGQR |
| EEF2 | EF2\_HUMAN | Elongation factor 2 | 567 | 560 | EEF2 | EF2\_HUMAN | Elongation factor 2 | 8C(125.0477) | NaN | X | NaN | NaN | NaN | NaN | 9.390575e+04 | 9.390575e+04 | NaN | NaN | DLEEDHACIPIK |
| GMDS | GMDS\_HUMAN | GDP-mannose 4,6 dehydratase | 92 | 82 | GMDS | GMDS\_HUMAN | GDP-mannose 4,6 dehydratase | 11C(125.0477) | NaN | X | NaN | NaN | NaN | NaN | 9.383810e+04 | 9.383810e+04 | NaN | NaN | LHYGDLTDSTCLVK |
| CLIC4 | CLIC4\_HUMAN | Chloride intracellular channel protein 4 | 234 | 228 | CLIC4 | CLIC4\_HUMAN | Chloride intracellular channel protein 4 | 7C(125.0477) | NaN | X | NaN | NaN | NaN | NaN | 9.381009e+04 | 9.381009e+04 | NaN | NaN | DEFTNTCPSDK |
| AHCY | SAHH\_HUMAN | Adenosylhomocysteinase | 195 | 189 | AHCY | SAHH\_HUMAN | Adenosylhomocysteinase | 7C(125.0477) | NaN | X | NaN | NaN | NaN | NaN | 9.375139e+04 | 8.113522e+04 | 1.261617e+04 | NaN | FDNLYGCR |
| CUL2 | CUL2\_HUMAN | Cullin-2 | 721 | 721 | CUL2 | CUL2\_HUMAN | Cullin-2 | 1C(125.0477) | NaN | X | NaN | NaN | NaN | NaN | 9.373838e+04 | 9.373838e+04 | NaN | NaN | CIEVLIDKQYIER |
| TATDN1 | TATD1\_HUMAN | Deoxyribonuclease TATDN1 | 76, 76 | 59 | TATDN1 | TATD1\_HUMAN | Deoxyribonuclease TATDN1 | 18C(15.9949) 18C(31.9988) | NaN | NaN | X | X | NaN | NaN | 9.373705e+04 | 9.373705e+04 | NaN | NaN | DALHLAQTNGMFFSTVGCHPTRCGEFEK |
| PGAM1 | PGAM1\_HUMAN | Phosphoglycerate mutase 1 | 55 | 47 | PGAM1 | PGAM1\_HUMAN | Phosphoglycerate mutase 1 | 9C(125.0477) | NaN | X | NaN | NaN | NaN | NaN | 9.358460e+04 | 9.358460e+04 | NaN | NaN | DAGYEFDICFTSVQK |
| CAST | ICAL\_HUMAN | Calpastatin | 328 | 325 | CAST | ICAL\_HUMAN | Calpastatin | 4C(125.0477) | NaN | X | NaN | NaN | NaN | NaN | 9.357778e+04 | 9.357778e+04 | NaN | NaN | LEKCGEDDETIPSEYR |
| PAAF1 | PAAF1\_HUMAN | Proteasomal ATPase-associated factor 1 | 265 | 263 | PAAF1 | PAAF1\_HUMAN | Proteasomal ATPase-associated factor 1 | 3C(125.0477) | NaN | X | NaN | NaN | NaN | NaN | 9.338004e+04 | 9.338004e+04 | NaN | NaN | LQCLGLQSR |
| PNPO | PNPO\_HUMAN | Pyridoxine-5'-phosphate oxidase | 156 | 149 | PNPO | PNPO\_HUMAN | Pyridoxine-5'-phosphate oxidase | 8C(125.0477) | NaN | X | NaN | NaN | NaN | NaN | 9.323425e+04 | 9.323425e+04 | NaN | NaN | LPEEEAECYFHSRPK |
| ADD2 | ADDB\_HUMAN | Beta-adducin | 177 | 174 | ADD2 | ADDB\_HUMAN | Beta-adducin | 4C(125.0477) | NaN | X | NaN | NaN | NaN | NaN | 9.308502e+04 | 9.308502e+04 | NaN | NaN | GVSCSEVTASSLIK |
| ASAP1 | ASAP1\_HUMAN | Arf-GAP with SH3 domain, ANK repeat and PH domain-containing protein 1 | 740 | 733 | ASAP1 | ASAP1\_HUMAN | Arf-GAP with SH3 domain, ANK repeat and PH domain-containing protein 1 | 8C(125.0477) | NaN | X | NaN | NaN | NaN | NaN | 9.295425e+04 | 9.295425e+04 | NaN | NaN | SPRPQSFCHSSSISPQDK |
| ENDOD1 | ENDD1\_HUMAN | Endonuclease domain-containing 1 protein | 34 | 23 | ENDOD1 | ENDD1\_HUMAN | Endonuclease domain-containing 1 protein | 12C(125.0477) | NaN | X | NaN | NaN | NaN | NaN | 9.294565e+04 | 7.687881e+04 | 1.606684e+04 | 0.0000 | LVGEEEAGFGECDK |
| MDH1 | MDHC\_HUMAN | Malate dehydrogenase, cytoplasmic | 137 | 126 | MDH1 | MDHC\_HUMAN | Malate dehydrogenase, cytoplasmic | 12C(125.0477) | NaN | X | NaN | NaN | NaN | NaN | 9.285716e+04 | NaN | 8.340959e+04 | 9447.5700 | VIVVGNPANTNCLTASK |
| IPO11 | IPO11\_HUMAN | Importin-11 | 366 | 354 | IPO11 | IPO11\_HUMAN | Importin-11 | 13C(125.0477) | NaN | X | NaN | NaN | NaN | NaN | 9.282929e+04 | 9.282929e+04 | NaN | NaN | MAFFTYPTLTEICR |
| THOP1 | THOP1\_HUMAN | Thimet oligopeptidase | 644 | 643 | THOP1 | THOP1\_HUMAN | Thimet oligopeptidase | 2C(125.0477) | NaN | X | NaN | NaN | NaN | NaN | 9.272402e+04 | 9.272402e+04 | NaN | NaN | SCILRPGGSEDASAMLR |
| WNK1 | WNK1\_HUMAN | Serine/threonine-protein kinase WNK1 | 1135 | 1132 | WNK1 | WNK1\_HUMAN | Serine/threonine-protein kinase WNK1 | 4C(125.0477) | NaN | X | NaN | NaN | NaN | NaN | 9.255887e+04 | NaN | 9.255887e+04 | NaN | VVECQLETHNR |
| ALDH9A1 | AL9A1\_HUMAN | 4-trimethylaminobutyraldehyde dehydrogenase | 267 | 259 | ALDH9A1 | AL9A1\_HUMAN | 4-trimethylaminobutyraldehyde dehydrogenase | 9C(125.0477) | NaN | X | NaN | NaN | NaN | NaN | 9.252428e+04 | 9.252428e+04 | NaN | NaN | SPLIIFSDCDMNNAVK |
| CLTC | CLH1\_HUMAN | Clathrin heavy chain 1 | 918 | 913 | CLTC | CLH1\_HUMAN | Clathrin heavy chain 1 | 6C(125.0477) | NaN | X | NaN | NaN | NaN | NaN | 9.243202e+04 | 8.358666e+04 | 8.845364e+03 | NaN | DPHLACVAYER |
| ALB | ALBU\_HUMAN | Albumin | 500 | 500 | ALB | ALBU\_HUMAN | Albumin | 1C(125.0477) | NaN | X | NaN | NaN | NaN | NaN | 9.239318e+04 | NaN | NaN | 92393.1800 | CCTESLVNR |
| FHIT | FHIT\_HUMAN | Bis(5'-adenosyl)-triphosphatase | 39 | 29 | FHIT | FHIT\_HUMAN | Bis(5'-adenosyl)-triphosphatase | 11C(125.0477) | NaN | X | NaN | NaN | NaN | NaN | 9.229727e+04 | 9.229727e+04 | NaN | NaN | KPVVPGHVLVCPLRPVER |
| IGHA1 | IGHA1\_HUMAN | Immunoglobulin heavy constant alpha 1 | 204, 204 | 201 | IGHA1 | IGHA1\_HUMAN | Immunoglobulin heavy constant alpha 1 | 4C(125.0477) 4C(57.0215) | X | X | NaN | NaN | NaN | NaN | 9.215225e+04 | NaN | 9.215225e+04 | NaN | TFTCTAAYPESK |
| KRT6A | K2C6A\_HUMAN | Keratin, type II cytoskeletal 6A | 77 | 70 | KRT6A | K2C6A\_HUMAN | Keratin, type II cytoskeletal 6A | 8C(125.0477) | NaN | X | NaN | NaN | NaN | NaN | 9.203419e+04 | 4.648308e+04 | 4.555111e+04 | NaN | ISIGGGSCAISGGYGSR |
| CTPS2 | PYRG2\_HUMAN | CTP synthase 2 | 176 | 172 | CTPS2 | PYRG2\_HUMAN | CTP synthase 2 | 5C(125.0477) | NaN | X | NaN | NaN | NaN | NaN | 9.201797e+04 | 9.201797e+04 | NaN | NaN | RENFCNIHVSLVPQLSATGEQK |
| PSMB7 | PSB7\_HUMAN | Proteasome subunit beta type-7 | 74 | 63 | PSMB7 | PSB7\_HUMAN | Proteasome subunit beta type-7 | 12C(125.0477) | NaN | X | NaN | NaN | NaN | NaN | 9.188782e+04 | 9.188782e+04 | NaN | NaN | ATEGMVVADKNCSK |
| MDH1 | MDHC\_HUMAN | Malate dehydrogenase, cytoplasmic | 154 | 126 | MDH1 | MDHC\_HUMAN | Malate dehydrogenase, cytoplasmic | 29C(15.9949) | NaN | NaN | X | NaN | NaN | NaN | 9.175879e+04 | 9.175879e+04 | NaN | NaN | VIVVGNPANTNCLTASKSAPSIPKENFSCLTR |
| CUL1 | CUL1\_HUMAN | Cullin-1 | 752 | 752 | CUL1 | CUL1\_HUMAN | Cullin-1 | 1C(125.0477) | NaN | X | NaN | NaN | NaN | NaN | 9.170284e+04 | 9.170284e+04 | NaN | NaN | CIDILIEKEYLER |
| HBA1 | HBA\_HUMAN | Hemoglobin subunit alpha | 105 | 94 | HBA1 | HBA\_HUMAN | Hemoglobin subunit alpha | 12C(125.0477) | NaN | X | NaN | NaN | NaN | NaN | 9.148586e+04 | NaN | 7.584288e+04 | 15642.9810 | VDPVNFKLLSHCLLVTLAAHLPAEFTPAVHASLDKFLASVSTVLTSK |
| ATP6V1B2 | VATB2\_HUMAN | V-type proton ATPase subunit B, brain isoform | 425 | 413 | ATP6V1B2 | VATB2\_HUMAN | V-type proton ATPase subunit B, brain isoform | 13C(125.0477) | NaN | X | NaN | NaN | NaN | NaN | 9.142216e+04 | 9.142216e+04 | NaN | NaN | KDHADVSNQLYACYAIGK |
| CLTC | CLH1\_HUMAN | Clathrin heavy chain 1 | 1528 | 1523 | CLTC | CLH1\_HUMAN | Clathrin heavy chain 1 | 6C(125.0477) | NaN | X | NaN | NaN | NaN | NaN | 9.138334e+04 | 9.138334e+04 | NaN | NaN | QSVELCK |
| CIAO1 | CIAO1\_HUMAN | Probable cytosolic iron-sulfur protein assembly protein CIAO1 | 97 | 91 | CIAO1 | CIAO1\_HUMAN | Probable cytosolic iron-sulfur protein assembly protein CIAO1 | 7C(125.0477) | NaN | X | NaN | NaN | NaN | NaN | 9.133209e+04 | 9.133209e+04 | NaN | NaN | NQDDFECVTTLEGHENEVK |
| PSMC2 | PRS7\_HUMAN | 26S proteasome regulatory subunit 7 | 236 | 233 | PSMC2 | PRS7\_HUMAN | 26S proteasome regulatory subunit 7 | 4C(125.0477) | NaN | X | NaN | NaN | NaN | NaN | 9.117405e+04 | NaN | 9.117405e+04 | NaN | TDACFIR |
| FTH1 | FRIH\_HUMAN | Ferritin heavy chain | 131 | 121 | FTH1 | FRIH\_HUMAN | Ferritin heavy chain | 11C(125.0477) | NaN | X | NaN | NaN | NaN | NaN | 9.116841e+04 | 9.116841e+04 | NaN | NaN | LATDKNDPHLCDFIETHYLNEQVK |
| DCAF11 | DCA11\_HUMAN | DDB1- and CUL4-associated factor 11 | 191 | 183 | DCAF11 | DCA11\_HUMAN | DDB1- and CUL4-associated factor 11 | 9C(125.0477) | NaN | X | NaN | NaN | NaN | NaN | 9.114212e+04 | 9.114212e+04 | NaN | NaN | DGQIFMSACQDQTIR |
| POTEI | POTEI\_HUMAN | POTE ankyrin domain family member I | 917, 917 | 916 | POTEI | POTEI\_HUMAN | POTE ankyrin domain family member I | 2C(15.9949) 2C(31.9988) | NaN | NaN | X | X | NaN | NaN | 9.114076e+04 | NaN | 9.114076e+04 | NaN | LCYVALDFEQEMAMAASSSSLEK |
| P4HB | PDIA1\_HUMAN | Protein disulfide-isomerase | 312 | 310 | P4HB | PDIA1\_HUMAN | Protein disulfide-isomerase | 3C(125.0477) | NaN | X | NaN | NaN | NaN | NaN | 9.110713e+04 | 5.478641e+04 | 3.632072e+04 | NaN | EECPAVR |
| SELENBP1 | SBP1\_HUMAN | Methanethiol oxidase | 80, 80, 83, 83 | 63 | SELENBP1 | SBP1\_HUMAN | Methanethiol oxidase | 18C(-34.0809) 18C(31.9988) 21C(125.0477) 21C(31.9988) | NaN | X | NaN | X | X | NaN | 9.108309e+04 | 9.108309e+04 | NaN | NaN | LPMPNLKDELHHSGWNTCSSCFGDSTK |
| GUK1 | KGUA\_HUMAN | Guanylate kinase | 98 | 97 | GUK1 | KGUA\_HUMAN | Guanylate kinase | 2C(125.0477) | NaN | X | NaN | NaN | NaN | NaN | 9.094913e+04 | 9.094913e+04 | NaN | NaN | ICVLDVDLQGVR |
| RAB4B | RAB4B\_HUMAN | Ras-related protein Rab-4B | 119 | 108 | RAB4B | RAB4B\_HUMAN | Ras-related protein Rab-4B | 12C(125.0477) | NaN | X | NaN | NaN | NaN | NaN | 9.094328e+04 | 9.094328e+04 | NaN | NaN | TLASPNIVVILCGNK |
| NAXD | NNRD\_HUMAN | ATP-dependent (S)-NAD(P)H-hydrate dehydratase | 316 | 315 | NAXD | NNRD\_HUMAN | ATP-dependent (S)-NAD(P)H-hydrate dehydratase | 2C(125.0477) | NaN | X | NaN | NaN | NaN | NaN | 9.088333e+04 | 9.088333e+04 | NaN | NaN | QCNHQAFQK |
| ANKRD28 | ANR28\_HUMAN | Serine/threonine-protein phosphatase 6 regulatory ankyrin repeat subunit A | 598 | 594 | ANKRD28 | ANR28\_HUMAN | Serine/threonine-protein phosphatase 6 regulatory ankyrin repeat subunit A | 5C(125.0477) | NaN | X | NaN | NaN | NaN | NaN | 9.085888e+04 | 9.085888e+04 | NaN | NaN | GHVECVDVLINQGASILVK |
| DDAH2 | DDAH2\_HUMAN | N(G),N(G)-dimethylarginine dimethylaminohydrolase 2 | 276 | 268 | DDAH2 | DDAH2\_HUMAN | N(G),N(G)-dimethylarginine dimethylaminohydrolase 2 | 9C(125.0477) | NaN | X | NaN | NaN | NaN | NaN | 9.084903e+04 | 9.084903e+04 | NaN | NaN | AGAGLSSLCLVLSTRPHS |
| LANCL1 | LANC1\_HUMAN | Glutathione S-transferase LANCL1 | 183 | 174 | LANCL1 | LANC1\_HUMAN | Glutathione S-transferase LANCL1 | 10C(125.0477) | NaN | X | NaN | NaN | NaN | NaN | 9.083198e+04 | 9.083198e+04 | NaN | NaN | IPQSHIQQICETILTSGENLAR |
| KPNA4 | IMA3\_HUMAN | Importin subunit alpha-3 | 228 | 219 | KPNA4 | IMA3\_HUMAN | Importin subunit alpha-3 | 10C(125.0477) | NaN | X | NaN | NaN | NaN | NaN | 9.076937e+04 | 9.076937e+04 | NaN | NaN | NVTWVMVNLCR |
| VPS35 | VPS35\_HUMAN | Vacuolar protein sorting-associated protein 35 | 640 | 640 | VPS35 | VPS35\_HUMAN | Vacuolar protein sorting-associated protein 35 | 1C(125.0477) | NaN | X | NaN | NaN | NaN | NaN | 9.058841e+04 | 9.058841e+04 | NaN | NaN | CFSEENHEPLR |
| PRKAR2B | KAP3\_HUMAN | cAMP-dependent protein kinase type II-beta regulatory subunit | 205 | 205 | PRKAR2B | KAP3\_HUMAN | cAMP-dependent protein kinase type II-beta regulatory subunit | 1C(15.9949) | NaN | NaN | X | NaN | NaN | NaN | 9.042588e+04 | 9.042588e+04 | NaN | NaN | CDGVGRCVGNYDNR |
| MARK3 | MARK3\_HUMAN | MAP/microtubule affinity-regulating kinase 3 | 280 | 272 | MARK3 | MARK3\_HUMAN | MAP/microtubule affinity-regulating kinase 3 | 9C(125.0477) | NaN | X | NaN | NaN | NaN | NaN | 9.030859e+04 | 9.030859e+04 | NaN | NaN | IPFYMSTDCENLLK |
| UBE4A | UBE4A\_HUMAN | Ubiquitin conjugation factor E4 A | 202 | 191 | UBE4A | UBE4A\_HUMAN | Ubiquitin conjugation factor E4 A | 12C(125.0477) | NaN | X | NaN | NaN | NaN | NaN | 9.023372e+04 | 9.023372e+04 | NaN | NaN | VPENLLPFAVQCR |
| DDB1 | DDB1\_HUMAN | DNA damage-binding protein 1 | 128 | 112 | DDB1 | DDB1\_HUMAN | DNA damage-binding protein 1 | 17C(125.0477) | NaN | X | NaN | NaN | NaN | NaN | 9.020718e+04 | 9.020718e+04 | NaN | NaN | IGRPSETGIIGIIDPECR |
| DGKA | DGKA\_HUMAN | Diacylglycerol kinase alpha | 707 | 692 | DGKA | DGKA\_HUMAN | Diacylglycerol kinase alpha | 16C(125.0477) | NaN | X | NaN | NaN | NaN | NaN | 9.016803e+04 | 9.016803e+04 | NaN | NaN | TLPMQIDGEPWMQTPCTIK |
| GSTO1 | GSTO1\_HUMAN | Glutathione S-transferase omega-1 | 32 | 31 | GSTO1 | GSTO1\_HUMAN | Glutathione S-transferase omega-1 | 2C(125.0477) | NaN | X | NaN | NaN | NaN | NaN | 9.012138e+04 | NaN | 9.012138e+04 | NaN | FCPFAER |
| CUL2 | CUL2\_HUMAN | Cullin-2 | 395 | 394 | CUL2 | CUL2\_HUMAN | Cullin-2 | 2C(125.0477) | NaN | X | NaN | NaN | NaN | NaN | 9.001336e+04 | 5.846914e+04 | 3.154422e+04 | NaN | YCDNLLK |
| CMBL | CMBL\_HUMAN | Carboxymethylenebutenolidase homolog | 198 | 194 | CMBL | CMBL\_HUMAN | Carboxymethylenebutenolidase homolog | 5C(125.0477) | NaN | X | NaN | NaN | NaN | NaN | 8.983560e+04 | 8.983560e+04 | NaN | NaN | LKEHCKVEYQIK |
| GNAS | GNAS1\_HUMAN | Guanine nucleotide-binding protein G(s) subunit alpha isoforms XLas | 1022 | 1018 | GNAS | GNAS1\_HUMAN | Guanine nucleotide-binding protein G(s) subunit alpha isoforms XLas | 5C(125.0477) | NaN | X | NaN | NaN | NaN | NaN | 8.979945e+04 | 8.979945e+04 | NaN | NaN | VFNDCRDIIQR |
| METTL7A | MET7A\_HUMAN | Putative methyltransferase-like protein 7A | 92 | 87 | METTL7A | MET7A\_HUMAN | Putative methyltransferase-like protein 7A | 6C(15.9949) | NaN | NaN | X | NaN | NaN | NaN | 8.975149e+04 | 8.975149e+04 | NaN | NaN | FYPPGCRVTCIDPNPNFEK |
| VCP | TERA\_HUMAN | Transitional endoplasmic reticulum ATPase | 209 | 192 | VCP | TERA\_HUMAN | Transitional endoplasmic reticulum ATPase | 18C(125.0477) | NaN | X | NaN | NaN | NaN | NaN | 8.950968e+04 | 7.145173e+04 | 1.805794e+04 | NaN | EDEEESLNEVGYDDIGGCR |
| CARHSP1 | CHSP1\_HUMAN | Calcium-regulated heat-stable protein 1 | 112 | 111 | CARHSP1 | CHSP1\_HUMAN | Calcium-regulated heat-stable protein 1 | 2C(125.0477) | NaN | X | NaN | NaN | NaN | NaN | 8.950192e+04 | 8.950192e+04 | NaN | NaN | MCSIPPKNEK |
| KRT81 | KRT81\_HUMAN | Keratin, type II cuticular Hb1 | 427, 427, 418 | 417 | KRT81 | KRT81\_HUMAN | Keratin, type II cuticular Hb1 | 11C(-34.0809) 11C(125.0477) 2C(57.0215) | X | X | NaN | NaN | X | NaN | 8.921403e+04 | 8.921403e+04 | NaN | NaN | LCEGIGAVNVCVSSSR |
| ETF1 | ERF1\_HUMAN | Eukaryotic peptide chain release factor subunit 1 | 335 | 331 | ETF1 | ERF1\_HUMAN | Eukaryotic peptide chain release factor subunit 1 | 5C(125.0477) | NaN | X | NaN | NaN | NaN | NaN | 8.899791e+04 | 8.899791e+04 | NaN | NaN | YVLHCQGTEEEK |
| AVEN | AVEN\_HUMAN | Cell death regulator Aven | 193 | 186 | AVEN | AVEN\_HUMAN | Cell death regulator Aven | 8C(125.0477) | NaN | X | NaN | NaN | NaN | NaN | 8.896391e+04 | 8.896391e+04 | NaN | NaN | ALQELPLCLR |
| HPRT1 | HPRT\_HUMAN | Hypoxanthine-guanine phosphoribosyltransferase | 106 | 104 | HPRT1 | HPRT\_HUMAN | Hypoxanthine-guanine phosphoribosyltransferase | 3C(125.0477) | NaN | X | NaN | NaN | NaN | NaN | 8.886218e+04 | NaN | 8.886218e+04 | NaN | SYCNDQSTGDIK |
| CISD2 | CISD2\_HUMAN | CDGSH iron-sulfur domain-containing protein 2 | 92 | 82 | CISD2 | CISD2\_HUMAN | CDGSH iron-sulfur domain-containing protein 2 | 11C(125.0477) | NaN | X | NaN | NaN | NaN | NaN | 8.879616e+04 | 8.879616e+04 | NaN | NaN | VVNEINIEDLCLTK |
| SPTB | SPTB1\_HUMAN | Spectrin beta chain, erythrocytic | 1892, 1892 | 1878 | SPTB | SPTB1\_HUMAN | Spectrin beta chain, erythrocytic | 15C(125.0477) 15C(31.9988) | NaN | X | NaN | X | NaN | NaN | 8.879034e+04 | NaN | 8.879034e+04 | NaN | EQEVSAAWQALLDACAGR |
| KRT1 | K2C1\_HUMAN | Keratin, type II cytoskeletal 1 | 497, 497 | 493 | KRT1 | K2C1\_HUMAN | Keratin, type II cytoskeletal 1 | 5C(125.0477) 5C(15.9949) | NaN | X | X | NaN | NaN | NaN | 8.868779e+04 | 8.868779e+04 | NaN | NaN | MSGECAPNVSVSVSTSHTTISGGGSR |
| PPIP5K2 | VIP2\_HUMAN | Inositol hexakisphosphate and diphosphoinositol-pentakisphosphate kinase 2 | 338 | 334 | PPIP5K2 | VIP2\_HUMAN | Inositol hexakisphosphate and diphosphoinositol-pentakisphosphate kinase 2 | 5C(125.0477) | NaN | X | NaN | NaN | NaN | NaN | 8.850669e+04 | 8.850669e+04 | NaN | NaN | YYDDCAK |
| ADD3 | ADDG\_HUMAN | Gamma-adducin | 73 | 69 | ADD3 | ADDG\_HUMAN | Gamma-adducin | 5C(125.0477) | NaN | X | NaN | NaN | NaN | NaN | 8.842095e+04 | NaN | 8.842095e+04 | NaN | EDLECLIQEQMK |
| ARPC4 | ARPC4\_HUMAN | Actin-related protein 2/3 complex subunit 4 | 21 | 14 | ARPC4 | ARPC4\_HUMAN | Actin-related protein 2/3 complex subunit 4 | 8C(125.0477) | NaN | X | NaN | NaN | NaN | NaN | 8.824364e+04 | 8.824364e+04 | NaN | NaN | ATLQAALCLENFSSQVVER |
| HSPA1A | HS71A\_HUMAN | Heat shock 70 kDa protein 1A | 603, 603 | 597 | HSPA1A | HS71A\_HUMAN | Heat shock 70 kDa protein 1A | 7C(125.0477) 7C(15.9949) | NaN | X | X | NaN | NaN | NaN | 8.793906e+04 | 8.793906e+04 | NaN | NaN | KELEQVCNPIISGLYQGAGGPGPGGFGAQGPK |
| IGHG1 | IGHG1\_HUMAN | Immunoglobulin heavy constant gamma 1 | 250 | 244 | IGHG1 | IGHG1\_HUMAN | Immunoglobulin heavy constant gamma 1 | 7C(125.0477) | NaN | X | NaN | NaN | NaN | NaN | 8.781067e+04 | 8.781067e+04 | NaN | NaN | NQVSLTCLVK |
| MYH9 | MYH9\_HUMAN | Myosin-9 | 1379 | 1373 | MYH9 | MYH9\_HUMAN | Myosin-9 | 7C(125.0477) | NaN | X | NaN | NaN | NaN | NaN | 8.775344e+04 | 8.775344e+04 | NaN | NaN | MEDSVGCLETAEEVKR |
| LDHB | LDHB\_HUMAN | L-lactate dehydrogenase B chain | 36 | 8 | LDHB | LDHB\_HUMAN | L-lactate dehydrogenase B chain | 29C(125.0477) | NaN | X | NaN | NaN | NaN | NaN | 8.774511e+04 | 8.774511e+04 | NaN | NaN | LIAPVAEEEATVPNNKITVVGVGQVGMACAISILGK |
| UBE2D2 | UB2D2\_HUMAN | Ubiquitin-conjugating enzyme E2 D2 | 85 | 73 | UBE2D2 | UB2D2\_HUMAN | Ubiquitin-conjugating enzyme E2 D2 | 13C(125.0477) | NaN | X | NaN | NaN | NaN | NaN | 8.737964e+04 | 8.737964e+04 | NaN | NaN | IYHPNINSNGSICLDILR |
| ATP1A1 | AT1A1\_HUMAN | Sodium/potassium-transporting ATPase subunit alpha-1 | 705 | 699 | ATP1A1 | AT1A1\_HUMAN | Sodium/potassium-transporting ATPase subunit alpha-1 | 7C(125.0477) | NaN | X | NaN | NaN | NaN | NaN | 8.736706e+04 | NaN | 8.736706e+04 | NaN | LIIVEGCQR |
| ALDH16A1 | A16A1\_HUMAN | Aldehyde dehydrogenase family 16 member A1 | 39 | 39 | ALDH16A1 | A16A1\_HUMAN | Aldehyde dehydrogenase family 16 member A1 | 1C(125.0477) | NaN | X | NaN | NaN | NaN | NaN | 8.730937e+04 | 8.730937e+04 | NaN | NaN | CLGHYVNGK |
| UROD | DCUP\_HUMAN | Uroporphyrinogen decarboxylase | 35 | 22 | UROD | DCUP\_HUMAN | Uroporphyrinogen decarboxylase | 14C(125.0477) | NaN | X | NaN | NaN | NaN | NaN | 8.723086e+04 | 8.723086e+04 | NaN | NaN | AAWGEETDYTPVWCMR |
| ELAC1 | RNZ1\_HUMAN | Zinc phosphodiesterase ELAC protein 1 | 249, 254 | 248 | ELAC1 | RNZ1\_HUMAN | Zinc phosphodiesterase ELAC protein 1 | 2C(31.9988) 7C(15.9949) | NaN | NaN | X | X | NaN | NaN | 8.712581e+04 | 8.712581e+04 | NaN | NaN | ICILGDCSGVVGDGGVK |
| GLUL | GLNA\_HUMAN | Glutamine synthetase | 346 | 341 | GLUL | GLNA\_HUMAN | Glutamine synthetase | 6C(125.0477) | NaN | X | NaN | NaN | NaN | NaN | 8.707685e+04 | 8.707685e+04 | NaN | NaN | RPSANCDPFSVTEALIR |
| HUWE1 | HUWE1\_HUMAN | E3 ubiquitin-protein ligase HUWE1 | 467 | 460 | HUWE1 | HUWE1\_HUMAN | E3 ubiquitin-protein ligase HUWE1 | 8C(125.0477) | NaN | X | NaN | NaN | NaN | NaN | 8.707422e+04 | 8.707422e+04 | NaN | NaN | LEHEVDLCRK |
| PPP1CA | PP1A\_HUMAN | Serine/threonine-protein phosphatase PP1-alpha catalytic subunit | 245 | 239 | PPP1CA | PP1A\_HUMAN | Serine/threonine-protein phosphatase PP1-alpha catalytic subunit | 7C(125.0477) | NaN | X | NaN | NaN | NaN | NaN | 8.700262e+04 | 8.700262e+04 | NaN | NaN | HDLDLICR |
| CUL4A | CUL4A\_HUMAN | Cullin-4A | 289 | 267 | CUL4A | CUL4A\_HUMAN | Cullin-4A | 23C(125.0477) | NaN | X | NaN | NaN | NaN | NaN | 8.692131e+04 | 8.692131e+04 | NaN | NaN | LEEEGDRVITYLDHSTQKPLIACVEK |
| ERI3 | ERI3\_HUMAN | ERI1 exoribonuclease 3 | 245 | 240 | ERI3 | ERI3\_HUMAN | ERI1 exoribonuclease 3 | 6C(125.0477) | NaN | X | NaN | NaN | NaN | NaN | 8.685950e+04 | 8.685950e+04 | NaN | NaN | SIFVTCGDWDLK |
| EIF3M | EIF3M\_HUMAN | Eukaryotic translation initiation factor 3 subunit M | 175 | 167 | EIF3M | EIF3M\_HUMAN | Eukaryotic translation initiation factor 3 subunit M | 9C(125.0477) | NaN | X | NaN | NaN | NaN | NaN | 8.673117e+04 | 8.673117e+04 | NaN | NaN | LLYEALVDCKK |
| HSPH1 | HS105\_HUMAN | Heat shock protein 105 kDa | 796 | 791 | HSPH1 | HS105\_HUMAN | Heat shock protein 105 kDa | 6C(125.0477) | NaN | X | NaN | NaN | NaN | NaN | 8.661958e+04 | 8.661958e+04 | NaN | NaN | ELNNTCEPVVTQPKPK |
| EIF5 | IF5\_HUMAN | Eukaryotic translation initiation factor 5 | 59 | 56 | EIF5 | IF5\_HUMAN | Eukaryotic translation initiation factor 5 | 4C(125.0477) | NaN | X | NaN | NaN | NaN | NaN | 8.654331e+04 | 8.654331e+04 | NaN | NaN | YFGCELGAQTQFDVK |
| TPI1 | TPIS\_HUMAN | Triosephosphate isomerase | 87 | 86 | TPI1 | TPIS\_HUMAN | Triosephosphate isomerase | 2C(125.0477) | NaN | X | NaN | NaN | NaN | NaN | 8.643108e+04 | 8.643108e+04 | NaN | NaN | DCGATWVVLGHSERR |
| LANCL2 | LANC2\_HUMAN | LanC-like protein 2 | 187 | 184 | LANCL2 | LANC2\_HUMAN | LanC-like protein 2 | 4C(125.0477) | NaN | X | NaN | NaN | NaN | NaN | 8.635858e+04 | 6.918672e+04 | 1.717186e+04 | NaN | SVVCQESDLPDELLYGR |
| RNF123 | RN123\_HUMAN | E3 ubiquitin-protein ligase RNF123 | 461 | 444 | RNF123 | RN123\_HUMAN | E3 ubiquitin-protein ligase RNF123 | 18C(125.0477) | NaN | X | NaN | NaN | NaN | NaN | 8.578630e+04 | 8.578630e+04 | NaN | NaN | VEEAGLQELIPTTWWPHCSSR |
| SPTB | SPTB1\_HUMAN | Spectrin beta chain, erythrocytic | 1167 | 1161 | SPTB | SPTB1\_HUMAN | Spectrin beta chain, erythrocytic | 7C(125.0477) | NaN | X | NaN | NaN | NaN | NaN | 8.556382e+04 | 8.556382e+04 | NaN | NaN | SHTLAQCLGFQEFQKDAK |
| ANKRD28 | ANR28\_HUMAN | Serine/threonine-protein phosphatase 6 regulatory ankyrin repeat subunit A | 487 | 487 | ANKRD28 | ANR28\_HUMAN | Serine/threonine-protein phosphatase 6 regulatory ankyrin repeat subunit A | 1C(125.0477) | NaN | X | NaN | NaN | NaN | NaN | 8.537873e+04 | 8.537873e+04 | NaN | NaN | CLEYLLR |
| ATG3 | ATG3\_HUMAN | Ubiquitin-like-conjugating enzyme ATG3 | 182 | 177 | ATG3 | ATG3\_HUMAN | Ubiquitin-like-conjugating enzyme ATG3 | 6C(125.0477) | NaN | X | NaN | NaN | NaN | NaN | 8.532512e+04 | 5.950853e+04 | 2.581659e+04 | NaN | KIVEACK |
| EEF2 | EF2\_HUMAN | Elongation factor 2 | 693 | 689 | EEF2 | EF2\_HUMAN | Elongation factor 2 | 5C(125.0477) | NaN | X | NaN | NaN | NaN | NaN | 8.529181e+04 | 8.529181e+04 | NaN | NaN | EGALCEENMR |
| BLMH | BLMH\_HUMAN | Bleomycin hydrolase | 326 | 318 | BLMH | BLMH\_HUMAN | Bleomycin hydrolase | 9C(125.0477) | NaN | X | NaN | NaN | NaN | NaN | 8.522766e+04 | 8.522766e+04 | NaN | NaN | DGEAVWFGCDVGK |
| RHOA | RHOA\_HUMAN | Transforming protein RhoA | 107, 107 | 105 | RHOA | RHOA\_HUMAN | Transforming protein RhoA | 3C(125.0477) 3C(15.9949) | NaN | X | X | NaN | NaN | NaN | 8.521384e+04 | 8.521384e+04 | NaN | NaN | HFCPNVPIILVGNK |
| GSS | GSHB\_HUMAN | Glutathione synthetase | 294 | 294 | GSS | GSHB\_HUMAN | Glutathione synthetase | 1C(125.0477) | NaN | X | NaN | NaN | NaN | NaN | 8.516685e+04 | 7.281802e+04 | 1.234883e+04 | NaN | CPDIATQLAGTK |
| ALB | ALBU\_HUMAN | Albumin | 289, 289 | 287 | ALB | ALBU\_HUMAN | Albumin | 3C(125.0477) 3C(57.0215) | X | X | NaN | NaN | NaN | NaN | 8.509208e+04 | NaN | NaN | 85092.0800 | YICENQDSISSK |
| EPB42 | EPB42\_HUMAN | Protein 4.2 | 428 | 428 | EPB42 | EPB42\_HUMAN | Protein 4.2 | 1C(125.0477) | NaN | X | NaN | NaN | NaN | NaN | 8.502907e+04 | 8.502907e+04 | NaN | NaN | CEDITQNYKYPEGSLQEKEVLER |
| UBR4 | UBR4\_HUMAN | E3 ubiquitin-protein ligase UBR4 | 4146 | 4143 | UBR4 | UBR4\_HUMAN | E3 ubiquitin-protein ligase UBR4 | 4C(125.0477) | NaN | X | NaN | NaN | NaN | NaN | 8.498438e+04 | 8.498438e+04 | NaN | NaN | QAACTIVEALATIPSR |
| DPYSL2 | DPYL2\_HUMAN | Dihydropyrimidinase-related protein 2 | 248 | 239 | DPYSL2 | DPYL2\_HUMAN | Dihydropyrimidinase-related protein 2 | 10C(125.0477) | NaN | X | NaN | NaN | NaN | NaN | 8.489012e+04 | 8.489012e+04 | NaN | NaN | AITIANQTNCPLYITK |
| SCRN1 | SCRN1\_HUMAN | Secernin-1 | 324 | 320 | SCRN1 | SCRN1\_HUMAN | Secernin-1 | 5C(125.0477) | NaN | X | NaN | NaN | NaN | NaN | 8.480162e+04 | 8.480162e+04 | NaN | NaN | TQSPCFGDDDPAKKEPR |
| CUL4A | CUL4A\_HUMAN | Cullin-4A | 94 | 81 | CUL4A | CUL4A\_HUMAN | Cullin-4A | 14C(125.0477) | NaN | X | NaN | NaN | NaN | NaN | 8.474812e+04 | 8.474812e+04 | NaN | NaN | YNLEELYQAVENLCSHK |
| RNPEP | AMPB\_HUMAN | Aminopeptidase B | 151 | 144 | RNPEP | AMPB\_HUMAN | Aminopeptidase B | 8C(125.0477) | NaN | X | NaN | NaN | NaN | NaN | 8.470994e+04 | 8.470994e+04 | NaN | NaN | VGEGPGVCWLAPEQTAGK |
| PYCR3 | P5CR3\_HUMAN | Pyrroline-5-carboxylate reductase 3 | 159 | 151 | PYCR3 | P5CR3\_HUMAN | Pyrroline-5-carboxylate reductase 3 | 9C(125.0477) | NaN | X | NaN | NaN | NaN | NaN | 8.446874e+04 | 8.446874e+04 | NaN | NaN | LLQHLLEACGR |
| CLNS1A | ICLN\_HUMAN | Methylosome subunit pICln | 73 | 69 | CLNS1A | ICLN\_HUMAN | Methylosome subunit pICln | 5C(125.0477) | NaN | X | NaN | NaN | NaN | NaN | 8.446042e+04 | 8.446042e+04 | NaN | NaN | DRSDCLGEHLYVMVNAK |
| CCT7 | TCPH\_HUMAN | T-complex protein 1 subunit eta | 511 | 500 | CCT7 | TCPH\_HUMAN | T-complex protein 1 subunit eta | 12C(125.0477) | NaN | X | NaN | NaN | NaN | NaN | 8.443105e+04 | 8.443105e+04 | NaN | NaN | INALTAASEAACLIVSVDETIK |
| MYH9 | MYH9\_HUMAN | Myosin-9 | 569 | 564 | MYH9 | MYH9\_HUMAN | Myosin-9 | 6C(125.0477) | NaN | X | NaN | NaN | NaN | NaN | 8.437024e+04 | 8.437024e+04 | NaN | NaN | DKADFCIIHYAGK |
| HUWE1 | HUWE1\_HUMAN | E3 ubiquitin-protein ligase HUWE1 | 689 | 684 | HUWE1 | HUWE1\_HUMAN | E3 ubiquitin-protein ligase HUWE1 | 6C(125.0477) | NaN | X | NaN | NaN | NaN | NaN | 8.417787e+04 | 8.417787e+04 | NaN | NaN | LLEEICNLGR |
| PSMD11 | PSD11\_HUMAN | 26S proteasome non-ATPase regulatory subunit 11 | 289 | 289 | PSMD11 | PSD11\_HUMAN | 26S proteasome non-ATPase regulatory subunit 11 | 1C(125.0477) | NaN | X | NaN | NaN | NaN | NaN | 8.411186e+04 | 8.411186e+04 | NaN | NaN | CVAQASK |
| CCT2 | TCPB\_HUMAN | T-complex protein 1 subunit beta | 371 | 359 | CCT2 | TCPB\_HUMAN | T-complex protein 1 subunit beta | 13C(125.0477) | NaN | X | NaN | NaN | NaN | NaN | 8.392595e+04 | 8.392595e+04 | NaN | NaN | LIHFSGVALGEACTIVLR |
| AP2B1 | AP2B1\_HUMAN | AP-2 complex subunit beta | 380 | 380 | AP2B1 | AP2B1\_HUMAN | AP-2 complex subunit beta | 1C(125.0477) | NaN | X | NaN | NaN | NaN | NaN | 8.387771e+04 | 8.387771e+04 | NaN | NaN | CAIKVEQSAER |
| PSMA6 | PSA6\_HUMAN | Proteasome subunit alpha type-6 | 47 | 44 | PSMA6 | PSA6\_HUMAN | Proteasome subunit alpha type-6 | 4C(125.0477) | NaN | X | NaN | NaN | NaN | NaN | 8.364955e+04 | 8.364955e+04 | NaN | NaN | GKDCAVIVTQK |
| ADD2 | ADDB\_HUMAN | Beta-adducin | 135 | 135 | ADD2 | ADDB\_HUMAN | Beta-adducin | 1C(125.0477) | NaN | X | NaN | NaN | NaN | NaN | 8.357497e+04 | 4.656208e+04 | 3.701289e+04 | NaN | CKISSVYR |
| ALDH1A1 | AL1A1\_HUMAN | Aldehyde dehydrogenase 1A1 | 50 | 38 | ALDH1A1 | AL1A1\_HUMAN | Aldehyde dehydrogenase 1A1 | 13C(125.0477) | NaN | X | NaN | NaN | NaN | NaN | 8.339674e+04 | 8.339674e+04 | NaN | NaN | FPVFNPATEEELCQVEEGDKEDVDKAVK |
| CLTC | CLH1\_HUMAN | Clathrin heavy chain 1 | 1205 | 1183 | CLTC | CLH1\_HUMAN | Clathrin heavy chain 1 | 23C(125.0477) | NaN | X | NaN | NaN | NaN | NaN | 8.339406e+04 | 8.339406e+04 | NaN | NaN | LAELEEFINGPNNAHIQQVGDRCYDEK |
| AKR1A1 | AK1A1\_HUMAN | Aldo-keto reductase family 1 member A1 | 46 | 43 | AKR1A1 | AK1A1\_HUMAN | Aldo-keto reductase family 1 member A1 | 4C(125.0477) | NaN | X | NaN | NaN | NaN | NaN | 8.324221e+04 | 8.324221e+04 | NaN | NaN | HIDCAAIYGNEPEIGEALKEDVGPGK |
| DSTN | DEST\_HUMAN | Destrin | 23 | 23 | DSTN | DEST\_HUMAN | Destrin | 1C(125.0477) | NaN | X | NaN | NaN | NaN | NaN | 8.322639e+04 | 8.322639e+04 | NaN | NaN | CSTPEEIKK |
| AGO2 | AGO2\_HUMAN | Protein argonaute-2 | 455 | 450 | AGO2 | AGO2\_HUMAN | Protein argonaute-2 | 6C(125.0477) | NaN | X | NaN | NaN | NaN | NaN | 8.320678e+04 | 8.320678e+04 | NaN | NaN | VWAIACFAPQR |
| AP2A1 | AP2A1\_HUMAN | AP-2 complex subunit alpha-1 | 941 | 936 | AP2A1 | AP2A1\_HUMAN | AP-2 complex subunit alpha-1 | 6C(125.0477) | NaN | X | NaN | NaN | NaN | NaN | 8.305910e+04 | 8.305910e+04 | NaN | NaN | ALQVGCLLR |
| ACO1 | ACOHC\_HUMAN | Cytoplasmic aconitate hydratase | 50 | 49 | ACO1 | ACOHC\_HUMAN | Cytoplasmic aconitate hydratase | 2C(125.0477) | NaN | X | NaN | NaN | NaN | NaN | 8.303345e+04 | 8.303345e+04 | NaN | NaN | NCDEFLVK |
| C3 | CO3\_HUMAN | Complement C3 | 1158, 1158 | 1156 | C3 | CO3\_HUMAN | Complement C3 | 3C(125.0477) 3C(57.0215) | X | X | NaN | NaN | NaN | NaN | 8.293496e+04 | NaN | 8.293496e+04 | NaN | DICEEQVNSLPGSITK |
| TMEM30A | CC50A\_HUMAN | Cell cycle control protein 50A | 17 | 9 | TMEM30A | CC50A\_HUMAN | Cell cycle control protein 50A | 9C(125.0477) | NaN | X | NaN | NaN | NaN | NaN | 8.284062e+04 | NaN | 8.284062e+04 | NaN | DEVDGGPPCAPGGTAK |
| ECPAS | ECM29\_HUMAN | Proteasome adapter and scaffold protein ECM29 | 1183 | 1180 | ECPAS | ECM29\_HUMAN | Proteasome adapter and scaffold protein ECM29 | 4C(125.0477) | NaN | X | NaN | NaN | NaN | NaN | 8.277705e+04 | 8.277705e+04 | NaN | NaN | ESSCLALNDLLR |
| EML2 | EMAL2\_HUMAN | Echinoderm microtubule-associated protein-like 2 | 395 | 390 | EML2 | EMAL2\_HUMAN | Echinoderm microtubule-associated protein-like 2 | 6C(125.0477) | NaN | X | NaN | NaN | NaN | NaN | 8.270962e+04 | 2.241474e+04 | 6.029488e+04 | NaN | AQFVTCGQDK |
| NUP153 | NU153\_HUMAN | Nuclear pore complex protein Nup153 | 585 | 556 | NUP153 | NU153\_HUMAN | Nuclear pore complex protein Nup153 | 30C(31.9988) | NaN | NaN | NaN | X | NaN | NaN | 8.269033e+04 | 8.269033e+04 | NaN | NaN | TAELSGSSSTLEPIISSSAHHVTTVNSTNCKK |
| IPO11 | IPO11\_HUMAN | Importin-11 | 621 | 621 | IPO11 | IPO11\_HUMAN | Importin-11 | 1C(125.0477) | NaN | X | NaN | NaN | NaN | NaN | 8.263970e+04 | 8.263970e+04 | NaN | NaN | CAILTTLIHLVQGLGADSK |
| SCYL1 | SCYL1\_HUMAN | N-terminal kinase-like protein | 210 | 208 | SCYL1 | SCYL1\_HUMAN | N-terminal kinase-like protein | 3C(125.0477) | NaN | X | NaN | NaN | NaN | NaN | 8.250962e+04 | 8.250962e+04 | NaN | NaN | LGCLIWEVFNGPLPR |
| L3HYPDH | T3HPD\_HUMAN | Trans-3-hydroxy-L-proline dehydratase | 39 | 34 | L3HYPDH | T3HPD\_HUMAN | Trans-3-hydroxy-L-proline dehydratase | 6C(125.0477) | NaN | X | NaN | NaN | NaN | NaN | 8.226963e+04 | 8.226963e+04 | NaN | NaN | IVLAGCPEVSGPTLLAK |
| DNAJA4 | DNJA4\_HUMAN | DnaJ homolog subfamily A member 4 | 303 | 303 | DNAJA4 | DNJA4\_HUMAN | DnaJ homolog subfamily A member 4 | 1C(125.0477) | NaN | X | NaN | NaN | NaN | NaN | 8.212751e+04 | 8.212751e+04 | NaN | NaN | CVRDEGMPIYK |
| SFN | 1433S\_HUMAN | 14-3-3 protein sigma | 38 | 33 | SFN | 1433S\_HUMAN | 14-3-3 protein sigma | 6C(125.0477) | NaN | X | NaN | NaN | NaN | NaN | 8.203492e+04 | NaN | 8.203492e+04 | NaN | GEELSCEER |
| FASN | FAS\_HUMAN | Fatty acid synthase | 1471 | 1471 | FASN | FAS\_HUMAN | Fatty acid synthase | 1C(125.0477) | NaN | X | NaN | NaN | NaN | NaN | 8.184753e+04 | 8.184753e+04 | NaN | NaN | CVLLSNLSSTSHVPEVDPGSAELQK |
| RNH1 | RINI\_HUMAN | Ribonuclease inhibitor | 266 | 260 | RNH1 | RINI\_HUMAN | Ribonuclease inhibitor | 7C(125.0477) | NaN | X | NaN | NaN | NaN | NaN | 8.177478e+04 | 8.177478e+04 | NaN | NaN | TLWIWECGITAK |
| PFAS | PUR4\_HUMAN | Phosphoribosylformylglycinamidine synthase | 606 | 598 | PFAS | PUR4\_HUMAN | Phosphoribosylformylglycinamidine synthase | 9C(125.0477) | NaN | X | NaN | NaN | NaN | NaN | 8.177444e+04 | 8.177444e+04 | NaN | NaN | IVLVDDRECPVRR |
| EIF2S3 | IF2G\_HUMAN | Eukaryotic translation initiation factor 2 subunit 3 | 236 | 230 | EIF2S3 | IF2G\_HUMAN | Eukaryotic translation initiation factor 2 subunit 3 | 7C(125.0477) | NaN | X | NaN | NaN | NaN | NaN | 8.168772e+04 | 8.168772e+04 | NaN | NaN | YNIEVVCEYIVK |
| PHAF1 | PHAF1\_HUMAN | Phagosome assembly factor 1 | 222 | 217 | PHAF1 | PHAF1\_HUMAN | Phagosome assembly factor 1 | 6C(125.0477) | NaN | X | NaN | NaN | NaN | NaN | 8.167378e+04 | 8.167378e+04 | NaN | NaN | LLAAGCGPGLLADAK |
| HUWE1 | HUWE1\_HUMAN | E3 ubiquitin-protein ligase HUWE1 | 1879 | 1873 | HUWE1 | HUWE1\_HUMAN | E3 ubiquitin-protein ligase HUWE1 | 7C(125.0477) | NaN | X | NaN | NaN | NaN | NaN | 8.149138e+04 | 8.149138e+04 | NaN | NaN | VLGPAACR |
| HNRNPA3 | ROA3\_HUMAN | Heterogeneous nuclear ribonucleoprotein A3 | 196 | 188 | HNRNPA3 | ROA3\_HUMAN | Heterogeneous nuclear ribonucleoprotein A3 | 9C(125.0477) | NaN | X | NaN | NaN | NaN | NaN | 8.137624e+04 | 8.137624e+04 | NaN | NaN | YHTINGHNCEVK |
| DNAJC17 | DJC17\_HUMAN | DnaJ homolog subfamily C member 17 | 38 | 35 | DNAJC17 | DJC17\_HUMAN | DnaJ homolog subfamily C member 17 | 4C(125.0477) | NaN | X | NaN | NaN | NaN | NaN | 8.136464e+04 | 8.136464e+04 | NaN | NaN | ALSCHPDKNPDNPR |
| GSTM4 | GSTM4\_HUMAN | Glutathione S-transferase Mu 4 | 87 | 83 | GSTM4 | GSTM4\_HUMAN | Glutathione S-transferase Mu 4 | 5C(125.0477) | NaN | X | NaN | NaN | NaN | NaN | 8.135028e+04 | 8.135028e+04 | NaN | NaN | KHNLCGETEEEK |
| EIF2S3 | IF2G\_HUMAN | Eukaryotic translation initiation factor 2 subunit 3 | 311 | 309 | EIF2S3 | IF2G\_HUMAN | Eukaryotic translation initiation factor 2 subunit 3 | 3C(125.0477) | NaN | X | NaN | NaN | NaN | NaN | 8.126338e+04 | 8.126338e+04 | NaN | NaN | LMCKPIFSK |
| RNH1 | RINI\_HUMAN | Ribonuclease inhibitor | 216, 216, 209 | 206 | RNH1 | RINI\_HUMAN | Ribonuclease inhibitor | 11C(125.0477) 11C(31.9988) 4C(31.9988) | NaN | X | NaN | X | NaN | NaN | 8.114408e+04 | 8.114408e+04 | NaN | NaN | LESCGVTSDNCR |
| CNP | CN37\_HUMAN | 2',3'-cyclic-nucleotide 3'-phosphodiesterase | 111 | 102 | CNP | CN37\_HUMAN | 2',3'-cyclic-nucleotide 3'-phosphodiesterase | 10C(125.0477) | NaN | X | NaN | NaN | NaN | NaN | 8.092915e+04 | 8.092915e+04 | NaN | NaN | RLDEDLAAYCR |
| GCLC | GSH1\_HUMAN | Glutamate--cysteine ligase catalytic subunit | 295 | 288 | GCLC | GSH1\_HUMAN | Glutamate--cysteine ligase catalytic subunit | 8C(125.0477) | NaN | X | NaN | NaN | NaN | NaN | 8.092555e+04 | NaN | 8.092555e+04 | NaN | GYVSDIDCR |
| TRAF2 | TRAF2\_HUMAN | TNF receptor-associated factor 2 | 124 | 120 | TRAF2 | TRAF2\_HUMAN | TNF receptor-associated factor 2 | 5C(125.0477) | NaN | X | NaN | NaN | NaN | NaN | 8.090670e+04 | 8.090670e+04 | NaN | NaN | EYESCHEGR |
| CCT8 | TCPQ\_HUMAN | T-complex protein 1 subunit theta | 187, 187 | 182 | CCT8 | TCPQ\_HUMAN | T-complex protein 1 subunit theta | 6C(125.0477) 6C(305.3076) | NaN | X | NaN | NaN | NaN | X | 8.086584e+04 | 8.086584e+04 | NaN | NaN | LIAQACVSIFPDSGHFNVDNIR |
| AGO2 | AGO2\_HUMAN | Protein argonaute-2 | 272, 272 | 267 | AGO2 | AGO2\_HUMAN | Protein argonaute-2 | 6C(125.0477) 6C(15.9949) | NaN | X | X | NaN | NaN | NaN | 8.078727e+04 | 8.078727e+04 | NaN | NaN | VEITHCGQMK |
| IPO13 | IPO13\_HUMAN | Importin-13 | 848 | 848 | IPO13 | IPO13\_HUMAN | Importin-13 | 1C(125.0477) | NaN | X | NaN | NaN | NaN | NaN | 8.078545e+04 | 8.078545e+04 | NaN | NaN | CGEVESVGK |
| AVEN | AVEN\_HUMAN | Cell death regulator Aven | 251 | 246 | AVEN | AVEN\_HUMAN | Cell death regulator Aven | 6C(125.0477) | NaN | X | NaN | NaN | NaN | NaN | 8.055635e+04 | 8.055635e+04 | NaN | NaN | SVAAGCPVLLGK |
| CIAPIN1 | CPIN1\_HUMAN | Anamorsin | 116 | 115 | CIAPIN1 | CPIN1\_HUMAN | Anamorsin | 2C(125.0477) | NaN | X | NaN | NaN | NaN | NaN | 8.052777e+04 | 8.052777e+04 | NaN | NaN | LCSALTLSGLVEVK |
| G6PD | G6PD\_HUMAN | Glucose-6-phosphate 1-dehydrogenase | 13 | 10 | G6PD | G6PD\_HUMAN | Glucose-6-phosphate 1-dehydrogenase | 4C(125.0477) | NaN | X | NaN | NaN | NaN | NaN | 8.052241e+04 | NaN | 8.052241e+04 | NaN | TQVCGILR |
| SND1 | SND1\_HUMAN | Staphylococcal nuclease domain-containing protein 1 | 152 | 149 | SND1 | SND1\_HUMAN | Staphylococcal nuclease domain-containing protein 1 | 4C(125.0477) | NaN | X | NaN | NaN | NaN | NaN | 8.043502e+04 | 5.788598e+04 | 2.254904e+04 | NaN | LSECEEQAK |
| ABCE1 | ABCE1\_HUMAN | ATP-binding cassette sub-family E member 1 | 65 | 64 | ABCE1 | ABCE1\_HUMAN | ATP-binding cassette sub-family E member 1 | 2C(125.0477) | NaN | X | NaN | NaN | NaN | NaN | 8.030917e+04 | 8.030917e+04 | NaN | NaN | KCPFGALSIVNLPSNLEK |
| FASN | FAS\_HUMAN | Fatty acid synthase | 1759 | 1759 | FASN | FAS\_HUMAN | Fatty acid synthase | 1C(125.0477) | NaN | X | NaN | NaN | NaN | NaN | 8.020754e+04 | 8.020754e+04 | NaN | NaN | CLATHGR |
| LDHB | LDHB\_HUMAN | L-lactate dehydrogenase B chain | 164 | 157 | LDHB | LDHB\_HUMAN | L-lactate dehydrogenase B chain | 8C(125.0477) | NaN | X | NaN | NaN | NaN | NaN | 8.011083e+04 | 8.011083e+04 | NaN | NaN | HRVIGSGCNLDSAR |
| PGD | 6PGD\_HUMAN | 6-phosphogluconate dehydrogenase, decarboxylating | 402 | 397 | PGD | 6PGD\_HUMAN | 6-phosphogluconate dehydrogenase, decarboxylating | 6C(125.0477) | NaN | X | NaN | NaN | NaN | NaN | 8.008061e+04 | NaN | 8.008061e+04 | NaN | SAVENCQDSWR |
| PA2G4 | PA2G4\_HUMAN | Proliferation-associated protein 2G4 | 149 | 145 | PA2G4 | PA2G4\_HUMAN | Proliferation-associated protein 2G4 | 5C(125.0477) | NaN | X | NaN | NaN | NaN | NaN | 7.992480e+04 | 7.992480e+04 | NaN | NaN | AAHLCAEAALR |
| UBR4 | UBR4\_HUMAN | E3 ubiquitin-protein ligase UBR4 | 934 | 932 | UBR4 | UBR4\_HUMAN | E3 ubiquitin-protein ligase UBR4 | 3C(125.0477) | NaN | X | NaN | NaN | NaN | NaN | 7.990271e+04 | 7.990271e+04 | NaN | NaN | FYCVLSPEASEDDLNR |
| PRMT5 | ANM5\_HUMAN | Protein arginine N-methyltransferase 5 | 518 | 506 | PRMT5 | ANM5\_HUMAN | Protein arginine N-methyltransferase 5 | 13C(125.0477) | NaN | X | NaN | NaN | NaN | NaN | 7.966095e+04 | 7.966095e+04 | NaN | NaN | LHNFHQLSAPQPCFTFSHPNR |
| ABRACL | ABRAL\_HUMAN | Costars family protein ABRACL | 39 | 36 | ABRACL | ABRAL\_HUMAN | Costars family protein ABRACL | 4C(125.0477) | NaN | X | NaN | NaN | NaN | NaN | 7.947584e+04 | 7.947584e+04 | NaN | NaN | DDKCANLFEALVGTLK |
| ERI3 | ERI3\_HUMAN | ERI1 exoribonuclease 3 | 258 | 252 | ERI3 | ERI3\_HUMAN | ERI1 exoribonuclease 3 | 7C(125.0477) | NaN | X | NaN | NaN | NaN | NaN | 7.943912e+04 | 7.943912e+04 | NaN | NaN | VMLPGQCQYLGLPVADYFK |
| GSN | GELS\_HUMAN | Gelsolin | 331 | 328 | GSN | GELS\_HUMAN | Gelsolin | 4C(125.0477) | NaN | X | NaN | NaN | NaN | NaN | 7.938552e+04 | 7.005450e+04 | 9.331019e+03 | NaN | SEDCFILDHGK |
| CTPS2 | PYRG2\_HUMAN | CTP synthase 2 | 176 | 173 | CTPS2 | PYRG2\_HUMAN | CTP synthase 2 | 4C(125.0477) | NaN | X | NaN | NaN | NaN | NaN | 7.923238e+04 | 7.923238e+04 | NaN | NaN | ENFCNIHVSLVPQLSATGEQK |
| EML2 | EMAL2\_HUMAN | Echinoderm microtubule-associated protein-like 2 | 301 | 285 | EML2 | EMAL2\_HUMAN | Echinoderm microtubule-associated protein-like 2 | 17C(125.0477) | NaN | X | NaN | NaN | NaN | NaN | 7.914830e+04 | 7.914830e+04 | NaN | NaN | ITQAVLGAHDGGVFGLCALR |
| PRKAR2A | KAP2\_HUMAN | cAMP-dependent protein kinase type II-alpha regulatory subunit | 374 | 370 | PRKAR2A | KAP2\_HUMAN | cAMP-dependent protein kinase type II-alpha regulatory subunit | 5C(125.0477) | NaN | X | NaN | NaN | NaN | NaN | 7.914801e+04 | 7.914801e+04 | NaN | NaN | LLGPCMDIMK |
| ERBIN | ERBIN\_HUMAN | Erbin | 464 | 458 | ERBIN | ERBIN\_HUMAN | Erbin | 7C(125.0477) | NaN | X | NaN | NaN | NaN | NaN | 7.909992e+04 | 7.909992e+04 | NaN | NaN | AQVAFECDEDKDER |
| PSMD13 | PSD13\_HUMAN | 26S proteasome non-ATPase regulatory subunit 13 | 114 | 106 | PSMD13 | PSD13\_HUMAN | 26S proteasome non-ATPase regulatory subunit 13 | 9C(125.0477) | NaN | X | NaN | NaN | NaN | NaN | 7.907587e+04 | NaN | 7.907587e+04 | NaN | SSDEAVILCK |
| PREP | PPCE\_HUMAN | Prolyl endopeptidase | 343 | 328 | PREP | PPCE\_HUMAN | Prolyl endopeptidase | 16C(125.0477) | NaN | X | NaN | NaN | NaN | NaN | 7.899869e+04 | 7.899869e+04 | NaN | NaN | VLVPEHEKDVLEWIACVR |
| DHX9 | DHX9\_HUMAN | ATP-dependent RNA helicase A | 578 | 567 | DHX9 | DHX9\_HUMAN | ATP-dependent RNA helicase A | 12C(125.0477) | NaN | X | NaN | NaN | NaN | NaN | 7.897094e+04 | 7.897094e+04 | NaN | NaN | TYPVQEYFLEDCIQMTHFVPPPKDK |
| AHSA1 | AHSA1\_HUMAN | Activator of 90 kDa heat shock protein ATPase homolog 1 | 56 | 56 | AHSA1 | AHSA1\_HUMAN | Activator of 90 kDa heat shock protein ATPase homolog 1 | 1C(125.0477) | NaN | X | NaN | NaN | NaN | NaN | 7.884689e+04 | 7.884689e+04 | NaN | NaN | CEVTEVSK |
| ARHGAP23 | RHG23\_HUMAN | Rho GTPase-activating protein 23 | 910 | 907 | ARHGAP23 | RHG23\_HUMAN | Rho GTPase-activating protein 23 | 4C(125.0477) | NaN | X | NaN | NaN | NaN | NaN | 7.874183e+04 | 7.101007e+04 | 7.731763e+03 | NaN | LEECQPATENQR |
| GART | PUR2\_HUMAN | Trifunctional purine biosynthetic protein adenosine-3 | 134 | 126 | GART | PUR2\_HUMAN | Trifunctional purine biosynthetic protein adenosine-3 | 9C(125.0477) | NaN | X | NaN | NaN | NaN | NaN | 7.872370e+04 | 7.872370e+04 | NaN | NaN | AFTKPEEACSFILSADFPALVVK |
| PRKCSH | GLU2B\_HUMAN | Glucosidase 2 subunit beta | 471 | 465 | PRKCSH | GLU2B\_HUMAN | Glucosidase 2 subunit beta | 7C(125.0477) | NaN | X | NaN | NaN | NaN | NaN | 7.870201e+04 | NaN | 7.870201e+04 | NaN | YEQGTGCWQGPNR |
| NUP43 | NUP43\_HUMAN | Nucleoporin Nup43 | 222 | 218 | NUP43 | NUP43\_HUMAN | Nucleoporin Nup43 | 5C(125.0477) | NaN | X | NaN | NaN | NaN | NaN | 7.867328e+04 | 7.867328e+04 | NaN | NaN | VPLHCVDR |
| FLOT1 | FLOT1\_HUMAN | Flotillin-1 | 85 | 79 | FLOT1 | FLOT1\_HUMAN | Flotillin-1 | 7C(125.0477) | NaN | X | NaN | NaN | NaN | NaN | 7.840231e+04 | 7.840231e+04 | NaN | NaN | EMLAAACQMFLGK |
| MIB1 | MIB1\_HUMAN | E3 ubiquitin-protein ligase MIB1 | 554 | 548 | MIB1 | MIB1\_HUMAN | E3 ubiquitin-protein ligase MIB1 | 7C(125.0477) | NaN | X | NaN | NaN | NaN | NaN | 7.833839e+04 | 7.833839e+04 | NaN | NaN | TLLDFGCHPSLQDSEGDTPLHDAISK |
| ACLY | ACLY\_HUMAN | ATP-citrate synthase | 1040 | 1039 | ACLY | ACLY\_HUMAN | ATP-citrate synthase | 2C(125.0477) | NaN | X | NaN | NaN | NaN | NaN | 7.819120e+04 | 7.819120e+04 | NaN | NaN | NCGSFTR |
| AMPD2 | AMPD2\_HUMAN | AMP deaminase 2 | 123 | 123 | AMPD2 | AMPD2\_HUMAN | AMP deaminase 2 | 1C(125.0477) | NaN | X | NaN | NaN | NaN | NaN | 7.804602e+04 | 7.804602e+04 | NaN | NaN | CKEIAEELFTR |
| CAPNS1 | CPNS1\_HUMAN | Calpain small subunit 1 | 144 | 127 | CAPNS1 | CPNS1\_HUMAN | Calpain small subunit 1 | 18C(125.0477) | NaN | X | NaN | NaN | NaN | NaN | 7.790302e+04 | 7.790302e+04 | NaN | NaN | VVTRHPDLKTDGFGIDTCR |
| VPS13C | VP13C\_HUMAN | Vacuolar protein sorting-associated protein 13C | 2395 | 2394 | VPS13C | VP13C\_HUMAN | Vacuolar protein sorting-associated protein 13C | 2C(125.0477) | NaN | X | NaN | NaN | NaN | NaN | 7.784141e+04 | 7.784141e+04 | NaN | NaN | SCLNVFNNLAK |
| C2CD5 | C2CD5\_HUMAN | C2 domain-containing protein 5 | 449 | 440 | C2CD5 | C2CD5\_HUMAN | C2 domain-containing protein 5 | 10C(125.0477) | NaN | X | NaN | NaN | NaN | NaN | 7.776945e+04 | 7.776945e+04 | NaN | NaN | FLQDGTVEGCLEQR |
| HSPBP1 | HPBP1\_HUMAN | Hsp70-binding protein 1 | 201 | 195 | HSPBP1 | HPBP1\_HUMAN | Hsp70-binding protein 1 | 7C(125.0477) | NaN | X | NaN | NaN | NaN | NaN | 7.775330e+04 | 7.775330e+04 | NaN | NaN | LLDRDACDTVR |
| HUWE1 | HUWE1\_HUMAN | E3 ubiquitin-protein ligase HUWE1 | 2721 | 2716 | HUWE1 | HUWE1\_HUMAN | E3 ubiquitin-protein ligase HUWE1 | 6C(125.0477) | NaN | X | NaN | NaN | NaN | NaN | 7.772320e+04 | 7.772320e+04 | NaN | NaN | DQSAQCTASK |
| GAPVD1 | GAPD1\_HUMAN | GTPase-activating protein and VPS9 domain-containing protein 1 | 789 | 787 | GAPVD1 | GAPD1\_HUMAN | GTPase-activating protein and VPS9 domain-containing protein 1 | 3C(125.0477) | NaN | X | NaN | NaN | NaN | NaN | 7.767393e+04 | 6.700098e+04 | 1.067294e+04 | NaN | SECSSDFGGK |
| STK24 | STK24\_HUMAN | Serine/threonine-protein kinase 24 | 394 | 391 | STK24 | STK24\_HUMAN | Serine/threonine-protein kinase 24 | 4C(125.0477) | NaN | X | NaN | NaN | NaN | NaN | 7.757355e+04 | 7.757355e+04 | NaN | NaN | SQACGGNLGSIEELR |
| CKAP5 | CKAP5\_HUMAN | Cytoskeleton-associated protein 5 | 144 | 140 | CKAP5 | CKAP5\_HUMAN | Cytoskeleton-associated protein 5 | 5C(125.0477) | NaN | X | NaN | NaN | NaN | NaN | 7.756461e+04 | 7.756461e+04 | NaN | NaN | IIVACIETLR |
| PFKM | PFKAM\_HUMAN | ATP-dependent 6-phosphofructokinase, muscle type | 709 | 700 | PFKM | PFKAM\_HUMAN | ATP-dependent 6-phosphofructokinase, muscle type | 10C(125.0477) | NaN | X | NaN | NaN | NaN | NaN | 7.746958e+04 | 7.746958e+04 | NaN | NaN | IFANTPDSGCVLGMR |
| PGD | 6PGD\_HUMAN | 6-phosphogluconate dehydrogenase, decarboxylating | 402, 402 | 397 | PGD | 6PGD\_HUMAN | 6-phosphogluconate dehydrogenase, decarboxylating | 6C(125.0477) 6C(57.0215) | X | X | NaN | NaN | NaN | NaN | 7.745078e+04 | 7.745078e+04 | NaN | NaN | SAVENCQDSWR |
| PCBD2 | PHS2\_HUMAN | Pterin-4-alpha-carbinolamine dehydratase 2 | 109 | 100 | PCBD2 | PHS2\_HUMAN | Pterin-4-alpha-carbinolamine dehydratase 2 | 10C(125.0477) | NaN | X | NaN | NaN | NaN | NaN | 7.719603e+04 | 7.719603e+04 | NaN | NaN | VQITLTSHDCGELTKK |
| CAPZB | CAPZB\_HUMAN | F-actin-capping protein subunit beta | 147, 147 | 146 | CAPZB | CAPZB\_HUMAN | F-actin-capping protein subunit beta | 2C(125.0477) 2C(31.9988) | NaN | X | NaN | X | NaN | NaN | 7.717071e+04 | 7.717071e+04 | NaN | NaN | GCWDSIHVVEVQEK |
| USP9X | USP9X\_HUMAN | Probable ubiquitin carboxyl-terminal hydrolase FAF-X | 540 | 535 | USP9X | USP9X\_HUMAN | Probable ubiquitin carboxyl-terminal hydrolase FAF-X | 6C(125.0477) | NaN | X | NaN | NaN | NaN | NaN | 7.716548e+04 | 7.716548e+04 | NaN | NaN | ILDYSCSQDRDTQK |
| S100A8 | S10A8\_HUMAN | Protein S100-A8 | 42 | 37 | S100A8 | S10A8\_HUMAN | Protein S100-A8 | 6C(125.0477) | NaN | X | NaN | NaN | NaN | NaN | 7.715886e+04 | NaN | 7.715886e+04 | NaN | LLETECPQYIR |
| USP24 | UBP24\_HUMAN | Ubiquitin carboxyl-terminal hydrolase 24 | 873 | 868 | USP24 | UBP24\_HUMAN | Ubiquitin carboxyl-terminal hydrolase 24 | 6C(125.0477) | NaN | X | NaN | NaN | NaN | NaN | 7.715704e+04 | 7.715704e+04 | NaN | NaN | KFIADCYTR |
| DUSP23 | DUS23\_HUMAN | Dual specificity protein phosphatase 23 | 95 | 88 | DUSP23 | DUS23\_HUMAN | Dual specificity protein phosphatase 23 | 8C(125.0477) | NaN | X | NaN | NaN | NaN | NaN | 7.712147e+04 | 7.712147e+04 | NaN | NaN | GEAVGVHCALGFGR |
| HUWE1 | HUWE1\_HUMAN | E3 ubiquitin-protein ligase HUWE1 | 4211 | 4205 | HUWE1 | HUWE1\_HUMAN | E3 ubiquitin-protein ligase HUWE1 | 7C(125.0477) | NaN | X | NaN | NaN | NaN | NaN | 7.692212e+04 | 7.692212e+04 | NaN | NaN | EYVHLVCQMR |
| RABL6 | RABL6\_HUMAN | Rab-like protein 6 | 491 | 483 | RABL6 | RABL6\_HUMAN | Rab-like protein 6 | 9C(125.0477) | NaN | X | NaN | NaN | NaN | NaN | 7.688944e+04 | 6.039471e+04 | 1.649473e+04 | NaN | GPAPAPQQCSEPETK |
| THOP1 | THOP1\_HUMAN | Thimet oligopeptidase | 231 | 231 | THOP1 | THOP1\_HUMAN | Thimet oligopeptidase | 1C(125.0477) | NaN | X | NaN | NaN | NaN | NaN | 7.675846e+04 | 4.522342e+04 | 3.153504e+04 | NaN | CHVPETR |
| PFAS | PUR4\_HUMAN | Phosphoribosylformylglycinamidine synthase | 606 | 598 | PFAS | PUR4\_HUMAN | Phosphoribosylformylglycinamidine synthase | 9C(125.0477) | NaN | X | NaN | NaN | NaN | NaN | 7.665372e+04 | 7.665372e+04 | NaN | NaN | IVLVDDRECPVR |
| RPN1 | RPN1\_HUMAN | Dolichyl-diphosphooligosaccharide--protein glycosyltransferase subunit 1 | 545 | 539 | RPN1 | RPN1\_HUMAN | Dolichyl-diphosphooligosaccharide--protein glycosyltransferase subunit 1 | 7C(125.0477) | NaN | X | NaN | NaN | NaN | NaN | 7.662477e+04 | NaN | 7.662477e+04 | NaN | TEGSDLCDR |
| EIF2S2 | IF2B\_HUMAN | Eukaryotic translation initiation factor 2 subunit 2 | 226 | 216 | EIF2S2 | IF2B\_HUMAN | Eukaryotic translation initiation factor 2 subunit 2 | 11C(125.0477) | NaN | X | NaN | NaN | NaN | NaN | 7.637430e+04 | 7.637430e+04 | NaN | NaN | KTSFVNFTDICK |
| STRADB | STRAB\_HUMAN | STE20-related kinase adapter protein beta | 96 | 90 | STRADB | STRAB\_HUMAN | STE20-related kinase adapter protein beta | 7C(125.0477) | NaN | X | NaN | NaN | NaN | NaN | 7.636914e+04 | 3.530289e+04 | 4.106625e+04 | NaN | ITNLENCNEER |
| PGM2L1 | PGM2L\_HUMAN | Glucose 1,6-bisphosphate synthase | 348 | 327 | PGM2L1 | PGM2L\_HUMAN | Glucose 1,6-bisphosphate synthase | 22C(125.0477) | NaN | X | NaN | NaN | NaN | NaN | 7.634567e+04 | 7.634567e+04 | NaN | NaN | VVLATDPDADRLAAAELQENGCWK |
| HSPBP1 | HPBP1\_HUMAN | Hsp70-binding protein 1 | 269 | 266 | HSPBP1 | HPBP1\_HUMAN | Hsp70-binding protein 1 | 4C(125.0477) | NaN | X | NaN | NaN | NaN | NaN | 7.625197e+04 | 7.625197e+04 | NaN | NaN | GTLCSMGMVQQLVALVR |
| GLRX3 | GLRX3\_HUMAN | Glutaredoxin-3 | 229 | 218 | GLRX3 | GLRX3\_HUMAN | Glutaredoxin-3 | 12C(125.0477) | NaN | X | NaN | NaN | NaN | NaN | 7.622240e+04 | 7.622240e+04 | NaN | NaN | ELEASEELDTICPK |
| EFNB1 | EFNB1\_HUMAN | Ephrin-B1 | 64 | 60 | EFNB1 | EFNB1\_HUMAN | Ephrin-B1 | 5C(125.0477) | NaN | X | NaN | NaN | NaN | NaN | 7.611910e+04 | NaN | 7.611910e+04 | NaN | LDIICPR |
| IPO9 | IPO9\_HUMAN | Importin-9 | 639 | 632 | IPO9 | IPO9\_HUMAN | Importin-9 | 8C(125.0477) | NaN | X | NaN | NaN | NaN | NaN | 7.611126e+04 | 7.611126e+04 | NaN | NaN | ELSQIEACQGPMQMR |
| SDF2L1 | SDF2L\_HUMAN | Stromal cell-derived factor 2-like protein 1 | 38 | 31 | SDF2L1 | SDF2L\_HUMAN | Stromal cell-derived factor 2-like protein 1 | 8C(125.0477) | NaN | X | NaN | NaN | NaN | NaN | 7.592531e+04 | NaN | 7.592531e+04 | NaN | TGAELVTCGSVLK |
| CFB | CFAB\_HUMAN | Complement factor B | 725 | 711 | CFB | CFAB\_HUMAN | Complement factor B | 15C(57.0215) | X | NaN | NaN | NaN | NaN | NaN | 7.588877e+04 | 7.588877e+04 | NaN | NaN | FIQVGVISWGVVDVCK |
| OTUB1 | OTUB1\_HUMAN | Ubiquitin thioesterase OTUB1 | 91 | 84 | OTUB1 | OTUB1\_HUMAN | Ubiquitin thioesterase OTUB1 | 8C(125.0477) | NaN | X | NaN | NaN | NaN | NaN | 7.588171e+04 | 7.588171e+04 | NaN | NaN | KTRPDGNCFYR |
| CAPN1 | CAN1\_HUMAN | Calpain-1 catalytic subunit | 49 | 49 | CAPN1 | CAN1\_HUMAN | Calpain-1 catalytic subunit | 1C(125.0477) | NaN | X | NaN | NaN | NaN | NaN | 7.571630e+04 | 7.571630e+04 | NaN | NaN | CLQSGTLFRDEAFPPVPQSLGYK |
| PCBD1 | PHS\_HUMAN | Pterin-4-alpha-carbinolamine dehydratase | 82 | 73 | PCBD1 | PHS\_HUMAN | Pterin-4-alpha-carbinolamine dehydratase | 10C(125.0477) | NaN | X | NaN | NaN | NaN | NaN | 7.549004e+04 | 7.549004e+04 | NaN | NaN | VHITLSTHECAGLSER |
| PSMD4 | PSMD4\_HUMAN | 26S proteasome non-ATPase regulatory subunit 4 | 58 | 43 | PSMD4 | PSMD4\_HUMAN | 26S proteasome non-ATPase regulatory subunit 4 | 16C(125.0477) | NaN | X | NaN | NaN | NaN | NaN | 7.547139e+04 | 7.547139e+04 | NaN | NaN | SNPENNVGLITLANDCEVLTTLTPDTGR |
| SCFD2 | SCFD2\_HUMAN | Sec1 family domain-containing protein 2 | 507 | 501 | SCFD2 | SCFD2\_HUMAN | Sec1 family domain-containing protein 2 | 7C(125.0477) | NaN | X | NaN | NaN | NaN | NaN | 7.538727e+04 | 7.538727e+04 | NaN | NaN | ALAQVFCEESGLSPLLQK |
| CAND1 | CAND1\_HUMAN | Cullin-associated NEDD8-dissociated protein 1 | 92 | 81 | CAND1 | CAND1\_HUMAN | Cullin-associated NEDD8-dissociated protein 1 | 12C(125.0477) | NaN | X | NaN | NaN | NaN | NaN | 7.538492e+04 | 7.538492e+04 | NaN | NaN | EYQVETIVDTLCTNMLSDK |
| THEM6 | THEM6\_HUMAN | Protein THEM6 | 146 | 142 | THEM6 | THEM6\_HUMAN | Protein THEM6 | 5C(125.0477) | NaN | X | NaN | NaN | NaN | NaN | 7.538465e+04 | 7.538465e+04 | NaN | NaN | DGFVCALLR |
| CLTC | CLH1\_HUMAN | Clathrin heavy chain 1 | 736 | 731 | CLTC | CLH1\_HUMAN | Clathrin heavy chain 1 | 6C(125.0477) | NaN | X | NaN | NaN | NaN | NaN | 7.527237e+04 | 4.961246e+04 | 2.565991e+04 | NaN | YIQAACK |
| FBXO4 | FBX4\_HUMAN | F-box only protein 4 | 283 | 282 | FBXO4 | FBX4\_HUMAN | F-box only protein 4 | 2C(125.0477) | NaN | X | NaN | NaN | NaN | NaN | 7.522043e+04 | 7.522043e+04 | NaN | NaN | VCEVVDGFIYVANAEAHKR |
| STRADB | STRAB\_HUMAN | STE20-related kinase adapter protein beta | 180 | 171 | STRADB | STRAB\_HUMAN | STE20-related kinase adapter protein beta | 10C(125.0477) | NaN | X | NaN | NaN | NaN | NaN | 7.506155e+04 | 7.506155e+04 | NaN | NaN | GLNYLHQNGCIHR |
| GLUL | GLNA\_HUMAN | Glutamine synthetase | 359 | 358 | GLUL | GLNA\_HUMAN | Glutamine synthetase | 2C(125.0477) | NaN | X | NaN | NaN | NaN | NaN | 7.505765e+04 | 7.505765e+04 | NaN | NaN | TCLLNETGDEPFQYK |
| DCXR | DCXR\_HUMAN | L-xylulose reductase | 138 | 126 | DCXR | DCXR\_HUMAN | L-xylulose reductase | 13C(125.0477) | NaN | X | NaN | NaN | NaN | NaN | 7.505438e+04 | 7.505438e+04 | NaN | NaN | GVPGAIVNVSSQCSQR |
| YARS1 | SYYC\_HUMAN | Tyrosine--tRNA ligase, cytoplasmic | 250 | 248 | YARS1 | SYYC\_HUMAN | Tyrosine--tRNA ligase, cytoplasmic | 3C(125.0477) | NaN | X | NaN | NaN | NaN | NaN | 7.503120e+04 | 7.503120e+04 | NaN | NaN | AFCEPGNVENNGVLSFIK |
| TLN2 | TLN2\_HUMAN | Talin-2 | 958 | 947 | TLN2 | TLN2\_HUMAN | Talin-2 | 12C(125.0477) | NaN | X | NaN | NaN | NaN | NaN | 7.502810e+04 | 7.502810e+04 | NaN | NaN | NPAAQQQLVQSCK |
| EIF3J | EIF3J\_HUMAN | Eukaryotic translation initiation factor 3 subunit J | 207 | 199 | EIF3J | EIF3J\_HUMAN | Eukaryotic translation initiation factor 3 subunit J | 9C(125.0477) | NaN | X | NaN | NaN | NaN | NaN | 7.494611e+04 | 7.494611e+04 | NaN | NaN | ITNSLTVLCSEK |
| CPNE3 | CPNE3\_HUMAN | Copine-3 | 15 | 9 | CPNE3 | CPNE3\_HUMAN | Copine-3 | 7C(125.0477) | NaN | X | NaN | NaN | NaN | NaN | 7.490452e+04 | 7.490452e+04 | NaN | NaN | VALNVSCANLLDKDIGSK |
| PABPC1 | PABP1\_HUMAN | Polyadenylate-binding protein 1 | 128 | 114 | PABPC1 | PABP1\_HUMAN | Polyadenylate-binding protein 1 | 15C(125.0477) | NaN | X | NaN | NaN | NaN | NaN | 7.484188e+04 | 7.484188e+04 | NaN | NaN | ALYDTFSAFGNILSCK |
| HECTD4 | HECD4\_HUMAN | Probable E3 ubiquitin-protein ligase HECTD4 | 1735 | 1729 | HECTD4 | HECD4\_HUMAN | Probable E3 ubiquitin-protein ligase HECTD4 | 7C(125.0477) | NaN | X | NaN | NaN | NaN | NaN | 7.469380e+04 | 7.469380e+04 | NaN | NaN | QAVAALCALGGFK |
| KBTBD8 | KBTB8\_HUMAN | Kelch repeat and BTB domain-containing protein 8 | 587 | 584 | KBTBD8 | KBTB8\_HUMAN | Kelch repeat and BTB domain-containing protein 8 | 4C(125.0477) | NaN | X | NaN | NaN | NaN | NaN | 7.466912e+04 | 7.466912e+04 | NaN | NaN | HFECAVAK |
| ROCK2 | ROCK2\_HUMAN | Rho-associated protein kinase 2 | 1257 | 1253 | ROCK2 | ROCK2\_HUMAN | Rho-associated protein kinase 2 | 5C(125.0477) | NaN | X | NaN | NaN | NaN | NaN | 7.461855e+04 | 7.461855e+04 | NaN | NaN | SNYICHK |
| GSTP1 | GSTP1\_HUMAN | Glutathione S-transferase P | 48 | 46 | GSTP1 | GSTP1\_HUMAN | Glutathione S-transferase P | 3C(125.0477) | NaN | X | NaN | NaN | NaN | NaN | 7.442755e+04 | 2.203816e+04 | 5.238939e+04 | NaN | ASCLYGQLPK |
| VAC14 | VAC14\_HUMAN | Protein VAC14 homolog | 79 | 71 | VAC14 | VAC14\_HUMAN | Protein VAC14 homolog | 9C(125.0477) | NaN | X | NaN | NaN | NaN | NaN | 7.415609e+04 | 7.415609e+04 | NaN | NaN | GGLIGLAACSIALGK |
| VPS4A | VPS4A\_HUMAN | Vacuolar protein sorting-associated protein 4A | 233 | 218 | VPS4A | VPS4A\_HUMAN | Vacuolar protein sorting-associated protein 4A | 16C(125.0477) | NaN | X | NaN | NaN | NaN | NaN | 7.412245e+04 | 7.412245e+04 | NaN | NaN | QHKPSIIFIDEVDSLCGSR |
| APEH | ACPH\_HUMAN | Acylamino-acid-releasing enzyme | 526 | 498 | APEH | ACPH\_HUMAN | Acylamino-acid-releasing enzyme | 29C(125.0477) | NaN | X | NaN | NaN | NaN | NaN | 7.407984e+04 | 7.407984e+04 | NaN | NaN | TQVPMVVMPHGGPHSSFVTAWMLFPAMLCK |
| XPO4 | XPO4\_HUMAN | Exportin-4 | 492 | 487 | XPO4 | XPO4\_HUMAN | Exportin-4 | 6C(125.0477) | NaN | X | NaN | NaN | NaN | NaN | 7.404512e+04 | 7.404512e+04 | NaN | NaN | IAAEHCIPLLTSLLEER |
| SPTB | SPTB1\_HUMAN | Spectrin beta chain, erythrocytic | 960, 963 | 956 | SPTB | SPTB1\_HUMAN | Spectrin beta chain, erythrocytic | 5C(15.9949) 8C(125.0477) | NaN | X | X | NaN | NaN | NaN | 7.402072e+04 | NaN | 7.402072e+04 | NaN | VHNYCVDCEETSKWITDK |
| NIT1 | NIT1\_HUMAN | Deaminated glutathione amidase | 145 | 142 | NIT1 | NIT1\_HUMAN | Deaminated glutathione amidase | 4C(125.0477) | NaN | X | NaN | NaN | NaN | NaN | 7.400343e+04 | 7.400343e+04 | NaN | NaN | IYNCHVLLNSK |
| DPP3 | DPP3\_HUMAN | Dipeptidyl peptidase 3 | 509 | 499 | DPP3 | DPP3\_HUMAN | Dipeptidyl peptidase 3 | 11C(125.0477) | NaN | X | NaN | NaN | NaN | NaN | 7.391572e+04 | 7.391572e+04 | NaN | NaN | FSTIASSYEECR |
| DIAPH1 | DIAP1\_HUMAN | Protein diaphanous homolog 1 | 1227 | 1224 | DIAPH1 | DIAP1\_HUMAN | Protein diaphanous homolog 1 | 4C(125.0477) | NaN | X | NaN | NaN | NaN | NaN | 7.368920e+04 | 7.368920e+04 | NaN | NaN | KAGCAVTSLLASELTK |
| VCP | TERA\_HUMAN | Transitional endoplasmic reticulum ATPase | 691 | 678 | VCP | TERA\_HUMAN | Transitional endoplasmic reticulum ATPase | 14C(125.0477) | NaN | X | NaN | NaN | NaN | NaN | 7.357644e+04 | 7.357644e+04 | NaN | NaN | MTNGFSGADLTEICQR |
| DFFA | DFFA\_HUMAN | DNA fragmentation factor subunit alpha | 289 | 281 | DFFA | DFFA\_HUMAN | DNA fragmentation factor subunit alpha | 9C(125.0477) | NaN | X | NaN | NaN | NaN | NaN | 7.354048e+04 | 7.354048e+04 | NaN | NaN | KTETVQEACER |
| PFAS | PUR4\_HUMAN | Phosphoribosylformylglycinamidine synthase | 831 | 819 | PFAS | PUR4\_HUMAN | Phosphoribosylformylglycinamidine synthase | 13C(125.0477) | NaN | X | NaN | NaN | NaN | NaN | 7.348660e+04 | 7.348660e+04 | NaN | NaN | APGSLVISAYAVCPDITATVTPDLK |
| TSNAX | TSNAX\_HUMAN | Translin-associated protein X | 202 | 201 | TSNAX | TSNAX\_HUMAN | Translin-associated protein X | 2C(125.0477) | NaN | X | NaN | NaN | NaN | NaN | 7.348332e+04 | 7.348332e+04 | NaN | NaN | MCINSVGNGDIDTPFEVSQFLR |
| SENP8 | SENP8\_HUMAN | Sentrin-specific protease 8 | 64 | 64 | SENP8 | SENP8\_HUMAN | Sentrin-specific protease 8 | 1C(125.0477) | NaN | X | NaN | NaN | NaN | NaN | 7.343712e+04 | 7.343712e+04 | NaN | NaN | CTSNPAEIAMFLEPLDLPNKR |
| AP1B1 | AP1B1\_HUMAN | AP-1 complex subunit beta-1 | 921 | 914 | AP1B1 | AP1B1\_HUMAN | AP-1 complex subunit beta-1 | 8C(125.0477) | NaN | X | NaN | NaN | NaN | NaN | 7.341494e+04 | 7.341494e+04 | NaN | NaN | IQPGNPSCTDLELSLK |
| LPIN2 | LPIN2\_HUMAN | Phosphatidate phosphatase LPIN2 | 790 | 786 | LPIN2 | LPIN2\_HUMAN | Phosphatidate phosphatase LPIN2 | 5C(125.0477) | NaN | X | NaN | NaN | NaN | NaN | 7.329921e+04 | 7.329921e+04 | NaN | NaN | FKIECLNDIK |
| NRAS | RASN\_HUMAN | GTPase NRas | 80 | 74 | NRAS | RASN\_HUMAN | GTPase NRas | 7C(125.0477) | NaN | X | NaN | NaN | NaN | NaN | 7.327227e+04 | 7.327227e+04 | NaN | NaN | TGEGFLCVFAINNSK |
| FKBP1A | FKB1A\_HUMAN | Peptidyl-prolyl cis-trans isomerase FKBP1A | 23 | 20 | FKBP1A | FKB1A\_HUMAN | Peptidyl-prolyl cis-trans isomerase FKBP1A | 4C(125.0477) | NaN | X | NaN | NaN | NaN | NaN | 7.317501e+04 | 7.317501e+04 | NaN | NaN | GQTCVVHYTGMLEDGK |
| GNAI3 | GNAI3\_HUMAN | Guanine nucleotide-binding protein G(i) subunit alpha-3 | 254 | 249 | GNAI3 | GNAI3\_HUMAN | Guanine nucleotide-binding protein G(i) subunit alpha-3 | 6C(125.0477) | NaN | X | NaN | NaN | NaN | NaN | 7.314426e+04 | NaN | 7.314426e+04 | NaN | LFDSICNNK |
| APIP | MTNB\_HUMAN | Methylthioribulose-1-phosphate dehydratase | 32 | 26 | APIP | MTNB\_HUMAN | Methylthioribulose-1-phosphate dehydratase | 7C(125.0477) | NaN | X | NaN | NaN | NaN | NaN | 7.311898e+04 | 7.311898e+04 | NaN | NaN | YLIPELCK |
| UCHL5 | UCHL5\_HUMAN | Ubiquitin carboxyl-terminal hydrolase isozyme L5 | 191 | 183 | UCHL5 | UCHL5\_HUMAN | Ubiquitin carboxyl-terminal hydrolase isozyme L5 | 9C(125.0477) | NaN | X | NaN | NaN | NaN | NaN | 7.305663e+04 | 7.305663e+04 | NaN | NaN | EGPIDLGACNQDDWISAVRPVIEK |
| ZFYVE1 | ZFYV1\_HUMAN | Zinc finger FYVE domain-containing protein 1 | 45 | 45 | ZFYVE1 | ZFYV1\_HUMAN | Zinc finger FYVE domain-containing protein 1 | 1C(125.0477) | NaN | X | NaN | NaN | NaN | NaN | 7.304780e+04 | 7.304780e+04 | NaN | NaN | CEEELHR |
| PSMB2 | PSB2\_HUMAN | Proteasome subunit beta type-2 | 163 | 163 | PSMB2 | PSB2\_HUMAN | Proteasome subunit beta type-2 | 1C(125.0477) | NaN | X | NaN | NaN | NaN | NaN | 7.294158e+04 | 7.294158e+04 | NaN | NaN | CLEELQKR |
| MYH8 | MYH8\_HUMAN | Myosin-8 | 1442 | 1438 | MYH8 | MYH8\_HUMAN | Myosin-8 | 5C(57.0215) | X | NaN | NaN | NaN | NaN | NaN | 7.292273e+04 | 7.292273e+04 | NaN | NaN | SNAACAALDKK |
| XPO7 | XPO7\_HUMAN | Exportin-7 | 525 | 508 | XPO7 | XPO7\_HUMAN | Exportin-7 | 18C(125.0477) | NaN | X | NaN | NaN | NaN | NaN | 7.291130e+04 | 7.291130e+04 | NaN | NaN | VSFASTDEQDAMDGELVCR |
| SNX5 | SNX5\_HUMAN | Sorting nexin-5 | 397 | 389 | SNX5 | SNX5\_HUMAN | Sorting nexin-5 | 9C(125.0477) | NaN | X | NaN | NaN | NaN | NaN | 7.290787e+04 | 7.290787e+04 | NaN | NaN | NNVSLLQSCIDLFK |
| ATG4B | ATG4B\_HUMAN | Cysteine protease ATG4B | 183 | 182 | ATG4B | ATG4B\_HUMAN | Cysteine protease ATG4B | 2C(15.9949) | NaN | NaN | X | NaN | NaN | NaN | 7.288322e+04 | 7.288322e+04 | NaN | NaN | LCRTSVPCAGATAFPADSDR |
| NCBP1 | NCBP1\_HUMAN | Nuclear cap-binding protein subunit 1 | 36 | 21 | NCBP1 | NCBP1\_HUMAN | Nuclear cap-binding protein subunit 1 | 16C(125.0477) | NaN | X | NaN | NaN | NaN | NaN | 7.288157e+04 | 7.288157e+04 | NaN | NaN | TSDANETEDHLESLICK |
| HSPA14 | HSP7E\_HUMAN | Heat shock 70 kDa protein 14 | 335 | 332 | HSPA14 | HSP7E\_HUMAN | Heat shock 70 kDa protein 14 | 4C(125.0477) | NaN | X | NaN | NaN | NaN | NaN | 7.281088e+04 | 7.281088e+04 | NaN | NaN | VVLCGGSSR |
| HSPA2 | HSP72\_HUMAN | Heat shock-related 70 kDa protein 2 | 606 | 605 | HSPA2 | HSP72\_HUMAN | Heat shock-related 70 kDa protein 2 | 2C(125.0477) | NaN | X | NaN | NaN | NaN | NaN | 7.279924e+04 | NaN | 7.279924e+04 | NaN | VCNPIISK |
| SOD1 | SODC\_HUMAN | Superoxide dismutase [Cu-Zn] | 147 | 138 | SOD1 | SODC\_HUMAN | Superoxide dismutase [Cu-Zn] | 10C(125.0477) | NaN | X | NaN | NaN | NaN | NaN | 7.251580e+04 | 7.251580e+04 | NaN | NaN | TGNAGSRLACGVIGIAQ |
| ZC3HAV1 | ZCCHV\_HUMAN | Zinc finger CCCH-type antiviral protein 1 | 798 | 792 | ZC3HAV1 | ZCCHV\_HUMAN | Zinc finger CCCH-type antiviral protein 1 | 7C(125.0477) | NaN | X | NaN | NaN | NaN | NaN | 7.250241e+04 | 7.250241e+04 | NaN | NaN | AYVESICSNNFDSFLHETHENK |
| VPS13C | VP13C\_HUMAN | Vacuolar protein sorting-associated protein 13C | 2588 | 2588 | VPS13C | VP13C\_HUMAN | Vacuolar protein sorting-associated protein 13C | 1C(125.0477) | NaN | X | NaN | NaN | NaN | NaN | 7.250205e+04 | 7.250205e+04 | NaN | NaN | CQLFIQPAGILEHQYK |
| BRCC3 | BRCC3\_HUMAN | Lys-63-specific deubiquitinase BRCC36 | 240 | 238 | BRCC3 | BRCC3\_HUMAN | Lys-63-specific deubiquitinase BRCC36 | 3C(125.0477) | NaN | X | NaN | NaN | NaN | NaN | 7.248872e+04 | 7.248872e+04 | NaN | NaN | ILCQEEQDAYRR |
| NASP | NASP\_HUMAN | Nuclear autoantigenic sperm protein | 254 | 244 | NASP | NASP\_HUMAN | Nuclear autoantigenic sperm protein | 11C(125.0477) | NaN | X | NaN | NaN | NaN | NaN | 7.243300e+04 | 7.243300e+04 | NaN | NaN | SVSGTDVQEECR |
| PSMD2 | PSMD2\_HUMAN | 26S proteasome non-ATPase regulatory subunit 2 | 598 | 589 | PSMD2 | PSMD2\_HUMAN | 26S proteasome non-ATPase regulatory subunit 2 | 10C(125.0477) | NaN | X | NaN | NaN | NaN | NaN | 7.241273e+04 | 7.241273e+04 | NaN | NaN | SFANTLVDVCAYAGSGNVLK |
| ADK | ADK\_HUMAN | Adenosine kinase | 106 | 100 | ADK | ADK\_HUMAN | Adenosine kinase | 7C(125.0477) | NaN | X | NaN | NaN | NaN | NaN | 7.239966e+04 | 7.239966e+04 | NaN | NaN | AATFFGCIGIDKFGEILK |
| GNL1 | GNL1\_HUMAN | Guanine nucleotide-binding protein-like 1 | 365 | 356 | GNL1 | GNL1\_HUMAN | Guanine nucleotide-binding protein-like 1 | 10C(125.0477) | NaN | X | NaN | NaN | NaN | NaN | 7.225923e+04 | 7.225923e+04 | NaN | NaN | YKDGVVTIGCVGFPNVGK |
| PRPSAP1 | KPRA\_HUMAN | Phosphoribosyl pyrophosphate synthase-associated protein 1 | 19 | 10 | PRPSAP1 | KPRA\_HUMAN | Phosphoribosyl pyrophosphate synthase-associated protein 1 | 10C(125.0477) | NaN | X | NaN | NaN | NaN | NaN | 7.220889e+04 | 6.566496e+04 | 6.543933e+03 | NaN | VFSANSTAACTELAK |
| CLTB | CLCB\_HUMAN | Clathrin light chain B | 199 | 195 | CLTB | CLCB\_HUMAN | Clathrin light chain B | 5C(125.0477) | NaN | X | NaN | NaN | NaN | NaN | 7.217194e+04 | NaN | 7.217194e+04 | NaN | VAQLCDFNPK |
| RAP1B | RAP1B\_HUMAN | Ras-related protein Rap-1b | 141 | 137 | RAP1B | RAP1B\_HUMAN | Ras-related protein Rap-1b | 5C(125.0477) | NaN | X | NaN | NaN | NaN | NaN | 7.210203e+04 | 7.210203e+04 | NaN | NaN | QWNNCAFLESSAK |
| CUL2 | CUL2\_HUMAN | Cullin-2 | 232 | 221 | CUL2 | CUL2\_HUMAN | Cullin-2 | 12C(125.0477) | NaN | X | NaN | NaN | NaN | NaN | 7.192931e+04 | 7.192931e+04 | NaN | NaN | QEASNLLQESNCSQYMEK |
| UBR1 | UBR1\_HUMAN | E3 ubiquitin-protein ligase UBR1 | 350 | 342 | UBR1 | UBR1\_HUMAN | E3 ubiquitin-protein ligase UBR1 | 9C(125.0477) | NaN | X | NaN | NaN | NaN | NaN | 7.185323e+04 | 7.185323e+04 | NaN | NaN | EEPDSENPCLISR |
| EZR | EZRI\_HUMAN | Ezrin | 117 | 108 | EZR | EZRI\_HUMAN | Ezrin | 10C(125.0477) | NaN | X | NaN | NaN | NaN | NaN | 7.170734e+04 | 7.170734e+04 | NaN | NaN | EGILSDEIYCPPETAVLLGSYAVQAK |
| PGD | 6PGD\_HUMAN | 6-phosphogluconate dehydrogenase, decarboxylating | 199 | 185 | PGD | 6PGD\_HUMAN | 6-phosphogluconate dehydrogenase, decarboxylating | 15C(125.0477) | NaN | X | NaN | NaN | NaN | NaN | 7.160462e+04 | 7.160462e+04 | NaN | NaN | MVHNGIEYGDMQLICEAYHLMK |
| GSTM2 | GSTM2\_HUMAN | Glutathione S-transferase Mu 2 | 87 | 84 | GSTM2 | GSTM2\_HUMAN | Glutathione S-transferase Mu 2 | 4C(125.0477) | NaN | X | NaN | NaN | NaN | NaN | 7.158584e+04 | 7.158584e+04 | NaN | NaN | HNLCGESEK |
| CAPN5 | CAN5\_HUMAN | Calpain-5 | 404 | 396 | CAPN5 | CAN5\_HUMAN | Calpain-5 | 9C(125.0477) | NaN | X | NaN | NaN | NaN | NaN | 7.154291e+04 | 7.154291e+04 | NaN | NaN | KPEDEVLICIQQRPK |
| QARS1 | SYQ\_HUMAN | Glutamine--tRNA ligase | 298 | 293 | QARS1 | SYQ\_HUMAN | Glutamine--tRNA ligase | 6C(125.0477) | NaN | X | NaN | NaN | NaN | NaN | 7.122846e+04 | 7.122846e+04 | NaN | NaN | ANNGICFLR |
| NAPA | SNAA\_HUMAN | Alpha-soluble NSF attachment protein | 103 | 95 | NAPA | SNAA\_HUMAN | Alpha-soluble NSF attachment protein | 9C(125.0477) | NaN | X | NaN | NaN | NaN | NaN | 7.119786e+04 | 7.119786e+04 | NaN | NaN | ADPQEAINCLMR |
| GLO1 | LGUL\_HUMAN | Lactoylglutathione lyase | 61 | 61 | GLO1 | LGUL\_HUMAN | Lactoylglutathione lyase | 1C(125.0477) | NaN | X | NaN | NaN | NaN | NaN | 7.118956e+04 | 7.118956e+04 | NaN | NaN | CDFPIMK |
| TTC38 | TTC38\_HUMAN | Tetratricopeptide repeat protein 38 | 380 | 374 | TTC38 | TTC38\_HUMAN | Tetratricopeptide repeat protein 38 | 7C(125.0477) | NaN | X | NaN | NaN | NaN | NaN | 7.117663e+04 | 7.117663e+04 | NaN | NaN | DVGLPLCQALVEAEDGNPDR |
| ATP11B | AT11B\_HUMAN | Phospholipid-transporting ATPase IF | 741 | 737 | ATP11B | AT11B\_HUMAN | Phospholipid-transporting ATPase IF | 5C(125.0477) | NaN | X | NaN | NaN | NaN | NaN | 7.110595e+04 | 3.147665e+04 | 3.962931e+04 | NaN | SDSECAEQLR |
| RAB3GAP2 | RBGPR\_HUMAN | Rab3 GTPase-activating protein non-catalytic subunit | 896 | 892 | RAB3GAP2 | RBGPR\_HUMAN | Rab3 GTPase-activating protein non-catalytic subunit | 5C(125.0477) | NaN | X | NaN | NaN | NaN | NaN | 7.110179e+04 | 7.110179e+04 | NaN | NaN | QLEDCLILQTLLHSK |
| PSMB2 | PSB2\_HUMAN | Proteasome subunit beta type-2 | 163 | 163 | PSMB2 | PSB2\_HUMAN | Proteasome subunit beta type-2 | 1C(125.0477) | NaN | X | NaN | NaN | NaN | NaN | 7.107974e+04 | 7.107974e+04 | NaN | NaN | CLEELQK |
| DIAPH1 | DIAP1\_HUMAN | Protein diaphanous homolog 1 | 267 | 262 | DIAPH1 | DIAP1\_HUMAN | Protein diaphanous homolog 1 | 6C(125.0477) | NaN | X | NaN | NaN | NaN | NaN | 7.100804e+04 | 7.100804e+04 | NaN | NaN | LLSALCILPQPEDMNER |
| PPP6R1 | PP6R1\_HUMAN | Serine/threonine-protein phosphatase 6 regulatory subunit 1 | 455 | 443 | PPP6R1 | PP6R1\_HUMAN | Serine/threonine-protein phosphatase 6 regulatory subunit 1 | 13C(125.0477) | NaN | X | NaN | NaN | NaN | NaN | 7.090365e+04 | 7.090365e+04 | NaN | NaN | ILTSWEENDRVQCAGGPR |
| HNRNPF | HNRPF\_HUMAN | Heterogeneous nuclear ribonucleoprotein F | 290 | 276 | HNRNPF | HNRPF\_HUMAN | Heterogeneous nuclear ribonucleoprotein F | 15C(125.0477) | NaN | X | NaN | NaN | NaN | NaN | 7.084689e+04 | 7.084689e+04 | NaN | NaN | YGDSEFTVQSTTGHCVHMR |
| GAPVD1 | GAPD1\_HUMAN | GTPase-activating protein and VPS9 domain-containing protein 1 | 374 | 373 | GAPVD1 | GAPD1\_HUMAN | GTPase-activating protein and VPS9 domain-containing protein 1 | 2C(125.0477) | NaN | X | NaN | NaN | NaN | NaN | 7.083438e+04 | 7.083438e+04 | NaN | NaN | SCVAAFLDVVIGGR |
| MTHFS | MTHFS\_HUMAN | 5-formyltetrahydrofolate cyclo-ligase | 160 | 160 | MTHFS | MTHFS\_HUMAN | 5-formyltetrahydrofolate cyclo-ligase | 1C(125.0477) | NaN | X | NaN | NaN | NaN | NaN | 7.070923e+04 | 7.070923e+04 | NaN | NaN | CLQHQEVKPYTLALAFK |
| UBR4 | UBR4\_HUMAN | E3 ubiquitin-protein ligase UBR4 | 4890 | 4874 | UBR4 | UBR4\_HUMAN | E3 ubiquitin-protein ligase UBR4 | 17C(125.0477) | NaN | X | NaN | NaN | NaN | NaN | 7.065028e+04 | 7.065028e+04 | NaN | NaN | QQGYSTVSHFNIVHYDCHLAAVR |
| ATG7 | ATG7\_HUMAN | Ubiquitin-like modifier-activating enzyme ATG7 | 406 | 398 | ATG7 | ATG7\_HUMAN | Ubiquitin-like modifier-activating enzyme ATG7 | 9C(125.0477) | NaN | X | NaN | NaN | NaN | NaN | 7.061821e+04 | 7.061821e+04 | NaN | NaN | QPLYEFEDCLGGGKPK |
| TMEM50B | TM50B\_HUMAN | Transmembrane protein 50B | 89 | 82 | TMEM50B | TM50B\_HUMAN | Transmembrane protein 50B | 8C(125.0477) | NaN | X | NaN | NaN | NaN | NaN | 7.052055e+04 | NaN | 7.052055e+04 | NaN | GDSYESGCLGR |
| UBA1 | UBA1\_HUMAN | Ubiquitin-like modifier-activating enzyme 1 | 906 | 890 | UBA1 | UBA1\_HUMAN | Ubiquitin-like modifier-activating enzyme 1 | 17C(125.0477) | NaN | X | NaN | NaN | NaN | NaN | 7.036259e+04 | 7.036259e+04 | NaN | NaN | IIPAIATTTAAVVGLVCLELYK |
| ZZEF1 | ZZEF1\_HUMAN | Zinc finger ZZ-type and EF-hand domain-containing protein 1 | 2466 | 2465 | ZZEF1 | ZZEF1\_HUMAN | Zinc finger ZZ-type and EF-hand domain-containing protein 1 | 2C(125.0477) | NaN | X | NaN | NaN | NaN | NaN | 7.013716e+04 | 7.013716e+04 | NaN | NaN | ICFLMAHDALNAPLHILR |
| HBD | HBD\_HUMAN | Hemoglobin subunit delta | 94 | 84 | HBD | HBD\_HUMAN | Hemoglobin subunit delta | 11C(125.0477) | NaN | X | NaN | NaN | NaN | NaN | 7.005749e+04 | NaN | 7.005749e+04 | NaN | GTFSQLSELHCDKLHVDPENFR |
| ENO1 | ENOA\_HUMAN | Alpha-enolase | 357, 357 | 344 | ENO1 | ENOA\_HUMAN | Alpha-enolase | 14C(125.0477) 14C(57.0215) | X | X | NaN | NaN | NaN | NaN | 6.999180e+04 | 4.518159e+04 | NaN | 24810.2070 | VNQIGSVTESLQACK |
| GPI | G6PI\_HUMAN | Glucose-6-phosphate isomerase | 404 | 401 | GPI | G6PI\_HUMAN | Glucose-6-phosphate isomerase | 4C(125.0477) | NaN | X | NaN | NaN | NaN | NaN | 6.993956e+04 | 6.993956e+04 | NaN | NaN | MIPCDFLIPVQTQHPIRK |
| HSP90AA1 | HS90A\_HUMAN | Heat shock protein HSP 90-alpha | 420 | 419 | HSP90AA1 | HS90A\_HUMAN | Heat shock protein HSP 90-alpha | 2C(125.0477) | NaN | X | NaN | NaN | NaN | NaN | 6.990920e+04 | 6.990920e+04 | NaN | NaN | KCLELFTELAEDKENYK |
| HDGF | HDGF\_HUMAN | Hepatoma-derived growth factor | 108 | 107 | HDGF | HDGF\_HUMAN | Hepatoma-derived growth factor | 2C(125.0477) | NaN | X | NaN | NaN | NaN | NaN | 6.972435e+04 | 4.763533e+04 | 2.208901e+04 | NaN | SCVEEPEPEPEAAEGDGDK |
| CPNE3 | CPNE3\_HUMAN | Copine-3 | 249 | 242 | CPNE3 | CPNE3\_HUMAN | Copine-3 | 8C(125.0477) | NaN | X | NaN | NaN | NaN | NaN | 6.971826e+04 | 2.848235e+04 | 4.123591e+04 | NaN | SSPVEFECINEK |
| UBE3C | UBE3C\_HUMAN | Ubiquitin-protein ligase E3C | 1051 | 1045 | UBE3C | UBE3C\_HUMAN | Ubiquitin-protein ligase E3C | 7C(125.0477) | NaN | X | NaN | NaN | NaN | NaN | 6.960675e+04 | 6.960675e+04 | NaN | NaN | LPTASTCMNLLK |
| FOCAD | FOCAD\_HUMAN | Focadhesin | 657 | 655 | FOCAD | FOCAD\_HUMAN | Focadhesin | 3C(125.0477) | NaN | X | NaN | NaN | NaN | NaN | 6.956950e+04 | 6.956950e+04 | NaN | NaN | LSCDTRPLILK |
| CAP1 | CAP1\_HUMAN | Adenylyl cyclase-associated protein 1 | 375 | 367 | CAP1 | CAP1\_HUMAN | Adenylyl cyclase-associated protein 1 | 9C(125.0477) | NaN | X | NaN | NaN | NaN | NaN | 6.953871e+04 | NaN | 6.953871e+04 | NaN | INSITVDNCK |
| USO1 | USO1\_HUMAN | General vesicular transport factor p115 | 303 | 289 | USO1 | USO1\_HUMAN | General vesicular transport factor p115 | 15C(125.0477) | NaN | X | NaN | NaN | NaN | NaN | 6.953718e+04 | 6.953718e+04 | NaN | NaN | VLVSPTNPPGATSSCQK |
| PGM2 | PGM2\_HUMAN | Phosphoglucomutase-2 | 86 | 73 | PGM2 | PGM2\_HUMAN | Phosphoglucomutase-2 | 14C(125.0477) | NaN | X | NaN | NaN | NaN | NaN | 6.951171e+04 | 6.951171e+04 | NaN | NaN | MNDLTIIQTTQGFCR |
| HINT1 | HINT1\_HUMAN | Adenosine 5'-monophosphoramidase HINT1 | 84 | 84 | HINT1 | HINT1\_HUMAN | Adenosine 5'-monophosphoramidase HINT1 | 1C(125.0477) | NaN | X | NaN | NaN | NaN | NaN | 6.941235e+04 | 6.941235e+04 | NaN | NaN | CAADLGLNKGYR |
| LXN | LXN\_HUMAN | Latexin | 70 | 68 | LXN | LXN\_HUMAN | Latexin | 3C(125.0477) | NaN | X | NaN | NaN | NaN | NaN | 6.932638e+04 | 6.932638e+04 | NaN | NaN | VNCTAEVLYPSTGQETAPEVNFTFEGETGKNPDEEDNTFYQR |
| RABGGTA | PGTA\_HUMAN | Geranylgeranyl transferase type-2 subunit alpha | 371 | 361 | RABGGTA | PGTA\_HUMAN | Geranylgeranyl transferase type-2 subunit alpha | 11C(125.0477) | NaN | X | NaN | NaN | NaN | NaN | 6.928227e+04 | 6.928227e+04 | NaN | NaN | STVLQSELESCK |
| LANCL1 | LANC1\_HUMAN | Glutathione S-transferase LANCL1 | 108 | 103 | LANCL1 | LANC1\_HUMAN | Glutathione S-transferase LANCL1 | 6C(125.0477) | NaN | X | NaN | NaN | NaN | NaN | 6.927777e+04 | 6.927777e+04 | NaN | NaN | SITFLCGDAGPLAVAAVLYHK |
| DSC1 | DSC1\_HUMAN | Desmocollin-1 | 196 | 190 | DSC1 | DSC1\_HUMAN | Desmocollin-1 | 7C(125.0477) | NaN | X | NaN | NaN | NaN | NaN | 6.926771e+04 | NaN | 4.305079e+04 | 26216.9160 | DTGDIFCTR |
| RALGAPA1 | RGPA1\_HUMAN | Ral GTPase-activating protein subunit alpha-1 | 1712 | 1699 | RALGAPA1 | RGPA1\_HUMAN | Ral GTPase-activating protein subunit alpha-1 | 14C(125.0477) | NaN | X | NaN | NaN | NaN | NaN | 6.926762e+04 | 6.926762e+04 | NaN | NaN | TGISLNIPAPQPVCISEK |
| DNAJC13 | DJC13\_HUMAN | DnaJ homolog subfamily C member 13 | 2082 | 2072 | DNAJC13 | DJC13\_HUMAN | DnaJ homolog subfamily C member 13 | 11C(125.0477) | NaN | X | NaN | NaN | NaN | NaN | 6.923583e+04 | 6.923583e+04 | NaN | NaN | VIHALSENELCVR |
| PPP1R21 | PPR21\_HUMAN | Protein phosphatase 1 regulatory subunit 21 | 200 | 197 | PPP1R21 | PPR21\_HUMAN | Protein phosphatase 1 regulatory subunit 21 | 4C(57.0215) | X | NaN | NaN | NaN | NaN | NaN | 6.919866e+04 | 6.919866e+04 | NaN | NaN | TEECQLQLK |
| ADK | ADK\_HUMAN | Adenosine kinase | 160 | 150 | ADK | ADK\_HUMAN | Adenosine kinase | 11C(125.0477) | NaN | X | NaN | NaN | NaN | NaN | 6.913334e+04 | 6.913334e+04 | NaN | NaN | SLIANLAAANCYKK |
| RNH1 | RINI\_HUMAN | Ribonuclease inhibitor | 209 | 206 | RNH1 | RINI\_HUMAN | Ribonuclease inhibitor | 4C(31.9988) | NaN | NaN | NaN | X | NaN | NaN | 6.888420e+04 | NaN | 6.888420e+04 | NaN | LESCGVTSDNCR |
| AMPD2 | AMPD2\_HUMAN | AMP deaminase 2 | 230 | 230 | AMPD2 | AMPD2\_HUMAN | AMP deaminase 2 | 1C(125.0477) | NaN | X | NaN | NaN | NaN | NaN | 6.878453e+04 | 6.878453e+04 | NaN | NaN | CGVPFTDLLDAAK |
| BPNT1 | BPNT1\_HUMAN | 3'(2'),5'-bisphosphate nucleotidase 1 | 206 | 202 | BPNT1 | BPNT1\_HUMAN | 3'(2'),5'-bisphosphate nucleotidase 1 | 5C(125.0477) | NaN | X | NaN | NaN | NaN | NaN | 6.875527e+04 | 6.875527e+04 | NaN | NaN | LVTDCVAAMNPDAVLR |
| IDE | IDE\_HUMAN | Insulin-degrading enzyme | 904 | 899 | IDE | IDE\_HUMAN | Insulin-degrading enzyme | 6C(125.0477) | NaN | X | NaN | NaN | NaN | NaN | 6.864088e+04 | 6.864088e+04 | NaN | NaN | KLSAECAK |
| CR1 | CR1\_HUMAN | Complement receptor type 1 | 1591 | 1591 | CR1 | CR1\_HUMAN | Complement receptor type 1 | 1C(125.0477) | NaN | X | NaN | NaN | NaN | NaN | 6.859440e+04 | NaN | 6.859440e+04 | NaN | CTAPEVENAIR |
| HSPA4 | HSP74\_HUMAN | Heat shock 70 kDa protein 4 | 245 | 235 | HSPA4 | HSP74\_HUMAN | Heat shock 70 kDa protein 4 | 11C(125.0477) | NaN | X | NaN | NaN | NaN | NaN | 6.858415e+04 | 6.858415e+04 | NaN | NaN | KFDEVLVNHFCEEFGK |
| PGK1 | PGK1\_HUMAN | Phosphoglycerate kinase 1 | 99 | 92 | PGK1 | PGK1\_HUMAN | Phosphoglycerate kinase 1 | 8C(125.0477) | NaN | X | NaN | NaN | NaN | NaN | 6.850750e+04 | 6.850750e+04 | NaN | NaN | DVLFLKDCVGPEVEK |
| PPP2R1B | 2AAB\_HUMAN | Serine/threonine-protein phosphatase 2A 65 kDa regulatory subunit A beta isoform | 389 | 371 | PPP2R1B | 2AAB\_HUMAN | Serine/threonine-protein phosphatase 2A 65 kDa regulatory subunit A beta isoform | 19C(125.0477) | NaN | X | NaN | NaN | NaN | NaN | 6.835436e+04 | 6.835436e+04 | NaN | NaN | ENTIEHLLPLFLAQLKDECPDVR |
| MYH9 | MYH9\_HUMAN | Myosin-9 | 931 | 924 | MYH9 | MYH9\_HUMAN | Myosin-9 | 8C(125.0477) | NaN | X | NaN | NaN | NaN | NaN | 6.816792e+04 | 6.816792e+04 | NaN | NaN | VEEEEERCQHLQAEKK |
| HNRNPH1 | HNRH1\_HUMAN | Heterogeneous nuclear ribonucleoprotein H | 22 | 17 | HNRNPH1 | HNRH1\_HUMAN | Heterogeneous nuclear ribonucleoprotein H | 6C(125.0477) | NaN | X | NaN | NaN | NaN | NaN | 6.812507e+04 | 6.812507e+04 | NaN | NaN | GLPWSCSADEVQR |
| CAP1 | CAP1\_HUMAN | Adenylyl cyclase-associated protein 1 | 375 | 365 | CAP1 | CAP1\_HUMAN | Adenylyl cyclase-associated protein 1 | 11C(125.0477) | NaN | X | NaN | NaN | NaN | NaN | 6.809197e+04 | 6.809197e+04 | NaN | NaN | GKINSITVDNCKK |
| CORO7 | CORO7\_HUMAN | Coronin-7 | 42 | 40 | CORO7 | CORO7\_HUMAN | Coronin-7 | 3C(125.0477) | NaN | X | NaN | NaN | NaN | NaN | 6.807809e+04 | 6.807809e+04 | NaN | NaN | SSCSLIAFNSDRPGVLGIVPLQGQGEDK |
| UROD | DCUP\_HUMAN | Uroporphyrinogen decarboxylase | 294 | 293 | UROD | DCUP\_HUMAN | Uroporphyrinogen decarboxylase | 2C(15.9949) | NaN | NaN | X | NaN | NaN | NaN | 6.799850e+04 | 6.799850e+04 | NaN | NaN | ECVGKTVTLQGNLDPCALYASEEEIGQLVK |
| ENOSF1 | ENOF1\_HUMAN | Mitochondrial enolase superfamily member 1 | 67 | 61 | ENOSF1 | ENOF1\_HUMAN | Mitochondrial enolase superfamily member 1 | 7C(125.0477) | NaN | X | NaN | NaN | NaN | NaN | 6.797503e+04 | 6.797503e+04 | NaN | NaN | GTEVVVCAVNALAHHVLNK |
| CLIC1 | CLIC1\_HUMAN | Chloride intracellular channel protein 1 | 191 | 184 | CLIC1 | CLIC1\_HUMAN | Chloride intracellular channel protein 1 | 8C(125.0477) | NaN | X | NaN | NaN | NaN | NaN | 6.796422e+04 | 6.796422e+04 | NaN | NaN | LHIVQVVCK |
| AGO3 | AGO3\_HUMAN | Protein argonaute-3 | 36 | 32 | AGO3 | AGO3\_HUMAN | Protein argonaute-3 | 5C(125.0477) | NaN | X | NaN | NaN | NaN | NaN | 6.786471e+04 | 6.786471e+04 | NaN | NaN | LLANCFQVEIPK |
| GET3 | GET3\_HUMAN | ATPase GET3 | 289, 289 | 270 | GET3 | GET3\_HUMAN | ATPase GET3 | 20C(125.0477) 20C(57.0215) | X | X | NaN | NaN | NaN | NaN | 6.779583e+04 | 6.779583e+04 | NaN | NaN | IDTHNIIVNQLVFPDPEKPCK |
| ENDOD1 | ENDD1\_HUMAN | Endonuclease domain-containing 1 protein | 190 | 185 | ENDOD1 | ENDD1\_HUMAN | Endonuclease domain-containing 1 protein | 6C(125.0477) | NaN | X | NaN | NaN | NaN | NaN | 6.777041e+04 | 6.777041e+04 | NaN | NaN | ALTPQCGSGEDLYILTGTVPSDYR |
| TRAPPC8 | TPPC8\_HUMAN | Trafficking protein particle complex subunit 8 | 505 | 505 | TRAPPC8 | TPPC8\_HUMAN | Trafficking protein particle complex subunit 8 | 1C(125.0477) | NaN | X | NaN | NaN | NaN | NaN | 6.775443e+04 | 6.775443e+04 | NaN | NaN | CVLLSAELLK |
| IGHM | IGHM\_HUMAN | Immunoglobulin heavy constant mu | 88 | 77 | IGHM | IGHM\_HUMAN | Immunoglobulin heavy constant mu | 12C(125.0477) | NaN | X | NaN | NaN | NaN | NaN | 6.769839e+04 | NaN | 6.769839e+04 | NaN | DVMQGTDEHVVCK |
| PSMA5 | PSA5\_HUMAN | Proteasome subunit alpha type-5 | 165 | 150 | PSMA5 | PSA5\_HUMAN | Proteasome subunit alpha type-5 | 16C(125.0477) | NaN | X | NaN | NaN | NaN | NaN | 6.755427e+04 | 6.755427e+04 | NaN | NaN | GPQLFHMDPSGTFVQCDAR |
| UBA1 | UBA1\_HUMAN | Ubiquitin-like modifier-activating enzyme 1 | 1039, 1040 | 1033 | UBA1 | UBA1\_HUMAN | Ubiquitin-like modifier-activating enzyme 1 | 7C(-34.0809) 8C(31.9988) | NaN | NaN | NaN | X | X | NaN | 6.749920e+04 | 6.749920e+04 | NaN | NaN | ALVLELCCNDESGEDVEVPYVR |
| VCPIP1 | VCIP1\_HUMAN | Deubiquitinating protein VCPIP1 | 397 | 391 | VCPIP1 | VCIP1\_HUMAN | Deubiquitinating protein VCPIP1 | 7C(125.0477) | NaN | X | NaN | NaN | NaN | NaN | 6.747884e+04 | 6.747884e+04 | NaN | NaN | LEEDGGCVIGGDR |
| ERMAP | ERMAP\_HUMAN | Erythroid membrane-associated protein | 310 | 305 | ERMAP | ERMAP\_HUMAN | Erythroid membrane-associated protein | 6C(125.0477) | NaN | X | NaN | NaN | NaN | NaN | 6.746075e+04 | 6.746075e+04 | NaN | NaN | WILGVCSESVSR |
| AMPD2 | AMPD2\_HUMAN | AMP deaminase 2 | 263 | 254 | AMPD2 | AMPD2\_HUMAN | AMP deaminase 2 | 10C(125.0477) | NaN | X | NaN | NaN | NaN | NaN | 6.740760e+04 | 6.740760e+04 | NaN | NaN | YMALSLQSFCPTTR |
| HSPH1 | HS105\_HUMAN | Heat shock protein 105 kDa | 650 | 647 | HSPH1 | HS105\_HUMAN | Heat shock protein 105 kDa | 4C(15.9949) | NaN | NaN | X | NaN | NaN | NaN | 6.735480e+04 | 6.735480e+04 | NaN | NaN | DKLCGPYEKFICEQDHQNFLR |
| ALDH9A1 | AL9A1\_HUMAN | 4-trimethylaminobutyraldehyde dehydrogenase | 355 | 353 | ALDH9A1 | AL9A1\_HUMAN | 4-trimethylaminobutyraldehyde dehydrogenase | 3C(125.0477) | NaN | X | NaN | NaN | NaN | NaN | 6.734462e+04 | 6.734462e+04 | NaN | NaN | VLCGGDIYVPEDPKLK |
| FKBP1A | FKB1A\_HUMAN | Peptidyl-prolyl cis-trans isomerase FKBP1A | 23 | 20 | FKBP1A | FKB1A\_HUMAN | Peptidyl-prolyl cis-trans isomerase FKBP1A | 4C(125.0477) | NaN | X | NaN | NaN | NaN | NaN | 6.731620e+04 | 6.731620e+04 | NaN | NaN | GQTCVVHYTGMLEDGKK |
| DDI2 | DDI2\_HUMAN | Protein DDI1 homolog 2 | 332 | 329 | DDI2 | DDI2\_HUMAN | Protein DDI1 homolog 2 | 4C(125.0477) | NaN | X | NaN | NaN | NaN | NaN | 6.718283e+04 | 6.718283e+04 | NaN | NaN | RHQCSIDLK |
| USP11 | UBP11\_HUMAN | Ubiquitin carboxyl-terminal hydrolase 11 | 428 | 423 | USP11 | UBP11\_HUMAN | Ubiquitin carboxyl-terminal hydrolase 11 | 6C(125.0477) | NaN | X | NaN | NaN | NaN | NaN | 6.714877e+04 | 6.714877e+04 | NaN | NaN | EYVELCDAAGRPDQEVAQEAWQNHK |
| PDE12 | PDE12\_HUMAN | 2',5'-phosphodiesterase 12 | 250 | 248 | PDE12 | PDE12\_HUMAN | 2',5'-phosphodiesterase 12 | 3C(125.0477) | NaN | X | NaN | NaN | NaN | NaN | 6.711642e+04 | 6.711642e+04 | NaN | NaN | LHCTPGDGQR |
| SNX1 | SNX1\_HUMAN | Sorting nexin-1 | 318 | 313 | SNX1 | SNX1\_HUMAN | Sorting nexin-1 | 6C(125.0477) | NaN | X | NaN | NaN | NaN | NaN | 6.710598e+04 | 6.710598e+04 | NaN | NaN | LQEVECEEQR |
| PSMA6 | PSA6\_HUMAN | Proteasome subunit alpha type-6 | 47 | 46 | PSMA6 | PSA6\_HUMAN | Proteasome subunit alpha type-6 | 2C(125.0477) | NaN | X | NaN | NaN | NaN | NaN | 6.692068e+04 | NaN | 6.692068e+04 | NaN | DCAVIVTQK |
| XK | XK\_HUMAN | Membrane transport protein XK | 405 | 401 | XK | XK\_HUMAN | Membrane transport protein XK | 5C(125.0477) | NaN | X | NaN | NaN | NaN | NaN | 6.691930e+04 | NaN | 6.691930e+04 | NaN | QQKPCEPIGK |
| LAMP1 | LAMP1\_HUMAN | Lysosome-associated membrane glycoprotein 1 | 338 | 338 | LAMP1 | LAMP1\_HUMAN | Lysosome-associated membrane glycoprotein 1 | 1C(125.0477) | NaN | X | NaN | NaN | NaN | NaN | 6.690555e+04 | NaN | 6.690555e+04 | NaN | CNAEEHVR |
| SERPINA1 | A1AT\_HUMAN | Alpha-1-antitrypsin | 256 | 247 | SERPINA1 | A1AT\_HUMAN | Alpha-1-antitrypsin | 10C(57.0215) | X | NaN | NaN | NaN | NaN | NaN | 6.670966e+04 | 6.670966e+04 | NaN | NaN | RLGMFNIQHCK |
| TTLL12 | TTL12\_HUMAN | Tubulin--tyrosine ligase-like protein 12 | 418 | 410 | TTLL12 | TTL12\_HUMAN | Tubulin--tyrosine ligase-like protein 12 | 9C(125.0477) | NaN | X | NaN | NaN | NaN | NaN | 6.662616e+04 | 6.662616e+04 | NaN | NaN | WGEDNHWICKPWNLAR |
| PIEZO2 | PIEZ2\_HUMAN | Piezo-type mechanosensitive ion channel component 2 | 375 | 372 | PIEZO2 | PIEZ2\_HUMAN | Piezo-type mechanosensitive ion channel component 2 | 4C(-34.0809) | NaN | NaN | NaN | NaN | X | NaN | 6.660776e+04 | NaN | NaN | 66607.7600 | ALACSPIQITAGR |
| BCAM | BCAM\_HUMAN | Basal cell adhesion molecule | 522 | 518 | BCAM | BCAM\_HUMAN | Basal cell adhesion molecule | 5C(125.0477) | NaN | X | NaN | NaN | NaN | NaN | 6.654974e+04 | NaN | 6.654974e+04 | NaN | DGISCEASNPHGNKR |
| CCT8 | TCPQ\_HUMAN | T-complex protein 1 subunit theta | 430 | 422 | CCT8 | TCPQ\_HUMAN | T-complex protein 1 subunit theta | 9C(125.0477) | NaN | X | NaN | NaN | NaN | NaN | 6.651061e+04 | 6.651061e+04 | NaN | NaN | QITSYGETCPGLEQYAIKK |
| IGHG4 | IGHG4\_HUMAN | Immunoglobulin heavy constant gamma 4 | 83 | 80 | IGHG4 | IGHG4\_HUMAN | Immunoglobulin heavy constant gamma 4 | 4C(57.0215) | X | NaN | NaN | NaN | NaN | NaN | 6.642224e+04 | 1.872831e+04 | 4.769393e+04 | NaN | TYTCNVDHKPSNTK |
| CDC42 | CDC42\_HUMAN | Cell division control protein 42 homolog | 18 | 17 | CDC42 | CDC42\_HUMAN | Cell division control protein 42 homolog | 2C(125.0477) | NaN | X | NaN | NaN | NaN | NaN | 6.633912e+04 | 6.633912e+04 | NaN | NaN | TCLLISYTTNK |
| PPP6R1 | PP6R1\_HUMAN | Serine/threonine-protein phosphatase 6 regulatory subunit 1 | 37 | 20 | PPP6R1 | PP6R1\_HUMAN | Serine/threonine-protein phosphatase 6 regulatory subunit 1 | 18C(125.0477) | NaN | X | NaN | NaN | NaN | NaN | 6.630965e+04 | 6.630965e+04 | NaN | NaN | EDLSLPELLDEEDVLQECK |
| SPTBN1 | SPTB2\_HUMAN | Spectrin beta chain, non-erythrocytic 1 | 1389 | 1382 | SPTBN1 | SPTB2\_HUMAN | Spectrin beta chain, non-erythrocytic 1 | 8C(125.0477) | NaN | X | NaN | NaN | NaN | NaN | 6.629373e+04 | NaN | 6.629373e+04 | NaN | AELFTQSCADLDK |
| THOP1 | THOP1\_HUMAN | Thimet oligopeptidase | 231 | 230 | THOP1 | THOP1\_HUMAN | Thimet oligopeptidase | 2C(125.0477) | NaN | X | NaN | NaN | NaN | NaN | 6.627843e+04 | 6.627843e+04 | NaN | NaN | KCHVPETR |
| DHX9 | DHX9\_HUMAN | ATP-dependent RNA helicase A | 12 | 6 | DHX9 | DHX9\_HUMAN | ATP-dependent RNA helicase A | 7C(125.0477) | NaN | X | NaN | NaN | NaN | NaN | 6.627138e+04 | 6.627138e+04 | NaN | NaN | NFLYAWCGK |
| ACTBL2 | ACTBL\_HUMAN | Beta-actin-like protein 2 | 53 | 52 | ACTBL2 | ACTBL\_HUMAN | Beta-actin-like protein 2 | 2C(15.9949) | NaN | NaN | X | NaN | NaN | NaN | 6.614894e+04 | 6.614894e+04 | NaN | NaN | DCYVGDEAQSK |
| NUBP1 | NUBP1\_HUMAN | Cytosolic Fe-S cluster assembly factor NUBP1 | 235 | 221 | NUBP1 | NUBP1\_HUMAN | Cytosolic Fe-S cluster assembly factor NUBP1 | 15C(125.0477) | NaN | X | NaN | NaN | NaN | NaN | 6.591331e+04 | 6.591331e+04 | NaN | NaN | LPIIGVVENMSGFICPK |
| ATP7A | ATP7A\_HUMAN | Copper-transporting ATPase 1 | 182 | 177 | ATP7A | ATP7A\_HUMAN | Copper-transporting ATPase 1 | 6C(15.9949) | NaN | NaN | X | NaN | NaN | NaN | 6.591010e+04 | NaN | NaN | 65910.1000 | VEGMTCHSCTSTIEGK |
| UBA6 | UBA6\_HUMAN | Ubiquitin-like modifier-activating enzyme 6 | 231 | 215 | UBA6 | UBA6\_HUMAN | Ubiquitin-like modifier-activating enzyme 6 | 17C(125.0477) | NaN | X | NaN | NaN | NaN | NaN | 6.575901e+04 | 6.575901e+04 | NaN | NaN | EIFISNITQANPGIVTCLENHPHK |
| CUL4A | CUL4A\_HUMAN | Cullin-4A | 289 | 266 | CUL4A | CUL4A\_HUMAN | Cullin-4A | 24C(125.0477) | NaN | X | NaN | NaN | NaN | NaN | 6.572216e+04 | 6.572216e+04 | NaN | NaN | RLEEEGDRVITYLDHSTQKPLIACVEK |
| AP2A1 | AP2A1\_HUMAN | AP-2 complex subunit alpha-1 | 830 | 812 | AP2A1 | AP2A1\_HUMAN | AP-2 complex subunit alpha-1 | 19C(125.0477) | NaN | X | NaN | NaN | NaN | NaN | 6.563066e+04 | 6.563066e+04 | NaN | NaN | VAAQVDGGAQVQQVLNIECLR |
| SGTA | SGTA\_HUMAN | Small glutamine-rich tetratricopeptide repeat-containing protein alpha | 148 | 138 | SGTA | SGTA\_HUMAN | Small glutamine-rich tetratricopeptide repeat-containing protein alpha | 11C(125.0477) | NaN | X | NaN | NaN | NaN | NaN | 6.551773e+04 | 4.682519e+04 | 1.869254e+04 | NaN | LGNYAGAVQDCER |
| SCYL1 | SCYL1\_HUMAN | N-terminal kinase-like protein | 309 | 300 | SCYL1 | SCYL1\_HUMAN | N-terminal kinase-like protein | 10C(125.0477) | NaN | X | NaN | NaN | NaN | NaN | 6.522995e+04 | 6.522995e+04 | NaN | NaN | SLDAFPEDFCR |
| ADPRS | ADPRS\_HUMAN | ADP-ribosylhydrolase ARH3 | 287 | 287 | ADPRS | ADPRS\_HUMAN | ADP-ribosylhydrolase ARH3 | 1C(125.0477) | NaN | X | NaN | NaN | NaN | NaN | 6.514260e+04 | 6.514260e+04 | NaN | NaN | CMEPDPEIPSAFNSLQR |
| HSPA1A | HS71A\_HUMAN | Heat shock 70 kDa protein 1A | 574 | 574 | HSPA1A | HS71A\_HUMAN | Heat shock 70 kDa protein 1A | 1C(125.0477) | NaN | X | NaN | NaN | NaN | NaN | 6.509586e+04 | 6.509586e+04 | NaN | NaN | CQEVISWLDANTLAEKDEFEHKR |
| KRT1 | K2C1\_HUMAN | Keratin, type II cytoskeletal 1 | 49 | 46 | KRT1 | K2C1\_HUMAN | Keratin, type II cytoskeletal 1 | 4C(125.0477) | NaN | X | NaN | NaN | NaN | NaN | 6.504727e+04 | NaN | 6.504727e+04 | NaN | FSSCGGGGGSFGAGGGFGSR |
| AKR1B1 | ALDR\_HUMAN | Aldo-keto reductase family 1 member B1 | 187 | 178 | AKR1B1 | ALDR\_HUMAN | Aldo-keto reductase family 1 member B1 | 10C(125.0477) | NaN | X | NaN | NaN | NaN | NaN | 6.497380e+04 | 6.497380e+04 | NaN | NaN | YKPAVNQIECHPYLTQEK |
| TPP2 | TPP2\_HUMAN | Tripeptidyl-peptidase 2 | 150 | 144 | TPP2 | TPP2\_HUMAN | Tripeptidyl-peptidase 2 | 7C(125.0477) | NaN | X | NaN | NaN | NaN | NaN | 6.491617e+04 | NaN | 6.491617e+04 | NaN | VALAEACR |
| TRIM58 | TRI58\_HUMAN | E3 ubiquitin-protein ligase TRIM58 | 334 | 329 | TRIM58 | TRI58\_HUMAN | E3 ubiquitin-protein ligase TRIM58 | 6C(125.0477) | NaN | X | NaN | NaN | NaN | NaN | 6.489257e+04 | 6.489257e+04 | NaN | NaN | FDTWPCILGLQSFSSGR |
| CFL1 | COF1\_HUMAN | Cofilin-1 | 39 | 34 | CFL1 | COF1\_HUMAN | Cofilin-1 | 6C(125.0477) | NaN | X | NaN | NaN | NaN | NaN | 6.484788e+04 | 6.484788e+04 | NaN | NaN | KAVLFCLSEDK |
| TARS1 | SYTC\_HUMAN | Threonine--tRNA ligase 1, cytoplasmic | 107 | 100 | TARS1 | SYTC\_HUMAN | Threonine--tRNA ligase 1, cytoplasmic | 8C(125.0477) | NaN | X | NaN | NaN | NaN | NaN | 6.474952e+04 | 6.474952e+04 | NaN | NaN | TTPYQIACGISQGLADNTVIAK |
| RPN1 | RPN1\_HUMAN | Dolichyl-diphosphooligosaccharide--protein glycosyltransferase subunit 1 | 545 | 537 | RPN1 | RPN1\_HUMAN | Dolichyl-diphosphooligosaccharide--protein glycosyltransferase subunit 1 | 9C(125.0477) | NaN | X | NaN | NaN | NaN | NaN | 6.468507e+04 | NaN | 6.468507e+04 | NaN | LKTEGSDLCDR |
| GNAI3 | GNAI3\_HUMAN | Guanine nucleotide-binding protein G(i) subunit alpha-3 | 139 | 133 | GNAI3 | GNAI3\_HUMAN | Guanine nucleotide-binding protein G(i) subunit alpha-3 | 7C(125.0477) | NaN | X | NaN | NaN | NaN | NaN | 6.451135e+04 | NaN | 6.451135e+04 | NaN | DGGVQACFSR |
| PFAS | PUR4\_HUMAN | Phosphoribosylformylglycinamidine synthase | 975 | 970 | PFAS | PUR4\_HUMAN | Phosphoribosylformylglycinamidine synthase | 6C(125.0477) | NaN | X | NaN | NaN | NaN | NaN | 6.448376e+04 | 6.448376e+04 | NaN | NaN | DAGLHCLELGHTGEAGPHAMVR |
| CLTC | CLH1\_HUMAN | Clathrin heavy chain 1 | 151 | 145 | CLTC | CLH1\_HUMAN | Clathrin heavy chain 1 | 7C(125.0477) | NaN | X | NaN | NaN | NaN | NaN | 6.442570e+04 | 6.442570e+04 | NaN | NaN | HSSLAGCQIINYR |
| PPA1 | IPYR\_HUMAN | Inorganic pyrophosphatase | 254 | 254 | PPA1 | IPYR\_HUMAN | Inorganic pyrophosphatase | 1C(125.0477) | NaN | X | NaN | NaN | NaN | NaN | 6.426662e+04 | 6.426662e+04 | NaN | NaN | CDPDAAR |
| ANXA7 | ANXA7\_HUMAN | Annexin A7 | 363 | 357 | ANXA7 | ANXA7\_HUMAN | Annexin A7 | 7C(125.0477) | NaN | X | NaN | NaN | NaN | NaN | 6.420169e+04 | 4.048337e+04 | 2.371832e+04 | NaN | LGTDESCFNMILATR |
| OSBP2 | OSBP2\_HUMAN | Oxysterol-binding protein 2 | 319 | 313 | OSBP2 | OSBP2\_HUMAN | Oxysterol-binding protein 2 | 7C(125.0477) | NaN | X | NaN | NaN | NaN | NaN | 6.409237e+04 | NaN | 6.409237e+04 | NaN | LDDLSTCNDLIAK |
| RHOA | RHOA\_HUMAN | Transforming protein RhoA | 16 | 8 | RHOA | RHOA\_HUMAN | Transforming protein RhoA | 9C(125.0477) | NaN | X | NaN | NaN | NaN | NaN | 6.407426e+04 | NaN | 6.407426e+04 | NaN | LVIVGDGACGK |
| HSPH1 | HS105\_HUMAN | Heat shock protein 105 kDa | 658 | 656 | HSPH1 | HS105\_HUMAN | Heat shock protein 105 kDa | 3C(125.0477) | NaN | X | NaN | NaN | NaN | NaN | 6.405519e+04 | 6.405519e+04 | NaN | NaN | FICEQDHQNFLR |
| RANGAP1 | RAGP1\_HUMAN | Ran GTPase-activating protein 1 | 303 | 295 | RANGAP1 | RAGP1\_HUMAN | Ran GTPase-activating protein 1 | 9C(125.0477) | NaN | X | NaN | NaN | NaN | NaN | 6.404502e+04 | 6.404502e+04 | NaN | NaN | LKELNLSFCEIK |
| MYH10 | MYH10\_HUMAN | Myosin-10 | 1011 | 1008 | MYH10 | MYH10\_HUMAN | Myosin-10 | 4C(125.0477) | NaN | X | NaN | NaN | NaN | NaN | 6.403900e+04 | 6.403900e+04 | NaN | NaN | IAECSSQLAEEEEKAK |
| GPX4 | GPX4\_HUMAN | Phospholipid hydroperoxide glutathione peroxidase | 102 | 97 | GPX4 | GPX4\_HUMAN | Phospholipid hydroperoxide glutathione peroxidase | 6C(125.0477) | NaN | X | NaN | NaN | NaN | NaN | 6.402574e+04 | 6.402574e+04 | NaN | NaN | ILAFPCNQFGK |
| SPTB | SPTB1\_HUMAN | Spectrin beta chain, erythrocytic | 1892 | 1871 | SPTB | SPTB1\_HUMAN | Spectrin beta chain, erythrocytic | 22C(125.0477) | NaN | X | NaN | NaN | NaN | NaN | 6.387144e+04 | 6.387144e+04 | NaN | NaN | AEAIQNKEQEVSAAWQALLDACAGRR |
| ALB | ALBU\_HUMAN | Albumin | 500 | 500 | ALB | ALBU\_HUMAN | Albumin | 1C(57.0215) | X | NaN | NaN | NaN | NaN | NaN | 6.386282e+04 | NaN | 6.386282e+04 | NaN | CCTESLVNR |
| PSMB6 | PSB6\_HUMAN | Proteasome subunit beta type-6 | 186 | 179 | PSMB6 | PSB6\_HUMAN | Proteasome subunit beta type-6 | 8C(125.0477) | NaN | X | NaN | NaN | NaN | NaN | 6.368152e+04 | 6.368152e+04 | NaN | NaN | EGMTKEECLQFTANALALAMER |
| LANCL1 | LANC1\_HUMAN | Glutathione S-transferase LANCL1 | 252 | 240 | LANCL1 | LANC1\_HUMAN | Glutathione S-transferase LANCL1 | 13C(125.0477) | NaN | X | NaN | NaN | NaN | NaN | 6.367529e+04 | 6.367529e+04 | NaN | NaN | LHSLVKPSVDYVCQLK |
| RAB14 | RAB14\_HUMAN | Ras-related protein Rab-14 | 26 | 25 | RAB14 | RAB14\_HUMAN | Ras-related protein Rab-14 | 2C(125.0477) | NaN | X | NaN | NaN | NaN | NaN | 6.362693e+04 | 6.362693e+04 | NaN | NaN | SCLLHQFTEK |
| STK38L | ST38L\_HUMAN | Serine/threonine-protein kinase 38-like | 235 | 229 | STK38L | ST38L\_HUMAN | Serine/threonine-protein kinase 38-like | 7C(125.0477) | NaN | X | NaN | NaN | NaN | NaN | 6.361677e+04 | 6.361677e+04 | NaN | NaN | LSDFGLCTGLK |
| CTPS1 | PYRG1\_HUMAN | CTP synthase 1 | 218 | 218 | CTPS1 | PYRG1\_HUMAN | CTP synthase 1 | 1C(125.0477) | NaN | X | NaN | NaN | NaN | NaN | 6.358855e+04 | 6.358855e+04 | NaN | NaN | CSNPLDTSVKEK |
| HNRNPC | HNRPC\_HUMAN | Heterogeneous nuclear ribonucleoproteins C1/C2 | 46 | 43 | HNRNPC | HNRPC\_HUMAN | Heterogeneous nuclear ribonucleoproteins C1/C2 | 4C(125.0477) | NaN | X | NaN | NaN | NaN | NaN | 6.340053e+04 | 6.340053e+04 | NaN | NaN | IVGCSVHK |
| HUWE1 | HUWE1\_HUMAN | E3 ubiquitin-protein ligase HUWE1 | 1401 | 1393 | HUWE1 | HUWE1\_HUMAN | E3 ubiquitin-protein ligase HUWE1 | 9C(125.0477) | NaN | X | NaN | NaN | NaN | NaN | 6.334932e+04 | 6.334932e+04 | NaN | NaN | AESPEEVACRKEEEER |
| AVEN | AVEN\_HUMAN | Cell death regulator Aven | 326 | 313 | AVEN | AVEN\_HUMAN | Cell death regulator Aven | 14C(125.0477) | NaN | X | NaN | NaN | NaN | NaN | 6.330981e+04 | 6.330981e+04 | NaN | NaN | SKEDGEVVQEEEVCAKPSVTEEK |
| OTUB1 | OTUB1\_HUMAN | Ubiquitin thioesterase OTUB1 | 91 | 85 | OTUB1 | OTUB1\_HUMAN | Ubiquitin thioesterase OTUB1 | 7C(125.0477) | NaN | X | NaN | NaN | NaN | NaN | 6.327766e+04 | 6.327766e+04 | NaN | NaN | TRPDGNCFYR |
| COPB1 | COPB\_HUMAN | Coatomer subunit beta | 248 | 248 | COPB1 | COPB\_HUMAN | Coatomer subunit beta | 1C(125.0477) | NaN | X | NaN | NaN | NaN | NaN | 6.294790e+04 | 6.294790e+04 | NaN | NaN | CIYNLLQSSSPAVK |
| PTPN11 | PTN11\_HUMAN | Tyrosine-protein phosphatase non-receptor type 11 | 104 | 92 | PTPN11 | PTN11\_HUMAN | Tyrosine-protein phosphatase non-receptor type 11 | 13C(125.0477) | NaN | X | NaN | NaN | NaN | NaN | 6.294543e+04 | 6.294543e+04 | NaN | NaN | NGDVIELKYPLNCADPTSER |
| LANCL1 | LANC1\_HUMAN | Glutathione S-transferase LANCL1 | 276 | 270 | LANCL1 | LANC1\_HUMAN | Glutathione S-transferase LANCL1 | 7C(125.0477) | NaN | X | NaN | NaN | NaN | NaN | 6.294261e+04 | 6.294261e+04 | NaN | NaN | DLLVHWCHGAPGVIYMLIQAYK |
| NME2P1 | NDK8\_HUMAN | Putative nucleoside diphosphate kinase | 130 | 129 | NME2P1 | NDK8\_HUMAN | Putative nucleoside diphosphate kinase | 2C(125.0477) | NaN | X | NaN | NaN | NaN | NaN | 6.292461e+04 | NaN | 6.292461e+04 | NaN | SCAHDWVYE |
| PCBP2 | PCBP2\_HUMAN | Poly(rC)-binding protein 2 | 109 | 102 | PCBP2 | PCBP2\_HUMAN | Poly(rC)-binding protein 2 | 8C(15.9949) | NaN | NaN | X | NaN | NaN | NaN | 6.288229e+04 | 6.288229e+04 | NaN | NaN | LVVPASQCGSLIGKGGCK |
| GSTM4 | GSTM4\_HUMAN | Glutathione S-transferase Mu 4 | 87 | 84 | GSTM4 | GSTM4\_HUMAN | Glutathione S-transferase Mu 4 | 4C(125.0477) | NaN | X | NaN | NaN | NaN | NaN | 6.280014e+04 | 6.280014e+04 | NaN | NaN | HNLCGETEEEK |
| HK1 | HXK1\_HUMAN | Hexokinase-1 | 158 | 148 | HK1 | HXK1\_HUMAN | Hexokinase-1 | 11C(125.0477) | NaN | X | NaN | NaN | NaN | NaN | 6.275533e+04 | 6.275533e+04 | NaN | NaN | LPVGFTFSFPCQQSK |
| STIP1 | STIP1\_HUMAN | Stress-induced-phosphoprotein 1 | 471, 461 | 454 | STIP1 | STIP1\_HUMAN | Stress-induced-phosphoprotein 1 | 18C(15.9949) 8C(31.9988) | NaN | NaN | X | X | NaN | NaN | 6.264209e+04 | 6.264209e+04 | NaN | NaN | ALDLDSSCKEAADGYQRCMMAQYNR |
| FAH | FAAA\_HUMAN | Fumarylacetoacetase | 396 | 382 | FAH | FAAA\_HUMAN | Fumarylacetoacetase | 15C(125.0477) | NaN | X | NaN | NaN | NaN | NaN | 6.264189e+04 | 6.264189e+04 | NaN | NaN | KFLLDGDEVIITGYCQGDGYR |
| IRGQ | IRGQ\_HUMAN | Immunity-related GTPase family Q protein | 28 | 21 | IRGQ | IRGQ\_HUMAN | Immunity-related GTPase family Q protein | 8C(125.0477) | NaN | X | NaN | NaN | NaN | NaN | 6.253964e+04 | 6.253964e+04 | NaN | NaN | SALIAALCDKDVETLEAPEGRPDSGVPSLR |
| HGS | HGS\_HUMAN | Hepatocyte growth factor-regulated tyrosine kinase substrate | 75 | 74 | HGS | HGS\_HUMAN | Hepatocyte growth factor-regulated tyrosine kinase substrate | 2C(125.0477) | NaN | X | NaN | NaN | NaN | NaN | 6.249959e+04 | NaN | 6.249959e+04 | NaN | NCGQTVHDEVANK |
| CNPY4 | CNPY4\_HUMAN | Protein canopy homolog 4 | 94 | 86 | CNPY4 | CNPY4\_HUMAN | Protein canopy homolog 4 | 9C(125.0477) | NaN | X | NaN | NaN | NaN | NaN | 6.242498e+04 | NaN | 6.242498e+04 | NaN | LEEALENLCER |
| CASP3 | CASP3\_HUMAN | Caspase-3 | 264 | 261 | CASP3 | CASP3\_HUMAN | Caspase-3 | 4C(125.0477) | NaN | X | NaN | NaN | NaN | NaN | 6.234334e+04 | 6.234334e+04 | NaN | NaN | QIPCIVSMLTK |
| TXNDC5 | TXND5\_HUMAN | Thioredoxin domain-containing protein 5 | 381 | 376 | TXNDC5 | TXND5\_HUMAN | Thioredoxin domain-containing protein 5 | 6C(125.0477) | NaN | X | NaN | NaN | NaN | NaN | 6.229777e+04 | NaN | 3.988585e+04 | 22411.9160 | IAEVDCTAER |
| IPO5 | IPO5\_HUMAN | Importin-5 | 180, 180 | 176 | IPO5 | IPO5\_HUMAN | Importin-5 | 5C(125.0477) 5C(57.0215) | X | X | NaN | NaN | NaN | NaN | 6.225530e+04 | 6.225530e+04 | NaN | NaN | MLVQCMQDQEHPSIR |
| LYPLA2 | LYPA2\_HUMAN | Acyl-protein thioesterase 2 | 171 | 165 | LYPLA2 | LYPA2\_HUMAN | Acyl-protein thioesterase 2 | 7C(125.0477) | NaN | X | NaN | NaN | NaN | NaN | 6.219844e+04 | 6.219844e+04 | NaN | NaN | DLAILQCHGELDPMVPVR |
| CAP1 | CAP1\_HUMAN | Adenylyl cyclase-associated protein 1 | 416 | 413 | CAP1 | CAP1\_HUMAN | Adenylyl cyclase-associated protein 1 | 4C(15.9949) | NaN | NaN | X | NaN | NaN | NaN | 6.215067e+04 | 6.215067e+04 | NaN | NaN | TDGCHAYLSKNSLDCEIVSAK |
| PPA1 | IPYR\_HUMAN | Inorganic pyrophosphatase | 254, 254 | 254 | PPA1 | IPYR\_HUMAN | Inorganic pyrophosphatase | 1C(15.9949) 1C(31.9988) | NaN | NaN | X | X | NaN | NaN | 6.214735e+04 | NaN | NaN | 62147.3480 | CDPDAAR |
| SLC19A1 | S19A1\_HUMAN | Reduced folate transporter | 220 | 220 | SLC19A1 | S19A1\_HUMAN | Reduced folate transporter | 1C(125.0477) | NaN | X | NaN | NaN | NaN | NaN | 6.210579e+04 | NaN | 6.210579e+04 | NaN | CETSASELER |
| DPP9 | DPP9\_HUMAN | Dipeptidyl peptidase 9 | 225 | 222 | DPP9 | DPP9\_HUMAN | Dipeptidyl peptidase 9 | 4C(125.0477) | NaN | X | NaN | NaN | NaN | NaN | 6.206299e+04 | 6.206299e+04 | NaN | NaN | LTFCHQGLSNVLDDPK |
| UBE2V1 | UB2V1\_HUMAN | Ubiquitin-conjugating enzyme E2 variant 1 | 71 | 64 | UBE2V1 | UB2V1\_HUMAN | Ubiquitin-conjugating enzyme E2 variant 1 | 8C(125.0477) | NaN | X | NaN | NaN | NaN | NaN | 6.197484e+04 | 6.197484e+04 | NaN | NaN | IYSLKIECGPK |
| RARS1 | SYRC\_HUMAN | Arginine--tRNA ligase, cytoplasmic | 638 | 635 | RARS1 | SYRC\_HUMAN | Arginine--tRNA ligase, cytoplasmic | 4C(125.0477) | NaN | X | NaN | NaN | NaN | NaN | 6.194781e+04 | 6.194781e+04 | NaN | NaN | MLLCEAVAAVMAK |
| RBKS | RBSK\_HUMAN | Ribokinase | 73 | 65 | RBKS | RBSK\_HUMAN | Ribokinase | 9C(125.0477) | NaN | X | NaN | NaN | NaN | NaN | 6.181759e+04 | 6.181759e+04 | NaN | NaN | LGAMTSMVCK |
| UBR4 | UBR4\_HUMAN | E3 ubiquitin-protein ligase UBR4 | 1274 | 1249 | UBR4 | UBR4\_HUMAN | E3 ubiquitin-protein ligase UBR4 | 26C(125.0477) | NaN | X | NaN | NaN | NaN | NaN | 6.179131e+04 | 6.179131e+04 | NaN | NaN | NAFANDTIPSESYISAVQAAHLGTLCSQSLPLAASLK |
| EIF3J | EIF3J\_HUMAN | Eukaryotic translation initiation factor 3 subunit J | 207 | 198 | EIF3J | EIF3J\_HUMAN | Eukaryotic translation initiation factor 3 subunit J | 10C(125.0477) | NaN | X | NaN | NaN | NaN | NaN | 6.169761e+04 | 6.169761e+04 | NaN | NaN | KITNSLTVLCSEK |
| TSC22D2 | T22D2\_HUMAN | TSC22 domain family protein 2 | 63 | 54 | TSC22D2 | T22D2\_HUMAN | TSC22 domain family protein 2 | 10C(125.0477) | NaN | X | NaN | NaN | NaN | NaN | 6.159764e+04 | 6.159764e+04 | NaN | NaN | ATDYGPEEVCER |
| PREP | PPCE\_HUMAN | Prolyl endopeptidase | 601 | 589 | PREP | PPCE\_HUMAN | Prolyl endopeptidase | 13C(125.0477) | NaN | X | NaN | NaN | NaN | NaN | 6.150396e+04 | 6.150396e+04 | NaN | NaN | YTIGHAWTTDYGCSDSK |
| DCTN1 | DCTN1\_HUMAN | Dynactin subunit 1 | 81 | 78 | DCTN1 | DCTN1\_HUMAN | Dynactin subunit 1 | 4C(125.0477) | NaN | X | NaN | NaN | NaN | NaN | 6.145952e+04 | 6.145952e+04 | NaN | NaN | YFTCDEGHGIFVR |
| FABP5 | FABP5\_HUMAN | Fatty acid-binding protein 5 | 87 | 83 | FABP5 | FABP5\_HUMAN | Fatty acid-binding protein 5 | 5C(125.0477) | NaN | X | NaN | NaN | NaN | NaN | 6.140806e+04 | 6.140806e+04 | NaN | NaN | TQTVCNFTDGALVQHQEWDGK |
| RAC1 | RAC1\_HUMAN | Ras-related C3 botulinum toxin substrate 1 | 178 | 175 | RAC1 | RAC1\_HUMAN | Ras-related C3 botulinum toxin substrate 1 | 4C(125.0477) | NaN | X | NaN | NaN | NaN | NaN | 6.138836e+04 | NaN | 6.138836e+04 | NaN | AVLCPPPVK |
| PSMA4 | PSA4\_HUMAN | Proteasome subunit alpha type-4 | 163 | 161 | PSMA4 | PSA4\_HUMAN | Proteasome subunit alpha type-4 | 3C(125.0477) | NaN | X | NaN | NaN | NaN | NaN | 6.134429e+04 | 3.128563e+04 | 3.005866e+04 | NaN | ATCIGNNSAAAVSMLK |
| C1QC | C1QC\_HUMAN | Complement C1q subcomponent subunit C | 179, 193 | 162 | C1QC | C1QC\_HUMAN | Complement C1q subcomponent subunit C | 18C(305.3076) 32C(15.9949) | NaN | NaN | X | NaN | NaN | X | 6.130436e+04 | NaN | 6.130436e+04 | NaN | VPGLYYFVYHASHTANLCVLLYRSGVKVVTFCGHTSK |
| XPO1 | XPO1\_HUMAN | Exportin-1 | 723 | 723 | XPO1 | XPO1\_HUMAN | Exportin-1 | 1C(125.0477) | NaN | X | NaN | NaN | NaN | NaN | 6.125457e+04 | 6.125457e+04 | NaN | NaN | CLSENISAAIQANGEMVTK |
| SDF2L1 | SDF2L\_HUMAN | Stromal cell-derived factor 2-like protein 1 | 149 | 149 | SDF2L1 | SDF2L\_HUMAN | Stromal cell-derived factor 2-like protein 1 | 1C(125.0477) | NaN | X | NaN | NaN | NaN | NaN | 6.125055e+04 | NaN | 6.125055e+04 | NaN | CSGQHWER |
| SELENBP1 | SBP1\_HUMAN | Methanethiol oxidase | 57 | 53 | SELENBP1 | SBP1\_HUMAN | Methanethiol oxidase | 5C(125.0477) | NaN | X | NaN | NaN | NaN | NaN | 6.124747e+04 | NaN | 6.124747e+04 | NaN | SPQYCQVIHR |
| PCBD2 | PHS2\_HUMAN | Pterin-4-alpha-carbinolamine dehydratase 2 | 109 | 100 | PCBD2 | PHS2\_HUMAN | Pterin-4-alpha-carbinolamine dehydratase 2 | 10C(125.0477) | NaN | X | NaN | NaN | NaN | NaN | 6.123977e+04 | 6.123977e+04 | NaN | NaN | VQITLTSHDCGELTK |
| PPIP5K2 | VIP2\_HUMAN | Inositol hexakisphosphate and diphosphoinositol-pentakisphosphate kinase 2 | 382 | 382 | PPIP5K2 | VIP2\_HUMAN | Inositol hexakisphosphate and diphosphoinositol-pentakisphosphate kinase 2 | 1C(125.0477) | NaN | X | NaN | NaN | NaN | NaN | 6.121644e+04 | 6.121644e+04 | NaN | NaN | CVIAVIR |
| DSTN | DEST\_HUMAN | Destrin | 135 | 133 | DSTN | DEST\_HUMAN | Destrin | 3C(125.0477) | NaN | X | NaN | NaN | NaN | NaN | 6.104108e+04 | 6.104108e+04 | NaN | NaN | HECQANGPEDLNR |
| EIF2S3 | IF2G\_HUMAN | Eukaryotic translation initiation factor 2 subunit 3 | 105 | 104 | EIF2S3 | IF2G\_HUMAN | Eukaryotic translation initiation factor 2 subunit 3 | 2C(125.0477) | NaN | X | NaN | NaN | NaN | NaN | 6.091272e+04 | 4.671783e+04 | 1.419489e+04 | NaN | SCGSSTPDEFPTDIPGTK |
| SOD1 | SODC\_HUMAN | Superoxide dismutase [Cu-Zn] | 7 | 5 | SOD1 | SODC\_HUMAN | Superoxide dismutase [Cu-Zn] | 3C(125.0477) | NaN | X | NaN | NaN | NaN | NaN | 6.090334e+04 | 6.090334e+04 | NaN | NaN | AVCVLKGDGPVQGIINFEQK |
| GNB2 | GBB2\_HUMAN | Guanine nucleotide-binding protein G(I)/G(S)/G(T) subunit beta-2 | 204 | 198 | GNB2 | GBB2\_HUMAN | Guanine nucleotide-binding protein G(I)/G(S)/G(T) subunit beta-2 | 7C(125.0477) | NaN | X | NaN | NaN | NaN | NaN | 6.084847e+04 | 2.158966e+04 | 3.925881e+04 | NaN | TFVSGACDASIK |
| CRKL | CRKL\_HUMAN | Crk-like protein | 44 | 40 | CRKL | CRKL\_HUMAN | Crk-like protein | 5C(125.0477) | NaN | X | NaN | NaN | NaN | NaN | 6.084236e+04 | 4.689533e+04 | 1.394703e+04 | NaN | DSSTCPGDYVLSVSENSR |
| FHOD1 | FHOD1\_HUMAN | FH1/FH2 domain-containing protein 1 | 43 | 40 | FHOD1 | FHOD1\_HUMAN | FH1/FH2 domain-containing protein 1 | 4C(125.0477) | NaN | X | NaN | NaN | NaN | NaN | 6.083801e+04 | 6.083801e+04 | NaN | NaN | APTCSLDGALPLGAQIPAVHR |
| GNB1 | GBB1\_HUMAN | Guanine nucleotide-binding protein G(I)/G(S)/G(T) subunit beta-1 | 25 | 24 | GNB1 | GBB1\_HUMAN | Guanine nucleotide-binding protein G(I)/G(S)/G(T) subunit beta-1 | 2C(125.0477) | NaN | X | NaN | NaN | NaN | NaN | 6.081620e+04 | 6.081620e+04 | NaN | NaN | ACADATLSQITNNIDPVGR |
| IREB2 | IREB2\_HUMAN | Iron-responsive element-binding protein 2 | 174, 178 | 170 | IREB2 | IREB2\_HUMAN | Iron-responsive element-binding protein 2 | 5C(15.9949) 9C(15.9949) | NaN | NaN | X | NaN | NaN | NaN | 6.079601e+04 | 6.079601e+04 | NaN | NaN | GQTTCRGSCDSGELGR |
| IPO7 | IPO7\_HUMAN | Importin-7 | 113 | 108 | IPO7 | IPO7\_HUMAN | Importin-7 | 6C(125.0477) | NaN | X | NaN | NaN | NaN | NaN | 6.079453e+04 | 6.079453e+04 | NaN | NaN | VQLTTCIHHIIK |
| PSMA1 | PSA1\_HUMAN | Proteasome subunit alpha type-1 | 182 | 175 | PSMA1 | PSA1\_HUMAN | Proteasome subunit alpha type-1 | 8C(125.0477) | NaN | X | NaN | NaN | NaN | NaN | 6.071838e+04 | 6.071838e+04 | NaN | NaN | HMSEFMECNLNELVK |
| WIPI2 | WIPI2\_HUMAN | WD repeat domain phosphoinositide-interacting protein 2 | 132 | 129 | WIPI2 | WIPI2\_HUMAN | WD repeat domain phosphoinositide-interacting protein 2 | 4C(125.0477) | NaN | X | NaN | NaN | NaN | NaN | 6.051522e+04 | 6.051522e+04 | NaN | NaN | LIVCLEESLYIHNIR |
| PSMG1 | PSMG1\_HUMAN | Proteasome assembly chaperone 1 | 157 | 153 | PSMG1 | PSMG1\_HUMAN | Proteasome assembly chaperone 1 | 5C(125.0477) | NaN | X | NaN | NaN | NaN | NaN | 6.039676e+04 | 6.039676e+04 | NaN | NaN | VFGSCPR |
| MYO18A | MY18A\_HUMAN | Unconventional myosin-XVIIIa | 250 | 244 | MYO18A | MY18A\_HUMAN | Unconventional myosin-XVIIIa | 7C(125.0477) | NaN | X | NaN | NaN | NaN | NaN | 6.038838e+04 | 6.038838e+04 | NaN | NaN | GPEGQACR |
| CAND1 | CAND1\_HUMAN | Cullin-associated NEDD8-dissociated protein 1 | 275 | 273 | CAND1 | CAND1\_HUMAN | Cullin-associated NEDD8-dissociated protein 1 | 3C(125.0477) | NaN | X | NaN | NaN | NaN | NaN | 6.038120e+04 | 6.038120e+04 | NaN | NaN | EYCIQAFESFVR |
| YKT6 | YKT6\_HUMAN | Synaptobrevin homolog YKT6 | 66 | 57 | YKT6 | YKT6\_HUMAN | Synaptobrevin homolog YKT6 | 10C(125.0477) | NaN | X | NaN | NaN | NaN | NaN | 6.028121e+04 | 6.028121e+04 | NaN | NaN | ASVKEQDYLCHVYVR |
| ANKRD28 | ANR28\_HUMAN | Serine/threonine-protein phosphatase 6 regulatory ankyrin repeat subunit A | 701 | 693 | ANKRD28 | ANR28\_HUMAN | Serine/threonine-protein phosphatase 6 regulatory ankyrin repeat subunit A | 9C(15.9949) | NaN | NaN | X | NaN | NaN | NaN | 6.024464e+04 | 6.024464e+04 | NaN | NaN | GAVTGHEECVDALLQHGAKCLLR |
| ATP11C | AT11C\_HUMAN | Phospholipid-transporting ATPase IG | 214 | 210 | ATP11C | AT11C\_HUMAN | Phospholipid-transporting ATPase IG | 5C(125.0477) | NaN | X | NaN | NaN | NaN | NaN | 6.021614e+04 | NaN | 6.021614e+04 | NaN | AAIECEQPQPDLYK |
| HBA1 | HBA\_HUMAN | Hemoglobin subunit alpha | 105 | 92 | HBA1 | HBA\_HUMAN | Hemoglobin subunit alpha | 14C(125.0477) | NaN | X | NaN | NaN | NaN | NaN | 6.013198e+04 | NaN | 4.845424e+04 | 11677.7420 | LRVDPVNFKLLSHCLLVTLAAHLPAEFTPAVHASLDK |
| UBR4 | UBR4\_HUMAN | E3 ubiquitin-protein ligase UBR4 | 3765 | 3749 | UBR4 | UBR4\_HUMAN | E3 ubiquitin-protein ligase UBR4 | 17C(125.0477) | NaN | X | NaN | NaN | NaN | NaN | 6.012627e+04 | 6.012627e+04 | NaN | NaN | VYHQLMGHRPQLENLLCK |
| GDI2 | GDIB\_HUMAN | Rab GDP dissociation inhibitor beta | 282 | 279 | GDI2 | GDIB\_HUMAN | Rab GDP dissociation inhibitor beta | 4C(125.0477) | NaN | X | NaN | NaN | NaN | NaN | 6.012108e+04 | NaN | 6.012108e+04 | NaN | QLICDPSYVK |
| LTA4H | LKHA4\_HUMAN | Leukotriene A-4 hydrolase | 543 | 542 | LTA4H | LKHA4\_HUMAN | Leukotriene A-4 hydrolase | 2C(125.0477) | NaN | X | NaN | NaN | NaN | NaN | 5.989024e+04 | 5.989024e+04 | NaN | NaN | LCIQSKWEDAIPLALK |
| MACROD1 | MACD1\_HUMAN | ADP-ribose glycohydrolase MACROD1 | 246 | 245 | MACROD1 | MACD1\_HUMAN | ADP-ribose glycohydrolase MACROD1 | 2C(125.0477) | NaN | X | NaN | NaN | NaN | NaN | 5.984406e+04 | 5.984406e+04 | NaN | NaN | SCYLSSLDLLLEHR |
| USP24 | UBP24\_HUMAN | Ubiquitin carboxyl-terminal hydrolase 24 | 172 | 163 | USP24 | UBP24\_HUMAN | Ubiquitin carboxyl-terminal hydrolase 24 | 10C(125.0477) | NaN | X | NaN | NaN | NaN | NaN | 5.982300e+04 | 5.982300e+04 | NaN | NaN | LGLSESDENCR |
| C2CD2L | C2C2L\_HUMAN | Phospholipid transfer protein C2CD2L | 385, 385 | 383 | C2CD2L | C2C2L\_HUMAN | Phospholipid transfer protein C2CD2L | 3C(125.0477) 3C(31.9988) | NaN | X | NaN | X | NaN | NaN | 5.979771e+04 | NaN | NaN | 59797.7100 | QLCPLTPGPGK |
| SLC19A1 | S19A1\_HUMAN | Reduced folate transporter | 246 | 244 | SLC19A1 | S19A1\_HUMAN | Reduced folate transporter | 3C(125.0477) | NaN | X | NaN | NaN | NaN | NaN | 5.978546e+04 | NaN | 5.978546e+04 | NaN | VACGDSVLAR |
| UBE2O | UBE2O\_HUMAN | (E3-independent) E2 ubiquitin-conjugating enzyme | 341 | 339 | UBE2O | UBE2O\_HUMAN | (E3-independent) E2 ubiquitin-conjugating enzyme | 3C(125.0477) | NaN | X | NaN | NaN | NaN | NaN | 5.971109e+04 | 5.971109e+04 | NaN | NaN | LGCFDHAQR |
| STAMBP | STABP\_HUMAN | STAM-binding protein | 390 | 379 | STAMBP | STABP\_HUMAN | STAM-binding protein | 12C(125.0477) | NaN | X | NaN | NaN | NaN | NaN | 5.956353e+04 | 4.288172e+04 | 1.668181e+04 | NaN | LTDHGLEEISSCR |
| FAM114A1 | NXP20\_HUMAN | Protein NOXP20 | 458 | 458 | FAM114A1 | NXP20\_HUMAN | Protein NOXP20 | 1C(57.0215) | X | NaN | NaN | NaN | NaN | NaN | 5.950419e+04 | 5.950419e+04 | NaN | NaN | CIEQLHKVAELILHGQEEEKPAQDQAK |
| SEMA7A | SEM7A\_HUMAN | Semaphorin-7A | 541 | 540 | SEMA7A | SEM7A\_HUMAN | Semaphorin-7A | 2C(125.0477) | NaN | X | NaN | NaN | NaN | NaN | 5.944081e+04 | NaN | 5.944081e+04 | NaN | ECPNPKPDK |
| DCTN1 | DCTN1\_HUMAN | Dynactin subunit 1 | 1252 | 1248 | DCTN1 | DCTN1\_HUMAN | Dynactin subunit 1 | 5C(125.0477) | NaN | X | NaN | NaN | NaN | NaN | 5.943141e+04 | 5.943141e+04 | NaN | NaN | VTFSCAAGFGQR |
| DNM2 | DYN2\_HUMAN | Dynamin-2 | 86 | 67 | DNM2 | DYN2\_HUMAN | Dynamin-2 | 20C(125.0477) | NaN | X | NaN | NaN | NaN | NaN | 5.941565e+04 | 5.941565e+04 | NaN | NaN | RPLILQLIFSKTEHAEFLHCK |
| DCTN4 | DCTN4\_HUMAN | Dynactin subunit 4 | 258 | 249 | DCTN4 | DCTN4\_HUMAN | Dynactin subunit 4 | 10C(125.0477) | NaN | X | NaN | NaN | NaN | NaN | 5.932502e+04 | 5.932502e+04 | NaN | NaN | LLQPDFQPVCASQLYPR |
| CLIC2 | CLIC2\_HUMAN | Chloride intracellular channel protein 2 | 229 | 223 | CLIC2 | CLIC2\_HUMAN | Chloride intracellular channel protein 2 | 7C(125.0477) | NaN | X | NaN | NaN | NaN | NaN | 5.928131e+04 | 5.928131e+04 | NaN | NaN | EEFTHTCPEDK |
| STK38L | ST38L\_HUMAN | Serine/threonine-protein kinase 38-like | 362 | 361 | STK38L | ST38L\_HUMAN | Serine/threonine-protein kinase 38-like | 2C(125.0477) | NaN | X | NaN | NaN | NaN | NaN | 5.926271e+04 | 5.926271e+04 | NaN | NaN | FCIDSENR |
| CFL1 | COF1\_HUMAN | Cofilin-1 | 139 | 133 | CFL1 | COF1\_HUMAN | Cofilin-1 | 7C(125.0477) | NaN | X | NaN | NaN | NaN | NaN | 5.916866e+04 | NaN | 5.916866e+04 | NaN | HELQANCYEEVK |
| UBE2R2 | UB2R2\_HUMAN | Ubiquitin-conjugating enzyme E2 R2 | 191 | 183 | UBE2R2 | UB2R2\_HUMAN | Ubiquitin-conjugating enzyme E2 R2 | 9C(125.0477) | NaN | X | NaN | NaN | NaN | NaN | 5.904256e+04 | 5.904256e+04 | NaN | NaN | VPTTLAEYCIK |
| RAPGEF2 | RPGF2\_HUMAN | Rap guanine nucleotide exchange factor 2 | 110 | 109 | RAPGEF2 | RPGF2\_HUMAN | Rap guanine nucleotide exchange factor 2 | 2C(125.0477) | NaN | X | NaN | NaN | NaN | NaN | 5.894132e+04 | 5.894132e+04 | NaN | NaN | DCLEKDPIDR |
| PSMC6 | PRS10\_HUMAN | 26S proteasome regulatory subunit 10B | 170 | 169 | PSMC6 | PRS10\_HUMAN | 26S proteasome regulatory subunit 10B | 2C(125.0477) | NaN | X | NaN | NaN | NaN | NaN | 5.891212e+04 | 5.891212e+04 | NaN | NaN | GCLLYGPPGTGK |
| PRPSAP1 | KPRA\_HUMAN | Phosphoribosyl pyrophosphate synthase-associated protein 1 | 19 | 10 | PRPSAP1 | KPRA\_HUMAN | Phosphoribosyl pyrophosphate synthase-associated protein 1 | 10C(125.0477) | NaN | X | NaN | NaN | NaN | NaN | 5.889235e+04 | 5.889235e+04 | NaN | NaN | VFSANSTAACTELAKR |
| DNAJA2 | DNJA2\_HUMAN | DnaJ homolog subfamily A member 2 | 308 | 303 | DNAJA2 | DNJA2\_HUMAN | DnaJ homolog subfamily A member 2 | 6C(125.0477) | NaN | X | NaN | NaN | NaN | NaN | 5.885429e+04 | NaN | 5.885429e+04 | NaN | VIEPGCVR |
| LAP3 | AMPL\_HUMAN | Cytosol aminopeptidase | 445 | 441 | LAP3 | AMPL\_HUMAN | Cytosol aminopeptidase | 5C(125.0477) | NaN | X | NaN | NaN | NaN | NaN | 5.881768e+04 | 5.881768e+04 | NaN | NaN | QVVDCQLADVNNIGK |
| PSMB2 | PSB2\_HUMAN | Proteasome subunit beta type-2 | 163 | 162 | PSMB2 | PSB2\_HUMAN | Proteasome subunit beta type-2 | 2C(125.0477) | NaN | X | NaN | NaN | NaN | NaN | 5.874947e+04 | 5.874947e+04 | NaN | NaN | KCLEELQK |
| EIF4A1 | IF4A1\_HUMAN | Eukaryotic initiation factor 4A-I | 66 | 62 | EIF4A1 | IF4A1\_HUMAN | Eukaryotic initiation factor 4A-I | 5C(125.0477) | NaN | X | NaN | NaN | NaN | NaN | 5.874548e+04 | NaN | 5.874548e+04 | NaN | AILPCIK |
| ADD2 | ADDB\_HUMAN | Beta-adducin | 177 | 174 | ADD2 | ADDB\_HUMAN | Beta-adducin | 4C(125.0477) | NaN | X | NaN | NaN | NaN | NaN | 5.871023e+04 | 5.871023e+04 | NaN | NaN | GVSCSEVTASSLIKVNILGEVVEK |
| GPS1 | CSN1\_HUMAN | COP9 signalosome complex subunit 1 | 202 | 197 | GPS1 | CSN1\_HUMAN | COP9 signalosome complex subunit 1 | 6C(125.0477) | NaN | X | NaN | NaN | NaN | NaN | 5.869712e+04 | 5.869712e+04 | NaN | NaN | HVINMCLNVIK |
| VCP | TERA\_HUMAN | Transitional endoplasmic reticulum ATPase | 105 | 96 | VCP | TERA\_HUMAN | Transitional endoplasmic reticulum ATPase | 10C(15.9949) | NaN | NaN | X | NaN | NaN | NaN | 5.868533e+04 | 5.868533e+04 | NaN | NaN | LGDVISIQPCPDVKYGK |
| PHKA2 | KPB2\_HUMAN | Phosphorylase b kinase regulatory subunit alpha, liver isoform | 760 | 753 | PHKA2 | KPB2\_HUMAN | Phosphorylase b kinase regulatory subunit alpha, liver isoform | 8C(125.0477) | NaN | X | NaN | NaN | NaN | NaN | 5.861161e+04 | 5.861161e+04 | NaN | NaN | DDHGDVDCEK |
| EEF2 | EF2\_HUMAN | Elongation factor 2 | 591 | 581 | EEF2 | EF2\_HUMAN | Elongation factor 2 | 11C(125.0477) | NaN | X | NaN | NaN | NaN | NaN | 5.856097e+04 | 3.427563e+04 | 2.428534e+04 | NaN | ETVSEESNVLCLSK |
| NR2C2AP | NR2CA\_HUMAN | Nuclear receptor 2C2-associated protein | 76 | 74 | NR2C2AP | NR2CA\_HUMAN | Nuclear receptor 2C2-associated protein | 3C(125.0477) | NaN | X | NaN | NaN | NaN | NaN | 5.855461e+04 | 5.855461e+04 | NaN | NaN | RGCLEGSQGTQALHK |
| TF | TRFE\_HUMAN | Serotransferrin | 596 | 588 | TF | TRFE\_HUMAN | Serotransferrin | 9C(57.0215) | X | NaN | NaN | NaN | NaN | NaN | 5.853406e+04 | 3.162392e+04 | 2.691014e+04 | NaN | KPVEEYANCHLAR |
| BNIP1 | SEC20\_HUMAN | Vesicle transport protein SEC20 | 33 | 32 | BNIP1 | SEC20\_HUMAN | Vesicle transport protein SEC20 | 2C(125.0477) | NaN | X | NaN | NaN | NaN | NaN | 5.849222e+04 | 5.849222e+04 | NaN | NaN | DCSGPLSALTELNTK |
| STXBP3 | STXB3\_HUMAN | Syntaxin-binding protein 3 | 557 | 556 | STXBP3 | STXB3\_HUMAN | Syntaxin-binding protein 3 | 2C(125.0477) | NaN | X | NaN | NaN | NaN | NaN | 5.842139e+04 | 5.842139e+04 | NaN | NaN | SCEVIIGSTHVLTPK |
| STIP1 | STIP1\_HUMAN | Stress-induced-phosphoprotein 1 | 370 | 367 | STIP1 | STIP1\_HUMAN | Stress-induced-phosphoprotein 1 | 4C(125.0477) | NaN | X | NaN | NaN | NaN | NaN | 5.842096e+04 | 5.842096e+04 | NaN | NaN | GNECFQK |
| ARL6IP1 | AR6P1\_HUMAN | ADP-ribosylation factor-like protein 6-interacting protein 1 | 109 | 105 | ARL6IP1 | AR6P1\_HUMAN | ADP-ribosylation factor-like protein 6-interacting protein 1 | 5C(125.0477) | NaN | X | NaN | NaN | NaN | NaN | 5.826424e+04 | 5.826424e+04 | NaN | NaN | FHEICSNLVK |
| TTLL12 | TTL12\_HUMAN | Tubulin--tyrosine ligase-like protein 12 | 528 | 525 | TTLL12 | TTL12\_HUMAN | Tubulin--tyrosine ligase-like protein 12 | 4C(125.0477) | NaN | X | NaN | NaN | NaN | NaN | 5.823062e+04 | 5.823062e+04 | NaN | NaN | QVHCEEFIPEFEK |
| HNRNPA1 | ROA1\_HUMAN | Heterogeneous nuclear ribonucleoprotein A1 | 175 | 167 | HNRNPA1 | ROA1\_HUMAN | Heterogeneous nuclear ribonucleoprotein A1 | 9C(125.0477) | NaN | X | NaN | NaN | NaN | NaN | 5.817949e+04 | 5.817949e+04 | NaN | NaN | YHTVNGHNCEVRK |
| EEFSEC | SELB\_HUMAN | Selenocysteine-specific elongation factor | 442 | 433 | EEFSEC | SELB\_HUMAN | Selenocysteine-specific elongation factor | 10C(125.0477) | NaN | X | NaN | NaN | NaN | NaN | 5.817448e+04 | 5.817448e+04 | NaN | NaN | LDADIHTNTCR |
| VCP | TERA\_HUMAN | Transitional endoplasmic reticulum ATPase | 415 | 390 | VCP | TERA\_HUMAN | Transitional endoplasmic reticulum ATPase | 26C(125.0477) | NaN | X | NaN | NaN | NaN | NaN | 5.812520e+04 | 5.812520e+04 | NaN | NaN | LADDVDLEQVANETHGHVGADLAALCSEAALQAIRK |
| XYLB | XYLB\_HUMAN | Xylulose kinase | 161 | 148 | XYLB | XYLB\_HUMAN | Xylulose kinase | 14C(125.0477) | NaN | X | NaN | NaN | NaN | NaN | 5.809230e+04 | 5.809230e+04 | NaN | NaN | QLEAAVGGAQALSCLTGSR |
| FLNA | FLNA\_HUMAN | Filamin-A | 574 | 570 | FLNA | FLNA\_HUMAN | Filamin-A | 5C(125.0477) | NaN | X | NaN | NaN | NaN | NaN | 5.809134e+04 | 5.809134e+04 | NaN | NaN | VGTECGNQK |
| SH3GL1 | SH3G1\_HUMAN | Endophilin-A2 | 96, 96 | 83 | SH3GL1 | SH3G1\_HUMAN | Endophilin-A2 | 14C(125.0477) 14C(57.0215) | X | X | NaN | NaN | NaN | NaN | 5.801098e+04 | 5.801098e+04 | NaN | NaN | NPGYPQSEGLLGECMIR |
| UBA1 | UBA1\_HUMAN | Ubiquitin-like modifier-activating enzyme 1 | 481 | 469 | UBA1 | UBA1\_HUMAN | Ubiquitin-like modifier-activating enzyme 1 | 13C(125.0477) | NaN | X | NaN | NaN | NaN | NaN | 5.780653e+04 | 5.780653e+04 | NaN | NaN | QKYFLVGAGAIGCELLK |
| DPP3 | DPP3\_HUMAN | Dipeptidyl peptidase 3 | 176 | 166 | DPP3 | DPP3\_HUMAN | Dipeptidyl peptidase 3 | 11C(125.0477) | NaN | X | NaN | NaN | NaN | NaN | 5.776638e+04 | 5.776638e+04 | NaN | NaN | EGITTYFSGNCTMEDAK |
| RAC1 | RAC1\_HUMAN | Ras-related C3 botulinum toxin substrate 1 | 157 | 154 | RAC1 | RAC1\_HUMAN | Ras-related C3 botulinum toxin substrate 1 | 4C(125.0477) | NaN | X | NaN | NaN | NaN | NaN | 5.772584e+04 | NaN | 5.772584e+04 | NaN | YLECSALTQR |
| RFESD | RFESD\_HUMAN | Rieske domain-containing protein | 19 | 14 | RFESD | RFESD\_HUMAN | Rieske domain-containing protein | 6C(125.0477) | NaN | X | NaN | NaN | NaN | NaN | 5.767766e+04 | 5.767766e+04 | NaN | NaN | EYSSVCVGREDDIKK |
| USP14 | UBP14\_HUMAN | Ubiquitin carboxyl-terminal hydrolase 14 | 277 | 268 | USP14 | UBP14\_HUMAN | Ubiquitin carboxyl-terminal hydrolase 14 | 10C(125.0477) | NaN | X | NaN | NaN | NaN | NaN | 5.753213e+04 | 5.753213e+04 | NaN | NaN | GKENQLQLSCFINQEVK |
| ZC3HAV1 | ZCCHV\_HUMAN | Zinc finger CCCH-type antiviral protein 1 | 219 | 204 | ZC3HAV1 | ZCCHV\_HUMAN | Zinc finger CCCH-type antiviral protein 1 | 16C(125.0477) | NaN | X | NaN | NaN | NaN | NaN | 5.744426e+04 | 5.744426e+04 | NaN | NaN | EHGLNPDVVQNIQDICNSK |
| GTPBP1 | GTPB1\_HUMAN | GTP-binding protein 1 | 313 | 310 | GTPBP1 | GTPB1\_HUMAN | GTP-binding protein 1 | 4C(125.0477) | NaN | X | NaN | NaN | NaN | NaN | 5.730658e+04 | 5.730658e+04 | NaN | NaN | IDMCPANILQETLK |
| ADD1 | ADDA\_HUMAN | Alpha-adducin | 525, 525 | 516 | ADD1 | ADDA\_HUMAN | Alpha-adducin | 10C(125.0477) 10C(15.9949) | NaN | X | X | NaN | NaN | NaN | 5.729012e+04 | 5.729012e+04 | NaN | NaN | TAGPQSQVLCGVVMDR |
| SMAD2 | SMAD2\_HUMAN | Mothers against decapentaplegic homolog 2 | 70 | 64 | SMAD2 | SMAD2\_HUMAN | Mothers against decapentaplegic homolog 2 | 7C(125.0477) | NaN | X | NaN | NaN | NaN | NaN | 5.727878e+04 | 5.727878e+04 | NaN | NaN | AITTQNCNTK |
| VPS13C | VP13C\_HUMAN | Vacuolar protein sorting-associated protein 13C | 1372 | 1366 | VPS13C | VP13C\_HUMAN | Vacuolar protein sorting-associated protein 13C | 7C(125.0477) | NaN | X | NaN | NaN | NaN | NaN | 5.721186e+04 | 5.721186e+04 | NaN | NaN | ILTENLCEGTEDLDKVKPR |
| KRT4 | K2C4\_HUMAN | Keratin, type II cytoskeletal 4 | 376 | 375 | KRT4 | K2C4\_HUMAN | Keratin, type II cytoskeletal 4 | 2C(125.0477) | NaN | X | NaN | NaN | NaN | NaN | 5.709350e+04 | NaN | 5.709350e+04 | NaN | QCQTLQVSVADAEQR |
| EEF2 | EF2\_HUMAN | Elongation factor 2 | 651, 651 | 649 | EEF2 | EF2\_HUMAN | Elongation factor 2 | 3C(125.0477) 3C(57.0215) | X | X | NaN | NaN | NaN | NaN | 5.698984e+04 | 5.698984e+04 | NaN | NaN | IWCFGPDGTGPNILTDITK |
| SEC31A | SC31A\_HUMAN | Protein transport protein Sec31A | 1130 | 1130 | SEC31A | SC31A\_HUMAN | Protein transport protein Sec31A | 1C(125.0477) | NaN | X | NaN | NaN | NaN | NaN | 5.695214e+04 | NaN | 5.695214e+04 | NaN | CLSSATDPQTK |
| EIF2S3 | IF2G\_HUMAN | Eukaryotic translation initiation factor 2 subunit 3 | 269 | 261 | EIF2S3 | IF2G\_HUMAN | Eukaryotic translation initiation factor 2 subunit 3 | 9C(125.0477) | NaN | X | NaN | NaN | NaN | NaN | 5.689401e+04 | 5.689401e+04 | NaN | NaN | SFDVNKPGCEVDDLKGGVAGGSILK |
| SUMF2 | SUMF2\_HUMAN | Inactive C-alpha-formylglycine-generating enzyme 2 | 290 | 290 | SUMF2 | SUMF2\_HUMAN | Inactive C-alpha-formylglycine-generating enzyme 2 | 1C(125.0477) | NaN | X | NaN | NaN | NaN | NaN | 5.687027e+04 | NaN | 5.687027e+04 | NaN | CAADAGRPPGEL |
| ACTR1B | ACTY\_HUMAN | Beta-centractin | 222 | 221 | ACTR1B | ACTY\_HUMAN | Beta-centractin | 2C(125.0477) | NaN | X | NaN | NaN | NaN | NaN | 5.683684e+04 | 5.683684e+04 | NaN | NaN | ACYLSINPQKDEALETEK |
| SEC31A | SC31A\_HUMAN | Protein transport protein Sec31A | 173 | 165 | SEC31A | SC31A\_HUMAN | Protein transport protein Sec31A | 9C(125.0477) | NaN | X | NaN | NaN | NaN | NaN | 5.683430e+04 | 5.683430e+04 | NaN | NaN | TQPPEDISCIAWNR |
| GORASP2 | GORS2\_HUMAN | Golgi reassembly-stacking protein 2 | 173 | 163 | GORASP2 | GORS2\_HUMAN | Golgi reassembly-stacking protein 2 | 11C(125.0477) | NaN | X | NaN | NaN | NaN | NaN | 5.680039e+04 | 5.680039e+04 | NaN | NaN | LYVYNTDTDNCR |
| TRIM58 | TRI58\_HUMAN | E3 ubiquitin-protein ligase TRIM58 | 240 | 240 | TRIM58 | TRI58\_HUMAN | E3 ubiquitin-protein ligase TRIM58 | 1C(125.0477) | NaN | X | NaN | NaN | NaN | NaN | 5.678805e+04 | 5.678805e+04 | NaN | NaN | CQRPALGLLEGVR |
| CMBL | CMBL\_HUMAN | Carboxymethylenebutenolidase homolog | 121 | 116 | CMBL | CMBL\_HUMAN | Carboxymethylenebutenolidase homolog | 6C(125.0477) | NaN | X | NaN | NaN | NaN | NaN | 5.676724e+04 | 5.676724e+04 | NaN | NaN | YLKQQCHAQK |
| GDI1 | GDIA\_HUMAN | Rab GDP dissociation inhibitor alpha | 302 | 300 | GDI1 | GDIA\_HUMAN | Rab GDP dissociation inhibitor alpha | 3C(125.0477) | NaN | X | NaN | NaN | NaN | NaN | 5.671109e+04 | 5.671109e+04 | NaN | NaN | IICILSHPIK |
| CLTC | CLH1\_HUMAN | Clathrin heavy chain 1 | 1102 | 1102 | CLTC | CLH1\_HUMAN | Clathrin heavy chain 1 | 1C(125.0477) | NaN | X | NaN | NaN | NaN | NaN | 5.667696e+04 | 5.667696e+04 | NaN | NaN | CNEPAVWSQLAK |
| RARS1 | SYRC\_HUMAN | Arginine--tRNA ligase, cytoplasmic | 369 | 363 | RARS1 | SYRC\_HUMAN | Arginine--tRNA ligase, cytoplasmic | 7C(125.0477) | NaN | X | NaN | NaN | NaN | NaN | 5.665607e+04 | 5.665607e+04 | NaN | NaN | IVFVPGCSIPLTIVK |
| RPTOR | RPTOR\_HUMAN | Regulatory-associated protein of mTOR | 86 | 84 | RPTOR | RPTOR\_HUMAN | Regulatory-associated protein of mTOR | 3C(125.0477) | NaN | X | NaN | NaN | NaN | NaN | 5.662367e+04 | 5.662367e+04 | NaN | NaN | LECWIDPLSMGPQK |
| WDR81 | WDR81\_HUMAN | WD repeat-containing protein 81 | 1653 | 1653 | WDR81 | WDR81\_HUMAN | WD repeat-containing protein 81 | 1C(125.0477) | NaN | X | NaN | NaN | NaN | NaN | 5.657516e+04 | 5.657516e+04 | NaN | NaN | CVAPLSSEDFFLSGSKDR |
| PUDP | HDHD1\_HUMAN | Pseudouridine-5'-phosphatase | 34 | 25 | PUDP | HDHD1\_HUMAN | Pseudouridine-5'-phosphatase | 10C(125.0477) | NaN | X | NaN | NaN | NaN | NaN | 5.655886e+04 | 5.655886e+04 | NaN | NaN | LYSVVFQEICNR |
| CHORDC1 | CHRD1\_HUMAN | Cysteine and histidine-rich domain-containing protein 1 | 176 | 165 | CHORDC1 | CHRD1\_HUMAN | Cysteine and histidine-rich domain-containing protein 1 | 12C(125.0477) | NaN | X | NaN | NaN | NaN | NaN | 5.654965e+04 | 5.654965e+04 | NaN | NaN | TYQGLESLEEVCVYHSGVPIFHEGMK |
| CAST | ICAL\_HUMAN | Calpastatin | 413, 413 | 410 | CAST | ICAL\_HUMAN | Calpastatin | 4C(15.9949) 4C(31.9988) | NaN | NaN | X | X | NaN | NaN | 5.652122e+04 | 5.652122e+04 | NaN | NaN | TSMCSIQSAPPEPATLK |
| ALDOA | ALDOA\_HUMAN | Fructose-bisphosphate aldolase A | 339, 339 | 332 | ALDOA | ALDOA\_HUMAN | Fructose-bisphosphate aldolase A | 8C(15.9949) 8C(31.9988) | NaN | NaN | X | X | NaN | NaN | 5.648566e+04 | 5.648566e+04 | NaN | NaN | ALANSLACQGKYTPSGQAGAAASESLFVSNHAY |
| PALS2 | PALS2\_HUMAN | Protein PALS2 | 432 | 431 | PALS2 | PALS2\_HUMAN | Protein PALS2 | 2C(125.0477) | NaN | X | NaN | NaN | NaN | NaN | 5.638023e+04 | 4.786761e+04 | 8.512621e+03 | NaN | TCILDVNPQALK |
| CCT7 | TCPH\_HUMAN | T-complex protein 1 subunit eta | 345 | 345 | CCT7 | TCPH\_HUMAN | T-complex protein 1 subunit eta | 1C(125.0477) | NaN | X | NaN | NaN | NaN | NaN | 5.633626e+04 | 2.418127e+04 | 3.215499e+04 | NaN | CQVFEETQIGGER |
| STK39 | STK39\_HUMAN | STE20/SPS1-related proline-alanine-rich protein kinase | 525 | 521 | STK39 | STK39\_HUMAN | STE20/SPS1-related proline-alanine-rich protein kinase | 5C(125.0477) | NaN | X | NaN | NaN | NaN | NaN | 5.620510e+04 | 5.620510e+04 | NaN | NaN | LASGCDGSEIPDEVK |
| DCTN4 | DCTN4\_HUMAN | Dynactin subunit 4 | 280 | 280 | DCTN4 | DCTN4\_HUMAN | Dynactin subunit 4 | 1C(125.0477) | NaN | X | NaN | NaN | NaN | NaN | 5.614004e+04 | 5.614004e+04 | NaN | NaN | CEHNLSKPEFNPTSIK |
| KRT5 | K2C5\_HUMAN | Keratin, type II cytoskeletal 5 | 407 | 406 | KRT5 | K2C5\_HUMAN | Keratin, type II cytoskeletal 5 | 2C(125.0477) | NaN | X | NaN | NaN | NaN | NaN | 5.604839e+04 | 5.604839e+04 | NaN | NaN | QCANLQNAIADAEQRGELALK |
| PSMD10 | PSD10\_HUMAN | 26S proteasome non-ATPase regulatory subunit 10 | 180 | 163 | PSMD10 | PSD10\_HUMAN | 26S proteasome non-ATPase regulatory subunit 10 | 18C(125.0477) | NaN | X | NaN | NaN | NaN | NaN | 5.600579e+04 | 5.600579e+04 | NaN | NaN | ASTNIQDTEGNTPLHLACDEER |
| ABCE1 | ABCE1\_HUMAN | ATP-binding cassette sub-family E member 1 | 65 | 65 | ABCE1 | ABCE1\_HUMAN | ATP-binding cassette sub-family E member 1 | 1C(125.0477) | NaN | X | NaN | NaN | NaN | NaN | 5.598454e+04 | 5.598454e+04 | NaN | NaN | CPFGALSIVNLPSNLEK |
| STK11 | STK11\_HUMAN | Serine/threonine-protein kinase STK11 | 73 | 65 | STK11 | STK11\_HUMAN | Serine/threonine-protein kinase STK11 | 9C(125.0477) | NaN | X | NaN | NaN | NaN | NaN | 5.597534e+04 | NaN | 5.597534e+04 | NaN | EVLDSETLCR |
| PSMC6 | PRS10\_HUMAN | 26S proteasome regulatory subunit 10B | 170, 170 | 169 | PSMC6 | PRS10\_HUMAN | 26S proteasome regulatory subunit 10B | 2C(-34.0809) 2C(125.0477) | NaN | X | NaN | NaN | X | NaN | 5.596820e+04 | NaN | 5.596820e+04 | NaN | GCLLYGPPGTGK |
| ITSN1 | ITSN1\_HUMAN | Intersectin-1 | 1629, 1621 | 1617 | ITSN1 | ITSN1\_HUMAN | Intersectin-1 | 13C(15.9949) 5C(57.0215) | X | NaN | X | NaN | NaN | NaN | 5.587963e+04 | NaN | 5.587963e+04 | NaN | SNPYCEVTMGSQCHITK |
| RPE | RPE\_HUMAN | Ribulose-phosphate 3-epimerase | 217 | 215 | RPE | RPE\_HUMAN | Ribulose-phosphate 3-epimerase | 3C(125.0477) | NaN | X | NaN | NaN | NaN | NaN | 5.584998e+04 | 5.584998e+04 | NaN | NaN | NVCSEAAQKR |
| CAPZB | CAPZB\_HUMAN | F-actin-capping protein subunit beta | 62 | 54 | CAPZB | CAPZB\_HUMAN | F-actin-capping protein subunit beta | 9C(125.0477) | NaN | X | NaN | NaN | NaN | NaN | 5.580280e+04 | 5.580280e+04 | NaN | NaN | VVGKDYLLCDYNR |
| WDR91 | WDR91\_HUMAN | WD repeat-containing protein 91 | 351 | 342 | WDR91 | WDR91\_HUMAN | WD repeat-containing protein 91 | 10C(125.0477) | NaN | X | NaN | NaN | NaN | NaN | 5.574130e+04 | NaN | 5.574130e+04 | NaN | ELFSTTTSQCAEK |
| USP15 | UBP15\_HUMAN | Ubiquitin carboxyl-terminal hydrolase 15 | 506 | 499 | USP15 | UBP15\_HUMAN | Ubiquitin carboxyl-terminal hydrolase 15 | 8C(125.0477) | NaN | X | NaN | NaN | NaN | NaN | 5.572221e+04 | 5.572221e+04 | NaN | NaN | IGNILDLCTALSALSGIPADK |
| RNF123 | RN123\_HUMAN | E3 ubiquitin-protein ligase RNF123 | 1187 | 1185 | RNF123 | RN123\_HUMAN | E3 ubiquitin-protein ligase RNF123 | 3C(125.0477) | NaN | X | NaN | NaN | NaN | NaN | 5.562833e+04 | 5.562833e+04 | NaN | NaN | SICYLLGQPEPPAPGTALPAPDRK |
| ACSL4 | ACSL4\_HUMAN | Long-chain-fatty-acid--CoA ligase 4 | 420 | 414 | ACSL4 | ACSL4\_HUMAN | Long-chain-fatty-acid--CoA ligase 4 | 7C(125.0477) | NaN | X | NaN | NaN | NaN | NaN | 5.550794e+04 | 5.550794e+04 | NaN | NaN | GYDAPLCNLLLFKK |
| ACTBL2 | ACTBL\_HUMAN | Beta-actin-like protein 2 | 53 | 52 | ACTBL2 | ACTBL\_HUMAN | Beta-actin-like protein 2 | 2C(15.9949) | NaN | NaN | X | NaN | NaN | NaN | 5.547531e+04 | 5.547531e+04 | NaN | NaN | DCYVGDEAQSKR |
| USP19 | UBP19\_HUMAN | Ubiquitin carboxyl-terminal hydrolase 19 | 1042 | 1039 | USP19 | UBP19\_HUMAN | Ubiquitin carboxyl-terminal hydrolase 19 | 4C(125.0477) | NaN | X | NaN | NaN | NaN | NaN | 5.545197e+04 | 4.638285e+04 | 9.069121e+03 | NaN | ELECAEDPGSAGEAAR |
| VAC14 | VAC14\_HUMAN | Protein VAC14 homolog | 114 | 111 | VAC14 | VAC14\_HUMAN | Protein VAC14 homolog | 4C(125.0477) | NaN | X | NaN | NaN | NaN | NaN | 5.544134e+04 | 5.544134e+04 | NaN | NaN | YYACEALYNIVK |
| FLG2 | FILA2\_HUMAN | Filaggrin-2 | 267 | 263 | FLG2 | FILA2\_HUMAN | Filaggrin-2 | 5C(125.0477) | NaN | X | NaN | NaN | NaN | NaN | 5.539917e+04 | 3.405657e+04 | 2.134261e+04 | NaN | LGSSCSGSGDSGR |
| ESD | ESTD\_HUMAN | S-formylglutathione hydrolase | 28 | 18 | ESD | ESTD\_HUMAN | S-formylglutathione hydrolase | 11C(125.0477) | NaN | X | NaN | NaN | NaN | NaN | 5.532222e+04 | NaN | 5.532222e+04 | NaN | VFEHDSVELNCK |
| COPG1 | COPG1\_HUMAN | Coatomer subunit gamma-1 | 296 | 284 | COPG1 | COPG1\_HUMAN | Coatomer subunit gamma-1 | 13C(125.0477) | NaN | X | NaN | NaN | NaN | NaN | 5.530958e+04 | 5.530958e+04 | NaN | NaN | ELAPAVSVLQLFCSSPK |
| PRPSAP2 | KPRB\_HUMAN | Phosphoribosyl pyrophosphate synthase-associated protein 2 | 31 | 18 | PRPSAP2 | KPRB\_HUMAN | Phosphoribosyl pyrophosphate synthase-associated protein 2 | 14C(125.0477) | NaN | X | NaN | NaN | NaN | NaN | 5.514168e+04 | 5.514168e+04 | NaN | NaN | GGLVLFSANSNSSCMELSK |
| LANCL1 | LANC1\_HUMAN | Glutathione S-transferase LANCL1 | 133 | 129 | LANCL1 | LANC1\_HUMAN | Glutathione S-transferase LANCL1 | 5C(125.0477) | NaN | X | NaN | NaN | NaN | NaN | 5.513799e+04 | 3.785306e+03 | 5.135269e+04 | NaN | QAEDCITR |
| TRAF2 | TRAF2\_HUMAN | TNF receptor-associated factor 2 | 287 | 279 | TRAF2 | TRAF2\_HUMAN | TNF receptor-associated factor 2 | 9C(125.0477) | NaN | X | NaN | NaN | NaN | NaN | 5.513757e+04 | 5.513757e+04 | NaN | NaN | TATFENIVCVLNR |
| CORO1A | COR1A\_HUMAN | Coronin-1A | 24 | 21 | CORO1A | COR1A\_HUMAN | Coronin-1A | 4C(125.0477) | NaN | X | NaN | NaN | NaN | NaN | 5.505159e+04 | 5.505159e+04 | NaN | NaN | ADQCYEDVR |
| PHKA2 | KPB2\_HUMAN | Phosphorylase b kinase regulatory subunit alpha, liver isoform | 841 | 834 | PHKA2 | KPB2\_HUMAN | Phosphorylase b kinase regulatory subunit alpha, liver isoform | 8C(125.0477) | NaN | X | NaN | NaN | NaN | NaN | 5.501923e+04 | 5.501923e+04 | NaN | NaN | VEVLAEACTDLLSHQK |
| GSTO1 | GSTO1\_HUMAN | Glutathione S-transferase omega-1 | 90 | 67 | GSTO1 | GSTO1\_HUMAN | Glutathione S-transferase omega-1 | 24C(125.0477) | NaN | X | NaN | NaN | NaN | NaN | 5.492875e+04 | 5.492875e+04 | NaN | NaN | NPFGLVPVLENSQGQLIYESAITCEYLDEAYPGKK |
| CNP | CN37\_HUMAN | 2',3'-cyclic-nucleotide 3'-phosphodiesterase | 111 | 103 | CNP | CN37\_HUMAN | 2',3'-cyclic-nucleotide 3'-phosphodiesterase | 9C(125.0477) | NaN | X | NaN | NaN | NaN | NaN | 5.485632e+04 | 5.485632e+04 | NaN | NaN | LDEDLAAYCR |
| CDC42 | CDC42\_HUMAN | Cell division control protein 42 homolog | 6 | 6 | CDC42 | CDC42\_HUMAN | Cell division control protein 42 homolog | 1C(15.9949) | NaN | NaN | X | NaN | NaN | NaN | 5.476914e+04 | 5.476914e+04 | NaN | NaN | CVVVGDGAVGKTCLLISYTTNK |
| CAND1 | CAND1\_HUMAN | Cullin-associated NEDD8-dissociated protein 1 | 92 | 79 | CAND1 | CAND1\_HUMAN | Cullin-associated NEDD8-dissociated protein 1 | 14C(125.0477) | NaN | X | NaN | NaN | NaN | NaN | 5.476434e+04 | 5.476434e+04 | NaN | NaN | VKEYQVETIVDTLCTNMLSDK |
| ASMTL | ASML\_HUMAN | Probable bifunctional dTTP/UTP pyrophosphatase/methyltransferase protein | 274 | 260 | ASMTL | ASML\_HUMAN | Probable bifunctional dTTP/UTP pyrophosphatase/methyltransferase protein | 15C(125.0477) | NaN | X | NaN | NaN | NaN | NaN | 5.464929e+04 | 5.464929e+04 | NaN | NaN | AEAGEAGQATAEAECHR |
| HSPBP1 | HPBP1\_HUMAN | Hsp70-binding protein 1 | 172, 172 | 165 | HSPBP1 | HPBP1\_HUMAN | Hsp70-binding protein 1 | 8C(125.0477) 8C(31.9988) | NaN | X | NaN | X | NaN | NaN | 5.463984e+04 | 5.463984e+04 | NaN | NaN | AAQLIGTCSQNVAAIQEQVLGLGALR |
| CTH | CGL\_HUMAN | Cystathionine gamma-lyase | 172 | 166 | CTH | CGL\_HUMAN | Cystathionine gamma-lyase | 7C(125.0477) | NaN | X | NaN | NaN | NaN | NaN | 5.460856e+04 | 5.460856e+04 | NaN | NaN | VIDIEGCAHIVHK |
| EPB41L2 | E41L2\_HUMAN | Band 4.1-like protein 2 | 424 | 421 | EPB41L2 | E41L2\_HUMAN | Band 4.1-like protein 2 | 4C(125.0477) | NaN | X | NaN | NaN | NaN | NaN | 5.445573e+04 | 5.445573e+04 | NaN | NaN | LGVCANGLLIYK |
| UGGT1 | UGGG1\_HUMAN | UDP-glucose:glycoprotein glucosyltransferase 1 | 1386 | 1372 | UGGT1 | UGGG1\_HUMAN | UDP-glucose:glycoprotein glucosyltransferase 1 | 15C(125.0477) | NaN | X | NaN | NaN | NaN | NaN | 5.440804e+04 | 5.440804e+04 | NaN | NaN | DFNLDGAPYGYTPFCDSR |
| ALAD | HEM2\_HUMAN | Delta-aminolevulinic acid dehydratase | 223 | 223 | ALAD | HEM2\_HUMAN | Delta-aminolevulinic acid dehydratase | 1C(125.0477) | NaN | X | NaN | NaN | NaN | NaN | 5.437921e+04 | NaN | 5.437921e+04 | NaN | CYQLPPGAR |
| ASAP1 | ASAP1\_HUMAN | Arf-GAP with SH3 domain, ANK repeat and PH domain-containing protein 1 | 685 | 682 | ASAP1 | ASAP1\_HUMAN | Arf-GAP with SH3 domain, ANK repeat and PH domain-containing protein 1 | 4C(125.0477) | NaN | X | NaN | NaN | NaN | NaN | 5.437067e+04 | NaN | 5.437067e+04 | NaN | ATQCEDLLSQAK |
| CMPK2 | CMPK2\_HUMAN | UMP-CMP kinase 2, mitochondrial | 422 | 417 | CMPK2 | CMPK2\_HUMAN | UMP-CMP kinase 2, mitochondrial | 6C(125.0477) | NaN | X | NaN | NaN | NaN | NaN | 5.436477e+04 | 5.436477e+04 | NaN | NaN | MENPGCHVVDASPSR |
| KRT10 | K1C10\_HUMAN | Keratin, type I cytoskeletal 10 | 427, 427 | 423 | KRT10 | K1C10\_HUMAN | Keratin, type I cytoskeletal 10 | 5C(125.0477) 5C(31.9988) | NaN | X | NaN | X | NaN | NaN | 5.432582e+04 | 5.432582e+04 | NaN | NaN | AETECQNTEYQQLLDIK |
| ZC3HAV1 | ZCCHV\_HUMAN | Zinc finger CCCH-type antiviral protein 1 | 174 | 171 | ZC3HAV1 | ZCCHV\_HUMAN | Zinc finger CCCH-type antiviral protein 1 | 4C(125.0477) | NaN | X | NaN | NaN | NaN | NaN | 5.432395e+04 | 5.432395e+04 | NaN | NaN | LHICDHFTR |
| PML | PML\_HUMAN | Protein PML | 151 | 151 | PML | PML\_HUMAN | Protein PML | 1C(125.0477) | NaN | X | NaN | NaN | NaN | NaN | 5.431489e+04 | 5.431489e+04 | NaN | NaN | CFEAHQWFLK |
| WDFY1 | WDFY1\_HUMAN | WD repeat and FYVE domain-containing protein 1 | 249 | 245 | WDFY1 | WDFY1\_HUMAN | WD repeat and FYVE domain-containing protein 1 | 5C(125.0477) | NaN | X | NaN | NaN | NaN | NaN | 5.428453e+04 | 5.428453e+04 | NaN | NaN | VQSLCYLQLTR |
| IGHM | IGHM\_HUMAN | Immunoglobulin heavy constant mu | 134, 134 | 132 | IGHM | IGHM\_HUMAN | Immunoglobulin heavy constant mu | 3C(125.0477) 3C(57.0215) | X | X | NaN | NaN | NaN | NaN | 5.428086e+04 | NaN | 5.428086e+04 | NaN | LICQATGFSPR |
| LRRFIP1 | LRRF1\_HUMAN | Leucine-rich repeat flightless-interacting protein 1 | 644 | 634 | LRRFIP1 | LRRF1\_HUMAN | Leucine-rich repeat flightless-interacting protein 1 | 11C(125.0477) | NaN | X | NaN | NaN | NaN | NaN | 5.421612e+04 | 5.421612e+04 | NaN | NaN | IAAESSENVDCPENPK |
| SLC43A2 | LAT4\_HUMAN | Large neutral amino acids transporter small subunit 4 | 392 | 389 | SLC43A2 | LAT4\_HUMAN | Large neutral amino acids transporter small subunit 4 | 4C(31.9988) | NaN | NaN | NaN | X | NaN | NaN | 5.416713e+04 | NaN | NaN | 54167.1300 | LKECEDASEEPEEK |
| C2CD5 | C2CD5\_HUMAN | C2 domain-containing protein 5 | 258 | 248 | C2CD5 | C2CD5\_HUMAN | C2 domain-containing protein 5 | 11C(125.0477) | NaN | X | NaN | NaN | NaN | NaN | 5.411147e+04 | 5.411147e+04 | NaN | NaN | LSSPAAFLPACNSPSK |
| WBP4 | WBP4\_HUMAN | WW domain-binding protein 4 | 18 | 18 | WBP4 | WBP4\_HUMAN | WW domain-binding protein 4 | 1C(125.0477) | NaN | X | NaN | NaN | NaN | NaN | 5.403341e+04 | 5.403341e+04 | NaN | NaN | CWIADNRPSVEFHER |
| METAP2 | MAP2\_HUMAN | Methionine aminopeptidase 2 | 263 | 260 | METAP2 | MAP2\_HUMAN | Methionine aminopeptidase 2 | 4C(125.0477) | NaN | X | NaN | NaN | NaN | NaN | 5.401630e+04 | 5.401630e+04 | NaN | NaN | IIDCAFTVTFNPK |
| ANKFY1 | ANFY1\_HUMAN | Rabankyrin-5 | 843 | 836 | ANKFY1 | ANFY1\_HUMAN | Rabankyrin-5 | 8C(125.0477) | NaN | X | NaN | NaN | NaN | NaN | 5.388183e+04 | 5.388183e+04 | NaN | NaN | QGLTPFACAMTFK |
| TGM2 | TGM2\_HUMAN | Protein-glutamine gamma-glutamyltransferase 2 | 269 | 264 | TGM2 | TGM2\_HUMAN | Protein-glutamine gamma-glutamyltransferase 2 | 6C(125.0477) | NaN | X | NaN | NaN | NaN | NaN | 5.379528e+04 | 5.379528e+04 | NaN | NaN | WKNHGCQR |
| SBF1 | MTMR5\_HUMAN | Myotubularin-related protein 5 | 896 | 890 | SBF1 | MTMR5\_HUMAN | Myotubularin-related protein 5 | 7C(125.0477) | NaN | X | NaN | NaN | NaN | NaN | 5.374632e+04 | 5.374632e+04 | NaN | NaN | LLPGEECVLDGLR |
| RANBP10 | RBP10\_HUMAN | Ran-binding protein 10 | 100 | 92 | RANBP10 | RBP10\_HUMAN | Ran-binding protein 10 | 9C(125.0477) | NaN | X | NaN | NaN | NaN | NaN | 5.371276e+04 | 5.371276e+04 | NaN | NaN | ATHPIPAACGIYYFEVK |
| WNK1 | WNK1\_HUMAN | Serine/threonine-protein kinase WNK1 | 459 | 454 | WNK1 | WNK1\_HUMAN | Serine/threonine-protein kinase WNK1 | 6C(125.0477) | NaN | X | NaN | NaN | NaN | NaN | 5.370327e+04 | NaN | 5.370327e+04 | NaN | EIIEGCIR |
| SSH3 | SSH3\_HUMAN | Protein phosphatase Slingshot homolog 3 | 413 | 404 | SSH3 | SSH3\_HUMAN | Protein phosphatase Slingshot homolog 3 | 10C(125.0477) | NaN | X | NaN | NaN | NaN | NaN | 5.367758e+04 | 5.367758e+04 | NaN | NaN | AQGTHVLVHCK |
| ACHE | ACES\_HUMAN | Acetylcholinesterase | 560 | 557 | ACHE | ACES\_HUMAN | Acetylcholinesterase | 4C(125.0477) | NaN | X | NaN | NaN | NaN | NaN | 5.366338e+04 | NaN | 5.366338e+04 | NaN | AQACAFWNR |
| CDK2 | CDK2\_HUMAN | Cyclin-dependent kinase 2 | 177 | 170 | CDK2 | CDK2\_HUMAN | Cyclin-dependent kinase 2 | 8C(125.0477) | NaN | X | NaN | NaN | NaN | NaN | 5.365056e+04 | 3.485300e+04 | 1.879756e+04 | NaN | APEILLGCK |
| WDR44 | WDR44\_HUMAN | WD repeat-containing protein 44 | 581 | 574 | WDR44 | WDR44\_HUMAN | WD repeat-containing protein 44 | 8C(125.0477) | NaN | X | NaN | NaN | NaN | NaN | 5.364496e+04 | 5.364496e+04 | NaN | NaN | SDTDTGVCSGTDEDPDDKNAPFR |
| ZDHHC20 | ZDH20\_HUMAN | Palmitoyltransferase ZDHHC20 | 131 | 131 | ZDHHC20 | ZDH20\_HUMAN | Palmitoyltransferase ZDHHC20 | 1C(125.0477) | NaN | X | NaN | NaN | NaN | NaN | 5.359325e+04 | NaN | 5.359325e+04 | NaN | CQLIKPDR |
| NAPA | SNAA\_HUMAN | Alpha-soluble NSF attachment protein | 84 | 79 | NAPA | SNAA\_HUMAN | Alpha-soluble NSF attachment protein | 6C(125.0477) | NaN | X | NaN | NaN | NaN | NaN | 5.353827e+04 | 5.353827e+04 | NaN | NaN | HDAATCFVDAGNAFK |
| CCT8 | TCPQ\_HUMAN | T-complex protein 1 subunit theta | 136 | 121 | CCT8 | TCPQ\_HUMAN | T-complex protein 1 subunit theta | 16C(125.0477) | NaN | X | NaN | NaN | NaN | NaN | 5.352902e+04 | 5.352902e+04 | NaN | NaN | IGLSVSEVIEGYEIACRK |
| PSMA3 | PSA3\_HUMAN | Proteasome subunit alpha type-3 | 187 | 184 | PSMA3 | PSA3\_HUMAN | Proteasome subunit alpha type-3 | 4C(125.0477) | NaN | X | NaN | NaN | NaN | NaN | 5.351904e+04 | 5.351904e+04 | NaN | NaN | EMTCRDIVKEVAK |
| ARHGAP23 | RHG23\_HUMAN | Rho GTPase-activating protein 23 | 601 | 596 | ARHGAP23 | RHG23\_HUMAN | Rho GTPase-activating protein 23 | 6C(125.0477) | NaN | X | NaN | NaN | NaN | NaN | 5.348853e+04 | 5.348853e+04 | 0.000000e+00 | NaN | HYSQDCSSIK |
| HBB | HBB\_HUMAN | Hemoglobin subunit beta | 113, 113 | 106 | HBB | HBB\_HUMAN | Hemoglobin subunit beta | 8C(125.0477) 8C(31.9988) | NaN | X | NaN | X | NaN | NaN | 5.343816e+04 | NaN | 5.343816e+04 | NaN | LLGNVLVCVLAHHFGKEFTPPVQAAYQK |
| EEF1A1 | EF1A1\_HUMAN | Elongation factor 1-alpha 1 | 234 | 220 | EEF1A1 | EF1A1\_HUMAN | Elongation factor 1-alpha 1 | 15C(57.0215) | X | NaN | NaN | NaN | NaN | NaN | 5.343428e+04 | 3.208460e+04 | 8.675872e+03 | 12673.7990 | DGNASGTTLLEALDCILPPTRPTDKPLR |
| RANGAP1 | RAGP1\_HUMAN | Ran GTPase-activating protein 1 | 303 | 297 | RANGAP1 | RAGP1\_HUMAN | Ran GTPase-activating protein 1 | 7C(125.0477) | NaN | X | NaN | NaN | NaN | NaN | 5.337773e+04 | 5.337773e+04 | NaN | NaN | ELNLSFCEIKR |
| MYH9 | MYH9\_HUMAN | Myosin-9 | 917 | 911 | MYH9 | MYH9\_HUMAN | Myosin-9 | 7C(125.0477) | NaN | X | NaN | NaN | NaN | NaN | 5.331922e+04 | 5.331922e+04 | NaN | NaN | QELEEICHDLEAR |
| ALB | ALBU\_HUMAN | Albumin | 148 | 139 | ALB | ALBU\_HUMAN | Albumin | 10C(57.0215) | X | NaN | NaN | NaN | NaN | NaN | 5.314049e+04 | 5.314049e+04 | NaN | NaN | LVRPEVDVMCTAFHDNEETFLK |
| KIF5B | KINH\_HUMAN | Kinesin-1 heavy chain | 421 | 420 | KIF5B | KINH\_HUMAN | Kinesin-1 heavy chain | 2C(125.0477) | NaN | X | NaN | NaN | NaN | NaN | 5.312328e+04 | 5.312328e+04 | NaN | NaN | KCEEEIAK |
| PHAF1 | PHAF1\_HUMAN | Phagosome assembly factor 1 | 345 | 336 | PHAF1 | PHAF1\_HUMAN | Phagosome assembly factor 1 | 10C(125.0477) | NaN | X | NaN | NaN | NaN | NaN | 5.309311e+04 | 5.309311e+04 | NaN | NaN | ENADGQTETCTTYSK |
| ERGIC3 | ERGI3\_HUMAN | Endoplasmic reticulum-Golgi intermediate compartment protein 3 | 182 | 175 | ERGIC3 | ERGI3\_HUMAN | Endoplasmic reticulum-Golgi intermediate compartment protein 3 | 8C(125.0477) | NaN | X | NaN | NaN | NaN | NaN | 5.309276e+04 | NaN | 5.309276e+04 | NaN | NPDTIEQCR |
| ALAD | HEM2\_HUMAN | Delta-aminolevulinic acid dehydratase | 75, 75 | 75 | ALAD | HEM2\_HUMAN | Delta-aminolevulinic acid dehydratase | 1C(125.0477) 1C(31.9988) | NaN | X | NaN | X | NaN | NaN | 5.305296e+04 | NaN | 5.305296e+04 | NaN | CVLIFGVPSR |
| SYNCRIP | HNRPQ\_HUMAN | Heterogeneous nuclear ribonucleoprotein Q | 211 | 204 | SYNCRIP | HNRPQ\_HUMAN | Heterogeneous nuclear ribonucleoprotein Q | 8C(125.0477) | NaN | X | NaN | NaN | NaN | NaN | 5.296286e+04 | 5.296286e+04 | NaN | NaN | GYAFVTFCTK |
| AMPD3 | AMPD3\_HUMAN | AMP deaminase 3 | 323 | 313 | AMPD3 | AMPD3\_HUMAN | AMP deaminase 3 | 11C(125.0477) | NaN | X | NaN | NaN | NaN | NaN | 5.294370e+04 | 5.294370e+04 | NaN | NaN | KVDTHIHAAACMNQK |
| DRG2 | DRG2\_HUMAN | Developmentally-regulated GTP-binding protein 2 | 241 | 237 | DRG2 | DRG2\_HUMAN | Developmentally-regulated GTP-binding protein 2 | 5C(125.0477) | NaN | X | NaN | NaN | NaN | NaN | 5.290298e+04 | 5.290298e+04 | NaN | NaN | VYMPCLYVYNK |
| VPS4B | VPS4B\_HUMAN | Vacuolar protein sorting-associated protein 4B | 240 | 225 | VPS4B | VPS4B\_HUMAN | Vacuolar protein sorting-associated protein 4B | 16C(125.0477) | NaN | X | NaN | NaN | NaN | NaN | 5.276841e+04 | 5.276841e+04 | NaN | NaN | ENKPSIIFIDEIDSLCGSR |
| CPOX | HEM6\_HUMAN | Oxygen-dependent coproporphyrinogen-III oxidase, mitochondrial | 373 | 372 | CPOX | HEM6\_HUMAN | Oxygen-dependent coproporphyrinogen-III oxidase, mitochondrial | 2C(125.0477) | NaN | X | NaN | NaN | NaN | NaN | 5.272260e+04 | 5.272260e+04 | NaN | NaN | HCDDSFTPQEK |
| PAFAH1B1 | LIS1\_HUMAN | Platelet-activating factor acetylhydrolase IB subunit beta | 168 | 164 | PAFAH1B1 | LIS1\_HUMAN | Platelet-activating factor acetylhydrolase IB subunit beta | 5C(125.0477) | NaN | X | NaN | NaN | NaN | NaN | 5.269693e+04 | 5.269693e+04 | NaN | NaN | LLASCSADMTIK |
| EIF2B5 | EI2BE\_HUMAN | Translation initiation factor eIF-2B subunit epsilon | 196 | 196 | EIF2B5 | EI2BE\_HUMAN | Translation initiation factor eIF-2B subunit epsilon | 1C(125.0477) | NaN | X | NaN | NaN | NaN | NaN | 5.269264e+04 | 5.269264e+04 | NaN | NaN | CHEDNVVVAVDSTTNR |
| THEM6 | THEM6\_HUMAN | Protein THEM6 | 168 | 163 | THEM6 | THEM6\_HUMAN | Protein THEM6 | 6C(125.0477) | NaN | X | NaN | NaN | NaN | NaN | 5.266092e+04 | 5.266092e+04 | NaN | NaN | VVQHLCQR |
| SPTLC2 | SPTC2\_HUMAN | Serine palmitoyltransferase 2 | 188 | 184 | SPTLC2 | SPTC2\_HUMAN | Serine palmitoyltransferase 2 | 5C(125.0477) | NaN | X | NaN | NaN | NaN | NaN | 5.261636e+04 | 5.261636e+04 | NaN | NaN | NTGSCQEAAAKVLEEYGAGVCSTR |
| SKIV2L | SKIV2\_HUMAN | Helicase SKI2W | 1014 | 1004 | SKIV2L | SKIV2\_HUMAN | Helicase SKI2W | 11C(125.0477) | NaN | X | NaN | NaN | NaN | NaN | 5.251683e+04 | 5.251683e+04 | NaN | NaN | KLEELIQGAQCVHSPR |
| MTMR12 | MTMRC\_HUMAN | Myotubularin-related protein 12 | 53 | 40 | MTMR12 | MTMRC\_HUMAN | Myotubularin-related protein 12 | 14C(125.0477) | NaN | X | NaN | NaN | NaN | NaN | 5.251414e+04 | 5.251414e+04 | NaN | NaN | EVTLHLLPGEQLLCEASTVLK |
| CCS | CCS\_HUMAN | Copper chaperone for superoxide dismutase | 227 | 225 | CCS | CCS\_HUMAN | Copper chaperone for superoxide dismutase | 3C(125.0477) | NaN | X | NaN | NaN | NaN | NaN | 5.248025e+04 | NaN | 5.248025e+04 | NaN | LACGIIAR |
| ACAP2 | ACAP2\_HUMAN | Arf-GAP with coiled-coil, ANK repeat and PH domain-containing protein 2 | 321 | 320 | ACAP2 | ACAP2\_HUMAN | Arf-GAP with coiled-coil, ANK repeat and PH domain-containing protein 2 | 2C(125.0477) | NaN | X | NaN | NaN | NaN | NaN | 5.233497e+04 | 5.233497e+04 | NaN | NaN | HCEDIERR |
| CAND1 | CAND1\_HUMAN | Cullin-associated NEDD8-dissociated protein 1 | 571 | 567 | CAND1 | CAND1\_HUMAN | Cullin-associated NEDD8-dissociated protein 1 | 5C(125.0477) | NaN | X | NaN | NaN | NaN | NaN | 5.231675e+04 | NaN | 5.231675e+04 | NaN | DLFTCTIK |
| ADD1 | ADDA\_HUMAN | Alpha-adducin | 430 | 406 | ADD1 | ADDA\_HUMAN | Alpha-adducin | 25C(125.0477) | NaN | X | NaN | NaN | NaN | NaN | 5.216898e+04 | 5.216898e+04 | NaN | NaN | KYSDVEVPASVTGYSFASDGDSGTCSPLR |
| PRPS1 | PRPS1\_HUMAN | Ribose-phosphate pyrophosphokinase 1 | 284 | 283 | PRPS1 | PRPS1\_HUMAN | Ribose-phosphate pyrophosphokinase 1 | 2C(125.0477) | NaN | X | NaN | NaN | NaN | NaN | 5.215282e+04 | 5.215282e+04 | NaN | NaN | HCSKIQVIDISMILAEAIRR |
| ARCN1 | COPD\_HUMAN | Coatomer subunit delta | 441 | 436 | ARCN1 | COPD\_HUMAN | Coatomer subunit delta | 6C(125.0477) | NaN | X | NaN | NaN | NaN | NaN | 5.212311e+04 | 5.212311e+04 | NaN | NaN | NTLEWCLPVIDAK |
| DPYD | DPYD\_HUMAN | Dihydropyrimidine dehydrogenase [NADP(+)] | 385 | 385 | DPYD | DPYD\_HUMAN | Dihydropyrimidine dehydrogenase [NADP(+)] | 1C(125.0477) | NaN | X | NaN | NaN | NaN | NaN | 5.212086e+04 | 5.212086e+04 | NaN | NaN | CEFLPFLSPR |
| CAPN5 | CAN5\_HUMAN | Calpain-5 | 495 | 487 | CAPN5 | CAN5\_HUMAN | Calpain-5 | 9C(125.0477) | NaN | X | NaN | NaN | NaN | NaN | 5.203727e+04 | NaN | 5.203727e+04 | NaN | VFTDVPSNCR |
| CST2 | CYTT\_HUMAN | Cystatin-SA | 104 | 97 | CST2 | CYTT\_HUMAN | Cystatin-SA | 8C(125.0477) | NaN | X | NaN | NaN | NaN | NaN | 5.198179e+04 | NaN | 5.198179e+04 | NaN | SQPNLDTCAFHEQPELQK |
| PSME2 | PSME2\_HUMAN | Proteasome activator complex subunit 2 | 118 | 116 | PSME2 | PSME2\_HUMAN | Proteasome activator complex subunit 2 | 3C(125.0477) | NaN | X | NaN | NaN | NaN | NaN | 5.197114e+04 | 5.197114e+04 | NaN | NaN | EKCILVITWIQHLIPK |
| DAZAP1 | DAZP1\_HUMAN | DAZ-associated protein 1 | 124, 124 | 115 | DAZAP1 | DAZP1\_HUMAN | DAZ-associated protein 1 | 10C(125.0477) 10C(15.9949) | NaN | X | X | NaN | NaN | NaN | 5.196115e+04 | 5.196115e+04 | NaN | NaN | IFVGGIPHNCGETELR |
| PCBP2 | PCBP2\_HUMAN | Poly(rC)-binding protein 2 | 109 | 102 | PCBP2 | PCBP2\_HUMAN | Poly(rC)-binding protein 2 | 8C(125.0477) | NaN | X | NaN | NaN | NaN | NaN | 5.188604e+04 | 5.188604e+04 | NaN | NaN | LVVPASQCGSLIGK |
| TGM3 | TGM3\_HUMAN | Protein-glutamine gamma-glutamyltransferase E | 367 | 361 | TGM3 | TGM3\_HUMAN | Protein-glutamine gamma-glutamyltransferase E | 7C(125.0477) | NaN | X | NaN | NaN | NaN | NaN | 5.173875e+04 | 5.173875e+04 | NaN | NaN | SQGVFQCGPASVIGVR |
| VCP | TERA\_HUMAN | Transitional endoplasmic reticulum ATPase | 209 | 192 | VCP | TERA\_HUMAN | Transitional endoplasmic reticulum ATPase | 18C(125.0477) | NaN | X | NaN | NaN | NaN | NaN | 5.169773e+04 | 5.169773e+04 | NaN | NaN | EDEEESLNEVGYDDIGGCRK |
| PPP2R5D | 2A5D\_HUMAN | Serine/threonine-protein phosphatase 2A 56 kDa regulatory subunit delta isoform | 484 | 480 | PPP2R5D | 2A5D\_HUMAN | Serine/threonine-protein phosphatase 2A 56 kDa regulatory subunit delta isoform | 5C(125.0477) | NaN | X | NaN | NaN | NaN | NaN | 5.164177e+04 | 2.844521e+04 | 2.319656e+04 | NaN | LFDDCTQQYK |
| COMP | COMP\_HUMAN | Cartilage oligomeric matrix protein | 726 | 722 | COMP | COMP\_HUMAN | Cartilage oligomeric matrix protein | 5C(57.0215) | X | NaN | NaN | NaN | NaN | NaN | 5.161942e+04 | 2.998478e+04 | 1.792460e+04 | 3710.0420 | LGVFCFSQENIIWANLR |
| ANKRD54 | ANR54\_HUMAN | Ankyrin repeat domain-containing protein 54 | 152 | 145 | ANKRD54 | ANR54\_HUMAN | Ankyrin repeat domain-containing protein 54 | 8C(125.0477) | NaN | X | NaN | NaN | NaN | NaN | 5.151992e+04 | 5.151992e+04 | NaN | NaN | TALHFASCNGNDQIVQLLLDHGADPNQR |
| DSP | DESP\_HUMAN | Desmoplakin | 2656 | 2650 | DSP | DESP\_HUMAN | Desmoplakin | 7C(125.0477) | NaN | X | NaN | NaN | NaN | NaN | 5.146877e+04 | 5.146877e+04 | NaN | NaN | LLEAQACTGGIIHPTTGQK |
| ARHGEF37 | ARH37\_HUMAN | Rho guanine nucleotide exchange factor 37 | 348 | 339 | ARHGEF37 | ARH37\_HUMAN | Rho guanine nucleotide exchange factor 37 | 10C(125.0477) | NaN | X | NaN | NaN | NaN | NaN | 5.145064e+04 | 5.145064e+04 | NaN | NaN | LEGLVWQPLCSLAK |
| CPT1A | CPT1A\_HUMAN | Carnitine O-palmitoyltransferase 1, liver isoform | 742 | 740 | CPT1A | CPT1A\_HUMAN | Carnitine O-palmitoyltransferase 1, liver isoform | 3C(125.0477) | NaN | X | NaN | NaN | NaN | NaN | 5.145061e+04 | 2.131184e+04 | 3.013877e+04 | NaN | FSCPETDSHR |
| SLC14A1 | UT1\_HUMAN | Urea transporter 1 | 25 | 18 | SLC14A1 | UT1\_HUMAN | Urea transporter 1 | 8C(125.0477) | NaN | X | NaN | NaN | NaN | NaN | 5.141398e+04 | NaN | 5.141398e+04 | NaN | GENQVSPCQGR |
| KRAS | RASK\_HUMAN | GTPase KRas | 80 | 74 | KRAS | RASK\_HUMAN | GTPase KRas | 7C(125.0477) | NaN | X | NaN | NaN | NaN | NaN | 5.140986e+04 | 5.140986e+04 | NaN | NaN | TGEGFLCVFAINNTK |
| ANKRD28 | ANR28\_HUMAN | Serine/threonine-protein phosphatase 6 regulatory ankyrin repeat subunit A | 487 | 471 | ANKRD28 | ANR28\_HUMAN | Serine/threonine-protein phosphatase 6 regulatory ankyrin repeat subunit A | 17C(15.9949) | NaN | NaN | X | NaN | NaN | NaN | 5.135284e+04 | 5.135284e+04 | NaN | NaN | GCTPLHYAATSDTDGKCLEYLLR |
| TMCC2 | TMCC2\_HUMAN | Transmembrane and coiled-coil domains protein 2 | 612 | 604 | TMCC2 | TMCC2\_HUMAN | Transmembrane and coiled-coil domains protein 2 | 9C(125.0477) | NaN | X | NaN | NaN | NaN | NaN | 5.134575e+04 | 5.134575e+04 | NaN | NaN | DIQEAVESCLTR |
| ARIH2 | ARI2\_HUMAN | E3 ubiquitin-protein ligase ARIH2 | 270 | 262 | ARIH2 | ARI2\_HUMAN | E3 ubiquitin-protein ligase ARIH2 | 9C(125.0477) | NaN | X | NaN | NaN | NaN | NaN | 5.131279e+04 | 5.131279e+04 | NaN | NaN | QMYHAPTDCATIR |
| ACOT7 | BACH\_HUMAN | Cytosolic acyl coenzyme A thioester hydrolase | 288 | 286 | ACOT7 | BACH\_HUMAN | Cytosolic acyl coenzyme A thioester hydrolase | 3C(125.0477) | NaN | X | NaN | NaN | NaN | NaN | 5.123106e+04 | 5.123106e+04 | NaN | NaN | KGCVITISGR |
| BROX | BROX\_HUMAN | BRO1 domain-containing protein BROX | 288 | 284 | BROX | BROX\_HUMAN | BRO1 domain-containing protein BROX | 5C(15.9949) | NaN | NaN | X | NaN | NaN | NaN | 5.120749e+04 | 5.120749e+04 | NaN | NaN | AEALCKEYGETKGPGPTVKPSGHLFFR |
| ABCC1 | MRP1\_HUMAN | Multidrug resistance-associated protein 1 | 1479 | 1474 | ABCC1 | MRP1\_HUMAN | Multidrug resistance-associated protein 1 | 6C(125.0477) | NaN | X | NaN | NaN | NaN | NaN | 5.118855e+04 | 5.118855e+04 | NaN | NaN | TQFEDCTVLTIAHR |
| SMAD2 | SMAD2\_HUMAN | Mothers against decapentaplegic homolog 2 | 161 | 157 | SMAD2 | SMAD2\_HUMAN | Mothers against decapentaplegic homolog 2 | 5C(125.0477) | NaN | X | NaN | NaN | NaN | NaN | 5.116764e+04 | 5.116764e+04 | NaN | NaN | KDEVCVNPYHYQR |
| GNB2 | GBB2\_HUMAN | Guanine nucleotide-binding protein G(I)/G(S)/G(T) subunit beta-2 | 317 | 315 | GNB2 | GBB2\_HUMAN | Guanine nucleotide-binding protein G(I)/G(S)/G(T) subunit beta-2 | 3C(125.0477) | NaN | X | NaN | NaN | NaN | NaN | 5.114153e+04 | 5.114153e+04 | NaN | NaN | VSCLGVTDDGMAVATGSWDSFLK |
| FKBP4 | FKBP4\_HUMAN | Peptidyl-prolyl cis-trans isomerase FKBP4 | 396 | 391 | FKBP4 | FKBP4\_HUMAN | Peptidyl-prolyl cis-trans isomerase FKBP4 | 6C(125.0477) | NaN | X | NaN | NaN | NaN | NaN | 5.110499e+04 | NaN | 5.110499e+04 | NaN | TQLAVCQQR |
| HK1 | HXK1\_HUMAN | Hexokinase-1 | 886 | 886 | HK1 | HXK1\_HUMAN | Hexokinase-1 | 1C(125.0477) | NaN | X | NaN | NaN | NaN | NaN | 5.106799e+04 | 5.106799e+04 | NaN | NaN | CNVSFLLSEDGSGK |
| TMEM50A | TM50A\_HUMAN | Transmembrane protein 50A | 87 | 80 | TMEM50A | TM50A\_HUMAN | Transmembrane protein 50A | 8C(125.0477) | NaN | X | NaN | NaN | NaN | NaN | 5.097555e+04 | NaN | 5.097555e+04 | NaN | GDSYSEGCLGQTGAR |
| LCP1 | PLSL\_HUMAN | Plastin-2 | 164 | 142 | LCP1 | PLSL\_HUMAN | Plastin-2 | 23C(125.0477) | NaN | X | NaN | NaN | NaN | NaN | 5.092225e+04 | 5.092225e+04 | NaN | NaN | HVIPMNPNTNDLFNAVGDGIVLCK |
| PFAS | PUR4\_HUMAN | Phosphoribosylformylglycinamidine synthase | 606 | 597 | PFAS | PUR4\_HUMAN | Phosphoribosylformylglycinamidine synthase | 10C(125.0477) | NaN | X | NaN | NaN | NaN | NaN | 5.084848e+04 | 5.084848e+04 | NaN | NaN | RIVLVDDRECPVR |
| RPTOR | RPTOR\_HUMAN | Regulatory-associated protein of mTOR | 283 | 283 | RPTOR | RPTOR\_HUMAN | Regulatory-associated protein of mTOR | 1C(125.0477) | NaN | X | NaN | NaN | NaN | NaN | 5.079530e+04 | 5.079530e+04 | NaN | NaN | CVSLVPGVTLDLIEK |
| F11R | JAM1\_HUMAN | Junctional adhesion molecule A | 153 | 149 | F11R | JAM1\_HUMAN | Junctional adhesion molecule A | 5C(125.0477) | NaN | X | NaN | NaN | NaN | NaN | 5.075510e+04 | 5.075510e+04 | NaN | NaN | AVLTCSEQDGSPPSEYTWFK |
| HMBS | HEM3\_HUMAN | Porphobilinogen deaminase | 211 | 202 | HMBS | HEM3\_HUMAN | Porphobilinogen deaminase | 10C(125.0477) | NaN | X | NaN | NaN | NaN | NaN | 5.072250e+04 | 5.072250e+04 | NaN | NaN | VGQILHPEECMYAVGQGALGVEVR |
| CWC27 | CWC27\_HUMAN | Spliceosome-associated protein CWC27 homolog | 164 | 163 | CWC27 | CWC27\_HUMAN | Spliceosome-associated protein CWC27 homolog | 2C(125.0477) | NaN | X | NaN | NaN | NaN | NaN | 5.068051e+04 | 5.068051e+04 | NaN | NaN | SCEVLFNPFDDIIPR |
| TPP2 | TPP2\_HUMAN | Tripeptidyl-peptidase 2 | 342 | 341 | TPP2 | TPP2\_HUMAN | Tripeptidyl-peptidase 2 | 2C(125.0477) | NaN | X | NaN | NaN | NaN | NaN | 5.061378e+04 | 5.061378e+04 | NaN | NaN | ICEVINEAVWK |
| CAND1 | CAND1\_HUMAN | Cullin-associated NEDD8-dissociated protein 1 | 954 | 949 | CAND1 | CAND1\_HUMAN | Cullin-associated NEDD8-dissociated protein 1 | 6C(125.0477) | NaN | X | NaN | NaN | NaN | NaN | 5.056288e+04 | NaN | 5.056288e+04 | NaN | NVVAECLGK |
| CPNE1 | CPNE1\_HUMAN | Copine-1 | 53 | 52 | CPNE1 | CPNE1\_HUMAN | Copine-1 | 2C(125.0477) | NaN | X | NaN | NaN | NaN | NaN | 5.054958e+04 | 5.054958e+04 | NaN | NaN | NCSSPEFSK |
| NIBAN2 | NIBA2\_HUMAN | Protein Niban 2 | 466 | 459 | NIBAN2 | NIBA2\_HUMAN | Protein Niban 2 | 8C(125.0477) | NaN | X | NaN | NaN | NaN | NaN | 5.049169e+04 | 5.049169e+04 | NaN | NaN | GPTKEELCK |
| NISCH | NISCH\_HUMAN | Nischarin | 436 | 432 | NISCH | NISCH\_HUMAN | Nischarin | 5C(125.0477) | NaN | X | NaN | NaN | NaN | NaN | 5.016148e+04 | 5.016148e+04 | NaN | NaN | ASEVCLDDTVTTEK |
| CDC23 | CDC23\_HUMAN | Cell division cycle protein 23 homolog | 109 | 102 | CDC23 | CDC23\_HUMAN | Cell division cycle protein 23 homolog | 8C(125.0477) | NaN | X | NaN | NaN | NaN | NaN | 5.013034e+04 | 5.013034e+04 | NaN | NaN | AAHFLHGCNSK |
| NCSTN | NICA\_HUMAN | Nicastrin | 586 | 584 | NCSTN | NICA\_HUMAN | Nicastrin | 3C(125.0477) | NaN | X | NaN | NaN | NaN | NaN | 5.011078e+04 | NaN | 5.011078e+04 | NaN | EQCQDPSK |
| KPNA4 | IMA3\_HUMAN | Importin subunit alpha-3 | 417 | 401 | KPNA4 | IMA3\_HUMAN | Importin subunit alpha-3 | 17C(125.0477) | NaN | X | NaN | NaN | NaN | NaN | 5.009695e+04 | 5.009695e+04 | NaN | NaN | KDQVAYLIQQNVIPPFCNLLTVK |
| EIF3J | EIF3J\_HUMAN | Eukaryotic translation initiation factor 3 subunit J | 188 | 186 | EIF3J | EIF3J\_HUMAN | Eukaryotic translation initiation factor 3 subunit J | 3C(125.0477) | NaN | X | NaN | NaN | NaN | NaN | 5.005331e+04 | 5.005331e+04 | NaN | NaN | DVCISLEIDDLKK |
| HBB | HBB\_HUMAN | Hemoglobin subunit beta | 113 | 97 | HBB | HBB\_HUMAN | Hemoglobin subunit beta | 17C(125.0477) | NaN | X | NaN | NaN | NaN | NaN | 5.004574e+04 | NaN | 5.004574e+04 | NaN | LHVDPENFRLLGNVLVCVLAHHFGKEFTPPVQAAYQK |
| GLOD4 | GLOD4\_HUMAN | Glyoxalase domain-containing protein 4 | 221 | 217 | GLOD4 | GLOD4\_HUMAN | Glyoxalase domain-containing protein 4 | 5C(125.0477) | NaN | X | NaN | NaN | NaN | NaN | 4.999816e+04 | 4.999816e+04 | NaN | NaN | IAFSCPQKELPDLEDLMK |
| NFX1 | NFX1\_HUMAN | Transcriptional repressor NF-X1 | 557 | 549 | NFX1 | NFX1\_HUMAN | Transcriptional repressor NF-X1 | 9C(125.0477) | NaN | X | NaN | NaN | NaN | NaN | 4.986637e+04 | 4.986637e+04 | NaN | NaN | SDGFGDFSCLK |
| VCPIP1 | VCIP1\_HUMAN | Deubiquitinating protein VCPIP1 | 940 | 939 | VCPIP1 | VCIP1\_HUMAN | Deubiquitinating protein VCPIP1 | 2C(125.0477) | NaN | X | NaN | NaN | NaN | NaN | 4.983436e+04 | 4.983436e+04 | NaN | NaN | HCTFPHLPGK |
| RIPOR1 | RIPR1\_HUMAN | Rho family-interacting cell polarization regulator 1 | 1191 | 1186 | RIPOR1 | RIPR1\_HUMAN | Rho family-interacting cell polarization regulator 1 | 6C(125.0477) | NaN | X | NaN | NaN | NaN | NaN | 4.982132e+04 | 4.982132e+04 | NaN | NaN | QSLQQCGEEGQSAHR |
| PDXP | PLPP\_HUMAN | Chronophin | 91 | 82 | PDXP | PLPP\_HUMAN | Chronophin | 10C(125.0477) | NaN | X | NaN | NaN | NaN | NaN | 4.974811e+04 | NaN | 4.974811e+04 | NaN | AEQLFSSALCAAR |
| GSTM2 | GSTM2\_HUMAN | Glutathione S-transferase Mu 2 | 87 | 83 | GSTM2 | GSTM2\_HUMAN | Glutathione S-transferase Mu 2 | 5C(125.0477) | NaN | X | NaN | NaN | NaN | NaN | 4.971597e+04 | 4.971597e+04 | NaN | NaN | KHNLCGESEK |
| EEF2 | EF2\_HUMAN | Elongation factor 2 | 567 | 560 | EEF2 | EF2\_HUMAN | Elongation factor 2 | 8C(125.0477) | NaN | X | NaN | NaN | NaN | NaN | 4.964446e+04 | 4.964446e+04 | NaN | NaN | DLEEDHACIPIKK |
| RAB7A | RAB7A\_HUMAN | Ras-related protein Rab-7a | 143 | 138 | RAB7A | RAB7A\_HUMAN | Ras-related protein Rab-7a | 6C(125.0477) | NaN | X | NaN | NaN | NaN | NaN | 4.962317e+04 | 4.962317e+04 | NaN | NaN | RAQAWCYSK |
| RNH1 | RINI\_HUMAN | Ribonuclease inhibitor | 266 | 258 | RNH1 | RINI\_HUMAN | Ribonuclease inhibitor | 9C(125.0477) | NaN | X | NaN | NaN | NaN | NaN | 4.952897e+04 | 4.952897e+04 | NaN | NaN | LRTLWIWECGITAK |
| LMAN2 | LMAN2\_HUMAN | Vesicular integral-membrane protein VIP36 | 239 | 238 | LMAN2 | LMAN2\_HUMAN | Vesicular integral-membrane protein VIP36 | 2C(125.0477) | NaN | X | NaN | NaN | NaN | NaN | 4.951611e+04 | NaN | 4.951611e+04 | NaN | NCIDITGVR |
| DTYMK | KTHY\_HUMAN | Thymidylate kinase | 31 | 25 | DTYMK | KTHY\_HUMAN | Thymidylate kinase | 7C(125.0477) | NaN | X | NaN | NaN | NaN | NaN | 4.947584e+04 | 4.947584e+04 | NaN | NaN | KLVEALCAAGHR |
| PLEKHF2 | PKHF2\_HUMAN | Pleckstrin homology domain-containing family F member 2 | 129 | 129 | PLEKHF2 | PKHF2\_HUMAN | Pleckstrin homology domain-containing family F member 2 | 1C(125.0477) | NaN | X | NaN | NaN | NaN | NaN | 4.946914e+04 | NaN | 4.946914e+04 | NaN | CVTDLLSK |
| ALB | ALBU\_HUMAN | Albumin | 302 | 301 | ALB | ALBU\_HUMAN | Albumin | 2C(57.0215) | X | NaN | NaN | NaN | NaN | NaN | 4.943236e+04 | NaN | 4.943236e+04 | NaN | ECCEKPLLEK |
| TXN | THIO\_HUMAN | Thioredoxin | 73 | 73 | TXN | THIO\_HUMAN | Thioredoxin | 1C(125.0477) | NaN | X | NaN | NaN | NaN | NaN | 4.942089e+04 | 4.942089e+04 | NaN | NaN | CMPTFQFFKK |
| USP24 | UBP24\_HUMAN | Ubiquitin carboxyl-terminal hydrolase 24 | 1239 | 1233 | USP24 | UBP24\_HUMAN | Ubiquitin carboxyl-terminal hydrolase 24 | 7C(125.0477) | NaN | X | NaN | NaN | NaN | NaN | 4.935602e+04 | 4.935602e+04 | NaN | NaN | QGVYSICLQLAR |
| PARP10 | PAR10\_HUMAN | Protein mono-ADP-ribosyltransferase PARP10 | 898 | 879 | PARP10 | PAR10\_HUMAN | Protein mono-ADP-ribosyltransferase PARP10 | 20C(125.0477) | NaN | X | NaN | NaN | NaN | NaN | 4.933895e+04 | 4.933895e+04 | NaN | NaN | RPVEQVLYHGTTAPAVPDICAHGFNR |
| ACP1 | PPAC\_HUMAN | Low molecular weight phosphotyrosine protein phosphatase | 63 | 42 | ACP1 | PPAC\_HUMAN | Low molecular weight phosphotyrosine protein phosphatase | 22C(31.9988) | NaN | NaN | NaN | X | NaN | NaN | 4.932076e+04 | 4.932076e+04 | NaN | NaN | VDSAATSGYEIGNPPDYRGQSCMK |
| PPP1CB | PP1B\_HUMAN | Serine/threonine-protein phosphatase PP1-beta catalytic subunit | 61 | 60 | PPP1CB | PP1B\_HUMAN | Serine/threonine-protein phosphatase PP1-beta catalytic subunit | 2C(125.0477) | NaN | X | NaN | NaN | NaN | NaN | 4.925947e+04 | 4.925947e+04 | NaN | NaN | ICGDIHGQYTDLLR |
| RIMOC1 | RIMC1\_HUMAN | RAB7A-interacting MON1-CCZ1 complex subunit 1 | 179 | 179 | RIMOC1 | RIMC1\_HUMAN | RAB7A-interacting MON1-CCZ1 complex subunit 1 | 1C(125.0477) | NaN | X | NaN | NaN | NaN | NaN | 4.915314e+04 | 4.915314e+04 | NaN | NaN | CPIQLNEGVSFQDLDTAK |
| LDHB | LDHB\_HUMAN | L-lactate dehydrogenase B chain | 164 | 159 | LDHB | LDHB\_HUMAN | L-lactate dehydrogenase B chain | 6C(125.0477) | NaN | X | NaN | NaN | NaN | NaN | 4.914765e+04 | 4.914765e+04 | NaN | NaN | VIGSGCNLDSARFR |
| STXBP3 | STXB3\_HUMAN | Syntaxin-binding protein 3 | 26 | 20 | STXBP3 | STXB3\_HUMAN | Syntaxin-binding protein 3 | 7C(125.0477) | NaN | X | NaN | NaN | NaN | NaN | 4.911763e+04 | 4.911763e+04 | NaN | NaN | ATVFDDCKK |
| GSS | GSHB\_HUMAN | Glutathione synthetase | 422 | 419 | GSS | GSHB\_HUMAN | Glutathione synthetase | 4C(125.0477) | NaN | X | NaN | NaN | NaN | NaN | 4.910170e+04 | 4.910170e+04 | NaN | NaN | VVQCISELGIFGVYVR |
| TUBA1B | TBA1B\_HUMAN | Tubulin alpha-1B chain | 295 | 281 | TUBA1B | TBA1B\_HUMAN | Tubulin alpha-1B chain | 15C(125.0477) | NaN | X | NaN | NaN | NaN | NaN | 4.909074e+04 | 4.909074e+04 | NaN | NaN | AYHEQLSVAEITNACFEPANQMVK |
| YWHAH | 1433F\_HUMAN | 14-3-3 protein eta | 97 | 92 | YWHAH | 1433F\_HUMAN | 14-3-3 protein eta | 6C(125.0477) | NaN | X | NaN | NaN | NaN | NaN | 4.908558e+04 | 4.908558e+04 | NaN | NaN | ELETVCNDVLSLLDK |
| GNL1 | GNL1\_HUMAN | Guanine nucleotide-binding protein-like 1 | 365 | 358 | GNL1 | GNL1\_HUMAN | Guanine nucleotide-binding protein-like 1 | 8C(125.0477) | NaN | X | NaN | NaN | NaN | NaN | 4.907308e+04 | 4.907308e+04 | NaN | NaN | DGVVTIGCVGFPNVGK |
| USP19 | UBP19\_HUMAN | Ubiquitin carboxyl-terminal hydrolase 19 | 833 | 831 | USP19 | UBP19\_HUMAN | Ubiquitin carboxyl-terminal hydrolase 19 | 3C(125.0477) | NaN | X | NaN | NaN | NaN | NaN | 4.904780e+04 | 4.904780e+04 | NaN | NaN | GLCRPENIGYPFLVSVPASR |
| CCT8 | TCPQ\_HUMAN | T-complex protein 1 subunit theta | 36 | 32 | CCT8 | TCPQ\_HUMAN | T-complex protein 1 subunit theta | 5C(125.0477) | NaN | X | NaN | NaN | NaN | NaN | 4.895359e+04 | 4.895359e+04 | NaN | NaN | NIQACKELAQTTR |
| SLC30A1 | ZNT1\_HUMAN | Zinc transporter 1 | 469 | 462 | SLC30A1 | ZNT1\_HUMAN | Zinc transporter 1 | 8C(125.0477) | NaN | X | NaN | NaN | NaN | NaN | 4.893809e+04 | 4.893809e+04 | NaN | NaN | TPAVSISCLELSNNLEK |
| SKP1 | SKP1\_HUMAN | S-phase kinase-associated protein 1 | 62 | 58 | SKP1 | SKP1\_HUMAN | S-phase kinase-associated protein 1 | 5C(125.0477) | NaN | X | NaN | NaN | NaN | NaN | 4.871100e+04 | 4.871100e+04 | NaN | NaN | VIQWCTHHKDDPPPPEDDENK |
| CAND1 | CAND1\_HUMAN | Cullin-associated NEDD8-dissociated protein 1 | 802 | 802 | CAND1 | CAND1\_HUMAN | Cullin-associated NEDD8-dissociated protein 1 | 1C(125.0477) | NaN | X | NaN | NaN | NaN | NaN | 4.868796e+04 | NaN | 4.868796e+04 | NaN | CVAALTR |
| VTN | VTNC\_HUMAN | Vitronectin | 180 | 177 | VTN | VTNC\_HUMAN | Vitronectin | 4C(125.0477) | NaN | X | NaN | NaN | NaN | NaN | 4.866739e+04 | NaN | 4.866739e+04 | NaN | GQYCYELDEK |
| JUP | PLAK\_HUMAN | Junction plakoglobin | 204 | 204 | JUP | PLAK\_HUMAN | Junction plakoglobin | 1C(125.0477) | NaN | X | NaN | NaN | NaN | NaN | 4.862142e+04 | 4.862142e+04 | NaN | NaN | CTTSILHNLSHHR |
| OTULIN | OTUL\_HUMAN | Ubiquitin thioesterase otulin | 129 | 123 | OTULIN | OTUL\_HUMAN | Ubiquitin thioesterase otulin | 7C(125.0477) | NaN | X | NaN | NaN | NaN | NaN | 4.860196e+04 | 4.860196e+04 | NaN | NaN | VRGDNYCALR |
| CUL2 | CUL2\_HUMAN | Cullin-2 | 103 | 97 | CUL2 | CUL2\_HUMAN | Cullin-2 | 7C(125.0477) | NaN | X | NaN | NaN | NaN | NaN | 4.848238e+04 | 4.848238e+04 | NaN | NaN | GADYMDCLYR |
| PAXX | PAXX\_HUMAN | Protein PAXX | 80 | 78 | PAXX | PAXX\_HUMAN | Protein PAXX | 3C(125.0477) | NaN | X | NaN | NaN | NaN | NaN | 4.845410e+04 | 4.845410e+04 | NaN | NaN | AACEQQAVALTLQEDR |
| FGG | FIBG\_HUMAN | Fibrinogen gamma chain | 365 | 365 | FGG | FIBG\_HUMAN | Fibrinogen gamma chain | 1C(57.0215) | X | NaN | NaN | NaN | NaN | NaN | 4.832564e+04 | 4.832564e+04 | NaN | NaN | CHAGHLNGVYYQGGTYSK |
| REXO2 | ORN\_HUMAN | Oligoribonuclease, mitochondrial | 97 | 85 | REXO2 | ORN\_HUMAN | Oligoribonuclease, mitochondrial | 13C(125.0477) | NaN | X | NaN | NaN | NaN | NaN | 4.816924e+04 | 4.816924e+04 | NaN | NaN | QPDELLDSMSDWCK |
| ASAP1 | ASAP1\_HUMAN | Arf-GAP with SH3 domain, ANK repeat and PH domain-containing protein 1 | 373 | 368 | ASAP1 | ASAP1\_HUMAN | Arf-GAP with SH3 domain, ANK repeat and PH domain-containing protein 1 | 6C(125.0477) | NaN | X | NaN | NaN | NaN | NaN | 4.816361e+04 | 4.816361e+04 | NaN | NaN | LNLLTCQVKPNAEDK |
| PPIB | PPIB\_HUMAN | Peptidyl-prolyl cis-trans isomerase B | 202 | 191 | PPIB | PPIB\_HUMAN | Peptidyl-prolyl cis-trans isomerase B | 12C(125.0477) | NaN | X | NaN | NaN | NaN | NaN | 4.816125e+04 | 4.816125e+04 | NaN | NaN | DKPLKDVIIADCGK |
| NRDC | NRDC\_HUMAN | Nardilysin | 1027 | 1024 | NRDC | NRDC\_HUMAN | Nardilysin | 4C(125.0477) | NaN | X | NaN | NaN | NaN | NaN | 4.815233e+04 | 4.815233e+04 | NaN | NaN | LKECEDTHLGEEVDR |
| CDC37 | CDC37\_HUMAN | Hsp90 co-chaperone Cdc37 | 336 | 336 | CDC37 | CDC37\_HUMAN | Hsp90 co-chaperone Cdc37 | 1C(125.0477) | NaN | X | NaN | NaN | NaN | NaN | 4.814534e+04 | 4.814534e+04 | NaN | NaN | CIDSGLWVPNSK |
| NUP153 | NU153\_HUMAN | Nuclear pore complex protein Nup153 | 21 | 21 | NUP153 | NU153\_HUMAN | Nuclear pore complex protein Nup153 | 1C(125.0477) | NaN | X | NaN | NaN | NaN | NaN | 4.812266e+04 | 4.812266e+04 | NaN | NaN | CHQGPIKPYQQGR |
| DHX9 | DHX9\_HUMAN | ATP-dependent RNA helicase A | 242 | 237 | DHX9 | DHX9\_HUMAN | ATP-dependent RNA helicase A | 6C(125.0477) | NaN | X | NaN | NaN | NaN | NaN | 4.810348e+04 | 4.810348e+04 | NaN | NaN | LAAQSCALSLVR |
| PPME1 | PPME1\_HUMAN | Protein phosphatase methylesterase 1 | 347 | 339 | PPME1 | PPME1\_HUMAN | Protein phosphatase methylesterase 1 | 9C(125.0477) | NaN | X | NaN | NaN | NaN | NaN | 4.810238e+04 | 4.810238e+04 | NaN | NaN | FQMQVLPQCGHAVHEDAPDK |
| IGHG3 | IGHG3\_HUMAN | Immunoglobulin heavy constant gamma 3 | 111 | 102 | IGHG3 | IGHG3\_HUMAN | Immunoglobulin heavy constant gamma 3 | 10C(57.0215) | X | NaN | NaN | NaN | NaN | NaN | 4.809587e+04 | NaN | 4.809587e+04 | NaN | TPLGDTTHTCPR |
| IGHG1 | IGHG1\_HUMAN | Immunoglobulin heavy constant gamma 1 | 250 | 244 | IGHG1 | IGHG1\_HUMAN | Immunoglobulin heavy constant gamma 1 | 7C(31.9988) | NaN | NaN | NaN | X | NaN | NaN | 4.791438e+04 | NaN | NaN | 47914.3750 | NQVSLTCLVK |
| CD44 | CD44\_HUMAN | CD44 antigen | 77 | 69 | CD44 | CD44\_HUMAN | CD44 antigen | 9C(125.0477) | NaN | X | NaN | NaN | NaN | NaN | 4.776672e+04 | NaN | 4.776672e+04 | NaN | ALSIGFETCR |
| VTA1 | VTA1\_HUMAN | Vacuolar protein sorting-associated protein VTA1 homolog | 38 | 30 | VTA1 | VTA1\_HUMAN | Vacuolar protein sorting-associated protein VTA1 homolog | 9C(125.0477) | NaN | X | NaN | NaN | NaN | NaN | 4.775902e+04 | 4.775902e+04 | NaN | NaN | RDPVVAYYCR |
| KRT33B | KT33B\_HUMAN | Keratin, type I cuticular Ha3-II | 365 | 358 | KRT33B | KT33B\_HUMAN | Keratin, type I cuticular Ha3-II | 8C(125.0477) | NaN | X | NaN | NaN | NaN | NaN | 4.772839e+04 | NaN | NaN | 47728.3950 | SLLESEDCK |
| ENO1 | ENOA\_HUMAN | Alpha-enolase | 389 | 373 | ENO1 | ENOA\_HUMAN | Alpha-enolase | 17C(15.9949) | NaN | NaN | X | NaN | NaN | NaN | 4.770264e+04 | 4.770264e+04 | NaN | NaN | SGETEDTFIADLVVGLCTGQIKTGAPCR |
| EIF2S1 | IF2A\_HUMAN | Eukaryotic translation initiation factor 2 subunit 1 | 98 | 90 | EIF2S1 | IF2A\_HUMAN | Eukaryotic translation initiation factor 2 subunit 1 | 9C(125.0477) | NaN | X | NaN | NaN | NaN | NaN | 4.770179e+04 | 4.770179e+04 | NaN | NaN | VSPEEAIKCEDKFTK |
| PRMT1 | ANM1\_HUMAN | Protein arginine N-methyltransferase 1 | 103 | 91 | PRMT1 | ANM1\_HUMAN | Protein arginine N-methyltransferase 1 | 13C(125.0477) | NaN | X | NaN | NaN | NaN | NaN | 4.769355e+04 | 4.769355e+04 | NaN | NaN | VVLDVGSGTGILCMFAAK |
| PEF1 | PEF1\_HUMAN | Peflin | 247 | 243 | PEF1 | PEF1\_HUMAN | Peflin | 5C(125.0477) | NaN | X | NaN | NaN | NaN | NaN | 4.768966e+04 | 4.768966e+04 | NaN | NaN | FIQVCTQLQVLTEAFR |
| FLNA | FLNA\_HUMAN | Filamin-A | 2102 | 2090 | FLNA | FLNA\_HUMAN | Filamin-A | 13C(125.0477) | NaN | X | NaN | NaN | NaN | NaN | 4.764411e+04 | 4.764411e+04 | NaN | NaN | VDINTEDLEDGTCR |
| CRYZ | QOR\_HUMAN | Quinone oxidoreductase | 145 | 139 | CRYZ | QOR\_HUMAN | Quinone oxidoreductase | 7C(125.0477) | NaN | X | NaN | NaN | NaN | NaN | 4.762516e+04 | 4.762516e+04 | NaN | NaN | ALIHSACVK |
| CASP6 | CASP6\_HUMAN | Caspase-6 | 277 | 274 | CASP6 | CASP6\_HUMAN | Caspase-6 | 4C(125.0477) | NaN | X | NaN | NaN | NaN | NaN | 4.751026e+04 | 4.751026e+04 | NaN | NaN | QVPCFASMLTK |
| GTPBP2 | GTPB2\_HUMAN | GTP-binding protein 2 | 249 | 244 | GTPBP2 | GTPB2\_HUMAN | GTP-binding protein 2 | 6C(125.0477) | NaN | X | NaN | NaN | NaN | NaN | 4.746142e+04 | 4.746142e+04 | NaN | NaN | TAEEICESSSK |
| GSTM3 | GSTM3\_HUMAN | Glutathione S-transferase Mu 3 | 119 | 118 | GSTM3 | GSTM3\_HUMAN | Glutathione S-transferase Mu 3 | 2C(125.0477) | NaN | X | NaN | NaN | NaN | NaN | 4.743023e+04 | 4.743023e+04 | NaN | NaN | LCYSSDHEK |
| CASP9 | CASP9\_HUMAN | Caspase-9 | 187 | 181 | CASP9 | CASP9\_HUMAN | Caspase-9 | 7C(125.0477) | NaN | X | NaN | NaN | NaN | NaN | 4.734549e+04 | 4.734549e+04 | NaN | NaN | TGSNIDCEK |
| TF | TRFE\_HUMAN | Serotransferrin | 58 | 47 | TF | TRFE\_HUMAN | Serotransferrin | 12C(57.0215) | X | NaN | NaN | NaN | NaN | NaN | 4.733714e+04 | NaN | 4.733714e+04 | NaN | SVIPSDGPSVACVK |
| GGACT | GGACT\_HUMAN | Gamma-glutamylaminecyclotransferase | 84 | 77 | GGACT | GGACT\_HUMAN | Gamma-glutamylaminecyclotransferase | 8C(125.0477) | NaN | X | NaN | NaN | NaN | NaN | 4.721711e+04 | 4.721711e+04 | NaN | NaN | FLDDFESCPALYQR |
| UBE4B | UBE4B\_HUMAN | Ubiquitin conjugation factor E4 B | 829 | 828 | UBE4B | UBE4B\_HUMAN | Ubiquitin conjugation factor E4 B | 2C(125.0477) | NaN | X | NaN | NaN | NaN | NaN | 4.715489e+04 | 4.715489e+04 | NaN | NaN | ACADAGLLDESFLR |
| PGM1 | PGM1\_HUMAN | Phosphoglucomutase-1 | 160 | 153 | PGM1 | PGM1\_HUMAN | Phosphoglucomutase-1 | 8C(125.0477) | NaN | X | NaN | NaN | NaN | NaN | 4.712802e+04 | 4.712802e+04 | NaN | NaN | TIEEYAVCPDLK |
| BCAM | BCAM\_HUMAN | Basal cell adhesion molecule | 522 | 518 | BCAM | BCAM\_HUMAN | Basal cell adhesion molecule | 5C(125.0477) | NaN | X | NaN | NaN | NaN | NaN | 4.711503e+04 | NaN | 4.002052e+04 | 7094.5130 | DGISCEASNPHGNK |
| ARHGAP4 | RHG04\_HUMAN | Rho GTPase-activating protein 4 | 527 | 512 | ARHGAP4 | RHG04\_HUMAN | Rho GTPase-activating protein 4 | 16C(125.0477) | NaN | X | NaN | NaN | NaN | NaN | 4.706405e+04 | 4.706405e+04 | NaN | NaN | FIQSSGQPVPLVVESCIR |
| TANGO2 | TNG2\_HUMAN | Transport and Golgi organization protein 2 homolog | 228 | 224 | TANGO2 | TNG2\_HUMAN | Transport and Golgi organization protein 2 homolog | 5C(125.0477) | NaN | X | NaN | NaN | NaN | NaN | 4.704256e+04 | NaN | 4.704256e+04 | NaN | YAAVCVR |
| RTCA | RTCA\_HUMAN | RNA 3'-terminal phosphate cyclase | 28 | 22 | RTCA | RTCA\_HUMAN | RNA 3'-terminal phosphate cyclase | 7C(125.0477) | NaN | X | NaN | NaN | NaN | NaN | 4.700395e+04 | 4.700395e+04 | NaN | NaN | VSTALSCLLGLPLR |
| MYH9 | MYH9\_HUMAN | Myosin-9 | 931 | 931 | MYH9 | MYH9\_HUMAN | Myosin-9 | 1C(125.0477) | NaN | X | NaN | NaN | NaN | NaN | 4.692852e+04 | 4.692852e+04 | NaN | NaN | CQHLQAEKK |
| PRKAR1B | KAP1\_HUMAN | cAMP-dependent protein kinase type I-beta regulatory subunit | 362 | 358 | PRKAR1B | KAP1\_HUMAN | cAMP-dependent protein kinase type I-beta regulatory subunit | 5C(125.0477) | NaN | X | NaN | NaN | NaN | NaN | 4.686116e+04 | NaN | 4.686116e+04 | NaN | VLGPCSEILK |
| UBR4 | UBR4\_HUMAN | E3 ubiquitin-protein ligase UBR4 | 2489 | 2478 | UBR4 | UBR4\_HUMAN | E3 ubiquitin-protein ligase UBR4 | 12C(125.0477) | NaN | X | NaN | NaN | NaN | NaN | 4.671112e+04 | 4.671112e+04 | NaN | NaN | LVVSSLEALESCFAVGPIIEK |
| GAPDH | G3P\_HUMAN | Glyceraldehyde-3-phosphate dehydrogenase | 247 | 228 | GAPDH | G3P\_HUMAN | Glyceraldehyde-3-phosphate dehydrogenase | 20C(125.0477) | NaN | X | NaN | NaN | NaN | NaN | 4.665083e+04 | 4.665083e+04 | NaN | NaN | LTGMAFRVPTANVSVVDLTCRLEKPAK |
| C3 | CO3\_HUMAN | Complement C3 | 1358 | 1352 | C3 | CO3\_HUMAN | Complement C3 | 7C(57.0215) | X | NaN | NaN | NaN | NaN | NaN | 4.664987e+04 | 1.737804e+04 | 2.927183e+04 | NaN | AKDQLTCNKFDLK |
| INPP5D | SHIP1\_HUMAN | Phosphatidylinositol 3,4,5-trisphosphate 5-phosphatase 1 | 138 | 130 | INPP5D | SHIP1\_HUMAN | Phosphatidylinositol 3,4,5-trisphosphate 5-phosphatase 1 | 9C(125.0477) | NaN | X | NaN | NaN | NaN | NaN | 4.658259e+04 | NaN | 4.658259e+04 | NaN | NIPLTASSCEAK |
| SCFD2 | SCFD2\_HUMAN | Sec1 family domain-containing protein 2 | 54 | 45 | SCFD2 | SCFD2\_HUMAN | Sec1 family domain-containing protein 2 | 10C(125.0477) | NaN | X | NaN | NaN | NaN | NaN | 4.654839e+04 | 4.654839e+04 | NaN | NaN | LLEAVGGPDCHLR |
| FAM114A2 | F1142\_HUMAN | Protein FAM114A2 | 438 | 434 | FAM114A2 | F1142\_HUMAN | Protein FAM114A2 | 5C(125.0477) | NaN | X | NaN | NaN | NaN | NaN | 4.651126e+04 | NaN | 4.651126e+04 | NaN | EFTTCLTTAGVK |
| ANK1 | ANK1\_HUMAN | Ankyrin-1 | 278 | 262 | ANK1 | ANK1\_HUMAN | Ankyrin-1 | 17C(125.0477) | NaN | X | NaN | NaN | NaN | NaN | 4.648577e+04 | 4.648577e+04 | NaN | NaN | GAQIETKTKDELTPLHCAAR |
| NECAP1 | NECP1\_HUMAN | Adaptin ear-binding coat-associated protein 1 | 162 | 161 | NECAP1 | NECP1\_HUMAN | Adaptin ear-binding coat-associated protein 1 | 2C(125.0477) | NaN | X | NaN | NaN | NaN | NaN | 4.644139e+04 | 4.644139e+04 | NaN | NaN | LCIGNITNK |
| ATP6V1B2 | VATB2\_HUMAN | V-type proton ATPase subunit B, brain isoform | 112 | 110 | ATP6V1B2 | VATB2\_HUMAN | V-type proton ATPase subunit B, brain isoform | 3C(125.0477) | NaN | X | NaN | NaN | NaN | NaN | 4.639256e+04 | 4.639256e+04 | NaN | NaN | TSCEFTGDILR |
| KRT10 | K1C10\_HUMAN | Keratin, type I cytoskeletal 10 | 66 | 60 | KRT10 | K1C10\_HUMAN | Keratin, type I cytoskeletal 10 | 7C(125.0477) | NaN | X | NaN | NaN | NaN | NaN | 4.632070e+04 | 1.965078e+04 | 2.666992e+04 | NaN | GSSGGGCFGGSSGGYGGLGGFGGGSFR |
| TBC1D5 | TBCD5\_HUMAN | TBC1 domain family member 5 | 96 | 93 | TBC1D5 | TBCD5\_HUMAN | TBC1 domain family member 5 | 4C(125.0477) | NaN | X | NaN | NaN | NaN | NaN | 4.631830e+04 | 4.631830e+04 | NaN | NaN | LFLCVLPQDK |
| GLTP | GLTP\_HUMAN | Glycolipid transfer protein | 176 | 168 | GLTP | GLTP\_HUMAN | Glycolipid transfer protein | 9C(125.0477) | NaN | X | NaN | NaN | NaN | NaN | 4.608529e+04 | 4.608529e+04 | NaN | NaN | GQNVTEEECLEK |
| NME2 | NDKB\_HUMAN | Nucleoside diphosphate kinase B | 145 | 144 | NME2 | NDKB\_HUMAN | Nucleoside diphosphate kinase B | 2C(125.0477) | NaN | X | NaN | NaN | NaN | NaN | 4.602485e+04 | 4.602485e+04 | NaN | NaN | SCAHDWVYE |
| EPS15 | EPS15\_HUMAN | Epidermal growth factor receptor substrate 15 | 586 | 585 | EPS15 | EPS15\_HUMAN | Epidermal growth factor receptor substrate 15 | 2C(125.0477) | NaN | X | NaN | NaN | NaN | NaN | 4.601098e+04 | NaN | 4.601098e+04 | NaN | VCSELDNNR |
| PRDX2 | PRDX2\_HUMAN | Peroxiredoxin-2 | 172 | 158 | PRDX2 | PRDX2\_HUMAN | Peroxiredoxin-2 | 15C(125.0477) | NaN | X | NaN | NaN | NaN | NaN | 4.599399e+04 | NaN | 4.599399e+04 | NaN | LVQAFQYTDEHGEVCPAGWKPGSDTIKPNVDDSKEYFSK |
| RRM1 | RIR1\_HUMAN | Ribonucleoside-diphosphate reductase large subunit | 492 | 480 | RRM1 | RIR1\_HUMAN | Ribonucleoside-diphosphate reductase large subunit | 13C(125.0477) | NaN | X | NaN | NaN | NaN | NaN | 4.581054e+04 | 4.581054e+04 | NaN | NaN | IIDINYYPVPEACLSNKR |
| A2M | A2MG\_HUMAN | Alpha-2-macroglobulin | 595 | 587 | A2M | A2MG\_HUMAN | Alpha-2-macroglobulin | 9C(57.0215) | X | NaN | NaN | NaN | NaN | NaN | 4.579680e+04 | NaN | 4.579680e+04 | NaN | VTAAPQSVCALR |
| PPP6R3 | PP6R3\_HUMAN | Serine/threonine-protein phosphatase 6 regulatory subunit 3 | 830 | 822 | PPP6R3 | PP6R3\_HUMAN | Serine/threonine-protein phosphatase 6 regulatory subunit 3 | 9C(15.9949) | NaN | NaN | X | NaN | NaN | NaN | 4.569433e+04 | 4.569433e+04 | NaN | NaN | LSTSQDAACKDAEECPETAEAK |
| PSMD6 | PSMD6\_HUMAN | 26S proteasome non-ATPase regulatory subunit 6 | 58 | 47 | PSMD6 | PSMD6\_HUMAN | 26S proteasome non-ATPase regulatory subunit 6 | 12C(125.0477) | NaN | X | NaN | NaN | NaN | NaN | 4.565239e+04 | 4.565239e+04 | NaN | NaN | DNNMAPYYEALCK |
| IPO5 | IPO5\_HUMAN | Importin-5 | 110 | 108 | IPO5 | IPO5\_HUMAN | Importin-5 | 3C(125.0477) | NaN | X | NaN | NaN | NaN | NaN | 4.551806e+04 | 4.551806e+04 | NaN | NaN | KVCDIAAELAR |
| PSMD2 | PSMD2\_HUMAN | 26S proteasome non-ATPase regulatory subunit 2 | 779 | 774 | PSMD2 | PSMD2\_HUMAN | 26S proteasome non-ATPase regulatory subunit 2 | 6C(125.0477) | NaN | X | NaN | NaN | NaN | NaN | 4.546809e+04 | 4.546809e+04 | NaN | NaN | GTLTLCPYHSDR |
| ATIC | PUR9\_HUMAN | Bifunctional purine biosynthesis protein ATIC | 146 | 142 | ATIC | PUR9\_HUMAN | Bifunctional purine biosynthesis protein ATIC | 5C(125.0477) | NaN | X | NaN | NaN | NaN | NaN | 4.543474e+04 | 4.543474e+04 | NaN | NaN | VTVVCEPEDYVVVSTEMQSSESK |
| IRGQ | IRGQ\_HUMAN | Immunity-related GTPase family Q protein | 340 | 331 | IRGQ | IRGQ\_HUMAN | Immunity-related GTPase family Q protein | 10C(125.0477) | NaN | X | NaN | NaN | NaN | NaN | 4.541783e+04 | 4.541783e+04 | NaN | NaN | TDGEGEDPECLGEGK |
| WIPI2 | WIPI2\_HUMAN | WD repeat domain phosphoinositide-interacting protein 2 | 328 | 325 | WIPI2 | WIPI2\_HUMAN | WD repeat domain phosphoinositide-interacting protein 2 | 4C(15.9949) | NaN | NaN | X | NaN | NaN | NaN | 4.528192e+04 | 4.528192e+04 | NaN | NaN | LPFCGHKNICSLATIQK |
| IPO5 | IPO5\_HUMAN | Importin-5 | 972 | 964 | IPO5 | IPO5\_HUMAN | Importin-5 | 9C(125.0477) | NaN | X | NaN | NaN | NaN | NaN | 4.526750e+04 | 4.526750e+04 | NaN | NaN | ENVNATENCISAVGK |
| KRT1 | K2C1\_HUMAN | Keratin, type II cytoskeletal 1 | 497 | 493 | KRT1 | K2C1\_HUMAN | Keratin, type II cytoskeletal 1 | 5C(15.9949) | NaN | NaN | X | NaN | NaN | NaN | 4.525577e+04 | NaN | 4.525577e+04 | NaN | MSGECAPNVSVSVSTSHTTISGGGSR |
| CTPS1 | PYRG1\_HUMAN | CTP synthase 1 | 176 | 173 | CTPS1 | PYRG1\_HUMAN | CTP synthase 1 | 4C(125.0477) | NaN | X | NaN | NaN | NaN | NaN | 4.506357e+04 | 4.506357e+04 | NaN | NaN | ENFCNIHVSLVPQPSSTGEQK |
| CTSB | CATB\_HUMAN | Cathepsin B | 319 | 315 | CTSB | CATB\_HUMAN | Cathepsin B | 5C(125.0477) | NaN | X | NaN | NaN | NaN | NaN | 4.503714e+04 | NaN | 4.503714e+04 | NaN | GQDHCGIESEVVAGIPR |
| MAP3K5 | M3K5\_HUMAN | Mitogen-activated protein kinase kinase kinase 5 | 928 | 927 | MAP3K5 | M3K5\_HUMAN | Mitogen-activated protein kinase kinase kinase 5 | 2C(125.0477) | NaN | X | NaN | NaN | NaN | NaN | 4.500162e+04 | 4.500162e+04 | NaN | NaN | ACANDLLVDEFLK |
| CSE1L | XPO2\_HUMAN | Exportin-2 | 272 | 269 | CSE1L | XPO2\_HUMAN | Exportin-2 | 4C(125.0477) | NaN | X | NaN | NaN | NaN | NaN | 4.494190e+04 | 4.494190e+04 | NaN | NaN | SQICDNAALYAQK |
| PPP5C | PPP5\_HUMAN | Serine/threonine-protein phosphatase 5 | 404 | 400 | PPP5C | PPP5\_HUMAN | Serine/threonine-protein phosphatase 5 | 5C(125.0477) | NaN | X | NaN | NaN | NaN | NaN | 4.493589e+04 | 4.493589e+04 | NaN | NaN | RGVSCQFGPDVTK |
| IAH1 | IAH1\_HUMAN | Isoamyl acetate-hydrolyzing esterase 1 homolog | 137, 145 | 128 | IAH1 | IAH1\_HUMAN | Isoamyl acetate-hydrolyzing esterase 1 homolog | 10C(125.0477) 18C(15.9949) | NaN | X | X | NaN | NaN | NaN | 4.492814e+04 | 4.492814e+04 | NaN | NaN | VILITPTPLCETAWEEQCIIQGCK |
| MYH9 | MYH9\_HUMAN | Myosin-9 | 1379 | 1373 | MYH9 | MYH9\_HUMAN | Myosin-9 | 7C(125.0477) | NaN | X | NaN | NaN | NaN | NaN | 4.486604e+04 | 4.486604e+04 | NaN | NaN | MEDSVGCLETAEEVK |
| ALAD | HEM2\_HUMAN | Delta-aminolevulinic acid dehydratase | 75 | 75 | ALAD | HEM2\_HUMAN | Delta-aminolevulinic acid dehydratase | 1C(125.0477) | NaN | X | NaN | NaN | NaN | NaN | 4.482097e+04 | 4.482097e+04 | NaN | NaN | CVLIFGVPSRVPK |
| CRKL | CRKL\_HUMAN | Crk-like protein | 249 | 247 | CRKL | CRKL\_HUMAN | Crk-like protein | 3C(125.0477) | NaN | X | NaN | NaN | NaN | NaN | 4.479752e+04 | NaN | 4.479752e+04 | NaN | VPCAYDK |
| PPP1CA | PP1A\_HUMAN | Serine/threonine-protein phosphatase PP1-alpha catalytic subunit | 245 | 235 | PPP1CA | PP1A\_HUMAN | Serine/threonine-protein phosphatase PP1-alpha catalytic subunit | 11C(125.0477) | NaN | X | NaN | NaN | NaN | NaN | 4.459836e+04 | 4.459836e+04 | NaN | NaN | FLHKHDLDLICR |
| NASP | NASP\_HUMAN | Nuclear autoantigenic sperm protein | 254 | 244 | NASP | NASP\_HUMAN | Nuclear autoantigenic sperm protein | 11C(125.0477) | NaN | X | NaN | NaN | NaN | NaN | 4.455797e+04 | 4.455797e+04 | NaN | NaN | SVSGTDVQEECREK |
| MTURN | MTURN\_HUMAN | Maturin | 14 | 13 | MTURN | MTURN\_HUMAN | Maturin | 2C(125.0477) | NaN | X | NaN | NaN | NaN | NaN | 4.445353e+04 | 4.445353e+04 | NaN | NaN | WCSNTPFELIATEETERR |
| THG1L | THG1\_HUMAN | Probable tRNA(His) guanylyltransferase | 279 | 272 | THG1L | THG1\_HUMAN | Probable tRNA(His) guanylyltransferase | 8C(125.0477) | NaN | X | NaN | NaN | NaN | NaN | 4.442994e+04 | 4.442994e+04 | NaN | NaN | TKPVPLHCDIIGDAFWK |
| CYB5R3 | NB5R3\_HUMAN | NADH-cytochrome b5 reductase 3 | 204 | 193 | CYB5R3 | NB5R3\_HUMAN | NADH-cytochrome b5 reductase 3 | 12C(125.0477) | NaN | X | NaN | NaN | NaN | NaN | 4.441933e+04 | 4.441933e+04 | NaN | NaN | AIMKDPDDHTVCHLLFANQTEK |
| PPIA | PPIA\_HUMAN | Peptidyl-prolyl cis-trans isomerase A | 52 | 45 | PPIA | PPIA\_HUMAN | Peptidyl-prolyl cis-trans isomerase A | 8C(125.0477) | NaN | X | NaN | NaN | NaN | NaN | 4.436956e+04 | 4.436956e+04 | NaN | NaN | GFGYKGSCFHR |
| MAT2B | MAT2B\_HUMAN | Methionine adenosyltransferase 2 subunit beta | 17 | 7 | MAT2B | MAT2B\_HUMAN | Methionine adenosyltransferase 2 subunit beta | 11C(125.0477) | NaN | X | NaN | NaN | NaN | NaN | 4.435563e+04 | 4.435563e+04 | NaN | NaN | ELSIHFVPGSCR |
| UBR4 | UBR4\_HUMAN | E3 ubiquitin-protein ligase UBR4 | 5121 | 5110 | UBR4 | UBR4\_HUMAN | E3 ubiquitin-protein ligase UBR4 | 12C(125.0477) | NaN | X | NaN | NaN | NaN | NaN | 4.434989e+04 | 4.434989e+04 | NaN | NaN | VPTSNTEGGWSCSLAEYIR |
| WNK1 | WNK1\_HUMAN | Serine/threonine-protein kinase WNK1 | 1135 | 1132 | WNK1 | WNK1\_HUMAN | Serine/threonine-protein kinase WNK1 | 4C(125.0477) | NaN | X | NaN | NaN | NaN | NaN | 4.433167e+04 | 4.433167e+04 | NaN | NaN | VVECQLETHNRK |
| CFHR1 | FHR1\_HUMAN | Complement factor H-related protein 1 | 300 | 292 | CFHR1 | FHR1\_HUMAN | Complement factor H-related protein 1 | 9C(57.0215) | X | NaN | NaN | NaN | NaN | NaN | 4.431845e+04 | NaN | 4.431845e+04 | NaN | TGESAEFVCK |
| GPX4 | GPX4\_HUMAN | Phospholipid hydroperoxide glutathione peroxidase | 175 | 168 | GPX4 | GPX4\_HUMAN | Phospholipid hydroperoxide glutathione peroxidase | 8C(125.0477) | NaN | X | NaN | NaN | NaN | NaN | 4.429591e+04 | 4.429591e+04 | NaN | NaN | FLIDKNGCVVK |
| XPO1 | XPO1\_HUMAN | Exportin-1 | 528 | 523 | XPO1 | XPO1\_HUMAN | Exportin-1 | 6C(125.0477) | NaN | X | NaN | NaN | NaN | NaN | 4.421752e+04 | 4.421752e+04 | NaN | NaN | DLLGLCEQKR |
| PPP2R1B | 2AAB\_HUMAN | Serine/threonine-protein phosphatase 2A 65 kDa regulatory subunit A beta isoform | 389 | 387 | PPP2R1B | 2AAB\_HUMAN | Serine/threonine-protein phosphatase 2A 65 kDa regulatory subunit A beta isoform | 3C(125.0477) | NaN | X | NaN | NaN | NaN | NaN | 4.421634e+04 | NaN | 4.421634e+04 | NaN | DECPDVR |
| PTER | PTER\_HUMAN | Phosphotriesterase-related protein | 289 | 283 | PTER | PTER\_HUMAN | Phosphotriesterase-related protein | 7C(125.0477) | NaN | X | NaN | NaN | NaN | NaN | 4.416662e+04 | 4.416662e+04 | NaN | NaN | LLVEEGCEDR |
| XPO4 | XPO4\_HUMAN | Exportin-4 | 988 | 983 | XPO4 | XPO4\_HUMAN | Exportin-4 | 6C(125.0477) | NaN | X | NaN | NaN | NaN | NaN | 4.415907e+04 | 4.415907e+04 | NaN | NaN | LITFICEIFPEK |
| PAICS | PUR6\_HUMAN | Multifunctional protein ADE2 | 266 | 256 | PAICS | PUR6\_HUMAN | Multifunctional protein ADE2 | 11C(125.0477) | NaN | X | NaN | NaN | NaN | NaN | 4.409190e+04 | 4.409190e+04 | NaN | NaN | VELLLKSESQCR |
| ROCK1 | ROCK1\_HUMAN | Rho-associated protein kinase 1 | 1300 | 1294 | ROCK1 | ROCK1\_HUMAN | Rho-associated protein kinase 1 | 7C(125.0477) | NaN | X | NaN | NaN | NaN | NaN | 4.408700e+04 | 4.408700e+04 | NaN | NaN | DMLLLACSQDEQKK |
| PRDX1 | PRDX1\_HUMAN | Peroxiredoxin-1 | 173 | 169 | PRDX1 | PRDX1\_HUMAN | Peroxiredoxin-1 | 5C(125.0477) | NaN | X | NaN | NaN | NaN | NaN | 4.404436e+04 | 4.404436e+04 | NaN | NaN | HGEVCPAGWKPGSDTIKPDVQKSK |
| BLMH | BLMH\_HUMAN | Bleomycin hydrolase | 164 | 164 | BLMH | BLMH\_HUMAN | Bleomycin hydrolase | 1C(125.0477) | NaN | X | NaN | NaN | NaN | NaN | 4.404293e+04 | 2.655406e+04 | 1.748887e+04 | NaN | CFPESYTTEATR |
| PHKB | KPBB\_HUMAN | Phosphorylase b kinase regulatory subunit beta | 348 | 348 | PHKB | KPBB\_HUMAN | Phosphorylase b kinase regulatory subunit beta | 1C(125.0477) | NaN | X | NaN | NaN | NaN | NaN | 4.385830e+04 | 4.385830e+04 | NaN | NaN | CYYKPAEIK |
| ALDH9A1 | AL9A1\_HUMAN | 4-trimethylaminobutyraldehyde dehydrogenase | 96 | 86 | ALDH9A1 | AL9A1\_HUMAN | 4-trimethylaminobutyraldehyde dehydrogenase | 11C(125.0477) | NaN | X | NaN | NaN | NaN | NaN | 4.381739e+04 | 4.381739e+04 | NaN | NaN | EREDEIATMECINNGK |
| TCEA1 | TCEA1\_HUMAN | Transcription elongation factor A protein 1 | 212 | 209 | TCEA1 | TCEA1\_HUMAN | Transcription elongation factor A protein 1 | 4C(125.0477) | NaN | X | NaN | NaN | NaN | NaN | 4.380314e+04 | 4.380314e+04 | NaN | NaN | NVLCGNIPPDLFAR |
| DDT | DOPD\_HUMAN | D-dopachrome decarboxylase | 24, 24, 24 | 22 | DDT | DOPD\_HUMAN | D-dopachrome decarboxylase | 3C(-34.0809) 3C(125.0477) 3C(31.9988) | NaN | X | NaN | X | X | NaN | 4.371463e+04 | 4.371463e+04 | NaN | NaN | RLCAAAASILGKPADR |
| HSPA8 | HSP7C\_HUMAN | Heat shock cognate 71 kDa protein | 574 | 574 | HSPA8 | HSP7C\_HUMAN | Heat shock cognate 71 kDa protein | 1C(125.0477) | NaN | X | NaN | NaN | NaN | NaN | 4.369997e+04 | 4.369997e+04 | NaN | NaN | CNEIINWLDK |
| RPS4X | RS4X\_HUMAN | 40S ribosomal protein S4, X isoform | 41 | 40 | RPS4X | RS4X\_HUMAN | 40S ribosomal protein S4, X isoform | 2C(125.0477) | NaN | X | NaN | NaN | NaN | NaN | 4.366176e+04 | 4.366176e+04 | NaN | NaN | ECLPLIIFLR |
| ASPSCR1 | ASPC1\_HUMAN | Tether containing UBX domain for GLUT4 | 40 | 26 | ASPSCR1 | ASPC1\_HUMAN | Tether containing UBX domain for GLUT4 | 15C(125.0477) | NaN | X | NaN | NaN | NaN | NaN | 4.365197e+04 | 4.365197e+04 | NaN | NaN | VTPSTVLLQVLEDTCRR |
| UGP2 | UGPA\_HUMAN | UTP--glucose-1-phosphate uridylyltransferase | 276 | 275 | UGP2 | UGPA\_HUMAN | UTP--glucose-1-phosphate uridylyltransferase | 2C(125.0477) | NaN | X | NaN | NaN | NaN | NaN | 4.345314e+04 | 4.345314e+04 | NaN | NaN | RCEFVMEVTNK |
| ACTB | ACTB\_HUMAN | Actin, cytoplasmic 1 | 217, 217 | 216 | ACTB | ACTB\_HUMAN | Actin, cytoplasmic 1 | 2C(125.0477) 2C(57.0215) | X | X | NaN | NaN | NaN | NaN | 4.332830e+04 | 4.332830e+04 | NaN | NaN | LCYVALDFEQEMATAASSSSLEK |
| GMDS | GMDS\_HUMAN | GDP-mannose 4,6 dehydratase | 142 | 141 | GMDS | GMDS\_HUMAN | GDP-mannose 4,6 dehydratase | 2C(125.0477) | NaN | X | NaN | NaN | NaN | NaN | 4.322746e+04 | 4.322746e+04 | NaN | NaN | TCGLINSVK |
| ZNF330 | ZN330\_HUMAN | Zinc finger protein 330 | 216 | 216 | ZNF330 | ZN330\_HUMAN | Zinc finger protein 330 | 1C(125.0477) | NaN | X | NaN | NaN | NaN | NaN | 4.320655e+04 | 4.320655e+04 | NaN | NaN | CGHETQETK |
| PLIN3 | PLIN3\_HUMAN | Perilipin-3 | 341 | 332 | PLIN3 | PLIN3\_HUMAN | Perilipin-3 | 10C(125.0477) | NaN | X | NaN | NaN | NaN | NaN | 4.313181e+04 | 4.313181e+04 | NaN | NaN | DIAQQLQATCTSLGSSIQGLPTNVK |
| PNP | PNPH\_HUMAN | Purine nucleoside phosphorylase | 231 | 230 | PNP | PNPH\_HUMAN | Purine nucleoside phosphorylase | 2C(125.0477) | NaN | X | NaN | NaN | NaN | NaN | 4.313132e+04 | 4.313132e+04 | NaN | NaN | HCGLRVFGFSLITNK |
| KRT1 | K2C1\_HUMAN | Keratin, type II cytoskeletal 1 | 497 | 484 | KRT1 | K2C1\_HUMAN | Keratin, type II cytoskeletal 1 | 14C(31.9988) | NaN | NaN | NaN | X | NaN | NaN | 4.311912e+04 | 4.311912e+04 | NaN | NaN | TLLEGEESRMSGECAPNVSVSVSTSHTTISGGGSR |
| PSMB5 | PSB5\_HUMAN | Proteasome subunit beta type-5 | 161 | 151 | PSMB5 | PSB5\_HUMAN | Proteasome subunit beta type-5 | 11C(125.0477) | NaN | X | NaN | NaN | NaN | NaN | 4.310463e+04 | 4.310463e+04 | NaN | NaN | GMGLSMGTMICGWDK |
| KRT5 | K2C5\_HUMAN | Keratin, type II cytoskeletal 5 | 407 | 406 | KRT5 | K2C5\_HUMAN | Keratin, type II cytoskeletal 5 | 2C(125.0477) | NaN | X | NaN | NaN | NaN | NaN | 4.310127e+04 | 4.310127e+04 | NaN | NaN | QCANLQNAIADAEQR |
| OTULIN | OTUL\_HUMAN | Ubiquitin thioesterase otulin | 92 | 80 | OTULIN | OTUL\_HUMAN | Ubiquitin thioesterase otulin | 13C(125.0477) | NaN | X | NaN | NaN | NaN | NaN | 4.306921e+04 | 4.306921e+04 | NaN | NaN | LSVAPEMDIMDYCKK |
| HUWE1 | HUWE1\_HUMAN | E3 ubiquitin-protein ligase HUWE1 | 19 | 10 | HUWE1 | HUWE1\_HUMAN | E3 ubiquitin-protein ligase HUWE1 | 10C(125.0477) | NaN | X | NaN | NaN | NaN | NaN | 4.305923e+04 | 4.305923e+04 | NaN | NaN | KTPTEAPADCR |
| NAA10 | NAA10\_HUMAN | N-alpha-acetyltransferase 10 | 194 | 184 | NAA10 | NAA10\_HUMAN | N-alpha-acetyltransferase 10 | 11C(125.0477) | NaN | X | NaN | NaN | NaN | NaN | 4.305627e+04 | 4.305627e+04 | NaN | NaN | GNSPPSSGEACR |
| IPO7 | IPO7\_HUMAN | Importin-7 | 415 | 411 | IPO7 | IPO7\_HUMAN | Importin-7 | 5C(125.0477) | NaN | X | NaN | NaN | NaN | NaN | 4.296047e+04 | 4.296047e+04 | NaN | NaN | TMGFCYQILTEPNADPR |
| MMS19 | MMS19\_HUMAN | MMS19 nucleotide excision repair protein homolog | 386 | 385 | MMS19 | MMS19\_HUMAN | MMS19 nucleotide excision repair protein homolog | 2C(125.0477) | NaN | X | NaN | NaN | NaN | NaN | 4.293268e+04 | 4.293268e+04 | NaN | NaN | ACDSVTSNVLPLLLEQFHK |
| WDSUB1 | WSDU1\_HUMAN | WD repeat, SAM and U-box domain-containing protein 1 | 410 | 399 | WDSUB1 | WSDU1\_HUMAN | WD repeat, SAM and U-box domain-containing protein 1 | 12C(125.0477) | NaN | X | NaN | NaN | NaN | NaN | 4.292860e+04 | 4.292860e+04 | NaN | NaN | SLSSGIPDEFICPITR |
| GPS1 | CSN1\_HUMAN | COP9 signalosome complex subunit 1 | 251 | 249 | GPS1 | CSN1\_HUMAN | COP9 signalosome complex subunit 1 | 3C(125.0477) | NaN | X | NaN | NaN | NaN | NaN | 4.290234e+04 | 4.290234e+04 | NaN | NaN | LKCAAGLAELAAR |
| NUMA1 | NUMA1\_HUMAN | Nuclear mitotic apparatus protein 1 | 65 | 61 | NUMA1 | NUMA1\_HUMAN | Nuclear mitotic apparatus protein 1 | 5C(125.0477) | NaN | X | NaN | NaN | NaN | NaN | 4.287918e+04 | 4.287918e+04 | NaN | NaN | LDFVCSFLQK |
| ARAP1 | ARAP1\_HUMAN | Arf-GAP with Rho-GAP domain, ANK repeat and PH domain-containing protein 1 | 900 | 893 | ARAP1 | ARAP1\_HUMAN | Arf-GAP with Rho-GAP domain, ANK repeat and PH domain-containing protein 1 | 8C(125.0477) | NaN | X | NaN | NaN | NaN | NaN | 4.283751e+04 | 4.283751e+04 | NaN | NaN | AVFPEGPCEEPLQLR |
| ROCK2 | ROCK2\_HUMAN | Rho-associated protein kinase 2 | 887 | 882 | ROCK2 | ROCK2\_HUMAN | Rho-associated protein kinase 2 | 6C(125.0477) | NaN | X | NaN | NaN | NaN | NaN | 4.281541e+04 | 4.281541e+04 | NaN | NaN | ELKEECEEKTK |
| UROD | DCUP\_HUMAN | Uroporphyrinogen decarboxylase | 308 | 298 | UROD | DCUP\_HUMAN | Uroporphyrinogen decarboxylase | 11C(125.0477) | NaN | X | NaN | NaN | NaN | NaN | 4.279318e+04 | 3.991886e+04 | 2.874319e+03 | NaN | TVTLQGNLDPCALYASEEEIGQLVK |
| ACSL4 | ACSL4\_HUMAN | Long-chain-fatty-acid--CoA ligase 4 | 420 | 413 | ACSL4 | ACSL4\_HUMAN | Long-chain-fatty-acid--CoA ligase 4 | 8C(125.0477) | NaN | X | NaN | NaN | NaN | NaN | 4.276678e+04 | 4.276678e+04 | NaN | NaN | KGYDAPLCNLLLFK |
| CFL1 | COF1\_HUMAN | Cofilin-1 | 80 | 79 | CFL1 | COF1\_HUMAN | Cofilin-1 | 2C(15.9949) | NaN | NaN | X | NaN | NaN | NaN | 4.272558e+04 | 4.272558e+04 | NaN | NaN | DCRYALYDATYETK |
| GLRX3 | GLRX3\_HUMAN | Glutaredoxin-3 | 146 | 139 | GLRX3 | GLRX3\_HUMAN | Glutaredoxin-3 | 8C(125.0477) | NaN | X | NaN | NaN | NaN | NaN | 4.271331e+04 | 4.271331e+04 | NaN | NaN | KLTHAAPCMLFMK |
| UBL4A | UBL4A\_HUMAN | Ubiquitin-like protein 4A | 13 | 12 | UBL4A | UBL4A\_HUMAN | Ubiquitin-like protein 4A | 2C(125.0477) | NaN | X | NaN | NaN | NaN | NaN | 4.270711e+04 | 4.270711e+04 | NaN | NaN | ECSLQVPEDELVSTLK |
| AKT2 | AKT2\_HUMAN | RAC-beta serine/threonine-protein kinase | 77 | 77 | AKT2 | AKT2\_HUMAN | RAC-beta serine/threonine-protein kinase | 1C(125.0477) | NaN | X | NaN | NaN | NaN | NaN | 4.264603e+04 | 4.264603e+04 | NaN | NaN | CLQWTTVIER |
| ARHGDIA | GDIR1\_HUMAN | Rho GDP-dissociation inhibitor 1 | 79 | 59 | ARHGDIA | GDIR1\_HUMAN | Rho GDP-dissociation inhibitor 1 | 21C(125.0477) | NaN | X | NaN | NaN | NaN | NaN | 4.254911e+04 | 4.254911e+04 | NaN | NaN | VAVSADPNVPNVVVTGLTLVCSSAPGPLELDLTGDLESFKK |
| CUL3 | CUL3\_HUMAN | Cullin-3 | 298 | 293 | CUL3 | CUL3\_HUMAN | Cullin-3 | 6C(125.0477) | NaN | X | NaN | NaN | NaN | NaN | 4.242704e+04 | NaN | 4.242704e+04 | NaN | TEDLGCMYK |
| PSMA7 | PSA7\_HUMAN | Proteasome subunit alpha type-7 | 91 | 89 | PSMA7 | PSA7\_HUMAN | Proteasome subunit alpha type-7 | 3C(125.0477) | NaN | X | NaN | NaN | NaN | NaN | 4.240104e+04 | 4.240104e+04 | NaN | NaN | VECQSHR |
| SOD1 | SODC\_HUMAN | Superoxide dismutase [Cu-Zn] | 112 | 93 | SOD1 | SODC\_HUMAN | Superoxide dismutase [Cu-Zn] | 20C(125.0477) | NaN | X | NaN | NaN | NaN | NaN | 4.239372e+04 | 4.239372e+04 | NaN | NaN | DGVADVSIEDSVISLSGDHCIIGR |
| NUDT5 | NUDT5\_HUMAN | ADP-sugar pyrophosphatase | 76 | 71 | NUDT5 | NUDT5\_HUMAN | ADP-sugar pyrophosphatase | 6C(125.0477) | NaN | X | NaN | NaN | NaN | NaN | 4.238533e+04 | 4.238533e+04 | NaN | NaN | TLHYECIVLVK |
| CCT5 | TCPE\_HUMAN | T-complex protein 1 subunit epsilon | 302 | 294 | CCT5 | TCPE\_HUMAN | T-complex protein 1 subunit epsilon | 9C(125.0477) | NaN | X | NaN | NaN | NaN | NaN | 4.236339e+04 | 4.236339e+04 | NaN | NaN | ETGANLAICQWGFDDEANHLLLQNNLPAVR |
| SNX12 | SNX12\_HUMAN | Sorting nexin-12 | 141 | 141 | SNX12 | SNX12\_HUMAN | Sorting nexin-12 | 1C(125.0477) | NaN | X | NaN | NaN | NaN | NaN | 4.235624e+04 | 4.235624e+04 | NaN | NaN | CLHMFLQEEAIDR |
| UBE2O | UBE2O\_HUMAN | (E3-independent) E2 ubiquitin-conjugating enzyme | 314 | 312 | UBE2O | UBE2O\_HUMAN | (E3-independent) E2 ubiquitin-conjugating enzyme | 3C(125.0477) | NaN | X | NaN | NaN | NaN | NaN | 4.232501e+04 | 4.232501e+04 | NaN | NaN | SFCPGGTDSVSPPPSVITQENLGR |
| AK1 | KAD1\_HUMAN | Adenylate kinase isoenzyme 1 | 25 | 10 | AK1 | KAD1\_HUMAN | Adenylate kinase isoenzyme 1 | 16C(125.0477) | NaN | X | NaN | NaN | NaN | NaN | 4.230675e+04 | 4.230675e+04 | NaN | NaN | IIFVVGGPGSGKGTQCEK |
| WDR20 | WDR20\_HUMAN | WD repeat-containing protein 20 | 237 | 234 | WDR20 | WDR20\_HUMAN | WD repeat-containing protein 20 | 4C(125.0477) | NaN | X | NaN | NaN | NaN | NaN | 4.224950e+04 | 4.224950e+04 | NaN | NaN | FLACVSQDGFLR |
| NMNAT3 | NMNA3\_HUMAN | Nicotinamide/nicotinic acid mononucleotide adenylyltransferase 3 | 12 | 5 | NMNAT3 | NMNA3\_HUMAN | Nicotinamide/nicotinic acid mononucleotide adenylyltransferase 3 | 8C(125.0477) | NaN | X | NaN | NaN | NaN | NaN | 4.222558e+04 | 4.222558e+04 | NaN | NaN | IPVVLLACGSFNPITNMHLR |
| PSMC5 | PRS8\_HUMAN | 26S proteasome regulatory subunit 8 | 112 | 95 | PSMC5 | PRS8\_HUMAN | 26S proteasome regulatory subunit 8 | 18C(125.0477) | NaN | X | NaN | NaN | NaN | NaN | 4.221390e+04 | 4.221390e+04 | NaN | NaN | FVVDVDKNIDINDVTPNCR |
| COPS2 | CSN2\_HUMAN | COP9 signalosome complex subunit 2 | 179 | 174 | COPS2 | CSN2\_HUMAN | COP9 signalosome complex subunit 2 | 6C(125.0477) | NaN | X | NaN | NaN | NaN | NaN | 4.218808e+04 | NaN | 4.218808e+04 | NaN | QLHQSCQTDDGEDDLKK |
| SLC44A2 | CTL2\_HUMAN | Choline transporter-like protein 2 | 172 | 172 | SLC44A2 | CTL2\_HUMAN | Choline transporter-like protein 2 | 1C(125.0477) | NaN | X | NaN | NaN | NaN | NaN | 4.213181e+04 | NaN | 4.213181e+04 | NaN | CFPAIHAYK |
| ACSL6 | ACSL6\_HUMAN | Long-chain-fatty-acid--CoA ligase 6 | 359 | 357 | ACSL6 | ACSL6\_HUMAN | Long-chain-fatty-acid--CoA ligase 6 | 3C(125.0477) | NaN | X | NaN | NaN | NaN | NaN | 4.200896e+04 | 4.200896e+04 | NaN | NaN | ALCPTIFPVVPR |
| EPS15 | EPS15\_HUMAN | Epidermal growth factor receptor substrate 15 | 657 | 649 | EPS15 | EPS15\_HUMAN | Epidermal growth factor receptor substrate 15 | 9C(125.0477) | NaN | X | NaN | NaN | NaN | NaN | 4.199254e+04 | 4.199254e+04 | NaN | NaN | GSDPFASDCFFR |
| COPB1 | COPB\_HUMAN | Coatomer subunit beta | 235 | 234 | COPB1 | COPB\_HUMAN | Coatomer subunit beta | 2C(125.0477) | NaN | X | NaN | NaN | NaN | NaN | 4.198574e+04 | 4.198574e+04 | NaN | NaN | VCHANPSER |
| UBR1 | UBR1\_HUMAN | E3 ubiquitin-protein ligase UBR1 | 1369 | 1367 | UBR1 | UBR1\_HUMAN | E3 ubiquitin-protein ligase UBR1 | 3C(125.0477) | NaN | X | NaN | NaN | NaN | NaN | 4.196369e+04 | 4.196369e+04 | NaN | NaN | ITCPQVLIQK |
| KRT13 | K1C13\_HUMAN | Keratin, type I cytoskeletal 13 | 383, 383 | 379 | KRT13 | K1C13\_HUMAN | Keratin, type I cytoskeletal 13 | 5C(125.0477) 5C(15.9949) | NaN | X | X | NaN | NaN | NaN | 4.193761e+04 | NaN | 4.193761e+04 | NaN | SEMECQNQEYK |
| USP9X | USP9X\_HUMAN | Probable ubiquitin carboxyl-terminal hydrolase FAF-X | 577 | 575 | USP9X | USP9X\_HUMAN | Probable ubiquitin carboxyl-terminal hydrolase FAF-X | 3C(125.0477) | NaN | X | NaN | NaN | NaN | NaN | 4.191517e+04 | 4.191517e+04 | NaN | NaN | EICSLFGEAPQNLSQTQR |
| LLGL1 | L2GL1\_HUMAN | Lethal(2) giant larvae protein homolog 1 | 947 | 941 | LLGL1 | L2GL1\_HUMAN | Lethal(2) giant larvae protein homolog 1 | 7C(125.0477) | NaN | X | NaN | NaN | NaN | NaN | 4.189800e+04 | 4.189800e+04 | NaN | NaN | NITEPLCSLDINWPR |
| NLN | NEUL\_HUMAN | Neurolysin, mitochondrial | 153 | 146 | NLN | NEUL\_HUMAN | Neurolysin, mitochondrial | 8C(125.0477) | NaN | X | NaN | NaN | NaN | NaN | 4.186248e+04 | 4.186248e+04 | NaN | NaN | IVHLQETCDLGK |
| PAFAH1B1 | LIS1\_HUMAN | Platelet-activating factor acetylhydrolase IB subunit beta | 184 | 176 | PAFAH1B1 | LIS1\_HUMAN | Platelet-activating factor acetylhydrolase IB subunit beta | 9C(125.0477) | NaN | X | NaN | NaN | NaN | NaN | 4.179256e+04 | 4.179256e+04 | NaN | NaN | LWDFQGFECIR |
| OSTC | OSTC\_HUMAN | Oligosaccharyltransferase complex subunit | 14 | 7 | OSTC | OSTC\_HUMAN | Oligosaccharyltransferase complex subunit | 8C(125.0477) | NaN | X | NaN | NaN | NaN | NaN | 4.178339e+04 | 4.178339e+04 | NaN | NaN | VPFLVLECPNLK |
| PGM2L1 | PGM2L\_HUMAN | Glucose 1,6-bisphosphate synthase | 236 | 230 | PGM2L1 | PGM2L\_HUMAN | Glucose 1,6-bisphosphate synthase | 7C(125.0477) | NaN | X | NaN | NaN | NaN | NaN | 4.168332e+04 | NaN | 4.168332e+04 | NaN | DPLQDICR |
| AFMID | KFA\_HUMAN | Kynurenine formamidase | 237 | 227 | AFMID | KFA\_HUMAN | Kynurenine formamidase | 11C(125.0477) | NaN | X | NaN | NaN | NaN | NaN | 4.168169e+04 | 4.168169e+04 | NaN | NaN | VAQAQPVDPTCR |
| SLC43A2 | LAT4\_HUMAN | Large neutral amino acids transporter small subunit 4 | 392 | 389 | SLC43A2 | LAT4\_HUMAN | Large neutral amino acids transporter small subunit 4 | 4C(125.0477) | NaN | X | NaN | NaN | NaN | NaN | 4.166607e+04 | NaN | 4.166607e+04 | NaN | LKECEDASEEPEEK |
| RIDA | RIDA\_HUMAN | 2-iminobutanoate/2-iminopropanoate deaminase | 71 | 68 | RIDA | RIDA\_HUMAN | 2-iminobutanoate/2-iminopropanoate deaminase | 4C(125.0477) | NaN | X | NaN | NaN | NaN | NaN | 4.159376e+04 | NaN | 4.159376e+04 | NaN | AAGCDFTNVVK |
| ARHGAP17 | RHG17\_HUMAN | Rho GTPase-activating protein 17 | 53 | 50 | ARHGAP17 | RHG17\_HUMAN | Rho GTPase-activating protein 17 | 4C(125.0477) | NaN | X | NaN | NaN | NaN | NaN | 4.158391e+04 | 4.158391e+04 | NaN | NaN | LVACFQGQHGTDAER |
| WNK1 | WNK1\_HUMAN | Serine/threonine-protein kinase WNK1 | 352 | 352 | WNK1 | WNK1\_HUMAN | Serine/threonine-protein kinase WNK1 | 1C(125.0477) | NaN | X | NaN | NaN | NaN | NaN | 4.143182e+04 | NaN | 4.143182e+04 | NaN | CDNIFITGPTGSVK |
| PGM2L1 | PGM2L\_HUMAN | Glucose 1,6-bisphosphate synthase | 583 | 578 | PGM2L1 | PGM2L\_HUMAN | Glucose 1,6-bisphosphate synthase | 6C(125.0477) | NaN | X | NaN | NaN | NaN | NaN | 4.131377e+04 | 4.131377e+04 | NaN | NaN | YYAEMCASPDQSDTALLEEELKK |
| IGHG1 | IGHG1\_HUMAN | Immunoglobulin heavy constant gamma 1 | 27 | 17 | IGHG1 | IGHG1\_HUMAN | Immunoglobulin heavy constant gamma 1 | 11C(125.0477) | NaN | X | NaN | NaN | NaN | NaN | 4.123280e+04 | 4.123280e+04 | NaN | NaN | STSGGTAALGCLVK |
| A2M | A2MG\_HUMAN | Alpha-2-macroglobulin | 563 | 558 | A2M | A2MG\_HUMAN | Alpha-2-macroglobulin | 6C(57.0215) | X | NaN | NaN | NaN | NaN | NaN | 4.123170e+04 | NaN | 4.123170e+04 | NaN | YDVENCLANK |
| GLIPR2 | GAPR1\_HUMAN | Golgi-associated plant pathogenesis-related protein 1 | 63 | 61 | GLIPR2 | GAPR1\_HUMAN | Golgi-associated plant pathogenesis-related protein 1 | 3C(125.0477) | NaN | X | NaN | NaN | NaN | NaN | 4.119788e+04 | 4.119788e+04 | NaN | NaN | GQCGENLAWASYDQTGK |
| PEPD | PEPD\_HUMAN | Xaa-Pro dipeptidase | 183 | 160 | PEPD | PEPD\_HUMAN | Xaa-Pro dipeptidase | 24C(125.0477) | NaN | X | NaN | NaN | NaN | NaN | 4.117780e+04 | 4.117780e+04 | NaN | NaN | EASFDGISKFEVNNTILHPEIVECR |
| PGP | PGP\_HUMAN | Glycerol-3-phosphate phosphatase | 297 | 291 | PGP | PGP\_HUMAN | Glycerol-3-phosphate phosphatase | 7C(125.0477) | NaN | X | NaN | NaN | NaN | NaN | 4.116994e+04 | 4.116994e+04 | NaN | NaN | NNQESDCVSKK |
| ENDOD1 | ENDD1\_HUMAN | Endonuclease domain-containing 1 protein | 34 | 23 | ENDOD1 | ENDD1\_HUMAN | Endonuclease domain-containing 1 protein | 12C(125.0477) | NaN | X | NaN | NaN | NaN | NaN | 4.112754e+04 | 4.112754e+04 | NaN | NaN | LVGEEEAGFGECDKFFYAGTPPAGLAADSHVK |
| WDFY1 | WDFY1\_HUMAN | WD repeat and FYVE domain-containing protein 1 | 401 | 392 | WDFY1 | WDFY1\_HUMAN | WD repeat and FYVE domain-containing protein 1 | 10C(125.0477) | NaN | X | NaN | NaN | NaN | NaN | 4.111513e+04 | 4.111513e+04 | NaN | NaN | IWDMTPVVGCSLATGFSPH |
| CNBP | CNBP\_HUMAN | CCHC-type zinc finger nucleic acid binding protein | 57 | 57 | CNBP | CNBP\_HUMAN | CCHC-type zinc finger nucleic acid binding protein | 1C(125.0477) | NaN | X | NaN | NaN | NaN | NaN | 4.110794e+04 | 4.110794e+04 | NaN | NaN | CGESGHLAK |
| NFKB1 | NFKB1\_HUMAN | Nuclear factor NF-kappa-B p105 subunit | 87 | 86 | NFKB1 | NFKB1\_HUMAN | Nuclear factor NF-kappa-B p105 subunit | 2C(125.0477) | NaN | X | NaN | NaN | NaN | NaN | 4.110061e+04 | 4.110061e+04 | NaN | NaN | ICNYVGPAK |
| GATD1 | GALD1\_HUMAN | Glutamine amidotransferase-like class 1 domain-containing protein 1 | 94 | 86 | GATD1 | GALD1\_HUMAN | Glutamine amidotransferase-like class 1 domain-containing protein 1 | 9C(125.0477) | NaN | X | NaN | NaN | NaN | NaN | 4.097717e+04 | 4.097717e+04 | NaN | NaN | YHALLIPSCPGALTDLASSGSLAR |
| KYAT3 | KAT3\_HUMAN | Kynurenine--oxoglutarate transaminase 3 | 416 | 407 | KYAT3 | KAT3\_HUMAN | Kynurenine--oxoglutarate transaminase 3 | 10C(125.0477) | NaN | X | NaN | NaN | NaN | NaN | 4.096562e+04 | 4.096562e+04 | NaN | NaN | LSAIPVSAFCNSETK |
| RPIA | RPIA\_HUMAN | Ribose-5-phosphate isomerase | 178 | 174 | RPIA | RPIA\_HUMAN | Ribose-5-phosphate isomerase | 5C(125.0477) | NaN | X | NaN | NaN | NaN | NaN | 4.093702e+04 | 4.093702e+04 | NaN | NaN | GGGGCLTQEKIVAGYASR |
| ALDOA | ALDOA\_HUMAN | Fructose-bisphosphate aldolase A | 202 | 202 | ALDOA | ALDOA\_HUMAN | Fructose-bisphosphate aldolase A | 1C(125.0477) | NaN | X | NaN | NaN | NaN | NaN | 4.083244e+04 | 4.083244e+04 | NaN | NaN | CQYVTEKVLAAVYK |
| SEMA7A | SEM7A\_HUMAN | Semaphorin-7A | 291, 291 | 287 | SEMA7A | SEM7A\_HUMAN | Semaphorin-7A | 5C(125.0477) 5C(57.0215) | X | X | NaN | NaN | NaN | NaN | 4.082892e+04 | NaN | 4.082892e+04 | NaN | AMLVCSDAATNK |
| SEPSECS | SPCS\_HUMAN | O-phosphoseryl-tRNA(Sec) selenium transferase | 262 | 262 | SEPSECS | SPCS\_HUMAN | O-phosphoseryl-tRNA(Sec) selenium transferase | 1C(125.0477) | NaN | X | NaN | NaN | NaN | NaN | 4.081336e+04 | 4.081336e+04 | NaN | NaN | CMHLIQQGAR |
| IPO9 | IPO9\_HUMAN | Importin-9 | 450 | 446 | IPO9 | IPO9\_HUMAN | Importin-9 | 5C(125.0477) | NaN | X | NaN | NaN | NaN | NaN | 4.077940e+04 | 4.077940e+04 | NaN | NaN | IHEACMLALGSVK |
| HBQ1 | HBAT\_HUMAN | Hemoglobin subunit theta-1 | 90, 105 | 74 | HBQ1 | HBAT\_HUMAN | Hemoglobin subunit theta-1 | 17C(15.9949) 32C(15.9949) | NaN | NaN | X | NaN | NaN | NaN | 4.061609e+04 | 4.061609e+04 | NaN | NaN | LDDLPHALSALSHLHACQLRVDPASFQLLGHCLLVTLAR |
| RAB4B | RAB4B\_HUMAN | Ras-related protein Rab-4B | 23 | 22 | RAB4B | RAB4B\_HUMAN | Ras-related protein Rab-4B | 2C(125.0477) | NaN | X | NaN | NaN | NaN | NaN | 4.061012e+04 | 4.061012e+04 | NaN | NaN | SCLLHQFIENK |
| EEA1 | EEA1\_HUMAN | Early endosome antigen 1 | 704 | 697 | EEA1 | EEA1\_HUMAN | Early endosome antigen 1 | 8C(125.0477) | NaN | X | NaN | NaN | NaN | NaN | 4.059732e+04 | 4.059732e+04 | NaN | NaN | LQDKQEHCSQLESHLK |
| C11orf68 | CK068\_HUMAN | UPF0696 protein C11orf68 | 240 | 240 | C11orf68 | CK068\_HUMAN | UPF0696 protein C11orf68 | 1C(125.0477) | NaN | X | NaN | NaN | NaN | NaN | 4.056687e+04 | 4.056687e+04 | NaN | NaN | CLLTYKPDVYTYLGIYR |
| IPO5 | IPO5\_HUMAN | Importin-5 | 1057 | 1053 | IPO5 | IPO5\_HUMAN | Importin-5 | 5C(125.0477) | NaN | X | NaN | NaN | NaN | NaN | 4.056466e+04 | 4.056466e+04 | NaN | NaN | HEDPCAK |
| HSPBP1 | HPBP1\_HUMAN | Hsp70-binding protein 1 | 172 | 165 | HSPBP1 | HPBP1\_HUMAN | Hsp70-binding protein 1 | 8C(125.0477) | NaN | X | NaN | NaN | NaN | NaN | 4.050832e+04 | 4.050832e+04 | NaN | NaN | AAQLIGTCSQNVAAIQEQVLGLGALRK |
| UBQLN4 | UBQL4\_HUMAN | Ubiquilin-4 | 29 | 22 | UBQLN4 | UBQL4\_HUMAN | Ubiquilin-4 | 8C(125.0477) | NaN | X | NaN | NaN | NaN | NaN | 4.046767e+04 | 4.046767e+04 | NaN | NaN | DKEEIVICDR |
| IGHG2 | IGHG2\_HUMAN | Immunoglobulin heavy constant gamma 2 | 14 | 5 | IGHG2 | IGHG2\_HUMAN | Immunoglobulin heavy constant gamma 2 | 10C(57.0215) | X | NaN | NaN | NaN | NaN | NaN | 4.045819e+04 | 4.045819e+04 | NaN | NaN | GPSVFPLAPCSR |
| GBF1 | GBF1\_HUMAN | Golgi-specific brefeldin A-resistance guanine nucleotide exchange factor 1 | 1588 | 1580 | GBF1 | GBF1\_HUMAN | Golgi-specific brefeldin A-resistance guanine nucleotide exchange factor 1 | 9C(125.0477) | NaN | X | NaN | NaN | NaN | NaN | 4.042576e+04 | 4.042576e+04 | NaN | NaN | LDALEWESCFNK |
| HTT | HD\_HUMAN | Huntingtin | 278 | 269 | HTT | HD\_HUMAN | Huntingtin | 10C(125.0477) | NaN | X | NaN | NaN | NaN | NaN | 4.033804e+04 | 4.033804e+04 | NaN | NaN | TAAGSAVSICQHSR |
| NIF3L1 | NIF3L\_HUMAN | NIF3-like protein 1 | 213 | 209 | NIF3L1 | NIF3L\_HUMAN | NIF3-like protein 1 | 5C(125.0477) | NaN | X | NaN | NaN | NaN | NaN | 4.024765e+04 | NaN | 4.024765e+04 | NaN | INLNCTQK |
| SMS | SPSY\_HUMAN | Spermine synthase | 237 | 235 | SMS | SPSY\_HUMAN | Spermine synthase | 3C(15.9949) | NaN | NaN | X | NaN | NaN | NaN | 4.023188e+04 | 4.023188e+04 | NaN | NaN | KTCGDVLDNLK |
| STX7 | STX7\_HUMAN | Syntaxin-7 | 28 | 25 | STX7 | STX7\_HUMAN | Syntaxin-7 | 4C(31.9988) | NaN | NaN | NaN | X | NaN | NaN | 4.023018e+04 | NaN | NaN | 40230.1800 | ITQCSVEIQR |
| GPS1 | CSN1\_HUMAN | COP9 signalosome complex subunit 1 | 175, 184 | 163 | GPS1 | CSN1\_HUMAN | COP9 signalosome complex subunit 1 | 13C(15.9949) 22C(15.9949) | NaN | NaN | X | NaN | NaN | NaN | 4.022060e+04 | 4.022060e+04 | NaN | NaN | RGHDDLGDHYLDCGDLSNALKCYSR |
| HTATIP2 | HTAI2\_HUMAN | Oxidoreductase HTATIP2 | 172 | 162 | HTATIP2 | HTAI2\_HUMAN | Oxidoreductase HTATIP2 | 11C(125.0477) | NaN | X | NaN | NaN | NaN | NaN | 4.015696e+04 | 4.015696e+04 | NaN | NaN | YSVFRPGVLLCDR |
| RAB3GAP1 | RB3GP\_HUMAN | Rab3 GTPase-activating protein catalytic subunit | 858 | 853 | RAB3GAP1 | RB3GP\_HUMAN | Rab3 GTPase-activating protein catalytic subunit | 6C(125.0477) | NaN | X | NaN | NaN | NaN | NaN | 4.015309e+04 | 4.015309e+04 | NaN | NaN | FGTEKCEQEEEKEDLER |
| ORM1 | A1AG1\_HUMAN | Alpha-1-acid glycoprotein 1 | 165 | 154 | ORM1 | A1AG1\_HUMAN | Alpha-1-acid glycoprotein 1 | 12C(57.0215) | X | NaN | NaN | NaN | NaN | NaN | 4.011956e+04 | 4.011956e+04 | NaN | NaN | EQLGEFYEALDCLR |
| TF | TRFE\_HUMAN | Serotransferrin | 67 | 62 | TF | TRFE\_HUMAN | Serotransferrin | 6C(57.0215) | X | NaN | NaN | NaN | NaN | NaN | 4.008206e+04 | NaN | 4.008206e+04 | NaN | ASYLDCIR |
| DUSP23 | DUS23\_HUMAN | Dual specificity protein phosphatase 23 | 53 | 46 | DUSP23 | DUS23\_HUMAN | Dual specificity protein phosphatase 23 | 8C(125.0477) | NaN | X | NaN | NaN | NaN | NaN | 4.000521e+04 | 4.000521e+04 | NaN | NaN | GPPHSDSCPGLTLHR |
| ANKS1A | ANS1A\_HUMAN | Ankyrin repeat and SAM domain-containing protein 1A | 965 | 957 | ANKS1A | ANS1A\_HUMAN | Ankyrin repeat and SAM domain-containing protein 1A | 9C(125.0477) | NaN | X | NaN | NaN | NaN | NaN | 3.999228e+04 | 3.999228e+04 | NaN | NaN | GTESTQDACAK |
| PSMA3 | PSA3\_HUMAN | Proteasome subunit alpha type-3 | 42 | 42 | PSMA3 | PSA3\_HUMAN | Proteasome subunit alpha type-3 | 1C(125.0477) | NaN | X | NaN | NaN | NaN | NaN | 3.995383e+04 | 3.995383e+04 | NaN | NaN | CKDGVVFGVEKLVLSK |
| UBE2L3 | UB2L3\_HUMAN | Ubiquitin-conjugating enzyme E2 L3 | 86 | 74 | UBE2L3 | UB2L3\_HUMAN | Ubiquitin-conjugating enzyme E2 L3 | 13C(125.0477) | NaN | X | NaN | NaN | NaN | NaN | 3.990478e+04 | 3.990478e+04 | NaN | NaN | IYHPNIDEKGQVCLPVISAENWKPATK |
| MDM4 | MDM4\_HUMAN | Protein Mdm4 | 477 | 470 | MDM4 | MDM4\_HUMAN | Protein Mdm4 | 8C(-34.0809) | NaN | NaN | NaN | NaN | X | NaN | 3.976838e+04 | 3.976838e+04 | NaN | NaN | AGASCPICKK |
| CHORDC1 | CHRD1\_HUMAN | Cysteine and histidine-rich domain-containing protein 1 | 59 | 47 | CHORDC1 | CHRD1\_HUMAN | Cysteine and histidine-rich domain-containing protein 1 | 13C(125.0477) | NaN | X | NaN | NaN | NaN | NaN | 3.971100e+04 | 3.971100e+04 | NaN | NaN | TTDFSDFLSIVGCTK |
| TSN | TSN\_HUMAN | Translin | 225 | 220 | TSN | TSN\_HUMAN | Translin | 6C(125.0477) | NaN | X | NaN | NaN | NaN | NaN | 3.962764e+04 | NaN | 3.962764e+04 | NaN | ETAAACVEK |
| GAPDH | G3P\_HUMAN | Glyceraldehyde-3-phosphate dehydrogenase | 247 | 235 | GAPDH | G3P\_HUMAN | Glyceraldehyde-3-phosphate dehydrogenase | 13C(125.0477) | NaN | X | NaN | NaN | NaN | NaN | 3.958225e+04 | 3.958225e+04 | NaN | NaN | VPTANVSVVDLTCRLEKPAKYDDIK |
| ALDH1A1 | AL1A1\_HUMAN | Aldehyde dehydrogenase 1A1 | 370 | 364 | ALDH1A1 | AL1A1\_HUMAN | Aldehyde dehydrogenase 1A1 | 7C(125.0477) | NaN | X | NaN | NaN | NaN | NaN | 3.943203e+04 | 3.943203e+04 | NaN | NaN | EGAKLECGGGPWGNK |
| ARFGEF2 | BIG2\_HUMAN | Brefeldin A-inhibited guanine nucleotide-exchange protein 2 | 1617 | 1614 | ARFGEF2 | BIG2\_HUMAN | Brefeldin A-inhibited guanine nucleotide-exchange protein 2 | 4C(125.0477) | NaN | X | NaN | NaN | NaN | NaN | 3.941789e+04 | 3.941789e+04 | NaN | NaN | LLDCLQESHSFSK |
| ATP7A | ATP7A\_HUMAN | Copper-transporting ATPase 1 | 160 | 157 | ATP7A | ATP7A\_HUMAN | Copper-transporting ATPase 1 | 4C(125.0477) | NaN | X | NaN | NaN | NaN | NaN | 3.939527e+04 | NaN | 3.939527e+04 | NaN | SGACEDHSMAQAGEVVLK |
| CARS1 | SYCC\_HUMAN | Cysteine--tRNA ligase, cytoplasmic | 405 | 393 | CARS1 | SYCC\_HUMAN | Cysteine--tRNA ligase, cytoplasmic | 13C(125.0477) | NaN | X | NaN | NaN | NaN | NaN | 3.935018e+04 | 3.935018e+04 | NaN | NaN | YFLHTGHLTIAGCK |
| SNX5 | SNX5\_HUMAN | Sorting nexin-5 | 250 | 239 | SNX5 | SNX5\_HUMAN | Sorting nexin-5 | 12C(125.0477) | NaN | X | NaN | NaN | NaN | NaN | 3.930422e+04 | 3.930422e+04 | NaN | NaN | NVADDYIHTAACLHSLALEEPTVIKK |
| AZGP1 | ZA2G\_HUMAN | Zinc-alpha-2-glycoprotein | 186 | 180 | AZGP1 | ZA2G\_HUMAN | Zinc-alpha-2-glycoprotein | 7C(125.0477) | NaN | X | NaN | NaN | NaN | NaN | 3.923171e+04 | NaN | 3.923171e+04 | NaN | AYLEEECPATLR |
| ST13 | F10A1\_HUMAN | Hsc70-interacting protein | 16 | 15 | ST13 | F10A1\_HUMAN | Hsc70-interacting protein | 2C(125.0477) | NaN | X | NaN | NaN | NaN | NaN | 3.922713e+04 | 3.922713e+04 | NaN | NaN | MCKQDPSVLHTEEMR |
| RNH1 | RINI\_HUMAN | Ribonuclease inhibitor | 191 | 189 | RNH1 | RINI\_HUMAN | Ribonuclease inhibitor | 3C(125.0477) | NaN | X | NaN | NaN | NaN | NaN | 3.919832e+04 | 3.919832e+04 | NaN | NaN | VLCQGLK |
| CD55 | DAF\_HUMAN | Complement decay-accelerating factor | 225 | 222 | CD55 | DAF\_HUMAN | Complement decay-accelerating factor | 4C(125.0477) | NaN | X | NaN | NaN | NaN | NaN | 3.918469e+04 | 3.918469e+04 | NaN | NaN | EIYCPAPPQIDNGIIQGER |
| TPP2 | TPP2\_HUMAN | Tripeptidyl-peptidase 2 | 710 | 709 | TPP2 | TPP2\_HUMAN | Tripeptidyl-peptidase 2 | 2C(125.0477) | NaN | X | NaN | NaN | NaN | NaN | 3.917193e+04 | 3.917193e+04 | NaN | NaN | FCSLPEKGTLTEAFPVLGGK |
| SARS1 | SYSC\_HUMAN | Serine--tRNA ligase, cytoplasmic | 300 | 294 | SARS1 | SYSC\_HUMAN | Serine--tRNA ligase, cytoplasmic | 7C(125.0477) | NaN | X | NaN | NaN | NaN | NaN | 3.910520e+04 | 3.910520e+04 | NaN | NaN | YAGLSTCFR |
| GAPVD1 | GAPD1\_HUMAN | GTPase-activating protein and VPS9 domain-containing protein 1 | 741 | 735 | GAPVD1 | GAPD1\_HUMAN | GTPase-activating protein and VPS9 domain-containing protein 1 | 7C(125.0477) | NaN | X | NaN | NaN | NaN | NaN | 3.910345e+04 | 3.354836e+04 | 5.555091e+03 | NaN | LQELESCSGLGSTSDDTDVR |
| THTPA | THTPA\_HUMAN | Thiamine-triphosphatase | 228 | 217 | THTPA | THTPA\_HUMAN | Thiamine-triphosphatase | 12C(125.0477) | NaN | X | NaN | NaN | NaN | NaN | 3.909074e+04 | 3.909074e+04 | NaN | NaN | ERPQETEDPDHCLG |
| MYO18A | MY18A\_HUMAN | Unconventional myosin-XVIIIa | 1155 | 1148 | MYO18A | MY18A\_HUMAN | Unconventional myosin-XVIIIa | 8C(125.0477) | NaN | X | NaN | NaN | NaN | NaN | 3.907665e+04 | 3.907665e+04 | NaN | NaN | AVEELLECLDLEK |
| TAB1 | TAB1\_HUMAN | TGF-beta-activated kinase 1 and MAP3K7-binding protein 1 | 235 | 230 | TAB1 | TAB1\_HUMAN | TGF-beta-activated kinase 1 and MAP3K7-binding protein 1 | 6C(125.0477) | NaN | X | NaN | NaN | NaN | NaN | 3.905647e+04 | 3.905647e+04 | NaN | NaN | QVGIICGQESTR |
| LPIN2 | LPIN2\_HUMAN | Phosphatidate phosphatase LPIN2 | 790 | 788 | LPIN2 | LPIN2\_HUMAN | Phosphatidate phosphatase LPIN2 | 3C(125.0477) | NaN | X | NaN | NaN | NaN | NaN | 3.904201e+04 | NaN | 3.904201e+04 | NaN | IECLNDIK |
| HBD | HBD\_HUMAN | Hemoglobin subunit delta | 94, 94 | 67 | HBD | HBD\_HUMAN | Hemoglobin subunit delta | 28C(125.0477) 28C(57.0215) | X | X | NaN | NaN | NaN | NaN | 3.899581e+04 | 3.899581e+04 | NaN | NaN | KVLGAFSDGLAHLDNLKGTFSQLSELHCDK |
| HPCAL1 | HPCL1\_HUMAN | Hippocalcin-like protein 1 | 38 | 37 | HPCAL1 | HPCL1\_HUMAN | Hippocalcin-like protein 1 | 2C(125.0477) | NaN | X | NaN | NaN | NaN | NaN | 3.897146e+04 | 3.897146e+04 | NaN | NaN | DCPTGHLTVDEFKK |
| GNB2 | GBB2\_HUMAN | Guanine nucleotide-binding protein G(I)/G(S)/G(T) subunit beta-2 | 25 | 23 | GNB2 | GBB2\_HUMAN | Guanine nucleotide-binding protein G(I)/G(S)/G(T) subunit beta-2 | 3C(15.9949) | NaN | NaN | X | NaN | NaN | NaN | 3.879576e+04 | 3.879576e+04 | NaN | NaN | KACGDSTLTQITAGLDPVGR |
| GPT | ALAT1\_HUMAN | Alanine aminotransferase 1 | 89 | 84 | GPT | ALAT1\_HUMAN | Alanine aminotransferase 1 | 6C(125.0477) | NaN | X | NaN | NaN | NaN | NaN | 3.877507e+04 | 3.877507e+04 | NaN | NaN | QVLALCVNPDLLSSPNFPDDAK |
| GGA3 | GGA3\_HUMAN | ADP-ribosylation factor-binding protein GGA3 | 255 | 250 | GGA3 | GGA3\_HUMAN | ADP-ribosylation factor-binding protein GGA3 | 6C(125.0477) | NaN | X | NaN | NaN | NaN | NaN | 3.875793e+04 | NaN | 3.875793e+04 | NaN | ELFDQCENK |
| AKT1S1 | AKTS1\_HUMAN | Proline-rich AKT1 substrate 1 | 61 | 61 | AKT1S1 | AKTS1\_HUMAN | Proline-rich AKT1 substrate 1 | 1C(125.0477) | NaN | X | NaN | NaN | NaN | NaN | 3.873029e+04 | 3.873029e+04 | NaN | NaN | CLHDIALAHR |
| ANK1 | ANK1\_HUMAN | Ankyrin-1 | 531 | 526 | ANK1 | ANK1\_HUMAN | Ankyrin-1 | 6C(125.0477) | NaN | X | NaN | NaN | NaN | NaN | 3.867558e+04 | 3.867558e+04 | NaN | NaN | EASQACMTKK |
| HSPA8 | HSP7C\_HUMAN | Heat shock cognate 71 kDa protein | 574 | 574 | HSPA8 | HSP7C\_HUMAN | Heat shock cognate 71 kDa protein | 1C(125.0477) | NaN | X | NaN | NaN | NaN | NaN | 3.864169e+04 | 3.864169e+04 | NaN | NaN | CNEIINWLDKNQTAEK |
| ARMC8 | ARMC8\_HUMAN | Armadillo repeat-containing protein 8 | 407 | 407 | ARMC8 | ARMC8\_HUMAN | Armadillo repeat-containing protein 8 | 1C(125.0477) | NaN | X | NaN | NaN | NaN | NaN | 3.862959e+04 | 3.862959e+04 | NaN | NaN | CLHSLSR |
| TRIM21 | RO52\_HUMAN | E3 ubiquitin-protein ligase TRIM21 | 92 | 92 | TRIM21 | RO52\_HUMAN | E3 ubiquitin-protein ligase TRIM21 | 1C(125.0477) | NaN | X | NaN | NaN | NaN | NaN | 3.861252e+04 | 3.861252e+04 | NaN | NaN | CAVHGER |
| HNRNPR | HNRPR\_HUMAN | Heterogeneous nuclear ribonucleoprotein R | 240 | 236 | HNRNPR | HNRPR\_HUMAN | Heterogeneous nuclear ribonucleoprotein R | 5C(125.0477) | NaN | X | NaN | NaN | NaN | NaN | 3.850836e+04 | 3.850836e+04 | NaN | NaN | HLGVCISVANNR |
| IPO9 | IPO9\_HUMAN | Importin-9 | 667 | 662 | IPO9 | IPO9\_HUMAN | Importin-9 | 6C(125.0477) | NaN | X | NaN | NaN | NaN | NaN | 3.850805e+04 | 3.850805e+04 | NaN | NaN | IPAGLCATAIDILTTVVR |
| ALAD | HEM2\_HUMAN | Delta-aminolevulinic acid dehydratase | 162 | 160 | ALAD | HEM2\_HUMAN | Delta-aminolevulinic acid dehydratase | 3C(125.0477) | NaN | X | NaN | NaN | NaN | NaN | 3.849031e+04 | 3.849031e+04 | NaN | NaN | AGCQVVAPSDMMDGRVEAIK |
| GSTM4 | GSTM4\_HUMAN | Glutathione S-transferase Mu 4 | 174 | 169 | GSTM4 | GSTM4\_HUMAN | Glutathione S-transferase Mu 4 | 6C(125.0477) | NaN | X | NaN | NaN | NaN | NaN | 3.844521e+04 | 3.844521e+04 | NaN | NaN | IFEPNCLDAFPNLK |
| RNH1 | RINI\_HUMAN | Ribonuclease inhibitor | 362 | 360 | RNH1 | RINI\_HUMAN | Ribonuclease inhibitor | 3C(125.0477) | NaN | X | NaN | NaN | NaN | NaN | 3.844138e+04 | 2.045463e+04 | 1.798675e+04 | NaN | ELCQGLGQPGSVLR |
| VPS26B | VP26B\_HUMAN | Vacuolar protein sorting-associated protein 26B | 334 | 325 | VPS26B | VP26B\_HUMAN | Vacuolar protein sorting-associated protein 26B | 10C(125.0477) | NaN | X | NaN | NaN | NaN | NaN | 3.837799e+04 | 3.837799e+04 | NaN | NaN | TPSQLSDNNCRQ |
| GSTM3 | GSTM3\_HUMAN | Glutathione S-transferase Mu 3 | 91 | 88 | GSTM3 | GSTM3\_HUMAN | Glutathione S-transferase Mu 3 | 4C(125.0477) | NaN | X | NaN | NaN | NaN | NaN | 3.833441e+04 | 3.833441e+04 | NaN | NaN | HNMCGETEEEK |
| CCRL2 | CCRL2\_HUMAN | C-C chemokine receptor-like 2 | 272 | 265 | CCRL2 | CCRL2\_HUMAN | C-C chemokine receptor-like 2 | 8C(125.0477) | NaN | X | NaN | NaN | NaN | NaN | 3.828327e+04 | NaN | 3.828327e+04 | NaN | EHFSLSDCK |
| CUL2 | CUL2\_HUMAN | Cullin-2 | 721 | 720 | CUL2 | CUL2\_HUMAN | Cullin-2 | 2C(125.0477) | NaN | X | NaN | NaN | NaN | NaN | 3.821799e+04 | 3.821799e+04 | NaN | NaN | KCIEVLIDKQYIER |
| ATP2B1 | AT2B1\_HUMAN | Plasma membrane calcium-transporting ATPase 1 | 60 | 49 | ATP2B1 | AT2B1\_HUMAN | Plasma membrane calcium-transporting ATPase 1 | 12C(125.0477) | NaN | X | NaN | NaN | NaN | NaN | 3.817294e+04 | 3.817294e+04 | NaN | NaN | IQESYGDVYGICTK |
| UBE4B | UBE4B\_HUMAN | Ubiquitin conjugation factor E4 B | 448 | 447 | UBE4B | UBE4B\_HUMAN | Ubiquitin conjugation factor E4 B | 2C(125.0477) | NaN | X | NaN | NaN | NaN | NaN | 3.816983e+04 | 3.816983e+04 | NaN | NaN | MCSQPAVSQLLSNIR |
| DSTN | DEST\_HUMAN | Destrin | 23 | 23 | DSTN | DEST\_HUMAN | Destrin | 1C(125.0477) | NaN | X | NaN | NaN | NaN | NaN | 3.811652e+04 | 3.811652e+04 | NaN | NaN | CSTPEEIK |
| PPIA | PPIA\_HUMAN | Peptidyl-prolyl cis-trans isomerase A | 62, 62 | 56 | PPIA | PPIA\_HUMAN | Peptidyl-prolyl cis-trans isomerase A | 7C(125.0477) 7C(57.0215) | X | X | NaN | NaN | NaN | NaN | 3.811481e+04 | NaN | 3.811481e+04 | NaN | IIPGFMCQGGDFTR |
| CR1 | CR1\_HUMAN | Complement receptor type 1 | 581 | 577 | CR1 | CR1\_HUMAN | Complement receptor type 1 | 5C(125.0477) | NaN | X | NaN | NaN | NaN | NaN | 3.810028e+04 | 2.907588e+04 | 9.024405e+03 | NaN | INYSCTTGHR |
| TCP1 | TCPA\_HUMAN | T-complex protein 1 subunit alpha | 236 | 234 | TCP1 | TCPA\_HUMAN | T-complex protein 1 subunit alpha | 3C(125.0477) | NaN | X | NaN | NaN | NaN | NaN | 3.808991e+04 | 3.808991e+04 | NaN | NaN | IACLDFSLQKTK |
| BRCC3 | BRCC3\_HUMAN | Lys-63-specific deubiquitinase BRCC36 | 240 | 238 | BRCC3 | BRCC3\_HUMAN | Lys-63-specific deubiquitinase BRCC36 | 3C(125.0477) | NaN | X | NaN | NaN | NaN | NaN | 3.807960e+04 | 3.807960e+04 | NaN | NaN | ILCQEEQDAYR |
| G3BP1 | G3BP1\_HUMAN | Ras GTPase-activating protein-binding protein 1 | 73 | 65 | G3BP1 | G3BP1\_HUMAN | Ras GTPase-activating protein-binding protein 1 | 9C(125.0477) | NaN | X | NaN | NaN | NaN | NaN | 3.802202e+04 | 3.802202e+04 | NaN | NaN | VMSQNFTNCHTK |
| ABCC4 | MRP4\_HUMAN | ATP-binding cassette sub-family C member 4 | 1186 | 1183 | ABCC4 | MRP4\_HUMAN | ATP-binding cassette sub-family C member 4 | 4C(125.0477) | NaN | X | NaN | NaN | NaN | NaN | 3.790829e+04 | NaN | 2.630149e+04 | 11606.8010 | QLVCLAR |
| RAB3GAP1 | RB3GP\_HUMAN | Rab3 GTPase-activating protein catalytic subunit | 873 | 870 | RAB3GAP1 | RB3GP\_HUMAN | Rab3 GTPase-activating protein catalytic subunit | 4C(125.0477) | NaN | X | NaN | NaN | NaN | NaN | 3.781466e+04 | 3.781466e+04 | NaN | NaN | FVSCLLEQPEVLVTGAGR |
| CD55 | DAF\_HUMAN | Complement decay-accelerating factor | 253 | 247 | CD55 | DAF\_HUMAN | Complement decay-accelerating factor | 7C(125.0477) | NaN | X | NaN | NaN | NaN | NaN | 3.778348e+04 | NaN | 3.778348e+04 | NaN | QSVTYACNK |
| GNB2 | GBB2\_HUMAN | Guanine nucleotide-binding protein G(I)/G(S)/G(T) subunit beta-2 | 25 | 24 | GNB2 | GBB2\_HUMAN | Guanine nucleotide-binding protein G(I)/G(S)/G(T) subunit beta-2 | 2C(125.0477) | NaN | X | NaN | NaN | NaN | NaN | 3.775007e+04 | 3.775007e+04 | NaN | NaN | ACGDSTLTQITAGLDPVGR |
| SPTA1 | SPTA1\_HUMAN | Spectrin alpha chain, erythrocytic 1 | 799 | 788 | SPTA1 | SPTA1\_HUMAN | Spectrin alpha chain, erythrocytic 1 | 12C(125.0477) | NaN | X | NaN | NaN | NaN | NaN | 3.772637e+04 | 3.772637e+04 | NaN | NaN | KLLDLLHLQLICR |
| PTPRA | PTPRA\_HUMAN | Receptor-type tyrosine-protein phosphatase alpha | 251 | 240 | PTPRA | PTPRA\_HUMAN | Receptor-type tyrosine-protein phosphatase alpha | 12C(125.0477) | NaN | X | NaN | NaN | NaN | NaN | 3.771049e+04 | 3.771049e+04 | NaN | NaN | LFREEFNALPACPIQATCEAASK |
| PARK7 | PARK7\_HUMAN | Parkinson disease protein 7 | 46 | 29 | PARK7 | PARK7\_HUMAN | Parkinson disease protein 7 | 18C(125.0477) | NaN | X | NaN | NaN | NaN | NaN | 3.769029e+04 | 3.769029e+04 | NaN | NaN | AGIKVTVAGLAGKDPVQCSR |
| HSPH1 | HS105\_HUMAN | Heat shock protein 105 kDa | 290 | 276 | HSPH1 | HS105\_HUMAN | Heat shock protein 105 kDa | 15C(125.0477) | NaN | X | NaN | NaN | NaN | NaN | 3.768346e+04 | 3.768346e+04 | NaN | NaN | LMSSNSTDLPLNIECFMNDK |
| KRT33B | KT33B\_HUMAN | Keratin, type I cuticular Ha3-II | 172 | 165 | KRT33B | KT33B\_HUMAN | Keratin, type I cuticular Ha3-II | 8C(31.9988) | NaN | NaN | NaN | X | NaN | NaN | 3.767423e+04 | 3.767423e+04 | NaN | NaN | ILDELTLCRSDLEAQMESLK |
| IPO9 | IPO9\_HUMAN | Importin-9 | 606 | 605 | IPO9 | IPO9\_HUMAN | Importin-9 | 2C(125.0477) | NaN | X | NaN | NaN | NaN | NaN | 3.765566e+04 | 3.765566e+04 | NaN | NaN | ICPFTIAIFLK |
| LDHC | LDHC\_HUMAN | L-lactate dehydrogenase C chain | 131 | 119 | LDHC | LDHC\_HUMAN | L-lactate dehydrogenase C chain | 13C(125.0477) | NaN | X | NaN | NaN | NaN | NaN | 3.764819e+04 | 3.764819e+04 | NaN | NaN | SIIPAIVHYSPDCK |
| LCP1 | PLSL\_HUMAN | Plastin-2 | 101 | 98 | LCP1 | PLSL\_HUMAN | Plastin-2 | 4C(125.0477) | NaN | X | NaN | NaN | NaN | NaN | 3.756278e+04 | 3.756278e+04 | NaN | NaN | EGICAIGGTSEQSSVGTQHSYSEEEK |
| SIRT2 | SIR2\_HUMAN | NAD-dependent protein deacetylase sirtuin-2 | 148 | 137 | SIRT2 | SIR2\_HUMAN | NAD-dependent protein deacetylase sirtuin-2 | 12C(125.0477) | NaN | X | NaN | NaN | NaN | NaN | 3.754243e+04 | 3.754243e+04 | NaN | NaN | ELYPGQFKPTICHYFMR |
| PPP1CB | PP1B\_HUMAN | Serine/threonine-protein phosphatase PP1-beta catalytic subunit | 244 | 234 | PPP1CB | PP1B\_HUMAN | Serine/threonine-protein phosphatase PP1-beta catalytic subunit | 11C(125.0477) | NaN | X | NaN | NaN | NaN | NaN | 3.753885e+04 | 3.753885e+04 | NaN | NaN | FLNRHDLDLICR |
| PSMB3 | PSB3\_HUMAN | Proteasome subunit beta type-3 | 19 | 16 | PSMB3 | PSB3\_HUMAN | Proteasome subunit beta type-3 | 4C(125.0477) | NaN | X | NaN | NaN | NaN | NaN | 3.751507e+04 | 3.751507e+04 | NaN | NaN | GKNCVAIAADRR |
| SHMT2 | GLYM\_HUMAN | Serine hydroxymethyltransferase, mitochondrial | 91 | 83 | SHMT2 | GLYM\_HUMAN | Serine hydroxymethyltransferase, mitochondrial | 9C(15.9949) | NaN | NaN | X | NaN | NaN | NaN | 3.744298e+04 | NaN | 3.744298e+04 | NaN | AALEALGSCLNNK |
| TRIM65 | TRI65\_HUMAN | Tripartite motif-containing protein 65 | 500 | 496 | TRIM65 | TRI65\_HUMAN | Tripartite motif-containing protein 65 | 5C(125.0477) | NaN | X | NaN | NaN | NaN | NaN | 3.743350e+04 | 3.743350e+04 | NaN | NaN | TLTLCHQPGAVFPLGPQEEVLS |
| PRKDC | PRKDC\_HUMAN | DNA-dependent protein kinase catalytic subunit | 795 | 791 | PRKDC | PRKDC\_HUMAN | DNA-dependent protein kinase catalytic subunit | 5C(125.0477) | NaN | X | NaN | NaN | NaN | NaN | 3.738734e+04 | 3.738734e+04 | NaN | NaN | DILPCLDGYLK |
| TP53I3 | QORX\_HUMAN | Quinone oxidoreductase PIG3 | 216 | 207 | TP53I3 | QORX\_HUMAN | Quinone oxidoreductase PIG3 | 10C(125.0477) | NaN | X | NaN | NaN | NaN | NaN | 3.734681e+04 | 3.734681e+04 | NaN | NaN | GAGVNLILDCIGGSYWEK |
| DCTN1 | DCTN1\_HUMAN | Dynactin subunit 1 | 677 | 670 | DCTN1 | DCTN1\_HUMAN | Dynactin subunit 1 | 8C(125.0477) | NaN | X | NaN | NaN | NaN | NaN | 3.727648e+04 | 3.727648e+04 | NaN | NaN | YEHALSQCSVDVYKK |
| UBR4 | UBR4\_HUMAN | E3 ubiquitin-protein ligase UBR4 | 4272 | 4262 | UBR4 | UBR4\_HUMAN | E3 ubiquitin-protein ligase UBR4 | 11C(125.0477) | NaN | X | NaN | NaN | NaN | NaN | 3.723527e+04 | 3.723527e+04 | NaN | NaN | LVGTVLNGYLCLR |
| CDK6 | CDK6\_HUMAN | Cyclin-dependent kinase 6 | 7 | 4 | CDK6 | CDK6\_HUMAN | Cyclin-dependent kinase 6 | 4C(15.9949) | NaN | NaN | X | NaN | NaN | NaN | 3.722556e+04 | 3.722556e+04 | NaN | NaN | DGLCRADQQYECVAEIGEGAYGK |
| HSP90AA1 | HS90A\_HUMAN | Heat shock protein HSP 90-alpha | 529 | 513 | HSP90AA1 | HS90A\_HUMAN | Heat shock protein HSP 90-alpha | 17C(125.0477) | NaN | X | NaN | NaN | NaN | NaN | 3.716175e+04 | 3.716175e+04 | NaN | NaN | KHGLEVIYMIEPIDEYCVQQLK |
| ALB | ALBU\_HUMAN | Albumin | 416 | 414 | ALB | ALBU\_HUMAN | Albumin | 3C(57.0215) | X | NaN | NaN | NaN | NaN | NaN | 3.713627e+04 | NaN | 3.713627e+04 | NaN | QNCELFEQLGEYK |
| TATDN1 | TATD1\_HUMAN | Deoxyribonuclease TATDN1 | 253 | 247 | TATDN1 | TATD1\_HUMAN | Deoxyribonuclease TATDN1 | 7C(125.0477) | NaN | X | NaN | NaN | NaN | NaN | 3.712628e+04 | 3.712628e+04 | NaN | NaN | KWESGHCLK |
| THTPA | THTPA\_HUMAN | Thiamine-triphosphatase | 66 | 66 | THTPA | THTPA\_HUMAN | Thiamine-triphosphatase | 1C(125.0477) | NaN | X | NaN | NaN | NaN | NaN | 3.710426e+04 | 3.710426e+04 | NaN | NaN | CPGAAGVLGPHTEYK |
| AGO2 | AGO2\_HUMAN | Protein argonaute-2 | 327 | 322 | AGO2 | AGO2\_HUMAN | Protein argonaute-2 | 6C(125.0477) | NaN | X | NaN | NaN | NaN | NaN | 3.709695e+04 | 3.709695e+04 | NaN | NaN | YPHLPCLQVGQEQK |
| SIRT2 | SIR2\_HUMAN | NAD-dependent protein deacetylase sirtuin-2 | 164 | 164 | SIRT2 | SIR2\_HUMAN | NAD-dependent protein deacetylase sirtuin-2 | 1C(125.0477) | NaN | X | NaN | NaN | NaN | NaN | 3.695343e+04 | NaN | 3.695343e+04 | NaN | CYTQNIDTLER |
| EIF6 | IF6\_HUMAN | Eukaryotic translation initiation factor 6 | 152 | 140 | EIF6 | IF6\_HUMAN | Eukaryotic translation initiation factor 6 | 13C(125.0477) | NaN | X | NaN | NaN | NaN | NaN | 3.694602e+04 | 3.694602e+04 | NaN | NaN | QTVADQVLVGSYCVFSNQGGLVHPK |
| FADD | FADD\_HUMAN | FAS-associated death domain protein | 168 | 167 | FADD | FADD\_HUMAN | FAS-associated death domain protein | 2C(125.0477) | NaN | X | NaN | NaN | NaN | NaN | 3.691904e+04 | 3.691904e+04 | NaN | NaN | SCQMNLVADLVQEVQQAR |
| DSG1 | DSG1\_HUMAN | Desmoglein-1 | 77 | 73 | DSG1 | DSG1\_HUMAN | Desmoglein-1 | 5C(125.0477) | NaN | X | NaN | NaN | NaN | NaN | 3.686752e+04 | 3.686752e+04 | NaN | NaN | IHSDCAANQQVTYR |
| PPP2R1A | 2AAA\_HUMAN | Serine/threonine-protein phosphatase 2A 65 kDa regulatory subunit A alpha isoform | 377 | 359 | PPP2R1A | 2AAA\_HUMAN | Serine/threonine-protein phosphatase 2A 65 kDa regulatory subunit A alpha isoform | 19C(15.9949) | NaN | NaN | X | NaN | NaN | NaN | 3.684648e+04 | 3.684648e+04 | NaN | NaN | DNTIEHLLPLFLAQLKDECPEVRLNIISNLDCVNEVIGIR |
| CLTC | CLH1\_HUMAN | Clathrin heavy chain 1 | 151 | 145 | CLTC | CLH1\_HUMAN | Clathrin heavy chain 1 | 7C(125.0477) | NaN | X | NaN | NaN | NaN | NaN | 3.684336e+04 | 3.684336e+04 | NaN | NaN | HSSLAGCQIINYRTDAK |
| UBR4 | UBR4\_HUMAN | E3 ubiquitin-protein ligase UBR4 | 1698 | 1697 | UBR4 | UBR4\_HUMAN | E3 ubiquitin-protein ligase UBR4 | 2C(125.0477) | NaN | X | NaN | NaN | NaN | NaN | 3.681372e+04 | 3.681372e+04 | NaN | NaN | VCHKDHEISYAK |
| STRAP | STRAP\_HUMAN | Serine-threonine kinase receptor-associated protein | 305 | 305 | STRAP | STRAP\_HUMAN | Serine-threonine kinase receptor-associated protein | 1C(125.0477) | NaN | X | NaN | NaN | NaN | NaN | 3.679246e+04 | 3.679246e+04 | NaN | NaN | CVLPEEDSGELAKPK |
| IGHM | IGHM\_HUMAN | Immunoglobulin heavy constant mu | 134 | 132 | IGHM | IGHM\_HUMAN | Immunoglobulin heavy constant mu | 3C(57.0215) | X | NaN | NaN | NaN | NaN | NaN | 3.675356e+04 | NaN | NaN | 36753.5600 | LICQATGFSPR |
| C1QC | C1QC\_HUMAN | Complement C1q subcomponent subunit C | 179 | 162 | C1QC | C1QC\_HUMAN | Complement C1q subcomponent subunit C | 18C(57.0215) | X | NaN | NaN | NaN | NaN | NaN | 3.669996e+04 | 3.669996e+04 | NaN | NaN | VPGLYYFVYHASHTANLCVLLYR |
| HGH1 | HGH1\_HUMAN | Protein HGH1 homolog | 138 | 134 | HGH1 | HGH1\_HUMAN | Protein HGH1 homolog | 5C(125.0477) | NaN | X | NaN | NaN | NaN | NaN | 3.654555e+04 | 3.654555e+04 | NaN | NaN | EPAPCAALMAALAAAEPADSGLER |
| HUWE1 | HUWE1\_HUMAN | E3 ubiquitin-protein ligase HUWE1 | 1278 | 1269 | HUWE1 | HUWE1\_HUMAN | E3 ubiquitin-protein ligase HUWE1 | 10C(125.0477) | NaN | X | NaN | NaN | NaN | NaN | 3.648716e+04 | 3.648716e+04 | NaN | NaN | MAESMLAILCHILR |
| SERINC3 | SERC3\_HUMAN | Serine incorporator 3 | 89, 89 | 88 | SERINC3 | SERC3\_HUMAN | Serine incorporator 3 | 2C(125.0477) 2C(57.0215) | X | X | NaN | NaN | NaN | NaN | 3.645029e+04 | NaN | 3.645029e+04 | NaN | DCDVLVGYK |
| IPO11 | IPO11\_HUMAN | Importin-11 | 252 | 246 | IPO11 | IPO11\_HUMAN | Importin-11 | 7C(125.0477) | NaN | X | NaN | NaN | NaN | NaN | 3.643191e+04 | 3.643191e+04 | NaN | NaN | LKQFLECSR |
| SKIV2L | SKIV2\_HUMAN | Helicase SKI2W | 877 | 870 | SKIV2L | SKIV2\_HUMAN | Helicase SKI2W | 8C(125.0477) | NaN | X | NaN | NaN | NaN | NaN | 3.642085e+04 | 3.642085e+04 | NaN | NaN | VFTTLVLCDKPLSQDPQDR |
| XPO7 | XPO7\_HUMAN | Exportin-7 | 606 | 606 | XPO7 | XPO7\_HUMAN | Exportin-7 | 1C(125.0477) | NaN | X | NaN | NaN | NaN | NaN | 3.635064e+04 | 3.635064e+04 | NaN | NaN | CEPITSK |
| GSK3A | GSK3A\_HUMAN | Glycogen synthase kinase-3 alpha | 170 | 166 | GSK3A | GSK3A\_HUMAN | Glycogen synthase kinase-3 alpha | 5C(125.0477) | NaN | X | NaN | NaN | NaN | NaN | 3.634987e+04 | 3.634987e+04 | NaN | NaN | KLDHCNIVR |
| A2M | A2MG\_HUMAN | Alpha-2-macroglobulin | 1321 | 1316 | A2M | A2MG\_HUMAN | Alpha-2-macroglobulin | 6C(57.0215) | X | NaN | NaN | NaN | NaN | NaN | 3.633843e+04 | 3.633843e+04 | NaN | NaN | VTGEGCVYLQTSLK |
| CCT7 | TCPH\_HUMAN | T-complex protein 1 subunit eta | 364 | 358 | CCT7 | TCPH\_HUMAN | T-complex protein 1 subunit eta | 7C(125.0477) | NaN | X | NaN | NaN | NaN | NaN | 3.623925e+04 | 1.631050e+04 | 1.992875e+04 | NaN | YNFFTGCPK |
| WDR77 | MEP50\_HUMAN | Methylosome protein 50 | 186 | 180 | WDR77 | MEP50\_HUMAN | Methylosome protein 50 | 7C(125.0477) | NaN | X | NaN | NaN | NaN | NaN | 3.621715e+04 | NaN | 3.621715e+04 | NaN | DSVFLSCSEDNR |
| PSMB3 | PSB3\_HUMAN | Proteasome subunit beta type-3 | 19 | 18 | PSMB3 | PSB3\_HUMAN | Proteasome subunit beta type-3 | 2C(125.0477) | NaN | X | NaN | NaN | NaN | NaN | 3.616638e+04 | NaN | 3.616638e+04 | NaN | NCVAIAADR |
| UBR4 | UBR4\_HUMAN | E3 ubiquitin-protein ligase UBR4 | 3392 | 3378 | UBR4 | UBR4\_HUMAN | E3 ubiquitin-protein ligase UBR4 | 15C(125.0477) | NaN | X | NaN | NaN | NaN | NaN | 3.616352e+04 | 3.616352e+04 | NaN | NaN | EKDGETSGSQEDQLCTALVNQLNK |
| VTA1 | VTA1\_HUMAN | Vacuolar protein sorting-associated protein VTA1 homolog | 56 | 49 | VTA1 | VTA1\_HUMAN | Vacuolar protein sorting-associated protein VTA1 homolog | 8C(125.0477) | NaN | X | NaN | NaN | NaN | NaN | 3.614912e+04 | 3.614912e+04 | NaN | NaN | IDSKTPECR |
| ATL3 | ATLA3\_HUMAN | Atlastin-3 | 429 | 423 | ATL3 | ATLA3\_HUMAN | Atlastin-3 | 7C(125.0477) | NaN | X | NaN | NaN | NaN | NaN | 3.608785e+04 | NaN | 3.608785e+04 | NaN | ELYENFCK |
| DSC1 | DSC1\_HUMAN | Desmocollin-1 | 255 | 246 | DSC1 | DSC1\_HUMAN | Desmocollin-1 | 10C(125.0477) | NaN | X | NaN | NaN | NaN | NaN | 3.597869e+04 | NaN | 3.597869e+04 | NaN | VTIFTVPENCR |
| NUP37 | NUP37\_HUMAN | Nucleoporin Nup37 | 149 | 136 | NUP37 | NUP37\_HUMAN | Nucleoporin Nup37 | 14C(125.0477) | NaN | X | NaN | NaN | NaN | NaN | 3.589561e+04 | 3.589561e+04 | NaN | NaN | EGQEIASVSDDHTCR |
| TNPO3 | TNPO3\_HUMAN | Transportin-3 | 634 | 621 | TNPO3 | TNPO3\_HUMAN | Transportin-3 | 14C(125.0477) | NaN | X | NaN | NaN | NaN | NaN | 3.587881e+04 | 3.587881e+04 | NaN | NaN | HTNPIVENGQTHPCQK |
| GPHN | GEPH\_HUMAN | Gephyrin | 212 | 209 | GPHN | GEPH\_HUMAN | Gephyrin | 4C(125.0477) | NaN | X | NaN | NaN | NaN | NaN | 3.584715e+04 | 3.584715e+04 | NaN | NaN | GVQCEEEEEEK |
| METAP2 | MAP2\_HUMAN | Methionine aminopeptidase 2 | 416 | 401 | METAP2 | MAP2\_HUMAN | Methionine aminopeptidase 2 | 16C(125.0477) | NaN | X | NaN | NaN | NaN | NaN | 3.575046e+04 | 3.575046e+04 | NaN | NaN | HLLNVINENFGTLAFCR |
| ZZEF1 | ZZEF1\_HUMAN | Zinc finger ZZ-type and EF-hand domain-containing protein 1 | 612 | 611 | ZZEF1 | ZZEF1\_HUMAN | Zinc finger ZZ-type and EF-hand domain-containing protein 1 | 2C(125.0477) | NaN | X | NaN | NaN | NaN | NaN | 3.567080e+04 | 3.567080e+04 | NaN | NaN | FCAEEHFK |
| RELCH | RELCH\_HUMAN | RAB11-binding protein RELCH | 526 | 526 | RELCH | RELCH\_HUMAN | RAB11-binding protein RELCH | 1C(125.0477) | NaN | X | NaN | NaN | NaN | NaN | 3.565997e+04 | 3.565997e+04 | NaN | NaN | CLPHIVPNVLLAK |
| LTA4H | LKHA4\_HUMAN | Leukotriene A-4 hydrolase | 147 | 143 | LTA4H | LKHA4\_HUMAN | Leukotriene A-4 hydrolase | 5C(125.0477) | NaN | X | NaN | NaN | NaN | NaN | 3.562368e+04 | NaN | 3.562368e+04 | NaN | AILPCQDTPSVK |
| SPAG9 | JIP4\_HUMAN | C-Jun-amino-terminal kinase-interacting protein 4 | 443 | 438 | SPAG9 | JIP4\_HUMAN | C-Jun-amino-terminal kinase-interacting protein 4 | 6C(125.0477) | NaN | X | NaN | NaN | NaN | NaN | 3.556453e+04 | 3.556453e+04 | NaN | NaN | VDELTCEK |
| BMP2K | BMP2K\_HUMAN | BMP-2-inducible protein kinase | 292 | 289 | BMP2K | BMP2K\_HUMAN | BMP-2-inducible protein kinase | 4C(125.0477) | NaN | X | NaN | NaN | NaN | NaN | 3.553750e+04 | 3.553750e+04 | NaN | NaN | NIHCLIR |
| CAND1 | CAND1\_HUMAN | Cullin-associated NEDD8-dissociated protein 1 | 1153 | 1145 | CAND1 | CAND1\_HUMAN | Cullin-associated NEDD8-dissociated protein 1 | 9C(125.0477) | NaN | X | NaN | NaN | NaN | NaN | 3.542602e+04 | 3.542602e+04 | NaN | NaN | LVEPLRATCTTK |
| SLC31A1 | COPT1\_HUMAN | High affinity copper uptake protein 1 | 189 | 180 | SLC31A1 | COPT1\_HUMAN | High affinity copper uptake protein 1 | 10C(125.0477) | NaN | X | NaN | NaN | NaN | NaN | 3.540682e+04 | NaN | 3.540682e+04 | NaN | AVVVDITEHCH |
| CLCN3 | CLCN3\_HUMAN | H(+)/Cl(-) exchange transporter 3 | 237 | 231 | CLCN3 | CLCN3\_HUMAN | H(+)/Cl(-) exchange transporter 3 | 7C(125.0477) | NaN | X | NaN | NaN | NaN | NaN | 3.534571e+04 | 3.534571e+04 | NaN | NaN | VFAPYACGSGIPEIK |
| TKFC | TKFC\_HUMAN | Triokinase/FMN cyclase | 382, 382 | 381 | TKFC | TKFC\_HUMAN | Triokinase/FMN cyclase | 2C(125.0477) 2C(15.9949) | NaN | X | X | NaN | NaN | NaN | 3.533208e+04 | 3.533208e+04 | NaN | NaN | VCSTLLGLEEHLNALDR |
| USP24 | UBP24\_HUMAN | Ubiquitin carboxyl-terminal hydrolase 24 | 2160 | 2155 | USP24 | UBP24\_HUMAN | Ubiquitin carboxyl-terminal hydrolase 24 | 6C(125.0477) | NaN | X | NaN | NaN | NaN | NaN | 3.530535e+04 | 3.530535e+04 | NaN | NaN | HPYYPCMAK |
| HSPA4 | HSP74\_HUMAN | Heat shock 70 kDa protein 4 | 245 | 236 | HSPA4 | HSP74\_HUMAN | Heat shock 70 kDa protein 4 | 10C(125.0477) | NaN | X | NaN | NaN | NaN | NaN | 3.529308e+04 | 3.529308e+04 | NaN | NaN | FDEVLVNHFCEEFGK |
| ATP6V0D1 | VA0D1\_HUMAN | V-type proton ATPase subunit d 1 | 326 | 321 | ATP6V0D1 | VA0D1\_HUMAN | V-type proton ATPase subunit d 1 | 6C(125.0477) | NaN | X | NaN | NaN | NaN | NaN | 3.512122e+04 | NaN | 3.512122e+04 | NaN | LKEQECR |
| MTOR | MTOR\_HUMAN | Serine/threonine-protein kinase mTOR | 606 | 605 | MTOR | MTOR\_HUMAN | Serine/threonine-protein kinase mTOR | 2C(125.0477) | NaN | X | NaN | NaN | NaN | NaN | 3.511331e+04 | 3.511331e+04 | NaN | NaN | HCADHFLNSEHK |
| TUBB4B | TBB4B\_HUMAN | Tubulin beta-4B chain | 239 | 217 | TUBB4B | TBB4B\_HUMAN | Tubulin beta-4B chain | 23C(57.0215) | X | NaN | NaN | NaN | NaN | NaN | 3.510234e+04 | 3.510234e+04 | NaN | NaN | LTTPTYGDLNHLVSATMSGVTTCLR |
| SLFN14 | SLN14\_HUMAN | Protein SLFN14 | 431 | 426 | SLFN14 | SLN14\_HUMAN | Protein SLFN14 | 6C(125.0477) | NaN | X | NaN | NaN | NaN | NaN | 3.507775e+04 | 3.507775e+04 | NaN | NaN | TLIHPCSQGIVIFSR |
| SPTA1 | SPTA1\_HUMAN | Spectrin alpha chain, erythrocytic 1 | 2388 | 2378 | SPTA1 | SPTA1\_HUMAN | Spectrin alpha chain, erythrocytic 1 | 11C(31.9988) | NaN | NaN | NaN | X | NaN | NaN | 3.500116e+04 | NaN | 3.500116e+04 | NaN | QALTPEQVSFCATHMQQYMDPR |
| ADSS2 | PURA2\_HUMAN | Adenylosuccinate synthetase isozyme 2 | 58 | 46 | ADSS2 | PURA2\_HUMAN | Adenylosuccinate synthetase isozyme 2 | 13C(125.0477) | NaN | X | NaN | NaN | NaN | NaN | 3.499716e+04 | 3.499716e+04 | NaN | NaN | VVDLLAQDADIVCR |
| AP1G1 | AP1G1\_HUMAN | AP-1 complex subunit gamma-1 | 392 | 384 | AP1G1 | AP1G1\_HUMAN | AP-1 complex subunit gamma-1 | 9C(125.0477) | NaN | X | NaN | NaN | NaN | NaN | 3.499414e+04 | 3.499414e+04 | NaN | NaN | ELLYFLDSCEPEFK |
| SPTB | SPTB1\_HUMAN | Spectrin beta chain, erythrocytic | 735 | 718 | SPTB | SPTB1\_HUMAN | Spectrin beta chain, erythrocytic | 18C(125.0477) | NaN | X | NaN | NaN | NaN | NaN | 3.497416e+04 | 3.497416e+04 | NaN | NaN | IKEVSAQWDQLKDLAAFCK |
| EML2 | EMAL2\_HUMAN | Echinoderm microtubule-associated protein-like 2 | 110 | 110 | EML2 | EMAL2\_HUMAN | Echinoderm microtubule-associated protein-like 2 | 1C(125.0477) | NaN | X | NaN | NaN | NaN | NaN | 3.488088e+04 | 3.488088e+04 | NaN | NaN | CLAIHPDMVTIATGQVAGTTK |
| ERLIN2 | ERLN2\_HUMAN | Erlin-2 | 262 | 258 | ERLIN2 | ERLN2\_HUMAN | Erlin-2 | 5C(125.0477) | NaN | X | NaN | NaN | NaN | NaN | 3.486944e+04 | NaN | 3.486944e+04 | NaN | ADAECYTAMK |
| MPP1 | EM55\_HUMAN | 55 kDa erythrocyte membrane protein | 94 | 88 | MPP1 | EM55\_HUMAN | 55 kDa erythrocyte membrane protein | 7C(125.0477) | NaN | X | NaN | NaN | NaN | NaN | 3.486756e+04 | 3.486756e+04 | NaN | NaN | LNEKQSCTVAR |
| ATP6V1G1 | VATG1\_HUMAN | V-type proton ATPase subunit G 1 | 69 | 59 | ATP6V1G1 | VATG1\_HUMAN | V-type proton ATPase subunit G 1 | 11C(125.0477) | NaN | X | NaN | NaN | NaN | NaN | 3.486634e+04 | 3.486634e+04 | NaN | NaN | EAAALGSRGSCSTEVEKETQEK |
| BPGM | PMGE\_HUMAN | Bisphosphoglycerate mutase | 145 | 144 | BPGM | PMGE\_HUMAN | Bisphosphoglycerate mutase | 2C(31.9988) | NaN | NaN | NaN | X | NaN | NaN | 3.482859e+04 | NaN | NaN | 34828.5860 | VCDVPLDQLPR |
| CISD2 | CISD2\_HUMAN | CDGSH iron-sulfur domain-containing protein 2 | 110 | 106 | CISD2 | CISD2\_HUMAN | CDGSH iron-sulfur domain-containing protein 2 | 5C(125.0477) | NaN | X | NaN | NaN | NaN | NaN | 3.482063e+04 | 3.482063e+04 | NaN | NaN | TFPACDGSHNK |
| ACTN4 | ACTN4\_HUMAN | Alpha-actinin-4 | 351 | 351 | ACTN4 | ACTN4\_HUMAN | Alpha-actinin-4 | 1C(125.0477) | NaN | X | NaN | NaN | NaN | NaN | 3.481302e+04 | 3.481302e+04 | NaN | NaN | CQLEINFNTLQTK |
| NAPRT | PNCB\_HUMAN | Nicotinate phosphoribosyltransferase | 254 | 246 | NAPRT | PNCB\_HUMAN | Nicotinate phosphoribosyltransferase | 9C(125.0477) | NaN | X | NaN | NaN | NaN | NaN | 3.479565e+04 | 3.479565e+04 | NaN | NaN | AQVWLEQVCAHLGLGVQEPHPGER |
| HSPH1 | HS105\_HUMAN | Heat shock protein 105 kDa | 34 | 34 | HSPH1 | HS105\_HUMAN | Heat shock protein 105 kDa | 1C(125.0477) | NaN | X | NaN | NaN | NaN | NaN | 3.478657e+04 | 3.478657e+04 | NaN | NaN | CTPSVISFGSK |
| CCT6A | TCPZ\_HUMAN | T-complex protein 1 subunit zeta | 282, 282 | 280 | CCT6A | TCPZ\_HUMAN | T-complex protein 1 subunit zeta | 3C(125.0477) 3C(15.9949) | NaN | X | X | NaN | NaN | NaN | 3.477669e+04 | 3.477669e+04 | NaN | NaN | KVCGDSDKGFVVINQK |
| WDR44 | WDR44\_HUMAN | WD repeat-containing protein 44 | 581 | 574 | WDR44 | WDR44\_HUMAN | WD repeat-containing protein 44 | 8C(125.0477) | NaN | X | NaN | NaN | NaN | NaN | 3.474718e+04 | 3.474718e+04 | NaN | NaN | SDTDTGVCSGTDEDPDDK |
| ERI3 | ERI3\_HUMAN | ERI1 exoribonuclease 3 | 313 | 297 | ERI3 | ERI3\_HUMAN | ERI1 exoribonuclease 3 | 17C(125.0477) | NaN | X | NaN | NaN | NaN | NaN | 3.474306e+04 | 3.474306e+04 | NaN | NaN | GLSLQHIGRPHSGIDDCK |
| ALB | ALBU\_HUMAN | Albumin | 340 | 338 | ALB | ALBU\_HUMAN | Albumin | 3C(57.0215) | X | NaN | NaN | NaN | NaN | NaN | 3.468996e+04 | 2.251735e+04 | 1.217261e+04 | NaN | DVCKNYAEAK |
| OTULIN | OTUL\_HUMAN | Ubiquitin thioesterase otulin | 92 | 80 | OTULIN | OTUL\_HUMAN | Ubiquitin thioesterase otulin | 13C(125.0477) | NaN | X | NaN | NaN | NaN | NaN | 3.458579e+04 | 3.458579e+04 | NaN | NaN | LSVAPEMDIMDYCK |
| TECPR1 | TCPR1\_HUMAN | Tectonin beta-propeller repeat-containing protein 1 | 462 | 453 | TECPR1 | TCPR1\_HUMAN | Tectonin beta-propeller repeat-containing protein 1 | 10C(125.0477) | NaN | X | NaN | NaN | NaN | NaN | 3.456893e+04 | NaN | 3.456893e+04 | NaN | TAEDTVEDACPAEGSR |
| PCBP1 | PCBP1\_HUMAN | Poly(rC)-binding protein 1 | 158 | 145 | PCBP1 | PCBP1\_HUMAN | Poly(rC)-binding protein 1 | 14C(125.0477) | NaN | X | NaN | NaN | NaN | NaN | 3.454973e+04 | NaN | 3.454973e+04 | NaN | AITIAGVPQSVTECVK |
| FLG2 | FILA2\_HUMAN | Filaggrin-2 | 366 | 359 | FLG2 | FILA2\_HUMAN | Filaggrin-2 | 8C(125.0477) | NaN | X | NaN | NaN | NaN | NaN | 3.452387e+04 | NaN | 3.452387e+04 | NaN | ENGQPQNCGGQWR |
| AMY1B | AMY1B\_HUMAN | Alpha-amylase 1B | 101 | 101 | AMY1B | AMY1B\_HUMAN | Alpha-amylase 1B | 1C(125.0477) | NaN | X | NaN | NaN | NaN | NaN | 3.450141e+04 | NaN | 3.450141e+04 | NaN | CNNVGVR |
| PSMC6 | PRS10\_HUMAN | 26S proteasome regulatory subunit 10B | 228 | 224 | PSMC6 | PRS10\_HUMAN | 26S proteasome regulatory subunit 10B | 5C(125.0477) | NaN | X | NaN | NaN | NaN | NaN | 3.447264e+04 | 3.447264e+04 | NaN | NaN | DHQPCIIFMDEIDAIGGRR |
| ZZEF1 | ZZEF1\_HUMAN | Zinc finger ZZ-type and EF-hand domain-containing protein 1 | 2229 | 2220 | ZZEF1 | ZZEF1\_HUMAN | Zinc finger ZZ-type and EF-hand domain-containing protein 1 | 10C(125.0477) | NaN | X | NaN | NaN | NaN | NaN | 3.439384e+04 | 3.439384e+04 | NaN | NaN | DVIATFTDHCIK |
| CSE1L | XPO2\_HUMAN | Exportin-2 | 853 | 849 | CSE1L | XPO2\_HUMAN | Exportin-2 | 5C(125.0477) | NaN | X | NaN | NaN | NaN | NaN | 3.438856e+04 | 3.438856e+04 | NaN | NaN | LLTECPPMMDTEYTK |
| WDR1 | WDR1\_HUMAN | WD repeat-containing protein 1 | 438 | 438 | WDR1 | WDR1\_HUMAN | WD repeat-containing protein 1 | 1C(125.0477) | NaN | X | NaN | NaN | NaN | NaN | 3.438708e+04 | 3.438708e+04 | NaN | NaN | CFSIDNPGYEPEVVAVHPGGDTVAIGGVDGNVR |
| HUWE1 | HUWE1\_HUMAN | E3 ubiquitin-protein ligase HUWE1 | 1401 | 1393 | HUWE1 | HUWE1\_HUMAN | E3 ubiquitin-protein ligase HUWE1 | 9C(125.0477) | NaN | X | NaN | NaN | NaN | NaN | 3.437832e+04 | 3.437832e+04 | NaN | NaN | AESPEEVACR |
| MAN2C1 | MA2C1\_HUMAN | Alpha-mannosidase 2C1 | 40 | 35 | MAN2C1 | MA2C1\_HUMAN | Alpha-mannosidase 2C1 | 6C(125.0477) | NaN | X | NaN | NaN | NaN | NaN | 3.434782e+04 | 3.434782e+04 | NaN | NaN | LFGASCPVAVLSSFLTPER |
| ABRAXAS2 | ABRX2\_HUMAN | BRISC complex subunit Abraxas 2 | 237 | 233 | ABRAXAS2 | ABRX2\_HUMAN | BRISC complex subunit Abraxas 2 | 5C(31.9988) | NaN | NaN | NaN | X | NaN | NaN | 3.431184e+04 | 3.431184e+04 | NaN | NaN | VVESCQAEVNK |
| MYH9 | MYH9\_HUMAN | Myosin-9 | 569 | 566 | MYH9 | MYH9\_HUMAN | Myosin-9 | 4C(125.0477) | NaN | X | NaN | NaN | NaN | NaN | 3.427447e+04 | 3.427447e+04 | NaN | NaN | ADFCIIHYAGK |
| CASP9 | CASP9\_HUMAN | Caspase-9 | 403 | 399 | CASP9 | CASP9\_HUMAN | Caspase-9 | 5C(125.0477) | NaN | X | NaN | NaN | NaN | NaN | 3.420488e+04 | 3.420488e+04 | NaN | NaN | QMPGCFNFLR |
| TKT | TKT\_HUMAN | Transketolase | 386 | 382 | TKT | TKT\_HUMAN | Transketolase | 5C(125.0477) | NaN | X | NaN | NaN | NaN | NaN | 3.412973e+04 | 3.412973e+04 | NaN | NaN | TVPFCSTFAAFFTR |
| PSMD10 | PSD10\_HUMAN | 26S proteasome non-ATPase regulatory subunit 10 | 180 | 163 | PSMD10 | PSD10\_HUMAN | 26S proteasome non-ATPase regulatory subunit 10 | 18C(125.0477) | NaN | X | NaN | NaN | NaN | NaN | 3.410876e+04 | 3.410876e+04 | NaN | NaN | ASTNIQDTEGNTPLHLACDEERVEEAK |
| KPNA6 | IMA7\_HUMAN | Importin subunit alpha-7 | 467 | 459 | KPNA6 | IMA7\_HUMAN | Importin subunit alpha-7 | 9C(125.0477) | NaN | X | NaN | NaN | NaN | NaN | 3.409162e+04 | 3.409162e+04 | NaN | NaN | SGSGVNPYCGLIEEAYGLDKIEFLQSHENQEIYQK |
| CUL3 | CUL3\_HUMAN | Cullin-3 | 187 | 185 | CUL3 | CUL3\_HUMAN | Cullin-3 | 3C(125.0477) | NaN | X | NaN | NaN | NaN | NaN | 3.407685e+04 | 3.407685e+04 | NaN | NaN | NACQMLMILGLEGR |
| PDXP | PLPP\_HUMAN | Chronophin | 171 | 168 | PDXP | PLPP\_HUMAN | Chronophin | 4C(125.0477) | NaN | X | NaN | NaN | NaN | NaN | 3.406618e+04 | NaN | 3.406618e+04 | NaN | DPECLLVATDR |
| NGLY1 | NGLY1\_HUMAN | Peptide-N(4)-(N-acetyl-beta-glucosaminyl)asparagine amidase | 197 | 194 | NGLY1 | NGLY1\_HUMAN | Peptide-N(4)-(N-acetyl-beta-glucosaminyl)asparagine amidase | 4C(125.0477) | NaN | X | NaN | NaN | NaN | NaN | 3.401071e+04 | 3.401071e+04 | NaN | NaN | ALACIPVQELKR |
| SEC24B | SC24B\_HUMAN | Protein transport protein Sec24B | 562 | 562 | SEC24B | SC24B\_HUMAN | Protein transport protein Sec24B | 1C(125.0477) | NaN | X | NaN | NaN | NaN | NaN | 3.391275e+04 | 3.391275e+04 | NaN | NaN | CTLTNIPQTQALLNK |
| PIGT | PIGT\_HUMAN | GPI transamidase component PIG-T | 272 | 267 | PIGT | PIGT\_HUMAN | GPI transamidase component PIG-T | 6C(125.0477) | NaN | X | NaN | NaN | NaN | NaN | 3.385039e+04 | NaN | 3.385039e+04 | NaN | TLTEPCPLASESR |
| CUL2 | CUL2\_HUMAN | Cullin-2 | 385 | 383 | CUL2 | CUL2\_HUMAN | Cullin-2 | 3C(125.0477) | NaN | X | NaN | NaN | NaN | NaN | 3.374511e+04 | 3.374511e+04 | NaN | NaN | SVCKAPELLAK |
| RAPGEF2 | RPGF2\_HUMAN | Rap guanine nucleotide exchange factor 2 | 1252 | 1247 | RAPGEF2 | RPGF2\_HUMAN | Rap guanine nucleotide exchange factor 2 | 6C(125.0477) | NaN | X | NaN | NaN | NaN | NaN | 3.374201e+04 | 3.374201e+04 | NaN | NaN | GSWTSCSSGSHDNIQTIQHQR |
| MAP2K3 | MP2K3\_HUMAN | Dual specificity mitogen-activated protein kinase kinase 3 | 305 | 294 | MAP2K3 | MP2K3\_HUMAN | Dual specificity mitogen-activated protein kinase kinase 3 | 12C(125.0477) | NaN | X | NaN | NaN | NaN | NaN | 3.372260e+04 | 3.372260e+04 | NaN | NaN | FSPEFVDFTAQCLR |
| CAST | ICAL\_HUMAN | Calpastatin | 381 | 364 | CAST | ICAL\_HUMAN | Calpastatin | 18C(125.0477) | NaN | X | NaN | NaN | NaN | NaN | 3.355428e+04 | 3.355428e+04 | NaN | NaN | SESELIDELSEDFDRSECKEKPSKPTEK |
| TBC1D24 | TBC24\_HUMAN | TBC1 domain family member 24 | 161 | 158 | TBC1D24 | TBC24\_HUMAN | TBC1 domain family member 24 | 4C(125.0477) | NaN | X | NaN | NaN | NaN | NaN | 3.352208e+04 | NaN | 3.352208e+04 | NaN | ILACNDPGR |
| EIF1AX | IF1AX\_HUMAN | Eukaryotic translation initiation factor 1A, X-chromosomal | 51 | 47 | EIF1AX | IF1AX\_HUMAN | Eukaryotic translation initiation factor 1A, X-chromosomal | 5C(125.0477) | NaN | X | NaN | NaN | NaN | NaN | 3.352187e+04 | 3.352187e+04 | NaN | NaN | LEAMCFDGVKR |
| HUWE1 | HUWE1\_HUMAN | E3 ubiquitin-protein ligase HUWE1 | 2666 | 2657 | HUWE1 | HUWE1\_HUMAN | E3 ubiquitin-protein ligase HUWE1 | 10C(125.0477) | NaN | X | NaN | NaN | NaN | NaN | 3.351117e+04 | 3.351117e+04 | NaN | NaN | VLDAESMHDCVSVVK |
| AK1 | KAD1\_HUMAN | Adenylate kinase isoenzyme 1 | 25 | 22 | AK1 | KAD1\_HUMAN | Adenylate kinase isoenzyme 1 | 4C(125.0477) | NaN | X | NaN | NaN | NaN | NaN | 3.350964e+04 | 3.350964e+04 | NaN | NaN | GTQCEKIVQKYGYTHLSTGDLLR |
| AP2A1 | AP2A1\_HUMAN | AP-2 complex subunit alpha-1 | 492 | 482 | AP2A1 | AP2A1\_HUMAN | AP-2 complex subunit alpha-1 | 11C(125.0477) | NaN | X | NaN | NaN | NaN | NaN | 3.350856e+04 | 3.350856e+04 | NaN | NaN | TVFEALQAPACHENMVK |
| HUWE1 | HUWE1\_HUMAN | E3 ubiquitin-protein ligase HUWE1 | 3239 | 3233 | HUWE1 | HUWE1\_HUMAN | E3 ubiquitin-protein ligase HUWE1 | 7C(125.0477) | NaN | X | NaN | NaN | NaN | NaN | 3.350854e+04 | NaN | 3.350854e+04 | NaN | SSESELCIETPK |
| TRIM25 | TRI25\_HUMAN | E3 ubiquitin/ISG15 ligase TRIM25 | 186 | 185 | TRIM25 | TRI25\_HUMAN | E3 ubiquitin/ISG15 ligase TRIM25 | 2C(125.0477) | NaN | X | NaN | NaN | NaN | NaN | 3.346317e+04 | 3.346317e+04 | NaN | NaN | TCSPASLSQASADLEATLR |
| ACTN4 | ACTN4\_HUMAN | Alpha-actinin-4 | 793 | 792 | ACTN4 | ACTN4\_HUMAN | Alpha-actinin-4 | 2C(125.0477) | NaN | X | NaN | NaN | NaN | NaN | 3.345372e+04 | 3.345372e+04 | NaN | NaN | ACLISLGYDVENDR |
| CD59 | CD59\_HUMAN | CD59 glycoprotein | 70, 64 | 56 | CD59 | CD59\_HUMAN | CD59 glycoprotein | 15C(31.9988) 9C(15.9949) | NaN | NaN | X | X | NaN | NaN | 3.344836e+04 | 3.344836e+04 | NaN | NaN | AGLQVYNKCWKFEHCNFNDVTTR |
| ARRB1 | ARRB1\_HUMAN | Beta-arrestin-1 | 150 | 139 | ARRB1 | ARRB1\_HUMAN | Beta-arrestin-1 | 12C(15.9949) | NaN | NaN | X | NaN | NaN | NaN | 3.344165e+04 | 3.344165e+04 | NaN | NaN | ACGVDYEVKAFCAENLEEK |
| PFKP | PFKAP\_HUMAN | ATP-dependent 6-phosphofructokinase, platelet type | 123 | 118 | PFKP | PFKAP\_HUMAN | ATP-dependent 6-phosphofructokinase, platelet type | 6C(125.0477) | NaN | X | NaN | NaN | NaN | NaN | 3.344116e+04 | 3.344116e+04 | NaN | NaN | GITNLCVIGGDGSLTGANLFR |
| BABAM2 | BABA2\_HUMAN | BRISC and BRCA1-A complex member 2 | 129 | 119 | BABAM2 | BABA2\_HUMAN | BRISC and BRCA1-A complex member 2 | 11C(125.0477) | NaN | X | NaN | NaN | NaN | NaN | 3.343776e+04 | 3.343776e+04 | NaN | NaN | ELVQQYHQFQCSR |
| LDHB | LDHB\_HUMAN | L-lactate dehydrogenase B chain | 294 | 280 | LDHB | LDHB\_HUMAN | L-lactate dehydrogenase B chain | 15C(125.0477) | NaN | X | NaN | NaN | NaN | NaN | 3.338190e+04 | 3.338190e+04 | NaN | NaN | GMYGIENEVFLSLPCILNAR |
| RTCA | RTCA\_HUMAN | RNA 3'-terminal phosphate cyclase | 63 | 61 | RTCA | RTCA\_HUMAN | RNA 3'-terminal phosphate cyclase | 3C(125.0477) | NaN | X | NaN | NaN | NaN | NaN | 3.337930e+04 | 3.337930e+04 | NaN | NaN | DLCDGQLEGAEIGSTEITFTPEK |
| PFAS | PUR4\_HUMAN | Phosphoribosylformylglycinamidine synthase | 831 | 819 | PFAS | PUR4\_HUMAN | Phosphoribosylformylglycinamidine synthase | 13C(125.0477) | NaN | X | NaN | NaN | NaN | NaN | 3.337104e+04 | 3.337104e+04 | NaN | NaN | APGSLVISAYAVCPDITATVTPDLKHPEGR |
| KBTBD8 | KBTB8\_HUMAN | Kelch repeat and BTB domain-containing protein 8 | 306 | 301 | KBTBD8 | KBTB8\_HUMAN | Kelch repeat and BTB domain-containing protein 8 | 6C(125.0477) | NaN | X | NaN | NaN | NaN | NaN | 3.334552e+04 | 3.334552e+04 | NaN | NaN | KQTVPCLDIVTGR |
| GPS1 | CSN1\_HUMAN | COP9 signalosome complex subunit 1 | 341 | 333 | GPS1 | CSN1\_HUMAN | COP9 signalosome complex subunit 1 | 9C(125.0477) | NaN | X | NaN | NaN | NaN | NaN | 3.322907e+04 | 3.322907e+04 | NaN | NaN | FYESKYASCLK |
| TPP2 | TPP2\_HUMAN | Tripeptidyl-peptidase 2 | 456 | 441 | TPP2 | TPP2\_HUMAN | Tripeptidyl-peptidase 2 | 16C(125.0477) | NaN | X | NaN | NaN | NaN | NaN | 3.320033e+04 | 3.320033e+04 | NaN | NaN | GTQLMNGTSMSSPNACGGIALILSGLK |
| DYNLL2 | DYL2\_HUMAN | Dynein light chain 2, cytoplasmic | 56 | 49 | DYNLL2 | DYL2\_HUMAN | Dynein light chain 2, cytoplasmic | 8C(125.0477) | NaN | X | NaN | NaN | NaN | NaN | 3.319805e+04 | 3.319805e+04 | NaN | NaN | KYNPTWHCIVGR |
| HSPA4 | HSP74\_HUMAN | Heat shock 70 kDa protein 4 | 213 | 197 | HSPA4 | HSP74\_HUMAN | Heat shock 70 kDa protein 4 | 17C(125.0477) | NaN | X | NaN | NaN | NaN | NaN | 3.318293e+04 | 3.318293e+04 | NaN | NaN | NVVFVDMGHSAYQVSVCAFNR |
| TRAPPC8 | TPPC8\_HUMAN | Trafficking protein particle complex subunit 8 | 1163 | 1163 | TRAPPC8 | TPPC8\_HUMAN | Trafficking protein particle complex subunit 8 | 1C(125.0477) | NaN | X | NaN | NaN | NaN | NaN | 3.317696e+04 | 3.317696e+04 | NaN | NaN | CEKEEAATQSSEK |
| TAGLN2 | TAGL2\_HUMAN | Transgelin-2 | 63 | 58 | TAGLN2 | TAGL2\_HUMAN | Transgelin-2 | 6C(125.0477) | NaN | X | NaN | NaN | NaN | NaN | 3.315918e+04 | 3.315918e+04 | NaN | NaN | DGTVLCELINALYPEGQAPVK |
| CYB5R3 | NB5R3\_HUMAN | NADH-cytochrome b5 reductase 3 | 204 | 193 | CYB5R3 | NB5R3\_HUMAN | NADH-cytochrome b5 reductase 3 | 12C(125.0477) | NaN | X | NaN | NaN | NaN | NaN | 3.315438e+04 | 3.315438e+04 | NaN | NaN | AIMKDPDDHTVCHLLFANQTEKDILLRPELEELR |
| FCHO2 | FCHO2\_HUMAN | F-BAR domain only protein 2 | 86 | 83 | FCHO2 | FCHO2\_HUMAN | F-BAR domain only protein 2 | 4C(125.0477) | NaN | X | NaN | NaN | NaN | NaN | 3.311275e+04 | 3.311275e+04 | NaN | NaN | LANCHLDLVR |
| RNH1 | RINI\_HUMAN | Ribonuclease inhibitor | 216, 199 | 196 | RNH1 | RINI\_HUMAN | Ribonuclease inhibitor | 21C(125.0477) 4C(15.9949) | NaN | X | X | NaN | NaN | NaN | 3.311143e+04 | 3.311143e+04 | NaN | NaN | DSPCQLEALKLESCGVTSDNCR |
| MBLAC2 | MBLC2\_HUMAN | Acyl-coenzyme A thioesterase MBLAC2 | 176 | 173 | MBLAC2 | MBLC2\_HUMAN | Acyl-coenzyme A thioesterase MBLAC2 | 4C(125.0477) | NaN | X | NaN | NaN | NaN | NaN | 3.309621e+04 | 3.309621e+04 | NaN | NaN | GSICLHDKDR |
| HEMGN | HEMGN\_HUMAN | Hemogen | 422 | 421 | HEMGN | HEMGN\_HUMAN | Hemogen | 2C(125.0477) | NaN | X | NaN | NaN | NaN | NaN | 3.307112e+04 | 3.307112e+04 | NaN | NaN | ECFPEPHQETGGPQGQDPK |
| NCAPD2 | CND1\_HUMAN | Condensin complex subunit 1 | 816 | 811 | NCAPD2 | CND1\_HUMAN | Condensin complex subunit 1 | 6C(125.0477) | NaN | X | NaN | NaN | NaN | NaN | 3.306507e+04 | 3.306507e+04 | NaN | NaN | LAQQVCHAIANISDR |
| MPP1 | EM55\_HUMAN | 55 kDa erythrocyte membrane protein | 242 | 228 | MPP1 | EM55\_HUMAN | 55 kDa erythrocyte membrane protein | 15C(125.0477) | NaN | X | NaN | NaN | NaN | NaN | 3.298011e+04 | 3.298011e+04 | NaN | NaN | VASMAQSAPSEAPSCSPFGK |
| ROCK1 | ROCK1\_HUMAN | Rho-associated protein kinase 1 | 1206 | 1195 | ROCK1 | ROCK1\_HUMAN | Rho-associated protein kinase 1 | 12C(125.0477) | NaN | X | NaN | NaN | NaN | NaN | 3.295248e+04 | 3.295248e+04 | NaN | NaN | IFQILYANEGECR |
| CNP | CN37\_HUMAN | 2',3'-cyclic-nucleotide 3'-phosphodiesterase | 418 | 412 | CNP | CN37\_HUMAN | 2',3'-cyclic-nucleotide 3'-phosphodiesterase | 7C(31.9988) | NaN | NaN | NaN | X | NaN | NaN | 3.295043e+04 | NaN | 3.295043e+04 | NaN | GGALQSCTII |
| DIAPH1 | DIAP1\_HUMAN | Protein diaphanous homolog 1 | 964 | 944 | DIAPH1 | DIAP1\_HUMAN | Protein diaphanous homolog 1 | 21C(125.0477) | NaN | X | NaN | NaN | NaN | NaN | 3.292027e+04 | 3.292027e+04 | NaN | NaN | LQFSEQVENIKPEIVSVTAACEELRK |
| SEC31A | SC31A\_HUMAN | Protein transport protein Sec31A | 1130 | 1130 | SEC31A | SC31A\_HUMAN | Protein transport protein Sec31A | 1C(125.0477) | NaN | X | NaN | NaN | NaN | NaN | 3.287136e+04 | 3.287136e+04 | NaN | NaN | CLSSATDPQTKR |
| ATP7B | ATP7B\_HUMAN | Copper-transporting ATPase 2 | 154 | 149 | ATP7B | ATP7B\_HUMAN | Copper-transporting ATPase 2 | 6C(125.0477) | NaN | X | NaN | NaN | NaN | NaN | 3.281808e+04 | 3.281808e+04 | NaN | NaN | VEGMTCQSCVSSIEGK |
| OLA1 | OLA1\_HUMAN | Obg-like ATPase 1 | 187 | 178 | OLA1 | OLA1\_HUMAN | Obg-like ATPase 1 | 10C(125.0477) | NaN | X | NaN | NaN | NaN | NaN | 3.279884e+04 | 3.279884e+04 | NaN | NaN | KLKPEYDIMCK |
| KCNAB2 | KCAB2\_HUMAN | Voltage-gated potassium channel subunit beta-2 | 248 | 236 | KCNAB2 | KCAB2\_HUMAN | Voltage-gated potassium channel subunit beta-2 | 13C(125.0477) | NaN | X | NaN | NaN | NaN | NaN | 3.272862e+04 | 3.272862e+04 | NaN | NaN | IGVGAMTWSPLACGIVSGK |
| ANKFY1 | ANFY1\_HUMAN | Rabankyrin-5 | 742 | 732 | ANKFY1 | ANFY1\_HUMAN | Rabankyrin-5 | 11C(15.9949) | NaN | NaN | X | NaN | NaN | NaN | 3.270666e+04 | 3.270666e+04 | NaN | NaN | AIDENNEPTACFLIRSGCDVNSPR |
| EDC4 | EDC4\_HUMAN | Enhancer of mRNA-decapping protein 4 | 838 | 836 | EDC4 | EDC4\_HUMAN | Enhancer of mRNA-decapping protein 4 | 3C(125.0477) | NaN | X | NaN | NaN | NaN | NaN | 3.267523e+04 | NaN | 3.267523e+04 | NaN | ETCSTLAESPR |
| IMPA1 | IMPA1\_HUMAN | Inositol monophosphatase 1 | 125 | 116 | IMPA1 | IMPA1\_HUMAN | Inositol monophosphatase 1 | 10C(125.0477) | NaN | X | NaN | NaN | NaN | NaN | 3.264147e+04 | 3.264147e+04 | NaN | NaN | KIEFGVVYSCVEGK |
| HEATR5B | HTR5B\_HUMAN | HEAT repeat-containing protein 5B | 223 | 200 | HEATR5B | HTR5B\_HUMAN | HEAT repeat-containing protein 5B | 24C(125.0477) | NaN | X | NaN | NaN | NaN | NaN | 3.259842e+04 | 3.259842e+04 | NaN | NaN | CLLELQNEAVFMWTAELENIATLCFK |
| ASAP1 | ASAP1\_HUMAN | Arf-GAP with SH3 domain, ANK repeat and PH domain-containing protein 1 | 219 | 213 | ASAP1 | ASAP1\_HUMAN | Arf-GAP with SH3 domain, ANK repeat and PH domain-containing protein 1 | 7C(125.0477) | NaN | X | NaN | NaN | NaN | NaN | 3.252151e+04 | 3.252151e+04 | NaN | NaN | LFQLQMCEYLIK |
| OSTF1 | OSTF1\_HUMAN | Osteoclast-stimulating factor 1 | 185 | 174 | OSTF1 | OSTF1\_HUMAN | Osteoclast-stimulating factor 1 | 12C(125.0477) | NaN | X | NaN | NaN | NaN | NaN | 3.251732e+04 | 3.251732e+04 | NaN | NaN | KLAFDMATNAACASLLK |
| RAB2B | RAB2B\_HUMAN | Ras-related protein Rab-2B | 21 | 20 | RAB2B | RAB2B\_HUMAN | Ras-related protein Rab-2B | 2C(125.0477) | NaN | X | NaN | NaN | NaN | NaN | 3.248394e+04 | 3.248394e+04 | NaN | NaN | SCLLLQFTDK |
| CUL2 | CUL2\_HUMAN | Cullin-2 | 282 | 270 | CUL2 | CUL2\_HUMAN | Cullin-2 | 13C(125.0477) | NaN | X | NaN | NaN | NaN | NaN | 3.243665e+04 | 3.243665e+04 | NaN | NaN | MVADHLQFLHAECHNIIR |
| PA2G4 | PA2G4\_HUMAN | Proliferation-associated protein 2G4 | 296 | 291 | PA2G4 | PA2G4\_HUMAN | Proliferation-associated protein 2G4 | 6C(125.0477) | NaN | X | NaN | NaN | NaN | NaN | 3.236969e+04 | 3.236969e+04 | NaN | NaN | MGVVECAK |
| CNP | CN37\_HUMAN | 2',3'-cyclic-nucleotide 3'-phosphodiesterase | 158 | 156 | CNP | CN37\_HUMAN | 2',3'-cyclic-nucleotide 3'-phosphodiesterase | 3C(125.0477) | NaN | X | NaN | NaN | NaN | NaN | 3.236341e+04 | 3.236341e+04 | NaN | NaN | LDCAQLK |
| CDC37 | CDC37\_HUMAN | Hsp90 co-chaperone Cdc37 | 308 | 308 | CDC37 | CDC37\_HUMAN | Hsp90 co-chaperone Cdc37 | 1C(125.0477) | NaN | X | NaN | NaN | NaN | NaN | 3.235239e+04 | 3.235239e+04 | NaN | NaN | CFDVKDVQMLQDAISK |
| TPM3 | TPM3\_HUMAN | Tropomyosin alpha-3 chain | 191 | 191 | TPM3 | TPM3\_HUMAN | Tropomyosin alpha-3 chain | 1C(125.0477) | NaN | X | NaN | NaN | NaN | NaN | 3.234716e+04 | NaN | 3.234716e+04 | NaN | CSELEEELK |
| ALB | ALBU\_HUMAN | Albumin | 125 | 118 | ALB | ALBU\_HUMAN | Albumin | 8C(57.0215) | X | NaN | NaN | NaN | NaN | NaN | 3.234670e+04 | 3.234670e+04 | NaN | NaN | QEPERNECFLQHK |
| AP1M1 | AP1M1\_HUMAN | AP-1 complex subunit mu-1 | 241 | 230 | AP1M1 | AP1M1\_HUMAN | AP-1 complex subunit mu-1 | 12C(125.0477) | NaN | X | NaN | NaN | NaN | NaN | 3.234152e+04 | 3.234152e+04 | NaN | NaN | SVELEDVKFHQCVR |
| ARMC8 | ARMC8\_HUMAN | Armadillo repeat-containing protein 8 | 661 | 652 | ARMC8 | ARMC8\_HUMAN | Armadillo repeat-containing protein 8 | 10C(125.0477) | NaN | X | NaN | NaN | NaN | NaN | 3.234062e+04 | NaN | 3.234062e+04 | NaN | LSQSPDSNLCDK |
| OXSR1 | OXSR1\_HUMAN | Serine/threonine-protein kinase | 191 | 185 | OXSR1 | OXSR1\_HUMAN | Serine/threonine-protein kinase | 7C(125.0477) | NaN | X | NaN | NaN | NaN | NaN | 3.232640e+04 | 3.232640e+04 | NaN | NaN | TFVGTPCWMAPEVMEQVR |
| STX12 | STX12\_HUMAN | Syntaxin-12 | 29 | 21 | STX12 | STX12\_HUMAN | Syntaxin-12 | 9C(125.0477) | NaN | X | NaN | NaN | NaN | NaN | 3.230720e+04 | 3.230720e+04 | NaN | NaN | DFSSIIQTCSGNIQR |
| ACTR1A | ACTZ\_HUMAN | Alpha-centractin | 34 | 33 | ACTR1A | ACTZ\_HUMAN | Alpha-centractin | 2C(125.0477) | NaN | X | NaN | NaN | NaN | NaN | 3.227869e+04 | 3.227869e+04 | NaN | NaN | YCFPNYVGRPK |
| TF | TRFE\_HUMAN | Serotransferrin | 503 | 495 | TF | TRFE\_HUMAN | Serotransferrin | 9C(57.0215) | X | NaN | NaN | NaN | NaN | NaN | 3.227687e+04 | NaN | 3.227687e+04 | NaN | FDEFFSEGCAPGSK |
| LEPROT | OBRG\_HUMAN | Leptin receptor gene-related protein | 66, 66 | 55 | LEPROT | OBRG\_HUMAN | Leptin receptor gene-related protein | 12C(125.0477) 12C(31.9988) | NaN | X | NaN | X | NaN | NaN | 3.226609e+04 | NaN | NaN | 32266.0940 | VTYDSDATSSACR |
| LRRC14 | LRC14\_HUMAN | Leucine-rich repeat-containing protein 14 | 221 | 209 | LRRC14 | LRC14\_HUMAN | Leucine-rich repeat-containing protein 14 | 13C(125.0477) | NaN | X | NaN | NaN | NaN | NaN | 3.219769e+04 | 3.219769e+04 | NaN | NaN | NTVALLQLLDAGCLR |
| GDPGP1 | GDPP1\_HUMAN | GDP-D-glucose phosphorylase 1 | 333 | 324 | GDPGP1 | GDPP1\_HUMAN | GDP-D-glucose phosphorylase 1 | 10C(125.0477) | NaN | X | NaN | NaN | NaN | NaN | 3.217367e+04 | 3.217367e+04 | NaN | NaN | DGEAFNVALCELAGHLPVK |
| DCUN1D1 | DCNL1\_HUMAN | DCN1-like protein 1 | 131 | 120 | DCUN1D1 | DCNL1\_HUMAN | DCN1-like protein 1 | 12C(125.0477) | NaN | X | NaN | NaN | NaN | NaN | 3.207883e+04 | 3.207883e+04 | NaN | NaN | QEFMDGMTELGCDSIEK |
| CAD | PYR1\_HUMAN | CAD protein | 1455 | 1451 | CAD | PYR1\_HUMAN | CAD protein | 5C(125.0477) | NaN | X | NaN | NaN | NaN | NaN | 3.206556e+04 | 3.206556e+04 | NaN | NaN | VHVDCMTSQK |
| LDHC | LDHC\_HUMAN | L-lactate dehydrogenase C chain | 21 | 12 | LDHC | LDHC\_HUMAN | L-lactate dehydrogenase C chain | 10C(125.0477) | NaN | X | NaN | NaN | NaN | NaN | 3.206430e+04 | 3.206430e+04 | NaN | NaN | LIEDDENSQCK |
| STK24 | STK24\_HUMAN | Serine/threonine-protein kinase 24 | 266 | 261 | STK24 | STK24\_HUMAN | Serine/threonine-protein kinase 24 | 6C(125.0477) | NaN | X | NaN | NaN | NaN | NaN | 3.199535e+04 | 3.199535e+04 | NaN | NaN | EFVEACLNKEPSFRPTAK |
| BMP2K | BMP2K\_HUMAN | BMP-2-inducible protein kinase | 197 | 183 | BMP2K | BMP2K\_HUMAN | BMP-2-inducible protein kinase | 15C(125.0477) | NaN | X | NaN | NaN | NaN | NaN | 3.198065e+04 | 3.198065e+04 | NaN | NaN | VENILLNDGGNYVLCDFGSATNK |
| GBE1 | GLGB\_HUMAN | 1,4-alpha-glucan-branching enzyme | 309 | 295 | GBE1 | GLGB\_HUMAN | 1,4-alpha-glucan-branching enzyme | 15C(125.0477) | NaN | X | NaN | NaN | NaN | NaN | 3.196365e+04 | 3.196365e+04 | NaN | NaN | NSADGLNMFDGTDSCYFHSGPR |
| ANK1 | ANK1\_HUMAN | Ankyrin-1 | 1070 | 1069 | ANK1 | ANK1\_HUMAN | Ankyrin-1 | 2C(125.0477) | NaN | X | NaN | NaN | NaN | NaN | 3.192827e+04 | 3.192827e+04 | NaN | NaN | LCQDYDTIGPEGGSLKSK |
| CMBL | CMBL\_HUMAN | Carboxymethylenebutenolidase homolog | 222 | 218 | CMBL | CMBL\_HUMAN | Carboxymethylenebutenolidase homolog | 5C(125.0477) | NaN | X | NaN | NaN | NaN | NaN | 3.189143e+04 | 3.189143e+04 | NaN | NaN | KREDCSPADKPYIDEAR |
| ATP6V0A1 | VPP1\_HUMAN | V-type proton ATPase 116 kDa subunit a 1 | 247 | 242 | ATP6V0A1 | VPP1\_HUMAN | V-type proton ATPase 116 kDa subunit a 1 | 6C(125.0477) | NaN | X | NaN | NaN | NaN | NaN | 3.187071e+04 | NaN | 3.187071e+04 | NaN | ASLYPCPETPQER |
| SCRN1 | SCRN1\_HUMAN | Secernin-1 | 290 | 287 | SCRN1 | SCRN1\_HUMAN | Secernin-1 | 4C(125.0477) | NaN | X | NaN | NaN | NaN | NaN | 3.184017e+04 | 3.184017e+04 | NaN | NaN | SSPCIHYFTGTPDPSR |
| WDR91 | WDR91\_HUMAN | WD repeat-containing protein 91 | 246 | 237 | WDR91 | WDR91\_HUMAN | WD repeat-containing protein 91 | 10C(125.0477) | NaN | X | NaN | NaN | NaN | NaN | 3.182721e+04 | 3.182721e+04 | NaN | NaN | LGDSELAMVCSQR |
| CAPZA2 | CAZA2\_HUMAN | F-actin-capping protein subunit alpha-2 | 141 | 130 | CAPZA2 | CAZA2\_HUMAN | F-actin-capping protein subunit alpha-2 | 12C(125.0477) | NaN | X | NaN | NaN | NaN | NaN | 3.181824e+04 | 3.181824e+04 | NaN | NaN | AYVKEHYPNGVCTVYGK |
| FLNA | FLNA\_HUMAN | Filamin-A | 1260 | 1247 | FLNA | FLNA\_HUMAN | Filamin-A | 14C(125.0477) | NaN | X | NaN | NaN | NaN | NaN | 3.174550e+04 | 3.174550e+04 | NaN | NaN | LQVEPAVDTSGVQCYGPGIEGQGVFR |
| USP24 | UBP24\_HUMAN | Ubiquitin carboxyl-terminal hydrolase 24 | 2281 | 2280 | USP24 | UBP24\_HUMAN | Ubiquitin carboxyl-terminal hydrolase 24 | 2C(125.0477) | NaN | X | NaN | NaN | NaN | NaN | 3.174116e+04 | 3.174116e+04 | NaN | NaN | NCAQYFFLFNTFVQK |
| AP2B1 | AP2B1\_HUMAN | AP-2 complex subunit beta | 129 | 129 | AP2B1 | AP2B1\_HUMAN | AP-2 complex subunit beta | 1C(125.0477) | NaN | X | NaN | NaN | NaN | NaN | 3.172019e+04 | 3.172019e+04 | NaN | NaN | CLKDEDPYVRK |
| GOT2 | AATM\_HUMAN | Aspartate aminotransferase, mitochondrial | 106 | 91 | GOT2 | AATM\_HUMAN | Aspartate aminotransferase, mitochondrial | 16C(125.0477) | NaN | X | NaN | NaN | NaN | NaN | 3.169213e+04 | 3.169213e+04 | NaN | NaN | NLDKEYLPIGGLAEFCK |
| TSNAX | TSNAX\_HUMAN | Translin-associated protein X | 257 | 253 | TSNAX | TSNAX\_HUMAN | Translin-associated protein X | 5C(125.0477) | NaN | X | NaN | NaN | NaN | NaN | 3.168433e+04 | NaN | 3.168433e+04 | NaN | VENACYALK |
| PAK2 | PAK2\_HUMAN | Serine/threonine-protein kinase PAK 2 | 390 | 384 | PAK2 | PAK2\_HUMAN | Serine/threonine-protein kinase PAK 2 | 7C(125.0477) | NaN | X | NaN | NaN | NaN | NaN | 3.158790e+04 | 3.158790e+04 | NaN | NaN | LTDFGFCAQITPEQSK |
| SEC62 | SEC62\_HUMAN | Translocation protein SEC62 | 82 | 76 | SEC62 | SEC62\_HUMAN | Translocation protein SEC62 | 7C(125.0477) | NaN | X | NaN | NaN | NaN | NaN | 3.158037e+04 | NaN | 3.158037e+04 | NaN | ESVVDYCNR |
| FTH1 | FRIH\_HUMAN | Ferritin heavy chain | 131 | 126 | FTH1 | FRIH\_HUMAN | Ferritin heavy chain | 6C(125.0477) | NaN | X | NaN | NaN | NaN | NaN | 3.156980e+04 | 3.156980e+04 | NaN | NaN | NDPHLCDFIETHYLNEQVK |
| AMPD3 | AMPD3\_HUMAN | AMP deaminase 3 | 747 | 743 | AMPD3 | AMPD3\_HUMAN | AMP deaminase 3 | 5C(125.0477) | NaN | X | NaN | NaN | NaN | NaN | 3.149518e+04 | 3.149518e+04 | NaN | NaN | YETLCNELSFLSDAMK |
| FCHO2 | FCHO2\_HUMAN | F-BAR domain only protein 2 | 679 | 679 | FCHO2 | FCHO2\_HUMAN | F-BAR domain only protein 2 | 1C(125.0477) | NaN | X | NaN | NaN | NaN | NaN | 3.147216e+04 | NaN | 3.147216e+04 | NaN | CSASTTDLR |
| ATP11A | AT11A\_HUMAN | Phospholipid-transporting ATPase IH | 625 | 616 | ATP11A | AT11A\_HUMAN | Phospholipid-transporting ATPase IH | 10C(125.0477) | NaN | X | NaN | NaN | NaN | NaN | 3.145696e+04 | NaN | 3.145696e+04 | NaN | LIQEEYEGICK |
| VAV3 | VAV3\_HUMAN | Guanine nucleotide exchange factor VAV3 | 44 | 39 | VAV3 | VAV3\_HUMAN | Guanine nucleotide exchange factor VAV3 | 6C(125.0477) | NaN | X | NaN | NaN | NaN | NaN | 3.143776e+04 | 3.143776e+04 | NaN | NaN | DGVLLCQLLNNLR |
| ANKRD28 | ANR28\_HUMAN | Serine/threonine-protein phosphatase 6 regulatory ankyrin repeat subunit A | 632 | 618 | ANKRD28 | ANR28\_HUMAN | Serine/threonine-protein phosphatase 6 regulatory ankyrin repeat subunit A | 15C(125.0477) | NaN | X | NaN | NaN | NaN | NaN | 3.138985e+04 | 3.138985e+04 | NaN | NaN | RTPIHAAATNGHSECLR |
| UROD | DCUP\_HUMAN | Uroporphyrinogen decarboxylase | 59 | 51 | UROD | DCUP\_HUMAN | Uroporphyrinogen decarboxylase | 9C(125.0477) | NaN | X | NaN | NaN | NaN | NaN | 3.137521e+04 | 2.009197e+04 | 1.128324e+04 | NaN | AAQDFFSTCR |
| ACTR2 | ARP2\_HUMAN | Actin-related protein 2 | 20 | 20 | ACTR2 | ARP2\_HUMAN | Actin-related protein 2 | 1C(125.0477) | NaN | X | NaN | NaN | NaN | NaN | 3.133796e+04 | 3.133796e+04 | NaN | NaN | CGYAGSNFPEHIFPALVGRPIIR |
| LRRC20 | LRC20\_HUMAN | Leucine-rich repeat-containing protein 20 | 32 | 16 | LRRC20 | LRC20\_HUMAN | Leucine-rich repeat-containing protein 20 | 17C(125.0477) | NaN | X | NaN | NaN | NaN | NaN | 3.118651e+04 | 3.118651e+04 | NaN | NaN | VNETVESGSDTLDLAECK |
| CAST | ICAL\_HUMAN | Calpastatin | 413 | 410 | CAST | ICAL\_HUMAN | Calpastatin | 4C(125.0477) | NaN | X | NaN | NaN | NaN | NaN | 3.115586e+04 | 3.115586e+04 | NaN | NaN | TSMCSIQSAPPEPATLKGTVPDDAVEALADSLGKK |
| CRNN | CRNN\_HUMAN | Cornulin | 425 | 425 | CRNN | CRNN\_HUMAN | Cornulin | 1C(125.0477) | NaN | X | NaN | NaN | NaN | NaN | 3.115551e+04 | NaN | 3.115551e+04 | NaN | CVTEGQGDR |
| HUWE1 | HUWE1\_HUMAN | E3 ubiquitin-protein ligase HUWE1 | 1421 | 1412 | HUWE1 | HUWE1\_HUMAN | E3 ubiquitin-protein ligase HUWE1 | 10C(125.0477) | NaN | X | NaN | NaN | NaN | NaN | 3.115015e+04 | 3.115015e+04 | NaN | NaN | EKQEEEEAKCLEK |
| PTP4A2 | TP4A2\_HUMAN | Protein tyrosine phosphatase type IVA 2 | 46 | 45 | PTP4A2 | TP4A2\_HUMAN | Protein tyrosine phosphatase type IVA 2 | 2C(125.0477) | NaN | X | NaN | NaN | NaN | NaN | 3.108479e+04 | 3.108479e+04 | NaN | NaN | VCDATYDKAPVEK |
| TOR1AIP2 | TOIP2\_HUMAN | Torsin-1A-interacting protein 2 | 382 | 381 | TOR1AIP2 | TOIP2\_HUMAN | Torsin-1A-interacting protein 2 | 2C(125.0477) | NaN | X | NaN | NaN | NaN | NaN | 3.106190e+04 | NaN | 3.106190e+04 | NaN | YCDHENAAFK |
| PPP2CB | PP2AB\_HUMAN | Serine/threonine-protein phosphatase 2A catalytic subunit beta isoform | 133 | 122 | PPP2CB | PP2AB\_HUMAN | Serine/threonine-protein phosphatase 2A catalytic subunit beta isoform | 12C(125.0477) | NaN | X | NaN | NaN | NaN | NaN | 3.105835e+04 | 3.105835e+04 | NaN | NaN | QITQVYGFYDECLRK |
| COMT | COMT\_HUMAN | Catechol O-methyltransferase | 223 | 213 | COMT | COMT\_HUMAN | Catechol O-methyltransferase | 11C(125.0477) | NaN | X | NaN | NaN | NaN | NaN | 3.103241e+04 | 3.103241e+04 | NaN | NaN | GTVLLADNVICPGAPDFLAHVR |
| INPP5D | SHIP1\_HUMAN | Phosphatidylinositol 3,4,5-trisphosphate 5-phosphatase 1 | 385 | 382 | INPP5D | SHIP1\_HUMAN | Phosphatidylinositol 3,4,5-trisphosphate 5-phosphatase 1 | 4C(125.0477) | NaN | X | NaN | NaN | NaN | NaN | 3.098062e+04 | 3.098062e+04 | NaN | NaN | EGFCQLLQQMK |
| CIB1 | CIB1\_HUMAN | Calcium and integrin-binding protein 1 | 134 | 131 | CIB1 | CIB1\_HUMAN | Calcium and integrin-binding protein 1 | 4C(125.0477) | NaN | X | NaN | NaN | NaN | NaN | 3.091924e+04 | NaN | 3.091924e+04 | NaN | LVNCLTGEGEDTR |
| AP1G1 | AP1G1\_HUMAN | AP-1 complex subunit gamma-1 | 160 | 157 | AP1G1 | AP1G1\_HUMAN | AP-1 complex subunit gamma-1 | 4C(125.0477) | NaN | X | NaN | NaN | NaN | NaN | 3.087644e+04 | 3.087644e+04 | NaN | NaN | AALCAVHVIR |
| RAB14 | RAB14\_HUMAN | Ras-related protein Rab-14 | 26 | 25 | RAB14 | RAB14\_HUMAN | Ras-related protein Rab-14 | 2C(125.0477) | NaN | X | NaN | NaN | NaN | NaN | 3.085913e+04 | 3.085913e+04 | NaN | NaN | SCLLHQFTEKK |
| PFKL | PFKAL\_HUMAN | ATP-dependent 6-phosphofructokinase, liver type | 708 | 699 | PFKL | PFKAL\_HUMAN | ATP-dependent 6-phosphofructokinase, liver type | 10C(125.0477) | NaN | X | NaN | NaN | NaN | NaN | 3.085143e+04 | 3.085143e+04 | NaN | NaN | VFANAPDSACVIGLKK |
| CDC42 | CDC42\_HUMAN | Cell division control protein 42 homolog | 81 | 67 | CDC42 | CDC42\_HUMAN | Cell division control protein 42 homolog | 15C(125.0477) | NaN | X | NaN | NaN | NaN | NaN | 3.084627e+04 | 3.084627e+04 | NaN | NaN | LRPLSYPQTDVFLVCFSVVSPSSFENVK |
| GC | VTDB\_HUMAN | Vitamin D-binding protein | 220 | 219 | GC | VTDB\_HUMAN | Vitamin D-binding protein | 2C(57.0215) | X | NaN | NaN | NaN | NaN | NaN | 3.081757e+04 | NaN | 3.081757e+04 | NaN | VCSQYAAYGEK |
| IPO5 | IPO5\_HUMAN | Importin-5 | 560 | 555 | IPO5 | IPO5\_HUMAN | Importin-5 | 6C(125.0477) | NaN | X | NaN | NaN | NaN | NaN | 3.080305e+04 | 3.080305e+04 | NaN | NaN | GKTIECISLIGLAVGKEK |
| MDH2 | MDHM\_HUMAN | Malate dehydrogenase, mitochondrial | 285 | 280 | MDH2 | MDHM\_HUMAN | Malate dehydrogenase, mitochondrial | 6C(125.0477) | NaN | X | NaN | NaN | NaN | NaN | 3.079575e+04 | 3.079575e+04 | NaN | NaN | SQETECTYFSTPLLLGK |
| RNPEP | AMPB\_HUMAN | Aminopeptidase B | 85 | 80 | RNPEP | AMPB\_HUMAN | Aminopeptidase B | 6C(125.0477) | NaN | X | NaN | NaN | NaN | NaN | 3.071228e+04 | 3.071228e+04 | NaN | NaN | LDSHPCLEVTAAALR |
| TXNRD1 | TRXR1\_HUMAN | Thioredoxin reductase 1, cytoplasmic | 625 | 613 | TXNRD1 | TRXR1\_HUMAN | Thioredoxin reductase 1, cytoplasmic | 13C(125.0477) | NaN | X | NaN | NaN | NaN | NaN | 3.069805e+04 | 3.069805e+04 | NaN | NaN | KQLDSTIGIHPVCAEVFTTLSVTKR |
| PSMA6 | PSA6\_HUMAN | Proteasome subunit alpha type-6 | 115 | 105 | PSMA6 | PSA6\_HUMAN | Proteasome subunit alpha type-6 | 11C(125.0477) | NaN | X | NaN | NaN | NaN | NaN | 3.069646e+04 | 3.069646e+04 | NaN | NaN | YGYEIPVDMLCKR |
| RNH1 | RINI\_HUMAN | Ribonuclease inhibitor | 95, 96 | 90 | RNH1 | RINI\_HUMAN | Ribonuclease inhibitor | 6C(125.0477) 7C(15.9949) | NaN | X | X | NaN | NaN | NaN | 3.068386e+04 | NaN | 3.068386e+04 | NaN | LSLQNCCLTGAGCGVLSSTLR |
| ERMAP | ERMAP\_HUMAN | Erythroid membrane-associated protein | 398 | 378 | ERMAP | ERMAP\_HUMAN | Erythroid membrane-associated protein | 21C(125.0477) | NaN | X | NaN | NaN | NaN | NaN | 3.064377e+04 | 3.064377e+04 | NaN | NaN | SHIFTFTHNFSGPLRPFFEPCLHDGGK |
| ALDH9A1 | AL9A1\_HUMAN | 4-trimethylaminobutyraldehyde dehydrogenase | 484 | 482 | ALDH9A1 | AL9A1\_HUMAN | 4-trimethylaminobutyraldehyde dehydrogenase | 3C(125.0477) | NaN | X | NaN | NaN | NaN | NaN | 3.056563e+04 | 3.056563e+04 | NaN | NaN | TVCVEMGDVESAF |
| ZFAND1 | ZFAN1\_HUMAN | AN1-type zinc finger protein 1 | 64 | 54 | ZFAND1 | ZFAN1\_HUMAN | AN1-type zinc finger protein 1 | 11C(125.0477) | NaN | X | NaN | NaN | NaN | NaN | 3.056431e+04 | 3.056431e+04 | NaN | NaN | LKTDQHTSYPCSFK |
| ATG4A | ATG4A\_HUMAN | Cysteine protease ATG4A | 334 | 329 | ATG4A | ATG4A\_HUMAN | Cysteine protease ATG4A | 6C(125.0477) | NaN | X | NaN | NaN | NaN | NaN | 3.054426e+04 | 3.054426e+04 | NaN | NaN | DFDNWCSLVQK |
| HERC1 | HERC1\_HUMAN | Probable E3 ubiquitin-protein ligase HERC1 | 2232 | 2229 | HERC1 | HERC1\_HUMAN | Probable E3 ubiquitin-protein ligase HERC1 | 4C(15.9949) | NaN | NaN | X | NaN | NaN | NaN | 3.051923e+04 | 3.051923e+04 | NaN | NaN | WTYCINK |
| NAA25 | NAA25\_HUMAN | N-alpha-acetyltransferase 25, NatB auxiliary subunit | 339 | 336 | NAA25 | NAA25\_HUMAN | N-alpha-acetyltransferase 25, NatB auxiliary subunit | 4C(125.0477) | NaN | X | NaN | NaN | NaN | NaN | 3.050975e+04 | 3.050975e+04 | NaN | NaN | SQGCNDEYK |
| AMPD3 | AMPD3\_HUMAN | AMP deaminase 3 | 323 | 314 | AMPD3 | AMPD3\_HUMAN | AMP deaminase 3 | 10C(125.0477) | NaN | X | NaN | NaN | NaN | NaN | 3.047274e+04 | 3.047274e+04 | NaN | NaN | VDTHIHAAACMNQK |
| ZFPL1 | ZFPL1\_HUMAN | Zinc finger protein-like 1 | 230 | 219 | ZFPL1 | ZFPL1\_HUMAN | Zinc finger protein-like 1 | 12C(125.0477) | NaN | X | NaN | NaN | NaN | NaN | 3.045805e+04 | 3.045805e+04 | NaN | NaN | DDDRTPGLHGDCDDDKYR |
| AP1B1 | AP1B1\_HUMAN | AP-1 complex subunit beta-1 | 57 | 46 | AP1B1 | AP1B1\_HUMAN | AP-1 complex subunit beta-1 | 12C(125.0477) | NaN | X | NaN | NaN | NaN | NaN | 3.045742e+04 | 3.045742e+04 | NaN | NaN | DVSALFPDVVNCMQTDNLELKK |
| PPP4C | PP4C\_HUMAN | Serine/threonine-protein phosphatase 4 catalytic subunit | 266 | 266 | PPP4C | PP4C\_HUMAN | Serine/threonine-protein phosphatase 4 catalytic subunit | 1C(125.0477) | NaN | X | NaN | NaN | NaN | NaN | 3.044487e+04 | 3.044487e+04 | NaN | NaN | CGNVAAILELDEHLQK |
| CTSB | CATB\_HUMAN | Cathepsin B | 211 | 210 | CTSB | CATB\_HUMAN | Cathepsin B | 2C(125.0477) | NaN | X | NaN | NaN | NaN | NaN | 3.043187e+04 | NaN | 3.043187e+04 | NaN | ICEPGYSPTYK |
| COPS3 | CSN3\_HUMAN | COP9 signalosome complex subunit 3 | 383 | 383 | COPS3 | CSN3\_HUMAN | COP9 signalosome complex subunit 3 | 1C(125.0477) | NaN | X | NaN | NaN | NaN | NaN | 3.042227e+04 | 3.042227e+04 | NaN | NaN | CIELDERLK |
| CCT5 | TCPE\_HUMAN | T-complex protein 1 subunit epsilon | 429 | 418 | CCT5 | TCPE\_HUMAN | T-complex protein 1 subunit epsilon | 12C(125.0477) | NaN | X | NaN | NaN | NaN | NaN | 3.041441e+04 | 3.041441e+04 | NaN | NaN | VVYGGGAAEISCALAVSQEADK |
| MTHFD1 | C1TC\_HUMAN | C-1-tetrahydrofolate synthase, cytoplasmic | 147, 143 | 138 | MTHFD1 | C1TC\_HUMAN | C-1-tetrahydrofolate synthase, cytoplasmic | 10C(15.9949) 6C(125.0477) | NaN | X | X | NaN | NaN | NaN | 3.041127e+04 | 3.041127e+04 | NaN | NaN | GDLNDCFIPCTPKGCLELIK |
| COASY | COASY\_HUMAN | Bifunctional coenzyme A synthase | 357 | 340 | COASY | COASY\_HUMAN | Bifunctional coenzyme A synthase | 18C(125.0477) | NaN | X | NaN | NaN | NaN | NaN | 3.037913e+04 | 3.037913e+04 | NaN | NaN | MLGNLLRPPYERPELPTCLYVIGLTGISGSGK |
| GART | PUR2\_HUMAN | Trifunctional purine biosynthetic protein adenosine-3 | 298 | 293 | GART | PUR2\_HUMAN | Trifunctional purine biosynthetic protein adenosine-3 | 6C(125.0477) | NaN | X | NaN | NaN | NaN | NaN | 3.035897e+04 | 3.035897e+04 | NaN | NaN | FGDPECQVILPLLK |
| PAIP1 | PAIP1\_HUMAN | Polyadenylate-binding protein-interacting protein 1 | 470 | 469 | PAIP1 | PAIP1\_HUMAN | Polyadenylate-binding protein-interacting protein 1 | 2C(125.0477) | NaN | X | NaN | NaN | NaN | NaN | 3.035888e+04 | 3.035888e+04 | NaN | NaN | FCLESER |
| ZZEF1 | ZZEF1\_HUMAN | Zinc finger ZZ-type and EF-hand domain-containing protein 1 | 2553 | 2553 | ZZEF1 | ZZEF1\_HUMAN | Zinc finger ZZ-type and EF-hand domain-containing protein 1 | 1C(125.0477) | NaN | X | NaN | NaN | NaN | NaN | 3.035184e+04 | 3.035184e+04 | NaN | NaN | CDQATAESNPVTQK |
| PSMD11 | PSD11\_HUMAN | 26S proteasome non-ATPase regulatory subunit 11 | 257 | 247 | PSMD11 | PSD11\_HUMAN | 26S proteasome non-ATPase regulatory subunit 11 | 11C(125.0477) | NaN | X | NaN | NaN | NaN | NaN | 3.033966e+04 | 3.033966e+04 | NaN | NaN | AITSLKYMLLCK |
| RP2 | XRP2\_HUMAN | Protein XRP2 | 236 | 229 | RP2 | XRP2\_HUMAN | Protein XRP2 | 8C(125.0477) | NaN | X | NaN | NaN | NaN | NaN | 3.028201e+04 | 3.028201e+04 | NaN | NaN | QKSSDESCLVVLFAGDYTIANAR |
| TF | TRFE\_HUMAN | Serotransferrin | 364 | 363 | TF | TRFE\_HUMAN | Serotransferrin | 2C(57.0215) | X | NaN | NaN | NaN | NaN | NaN | 3.023690e+04 | 3.023690e+04 | NaN | NaN | WCALSHHER |
| CR1 | CR1\_HUMAN | Complement receptor type 1 | 1350 | 1345 | CR1 | CR1\_HUMAN | Complement receptor type 1 | 6C(125.0477) | NaN | X | NaN | NaN | NaN | NaN | 3.022463e+04 | NaN | 3.022463e+04 | NaN | EISYTCDPHPDR |
| LSM7 | LSM7\_HUMAN | U6 snRNA-associated Sm-like protein LSm7 | 76 | 70 | LSM7 | LSM7\_HUMAN | U6 snRNA-associated Sm-like protein LSm7 | 7C(125.0477) | NaN | X | NaN | NaN | NaN | NaN | 3.022415e+04 | 3.022415e+04 | NaN | NaN | QLGLVVCR |
| GLOD4 | GLOD4\_HUMAN | Glyoxalase domain-containing protein 4 | 197 | 188 | GLOD4 | GLOD4\_HUMAN | Glyoxalase domain-containing protein 4 | 10C(125.0477) | NaN | X | NaN | NaN | NaN | NaN | 3.020506e+04 | 3.020506e+04 | NaN | NaN | ALLGYADNQCKLELQGVK |
| CUL4A | CUL4A\_HUMAN | Cullin-4A | 633 | 627 | CUL4A | CUL4A\_HUMAN | Cullin-4A | 7C(57.0215) | X | NaN | NaN | NaN | NaN | NaN | 3.019598e+04 | NaN | NaN | 30195.9790 | TLQSLACGK |
| PDXP | PLPP\_HUMAN | Chronophin | 163 | 161 | PDXP | PLPP\_HUMAN | Chronophin | 3C(125.0477) | NaN | X | NaN | NaN | NaN | NaN | 3.019103e+04 | 3.019103e+04 | NaN | NaN | EACAHLR |
| RABGAP1L | RBG1L\_HUMAN | Rab GTPase-activating protein 1-like | 777 | 771 | RABGAP1L | RBG1L\_HUMAN | Rab GTPase-activating protein 1-like | 7C(125.0477) | NaN | X | NaN | NaN | NaN | NaN | 3.016321e+04 | 3.016321e+04 | NaN | NaN | RLMEQACNIK |
| ZFAND1 | ZFAN1\_HUMAN | AN1-type zinc finger protein 1 | 98 | 92 | ZFAND1 | ZFAN1\_HUMAN | AN1-type zinc finger protein 1 | 7C(125.0477) | NaN | X | NaN | NaN | NaN | NaN | 3.015758e+04 | 3.015758e+04 | NaN | NaN | HQSDHECEKLEIPKPR |
| EIF2S2 | IF2B\_HUMAN | Eukaryotic translation initiation factor 2 subunit 2 | 302, 305 | 297 | EIF2S2 | IF2B\_HUMAN | Eukaryotic translation initiation factor 2 subunit 2 | 6C(125.0477) 9C(15.9949) | NaN | X | X | NaN | NaN | NaN | 3.014102e+04 | 3.014102e+04 | NaN | NaN | LYFLQCETCHSRCSVASIK |
| TUBA1A | TBA1A\_HUMAN | Tubulin alpha-1A chain | 347, 347 | 340 | TUBA1A | TBA1A\_HUMAN | Tubulin alpha-1A chain | 8C(125.0477) 8C(57.0215) | X | X | NaN | NaN | NaN | NaN | 3.012539e+04 | 3.012539e+04 | NaN | NaN | TIQFVDWCPTGFK |
| PFAS | PUR4\_HUMAN | Phosphoribosylformylglycinamidine synthase | 697 | 687 | PFAS | PUR4\_HUMAN | Phosphoribosylformylglycinamidine synthase | 11C(125.0477) | NaN | X | NaN | NaN | NaN | NaN | 3.007862e+04 | 3.007862e+04 | NaN | NaN | SVGGLVAQQQCVGPLQTPLADVAVVALSHEELIGAATALGEQPVK |
| ADAL | ADAL\_HUMAN | Adenosine deaminase-like protein | 260 | 255 | ADAL | ADAL\_HUMAN | Adenosine deaminase-like protein | 6C(125.0477) | NaN | X | NaN | NaN | NaN | NaN | 3.007438e+04 | 3.007438e+04 | NaN | NaN | IPLELCLTSNVK |
| DRG2 | DRG2\_HUMAN | Developmentally-regulated GTP-binding protein 2 | 319 | 312 | DRG2 | DRG2\_HUMAN | Developmentally-regulated GTP-binding protein 2 | 8C(125.0477) | NaN | X | NaN | NaN | NaN | NaN | 3.007282e+04 | 3.007282e+04 | NaN | NaN | GASVEHVCHR |
| PRDX2 | PRDX2\_HUMAN | Peroxiredoxin-2 | 51 | 37 | PRDX2 | PRDX2\_HUMAN | Peroxiredoxin-2 | 15C(125.0477) | NaN | X | NaN | NaN | NaN | NaN | 3.003543e+04 | 3.003543e+04 | NaN | NaN | YVVLFFYPLDFTFVCPTEIIAFSNR |
| SHPK | SHPK\_HUMAN | Sedoheptulokinase | 72 | 65 | SHPK | SHPK\_HUMAN | Sedoheptulokinase | 8C(125.0477) | NaN | X | NaN | NaN | NaN | NaN | 3.000813e+04 | 3.000813e+04 | NaN | NaN | ILQALHECLAALPRPQLR |
| MYH2 | MYH2\_HUMAN | Myosin-2 | 1346 | 1344 | MYH2 | MYH2\_HUMAN | Myosin-2 | 3C(57.0215) | X | NaN | NaN | NaN | NaN | NaN | 2.998074e+04 | 2.998074e+04 | NaN | NaN | HDCDLLR |
| PA2G4 | PA2G4\_HUMAN | Proliferation-associated protein 2G4 | 49 | 34 | PA2G4 | PA2G4\_HUMAN | Proliferation-associated protein 2G4 | 16C(125.0477) | NaN | X | NaN | NaN | NaN | NaN | 2.989057e+04 | 2.989057e+04 | NaN | NaN | SLVEASSSGVSVLSLCEK |
| TLN2 | TLN2\_HUMAN | Talin-2 | 1427 | 1418 | TLN2 | TLN2\_HUMAN | Talin-2 | 10C(125.0477) | NaN | X | NaN | NaN | NaN | NaN | 2.987741e+04 | 2.987741e+04 | NaN | NaN | TGDLPAFGECVGIASK |
| SNX2 | SNX2\_HUMAN | Sorting nexin-2 | 332 | 323 | SNX2 | SNX2\_HUMAN | Sorting nexin-2 | 10C(125.0477) | NaN | X | NaN | NaN | NaN | NaN | 2.987290e+04 | 2.987290e+04 | NaN | NaN | LHVSVEALVCHR |
| PITHD1 | PITH1\_HUMAN | PITH domain-containing protein 1 | 187 | 182 | PITHD1 | PITH1\_HUMAN | PITH domain-containing protein 1 | 6C(125.0477) | NaN | X | NaN | NaN | NaN | NaN | 2.985912e+04 | NaN | 2.985912e+04 | NaN | HEVTICNYEASANPADHR |
| EHBP1L1 | EH1L1\_HUMAN | EH domain-binding protein 1-like protein 1 | 346 | 340 | EHBP1L1 | EH1L1\_HUMAN | EH domain-binding protein 1-like protein 1 | 7C(125.0477) | NaN | X | NaN | NaN | NaN | NaN | 2.982825e+04 | NaN | 2.982825e+04 | NaN | ETQAQACPQEGTEAHGAR |
| GALK2 | GALK2\_HUMAN | N-acetylgalactosamine kinase | 113 | 103 | GALK2 | GALK2\_HUMAN | N-acetylgalactosamine kinase | 11C(125.0477) | NaN | X | NaN | NaN | NaN | NaN | 2.982562e+04 | 2.982562e+04 | NaN | NaN | TKPLWHNYFLCGLK |
| RHOA | RHOA\_HUMAN | Transforming protein RhoA | 16 | 6 | RHOA | RHOA\_HUMAN | Transforming protein RhoA | 11C(125.0477) | NaN | X | NaN | NaN | NaN | NaN | 2.982033e+04 | 2.982033e+04 | NaN | NaN | KKLVIVGDGACGK |
| XPO7 | XPO7\_HUMAN | Exportin-7 | 835 | 829 | XPO7 | XPO7\_HUMAN | Exportin-7 | 7C(125.0477) | NaN | X | NaN | NaN | NaN | NaN | 2.980663e+04 | 2.980663e+04 | NaN | NaN | LKGISICFSMLK |
| PI4K2A | P4K2A\_HUMAN | Phosphatidylinositol 4-kinase type 2-alpha | 320 | 318 | PI4K2A | P4K2A\_HUMAN | Phosphatidylinositol 4-kinase type 2-alpha | 3C(125.0477) | NaN | X | NaN | NaN | NaN | NaN | 2.976559e+04 | NaN | 2.976559e+04 | NaN | YDCPMDSSSSR |
| MYH9 | MYH9\_HUMAN | Myosin-9 | 896 | 883 | MYH9 | MYH9\_HUMAN | Myosin-9 | 14C(125.0477) | NaN | X | NaN | NaN | NaN | NaN | 2.975846e+04 | 2.975846e+04 | NaN | NaN | LQLQEQLQAETELCAEAEELR |
| PTPA | PTPA\_HUMAN | Serine/threonine-protein phosphatase 2A activator | 334 | 329 | PTPA | PTPA\_HUMAN | Serine/threonine-protein phosphatase 2A activator | 6C(125.0477) | NaN | X | NaN | NaN | NaN | NaN | 2.974236e+04 | 2.974236e+04 | NaN | NaN | MYKAECLEK |
| CAPN1 | CAN1\_HUMAN | Calpain-1 catalytic subunit | 417 | 401 | CAPN1 | CAN1\_HUMAN | Calpain-1 catalytic subunit | 17C(125.0477) | NaN | X | NaN | NaN | NaN | NaN | 2.973734e+04 | 2.973734e+04 | NaN | NaN | LDETDDPDDYGDRESGCSFVLALMQK |
| PELO | PELO\_HUMAN | Protein pelota homolog | 204 | 204 | PELO | PELO\_HUMAN | Protein pelota homolog | 1C(125.0477) | NaN | X | NaN | NaN | NaN | NaN | 2.969871e+04 | 2.969871e+04 | NaN | NaN | CILVASPGFVR |
| TLN1 | TLN1\_HUMAN | Talin-1 | 2161 | 2155 | TLN1 | TLN1\_HUMAN | Talin-1 | 7C(125.0477) | NaN | X | NaN | NaN | NaN | NaN | 2.969525e+04 | 2.969525e+04 | NaN | NaN | QELAVFCSPEPPAK |
| SLK | SLK\_HUMAN | STE20-like serine/threonine-protein kinase | 1153 | 1153 | SLK | SLK\_HUMAN | STE20-like serine/threonine-protein kinase | 1C(125.0477) | NaN | X | NaN | NaN | NaN | NaN | 2.965315e+04 | 2.965315e+04 | NaN | NaN | CHLLVEHETQK |
| BMP2K | BMP2K\_HUMAN | BMP-2-inducible protein kinase | 32 | 10 | BMP2K | BMP2K\_HUMAN | BMP-2-inducible protein kinase | 23C(125.0477) | NaN | X | NaN | NaN | NaN | NaN | 2.964828e+04 | 1.318798e+04 | 1.646029e+04 | NaN | SEGGSGGGAAGGGAGGAGAGAGCGSGGSSVGVR |
| ZZEF1 | ZZEF1\_HUMAN | Zinc finger ZZ-type and EF-hand domain-containing protein 1 | 2261 | 2261 | ZZEF1 | ZZEF1\_HUMAN | Zinc finger ZZ-type and EF-hand domain-containing protein 1 | 1C(125.0477) | NaN | X | NaN | NaN | NaN | NaN | 2.964437e+04 | 2.964437e+04 | NaN | NaN | CVYMDNANEPHNVIILK |
| DPYD | DPYD\_HUMAN | Dihydropyrimidine dehydrogenase [NADP(+)] | 29 | 22 | DPYD | DPYD\_HUMAN | Dihydropyrimidine dehydrogenase [NADP(+)] | 8C(125.0477) | NaN | X | NaN | NaN | NaN | NaN | 2.963953e+04 | 2.963953e+04 | NaN | NaN | TQTHATLCSTSAK |
| CTPS1 | PYRG1\_HUMAN | CTP synthase 1 | 491 | 491 | CTPS1 | PYRG1\_HUMAN | CTP synthase 1 | 1C(125.0477) | NaN | X | NaN | NaN | NaN | NaN | 2.961414e+04 | NaN | 2.961414e+04 | NaN | CLEEQGLK |
| NPTN | NPTN\_HUMAN | Neuroplastin | 259 | 246 | NPTN | NPTN\_HUMAN | Neuroplastin | 14C(125.0477) | NaN | X | NaN | NaN | NaN | NaN | 2.960278e+04 | NaN | 2.960278e+04 | NaN | SENKNEGQDATMYCK |
| TNS1 | TENS1\_HUMAN | Tensin-1 | 924 | 920 | TNS1 | TENS1\_HUMAN | Tensin-1 | 5C(125.0477) | NaN | X | NaN | NaN | NaN | NaN | 2.945604e+04 | 2.945604e+04 | NaN | NaN | SPVQCVSPELALTIALNPGGRPK |
| INF2 | INF2\_HUMAN | Inverted formin-2 | 332 | 322 | INF2 | INF2\_HUMAN | Inverted formin-2 | 11C(125.0477) | NaN | X | NaN | NaN | NaN | NaN | 2.945060e+04 | 2.945060e+04 | NaN | NaN | AVLLASDAQECTLEEVVER |
| ZFAND1 | ZFAN1\_HUMAN | AN1-type zinc finger protein 1 | 64 | 56 | ZFAND1 | ZFAN1\_HUMAN | AN1-type zinc finger protein 1 | 9C(125.0477) | NaN | X | NaN | NaN | NaN | NaN | 2.944071e+04 | 2.944071e+04 | NaN | NaN | TDQHTSYPCSFK |
| MARK2 | MARK2\_HUMAN | Serine/threonine-protein kinase MARK2 | 723 | 716 | MARK2 | MARK2\_HUMAN | Serine/threonine-protein kinase MARK2 | 8C(125.0477) | NaN | X | NaN | NaN | NaN | NaN | 2.942546e+04 | 2.942546e+04 | NaN | NaN | KVLDANSCQSELHEK |
| GSTM3 | GSTM3\_HUMAN | Glutathione S-transferase Mu 3 | 91 | 88 | GSTM3 | GSTM3\_HUMAN | Glutathione S-transferase Mu 3 | 4C(125.0477) | NaN | X | NaN | NaN | NaN | NaN | 2.939650e+04 | 2.939650e+04 | NaN | NaN | HNMCGETEEEKIR |
| ADSL | PUR8\_HUMAN | Adenylosuccinate lyase | 113 | 102 | ADSL | PUR8\_HUMAN | Adenylosuccinate lyase | 12C(125.0477) | NaN | X | NaN | NaN | NaN | NaN | 2.939147e+04 | 2.939147e+04 | NaN | NaN | AAGIIHLGATSCYVGDNTDLIILR |
| FASN | FAS\_HUMAN | Fatty acid synthase | 223 | 214 | FASN | FAS\_HUMAN | Fatty acid synthase | 10C(125.0477) | NaN | X | NaN | NaN | NaN | NaN | 2.939037e+04 | 2.939037e+04 | NaN | NaN | AFDTAGNGYCR |
| DCTN4 | DCTN4\_HUMAN | Dynactin subunit 4 | 418 | 406 | DCTN4 | DCTN4\_HUMAN | Dynactin subunit 4 | 13C(125.0477) | NaN | X | NaN | NaN | NaN | NaN | 2.938654e+04 | 2.938654e+04 | NaN | NaN | VTPQREEGEVTVCFK |
| ABCG2 | ABCG2\_HUMAN | Broad substrate specificity ATP-binding cassette transporter ABCG2 | 55 | 50 | ABCG2 | ABCG2\_HUMAN | Broad substrate specificity ATP-binding cassette transporter ABCG2 | 6C(125.0477) | NaN | X | NaN | NaN | NaN | NaN | 2.938419e+04 | NaN | 2.938419e+04 | NaN | SGFLPCR |
| GAK | GAK\_HUMAN | Cyclin-G-associated kinase | 691 | 686 | GAK | GAK\_HUMAN | Cyclin-G-associated kinase | 6C(125.0477) | NaN | X | NaN | NaN | NaN | NaN | 2.937636e+04 | NaN | 2.937636e+04 | NaN | YDLDACDIQEK |
| LACTB2 | LACB2\_HUMAN | Endoribonuclease LACTB2 | 90 | 81 | LACTB2 | LACB2\_HUMAN | Endoribonuclease LACTB2 | 10C(125.0477) | NaN | X | NaN | NaN | NaN | NaN | 2.933454e+04 | 2.933454e+04 | NaN | NaN | DHSGGIGDICK |
| S100A7 | S10A7\_HUMAN | Protein S100-A7 | 96 | 89 | S100A7 | S10A7\_HUMAN | Protein S100-A7 | 8C(125.0477) | NaN | X | NaN | NaN | NaN | NaN | 2.930116e+04 | NaN | 2.930116e+04 | NaN | QSHGAAPCSGGSQ |
| INF2 | INF2\_HUMAN | Inverted formin-2 | 1029 | 1029 | INF2 | INF2\_HUMAN | Inverted formin-2 | 1C(125.0477) | NaN | X | NaN | NaN | NaN | NaN | 2.928115e+04 | 2.204466e+04 | 7.236498e+03 | NaN | CPASEPGLDATTASESR |
| HUWE1 | HUWE1\_HUMAN | E3 ubiquitin-protein ligase HUWE1 | 1462 | 1461 | HUWE1 | HUWE1\_HUMAN | E3 ubiquitin-protein ligase HUWE1 | 2C(125.0477) | NaN | X | NaN | NaN | NaN | NaN | 2.927316e+04 | 2.927316e+04 | NaN | NaN | VCDLIMTAIK |
| CCT5 | TCPE\_HUMAN | T-complex protein 1 subunit epsilon | 181 | 177 | CCT5 | TCPE\_HUMAN | T-complex protein 1 subunit epsilon | 5C(125.0477) | NaN | X | NaN | NaN | NaN | NaN | 2.922628e+04 | 2.922628e+04 | NaN | NaN | VVNSCHR |
| KRT85 | KRT85\_HUMAN | Keratin, type II cuticular Hb5 | 12 | 8 | KRT85 | KRT85\_HUMAN | Keratin, type II cuticular Hb5 | 5C(125.0477) | NaN | X | NaN | NaN | NaN | NaN | 2.922434e+04 | NaN | NaN | 29224.3420 | ISSGCGVTR |
| SPTB | SPTB1\_HUMAN | Spectrin beta chain, erythrocytic | 1962 | 1956 | SPTB | SPTB1\_HUMAN | Spectrin beta chain, erythrocytic | 7C(125.0477) | NaN | X | NaN | NaN | NaN | NaN | 2.922156e+04 | 2.922156e+04 | NaN | NaN | SKNFSACLELGESLLQR |
| CASP9 | CASP9\_HUMAN | Caspase-9 | 287 | 281 | CASP9 | CASP9\_HUMAN | Caspase-9 | 7C(125.0477) | NaN | X | NaN | NaN | NaN | NaN | 2.921576e+04 | 2.921576e+04 | NaN | NaN | LFFIQACGGEQK |
| KPRP | KPRP\_HUMAN | Keratinocyte proline-rich protein | 16, 17 | 11 | KPRP | KPRP\_HUMAN | Keratinocyte proline-rich protein | 6C(57.0215) 7C(31.9988) | X | NaN | NaN | X | NaN | NaN | 2.920728e+04 | NaN | NaN | 29207.2750 | LPLQQCCVK |
| PSMB2 | PSB2\_HUMAN | Proteasome subunit beta type-2 | 163 | 162 | PSMB2 | PSB2\_HUMAN | Proteasome subunit beta type-2 | 2C(125.0477) | NaN | X | NaN | NaN | NaN | NaN | 2.919007e+04 | 2.919007e+04 | NaN | NaN | KCLEELQKR |
| DTYMK | KTHY\_HUMAN | Thymidylate kinase | 163 | 163 | DTYMK | KTHY\_HUMAN | Thymidylate kinase | 1C(125.0477) | NaN | X | NaN | NaN | NaN | NaN | 2.915291e+04 | 2.915291e+04 | NaN | NaN | CFHQLMK |
| CFB | CFAB\_HUMAN | Complement factor B | 191 | 183 | CFB | CFAB\_HUMAN | Complement factor B | 9C(57.0215) | X | NaN | NaN | NaN | NaN | NaN | 2.910431e+04 | NaN | 2.910431e+04 | NaN | LEDSVTYHCSR |
| CLIC1 | CLIC1\_HUMAN | Chloride intracellular channel protein 1 | 59 | 58 | CLIC1 | CLIC1\_HUMAN | Chloride intracellular channel protein 1 | 2C(125.0477) | NaN | X | NaN | NaN | NaN | NaN | 2.908871e+04 | 2.908871e+04 | NaN | NaN | LCPGGQLPFLLYGTEVHTDTNK |
| SPTB | SPTB1\_HUMAN | Spectrin beta chain, erythrocytic | 1552 | 1541 | SPTB | SPTB1\_HUMAN | Spectrin beta chain, erythrocytic | 12C(125.0477) | NaN | X | NaN | NaN | NaN | NaN | 2.905742e+04 | 2.905742e+04 | NaN | NaN | GQQLVEAAEIDCQDLEER |
| TEPSIN | AP4AT\_HUMAN | AP-4 complex accessory subunit Tepsin | 31 | 23 | TEPSIN | AP4AT\_HUMAN | AP-4 complex accessory subunit Tepsin | 9C(125.0477) | NaN | X | NaN | NaN | NaN | NaN | 2.897338e+04 | 2.897338e+04 | NaN | NaN | GTSDDDVPCPGYLFEEIAK |
| CNPY3 | CNPY3\_HUMAN | Protein canopy homolog 3 | 206 | 203 | CNPY3 | CNPY3\_HUMAN | Protein canopy homolog 3 | 4C(125.0477) | NaN | X | NaN | NaN | NaN | NaN | 2.894932e+04 | NaN | 2.894932e+04 | NaN | DTSCLAEQWSGK |
| PANK4 | PANK4\_HUMAN | 4'-phosphopantetheine phosphatase | 699 | 688 | PANK4 | PANK4\_HUMAN | 4'-phosphopantetheine phosphatase | 12C(125.0477) | NaN | X | NaN | NaN | NaN | NaN | 2.889644e+04 | 2.889644e+04 | NaN | NaN | LLLVQTGSSSPCLDLSR |
| DLG1 | DLG1\_HUMAN | Disks large homolog 1 | 275 | 272 | DLG1 | DLG1\_HUMAN | Disks large homolog 1 | 4C(125.0477) | NaN | X | NaN | NaN | NaN | NaN | 2.886907e+04 | NaN | 2.886907e+04 | NaN | VNDCILR |
| KIF1B | KIF1B\_HUMAN | Kinesin-like protein KIF1B | 356 | 356 | KIF1B | KIF1B\_HUMAN | Kinesin-like protein KIF1B | 1C(125.0477) | NaN | X | NaN | NaN | NaN | NaN | 2.884077e+04 | 2.884077e+04 | NaN | NaN | CNAVINEDPNAK |
| DARS1 | SYDC\_HUMAN | Aspartate--tRNA ligase, cytoplasmic | 203 | 198 | DARS1 | SYDC\_HUMAN | Aspartate--tRNA ligase, cytoplasmic | 6C(125.0477) | NaN | X | NaN | NaN | NaN | NaN | 2.883676e+04 | 2.883676e+04 | NaN | NaN | LQSGICHLFR |
| COPS6 | CSN6\_HUMAN | COP9 signalosome complex subunit 6 | 299 | 298 | COPS6 | CSN6\_HUMAN | COP9 signalosome complex subunit 6 | 2C(15.9949) | NaN | NaN | X | NaN | NaN | NaN | 2.882007e+04 | 2.882007e+04 | NaN | NaN | TCNTMNQFVNK |
| PFKM | PFKAM\_HUMAN | ATP-dependent 6-phosphofructokinase, muscle type | 114 | 108 | PFKM | PFKAM\_HUMAN | ATP-dependent 6-phosphofructokinase, muscle type | 7C(125.0477) | NaN | X | NaN | NaN | NaN | NaN | 2.875824e+04 | 2.875824e+04 | NaN | NaN | RGITNLCVIGGDGSLTGADTFR |
| RUVBL1 | RUVB1\_HUMAN | RuvB-like 1 | 94 | 91 | RUVBL1 | RUVB1\_HUMAN | RuvB-like 1 | 4C(125.0477) | NaN | X | NaN | NaN | NaN | NaN | 2.875752e+04 | 2.875752e+04 | NaN | NaN | VPFCPMVGSEVYSTEIK |
| LMAN1 | LMAN1\_HUMAN | Protein ERGIC-53 | 230 | 225 | LMAN1 | LMAN1\_HUMAN | Protein ERGIC-53 | 6C(125.0477) | NaN | X | NaN | NaN | NaN | NaN | 2.873191e+04 | NaN | 2.873191e+04 | NaN | NDYEFCAK |
| WDR45B | WIPI3\_HUMAN | WD repeat domain phosphoinositide-interacting protein 3 | 63 | 63 | WDR45B | WIPI3\_HUMAN | WD repeat domain phosphoinositide-interacting protein 3 | 1C(125.0477) | NaN | X | NaN | NaN | NaN | NaN | 2.872312e+04 | 2.872312e+04 | NaN | NaN | CNYLALVGGGK |
| SEMA7A | SEM7A\_HUMAN | Semaphorin-7A | 126 | 125 | SEMA7A | SEM7A\_HUMAN | Semaphorin-7A | 2C(125.0477) | NaN | X | NaN | NaN | NaN | NaN | 2.871907e+04 | NaN | 2.871907e+04 | NaN | DCENYITLLER |
| POMGNT2 | PMGT2\_HUMAN | Protein O-linked-mannose beta-1,4-N-acetylglucosaminyltransferase 2 | 490 | 490 | POMGNT2 | PMGT2\_HUMAN | Protein O-linked-mannose beta-1,4-N-acetylglucosaminyltransferase 2 | 1C(125.0477) | NaN | X | NaN | NaN | NaN | NaN | 2.853349e+04 | NaN | 2.853349e+04 | NaN | CQASVHGASEAR |
| UBR4 | UBR4\_HUMAN | E3 ubiquitin-protein ligase UBR4 | 260 | 259 | UBR4 | UBR4\_HUMAN | E3 ubiquitin-protein ligase UBR4 | 2C(125.0477) | NaN | X | NaN | NaN | NaN | NaN | 2.850213e+04 | 2.850213e+04 | NaN | NaN | VCLNLPYFLR |
| TXNRD1 | TRXR1\_HUMAN | Thioredoxin reductase 1, cytoplasmic | 625 | 613 | TXNRD1 | TRXR1\_HUMAN | Thioredoxin reductase 1, cytoplasmic | 13C(125.0477) | NaN | X | NaN | NaN | NaN | NaN | 2.848975e+04 | 2.848975e+04 | NaN | NaN | KQLDSTIGIHPVCAEVFTTLSVTK |
| CLASP2 | CLAP2\_HUMAN | CLIP-associating protein 2 | 1184 | 1182 | CLASP2 | CLAP2\_HUMAN | CLIP-associating protein 2 | 3C(125.0477) | NaN | X | NaN | NaN | NaN | NaN | 2.844650e+04 | 2.844650e+04 | NaN | NaN | VLCPIIQTADYPINLAAIK |
| RPS6KA3 | KS6A3\_HUMAN | Ribosomal protein S6 kinase alpha-3 | 229 | 225 | RPS6KA3 | KS6A3\_HUMAN | Ribosomal protein S6 kinase alpha-3 | 5C(125.0477) | NaN | X | NaN | NaN | NaN | NaN | 2.839493e+04 | 2.839493e+04 | NaN | NaN | AYSFCGTVEYMAPEVVNR |
| PHLPP2 | PHLP2\_HUMAN | PH domain leucine-rich repeat-containing protein phosphatase 2 | 402 | 396 | PHLPP2 | PHLP2\_HUMAN | PH domain leucine-rich repeat-containing protein phosphatase 2 | 7C(125.0477) | NaN | X | NaN | NaN | NaN | NaN | 2.834839e+04 | 2.834839e+04 | NaN | NaN | VVMAGNCLEVLNLGVLNR |
| SWAP70 | SWP70\_HUMAN | Switch-associated protein 70 | 40 | 34 | SWAP70 | SWP70\_HUMAN | Switch-associated protein 70 | 7C(125.0477) | NaN | X | NaN | NaN | NaN | NaN | 2.834631e+04 | 2.834631e+04 | NaN | NaN | VLSHNLCTVLK |
| PLOD3 | PLOD3\_HUMAN | Multifunctional procollagen lysine hydroxylase and glycosyltransferase LH3 | 691 | 683 | PLOD3 | PLOD3\_HUMAN | Multifunctional procollagen lysine hydroxylase and glycosyltransferase LH3 | 9C(125.0477) | NaN | X | NaN | NaN | NaN | NaN | 2.832460e+04 | NaN | 2.832460e+04 | NaN | GLDYEGGGCR |
| NF2 | MERL\_HUMAN | Merlin | 51 | 45 | NF2 | MERL\_HUMAN | Merlin | 7C(125.0477) | NaN | X | NaN | NaN | NaN | NaN | 2.823706e+04 | 2.823706e+04 | NaN | NaN | DLFDLVCR |
| GBF1 | GBF1\_HUMAN | Golgi-specific brefeldin A-resistance guanine nucleotide exchange factor 1 | 1173 | 1171 | GBF1 | GBF1\_HUMAN | Golgi-specific brefeldin A-resistance guanine nucleotide exchange factor 1 | 3C(125.0477) | NaN | X | NaN | NaN | NaN | NaN | 2.821763e+04 | 2.821763e+04 | NaN | NaN | VGCVWQTVR |
| MACROD1 | MACD1\_HUMAN | ADP-ribose glycohydrolase MACROD1 | 205 | 190 | MACROD1 | MACD1\_HUMAN | ADP-ribose glycohydrolase MACROD1 | 16C(15.9949) | NaN | NaN | X | NaN | NaN | NaN | 2.819351e+04 | 2.819351e+04 | NaN | NaN | AAGPLLTDECRTLQSCK |
| IPO13 | IPO13\_HUMAN | Importin-13 | 821 | 817 | IPO13 | IPO13\_HUMAN | Importin-13 | 5C(125.0477) | NaN | X | NaN | NaN | NaN | NaN | 2.819283e+04 | 2.819283e+04 | NaN | NaN | AVFQCAVLALK |
| ANKRD54 | ANR54\_HUMAN | Ankyrin repeat domain-containing protein 54 | 265 | 259 | ANKRD54 | ANR54\_HUMAN | Ankyrin repeat domain-containing protein 54 | 7C(125.0477) | NaN | X | NaN | NaN | NaN | NaN | 2.818103e+04 | 2.818103e+04 | NaN | NaN | ERLDDLCTR |
| CRTAP | CRTAP\_HUMAN | Cartilage-associated protein | 16, 16, 19, 19 | 7 | CRTAP | CRTAP\_HUMAN | Cartilage-associated protein | 10C(15.9949) 10C(31.9988) 13C(15.9949) 13C(31.9988) | NaN | NaN | X | X | NaN | NaN | 2.817200e+04 | NaN | 2.817200e+04 | NaN | GAAALLALLCVACALRAGR |
| SUMF1 | SUMF1\_HUMAN | Formylglycine-generating enzyme | 235 | 225 | SUMF1 | SUMF1\_HUMAN | Formylglycine-generating enzyme | 11C(125.0477) | NaN | X | NaN | NaN | NaN | NaN | 2.815096e+04 | NaN | 2.815096e+04 | NaN | LPTEAEWEYSCR |
| GMPR2 | GMPR2\_HUMAN | GMP reductase 2 | 127 | 125 | GMPR2 | GMPR2\_HUMAN | GMP reductase 2 | 3C(125.0477) | NaN | X | NaN | NaN | NaN | NaN | 2.814878e+04 | 2.814878e+04 | NaN | NaN | YICLDVANGYSEHFVEFVK |
| PYCR3 | P5CR3\_HUMAN | Pyrroline-5-carboxylate reductase 3 | 56 | 47 | PYCR3 | P5CR3\_HUMAN | Pyrroline-5-carboxylate reductase 3 | 10C(125.0477) | NaN | X | NaN | NaN | NaN | NaN | 2.812525e+04 | 2.812525e+04 | NaN | NaN | NLCHFQALGCR |
| DDI2 | DDI2\_HUMAN | Protein DDI1 homolog 2 | 361 | 339 | DDI2 | DDI2\_HUMAN | Protein DDI1 homolog 2 | 23C(15.9949) | NaN | NaN | X | NaN | NaN | NaN | 2.811763e+04 | 2.811763e+04 | NaN | NaN | NVLVIGTTGSQTTFLPEGELPECARLAYGAGR |
| VPS13A | VP13A\_HUMAN | Vacuolar protein sorting-associated protein 13A | 1979 | 1976 | VPS13A | VP13A\_HUMAN | Vacuolar protein sorting-associated protein 13A | 4C(125.0477) | NaN | X | NaN | NaN | NaN | NaN | 2.811358e+04 | NaN | 2.811358e+04 | NaN | SIVCQIDTVEGSK |
| NUP50 | NUP50\_HUMAN | Nuclear pore complex protein Nup50 | 181 | 174 | NUP50 | NUP50\_HUMAN | Nuclear pore complex protein Nup50 | 8C(125.0477) | NaN | X | NaN | NaN | NaN | NaN | 2.809847e+04 | 2.809847e+04 | NaN | NaN | HVNTNPLCDLTPIFK |
| XPO1 | XPO1\_HUMAN | Exportin-1 | 119 | 113 | XPO1 | XPO1\_HUMAN | Exportin-1 | 7C(125.0477) | NaN | X | NaN | NaN | NaN | NaN | 2.803312e+04 | NaN | 2.803312e+04 | NaN | TSSDPTCVEK |
| TRIM10 | TRI10\_HUMAN | Tripartite motif-containing protein 10 | 353 | 351 | TRIM10 | TRI10\_HUMAN | Tripartite motif-containing protein 10 | 3C(125.0477) | NaN | X | NaN | NaN | NaN | NaN | 2.802933e+04 | 2.802933e+04 | NaN | NaN | ATCVLAHTGITGGR |
| C3 | CO3\_HUMAN | Complement C3 | 627 | 623 | C3 | CO3\_HUMAN | Complement C3 | 5C(57.0215) | X | NaN | NaN | NaN | NaN | NaN | 2.796765e+04 | NaN | 2.796765e+04 | NaN | ADIGCTPGSGK |
| GTPBP1 | GTPB1\_HUMAN | GTP-binding protein 1 | 515 | 509 | GTPBP1 | GTPB1\_HUMAN | GTP-binding protein 1 | 7C(125.0477) | NaN | X | NaN | NaN | NaN | NaN | 2.793459e+04 | 2.793459e+04 | NaN | NaN | YQAMVHCGSIR |
| FRYL | FRYL\_HUMAN | Protein furry homolog-like | 2369 | 2362 | FRYL | FRYL\_HUMAN | Protein furry homolog-like | 8C(125.0477) | NaN | X | NaN | NaN | NaN | NaN | 2.793269e+04 | 2.793269e+04 | NaN | NaN | LMNVLSLCGPESGLPK |
| JMJD6 | JMJD6\_HUMAN | Bifunctional arginine demethylase and lysyl-hydroxylase JMJD6 | 101 | 101 | JMJD6 | JMJD6\_HUMAN | Bifunctional arginine demethylase and lysyl-hydroxylase JMJD6 | 1C(125.0477) | NaN | X | NaN | NaN | NaN | NaN | 2.791951e+04 | 2.791951e+04 | NaN | NaN | CGEDNDGYSVK |
| NHLRC2 | NHLC2\_HUMAN | NHL repeat-containing protein 2 | 716 | 700 | NHLRC2 | NHLC2\_HUMAN | NHL repeat-containing protein 2 | 17C(125.0477) | NaN | X | NaN | NaN | NaN | NaN | 2.779824e+04 | 2.779824e+04 | NaN | NaN | AILFSQPLQITDTQQGCIAPVELR |
| SKIV2L | SKIV2\_HUMAN | Helicase SKI2W | 719 | 708 | SKIV2L | SKIV2\_HUMAN | Helicase SKI2W | 12C(125.0477) | NaN | X | NaN | NaN | NaN | NaN | 2.779083e+04 | 2.779083e+04 | NaN | NaN | GLDPTGTVILLCK |
| UBE2V2 | UB2V2\_HUMAN | Ubiquitin-conjugating enzyme E2 variant 2 | 69 | 62 | UBE2V2 | UB2V2\_HUMAN | Ubiquitin-conjugating enzyme E2 variant 2 | 8C(125.0477) | NaN | X | NaN | NaN | NaN | NaN | 2.771891e+04 | 2.771891e+04 | NaN | NaN | IYSLKVECGPK |
| MTOR | MTOR\_HUMAN | Serine/threonine-protein kinase mTOR | 216 | 215 | MTOR | MTOR\_HUMAN | Serine/threonine-protein kinase mTOR | 2C(125.0477) | NaN | X | NaN | NaN | NaN | NaN | 2.769626e+04 | 2.769626e+04 | NaN | NaN | ACLILTTQREPK |
| C11orf68 | CK068\_HUMAN | UPF0696 protein C11orf68 | 214 | 211 | C11orf68 | CK068\_HUMAN | UPF0696 protein C11orf68 | 4C(125.0477) | NaN | X | NaN | NaN | NaN | NaN | 2.767655e+04 | 2.767655e+04 | NaN | NaN | QVICVYTDDFTDR |
| YWHAQ | 1433T\_HUMAN | 14-3-3 protein theta | 94 | 92 | YWHAQ | 1433T\_HUMAN | 14-3-3 protein theta | 3C(125.0477) | NaN | X | NaN | NaN | NaN | NaN | 2.764482e+04 | 2.764482e+04 | NaN | NaN | SICTTVLELLDK |
| RAB3IL1 | R3GEF\_HUMAN | Guanine nucleotide exchange factor for Rab-3A | 29 | 25 | RAB3IL1 | R3GEF\_HUMAN | Guanine nucleotide exchange factor for Rab-3A | 5C(125.0477) | NaN | X | NaN | NaN | NaN | NaN | 2.757915e+04 | 2.757915e+04 | NaN | NaN | STDPCQGHR |
| TTLL12 | TTL12\_HUMAN | Tubulin--tyrosine ligase-like protein 12 | 361 | 347 | TTLL12 | TTL12\_HUMAN | Tubulin--tyrosine ligase-like protein 12 | 15C(125.0477) | NaN | X | NaN | NaN | NaN | NaN | 2.756303e+04 | 2.756303e+04 | NaN | NaN | LSQERPGVLLNQFPCENLLTVK |
| C8A | CO8A\_HUMAN | Complement component C8 alpha chain | 528 | 521 | C8A | CO8A\_HUMAN | Complement component C8 alpha chain | 8C(57.0215) | X | NaN | NaN | NaN | NaN | NaN | 2.755309e+04 | NaN | 2.755309e+04 | NaN | LGSLGAACEQTQTEGAK |
| PTPN4 | PTN4\_HUMAN | Tyrosine-protein phosphatase non-receptor type 4 | 301 | 296 | PTPN4 | PTN4\_HUMAN | Tyrosine-protein phosphatase non-receptor type 4 | 6C(31.9988) | NaN | NaN | NaN | X | NaN | NaN | 2.753403e+04 | 2.753403e+04 | NaN | NaN | NLWKACVEHHTFFR |
| UBR1 | UBR1\_HUMAN | E3 ubiquitin-protein ligase UBR1 | 589 | 584 | UBR1 | UBR1\_HUMAN | E3 ubiquitin-protein ligase UBR1 | 6C(125.0477) | NaN | X | NaN | NaN | NaN | NaN | 2.752229e+04 | 2.752229e+04 | NaN | NaN | TVVQSCGHSLETK |
| RABEP2 | RABE2\_HUMAN | Rab GTPase-binding effector protein 2 | 115 | 109 | RABEP2 | RABE2\_HUMAN | Rab GTPase-binding effector protein 2 | 7C(125.0477) | NaN | X | NaN | NaN | NaN | NaN | 2.748407e+04 | 2.748407e+04 | NaN | NaN | QQQQQDCEEKER |
| VAC14 | VAC14\_HUMAN | Protein VAC14 homolog | 674 | 668 | VAC14 | VAC14\_HUMAN | Protein VAC14 homolog | 7C(125.0477) | NaN | X | NaN | NaN | NaN | NaN | 2.730734e+04 | 2.730734e+04 | NaN | NaN | LVQLIECPIFTYLR |
| NCAPG | CND3\_HUMAN | Condensin complex subunit 3 | 202 | 198 | NCAPG | CND3\_HUMAN | Condensin complex subunit 3 | 5C(125.0477) | NaN | X | NaN | NaN | NaN | NaN | 2.729244e+04 | 2.729244e+04 | NaN | NaN | AVLSCIAPSAK |
| LCP1 | PLSL\_HUMAN | Plastin-2 | 336 | 334 | LCP1 | PLSL\_HUMAN | Plastin-2 | 3C(125.0477) | NaN | X | NaN | NaN | NaN | NaN | 2.726256e+04 | NaN | 2.726256e+04 | NaN | AECMLQQAER |
| ACTBL2 | ACTBL\_HUMAN | Beta-actin-like protein 2 | 53 | 41 | ACTBL2 | ACTBL\_HUMAN | Beta-actin-like protein 2 | 13C(15.9949) | NaN | NaN | X | NaN | NaN | NaN | 2.726142e+04 | NaN | NaN | 27261.4220 | HQGVMVGMGQKDCYVGDEAQSK |
| TTC38 | TTC38\_HUMAN | Tetratricopeptide repeat protein 38 | 128 | 127 | TTC38 | TTC38\_HUMAN | Tetratricopeptide repeat protein 38 | 2C(125.0477) | NaN | X | NaN | NaN | NaN | NaN | 2.725878e+04 | 2.725878e+04 | NaN | NaN | ACELWEQILQDHPTDMLALK |
| IMPACT | IMPCT\_HUMAN | Protein IMPACT | 236 | 230 | IMPACT | IMPCT\_HUMAN | Protein IMPACT | 7C(125.0477) | NaN | X | NaN | NaN | NaN | NaN | 2.723634e+04 | 2.723634e+04 | NaN | NaN | QTFLQDCEDDGETAAGGR |
| CAP1 | CAP1\_HUMAN | Adenylyl cyclase-associated protein 1 | 375 | 365 | CAP1 | CAP1\_HUMAN | Adenylyl cyclase-associated protein 1 | 11C(125.0477) | NaN | X | NaN | NaN | NaN | NaN | 2.719535e+04 | 2.719535e+04 | NaN | NaN | GKINSITVDNCK |
| EIF3M | EIF3M\_HUMAN | Eukaryotic translation initiation factor 3 subunit M | 134 | 130 | EIF3M | EIF3M\_HUMAN | Eukaryotic translation initiation factor 3 subunit M | 5C(125.0477) | NaN | X | NaN | NaN | NaN | NaN | 2.719383e+04 | 2.719383e+04 | NaN | NaN | VAASCGAIQYIPTELDQVR |
| A1BG | A1BG\_HUMAN | Alpha-1B-glycoprotein | 279 | 262 | A1BG | A1BG\_HUMAN | Alpha-1B-glycoprotein | 18C(57.0215) | X | NaN | NaN | NaN | NaN | NaN | 2.717669e+04 | 2.717669e+04 | NaN | NaN | IFFHLNAVALGDGGHYTCR |
| EIF2S1 | IF2A\_HUMAN | Eukaryotic translation initiation factor 2 subunit 1 | 70 | 68 | EIF2S1 | IF2A\_HUMAN | Eukaryotic translation initiation factor 2 subunit 1 | 3C(125.0477) | NaN | X | NaN | NaN | NaN | NaN | 2.716572e+04 | NaN | 2.716572e+04 | NaN | NECVVVIR |
| TATDN1 | TATD1\_HUMAN | Deoxyribonuclease TATDN1 | 150 | 144 | TATDN1 | TATD1\_HUMAN | Deoxyribonuclease TATDN1 | 7C(125.0477) | NaN | X | NaN | NaN | NaN | NaN | 2.716497e+04 | 2.716497e+04 | NaN | NaN | LPMFLHCR |
| CXADR | CXAR\_HUMAN | Coxsackievirus and adenovirus receptor | 223 | 218 | CXADR | CXAR\_HUMAN | Coxsackievirus and adenovirus receptor | 6C(125.0477) | NaN | X | NaN | NaN | NaN | NaN | 2.714390e+04 | NaN | 2.714390e+04 | NaN | VGSDQCLLR |
| VPS13C | VP13C\_HUMAN | Vacuolar protein sorting-associated protein 13C | 2925 | 2922 | VPS13C | VP13C\_HUMAN | Vacuolar protein sorting-associated protein 13C | 4C(125.0477) | NaN | X | NaN | NaN | NaN | NaN | 2.709519e+04 | 2.709519e+04 | NaN | NaN | VVGCEGSSKPFFYNR |
| ARF6 | ARF6\_HUMAN | ADP-ribosylation factor 6 | 90 | 76 | ARF6 | ARF6\_HUMAN | ADP-ribosylation factor 6 | 15C(125.0477) | NaN | X | NaN | NaN | NaN | NaN | 2.707691e+04 | 2.707691e+04 | NaN | NaN | HYYTGTQGLIFVVDCADRDR |
| AKR1A1 | AK1A1\_HUMAN | Aldo-keto reductase family 1 member A1 | 46 | 43 | AKR1A1 | AK1A1\_HUMAN | Aldo-keto reductase family 1 member A1 | 4C(125.0477) | NaN | X | NaN | NaN | NaN | NaN | 2.705948e+04 | 2.705948e+04 | NaN | NaN | HIDCAAIYGNEPEIGEALK |
| ISG20 | ISG20\_HUMAN | Interferon-stimulated gene 20 kDa protein | 28 | 28 | ISG20 | ISG20\_HUMAN | Interferon-stimulated gene 20 kDa protein | 1C(125.0477) | NaN | X | NaN | NaN | NaN | NaN | 2.699602e+04 | 2.699602e+04 | NaN | NaN | CSLVNVHGAVLYDK |
| NCAPG | CND3\_HUMAN | Condensin complex subunit 3 | 509 | 504 | NCAPG | CND3\_HUMAN | Condensin complex subunit 3 | 6C(125.0477) | NaN | X | NaN | NaN | NaN | NaN | 2.696554e+04 | 2.696554e+04 | NaN | NaN | EALENCITLQDFNR |
| EIF2S1 | IF2A\_HUMAN | Eukaryotic translation initiation factor 2 subunit 1 | 70 | 65 | EIF2S1 | IF2A\_HUMAN | Eukaryotic translation initiation factor 2 subunit 1 | 6C(125.0477) | NaN | X | NaN | NaN | NaN | NaN | 2.694792e+04 | 2.694792e+04 | NaN | NaN | IGRNECVVVIR |
| WASF2 | WASF2\_HUMAN | Wiskott-Aldrich syndrome protein family member 2 | 27 | 23 | WASF2 | WASF2\_HUMAN | Wiskott-Aldrich syndrome protein family member 2 | 5C(125.0477) | NaN | X | NaN | NaN | NaN | NaN | 2.689926e+04 | 2.689926e+04 | NaN | NaN | SELECVTNITLANVIR |
| CAP1 | CAP1\_HUMAN | Adenylyl cyclase-associated protein 1 | 93 | 85 | CAP1 | CAP1\_HUMAN | Adenylyl cyclase-associated protein 1 | 9C(125.0477) | NaN | X | NaN | NaN | NaN | NaN | 2.687642e+04 | 2.687642e+04 | NaN | NaN | ALLVTASQCQQPAENK |
| PUS1 | TRUA\_HUMAN | tRNA pseudouridine synthase A | 260 | 253 | PUS1 | TRUA\_HUMAN | tRNA pseudouridine synthase A | 8C(125.0477) | NaN | X | NaN | NaN | NaN | NaN | 2.685993e+04 | 2.685993e+04 | NaN | NaN | GPQDPSACR |
| OSTF1 | OSTF1\_HUMAN | Osteoclast-stimulating factor 1 | 185 | 175 | OSTF1 | OSTF1\_HUMAN | Osteoclast-stimulating factor 1 | 11C(125.0477) | NaN | X | NaN | NaN | NaN | NaN | 2.682999e+04 | 2.682999e+04 | NaN | NaN | LAFDMATNAACASLLK |
| ATG7 | ATG7\_HUMAN | Ubiquitin-like modifier-activating enzyme ATG7 | 322 | 316 | ATG7 | ATG7\_HUMAN | Ubiquitin-like modifier-activating enzyme ATG7 | 7C(125.0477) | NaN | X | NaN | NaN | NaN | NaN | 2.678813e+04 | 2.678813e+04 | NaN | NaN | MVNLSECMDPKR |
| TCP1 | TCPA\_HUMAN | T-complex protein 1 subunit alpha | 385 | 379 | TCP1 | TCPA\_HUMAN | T-complex protein 1 subunit alpha | 7C(31.9988) | NaN | NaN | NaN | X | NaN | NaN | 2.676087e+04 | 2.676087e+04 | NaN | NaN | GANDFMCDEMERSLHDALCVVK |
| SERPINA3 | AACT\_HUMAN | Alpha-1-antichymotrypsin | 261 | 256 | SERPINA3 | AACT\_HUMAN | Alpha-1-antichymotrypsin | 6C(57.0215) | X | NaN | NaN | NaN | NaN | NaN | 2.674894e+04 | NaN | 2.674894e+04 | NaN | DEELSCTVVELK |
| NAPB | SNAB\_HUMAN | Beta-soluble NSF attachment protein | 212 | 209 | NAPB | SNAB\_HUMAN | Beta-soluble NSF attachment protein | 4C(125.0477) | NaN | X | NaN | NaN | NaN | NaN | 2.672334e+04 | 2.672334e+04 | NaN | NaN | AALCHFIVDELNAK |
| SLC19A1 | S19A1\_HUMAN | Reduced folate transporter | 220, 220 | 220 | SLC19A1 | S19A1\_HUMAN | Reduced folate transporter | 1C(125.0477) 1C(31.9988) | NaN | X | NaN | X | NaN | NaN | 2.671746e+04 | NaN | NaN | 26717.4550 | CETSASELER |
| UBR1 | UBR1\_HUMAN | E3 ubiquitin-protein ligase UBR1 | 180 | 180 | UBR1 | UBR1\_HUMAN | E3 ubiquitin-protein ligase UBR1 | 1C(125.0477) | NaN | X | NaN | NaN | NaN | NaN | 2.669870e+04 | 2.669870e+04 | NaN | NaN | CPLNEEVIVQAR |
| INPP5D | SHIP1\_HUMAN | Phosphatidylinositol 3,4,5-trisphosphate 5-phosphatase 1 | 1088 | 1085 | INPP5D | SHIP1\_HUMAN | Phosphatidylinositol 3,4,5-trisphosphate 5-phosphatase 1 | 4C(125.0477) | NaN | X | NaN | NaN | NaN | NaN | 2.669738e+04 | NaN | 2.669738e+04 | NaN | SFTCSSSAEGR |
| HTT | HD\_HUMAN | Huntingtin | 664 | 651 | HTT | HD\_HUMAN | Huntingtin | 14C(125.0477) | NaN | X | NaN | NaN | NaN | NaN | 2.665229e+04 | 2.665229e+04 | NaN | NaN | DEATEPGDQENKPCR |
| CCT3 | TCPG\_HUMAN | T-complex protein 1 subunit gamma | 366 | 354 | CCT3 | TCPG\_HUMAN | T-complex protein 1 subunit gamma | 13C(125.0477) | NaN | X | NaN | NaN | NaN | NaN | 2.662421e+04 | 2.662421e+04 | NaN | NaN | KIGDEYFTFITDCKDPK |
| KRT86 | KRT86\_HUMAN | Keratin, type II cuticular Hb6 | 217 | 214 | KRT86 | KRT86\_HUMAN | Keratin, type II cuticular Hb6 | 4C(125.0477) | NaN | X | NaN | NaN | NaN | NaN | 2.660538e+04 | NaN | NaN | 26605.3750 | DVDCAYLR |
| VTA1 | VTA1\_HUMAN | Vacuolar protein sorting-associated protein VTA1 homolog | 56 | 49 | VTA1 | VTA1\_HUMAN | Vacuolar protein sorting-associated protein VTA1 homolog | 8C(125.0477) | NaN | X | NaN | NaN | NaN | NaN | 2.654882e+04 | 2.654882e+04 | NaN | NaN | IDSKTPECRK |
| CRYL1 | CRYL1\_HUMAN | Lambda-crystallin homolog | 141 | 140 | CRYL1 | CRYL1\_HUMAN | Lambda-crystallin homolog | 2C(125.0477) | NaN | X | NaN | NaN | NaN | NaN | 2.652288e+04 | 2.652288e+04 | NaN | NaN | QCIVAHPVNPPYYIPLVELVPHPETAPTTVDR |
| CAPZA1 | CAZA1\_HUMAN | F-actin-capping protein subunit alpha-1 | 141 | 130 | CAPZA1 | CAZA1\_HUMAN | F-actin-capping protein subunit alpha-1 | 12C(125.0477) | NaN | X | NaN | NaN | NaN | NaN | 2.651881e+04 | 2.651881e+04 | NaN | NaN | AYVKDHYSNGFCTVYAK |
| DDX19B | DD19B\_HUMAN | ATP-dependent RNA helicase DDX19B | 393 | 386 | DDX19B | DD19B\_HUMAN | ATP-dependent RNA helicase DDX19B | 8C(125.0477) | NaN | X | NaN | NaN | NaN | NaN | 2.651649e+04 | 2.651649e+04 | NaN | NaN | VLVTTNVCAR |
| ASAP1 | ASAP1\_HUMAN | Arf-GAP with SH3 domain, ANK repeat and PH domain-containing protein 1 | 248 | 243 | ASAP1 | ASAP1\_HUMAN | Arf-GAP with SH3 domain, ANK repeat and PH domain-containing protein 1 | 6C(125.0477) | NaN | X | NaN | NaN | NaN | NaN | 2.650908e+04 | 2.650908e+04 | NaN | NaN | YYHAQCNFFQDGLK |
| ADRM1 | ADRM1\_HUMAN | Proteasomal ubiquitin receptor ADRM1 | 80 | 65 | ADRM1 | ADRM1\_HUMAN | Proteasomal ubiquitin receptor ADRM1 | 16C(15.9949) | NaN | NaN | X | NaN | NaN | NaN | 2.650858e+04 | 2.650858e+04 | NaN | NaN | TSGNVEDDLIIFPDDCEFKRVPQCPSGR |
| DCTPP1 | DCTP1\_HUMAN | dCTP pyrophosphatase 1 | 110 | 110 | DCTPP1 | DCTP1\_HUMAN | dCTP pyrophosphatase 1 | 1C(125.0477) | NaN | X | NaN | NaN | NaN | NaN | 2.643920e+04 | 2.643920e+04 | NaN | NaN | CRVDLPLAVLSK |
| HSPD1 | CH60\_HUMAN | 60 kDa heat shock protein, mitochondrial | 447 | 447 | HSPD1 | CH60\_HUMAN | 60 kDa heat shock protein, mitochondrial | 1C(125.0477) | NaN | X | NaN | NaN | NaN | NaN | 2.639624e+04 | 2.639624e+04 | NaN | NaN | CIPALDSLTPANEDQK |
| SEPTIN2 | SEPT2\_HUMAN | Septin-2 | 111 | 97 | SEPTIN2 | SEPT2\_HUMAN | Septin-2 | 15C(31.9988) | NaN | NaN | NaN | X | NaN | NaN | 2.632213e+04 | NaN | 2.632213e+04 | NaN | LTVVDTPGYGDAINCR |
| IPO11 | IPO11\_HUMAN | Importin-11 | 320 | 316 | IPO11 | IPO11\_HUMAN | Importin-11 | 5C(125.0477) | NaN | X | NaN | NaN | NaN | NaN | 2.631237e+04 | 2.631237e+04 | NaN | NaN | FIVQCMNLIK |
| NME1 | NDKA\_HUMAN | Nucleoside diphosphate kinase A | 109 | 106 | NME1 | NDKA\_HUMAN | Nucleoside diphosphate kinase A | 4C(125.0477) | NaN | X | NaN | NaN | NaN | NaN | 2.627410e+04 | 2.627410e+04 | NaN | NaN | GDFCIQVGR |
| GBF1 | GBF1\_HUMAN | Golgi-specific brefeldin A-resistance guanine nucleotide exchange factor 1 | 1414 | 1402 | GBF1 | GBF1\_HUMAN | Golgi-specific brefeldin A-resistance guanine nucleotide exchange factor 1 | 13C(125.0477) | NaN | X | NaN | NaN | NaN | NaN | 2.626746e+04 | 2.626746e+04 | NaN | NaN | DAAHITPDNFELCVK |
| PUDP | HDHD1\_HUMAN | Pseudouridine-5'-phosphatase | 34 | 25 | PUDP | HDHD1\_HUMAN | Pseudouridine-5'-phosphatase | 10C(125.0477) | NaN | X | NaN | NaN | NaN | NaN | 2.625014e+04 | 2.625014e+04 | NaN | NaN | LYSVVFQEICNRYDK |
| USP24 | UBP24\_HUMAN | Ubiquitin carboxyl-terminal hydrolase 24 | 172 | 163 | USP24 | UBP24\_HUMAN | Ubiquitin carboxyl-terminal hydrolase 24 | 10C(125.0477) | NaN | X | NaN | NaN | NaN | NaN | 2.624310e+04 | 2.624310e+04 | NaN | NaN | LGLSESDENCRR |
| SEMA7A | SEM7A\_HUMAN | Semaphorin-7A | 518 | 518 | SEMA7A | SEM7A\_HUMAN | Semaphorin-7A | 1C(125.0477) | NaN | X | NaN | NaN | NaN | NaN | 2.621374e+04 | NaN | 2.621374e+04 | NaN | CISIYSSER |
| UGGT1 | UGGG1\_HUMAN | UDP-glucose:glycoprotein glucosyltransferase 1 | 880 | 870 | UGGT1 | UGGG1\_HUMAN | UDP-glucose:glycoprotein glucosyltransferase 1 | 11C(125.0477) | NaN | X | NaN | NaN | NaN | NaN | 2.618429e+04 | 2.618429e+04 | NaN | NaN | MDFILSHAVYCR |
| UBE3C | UBE3C\_HUMAN | Ubiquitin-protein ligase E3C | 1018 | 1014 | UBE3C | UBE3C\_HUMAN | Ubiquitin-protein ligase E3C | 5C(125.0477) | NaN | X | NaN | NaN | NaN | NaN | 2.604938e+04 | 2.604938e+04 | NaN | NaN | FVTSCSRPPLLGFK |
| DDX17 | DDX17\_HUMAN | Probable ATP-dependent RNA helicase DDX17 | 247 | 242 | DDX17 | DDX17\_HUMAN | Probable ATP-dependent RNA helicase DDX17 | 6C(125.0477) | NaN | X | NaN | NaN | NaN | NaN | 2.603987e+04 | 2.603987e+04 | NaN | NaN | GDGPICLVLAPTR |
| KRT72 | K2C72\_HUMAN | Keratin, type II cytoskeletal 72 | 364, 364 | 363 | KRT72 | K2C72\_HUMAN | Keratin, type II cytoskeletal 72 | 2C(15.9949) 2C(57.0215) | X | NaN | X | NaN | NaN | NaN | 2.601026e+04 | NaN | NaN | 26010.2580 | QCADLETAIADAEQR |
| DSTN | DEST\_HUMAN | Destrin | 135 | 128 | DSTN | DEST\_HUMAN | Destrin | 8C(15.9949) | NaN | NaN | X | NaN | NaN | NaN | 2.594682e+04 | 2.594682e+04 | NaN | NaN | FQGIKHECQANGPEDLNRACIAEK |
| TGM2 | TGM2\_HUMAN | Protein-glutamine gamma-glutamyltransferase 2 | 230, 230 | 223 | TGM2 | TGM2\_HUMAN | Protein-glutamine gamma-glutamyltransferase 2 | 8C(125.0477) 8C(15.9949) | NaN | X | X | NaN | NaN | NaN | 2.593485e+04 | 2.593485e+04 | NaN | NaN | VVSGMVNCNDDQGVLLGR |
| SERINC3 | SERC3\_HUMAN | Serine incorporator 3 | 72 | 68 | SERINC3 | SERC3\_HUMAN | Serine incorporator 3 | 5C(125.0477) | NaN | X | NaN | NaN | NaN | NaN | 2.586943e+04 | NaN | 2.586943e+04 | NaN | IPGFCEGGFK |
| ZW10 | ZW10\_HUMAN | Centromere/kinetochore protein zw10 homolog | 650 | 636 | ZW10 | ZW10\_HUMAN | Centromere/kinetochore protein zw10 homolog | 15C(125.0477) | NaN | X | NaN | NaN | NaN | NaN | 2.586658e+04 | 2.586658e+04 | NaN | NaN | LGIVWQDVLPVNIYCK |
| HBG2 | HBG2\_HUMAN | Hemoglobin subunit gamma-2 | 94 | 68 | HBG2 | HBG2\_HUMAN | Hemoglobin subunit gamma-2 | 27C(125.0477) | NaN | X | NaN | NaN | NaN | NaN | 2.582515e+04 | 2.582515e+04 | NaN | NaN | VLTSLGDAIKHLDDLKGTFAQLSELHCDK |
| DNM1L | DNM1L\_HUMAN | Dynamin-1-like protein | 446 | 442 | DNM1L | DNM1L\_HUMAN | Dynamin-1-like protein | 5C(125.0477) | NaN | X | NaN | NaN | NaN | NaN | 2.581778e+04 | 2.581778e+04 | NaN | NaN | IIQHCSNYSTQELLR |
| PFAS | PUR4\_HUMAN | Phosphoribosylformylglycinamidine synthase | 502 | 501 | PFAS | PUR4\_HUMAN | Phosphoribosylformylglycinamidine synthase | 2C(125.0477) | NaN | X | NaN | NaN | NaN | NaN | 2.581085e+04 | 2.581085e+04 | NaN | NaN | ACVEAPK |
| PABPC1 | PABP1\_HUMAN | Polyadenylate-binding protein 1 | 339 | 334 | PABPC1 | PABP1\_HUMAN | Polyadenylate-binding protein 1 | 6C(125.0477) | NaN | X | NaN | NaN | NaN | NaN | 2.580450e+04 | 2.580450e+04 | NaN | NaN | GFGFVCFSSPEEATK |
| PTPRA | PTPRA\_HUMAN | Receptor-type tyrosine-protein phosphatase alpha | 743 | 739 | PTPRA | PTPRA\_HUMAN | Receptor-type tyrosine-protein phosphatase alpha | 5C(125.0477) | NaN | X | NaN | NaN | NaN | NaN | 2.578012e+04 | 2.578012e+04 | NaN | NaN | TGTFCALSTVLER |
| WDFY1 | WDFY1\_HUMAN | WD repeat and FYVE domain-containing protein 1 | 204 | 197 | WDFY1 | WDFY1\_HUMAN | WD repeat and FYVE domain-containing protein 1 | 8C(125.0477) | NaN | X | NaN | NaN | NaN | NaN | 2.576562e+04 | 2.576562e+04 | NaN | NaN | GHEGSVACLWWDPIQR |
| ATP6V1B2 | VATB2\_HUMAN | V-type proton ATPase subunit B, brain isoform | 162 | 142 | ATP6V1B2 | VATB2\_HUMAN | V-type proton ATPase subunit B, brain isoform | 21C(125.0477) | NaN | X | NaN | NaN | NaN | NaN | 2.575503e+04 | 2.575503e+04 | NaN | NaN | GPVVLAEDFLDIMGQPINPQCR |
| LRSAM1 | LRSM1\_HUMAN | E3 ubiquitin-protein ligase LRSAM1 | 75 | 74 | LRSAM1 | LRSM1\_HUMAN | E3 ubiquitin-protein ligase LRSAM1 | 2C(125.0477) | NaN | X | NaN | NaN | NaN | NaN | 2.571522e+04 | 2.571522e+04 | NaN | NaN | SCSLLSLATIK |
| MARK2 | MARK2\_HUMAN | Serine/threonine-protein kinase MARK2 | 723 | 717 | MARK2 | MARK2\_HUMAN | Serine/threonine-protein kinase MARK2 | 7C(125.0477) | NaN | X | NaN | NaN | NaN | NaN | 2.569331e+04 | 2.569331e+04 | NaN | NaN | VLDANSCQSELHEK |
| ATP1B2 | AT1B2\_HUMAN | Sodium/potassium-transporting ATPase subunit beta-2 | 10 | 9 | ATP1B2 | AT1B2\_HUMAN | Sodium/potassium-transporting ATPase subunit beta-2 | 2C(125.0477) | NaN | X | NaN | NaN | NaN | NaN | 2.564778e+04 | NaN | 2.564778e+04 | NaN | SCGQVVEEWK |
| C4A | CO4A\_HUMAN | Complement C4-A | 820 | 818 | C4A | CO4A\_HUMAN | Complement C4-A | 3C(57.0215) | X | NaN | NaN | NaN | NaN | NaN | 2.561209e+04 | NaN | 2.561209e+04 | NaN | GLCVATPVQLR |
| NCBP1 | NCBP1\_HUMAN | Nuclear cap-binding protein subunit 1 | 320 | 312 | NCBP1 | NCBP1\_HUMAN | Nuclear cap-binding protein subunit 1 | 9C(125.0477) | NaN | X | NaN | NaN | NaN | NaN | 2.559591e+04 | 2.559591e+04 | NaN | NaN | FVIEENLHCIIK |
| GSTO1 | GSTO1\_HUMAN | Glutathione S-transferase omega-1 | 90 | 66 | GSTO1 | GSTO1\_HUMAN | Glutathione S-transferase omega-1 | 25C(125.0477) | NaN | X | NaN | NaN | NaN | NaN | 2.557825e+04 | 2.557825e+04 | NaN | NaN | KNPFGLVPVLENSQGQLIYESAITCEYLDEAYPGK |
| AMBP | AMBP\_HUMAN | Protein AMBP | 188 | 186 | AMBP | AMBP\_HUMAN | Protein AMBP | 3C(125.0477) | NaN | X | NaN | NaN | NaN | NaN | 2.556476e+04 | NaN | 2.556476e+04 | NaN | GECVPGEQEPEPILIPR |
| NAXD | NNRD\_HUMAN | ATP-dependent (S)-NAD(P)H-hydrate dehydratase | 82 | 76 | NAXD | NNRD\_HUMAN | ATP-dependent (S)-NAD(P)H-hydrate dehydratase | 7C(125.0477) | NaN | X | NaN | NaN | NaN | NaN | 2.554994e+04 | 2.554994e+04 | NaN | NaN | IGVVGGCQEYTGAPYFAAISALK |
| TGM2 | TGM2\_HUMAN | Protein-glutamine gamma-glutamyltransferase 2 | 27 | 20 | TGM2 | TGM2\_HUMAN | Protein-glutamine gamma-glutamyltransferase 2 | 8C(125.0477) | NaN | X | NaN | NaN | NaN | NaN | 2.554104e+04 | NaN | 2.554104e+04 | NaN | DHHTADLCR |
| VCP | TERA\_HUMAN | Transitional endoplasmic reticulum ATPase | 77 | 66 | VCP | TERA\_HUMAN | Transitional endoplasmic reticulum ATPase | 12C(31.9988) | NaN | NaN | NaN | X | NaN | NaN | 2.549532e+04 | 2.549532e+04 | NaN | NaN | EAVCIVLSDDTCSDEK |
| RHOA | RHOA\_HUMAN | Transforming protein RhoA | 159 | 151 | RHOA | RHOA\_HUMAN | Transforming protein RhoA | 9C(125.0477) | NaN | X | NaN | NaN | NaN | NaN | 2.546881e+04 | 2.546881e+04 | NaN | NaN | IGAFGYMECSAK |
| AKR1A1 | AK1A1\_HUMAN | Aldo-keto reductase family 1 member A1 | 134 | 128 | AKR1A1 | AK1A1\_HUMAN | Aldo-keto reductase family 1 member A1 | 7C(125.0477) | NaN | X | NaN | NaN | NaN | NaN | 2.540928e+04 | 2.540928e+04 | NaN | NaN | NADGTICYDSTHYKETWK |
| NFX1 | NFX1\_HUMAN | Transcriptional repressor NF-X1 | 92 | 82 | NFX1 | NFX1\_HUMAN | Transcriptional repressor NF-X1 | 11C(125.0477) | NaN | X | NaN | NaN | NaN | NaN | 2.540528e+04 | 2.540528e+04 | NaN | NaN | SQQTSFQSSPCNK |
| PTPN7 | PTN7\_HUMAN | Tyrosine-protein phosphatase non-receptor type 7 | 324 | 315 | PTPN7 | PTN7\_HUMAN | Tyrosine-protein phosphatase non-receptor type 7 | 10C(125.0477) | NaN | X | NaN | NaN | NaN | NaN | 2.539761e+04 | 2.539761e+04 | NaN | NaN | GEVDILGIVCQLR |
| ARMT1 | ARMT1\_HUMAN | Damage-control phosphatase ARMT1 | 159 | 145 | ARMT1 | ARMT1\_HUMAN | Damage-control phosphatase ARMT1 | 15C(125.0477) | NaN | X | NaN | NaN | NaN | NaN | 2.536239e+04 | 2.536239e+04 | NaN | NaN | EQNFYGSQESIIALCTHLQQLIR |
| RUFY1 | RUFY1\_HUMAN | RUN and FYVE domain-containing protein 1 | 603 | 602 | RUFY1 | RUFY1\_HUMAN | RUN and FYVE domain-containing protein 1 | 2C(125.0477) | NaN | X | NaN | NaN | NaN | NaN | 2.535691e+04 | 2.535691e+04 | NaN | NaN | ICEEQEQALQEMGLHLSQSK |
| CFL1 | COF1\_HUMAN | Cofilin-1 | 39 | 35 | CFL1 | COF1\_HUMAN | Cofilin-1 | 5C(125.0477) | NaN | X | NaN | NaN | NaN | NaN | 2.533494e+04 | NaN | 2.533494e+04 | NaN | AVLFCLSEDK |
| PPIL3 | PPIL3\_HUMAN | Peptidyl-prolyl cis-trans isomerase-like 3 | 18 | 14 | PPIL3 | PPIL3\_HUMAN | Peptidyl-prolyl cis-trans isomerase-like 3 | 5C(125.0477) | NaN | X | NaN | NaN | NaN | NaN | 2.532460e+04 | NaN | 2.532460e+04 | NaN | IEVFCER |
| HPRT1 | HPRT\_HUMAN | Hypoxanthine-guanine phosphoribosyltransferase | 106 | 102 | HPRT1 | HPRT\_HUMAN | Hypoxanthine-guanine phosphoribosyltransferase | 5C(125.0477) | NaN | X | NaN | NaN | NaN | NaN | 2.531838e+04 | 2.531838e+04 | NaN | NaN | LKSYCNDQSTGDIK |
| OXSR1 | OXSR1\_HUMAN | Serine/threonine-protein kinase | 271 | 267 | OXSR1 | OXSR1\_HUMAN | Serine/threonine-protein kinase | 5C(125.0477) | NaN | X | NaN | NaN | NaN | NaN | 2.529220e+04 | 2.529220e+04 | NaN | NaN | MISLCLQKDPEK |
| HARS1 | HARS1\_HUMAN | Histidine--tRNA ligase, cytoplasmic | 235 | 233 | HARS1 | HARS1\_HUMAN | Histidine--tRNA ligase, cytoplasmic | 3C(125.0477) | NaN | X | NaN | NaN | NaN | NaN | 2.527153e+04 | 2.527153e+04 | NaN | NaN | TICSSVDKLDK |
| HDDC2 | HDDC2\_HUMAN | 5'-deoxynucleotidase HDDC2 | 82 | 73 | HDDC2 | HDDC2\_HUMAN | 5'-deoxynucleotidase HDDC2 | 10C(125.0477) | NaN | X | NaN | NaN | NaN | NaN | 2.526281e+04 | 2.526281e+04 | NaN | NaN | LALVHDMAECIVGDIAPADNIPKEEK |
| TSC22D4 | T22D4\_HUMAN | TSC22 domain family protein 4 | 92 | 90 | TSC22D4 | T22D4\_HUMAN | TSC22 domain family protein 4 | 3C(125.0477) | NaN | X | NaN | NaN | NaN | NaN | 2.522463e+04 | 2.522463e+04 | NaN | NaN | WTCVDVYER |
| PLEKHF2 | PKHF2\_HUMAN | Pleckstrin homology domain-containing family F member 2 | 21 | 14 | PLEKHF2 | PKHF2\_HUMAN | Pleckstrin homology domain-containing family F member 2 | 8C(-34.0809) | NaN | NaN | NaN | NaN | X | NaN | 2.521554e+04 | 2.521554e+04 | NaN | NaN | RISIVENCFGAAGQPLTIPGR |
| CFL2 | COF2\_HUMAN | Cofilin-2 | 39 | 34 | CFL2 | COF2\_HUMAN | Cofilin-2 | 6C(125.0477) | NaN | X | NaN | NaN | NaN | NaN | 2.521512e+04 | 2.521512e+04 | NaN | NaN | KAVLFCLSDDKR |
| BLVRB | BLVRB\_HUMAN | Flavin reductase (NADPH) | 109 | 93 | BLVRB | BLVRB\_HUMAN | Flavin reductase (NADPH) | 17C(-34.0809) | NaN | NaN | NaN | NaN | X | NaN | 2.516462e+04 | 2.516462e+04 | NaN | NaN | NIVAAMKAHGVDKVVACTSAFLLWDPTK |
| INPP5D | SHIP1\_HUMAN | Phosphatidylinositol 3,4,5-trisphosphate 5-phosphatase 1 | 217 | 211 | INPP5D | SHIP1\_HUMAN | Phosphatidylinositol 3,4,5-trisphosphate 5-phosphatase 1 | 7C(125.0477) | NaN | X | NaN | NaN | NaN | NaN | 2.505037e+04 | 2.505037e+04 | NaN | NaN | KLTTLLCK |
| AGO2 | AGO2\_HUMAN | Protein argonaute-2 | 352 | 336 | AGO2 | AGO2\_HUMAN | Protein argonaute-2 | 17C(15.9949) | NaN | NaN | X | NaN | NaN | NaN | 2.503892e+04 | 2.503892e+04 | NaN | NaN | HTYLPLEVCNIVAGQRCIKK |
| RDH11 | RDH11\_HUMAN | Retinol dehydrogenase 11 | 33 | 27 | RDH11 | RDH11\_HUMAN | Retinol dehydrogenase 11 | 7C(125.0477) | NaN | X | NaN | NaN | NaN | NaN | 2.503472e+04 | 2.503472e+04 | NaN | NaN | MLSSGVCTSTVQLPGK |
| MON2 | MON2\_HUMAN | Protein MON2 homolog | 212 | 205 | MON2 | MON2\_HUMAN | Protein MON2 homolog | 8C(125.0477) | NaN | X | NaN | NaN | NaN | NaN | 2.496458e+04 | 2.496458e+04 | NaN | NaN | SVSTLKPCAK |
| ANKRD28 | ANR28\_HUMAN | Serine/threonine-protein phosphatase 6 regulatory ankyrin repeat subunit A | 334 | 322 | ANKRD28 | ANR28\_HUMAN | Serine/threonine-protein phosphatase 6 regulatory ankyrin repeat subunit A | 13C(125.0477) | NaN | X | NaN | NaN | NaN | NaN | 2.494470e+04 | 2.494470e+04 | NaN | NaN | SQTIIQSGAVIDCEDK |
| CAPZA2 | CAZA2\_HUMAN | F-actin-capping protein subunit alpha-2 | 157 | 147 | CAPZA2 | CAZA2\_HUMAN | F-actin-capping protein subunit alpha-2 | 11C(125.0477) | NaN | X | NaN | NaN | NaN | NaN | 2.492679e+04 | 2.492679e+04 | NaN | NaN | KIDGQQTIIACIESHQFQAK |
| FLOT2 | FLOT2\_HUMAN | Flotillin-2 | 56 | 56 | FLOT2 | FLOT2\_HUMAN | Flotillin-2 | 1C(125.0477) | NaN | X | NaN | NaN | NaN | NaN | 2.488822e+04 | NaN | 2.488822e+04 | NaN | CEDVETAEGVALTVTGVAQVK |
| PNP | PNPH\_HUMAN | Purine nucleoside phosphorylase | 78 | 77 | PNP | PNPH\_HUMAN | Purine nucleoside phosphorylase | 2C(125.0477) | NaN | X | NaN | NaN | NaN | NaN | 2.488599e+04 | 2.488599e+04 | NaN | NaN | ACVMMQGR |
| STXBP3 | STXB3\_HUMAN | Syntaxin-binding protein 3 | 90 | 87 | STXBP3 | STXB3\_HUMAN | Syntaxin-binding protein 3 | 4C(125.0477) | NaN | X | NaN | NaN | NaN | NaN | 2.485935e+04 | 2.485935e+04 | NaN | NaN | SVDCFLHDFASK |
| PRKD2 | KPCD2\_HUMAN | Serine/threonine-protein kinase D2 | 694 | 693 | PRKD2 | KPCD2\_HUMAN | Serine/threonine-protein kinase D2 | 2C(125.0477) | NaN | X | NaN | NaN | NaN | NaN | 2.484675e+04 | 2.484675e+04 | NaN | NaN | LCDFGFAR |
| TUBB4B | TBB4B\_HUMAN | Tubulin beta-4B chain | 354 | 351 | TUBB4B | TBB4B\_HUMAN | Tubulin beta-4B chain | 4C(125.0477) | NaN | X | NaN | NaN | NaN | NaN | 2.484076e+04 | NaN | 2.484076e+04 | NaN | TAVCDIPPR |
| AHSA1 | AHSA1\_HUMAN | Activator of 90 kDa heat shock protein ATPase homolog 1 | 301 | 295 | AHSA1 | AHSA1\_HUMAN | Activator of 90 kDa heat shock protein ATPase homolog 1 | 7C(125.0477) | NaN | X | NaN | NaN | NaN | NaN | 2.482420e+04 | 2.482420e+04 | NaN | NaN | NGETELCMEGR |
| UBE2V1 | UB2V1\_HUMAN | Ubiquitin-conjugating enzyme E2 variant 1 | 71 | 69 | UBE2V1 | UB2V1\_HUMAN | Ubiquitin-conjugating enzyme E2 variant 1 | 3C(125.0477) | NaN | X | NaN | NaN | NaN | NaN | 2.482019e+04 | 2.482019e+04 | NaN | NaN | IECGPKYPEAPPFVR |
| SPTA1 | SPTA1\_HUMAN | Spectrin alpha chain, erythrocytic 1 | 1879 | 1875 | SPTA1 | SPTA1\_HUMAN | Spectrin alpha chain, erythrocytic 1 | 5C(125.0477) | NaN | X | NaN | NaN | NaN | NaN | 2.481949e+04 | 2.481949e+04 | NaN | NaN | VQNVCAQGEDILNKVLQEESQNKEISSK |
| SPTBN1 | SPTB2\_HUMAN | Spectrin beta chain, non-erythrocytic 1 | 1900 | 1895 | SPTBN1 | SPTB2\_HUMAN | Spectrin beta chain, non-erythrocytic 1 | 6C(125.0477) | NaN | X | NaN | NaN | NaN | NaN | 2.480449e+04 | NaN | 2.480449e+04 | NaN | SLLDACESR |
| VPS50 | VPS50\_HUMAN | Syndetin | 851 | 839 | VPS50 | VPS50\_HUMAN | Syndetin | 13C(125.0477) | NaN | X | NaN | NaN | NaN | NaN | 2.476410e+04 | 2.476410e+04 | NaN | NaN | IPLPVSNILWEHCIR |
| VTA1 | VTA1\_HUMAN | Vacuolar protein sorting-associated protein VTA1 homolog | 38 | 23 | VTA1 | VTA1\_HUMAN | Vacuolar protein sorting-associated protein VTA1 homolog | 16C(125.0477) | NaN | X | NaN | NaN | NaN | NaN | 2.476059e+04 | 2.476059e+04 | NaN | NaN | TAQEHDKRDPVVAYYCR |
| XPO4 | XPO4\_HUMAN | Exportin-4 | 54 | 49 | XPO4 | XPO4\_HUMAN | Exportin-4 | 6C(125.0477) | NaN | X | NaN | NaN | NaN | NaN | 2.472824e+04 | 2.472824e+04 | NaN | NaN | SPFAVCK |
| PSMB2 | PSB2\_HUMAN | Proteasome subunit beta type-2 | 91 | 87 | PSMB2 | PSB2\_HUMAN | Proteasome subunit beta type-2 | 5C(125.0477) | NaN | X | NaN | NaN | NaN | NaN | 2.471664e+04 | NaN | 2.471664e+04 | NaN | NLADCLR |
| ATP6V0A1 | VPP1\_HUMAN | V-type proton ATPase 116 kDa subunit a 1 | 58 | 57 | ATP6V0A1 | VPP1\_HUMAN | V-type proton ATPase 116 kDa subunit a 1 | 2C(125.0477) | NaN | X | NaN | NaN | NaN | NaN | 2.471019e+04 | 2.471019e+04 | NaN | NaN | RCEEMDR |
| EIF2B5 | EI2BE\_HUMAN | Translation initiation factor eIF-2B subunit epsilon | 335 | 317 | EIF2B5 | EI2BE\_HUMAN | Translation initiation factor eIF-2B subunit epsilon | 19C(125.0477) | NaN | X | NaN | NaN | NaN | NaN | 2.466619e+04 | 2.466619e+04 | NaN | NaN | WVYPLTPEANFTDSTTQSCTHSR |
| ATP2B4 | AT2B4\_HUMAN | Plasma membrane calcium-transporting ATPase 4 | 632 | 625 | ATP2B4 | AT2B4\_HUMAN | Plasma membrane calcium-transporting ATPase 4 | 8C(15.9949) | NaN | NaN | X | NaN | NaN | NaN | 2.461788e+04 | NaN | NaN | 24617.8800 | TVIEPMACDGLR |
| UBA1 | UBA1\_HUMAN | Ubiquitin-like modifier-activating enzyme 1 | 494 | 486 | UBA1 | UBA1\_HUMAN | Ubiquitin-like modifier-activating enzyme 1 | 9C(125.0477) | NaN | X | NaN | NaN | NaN | NaN | 2.461062e+04 | 2.461062e+04 | NaN | NaN | NFAMIGLGCGEGGEIIVTDMDTIEK |
| KPNA4 | IMA3\_HUMAN | Importin subunit alpha-3 | 325 | 307 | KPNA4 | IMA3\_HUMAN | Importin subunit alpha-3 | 19C(125.0477) | NaN | X | NaN | NaN | NaN | NaN | 2.459226e+04 | 2.459226e+04 | NaN | NaN | AVGNIVTGTDEQTQVVLNCDALSHFPALLTHPK |
| HBB | HBB\_HUMAN | Hemoglobin subunit beta | 113 | 106 | HBB | HBB\_HUMAN | Hemoglobin subunit beta | 8C(125.0477) | NaN | X | NaN | NaN | NaN | NaN | 2.457603e+04 | 2.457603e+04 | NaN | NaN | LLGNVLVCVLAHHFGKEFTPPVQAAYQKVVAGVANALAHK |
| LRRC57 | LRC57\_HUMAN | Leucine-rich repeat-containing protein 57 | 128 | 112 | LRRC57 | LRC57\_HUMAN | Leucine-rich repeat-containing protein 57 | 17C(125.0477) | NaN | X | NaN | NaN | NaN | NaN | 2.455434e+04 | 2.455434e+04 | NaN | NaN | TLSLSGNQLGALPPQLCSLR |
| EPHX2 | HYES\_HUMAN | Bifunctional epoxide hydrolase 2 | 423 | 422 | EPHX2 | HYES\_HUMAN | Bifunctional epoxide hydrolase 2 | 2C(125.0477) | NaN | X | NaN | NaN | NaN | NaN | 2.452625e+04 | 2.452625e+04 | NaN | NaN | VCEAGGLFVNSPEEPSLSR |
| RIC8A | RIC8A\_HUMAN | Synembryn-A | 79 | 78 | RIC8A | RIC8A\_HUMAN | Synembryn-A | 2C(125.0477) | NaN | X | NaN | NaN | NaN | NaN | 2.452549e+04 | NaN | 2.452549e+04 | NaN | NCLDPFTSR |
| PEBP1 | PEBP1\_HUMAN | Phosphatidylethanolamine-binding protein 1 | 133 | 120 | PEBP1 | PEBP1\_HUMAN | Phosphatidylethanolamine-binding protein 1 | 14C(125.0477) | NaN | X | NaN | NaN | NaN | NaN | 2.449727e+04 | 2.449727e+04 | NaN | NaN | YVWLVYEQDRPLKCDEPILSNR |
| UBE2K | UBE2K\_HUMAN | Ubiquitin-conjugating enzyme E2 K | 170 | 165 | UBE2K | UBE2K\_HUMAN | Ubiquitin-conjugating enzyme E2 K | 6C(125.0477) | NaN | X | NaN | NaN | NaN | NaN | 2.446872e+04 | 2.446872e+04 | NaN | NaN | KIENLCAMGFDR |
| CKAP5 | CKAP5\_HUMAN | Cytoskeleton-associated protein 5 | 427 | 426 | CKAP5 | CKAP5\_HUMAN | Cytoskeleton-associated protein 5 | 2C(125.0477) | NaN | X | NaN | NaN | NaN | NaN | 2.446522e+04 | 2.446522e+04 | NaN | NaN | HCTASTLPK |
| GLMN | GLMN\_HUMAN | Glomulin | 36 | 36 | GLMN | GLMN\_HUMAN | Glomulin | 1C(125.0477) | NaN | X | NaN | NaN | NaN | NaN | 2.442894e+04 | 2.442894e+04 | NaN | NaN | CIEEGHTDQLLEIIQNEK |
| HARS1 | HARS1\_HUMAN | Histidine--tRNA ligase, cytoplasmic | 379 | 376 | HARS1 | HARS1\_HUMAN | Histidine--tRNA ligase, cytoplasmic | 4C(125.0477) | NaN | X | NaN | NaN | NaN | NaN | 2.441109e+04 | 2.441109e+04 | NaN | NaN | KVPCVGLSIGVER |
| TLN1 | TLN1\_HUMAN | Talin-1 | 1927 | 1920 | TLN1 | TLN1\_HUMAN | Talin-1 | 8C(125.0477) | NaN | X | NaN | NaN | NaN | NaN | 2.438530e+04 | 2.438530e+04 | NaN | NaN | VQELGHGCAALVTK |
| RANGAP1 | RAGP1\_HUMAN | Ran GTPase-activating protein 1 | 152 | 149 | RANGAP1 | RAGP1\_HUMAN | Ran GTPase-activating protein 1 | 4C(125.0477) | NaN | X | NaN | NaN | NaN | NaN | 2.438496e+04 | 2.438496e+04 | NaN | NaN | LNNCGMGIGGGK |
| PSME2 | PSME2\_HUMAN | Proteasome activator complex subunit 2 | 118 | 118 | PSME2 | PSME2\_HUMAN | Proteasome activator complex subunit 2 | 1C(125.0477) | NaN | X | NaN | NaN | NaN | NaN | 2.438234e+04 | 2.438234e+04 | NaN | NaN | CILVITWIQHLIPK |
| APOB | APOB\_HUMAN | Apolipoprotein B-100 | 1505 | 1499 | APOB | APOB\_HUMAN | Apolipoprotein B-100 | 7C(57.0215) | X | NaN | NaN | NaN | NaN | NaN | 2.437819e+04 | NaN | 2.437819e+04 | NaN | GTYGLSCQR |
| EPS15 | EPS15\_HUMAN | Epidermal growth factor receptor substrate 15 | 824 | 817 | EPS15 | EPS15\_HUMAN | Epidermal growth factor receptor substrate 15 | 8C(125.0477) | NaN | X | NaN | NaN | NaN | NaN | 2.437805e+04 | 2.437805e+04 | NaN | NaN | EKDPEIFCDPFTSATTTTNK |
| ANP32E | AN32E\_HUMAN | Acidic leucine-rich nuclear phosphoprotein 32 family member E | 87 | 87 | ANP32E | AN32E\_HUMAN | Acidic leucine-rich nuclear phosphoprotein 32 family member E | 1C(125.0477) | NaN | X | NaN | NaN | NaN | NaN | 2.436248e+04 | 2.436248e+04 | NaN | NaN | CPNLTYLNLSGNK |
| ITPA | ITPA\_HUMAN | Inosine triphosphate pyrophosphatase | 116 | 111 | ITPA | ITPA\_HUMAN | Inosine triphosphate pyrophosphatase | 6C(125.0477) | NaN | X | NaN | NaN | NaN | NaN | 2.434843e+04 | 2.434843e+04 | NaN | NaN | SAYALCTFALSTGDPSQPVR |
| ARFGEF1 | BIG1\_HUMAN | Brefeldin A-inhibited guanine nucleotide-exchange protein 1 | 1685 | 1682 | ARFGEF1 | BIG1\_HUMAN | Brefeldin A-inhibited guanine nucleotide-exchange protein 1 | 4C(125.0477) | NaN | X | NaN | NaN | NaN | NaN | 2.433076e+04 | 2.433076e+04 | NaN | NaN | LLDCLLESHR |
| HAGH | GLO2\_HUMAN | Hydroxyacylglutathione hydrolase, mitochondrial | 201 | 192 | HAGH | GLO2\_HUMAN | Hydroxyacylglutathione hydrolase, mitochondrial | 10C(125.0477) | NaN | X | NaN | NaN | NaN | NaN | 2.430839e+04 | 2.430839e+04 | NaN | NaN | FYEGTADEMCK |
| ANXA7 | ANXA7\_HUMAN | Annexin A7 | 298 | 298 | ANXA7 | ANXA7\_HUMAN | Annexin A7 | 1C(125.0477) | NaN | X | NaN | NaN | NaN | NaN | 2.424438e+04 | 2.424438e+04 | NaN | NaN | CYQSEFGRDLEKDIR |
| EIF5A | IF5A1\_HUMAN | Eukaryotic translation initiation factor 5A-1 | 73 | 56 | EIF5A | IF5A1\_HUMAN | Eukaryotic translation initiation factor 5A-1 | 18C(125.0477) | NaN | X | NaN | NaN | NaN | NaN | 2.424285e+04 | 2.424285e+04 | NaN | NaN | VHLVGIDIFTGKKYEDICPSTHNMDVPNIK |
| AMBRA1 | AMRA1\_HUMAN | Activating molecule in BECN1-regulated autophagy protein 1 | 87 | 87 | AMBRA1 | AMRA1\_HUMAN | Activating molecule in BECN1-regulated autophagy protein 1 | 1C(125.0477) | NaN | X | NaN | NaN | NaN | NaN | 2.422769e+04 | 2.422769e+04 | NaN | NaN | CVHSLIGHR |
| STAT6 | STAT6\_HUMAN | Signal transducer and activator of transcription 6 | 272 | 267 | STAT6 | STAT6\_HUMAN | Signal transducer and activator of transcription 6 | 6C(125.0477) | NaN | X | NaN | NaN | NaN | NaN | 2.413178e+04 | 2.413178e+04 | NaN | NaN | TLVTSCFLVEK |
| GUCD1 | GUCD1\_HUMAN | Protein GUCD1 | 85 | 84 | GUCD1 | GUCD1\_HUMAN | Protein GUCD1 | 2C(125.0477) | NaN | X | NaN | NaN | NaN | NaN | 2.412284e+04 | NaN | 2.412284e+04 | NaN | FCTQTLGVDK |
| SURF4 | SURF4\_HUMAN | Surfeit locus protein 4 | 32 | 31 | SURF4 | SURF4\_HUMAN | Surfeit locus protein 4 | 2C(125.0477) | NaN | X | NaN | NaN | NaN | NaN | 2.411080e+04 | 2.411080e+04 | NaN | NaN | LCLISTFLEDGIR |
| APMAP | APMAP\_HUMAN | Adipocyte plasma membrane-associated protein | 140 | 135 | APMAP | APMAP\_HUMAN | Adipocyte plasma membrane-associated protein | 6C(125.0477) | NaN | X | NaN | NaN | NaN | NaN | 2.410992e+04 | NaN | 2.410992e+04 | NaN | FGSGPCK |
| RRAGA | RRAGA\_HUMAN | Ras-related GTP-binding protein A | 219 | 209 | RRAGA | RRAGA\_HUMAN | Ras-related GTP-binding protein A | 11C(125.0477) | NaN | X | NaN | NaN | NaN | NaN | 2.408879e+04 | 2.408879e+04 | NaN | NaN | ATFLVISHYQCK |
| EIF4E | IF4E\_HUMAN | Eukaryotic translation initiation factor 4E | 170 | 163 | EIF4E | IF4E\_HUMAN | Eukaryotic translation initiation factor 4E | 8C(125.0477) | NaN | X | NaN | NaN | NaN | NaN | 2.407203e+04 | 2.407203e+04 | NaN | NaN | IAIWTTECENREAVTHIGR |
| PTPN7 | PTN7\_HUMAN | Tyrosine-protein phosphatase non-receptor type 7 | 300 | 298 | PTPN7 | PTN7\_HUMAN | Tyrosine-protein phosphatase non-receptor type 7 | 3C(125.0477) | NaN | X | NaN | NaN | NaN | NaN | 2.403393e+04 | NaN | 2.403393e+04 | NaN | TGCFIATR |
| INF2 | INF2\_HUMAN | Inverted formin-2 | 745 | 738 | INF2 | INF2\_HUMAN | Inverted formin-2 | 8C(125.0477) | NaN | X | NaN | NaN | NaN | NaN | 2.395333e+04 | 2.395333e+04 | NaN | NaN | AQLVLAACESLLTSR |
| SNX9 | SNX9\_HUMAN | Sorting nexin-9 | 492 | 484 | SNX9 | SNX9\_HUMAN | Sorting nexin-9 | 9C(125.0477) | NaN | X | NaN | NaN | NaN | NaN | 2.390385e+04 | 2.390385e+04 | NaN | NaN | KDLHFLMECNHEYK |
| HSPA1A | HS71A\_HUMAN | Heat shock 70 kDa protein 1A | 603 | 598 | HSPA1A | HS71A\_HUMAN | Heat shock 70 kDa protein 1A | 6C(125.0477) | NaN | X | NaN | NaN | NaN | NaN | 2.390249e+04 | 2.390249e+04 | NaN | NaN | ELEQVCNPIISGLYQGAGGPGPGGFGAQGPK |
| FTO | FTO\_HUMAN | Alpha-ketoglutarate-dependent dioxygenase FTO | 171 | 163 | FTO | FTO\_HUMAN | Alpha-ketoglutarate-dependent dioxygenase FTO | 9C(125.0477) | NaN | X | NaN | NaN | NaN | NaN | 2.387207e+04 | 2.387207e+04 | NaN | NaN | ANEDAVPLCMSADFPR |
| SLC43A2 | LAT4\_HUMAN | Large neutral amino acids transporter small subunit 4 | 392 | 389 | SLC43A2 | LAT4\_HUMAN | Large neutral amino acids transporter small subunit 4 | 4C(125.0477) | NaN | X | NaN | NaN | NaN | NaN | 2.383881e+04 | 2.383881e+04 | NaN | NaN | LKECEDASEEPEEKDANQGEK |
| A2M | A2MG\_HUMAN | Alpha-2-macroglobulin | 431 | 429 | A2M | A2MG\_HUMAN | Alpha-2-macroglobulin | 3C(57.0215) | X | NaN | NaN | NaN | NaN | NaN | 2.379774e+04 | 2.379774e+04 | NaN | NaN | SPCYGYQWVSEEHEEAHHTAYLVFSPSK |
| ADRM1 | ADRM1\_HUMAN | Proteasomal ubiquitin receptor ADRM1 | 80 | 65 | ADRM1 | ADRM1\_HUMAN | Proteasomal ubiquitin receptor ADRM1 | 16C(125.0477) | NaN | X | NaN | NaN | NaN | NaN | 2.379223e+04 | 2.379223e+04 | NaN | NaN | TSGNVEDDLIIFPDDCEFKR |
| CUL3 | CUL3\_HUMAN | Cullin-3 | 251 | 248 | CUL3 | CUL3\_HUMAN | Cullin-3 | 4C(125.0477) | NaN | X | NaN | NaN | NaN | NaN | 2.378102e+04 | 2.378102e+04 | NaN | NaN | VMHCLDKSTEEPIVK |
| LZIC | LZIC\_HUMAN | Protein LZIC | 33 | 23 | LZIC | LZIC\_HUMAN | Protein LZIC | 11C(125.0477) | NaN | X | NaN | NaN | NaN | NaN | 2.372820e+04 | 2.372820e+04 | NaN | NaN | LMQQLQDLEECREELDTDEYEETKK |
| REXO2 | ORN\_HUMAN | Oligoribonuclease, mitochondrial | 176 | 166 | REXO2 | ORN\_HUMAN | Oligoribonuclease, mitochondrial | 11C(125.0477) | NaN | X | NaN | NaN | NaN | NaN | 2.371515e+04 | 2.371515e+04 | NaN | NaN | IIDVSTVKELCRR |
| ZFPL1 | ZFPL1\_HUMAN | Zinc finger protein-like 1 | 16 | 11 | ZFPL1 | ZFPL1\_HUMAN | Zinc finger protein-like 1 | 6C(125.0477) | NaN | X | NaN | NaN | NaN | NaN | 2.370759e+04 | 2.370759e+04 | NaN | NaN | VTNLFCFEHR |
| WDR81 | WDR81\_HUMAN | WD repeat-containing protein 81 | 76 | 70 | WDR81 | WDR81\_HUMAN | WD repeat-containing protein 81 | 7C(125.0477) | NaN | X | NaN | NaN | NaN | NaN | 2.365183e+04 | 2.365183e+04 | NaN | NaN | RLPLGPCPR |
| ALB | ALBU\_HUMAN | Albumin | 125 | 118 | ALB | ALBU\_HUMAN | Albumin | 8C(57.0215) | X | NaN | NaN | NaN | NaN | NaN | 2.362656e+04 | 2.362656e+04 | NaN | NaN | QEPERNECFLQHKDDNPNLPR |
| ACO1 | ACOHC\_HUMAN | Cytoplasmic aconitate hydratase | 118 | 114 | ACO1 | ACOHC\_HUMAN | Cytoplasmic aconitate hydratase | 5C(125.0477) | NaN | X | NaN | NaN | NaN | NaN | 2.351312e+04 | 2.351312e+04 | NaN | NaN | INPVCPADLVIDHSIQVDFNR |
| SUGT1 | SGT1\_HUMAN | Protein SGT1 homolog | 154 | 154 | SUGT1 | SGT1\_HUMAN | Protein SGT1 homolog | 1C(125.0477) | NaN | X | NaN | NaN | NaN | NaN | 2.348387e+04 | 2.348387e+04 | NaN | NaN | CQEAQNGSESEVWTHQSK |
| AMBP | AMBP\_HUMAN | Protein AMBP | 287 | 283 | AMBP | AMBP\_HUMAN | Protein AMBP | 5C(57.0215) | X | NaN | NaN | NaN | NaN | NaN | 2.345292e+04 | NaN | 2.345292e+04 | NaN | TVAACNLPIVR |
| DDX58 | DDX58\_HUMAN | Antiviral innate immune response receptor RIG-I | 738 | 738 | DDX58 | DDX58\_HUMAN | Antiviral innate immune response receptor RIG-I | 1C(125.0477) | NaN | X | NaN | NaN | NaN | NaN | 2.341752e+04 | 2.341752e+04 | NaN | NaN | CFLLTSNAGVIEK |
| VPS13C | VP13C\_HUMAN | Vacuolar protein sorting-associated protein 13C | 3723 | 3722 | VPS13C | VP13C\_HUMAN | Vacuolar protein sorting-associated protein 13C | 2C(125.0477) | NaN | X | NaN | NaN | NaN | NaN | 2.340326e+04 | NaN | 2.340326e+04 | NaN | ACNAIEDAQSTR |
| CLTC | CLH1\_HUMAN | Clathrin heavy chain 1 | 491 | 482 | CLTC | CLH1\_HUMAN | Clathrin heavy chain 1 | 10C(125.0477) | NaN | X | NaN | NaN | NaN | NaN | 2.339653e+04 | 2.339653e+04 | NaN | NaN | ANVPNKVIQCFAETGQVQK |
| HTT | HD\_HUMAN | Huntingtin | 1152 | 1148 | HTT | HD\_HUMAN | Huntingtin | 5C(125.0477) | NaN | X | NaN | NaN | NaN | NaN | 2.337302e+04 | 2.337302e+04 | NaN | NaN | VINICAHVLDDVAPGPAIK |
| GNB4 | GBB4\_HUMAN | Guanine nucleotide-binding protein subunit beta-4 | 204 | 198 | GNB4 | GBB4\_HUMAN | Guanine nucleotide-binding protein subunit beta-4 | 7C(125.0477) | NaN | X | NaN | NaN | NaN | NaN | 2.332622e+04 | NaN | 2.332622e+04 | NaN | TFVSGACDASSK |
| HNRNPH1 | HNRH1\_HUMAN | Heterogeneous nuclear ribonucleoprotein H | 290 | 276 | HNRNPH1 | HNRH1\_HUMAN | Heterogeneous nuclear ribonucleoprotein H | 15C(125.0477) | NaN | X | NaN | NaN | NaN | NaN | 2.330082e+04 | 2.330082e+04 | NaN | NaN | YGDGGSTFQSTTGHCVHMR |
| EIF2B5 | EI2BE\_HUMAN | Translation initiation factor eIF-2B subunit epsilon | 414 | 402 | EIF2B5 | EI2BE\_HUMAN | Translation initiation factor eIF-2B subunit epsilon | 13C(125.0477) | NaN | X | NaN | NaN | NaN | NaN | 2.324840e+04 | 2.324840e+04 | NaN | NaN | VAAGAQIHQSLLCDNAEVK |
| PSME4 | PSME4\_HUMAN | Proteasome activator complex subunit 4 | 554 | 554 | PSME4 | PSME4\_HUMAN | Proteasome activator complex subunit 4 | 1C(125.0477) | NaN | X | NaN | NaN | NaN | NaN | 2.323954e+04 | 2.323954e+04 | NaN | NaN | CFGLIESSTLEQTR |
| LDHA | LDHA\_HUMAN | L-lactate dehydrogenase A chain | 293 | 285 | LDHA | LDHA\_HUMAN | L-lactate dehydrogenase A chain | 9C(125.0477) | NaN | X | NaN | NaN | NaN | NaN | 2.322457e+04 | 2.322457e+04 | NaN | NaN | DDVFLSVPCILGQNGISDLVK |
| EIF2S3 | IF2G\_HUMAN | Eukaryotic translation initiation factor 2 subunit 3 | 269 | 261 | EIF2S3 | IF2G\_HUMAN | Eukaryotic translation initiation factor 2 subunit 3 | 9C(125.0477) | NaN | X | NaN | NaN | NaN | NaN | 2.316288e+04 | 2.316288e+04 | NaN | NaN | SFDVNKPGCEVDDLK |
| TCP1 | TCPA\_HUMAN | T-complex protein 1 subunit alpha | 125 | 112 | TCP1 | TCPA\_HUMAN | T-complex protein 1 subunit alpha | 14C(15.9949) | NaN | NaN | X | NaN | NaN | NaN | 2.315986e+04 | 2.315986e+04 | NaN | NaN | IHPTSVISGYRLACK |
| PSMB6 | PSB6\_HUMAN | Proteasome subunit beta type-6 | 186 | 184 | PSMB6 | PSB6\_HUMAN | Proteasome subunit beta type-6 | 3C(125.0477) | NaN | X | NaN | NaN | NaN | NaN | 2.315158e+04 | 2.315158e+04 | NaN | NaN | EECLQFTANALALAMER |
| SGTA | SGTA\_HUMAN | Small glutamine-rich tetratricopeptide repeat-containing protein alpha | 153 | 151 | SGTA | SGTA\_HUMAN | Small glutamine-rich tetratricopeptide repeat-containing protein alpha | 3C(125.0477) | NaN | X | NaN | NaN | NaN | NaN | 2.306582e+04 | NaN | 2.306582e+04 | NaN | AICIDPAYSK |
| DSTN | DEST\_HUMAN | Destrin | 163 | 152 | DSTN | DEST\_HUMAN | Destrin | 12C(125.0477) | NaN | X | NaN | NaN | NaN | NaN | 2.305767e+04 | 2.305767e+04 | NaN | NaN | LGGSLIVAFEGCPV |
| DDAH2 | DDAH2\_HUMAN | N(G),N(G)-dimethylarginine dimethylaminohydrolase 2 | 262 | 252 | DDAH2 | DDAH2\_HUMAN | N(G),N(G)-dimethylarginine dimethylaminohydrolase 2 | 11C(15.9949) | NaN | NaN | X | NaN | NaN | NaN | 2.304630e+04 | 2.304630e+04 | NaN | NaN | LSDVTLVPVSCSELEKAGAGLSSLCLVLSTRPHS |
| CFH | CFAH\_HUMAN | Complement factor H | 1018 | 1010 | CFH | CFAH\_HUMAN | Complement factor H | 9C(57.0215) | X | NaN | NaN | NaN | NaN | NaN | 2.300062e+04 | NaN | 2.300062e+04 | NaN | AGEQVTYTCATYYK |
| XPO4 | XPO4\_HUMAN | Exportin-4 | 186 | 173 | XPO4 | XPO4\_HUMAN | Exportin-4 | 14C(125.0477) | NaN | X | NaN | NaN | NaN | NaN | 2.298854e+04 | 2.298854e+04 | NaN | NaN | TSNIGLSMEFHGNCK |
| FHIP2A | FHI2A\_HUMAN | FHF complex subunit HOOK interacting protein 2A | 624 | 624 | FHIP2A | FHI2A\_HUMAN | FHF complex subunit HOOK interacting protein 2A | 1C(125.0477) | NaN | X | NaN | NaN | NaN | NaN | 2.298046e+04 | 2.298046e+04 | NaN | NaN | CNLEAAFFEGHFLK |
| TRAF2 | TRAF2\_HUMAN | TNF receptor-associated factor 2 | 49 | 43 | TRAF2 | TRAF2\_HUMAN | TNF receptor-associated factor 2 | 7C(125.0477) | NaN | X | NaN | NaN | NaN | NaN | 2.295253e+04 | 2.295253e+04 | NaN | NaN | RPFQAQCGHR |
| TES | TES\_HUMAN | Testin | 294 | 292 | TES | TES\_HUMAN | Testin | 3C(125.0477) | NaN | X | NaN | NaN | NaN | NaN | 2.288834e+04 | 2.288834e+04 | NaN | NaN | HYCDSEKPR |
| PFKM | PFKAM\_HUMAN | ATP-dependent 6-phosphofructokinase, muscle type | 334 | 316 | PFKM | PFKAM\_HUMAN | ATP-dependent 6-phosphofructokinase, muscle type | 19C(125.0477) | NaN | X | NaN | NaN | NaN | NaN | 2.284650e+04 | 2.284650e+04 | NaN | NaN | MGVEAVMALLEGTPDTPACVVSLSGNQAVR |
| NT5DC1 | NT5D1\_HUMAN | 5'-nucleotidase domain-containing protein 1 | 197 | 193 | NT5DC1 | NT5D1\_HUMAN | 5'-nucleotidase domain-containing protein 1 | 5C(125.0477) | NaN | X | NaN | NaN | NaN | NaN | 2.283915e+04 | 2.283915e+04 | NaN | NaN | YLHSCPESVKK |
| XPO5 | XPO5\_HUMAN | Exportin-5 | 1052 | 1047 | XPO5 | XPO5\_HUMAN | Exportin-5 | 6C(125.0477) | NaN | X | NaN | NaN | NaN | NaN | 2.282165e+04 | 2.282165e+04 | NaN | NaN | TTSQLCWPLLK |
| CLIC1 | CLIC1\_HUMAN | Chloride intracellular channel protein 1 | 223 | 217 | CLIC1 | CLIC1\_HUMAN | Chloride intracellular channel protein 1 | 7C(125.0477) | NaN | X | NaN | NaN | NaN | NaN | 2.281286e+04 | 2.281286e+04 | NaN | NaN | EEFASTCPDDEEIELAYEQVAK |
| APEH | ACPH\_HUMAN | Acylamino-acid-releasing enzyme | 292 | 280 | APEH | ACPH\_HUMAN | Acylamino-acid-releasing enzyme | 13C(125.0477) | NaN | X | NaN | NaN | NaN | NaN | 2.274631e+04 | 2.274631e+04 | NaN | NaN | SALYYVDLIGGKCELLSDDSLAVSSPR |
| CAPZB | CAPZB\_HUMAN | F-actin-capping protein subunit beta | 62 | 52 | CAPZB | CAPZB\_HUMAN | F-actin-capping protein subunit beta | 11C(125.0477) | NaN | X | NaN | NaN | NaN | NaN | 2.271826e+04 | 2.271826e+04 | NaN | NaN | DKVVGKDYLLCDYNR |
| TTC37 | TTC37\_HUMAN | Tetratricopeptide repeat protein 37 | 497 | 495 | TTC37 | TTC37\_HUMAN | Tetratricopeptide repeat protein 37 | 3C(125.0477) | NaN | X | NaN | NaN | NaN | NaN | 2.269960e+04 | 2.269960e+04 | NaN | NaN | VFCYLGHYYR |
| APEH | ACPH\_HUMAN | Acylamino-acid-releasing enzyme | 312 | 307 | APEH | ACPH\_HUMAN | Acylamino-acid-releasing enzyme | 6C(125.0477) | NaN | X | NaN | NaN | NaN | NaN | 2.267441e+04 | NaN | 2.267441e+04 | NaN | LSPDQCR |
| LANCL1 | LANC1\_HUMAN | Glutathione S-transferase LANCL1 | 363, 349, 355 | 348 | LANCL1 | LANC1\_HUMAN | Glutathione S-transferase LANCL1 | 16C(125.0477) 2C(15.9949) 8C(31.9988) | NaN | X | X | X | NaN | NaN | 2.262362e+04 | 2.262362e+04 | NaN | NaN | ACKFAEWCLEYGEHGCR |
| RNF123 | RN123\_HUMAN | E3 ubiquitin-protein ligase RNF123 | 931 | 920 | RNF123 | RN123\_HUMAN | E3 ubiquitin-protein ligase RNF123 | 12C(125.0477) | NaN | X | NaN | NaN | NaN | NaN | 2.257604e+04 | 2.257604e+04 | NaN | NaN | DSLMQALASYVCYPHSLR |
| ABCG2 | ABCG2\_HUMAN | Broad substrate specificity ATP-binding cassette transporter ABCG2 | 374 | 366 | ABCG2 | ABCG2\_HUMAN | Broad substrate specificity ATP-binding cassette transporter ABCG2 | 9C(125.0477) | NaN | X | NaN | NaN | NaN | NaN | 2.254937e+04 | 2.254937e+04 | NaN | NaN | EISYTTSFCHQLR |
| SPTB | SPTB1\_HUMAN | Spectrin beta chain, erythrocytic | 735 | 720 | SPTB | SPTB1\_HUMAN | Spectrin beta chain, erythrocytic | 16C(125.0477) | NaN | X | NaN | NaN | NaN | NaN | 2.250679e+04 | 2.250679e+04 | NaN | NaN | EVSAQWDQLKDLAAFCK |
| FTO | FTO\_HUMAN | Alpha-ketoglutarate-dependent dioxygenase FTO | 326 | 323 | FTO | FTO\_HUMAN | Alpha-ketoglutarate-dependent dioxygenase FTO | 4C(125.0477) | NaN | X | NaN | NaN | NaN | NaN | 2.246282e+04 | 2.246282e+04 | NaN | NaN | VAECSTGTLDYILQR |
| ANAPC4 | APC4\_HUMAN | Anaphase-promoting complex subunit 4 | 732 | 730 | ANAPC4 | APC4\_HUMAN | Anaphase-promoting complex subunit 4 | 3C(125.0477) | NaN | X | NaN | NaN | NaN | NaN | 2.242752e+04 | 2.242752e+04 | NaN | NaN | VSCVLSSNLR |
| TNPO3 | TNPO3\_HUMAN | Transportin-3 | 511 | 509 | TNPO3 | TNPO3\_HUMAN | Transportin-3 | 3C(125.0477) | NaN | X | NaN | NaN | NaN | NaN | 2.242319e+04 | 2.242319e+04 | NaN | NaN | GLCEKPLASAAAK |
| TES | TES\_HUMAN | Testin | 46 | 46 | TES | TES\_HUMAN | Testin | 1C(125.0477) | NaN | X | NaN | NaN | NaN | NaN | 2.241192e+04 | 2.241192e+04 | NaN | NaN | CGQEEHDVLLSNEEDRK |
| USP14 | UBP14\_HUMAN | Ubiquitin carboxyl-terminal hydrolase 14 | 359 | 346 | USP14 | UBP14\_HUMAN | Ubiquitin carboxyl-terminal hydrolase 14 | 14C(125.0477) | NaN | X | NaN | NaN | NaN | NaN | 2.238544e+04 | 2.238544e+04 | NaN | NaN | DVKFPLMLDMYELCTPELQEK |
| NAA50 | NAA50\_HUMAN | N-alpha-acetyltransferase 50 | 100 | 92 | NAA50 | NAA50\_HUMAN | N-alpha-acetyltransferase 50 | 9C(125.0477) | NaN | X | NaN | NaN | NaN | NaN | 2.235285e+04 | 2.235285e+04 | NaN | NaN | MLNHVLNICEK |
| SLC12A6 | S12A6\_HUMAN | Solute carrier family 12 member 6 | 390 | 386 | SLC12A6 | S12A6\_HUMAN | Solute carrier family 12 member 6 | 5C(125.0477) | NaN | X | NaN | NaN | NaN | NaN | 2.234106e+04 | NaN | 2.234106e+04 | NaN | HIDVCSK |
| TKFC | TKFC\_HUMAN | Triokinase/FMN cyclase | 155 | 152 | TKFC | TKFC\_HUMAN | Triokinase/FMN cyclase | 4C(125.0477) | NaN | X | NaN | NaN | NaN | NaN | 2.233722e+04 | 2.233722e+04 | NaN | NaN | RGLCGTVLIHK |
| LCP1 | PLSL\_HUMAN | Plastin-2 | 42 | 40 | LCP1 | PLSL\_HUMAN | Plastin-2 | 3C(125.0477) | NaN | X | NaN | NaN | NaN | NaN | 2.222588e+04 | NaN | 2.222588e+04 | NaN | AACLPLPGYR |
| PSMC3 | PRS6A\_HUMAN | 26S proteasome regulatory subunit 6A | 400 | 398 | PSMC3 | PRS6A\_HUMAN | 26S proteasome regulatory subunit 6A | 3C(125.0477) | NaN | X | NaN | NaN | NaN | NaN | 2.219978e+04 | 2.219978e+04 | NaN | NaN | AVCVEAGMIALRR |
| CAB39 | CAB39\_HUMAN | Calcium-binding protein 39 | 150 | 149 | CAB39 | CAB39\_HUMAN | Calcium-binding protein 39 | 2C(125.0477) | NaN | X | NaN | NaN | NaN | NaN | 2.218381e+04 | 2.218381e+04 | NaN | NaN | ECIRHEPLAK |
| CNP | CN37\_HUMAN | 2',3'-cyclic-nucleotide 3'-phosphodiesterase | 158 | 156 | CNP | CN37\_HUMAN | 2',3'-cyclic-nucleotide 3'-phosphodiesterase | 3C(125.0477) | NaN | X | NaN | NaN | NaN | NaN | 2.216848e+04 | 2.216848e+04 | NaN | NaN | LDCAQLKEK |
| RPE | RPE\_HUMAN | Ribulose-phosphate 3-epimerase | 186 | 186 | RPE | RPE\_HUMAN | Ribulose-phosphate 3-epimerase | 1C(125.0477) | NaN | X | NaN | NaN | NaN | NaN | 2.216453e+04 | 2.216453e+04 | NaN | NaN | CAEAGANMIVSGSAIMR |
| DOCK11 | DOC11\_HUMAN | Dedicator of cytokinesis protein 11 | 160 | 153 | DOCK11 | DOC11\_HUMAN | Dedicator of cytokinesis protein 11 | 8C(125.0477) | NaN | X | NaN | NaN | NaN | NaN | 2.213317e+04 | 2.213317e+04 | NaN | NaN | DEDSSSLCSQK |
| CCT3 | TCPG\_HUMAN | T-complex protein 1 subunit gamma | 366 | 354 | CCT3 | TCPG\_HUMAN | T-complex protein 1 subunit gamma | 13C(125.0477) | NaN | X | NaN | NaN | NaN | NaN | 2.210720e+04 | 2.210720e+04 | NaN | NaN | KIGDEYFTFITDCK |
| PGK1 | PGK1\_HUMAN | Phosphoglycerate kinase 1 | 108 | 107 | PGK1 | PGK1\_HUMAN | Phosphoglycerate kinase 1 | 2C(125.0477) | NaN | X | NaN | NaN | NaN | NaN | 2.210435e+04 | NaN | 2.210435e+04 | NaN | ACANPAAGSVILLENLR |
| TXNDC5 | TXND5\_HUMAN | Thioredoxin domain-containing protein 5 | 128, 121, 121 | 119 | TXNDC5 | TXND5\_HUMAN | Thioredoxin domain-containing protein 5 | 10C(15.9949) 3C(125.0477) 3C(31.9988) | NaN | X | X | X | NaN | NaN | 2.203853e+04 | NaN | 2.203853e+04 | NaN | VDCTAHSDVCSAQGVR |
| INPP5D | SHIP1\_HUMAN | Phosphatidylinositol 3,4,5-trisphosphate 5-phosphatase 1 | 819 | 808 | INPP5D | SHIP1\_HUMAN | Phosphatidylinositol 3,4,5-trisphosphate 5-phosphatase 1 | 12C(125.0477) | NaN | X | NaN | NaN | NaN | NaN | 2.202206e+04 | 2.202206e+04 | NaN | NaN | SSDSDESYGEGCIALR |
| CCT2 | TCPB\_HUMAN | T-complex protein 1 subunit beta | 289 | 282 | CCT2 | TCPB\_HUMAN | T-complex protein 1 subunit beta | 8C(125.0477) | NaN | X | NaN | NaN | NaN | NaN | 2.201597e+04 | 2.201597e+04 | NaN | NaN | ILKHGINCFINR |
| PSPH | SERB\_HUMAN | Phosphoserine phosphatase | 38 | 37 | PSPH | SERB\_HUMAN | Phosphoserine phosphatase | 2C(125.0477) | NaN | X | NaN | NaN | NaN | NaN | 2.199429e+04 | 2.199429e+04 | NaN | NaN | ICGVEDAVSEMTR |
| ATP6V1A | VATA\_HUMAN | V-type proton ATPase catalytic subunit A | 138 | 133 | ATP6V1A | VATA\_HUMAN | V-type proton ATPase catalytic subunit A | 6C(125.0477) | NaN | X | NaN | NaN | NaN | NaN | 2.197494e+04 | NaN | 2.197494e+04 | NaN | WDFTPCK |
| UBE4A | UBE4A\_HUMAN | Ubiquitin conjugation factor E4 A | 490 | 484 | UBE4A | UBE4A\_HUMAN | Ubiquitin conjugation factor E4 A | 7C(125.0477) | NaN | X | NaN | NaN | NaN | NaN | 2.196648e+04 | 2.196648e+04 | NaN | NaN | GLDKETCLIPAVQEPK |
| PIK3AP1 | BCAP\_HUMAN | Phosphoinositide 3-kinase adapter protein 1 | 147 | 140 | PIK3AP1 | BCAP\_HUMAN | Phosphoinositide 3-kinase adapter protein 1 | 8C(125.0477) | NaN | X | NaN | NaN | NaN | NaN | 2.192440e+04 | 2.192440e+04 | NaN | NaN | AISEDSGCDSVTDTEPEDEK |
| CDC42 | CDC42\_HUMAN | Cell division control protein 42 homolog | 157 | 154 | CDC42 | CDC42\_HUMAN | Cell division control protein 42 homolog | 4C(125.0477) | NaN | X | NaN | NaN | NaN | NaN | 2.191924e+04 | NaN | 2.191924e+04 | NaN | YVECSALTQK |
| TERF2IP | TE2IP\_HUMAN | Telomeric repeat-binding factor 2-interacting protein 1 | 54 | 42 | TERF2IP | TE2IP\_HUMAN | Telomeric repeat-binding factor 2-interacting protein 1 | 13C(125.0477) | NaN | X | NaN | NaN | NaN | NaN | 2.191806e+04 | 2.191806e+04 | NaN | NaN | LSTLILHGGGTVCR |
| HBA1 | HBA\_HUMAN | Hemoglobin subunit alpha | 105 | 101 | HBA1 | HBA\_HUMAN | Hemoglobin subunit alpha | 5C(125.0477) | NaN | X | NaN | NaN | NaN | NaN | 2.187732e+04 | NaN | 2.187732e+04 | 0.0000 | LLSHCLLVTLAAHLPAEFTPAVHASLDKFLASVSTVLTSKYR |
| TSN | TSN\_HUMAN | Translin | 225 | 216 | TSN | TSN\_HUMAN | Translin | 10C(125.0477) | NaN | X | NaN | NaN | NaN | NaN | 2.185758e+04 | 2.185758e+04 | NaN | NaN | GFNKETAAACVEK |
| GLUL | GLNA\_HUMAN | Glutamine synthetase | 99 | 91 | GLUL | GLNA\_HUMAN | Glutamine synthetase | 9C(125.0477) | NaN | X | NaN | NaN | NaN | NaN | 2.182654e+04 | 2.182654e+04 | NaN | NaN | KDPNKLVLCEVFK |
| RAB3GAP1 | RB3GP\_HUMAN | Rab3 GTPase-activating protein catalytic subunit | 218 | 216 | RAB3GAP1 | RB3GP\_HUMAN | Rab3 GTPase-activating protein catalytic subunit | 3C(125.0477) | NaN | X | NaN | NaN | NaN | NaN | 2.174503e+04 | 2.174503e+04 | NaN | NaN | IGCPLTPLPPVSIAIR |
| VPS35 | VPS35\_HUMAN | Vacuolar protein sorting-associated protein 35 | 673 | 669 | VPS35 | VPS35\_HUMAN | Vacuolar protein sorting-associated protein 35 | 5C(125.0477) | NaN | X | NaN | NaN | NaN | NaN | 2.167869e+04 | 2.167869e+04 | NaN | NaN | AVSTCAHLFWSGR |
| TFCP2 | TFCP2\_HUMAN | Alpha-globin transcription factor CP2 | 369 | 363 | TFCP2 | TFCP2\_HUMAN | Alpha-globin transcription factor CP2 | 7C(125.0477) | NaN | X | NaN | NaN | NaN | NaN | 2.167247e+04 | 2.167247e+04 | NaN | NaN | DDVIQICGPADGIR |
| ANK2 | ANK2\_HUMAN | Ankyrin-2 | 305 | 296 | ANK2 | ANK2\_HUMAN | Ankyrin-2 | 10C(125.0477) | NaN | X | NaN | NaN | NaN | NaN | 2.161758e+04 | 2.161758e+04 | NaN | NaN | TRDGLTPLHCAAR |
| PTPN23 | PTN23\_HUMAN | Tyrosine-protein phosphatase non-receptor type 23 | 425 | 425 | PTPN23 | PTN23\_HUMAN | Tyrosine-protein phosphatase non-receptor type 23 | 1C(125.0477) | NaN | X | NaN | NaN | NaN | NaN | 2.153994e+04 | NaN | 2.153994e+04 | NaN | CAALSVRPDTVR |
| ASPSCR1 | ASPC1\_HUMAN | Tether containing UBX domain for GLUT4 | 325 | 316 | ASPSCR1 | ASPC1\_HUMAN | Tether containing UBX domain for GLUT4 | 10C(125.0477) | NaN | X | NaN | NaN | NaN | NaN | 2.152219e+04 | 2.152219e+04 | NaN | NaN | EPVDREPVVCHPDLEER |
| GAN | GAN\_HUMAN | Gigaxonin | 490 | 480 | GAN | GAN\_HUMAN | Gigaxonin | 11C(125.0477) | NaN | X | NaN | NaN | NaN | NaN | 2.152110e+04 | 2.152110e+04 | NaN | NaN | EDAQGSEMVTCK |
| EEF2 | EF2\_HUMAN | Elongation factor 2 | 466 | 457 | EEF2 | EF2\_HUMAN | Elongation factor 2 | 10C(125.0477) | NaN | X | NaN | NaN | NaN | NaN | 2.151898e+04 | 2.151898e+04 | NaN | NaN | YVEPIEDVPCGNIVGLVGVDQFLVK |
| ACTBL2 | ACTBL\_HUMAN | Beta-actin-like protein 2 | 53 | 41 | ACTBL2 | ACTBL\_HUMAN | Beta-actin-like protein 2 | 13C(15.9949) | NaN | NaN | X | NaN | NaN | NaN | 2.151462e+04 | NaN | 2.151462e+04 | NaN | HQGVMVGMGQKDCYVGDEAQSKR |
| PLS3 | PLST\_HUMAN | Plastin-3 | 167 | 145 | PLS3 | PLST\_HUMAN | Plastin-3 | 23C(125.0477) | NaN | X | NaN | NaN | NaN | NaN | 2.151127e+04 | 2.151127e+04 | NaN | NaN | HVIPMNPNTDDLFKAVGDGIVLCK |
| LARP1 | LARP1\_HUMAN | La-related protein 1 | 953 | 949 | LARP1 | LARP1\_HUMAN | La-related protein 1 | 5C(125.0477) | NaN | X | NaN | NaN | NaN | NaN | 2.147543e+04 | 2.147543e+04 | NaN | NaN | YGLECLFR |
| SPTLC2 | SPTC2\_HUMAN | Serine palmitoyltransferase 2 | 188 | 184 | SPTLC2 | SPTC2\_HUMAN | Serine palmitoyltransferase 2 | 5C(125.0477) | NaN | X | NaN | NaN | NaN | NaN | 2.143966e+04 | 2.143966e+04 | NaN | NaN | NTGSCQEAAAK |
| SNX6 | SNX6\_HUMAN | Sorting nexin-6 | 149 | 140 | SNX6 | SNX6\_HUMAN | Sorting nexin-6 | 10C(125.0477) | NaN | X | NaN | NaN | NaN | NaN | 2.141236e+04 | 2.141236e+04 | NaN | NaN | TVAMHEVFLCR |
| PRKAR1A | KAP0\_HUMAN | cAMP-dependent protein kinase type I-alpha regulatory subunit | 362 | 358 | PRKAR1A | KAP0\_HUMAN | cAMP-dependent protein kinase type I-alpha regulatory subunit | 5C(125.0477) | NaN | X | NaN | NaN | NaN | NaN | 2.138145e+04 | 2.138145e+04 | NaN | NaN | VLGPCSDILKR |
| CD38 | CD38\_HUMAN | ADP-ribosyl cyclase/cyclic ADP-ribose hydrolase 1 | 119 | 112 | CD38 | CD38\_HUMAN | ADP-ribosyl cyclase/cyclic ADP-ribose hydrolase 1 | 8C(125.0477) | NaN | X | NaN | NaN | NaN | NaN | 2.135803e+04 | NaN | 2.135803e+04 | NaN | LGTQTVPCNK |
| DDB1 | DDB1\_HUMAN | DNA damage-binding protein 1 | 378 | 370 | DDB1 | DDB1\_HUMAN | DNA damage-binding protein 1 | 9C(125.0477) | NaN | X | NaN | NaN | NaN | NaN | 2.135322e+04 | 1.500870e+04 | 6.344520e+03 | NaN | QGQGQLVTCSGAFK |
| SERPINB1 | ILEU\_HUMAN | Leukocyte elastase inhibitor | 214 | 204 | SERPINB1 | ILEU\_HUMAN | Leukocyte elastase inhibitor | 11C(125.0477) | NaN | X | NaN | NaN | NaN | NaN | 2.133131e+04 | 2.133131e+04 | NaN | NaN | FAYGYIEDLKCR |
| NECAP2 | NECP2\_HUMAN | Adaptin ear-binding coat-associated protein 2 | 133 | 131 | NECAP2 | NECP2\_HUMAN | Adaptin ear-binding coat-associated protein 2 | 3C(125.0477) | NaN | X | NaN | NaN | NaN | NaN | 2.132824e+04 | NaN | 2.132824e+04 | NaN | QQCEFAK |
| NISCH | NISCH\_HUMAN | Nischarin | 403 | 397 | NISCH | NISCH\_HUMAN | Nischarin | 7C(125.0477) | NaN | X | NaN | NaN | NaN | NaN | 2.132378e+04 | 2.132378e+04 | NaN | NaN | SIGSLPCLEHVSLLNNPLSIIPDYR |
| CAPN2 | CAN2\_HUMAN | Calpain-2 catalytic subunit | 341 | 338 | CAPN2 | CAN2\_HUMAN | Calpain-2 catalytic subunit | 4C(125.0477) | NaN | X | NaN | NaN | NaN | NaN | 2.128670e+04 | 2.128670e+04 | NaN | NaN | LEICNLTPDTLTSDTYK |
| FADD | FADD\_HUMAN | FAS-associated death domain protein | 98 | 79 | FADD | FADD\_HUMAN | FAS-associated death domain protein | 20C(125.0477) | NaN | X | NaN | NaN | NaN | NaN | 2.128662e+04 | 2.128662e+04 | NaN | NaN | VDDFEAGAAAGAAPGEEDLCAAFNVICDNVGK |
| FCSK | FCSK\_HUMAN | L-fucose kinase | 484 | 470 | FCSK | FCSK\_HUMAN | L-fucose kinase | 15C(125.0477) | NaN | X | NaN | NaN | NaN | NaN | 2.126716e+04 | 2.126716e+04 | NaN | NaN | AWDLWDPETLPAEYCLPSAR |
| ALB | ALBU\_HUMAN | Albumin | 591 | 585 | ALB | ALBU\_HUMAN | Albumin | 7C(125.0477) | NaN | X | NaN | NaN | NaN | NaN | 2.122308e+04 | NaN | 2.122308e+04 | NaN | ADDKETCFAEEGK |
| AMPD3 | AMPD3\_HUMAN | AMP deaminase 3 | 47 | 31 | AMPD3 | AMPD3\_HUMAN | AMP deaminase 3 | 17C(125.0477) | NaN | X | NaN | NaN | NaN | NaN | 2.121603e+04 | 2.121603e+04 | NaN | NaN | EEDSKDALSLFTVPEDCPIGQK |
| PLCB3 | PLCB3\_HUMAN | 1-phosphatidylinositol 4,5-bisphosphate phosphodiesterase beta-3 | 193 | 185 | PLCB3 | PLCB3\_HUMAN | 1-phosphatidylinositol 4,5-bisphosphate phosphodiesterase beta-3 | 9C(125.0477) | NaN | X | NaN | NaN | NaN | NaN | 2.117362e+04 | 2.117362e+04 | NaN | NaN | RVETALESCGLK |
| ITIH1 | ITIH1\_HUMAN | Inter-alpha-trypsin inhibitor heavy chain H1 | 60 | 55 | ITIH1 | ITIH1\_HUMAN | Inter-alpha-trypsin inhibitor heavy chain H1 | 6C(57.0215) | X | NaN | NaN | NaN | NaN | NaN | 2.116319e+04 | 2.116319e+04 | NaN | NaN | SLKVNCKVTSR |
| LTF | TRFL\_HUMAN | Lactotransferrin | 367 | 364 | LTF | TRFL\_HUMAN | Lactotransferrin | 4C(-34.0809) | NaN | NaN | NaN | NaN | X | NaN | 2.114596e+04 | NaN | NaN | 21145.9570 | VVWCAVGEQELR |
| DDX58 | DDX58\_HUMAN | Antiviral innate immune response receptor RIG-I | 268, 263 | 259 | DDX58 | DDX58\_HUMAN | Antiviral innate immune response receptor RIG-I | 10C(15.9949) 5C(-34.0809) | NaN | NaN | X | NaN | X | NaN | 2.112214e+04 | NaN | 2.112214e+04 | NaN | NTIICAPTGCGK |
| ACTR2 | ARP2\_HUMAN | Actin-related protein 2 | 221 | 220 | ACTR2 | ARP2\_HUMAN | Actin-related protein 2 | 2C(125.0477) | NaN | X | NaN | NaN | NaN | NaN | 2.111394e+04 | 2.111394e+04 | NaN | NaN | LCYVGYNIEQEQK |
| MVD | MVD1\_HUMAN | Diphosphomevalonate decarboxylase | 108 | 98 | MVD | MVD1\_HUMAN | Diphosphomevalonate decarboxylase | 11C(125.0477) | NaN | X | NaN | NaN | NaN | NaN | 2.110386e+04 | 7.434049e+03 | 1.366981e+04 | NaN | DGDPLPSSLSCK |
| PSMC6 | PRS10\_HUMAN | 26S proteasome regulatory subunit 10B | 193 | 186 | PSMC6 | PRS10\_HUMAN | 26S proteasome regulatory subunit 10B | 8C(125.0477) | NaN | X | NaN | NaN | NaN | NaN | 2.108635e+04 | NaN | 2.108635e+04 | NaN | AVASQLDCNFLK |
| DSP | DESP\_HUMAN | Desmoplakin | 682 | 678 | DSP | DESP\_HUMAN | Desmoplakin | 5C(125.0477) | NaN | X | NaN | NaN | NaN | NaN | 2.106860e+04 | 1.069688e+04 | 1.037172e+04 | NaN | QIEHCEGR |
| APRT | APT\_HUMAN | Adenine phosphoribosyltransferase | 83 | 68 | APRT | APT\_HUMAN | Adenine phosphoribosyltransferase | 16C(125.0477) | NaN | X | NaN | NaN | NaN | NaN | 2.105819e+04 | 2.105819e+04 | NaN | NaN | GFLFGPSLAQELGLGCVLIR |
| CAST | ICAL\_HUMAN | Calpastatin | 408 | 397 | CAST | ICAL\_HUMAN | Calpastatin | 12C(31.9988) | NaN | NaN | NaN | X | NaN | NaN | 2.103671e+04 | 2.103671e+04 | NaN | NaN | AAAPAPVSEAVCRTSMCSIQSAPPEPATLKGTVPDDAVEALADSLGK |
| MAT2B | MAT2B\_HUMAN | Methionine adenosyltransferase 2 subunit beta | 93 | 67 | MAT2B | MAT2B\_HUMAN | Methionine adenosyltransferase 2 subunit beta | 27C(125.0477) | NaN | X | NaN | NaN | NaN | NaN | 2.100313e+04 | 2.100313e+04 | NaN | NaN | FEQVNLLDSNAVHHIIHDFQPHVIVHCAAER |
| ABRAXAS2 | ABRX2\_HUMAN | BRISC complex subunit Abraxas 2 | 224 | 220 | ABRAXAS2 | ABRX2\_HUMAN | BRISC complex subunit Abraxas 2 | 5C(125.0477) | NaN | X | NaN | NaN | NaN | NaN | 2.093651e+04 | 2.093651e+04 | NaN | NaN | VQAVCADVEKSER |
| MAP2K2 | MP2K2\_HUMAN | Dual specificity mitogen-activated protein kinase kinase 2 | 384 | 371 | MAP2K2 | MP2K2\_HUMAN | Dual specificity mitogen-activated protein kinase kinase 2 | 14C(125.0477) | NaN | X | NaN | NaN | NaN | NaN | 2.093291e+04 | 2.093291e+04 | NaN | NaN | RSEVEEVDFAGWLCK |
| GLMN | GLMN\_HUMAN | Glomulin | 13 | 13 | GLMN | GLMN\_HUMAN | Glomulin | 1C(125.0477) | NaN | X | NaN | NaN | NaN | NaN | 2.088962e+04 | 2.088962e+04 | NaN | NaN | CQILEEQDFKEEDFGLFQLAGQR |
| UBE3A | UBE3A\_HUMAN | Ubiquitin-protein ligase E3A | 843 | 837 | UBE3A | UBE3A\_HUMAN | Ubiquitin-protein ligase E3A | 7C(125.0477) | NaN | X | NaN | NaN | NaN | NaN | 2.086906e+04 | 2.086906e+04 | NaN | NaN | LPTSHTCFNVLLLPEYSSK |
| CLNS1A | ICLN\_HUMAN | Methylosome subunit pICln | 73 | 71 | CLNS1A | ICLN\_HUMAN | Methylosome subunit pICln | 3C(125.0477) | NaN | X | NaN | NaN | NaN | NaN | 2.079078e+04 | 2.079078e+04 | NaN | NaN | SDCLGEHLYVMVNAK |
| SMAD2 | SMAD2\_HUMAN | Mothers against decapentaplegic homolog 2 | 161 | 158 | SMAD2 | SMAD2\_HUMAN | Mothers against decapentaplegic homolog 2 | 4C(125.0477) | NaN | X | NaN | NaN | NaN | NaN | 2.069047e+04 | 2.069047e+04 | NaN | NaN | DEVCVNPYHYQR |
| FASN | FAS\_HUMAN | Fatty acid synthase | 2468 | 2450 | FASN | FAS\_HUMAN | Fatty acid synthase | 19C(125.0477) | NaN | X | NaN | NaN | NaN | NaN | 2.067153e+04 | 2.067153e+04 | NaN | NaN | TGGAYGEDLGADYNLSQVCDGK |
| PFKP | PFKAP\_HUMAN | ATP-dependent 6-phosphofructokinase, platelet type | 411 | 409 | PFKP | PFKAP\_HUMAN | ATP-dependent 6-phosphofructokinase, platelet type | 3C(31.9988) | NaN | NaN | NaN | X | NaN | NaN | 2.065688e+04 | 2.065688e+04 | NaN | NaN | TNCNVAVINVGAPAAGMNAAVR |
| UNC45A | UN45A\_HUMAN | Protein unc-45 homolog A | 118 | 108 | UNC45A | UN45A\_HUMAN | Protein unc-45 homolog A | 11C(15.9949) | NaN | NaN | X | NaN | NaN | NaN | 2.055896e+04 | 2.055896e+04 | NaN | NaN | LDQAVLDLQRCVSLEPKNK |
| TTLL12 | TTL12\_HUMAN | Tubulin--tyrosine ligase-like protein 12 | 98 | 89 | TTLL12 | TTL12\_HUMAN | Tubulin--tyrosine ligase-like protein 12 | 10C(125.0477) | NaN | X | NaN | NaN | NaN | NaN | 2.055841e+04 | 2.055841e+04 | NaN | NaN | QQPNPGNELCYK |
| CCT5 | TCPE\_HUMAN | T-complex protein 1 subunit epsilon | 181 | 171 | CCT5 | TCPE\_HUMAN | T-complex protein 1 subunit epsilon | 11C(125.0477) | NaN | X | NaN | NaN | NaN | NaN | 2.054869e+04 | 2.054869e+04 | NaN | NaN | TTLGSKVVNSCHR |
| PPP6R1 | PP6R1\_HUMAN | Serine/threonine-protein phosphatase 6 regulatory subunit 1 | 326 | 324 | PPP6R1 | PP6R1\_HUMAN | Serine/threonine-protein phosphatase 6 regulatory subunit 1 | 3C(125.0477) | NaN | X | NaN | NaN | NaN | NaN | 2.053195e+04 | 2.053195e+04 | NaN | NaN | LSCFHQLLLEPPK |
| ENOSF1 | ENOF1\_HUMAN | Mitochondrial enolase superfamily member 1 | 52 | 51 | ENOSF1 | ENOF1\_HUMAN | Mitochondrial enolase superfamily member 1 | 2C(15.9949) | NaN | NaN | X | NaN | NaN | NaN | 2.049010e+04 | 2.049010e+04 | NaN | NaN | GCGITFTLGKGTEVVVCAVNALAHHVLNK |
| WDR37 | WDR37\_HUMAN | WD repeat-containing protein 37 | 8 | 2 | WDR37 | WDR37\_HUMAN | WD repeat-containing protein 37 | 7C(125.0477) | NaN | X | NaN | NaN | NaN | NaN | 2.048938e+04 | 2.048938e+04 | NaN | NaN | PTESASCSTAR |
| GMPR | GMPR1\_HUMAN | GMP reductase 1 | 95 | 87 | GMPR | GMPR1\_HUMAN | GMP reductase 1 | 9C(125.0477) | NaN | X | NaN | NaN | NaN | NaN | 2.048777e+04 | 2.048777e+04 | NaN | NaN | LFATNHPECLQNVAVSSGSGQNDLEK |
| ARHGAP12 | RHG12\_HUMAN | Rho GTPase-activating protein 12 | 199 | 191 | ARHGAP12 | RHG12\_HUMAN | Rho GTPase-activating protein 12 | 9C(125.0477) | NaN | X | NaN | NaN | NaN | NaN | 2.042104e+04 | 2.042104e+04 | NaN | NaN | TSFSQEQSCDSAGEGSER |
| USP24 | UBP24\_HUMAN | Ubiquitin carboxyl-terminal hydrolase 24 | 1556 | 1553 | USP24 | UBP24\_HUMAN | Ubiquitin carboxyl-terminal hydrolase 24 | 4C(125.0477) | NaN | X | NaN | NaN | NaN | NaN | 2.040173e+04 | 2.040173e+04 | NaN | NaN | TAECETSEADNILLAGHLR |
| ABRAXAS2 | ABRX2\_HUMAN | BRISC complex subunit Abraxas 2 | 237 | 233 | ABRAXAS2 | ABRX2\_HUMAN | BRISC complex subunit Abraxas 2 | 5C(125.0477) | NaN | X | NaN | NaN | NaN | NaN | 2.036692e+04 | 2.036692e+04 | NaN | NaN | VVESCQAEVNKLR |
| GMPR | GMPR1\_HUMAN | GMP reductase 1 | 224, 224, 222, 222 | 214 | GMPR | GMPR1\_HUMAN | GMP reductase 1 | 11C(31.9988) 11C(57.0215) 9C(125.0477) 9C(15.9949) | X | X | X | X | NaN | NaN | 2.032044e+04 | 2.032044e+04 | NaN | NaN | GHIISDGGCTCPGDVAK |
| IQGAP2 | IQGA2\_HUMAN | Ras GTPase-activating-like protein IQGAP2 | 42 | 34 | IQGAP2 | IQGA2\_HUMAN | Ras GTPase-activating-like protein IQGAP2 | 9C(125.0477) | NaN | X | NaN | NaN | NaN | NaN | 2.028866e+04 | 2.028866e+04 | NaN | NaN | QNIAYEYLCHLEEAK |
| PPP1CA | PP1A\_HUMAN | Serine/threonine-protein phosphatase PP1-alpha catalytic subunit | 62 | 61 | PPP1CA | PP1A\_HUMAN | Serine/threonine-protein phosphatase PP1-alpha catalytic subunit | 2C(125.0477) | NaN | X | NaN | NaN | NaN | NaN | 2.023290e+04 | 2.023290e+04 | NaN | NaN | ICGDIHGQYYDLLR |
| PGD | 6PGD\_HUMAN | 6-phosphogluconate dehydrogenase, decarboxylating | 402 | 397 | PGD | 6PGD\_HUMAN | 6-phosphogluconate dehydrogenase, decarboxylating | 6C(125.0477) | NaN | X | NaN | NaN | NaN | NaN | 2.023149e+04 | 2.023149e+04 | NaN | NaN | SAVENCQDSWRR |
| NT5DC1 | NT5D1\_HUMAN | 5'-nucleotidase domain-containing protein 1 | 226 | 214 | NT5DC1 | NT5D1\_HUMAN | 5'-nucleotidase domain-containing protein 1 | 13C(125.0477) | NaN | X | NaN | NaN | NaN | NaN | 2.020140e+04 | 2.020140e+04 | NaN | NaN | ILLLITSSHSDYCR |
| CRK | CRK\_HUMAN | Adapter molecule crk | 279 | 269 | CRK | CRK\_HUMAN | Adapter molecule crk | 11C(125.0477) | NaN | X | NaN | NaN | NaN | NaN | 2.018397e+04 | 2.018397e+04 | NaN | NaN | INVSGQWEGECNGK |
| CAPN7 | CAN7\_HUMAN | Calpain-7 | 548 | 545 | CAPN7 | CAN7\_HUMAN | Calpain-7 | 4C(125.0477) | NaN | X | NaN | NaN | NaN | NaN | 2.016467e+04 | 2.016467e+04 | NaN | NaN | ESTCIHSTWDAK |
| FLOT2 | FLOT2\_HUMAN | Flotillin-2 | 197 | 178 | FLOT2 | FLOT2\_HUMAN | Flotillin-2 | 20C(125.0477) | NaN | X | NaN | NaN | NaN | NaN | 2.014475e+04 | 2.014475e+04 | NaN | NaN | DADIGVAEAERDAGIREAECK |
| ASPSCR1 | ASPC1\_HUMAN | Tether containing UBX domain for GLUT4 | 48 | 42 | ASPSCR1 | ASPC1\_HUMAN | Tether containing UBX domain for GLUT4 | 7C(125.0477) | NaN | X | NaN | NaN | NaN | NaN | 2.013982e+04 | 2.013982e+04 | NaN | NaN | RQDFNPCEYDLK |
| DNAJB4 | DNJB4\_HUMAN | DnaJ homolog subfamily B member 4 | 175 | 166 | DNAJB4 | DNJB4\_HUMAN | DnaJ homolog subfamily B member 4 | 10C(125.0477) | NaN | X | NaN | NaN | NaN | NaN | 2.011807e+04 | 2.011807e+04 | NaN | NaN | VSLEEIYSGCTKR |
| ACTR3 | ARP3\_HUMAN | Actin-related protein 3 | 408 | 399 | ACTR3 | ARP3\_HUMAN | Actin-related protein 3 | 10C(125.0477) | NaN | X | NaN | NaN | NaN | NaN | 2.011046e+04 | NaN | 2.011046e+04 | NaN | DYEEIGPSICR |
| HDGF | HDGF\_HUMAN | Hepatoma-derived growth factor | 12 | 12 | HDGF | HDGF\_HUMAN | Hepatoma-derived growth factor | 1C(125.0477) | NaN | X | NaN | NaN | NaN | NaN | 2.004453e+04 | NaN | 2.004453e+04 | NaN | CGDLVFAK |
| FN1 | FINC\_HUMAN | Fibronectin | 401 | 398 | FN1 | FINC\_HUMAN | Fibronectin | 4C(57.0215) | X | NaN | NaN | NaN | NaN | NaN | 1.992398e+04 | 1.992398e+04 | NaN | NaN | YSFCTDHTVLVQTR |
| FLNA | FLNA\_HUMAN | Filamin-A | 2543 | 2541 | FLNA | FLNA\_HUMAN | Filamin-A | 3C(125.0477) | NaN | X | NaN | NaN | NaN | NaN | 1.991481e+04 | 1.991481e+04 | NaN | NaN | ATCAPQHGAPGPGPADASK |
| CCNY | CCNY\_HUMAN | Cyclin-Y | 39 | 35 | CCNY | CCNY\_HUMAN | Cyclin-Y | 5C(125.0477) | NaN | X | NaN | NaN | NaN | NaN | 1.989336e+04 | NaN | 1.989336e+04 | NaN | EDTGCNLQHISDR |
| ULK3 | ULK3\_HUMAN | Serine/threonine-protein kinase ULK3 | 243 | 242 | ULK3 | ULK3\_HUMAN | Serine/threonine-protein kinase ULK3 | 2C(125.0477) | NaN | X | NaN | NaN | NaN | NaN | 1.985039e+04 | 1.985039e+04 | NaN | NaN | DCRDLLQR |
| TTC38 | TTC38\_HUMAN | Tetratricopeptide repeat protein 38 | 367 | 358 | TTC38 | TTC38\_HUMAN | Tetratricopeptide repeat protein 38 | 10C(125.0477) | NaN | X | NaN | NaN | NaN | NaN | 1.983624e+04 | 1.983624e+04 | NaN | NaN | DASESPGENCQHLLAR |
| NPEPPS | PSA\_HUMAN | Puromycin-sensitive aminopeptidase | 66 | 54 | NPEPPS | PSA\_HUMAN | Puromycin-sensitive aminopeptidase | 13C(125.0477) | NaN | X | NaN | NaN | NaN | NaN | 1.983473e+04 | 1.983473e+04 | NaN | NaN | LPADVSPINYSLCLKPDLLDFTFEGKLEAAAQVR |
| ATP2B1 | AT2B1\_HUMAN | Plasma membrane calcium-transporting ATPase 1 | 547 | 545 | ATP2B1 | AT2B1\_HUMAN | Plasma membrane calcium-transporting ATPase 1 | 3C(125.0477) | NaN | X | NaN | NaN | NaN | NaN | 1.982761e+04 | 1.982761e+04 | NaN | NaN | TECALLGLLLDLKR |
| SLC43A2 | LAT4\_HUMAN | Large neutral amino acids transporter small subunit 4 | 392 | 391 | SLC43A2 | LAT4\_HUMAN | Large neutral amino acids transporter small subunit 4 | 2C(125.0477) | NaN | X | NaN | NaN | NaN | NaN | 1.982637e+04 | 1.982637e+04 | NaN | NaN | ECEDASEEPEEKDANQGEK |
| TPI1 | TPIS\_HUMAN | Triosephosphate isomerase | 42 | 34 | TPI1 | TPIS\_HUMAN | Triosephosphate isomerase | 9C(125.0477) | NaN | X | NaN | NaN | NaN | NaN | 1.981471e+04 | NaN | 1.981471e+04 | NaN | VPADTEVVCAPPTAYIDFAR |
| XPOT | XPOT\_HUMAN | Exportin-T | 38 | 30 | XPOT | XPOT\_HUMAN | Exportin-T | 9C(125.0477) | NaN | X | NaN | NaN | NaN | NaN | 1.976004e+04 | 1.976004e+04 | NaN | NaN | ISPDAWQVCAEALAQR |
| VAPA | VAPA\_HUMAN | Vesicle-associated membrane protein-associated protein A | 179 | 174 | VAPA | VAPA\_HUMAN | Vesicle-associated membrane protein-associated protein A | 6C(125.0477) | NaN | X | NaN | NaN | NaN | NaN | 1.974006e+04 | NaN | 1.974006e+04 | NaN | KLMEECK |
| APRT | APT\_HUMAN | Adenine phosphoribosyltransferase | 153 | 146 | APRT | APT\_HUMAN | Adenine phosphoribosyltransferase | 8C(125.0477) | NaN | X | NaN | NaN | NaN | NaN | 1.973901e+04 | 1.973901e+04 | NaN | NaN | LQAEVLECVSLVELTSLKGR |
| IMPACT | IMPCT\_HUMAN | Protein IMPACT | 284 | 280 | IMPACT | IMPCT\_HUMAN | Protein IMPACT | 5C(31.9988) | NaN | NaN | NaN | X | NaN | NaN | 1.973742e+04 | 1.973742e+04 | NaN | NaN | HINNCAR |
| CHORDC1 | CHRD1\_HUMAN | Cysteine and histidine-rich domain-containing protein 1 | 211 | 198 | CHORDC1 | CHRD1\_HUMAN | Cysteine and histidine-rich domain-containing protein 1 | 14C(125.0477) | NaN | X | NaN | NaN | NaN | NaN | 1.973523e+04 | 1.973523e+04 | NaN | NaN | KTSDFNTFLAQEGCTK |
| MARCHF8 | MARH8\_HUMAN | E3 ubiquitin-protein ligase MARCHF8 | 241 | 235 | MARCHF8 | MARH8\_HUMAN | E3 ubiquitin-protein ligase MARCHF8 | 7C(31.9988) | NaN | NaN | NaN | X | NaN | NaN | 1.969528e+04 | NaN | NaN | 19695.2810 | VIYVQNCPETSK |
| DNAJA4 | DNJA4\_HUMAN | DnaJ homolog subfamily A member 4 | 368 | 366 | DNAJA4 | DNJA4\_HUMAN | DnaJ homolog subfamily A member 4 | 3C(125.0477) | NaN | X | NaN | NaN | NaN | NaN | 1.969490e+04 | NaN | 1.969490e+04 | NaN | EFCPNEQNWR |
| ENOPH1 | ENOPH\_HUMAN | Enolase-phosphatase E1 | 202 | 196 | ENOPH1 | ENOPH\_HUMAN | Enolase-phosphatase E1 | 7C(125.0477) | NaN | X | NaN | NaN | NaN | NaN | 1.968630e+04 | 1.968630e+04 | NaN | NaN | IADSIGCSTNNILFLTDVTR |
| DNAJC13 | DJC13\_HUMAN | DnaJ homolog subfamily C member 13 | 1770 | 1762 | DNAJC13 | DJC13\_HUMAN | DnaJ homolog subfamily C member 13 | 9C(125.0477) | NaN | X | NaN | NaN | NaN | NaN | 1.968345e+04 | 1.968345e+04 | NaN | NaN | YNPGSESECIGHFK |
| EIF5 | IF5\_HUMAN | Eukaryotic translation initiation factor 5 | 320 | 313 | EIF5 | IF5\_HUMAN | Eukaryotic translation initiation factor 5 | 8C(125.0477) | NaN | X | NaN | NaN | NaN | NaN | 1.959865e+04 | 1.959865e+04 | NaN | NaN | YLLHGLECVVAMHQAQLISK |
| DHX9 | DHX9\_HUMAN | ATP-dependent RNA helicase A | 438 | 435 | DHX9 | DHX9\_HUMAN | ATP-dependent RNA helicase A | 4C(125.0477) | NaN | X | NaN | NaN | NaN | NaN | 1.959375e+04 | NaN | 1.959375e+04 | NaN | AAECNIVVTQPR |
| SCYL1 | SCYL1\_HUMAN | N-terminal kinase-like protein | 512 | 485 | SCYL1 | SCYL1\_HUMAN | N-terminal kinase-like protein | 28C(31.9988) | NaN | NaN | NaN | X | NaN | NaN | 1.956594e+04 | 1.956594e+04 | NaN | NaN | VAGVLGFAATHNLYSMNDCAQKILPVLCGLTVDPEK |
| LCP1 | PLSL\_HUMAN | Plastin-2 | 460 | 457 | LCP1 | PLSL\_HUMAN | Plastin-2 | 4C(125.0477) | NaN | X | NaN | NaN | NaN | NaN | 1.954254e+04 | NaN | 1.954254e+04 | NaN | LENCNYAVELGK |
| PYCR3 | P5CR3\_HUMAN | Pyrroline-5-carboxylate reductase 3 | 266 | 255 | PYCR3 | P5CR3\_HUMAN | Pyrroline-5-carboxylate reductase 3 | 12C(125.0477) | NaN | X | NaN | NaN | NaN | NaN | 1.953238e+04 | NaN | 1.953238e+04 | NaN | AATMSAVEAATCR |
| PFN1 | PROF1\_HUMAN | Profilin-1 | 128 | 128 | PFN1 | PROF1\_HUMAN | Profilin-1 | 1C(125.0477) | NaN | X | NaN | NaN | NaN | NaN | 1.952494e+04 | 1.952494e+04 | NaN | NaN | CYEMASHLRR |
| WDHD1 | WDHD1\_HUMAN | WD repeat and HMG-box DNA-binding protein 1 | 304 | 288 | WDHD1 | WDHD1\_HUMAN | WD repeat and HMG-box DNA-binding protein 1 | 17C(125.0477) | NaN | X | NaN | NaN | NaN | NaN | 1.952469e+04 | 1.952469e+04 | NaN | NaN | ISYTDAEGNLGLLENVCDPSGK |
| GPX4 | GPX4\_HUMAN | Phospholipid hydroperoxide glutathione peroxidase | 175 | 168 | GPX4 | GPX4\_HUMAN | Phospholipid hydroperoxide glutathione peroxidase | 8C(125.0477) | NaN | X | NaN | NaN | NaN | NaN | 1.951892e+04 | 1.951892e+04 | NaN | NaN | FLIDKNGCVVKR |
| DBNL | DBNL\_HUMAN | Drebrin-like protein | 97 | 95 | DBNL | DBNL\_HUMAN | Drebrin-like protein | 3C(125.0477) | NaN | X | NaN | NaN | NaN | NaN | 1.947096e+04 | 1.947096e+04 | NaN | NaN | GACASHVSTMASFLK |
| DERA | DEOC\_HUMAN | Deoxyribose-phosphate aldolase | 182 | 180 | DERA | DEOC\_HUMAN | Deoxyribose-phosphate aldolase | 3C(125.0477) | NaN | X | NaN | NaN | NaN | NaN | 1.947080e+04 | 1.947080e+04 | NaN | NaN | KACGEAHLK |
| LYZ | LYSC\_HUMAN | Lysozyme C | 83 | 81 | LYZ | LYSC\_HUMAN | Lysozyme C | 3C(125.0477) | NaN | X | NaN | NaN | NaN | NaN | 1.944587e+04 | NaN | 1.944587e+04 | NaN | YWCNDGK |
| DSG1 | DSG1\_HUMAN | Desmoglein-1 | 127 | 118 | DSG1 | DSG1\_HUMAN | Desmoglein-1 | 10C(125.0477) | NaN | X | NaN | NaN | NaN | NaN | 1.942187e+04 | 1.942187e+04 | NaN | NaN | EVTPFFIIYCR |
| PIK3CA | PK3CA\_HUMAN | Phosphatidylinositol 4,5-bisphosphate 3-kinase catalytic subunit alpha isoform | 558 | 556 | PIK3CA | PK3CA\_HUMAN | Phosphatidylinositol 4,5-bisphosphate 3-kinase catalytic subunit alpha isoform | 3C(125.0477) | NaN | X | NaN | NaN | NaN | NaN | 1.939853e+04 | 1.939853e+04 | NaN | NaN | HYCVTIPEILPK |
| PPP6R3 | PP6R3\_HUMAN | Serine/threonine-protein phosphatase 6 regulatory subunit 3 | 216 | 208 | PPP6R3 | PP6R3\_HUMAN | Serine/threonine-protein phosphatase 6 regulatory subunit 3 | 9C(125.0477) | NaN | X | NaN | NaN | NaN | NaN | 1.937115e+04 | 1.937115e+04 | NaN | NaN | HSNASQSLCEIVR |
| UNC13D | UN13D\_HUMAN | Protein unc-13 homolog D | 636 | 626 | UNC13D | UN13D\_HUMAN | Protein unc-13 homolog D | 11C(125.0477) | NaN | X | NaN | NaN | NaN | NaN | 1.936036e+04 | 1.936036e+04 | NaN | NaN | HSTSAVDLSTCFAQISHTAR |
| GDI2 | GDIB\_HUMAN | Rab GDP dissociation inhibitor beta | 317 | 310 | GDI2 | GDIB\_HUMAN | Rab GDP dissociation inhibitor beta | 8C(125.0477) | NaN | X | NaN | NaN | NaN | NaN | 1.935651e+04 | NaN | 1.935651e+04 | NaN | NTNDANSCQIIIPQNQVNR |
| TBC1D5 | TBCD5\_HUMAN | TBC1 domain family member 5 | 706 | 698 | TBC1D5 | TBCD5\_HUMAN | TBC1 domain family member 5 | 9C(125.0477) | NaN | X | NaN | NaN | NaN | NaN | 1.933918e+04 | 1.933918e+04 | NaN | NaN | QASSETPGCTDR |
| PIGR | PIGR\_HUMAN | Polymeric immunoglobulin receptor | 279 | 273 | PIGR | PIGR\_HUMAN | Polymeric immunoglobulin receptor | 7C(125.0477) | NaN | X | NaN | NaN | NaN | NaN | 1.933290e+04 | NaN | 1.933290e+04 | NaN | QSSGENCDVVVNTLGK |
| MAN2C1 | MA2C1\_HUMAN | Alpha-mannosidase 2C1 | 297 | 290 | MAN2C1 | MA2C1\_HUMAN | Alpha-mannosidase 2C1 | 8C(125.0477) | NaN | X | NaN | NaN | NaN | NaN | 1.932868e+04 | 1.932868e+04 | NaN | NaN | NPEFIFACSQAQQLEWVK |
| CDSN | CDSN\_HUMAN | Corneodesmosin | 448 | 443 | CDSN | CDSN\_HUMAN | Corneodesmosin | 6C(125.0477) | NaN | X | NaN | NaN | NaN | NaN | 1.931352e+04 | NaN | 1.931352e+04 | NaN | IILQPCGSK |
| RIMOC1 | RIMC1\_HUMAN | RAB7A-interacting MON1-CCZ1 complex subunit 1 | 143 | 136 | RIMOC1 | RIMC1\_HUMAN | RAB7A-interacting MON1-CCZ1 complex subunit 1 | 8C(125.0477) | NaN | X | NaN | NaN | NaN | NaN | 1.928971e+04 | 1.928971e+04 | NaN | NaN | GALLYMYCHSLTK |
| CASP8 | CASP8\_HUMAN | Caspase-8 | 236 | 234 | CASP8 | CASP8\_HUMAN | Caspase-8 | 3C(125.0477) | NaN | X | NaN | NaN | NaN | NaN | 1.924115e+04 | 1.924115e+04 | NaN | NaN | GYCLIINNHNFAK |
| ACSL3 | ACSL3\_HUMAN | Fatty acid CoA ligase Acsl3 | 85 | 70 | ACSL3 | ACSL3\_HUMAN | Fatty acid CoA ligase Acsl3 | 16C(125.0477) | NaN | X | NaN | NaN | NaN | NaN | 1.922331e+04 | 1.922331e+04 | NaN | NaN | SVNSLDGLASVLYPGCDTLDK |
| AGT | ANGT\_HUMAN | Angiotensinogen | 256 | 254 | AGT | ANGT\_HUMAN | Angiotensinogen | 3C(57.0215) | X | NaN | NaN | NaN | NaN | NaN | 1.921657e+04 | 1.921657e+04 | NaN | NaN | TGCSLMGASVDSTLAFNTYVHFQGK |
| ILF3 | ILF3\_HUMAN | Interleukin enhancer-binding factor 3 | 255 | 248 | ILF3 | ILF3\_HUMAN | Interleukin enhancer-binding factor 3 | 8C(125.0477) | NaN | X | NaN | NaN | NaN | NaN | 1.915073e+04 | 1.915073e+04 | NaN | NaN | GWPLELLCEK |
| TPP2 | TPP2\_HUMAN | Tripeptidyl-peptidase 2 | 209 | 208 | TPP2 | TPP2\_HUMAN | Tripeptidyl-peptidase 2 | 2C(125.0477) | NaN | X | NaN | NaN | NaN | NaN | 1.913993e+04 | NaN | 1.913993e+04 | NaN | ACIDSNEDGDLSK |
| BLVRB | BLVRB\_HUMAN | Flavin reductase (NADPH) | 109 | 106 | BLVRB | BLVRB\_HUMAN | Flavin reductase (NADPH) | 4C(125.0477) | NaN | X | NaN | NaN | NaN | NaN | 1.913555e+04 | NaN | 1.913555e+04 | NaN | VVACTSAFLLWDPTKVPPR |
| FLNB | FLNB\_HUMAN | Filamin-B | 604 | 594 | FLNB | FLNB\_HUMAN | Filamin-B | 11C(125.0477) | NaN | X | NaN | NaN | NaN | NaN | 1.911006e+04 | 1.911006e+04 | NaN | NaN | IEYNDQNDGSCDVK |
| USP15 | UBP15\_HUMAN | Ubiquitin carboxyl-terminal hydrolase 15 | 791 | 788 | USP15 | UBP15\_HUMAN | Ubiquitin carboxyl-terminal hydrolase 15 | 4C(125.0477) | NaN | X | NaN | NaN | NaN | NaN | 1.908153e+04 | 1.908153e+04 | NaN | NaN | LKDCIELFTTK |
| UBE4A | UBE4A\_HUMAN | Ubiquitin conjugation factor E4 A | 710 | 708 | UBE4A | UBE4A\_HUMAN | Ubiquitin conjugation factor E4 A | 3C(125.0477) | NaN | X | NaN | NaN | NaN | NaN | 1.907920e+04 | 1.907920e+04 | NaN | NaN | VFCNFQYAPQLAEALIK |
| KRT13 | K1C13\_HUMAN | Keratin, type I cytoskeletal 13 | 383 | 379 | KRT13 | K1C13\_HUMAN | Keratin, type I cytoskeletal 13 | 5C(125.0477) | NaN | X | NaN | NaN | NaN | NaN | 1.906568e+04 | 1.906568e+04 | NaN | NaN | SEMECQNQEYK |
| ACSL6 | ACSL6\_HUMAN | Long-chain-fatty-acid--CoA ligase 6 | 625 | 616 | ACSL6 | ACSL6\_HUMAN | Long-chain-fatty-acid--CoA ligase 6 | 10C(15.9949) | NaN | NaN | X | NaN | NaN | NaN | 1.906316e+04 | NaN | 1.906316e+04 | NaN | GIEGTYADLCTNK |
| GFUS | FCL\_HUMAN | GDP-L-fucose synthase | 309 | 307 | GFUS | FCL\_HUMAN | GDP-L-fucose synthase | 3C(125.0477) | NaN | X | NaN | NaN | NaN | NaN | 1.905640e+04 | 1.905640e+04 | NaN | NaN | ETCAWFTDNYEQARK |
| FGB | FIBB\_HUMAN | Fibrinogen beta chain | 223 | 207 | FGB | FIBB\_HUMAN | Fibrinogen beta chain | 17C(15.9949) | NaN | NaN | X | NaN | NaN | NaN | 1.903957e+04 | 1.903957e+04 | NaN | NaN | SKIQKLESDVSAQMEYCR |
| TXNRD1 | TRXR1\_HUMAN | Thioredoxin reductase 1, cytoplasmic | 571 | 571 | TXNRD1 | TRXR1\_HUMAN | Thioredoxin reductase 1, cytoplasmic | 1C(15.9949) | NaN | NaN | X | NaN | NaN | NaN | 1.901733e+04 | 1.901733e+04 | NaN | NaN | CYAKIICNTKDNER |
| NIT1 | NIT1\_HUMAN | Deaminated glutathione amidase | 120 | 119 | NIT1 | NIT1\_HUMAN | Deaminated glutathione amidase | 2C(125.0477) | NaN | X | NaN | NaN | NaN | NaN | 1.901549e+04 | 1.901549e+04 | NaN | NaN | ECGLWLSLGGFHER |
| DDT | DOPD\_HUMAN | D-dopachrome decarboxylase | 57 | 38 | DDT | DOPD\_HUMAN | D-dopachrome decarboxylase | 20C(125.0477) | NaN | X | NaN | NaN | NaN | NaN | 1.900877e+04 | 1.900877e+04 | NaN | NaN | VNVTVRPGLAMALSGSTEPCAQLSISSIGVVGTAEDNR |
| RNF123 | RN123\_HUMAN | E3 ubiquitin-protein ligase RNF123 | 1277 | 1273 | RNF123 | RN123\_HUMAN | E3 ubiquitin-protein ligase RNF123 | 5C(15.9949) | NaN | NaN | X | NaN | NaN | NaN | 1.897684e+04 | 1.897684e+04 | NaN | NaN | SCKACINQHLMNNK |
| PGK1 | PGK1\_HUMAN | Phosphoglycerate kinase 1 | 99 | 92 | PGK1 | PGK1\_HUMAN | Phosphoglycerate kinase 1 | 8C(15.9949) | NaN | NaN | X | NaN | NaN | NaN | 1.896830e+04 | 1.896830e+04 | NaN | NaN | DVLFLKDCVGPEVEKACANPAAGSVILLENLR |
| PSMC3 | PRS6A\_HUMAN | 26S proteasome regulatory subunit 6A | 387 | 387 | PSMC3 | PRS6A\_HUMAN | 26S proteasome regulatory subunit 6A | 1C(31.9988) | NaN | NaN | NaN | X | NaN | NaN | 1.892407e+04 | 1.892407e+04 | NaN | NaN | CTDDFNGAQCK |
| TMED7 | TMED7\_HUMAN | Transmembrane emp24 domain-containing protein 7 | 109 | 108 | TMED7 | TMED7\_HUMAN | Transmembrane emp24 domain-containing protein 7 | 2C(125.0477) | NaN | X | NaN | NaN | NaN | NaN | 1.887956e+04 | 1.887956e+04 | NaN | NaN | FCFSNEFSTFTHK |
| SERPINA1 | A1AT\_HUMAN | Alpha-1-antitrypsin | 256 | 248 | SERPINA1 | A1AT\_HUMAN | Alpha-1-antitrypsin | 9C(57.0215) | X | NaN | NaN | NaN | NaN | NaN | 1.887765e+04 | 1.887765e+04 | NaN | NaN | LGMFNIQHCKKLSSWVLLMK |
| IGHV3-7 | HV307\_HUMAN | Immunoglobulin heavy variable 3-7 | 115 | 107 | IGHV3-7 | HV307\_HUMAN | Immunoglobulin heavy variable 3-7 | 9C(57.0215) | X | NaN | NaN | NaN | NaN | NaN | 1.887760e+04 | NaN | 1.887760e+04 | NaN | AEDTAVYYCAR |
| TTC37 | TTC37\_HUMAN | Tetratricopeptide repeat protein 37 | 705 | 699 | TTC37 | TTC37\_HUMAN | Tetratricopeptide repeat protein 37 | 7C(125.0477) | NaN | X | NaN | NaN | NaN | NaN | 1.884093e+04 | 1.884093e+04 | NaN | NaN | ALEYFTCALQHR |
| NISCH | NISCH\_HUMAN | Nischarin | 227 | 218 | NISCH | NISCH\_HUMAN | Nischarin | 10C(125.0477) | NaN | X | NaN | NaN | NaN | NaN | 1.883500e+04 | 1.883500e+04 | NaN | NaN | SLHQVEISHCDAK |
| USP7 | UBP7\_HUMAN | Ubiquitin carboxyl-terminal hydrolase 7 | 799 | 794 | USP7 | UBP7\_HUMAN | Ubiquitin carboxyl-terminal hydrolase 7 | 6C(125.0477) | NaN | X | NaN | NaN | NaN | NaN | 1.882238e+04 | NaN | 1.882238e+04 | NaN | VDVIFCDK |
| LRRC59 | LRC59\_HUMAN | Leucine-rich repeat-containing protein 59 | 48 | 41 | LRRC59 | LRC59\_HUMAN | Leucine-rich repeat-containing protein 59 | 8C(125.0477) | NaN | X | NaN | NaN | NaN | NaN | 1.873305e+04 | NaN | 1.873305e+04 | NaN | ATILDLSCNK |
| ENO1 | ENOA\_HUMAN | Alpha-enolase | 119 | 106 | ENO1 | ENOA\_HUMAN | Alpha-enolase | 14C(125.0477) | NaN | X | NaN | NaN | NaN | NaN | 1.872645e+04 | 1.872645e+04 | NaN | NaN | FGANAILGVSLAVCKAGAVEK |
| TTC37 | TTC37\_HUMAN | Tetratricopeptide repeat protein 37 | 1162 | 1162 | TTC37 | TTC37\_HUMAN | Tetratricopeptide repeat protein 37 | 1C(125.0477) | NaN | X | NaN | NaN | NaN | NaN | 1.872285e+04 | 1.872285e+04 | NaN | NaN | CLLTSAIYALQGR |
| CAST | ICAL\_HUMAN | Calpastatin | 381 | 364 | CAST | ICAL\_HUMAN | Calpastatin | 18C(125.0477) | NaN | X | NaN | NaN | NaN | NaN | 1.869546e+04 | 1.869546e+04 | NaN | NaN | SESELIDELSEDFDRSECK |
| AKR1B1 | ALDR\_HUMAN | Aldo-keto reductase family 1 member B1 | 81 | 79 | AKR1B1 | ALDR\_HUMAN | Aldo-keto reductase family 1 member B1 | 3C(125.0477) | NaN | X | NaN | NaN | NaN | NaN | 1.864036e+04 | 1.864036e+04 | NaN | NaN | LWCTYHEK |
| WIPI1 | WIPI1\_HUMAN | WD repeat domain phosphoinositide-interacting protein 1 | 146 | 134 | WIPI1 | WIPI1\_HUMAN | WD repeat domain phosphoinositide-interacting protein 1 | 13C(125.0477) | NaN | X | NaN | NaN | NaN | NaN | 1.861350e+04 | 1.861350e+04 | NaN | NaN | TLLDIPANPTGLCALSINHSNSYLAYPGSLTSGEIVLYDGNSLK |
| SPTB | SPTB1\_HUMAN | Spectrin beta chain, erythrocytic | 960 | 956 | SPTB | SPTB1\_HUMAN | Spectrin beta chain, erythrocytic | 5C(31.9988) | NaN | NaN | NaN | X | NaN | NaN | 1.860900e+04 | 1.860900e+04 | NaN | NaN | VHNYCVDCEETSK |
| UBE2O | UBE2O\_HUMAN | (E3-independent) E2 ubiquitin-conjugating enzyme | 598 | 594 | UBE2O | UBE2O\_HUMAN | (E3-independent) E2 ubiquitin-conjugating enzyme | 5C(125.0477) | NaN | X | NaN | NaN | NaN | NaN | 1.848115e+04 | 1.848115e+04 | NaN | NaN | RVQSCPDPAVYGVVQSGDHIGR |
| PDE12 | PDE12\_HUMAN | 2',5'-phosphodiesterase 12 | 348 | 338 | PDE12 | PDE12\_HUMAN | 2',5'-phosphodiesterase 12 | 11C(125.0477) | NaN | X | NaN | NaN | NaN | NaN | 1.847691e+04 | 1.847691e+04 | NaN | NaN | ELTGYNADVICLQEVDR |
| TMEM94 | TMM94\_HUMAN | Transmembrane protein 94 | 1206 | 1202 | TMEM94 | TMM94\_HUMAN | Transmembrane protein 94 | 5C(-34.0809) | NaN | NaN | NaN | NaN | X | NaN | 1.845495e+04 | NaN | 1.845495e+04 | NaN | NLTNCSSVMLPSNDDR |
| CHORDC1 | CHRD1\_HUMAN | Cysteine and histidine-rich domain-containing protein 1 | 59 | 46 | CHORDC1 | CHRD1\_HUMAN | Cysteine and histidine-rich domain-containing protein 1 | 14C(125.0477) | NaN | X | NaN | NaN | NaN | NaN | 1.845296e+04 | 1.845296e+04 | NaN | NaN | RTTDFSDFLSIVGCTK |
| SELENBP1 | SBP1\_HUMAN | Methanethiol oxidase | 80, 83 | 70 | SELENBP1 | SBP1\_HUMAN | Methanethiol oxidase | 11C(15.9949) 14C(125.0477) | NaN | X | X | NaN | NaN | NaN | 1.844544e+04 | 1.844544e+04 | NaN | NaN | DELHHSGWNTCSSCFGDSTKSR |
| ESD | ESTD\_HUMAN | S-formylglutathione hydrolase | 243 | 225 | ESD | ESTD\_HUMAN | S-formylglutathione hydrolase | 19C(125.0477) | NaN | X | NaN | NaN | NaN | NaN | 1.840187e+04 | 1.840187e+04 | NaN | NaN | DDQFLLDGQLLPDNFIAACTEK |
| WDFY1 | WDFY1\_HUMAN | WD repeat and FYVE domain-containing protein 1 | 290 | 290 | WDFY1 | WDFY1\_HUMAN | WD repeat and FYVE domain-containing protein 1 | 1C(125.0477) | NaN | X | NaN | NaN | NaN | NaN | 1.839109e+04 | 1.839109e+04 | NaN | NaN | CEQPFFWNIK |
| PLD1 | PLD1\_HUMAN | Phospholipase D1 | 1004 | 1004 | PLD1 | PLD1\_HUMAN | Phospholipase D1 | 1C(125.0477) | NaN | X | NaN | NaN | NaN | NaN | 1.837167e+04 | 1.837167e+04 | NaN | NaN | CLPNDEVHNLIQLR |
| MMS19 | MMS19\_HUMAN | MMS19 nucleotide excision repair protein homolog | 549 | 535 | MMS19 | MMS19\_HUMAN | MMS19 nucleotide excision repair protein homolog | 15C(125.0477) | NaN | X | NaN | NaN | NaN | NaN | 1.836384e+04 | 1.836384e+04 | NaN | NaN | VGESNLTNGDEPTQCSR |
| KDELR1 | ERD21\_HUMAN | ER lumen protein-retaining receptor 1 | 29 | 28 | KDELR1 | ERD21\_HUMAN | ER lumen protein-retaining receptor 1 | 2C(125.0477) | NaN | X | NaN | NaN | NaN | NaN | 1.835694e+04 | NaN | 1.835694e+04 | NaN | SCAGISGK |
| PGRMC2 | PGRC2\_HUMAN | Membrane-associated progesterone receptor component 2 | 159 | 154 | PGRMC2 | PGRC2\_HUMAN | Membrane-associated progesterone receptor component 2 | 6C(125.0477) | NaN | X | NaN | NaN | NaN | NaN | 1.828391e+04 | NaN | 1.828391e+04 | NaN | GLATFCLDK |
| COPS5 | CSN5\_HUMAN | COP9 signalosome complex subunit 5 | 218 | 192 | COPS5 | CSN5\_HUMAN | COP9 signalosome complex subunit 5 | 27C(125.0477) | NaN | X | NaN | NaN | NaN | NaN | 1.822204e+04 | 1.822204e+04 | NaN | NaN | GYKPPDEGPSEYQTIPLNKIEDFGVHCK |
| ROCK2 | ROCK2\_HUMAN | Rho-associated protein kinase 2 | 428 | 418 | ROCK2 | ROCK2\_HUMAN | Rho-associated protein kinase 2 | 11C(125.0477) | NaN | X | NaN | NaN | NaN | NaN | 1.821079e+04 | NaN | 1.821079e+04 | NaN | ENLLLSDSPSCR |
| GMPS | GUAA\_HUMAN | GMP synthase [glutamine-hydrolyzing] | 530 | 525 | GMPS | GUAA\_HUMAN | GMP synthase [glutamine-hydrolyzing] | 6C(125.0477) | NaN | X | NaN | NaN | NaN | NaN | 1.820380e+04 | 1.820380e+04 | NaN | NaN | SYSYVCGISSK |
| IGLL1 | IGLL1\_HUMAN | Immunoglobulin lambda-like polypeptide 1 | 194 | 191 | IGLL1 | IGLL1\_HUMAN | Immunoglobulin lambda-like polypeptide 1 | 4C(15.9949) | NaN | NaN | X | NaN | NaN | NaN | 1.819962e+04 | 0.000000e+00 | 1.568949e+04 | 2510.1316 | SYSCQVMHEGSTVEK |
| ABCC4 | MRP4\_HUMAN | ATP-binding cassette sub-family C member 4 | 1226 | 1221 | ABCC4 | MRP4\_HUMAN | ATP-binding cassette sub-family C member 4 | 6C(125.0477) | NaN | X | NaN | NaN | NaN | NaN | 1.816508e+04 | 1.816508e+04 | NaN | NaN | EKFAHCTVLTIAHR |
| GBF1 | GBF1\_HUMAN | Golgi-specific brefeldin A-resistance guanine nucleotide exchange factor 1 | 403 | 392 | GBF1 | GBF1\_HUMAN | Golgi-specific brefeldin A-resistance guanine nucleotide exchange factor 1 | 12C(125.0477) | NaN | X | NaN | NaN | NaN | NaN | 1.816062e+04 | 1.816062e+04 | NaN | NaN | EGTALVPYGLPCIR |
| WBP2 | WBP2\_HUMAN | WW domain-binding protein 2 | 80 | 79 | WBP2 | WBP2\_HUMAN | WW domain-binding protein 2 | 2C(125.0477) | NaN | X | NaN | NaN | NaN | NaN | 1.803490e+04 | 1.803490e+04 | NaN | NaN | DCEIKQPVFGANYIK |
| SORD | DHSO\_HUMAN | Sorbitol dehydrogenase | 106 | 92 | SORD | DHSO\_HUMAN | Sorbitol dehydrogenase | 15C(125.0477) | NaN | X | NaN | NaN | NaN | NaN | 1.801522e+04 | 1.801522e+04 | NaN | NaN | VAIEPGAPRENDEFCK |
| CAND1 | CAND1\_HUMAN | Cullin-associated NEDD8-dissociated protein 1 | 71, 71 | 71 | CAND1 | CAND1\_HUMAN | Cullin-associated NEDD8-dissociated protein 1 | 1C(-34.0809) 1C(125.0477) | NaN | X | NaN | NaN | X | NaN | 1.793918e+04 | NaN | NaN | 17939.1820 | CLGPLVSK |
| PGK2 | PGK2\_HUMAN | Phosphoglycerate kinase 2 | 108, 108, 108 | 107 | PGK2 | PGK2\_HUMAN | Phosphoglycerate kinase 2 | 2C(125.0477) 2C(15.9949) 2C(31.9988) | NaN | X | X | X | NaN | NaN | 1.790754e+04 | 1.790754e+04 | NaN | NaN | ACANPAPGSVILLENLR |
| AARS1 | SYAC\_HUMAN | Alanine--tRNA ligase, cytoplasmic | 723 | 696 | AARS1 | SYAC\_HUMAN | Alanine--tRNA ligase, cytoplasmic | 28C(125.0477) | NaN | X | NaN | NaN | NaN | NaN | 1.790343e+04 | 1.790343e+04 | NaN | NaN | VVSIGVPVSELLDDPSGPAGSLTSVEFCGGTHLR |
| GPT | ALAT1\_HUMAN | Alanine aminotransferase 1 | 238 | 236 | GPT | ALAT1\_HUMAN | Alanine aminotransferase 1 | 3C(125.0477) | NaN | X | NaN | NaN | NaN | NaN | 1.788704e+04 | 1.788704e+04 | NaN | NaN | ALCVINPGNPTGQVQTR |
| IGHM | IGHM\_HUMAN | Immunoglobulin heavy constant mu | 303 | 301 | IGHM | IGHM\_HUMAN | Immunoglobulin heavy constant mu | 3C(57.0215) | X | NaN | NaN | NaN | NaN | NaN | 1.788155e+04 | NaN | 1.788155e+04 | NaN | FTCTVTHTDLPSPLK |
| STAT5B | STA5B\_HUMAN | Signal transducer and activator of transcription 5B | 688 | 682 | STAT5B | STA5B\_HUMAN | Signal transducer and activator of transcription 5B | 7C(125.0477) | NaN | X | NaN | NaN | NaN | NaN | 1.786304e+04 | NaN | 1.786304e+04 | NaN | YYTPVPCESATAK |
| ANKRD54 | ANR54\_HUMAN | Ankyrin repeat domain-containing protein 54 | 137 | 114 | ANKRD54 | ANR54\_HUMAN | Ankyrin repeat domain-containing protein 54 | 24C(125.0477) | NaN | X | NaN | NaN | NaN | NaN | 1.785910e+04 | 1.785910e+04 | NaN | NaN | LRDSANANDVETVQQLLEDGADPCAADDKGR |
| UBXN6 | UBXN6\_HUMAN | UBX domain-containing protein 6 | 347 | 343 | UBXN6 | UBXN6\_HUMAN | UBX domain-containing protein 6 | 5C(125.0477) | NaN | X | NaN | NaN | NaN | NaN | 1.782774e+04 | 1.782774e+04 | NaN | NaN | LPDGCLLQGTFYAR |
| RANBP1 | RANG\_HUMAN | Ran-specific GTPase-activating protein | 158 | 153 | RANBP1 | RANG\_HUMAN | Ran-specific GTPase-activating protein | 6C(125.0477) | NaN | X | NaN | NaN | NaN | NaN | 1.780630e+04 | 1.780630e+04 | NaN | NaN | TKFEECR |
| FABP5 | FABP5\_HUMAN | Fatty acid-binding protein 5 | 87 | 82 | FABP5 | FABP5\_HUMAN | Fatty acid-binding protein 5 | 6C(125.0477) | NaN | X | NaN | NaN | NaN | NaN | 1.779845e+04 | 1.779845e+04 | NaN | NaN | KTQTVCNFTDGALVQHQEWDGK |
| GNAI3 | GNAI3\_HUMAN | Guanine nucleotide-binding protein G(i) subunit alpha-3 | 325 | 315 | GNAI3 | GNAI3\_HUMAN | Guanine nucleotide-binding protein G(i) subunit alpha-3 | 11C(125.0477) | NaN | X | NaN | NaN | NaN | NaN | 1.772323e+04 | 1.772323e+04 | NaN | NaN | DTKEIYTHFTCATDTK |
| PLAA | PLAP\_HUMAN | Phospholipase A-2-activating protein | 116 | 113 | PLAA | PLAP\_HUMAN | Phospholipase A-2-activating protein | 4C(125.0477) | NaN | X | NaN | NaN | NaN | NaN | 1.768442e+04 | NaN | 1.768442e+04 | NaN | NTVCSLSSGK |
| GNAI2 | GNAI2\_HUMAN | Guanine nucleotide-binding protein G(i) subunit alpha-2 | 287 | 279 | GNAI2 | GNAI2\_HUMAN | Guanine nucleotide-binding protein G(i) subunit alpha-2 | 9C(125.0477) | NaN | X | NaN | NaN | NaN | NaN | 1.761457e+04 | 1.761457e+04 | NaN | NaN | ITHSPLTICFPEYTGANK |
| CERT1 | CERT\_HUMAN | Ceramide transfer protein | 65 | 57 | CERT1 | CERT\_HUMAN | Ceramide transfer protein | 9C(125.0477) | NaN | X | NaN | NaN | NaN | NaN | 1.761283e+04 | 1.761283e+04 | NaN | NaN | SEDETEYGCR |
| ANXA6 | ANXA6\_HUMAN | Annexin A6 | 114 | 114 | ANXA6 | ANXA6\_HUMAN | Annexin A6 | 1C(125.0477) | NaN | X | NaN | NaN | NaN | NaN | 1.760491e+04 | 1.760491e+04 | NaN | NaN | CLIEILASR |
| SH3GL1 | SH3G1\_HUMAN | Endophilin-A2 | 147 | 137 | SH3GL1 | SH3G1\_HUMAN | Endophilin-A2 | 11C(125.0477) | NaN | X | NaN | NaN | NaN | NaN | 1.757305e+04 | 1.757305e+04 | NaN | NaN | QNFIDPLQNLCEK |
| CUL4A | CUL4A\_HUMAN | Cullin-4A | 110 | 108 | CUL4A | CUL4A\_HUMAN | Cullin-4A | 3C(125.0477) | NaN | X | NaN | NaN | NaN | NaN | 1.752425e+04 | 1.752425e+04 | NaN | NaN | QACEDHVQAQILPFR |
| PPP6R3 | PP6R3\_HUMAN | Serine/threonine-protein phosphatase 6 regulatory subunit 3 | 836 | 832 | PPP6R3 | PP6R3\_HUMAN | Serine/threonine-protein phosphatase 6 regulatory subunit 3 | 5C(15.9949) | NaN | NaN | X | NaN | NaN | NaN | 1.745819e+04 | 1.745819e+04 | NaN | NaN | DAEECPETAEAKCAAPRPPSSSPEQR |
| STUB1 | CHIP\_HUMAN | E3 ubiquitin-protein ligase CHIP | 180 | 174 | STUB1 | CHIP\_HUMAN | E3 ubiquitin-protein ligase CHIP | 7C(125.0477) | NaN | X | NaN | NaN | NaN | NaN | 1.745233e+04 | NaN | 1.745233e+04 | NaN | ERELEECQR |
| ALAD | HEM2\_HUMAN | Delta-aminolevulinic acid dehydratase | 162 | 160 | ALAD | HEM2\_HUMAN | Delta-aminolevulinic acid dehydratase | 3C(125.0477) | NaN | X | NaN | NaN | NaN | NaN | 1.744312e+04 | 1.744312e+04 | NaN | NaN | AGCQVVAPSDMMDGRVEAIKEALMAHGLGNR |
| CTBP1 | CTBP1\_HUMAN | C-terminal-binding protein 1 | 54 | 47 | CTBP1 | CTBP1\_HUMAN | C-terminal-binding protein 1 | 8C(125.0477) | NaN | X | NaN | NaN | NaN | NaN | 1.741627e+04 | 1.741627e+04 | NaN | NaN | DVATVAFCDAQSTQEIHEK |
| ARPC1B | ARC1B\_HUMAN | Actin-related protein 2/3 complex subunit 1B | 227 | 217 | ARPC1B | ARC1B\_HUMAN | Actin-related protein 2/3 complex subunit 1B | 11C(125.0477) | NaN | X | NaN | NaN | NaN | NaN | 1.740048e+04 | 1.740048e+04 | NaN | NaN | VAWVSHDSTVCLADADKK |
| SEC14L2 | S14L2\_HUMAN | SEC14-like protein 2 | 301 | 287 | SEC14L2 | S14L2\_HUMAN | SEC14-like protein 2 | 15C(125.0477) | NaN | X | NaN | NaN | NaN | NaN | 1.736256e+04 | 1.736256e+04 | NaN | NaN | GSSHQVEYEILFPGCVLR |
| MPZL1 | MPZL1\_HUMAN | Myelin protein zero-like protein 1 | 203 | 199 | MPZL1 | MPZL1\_HUMAN | Myelin protein zero-like protein 1 | 5C(125.0477) | NaN | X | NaN | NaN | NaN | NaN | 1.728042e+04 | NaN | 1.728042e+04 | NaN | DYTGCSTSESLSPVK |
| MPI | MPI\_HUMAN | Mannose-6-phosphate isomerase | 309 | 302 | MPI | MPI\_HUMAN | Mannose-6-phosphate isomerase | 8C(125.0477) | NaN | X | NaN | NaN | NaN | NaN | 1.727650e+04 | 1.727650e+04 | NaN | NaN | FIDVPTLCEMLSYTPSSSK |
| NUP205 | NU205\_HUMAN | Nuclear pore complex protein Nup205 | 1486 | 1484 | NUP205 | NU205\_HUMAN | Nuclear pore complex protein Nup205 | 3C(125.0477) | NaN | X | NaN | NaN | NaN | NaN | 1.727020e+04 | 1.727020e+04 | NaN | NaN | DACDGHEIGR |
| AFM | AFAM\_HUMAN | Afamin | 124 | 122 | AFM | AFAM\_HUMAN | Afamin | 3C(57.0215) | X | NaN | NaN | NaN | NaN | NaN | 1.726520e+04 | 1.726520e+04 | NaN | NaN | RLCFFYNKK |
| STXBP2 | STXB2\_HUMAN | Syntaxin-binding protein 2 | 353 | 343 | STXBP2 | STXB2\_HUMAN | Syntaxin-binding protein 2 | 11C(125.0477) | NaN | X | NaN | NaN | NaN | NaN | 1.724193e+04 | 1.724193e+04 | NaN | NaN | YSTHLHLADDCMK |
| CCT4 | TCPD\_HUMAN | T-complex protein 1 subunit delta | 295 | 292 | CCT4 | TCPD\_HUMAN | T-complex protein 1 subunit delta | 4C(125.0477) | NaN | X | NaN | NaN | NaN | NaN | 1.723382e+04 | 1.723382e+04 | NaN | NaN | KTGCNVLLIQK |
| PREP | PPCE\_HUMAN | Prolyl endopeptidase | 175 | 173 | PREP | PPCE\_HUMAN | Prolyl endopeptidase | 3C(125.0477) | NaN | X | NaN | NaN | NaN | NaN | 1.721101e+04 | 1.721101e+04 | NaN | NaN | FSCMAWTHDGK |
| CR1 | CR1\_HUMAN | Complement receptor type 1 | 1585 | 1585 | CR1 | CR1\_HUMAN | Complement receptor type 1 | 1C(15.9949) | NaN | NaN | X | NaN | NaN | NaN | 1.715744e+04 | NaN | NaN | 17157.4360 | CISTNKCTAPEVENAIR |
| ACHE | ACES\_HUMAN | Acetylcholinesterase | 440 | 427 | ACHE | ACES\_HUMAN | Acetylcholinesterase | 14C(125.0477) | NaN | X | NaN | NaN | NaN | NaN | 1.713550e+04 | 1.713550e+04 | NaN | NaN | EALSDVVGDHNVVCPVAQLAGR |
| NME2 | NDKB\_HUMAN | Nucleoside diphosphate kinase B | 109 | 106 | NME2 | NDKB\_HUMAN | Nucleoside diphosphate kinase B | 4C(125.0477) | NaN | X | NaN | NaN | NaN | NaN | 1.705552e+04 | NaN | 1.705552e+04 | NaN | GDFCIQVGR |
| ZZEF1 | ZZEF1\_HUMAN | Zinc finger ZZ-type and EF-hand domain-containing protein 1 | 162 | 158 | ZZEF1 | ZZEF1\_HUMAN | Zinc finger ZZ-type and EF-hand domain-containing protein 1 | 5C(125.0477) | NaN | X | NaN | NaN | NaN | NaN | 1.702672e+04 | 1.702672e+04 | NaN | NaN | QLQACSLVPGFTDIFSESK |
| MYLPF | MLRS\_HUMAN | Myosin regulatory light chain 2, skeletal muscle isoform | 157 | 155 | MYLPF | MLRS\_HUMAN | Myosin regulatory light chain 2, skeletal muscle isoform | 3C(57.0215) | X | NaN | NaN | NaN | NaN | NaN | 1.701073e+04 | 1.701073e+04 | NaN | NaN | NICYVITHGDAK |
| KPNB1 | IMB1\_HUMAN | Importin subunit beta-1 | 345 | 333 | KPNB1 | IMB1\_HUMAN | Importin subunit beta-1 | 13C(125.0477) | NaN | X | NaN | NaN | NaN | NaN | 1.700262e+04 | 1.700262e+04 | NaN | NaN | QDENDDDDDWNPCK |
| CSNK1A1 | KC1A\_HUMAN | Casein kinase I isoform alpha | 249 | 241 | CSNK1A1 | KC1A\_HUMAN | Casein kinase I isoform alpha | 9C(125.0477) | NaN | X | NaN | NaN | NaN | NaN | 1.696112e+04 | NaN | 1.696112e+04 | NaN | MSTPVEVLCK |
| DSP | DESP\_HUMAN | Desmoplakin | 954 | 950 | DSP | DESP\_HUMAN | Desmoplakin | 5C(125.0477) | NaN | X | NaN | NaN | NaN | NaN | 1.695311e+04 | NaN | 1.695311e+04 | NaN | IAELCANSIK |
| DPP3 | DPP3\_HUMAN | Dipeptidyl peptidase 3 | 147 | 142 | DPP3 | DPP3\_HUMAN | Dipeptidyl peptidase 3 | 6C(125.0477) | NaN | X | NaN | NaN | NaN | NaN | 1.692236e+04 | 1.692236e+04 | NaN | NaN | GLWQTCGELMFSLEPR |
| PREP | PPCE\_HUMAN | Prolyl endopeptidase | 264 | 261 | PREP | PPCE\_HUMAN | Prolyl endopeptidase | 4C(125.0477) | NaN | X | NaN | NaN | NaN | NaN | 1.690718e+04 | 1.690718e+04 | NaN | NaN | LWYCDLQQESSGIAGILK |
| UBE2V1 | UB2V1\_HUMAN | Ubiquitin-conjugating enzyme E2 variant 1 | 71 | 64 | UBE2V1 | UB2V1\_HUMAN | Ubiquitin-conjugating enzyme E2 variant 1 | 8C(125.0477) | NaN | X | NaN | NaN | NaN | NaN | 1.686741e+04 | 1.686741e+04 | NaN | NaN | IYSLKIECGPKYPEAPPFVR |
| DYNLL2 | DYL2\_HUMAN | Dynein light chain 2, cytoplasmic | 56 | 45 | DYNLL2 | DYL2\_HUMAN | Dynein light chain 2, cytoplasmic | 12C(125.0477) | NaN | X | NaN | NaN | NaN | NaN | 1.682872e+04 | 1.682872e+04 | NaN | NaN | EFDKKYNPTWHCIVGR |
| EPB42 | EPB42\_HUMAN | Protein 4.2 | 10 | 9 | EPB42 | EPB42\_HUMAN | Protein 4.2 | 2C(125.0477) | NaN | X | NaN | NaN | NaN | NaN | 1.680946e+04 | 1.680946e+04 | NaN | NaN | SCDFQAARNNEEHHTK |
| RAB3GAP2 | RBGPR\_HUMAN | Rab3 GTPase-activating protein non-catalytic subunit | 636 | 622 | RAB3GAP2 | RBGPR\_HUMAN | Rab3 GTPase-activating protein non-catalytic subunit | 15C(125.0477) | NaN | X | NaN | NaN | NaN | NaN | 1.679472e+04 | 1.679472e+04 | NaN | NaN | SQELESVDEGLLQFCANK |
| SCRN2 | SCRN2\_HUMAN | Secernin-2 | 59 | 57 | SCRN2 | SCRN2\_HUMAN | Secernin-2 | 3C(125.0477) | NaN | X | NaN | NaN | NaN | NaN | 1.677367e+04 | 1.677367e+04 | NaN | NaN | LQCTYIEVEQVSK |
| DNAJA4 | DNJA4\_HUMAN | DnaJ homolog subfamily A member 4 | 138 | 138 | DNAJA4 | DNJA4\_HUMAN | DnaJ homolog subfamily A member 4 | 1C(125.0477) | NaN | X | NaN | NaN | NaN | NaN | 1.675665e+04 | 1.675665e+04 | NaN | NaN | CEGVGGKK |
| RNF123 | RN123\_HUMAN | E3 ubiquitin-protein ligase RNF123 | 1180 | 1163 | RNF123 | RN123\_HUMAN | E3 ubiquitin-protein ligase RNF123 | 18C(125.0477) | NaN | X | NaN | NaN | NaN | NaN | 1.672113e+04 | 1.672113e+04 | NaN | NaN | GPASEREQATSVLLADPCFQLR |
| BPNT1 | BPNT1\_HUMAN | 3'(2'),5'-bisphosphate nucleotidase 1 | 249 | 245 | BPNT1 | BPNT1\_HUMAN | 3'(2'),5'-bisphosphate nucleotidase 1 | 5C(125.0477) | NaN | X | NaN | NaN | NaN | NaN | 1.665510e+04 | 1.665510e+04 | NaN | NaN | KWDTCAPEVILHAVGGK |
| HMBS | HEM3\_HUMAN | Porphobilinogen deaminase | 114 | 80 | HMBS | HEM3\_HUMAN | Porphobilinogen deaminase | 35C(125.0477) | NaN | X | NaN | NaN | NaN | NaN | 1.662622e+04 | 1.662622e+04 | NaN | NaN | ELEHALEKNEVDLVVHSLKDLPTVLPPGFTIGAICK |
| GART | PUR2\_HUMAN | Trifunctional purine biosynthetic protein adenosine-3 | 62 | 45 | GART | PUR2\_HUMAN | Trifunctional purine biosynthetic protein adenosine-3 | 18C(125.0477) | NaN | X | NaN | NaN | NaN | NaN | 1.662010e+04 | 1.662010e+04 | NaN | NaN | ISNTAISISDHTALAQFCKEK |
| DAAM1 | DAAM1\_HUMAN | Disheveled-associated activator of morphogenesis 1 | 696 | 693 | DAAM1 | DAAM1\_HUMAN | Disheveled-associated activator of morphogenesis 1 | 4C(125.0477) | NaN | X | NaN | NaN | NaN | NaN | 1.661438e+04 | NaN | 1.661438e+04 | NaN | AQNCNILLSR |
| CUL1 | CUL1\_HUMAN | Cullin-1 | 752 | 751 | CUL1 | CUL1\_HUMAN | Cullin-1 | 2C(125.0477) | NaN | X | NaN | NaN | NaN | NaN | 1.660284e+04 | 1.660284e+04 | NaN | NaN | KCIDILIEK |
| PPP6C | PPP6\_HUMAN | Serine/threonine-protein phosphatase 6 catalytic subunit | 265 | 265 | PPP6C | PPP6\_HUMAN | Serine/threonine-protein phosphatase 6 catalytic subunit | 1C(125.0477) | NaN | X | NaN | NaN | NaN | NaN | 1.657201e+04 | 1.657201e+04 | NaN | NaN | CGNIASIMVFKDVNTR |
| C3 | CO3\_HUMAN | Complement C3 | 559 | 557 | C3 | CO3\_HUMAN | Complement C3 | 3C(57.0215) | X | NaN | NaN | NaN | NaN | NaN | 1.655581e+04 | NaN | 1.655581e+04 | NaN | DSCVGSLVVK |
| VAC14 | VAC14\_HUMAN | Protein VAC14 homolog | 79 | 70 | VAC14 | VAC14\_HUMAN | Protein VAC14 homolog | 10C(125.0477) | NaN | X | NaN | NaN | NaN | NaN | 1.652818e+04 | 1.652818e+04 | NaN | NaN | KGGLIGLAACSIALGK |
| PRKDC | PRKDC\_HUMAN | DNA-dependent protein kinase catalytic subunit | 90 | 88 | PRKDC | PRKDC\_HUMAN | DNA-dependent protein kinase catalytic subunit | 3C(125.0477) | NaN | X | NaN | NaN | NaN | NaN | 1.652006e+04 | 1.652006e+04 | NaN | NaN | FLCIFLEK |
| DCAF8 | DCAF8\_HUMAN | DDB1- and CUL4-associated factor 8 | 377 | 376 | DCAF8 | DCAF8\_HUMAN | DDB1- and CUL4-associated factor 8 | 2C(125.0477) | NaN | X | NaN | NaN | NaN | NaN | 1.648770e+04 | 1.648770e+04 | NaN | NaN | FCPHHLVNSESK |
| PIEZO1 | PIEZ1\_HUMAN | Piezo-type mechanosensitive ion channel component 1 | 1151 | 1135 | PIEZO1 | PIEZ1\_HUMAN | Piezo-type mechanosensitive ion channel component 1 | 17C(125.0477) | NaN | X | NaN | NaN | NaN | NaN | 1.643025e+04 | 1.643025e+04 | NaN | NaN | LEPLRGEPNPVPNFIHCR |
| IAH1 | IAH1\_HUMAN | Isoamyl acetate-hydrolyzing esterase 1 homolog | 4 | 2 | IAH1 | IAH1\_HUMAN | Isoamyl acetate-hydrolyzing esterase 1 homolog | 3C(57.0215) | X | NaN | NaN | NaN | NaN | NaN | 1.642004e+04 | 1.642004e+04 | NaN | NaN | ALCEAAGCGSALLWPR |
| SELENOO | SELO\_HUMAN | Protein adenylyltransferase SelO, mitochondrial | 219 | 216 | SELENOO | SELO\_HUMAN | Protein adenylyltransferase SelO, mitochondrial | 4C(125.0477) | NaN | X | NaN | NaN | NaN | NaN | 1.636144e+04 | NaN | 1.636144e+04 | NaN | AGACVTSESTVVR |
| GMPPB | GMPPB\_HUMAN | Mannose-1-phosphate guanyltransferase beta | 149 | 138 | GMPPB | GMPPB\_HUMAN | Mannose-1-phosphate guanyltransferase beta | 12C(125.0477) | NaN | X | NaN | NaN | NaN | NaN | 1.633417e+04 | 1.633417e+04 | NaN | NaN | VEEPSKYGVVVCEADTGR |
| PEPD | PEPD\_HUMAN | Xaa-Pro dipeptidase | 58 | 57 | PEPD | PEPD\_HUMAN | Xaa-Pro dipeptidase | 2C(125.0477) | NaN | X | NaN | NaN | NaN | NaN | 1.631031e+04 | NaN | 1.631031e+04 | NaN | YCTDTGVLFR |
| EIF1AX | IF1AX\_HUMAN | Eukaryotic translation initiation factor 1A, X-chromosomal | 51 | 47 | EIF1AX | IF1AX\_HUMAN | Eukaryotic translation initiation factor 1A, X-chromosomal | 5C(125.0477) | NaN | X | NaN | NaN | NaN | NaN | 1.630988e+04 | 1.630988e+04 | NaN | NaN | LEAMCFDGVK |
| MTMR12 | MTMRC\_HUMAN | Myotubularin-related protein 12 | 67 | 61 | MTMR12 | MTMRC\_HUMAN | Myotubularin-related protein 12 | 7C(125.0477) | NaN | X | NaN | NaN | NaN | NaN | 1.628445e+04 | 1.628445e+04 | NaN | NaN | YVQEDSCQHGVYGR |
| PTPRA | PTPRA\_HUMAN | Receptor-type tyrosine-protein phosphatase alpha | 668 | 664 | PTPRA | PTPRA\_HUMAN | Receptor-type tyrosine-protein phosphatase alpha | 5C(125.0477) | NaN | X | NaN | NaN | NaN | NaN | 1.626676e+04 | 1.626676e+04 | NaN | NaN | KEEECESYTVR |
| LYPLA2 | LYPA2\_HUMAN | Acyl-protein thioesterase 2 | 56 | 54 | LYPLA2 | LYPA2\_HUMAN | Acyl-protein thioesterase 2 | 3C(125.0477) | NaN | X | NaN | NaN | NaN | NaN | 1.621949e+04 | 1.621949e+04 | NaN | NaN | YICPHAPR |
| KIF5B | KINH\_HUMAN | Kinesin-1 heavy chain | 608 | 608 | KIF5B | KINH\_HUMAN | Kinesin-1 heavy chain | 1C(125.0477) | NaN | X | NaN | NaN | NaN | NaN | 1.620808e+04 | 1.620808e+04 | NaN | NaN | CKQLESTQTESNK |
| STK16 | STK16\_HUMAN | Serine/threonine-protein kinase 16 | 53 | 51 | STK16 | STK16\_HUMAN | Serine/threonine-protein kinase 16 | 3C(125.0477) | NaN | X | NaN | NaN | NaN | NaN | 1.619392e+04 | 1.619392e+04 | NaN | NaN | ILCHEQQDREEAQR |
| NME7 | NDK7\_HUMAN | Nucleoside diphosphate kinase 7 | 356 | 350 | NME7 | NDK7\_HUMAN | Nucleoside diphosphate kinase 7 | 7C(125.0477) | NaN | X | NaN | NaN | NaN | NaN | 1.618153e+04 | 1.618153e+04 | NaN | NaN | IQNAVHCTDLPEDGLLEVQYFFK |
| ERP44 | ERP44\_HUMAN | Endoplasmic reticulum resident protein 44 | 189 | 181 | ERP44 | ERP44\_HUMAN | Endoplasmic reticulum resident protein 44 | 9C(125.0477) | NaN | X | NaN | NaN | NaN | NaN | 1.617131e+04 | 1.617131e+04 | NaN | NaN | VANILHDDCAFLSAFGDVSKPER |
| IPO5 | IPO5\_HUMAN | Importin-5 | 733 | 721 | IPO5 | IPO5\_HUMAN | Importin-5 | 13C(125.0477) | NaN | X | NaN | NaN | NaN | NaN | 1.616895e+04 | 1.616895e+04 | NaN | NaN | VAAAESMPLLLECAR |
| CRNN | CRNN\_HUMAN | Cornulin | 425 | 424 | CRNN | CRNN\_HUMAN | Cornulin | 2C(125.0477) | NaN | X | NaN | NaN | NaN | NaN | 1.615543e+04 | NaN | 1.615543e+04 | NaN | RCVTEGQGDR |
| FSCN1 | FSCN1\_HUMAN | Fascin | 260 | 251 | FSCN1 | FSCN1\_HUMAN | Fascin | 10C(125.0477) | NaN | X | NaN | NaN | NaN | NaN | 1.615047e+04 | 1.615047e+04 | NaN | NaN | DELFALEQSCAQVVLQAANER |
| NUDT2 | AP4A\_HUMAN | Bis(5'-nucleosyl)-tetraphosphatase [asymmetrical] | 123 | 116 | NUDT2 | AP4A\_HUMAN | Bis(5'-nucleosyl)-tetraphosphatase [asymmetrical] | 8C(125.0477) | NaN | X | NaN | NaN | NaN | NaN | 1.609012e+04 | 1.609012e+04 | NaN | NaN | WLGLEEACQLAQFK |
| CFHR1 | FHR1\_HUMAN | Complement factor H-related protein 1 | 300 | 292 | CFHR1 | FHR1\_HUMAN | Complement factor H-related protein 1 | 9C(57.0215) | X | NaN | NaN | NaN | NaN | NaN | 1.602202e+04 | NaN | 1.602202e+04 | NaN | TGESAEFVCKR |
| EFHD2 | EFHD2\_HUMAN | EF-hand domain-containing protein D2 | 172 | 160 | EFHD2 | EFHD2\_HUMAN | EF-hand domain-containing protein D2 | 13C(125.0477) | NaN | X | NaN | NaN | NaN | NaN | 1.600069e+04 | 1.600069e+04 | NaN | NaN | AAAGELQEDSGLCVLAR |
| MAT2B | MAT2B\_HUMAN | Methionine adenosyltransferase 2 subunit beta | 58 | 43 | MAT2B | MAT2B\_HUMAN | Methionine adenosyltransferase 2 subunit beta | 16C(125.0477) | NaN | X | NaN | NaN | NaN | NaN | 1.598527e+04 | 1.598527e+04 | NaN | NaN | AVHKEFQQNNWHAVGCGFR |
| NRDC | NRDC\_HUMAN | Nardilysin | 674 | 671 | NRDC | NRDC\_HUMAN | Nardilysin | 4C(125.0477) | NaN | X | NaN | NaN | NaN | NaN | 1.597400e+04 | 1.597400e+04 | NaN | NaN | AFDCPETEYPVK |
| GMPS | GUAA\_HUMAN | GMP synthase [glutamine-hydrolyzing] | 554 | 553 | GMPS | GUAA\_HUMAN | GMP synthase [glutamine-hydrolyzing] | 2C(125.0477) | NaN | X | NaN | NaN | NaN | NaN | 1.594716e+04 | 1.594716e+04 | NaN | NaN | MCHNVNR |
| RAN | RAN\_HUMAN | GTP-binding nuclear protein Ran | 85 | 77 | RAN | RAN\_HUMAN | GTP-binding nuclear protein Ran | 9C(125.0477) | NaN | X | NaN | NaN | NaN | NaN | 1.594188e+04 | 1.594188e+04 | NaN | NaN | DGYYIQAQCAIIMFDVTSR |
| AMBP | AMBP\_HUMAN | Protein AMBP | 91 | 89 | AMBP | AMBP\_HUMAN | Protein AMBP | 3C(125.0477) | NaN | X | NaN | NaN | NaN | NaN | 1.592639e+04 | NaN | 1.592639e+04 | NaN | GVCEETSGAYEK |
| USO1 | USO1\_HUMAN | General vesicular transport factor p115 | 478 | 461 | USO1 | USO1\_HUMAN | General vesicular transport factor p115 | 18C(125.0477) | NaN | X | NaN | NaN | NaN | NaN | 1.592603e+04 | 1.592603e+04 | NaN | NaN | VQLATSIGNPPVSLLQQCTNILSQGSK |
| CFL1 | COF1\_HUMAN | Cofilin-1 | 39 | 33 | CFL1 | COF1\_HUMAN | Cofilin-1 | 7C(125.0477) | NaN | X | NaN | NaN | NaN | NaN | 1.589808e+04 | 1.589808e+04 | NaN | NaN | KKAVLFCLSEDK |
| MPO | PERM\_HUMAN | Myeloperoxidase | 730 | 726 | MPO | PERM\_HUMAN | Myeloperoxidase | 5C(57.0215) | X | NaN | NaN | NaN | NaN | NaN | 1.589653e+04 | 1.589653e+04 | NaN | NaN | DFVNCSTLPALNLASWR |
| OPTN | OPTN\_HUMAN | Optineurin | 472 | 466 | OPTN | OPTN\_HUMAN | Optineurin | 7C(125.0477) | NaN | X | NaN | NaN | NaN | NaN | 1.584646e+04 | 1.584646e+04 | NaN | NaN | AQMEVYCSDFHAER |
| DNASE1L1 | DNSL1\_HUMAN | Deoxyribonuclease-1-like 1 | 224 | 220 | DNASE1L1 | DNSL1\_HUMAN | Deoxyribonuclease-1-like 1 | 5C(125.0477) | NaN | X | NaN | NaN | NaN | NaN | 1.583957e+04 | NaN | 1.583957e+04 | NaN | ASTHCTYDR |
| UGP2 | UGPA\_HUMAN | UTP--glucose-1-phosphate uridylyltransferase | 181 | 174 | UGP2 | UGPA\_HUMAN | UTP--glucose-1-phosphate uridylyltransferase | 8C(125.0477) | NaN | X | NaN | NaN | NaN | NaN | 1.583824e+04 | 1.583824e+04 | NaN | NaN | ILQKYNHCR |
| TUBA4A | TBA4A\_HUMAN | Tubulin alpha-4A chain | 347 | 340 | TUBA4A | TBA4A\_HUMAN | Tubulin alpha-4A chain | 8C(57.0215) | X | NaN | NaN | NaN | NaN | NaN | 1.582496e+04 | NaN | NaN | 15824.9570 | SIQFVDWCPTGFK |
| CAPN1 | CAN1\_HUMAN | Calpain-1 catalytic subunit | 351 | 348 | CAPN1 | CAN1\_HUMAN | Calpain-1 catalytic subunit | 4C(125.0477) | NaN | X | NaN | NaN | NaN | NaN | 1.581381e+04 | 1.581381e+04 | NaN | NaN | LEICNLTPDALKSR |
| SLC4A1 | B3AT\_HUMAN | Band 3 anion transport protein | 843 | 833 | SLC4A1 | B3AT\_HUMAN | Band 3 anion transport protein | 11C(15.9949) | NaN | NaN | X | NaN | NaN | NaN | 1.579978e+04 | 1.579978e+04 | NaN | NaN | MHLFTGIQIICLAVLWVVK |
| WNK1 | WNK1\_HUMAN | Serine/threonine-protein kinase WNK1 | 296 | 296 | WNK1 | WNK1\_HUMAN | Serine/threonine-protein kinase WNK1 | 1C(125.0477) | NaN | X | NaN | NaN | NaN | NaN | 1.579945e+04 | 1.579945e+04 | NaN | NaN | CIVLVTELMTSGTLK |
| ANKFY1 | ANFY1\_HUMAN | Rabankyrin-5 | 34 | 34 | ANKFY1 | ANFY1\_HUMAN | Rabankyrin-5 | 1C(125.0477) | NaN | X | NaN | NaN | NaN | NaN | 1.579107e+04 | 1.579107e+04 | NaN | NaN | CALLAAQANKESSSESFISR |
| UBR4 | UBR4\_HUMAN | E3 ubiquitin-protein ligase UBR4 | 3414 | 3414 | UBR4 | UBR4\_HUMAN | E3 ubiquitin-protein ligase UBR4 | 1C(125.0477) | NaN | X | NaN | NaN | NaN | NaN | 1.578827e+04 | 1.578827e+04 | NaN | NaN | CFLLESNSSSVR |
| AP4B1 | AP4B1\_HUMAN | AP-4 complex subunit beta-1 | 320 | 318 | AP4B1 | AP4B1\_HUMAN | AP-4 complex subunit beta-1 | 3C(125.0477) | NaN | X | NaN | NaN | NaN | NaN | 1.577065e+04 | 1.577065e+04 | NaN | NaN | FFCSYSEPHYIK |
| DDX3X | DDX3X\_HUMAN | ATP-dependent RNA helicase DDX3X | 317 | 316 | DDX3X | DDX3X\_HUMAN | ATP-dependent RNA helicase DDX3X | 2C(125.0477) | NaN | X | NaN | NaN | NaN | NaN | 1.576251e+04 | 1.576251e+04 | NaN | NaN | GCHLLVATPGR |
| METTL26 | MTL26\_HUMAN | Methyltransferase-like 26 | 63 | 63 | METTL26 | MTL26\_HUMAN | Methyltransferase-like 26 | 1C(125.0477) | NaN | X | NaN | NaN | NaN | NaN | 1.566921e+04 | 1.566921e+04 | NaN | NaN | CLDSIAATTQAQGLTNVK |
| FABP5 | FABP5\_HUMAN | Fatty acid-binding protein 5 | 67 | 62 | FABP5 | FABP5\_HUMAN | Fatty acid-binding protein 5 | 6C(125.0477) | NaN | X | NaN | NaN | NaN | NaN | 1.566425e+04 | 1.566425e+04 | NaN | NaN | TTQFSCTLGEKFEETTADGRK |
| USP47 | UBP47\_HUMAN | Ubiquitin carboxyl-terminal hydrolase 47 | 507 | 483 | USP47 | UBP47\_HUMAN | Ubiquitin carboxyl-terminal hydrolase 47 | 25C(125.0477) | NaN | X | NaN | NaN | NaN | NaN | 1.563569e+04 | 1.563569e+04 | NaN | NaN | NSLIYELFSVMVHSGSAAGGHYYACIK |
| RNPEP | AMPB\_HUMAN | Aminopeptidase B | 85 | 80 | RNPEP | AMPB\_HUMAN | Aminopeptidase B | 6C(125.0477) | NaN | X | NaN | NaN | NaN | NaN | 1.563360e+04 | 1.563360e+04 | NaN | NaN | LDSHPCLEVTAAALRR |
| EHBP1L1 | EH1L1\_HUMAN | EH domain-binding protein 1-like protein 1 | 1364 | 1356 | EHBP1L1 | EH1L1\_HUMAN | EH domain-binding protein 1-like protein 1 | 9C(125.0477) | NaN | X | NaN | NaN | NaN | NaN | 1.560098e+04 | 1.560098e+04 | NaN | NaN | FQDTSQYVCAELQALEQEQR |
| CSNK2A2 | CSK22\_HUMAN | Casein kinase II subunit alpha' | 336 | 331 | CSNK2A2 | CSK22\_HUMAN | Casein kinase II subunit alpha' | 6C(125.0477) | NaN | X | NaN | NaN | NaN | NaN | 1.558096e+04 | 1.558096e+04 | NaN | NaN | EQSQPCADNAVLSSGLTAAR |
| TBC1D15 | TBC15\_HUMAN | TBC1 domain family member 15 | 24 | 11 | TBC1D15 | TBC15\_HUMAN | TBC1 domain family member 15 | 14C(125.0477) | NaN | X | NaN | NaN | NaN | NaN | 1.557263e+04 | 1.557263e+04 | NaN | NaN | IIYEQEGVYIHSSCGK |
| NBEAL2 | NBEL2\_HUMAN | Neurobeachin-like protein 2 | 1661 | 1661 | NBEAL2 | NBEL2\_HUMAN | Neurobeachin-like protein 2 | 1C(125.0477) | NaN | X | NaN | NaN | NaN | NaN | 1.552818e+04 | 1.552818e+04 | NaN | NaN | CSWLVPLVR |
| HBD | HBD\_HUMAN | Hemoglobin subunit delta | 113 | 97 | HBD | HBD\_HUMAN | Hemoglobin subunit delta | 17C(125.0477) | NaN | X | NaN | NaN | NaN | NaN | 1.548682e+04 | 1.548682e+04 | NaN | NaN | LHVDPENFRLLGNVLVCVLARNFGK |
| CP | CERU\_HUMAN | Ceruloplasmin | 534 | 524 | CP | CERU\_HUMAN | Ceruloplasmin | 11C(125.0477) | NaN | X | NaN | NaN | NaN | NaN | 1.545157e+04 | NaN | 1.545157e+04 | NaN | EVGPTNADPVCLAK |
| RAN | RAN\_HUMAN | GTP-binding nuclear protein Ran | 85 | 72 | RAN | RAN\_HUMAN | GTP-binding nuclear protein Ran | 14C(125.0477) | NaN | X | NaN | NaN | NaN | NaN | 1.544463e+04 | 1.544463e+04 | NaN | NaN | FGGLRDGYYIQAQCAIIMFDVTSRVTYK |
| CMPK2 | CMPK2\_HUMAN | UMP-CMP kinase 2, mitochondrial | 189 | 185 | CMPK2 | CMPK2\_HUMAN | UMP-CMP kinase 2, mitochondrial | 5C(125.0477) | NaN | X | NaN | NaN | NaN | NaN | 1.542959e+04 | 1.542959e+04 | NaN | NaN | LQVGCAQVVPVPEPPLHPVVPDLPSSVVFPDR |
| G6PD | G6PD\_HUMAN | Glucose-6-phosphate 1-dehydrogenase | 158 | 153 | G6PD | G6PD\_HUMAN | Glucose-6-phosphate 1-dehydrogenase | 6C(125.0477) | NaN | X | NaN | NaN | NaN | NaN | 1.540958e+04 | 1.540958e+04 | NaN | NaN | NIHESCMSQIGWNR |
| C20orf27 | CT027\_HUMAN | UPF0687 protein C20orf27 | 156 | 156 | C20orf27 | CT027\_HUMAN | UPF0687 protein C20orf27 | 1C(125.0477) | NaN | X | NaN | NaN | NaN | NaN | 1.539201e+04 | 1.539201e+04 | NaN | NaN | CVGAELEYDSEHSDWHGFD |
| SDSL | SDSL\_HUMAN | Serine dehydratase-like | 56 | 51 | SDSL | SDSL\_HUMAN | Serine dehydratase-like | 6C(125.0477) | NaN | X | NaN | NaN | NaN | NaN | 1.535818e+04 | 1.535818e+04 | NaN | NaN | GIGHFCQEMAK |
| PPP1R21 | PPR21\_HUMAN | Protein phosphatase 1 regulatory subunit 21 | 696 | 689 | PPP1R21 | PPR21\_HUMAN | Protein phosphatase 1 regulatory subunit 21 | 8C(125.0477) | NaN | X | NaN | NaN | NaN | NaN | 1.533843e+04 | 1.533843e+04 | NaN | NaN | SVHFYAECR |
| UBR4 | UBR4\_HUMAN | E3 ubiquitin-protein ligase UBR4 | 3430 | 3426 | UBR4 | UBR4\_HUMAN | E3 ubiquitin-protein ligase UBR4 | 5C(125.0477) | NaN | X | NaN | NaN | NaN | NaN | 1.532465e+04 | 1.532465e+04 | NaN | NaN | WQAHCLTLHIYR |
| PRMT5 | ANM5\_HUMAN | Protein arginine N-methyltransferase 5 | 22 | 19 | PRMT5 | ANM5\_HUMAN | Protein arginine N-methyltransferase 5 | 4C(125.0477) | NaN | X | NaN | NaN | NaN | NaN | 1.532347e+04 | 1.023460e+04 | NaN | 5088.8687 | DLNCVPEIADTLGAVAK |
| SEC31A | SC31A\_HUMAN | Protein transport protein Sec31A | 60 | 59 | SEC31A | SC31A\_HUMAN | Protein transport protein Sec31A | 2C(125.0477) | NaN | X | NaN | NaN | NaN | NaN | 1.528193e+04 | 1.528193e+04 | NaN | NaN | SCATFSSSHR |
| ALB | ALBU\_HUMAN | Albumin | 192 | 187 | ALB | ALBU\_HUMAN | Albumin | 6C(57.0215) | X | NaN | NaN | NaN | NaN | NaN | 1.525844e+04 | NaN | 1.525844e+04 | NaN | AAFTECCQAADK |
| CKAP5 | CKAP5\_HUMAN | Cytoskeleton-associated protein 5 | 343 | 343 | CKAP5 | CKAP5\_HUMAN | Cytoskeleton-associated protein 5 | 1C(125.0477) | NaN | X | NaN | NaN | NaN | NaN | 1.525231e+04 | 1.525231e+04 | NaN | NaN | CLTGLAVGLR |
| ABCC4 | MRP4\_HUMAN | ATP-binding cassette sub-family C member 4 | 1226 | 1223 | ABCC4 | MRP4\_HUMAN | ATP-binding cassette sub-family C member 4 | 4C(125.0477) | NaN | X | NaN | NaN | NaN | NaN | 1.516402e+04 | 1.516402e+04 | NaN | NaN | FAHCTVLTIAHR |
| DSP | DESP\_HUMAN | Desmoplakin | 682 | 677 | DSP | DESP\_HUMAN | Desmoplakin | 6C(125.0477) | NaN | X | NaN | NaN | NaN | NaN | 1.510518e+04 | 1.510518e+04 | NaN | NaN | RQIEHCEGR |
| APEX1 | APEX1\_HUMAN | DNA-(apurinic or apyrimidinic site) endonuclease | 296 | 282 | APEX1 | APEX1\_HUMAN | DNA-(apurinic or apyrimidinic site) endonuclease | 15C(125.0477) | NaN | X | NaN | NaN | NaN | NaN | 1.507696e+04 | 1.507696e+04 | NaN | NaN | LDYFLLSHSLLPALCDSK |
| PDZD8 | PDZD8\_HUMAN | PDZ domain-containing protein 8 | 743 | 736 | PDZD8 | PDZD8\_HUMAN | PDZ domain-containing protein 8 | 8C(125.0477) | NaN | X | NaN | NaN | NaN | NaN | 1.507547e+04 | 1.507547e+04 | NaN | NaN | LEDVALGCLATSNTEYLSK |
| LMAN2 | LMAN2\_HUMAN | Vesicular integral-membrane protein VIP36 | 239 | 238 | LMAN2 | LMAN2\_HUMAN | Vesicular integral-membrane protein VIP36 | 2C(31.9988) | NaN | NaN | NaN | X | NaN | NaN | 1.505869e+04 | NaN | NaN | 15058.6875 | NCIDITGVR |
| TUBA1B | TBA1B\_HUMAN | Tubulin alpha-1B chain | 347 | 340 | TUBA1B | TBA1B\_HUMAN | Tubulin alpha-1B chain | 8C(57.0215) | X | NaN | NaN | NaN | NaN | NaN | 1.505612e+04 | NaN | 1.505612e+04 | NaN | SIQFVDWCPTGFK |
| AP2B1 | AP2B1\_HUMAN | AP-2 complex subunit beta | 57 | 46 | AP2B1 | AP2B1\_HUMAN | AP-2 complex subunit beta | 12C(125.0477) | NaN | X | NaN | NaN | NaN | NaN | 1.501291e+04 | 1.501291e+04 | NaN | NaN | DVSSLFPDVVNCMQTDNLELKK |
| HSP90AA1 | HS90A\_HUMAN | Heat shock protein HSP 90-alpha | 420 | 420 | HSP90AA1 | HS90A\_HUMAN | Heat shock protein HSP 90-alpha | 1C(125.0477) | NaN | X | NaN | NaN | NaN | NaN | 1.499480e+04 | 1.499480e+04 | NaN | NaN | CLELFTELAEDK |
| MPP1 | EM55\_HUMAN | 55 kDa erythrocyte membrane protein | 454 | 446 | MPP1 | EM55\_HUMAN | 55 kDa erythrocyte membrane protein | 9C(125.0477) | NaN | X | NaN | NaN | NaN | NaN | 1.498567e+04 | 1.498567e+04 | NaN | NaN | LQEAFDQACSSPQWVPVSWVY |
| KEAP1 | KEAP1\_HUMAN | Kelch-like ECH-associated protein 1 | 434 | 416 | KEAP1 | KEAP1\_HUMAN | Kelch-like ECH-associated protein 1 | 19C(125.0477) | NaN | X | NaN | NaN | NaN | NaN | 1.498201e+04 | 1.498201e+04 | NaN | NaN | IGVGVIDGHIYAVGGSHGCIHHNSVER |
| STX2 | STX2\_HUMAN | Syntaxin-2 | 11 | 5 | STX2 | STX2\_HUMAN | Syntaxin-2 | 7C(125.0477) | NaN | X | NaN | NaN | NaN | NaN | 1.497579e+04 | NaN | 1.497579e+04 | NaN | LPDLTACR |
| MAP3K5 | M3K5\_HUMAN | Mitogen-activated protein kinase kinase kinase 5 | 120 | 118 | MAP3K5 | M3K5\_HUMAN | Mitogen-activated protein kinase kinase kinase 5 | 3C(125.0477) | NaN | X | NaN | NaN | NaN | NaN | 1.495600e+04 | 1.495600e+04 | NaN | NaN | EACETVGATLETLHFGK |
| SPTB | SPTB1\_HUMAN | Spectrin beta chain, erythrocytic | 603 | 594 | SPTB | SPTB1\_HUMAN | Spectrin beta chain, erythrocytic | 10C(125.0477) | NaN | X | NaN | NaN | NaN | NaN | 1.495095e+04 | NaN | 1.495095e+04 | NaN | FTEGKGYQPCDPQVIQDR |
| RANGAP1 | RAGP1\_HUMAN | Ran GTPase-activating protein 1 | 82 | 82 | RANGAP1 | RAGP1\_HUMAN | Ran GTPase-activating protein 1 | 1C(125.0477) | NaN | X | NaN | NaN | NaN | NaN | 1.492103e+04 | 1.492103e+04 | NaN | NaN | CHWSDMFTGR |
| HUWE1 | HUWE1\_HUMAN | E3 ubiquitin-protein ligase HUWE1 | 3259 | 3258 | HUWE1 | HUWE1\_HUMAN | E3 ubiquitin-protein ligase HUWE1 | 2C(125.0477) | NaN | X | NaN | NaN | NaN | NaN | 1.488813e+04 | 1.488813e+04 | NaN | NaN | SCGSSSHENRPLDLLHK |
| EIF3I | EIF3I\_HUMAN | Eukaryotic translation initiation factor 3 subunit I | 76 | 66 | EIF3I | EIF3I\_HUMAN | Eukaryotic translation initiation factor 3 subunit I | 11C(125.0477) | NaN | X | NaN | NaN | NaN | NaN | 1.485656e+04 | 1.485656e+04 | NaN | NaN | HVLTGSADNSCR |
| USP15 | UBP15\_HUMAN | Ubiquitin carboxyl-terminal hydrolase 15 | 791 | 788 | USP15 | UBP15\_HUMAN | Ubiquitin carboxyl-terminal hydrolase 15 | 4C(125.0477) | NaN | X | NaN | NaN | NaN | NaN | 1.482100e+04 | 1.482100e+04 | NaN | NaN | LKDCIELFTTKEK |
| CSRP1 | CSRP1\_HUMAN | Cysteine and glycine-rich protein 1 | 167 | 152 | CSRP1 | CSRP1\_HUMAN | Cysteine and glycine-rich protein 1 | 16C(125.0477) | NaN | X | NaN | NaN | NaN | NaN | 1.480128e+04 | 1.480128e+04 | NaN | NaN | GLESTTLADKDGEIYCK |
| MAT2B | MAT2B\_HUMAN | Methionine adenosyltransferase 2 subunit beta | 58 | 47 | MAT2B | MAT2B\_HUMAN | Methionine adenosyltransferase 2 subunit beta | 12C(125.0477) | NaN | X | NaN | NaN | NaN | NaN | 1.479527e+04 | 1.479527e+04 | NaN | NaN | EFQQNNWHAVGCGFR |
| QDPR | DHPR\_HUMAN | Dihydropteridine reductase | 85 | 74 | QDPR | DHPR\_HUMAN | Dihydropteridine reductase | 12C(125.0477) | NaN | X | NaN | NaN | NaN | NaN | 1.474613e+04 | 1.474613e+04 | NaN | NaN | LLGEEKVDAILCVAGGWAGGNAK |
| GSPT1 | ERF3A\_HUMAN | Eukaryotic peptide chain release factor GTP-binding subunit ERF3A | 237 | 222 | GSPT1 | ERF3A\_HUMAN | Eukaryotic peptide chain release factor GTP-binding subunit ERF3A | 16C(125.0477) | NaN | X | NaN | NaN | NaN | NaN | 1.473996e+04 | 1.473996e+04 | NaN | NaN | MDDPTVNWSNERYEECKEK |
| VIM | VIME\_HUMAN | Vimentin | 328 | 322 | VIM | VIME\_HUMAN | Vimentin | 7C(15.9949) | NaN | NaN | X | NaN | NaN | NaN | 1.472466e+04 | 1.472466e+04 | NaN | NaN | QVQSLTCEVDALK |
| SERPINA12 | SPA12\_HUMAN | Serpin A12 | 257 | 255 | SERPINA12 | SPA12\_HUMAN | Serpin A12 | 3C(125.0477) | NaN | X | NaN | NaN | NaN | NaN | 1.469081e+04 | 1.469081e+04 | NaN | NaN | LSCTILEIPYQK |
| KRT14 | K1C14\_HUMAN | Keratin, type I cytoskeletal 14 | 389 | 389 | KRT14 | K1C14\_HUMAN | Keratin, type I cytoskeletal 14 | 1C(15.9949) | NaN | NaN | X | NaN | NaN | NaN | 1.461454e+04 | 1.461454e+04 | NaN | NaN | CEMEQQNQEYK |
| RNF213 | RN213\_HUMAN | E3 ubiquitin-protein ligase RNF213 | 4937 | 4928 | RNF213 | RN213\_HUMAN | E3 ubiquitin-protein ligase RNF213 | 10C(125.0477) | NaN | X | NaN | NaN | NaN | NaN | 1.454262e+04 | 1.454262e+04 | NaN | NaN | DLTPLILSNCQYQVEEGR |
| UROD | DCUP\_HUMAN | Uroporphyrinogen decarboxylase | 66, 59 | 51 | UROD | DCUP\_HUMAN | Uroporphyrinogen decarboxylase | 16C(125.0477) 9C(15.9949) | NaN | X | X | NaN | NaN | NaN | 1.454217e+04 | 1.454217e+04 | NaN | NaN | AAQDFFSTCRSPEACCELTLQPLRR |
| EIF4G1 | IF4G1\_HUMAN | Eukaryotic translation initiation factor 4 gamma 1 | 662 | 657 | EIF4G1 | IF4G1\_HUMAN | Eukaryotic translation initiation factor 4 gamma 1 | 6C(125.0477) | NaN | X | NaN | NaN | NaN | NaN | 1.453851e+04 | 1.453851e+04 | NaN | NaN | LQGINCGPDFTPSFANLGR |
| IPO13 | IPO13\_HUMAN | Importin-13 | 838 | 836 | IPO13 | IPO13\_HUMAN | Importin-13 | 3C(125.0477) | NaN | X | NaN | NaN | NaN | NaN | 1.452249e+04 | 1.452249e+04 | NaN | NaN | ASCGFFTELLPR |
| CDC42 | CDC42\_HUMAN | Cell division control protein 42 homolog | 157 | 151 | CDC42 | CDC42\_HUMAN | Cell division control protein 42 homolog | 7C(125.0477) | NaN | X | NaN | NaN | NaN | NaN | 1.451798e+04 | 1.451798e+04 | NaN | NaN | AVKYVECSALTQK |
| KCMF1 | KCMF1\_HUMAN | E3 ubiquitin-protein ligase KCMF1 | 9 | 4 | KCMF1 | KCMF1\_HUMAN | E3 ubiquitin-protein ligase KCMF1 | 6C(31.9988) | NaN | NaN | NaN | X | NaN | NaN | 1.448478e+04 | 1.448478e+04 | NaN | NaN | HEGVSCDACLK |
| COPS3 | CSN3\_HUMAN | COP9 signalosome complex subunit 3 | 298 | 297 | COPS3 | CSN3\_HUMAN | COP9 signalosome complex subunit 3 | 2C(125.0477) | NaN | X | NaN | NaN | NaN | NaN | 1.448110e+04 | 1.448110e+04 | NaN | NaN | QCLSSLYKK |
| FKBP2 | FKBP2\_HUMAN | Peptidyl-prolyl cis-trans isomerase FKBP2 | 42 | 39 | FKBP2 | FKBP2\_HUMAN | Peptidyl-prolyl cis-trans isomerase FKBP2 | 4C(125.0477) | NaN | X | NaN | NaN | NaN | NaN | 1.445073e+04 | NaN | 1.445073e+04 | NaN | VDHCPIK |
| HMGXB3 | HMGX3\_HUMAN | HMG domain-containing protein 3 | 1255 | 1247 | HMGXB3 | HMGX3\_HUMAN | HMG domain-containing protein 3 | 9C(57.0215) | X | NaN | NaN | NaN | NaN | NaN | 1.442958e+04 | 1.442958e+04 | NaN | NaN | QIHDIVQSCQPGEVVIR |
| PPP1CA | PP1A\_HUMAN | Serine/threonine-protein phosphatase PP1-alpha catalytic subunit | 105 | 97 | PPP1CA | PP1A\_HUMAN | Serine/threonine-protein phosphatase PP1-alpha catalytic subunit | 9C(125.0477) | NaN | X | NaN | NaN | NaN | NaN | 1.442956e+04 | 1.442956e+04 | NaN | NaN | GKQSLETICLLLAYK |
| HSP90AB1 | HS90B\_HUMAN | Heat shock protein HSP 90-beta | 412 | 412 | HSP90AB1 | HS90B\_HUMAN | Heat shock protein HSP 90-beta | 1C(125.0477) | NaN | X | NaN | NaN | NaN | NaN | 1.441100e+04 | 1.441100e+04 | NaN | NaN | CLELFSELAEDK |
| SBF1 | MTMR5\_HUMAN | Myotubularin-related protein 5 | 987 | 986 | SBF1 | MTMR5\_HUMAN | Myotubularin-related protein 5 | 2C(125.0477) | NaN | X | NaN | NaN | NaN | NaN | 1.437680e+04 | 1.437680e+04 | NaN | NaN | SCTFQLLK |
| PTPA | PTPA\_HUMAN | Serine/threonine-protein phosphatase 2A activator | 291 | 278 | PTPA | PTPA\_HUMAN | Serine/threonine-protein phosphatase 2A activator | 14C(125.0477) | NaN | X | NaN | NaN | NaN | NaN | 1.432285e+04 | 1.432285e+04 | NaN | NaN | AVNENHKDYMFLECILFITEMK |
| QTRT2 | QTRT2\_HUMAN | Queuine tRNA-ribosyltransferase accessory subunit 2 | 174 | 167 | QTRT2 | QTRT2\_HUMAN | Queuine tRNA-ribosyltransferase accessory subunit 2 | 8C(125.0477) | NaN | X | NaN | NaN | NaN | NaN | 1.429920e+04 | 1.429920e+04 | NaN | NaN | SLLFLDNCLR |
| OSBP2 | OSBP2\_HUMAN | Oxysterol-binding protein 2 | 621 | 619 | OSBP2 | OSBP2\_HUMAN | Oxysterol-binding protein 2 | 3C(125.0477) | NaN | X | NaN | NaN | NaN | NaN | 1.428067e+04 | 1.428067e+04 | NaN | NaN | SLCEQVSHHPPSAAHYVFSK |
| RANBP3 | RANB3\_HUMAN | Ran-binding protein 3 | 249 | 238 | RANBP3 | RANB3\_HUMAN | Ran-binding protein 3 | 12C(125.0477) | NaN | X | NaN | NaN | NaN | NaN | 1.427716e+04 | 1.427716e+04 | NaN | NaN | NESSNASEEEACEK |
| BCAP29 | BAP29\_HUMAN | B-cell receptor-associated protein 29 | 171 | 164 | BCAP29 | BAP29\_HUMAN | B-cell receptor-associated protein 29 | 8C(125.0477) | NaN | X | NaN | NaN | NaN | NaN | 1.422520e+04 | 1.422520e+04 | NaN | NaN | SHGKDEECVLEAENKK |
| TEPSIN | AP4AT\_HUMAN | AP-4 complex accessory subunit Tepsin | 243 | 235 | TEPSIN | AP4AT\_HUMAN | AP-4 complex accessory subunit Tepsin | 9C(125.0477) | NaN | X | NaN | NaN | NaN | NaN | 1.415552e+04 | 1.415552e+04 | NaN | NaN | VEVVALSDCQQELSLVR |
| KRT78 | K2C78\_HUMAN | Keratin, type II cytoskeletal 78 | 394 | 392 | KRT78 | K2C78\_HUMAN | Keratin, type II cytoskeletal 78 | 3C(125.0477) | NaN | X | NaN | NaN | NaN | NaN | 1.412088e+04 | NaN | 1.412088e+04 | NaN | LLCEYQELTSTK |
| C4A | CO4A\_HUMAN | Complement C4-A | 567 | 560 | C4A | CO4A\_HUMAN | Complement C4-A | 8C(57.0215) | X | NaN | NaN | NaN | NaN | NaN | 1.411593e+04 | NaN | 1.411593e+04 | NaN | VDVQAGACEGK |
| GSTM2 | GSTM2\_HUMAN | Glutathione S-transferase Mu 2 | 87 | 83 | GSTM2 | GSTM2\_HUMAN | Glutathione S-transferase Mu 2 | 5C(125.0477) | NaN | X | NaN | NaN | NaN | NaN | 1.410396e+04 | 1.410396e+04 | NaN | NaN | KHNLCGESEKEQIR |
| EIF5 | IF5\_HUMAN | Eukaryotic translation initiation factor 5 | 122 | 115 | EIF5 | IF5\_HUMAN | Eukaryotic translation initiation factor 5 | 8C(125.0477) | NaN | X | NaN | NaN | NaN | NaN | 1.409408e+04 | 1.409408e+04 | NaN | NaN | KQTIGNSCK |
| ACTB | ACTB\_HUMAN | Actin, cytoplasmic 1 | 217, 217 | 216 | ACTB | ACTB\_HUMAN | Actin, cytoplasmic 1 | 2C(31.9988) 2C(57.0215) | X | NaN | NaN | X | NaN | NaN | 1.408233e+04 | NaN | 1.408233e+04 | NaN | LCYVALDFEQEMATAASSSSLEK |
| SERPINA6 | CBG\_HUMAN | Corticosteroid-binding globulin | 82 | 61 | SERPINA6 | CBG\_HUMAN | Corticosteroid-binding globulin | 22C(57.0215) | X | NaN | NaN | NaN | NaN | NaN | 1.406572e+04 | 1.406572e+04 | NaN | NaN | KNIFISPVSISMALAMLSLGTCGHTR |
| CHERP | CHERP\_HUMAN | Calcium homeostasis endoplasmic reticulum protein | 246 | 240 | CHERP | CHERP\_HUMAN | Calcium homeostasis endoplasmic reticulum protein | 7C(125.0477) | NaN | X | NaN | NaN | NaN | NaN | 1.396326e+04 | 1.396326e+04 | NaN | NaN | VVVPIYCTSFLAVEEDKQQK |
| WDR81 | WDR81\_HUMAN | WD repeat-containing protein 81 | 1226 | 1220 | WDR81 | WDR81\_HUMAN | WD repeat-containing protein 81 | 7C(125.0477) | NaN | X | NaN | NaN | NaN | NaN | 1.395865e+04 | NaN | 1.395865e+04 | NaN | ILLDTACK |
| LTA4H | LKHA4\_HUMAN | Leukotriene A-4 hydrolase | 200 | 196 | LTA4H | LKHA4\_HUMAN | Leukotriene A-4 hydrolase | 5C(125.0477) | NaN | X | NaN | NaN | NaN | NaN | 1.395223e+04 | 1.395223e+04 | NaN | NaN | VPIPCYLIALVVGALESR |
| KPRP | KPRP\_HUMAN | Keratinocyte proline-rich protein | 343 | 343 | KPRP | KPRP\_HUMAN | Keratinocyte proline-rich protein | 1C(125.0477) | NaN | X | NaN | NaN | NaN | NaN | 1.391901e+04 | NaN | 1.391901e+04 | NaN | CPVEIPPIR |
| RANGAP1 | RAGP1\_HUMAN | Ran GTPase-activating protein 1 | 274 | 265 | RANGAP1 | RAGP1\_HUMAN | Ran GTPase-activating protein 1 | 10C(125.0477) | NaN | X | NaN | NaN | NaN | NaN | 1.390679e+04 | 1.390679e+04 | NaN | NaN | QVEVINFGDCLVR |
| WIPI2 | WIPI2\_HUMAN | WD repeat domain phosphoinositide-interacting protein 2 | 430 | 417 | WIPI2 | WIPI2\_HUMAN | WD repeat domain phosphoinositide-interacting protein 2 | 14C(125.0477) | NaN | X | NaN | NaN | NaN | NaN | 1.383394e+04 | 1.383394e+04 | NaN | NaN | LAYTDDLGAVGGACLEDEASALR |
| IGHA1 | IGHA1\_HUMAN | Immunoglobulin heavy constant alpha 1 | 204 | 201 | IGHA1 | IGHA1\_HUMAN | Immunoglobulin heavy constant alpha 1 | 4C(57.0215) | X | NaN | NaN | NaN | NaN | NaN | 1.380704e+04 | NaN | NaN | 13807.0440 | TFTCTAAYPESK |
| GGA3 | GGA3\_HUMAN | ADP-ribosylation factor-binding protein GGA3 | 255 | 250 | GGA3 | GGA3\_HUMAN | ADP-ribosylation factor-binding protein GGA3 | 6C(31.9988) | NaN | NaN | NaN | X | NaN | NaN | 1.377919e+04 | NaN | NaN | 13779.1920 | ELFDQCENK |
| CAPN1 | CAN1\_HUMAN | Calpain-1 catalytic subunit | 417 | 414 | CAPN1 | CAN1\_HUMAN | Calpain-1 catalytic subunit | 4C(125.0477) | NaN | X | NaN | NaN | NaN | NaN | 1.376612e+04 | 1.376612e+04 | NaN | NaN | ESGCSFVLALMQK |
| GTF3C5 | TF3C5\_HUMAN | General transcription factor 3C polypeptide 5 | 432 | 425 | GTF3C5 | TF3C5\_HUMAN | General transcription factor 3C polypeptide 5 | 8C(125.0477) | NaN | X | NaN | NaN | NaN | NaN | 1.374751e+04 | 1.374751e+04 | NaN | NaN | NDGAENSCTER |
| CKM | KCRM\_HUMAN | Creatine kinase M-type | 254 | 253 | CKM | KCRM\_HUMAN | Creatine kinase M-type | 2C(125.0477) | NaN | X | NaN | NaN | NaN | NaN | 1.372946e+04 | 1.372946e+04 | NaN | NaN | FCVGLQK |
| PSMC5 | PRS8\_HUMAN | 26S proteasome regulatory subunit 8 | 112 | 102 | PSMC5 | PRS8\_HUMAN | 26S proteasome regulatory subunit 8 | 11C(125.0477) | NaN | X | NaN | NaN | NaN | NaN | 1.366808e+04 | NaN | 1.366808e+04 | NaN | NIDINDVTPNCR |
| NAXD | NNRD\_HUMAN | ATP-dependent (S)-NAD(P)H-hydrate dehydratase | 261 | 250 | NAXD | NNRD\_HUMAN | ATP-dependent (S)-NAD(P)H-hydrate dehydratase | 12C(125.0477) | NaN | X | NaN | NaN | NaN | NaN | 1.366282e+04 | 1.366282e+04 | NaN | NaN | DILSNGQQVLVCSQEGSSR |
| TF | TRFE\_HUMAN | Serotransferrin | 260 | 252 | TF | TRFE\_HUMAN | Serotransferrin | 9C(57.0215) | X | NaN | NaN | NaN | NaN | NaN | 1.363507e+04 | 1.363507e+04 | NaN | NaN | KPVDEYKDCHLAQVPSHTVVAR |
| NF2 | MERL\_HUMAN | Merlin | 133 | 131 | NF2 | MERL\_HUMAN | Merlin | 3C(125.0477) | NaN | X | NaN | NaN | NaN | NaN | 1.363359e+04 | 1.363359e+04 | NaN | NaN | IYCPPEASVLLASYAVQAK |
| PEPD | PEPD\_HUMAN | Xaa-Pro dipeptidase | 230 | 219 | PEPD | PEPD\_HUMAN | Xaa-Pro dipeptidase | 12C(125.0477) | NaN | X | NaN | NaN | NaN | NaN | 1.358562e+04 | 1.358562e+04 | NaN | NaN | EYELESLFEHYCYSR |
| TF | TRFE\_HUMAN | Serotransferrin | 246 | 240 | TF | TRFE\_HUMAN | Serotransferrin | 7C(57.0215) | X | NaN | NaN | NaN | NaN | NaN | 1.354168e+04 | NaN | 1.354168e+04 | NaN | DQYELLCLDNTR |
| IQGAP2 | IQGA2\_HUMAN | Ras GTPase-activating-like protein IQGAP2 | 575 | 563 | IQGAP2 | IQGA2\_HUMAN | Ras GTPase-activating-like protein IQGAP2 | 13C(125.0477) | NaN | X | NaN | NaN | NaN | NaN | 1.351967e+04 | NaN | 1.351967e+04 | NaN | SSTSNANDIIPECADK |
| NUP93 | NUP93\_HUMAN | Nuclear pore complex protein Nup93 | 422 | 418 | NUP93 | NUP93\_HUMAN | Nuclear pore complex protein Nup93 | 5C(125.0477) | NaN | X | NaN | NaN | NaN | NaN | 1.350704e+04 | 1.350704e+04 | NaN | NaN | LNQVCFDDDGTSSPQDR |
| STUB1 | CHIP\_HUMAN | E3 ubiquitin-protein ligase CHIP | 232 | 226 | STUB1 | CHIP\_HUMAN | E3 ubiquitin-protein ligase CHIP | 7C(125.0477) | NaN | X | NaN | NaN | NaN | NaN | 1.346541e+04 | NaN | 1.346541e+04 | NaN | DIPDYLCGK |
| HACE1 | HACE1\_HUMAN | E3 ubiquitin-protein ligase HACE1 | 481 | 476 | HACE1 | HACE1\_HUMAN | E3 ubiquitin-protein ligase HACE1 | 6C(125.0477) | NaN | X | NaN | NaN | NaN | NaN | 1.340408e+04 | 1.340408e+04 | NaN | NaN | FIEFVCKHDEVLK |
| DIP2B | DIP2B\_HUMAN | Disco-interacting protein 2 homolog B | 895 | 881 | DIP2B | DIP2B\_HUMAN | Disco-interacting protein 2 homolog B | 15C(125.0477) | NaN | X | NaN | NaN | NaN | NaN | 1.338483e+04 | 1.338483e+04 | NaN | NaN | VLQAIDSIHQVGVYCLALVPANTLPK |
| PPIP5K2 | VIP2\_HUMAN | Inositol hexakisphosphate and diphosphoinositol-pentakisphosphate kinase 2 | 900 | 899 | PPIP5K2 | VIP2\_HUMAN | Inositol hexakisphosphate and diphosphoinositol-pentakisphosphate kinase 2 | 2C(125.0477) | NaN | X | NaN | NaN | NaN | NaN | 1.335425e+04 | 1.335425e+04 | NaN | NaN | GCEEDKNLPSGYGYRPASR |
| HBD | HBD\_HUMAN | Hemoglobin subunit delta | 94, 94 | 68 | HBD | HBD\_HUMAN | Hemoglobin subunit delta | 27C(125.0477) 27C(15.9949) | NaN | X | X | NaN | NaN | NaN | 1.331762e+04 | 1.331762e+04 | NaN | NaN | VLGAFSDGLAHLDNLKGTFSQLSELHCDKLHVDPENFR |
| MYH9 | MYH9\_HUMAN | Myosin-9 | 91 | 83 | MYH9 | MYH9\_HUMAN | Myosin-9 | 9C(125.0477) | NaN | X | NaN | NaN | NaN | NaN | 1.323004e+04 | 1.323004e+04 | NaN | NaN | VEDMAELTCLNEASVLHNLKER |
| HPRT1 | HPRT\_HUMAN | Hypoxanthine-guanine phosphoribosyltransferase | 206 | 201 | HPRT1 | HPRT\_HUMAN | Hypoxanthine-guanine phosphoribosyltransferase | 6C(125.0477) | NaN | X | NaN | NaN | NaN | NaN | 1.319087e+04 | 1.319087e+04 | NaN | NaN | DLNHVCVISETGKAK |
| HAGH | GLO2\_HUMAN | Hydroxyacylglutathione hydrolase, mitochondrial | 34 | 24 | HAGH | GLO2\_HUMAN | Hydroxyacylglutathione hydrolase, mitochondrial | 11C(125.0477) | NaN | X | NaN | NaN | NaN | NaN | 1.317159e+04 | 1.317159e+04 | NaN | NaN | GLGPALLGVFCHTDLR |
| MTAP | MTAP\_HUMAN | S-methyl-5'-thioadenosine phosphorylase | 55 | 52 | MTAP | MTAP\_HUMAN | S-methyl-5'-thioadenosine phosphorylase | 4C(125.0477) | NaN | X | NaN | NaN | NaN | NaN | 1.315366e+04 | NaN | 1.315366e+04 | NaN | NVDCVLLAR |
| SERPINA1 | A1AT\_HUMAN | Alpha-1-antitrypsin | 256 | 248 | SERPINA1 | A1AT\_HUMAN | Alpha-1-antitrypsin | 9C(57.0215) | X | NaN | NaN | NaN | NaN | NaN | 1.315025e+04 | 1.315025e+04 | NaN | NaN | LGMFNIQHCK |
| FBXO4 | FBX4\_HUMAN | F-box only protein 4 | 283 | 282 | FBXO4 | FBX4\_HUMAN | F-box only protein 4 | 2C(125.0477) | NaN | X | NaN | NaN | NaN | NaN | 1.312104e+04 | 1.312104e+04 | NaN | NaN | VCEVVDGFIYVANAEAHK |
| NFKB1 | NFKB1\_HUMAN | Nuclear factor NF-kappa-B p105 subunit | 61 | 59 | NFKB1 | NFKB1\_HUMAN | Nuclear factor NF-kappa-B p105 subunit | 3C(125.0477) | NaN | X | NaN | NaN | NaN | NaN | 1.310223e+04 | 1.310223e+04 | NaN | NaN | YVCEGPSHGGLPGASSEK |
| ITGB1 | ITB1\_HUMAN | Integrin beta-1 | 64 | 64 | ITGB1 | ITB1\_HUMAN | Integrin beta-1 | 1C(125.0477) | NaN | X | NaN | NaN | NaN | NaN | 1.310133e+04 | NaN | 1.310133e+04 | NaN | CDDLEALK |
| ETF1 | ERF1\_HUMAN | Eukaryotic peptide chain release factor subunit 1 | 127 | 110 | ETF1 | ERF1\_HUMAN | Eukaryotic peptide chain release factor subunit 1 | 18C(125.0477) | NaN | X | NaN | NaN | NaN | NaN | 1.308831e+04 | 1.308831e+04 | NaN | NaN | VNIDFEPFKPINTSLYLCDNK |
| ADK | ADK\_HUMAN | Adenosine kinase | 182 | 181 | ADK | ADK\_HUMAN | Adenosine kinase | 2C(125.0477) | NaN | X | NaN | NaN | NaN | NaN | 1.306544e+04 | 1.306544e+04 | NaN | NaN | VCYIAGFFLTVSPESVLK |
| ANKRD28 | ANR28\_HUMAN | Serine/threonine-protein phosphatase 6 regulatory ankyrin repeat subunit A | 472 | 471 | ANKRD28 | ANR28\_HUMAN | Serine/threonine-protein phosphatase 6 regulatory ankyrin repeat subunit A | 2C(125.0477) | NaN | X | NaN | NaN | NaN | NaN | 1.306073e+04 | 1.306073e+04 | NaN | NaN | GCTPLHYAATSDTDGK |
| ACSL6 | ACSL6\_HUMAN | Long-chain-fatty-acid--CoA ligase 6 | 142 | 130 | ACSL6 | ACSL6\_HUMAN | Long-chain-fatty-acid--CoA ligase 6 | 13C(125.0477) | NaN | X | NaN | NaN | NaN | NaN | 1.305464e+04 | 1.305464e+04 | NaN | NaN | AEFLGSGLLQHNCK |
| RAB21 | RAB21\_HUMAN | Ras-related protein Rab-21 | 177 | 168 | RAB21 | RAB21\_HUMAN | Ras-related protein Rab-21 | 10C(125.0477) | NaN | X | NaN | NaN | NaN | NaN | 1.298313e+04 | 1.298313e+04 | NaN | NaN | GIEELFLDLCKR |
| USP24 | UBP24\_HUMAN | Ubiquitin carboxyl-terminal hydrolase 24 | 1333 | 1317 | USP24 | UBP24\_HUMAN | Ubiquitin carboxyl-terminal hydrolase 24 | 17C(125.0477) | NaN | X | NaN | NaN | NaN | NaN | 1.293721e+04 | 1.293721e+04 | NaN | NaN | VAIQTMEVSDFTSTVACFMR |
| ALDOA | ALDOA\_HUMAN | Fructose-bisphosphate aldolase A | 178 | 174 | ALDOA | ALDOA\_HUMAN | Fructose-bisphosphate aldolase A | 5C(57.0215) | X | NaN | NaN | NaN | NaN | NaN | 1.293295e+04 | NaN | 5.931594e+03 | 7001.3570 | YASICQQNGIVPIVEPEILPDGDHDLK |
| CAD | PYR1\_HUMAN | CAD protein | 1613 | 1609 | CAD | PYR1\_HUMAN | CAD protein | 5C(125.0477) | NaN | X | NaN | NaN | NaN | NaN | 1.292606e+04 | 1.292606e+04 | NaN | NaN | SVHICHVAR |
| GNAI3 | GNAI3\_HUMAN | Guanine nucleotide-binding protein G(i) subunit alpha-3 | 254 | 249 | GNAI3 | GNAI3\_HUMAN | Guanine nucleotide-binding protein G(i) subunit alpha-3 | 6C(125.0477) | NaN | X | NaN | NaN | NaN | NaN | 1.290056e+04 | 1.290056e+04 | NaN | NaN | LFDSICNNKWFTETSIILFLNKK |
| TALDO1 | TALDO\_HUMAN | Transaldolase | 250 | 246 | TALDO1 | TALDO\_HUMAN | Transaldolase | 5C(125.0477) | NaN | X | NaN | NaN | NaN | NaN | 1.288955e+04 | NaN | 1.288955e+04 | NaN | ALAGCDFLTISPK |
| GUCD1 | GUCD1\_HUMAN | Protein GUCD1 | 85 | 84 | GUCD1 | GUCD1\_HUMAN | Protein GUCD1 | 2C(31.9988) | NaN | NaN | NaN | X | NaN | NaN | 1.287118e+04 | NaN | NaN | 12871.1840 | FCTQTLGVDK |
| FYCO1 | FYCO1\_HUMAN | FYVE and coiled-coil domain-containing protein 1 | 737 | 732 | FYCO1 | FYCO1\_HUMAN | FYVE and coiled-coil domain-containing protein 1 | 6C(125.0477) | NaN | X | NaN | NaN | NaN | NaN | 1.279863e+04 | 1.279863e+04 | NaN | NaN | ALESQCQQQTQLIEVLTAEK |
| DRG2 | DRG2\_HUMAN | Developmentally-regulated GTP-binding protein 2 | 319 | 311 | DRG2 | DRG2\_HUMAN | Developmentally-regulated GTP-binding protein 2 | 9C(125.0477) | NaN | X | NaN | NaN | NaN | NaN | 1.275184e+04 | 1.275184e+04 | NaN | NaN | KGASVEHVCHR |
| MBLAC2 | MBLC2\_HUMAN | Acyl-coenzyme A thioesterase MBLAC2 | 176 | 173 | MBLAC2 | MBLC2\_HUMAN | Acyl-coenzyme A thioesterase MBLAC2 | 4C(125.0477) | NaN | X | NaN | NaN | NaN | NaN | 1.274690e+04 | 1.274690e+04 | NaN | NaN | GSICLHDKDRK |
| USP15 | UBP15\_HUMAN | Ubiquitin carboxyl-terminal hydrolase 15 | 139 | 138 | USP15 | UBP15\_HUMAN | Ubiquitin carboxyl-terminal hydrolase 15 | 2C(125.0477) | NaN | X | NaN | NaN | NaN | NaN | 1.269940e+04 | NaN | 1.269940e+04 | NaN | LCENGNMNNVVTR |
| CLTC | CLH1\_HUMAN | Clathrin heavy chain 1 | 870 | 866 | CLTC | CLH1\_HUMAN | Clathrin heavy chain 1 | 5C(125.0477) | NaN | X | NaN | NaN | NaN | NaN | 1.264940e+04 | NaN | 1.264940e+04 | NaN | IHEGCEEPATHNALAK |
| DOCK11 | DOC11\_HUMAN | Dedicator of cytokinesis protein 11 | 667 | 663 | DOCK11 | DOC11\_HUMAN | Dedicator of cytokinesis protein 11 | 5C(-34.0809) | NaN | NaN | NaN | NaN | X | NaN | 1.264652e+04 | NaN | NaN | 12646.5160 | NIAVCVEFR |
| UBAC1 | UBAC1\_HUMAN | Ubiquitin-associated domain-containing protein 1 | 19 | 16 | UBAC1 | UBAC1\_HUMAN | Ubiquitin-associated domain-containing protein 1 | 4C(125.0477) | NaN | X | NaN | NaN | NaN | NaN | 1.264392e+04 | 1.264392e+04 | NaN | NaN | LHICASDGAEWLEEATEDTSVEKLKER |
| UBE2O | UBE2O\_HUMAN | (E3-independent) E2 ubiquitin-conjugating enzyme | 1040 | 1029 | UBE2O | UBE2O\_HUMAN | (E3-independent) E2 ubiquitin-conjugating enzyme | 12C(125.0477) | NaN | X | NaN | NaN | NaN | NaN | 1.259353e+04 | 1.259353e+04 | NaN | NaN | LNPNLYDNGKVCVSLLGTWIGKGTER |
| GBF1 | GBF1\_HUMAN | Golgi-specific brefeldin A-resistance guanine nucleotide exchange factor 1 | 1430 | 1420 | GBF1 | GBF1\_HUMAN | Golgi-specific brefeldin A-resistance guanine nucleotide exchange factor 1 | 11C(31.9988) | NaN | NaN | NaN | X | NaN | NaN | 1.254694e+04 | NaN | NaN | 12546.9400 | IFVEASLNGGCK |
| ATP1A1 | AT1A1\_HUMAN | Sodium/potassium-transporting ATPase subunit alpha-1 | 374 | 360 | ATP1A1 | AT1A1\_HUMAN | Sodium/potassium-transporting ATPase subunit alpha-1 | 15C(125.0477) | NaN | X | NaN | NaN | NaN | NaN | 1.250454e+04 | NaN | 1.250454e+04 | NaN | NLEAVETLGSTSTICSDK |
| HECTD3 | HECD3\_HUMAN | E3 ubiquitin-protein ligase HECTD3 | 112 | 106 | HECTD3 | HECD3\_HUMAN | E3 ubiquitin-protein ligase HECTD3 | 7C(125.0477) | NaN | X | NaN | NaN | NaN | NaN | 1.248993e+04 | 1.248993e+04 | NaN | NaN | TTGEELCNGHGLWVK |
| DIS3L2 | DI3L2\_HUMAN | DIS3-like exonuclease 2 | 216 | 212 | DIS3L2 | DI3L2\_HUMAN | DIS3-like exonuclease 2 | 5C(125.0477) | NaN | X | NaN | NaN | NaN | NaN | 1.239388e+04 | 1.239388e+04 | NaN | NaN | DETTCISQDTR |
| TBC1D13 | TBC13\_HUMAN | TBC1 domain family member 13 | 145 | 140 | TBC1D13 | TBC13\_HUMAN | TBC1 domain family member 13 | 6C(125.0477) | NaN | X | NaN | NaN | NaN | NaN | 1.228776e+04 | 1.228776e+04 | NaN | NaN | ATDYPCLLILDPQNEFETLR |
| RPLP0 | RLA0\_HUMAN | 60S acidic ribosomal protein P0 | 119 | 113 | RPLP0 | RLA0\_HUMAN | 60S acidic ribosomal protein P0 | 7C(125.0477) | NaN | X | NaN | NaN | NaN | NaN | 1.227381e+04 | 1.227381e+04 | NaN | NaN | AGAIAPCEVTVPAQNTGLGPEK |
| HUWE1 | HUWE1\_HUMAN | E3 ubiquitin-protein ligase HUWE1 | 2721 | 2713 | HUWE1 | HUWE1\_HUMAN | E3 ubiquitin-protein ligase HUWE1 | 9C(125.0477) | NaN | X | NaN | NaN | NaN | NaN | 1.221089e+04 | 1.221089e+04 | NaN | NaN | ENRDQSAQCTASK |
| ARL15 | ARL15\_HUMAN | ADP-ribosylation factor-like protein 15 | 133 | 121 | ARL15 | ARL15\_HUMAN | ADP-ribosylation factor-like protein 15 | 13C(125.0477) | NaN | X | NaN | NaN | NaN | NaN | 1.216067e+04 | 1.216067e+04 | NaN | NaN | NELHSALQHPQLCTLPFLILANHQDKPAAR |
| RPS3A | RS3A\_HUMAN | 40S ribosomal protein S3a | 201 | 200 | RPS3A | RS3A\_HUMAN | 40S ribosomal protein S3a | 2C(125.0477) | NaN | X | NaN | NaN | NaN | NaN | 1.204396e+04 | 1.204396e+04 | NaN | NaN | ACQSIYPLHDVFVR |
| SLC12A7 | S12A7\_HUMAN | Solute carrier family 12 member 7 | 323 | 319 | SLC12A7 | S12A7\_HUMAN | Solute carrier family 12 member 7 | 5C(125.0477) | NaN | X | NaN | NaN | NaN | NaN | 1.202212e+04 | NaN | 1.202212e+04 | NaN | SFDACVK |
| VASP | VASP\_HUMAN | Vasodilator-stimulated phosphoprotein | 334 | 322 | VASP | VASP\_HUMAN | Vasodilator-stimulated phosphoprotein | 13C(125.0477) | NaN | X | NaN | NaN | NaN | NaN | 1.191419e+04 | 1.191419e+04 | NaN | NaN | SSSSVTTSETQPCTPSSSDYSDLQR |
| PANK2 | PANK2\_HUMAN | Pantothenate kinase 2, mitochondrial | 276 | 265 | PANK2 | PANK2\_HUMAN | Pantothenate kinase 2, mitochondrial | 12C(125.0477) | NaN | X | NaN | NaN | NaN | NaN | 1.188370e+04 | 1.188370e+04 | NaN | NaN | DVHLELKDLTLCGR |
| AP2A1 | AP2A1\_HUMAN | AP-2 complex subunit alpha-1 | 171 | 154 | AP2A1 | AP2A1\_HUMAN | AP-2 complex subunit alpha-1 | 18C(125.0477) | NaN | X | NaN | NaN | NaN | NaN | 1.186015e+04 | 1.186015e+04 | NaN | NaN | ILVAGDSMDSVKQSAALCLLRLYK |
| EIF2S3 | IF2G\_HUMAN | Eukaryotic translation initiation factor 2 subunit 3 | 434 | 427 | EIF2S3 | IF2G\_HUMAN | Eukaryotic translation initiation factor 2 subunit 3 | 8C(125.0477) | NaN | X | NaN | NaN | NaN | NaN | 1.184842e+04 | NaN | 1.184842e+04 | NaN | IVLTNPVCTEVGEK |
| MAPRE1 | MARE1\_HUMAN | Microtubule-associated protein RP/EB family member 1 | 228 | 223 | MAPRE1 | MARE1\_HUMAN | Microtubule-associated protein RP/EB family member 1 | 6C(125.0477) | NaN | X | NaN | NaN | NaN | NaN | 1.182886e+04 | NaN | 1.182886e+04 | NaN | NIELICQENEGENDPVLQR |
| RNF123 | RN123\_HUMAN | E3 ubiquitin-protein ligase RNF123 | 931 | 913 | RNF123 | RN123\_HUMAN | E3 ubiquitin-protein ligase RNF123 | 19C(125.0477) | NaN | X | NaN | NaN | NaN | NaN | 1.181051e+04 | 1.181051e+04 | NaN | NaN | IVGTDIRDSLMQALASYVCYPHSLR |
| RHAG | RHAG\_HUMAN | Ammonium transporter Rh type A | 250 | 239 | RHAG | RHAG\_HUMAN | Ammonium transporter Rh type A | 12C(125.0477) | NaN | X | NaN | NaN | NaN | NaN | 1.179069e+04 | 1.179069e+04 | NaN | NaN | AIVNTYFSLAACVLTAFAFSSLVEHR |
| PRKCI | KPCI\_HUMAN | Protein kinase C iota type | 137 | 135 | PRKCI | KPCI\_HUMAN | Protein kinase C iota type | 3C(125.0477) | NaN | X | NaN | NaN | NaN | NaN | 1.174483e+04 | 1.174483e+04 | NaN | NaN | LYCANGHTFQAK |
| ACLY | ACLY\_HUMAN | ATP-citrate synthase | 20 | 18 | ACLY | ACLY\_HUMAN | ATP-citrate synthase | 3C(125.0477) | NaN | X | NaN | NaN | NaN | NaN | 1.172921e+04 | NaN | 1.172921e+04 | NaN | FICTTSAIQNR |
| PNPLA6 | PLPL6\_HUMAN | Patatin-like phospholipase domain-containing protein 6 | 928 | 926 | PNPLA6 | PLPL6\_HUMAN | Patatin-like phospholipase domain-containing protein 6 | 3C(125.0477) | NaN | X | NaN | NaN | NaN | NaN | 1.170728e+04 | 1.170728e+04 | NaN | NaN | SWCSGHLHLR |
| IRF3 | IRF3\_HUMAN | Interferon regulatory factor 3 | 267 | 263 | IRF3 | IRF3\_HUMAN | Interferon regulatory factor 3 | 5C(125.0477) | NaN | X | NaN | NaN | NaN | NaN | 1.166308e+04 | 1.166308e+04 | NaN | NaN | HVLSCLGGGLALWR |
| EEF2 | EF2\_HUMAN | Elongation factor 2 | 41 | 33 | EEF2 | EF2\_HUMAN | Elongation factor 2 | 9C(125.0477) | NaN | X | NaN | NaN | NaN | NaN | 1.162987e+04 | NaN | 1.162987e+04 | NaN | STLTDSLVCK |
| KARS1 | SYK\_HUMAN | Lysine--tRNA ligase | 427 | 422 | KARS1 | SYK\_HUMAN | Lysine--tRNA ligase | 6C(31.9988) | NaN | NaN | NaN | X | NaN | NaN | 1.157244e+04 | NaN | NaN | 11572.4380 | ILDDICVAK |
| NRDC | NRDC\_HUMAN | Nardilysin | 269 | 258 | NRDC | NRDC\_HUMAN | Nardilysin | 12C(125.0477) | NaN | X | NaN | NaN | NaN | NaN | 1.156599e+04 | 1.156599e+04 | NaN | NaN | KHGGSDNASTDCER |
| EHD1 | EHD1\_HUMAN | EH domain-containing protein 1 | 138 | 136 | EHD1 | EHD1\_HUMAN | EH domain-containing protein 1 | 3C(125.0477) | NaN | X | NaN | NaN | NaN | NaN | 1.150001e+04 | 1.150001e+04 | NaN | NaN | FMCAQLPNPVLDSISIIDTPGILSGEK |
| KIF5B | KINH\_HUMAN | Kinesin-1 heavy chain | 421 | 419 | KIF5B | KINH\_HUMAN | Kinesin-1 heavy chain | 3C(125.0477) | NaN | X | NaN | NaN | NaN | NaN | 1.148033e+04 | 1.148033e+04 | NaN | NaN | RKCEEEIAK |
| VPS28 | VPS28\_HUMAN | Vacuolar protein sorting-associated protein 28 homolog | 96 | 83 | VPS28 | VPS28\_HUMAN | Vacuolar protein sorting-associated protein 28 homolog | 14C(125.0477) | NaN | X | NaN | NaN | NaN | NaN | 1.147649e+04 | 1.147649e+04 | NaN | NaN | QVQGSEISSIDEFCRK |
| CYFIP1 | CYFP1\_HUMAN | Cytoplasmic FMR1-interacting protein 1 | 1180 | 1174 | CYFIP1 | CYFP1\_HUMAN | Cytoplasmic FMR1-interacting protein 1 | 7C(125.0477) | NaN | X | NaN | NaN | NaN | NaN | 1.142836e+04 | 1.142836e+04 | NaN | NaN | FAVLDFCYHLLK |
| ARCN1 | COPD\_HUMAN | Coatomer subunit delta | 479 | 476 | ARCN1 | COPD\_HUMAN | Coatomer subunit delta | 4C(125.0477) | NaN | X | NaN | NaN | NaN | NaN | 1.141510e+04 | 1.141510e+04 | NaN | NaN | KNYCNIQVTK |
| INPP5D | SHIP1\_HUMAN | Phosphatidylinositol 3,4,5-trisphosphate 5-phosphatase 1 | 217 | 212 | INPP5D | SHIP1\_HUMAN | Phosphatidylinositol 3,4,5-trisphosphate 5-phosphatase 1 | 6C(125.0477) | NaN | X | NaN | NaN | NaN | NaN | 1.138735e+04 | NaN | 1.138735e+04 | NaN | LTTLLCK |
| PPIA | PPIA\_HUMAN | Peptidyl-prolyl cis-trans isomerase A | 62 | 56 | PPIA | PPIA\_HUMAN | Peptidyl-prolyl cis-trans isomerase A | 7C(57.0215) | X | NaN | NaN | NaN | NaN | NaN | 1.131725e+04 | NaN | NaN | 11317.2470 | IIPGFMCQGGDFTR |
| CARS1 | SYCC\_HUMAN | Cysteine--tRNA ligase, cytoplasmic | 182 | 174 | CARS1 | SYCC\_HUMAN | Cysteine--tRNA ligase, cytoplasmic | 9C(125.0477) | NaN | X | NaN | NaN | NaN | NaN | 1.131300e+04 | 1.131300e+04 | NaN | NaN | LTGEEVNSCVEVLLEEAK |
| UBR4 | UBR4\_HUMAN | E3 ubiquitin-protein ligase UBR4 | 1975 | 1974 | UBR4 | UBR4\_HUMAN | E3 ubiquitin-protein ligase UBR4 | 2C(125.0477) | NaN | X | NaN | NaN | NaN | NaN | 1.129985e+04 | 1.129985e+04 | NaN | NaN | DCHVLTFSSSGSVSDHLVLHPQLATGNFIIK |
| PI4KA | PI4KA\_HUMAN | Phosphatidylinositol 4-kinase alpha | 1832 | 1826 | PI4KA | PI4KA\_HUMAN | Phosphatidylinositol 4-kinase alpha | 7C(125.0477) | NaN | X | NaN | NaN | NaN | NaN | 1.122421e+04 | 1.122421e+04 | NaN | NaN | SDSEDECSTQEADGQK |
| HSPA1A | HS71A\_HUMAN | Heat shock 70 kDa protein 1A | 574 | 574 | HSPA1A | HS71A\_HUMAN | Heat shock 70 kDa protein 1A | 1C(125.0477) | NaN | X | NaN | NaN | NaN | NaN | 1.120000e+04 | 1.120000e+04 | NaN | NaN | CQEVISWLDANTLAEK |
| UBR2 | UBR2\_HUMAN | E3 ubiquitin-protein ligase UBR2 | 99 | 89 | UBR2 | UBR2\_HUMAN | E3 ubiquitin-protein ligase UBR2 | 11C(125.0477) | NaN | X | NaN | NaN | NaN | NaN | 1.119390e+04 | 1.119390e+04 | NaN | NaN | LEQANKPSHLCGR |
| DDB1 | DDB1\_HUMAN | DNA damage-binding protein 1 | 903 | 901 | DDB1 | DDB1\_HUMAN | DNA damage-binding protein 1 | 3C(125.0477) | NaN | X | NaN | NaN | NaN | NaN | 1.116628e+04 | 1.116628e+04 | NaN | NaN | TECNHYNNIMALYLK |
| CTPS1 | PYRG1\_HUMAN | CTP synthase 1 | 216 | 206 | CTPS1 | PYRG1\_HUMAN | CTP synthase 1 | 11C(125.0477) | NaN | X | NaN | NaN | NaN | NaN | 1.115920e+04 | NaN | 1.115920e+04 | NaN | GLGLSPDLVVCR |
| ANKRD13A | AN13A\_HUMAN | Ankyrin repeat domain-containing protein 13A | 540 | 527 | ANKRD13A | AN13A\_HUMAN | Ankyrin repeat domain-containing protein 13A | 14C(125.0477) | NaN | X | NaN | NaN | NaN | NaN | 1.113556e+04 | 1.113556e+04 | NaN | NaN | AIQESLLTSTEGLCPSALSETSR |
| TARS1 | SYTC\_HUMAN | Threonine--tRNA ligase 1, cytoplasmic | 656 | 646 | TARS1 | SYTC\_HUMAN | Threonine--tRNA ligase 1, cytoplasmic | 11C(125.0477) | NaN | X | NaN | NaN | NaN | NaN | 1.110128e+04 | 1.110128e+04 | NaN | NaN | FMADIDLDPGCTLNK |
| NBEAL2 | NBEL2\_HUMAN | Neurobeachin-like protein 2 | 207 | 202 | NBEAL2 | NBEL2\_HUMAN | Neurobeachin-like protein 2 | 6C(125.0477) | NaN | X | NaN | NaN | NaN | NaN | 1.109253e+04 | 1.109253e+04 | NaN | NaN | LIHLFCAVLAGGK |
| ATP11B | AT11B\_HUMAN | Phospholipid-transporting ATPase IF | 720 | 710 | ATP11B | AT11B\_HUMAN | Phospholipid-transporting ATPase IF | 11C(-34.0809) | NaN | NaN | NaN | NaN | X | NaN | 1.108546e+04 | 1.108546e+04 | NaN | NaN | HETAVSVSLSCGHFHR |
| VCPIP1 | VCIP1\_HUMAN | Deubiquitinating protein VCPIP1 | 816 | 807 | VCPIP1 | VCIP1\_HUMAN | Deubiquitinating protein VCPIP1 | 10C(125.0477) | NaN | X | NaN | NaN | NaN | NaN | 1.103982e+04 | 1.103982e+04 | NaN | NaN | EFNIPPYLQCIR |
| RABGGTA | PGTA\_HUMAN | Geranylgeranyl transferase type-2 subunit alpha | 457 | 445 | RABGGTA | PGTA\_HUMAN | Geranylgeranyl transferase type-2 subunit alpha | 13C(125.0477) | NaN | X | NaN | NaN | NaN | NaN | 1.101797e+04 | 1.101797e+04 | NaN | NaN | VLHLAHKDLTVLCHLEQLLLVTHLDLSHNR |
| CAST | ICAL\_HUMAN | Calpastatin | 381 | 379 | CAST | ICAL\_HUMAN | Calpastatin | 3C(125.0477) | NaN | X | NaN | NaN | NaN | NaN | 1.101088e+04 | 1.101088e+04 | NaN | NaN | SECKEKPSKPTEKTEESK |
| ZZEF1 | ZZEF1\_HUMAN | Zinc finger ZZ-type and EF-hand domain-containing protein 1 | 1636 | 1625 | ZZEF1 | ZZEF1\_HUMAN | Zinc finger ZZ-type and EF-hand domain-containing protein 1 | 12C(125.0477) | NaN | X | NaN | NaN | NaN | NaN | 1.099069e+04 | 1.099069e+04 | NaN | NaN | DFTNYFGHLEGCGADLHK |
| COPS6 | CSN6\_HUMAN | COP9 signalosome complex subunit 6 | 283 | 277 | COPS6 | CSN6\_HUMAN | COP9 signalosome complex subunit 6 | 7C(125.0477) | NaN | X | NaN | NaN | NaN | NaN | 1.095990e+04 | 1.095990e+04 | NaN | NaN | TDFYDQCNDVGLMAYLGTITK |
| CNPY3 | CNPY3\_HUMAN | Protein canopy homolog 3 | 104 | 96 | CNPY3 | CNPY3\_HUMAN | Protein canopy homolog 3 | 9C(125.0477) | NaN | X | NaN | NaN | NaN | NaN | 1.093802e+04 | NaN | 1.093802e+04 | NaN | LIEVTETICK |
| NUP43 | NUP43\_HUMAN | Nucleoporin Nup43 | 280 | 259 | NUP43 | NUP43\_HUMAN | Nucleoporin Nup43 | 22C(125.0477) | NaN | X | NaN | NaN | NaN | NaN | 1.088611e+04 | 1.088611e+04 | NaN | NaN | AHEAEMWEVHFHPSNPEHLFTCSEDGSLWHWDASTDVPEK |
| ANK1 | ANK1\_HUMAN | Ankyrin-1 | 41 | 28 | ANK1 | ANK1\_HUMAN | Ankyrin-1 | 14C(125.0477) | NaN | X | NaN | NaN | NaN | NaN | 1.087101e+04 | 1.087101e+04 | NaN | NaN | ALDHLRNGVDINTCNQNGLNGLHLASK |
| PSMA6 | PSA6\_HUMAN | Proteasome subunit alpha type-6 | 47 | 44 | PSMA6 | PSA6\_HUMAN | Proteasome subunit alpha type-6 | 4C(125.0477) | NaN | X | NaN | NaN | NaN | NaN | 1.084112e+04 | 1.084112e+04 | NaN | NaN | GKDCAVIVTQKK |
| KRT14 | K1C14\_HUMAN | Keratin, type I cytoskeletal 14 | 389, 389 | 389 | KRT14 | K1C14\_HUMAN | Keratin, type I cytoskeletal 14 | 1C(125.0477) 1C(15.9949) | NaN | X | X | NaN | NaN | NaN | 1.084049e+04 | NaN | 1.084049e+04 | NaN | CEMEQQNQEYK |
| KRT72 | K2C72\_HUMAN | Keratin, type II cytoskeletal 72 | 364 | 363 | KRT72 | K2C72\_HUMAN | Keratin, type II cytoskeletal 72 | 2C(57.0215) | X | NaN | NaN | NaN | NaN | NaN | 1.082666e+04 | NaN | 1.082666e+04 | NaN | QCADLETAIADAEQR |
| GFUS | FCL\_HUMAN | GDP-L-fucose synthase | 112, 116 | 108 | GFUS | FCL\_HUMAN | GDP-L-fucose synthase | 5C(15.9949) 9C(125.0477) | NaN | X | X | NaN | NaN | NaN | 1.079759e+04 | 1.079759e+04 | NaN | NaN | KVVSCLSTCIFPDK |
| TNPO3 | TNPO3\_HUMAN | Transportin-3 | 60 | 55 | TNPO3 | TNPO3\_HUMAN | Transportin-3 | 6C(125.0477) | NaN | X | NaN | NaN | NaN | NaN | 1.071418e+04 | 1.071418e+04 | NaN | NaN | QDVESCYFAAQTMK |
| EEF1A1 | EF1A1\_HUMAN | Elongation factor 1-alpha 1 | 411 | 396 | EEF1A1 | EF1A1\_HUMAN | Elongation factor 1-alpha 1 | 16C(57.0215) | X | NaN | NaN | NaN | NaN | NaN | 1.068712e+04 | NaN | 5.486520e+03 | 5200.5980 | SGDAAIVDMVPGKPMCVESFSDYPPLGR |
| SLC30A1 | ZNT1\_HUMAN | Zinc transporter 1 | 390 | 390 | SLC30A1 | ZNT1\_HUMAN | Zinc transporter 1 | 1C(125.0477) | NaN | X | NaN | NaN | NaN | NaN | 1.064925e+04 | NaN | 1.064925e+04 | NaN | CEDPTSYMEVAK |
| EPS15L1 | EP15R\_HUMAN | Epidermal growth factor receptor substrate 15-like 1 | 470 | 470 | EPS15L1 | EP15R\_HUMAN | Epidermal growth factor receptor substrate 15-like 1 | 1C(125.0477) | NaN | X | NaN | NaN | NaN | NaN | 1.060703e+04 | NaN | 1.060703e+04 | NaN | CQDETQMISSLK |
| GNB4 | GBB4\_HUMAN | Guanine nucleotide-binding protein subunit beta-4 | 25 | 24 | GNB4 | GBB4\_HUMAN | Guanine nucleotide-binding protein subunit beta-4 | 2C(125.0477) | NaN | X | NaN | NaN | NaN | NaN | 1.056935e+04 | 1.056935e+04 | NaN | NaN | ACNDATLVQITSNMDSVGR |
| STK11 | STK11\_HUMAN | Serine/threonine-protein kinase STK11 | 391 | 389 | STK11 | STK11\_HUMAN | Serine/threonine-protein kinase STK11 | 3C(125.0477) | NaN | X | NaN | NaN | NaN | NaN | 1.049925e+04 | 1.049925e+04 | NaN | NaN | AVCMNGTEAAQLSTK |
| HBB | HBB\_HUMAN | Hemoglobin subunit beta | 113 | 97 | HBB | HBB\_HUMAN | Hemoglobin subunit beta | 17C(125.0477) | NaN | X | NaN | NaN | NaN | NaN | 1.044743e+04 | NaN | 1.044743e+04 | NaN | LHVDPENFRLLGNVLVCVLAHHFGK |
| SWAP70 | SWP70\_HUMAN | Switch-associated protein 70 | 40 | 34 | SWAP70 | SWP70\_HUMAN | Switch-associated protein 70 | 7C(57.0215) | X | NaN | NaN | NaN | NaN | NaN | 1.043661e+04 | NaN | NaN | 10436.6140 | VLSHNLCTVLK |
| ARPC1B | ARC1B\_HUMAN | Actin-related protein 2/3 complex subunit 1B | 26 | 21 | ARPC1B | ARC1B\_HUMAN | Actin-related protein 2/3 complex subunit 1B | 6C(125.0477) | NaN | X | NaN | NaN | NaN | NaN | 1.035112e+04 | 1.035112e+04 | NaN | NaN | TQIAICPNNHEVHIYEK |
| VPS13A | VP13A\_HUMAN | Vacuolar protein sorting-associated protein 13A | 582 | 582 | VPS13A | VP13A\_HUMAN | Vacuolar protein sorting-associated protein 13A | 1C(125.0477) | NaN | X | NaN | NaN | NaN | NaN | 1.031606e+04 | 1.031606e+04 | NaN | NaN | CIIEAEPLEIIYDAR |
| MAP2K2 | MP2K2\_HUMAN | Dual specificity mitogen-activated protein kinase kinase 2 | 211 | 210 | MAP2K2 | MP2K2\_HUMAN | Dual specificity mitogen-activated protein kinase kinase 2 | 2C(125.0477) | NaN | X | NaN | NaN | NaN | NaN | 1.029152e+04 | 1.029152e+04 | NaN | NaN | LCDFGVSGQLIDSMANSFVGTR |
| CASP14 | CASPE\_HUMAN | Caspase-14 | 132 | 126 | CASP14 | CASPE\_HUMAN | Caspase-14 | 7C(125.0477) | NaN | X | NaN | NaN | NaN | NaN | 1.027430e+04 | NaN | 1.027430e+04 | NaN | VYIIQACR |
| DYNLL2 | DYL2\_HUMAN | Dynein light chain 2, cytoplasmic | 24 | 10 | DYNLL2 | DYL2\_HUMAN | Dynein light chain 2, cytoplasmic | 15C(125.0477) | NaN | X | NaN | NaN | NaN | NaN | 1.016395e+04 | 1.016395e+04 | NaN | NaN | NADMSEDMQQDAVDCATQAMEK |
| ATP5PO | ATPO\_HUMAN | ATP synthase subunit O, mitochondrial | 141 | 137 | ATP5PO | ATPO\_HUMAN | ATP synthase subunit O, mitochondrial | 5C(125.0477) | NaN | X | NaN | NaN | NaN | NaN | 1.009358e+04 | 1.009358e+04 | NaN | NaN | GEVPCTVTSASPLEEATLSELK |
| PRPS2 | PRPS2\_HUMAN | Ribose-phosphate pyrophosphokinase 2 | 165 | 164 | PRPS2 | PRPS2\_HUMAN | Ribose-phosphate pyrophosphokinase 2 | 2C(-34.0809) | NaN | NaN | NaN | NaN | X | NaN | 1.006174e+04 | 1.006174e+04 | NaN | NaN | NCIIVSPDAGGAK |
| ERP44 | ERP44\_HUMAN | Endoplasmic reticulum resident protein 44 | 92 | 90 | ERP44 | ERP44\_HUMAN | Endoplasmic reticulum resident protein 44 | 3C(31.9988) | NaN | NaN | NaN | X | NaN | NaN | 1.005978e+04 | NaN | NaN | 10059.7810 | VDCDQHSDIAQR |
| PPCDC | COAC\_HUMAN | Phosphopantothenoylcysteine decarboxylase | 122, 116 | 111 | PPCDC | COAC\_HUMAN | Phosphopantothenoylcysteine decarboxylase | 12C(-34.0809) 6C(57.0215) | X | NaN | NaN | NaN | X | NaN | 1.002859e+04 | NaN | 1.002859e+04 | NaN | VASGICDNLLTCVMR |
| ANXA2 | ANXA2\_HUMAN | Annexin A2 | 133 | 120 | ANXA2 | ANXA2\_HUMAN | Annexin A2 | 14C(125.0477) | NaN | X | NaN | NaN | NaN | NaN | 9.993023e+03 | 9.993023e+03 | NaN | NaN | GLGTDEDSLIEIICSR |
| SPTA1 | SPTA1\_HUMAN | Spectrin alpha chain, erythrocytic 1 | 2059 | 2052 | SPTA1 | SPTA1\_HUMAN | Spectrin alpha chain, erythrocytic 1 | 8C(125.0477) | NaN | X | NaN | NaN | NaN | NaN | 9.968184e+03 | NaN | NaN | 9968.1840 | ASALNNWCEK |
| HBQ1 | HBAT\_HUMAN | Hemoglobin subunit theta-1 | 90 | 63 | HBQ1 | HBAT\_HUMAN | Hemoglobin subunit theta-1 | 28C(125.0477) | NaN | X | NaN | NaN | NaN | NaN | 9.928384e+03 | 9.928384e+03 | NaN | NaN | VADALSLAVERLDDLPHALSALSHLHACQLR |
| APEH | ACPH\_HUMAN | Acylamino-acid-releasing enzyme | 30 | 23 | APEH | ACPH\_HUMAN | Acylamino-acid-releasing enzyme | 8C(125.0477) | NaN | X | NaN | NaN | NaN | NaN | 9.877901e+03 | 9.877901e+03 | NaN | NaN | QPALSAACLGPEVTTQYGGQYR |
| VCPIP1 | VCIP1\_HUMAN | Deubiquitinating protein VCPIP1 | 160 | 151 | VCPIP1 | VCIP1\_HUMAN | Deubiquitinating protein VCPIP1 | 10C(125.0477) | NaN | X | NaN | NaN | NaN | NaN | 9.860685e+03 | 9.860685e+03 | NaN | NaN | DMNQGELFDCALLGDR |
| PSMB2 | PSB2\_HUMAN | Proteasome subunit beta type-2 | 46 | 42 | PSMB2 | PSB2\_HUMAN | Proteasome subunit beta type-2 | 5C(125.0477) | NaN | X | NaN | NaN | NaN | NaN | 9.749344e+03 | 9.749344e+03 | NaN | NaN | ILLLCVGEAGDTVQFAEYIQK |
| TGM2 | TGM2\_HUMAN | Protein-glutamine gamma-glutamyltransferase 2 | 27 | 20 | TGM2 | TGM2\_HUMAN | Protein-glutamine gamma-glutamyltransferase 2 | 8C(125.0477) | NaN | X | NaN | NaN | NaN | NaN | 9.748321e+03 | 9.748321e+03 | NaN | NaN | DHHTADLCREK |
| ACOT7 | BACH\_HUMAN | Cytosolic acyl coenzyme A thioester hydrolase | 100 | 91 | ACOT7 | BACH\_HUMAN | Cytosolic acyl coenzyme A thioester hydrolase | 10C(15.9949) | NaN | NaN | X | NaN | NaN | NaN | 9.726032e+03 | 9.726032e+03 | NaN | NaN | HCNSQNGERCVAALAR |
| AKR1B1 | ALDR\_HUMAN | Aldo-keto reductase family 1 member B1 | 45 | 42 | AKR1B1 | ALDR\_HUMAN | Aldo-keto reductase family 1 member B1 | 4C(125.0477) | NaN | X | NaN | NaN | NaN | NaN | 9.686852e+03 | 9.686852e+03 | NaN | NaN | HIDCAHVYQNENEVGVAIQEK |
| TF | TRFE\_HUMAN | Serotransferrin | 582 | 577 | TF | TRFE\_HUMAN | Serotransferrin | 6C(57.0215) | X | NaN | NaN | NaN | NaN | NaN | 9.665851e+03 | NaN | 9.665851e+03 | NaN | DYELLCLDGTR |
| CA8 | CAH8\_HUMAN | Carbonic anhydrase-related protein | 80 | 79 | CA8 | CAH8\_HUMAN | Carbonic anhydrase-related protein | 2C(125.0477) | NaN | X | NaN | NaN | NaN | NaN | 9.611432e+03 | 9.611432e+03 | NaN | NaN | DCEVTNDGHTIQVILK |
| UBA5 | UBA5\_HUMAN | Ubiquitin-like modifier-activating enzyme 5 | 250 | 247 | UBA5 | UBA5\_HUMAN | Ubiquitin-like modifier-activating enzyme 5 | 4C(125.0477) | NaN | X | NaN | NaN | NaN | NaN | 9.591880e+03 | 9.591880e+03 | NaN | NaN | EGVCAASLPTTMGVVAGILVQNVLK |
| SELENBP1 | SBP1\_HUMAN | Methanethiol oxidase | 268 | 255 | SELENBP1 | SBP1\_HUMAN | Methanethiol oxidase | 14C(125.0477) | NaN | X | NaN | NaN | NaN | NaN | 9.553940e+03 | 9.553940e+03 | NaN | NaN | FLHNPDAAQGFVGCALSSTIQRFYKNEGGTWSVEK |
| PARK7 | PARK7\_HUMAN | Parkinson disease protein 7 | 53, 53 | 49 | PARK7 | PARK7\_HUMAN | Parkinson disease protein 7 | 5C(125.0477) 5C(15.9949) | NaN | X | X | NaN | NaN | NaN | 9.533595e+03 | 9.533595e+03 | NaN | NaN | DVVICPDASLEDAKKEGPYDVVVLPGGNLGAQNLSESAAVK |
| C5 | CO5\_HUMAN | Complement C5 | 1405 | 1401 | C5 | CO5\_HUMAN | Complement C5 | 5C(57.0215) | X | NaN | NaN | NaN | NaN | NaN | 9.457586e+03 | 9.457586e+03 | NaN | NaN | RIVACASYKPSR |
| MYO18A | MY18A\_HUMAN | Unconventional myosin-XVIIIa | 811 | 804 | MYO18A | MY18A\_HUMAN | Unconventional myosin-XVIIIa | 8C(125.0477) | NaN | X | NaN | NaN | NaN | NaN | 9.359191e+03 | 9.359191e+03 | NaN | NaN | GASFEELCHNYTQDR |
| MTHFD1 | C1TC\_HUMAN | C-1-tetrahydrofolate synthase, cytoplasmic | 152 | 151 | MTHFD1 | C1TC\_HUMAN | C-1-tetrahydrofolate synthase, cytoplasmic | 2C(125.0477) | NaN | X | NaN | NaN | NaN | NaN | 9.343981e+03 | 9.343981e+03 | NaN | NaN | GCLELIKETGVPIAGR |
| EPB41 | EPB41\_HUMAN | Protein 4.1 | 224 | 214 | EPB41 | EPB41\_HUMAN | Protein 4.1 | 11C(125.0477) | NaN | X | NaN | NaN | NaN | NaN | 9.329807e+03 | 9.329807e+03 | NaN | NaN | VSLLDDTVYECVVEKHAK |
| PFKL | PFKAL\_HUMAN | ATP-dependent 6-phosphofructokinase, liver type | 630 | 630 | PFKL | PFKAL\_HUMAN | ATP-dependent 6-phosphofructokinase, liver type | 1C(125.0477) | NaN | X | NaN | NaN | NaN | NaN | 9.280217e+03 | 9.280217e+03 | NaN | NaN | CHDYYTTEFLYNLYSSEGK |
| PPM1F | PPM1F\_HUMAN | Protein phosphatase 1F | 359 | 347 | PPM1F | PPM1F\_HUMAN | Protein phosphatase 1F | 13C(125.0477) | NaN | X | NaN | NaN | NaN | NaN | 9.279960e+03 | 9.279960e+03 | NaN | NaN | ALTGSEDYLLLACDGFFDVVPHQEVVGLVQSHLTR |
| LTA4H | LKHA4\_HUMAN | Leukotriene A-4 hydrolase | 136 | 128 | LTA4H | LKHA4\_HUMAN | Leukotriene A-4 hydrolase | 9C(15.9949) | NaN | NaN | X | NaN | NaN | NaN | 9.274068e+03 | 9.274068e+03 | NaN | NaN | EHPYLFSQCQAIHCR |
| MRI1 | MTNA\_HUMAN | Methylthioribose-1-phosphate isomerase | 168 | 162 | MRI1 | MTNA\_HUMAN | Methylthioribose-1-phosphate isomerase | 7C(31.9988) | NaN | NaN | NaN | X | NaN | NaN | 9.270074e+03 | 9.270074e+03 | NaN | NaN | VTVLTHCNTGALATAGYGTALGVIR |
| HUWE1 | HUWE1\_HUMAN | E3 ubiquitin-protein ligase HUWE1 | 1628 | 1627 | HUWE1 | HUWE1\_HUMAN | E3 ubiquitin-protein ligase HUWE1 | 2C(125.0477) | NaN | X | NaN | NaN | NaN | NaN | 9.263761e+03 | 9.263761e+03 | NaN | NaN | WCSYSASNNSTIDSAWK |
| ANKRD28 | ANR28\_HUMAN | Serine/threonine-protein phosphatase 6 regulatory ankyrin repeat subunit A | 334 | 322 | ANKRD28 | ANR28\_HUMAN | Serine/threonine-protein phosphatase 6 regulatory ankyrin repeat subunit A | 13C(125.0477) | NaN | X | NaN | NaN | NaN | NaN | 9.248902e+03 | 9.248902e+03 | NaN | NaN | SQTIIQSGAVIDCEDKNGNTPLHIAAR |
| TMED7 | TMED7\_HUMAN | Transmembrane emp24 domain-containing protein 7 | 48 | 47 | TMED7 | TMED7\_HUMAN | Transmembrane emp24 domain-containing protein 7 | 2C(125.0477) | NaN | X | NaN | NaN | NaN | NaN | 9.248160e+03 | NaN | 9.248160e+03 | NaN | QCFYEDIAQGTK |
| YWHAG | 1433G\_HUMAN | 14-3-3 protein gamma | 112 | 111 | YWHAG | 1433G\_HUMAN | 14-3-3 protein gamma | 2C(125.0477) | NaN | X | NaN | NaN | NaN | NaN | 9.180344e+03 | 9.180344e+03 | NaN | NaN | NCSETQYESKVFYLK |
| IGHG1 | IGHG1\_HUMAN | Immunoglobulin heavy constant gamma 1 | 144, 144 | 139 | IGHG1 | IGHG1\_HUMAN | Immunoglobulin heavy constant gamma 1 | 6C(125.0477) 6C(57.0215) | X | X | NaN | NaN | NaN | NaN | 9.126116e+03 | 9.126116e+03 | NaN | NaN | TPEVTCVVVDVSHEDPEVK |
| MYH9 | MYH9\_HUMAN | Myosin-9 | 1379 | 1372 | MYH9 | MYH9\_HUMAN | Myosin-9 | 8C(125.0477) | NaN | X | NaN | NaN | NaN | NaN | 9.118394e+03 | 9.118394e+03 | NaN | NaN | KMEDSVGCLETAEEVKR |
| ECPAS | ECM29\_HUMAN | Proteasome adapter and scaffold protein ECM29 | 1420 | 1419 | ECPAS | ECM29\_HUMAN | Proteasome adapter and scaffold protein ECM29 | 2C(125.0477) | NaN | X | NaN | NaN | NaN | NaN | 9.093462e+03 | 9.093462e+03 | NaN | NaN | SCAFAMGHLVR |
| GNAQ | GNAQ\_HUMAN | Guanine nucleotide-binding protein G(q) subunit alpha | 330 | 323 | GNAQ | GNAQ\_HUMAN | Guanine nucleotide-binding protein G(q) subunit alpha | 8C(125.0477) | NaN | X | NaN | NaN | NaN | NaN | 9.065652e+03 | 9.065652e+03 | NaN | NaN | IIYSHFTCATDTENIR |
| EFL1 | EFL1\_HUMAN | Elongation factor-like GTPase 1 | 23 | 21 | EFL1 | EFL1\_HUMAN | Elongation factor-like GTPase 1 | 3C(15.9949) | NaN | NaN | X | NaN | NaN | NaN | 9.042478e+03 | 9.042478e+03 | NaN | NaN | NICVLAHVDHGK |
| NAPA | SNAA\_HUMAN | Alpha-soluble NSF attachment protein | 66 | 57 | NAPA | SNAA\_HUMAN | Alpha-soluble NSF attachment protein | 10C(125.0477) | NaN | X | NaN | NaN | NaN | NaN | 9.027129e+03 | 9.027129e+03 | NaN | NaN | NWSAAGNAFCQAAQLHLQLQSK |
| MAP2K4 | MP2K4\_HUMAN | Dual specificity mitogen-activated protein kinase kinase 4 | 347 | 335 | MAP2K4 | MP2K4\_HUMAN | Dual specificity mitogen-activated protein kinase kinase 4 | 13C(125.0477) | NaN | X | NaN | NaN | NaN | NaN | 8.998251e+03 | 8.998251e+03 | NaN | NaN | EFSPSFINFVNLCLTK |
| COL6A1 | CO6A1\_HUMAN | Collagen alpha-1(VI) chain | 745 | 737 | COL6A1 | CO6A1\_HUMAN | Collagen alpha-1(VI) chain | 9C(57.0215) | X | NaN | NaN | NaN | NaN | NaN | 8.989970e+03 | 8.989970e+03 | NaN | NaN | DTTPLNVLCSPGIQVVSVGIK |
| METAP1 | MAP11\_HUMAN | Methionine aminopeptidase 1 | 174 | 148 | METAP1 | MAP11\_HUMAN | Methionine aminopeptidase 1 | 27C(125.0477) | NaN | X | NaN | NaN | NaN | NaN | 8.970307e+03 | 8.970307e+03 | NaN | NaN | EVLDVAAGMIKPGVTTEEIDHAVHLACIAR |
| DENND1A | DEN1A\_HUMAN | DENN domain-containing protein 1A | 459 | 453 | DENND1A | DEN1A\_HUMAN | DENN domain-containing protein 1A | 7C(125.0477) | NaN | X | NaN | NaN | NaN | NaN | 8.957241e+03 | 8.957241e+03 | NaN | NaN | DIAENGCAPTPEEQLPK |
| PSMD1 | PSMD1\_HUMAN | 26S proteasome non-ATPase regulatory subunit 1 | 591 | 581 | PSMD1 | PSMD1\_HUMAN | 26S proteasome non-ATPase regulatory subunit 1 | 11C(125.0477) | NaN | X | NaN | NaN | NaN | NaN | 8.955439e+03 | 8.955439e+03 | NaN | NaN | SGMYTVAMAYCGSGNNK |
| OSBP2 | OSBP2\_HUMAN | Oxysterol-binding protein 2 | 224 | 216 | OSBP2 | OSBP2\_HUMAN | Oxysterol-binding protein 2 | 9C(125.0477) | NaN | X | NaN | NaN | NaN | NaN | 8.933600e+03 | 8.933600e+03 | NaN | NaN | NQGEMAHTCR |
| SACM1L | SAC1\_HUMAN | Phosphatidylinositol-3-phosphatase SAC1 | 445 | 436 | SACM1L | SAC1\_HUMAN | Phosphatidylinositol-3-phosphatase SAC1 | 10C(125.0477) | NaN | X | NaN | NaN | NaN | NaN | 8.910438e+03 | NaN | 8.910438e+03 | NaN | NAWADNANACAK |
| AMPD3 | AMPD3\_HUMAN | AMP deaminase 3 | 587 | 577 | AMPD3 | AMPD3\_HUMAN | AMP deaminase 3 | 11C(125.0477) | NaN | X | NaN | NaN | NaN | NaN | 8.889171e+03 | 8.889171e+03 | NaN | NaN | GLSTFLFRPHCGEAGSITHLVSAFLTADNISHGLLLK |
| MKI67 | KI67\_HUMAN | Proliferation marker protein Ki-67 | 27 | 13 | MKI67 | KI67\_HUMAN | Proliferation marker protein Ki-67 | 15C(125.0477) | NaN | X | NaN | NaN | NaN | NaN | 8.888881e+03 | 8.888881e+03 | NaN | NaN | SGVDGPHFPLSLSTCLFGR |
| THEM6 | THEM6\_HUMAN | Protein THEM6 | 29 | 27 | THEM6 | THEM6\_HUMAN | Protein THEM6 | 3C(125.0477) | NaN | X | NaN | NaN | NaN | NaN | 8.768323e+03 | NaN | 8.768323e+03 | NaN | LPCAVLR |
| THEM6 | THEM6\_HUMAN | Protein THEM6 | 85 | 85 | THEM6 | THEM6\_HUMAN | Protein THEM6 | 1C(125.0477) | NaN | X | NaN | NaN | NaN | NaN | 8.720959e+03 | NaN | 8.720959e+03 | NaN | CGVLGALR |
| PKM | KPYM\_HUMAN | Pyruvate kinase PKM | 49 | 44 | PKM | KPYM\_HUMAN | Pyruvate kinase PKM | 6C(125.0477) | NaN | X | NaN | NaN | NaN | NaN | 8.661328e+03 | NaN | NaN | 8661.3280 | NTGIICTIGPASR |
| SERPIND1 | HEP2\_HUMAN | Heparin cofactor 2 | 486 | 469 | SERPIND1 | HEP2\_HUMAN | Heparin cofactor 2 | 18C(57.0215) | X | NaN | NaN | NaN | NaN | NaN | 8.595938e+03 | 8.595938e+03 | NaN | NaN | FTVDRPFLFLIYEHRTSCLLFMGR |
| EPB41L4A | E41LA\_HUMAN | Band 4.1-like protein 4A | 287, 287 | 287 | EPB41L4A | E41LA\_HUMAN | Band 4.1-like protein 4A | 1C(125.0477) 1C(57.0215) | X | X | NaN | NaN | NaN | NaN | 8.560176e+03 | NaN | 8.560176e+03 | NaN | CSVEHHTFFR |
| HSP90AA1 | HS90A\_HUMAN | Heat shock protein HSP 90-alpha | 481 | 465 | HSP90AA1 | HS90A\_HUMAN | Heat shock protein HSP 90-alpha | 17C(125.0477) | NaN | X | NaN | NaN | NaN | NaN | 8.533048e+03 | 8.533048e+03 | NaN | NaN | YYTSASGDEMVSLKDYCTR |
| RABGAP1 | RBGP1\_HUMAN | Rab GTPase-activating protein 1 | 346 | 346 | RABGAP1 | RBGP1\_HUMAN | Rab GTPase-activating protein 1 | 1C(125.0477) | NaN | X | NaN | NaN | NaN | NaN | 8.532751e+03 | 8.532751e+03 | NaN | NaN | CFGLLLSPGK |
| MYO18A | MY18A\_HUMAN | Unconventional myosin-XVIIIa | 1046 | 1041 | MYO18A | MY18A\_HUMAN | Unconventional myosin-XVIIIa | 6C(125.0477) | NaN | X | NaN | NaN | NaN | NaN | 8.423706e+03 | 8.423706e+03 | NaN | NaN | LHFVHCFLPVAEGWAGEPR |
| PHAF1 | PHAF1\_HUMAN | Phagosome assembly factor 1 | 385 | 371 | PHAF1 | PHAF1\_HUMAN | Phagosome assembly factor 1 | 15C(125.0477) | NaN | X | NaN | NaN | NaN | NaN | 8.407617e+03 | 8.407617e+03 | NaN | NaN | SSSPNNTNPFGSTFCFGLQR |
| GPX1 | GPX1\_HUMAN | Glutathione peroxidase 1 | 202 | 183 | GPX1 | GPX1\_HUMAN | Glutathione peroxidase 1 | 20C(125.0477) | NaN | X | NaN | NaN | NaN | NaN | 8.391027e+03 | 8.391027e+03 | NaN | NaN | FQTIDIEPDIEALLSQGPSCA |
| SEMA7A | SEM7A\_HUMAN | Semaphorin-7A | 291 | 287 | SEMA7A | SEM7A\_HUMAN | Semaphorin-7A | 5C(57.0215) | X | NaN | NaN | NaN | NaN | NaN | 8.299744e+03 | NaN | NaN | 8299.7440 | AMLVCSDAATNK |
| EHBP1L1 | EH1L1\_HUMAN | EH domain-binding protein 1-like protein 1 | 422 | 412 | EHBP1L1 | EH1L1\_HUMAN | EH domain-binding protein 1-like protein 1 | 11C(125.0477) | NaN | X | NaN | NaN | NaN | NaN | 8.269528e+03 | NaN | 8.269528e+03 | NaN | AGEAEESSAVCQVDAEQR |
| RPLP0 | RLA0\_HUMAN | 60S acidic ribosomal protein P0 | 27 | 27 | RPLP0 | RLA0\_HUMAN | 60S acidic ribosomal protein P0 | 1C(125.0477) | NaN | X | NaN | NaN | NaN | NaN | 8.228138e+03 | NaN | 8.228138e+03 | NaN | CFIVGADNVGSK |
| FLNB | FLNB\_HUMAN | Filamin-B | 1617 | 1617 | FLNB | FLNB\_HUMAN | Filamin-B | 1C(125.0477) | NaN | X | NaN | NaN | NaN | NaN | 8.225806e+03 | NaN | 8.225806e+03 | NaN | CLATGPGIASTVK |
| KRT86 | KRT86\_HUMAN | Keratin, type II cuticular Hb6 | 341, 341, 341 | 341 | KRT86 | KRT86\_HUMAN | Keratin, type II cuticular Hb6 | 1C(15.9949) 1C(31.9988) 1C(57.0215) | X | NaN | X | X | NaN | NaN | 8.190934e+03 | 8.190934e+03 | NaN | NaN | CQNSKLEAAVAQSEQQGEAALSDARCK |
| PGP | PGP\_HUMAN | Glycerol-3-phosphate phosphatase | 243 | 239 | PGP | PGP\_HUMAN | Glycerol-3-phosphate phosphatase | 5C(125.0477) | NaN | X | NaN | NaN | NaN | NaN | 8.184234e+03 | 8.184234e+03 | NaN | NaN | FIFDCVSQEYGINPER |
| FH | FUMH\_HUMAN | Fumarate hydratase, mitochondrial | 377 | 372 | FH | FUMH\_HUMAN | Fumarate hydratase, mitochondrial | 6C(125.0477) | NaN | X | NaN | NaN | NaN | NaN | 8.182887e+03 | 8.182887e+03 | NaN | NaN | VNPTQCEAMTMVAAQVMGNHVAVTVGGSNGHFELNVFKPMMIK |
| ADH5 | ADHX\_HUMAN | Alcohol dehydrogenase class-3 | 60, 45 | 39 | ADH5 | ADHX\_HUMAN | Alcohol dehydrogenase class-3 | 22C(31.9988) 7C(-34.0809) | NaN | NaN | NaN | X | X | NaN | 8.171291e+03 | 8.171291e+03 | NaN | NaN | IIATAVCHTDAYTLSGADPEGCFPVILGHEGAGIVESVGEGVTK |
| JCHAIN | IGJ\_HUMAN | Immunoglobulin J chain | 131 | 131 | JCHAIN | IGJ\_HUMAN | Immunoglobulin J chain | 1C(125.0477) | NaN | X | NaN | NaN | NaN | NaN | 8.157279e+03 | NaN | 8.157279e+03 | NaN | CYTAVVPLVYGGETK |
| SEC24A | SC24A\_HUMAN | Protein transport protein Sec24A | 704 | 675 | SEC24A | SC24A\_HUMAN | Protein transport protein Sec24A | 30C(15.9949) | NaN | NaN | X | NaN | NaN | NaN | 8.155917e+03 | 8.155917e+03 | NaN | NaN | KLALDCSGQQVAVDLFLLSGQYSDLASLGCISR |
| RUVBL1 | RUVB1\_HUMAN | RuvB-like 1 | 141 | 129 | RUVBL1 | RUVB1\_HUMAN | RuvB-like 1 | 13C(125.0477) | NaN | X | NaN | NaN | NaN | NaN | 8.147385e+03 | 8.147385e+03 | NaN | NaN | EVYEGEVTELTPCETENPMGGYGK |
| USP24 | UBP24\_HUMAN | Ubiquitin carboxyl-terminal hydrolase 24 | 679 | 671 | USP24 | UBP24\_HUMAN | Ubiquitin carboxyl-terminal hydrolase 24 | 9C(125.0477) | NaN | X | NaN | NaN | NaN | NaN | 8.129920e+03 | 8.129920e+03 | NaN | NaN | LVTGSLIACHR |
| IDE | IDE\_HUMAN | Insulin-degrading enzyme | 257 | 244 | IDE | IDE\_HUMAN | Insulin-degrading enzyme | 14C(125.0477) | NaN | X | NaN | NaN | NaN | NaN | 8.012158e+03 | 8.012158e+03 | NaN | NaN | FHSAYYSSNLMAVCVLGR |
| UBA6 | UBA6\_HUMAN | Ubiquitin-like modifier-activating enzyme 6 | 546 | 545 | UBA6 | UBA6\_HUMAN | Ubiquitin-like modifier-activating enzyme 6 | 2C(125.0477) | NaN | X | NaN | NaN | NaN | NaN | 8.002574e+03 | 8.002574e+03 | NaN | NaN | VCPTTETIYNDEFYTK |
| NaN | YJ005\_HUMAN | Uncharacterized protein FLJ45252 | 251 | 249 | NaN | YJ005\_HUMAN | Uncharacterized protein FLJ45252 | 3C(125.0477) | NaN | X | NaN | NaN | NaN | NaN | 7.996490e+03 | 7.996490e+03 | NaN | NaN | GSCLESGGHSNDR |
| KRT78 | K2C78\_HUMAN | Keratin, type II cytoskeletal 78 | 190 | 189 | KRT78 | K2C78\_HUMAN | Keratin, type II cytoskeletal 78 | 2C(125.0477) | NaN | X | NaN | NaN | NaN | NaN | 7.958460e+03 | 7.958460e+03 | NaN | NaN | ACRDQEEEYKSK |
| DENND4C | DEN4C\_HUMAN | DENN domain-containing protein 4C | 1211 | 1195 | DENND4C | DEN4C\_HUMAN | DENN domain-containing protein 4C | 17C(125.0477) | NaN | X | NaN | NaN | NaN | NaN | 7.950061e+03 | 7.950061e+03 | NaN | NaN | EDVETGLDPLSLLATECTGGK |
| IGHG4 | IGHG4\_HUMAN | Immunoglobulin heavy constant gamma 4 | 106 | 102 | IGHG4 | IGHG4\_HUMAN | Immunoglobulin heavy constant gamma 4 | 5C(15.9949) | NaN | NaN | X | NaN | NaN | NaN | 7.906596e+03 | 7.906596e+03 | NaN | NaN | YGPPCPSCPAPEFLGGPSVFLFPPKPK |
| TGM2 | TGM2\_HUMAN | Protein-glutamine gamma-glutamyltransferase 2 | 524 | 513 | TGM2 | TGM2\_HUMAN | Protein-glutamine gamma-glutamyltransferase 2 | 12C(57.0215) | X | NaN | NaN | NaN | NaN | NaN | 7.902300e+03 | 7.902300e+03 | NaN | NaN | TVSYNGILGPECGTKYLLNLNLEPFSEK |
| CAPZB | CAPZB\_HUMAN | F-actin-capping protein subunit beta | 206 | 200 | CAPZB | CAPZB\_HUMAN | F-actin-capping protein subunit beta | 7C(125.0477) | NaN | X | NaN | NaN | NaN | NaN | 7.865032e+03 | 7.865032e+03 | NaN | NaN | DETVSDCSPHIANIGR |
| AP2A1 | AP2A1\_HUMAN | AP-2 complex subunit alpha-1 | 397 | 389 | AP2A1 | AP2A1\_HUMAN | AP-2 complex subunit alpha-1 | 9C(125.0477) | NaN | X | NaN | NaN | NaN | NaN | 7.752087e+03 | NaN | 7.752087e+03 | NaN | AADLLYAMCDR |
| MSN | MOES\_HUMAN | Moesin | 117 | 108 | MSN | MOES\_HUMAN | Moesin | 10C(125.0477) | NaN | X | NaN | NaN | NaN | NaN | 7.747557e+03 | 7.747557e+03 | NaN | NaN | EGILNDDIYCPPETAVLLASYAVQSK |
| TRAPPC8 | TPPC8\_HUMAN | Trafficking protein particle complex subunit 8 | 548 | 538 | TRAPPC8 | TPPC8\_HUMAN | Trafficking protein particle complex subunit 8 | 11C(125.0477) | NaN | X | NaN | NaN | NaN | NaN | 7.697904e+03 | 7.697904e+03 | NaN | NaN | SALLLEQAAHCFINMK |
| ALB | ALBU\_HUMAN | Albumin | 416 | 397 | ALB | ALBU\_HUMAN | Albumin | 20C(57.0215) | X | NaN | NaN | NaN | NaN | NaN | 7.680232e+03 | 7.680232e+03 | NaN | NaN | VFDEFKPLVEEPQNLIKQNCELFEQLGEYKFQNALLVR |
| MAEA | MAEA\_HUMAN | E3 ubiquitin-protein transferase MAEA | 314 | 311 | MAEA | MAEA\_HUMAN | E3 ubiquitin-protein transferase MAEA | 4C(57.0215) | X | NaN | NaN | NaN | NaN | NaN | 7.645973e+03 | NaN | 7.645973e+03 | NaN | SPDCPVCSRSLNK |
| PPP6R1 | PP6R1\_HUMAN | Serine/threonine-protein phosphatase 6 regulatory subunit 1 | 795 | 788 | PPP6R1 | PP6R1\_HUMAN | Serine/threonine-protein phosphatase 6 regulatory subunit 1 | 8C(125.0477) | NaN | X | NaN | NaN | NaN | NaN | 7.641440e+03 | 7.641440e+03 | NaN | NaN | VTEPSAPCQALVSIGDLQATFHGIR |
| TLN1 | TLN1\_HUMAN | Talin-1 | 1927 | 1920 | TLN1 | TLN1\_HUMAN | Talin-1 | 8C(31.9988) | NaN | NaN | NaN | X | NaN | NaN | 7.615406e+03 | NaN | 7.615406e+03 | NaN | VQELGHGCAALVTK |
| CIAO1 | CIAO1\_HUMAN | Probable cytosolic iron-sulfur protein assembly protein CIAO1 | 234 | 223 | CIAO1 | CIAO1\_HUMAN | Probable cytosolic iron-sulfur protein assembly protein CIAO1 | 12C(125.0477) | NaN | X | NaN | NaN | NaN | NaN | 7.560977e+03 | 7.560977e+03 | NaN | NaN | QYLPGNEQGVACSGSDPSWK |
| CLTCL1 | CLH2\_HUMAN | Clathrin heavy chain 2 | 934, 926 | 924 | CLTCL1 | CLH2\_HUMAN | Clathrin heavy chain 2 | 11C(15.9949) 3C(-34.0809) | NaN | NaN | X | NaN | X | NaN | 7.531348e+03 | 7.531348e+03 | NaN | NaN | GQCDLELIKVCNENSLFK |
| PRDX2 | PRDX2\_HUMAN | Peroxiredoxin-2 | 70 | 68 | PRDX2 | PRDX2\_HUMAN | Peroxiredoxin-2 | 3C(125.0477) | NaN | X | NaN | NaN | NaN | NaN | 7.513773e+03 | NaN | 7.513773e+03 | NaN | LGCEVLGVSVDSQFTHLAWINTPR |
| ATP11C | AT11C\_HUMAN | Phospholipid-transporting ATPase IG | 200 | 195 | ATP11C | AT11C\_HUMAN | Phospholipid-transporting ATPase IG | 6C(125.0477) | NaN | X | NaN | NaN | NaN | NaN | 7.483818e+03 | NaN | 7.483818e+03 | NaN | DTIALCTAESIDTLR |
| CAPZB | CAPZB\_HUMAN | F-actin-capping protein subunit beta | 62 | 58 | CAPZB | CAPZB\_HUMAN | F-actin-capping protein subunit beta | 5C(125.0477) | NaN | X | NaN | NaN | NaN | NaN | 7.447746e+03 | NaN | 7.447746e+03 | NaN | DYLLCDYNR |
| SORD | DHSO\_HUMAN | Sorbitol dehydrogenase | 45 | 39 | SORD | DHSO\_HUMAN | Sorbitol dehydrogenase | 7C(125.0477) | NaN | X | NaN | NaN | NaN | NaN | 7.405621e+03 | 7.405621e+03 | NaN | NaN | MHSVGICGSDVHYWEYGR |
| HECTD4 | HECD4\_HUMAN | Probable E3 ubiquitin-protein ligase HECTD4 | 3673 | 3661 | HECTD4 | HECD4\_HUMAN | Probable E3 ubiquitin-protein ligase HECTD4 | 13C(125.0477) | NaN | X | NaN | NaN | NaN | NaN | 7.390553e+03 | 7.390553e+03 | NaN | NaN | ELQSSSLSLLLLCPSSAVNK |
| LRRC20 | LRC20\_HUMAN | Leucine-rich repeat-containing protein 20 | 32 | 15 | LRRC20 | LRC20\_HUMAN | Leucine-rich repeat-containing protein 20 | 18C(125.0477) | NaN | X | NaN | NaN | NaN | NaN | 7.303076e+03 | 7.303076e+03 | NaN | NaN | KVNETVESGSDTLDLAECK |
| HSPA1A | HS71A\_HUMAN | Heat shock 70 kDa protein 1A | 306 | 302 | HSPA1A | HS71A\_HUMAN | Heat shock 70 kDa protein 1A | 5C(57.0215) | X | NaN | NaN | NaN | NaN | NaN | 7.289648e+03 | NaN | NaN | 7289.6484 | FEELCSDLFR |
| PHAX | PHAX\_HUMAN | Phosphorylated adapter RNA export protein | 51 | 43 | PHAX | PHAX\_HUMAN | Phosphorylated adapter RNA export protein | 9C(125.0477) | NaN | X | NaN | NaN | NaN | NaN | 7.258912e+03 | 7.258912e+03 | NaN | NaN | AFQNTATACAPVSHYR |
| PFKM | PFKAM\_HUMAN | ATP-dependent 6-phosphofructokinase, muscle type | 631 | 631 | PFKM | PFKAM\_HUMAN | ATP-dependent 6-phosphofructokinase, muscle type | 1C(125.0477) | NaN | X | NaN | NaN | NaN | NaN | 7.097955e+03 | 7.097955e+03 | NaN | NaN | CNENYTTDFIFNLYSEEGK |
| VPS13A | VP13A\_HUMAN | Vacuolar protein sorting-associated protein 13A | 2377 | 2374 | VPS13A | VP13A\_HUMAN | Vacuolar protein sorting-associated protein 13A | 4C(125.0477) | NaN | X | NaN | NaN | NaN | NaN | 7.044828e+03 | NaN | 7.044828e+03 | NaN | QENCILLR |
| GGCT | GGCT\_HUMAN | Gamma-glutamylcyclotransferase | 144 | 142 | GGCT | GGCT\_HUMAN | Gamma-glutamylcyclotransferase | 3C(125.0477) | NaN | X | NaN | NaN | NaN | NaN | 7.038349e+03 | 7.038349e+03 | NaN | NaN | IICMGAK |
| ABCG2 | ABCG2\_HUMAN | Broad substrate specificity ATP-binding cassette transporter ABCG2 | 43 | 30 | ABCG2 | ABCG2\_HUMAN | Broad substrate specificity ATP-binding cassette transporter ABCG2 | 14C(125.0477) | NaN | X | NaN | NaN | NaN | NaN | 7.026120e+03 | 7.026120e+03 | NaN | NaN | AFTEGAVLSFHNICYR |
| PKM | KPYM\_HUMAN | Pyruvate kinase PKM | 49 | 44 | PKM | KPYM\_HUMAN | Pyruvate kinase PKM | 6C(-34.0809) | NaN | NaN | NaN | NaN | X | NaN | 6.980791e+03 | NaN | 6.980791e+03 | NaN | NTGIICTIGPASR |
| UBR4 | UBR4\_HUMAN | E3 ubiquitin-protein ligase UBR4 | 2222 | 2219 | UBR4 | UBR4\_HUMAN | E3 ubiquitin-protein ligase UBR4 | 4C(125.0477) | NaN | X | NaN | NaN | NaN | NaN | 6.929389e+03 | 6.929389e+03 | NaN | NaN | HTACNEQQR |
| GTPBP1 | GTPB1\_HUMAN | GTP-binding protein 1 | 531 | 520 | GTPBP1 | GTPB1\_HUMAN | GTP-binding protein 1 | 12C(125.0477) | NaN | X | NaN | NaN | NaN | NaN | 6.749570e+03 | 6.749570e+03 | NaN | NaN | QTATILSMDKDCLR |
| AGO2 | AGO2\_HUMAN | Protein argonaute-2 | 272 | 267 | AGO2 | AGO2\_HUMAN | Protein argonaute-2 | 6C(125.0477) | NaN | X | NaN | NaN | NaN | NaN | 6.689697e+03 | NaN | 6.689697e+03 | NaN | VEITHCGQMK |
| TUBA1A | TBA1A\_HUMAN | Tubulin alpha-1A chain | 347 | 340 | TUBA1A | TBA1A\_HUMAN | Tubulin alpha-1A chain | 8C(57.0215) | X | NaN | NaN | NaN | NaN | NaN | 6.657459e+03 | NaN | NaN | 6657.4590 | TIQFVDWCPTGFK |
| DHRS11 | DHR11\_HUMAN | Dehydrogenase/reductase SDR family member 11 | 41 | 30 | DHRS11 | DHR11\_HUMAN | Dehydrogenase/reductase SDR family member 11 | 12C(125.0477) | NaN | X | NaN | NaN | NaN | NaN | 6.654804e+03 | 6.654804e+03 | NaN | NaN | ALVQQGLKVVGCAR |
| GAPDH | G3P\_HUMAN | Glyceraldehyde-3-phosphate dehydrogenase | 247 | 235 | GAPDH | G3P\_HUMAN | Glyceraldehyde-3-phosphate dehydrogenase | 13C(57.0215) | X | NaN | NaN | NaN | NaN | NaN | 6.605717e+03 | NaN | NaN | 6605.7170 | VPTANVSVVDLTCR |
| DAZAP1 | DAZP1\_HUMAN | DAZ-associated protein 1 | 124 | 115 | DAZAP1 | DAZP1\_HUMAN | DAZ-associated protein 1 | 10C(15.9949) | NaN | NaN | X | NaN | NaN | NaN | 6.571146e+03 | NaN | 6.571146e+03 | NaN | IFVGGIPHNCGETELR |
| RAN | RAN\_HUMAN | GTP-binding nuclear protein Ran | 112 | 111 | RAN | RAN\_HUMAN | GTP-binding nuclear protein Ran | 2C(125.0477) | NaN | X | NaN | NaN | NaN | NaN | 6.520744e+03 | 6.520744e+03 | NaN | NaN | VCENIPIVLCGNKVDIK |
| MINPP1 | MINP1\_HUMAN | Multiple inositol polyphosphate phosphatase 1 | 302 | 299 | MINPP1 | MINP1\_HUMAN | Multiple inositol polyphosphate phosphatase 1 | 4C(125.0477) | NaN | X | NaN | NaN | NaN | NaN | 6.512272e+03 | NaN | 6.512272e+03 | NaN | SPWCDVFDIDDAK |
| ACTBL2 | ACTBL\_HUMAN | Beta-actin-like protein 2 | 53 | 52 | ACTBL2 | ACTBL\_HUMAN | Beta-actin-like protein 2 | 2C(125.0477) | NaN | X | NaN | NaN | NaN | NaN | 6.422804e+03 | NaN | NaN | 6422.8037 | DCYVGDEAQSKR |
| UBR4 | UBR4\_HUMAN | E3 ubiquitin-protein ligase UBR4 | 4642 | 4638 | UBR4 | UBR4\_HUMAN | E3 ubiquitin-protein ligase UBR4 | 5C(125.0477) | NaN | X | NaN | NaN | NaN | NaN | 6.385310e+03 | 6.385310e+03 | NaN | NaN | FKPYCNFDKYDEDHSGDDK |
| PRDX4 | PRDX4\_HUMAN | Peroxiredoxin-4 | 148, 148 | 140 | PRDX4 | PRDX4\_HUMAN | Peroxiredoxin-4 | 9C(15.9949) 9C(31.9988) | NaN | NaN | X | X | NaN | NaN | 6.326515e+03 | 6.326515e+03 | NaN | NaN | SINTEVVACSVDSQFTHLAWINTPR |
| ACSL6 | ACSL6\_HUMAN | Long-chain-fatty-acid--CoA ligase 6 | 336 | 329 | ACSL6 | ACSL6\_HUMAN | Long-chain-fatty-acid--CoA ligase 6 | 8C(125.0477) | NaN | X | NaN | NaN | NaN | NaN | 6.283377e+03 | 6.283377e+03 | NaN | NaN | VIQSVVYCHGGR |
| VAC14 | VAC14\_HUMAN | Protein VAC14 homolog | 372 | 360 | VAC14 | VAC14\_HUMAN | Protein VAC14 homolog | 13C(125.0477) | NaN | X | NaN | NaN | NaN | NaN | 6.283248e+03 | 6.283248e+03 | NaN | NaN | QEGTASGGPDGSCDSSFSSGISVFTAASTER |
| NIT1 | NIT1\_HUMAN | Deaminated glutathione amidase | 285, 292 | 285 | NIT1 | NIT1\_HUMAN | Deaminated glutathione amidase | 1C(57.0215) 8C(15.9949) | X | NaN | X | NaN | NaN | NaN | 6.217816e+03 | NaN | NaN | 6217.8164 | CSEGPGLCLAR |
| UHRF1BP1 | URFB1\_HUMAN | UHRF1-binding protein 1 | 729 | 713 | UHRF1BP1 | URFB1\_HUMAN | UHRF1-binding protein 1 | 17C(15.9949) | NaN | NaN | X | NaN | NaN | NaN | 6.181510e+03 | 6.181510e+03 | NaN | NaN | GHTLNFVAPFPLSIWACLPLRWQQAQAR |
| RIC8A | RIC8A\_HUMAN | Synembryn-A | 341, 341 | 329 | RIC8A | RIC8A\_HUMAN | Synembryn-A | 13C(-34.0809) 13C(125.0477) | NaN | X | NaN | NaN | X | NaN | 5.937707e+03 | NaN | 5.937707e+03 | NaN | ESVAPVLSVLTECAR |
| UBR1 | UBR1\_HUMAN | E3 ubiquitin-protein ligase UBR1 | 251 | 248 | UBR1 | UBR1\_HUMAN | E3 ubiquitin-protein ligase UBR1 | 4C(125.0477) | NaN | X | NaN | NaN | NaN | NaN | 5.548279e+03 | 5.548279e+03 | NaN | NaN | ALDCELAEAQLHTTAIDK |
| FGB | FIBB\_HUMAN | Fibrinogen beta chain | 437 | 437 | FGB | FIBB\_HUMAN | Fibrinogen beta chain | 1C(125.0477) | NaN | X | NaN | NaN | NaN | NaN | 5.501960e+03 | NaN | 5.501960e+03 | NaN | CHAANPNGR |
| WIPI2 | WIPI2\_HUMAN | WD repeat domain phosphoinositide-interacting protein 2 | 111 | 106 | WIPI2 | WIPI2\_HUMAN | WD repeat domain phosphoinositide-interacting protein 2 | 6C(125.0477) | NaN | X | NaN | NaN | NaN | NaN | 5.497119e+03 | 5.497119e+03 | NaN | NaN | KGTEICNYSYSNTILAVK |
| AHSG | FETUA\_HUMAN | Alpha-2-HS-glycoprotein | 132 | 132 | AHSG | FETUA\_HUMAN | Alpha-2-HS-glycoprotein | 1C(57.0215) | X | NaN | NaN | NaN | NaN | NaN | 5.496895e+03 | 5.496895e+03 | NaN | NaN | CDSSPDSAEDVR |
| UBE3A | UBE3A\_HUMAN | Ubiquitin-protein ligase E3A | 198 | 196 | UBE3A | UBE3A\_HUMAN | Ubiquitin-protein ligase E3A | 3C(125.0477) | NaN | X | NaN | NaN | NaN | NaN | 5.437889e+03 | 5.437889e+03 | NaN | NaN | AACSAAAMEEDSEASSSR |
| RAB21 | RAB21\_HUMAN | Ras-related protein Rab-21 | 128 | 120 | RAB21 | RAB21\_HUMAN | Ras-related protein Rab-21 | 9C(125.0477) | NaN | X | NaN | NaN | NaN | NaN | 5.380835e+03 | NaN | 5.380835e+03 | NaN | MLGNEICLCIVGNK |
| GCLC | GSH1\_HUMAN | Glutamate--cysteine ligase catalytic subunit | 142 | 130 | GCLC | GSH1\_HUMAN | Glutamate--cysteine ligase catalytic subunit | 13C(125.0477) | NaN | X | NaN | NaN | NaN | NaN | 5.244865e+03 | NaN | 5.244865e+03 | NaN | EATSILEENQALCTITSFPR |
| HBB | HBB\_HUMAN | Hemoglobin subunit beta | 94 | 68 | HBB | HBB\_HUMAN | Hemoglobin subunit beta | 27C(125.0477) | NaN | X | NaN | NaN | NaN | NaN | 5.173488e+03 | NaN | 5.173488e+03 | NaN | VLGAFSDGLAHLDNLKGTFATLSELHCDKLHVDPENFR |
| HNRNPA1 | ROA1\_HUMAN | Heterogeneous nuclear ribonucleoprotein A1 | 175 | 167 | HNRNPA1 | ROA1\_HUMAN | Heterogeneous nuclear ribonucleoprotein A1 | 9C(57.0215) | X | NaN | NaN | NaN | NaN | NaN | 5.111931e+03 | NaN | 5.111931e+03 | NaN | YHTVNGHNCEVR |
| NGLY1 | NGLY1\_HUMAN | Peptide-N(4)-(N-acetyl-beta-glucosaminyl)asparagine amidase | 644 | 635 | NGLY1 | NGLY1\_HUMAN | Peptide-N(4)-(N-acetyl-beta-glucosaminyl)asparagine amidase | 10C(125.0477) | NaN | X | NaN | NaN | NaN | NaN | 5.017611e+03 | 5.017611e+03 | NaN | NaN | QSLNDHEENCLEIIIK |
| DDB1 | DDB1\_HUMAN | DNA damage-binding protein 1 | 725 | 724 | DDB1 | DDB1\_HUMAN | DNA damage-binding protein 1 | 2C(125.0477) | NaN | X | NaN | NaN | NaN | NaN | 4.969502e+03 | 4.969502e+03 | NaN | NaN | ICYQEVSQCFGVLSSR |
| MAP4K4 | M4K4\_HUMAN | Mitogen-activated protein kinase kinase kinase kinase 4 | 496 | 478 | MAP4K4 | M4K4\_HUMAN | Mitogen-activated protein kinase kinase kinase kinase 4 | 19C(15.9949) | NaN | NaN | X | NaN | NaN | NaN | 4.913195e+03 | NaN | 4.913195e+03 | NaN | HLEVLQQQLLQEQAMLLECRWR |
| ALB | ALBU\_HUMAN | Albumin | 86 | 76 | ALB | ALBU\_HUMAN | Albumin | 11C(57.0215) | X | NaN | NaN | NaN | NaN | NaN | 4.771747e+03 | 4.771747e+03 | NaN | NaN | TCVADESAENCDK |
| USP9X | USP9X\_HUMAN | Probable ubiquitin carboxyl-terminal hydrolase FAF-X | 159 | 155 | USP9X | USP9X\_HUMAN | Probable ubiquitin carboxyl-terminal hydrolase FAF-X | 5C(125.0477) | NaN | X | NaN | NaN | NaN | NaN | 4.703253e+03 | NaN | 4.703253e+03 | NaN | LVELCVAK |
| ABRACL | ABRAL\_HUMAN | Costars family protein ABRACL | 39 | 39 | ABRACL | ABRAL\_HUMAN | Costars family protein ABRACL | 1C(125.0477) | NaN | X | NaN | NaN | NaN | NaN | 4.614652e+03 | 4.614652e+03 | NaN | NaN | CANLFEALVGTLK |
| RNH1 | RINI\_HUMAN | Ribonuclease inhibitor | 30 | 20 | RNH1 | RINI\_HUMAN | Ribonuclease inhibitor | 11C(125.0477) | NaN | X | NaN | NaN | NaN | NaN | 4.491818e+03 | NaN | 4.491818e+03 | NaN | WAELLPLLQQCQVVR |
| ISG15 | ISG15\_HUMAN | Ubiquitin-like protein ISG15 | 78 | 78 | ISG15 | ISG15\_HUMAN | Ubiquitin-like protein ISG15 | 1C(125.0477) | NaN | X | NaN | NaN | NaN | NaN | 4.354821e+03 | NaN | 4.354821e+03 | NaN | CDEPLSILVR |
| PPIA | PPIA\_HUMAN | Peptidyl-prolyl cis-trans isomerase A | 161 | 156 | PPIA | PPIA\_HUMAN | Peptidyl-prolyl cis-trans isomerase A | 6C(57.0215) | X | NaN | NaN | NaN | NaN | NaN | 4.344134e+03 | NaN | NaN | 4344.1340 | ITIADCGQLE |
| ANKRD13A | AN13A\_HUMAN | Ankyrin repeat domain-containing protein 13A | 146 | 140 | ANKRD13A | AN13A\_HUMAN | Ankyrin repeat domain-containing protein 13A | 7C(57.0215) | X | NaN | NaN | NaN | NaN | NaN | 4.218723e+03 | NaN | 4.218723e+03 | NaN | ICPNDVCR |
| WDR61 | WDR61\_HUMAN | WD repeat-containing protein 61 | 266 | 265 | WDR61 | WDR61\_HUMAN | WD repeat-containing protein 61 | 2C(125.0477) | NaN | X | NaN | NaN | NaN | NaN | 4.199070e+03 | 4.199070e+03 | NaN | NaN | TCVHTFFDHQDQVWGVK |
| EIF2B3 | EI2BG\_HUMAN | Translation initiation factor eIF-2B subunit gamma | 334 | 329 | EIF2B3 | EI2BG\_HUMAN | Translation initiation factor eIF-2B subunit gamma | 6C(125.0477) | NaN | X | NaN | NaN | NaN | NaN | 4.137141e+03 | 4.137141e+03 | NaN | NaN | LLSALCPEEPPVHSSAQIVSK |
| GPHN | GEPH\_HUMAN | Gephyrin | 154 | 150 | GPHN | GEPH\_HUMAN | Gephyrin | 5C(125.0477) | NaN | X | NaN | NaN | NaN | NaN | 4.133957e+03 | 4.133957e+03 | NaN | NaN | GSQECFQFILPALPHAIDLLR |
| VAC14 | VAC14\_HUMAN | Protein VAC14 homolog | 79 | 70 | VAC14 | VAC14\_HUMAN | Protein VAC14 homolog | 10C(15.9949) | NaN | NaN | X | NaN | NaN | NaN | 4.115890e+03 | 4.115890e+03 | NaN | NaN | KGGLIGLAACSIALGKDSGLYLK |
| NANP | NANP\_HUMAN | N-acylneuraminate-9-phosphatase | 242 | 219 | NANP | NANP\_HUMAN | N-acylneuraminate-9-phosphatase | 24C(125.0477) | NaN | X | NaN | NaN | NaN | NaN | 4.018916e+03 | 4.018916e+03 | NaN | NaN | SSPVPHYMVSSVLELPALLQSIDCK |
| EEF1G | EF1G\_HUMAN | Elongation factor 1-gamma | 194 | 190 | EEF1G | EF1G\_HUMAN | Elongation factor 1-gamma | 5C(57.0215) | X | NaN | NaN | NaN | NaN | NaN | 3.987357e+03 | NaN | NaN | 3987.3567 | WFLTCINQPQFR |
| KRT72 | K2C72\_HUMAN | Keratin, type II cytoskeletal 72 | 364 | 362 | KRT72 | K2C72\_HUMAN | Keratin, type II cytoskeletal 72 | 3C(15.9949) | NaN | NaN | X | NaN | NaN | NaN | 3.952530e+03 | NaN | NaN | 3952.5303 | KQCADLETAIADAEQR |
| STIP1 | STIP1\_HUMAN | Stress-induced-phosphoprotein 1 | 26 | 14 | STIP1 | STIP1\_HUMAN | Stress-induced-phosphoprotein 1 | 13C(125.0477) | NaN | X | NaN | NaN | NaN | NaN | 3.844616e+03 | NaN | 3.844616e+03 | NaN | ALSVGNIDDALQCYSEAIK |
| GMPR2 | GMPR2\_HUMAN | GMP reductase 2 | 224, 222 | 214 | GMPR2 | GMPR2\_HUMAN | GMP reductase 2 | 11C(57.0215) 9C(125.0477) | X | X | NaN | NaN | NaN | NaN | 3.733873e+03 | 3.733873e+03 | NaN | NaN | GHIISDGGCSCPGDVAK |
| HSPA1A | HS71A\_HUMAN | Heat shock 70 kDa protein 1A | 603 | 598 | HSPA1A | HS71A\_HUMAN | Heat shock 70 kDa protein 1A | 6C(57.0215) | X | NaN | NaN | NaN | NaN | NaN | 3.584678e+03 | NaN | 0.000000e+00 | 3584.6780 | ELEQVCNPIISGLYQGAGGPGPGGFGAQGPK |
| ANKRD54 | ANR54\_HUMAN | Ankyrin repeat domain-containing protein 54 | 230 | 220 | ANKRD54 | ANR54\_HUMAN | Ankyrin repeat domain-containing protein 54 | 11C(125.0477) | NaN | X | NaN | NaN | NaN | NaN | 3.505113e+03 | 3.505113e+03 | NaN | NaN | LNILQEGHAQCLEAVR |
| NUTF2 | NTF2\_HUMAN | Nuclear transport factor 2 | 38 | 27 | NUTF2 | NTF2\_HUMAN | Nuclear transport factor 2 | 12C(125.0477) | NaN | X | NaN | NaN | NaN | NaN | 3.490372e+03 | 3.490372e+03 | NaN | NaN | TQLGAIYIDASCLTWEGQQFQGK |
| KPNB1 | IMB1\_HUMAN | Importin subunit beta-1 | 585 | 575 | KPNB1 | IMB1\_HUMAN | Importin subunit beta-1 | 11C(125.0477) | NaN | X | NaN | NaN | NaN | NaN | 3.469045e+03 | 3.469045e+03 | 0.000000e+00 | NaN | IQFNDLQSLLCATLQNVLR |
| AP2A2 | AP2A2\_HUMAN | AP-2 complex subunit alpha-2 | 134 | 124 | AP2A2 | AP2A2\_HUMAN | AP-2 complex subunit alpha-2 | 11C(125.0477) | NaN | X | NaN | NaN | NaN | NaN | 3.419798e+03 | 3.419798e+03 | NaN | NaN | NPTFMGLALHCIASVGSR |
| ABCC5 | MRP5\_HUMAN | ATP-binding cassette sub-family C member 5 | 146 | 139 | ABCC5 | MRP5\_HUMAN | ATP-binding cassette sub-family C member 5 | 8C(31.9988) | NaN | NaN | NaN | X | NaN | NaN | 3.363122e+03 | 3.363122e+03 | NaN | NaN | HESSDVNCRR |
| MMS19 | MMS19\_HUMAN | MMS19 nucleotide excision repair protein homolog | 331 | 317 | MMS19 | MMS19\_HUMAN | MMS19 nucleotide excision repair protein homolog | 15C(125.0477) | NaN | X | NaN | NaN | NaN | NaN | 3.132783e+03 | 3.132783e+03 | NaN | NaN | VEAEGLAALHSLTACLSR |
| SPRR3 | SPRR3\_HUMAN | Small proline-rich protein 3 | 153 | 149 | SPRR3 | SPRR3\_HUMAN | Small proline-rich protein 3 | 5C(125.0477) | NaN | X | NaN | NaN | NaN | NaN | 3.054923e+03 | NaN | 3.054923e+03 | NaN | LPEPCPSTVTPGPAQQK |
| EPS15 | EPS15\_HUMAN | Epidermal growth factor receptor substrate 15 | 824 | 819 | EPS15 | EPS15\_HUMAN | Epidermal growth factor receptor substrate 15 | 6C(125.0477) | NaN | X | NaN | NaN | NaN | NaN | 2.953620e+03 | 2.953620e+03 | NaN | NaN | DPEIFCDPFTSATTTTNK |
| KPNB1 | IMB1\_HUMAN | Importin subunit beta-1 | 228, 223 | 216 | KPNB1 | IMB1\_HUMAN | Importin subunit beta-1 | 13C(125.0477) 8C(15.9949) | NaN | X | X | NaN | NaN | NaN | 2.940119e+03 | NaN | 2.940119e+03 | NaN | HFIMQVVCEATQCPDTR |
| BCAP29 | BAP29\_HUMAN | B-cell receptor-associated protein 29 | 171 | 164 | BCAP29 | BAP29\_HUMAN | B-cell receptor-associated protein 29 | 8C(125.0477) | NaN | X | NaN | NaN | NaN | NaN | 2.852821e+03 | 2.852821e+03 | NaN | NaN | SHGKDEECVLEAENK |
| KPNB1 | IMB1\_HUMAN | Importin subunit beta-1 | 158 | 148 | KPNB1 | IMB1\_HUMAN | Importin subunit beta-1 | 11C(125.0477) | NaN | X | NaN | NaN | NaN | NaN | 2.807593e+03 | 2.807593e+03 | NaN | NaN | ESTLEAIGYICQDIDPEQLQDK |
| SPTA1 | SPTA1\_HUMAN | Spectrin alpha chain, erythrocytic 1 | 1568, 1571 | 1552 | SPTA1 | SPTA1\_HUMAN | Spectrin alpha chain, erythrocytic 1 | 17C(-34.0809) 20C(31.9988) | NaN | NaN | NaN | X | X | NaN | 2.614136e+03 | 2.614136e+03 | NaN | NaN | SEQVHGVINLGNSLIECSACDGNEEAMKEQLEQLK |
| ESD | ESTD\_HUMAN | S-formylglutathione hydrolase | 176 | 168 | ESD | ESTD\_HUMAN | S-formylglutathione hydrolase | 9C(-34.0809) | NaN | NaN | NaN | NaN | X | NaN | 2.561631e+03 | NaN | 2.561631e+03 | NaN | SVSAFAPICNPVLCPWGK |
| USP5 | UBP5\_HUMAN | Ubiquitin carboxyl-terminal hydrolase 5 | 219 | 206 | USP5 | UBP5\_HUMAN | Ubiquitin carboxyl-terminal hydrolase 5 | 14C(125.0477) | NaN | X | NaN | NaN | NaN | NaN | 2.493089e+03 | 2.493089e+03 | NaN | NaN | ENLWLNLTDGSILCGR |
| CPPED1 | CPPED\_HUMAN | Serine/threonine-protein phosphatase CPPED1 | 88 | 84 | CPPED1 | CPPED\_HUMAN | Serine/threonine-protein phosphatase CPPED1 | 5C(125.0477) | NaN | X | NaN | NaN | NaN | NaN | 2.428635e+03 | 2.428635e+03 | NaN | NaN | FFVLCGDLIHAMPGKPWR |
| MOB2 | MOB2\_HUMAN | MOB kinase activator 2 | 148 | 147 | MOB2 | MOB2\_HUMAN | MOB kinase activator 2 | 2C(57.0215) | X | NaN | NaN | NaN | NaN | NaN | 2.234993e+03 | 2.234993e+03 | NaN | NaN | ICRHLFHVLAHIYWAHFK |
| RCC2 | RCC2\_HUMAN | Protein RCC2 | 158 | 152 | RCC2 | RCC2\_HUMAN | Protein RCC2 | 7C(125.0477) | NaN | X | NaN | NaN | NaN | NaN | 2.095001e+03 | 2.095001e+03 | NaN | NaN | TVVSGSCAAHSLLITTEGK |
| CCT8 | TCPQ\_HUMAN | T-complex protein 1 subunit theta | 244 | 239 | CCT8 | TCPQ\_HUMAN | T-complex protein 1 subunit theta | 6C(125.0477) | NaN | X | NaN | NaN | NaN | NaN | 2.088405e+03 | NaN | 2.088405e+03 | NaN | IAVYSCPFDGMITETK |
| KPNA6 | IMA7\_HUMAN | Importin subunit alpha-7 | 357 | 355 | KPNA6 | IMA7\_HUMAN | Importin subunit alpha-7 | 3C(125.0477) | NaN | X | NaN | NaN | NaN | NaN | 1.627340e+03 | 1.627340e+03 | NaN | NaN | EACWTISNITAGNR |
| BABAM1 | BABA1\_HUMAN | BRISC and BRCA1-A complex member 1 | 294 | 280 | BABAM1 | BABA1\_HUMAN | BRISC and BRCA1-A complex member 1 | 15C(125.0477) | NaN | X | NaN | NaN | NaN | NaN | 1.463308e+03 | 1.463308e+03 | NaN | NaN | YEVALAGPALELHNCMAK |
| IPO5 | IPO5\_HUMAN | Importin-5 | 473 | 458 | IPO5 | IPO5\_HUMAN | Importin-5 | 16C(125.0477) | NaN | X | NaN | NaN | NaN | NaN | 1.389215e+03 | 1.389215e+03 | NaN | NaN | VQAHAAAALINFTEDCPK |
| EHBP1L1 | EH1L1\_HUMAN | EH domain-binding protein 1-like protein 1 | 1141 | 1139 | EHBP1L1 | EH1L1\_HUMAN | EH domain-binding protein 1-like protein 1 | 3C(125.0477) | NaN | X | NaN | NaN | NaN | NaN | 0.000000e+00 | 0.000000e+00 | NaN | NaN | AFCTGQELQLVQLEGGGGAGTYR |
| ATP2A2 | AT2A2\_HUMAN | Sarcoplasmic/endoplasmic reticulum calcium ATPase 2 | 669 | 651 | ATP2A2 | AT2A2\_HUMAN | Sarcoplasmic/endoplasmic reticulum calcium ATPase 2 | 19C(125.0477) | NaN | X | NaN | NaN | NaN | NaN | 0.000000e+00 | 0.000000e+00 | NaN | NaN | AFTGREFDELNPSAQRDACLNAR |
| NAPRT | PNCB\_HUMAN | Nicotinate phosphoribosyltransferase | 140 | 108 | NAPRT | PNCB\_HUMAN | Nicotinate phosphoribosyltransferase | 33C(125.0477) | NaN | X | NaN | NaN | NaN | NaN | 0.000000e+00 | 0.000000e+00 | NaN | NaN | ALPEGSLAFPGVPLLQVSGPLLVVQLLETPLLCLVSYASLVATNAAR |
| TTPAL | TTPAL\_HUMAN | Alpha-tocopherol transfer protein-like | 105 | 95 | TTPAL | TTPAL\_HUMAN | Alpha-tocopherol transfer protein-like | 11C(125.0477) | NaN | X | NaN | NaN | NaN | NaN | 0.000000e+00 | 0.000000e+00 | NaN | NaN | ALQLLVNYHSCR |
| ABCA7 | ABCA7\_HUMAN | Phospholipid-transporting ATPase ABCA7 | 674, 659 | 653 | ABCA7 | ABCA7\_HUMAN | Phospholipid-transporting ATPase ABCA7 | 22C(15.9949) 7C(125.0477) | NaN | X | X | NaN | NaN | NaN | 0.000000e+00 | 0.000000e+00 | NaN | NaN | ANLAAACGGLAYFSLYLPYVLCVAWR |
| CA1 | CAH1\_HUMAN | Carbonic anhydrase 1 | 213 | 175 | CA1 | CAH1\_HUMAN | Carbonic anhydrase 1 | 39C(125.0477) | NaN | X | NaN | NaN | NaN | NaN | 0.000000e+00 | 0.000000e+00 | NaN | NaN | APFTNFDPSTLLPSSLDFWTYPGSLTHPPLYESVTWIICK |
| SPTA1 | SPTA1\_HUMAN | Spectrin alpha chain, erythrocytic 1 | 2072 | 2052 | SPTA1 | SPTA1\_HUMAN | Spectrin alpha chain, erythrocytic 1 | 21C(31.9988) | NaN | NaN | NaN | X | NaN | NaN | 0.000000e+00 | 0.000000e+00 | NaN | NaN | ASALNNWCEKMEENLSEPVHCVSLNEIR |
| ZZEF1 | ZZEF1\_HUMAN | Zinc finger ZZ-type and EF-hand domain-containing protein 1 | 2084 | 2072 | ZZEF1 | ZZEF1\_HUMAN | Zinc finger ZZ-type and EF-hand domain-containing protein 1 | 13C(125.0477) | NaN | X | NaN | NaN | NaN | NaN | 0.000000e+00 | 0.000000e+00 | NaN | NaN | AVTPSPEQVFAECSQK |
| ICAM4 | ICAM4\_HUMAN | Intercellular adhesion molecule 4 | 117, 113 | 106 | ICAM4 | ICAM4\_HUMAN | Intercellular adhesion molecule 4 | 12C(31.9988) 8C(15.9949) | NaN | NaN | X | X | NaN | NaN | 0.000000e+00 | 0.000000e+00 | NaN | NaN | AWSSLAHCLVTCAGK |
| ATP6V1A | VATA\_HUMAN | V-type proton ATPase catalytic subunit A | 394 | 394 | ATP6V1A | VATA\_HUMAN | V-type proton ATPase catalytic subunit A | 1C(125.0477) | NaN | X | NaN | NaN | NaN | NaN | 0.000000e+00 | 0.000000e+00 | NaN | NaN | CLGNPER |
| RHCE | RHCE\_HUMAN | Blood group Rh(CE) polypeptide | 12 | 12 | RHCE | RHCE\_HUMAN | Blood group Rh(CE) polypeptide | 1C(125.0477) | NaN | X | NaN | NaN | NaN | NaN | 0.000000e+00 | 0.000000e+00 | NaN | NaN | CLPLWALTLEAALILLFYFFTHYDASLEDQK |
| HDAC6 | HDAC6\_HUMAN | Histone deacetylase 6 | 523 | 523 | HDAC6 | HDAC6\_HUMAN | Histone deacetylase 6 | 1C(15.9949) | NaN | NaN | X | NaN | NaN | NaN | 0.000000e+00 | 0.000000e+00 | NaN | NaN | CLTLTPRPATEAELLTCHSAEYVGHLR |
| STIP1 | STIP1\_HUMAN | Stress-induced-phosphoprotein 1 | 471 | 471 | STIP1 | STIP1\_HUMAN | Stress-induced-phosphoprotein 1 | 1C(125.0477) | NaN | X | NaN | NaN | NaN | NaN | 0.000000e+00 | 0.000000e+00 | NaN | NaN | CMMAQYNRHDSPEDVKR |
| UBR4 | UBR4\_HUMAN | E3 ubiquitin-protein ligase UBR4 | 4916, 4935 | 4916 | UBR4 | UBR4\_HUMAN | E3 ubiquitin-protein ligase UBR4 | 1C(57.0215) 20C(125.0477) | X | X | NaN | NaN | NaN | NaN | 0.000000e+00 | 0.000000e+00 | NaN | NaN | CNGLLPVWGPHVPESAFATCLAR |
| NAMPT | NAMPT\_HUMAN | Nicotinamide phosphoribosyltransferase | 401 | 401 | NAMPT | NAMPT\_HUMAN | Nicotinamide phosphoribosyltransferase | 1C(125.0477) | NaN | X | NaN | NaN | NaN | NaN | 0.000000e+00 | 0.000000e+00 | NaN | NaN | CSYVVTNGLGINVFKDPVADPNKR |
| ADD3 | ADDG\_HUMAN | Gamma-adducin | 228 | 228 | ADD3 | ADDG\_HUMAN | Gamma-adducin | 1C(125.0477) | NaN | X | NaN | NaN | NaN | NaN | 0.000000e+00 | 0.000000e+00 | NaN | NaN | CVIHIHTLATAAVSSMK |
| FLNA | FLNA\_HUMAN | Filamin-A | 1912, 1920 | 1892 | FLNA | FLNA\_HUMAN | Filamin-A | 21C(-34.0809) 29C(31.9988) | NaN | NaN | NaN | X | X | NaN | 0.000000e+00 | 0.000000e+00 | NaN | NaN | DAGEGGLSLAIEGPSKAEISCTDNQDGTCSVSYLPVLPGDYSILVK |
| TGM2 | TGM2\_HUMAN | Protein-glutamine gamma-glutamyltransferase 2 | 554 | 553 | TGM2 | TGM2\_HUMAN | Protein-glutamine gamma-glutamyltransferase 2 | 2C(125.0477) | NaN | X | NaN | NaN | NaN | NaN | 0.000000e+00 | 0.000000e+00 | NaN | NaN | DCLTESNLIKVR |
| CUL4A | CUL4A\_HUMAN | Cullin-4A | 379 | 370 | CUL4A | CUL4A\_HUMAN | Cullin-4A | 10C(125.0477) | NaN | X | NaN | NaN | NaN | NaN | 0.000000e+00 | 0.000000e+00 | NaN | NaN | DKVDHVIEVCFQK |
| RARS1 | SYRC\_HUMAN | Arginine--tRNA ligase, cytoplasmic | 502 | 483 | RARS1 | SYRC\_HUMAN | Arginine--tRNA ligase, cytoplasmic | 20C(125.0477) | NaN | X | NaN | NaN | NaN | NaN | 0.000000e+00 | 0.000000e+00 | NaN | NaN | DKVLTAEELNAAQTSVAYGCIK |
| TATDN1 | TATD1\_HUMAN | Deoxyribonuclease TATDN1 | 168 | 166 | TATDN1 | TATD1\_HUMAN | Deoxyribonuclease TATDN1 | 3C(125.0477) | NaN | X | NaN | NaN | NaN | NaN | 0.000000e+00 | 0.000000e+00 | NaN | NaN | DRCVGGVVHSFDGTK |
| PFAS | PUR4\_HUMAN | Phosphoribosylformylglycinamidine synthase | 336 | 333 | PFAS | PUR4\_HUMAN | Phosphoribosylformylglycinamidine synthase | 4C(125.0477) | NaN | X | NaN | NaN | NaN | NaN | 0.000000e+00 | 0.000000e+00 | NaN | NaN | DVQCTGR |
| PCDHGC4 | PCDGL\_HUMAN | Protocadherin gamma-C4 | 102, 96 | 93 | PCDHGC4 | PCDGL\_HUMAN | Protocadherin gamma-C4 | 10C(-34.0809) 4C(31.9988) | NaN | NaN | NaN | X | X | NaN | 0.000000e+00 | 0.000000e+00 | NaN | NaN | EALCGLSASCIVPLEFVTEGPLEMYR |
| STAT5A | STA5A\_HUMAN | Signal transducer and activator of transcription 5A | 126 | 122 | STAT5A | STA5A\_HUMAN | Signal transducer and activator of transcription 5A | 5C(125.0477) | NaN | X | NaN | NaN | NaN | NaN | 0.000000e+00 | 0.000000e+00 | NaN | NaN | EANNCSSPAGILVDAMSQK |
| CAPZA2 | CAZA2\_HUMAN | F-actin-capping protein subunit alpha-2 | 111 | 104 | CAPZA2 | CAZA2\_HUMAN | F-actin-capping protein subunit alpha-2 | 8C(31.9988) | NaN | NaN | NaN | X | NaN | NaN | 0.000000e+00 | 0.000000e+00 | NaN | NaN | EATDPRPCEVENAVESWR |
| EIF3G | EIF3G\_HUMAN | Eukaryotic translation initiation factor 3 subunit G | 139 | 135 | EIF3G | EIF3G\_HUMAN | Eukaryotic translation initiation factor 3 subunit G | 5C(125.0477) | NaN | X | NaN | NaN | NaN | NaN | 0.000000e+00 | 0.000000e+00 | NaN | NaN | EDLNCQEEEDPMNK |
| CAPZA2 | CAZA2\_HUMAN | F-actin-capping protein subunit alpha-2 | 141 | 134 | CAPZA2 | CAZA2\_HUMAN | F-actin-capping protein subunit alpha-2 | 8C(125.0477) | NaN | X | NaN | NaN | NaN | NaN | 0.000000e+00 | 0.000000e+00 | NaN | NaN | EHYPNGVCTVYGK |
| RNH1 | RINI\_HUMAN | Ribonuclease inhibitor | 409 | 402 | RNH1 | RINI\_HUMAN | Ribonuclease inhibitor | 8C(125.0477) | NaN | X | NaN | NaN | NaN | NaN | 0.000000e+00 | 0.000000e+00 | NaN | NaN | ELDLSNNCLGDAGILQLVESVR |
| PNP | PNPH\_HUMAN | Purine nucleoside phosphorylase | 206 | 186 | PNP | PNPH\_HUMAN | Purine nucleoside phosphorylase | 21C(125.0477) | NaN | X | NaN | NaN | NaN | NaN | 0.000000e+00 | 0.000000e+00 | NaN | NaN | ELQEGTYVMVAGPSFETVAECR |
| UBA3 | UBA3\_HUMAN | NEDD8-activating enzyme E1 catalytic subunit | 364 | 362 | UBA3 | UBA3\_HUMAN | NEDD8-activating enzyme E1 catalytic subunit | 3C(125.0477) | NaN | X | NaN | NaN | NaN | NaN | 0.000000e+00 | 0.000000e+00 | NaN | NaN | ENCPACSQLPQNIQFSPSAK |
| YKT6 | YKT6\_HUMAN | Synaptobrevin homolog YKT6 | 66 | 61 | YKT6 | YKT6\_HUMAN | Synaptobrevin homolog YKT6 | 6C(125.0477) | NaN | X | NaN | NaN | NaN | NaN | 0.000000e+00 | 0.000000e+00 | NaN | NaN | EQDYLCHVYVR |
| SPTB | SPTB1\_HUMAN | Spectrin beta chain, erythrocytic | 1892 | 1878 | SPTB | SPTB1\_HUMAN | Spectrin beta chain, erythrocytic | 15C(-34.0809) | NaN | NaN | NaN | NaN | X | NaN | 0.000000e+00 | 0.000000e+00 | NaN | NaN | EQEVSAAWQALLDACAGRR |
| LCMT1 | LCMT1\_HUMAN | Leucine carboxyl methyltransferase 1 | 13 | 7 | LCMT1 | LCMT1\_HUMAN | Leucine carboxyl methyltransferase 1 | 7C(57.0215) | X | NaN | NaN | NaN | NaN | NaN | 0.000000e+00 | 0.000000e+00 | NaN | NaN | ESSITSCCSTSSCDADDEGVR |
| AIFM1 | AIFM1\_HUMAN | Apoptosis-inducing factor 1, mitochondrial | 317 | 299 | AIFM1 | AIFM1\_HUMAN | Apoptosis-inducing factor 1, mitochondrial | 19C(-34.0809) | NaN | NaN | NaN | NaN | X | NaN | 0.000000e+00 | 0.000000e+00 | NaN | NaN | EVKSITIIGGGFLGSELACALGR |
| OGT | OGT1\_HUMAN | UDP-N-acetylglucosamine--peptide N-acetylglucosaminyltransferase 110 kDa subunit | 590 | 586 | OGT | OGT1\_HUMAN | UDP-N-acetylglucosamine--peptide N-acetylglucosaminyltransferase 110 kDa subunit | 5C(15.9949) | NaN | NaN | X | NaN | NaN | NaN | 0.000000e+00 | 0.000000e+00 | NaN | NaN | FEVFCYALSPDDGTNFRVK |
| USP9X | USP9X\_HUMAN | Probable ubiquitin carboxyl-terminal hydrolase FAF-X | 1451 | 1447 | USP9X | USP9X\_HUMAN | Probable ubiquitin carboxyl-terminal hydrolase FAF-X | 5C(125.0477) | NaN | X | NaN | NaN | NaN | NaN | 0.000000e+00 | 0.000000e+00 | NaN | NaN | FHIGCEK |
| OLA1 | OLA1\_HUMAN | Obg-like ATPase 1 | 117 | 99 | OLA1 | OLA1\_HUMAN | Obg-like ATPase 1 | 19C(125.0477) | NaN | X | NaN | NaN | NaN | NaN | 0.000000e+00 | 0.000000e+00 | NaN | NaN | GAHNGQGLGNAFLSHISACDGIFHLTR |
| PWP1 | PWP1\_HUMAN | Periodic tryptophan protein 1 homolog | 400 | 399 | PWP1 | PWP1\_HUMAN | Periodic tryptophan protein 1 homolog | 2C(125.0477) | NaN | X | NaN | NaN | NaN | NaN | 0.000000e+00 | 0.000000e+00 | NaN | NaN | GCLVTASADKYVK |
| QARS1 | SYQ\_HUMAN | Glutamine--tRNA ligase | 536 | 526 | QARS1 | SYQ\_HUMAN | Glutamine--tRNA ligase | 11C(31.9988) | NaN | NaN | NaN | X | NaN | NaN | 0.000000e+00 | 0.000000e+00 | NaN | NaN | GFPPEAINNFCAR |
| PI4KA | PI4KA\_HUMAN | Phosphatidylinositol 4-kinase alpha | 19 | 8 | PI4KA | PI4KA\_HUMAN | Phosphatidylinositol 4-kinase alpha | 12C(125.0477) | NaN | X | NaN | NaN | NaN | NaN | 0.000000e+00 | 0.000000e+00 | NaN | NaN | GGGGGGGGGGGCSGSGSSASR |
| DCAF8 | DCAF8\_HUMAN | DDB1- and CUL4-associated factor 8 | 483 | 478 | DCAF8 | DCAF8\_HUMAN | DDB1- and CUL4-associated factor 8 | 6C(31.9988) | NaN | NaN | NaN | X | NaN | NaN | 0.000000e+00 | 0.000000e+00 | NaN | NaN | GGVVNCLEPHPHLPVLATSGLDHDVK |
| ERP44 | ERP44\_HUMAN | Endoplasmic reticulum resident protein 44 | 301 | 292 | ERP44 | ERP44\_HUMAN | Endoplasmic reticulum resident protein 44 | 10C(125.0477) | NaN | X | NaN | NaN | NaN | NaN | 0.000000e+00 | 0.000000e+00 | NaN | NaN | GTINFLHADCDKFR |
| PEPD | PEPD\_HUMAN | Xaa-Pro dipeptidase | 158, 183 | 149 | PEPD | PEPD\_HUMAN | Xaa-Pro dipeptidase | 10C(15.9949) 35C(125.0477) | NaN | X | X | NaN | NaN | NaN | 0.000000e+00 | 0.000000e+00 | NaN | NaN | GVNTDSGSVCREASFDGISKFEVNNTILHPEIVECR |
| ALDH1A1 | AL1A1\_HUMAN | Aldehyde dehydrogenase 1A1 | 163 | 157 | ALDH1A1 | AL1A1\_HUMAN | Aldehyde dehydrogenase 1A1 | 7C(125.0477) | NaN | X | NaN | NaN | NaN | NaN | 0.000000e+00 | 0.000000e+00 | NaN | NaN | HEPIGVCGQIIPWNFPLVMLIWK |
| RANBP10 | RBP10\_HUMAN | Ran-binding protein 10 | 389 | 372 | RANBP10 | RBP10\_HUMAN | Ran-binding protein 10 | 18C(125.0477) | NaN | X | NaN | NaN | NaN | NaN | 0.000000e+00 | 0.000000e+00 | NaN | NaN | HGPSSSHMHNTGADSPSCSNGVASTK |
| PEPD | PEPD\_HUMAN | Xaa-Pro dipeptidase | 243, 245 | 238 | PEPD | PEPD\_HUMAN | Xaa-Pro dipeptidase | 6C(-34.0809) 8C(31.9988) | NaN | NaN | NaN | X | X | NaN | 0.000000e+00 | 0.000000e+00 | NaN | NaN | HSSYTCICGSGENSAVLHYGHAGAPNDR |
| ANK1 | ANK1\_HUMAN | Ankyrin-1 | 1529 | 1499 | ANK1 | ANK1\_HUMAN | Ankyrin-1 | 31C(125.0477) | NaN | X | NaN | NaN | NaN | NaN | 0.000000e+00 | 0.000000e+00 | NaN | NaN | HTDRDYSLSPSQMNGYSSLQDELLSPASLGCALSSPLR |
| KRT10 | K1C10\_HUMAN | Keratin, type I cytoskeletal 10 | 25 | 10 | KRT10 | K1C10\_HUMAN | Keratin, type I cytoskeletal 10 | 16C(125.0477) | NaN | X | NaN | NaN | NaN | NaN | 0.000000e+00 | 0.000000e+00 | NaN | NaN | HYSSSRSGGGGGGGGCGGGGGVSSLR |
| ARF6 | ARF6\_HUMAN | ADP-ribosylation factor 6 | 90 | 76 | ARF6 | ARF6\_HUMAN | ADP-ribosylation factor 6 | 15C(125.0477) | NaN | X | NaN | NaN | NaN | NaN | 0.000000e+00 | 0.000000e+00 | NaN | NaN | HYYTGTQGLIFVVDCADRDRIDEAR |
| PSMA7 | PSA7\_HUMAN | Proteasome subunit alpha type-7 | 63, 63, 70, 70 | 62 | PSMA7 | PSA7\_HUMAN | Proteasome subunit alpha type-7 | 2C(125.0477) 2C(15.9949) 9C(125.0477) 9C(15.9949) | NaN | X | X | NaN | NaN | NaN | 0.000000e+00 | 0.000000e+00 | NaN | NaN | ICALDDNVCMAFAGLTADAR |
| PDE12 | PDE12\_HUMAN | 2',5'-phosphodiesterase 12 | 451 | 450 | PDE12 | PDE12\_HUMAN | 2',5'-phosphodiesterase 12 | 2C(125.0477) | NaN | X | NaN | NaN | NaN | NaN | 0.000000e+00 | 0.000000e+00 | NaN | NaN | ICVANTHLYWHPK |
| RBM42 | RBM42\_HUMAN | RNA-binding protein 42 | 385 | 383 | RBM42 | RBM42\_HUMAN | RNA-binding protein 42 | 3C(125.0477) | NaN | X | NaN | NaN | NaN | NaN | 0.000000e+00 | 0.000000e+00 | NaN | NaN | IFCGDLGNEVNDDILAR |
| SEPTIN11 | SEP11\_HUMAN | Septin-11 | 150 | 147 | SEPTIN11 | SEP11\_HUMAN | Septin-11 | 4C(15.9949) | NaN | NaN | X | NaN | NaN | NaN | 0.000000e+00 | 0.000000e+00 | NaN | NaN | IHACLYFIAPTGHSLK |
| WNK1 | WNK1\_HUMAN | Serine/threonine-protein kinase WNK1 | 1135 | 1122 | WNK1 | WNK1\_HUMAN | Serine/threonine-protein kinase WNK1 | 14C(125.0477) | NaN | X | NaN | NaN | NaN | NaN | 0.000000e+00 | 0.000000e+00 | NaN | NaN | ILNVSNKGDRVVECQLETHNR |
| RNH1 | RINI\_HUMAN | Ribonuclease inhibitor | 96, 102 | 87 | RNH1 | RINI\_HUMAN | Ribonuclease inhibitor | 10C(-34.0809) 16C(57.0215) | X | NaN | NaN | NaN | X | NaN | 0.000000e+00 | 0.000000e+00 | NaN | NaN | IQKLSLQNCCLTGAGCGVLSSTLR |
| TKT | TKT\_HUMAN | Transketolase | 41, 42 | 24 | TKT | TKT\_HUMAN | Transketolase | 18C(-34.0809) 19C(31.9988) | NaN | NaN | NaN | X | X | NaN | 0.000000e+00 | 0.000000e+00 | NaN | NaN | ISSIQATTAAGSGHPTSCCSAAEIMAVLFFHTMR |
| LDHA | LDHA\_HUMAN | L-lactate dehydrogenase A chain | 35 | 23 | LDHA | LDHA\_HUMAN | L-lactate dehydrogenase A chain | 13C(125.0477) | NaN | X | NaN | NaN | NaN | NaN | 0.000000e+00 | 0.000000e+00 | NaN | NaN | ITVVGVGAVGMACAISILMK |
| TMEM225 | TM225\_HUMAN | Transmembrane protein 225 | 154, 162 | 137 | TMEM225 | TM225\_HUMAN | Transmembrane protein 225 | 18C(125.0477) 26C(31.9988) | NaN | X | NaN | X | NaN | NaN | 0.000000e+00 | 0.000000e+00 | NaN | NaN | ITWIMYTAYLNVFFLSVCGVLSLLECK |
| PPP1CA | PP1A\_HUMAN | Serine/threonine-protein phosphatase PP1-alpha catalytic subunit | 140 | 133 | PPP1CA | PP1A\_HUMAN | Serine/threonine-protein phosphatase PP1-alpha catalytic subunit | 8C(125.0477) | NaN | X | NaN | NaN | NaN | NaN | 0.000000e+00 | 0.000000e+00 | NaN | NaN | IYGFYDECKR |
| GMDS | GMDS\_HUMAN | GDP-mannose 4,6 dehydratase | 237 | 230 | GMDS | GMDS\_HUMAN | GDP-mannose 4,6 dehydratase | 8C(125.0477) | NaN | X | NaN | NaN | NaN | NaN | 0.000000e+00 | 0.000000e+00 | NaN | NaN | IYLGQLECFSLGNLDAK |
| ACIN1 | ACINU\_HUMAN | Apoptotic chromatin condensation inducer in the nucleus | 733 | 732 | ACIN1 | ACINU\_HUMAN | Apoptotic chromatin condensation inducer in the nucleus | 2C(125.0477) | NaN | X | NaN | NaN | NaN | NaN | 0.000000e+00 | 0.000000e+00 | NaN | NaN | KCEAEEAEPPAATQPQTSETQTSHLPESER |
| PSMA7 | PSA7\_HUMAN | Proteasome subunit alpha type-7 | 70, 63 | 61 | PSMA7 | PSA7\_HUMAN | Proteasome subunit alpha type-7 | 10C(31.9988) 3C(125.0477) | NaN | X | NaN | X | NaN | NaN | 0.000000e+00 | 0.000000e+00 | NaN | NaN | KICALDDNVCMAFAGLTADAR |
| PHKA2 | KPB2\_HUMAN | Phosphorylase b kinase regulatory subunit alpha, liver isoform | 841 | 832 | PHKA2 | KPB2\_HUMAN | Phosphorylase b kinase regulatory subunit alpha, liver isoform | 10C(125.0477) | NaN | X | NaN | NaN | NaN | NaN | 0.000000e+00 | 0.000000e+00 | NaN | NaN | KKVEVLAEACTDLLSHQK |
| SEC14L2 | S14L2\_HUMAN | SEC14-like protein 2 | 155 | 145 | SEC14L2 | S14L2\_HUMAN | SEC14-like protein 2 | 11C(125.0477) | NaN | X | NaN | NaN | NaN | NaN | 0.000000e+00 | 0.000000e+00 | NaN | NaN | KVETITIIYDCEGLGLK |
| DHX29 | DHX29\_HUMAN | ATP-dependent RNA helicase DHX29 | 1245 | 1243 | DHX29 | DHX29\_HUMAN | ATP-dependent RNA helicase DHX29 | 3C(125.0477) | NaN | X | NaN | NaN | NaN | NaN | 0.000000e+00 | 0.000000e+00 | NaN | NaN | LACIVETAQGKAQVHPSSVNR |
| ACAN | PGCA\_HUMAN | Aggrecan core protein | 297 | 280 | ACAN | PGCA\_HUMAN | Aggrecan core protein | 18C(57.0215) | X | NaN | NaN | NaN | NaN | NaN | 0.000000e+00 | 0.000000e+00 | 0.000000e+00 | NaN | LATTGQLYLAWQAGMDMCSAGWLADR |
| HECTD4 | HECD4\_HUMAN | Probable E3 ubiquitin-protein ligase HECTD4 | 1299 | 1298 | HECTD4 | HECD4\_HUMAN | Probable E3 ubiquitin-protein ligase HECTD4 | 2C(125.0477) | NaN | X | NaN | NaN | NaN | NaN | 0.000000e+00 | 0.000000e+00 | NaN | NaN | LCSLSNQTESSSSEK |
| WIPI2 | WIPI2\_HUMAN | WD repeat domain phosphoinositide-interacting protein 2 | 393 | 375 | WIPI2 | WIPI2\_HUMAN | WD repeat domain phosphoinositide-interacting protein 2 | 19C(125.0477) | NaN | X | NaN | NaN | NaN | NaN | 0.000000e+00 | 0.000000e+00 | NaN | NaN | LDGSLETTNEILDSASHDCPLVTQTYGAAAGK |
| CDK6 | CDK6\_HUMAN | Cyclin-dependent kinase 6 | 83 | 79 | CDK6 | CDK6\_HUMAN | Cyclin-dependent kinase 6 | 5C(125.0477) | NaN | X | NaN | NaN | NaN | NaN | 0.000000e+00 | 0.000000e+00 | NaN | NaN | LFDVCTVSR |
| GAK | GAK\_HUMAN | Cyclin-G-associated kinase | 544 | 528 | GAK | GAK\_HUMAN | Cyclin-G-associated kinase | 17C(57.0215) | X | NaN | NaN | NaN | NaN | NaN | 0.000000e+00 | 0.000000e+00 | NaN | NaN | LFSTAEAAVYMFSMKRCPPGIWPSHK |
| SMAD7 | SMAD7\_HUMAN | Mothers against decapentaplegic homolog 7 | 160 | 136 | SMAD7 | SMAD7\_HUMAN | Mothers against decapentaplegic homolog 7 | 25C(125.0477) | NaN | X | NaN | NaN | NaN | NaN | 0.000000e+00 | 0.000000e+00 | NaN | NaN | LGPGAPAGAQPAQPPSSYSLPLLLCKVFR |
| HPS1 | HPS1\_HUMAN | Hermansky-Pudlak syndrome 1 protein | 183, 180 | 174 | HPS1 | HPS1\_HUMAN | Hermansky-Pudlak syndrome 1 protein | 10C(-34.0809) 7C(31.9988) | NaN | NaN | NaN | X | X | NaN | 0.000000e+00 | 0.000000e+00 | NaN | NaN | LIHPQLCELCIEALER |
| COMT | COMT\_HUMAN | Catechol O-methyltransferase | 145 | 136 | COMT | COMT\_HUMAN | Catechol O-methyltransferase | 10C(125.0477) | NaN | X | NaN | NaN | NaN | NaN | 0.000000e+00 | 0.000000e+00 | NaN | NaN | LITIEINPDCAAITQR |
| RBM38 | RBM38\_HUMAN | RNA-binding protein 38 | 8 | 2 | RBM38 | RBM38\_HUMAN | RNA-binding protein 38 | 7C(57.0215) | X | NaN | NaN | NaN | NaN | NaN | 0.000000e+00 | 0.000000e+00 | NaN | NaN | LLQPAPCAPSAGFPRPLAAPGAMHGSQKDTTFTK |
| SEC24C | SC24C\_HUMAN | Protein transport protein Sec24C | 809, 816 | 799 | SEC24C | SC24C\_HUMAN | Protein transport protein Sec24C | 11C(125.0477) 18C(31.9988) | NaN | X | NaN | X | NaN | NaN | 0.000000e+00 | 0.000000e+00 | NaN | NaN | LNEESGALLQCALLYTSCAGQRR |
| PGM3 | AGM1\_HUMAN | Phosphoacetylglucosamine mutase | 249 | 245 | PGM3 | AGM1\_HUMAN | Phosphoacetylglucosamine mutase | 5C(125.0477) | NaN | X | NaN | NaN | NaN | NaN | 0.000000e+00 | 0.000000e+00 | NaN | NaN | LNHLCGADFVK |
| LAP3 | AMPL\_HUMAN | Cytosol aminopeptidase | 335 | 322 | LAP3 | AMPL\_HUMAN | Cytosol aminopeptidase | 14C(-34.0809) | NaN | NaN | NaN | NaN | X | NaN | 0.000000e+00 | 0.000000e+00 | NaN | NaN | LNLPINIIGLAPLCENMPSGKANKPGDVVR |
| EPRS1 | SYEP\_HUMAN | Bifunctional glutamate/proline--tRNA ligase | 1148 | 1144 | EPRS1 | SYEP\_HUMAN | Bifunctional glutamate/proline--tRNA ligase | 5C(125.0477) | NaN | X | NaN | NaN | NaN | NaN | 0.000000e+00 | 0.000000e+00 | NaN | NaN | LNQWCNVVR |
| PAICS | PUR6\_HUMAN | Multifunctional protein ADE2 | 374 | 368 | PAICS | PUR6\_HUMAN | Multifunctional protein ADE2 | 7C(125.0477) | NaN | X | NaN | NaN | NaN | NaN | 0.000000e+00 | 0.000000e+00 | NaN | NaN | LPSGLGCSTVLSPEGSAQFAAQIFGLSNHLVWSK |
| NUMB | NUMB\_HUMAN | Protein numb homolog | 165 | 154 | NUMB | NUMB\_HUMAN | Protein numb homolog | 12C(125.0477) | NaN | X | NaN | NaN | NaN | NaN | 0.000000e+00 | 0.000000e+00 | NaN | NaN | LSHAVGCAFAACLER |
| RPS12 | RS12\_HUMAN | 40S ribosomal protein S12 | 69 | 64 | RPS12 | RS12\_HUMAN | 40S ribosomal protein S12 | 6C(125.0477) | NaN | X | NaN | NaN | NaN | NaN | 0.000000e+00 | 0.000000e+00 | NaN | NaN | LVEALCAEHQINLIK |
| TPRG1L | TPRGL\_HUMAN | Tumor protein p63-regulated gene 1-like protein | 120 | 109 | TPRG1L | TPRGL\_HUMAN | Tumor protein p63-regulated gene 1-like protein | 12C(125.0477) | NaN | X | NaN | NaN | NaN | NaN | 0.000000e+00 | 0.000000e+00 | NaN | NaN | LVLVTEQSLLICK |
| GNAI2 | GNAI2\_HUMAN | Guanine nucleotide-binding protein G(i) subunit alpha-2 | 140 | 131 | GNAI2 | GNAI2\_HUMAN | Guanine nucleotide-binding protein G(i) subunit alpha-2 | 10C(125.0477) | NaN | X | NaN | NaN | NaN | NaN | 0.000000e+00 | 0.000000e+00 | NaN | NaN | LWADHGVQACFGR |
| SELENBP1 | SBP1\_HUMAN | Methanethiol oxidase | 5, 8 | 1 | SELENBP1 | SBP1\_HUMAN | Methanethiol oxidase | 5C(31.9988) 8C(31.9988) | NaN | NaN | NaN | X | NaN | NaN | 0.000000e+00 | 0.000000e+00 | NaN | NaN | MATKCGNCGPGYSTPLEAMK |
| AP2M1 | AP2M1\_HUMAN | AP-2 complex subunit mu | 86 | 85 | AP2M1 | AP2M1\_HUMAN | AP-2 complex subunit mu | 2C(125.0477) | NaN | X | NaN | NaN | NaN | NaN | 0.000000e+00 | 0.000000e+00 | NaN | NaN | MCDVMAAYFGK |
| GTPBP1 | GTPB1\_HUMAN | GTP-binding protein 1 | 360 | 359 | GTPBP1 | GTPB1\_HUMAN | GTP-binding protein 1 | 2C(125.0477) | NaN | X | NaN | NaN | NaN | NaN | 0.000000e+00 | 0.000000e+00 | NaN | NaN | MCPIFQISNVTGENLDLLK |
| SPTA1 | SPTA1\_HUMAN | Spectrin alpha chain, erythrocytic 1 | 2072 | 2062 | SPTA1 | SPTA1\_HUMAN | Spectrin alpha chain, erythrocytic 1 | 11C(-34.0809) | NaN | NaN | NaN | NaN | X | NaN | 0.000000e+00 | 0.000000e+00 | NaN | NaN | MEENLSEPVHCVSLNEIRQLQKDHEDFLASLAR |
| CCT5 | TCPE\_HUMAN | T-complex protein 1 subunit epsilon | 377 | 371 | CCT5 | TCPE\_HUMAN | T-complex protein 1 subunit epsilon | 7C(125.0477) | NaN | X | NaN | NaN | NaN | NaN | 0.000000e+00 | 0.000000e+00 | NaN | NaN | MLVIEQCKNSR |
| GAPVD1 | GAPD1\_HUMAN | GTPase-activating protein and VPS9 domain-containing protein 1 | 1160 | 1145 | GAPVD1 | GAPD1\_HUMAN | GTPase-activating protein and VPS9 domain-containing protein 1 | 16C(125.0477) | NaN | X | NaN | NaN | NaN | NaN | 0.000000e+00 | 0.000000e+00 | NaN | NaN | NGLPDHTDPEDNEIVCFLK |
| DDB1 | DDB1\_HUMAN | DNA damage-binding protein 1 | 503 | 497 | DDB1 | DDB1\_HUMAN | DNA damage-binding protein 1 | 7C(125.0477) | NaN | X | NaN | NaN | NaN | NaN | 0.000000e+00 | 0.000000e+00 | NaN | NaN | NISVASCNSSQVVVAVGR |
| OSBP2 | OSBP2\_HUMAN | Oxysterol-binding protein 2 | 319 | 308 | OSBP2 | OSBP2\_HUMAN | Oxysterol-binding protein 2 | 12C(125.0477) | NaN | X | NaN | NaN | NaN | NaN | 0.000000e+00 | 0.000000e+00 | NaN | NaN | NLSLKLDDLSTCNDLIAK |
| HBE1 | HBE\_HUMAN | Hemoglobin subunit epsilon | 94 | 78 | HBE1 | HBE\_HUMAN | Hemoglobin subunit epsilon | 17C(125.0477) | NaN | X | NaN | NaN | NaN | NaN | 0.000000e+00 | 0.000000e+00 | NaN | NaN | NMDNLKPAFAKLSELHCDK |
| HBE1 | HBE\_HUMAN | Hemoglobin subunit epsilon | 94 | 78 | HBE1 | HBE\_HUMAN | Hemoglobin subunit epsilon | 17C(125.0477) | NaN | X | NaN | NaN | NaN | NaN | 0.000000e+00 | 0.000000e+00 | NaN | NaN | NMDNLKPAFAKLSELHCDKLHVDPENFK |
| RAB23 | RAB23\_HUMAN | Ras-related protein Rab-23 | 234 | 229 | RAB23 | RAB23\_HUMAN | Ras-related protein Rab-23 | 6C(31.9988) | NaN | NaN | NaN | X | NaN | NaN | 0.000000e+00 | 0.000000e+00 | NaN | NaN | NPFSSCSIP |
| ZC3HAV1 | ZCCHV\_HUMAN | Zinc finger CCCH-type antiviral protein 1 | 645 | 630 | ZC3HAV1 | ZCCHV\_HUMAN | Zinc finger CCCH-type antiviral protein 1 | 16C(125.0477) | NaN | X | NaN | NaN | NaN | NaN | 0.000000e+00 | 0.000000e+00 | NaN | NaN | NSNVDSSYLESLYQSCPR |
| ELANE | ELNE\_HUMAN | Neutrophil elastase | 208, 198 | 194 | ELANE | ELNE\_HUMAN | Neutrophil elastase | 15C(31.9988) 5C(-34.0809) | NaN | NaN | NaN | X | X | NaN | 0.000000e+00 | 0.000000e+00 | NaN | NaN | QAGVCFGDSGSPLVCNGLIHGIASFVR |
| MYH1 | MYH1\_HUMAN | Myosin-1 | 1464 | 1462 | MYH1 | MYH1\_HUMAN | Myosin-1 | 3C(31.9988) | NaN | NaN | NaN | X | NaN | NaN | 0.000000e+00 | 0.000000e+00 | NaN | NaN | QKCEETHAELEASQKESR |
| PIEZO1 | PIEZ1\_HUMAN | Piezo-type mechanosensitive ion channel component 1 | 982 | 973 | PIEZO1 | PIEZ1\_HUMAN | Piezo-type mechanosensitive ion channel component 1 | 10C(125.0477) | NaN | X | NaN | NaN | NaN | NaN | 0.000000e+00 | 0.000000e+00 | NaN | NaN | QQLDQDLLGCLK |
| TLN1 | TLN1\_HUMAN | Talin-1 | 2442 | 2430 | TLN1 | TLN1\_HUMAN | Talin-1 | 13C(125.0477) | NaN | X | NaN | NaN | NaN | NaN | 0.000000e+00 | 0.000000e+00 | NaN | NaN | QVAASTAQLLVACKVK |
| NAPRT | PNCB\_HUMAN | Nicotinate phosphoribosyltransferase | 48 | 47 | NAPRT | PNCB\_HUMAN | Nicotinate phosphoribosyltransferase | 2C(-34.0809) | NaN | NaN | NaN | NaN | X | NaN | 0.000000e+00 | 0.000000e+00 | NaN | NaN | RCPFGGAFALAAGLR |
| RPS17 | RS17\_HUMAN | 40S ribosomal protein S17 | 35 | 33 | RPS17 | RS17\_HUMAN | 40S ribosomal protein S17 | 3C(-34.0809) | NaN | NaN | NaN | NaN | X | NaN | 0.000000e+00 | 0.000000e+00 | NaN | NaN | RVCEEIAIIPSKK |
| NAP1L1 | NP1L1\_HUMAN | Nucleosome assembly protein 1-like 1 | 255 | 237 | NAP1L1 | NP1L1\_HUMAN | Nucleosome assembly protein 1-like 1 | 19C(15.9949) | NaN | NaN | X | NaN | NaN | NaN | 0.000000e+00 | 0.000000e+00 | NaN | NaN | SEPDDSDPFSFDGPEIMGCTGCQIDWK |
| ENO1 | ENOA\_HUMAN | Alpha-enolase | 389 | 373 | ENO1 | ENOA\_HUMAN | Alpha-enolase | 17C(57.0215) | X | NaN | NaN | NaN | NaN | NaN | 0.000000e+00 | 0.000000e+00 | NaN | NaN | SGETEDTFIADLVVGLCTGQIKTGAPCRSER |
| DNAJC13 | DJC13\_HUMAN | DnaJ homolog subfamily C member 13 | 2181 | 2169 | DNAJC13 | DJC13\_HUMAN | DnaJ homolog subfamily C member 13 | 13C(125.0477) | NaN | X | NaN | NaN | NaN | NaN | 0.000000e+00 | 0.000000e+00 | NaN | NaN | SLQYGEQVNEILCR |
| ADPGK | ADPGK\_HUMAN | ADP-dependent glucokinase | 40 | 34 | ADPGK | ADPGK\_HUMAN | ADP-dependent glucokinase | 7C(15.9949) | NaN | NaN | X | NaN | NaN | NaN | 0.000000e+00 | 0.000000e+00 | NaN | NaN | SLWSSLCLGPAPAPPGPVSPEGR |
| CORO7 | CORO7\_HUMAN | Coronin-7 | 42 | 40 | CORO7 | CORO7\_HUMAN | Coronin-7 | 3C(125.0477) | NaN | X | NaN | NaN | NaN | NaN | 0.000000e+00 | 0.000000e+00 | NaN | NaN | SSCSLIAFNSDRPGVLGIVPLQGQGEDKR |
| ESD | ESTD\_HUMAN | S-formylglutathione hydrolase | 181, 176 | 168 | ESD | ESTD\_HUMAN | S-formylglutathione hydrolase | 14C(31.9988) 9C(125.0477) | NaN | X | NaN | X | NaN | NaN | 0.000000e+00 | 0.000000e+00 | NaN | NaN | SVSAFAPICNPVLCPWGKK |
| PTRH2 | PTH2\_HUMAN | Peptidyl-tRNA hydrolase 2, mitochondrial | 86 | 82 | PTRH2 | PTH2\_HUMAN | Peptidyl-tRNA hydrolase 2, mitochondrial | 5C(125.0477) | NaN | X | NaN | NaN | NaN | NaN | 0.000000e+00 | 0.000000e+00 | NaN | NaN | VAAQCSHAAVSAYK |
| RPS17 | RS17\_HUMAN | 40S ribosomal protein S17 | 35 | 34 | RPS17 | RS17\_HUMAN | 40S ribosomal protein S17 | 2C(125.0477) | NaN | X | NaN | NaN | NaN | NaN | 0.000000e+00 | 0.000000e+00 | NaN | NaN | VCEEIAIIPSKKLR |
| MYH11 | MYH11\_HUMAN | Myosin-11 | 382 | 381 | MYH11 | MYH11\_HUMAN | Myosin-11 | 2C(-34.0809) | NaN | NaN | NaN | NaN | X | NaN | 0.000000e+00 | 0.000000e+00 | NaN | NaN | VCHLMGINVTDFTR |
| UBE2O | UBE2O\_HUMAN | (E3-independent) E2 ubiquitin-conjugating enzyme | 1040 | 1039 | UBE2O | UBE2O\_HUMAN | (E3-independent) E2 ubiquitin-conjugating enzyme | 2C(125.0477) | NaN | X | NaN | NaN | NaN | NaN | 0.000000e+00 | 0.000000e+00 | NaN | NaN | VCVSLLGTWIGK |
| OSBP | OSBP1\_HUMAN | Oxysterol-binding protein 1 | 224 | 218 | OSBP | OSBP1\_HUMAN | Oxysterol-binding protein 1 | 7C(125.0477) | NaN | X | NaN | NaN | NaN | NaN | 0.000000e+00 | 0.000000e+00 | NaN | NaN | VEDLSTCNDLIAK |
| KIF5B | KINH\_HUMAN | Kinesin-1 heavy chain | 65 | 51 | KIF5B | KINH\_HUMAN | Kinesin-1 heavy chain | 15C(15.9949) | NaN | NaN | X | NaN | NaN | NaN | 0.000000e+00 | 0.000000e+00 | NaN | NaN | VFQSSTSQEQVYNDCAKK |
| ECPAS | ECM29\_HUMAN | Proteasome adapter and scaffold protein ECM29 | 243 | 229 | ECPAS | ECM29\_HUMAN | Proteasome adapter and scaffold protein ECM29 | 15C(125.0477) | NaN | X | NaN | NaN | NaN | NaN | 0.000000e+00 | 0.000000e+00 | NaN | NaN | VIGDNPWTPEQLEQCK |
| HBE1 | HBE\_HUMAN | Hemoglobin subunit epsilon | 94 | 68 | HBE1 | HBE\_HUMAN | Hemoglobin subunit epsilon | 27C(125.0477) | NaN | X | NaN | NaN | NaN | NaN | 0.000000e+00 | 0.000000e+00 | NaN | NaN | VLTSFGDAIKNMDNLKPAFAKLSELHCDK |
| HECTD4 | HECD4\_HUMAN | Probable E3 ubiquitin-protein ligase HECTD4 | 2597 | 2581 | HECTD4 | HECD4\_HUMAN | Probable E3 ubiquitin-protein ligase HECTD4 | 17C(31.9988) | NaN | NaN | NaN | X | NaN | NaN | 0.000000e+00 | 0.000000e+00 | NaN | NaN | VNMDVNFPGAAFVVVSCK |
| ETF1 | ERF1\_HUMAN | Eukaryotic peptide chain release factor subunit 1 | 97 | 88 | ETF1 | ERF1\_HUMAN | Eukaryotic peptide chain release factor subunit 1 | 10C(125.0477) | NaN | X | NaN | NaN | NaN | NaN | 0.000000e+00 | 0.000000e+00 | NaN | NaN | VPPNGLVVYCGTIVTEEGKEK |
| GAPDHS | G3PT\_HUMAN | Glyceraldehyde-3-phosphate dehydrogenase, testis-specific | 319 | 307 | GAPDHS | G3PT\_HUMAN | Glyceraldehyde-3-phosphate dehydrogenase, testis-specific | 13C(125.0477) | NaN | X | NaN | NaN | NaN | NaN | 0.000000e+00 | 0.000000e+00 | NaN | NaN | VPTPDVSVVDLTCR |
| TATDN1 | TATD1\_HUMAN | Deoxyribonuclease TATDN1 | 113 | 107 | TATDN1 | TATD1\_HUMAN | Deoxyribonuclease TATDN1 | 7C(125.0477) | NaN | X | NaN | NaN | NaN | NaN | 0.000000e+00 | 0.000000e+00 | NaN | NaN | VVAIGECGLDFDR |
| VCP | TERA\_HUMAN | Transitional endoplasmic reticulum ATPase | 174, 184 | 165 | VCP | TERA\_HUMAN | Transitional endoplasmic reticulum ATPase | 10C(125.0477) 20C(-34.0809) | NaN | X | NaN | NaN | X | NaN | 0.000000e+00 | 0.000000e+00 | NaN | NaN | VVETDPSPYCIVAPDTVIHCEGEPIKR |
| WDR53 | WDR53\_HUMAN | WD repeat-containing protein 53 | 15 | 5 | WDR53 | WDR53\_HUMAN | WD repeat-containing protein 53 | 11C(-34.0809) | NaN | NaN | NaN | NaN | X | NaN | 0.000000e+00 | 0.000000e+00 | NaN | NaN | WTGGHSSPVLCLNASK |
| CUL1 | CUL1\_HUMAN | Cullin-1 | 426 | 425 | CUL1 | CUL1\_HUMAN | Cullin-1 | 2C(125.0477) | NaN | X | NaN | NaN | NaN | NaN | 0.000000e+00 | 0.000000e+00 | NaN | NaN | YCDSLLKK |
| TMCC2 | TMCC2\_HUMAN | Transmembrane and coiled-coil domains protein 2 | 479 | 473 | TMCC2 | TMCC2\_HUMAN | Transmembrane and coiled-coil domains protein 2 | 7C(125.0477) | NaN | X | NaN | NaN | NaN | NaN | 0.000000e+00 | 0.000000e+00 | NaN | NaN | YGSDDECSSASASSAGAGSNSGAGPGGALGSPK |
| DYNLL2 | DYL2\_HUMAN | Dynein light chain 2, cytoplasmic | 56 | 50 | DYNLL2 | DYL2\_HUMAN | Dynein light chain 2, cytoplasmic | 7C(125.0477) | NaN | X | NaN | NaN | NaN | NaN | 0.000000e+00 | 0.000000e+00 | NaN | NaN | YNPTWHCIVGR |
| IPO9 | IPO9\_HUMAN | Importin-9 | 639 | 616 | IPO9 | IPO9\_HUMAN | Importin-9 | 24C(57.0215) | X | NaN | NaN | NaN | NaN | NaN | 0.000000e+00 | 0.000000e+00 | NaN | NaN | YSNDPVVASLAQDIFKELSQIEACQGPMQMR |
| ALB | ALBU\_HUMAN | Albumin | 58 | 45 | ALB | ALBU\_HUMAN | Albumin | 14C(57.0215) | X | NaN | NaN | NaN | NaN | NaN | 0.000000e+00 | NaN | 0.000000e+00 | 0.0000 | ALVLIAFAQYLQQCPFEDHVK |
| CR1 | CR1\_HUMAN | Complement receptor type 1 | 1370 | 1370 | CR1 | CR1\_HUMAN | Complement receptor type 1 | 1C(15.9949) | NaN | NaN | X | NaN | NaN | NaN | 0.000000e+00 | NaN | NaN | 0.0000 | CTSDPHGNGVWSSPAPR |
| RPTOR | RPTOR\_HUMAN | Regulatory-associated protein of mTOR | 1037 | 1031 | RPTOR | RPTOR\_HUMAN | Regulatory-associated protein of mTOR | 7C(15.9949) | NaN | NaN | X | NaN | NaN | NaN | 0.000000e+00 | NaN | NaN | 0.0000 | FHPFTPCIAVADK |
| KRT6A | K2C6A\_HUMAN | Keratin, type II cytoskeletal 6A | 402 | 400 | KRT6A | K2C6A\_HUMAN | Keratin, type II cytoskeletal 6A | 3C(-34.0809) | NaN | NaN | NaN | NaN | X | NaN | 0.000000e+00 | NaN | NaN | 0.0000 | KQCANLQAAIADAEQR |
| CLIC1 | CLIC1\_HUMAN | Chloride intracellular channel protein 1 | 59 | 58 | CLIC1 | CLIC1\_HUMAN | Chloride intracellular channel protein 1 | 2C(57.0215) | X | NaN | NaN | NaN | NaN | NaN | 0.000000e+00 | NaN | NaN | 0.0000 | LCPGGQLPFLLYGTEVHTDTNK |
| ACTB | ACTB\_HUMAN | Actin, cytoplasmic 1 | 217 | 216 | ACTB | ACTB\_HUMAN | Actin, cytoplasmic 1 | 2C(57.0215) | X | NaN | NaN | NaN | NaN | NaN | 0.000000e+00 | NaN | NaN | 0.0000 | LCYVALDFEQEMATAASSSSLEK |
| CCT4 | TCPD\_HUMAN | T-complex protein 1 subunit delta | 379 | 373 | CCT4 | TCPD\_HUMAN | T-complex protein 1 subunit delta | 7C(15.9949) | NaN | NaN | X | NaN | NaN | NaN | 0.000000e+00 | NaN | NaN | 0.0000 | LLKITGCASPGK |
| ATP1A4 | AT1A4\_HUMAN | Sodium/potassium-transporting ATPase subunit alpha-4 | 251 | 249 | ATP1A4 | AT1A4\_HUMAN | Sodium/potassium-transporting ATPase subunit alpha-4 | 3C(15.9949) | NaN | NaN | X | NaN | NaN | NaN | 0.000000e+00 | NaN | NaN | 0.0000 | NICFFSTNCVEGTAR |
| EEF2 | EF2\_HUMAN | Elongation factor 2 | 290 | 288 | EEF2 | EF2\_HUMAN | Elongation factor 2 | 3C(57.0215) | X | NaN | NaN | NaN | NaN | NaN | 0.000000e+00 | NaN | NaN | 0.0000 | TFCQLILDPIFK |
| ZDHHC5 | ZDHC5\_HUMAN | Palmitoyltransferase ZDHHC5 | 245 | 243 | ZDHHC5 | ZDHC5\_HUMAN | Palmitoyltransferase ZDHHC5 | 3C(31.9988) | NaN | NaN | NaN | X | NaN | NaN | 0.000000e+00 | NaN | NaN | 0.0000 | VLCSSPAPR |
| IER3IP1 | IR3IP\_HUMAN | Immediate early response 3-interacting protein 1 | 15 | 2 | IER3IP1 | IR3IP\_HUMAN | Immediate early response 3-interacting protein 1 | 14C(305.3076) | NaN | NaN | NaN | NaN | NaN | X | 0.000000e+00 | NaN | 0.000000e+00 | NaN | AFTLYSLLQAALLCVNAIAVLHEER |
| PARP12 | PAR12\_HUMAN | Protein mono-ADP-ribosyltransferase PARP12 | 95, 85 | 77 | PARP12 | PAR12\_HUMAN | Protein mono-ADP-ribosyltransferase PARP12 | 19C(-34.0809) 9C(31.9988) | NaN | NaN | NaN | X | X | NaN | 0.000000e+00 | NaN | 0.000000e+00 | NaN | AHQGSKPGCVGLCAQLHLCR |
| FASN | FAS\_HUMAN | Fatty acid synthase | 1459, 1448 | 1445 | FASN | FAS\_HUMAN | Fatty acid synthase | 15C(-34.0809) 4C(15.9949) | NaN | NaN | X | NaN | X | NaN | 0.000000e+00 | NaN | 0.000000e+00 | NaN | AINCATSGVVGLVNCLR |
| SPTA1 | SPTA1\_HUMAN | Spectrin alpha chain, erythrocytic 1 | 965 | 957 | SPTA1 | SPTA1\_HUMAN | Spectrin alpha chain, erythrocytic 1 | 9C(31.9988) | NaN | NaN | NaN | X | NaN | NaN | 0.000000e+00 | NaN | 0.000000e+00 | NaN | ALRNQANACQQQQAAPVEGVAGEQR |
| MYO18A | MY18A\_HUMAN | Unconventional myosin-XVIIIa | 1155 | 1148 | MYO18A | MY18A\_HUMAN | Unconventional myosin-XVIIIa | 8C(15.9949) | NaN | NaN | X | NaN | NaN | NaN | 0.000000e+00 | NaN | 0.000000e+00 | NaN | AVEELLECLDLEK |
| CORO1C | COR1C\_HUMAN | Coronin-1C | 420 | 420 | CORO1C | COR1C\_HUMAN | Coronin-1C | 1C(125.0477) | NaN | X | NaN | NaN | NaN | NaN | 0.000000e+00 | NaN | 0.000000e+00 | NaN | CDLISIPK |
| CUL1 | CUL1\_HUMAN | Cullin-1 | 752 | 752 | CUL1 | CUL1\_HUMAN | Cullin-1 | 1C(31.9988) | NaN | NaN | NaN | X | NaN | NaN | 0.000000e+00 | NaN | 0.000000e+00 | NaN | CIDILIEKEYLERVDGEK |
| PRSS27 | PRS27\_HUMAN | Serine protease 27 | 191 | 191 | PRSS27 | PRS27\_HUMAN | Serine protease 27 | 1C(15.9949) | NaN | NaN | X | NaN | NaN | NaN | 0.000000e+00 | NaN | 0.000000e+00 | NaN | CNLLYSKDTEFGYQPK |
| KIF26A | KI26A\_HUMAN | Kinesin-like protein KIF26A | 838 | 838 | KIF26A | KI26A\_HUMAN | Kinesin-like protein KIF26A | 1C(57.0215) | X | NaN | NaN | NaN | NaN | NaN | 0.000000e+00 | NaN | 0.000000e+00 | NaN | CSTFAELQER |
| CR1 | CR1\_HUMAN | Complement receptor type 1 | 470 | 470 | CR1 | CR1\_HUMAN | Complement receptor type 1 | 1C(125.0477) | NaN | X | NaN | NaN | NaN | NaN | 0.000000e+00 | NaN | 0.000000e+00 | NaN | CTSDPQGNGVWSSPAPR |
| CD59 | CD59\_HUMAN | CD59 glycoprotein | 64, 70 | 64 | CD59 | CD59\_HUMAN | CD59 glycoprotein | 1C(125.0477) 7C(57.0215) | X | X | NaN | NaN | NaN | NaN | 0.000000e+00 | NaN | 0.000000e+00 | NaN | CWKFEHCNFNDVTTR |
| CFH | CFAH\_HUMAN | Complement factor H | 505 | 497 | CFH | CFAH\_HUMAN | Complement factor H | 9C(57.0215) | X | NaN | NaN | NaN | NaN | NaN | 0.000000e+00 | NaN | 0.000000e+00 | NaN | DGWSAQPTCIK |
| EIF3J | EIF3J\_HUMAN | Eukaryotic translation initiation factor 3 subunit J | 188 | 186 | EIF3J | EIF3J\_HUMAN | Eukaryotic translation initiation factor 3 subunit J | 3C(-34.0809) | NaN | NaN | NaN | NaN | X | NaN | 0.000000e+00 | NaN | 0.000000e+00 | NaN | DVCISLEIDDLK |
| SPTLC2 | SPTC2\_HUMAN | Serine palmitoyltransferase 2 | 438 | 437 | SPTLC2 | SPTC2\_HUMAN | Serine palmitoyltransferase 2 | 2C(125.0477) | NaN | X | NaN | NaN | NaN | NaN | 0.000000e+00 | NaN | 0.000000e+00 | NaN | ECVQQLAENTR |
| POTEI | POTEI\_HUMAN | POTE ankyrin domain family member I | 917 | 914 | POTEI | POTEI\_HUMAN | POTE ankyrin domain family member I | 4C(31.9988) | NaN | NaN | NaN | X | NaN | NaN | 0.000000e+00 | NaN | 0.000000e+00 | NaN | EKLCYVALDFEQEMAMAASSSSLEK |
| CTTNBP2 | CTTB2\_HUMAN | Cortactin-binding protein 2 | 1595 | 1583 | CTTNBP2 | CTTB2\_HUMAN | Cortactin-binding protein 2 | 13C(125.0477) | NaN | X | NaN | NaN | NaN | NaN | 0.000000e+00 | NaN | 0.000000e+00 | NaN | EVSPLSSHQTTECSNSK |
| ANKRD44 | ANR44\_HUMAN | Serine/threonine-protein phosphatase 6 regulatory ankyrin repeat subunit B | 420, 408 | 404 | ANKRD44 | ANR44\_HUMAN | Serine/threonine-protein phosphatase 6 regulatory ankyrin repeat subunit B | 17C(125.0477) 5C(31.9988) | NaN | X | NaN | X | NaN | NaN | 0.000000e+00 | NaN | 0.000000e+00 | NaN | FGRTCLHAAAAGGNVECIK |
| CSNK2A1 | CSK21\_HUMAN | Casein kinase II subunit alpha | 147 | 135 | CSNK2A1 | CSK21\_HUMAN | Casein kinase II subunit alpha | 13C(15.9949) | NaN | NaN | X | NaN | NaN | NaN | 0.000000e+00 | NaN | 0.000000e+00 | NaN | FYMYEILKALDYCHSMGIMHR |
| SLC4A1 | B3AT\_HUMAN | Band 3 anion transport protein | 317 | 305 | SLC4A1 | B3AT\_HUMAN | Band 3 anion transport protein | 13C(125.0477) | NaN | X | NaN | NaN | NaN | NaN | 0.000000e+00 | NaN | 0.000000e+00 | NaN | GELLHSLEGFLDCSLVLPPTDAPSEQALLSLVPVQR |
| HBG1 | HBG1\_HUMAN | Hemoglobin subunit gamma-1 | 94 | 78 | HBG1 | HBG1\_HUMAN | Hemoglobin subunit gamma-1 | 17C(125.0477) | NaN | X | NaN | NaN | NaN | NaN | 0.000000e+00 | NaN | 0.000000e+00 | NaN | HLDDLKGTFAQLSELHCDKLHVDPENFK |
| PPIA | PPIA\_HUMAN | Peptidyl-prolyl cis-trans isomerase A | 115 | 92 | PPIA | PPIA\_HUMAN | Peptidyl-prolyl cis-trans isomerase A | 24C(57.0215) | X | NaN | NaN | NaN | NaN | NaN | 0.000000e+00 | NaN | 0.000000e+00 | NaN | HTGPGILSMANAGPNTNGSQFFICTAK |
| TMEM63B | CSCL2\_HUMAN | CSC1-like protein 2 | 691 | 676 | TMEM63B | CSCL2\_HUMAN | CSC1-like protein 2 | 16C(57.0215) | X | NaN | NaN | NaN | NaN | NaN | 0.000000e+00 | NaN | 0.000000e+00 | NaN | IHSGAVNQVVAAPILCLFWLLFFSTMR |
| LDHB | LDHB\_HUMAN | L-lactate dehydrogenase B chain | 36 | 24 | LDHB | LDHB\_HUMAN | L-lactate dehydrogenase B chain | 13C(125.0477) | NaN | X | NaN | NaN | NaN | NaN | 0.000000e+00 | NaN | 0.000000e+00 | NaN | ITVVGVGQVGMACAISILGK |
| GNB1 | GBB1\_HUMAN | Guanine nucleotide-binding protein G(I)/G(S)/G(T) subunit beta-1 | 25 | 23 | GNB1 | GBB1\_HUMAN | Guanine nucleotide-binding protein G(I)/G(S)/G(T) subunit beta-1 | 3C(15.9949) | NaN | NaN | X | NaN | NaN | NaN | 0.000000e+00 | NaN | 0.000000e+00 | NaN | KACADATLSQITNNIDPVGR |
| PIEZO1 | PIEZ1\_HUMAN | Piezo-type mechanosensitive ion channel component 1 | 95 | 87 | PIEZO1 | PIEZ1\_HUMAN | Piezo-type mechanosensitive ion channel component 1 | 9C(125.0477) | NaN | X | NaN | NaN | NaN | NaN | 0.000000e+00 | NaN | 0.000000e+00 | NaN | LDQLLGPSCSR |
| MUC5B | MUC5B\_HUMAN | Mucin-5B | 5328 | 5324 | MUC5B | MUC5B\_HUMAN | Mucin-5B | 5C(31.9988) | NaN | NaN | NaN | X | NaN | NaN | 0.000000e+00 | NaN | 0.000000e+00 | NaN | LEVPCQSLEAYAELCR |
| TUBB1 | TBB1\_HUMAN | Tubulin beta-1 chain | 340 | 337 | TUBB1 | TBB1\_HUMAN | Tubulin beta-1 chain | 4C(57.0215) | X | NaN | NaN | NaN | NaN | NaN | 0.000000e+00 | NaN | 0.000000e+00 | NaN | NSSCFVEWIPNNVK |
| NRBF2 | NRBF2\_HUMAN | Nuclear receptor-binding factor 2 | 155 | 149 | NRBF2 | NRBF2\_HUMAN | Nuclear receptor-binding factor 2 | 7C(57.0215) | X | NaN | NaN | NaN | NaN | NaN | 0.000000e+00 | NaN | 0.000000e+00 | NaN | SEPAEPCIGSK |
| RNH1 | RINI\_HUMAN | Ribonuclease inhibitor | 75, 85 | 65 | RNH1 | RINI\_HUMAN | Ribonuclease inhibitor | 11C(-34.0809) 21C(15.9949) | NaN | NaN | X | NaN | X | NaN | 0.000000e+00 | NaN | 0.000000e+00 | NaN | SNELGDVGVHCVLQGLQTPSCK |
| TPP2 | TPP2\_HUMAN | Tripeptidyl-peptidase 2 | 1125 | 1118 | TPP2 | TPP2\_HUMAN | Tripeptidyl-peptidase 2 | 8C(125.0477) | NaN | X | NaN | NaN | NaN | NaN | 0.000000e+00 | NaN | 0.000000e+00 | NaN | STLVDALCR |
| IPO5 | IPO5\_HUMAN | Importin-5 | 110 | 109 | IPO5 | IPO5\_HUMAN | Importin-5 | 2C(125.0477) | NaN | X | NaN | NaN | NaN | NaN | 0.000000e+00 | NaN | 0.000000e+00 | NaN | VCDIAAELAR |
| RIPOR1 | RIPR1\_HUMAN | Rho family-interacting cell polarization regulator 1 | 945 | 940 | RIPOR1 | RIPR1\_HUMAN | Rho family-interacting cell polarization regulator 1 | 6C(57.0215) | X | NaN | NaN | NaN | NaN | NaN | 0.000000e+00 | NaN | 0.000000e+00 | NaN | VLEAVCEFSR |
| SPTAN1 | SPTN1\_HUMAN | Spectrin alpha chain, non-erythrocytic 1 | 1930 | 1926 | SPTAN1 | SPTN1\_HUMAN | Spectrin alpha chain, non-erythrocytic 1 | 5C(125.0477) | NaN | X | NaN | NaN | NaN | NaN | 0.000000e+00 | NaN | 0.000000e+00 | NaN | VNDVCTNGQDLIK |
| PPP6R1 | PP6R1\_HUMAN | Serine/threonine-protein phosphatase 6 regulatory subunit 1 | 455 | 453 | PPP6R1 | PP6R1\_HUMAN | Serine/threonine-protein phosphatase 6 regulatory subunit 1 | 3C(125.0477) | NaN | X | NaN | NaN | NaN | NaN | 0.000000e+00 | NaN | 0.000000e+00 | NaN | VQCAGGPR |
| USP15 | UBP15\_HUMAN | Ubiquitin carboxyl-terminal hydrolase 15 | 127 | 117 | USP15 | UBP15\_HUMAN | Ubiquitin carboxyl-terminal hydrolase 15 | 11C(125.0477) | NaN | X | NaN | NaN | NaN | NaN | 0.000000e+00 | NaN | 0.000000e+00 | NaN | VVEQGMFVKHCK |
| VCP | TERA\_HUMAN | Transitional endoplasmic reticulum ATPase | 174, 184 | 165 | VCP | TERA\_HUMAN | Transitional endoplasmic reticulum ATPase | 10C(31.9988) 20C(125.0477) | NaN | X | NaN | X | NaN | NaN | 0.000000e+00 | NaN | 0.000000e+00 | NaN | VVETDPSPYCIVAPDTVIHCEGEPIK |
| SELENOO | SELO\_HUMAN | Protein adenylyltransferase SelO, mitochondrial | 241 | 238 | SELENOO | SELO\_HUMAN | Protein adenylyltransferase SelO, mitochondrial | 4C(31.9988) | NaN | NaN | NaN | X | NaN | NaN | 0.000000e+00 | NaN | 0.000000e+00 | NaN | YEQCTVVLR |
| NDUFV2 | NDUV2\_HUMAN | NADH dehydrogenase [ubiquinone] flavoprotein 2, mitochondrial | 140 | 130 | NDUFV2 | NDUV2\_HUMAN | NADH dehydrogenase [ubiquinone] flavoprotein 2, mitochondrial | 11C(15.9949) | NaN | NaN | X | NaN | NaN | NaN | 0.000000e+00 | NaN | 0.000000e+00 | NaN | YHIQVCTTTPCMLR |
