## Supplementary Resource - Deep Red for "Deep Red Blood Cell Proteome Defines the Band 3 N-Terminus Interactome as a Regulator of Hypoxic Adaptation via BLVRB-Dependent *S*-Nitroso Transfer": Phosphoproteomics 2025.html

RBC Phosphopeptide Analysis

### RBC Phosphopeptide Analysis

| Protein | Uniprot ID | Description | Peptide | Modified Residue | Modified Position | Total | Cell | sECM | iECM |
| --- | --- | --- | --- | --- | --- | --- | --- | --- | --- |
| SPTB | SPTB1\_HUMAN | Spectrin beta chain, erythrocytic | T[79.9663]SPVSLWSR | T | 2113.0 | 98934276.12 | 0.0 | 65676491.5 | 33257784.62 |
| SLC4A1 | B3AT\_HUMAN | Band 3 anion transport protein | YQSSPAKPDS[79.9663]SFYK | S | 356.0 | 50325787.64 | 0.0 | 25517735.86 | 24808051.78 |
| SPTB | SPTB1\_HUMAN | Spectrin beta chain, erythrocytic | TS[79.9663]PVSLWSR | S | 2114.0 | 40585653.94 | 0.0 | 9919310.571 | 30666343.37 |
| EPB41 | EPB41\_HUMAN | Isoform 5 of Protein 4.1 | S[79.9663]LDGAAAVDSADR | S | 542.0 | 33647722.21 | 0.0 | 30513865.93 | 3133856.275 |
| ADD1 | ADDA\_HUMAN | Isoform 3 of Alpha-adducin | AAVVTS[79.9663]PPPTTAPHK | S | 12.0 | 22029530.61 | 2267602.65 | 17333897.24 | 2428030.72 |
| DMTN | DEMA\_HUMAN | Isoform 4 of Dematin | S[79.9663]SSLPAYGR | S | 287.0 | 21246265.03 | 0.0 | 20432812.97 | 813452.06 |
| DMTN | DEMA\_HUMAN | Isoform 4 of Dematin | SSS[79.9663]LPAYGR | S | 289.0 | 20489165.18 | 0.0 | 19183114.4 | 1306050.783 |
| SLC4A1 | B3AT\_HUMAN | Band 3 anion transport protein | YQSS[79.9663]PAKPDSSFYK | S | 350.0 | 20073081.42 | 0.0 | 5259331.89 | 14813749.53 |
| DMTN | DEMA\_HUMAN | Isoform 4 of Dematin | HLS[79.9663]AEDFSR | S | 372.0 | 19478588.78 | 0.0 | 19478588.78 | 0.0 |
| ADD1 | ADDA\_HUMAN | Isoform 3 of Alpha-adducin | AAVVT[79.9663]SPPPTTAPHK | T | 11.0 | 18904297.12 | 0.0 | 16640193.54 | 2264103.58 |
| ANK1 | ANK1\_HUMAN | Isoform Br21 of Ankyrin-1 | ITHS[79.9663]PTVSQVTER | S | 1686.0 | 18150188.42 | 0.0 | 13011229.01 | 5138959.409 |
| DMTN | DEMA\_HUMAN | Isoform 4 of Dematin | RGAEEEEEEEDDDS[79.9663]GEEM[15.9949]K | S |  | 16016581.39 | 0.0 | 15475380.11 | 541201.2793 |
| SLC4A1 | B3AT\_HUMAN | Band 3 anion transport protein | YQSSPAKPDSS[79.9663]FYK | S | 357.0 | 15531761.57 | 0.0 | 8432718.36 | 7099043.21 |
| ADD1 | ADDA\_HUMAN | Isoform 3 of Alpha-adducin | SPGS[79.9663]PVGEGTGSPPK | S | 358.0 | 13627650.05 | 0.0 | 12047906.03 | 1579744.024 |
| DMTN | DEMA\_HUMAN | Isoform 4 of Dematin | GAEEEEEEEDDDS[79.9663]GEEM[15.9949]K | S |  | 12905399.18 | 0.0 | 12145596.14 | 759803.04 |
| ANK1 | ANK1\_HUMAN | Isoform Br21 of Ankyrin-1 | LS[79.9663]TPPPLAEEEGLASR | S | 960.0 | 12108510.92 | 0.0 | 10785034.29 | 1323476.627 |
| ANK1 | ANK1\_HUMAN | Isoform Br21 of Ankyrin-1 | RRT[79.9663]PTPLALR | T | 1378.0 | 11224216.32 | 0.0 | 11105506.04 | 118710.28 |
| SPTB | SPTB1\_HUMAN | Spectrin beta chain, erythrocytic | T[79.9663]SATEFENVGNQPPYSR | T | 2.0 | 10534208.66 | 0.0 | 8988867.656 | 1545341.0 |
| SLC14A1 | UT1\_HUMAN | Urea transporter 1 | VDS[79.9663]PTM[15.9949]VR | S |  | 10170197.08 | 0.0 | 1700597.875 | 8469599.207 |
| MINK1 | MINK1\_HUMAN | Isoform 1 of Misshapen-like kinase 1 | QLQQEHAY[79.9663]LK | Y | 475.0 | 8855224.17 | 0.0 | 8855224.17 | 0.0 |
| SPTB | SPTB1\_HUMAN | Spectrin beta chain, erythrocytic | WDAPDDELDNDNS[79.9663]SAR | S | 35.0 | 8090530.122 | 0.0 | 7761282.328 | 329247.794 |
| SPTB | SPTB1\_HUMAN | Spectrin beta chain, erythrocytic | WDAPDDELDNDNSS[79.9663]AR | S | 36.0 | 8068498.814 | 0.0 | 7747970.486 | 320528.328 |
| ANK1 | ANK1\_HUMAN | Isoform Br21 of Ankyrin-1 | T[79.9663]PTPLALR | T | 1378.0 | 7331898.056 | 0.0 | 3555359.156 | 3776538.9 |
| ANK1 | ANK1\_HUMAN | Isoform Br21 of Ankyrin-1 | RTPT[79.9663]PLALR | T | 1380.0 | 7298820.687 | 0.0 | 5963239.017 | 1335581.67 |
| EPB41 | EPB41\_HUMAN | Isoform 5 of Protein 4.1 | SLDGAAAVDSADRS[79.9663]PR | S | 555.0 | 7025503.596 | 0.0 | 6821790.013 | 203713.583 |
| SLC4A1 | B3AT\_HUMAN | Band 3 anion transport protein | RYQSSPAKPDS[79.9663]SFYK | S | 356.0 | 6884682.032 | 0.0 | 3389058.092 | 3495623.94 |
| SPTB | SPTB1\_HUMAN | Spectrin beta chain, erythrocytic | PAEETGPQEEEGETAGEAPVSHHAAT[79.9663]ER | T | 2110.0 | 6629938.195 | 0.0 | 6573984.065 | 55954.13 |
| ANK1 | ANK1\_HUMAN | Isoform Br21 of Ankyrin-1 | IT[79.9663]HSPTVSQVTER | T | 1684.0 | 6585464.039 | 0.0 | 4042337.117 | 2543126.922 |
| EPB41 | EPB41\_HUMAN | Isoform 5 of Protein 4.1 | LSTHS[79.9663]PFR | S | 712.0 | 6496979.282 | 0.0 | 6059679.082 | 437300.2 |
| DMTN | DEMA\_HUMAN | Isoform 4 of Dematin | DSSVPGSPS[79.9663]SIVAK | S | 28.0 | 6271118.7 | 0.0 | 5867471.3 | 403647.4 |
| SPTB | SPTB1\_HUMAN | Spectrin beta chain, erythrocytic | LS[79.9663]SSWESLQPEPSHPY | S | 2123.0 | 5956547.239 | 0.0 | 5788354.679 | 168192.56 |
| ANK1 | ANK1\_HUMAN | Isoform Br21 of Ankyrin-1 | LST[79.9663]PPPLAEEEGLASR | T | 961.0 | 5558713.5 | 0.0 | 4302320.0 | 1256393.5 |
| ADD2 | ADDB\_HUMAN | Beta-adducin | TESVTSGPM[15.9949]SPEGSPSKS[79.9663]PSK | S | 617.0 | 5314249.226 | 0.0 | 5314249.226 | 0.0 |
| DMTN | DEMA\_HUMAN | Isoform 4 of Dematin | DSSVPGS[79.9663]PSSIVAK | S | 26.0 | 5064959.09 | 0.0 | 5064959.09 | 0.0 |
| SPTB | SPTB1\_HUMAN | Spectrin beta chain, erythrocytic | LSSSWES[79.9663]LQPEPSHPY | S | 2128.0 | 4932044.305 | 0.0 | 4932044.305 | 0.0 |
| DMTN | DEMA\_HUMAN | Isoform 4 of Dematin | DSSVPGSPSS[79.9663]IVAK | S | 29.0 | 4513867.5 | 0.0 | 4513867.5 | 0.0 |
| SPTB | SPTB1\_HUMAN | Spectrin beta chain, erythrocytic | STAS[79.9663]WAER | S | 2060.0 | 4389855.448 | 0.0 | 2948129.448 | 1441726.0 |
| SLC4A1 | B3AT\_HUMAN | Band 3 anion transport protein | YQS[79.9663]SPAKPDSSFYK | S | 349.0 | 4307463.556 | 0.0 | 1668208.37 | 2639255.186 |
| ADD1 | ADDA\_HUMAN | Isoform 3 of Alpha-adducin | S[79.9663]PGSPVGEGTGSPPK | S | 355.0 | 4005014.696 | 0.0 | 2439826.63 | 1565188.066 |
| ADD2 | ADDB\_HUMAN | Beta-adducin | TESVTSGPM[15.9949]SPEGSPS[79.9663]KSPSK | S | 699.0 | 3762100.401 | 0.0 | 3736895.296 | 25205.105 |
| EPB41 | EPB41\_HUMAN | Isoform 5 of Protein 4.1 | SLDGAAAVDS[79.9663]ADRSPR | S | 551.0 | 3716304.098 | 0.0 | 3716304.098 | 0.0 |
| ANK1 | ANK1\_HUMAN | Isoform Br21 of Ankyrin-1 | TPT[79.9663]PLALR | T | 1380.0 | 3573456.94 | 0.0 | 3573456.94 | 0.0 |
| SLC4A1 | B3AT\_HUMAN | Band 3 anion transport protein | AS[79.9663]TPGAAAQIQEVK | S | 745.0 | 3502793.734 | 0.0 | 990340.57 | 2512453.164 |
| ANK1 | ANK1\_HUMAN | Isoform Br21 of Ankyrin-1 | LEGALS[79.9663]EEPR | S | 1607.0 | 3315823.578 | 0.0 | 2780311.398 | 535512.18 |
| DMTN | DEMA\_HUMAN | Isoform 4 of Dematin | QRES[79.9663]VGGSPQTK | S | 152.0 | 3307819.707 | 0.0 | 2966028.44 | 341791.267 |
| ADD1 | ADDA\_HUMAN | Isoform 3 of Alpha-adducin | QKGS[79.9663]EENLDEAR | S | 586.0 | 3273929.363 | 0.0 | 3032074.893 | 241854.47 |
| SPTB | SPTB1\_HUMAN | Spectrin beta chain, erythrocytic | TSPVS[79.9663]LWSR | S | 2117.0 | 3181977.96 | 0.0 | 2966250.3 | 215727.66 |
| STOM | STOM\_HUMAN | Stomatin | LPDS[79.9663]FKDSPSK | S | 18.0 | 2845978.33 | 0.0 | 899433.13 | 1946545.2 |
| DMTN | DEMA\_HUMAN | Isoform 4 of Dematin | S[79.9663]TSPPPSPEVWADSR | S | 90.0 | 2751164.78 | 0.0 | 2751164.78 | 0.0 |
| EPB41L2 | E41L2\_HUMAN | Band 4.1-like protein 2 | GLS[79.9663]PAQADS[79.9663]QFLENAK | S |  | 2532098.81 | 0.0 | 2532098.81 | 0.0 |
| PIEZO1 | PIEZ1\_HUMAN | Piezo-type mechanosensitive ion channel component 1 | S[79.9663]GSEEAVTDPGER | S | 1619.0 | 2474425.847 | 0.0 | 300785.792 | 2173640.055 |
| PSMF1 | PSMF1\_HUMAN | Proteasome inhibitor PI31 subunit | ANVS[79.9663]SPHREFPPATAR | S | 152.0 | 2350349.1 | 2350349.1 | 0.0 | 0.0 |
| DMTN | DEMA\_HUMAN | Isoform 4 of Dematin | S[79.9663]TSPPPS[79.9663]PEVWADSR | S |  | 2345147.76 | 0.0 | 1947374.92 | 397772.84 |
| SPTB | SPTB1\_HUMAN | Spectrin beta chain, erythrocytic | PAEETGPQEEEGETAGEAPVS[79.9663]HHAATER | S | 2105.0 | 2338763.074 | 0.0 | 2296529.4 | 42233.674 |
| SMIM1 | SMIM1\_HUMAN | Small integral membrane protein 1 | DGVS[79.9663]LGAVS[79.9663]STEEASR | S |  | 2328570.01 | 0.0 | 598123.93 | 1730446.08 |
| IREB2 | IREB2\_HUMAN | Iron-responsive element-binding protein 2 | AVLAESY[79.9663]EK | Y | 875.0 | 2328161.8 | 0.0 | 0.0 | 2328161.8 |
| ERMAP | ERMAP\_HUMAN | Erythroid membrane-associated protein | S[79.9663]EESIVPRPEGK | S | 418.0 | 2207558.467 | 0.0 | 1355420.33 | 852138.137 |
| SLC4A1 | B3AT\_HUMAN | Band 3 anion transport protein | IDAY[79.9663]M[15.9949]AQSR | Y |  | 2184542.713 | 0.0 | 45311.684 | 2139231.029 |
| ANK1 | ANK1\_HUMAN | Isoform Br21 of Ankyrin-1 | ITHS[79.9663]PTVS[79.9663]QVTER | S |  | 2182363.231 | 0.0 | 2182363.231 | 0.0 |
| DMTN | DEMA\_HUMAN | Isoform 4 of Dematin | SPGIIS[79.9663]QASAPR | S | 110.0 | 2181990.715 | 0.0 | 2043532.555 | 138458.16 |
| PIEZO1 | PIEZ1\_HUMAN | Piezo-type mechanosensitive ion channel component 1 | TAS[79.9663]ELLLDR | S | 1646.0 | 2177199.62 | 0.0 | 414074.644 | 1763124.976 |
| PSMF1 | PSMF1\_HUMAN | Proteasome inhibitor PI31 subunit | ANVSS[79.9663]PHREFPPATAR | S | 153.0 | 2120402.56 | 2120402.56 | 0.0 | 0.0 |
| HBB | HBB\_HUMAN | Hemoglobin subunit beta | EFTPPVQAAY[79.9663]QK | Y | 131.0 | 2023145.23 | 0.0 | 586149.64 | 1436995.59 |
| EPB41 | EPB41\_HUMAN | Isoform 5 of Protein 4.1 | S[79.9663]LDGAAAVDSADRSPR | S | 542.0 | 1982520.84 | 0.0 | 1982520.84 | 0.0 |
| SMIM1 | SMIM1\_HUMAN | Small integral membrane protein 1 | DGVSLGAVS[79.9663]STEEASR | S | 27.0 | 1974860.038 | 0.0 | 253008.339 | 1721851.699 |
| ADD2 | ADDB\_HUMAN | Beta-adducin | SPSTES[79.9663]QLM[15.9949]SK | S |  | 1915435.301 | 0.0 | 1824415.693 | 91019.608 |
| SPTB | SPTB1\_HUMAN | Spectrin beta chain, erythrocytic | VLS[79.9663]PVDSGNK | S | 1226.0 | 1871858.82 | 0.0 | 1547392.04 | 324466.78 |
| PIEZO1 | PIEZ1\_HUMAN | Piezo-type mechanosensitive ion channel component 1 | T[79.9663]ASELLLDR | T | 1644.0 | 1843655.475 | 0.0 | 378603.75 | 1465051.725 |
| SPTB | SPTB1\_HUMAN | Spectrin beta chain, erythrocytic | n[42.0106]TS[79.9663]ATEFENVGNQPPYSR | S | 3.0 | 1808056.846 | 0.0 | 1606723.506 | 201333.34 |
| SPTA1 | SPTA1\_HUMAN | Spectrin alpha chain, erythrocytic 1 | DRKES[79.9663]LNEAQK | S | 1284.0 | 1802238.89 | 0.0 | 1664931.41 | 137307.48 |
| PIEZO1 | PIEZ1\_HUMAN | Piezo-type mechanosensitive ion channel component 1 | SGS[79.9663]EEAVTDPGER | S | 1621.0 | 1716807.5 | 0.0 | 0.0 | 1716807.5 |
| SPTB | SPTB1\_HUMAN | Spectrin beta chain, erythrocytic | PAEETGPQEEEGET[79.9663]AGEAPVSHHAATER | T | 2098.0 | 1716045.12 | 0.0 | 1716045.12 | 0.0 |
| DMTN | DEMA\_HUMAN | Isoform 4 of Dematin | STSPPPS[79.9663]PEVWADSR | S | 96.0 | 1626288.9 | 0.0 | 1626288.9 | 0.0 |
| SPTA1 | SPTA1\_HUMAN | Spectrin alpha chain, erythrocytic 1 | HEIDS[79.9663]YDDR | S | 421.0 | 1515980.778 | 0.0 | 1456231.11 | 59749.668 |
| ANK1 | ANK1\_HUMAN | Isoform Br21 of Ankyrin-1 | ITHSPTVS[79.9663]QVTER | S | 1690.0 | 1475332.0 | 0.0 | 1475332.0 | 0.0 |
| EPB41 | EPB41\_HUMAN | Isoform 5 of Protein 4.1 | HHAS[79.9663]ISELKK | S | 684.0 | 1447993.266 | 0.0 | 1410179.704 | 37813.562 |
| ANK1 | ANK1\_HUMAN | Isoform Br21 of Ankyrin-1 | GAS[79.9663]PNVSNVK | S | 429.0 | 1418594.211 | 0.0 | 1083669.345 | 334924.866 |
| ANK1 | ANK1\_HUMAN | Isoform Br21 of Ankyrin-1 | AEDS[79.9663]DATGHEWK | S | 1593.0 | 1404766.529 | 393976.866 | 939282.233 | 71507.43 |
| SPTB | SPTB1\_HUMAN | Spectrin beta chain, erythrocytic | PAEETGPQEEEGETAGEAPVS[79.9663]HHAAT[79.9663]ER | S |  | 1402656.996 | 0.0 | 1402656.996 | 0.0 |
| ADD3 | ADDG\_HUMAN | Isoform 1 of Gamma-adducin | IEEVLSPEGSPS[79.9663]KSPSK | S | 679.0 | 1401941.3 | 0.0 | 1331337.2 | 70604.1 |
| ADD3 | ADDG\_HUMAN | Isoform 1 of Gamma-adducin | IEEVLSPEGSPSKS[79.9663]PSK | S | 677.0 | 1386526.49 | 0.0 | 1316628.2 | 69898.29 |
| SPTA1 | SPTA1\_HUMAN | Spectrin alpha chain, erythrocytic 1 | HEIDSY[79.9663]DDR | Y | 422.0 | 1301590.1 | 0.0 | 1301590.1 | 0.0 |
| DMTN | DEMA\_HUMAN | Isoform 4 of Dematin | Q[-17.0265]RES[79.9663]VGGSPQTK | S | 152.0 | 1288685.549 | 0.0 | 1161383.739 | 127301.81 |
| ADD2 | ADDB\_HUMAN | Beta-adducin | SPS[79.9663]TES[79.9663]QLM[15.9949]SK | S |  | 1271569.6 | 0.0 | 1255174.916 | 16394.684 |
| ADD2 | ADDB\_HUMAN | Beta-adducin | DKTESVTSGPM[15.9949]SPEGSPSKS[79.9663]PSK | S | 617.0 | 1271526.71 | 0.0 | 1271526.71 | 0.0 |
| DMTN | DEMA\_HUMAN | Isoform 4 of Dematin | DS[79.9663]SVPGSPSSIVAK | S | 21.0 | 1271184.117 | 0.0 | 1271184.117 | 0.0 |
| EPB41 | EPB41\_HUMAN | Isoform 5 of Protein 4.1 | SLDGAAAVDS[79.9663]ADR | S | 551.0 | 1249141.318 | 0.0 | 1189817.399 | 59323.919 |
| ADD2 | ADDB\_HUMAN | Beta-adducin | DKTESVTSGPM[15.9949]SPEGSPS[79.9663]KSPSK | S | 699.0 | 1247747.493 | 0.0 | 1247747.493 | 0.0 |
| ADD1 | ADDA\_HUMAN | Isoform 3 of Alpha-adducin | Q[-17.0265]KGS[79.9663]EENLDEAR | S | 586.0 | 1235631.936 | 0.0 | 1143926.236 | 91705.7 |
| DMTN | DEMA\_HUMAN | Isoform 4 of Dematin | QPLTSPGSVS[79.9663]PSR | S | 16.0 | 1201427.449 | 0.0 | 831259.389 | 370168.06 |
| DMTN | DEMA\_HUMAN | Isoform 4 of Dematin | QPLTSPGS[79.9663]VSPSR | S | 14.0 | 1198825.454 | 0.0 | 1112910.264 | 85915.19 |
| SLC4A1 | B3AT\_HUMAN | Band 3 anion transport protein | Y[79.9663]HPDVPYVK | Y | 818.0 | 1190818.75 | 0.0 | 235850.5 | 954968.25 |
| SPTBN4 | SPTN4\_HUMAN | Isoform 4 of Spectrin beta chain, non-erythrocytic 4 | VHLENVGS[79.9663]HDIVDGNHR | S | 142.0 | 1180822.395 | 0.0 | 1134498.559 | 46323.836 |
| EPB41 | EPB41\_HUMAN | Isoform 5 of Protein 4.1 | SQVS[79.9663]EEEGKEVESDKEK | S | 95.0 | 1173477.36 | 0.0 | 1135267.844 | 38209.516 |
| ADD2 | ADDB\_HUMAN | Beta-adducin | S[79.9663]PSTES[79.9663]QLM[15.9949]SK | S |  | 1159400.04 | 0.0 | 1159400.04 | 0.0 |
| ADD1 | ADDA\_HUMAN | Isoform 3 of Alpha-adducin | SPGSPVGEGT[79.9663]GSPPK | T | 364.0 | 1141957.808 | 0.0 | 1113201.886 | 28755.922 |
| ADD1 | ADDA\_HUMAN | Isoform 3 of Alpha-adducin | SPGSPVGEGTGS[79.9663]PPK | S | 366.0 | 1116679.792 | 0.0 | 1116679.792 | 0.0 |
| ANK1 | ANK1\_HUMAN | Isoform Br21 of Ankyrin-1 | RT[79.9663]PTPLALR | T | 1378.0 | 1109815.666 | 0.0 | 29087.96 | 1080727.706 |
| KRT13 | K1C13\_HUMAN | Keratin, type I cytoskeletal 13 | M[15.9949]IGFPSSAGSVS[79.9663]PR | S | 427.0 | 1089122.883 | 0.0 | 935763.207 | 153359.676 |
| SPTB | SPTB1\_HUMAN | Spectrin beta chain, erythrocytic | DKVLS[79.9663]PVDSGNK | S | 1226.0 | 1060332.241 | 0.0 | 945161.373 | 115170.868 |
| HBB | HBB\_HUMAN | Hemoglobin subunit beta | LLVVY[79.9663]PWTQR | Y | 36.0 | 1051419.944 | 0.0 | 61557.301 | 989862.643 |
| ADD2 | ADDB\_HUMAN | Beta-adducin | SPLVSPSKS[79.9663]LEEGTK | S | 621.0 | 1012203.2 | 0.0 | 1012203.2 | 0.0 |
| ADD2 | ADDB\_HUMAN | Beta-adducin | SPLVSPS[79.9663]KSLEEGTK | S | 619.0 | 978335.75 | 0.0 | 978335.75 | 0.0 |
| ANK1 | ANK1\_HUMAN | Isoform Br21 of Ankyrin-1 | LGYIS[79.9663]VTDVLK | S | 781.0 | 957165.41 | 0.0 | 451694.55 | 505470.86 |
| SPTB | SPTB1\_HUMAN | Spectrin beta chain, erythrocytic | LSSS[79.9663]WESLQPEPSHPY | S | 2125.0 | 954650.04 | 0.0 | 954650.04 | 0.0 |
| ADD2 | ADDB\_HUMAN | Beta-adducin | TESVTSGPM[15.9949]SPEGS[79.9663]PSK | S | 617.0 | 952301.367 | 0.0 | 873373.177 | 78928.19 |
| GYPA | GLPA\_HUMAN | Isoform 2 of Glycophorin-A | SPSDVKPLPSPDTDVPLS[79.9663]SVEIENPETSDQ | S | 138.0 | 926812.13 | 0.0 | 926812.13 | 0.0 |
| DMTN | DEMA\_HUMAN | Isoform 4 of Dematin | SPGIISQAS[79.9663]APR | S | 113.0 | 863477.501 | 0.0 | 715630.066 | 147847.435 |
| SPTB | SPTB1\_HUMAN | Spectrin beta chain, erythrocytic | TSAT[79.9663]EFENVGNQPPYSR | T | 5.0 | 858047.1 | 0.0 | 858047.1 | 0.0 |
| GYPA | GLPA\_HUMAN | Isoform 2 of Glycophorin-A | SPSDVKPLPSPDTDVPLSS[79.9663]VEIENPETSDQ | S | 139.0 | 854428.663 | 0.0 | 854428.663 | 0.0 |
| TLN2 | TLN2\_HUMAN | Talin-2 | ALSDLISAT[79.9663]KGAASK | T | 2099.0 | 847150.004 | 0.0 | 835244.652 | 11905.352 |
| SPTA1 | SPTA1\_HUMAN | Spectrin alpha chain, erythrocytic 1 | DSEQVDS[79.9663]WMSR | S | 490.0 | 846025.175 | 0.0 | 782896.14 | 63129.035 |
| EPB41 | EPB41\_HUMAN | Isoform 5 of Protein 4.1 | LTSTDT[79.9663]IPK | T | 494.0 | 822383.76 | 0.0 | 822383.76 | 0.0 |
| ADD2 | ADDB\_HUMAN | Beta-adducin | SAPAS[79.9663]PVQSPAK | S | 600.0 | 820791.1 | 0.0 | 750000.5 | 70790.6 |
| HBA1 | HBA\_HUMAN | Hemoglobin subunit alpha | VLS[79.9663]PADKTNVK | S | 4.0 | 815736.017 | 0.0 | 414588.837 | 401147.18 |
| ADD2 | ADDB\_HUMAN | Beta-adducin | ETAPEEPGS[79.9663]PAK | S | 592.0 | 801221.505 | 0.0 | 721722.305 | 79499.2 |
| HBB | HBB\_HUMAN | Hemoglobin subunit beta | EFT[79.9663]PPVQAAYQK | T | 124.0 | 795999.6907 | 0.0 | 675689.2 | 120310.4907 |
| SPTA1 | SPTA1\_HUMAN | Spectrin alpha chain, erythrocytic 1 | DSEQVDSWMS[79.9663]R | S | 105.0 | 787841.367 | 0.0 | 771373.244 | 16468.123 |
| HBA1 | HBA\_HUMAN | Hemoglobin subunit alpha | M[15.9949]FLS[79.9663]FPTTK | S | 36.0 | 778631.221 | 0.0 | 729146.346 | 49484.875 |
| CARHSP1 | CHSP1\_HUMAN | Calcium-regulated heat-stable protein 1 | GNVVPS[79.9663]PLPTR | S | 41.0 | 770738.8211 | 0.0 | 349026.0411 | 421712.78 |
| ADD2 | ADDB\_HUMAN | Beta-adducin | TESVTSGPM[15.9949]SPEGSPSKS[79.9663]PSKK | S | 701.0 | 764848.43 | 0.0 | 764848.43 | 0.0 |
| SMIM1 | SMIM1\_HUMAN | Small integral membrane protein 1 | DGVS[79.9663]LGAVSS[79.9663]TEEASR | S |  | 760597.783 | 0.0 | 232599.86 | 527997.923 |
| ADD1 | ADDA\_HUMAN | Isoform 3 of Alpha-adducin | AAVVTS[79.9663]PPPTTAPHKER | S | 12.0 | 758005.801 | 758005.801 | 0.0 | 0.0 |
| ANK1 | ANK1\_HUMAN | Isoform Br21 of Ankyrin-1 | QDDATGAGQDS[79.9663]ENEVSLVSGHQR | S | 1666.0 | 744435.305 | 0.0 | 643921.055 | 100514.25 |
| KRT13 | K1C13\_HUMAN | Keratin, type I cytoskeletal 13 | M[15.9949]IGFPSSAGS[79.9663]VSPR | S | 425.0 | 710689.039 | 0.0 | 538437.773 | 172251.266 |
| ADD1 | ADDA\_HUMAN | Isoform 3 of Alpha-adducin | SRSPGS[79.9663]PVGEGTGSPPK | S | 358.0 | 707481.38 | 0.0 | 707481.38 | 0.0 |
| ANK1 | ANK1\_HUMAN | Isoform Br21 of Ankyrin-1 | ITHSPT[79.9663]VSQVTER | T | 1688.0 | 697499.688 | 0.0 | 572496.158 | 125003.53 |
| ERMAP | ERMAP\_HUMAN | Erythroid membrane-associated protein | S[79.9663]EESIVPR | S | 418.0 | 695937.826 | 0.0 | 109353.766 | 586584.06 |
| ADD2 | ADDB\_HUMAN | Beta-adducin | TESVTSGPM[15.9949]S[79.9663]PEGSPSK | S | 693.0 | 687085.0 | 0.0 | 687085.0 | 0.0 |
| ANK1 | ANK1\_HUMAN | Isoform Br21 of Ankyrin-1 | NGAS[79.9663]PNEVSSDGTTPLAIAK | S | 759.0 | 685729.9465 | 0.0 | 685729.9465 | 0.0 |
| SLC4A1 | B3AT\_HUMAN | Band 3 anion transport protein | YQSSPAKPDS[79.9663]S[79.9663]FYK | S |  | 682675.84 | 0.0 | 243943.0 | 438732.84 |
| ADD2 | ADDB\_HUMAN | Beta-adducin | TESVTSGPM[15.9949]SPEGSPS[79.9663]KSPSKK | S | 699.0 | 666165.62 | 0.0 | 666165.62 | 0.0 |
| SLC4A1 | B3AT\_HUMAN | Band 3 anion transport protein | IDAY[79.9663]MAQSR | Y | 299.0 | 658329.728 | 0.0 | 0.0 | 658329.728 |
| KEL | KELL\_HUMAN | Kell blood group glycoprotein | M[15.9949]EGGDQS[79.9663]EEEPR | S | 7.0 | 653863.32 | 0.0 | 256229.22 | 397634.1 |
| CCNY | CCNY\_HUMAN | Cyclin-Y | SAS[79.9663]ADNLTLPR | S | 326.0 | 647904.56 | 0.0 | 131170.16 | 516734.4 |
| SLC4A1 | B3AT\_HUMAN | Band 3 anion transport protein | RYQSS[79.9663]PAKPDSSFYK | S | 350.0 | 640405.346 | 0.0 | 314275.992 | 326129.354 |
| SLC4A1 | B3AT\_HUMAN | Band 3 anion transport protein | IDAYM[15.9949]AQS[79.9663]R | S | 303.0 | 637479.908 | 0.0 | 32789.523 | 604690.385 |
| FLOT1 | FLOT1\_HUMAN | Flotillin-1 | ITLVSSGS[79.9663]GTM[15.9949]GAAK | S |  | 629350.559 | 0.0 | 185296.52 | 444054.039 |
| CCNY | CCNY\_HUMAN | Cyclin-Y | S[79.9663]ASADNLTLPR | S | 324.0 | 628618.002 | 0.0 | 127477.662 | 501140.34 |
| SPTA1 | SPTA1\_HUMAN | Spectrin alpha chain, erythrocytic 1 | HQS[79.9663]LEAEVQTK | S | 96.0 | 611512.091 | 0.0 | 576590.325 | 34921.766 |
| ADD2 | ADDB\_HUMAN | Beta-adducin | TES[79.9663]VTSGPM[15.9949]SPEGSPSK | S |  | 591044.4 | 0.0 | 591044.4 | 0.0 |
| ADD2 | ADDB\_HUMAN | Beta-adducin | S[79.9663]APASPVQSPAK | S | 596.0 | 580446.22 | 0.0 | 542142.94 | 38303.28 |
| EPB41 | EPB41\_HUMAN | Isoform 5 of Protein 4.1 | RAS[79.9663]RS[79.9663]LDGAAAVDSADR | S |  | 576030.75 | 0.0 | 576030.75 | 0.0 |
| ANK1 | ANK1\_HUMAN | Isoform Br21 of Ankyrin-1 | LDQVVES[79.9663]PAIPR | S | 856.0 | 570361.577 | 0.0 | 507907.3 | 62454.277 |
| ADD2 | ADDB\_HUMAN | Beta-adducin | T[79.9663]ESVTSGPM[15.9949]SPEGSPSK | T |  | 568759.037 | 0.0 | 518757.084 | 50001.953 |
| ANK1 | ANK1\_HUMAN | Isoform Br21 of Ankyrin-1 | IT[79.9663]HSPTVS[79.9663]QVTER | T |  | 558920.006 | 0.0 | 542954.292 | 15965.714 |
| EPB41 | EPB41\_HUMAN | Isoform 5 of Protein 4.1 | SPRPT[79.9663]SAPAITQGQVAEGGVLDASAK | T | 559.0 | 555693.0 | 0.0 | 555693.0 | 0.0 |
| ACTA1 | ACTS\_HUMAN | Actin, alpha skeletal muscle | QEY[79.9663]DEAGPSIVHR | Y | 364.0 | 548658.2774 | 0.0 | 548658.2774 | 0.0 |
| EPB41 | EPB41\_HUMAN | Isoform 5 of Protein 4.1 | S[79.9663]QVSEEEGKEVESDKEK | S | 92.0 | 543855.49 | 0.0 | 528467.487 | 15388.003 |
| ADD2 | ADDB\_HUMAN | Beta-adducin | DKTESVTSGPM[15.9949]SPEGS[79.9663]PSKSPSK | S | 697.0 | 541278.92 | 0.0 | 541278.92 | 0.0 |
| SPTA1 | SPTA1\_HUMAN | Spectrin alpha chain, erythrocytic 1 | ALS[79.9663]NAANLQR | S | 261.0 | 539806.67 | 0.0 | 539806.67 | 0.0 |
| ADD1 | ADDA\_HUMAN | Isoform 3 of Alpha-adducin | AAVVT[79.9663]SPPPTTAPHKER | T | 11.0 | 525412.933 | 525412.933 | 0.0 | 0.0 |
| ADD1 | ADDA\_HUMAN | Isoform 3 of Alpha-adducin | S[79.9663]RSPGSPVGEGTGSPPK | S | 353.0 | 520324.754 | 0.0 | 520324.754 | 0.0 |
| HBA1 | HBA\_HUMAN | Hemoglobin subunit alpha | VGAHAGEY[79.9663]GAEALER | Y | 25.0 | 513857.879 | 0.0 | 165110.5725 | 348747.3065 |
| SPTA1 | SPTA1\_HUMAN | Spectrin alpha chain, erythrocytic 1 | GVS[79.9663]EETLKEFSTIYK | S | 2269.0 | 509548.22 | 0.0 | 509548.22 | 0.0 |
| ADD2 | ADDB\_HUMAN | Beta-adducin | S[79.9663]RSPSTESQLM[15.9949]SK | S |  | 504135.137 | 0.0 | 504135.137 | 0.0 |
| KRT4 | K2C4\_HUMAN | Keratin, type II cytoskeletal 4 | GAFSSVS[79.9663]M[15.9949]SGGAGR | S |  | 502959.753 | 0.0 | 431109.093 | 71850.66 |
| SLC4A1 | B3AT\_HUMAN | Band 3 anion transport protein | RYQSSPAKPDSS[79.9663]FYK | S | 357.0 | 490493.664 | 0.0 | 100333.164 | 390160.5 |
| SLC43A3 | S43A3\_HUMAN | Isoform 2 of Equilibrative nucleobase transporter 1 | ELQS[79.9663]KEFLSAK | S | 248.0 | 474918.42 | 0.0 | 159990.39 | 314928.03 |
| SPTB | SPTB1\_HUMAN | Spectrin beta chain, erythrocytic | LVAEGNLY[79.9663]SDK | Y | 1241.0 | 468552.97 | 0.0 | 468552.97 | 0.0 |
| CFL1 | COF1\_HUMAN | Cofilin-1 | n[42.0106]AS[79.9663]GVAVSDGVIK | S | 3.0 | 467484.492 | 0.0 | 304516.66 | 162967.832 |
| ADD2 | ADDB\_HUMAN | Beta-adducin | TESVTSGPM[15.9949]SPEGS[79.9663]PSKSPSKK | S | 697.0 | 456561.83 | 0.0 | 456561.83 | 0.0 |
| SPTA1 | SPTA1\_HUMAN | Spectrin alpha chain, erythrocytic 1 | LY[79.9663]EDSDDLK | Y | 588.0 | 455004.877 | 0.0 | 408517.204 | 46487.673 |
| EPB41 | EPB41\_HUMAN | Isoform 5 of Protein 4.1 | SPRPTS[79.9663]APAITQGQVAEGGVLDASAK | S | 560.0 | 444817.3 | 0.0 | 444817.3 | 0.0 |
| EPB41 | EPB41\_HUMAN | Isoform 5 of Protein 4.1 | LDGENIY[79.9663]IR | Y | 660.0 | 441926.55 | 0.0 | 441926.55 | 0.0 |
| CD44 | CD44\_HUMAN | Isoform 10 of CD44 antigen | S[79.9663]QEM[15.9949]VHLVNK | S |  | 435811.415 | 0.0 | 139314.06 | 296497.355 |
| SLC4A1 | B3AT\_HUMAN | Band 3 anion transport protein | Y[79.9663]QSSPAKPDSSFYK | Y | 347.0 | 429313.456 | 0.0 | 20462.816 | 408850.64 |
| PIEZO1 | PIEZ1\_HUMAN | Piezo-type mechanosensitive ion channel component 1 | QDAVS[79.9663]GTPLLR | S | 732.0 | 425726.77 | 0.0 | 71275.72 | 354451.05 |
| EPB41 | EPB41\_HUMAN | Isoform 5 of Protein 4.1 | S[79.9663]PRPTSAPAITQGQVAEGGVLDASAK | S | 555.0 | 423911.25 | 0.0 | 423911.25 | 0.0 |
| BAG6 | BAG6\_HUMAN | Isoform 2 of Large proline-rich protein BAG6 | APPQTHLPSGASSGTGSASATHGGGS[79.9663]PPGTR | S | 113.0 | 420206.6 | 420206.6 | 0.0 | 0.0 |
| EPB41 | EPB41\_HUMAN | Isoform 5 of Protein 4.1 | SLDGAAAVDS[79.9663]ADRSPRPTSAPAITQGQVAEGGVLDASAK | S | 551.0 | 408350.841 | 0.0 | 408350.841 | 0.0 |
| SLC4A1 | B3AT\_HUMAN | Band 3 anion transport protein | RYQS[79.9663]SPAKPDSSFYK | S | 349.0 | 408347.41 | 0.0 | 350645.0 | 57702.41 |
| SLC4A1 | B3AT\_HUMAN | Band 3 anion transport protein | YQSSPAKPDSSFY[79.9663]K | Y | 359.0 | 405894.633 | 0.0 | 131074.085 | 274820.548 |
| OR51D1 | O51D1\_HUMAN | Olfactory receptor 51D1 | VNVVYGLFIILS[79.9663]VMGVDSLFIGFSYILILWAVLELSSR | S | 218.0 | 401451.34 | 0.0 | 0.0 | 401451.34 |
| UBR4 | UBR4\_HUMAN | Isoform 2 of E3 ubiquitin-protein ligase UBR4 | HVTLPSS[79.9663]PR | S | 620.0 | 399182.38 | 0.0 | 0.0 | 399182.38 |
| SMIM1 | SMIM1\_HUMAN | Small integral membrane protein 1 | M[15.9949]QPQES[79.9663]HVHYSR | S | 6.0 | 397669.193 | 0.0 | 136543.27 | 261125.923 |
| TMBIM1 | LFG3\_HUMAN | Protein lifeguard 3 | SNPS[79.9663]APPPYEDR | S | 5.0 | 392576.2 | 0.0 | 0.0 | 392576.2 |
| SLC43A2 | LAT4\_HUMAN | Isoform 3 of Large neutral amino acids transporter small subunit 4 | RLS[79.9663]VGSSM[15.9949]R | S |  | 387672.2 | 0.0 | 0.0 | 387672.2 |
| SPTBN4 | SPTN4\_HUMAN | Isoform 4 of Spectrin beta chain, non-erythrocytic 4 | IHS[79.9663]LENVDK | S | 119.0 | 384708.469 | 0.0 | 384708.469 | 0.0 |
| ADD1 | ADDA\_HUMAN | Isoform 3 of Alpha-adducin | S[79.9663]RSPGS[79.9663]PVGEGTGSPPK | S |  | 380676.283 | 0.0 | 380676.283 | 0.0 |
| XK | XK\_HUMAN | Endoplasmic reticulum membrane adapter protein XK | EDLQS[79.9663]SRDRDETPSSSK | S | 415.0 | 378770.2 | 0.0 | 0.0 | 378770.2 |
| ADD2 | ADDB\_HUMAN | Beta-adducin | DKTESVTSGPM[15.9949]SPEGS[79.9663]PSK | S | 617.0 | 370234.237 | 0.0 | 357121.854 | 13112.383 |
| TNS1 | TENS1\_HUMAN | Tensin-1 | VATTPGS[79.9663]PSLGR | S | 1373.0 | 368391.987 | 0.0 | 157400.75 | 210991.237 |
| ADD2 | ADDB\_HUMAN | Beta-adducin | TESVTSGPM[15.9949]SPEGS[79.9663]PSKSPSK | S | 697.0 | 364298.373 | 0.0 | 338283.75 | 26014.623 |
| EPB41 | EPB41\_HUMAN | Isoform 5 of Protein 4.1 | SQVSEEEGKEVES[79.9663]DKEK | S | 104.0 | 362647.54 | 0.0 | 362647.54 | 0.0 |
| SPTB | SPTB1\_HUMAN | Spectrin beta chain, erythrocytic | ET[79.9663]WLSENQR | T | 436.0 | 361007.14 | 0.0 | 361007.14 | 0.0 |
| SPTB | SPTB1\_HUMAN | Spectrin beta chain, erythrocytic | T[79.9663]SPVSLWS[79.9663]R | T |  | 356955.5 | 0.0 | 356955.5 | 0.0 |
| SPTB | SPTB1\_HUMAN | Spectrin beta chain, erythrocytic | QIAERPAEETGPQEEEGETAGEAPVS[79.9663]HHAATER | S | 2105.0 | 356668.77 | 0.0 | 356668.77 | 0.0 |
| SPTA1 | SPTA1\_HUMAN | Spectrin alpha chain, erythrocytic 1 | EDLVS[79.9663]SWEHIR | S | 350.0 | 353160.16 | 0.0 | 353160.16 | 0.0 |
| ADD2 | ADDB\_HUMAN | Beta-adducin | TESVTSGPMSPEGS[79.9663]PSK | S | 617.0 | 346525.8 | 0.0 | 346525.8 | 0.0 |
| HBA1 | HBA\_HUMAN | Hemoglobin subunit alpha | VLS[79.9663]PADK | S | 4.0 | 345158.72 | 0.0 | 204556.84 | 140601.88 |
| ADD2 | ADDB\_HUMAN | Beta-adducin | SPLVSPSKS[79.9663]LEEGTKK | S | 621.0 | 338627.4 | 0.0 | 338627.4 | 0.0 |
| ADD1 | ADDA\_HUMAN | Isoform 3 of Alpha-adducin | S[79.9663]PGSPVGEGT[79.9663]GSPPK | S |  | 338619.45 | 0.0 | 338619.45 | 0.0 |
| EPB41 | EPB41\_HUMAN | Isoform 5 of Protein 4.1 | S[79.9663]LDGAAAVDSADRSPRPTSAPAITQGQVAEGGVLDASAK | S | 542.0 | 337458.4203 | 0.0 | 337458.4203 | 0.0 |
| DMTN | DEMA\_HUMAN | Isoform 4 of Dematin | TPFHTS[79.9663]LHQGTSK | S | 279.0 | 333700.384 | 0.0 | 333700.384 | 0.0 |
| SPTB | SPTB1\_HUMAN | Spectrin beta chain, erythrocytic | QIAERPAEETGPQEEEGETAGEAPVSHHAAT[79.9663]ER | T | 2110.0 | 330870.78 | 0.0 | 330870.78 | 0.0 |
| PI4K2A | P4K2A\_HUMAN | Phosphatidylinositol 4-kinase type 2-alpha | VAAAAGSGPS[79.9663]PPGSPGHDR | S | 47.0 | 320136.224 | 0.0 | 0.0 | 320136.224 |
| HBB | HBB\_HUMAN | Hemoglobin subunit beta | VHLT[79.9663]PEEK | T | 5.0 | 318444.485 | 0.0 | 241797.345 | 76647.14 |
| SYNJ1 | SYNJ1\_HUMAN | Synaptojanin-1 | ASAGRLT[79.9663]PESQSK | T | 1220.0 | 318077.278 | 300300.1 | 17777.178 | 0.0 |
| PGK1 | PGK1\_HUMAN | Phosphoglycerate kinase 1 | ALES[79.9663]PERPFLAILGGAK | S | 203.0 | 312342.3 | 312342.3 | 0.0 | 0.0 |
| ADD1 | ADDA\_HUMAN | Isoform 3 of Alpha-adducin | SRS[79.9663]PGS[79.9663]PVGEGTGSPPK | S |  | 307359.88 | 0.0 | 307359.88 | 0.0 |
| ADD1 | ADDA\_HUMAN | Isoform 3 of Alpha-adducin | S[79.9663]PGSPVGEGTGS[79.9663]PPK | S |  | 304373.72 | 0.0 | 304373.72 | 0.0 |
| BSG | BASI\_HUMAN | Isoform 2 of Basigin | KPEDVLDDDDAGS[79.9663]APLK | S | 362.0 | 302355.25 | 0.0 | 302355.25 | 0.0 |
| TMCC2 | TMCC2\_HUMAN | Isoform 2 of Transmembrane and coiled-coil domains protein 2 | ALSGS[79.9663]ATLVSSPK | S | 464.0 | 300326.71 | 0.0 | 59889.61 | 240437.1 |
| EPB41 | EPB41\_HUMAN | Isoform 5 of Protein 4.1 | KEDEPPEQAEPEPT[79.9663]EAWK | T | 611.0 | 299204.2 | 0.0 | 299204.2 | 0.0 |
| SPTAN1 | SPTN1\_HUMAN | Isoform 3 of Spectrin alpha chain, non-erythrocytic 1 | ADVVES[79.9663]WIGEK | S | 1990.0 | 299030.3 | 0.0 | 299030.3 | 0.0 |
| ADD1 | ADDA\_HUMAN | Isoform 3 of Alpha-adducin | KQKGS[79.9663]EENLDEAR | S | 586.0 | 298907.607 | 0.0 | 273670.705 | 25236.902 |
| ANK1 | ANK1\_HUMAN | Isoform Br21 of Ankyrin-1 | EADAATS[79.9663]FLR | S | 15.0 | 298017.774 | 0.0 | 279328.44 | 18689.334 |
| EPB41 | EPB41\_HUMAN | Isoform 5 of Protein 4.1 | PT[79.9663]SAPAITQGQVAEGGVLDASAK | T | 559.0 | 297680.5002 | 0.0 | 297680.5002 | 0.0 |
| GYPA | GLPA\_HUMAN | Isoform 2 of Glycophorin-A | KSPSDVKPLPSPDTDVPLSSVEIENPET[79.9663]SDQ | T | 147.0 | 296189.7896 | 0.0 | 296189.7896 | 0.0 |
| ADD3 | ADDG\_HUMAN | Isoform 1 of Gamma-adducin | IEEVLSPEGS[79.9663]PSK | S | 677.0 | 296049.474 | 0.0 | 288480.487 | 7568.987 |
| ADD2 | ADDB\_HUMAN | Beta-adducin | n[42.0106]SEETVPEAAS[79.9663]PPPPQGQPYFDR | S | 11.0 | 295993.94 | 0.0 | 295993.94 | 0.0 |
| GYPA | GLPA\_HUMAN | Isoform 2 of Glycophorin-A | SPSDVKPLPSPDT[79.9663]DVPLSSVEIENPETSDQ | T | 133.0 | 295037.382 | 0.0 | 295037.382 | 0.0 |
| SPTA1 | SPTA1\_HUMAN | Spectrin alpha chain, erythrocytic 1 | LS[79.9663]ESHPDATEDLQR | S | 1250.0 | 293589.797 | 0.0 | 293589.797 | 0.0 |
| DMTN | DEMA\_HUMAN | Isoform 4 of Dematin | TPFHTSLHQGT[79.9663]SK | T | 284.0 | 290308.44 | 0.0 | 290308.44 | 0.0 |
| ADD1 | ADDA\_HUMAN | Isoform 3 of Alpha-adducin | SPGS[79.9663]PVGEGT[79.9663]GSPPK | S |  | 290294.7 | 0.0 | 290294.7 | 0.0 |
| SPTA1 | SPTA1\_HUMAN | Spectrin alpha chain, erythrocytic 1 | EDLVSS[79.9663]WEHIR | S | 351.0 | 285137.03 | 0.0 | 285137.03 | 0.0 |
| ADD1 | ADDA\_HUMAN | Isoform 3 of Alpha-adducin | S[79.9663]PGS[79.9663]PVGEGTGSPPK | S |  | 280928.8 | 0.0 | 280928.8 | 0.0 |
| ADD2 | ADDB\_HUMAN | Beta-adducin | GLSQM[15.9949]TTSADT[79.9663]DVDTSK | T | 675.0 | 280037.371 | 0.0 | 237152.504 | 42884.867 |
| SPTB | SPTB1\_HUMAN | Spectrin beta chain, erythrocytic | AWES[79.9663]LEEAEYR | S | 398.0 | 276452.602 | 0.0 | 276452.602 | 0.0 |
| SPTA1 | SPTA1\_HUMAN | Spectrin alpha chain, erythrocytic 1 | AY[79.9663]FLDGSLLK | Y | 2179.0 | 275113.47 | 0.0 | 275113.47 | 0.0 |
| MAP2K2 | MP2K2\_HUMAN | Dual specificity mitogen-activated protein kinase kinase 2 | LNQPGT[79.9663]PTR | T | 394.0 | 271065.596 | 0.0 | 215007.6 | 56057.996 |
| STK11 | STK11\_HUMAN | Isoform 2 of Serine/threonine-protein kinase STK11 | IDS[79.9663]TEVIYQPR | S | 31.0 | 270335.157 | 0.0 | 115004.914 | 155330.243 |
| C1orf198 | CA198\_HUMAN | Uncharacterized protein C1orf198 | S[79.9663]SSLDALGPTR | S | 173.0 | 264583.529 | 0.0 | 225297.236 | 39286.293 |
| DENND1A | DEN1A\_HUMAN | DENN domain-containing protein 1A | TSVPS[79.9663]PEQPQPYR | S | 523.0 | 259147.06 | 0.0 | 182028.45 | 77118.61 |
| ADD1 | ADDA\_HUMAN | Isoform 3 of Alpha-adducin | SRS[79.9663]PGSPVGEGTGSPPK | S | 355.0 | 256419.67 | 0.0 | 256419.67 | 0.0 |
| PIEZO1 | PIEZ1\_HUMAN | Piezo-type mechanosensitive ion channel component 1 | QDAVSGT[79.9663]PLLR | T | 734.0 | 254760.9 | 0.0 | 0.0 | 254760.9 |
| PGM2L1 | PGM2L\_HUMAN | Glucose 1,6-bisphosphate synthase | AVAGVM[15.9949]ITAS[79.9663]HNR | S | 175.0 | 254379.137 | 0.0 | 216583.0 | 37796.137 |
| TAS2R8 | TA2R8\_HUMAN | Taste receptor type 2 member 8 | ISTVDYILTNLVIARIC[57.0215]LIS[79.9663]VM[15.9949]VVNGIVIVLNPDVYTK | S |  | 251256.45 | 0.0 | 0.0 | 251256.45 |
| ADD2 | ADDB\_HUMAN | Beta-adducin | GLSQM[15.9949]TTSADT[79.9663]DVDTSKDK | T | 675.0 | 250376.069 | 0.0 | 239305.861 | 11070.208 |
| SLC4A1 | B3AT\_HUMAN | Band 3 anion transport protein | YQS[79.9663]SPAKPDS[79.9663]SFYK | S |  | 240859.885 | 0.0 | 0.0 | 240859.885 |
| ADD2 | ADDB\_HUMAN | Beta-adducin | SRS[79.9663]PSTESQLM[15.9949]SK | S |  | 237616.352 | 0.0 | 219806.43 | 17809.922 |
| EPB41 | EPB41\_HUMAN | Isoform 5 of Protein 4.1 | HHAS[79.9663]ISELK | S | 684.0 | 236403.22 | 0.0 | 236403.22 | 0.0 |
| KRT2 | K22E\_HUMAN | Keratin, type II cytoskeletal 2 epidermal | S[79.9663]LVGLGGTK | S | 62.0 | 235721.0 | 0.0 | 0.0 | 235721.0 |
| SPTA1 | SPTA1\_HUMAN | Spectrin alpha chain, erythrocytic 1 | HFDENLT[79.9663]GR | T | 2288.0 | 234360.06 | 0.0 | 234360.06 | 0.0 |
| FLOT1 | FLOT1\_HUMAN | Flotillin-1 | S[79.9663]PPVM[15.9949]VAGGR | S |  | 231494.206 | 0.0 | 70324.34 | 161169.866 |
| SPTB | SPTB1\_HUMAN | Spectrin beta chain, erythrocytic | EQIYS[79.9663]SLDYGK | S | 661.0 | 229505.23 | 0.0 | 229505.23 | 0.0 |
| TSC22D4 | T22D4\_HUMAN | TSC22 domain family protein 4 | AEKPPLSASSPQQRPPEPETGESAGT[79.9663]SR | T | 206.0 | 225812.08 | 225812.08 | 0.0 | 0.0 |
| SPTA1 | SPTA1\_HUMAN | Spectrin alpha chain, erythrocytic 1 | QDTLDAS[79.9663]LQSFQQER | S | 1976.0 | 225676.825 | 0.0 | 216703.977 | 8972.848 |
| GYPA | GLPA\_HUMAN | Isoform 2 of Glycophorin-A | KSPSDVKPLPSPDTDVPLSSVEIENPETS[79.9663]DQ | S | 148.0 | 224170.442 | 0.0 | 224170.442 | 0.0 |
| TMEM9 | TMEM9\_HUMAN | Proton-transporting V-type ATPase complex assembly regulator TMEM9 | S[79.9663]M[15.9949]AAAAASLGGPR | S |  | 221547.281 | 0.0 | 55498.445 | 166048.836 |
| ECPAS | ECM29\_HUMAN | Proteasome adapter and scaffold protein ECM29 | AQGAIAMAS[79.9663]IAK | S | 1553.0 | 220560.13 | 0.0 | 220560.13 | 0.0 |
| SLC4A1 | B3AT\_HUMAN | Band 3 anion transport protein | YQSS[79.9663]PAKPDSS[79.9663]FYK | S |  | 220052.44 | 0.0 | 0.0 | 220052.44 |
| SLC29A1 | S29A1\_HUMAN | Isoform 2 of Equilibrative nucleoside transporter 1 | LDLIS[79.9663]KGEEPR | S | 254.0 | 219936.846 | 0.0 | 119093.286 | 100843.56 |
| DMTN | DEMA\_HUMAN | Isoform 4 of Dematin | RGAEEEEEEEDDDS[79.9663]GEEM[15.9949]KALR | S |  | 217416.47 | 0.0 | 217416.47 | 0.0 |
| ADD2 | ADDB\_HUMAN | Beta-adducin | DKTES[79.9663]VTSGPM[15.9949]SPEGSPSK | S |  | 216437.6 | 0.0 | 216437.6 | 0.0 |
| SLC4A1 | B3AT\_HUMAN | Band 3 anion transport protein | YQSSPAKPDS[79.9663]SFY[79.9663]K | S |  | 216342.52 | 0.0 | 216342.52 | 0.0 |
| DBNL | DBNL\_HUMAN | Isoform 2 of Drebrin-like protein | AM[15.9949]S[79.9663]TTSISSPQPGK | S | 269.0 | 215389.224 | 215389.224 | 0.0 | 0.0 |
| DMTN | DEMA\_HUMAN | Isoform 4 of Dematin | EEM[15.9949]EKS[79.9663]LPIR | S | 260.0 | 215021.28 | 0.0 | 215021.28 | 0.0 |
| TMBIM1 | LFG3\_HUMAN | Protein lifeguard 3 | n[42.0106]SNPS[79.9663]APPPYEDR | S | 5.0 | 212465.16 | 0.0 | 0.0 | 212465.16 |
| KLC1 | KLC1\_HUMAN | Isoform J of Kinesin light chain 1 | ALSAS[79.9663]HTDLAH | S |  | 208858.84 | 208858.84 | 0.0 | 0.0 |
| ADD2 | ADDB\_HUMAN | Beta-adducin | ADEVEKSS[79.9663]SGM[15.9949]PIR | S |  | 206673.378 | 46775.664 | 159897.714 | 0.0 |
| KLC1 | KLC1\_HUMAN | Isoform J of Kinesin light chain 1 | ALS[79.9663]ASHTDLAH | S |  | 205908.69 | 205908.69 | 0.0 | 0.0 |
| ACTB | ACTB\_HUMAN | Actin, cytoplasmic 1 | QEY[79.9663]DESGPSIVHR | Y | 362.0 | 205903.867 | 0.0 | 205903.867 | 0.0 |
| ANK1 | ANK1\_HUMAN | Isoform Br21 of Ankyrin-1 | RQDDATGAGQDS[79.9663]ENEVSLVSGHQR | S | 1666.0 | 205269.98 | 0.0 | 205269.98 | 0.0 |
| ADD2 | ADDB\_HUMAN | Beta-adducin | SRSPS[79.9663]TESQLM[15.9949]SK | S |  | 204228.8 | 0.0 | 204228.8 | 0.0 |
| SPTB | SPTB1\_HUMAN | Spectrin beta chain, erythrocytic | LVAQDNFGY[79.9663]DLAAVEAAK | Y | 452.0 | 203792.14 | 0.0 | 203792.14 | 0.0 |
| SPTA1 | SPTA1\_HUMAN | Spectrin alpha chain, erythrocytic 1 | DSEQVDS[79.9663]WM[15.9949]SR | S |  | 203075.177 | 0.0 | 184272.136 | 18803.041 |
| PSMA5 | PSA5\_HUMAN | Proteasome subunit alpha type-5 | IT[79.9663]SPLM[15.9949]EPSSIEK | T |  | 202349.003 | 0.0 | 145897.27 | 56451.733 |
| SPTB | SPTB1\_HUMAN | Spectrin beta chain, erythrocytic | QIAERPAEETGPQEEEGETAGEAPVS[79.9663]HHAAT[79.9663]ER | S |  | 201321.28 | 0.0 | 201321.28 | 0.0 |
| SPTA1 | SPTA1\_HUMAN | Spectrin alpha chain, erythrocytic 1 | LGDY[79.9663]ANLK | Y | 1499.0 | 200835.27 | 0.0 | 200835.27 | 0.0 |
| GYPA | GLPA\_HUMAN | Isoform 2 of Glycophorin-A | KSPSDVKPLPSPDTDVPLS[79.9663]SVEIENPETSDQ | S | 138.0 | 199493.475 | 0.0 | 199493.475 | 0.0 |
| GYPA | GLPA\_HUMAN | Isoform 2 of Glycophorin-A | SPSDVKPLPS[79.9663]PDTDVPLSSVEIENPETSDQ | S | 130.0 | 195524.481 | 0.0 | 195524.481 | 0.0 |
| SPTB | SPTB1\_HUMAN | Spectrin beta chain, erythrocytic | Y[79.9663]HQGINAEIETR | Y | 1944.0 | 192327.704 | 0.0 | 192327.704 | 0.0 |
| SPTB | SPTB1\_HUMAN | Spectrin beta chain, erythrocytic | EQIY[79.9663]SSLDYGK | Y | 660.0 | 191934.53 | 0.0 | 191934.53 | 0.0 |
| EPB41 | EPB41\_HUMAN | Isoform 5 of Protein 4.1 | RLS[79.9663]THSPFR | S | 709.0 | 191681.47 | 0.0 | 191681.47 | 0.0 |
| HSPB1 | HSPB1\_HUMAN | Heat shock protein beta-1 | QLS[79.9663]SGVSEIR | S | 82.0 | 187541.864 | 0.0 | 130785.36 | 56756.504 |
| NRBP1 | NRBP\_HUMAN | Nuclear receptor-binding protein | T[79.9663]PTPEPAEVETR | T | 431.0 | 185622.0 | 0.0 | 141842.53 | 43779.47 |
| TMEM237 | TM237\_HUMAN | Isoform 2 of Transmembrane protein 237 | RPS[79.9663]EGNEPSTK | S | 62.0 | 178224.921 | 0.0 | 29998.691 | 148226.23 |
| SPTA1 | SPTA1\_HUMAN | Spectrin alpha chain, erythrocytic 1 | GVS[79.9663]EETLK | S | 2269.0 | 178147.03 | 0.0 | 0.0 | 178147.03 |
| C1orf198 | CA198\_HUMAN | Uncharacterized protein C1orf198 | SSS[79.9663]LDALGPTR | S | 175.0 | 177917.96 | 0.0 | 177917.96 | 0.0 |
| ANK1 | ANK1\_HUMAN | Isoform Br21 of Ankyrin-1 | ITHSPTVSQVTERS[79.9663]QDR | S | 1696.0 | 177072.94 | 0.0 | 177072.94 | 0.0 |
| CSNK1A1 | KC1A\_HUMAN | Isoform 3 of Casein kinase I isoform alpha | AAQQAASSSGQGQQAQT[79.9663]PTGK | T | 321.0 | 173475.12 | 173475.12 | 0.0 | 0.0 |
| EPB41 | EPB41\_HUMAN | Isoform 5 of Protein 4.1 | LS[79.9663]MYGVDLHK | S | 394.0 | 172906.5514 | 0.0 | 172906.5514 | 0.0 |
| ADD2 | ADDB\_HUMAN | Beta-adducin | GLSQM[15.9949]TTS[79.9663]ADTDVDTSKDK | S | 672.0 | 172840.92 | 0.0 | 172840.92 | 0.0 |
| CCRL2 | CCRL2\_HUMAN | Isoform 2 of C-C chemokine receptor-like 2 | EEPDHS[79.9663]TEV | S | 341.0 | 169564.48 | 0.0 | 0.0 | 169564.48 |
| ADD2 | ADDB\_HUMAN | Beta-adducin | KLELDGEKETAPEEPGS[79.9663]PAK | S | 592.0 | 167899.2 | 0.0 | 167899.2 | 0.0 |
| SPTB | SPTB1\_HUMAN | Spectrin beta chain, erythrocytic | WIS[79.9663]AM[15.9949]EDQLR | S |  | 166917.38 | 0.0 | 166917.38 | 0.0 |
| EPB41 | EPB41\_HUMAN | Isoform 5 of Protein 4.1 | SLDGAAAVDSADRS[79.9663]PRPTSAPAITQGQVAEGGVLDASAK | S | 555.0 | 160541.84 | 0.0 | 160541.84 | 0.0 |
| DMTN | DEMA\_HUMAN | Isoform 4 of Dematin | TPFHT[79.9663]SLHQGTSK | T | 278.0 | 160321.27 | 0.0 | 160321.27 | 0.0 |
| EPB41 | EPB41\_HUMAN | Isoform 5 of Protein 4.1 | IEVKEES[79.9663]PQSK | S | 188.0 | 160106.363 | 0.0 | 160106.363 | 0.0 |
| ANK1 | ANK1\_HUMAN | Isoform Br21 of Ankyrin-1 | ITHS[79.9663]PTVSQVTERSQDR | S | 1686.0 | 157430.39 | 0.0 | 157430.39 | 0.0 |
| TMEM237 | TM237\_HUMAN | Isoform 2 of Transmembrane protein 237 | NTPASAS[79.9663]LEGLAQTAGR | S | 49.0 | 153796.58 | 0.0 | 0.0 | 153796.58 |
| ACTG2 | ACTH\_HUMAN | Actin, gamma-enteric smooth muscle | PEY[79.9663]DEAGPSIVHR | Y | 363.0 | 153488.887 | 0.0 | 153488.887 | 0.0 |
| EPB41 | EPB41\_HUMAN | Isoform 5 of Protein 4.1 | TQT[79.9663]VTISDNANAVK | T | 736.0 | 151536.72 | 0.0 | 151536.72 | 0.0 |
| MICALL2 | MILK2\_HUMAN | MICAL-like protein 2 | LPS[79.9663]PAPAR | S | 143.0 | 145360.449 | 0.0 | 83238.164 | 62122.285 |
| ADD2 | ADDB\_HUMAN | Beta-adducin | LELDGEKETAPEEPGS[79.9663]PAK | S | 592.0 | 145096.83 | 0.0 | 145096.83 | 0.0 |
| SPTA1 | SPTA1\_HUMAN | Spectrin alpha chain, erythrocytic 1 | LT[79.9663]LSHPSDAPQIQEM[15.9949]K | T |  | 144893.874 | 0.0 | 144893.874 | 0.0 |
| RAN | RAN\_HUMAN | GTP-binding nuclear protein Ran | AKS[79.9663]IVFHR | S | 135.0 | 144556.335 | 107800.03 | 36756.305 | 0.0 |
| ZDHHC5 | ZDHC5\_HUMAN | Isoform 2 of Palmitoyltransferase ZDHHC5 | DSPPT[79.9663]PTM[15.9949]YK | T |  | 142719.738 | 0.0 | 0.0 | 142719.738 |
| SMIM1 | SMIM1\_HUMAN | Small integral membrane protein 1 | DGVS[79.9663]LGAVSSTEEASR | S | 22.0 | 142648.44 | 0.0 | 0.0 | 142648.44 |
| SLC4A1 | B3AT\_HUMAN | Band 3 anion transport protein | T[79.9663]YNYNVLM[15.9949]VPK | T |  | 141547.2983 | 0.0 | 77335.2 | 64212.0983 |
| TBC1D10B | TB10B\_HUMAN | TBC1 domain family member 10B | AAGGAPS[79.9663]PPPPVR | S | 678.0 | 140229.884 | 0.0 | 44090.434 | 96139.45 |
| SPTA1 | SPTA1\_HUMAN | Spectrin alpha chain, erythrocytic 1 | ALATS[79.9663]RYEK | S | 361.0 | 139895.03 | 0.0 | 139895.03 | 0.0 |
| EPB41 | EPB41\_HUMAN | Isoform 5 of Protein 4.1 | HHASIS[79.9663]ELKK | S | 686.0 | 139847.94 | 0.0 | 139847.94 | 0.0 |
| ADD2 | ADDB\_HUMAN | Beta-adducin | DKTESVTSGPMSPEGS[79.9663]PSK | S | 617.0 | 139118.12 | 0.0 | 139118.12 | 0.0 |
| RABEP1 | RABE1\_HUMAN | Isoform 2 of Rab GTPase-binding effector protein 1 | AQS[79.9663]TDSLGTSGSLQSK | S | 407.0 | 138849.551 | 131551.573 | 7297.978 | 0.0 |
| ADD2 | ADDB\_HUMAN | Beta-adducin | SRSPS[79.9663]TES[79.9663]QLM[15.9949]SK | S |  | 138588.78 | 0.0 | 138588.78 | 0.0 |
| LPIN2 | LPIN2\_HUMAN | Phosphatidate phosphatase LPIN2 | T[79.9663]ATITPSENTHFR | T | 287.0 | 138345.02 | 0.0 | 138345.02 | 0.0 |
| SPTA1 | SPTA1\_HUMAN | Spectrin alpha chain, erythrocytic 1 | LADDEDY[79.9663]KDIQNLK | Y | 610.0 | 135330.233 | 0.0 | 135330.233 | 0.0 |
| ADD2 | ADDB\_HUMAN | Beta-adducin | SPSTES[79.9663]QLMSK | S | 535.0 | 134488.526 | 0.0 | 134488.526 | 0.0 |
| KRT4 | K2C4\_HUMAN | Keratin, type II cytoskeletal 4 | IISTTT[79.9663]LNK | T | 515.0 | 132709.77 | 0.0 | 132709.77 | 0.0 |
| HRNR | HORN\_HUMAN | Hornerin | QSLGHGQHGSGSGQSPS[79.9663]PSR | S | 542.0 | 132655.162 | 0.0 | 82064.85 | 50590.312 |
| EPB41 | EPB41\_HUMAN | Isoform 5 of Protein 4.1 | LTS[79.9663]TDTIPK | S | 491.0 | 131094.1 | 0.0 | 131094.1 | 0.0 |
| ERMAP | ERMAP\_HUMAN | Erythroid membrane-associated protein | SEES[79.9663]IVPRPEGK | S | 421.0 | 130870.328 | 0.0 | 69835.814 | 61034.514 |
| ADD2 | ADDB\_HUMAN | Beta-adducin | TESVTSGPMSPEGSPSKS[79.9663]PSK | S | 617.0 | 129393.74 | 0.0 | 129393.74 | 0.0 |
| EPB41 | EPB41\_HUMAN | Isoform 5 of Protein 4.1 | EQHPDM[15.9949]S[79.9663]VTK | S | 849.0 | 128842.24 | 0.0 | 128842.24 | 0.0 |
| EPB41 | EPB41\_HUMAN | Isoform 5 of Protein 4.1 | SQVS[79.9663]EEEGKEVESDK | S | 95.0 | 128385.383 | 0.0 | 128385.383 | 0.0 |
| ADD2 | ADDB\_HUMAN | Beta-adducin | S[79.9663]APAS[79.9663]PVQSPAK | S |  | 128231.042 | 0.0 | 128231.042 | 0.0 |
| ADD1 | ADDA\_HUMAN | Isoform 3 of Alpha-adducin | T[79.9663]STSAVPNLFVPLNTNPK | T | 480.0 | 127417.78 | 0.0 | 127417.78 | 0.0 |
| ADD2 | ADDB\_HUMAN | Beta-adducin | TESVT[79.9663]SGPM[15.9949]SPEGSPSK | T |  | 127065.34 | 0.0 | 127065.34 | 0.0 |
| ADD2 | ADDB\_HUMAN | Beta-adducin | DKT[79.9663]ESVTSGPM[15.9949]SPEGSPSK | T |  | 124576.766 | 0.0 | 124576.766 | 0.0 |
| ADD2 | ADDB\_HUMAN | Beta-adducin | T[79.9663]ESVTSGPMSPEGSPSK | T | 684.0 | 123249.875 | 0.0 | 123249.875 | 0.0 |
| ARHGAP1 | RHG01\_HUMAN | Rho GTPase-activating protein 1 | SSS[79.9663]PELVTHLK | S | 51.0 | 123016.058 | 0.0 | 95428.74 | 27587.318 |
| SLC4A1 | B3AT\_HUMAN | Band 3 anion transport protein | TYNY[79.9663]NVLM[15.9949]VPK | Y |  | 122675.7 | 0.0 | 73435.16 | 49240.54 |
| KRT13 | K1C13\_HUMAN | Keratin, type I cytoskeletal 13 | M[15.9949]IGFPS[79.9663]SAGSVSPR | S | 421.0 | 120872.036 | 0.0 | 28118.086 | 92753.95 |
| ANK1 | ANK1\_HUMAN | Isoform Br21 of Ankyrin-1 | ITHSPT[79.9663]VS[79.9663]QVTER | T |  | 120233.48 | 0.0 | 120233.48 | 0.0 |
| TLN2 | TLN2\_HUMAN | Talin-2 | ALSDLISATKGAAS[79.9663]K | S | 107.0 | 119818.24 | 0.0 | 119818.24 | 0.0 |
| NAA15 | NAA15\_HUMAN | N-alpha-acetyltransferase 15, NatA auxiliary subunit | IYEEAWT[79.9663]K | T | 229.0 | 119574.19 | 0.0 | 0.0 | 119574.19 |
| KRT1 | K2C1\_HUMAN | Keratin, type II cytoskeletal 1 | S[79.9663]LVNLGGSK | S | 66.0 | 119034.05 | 0.0 | 0.0 | 119034.05 |
| DMTN | DEMA\_HUMAN | Isoform 4 of Dematin | VFAM[15.9949]S[79.9663]PEEFGK | S | 383.0 | 118460.037 | 0.0 | 118460.037 | 0.0 |
| SPTA1 | SPTA1\_HUMAN | Spectrin alpha chain, erythrocytic 1 | DSEQVDSWM[15.9949]S[79.9663]R | S | 105.0 | 118057.56 | 0.0 | 118057.56 | 0.0 |
| CSNK1A1 | KC1A\_HUMAN | Isoform 3 of Casein kinase I isoform alpha | AAQQAASS[79.9663]SGQGQQAQTPTGK | S | 312.0 | 117665.81 | 117665.81 | 0.0 | 0.0 |
| SLC4A1 | B3AT\_HUMAN | Band 3 anion transport protein | YHPDVPY[79.9663]VK | Y | 824.0 | 117540.2 | 0.0 | 0.0 | 117540.2 |
| TBCA | TBCA\_HUMAN | Tubulin-specific chaperone A | AEDGENY[79.9663]DIKK | Y | 48.0 | 116294.09 | 0.0 | 0.0 | 116294.09 |
| ARHGAP1 | RHG01\_HUMAN | Rho GTPase-activating protein 1 | S[79.9663]SSPELVTHLK | S | 49.0 | 116286.76 | 0.0 | 116286.76 | 0.0 |
| BRSK1 | BRSK1\_HUMAN | Isoform 2 of Serine/threonine-protein kinase BRSK1 | VDS[79.9663]PM[15.9949]LSR | S |  | 116204.305 | 0.0 | 0.0 | 116204.305 |
| DMTN | DEMA\_HUMAN | Isoform 4 of Dematin | STS[79.9663]PPPSPEVWADSR | S | 92.0 | 114606.82 | 0.0 | 114606.82 | 0.0 |
| HSP90AB1 | HS90B\_HUMAN | Heat shock protein HSP 90-beta | IEDVGS[79.9663]DEEDDSGKDKK | S | 255.0 | 114508.49 | 0.0 | 98136.625 | 16371.865 |
| ADD3 | ADDG\_HUMAN | Isoform 1 of Gamma-adducin | IEEVLSPEGS[79.9663]PSKSPSK | S | 677.0 | 114462.561 | 0.0 | 44633.616 | 69828.945 |
| SPTA1 | SPTA1\_HUMAN | Spectrin alpha chain, erythrocytic 1 | KES[79.9663]LNEAQK | S | 1284.0 | 113987.647 | 0.0 | 49254.35 | 64733.297 |
| CYBRD1 | CYBR1\_HUMAN | Plasma membrane ascorbate-dependent reductase CYBRD1 | NLALDEAGQRS[79.9663]TM[15.9949] | S |  | 113926.14 | 0.0 | 38466.58 | 75459.56 |
| SLC16A1 | MOT1\_HUMAN | Monocarboxylate transporter 1 | DTDGGPKEEES[79.9663]PV | S | 154.0 | 113785.12 | 0.0 | 0.0 | 113785.12 |
| SPTB | SPTB1\_HUMAN | Spectrin beta chain, erythrocytic | LGHLQSS[79.9663]WDR | S | 1565.0 | 113514.81 | 0.0 | 113514.81 | 0.0 |
| ANK1 | ANK1\_HUMAN | Isoform Br21 of Ankyrin-1 | YSILS[79.9663]ESTPGSLSGTEQAEM[15.9949]K | S |  | 112992.998 | 0.0 | 112992.998 | 0.0 |
| ATP7A | ATP7A\_HUMAN | Isoform 1 of Copper-transporting ATPase 1 | YNASSVT[79.9663]PESLR | T | 327.0 | 112308.883 | 0.0 | 36514.508 | 75794.375 |
| HSP90AB1 | HS90B\_HUMAN | Heat shock protein HSP 90-beta | IEDVGS[79.9663]DEEDDSGKDK | S | 255.0 | 111904.9385 | 0.0 | 82795.6093 | 29109.3292 |
| SLC2A1 | GTR1\_HUMAN | Solute carrier family 2, facilitated glucose transporter member 1 | T[79.9663]FDEIASGFR | T | 459.0 | 111897.44 | 0.0 | 111897.44 | 0.0 |
| EPB41 | EPB41\_HUMAN | Isoform 5 of Protein 4.1 | LSMY[79.9663]GVDLHK | Y | 396.0 | 111096.734 | 0.0 | 111096.734 | 0.0 |
| ADD1 | ADDA\_HUMAN | Isoform 3 of Alpha-adducin | AKSRSPGS[79.9663]PVGEGTGSPPK | S | 358.0 | 110272.94 | 110272.94 | 0.0 | 0.0 |
| PSMA3 | PSA3\_HUMAN | Isoform 2 of Proteasome subunit alpha type-3 | ESLKEEDES[79.9663]DDDNM[15.9949] | S |  | 109752.4995 | 0.0 | 91241.9795 | 18510.52 |
| TRAPPC12 | TPC12\_HUMAN | Trafficking protein particle complex subunit 12 | VM[15.9949]Y[79.9663]SMANC[57.0215]LLLM[15.9949]K | Y |  | 108027.58 | 0.0 | 108027.58 | 0.0 |
| SERINC1 | SERC1\_HUMAN | Serine incorporator 1 | S[79.9663]DGSLEDGDDVHR | S | 361.0 | 107579.392 | 0.0 | 21428.344 | 86151.048 |
| EPB41 | EPB41\_HUMAN | Isoform 5 of Protein 4.1 | RAS[79.9663]RSLDGAAAVDSADR | S | 540.0 | 107293.01 | 0.0 | 107293.01 | 0.0 |
| EPB41 | EPB41\_HUMAN | Isoform 5 of Protein 4.1 | IRPGEQEQY[79.9663]ESTIGFK | Y | 459.0 | 106993.941 | 0.0 | 106993.941 | 0.0 |
| EPB41 | EPB41\_HUMAN | Isoform 5 of Protein 4.1 | DLDKS[79.9663]QEEIKK | S | 674.0 | 106551.3 | 0.0 | 106551.3 | 0.0 |
| LPIN2 | LPIN2\_HUMAN | Phosphatidate phosphatase LPIN2 | TATIT[79.9663]PSENTHFR | T | 291.0 | 106213.835 | 0.0 | 68035.94 | 38177.895 |
| SPTA1 | SPTA1\_HUMAN | Spectrin alpha chain, erythrocytic 1 | VM[15.9949]ALY[79.9663]DFQAR | Y | 986.0 | 105387.613 | 0.0 | 95003.666 | 10383.947 |
| CYBRD1 | CYBR1\_HUMAN | Plasma membrane ascorbate-dependent reductase CYBRD1 | S[79.9663]DSELNSEVAAR | S | 260.0 | 104145.173 | 0.0 | 27434.969 | 76710.204 |
| ANK1 | ANK1\_HUMAN | Isoform Br21 of Ankyrin-1 | YSILSES[79.9663]TPGSLSGTEQAEM[15.9949]K | S |  | 104036.748 | 0.0 | 104036.748 | 0.0 |
| DENND1A | DEN1A\_HUMAN | DENN domain-containing protein 1A | TAPS[79.9663]PLVEAK | S | 473.0 | 103934.141 | 0.0 | 55147.227 | 48786.914 |
| HBA1 | HBA\_HUMAN | Hemoglobin subunit alpha | VLSPADKT[79.9663]NVK | T | 9.0 | 102285.47 | 0.0 | 102285.47 | 0.0 |
| GYPA | GLPA\_HUMAN | Isoform 2 of Glycophorin-A | KSPSDVKPLPS[79.9663]PDTDVPLSSVEIENPETSDQ | S | 130.0 | 102167.046 | 0.0 | 102167.046 | 0.0 |
| SPTAN1 | SPTN1\_HUMAN | Isoform 3 of Spectrin alpha chain, non-erythrocytic 1 | VNS[79.9663]LGETAER | S | 1291.0 | 101166.2 | 0.0 | 101166.2 | 0.0 |
| HUWE1 | HUWE1\_HUMAN | Isoform 2 of E3 ubiquitin-protein ligase HUWE1 | AGS[79.9663]STPGDAPPAVAEVQGR | S | 2887.0 | 100309.01 | 100309.01 | 0.0 | 0.0 |
| SPTA1 | SPTA1\_HUMAN | Spectrin alpha chain, erythrocytic 1 | SLNQNM[15.9949]ES[79.9663]LR | S | 883.0 | 99947.317 | 0.0 | 99947.317 | 0.0 |
| GYPA | GLPA\_HUMAN | Isoform 2 of Glycophorin-A | KSPSDVKPLPSPDT[79.9663]DVPLSSVEIENPETSDQ | T | 133.0 | 99759.12 | 0.0 | 99759.12 | 0.0 |
| PGM2 | PGM2\_HUMAN | Phosphopentomutase | AAPEGS[79.9663]GLGEDAR | S | 7.0 | 99717.484 | 99717.484 | 0.0 | 0.0 |
| HBA1 | HBA\_HUMAN | Hemoglobin subunit alpha | FLASVS[79.9663]TVLTSK | S | 134.0 | 99618.21 | 0.0 | 99618.21 | 0.0 |
| KRT26 | K1C26\_HUMAN | Keratin, type I cytoskeletal 26 | AEY[79.9663]EDLAEQNR | Y | 263.0 | 99263.47 | 0.0 | 99263.47 | 0.0 |
| KRT1 | K2C1\_HUMAN | Keratin, type II cytoskeletal 1 | SGGGFSSGS[79.9663]AGIINYQR | S | 21.0 | 98283.953 | 0.0 | 88572.677 | 9711.276 |
| SMIM1 | SMIM1\_HUMAN | Small integral membrane protein 1 | DGVSLGAVSS[79.9663]TEEASR | S | 28.0 | 97744.0 | 0.0 | 0.0 | 97744.0 |
| HUWE1 | HUWE1\_HUMAN | Isoform 2 of E3 ubiquitin-protein ligase HUWE1 | AGSST[79.9663]PGDAPPAVAEVQGR | T | 2889.0 | 97534.88 | 97534.88 | 0.0 | 0.0 |
| EPB41 | EPB41\_HUMAN | Isoform 5 of Protein 4.1 | IRPGEQEQYES[79.9663]TIGFK | S | 461.0 | 97005.58 | 0.0 | 97005.58 | 0.0 |
| EPB41 | EPB41\_HUMAN | Isoform 5 of Protein 4.1 | S[79.9663]LDGAAAVDSADRS[79.9663]PR | S |  | 96845.16 | 0.0 | 96845.16 | 0.0 |
| EPB41 | EPB41\_HUMAN | Isoform 5 of Protein 4.1 | RLSTHS[79.9663]PFR | S | 712.0 | 96724.9 | 0.0 | 96724.9 | 0.0 |
| TNS1 | TENS1\_HUMAN | Tensin-1 | QGS[79.9663]PTPALPEK | S | 1485.0 | 96595.775 | 0.0 | 74848.4 | 21747.375 |
| SPTB | SPTB1\_HUMAN | Spectrin beta chain, erythrocytic | T[79.9663]SPVS[79.9663]LWSR | T |  | 96559.87 | 0.0 | 96559.87 | 0.0 |
| STX7 | STX7\_HUMAN | Isoform 2 of Syntaxin-7 | VS[79.9663]GSFPEDSSK | S | 129.0 | 95773.52 | 0.0 | 0.0 | 95773.52 |
| ADD2 | ADDB\_HUMAN | Beta-adducin | S[79.9663]PSTESQLM[15.9949]SK | S |  | 95515.545 | 0.0 | 60606.33 | 34909.215 |
| SPTA1 | SPTA1\_HUMAN | Spectrin alpha chain, erythrocytic 1 | SDDKSS[79.9663]LDSLEALM[15.9949]K | S |  | 95157.46 | 0.0 | 95157.46 | 0.0 |
| ADD2 | ADDB\_HUMAN | Beta-adducin | SPS[79.9663]TESQLM[15.9949]SK | S |  | 94951.95 | 0.0 | 94951.95 | 0.0 |
| ADD2 | ADDB\_HUMAN | Beta-adducin | ADEVEKS[79.9663]SSGMPIR | S | 471.0 | 94635.484 | 0.0 | 94635.484 | 0.0 |
| TSC22D4 | T22D4\_HUMAN | TSC22 domain family protein 4 | AAT[79.9663]PLPSLR | T | 211.0 | 94420.887 | 0.0 | 94420.887 | 0.0 |
| RILP | RILP\_HUMAN | Rab-interacting lysosomal protein | AES[79.9663]SEDETSSPAPSK | S | 354.0 | 94355.1592 | 70383.9062 | 18939.766 | 5031.487 |
| EPB41 | EPB41\_HUMAN | Isoform 5 of Protein 4.1 | EES[79.9663]PQS[79.9663]KAETELK | S |  | 92554.55 | 0.0 | 92554.55 | 0.0 |
| ACTB | ACTB\_HUMAN | Actin, cytoplasmic 1 | QEYDES[79.9663]GPSIVHR | S | 365.0 | 91710.478 | 0.0 | 91710.478 | 0.0 |
| SLC66A2 | S66A2\_HUMAN | Solute carrier family 66 member 2 | S[79.9663]FTAADSKDEEVK | S | 110.0 | 91228.0 | 0.0 | 32707.594 | 58520.406 |
| GYPA | GLPA\_HUMAN | Isoform 2 of Glycophorin-A | S[79.9663]PSDVKPLPSPDTDVPLSSVEIENPETSDQ | S | 121.0 | 91102.469 | 0.0 | 91102.469 | 0.0 |
| OR51D1 | O51D1\_HUMAN | Olfactory receptor 51D1 | VNVVY[79.9663]GLFIILSVMGVDSLFIGFSYILILWAVLELSSR | Y | 211.0 | 90738.63 | 0.0 | 0.0 | 90738.63 |
| SLC2A1 | GTR1\_HUMAN | Solute carrier family 2, facilitated glucose transporter member 1 | TPEELFHPLGADS[79.9663]QV | S | 490.0 | 89626.681 | 0.0 | 57675.023 | 31951.658 |
| EPB41 | EPB41\_HUMAN | Isoform 5 of Protein 4.1 | S[79.9663]QVS[79.9663]EEEGKEVESDKEK | S |  | 88872.945 | 0.0 | 88872.945 | 0.0 |
| WNK1 | WNK1\_HUMAN | Isoform 2 of Serine/threonine-protein kinase WNK1 | FIVS[79.9663]PVPESR | S | 1261.0 | 88844.82 | 0.0 | 57131.863 | 31712.957 |
| CALR | CALR\_HUMAN | Calreticulin | IKDPDAS[79.9663]KPEDWDER | S | 214.0 | 88717.71 | 0.0 | 88717.71 | 0.0 |
| SLC2A1 | GTR1\_HUMAN | Solute carrier family 2, facilitated glucose transporter member 1 | GT[79.9663]ADVTHDLQEM[15.9949]K | T |  | 87688.854 | 0.0 | 67006.36 | 20682.494 |
| ANK1 | ANK1\_HUMAN | Isoform Br21 of Ankyrin-1 | Y[79.9663]SILSESTPGSLSGTEQAEM[15.9949]K | Y |  | 84955.61 | 0.0 | 84955.61 | 0.0 |
| HBA1 | HBA\_HUMAN | Hemoglobin subunit alpha | T[79.9663]YFPHFDLSHGSAQVK | T | 42.0 | 84770.555 | 0.0 | 84770.555 | 0.0 |
| DMTN | DEMA\_HUMAN | Isoform 4 of Dematin | DSS[79.9663]VPGSPSSIVAK | S | 22.0 | 83855.3 | 0.0 | 0.0 | 83855.3 |
| FLOT1 | FLOT1\_HUMAN | Flotillin-1 | ITLVSSGSGT[79.9663]M[15.9949]GAAK | T |  | 83758.08 | 0.0 | 0.0 | 83758.08 |
| SPTB | SPTB1\_HUMAN | Spectrin beta chain, erythrocytic | VYT[79.9663]PHDGK | T | 382.0 | 82564.625 | 0.0 | 82564.625 | 0.0 |
| TMC8 | TMC8\_HUMAN | Transmembrane channel-like protein 8 | LLPEPGPSDS[79.9663]PGPK | S | 658.0 | 82554.78 | 0.0 | 0.0 | 82554.78 |
| SYNJ1 | SYNJ1\_HUMAN | Synaptojanin-1 | AQLSVQTS[79.9663]PVPTPDPK | S | 1345.0 | 81963.5 | 81963.5 | 0.0 | 0.0 |
| ARHGAP23 | RHG23\_HUMAN | Rho GTPase-activating protein 23 | S[79.9663]AEALGPGALVSPR | S | 361.0 | 81455.446 | 0.0 | 71628.164 | 9827.282 |
| SPTA1 | SPTA1\_HUMAN | Spectrin alpha chain, erythrocytic 1 | TGQEMIEGGHY[79.9663]ASDNVTTR | Y | 653.0 | 81390.09 | 0.0 | 81390.09 | 0.0 |
| ACTB | ACTB\_HUMAN | Actin, cytoplasmic 1 | GY[79.9663]SFTTTAER | Y | 198.0 | 81341.353 | 0.0 | 54112.002 | 27229.351 |
| GNG5 | GBG5\_HUMAN | Guanine nucleotide-binding protein G(I)/G(S)/G(O) subunit gamma-5 | n[42.0106]SGS[79.9663]SSVAAM[15.9949]K | S |  | 81308.78 | 0.0 | 81308.78 | 0.0 |
| KRT13 | K1C13\_HUMAN | Keratin, type I cytoskeletal 13 | EVS[79.9663]TNTAM[15.9949]IQTSK | S |  | 81280.406 | 0.0 | 64309.64 | 16970.766 |
| TNS1 | TENS1\_HUMAN | Tensin-1 | AQFSVAGVHTVPGS[79.9663]PQAR | S | 1281.0 | 81254.53 | 81254.53 | 0.0 | 0.0 |
| CSNK1A1 | KC1A\_HUMAN | Isoform 3 of Casein kinase I isoform alpha | AAQQAAS[79.9663]SSGQGQQAQTPTGK | S | 311.0 | 80948.338 | 80948.338 | 0.0 | 0.0 |
| BCAM | BCAM\_HUMAN | Basal cell adhesion molecule | VEDY[79.9663]DAADDVQLSK | Y | 342.0 | 80772.06 | 0.0 | 0.0 | 80772.06 |
| EPB41 | EPB41\_HUMAN | Isoform 5 of Protein 4.1 | VVVHQET[79.9663]EIADE | T | 859.0 | 80768.463 | 0.0 | 80768.463 | 0.0 |
| SLC2A1 | GTR1\_HUMAN | Solute carrier family 2, facilitated glucose transporter member 1 | Q[-17.0265]GGASQSDKT[79.9663]PEELFHPLGADSQV | T | 478.0 | 80725.13 | 0.0 | 0.0 | 80725.13 |
| ADD2 | ADDB\_HUMAN | Beta-adducin | SPLVSPS[79.9663]KSLEEGTKK | S | 619.0 | 80174.32 | 0.0 | 80174.32 | 0.0 |
| SPTB | SPTB1\_HUMAN | Spectrin beta chain, erythrocytic | LLTSQDVSY[79.9663]DEAR | Y | 1302.0 | 80115.91 | 0.0 | 43296.176 | 36819.734 |
| KRT3 | K2C3\_HUMAN | Keratin, type II cytoskeletal 3 | S[79.9663]LYNLGGNK | S | 56.0 | 79569.41 | 0.0 | 30989.44 | 48579.97 |
| MPP1 | EM55\_HUMAN | Isoform 2 of 55 kDa erythrocyte membrane protein | ASEGESGGSM[15.9949]HTALS[79.9663]DLYLEHLLQK | S | 19.0 | 79116.586 | 79116.586 | 0.0 | 0.0 |
| EHBP1L1 | EH1L1\_HUMAN | EH domain-binding protein 1-like protein 1 | ANEAGGQVGPEAPRPPET[79.9663]SPEMR | T | 284.0 | 77802.98 | 77802.98 | 0.0 | 0.0 |
| CA2 | CAH2\_HUMAN | Carbonic anhydrase 2 | QSPVDIDTHT[79.9663]AK | T | 37.0 | 77751.72 | 0.0 | 77751.72 | 0.0 |
| RAD23A | RD23A\_HUMAN | Isoform 2 of UV excision repair protein RAD23 homolog A | EDKS[79.9663]PSEESAPTTSPESVSGSVPSSGSSGR | S | 123.0 | 77567.33 | 0.0 | 77567.33 | 0.0 |
| KRT5 | K2C5\_HUMAN | Keratin, type II cytoskeletal 5 | ISIS[79.9663]TSGGSFR | S | 77.0 | 77513.641 | 0.0 | 40930.016 | 36583.625 |
| CYBRD1 | CYBR1\_HUMAN | Plasma membrane ascorbate-dependent reductase CYBRD1 | GSM[15.9949]PAYS[79.9663]GNNM[15.9949]DK | S |  | 77398.418 | 0.0 | 18164.828 | 59233.59 |
| RAD23A | RD23A\_HUMAN | Isoform 2 of UV excision repair protein RAD23 homolog A | EDKSPSEES[79.9663]APTTSPESVSGSVPSSGSSGR | S | 128.0 | 77244.48 | 0.0 | 77244.48 | 0.0 |
| SCAMP2 | SCAM2\_HUMAN | Secretory carrier-associated membrane protein 2 | AAS[79.9663]SAAQGAFQGN | S | 319.0 | 76946.57 | 0.0 | 0.0 | 76946.57 |
| SPTB | SPTB1\_HUMAN | Spectrin beta chain, erythrocytic | LLTSQDVS[79.9663]YDEAR | S | 1301.0 | 76914.2685 | 0.0 | 76914.2685 | 0.0 |
| SPTB | SPTB1\_HUMAN | Spectrin beta chain, erythrocytic | WLAEMEMPDT[79.9663]LEDLEVVQHR | T | 879.0 | 76847.5 | 0.0 | 76847.5 | 0.0 |
| ASPSCR1 | ASPC1\_HUMAN | Isoform 2 of Tether containing UBX domain for GLUT4 | AAGS[79.9663]PSPLPAPDPAPK | S | 500.0 | 75768.016 | 75768.016 | 0.0 | 0.0 |
| HUWE1 | HUWE1\_HUMAN | Isoform 2 of E3 ubiquitin-protein ligase HUWE1 | GSGTAS[79.9663]DDEFENLR | S | 1907.0 | 75324.265 | 0.0 | 11943.129 | 63381.136 |
| ITSN1 | ITSN1\_HUMAN | Isoform 10 of Intersectin-1 | SAFTPATATGSSPS[79.9663]PVLGQGEK | S | 904.0 | 75225.414 | 0.0 | 75225.414 | 0.0 |
| SPTA1 | SPTA1\_HUMAN | Spectrin alpha chain, erythrocytic 1 | ETGTLES[79.9663]QLEANK | S | 2194.0 | 74969.336 | 0.0 | 74969.336 | 0.0 |
| DMTN | DEMA\_HUMAN | Isoform 4 of Dematin | HLSAEDFS[79.9663]R | S | 18.0 | 74690.8957 | 0.0 | 74690.8957 | 0.0 |
| AGFG1 | AGFG1\_HUMAN | Isoform 2 of Arf-GAP domain and FG repeat-containing protein 1 | GTPSQS[79.9663]PVVGR | S | 181.0 | 74281.34 | 0.0 | 74281.34 | 0.0 |
| KRT1 | K2C1\_HUMAN | Keratin, type II cytoskeletal 1 | S[79.9663]KAEAESLYQSK | S | 365.0 | 73706.971 | 0.0 | 0.0 | 73706.971 |
| CYBRD1 | CYBR1\_HUMAN | Plasma membrane ascorbate-dependent reductase CYBRD1 | NLALDEAGQRST[79.9663]M[15.9949] | T |  | 73541.055 | 0.0 | 0.0 | 73541.055 |
| SPTB | SPTB1\_HUMAN | Spectrin beta chain, erythrocytic | n[42.0106]TSATEFENVGNQPPY[79.9663]SR | Y | 16.0 | 73498.667 | 0.0 | 65105.078 | 8393.589 |
| ADD2 | ADDB\_HUMAN | Beta-adducin | EAET[79.9663]KSPLVSPSK | T | 611.0 | 73180.81 | 0.0 | 73180.81 | 0.0 |
| C2CD2L | C2C2L\_HUMAN | Phospholipid transfer protein C2CD2L | NLGTPTSSTPRPS[79.9663]ITPTK | S | 426.0 | 72262.16 | 0.0 | 72262.16 | 0.0 |
| ADD3 | ADDG\_HUMAN | Isoform 1 of Gamma-adducin | IEEVLSPEGSPSKS[79.9663]PSKK | S | 681.0 | 71648.73 | 0.0 | 71648.73 | 0.0 |
| ARHGAP23 | RHG23\_HUMAN | Rho GTPase-activating protein 23 | ARS[79.9663]DDYLSR | S | 351.0 | 71254.35 | 71254.35 | 0.0 | 0.0 |
| C2CD2L | C2C2L\_HUMAN | Phospholipid transfer protein C2CD2L | EAGLS[79.9663]QSHDDLSNATATPSVR | S | 660.0 | 70971.625 | 0.0 | 70971.625 | 0.0 |
| DMTN | DEMA\_HUMAN | Isoform 4 of Dematin | SPGIIS[79.9663]QAS[79.9663]APR | S |  | 70760.31 | 0.0 | 70760.31 | 0.0 |
| AQP1 | AQP1\_HUMAN | Aquaporin-1 | Y[79.9663]PVGNNQTAVQDNVK | Y | 37.0 | 70710.146 | 0.0 | 26482.436 | 44227.71 |
| SLC43A1 | LAT3\_HUMAN | Large neutral amino acids transporter small subunit 3 | APS[79.9663]LEDGSDAFM[15.9949]SPQDVR | S |  | 70455.1276 | 0.0 | 27985.844 | 42469.2836 |
| KRT6A | K2C6A\_HUMAN | Keratin, type II cytoskeletal 6A | AIGGGLS[79.9663]SVGGGSSTIK | S | 540.0 | 70217.015 | 0.0 | 70217.015 | 0.0 |
| ABCC4 | MRP4\_HUMAN | ATP-binding cassette sub-family C member 4 | DNEESEQPPVPGT[79.9663]PTLR | T | 646.0 | 70200.246 | 0.0 | 26648.26 | 43551.986 |
| VAMP3 | VAMP3\_HUMAN | Vesicle-associated membrane protein 3 | ADALQAGAS[79.9663]QFETSAAK | S | 58.0 | 69953.97 | 0.0 | 0.0 | 69953.97 |
| PSMA5 | PSA5\_HUMAN | Proteasome subunit alpha type-5 | ITS[79.9663]PLM[15.9949]EPSSIEK | S |  | 69879.01 | 0.0 | 69879.01 | 0.0 |
| MARVELD2 | MALD2\_HUMAN | MARVEL domain-containing protein 2 | EYM[15.9949]EQQEINEPS[79.9663]LSSK | S | 387.0 | 68984.3747 | 0.0 | 6747.0117 | 62237.363 |
| SPTA1 | SPTA1\_HUMAN | Spectrin alpha chain, erythrocytic 1 | S[79.9663]YEDPTNIQGK | S | 80.0 | 68860.998 | 0.0 | 53121.802 | 15739.196 |
| ACTA1 | ACTS\_HUMAN | Actin, alpha skeletal muscle | Q[-17.0265]EY[79.9663]DEAGPSIVHR | Y | 364.0 | 68736.228 | 0.0 | 68736.228 | 0.0 |
| ABCC1 | MRP1\_HUMAN | Isoform 2 of Multidrug resistance-associated protein 1 | QLSSSS[79.9663]SYSGDISR | S | 918.0 | 68657.848 | 0.0 | 0.0 | 68657.848 |
| ITGB1 | ITB1\_HUMAN | Integrin beta-1 | SAVTT[79.9663]VVNPK | T | 789.0 | 67735.766 | 0.0 | 67735.766 | 0.0 |
| MARK3 | MARK3\_HUMAN | Isoform 4 of MAP/microtubule affinity-regulating kinase 3 | SRGS[79.9663]TNLFSK | S | 601.0 | 67369.664 | 0.0 | 67369.664 | 0.0 |
| SLC43A2 | LAT4\_HUMAN | Isoform 3 of Large neutral amino acids transporter small subunit 4 | LSVGS[79.9663]SM[15.9949]R | S |  | 67166.36 | 0.0 | 0.0 | 67166.36 |
| SPTB | SPTB1\_HUMAN | Spectrin beta chain, erythrocytic | DVSS[79.9663]VELLM[15.9949]K | S |  | 66794.16 | 0.0 | 66794.16 | 0.0 |
| GYPA | GLPA\_HUMAN | Isoform 2 of Glycophorin-A | PLPSPDTDVPLS[79.9663]SVEIENPETSDQ | S | 138.0 | 66185.807 | 0.0 | 66185.807 | 0.0 |
| KRT6B | K2C6B\_HUMAN | Keratin, type II cytoskeletal 6B | S[79.9663]LYGLGGSK | S | 60.0 | 65784.21 | 0.0 | 0.0 | 65784.21 |
| SPTA1 | SPTA1\_HUMAN | Spectrin alpha chain, erythrocytic 1 | SSLDS[79.9663]LEALM[15.9949]K | S |  | 65516.043 | 0.0 | 65516.043 | 0.0 |
| KRT1 | K2C1\_HUMAN | Keratin, type II cytoskeletal 1 | SGGGFSS[79.9663]GSAGIINYQR | S | 19.0 | 65477.852 | 0.0 | 57071.773 | 8406.079 |
| SPTA1 | SPTA1\_HUMAN | Spectrin alpha chain, erythrocytic 1 | ETVVESS[79.9663]GPK | S | 13.0 | 65239.332 | 0.0 | 65239.332 | 0.0 |
| ADD2 | ADDB\_HUMAN | Beta-adducin | DKTESVTSGPM[15.9949]SPEGS[79.9663]PSKS[79.9663]PSK | S |  | 64980.656 | 0.0 | 64980.656 | 0.0 |
| ARHGDIA | GDIR1\_HUMAN | Rho GDP-dissociation inhibitor 1 | AEQEPT[79.9663]AEQLAQIAAENEEDEHSVNYKPPAQK | T | 7.0 | 64405.797 | 64405.797 | 0.0 | 0.0 |
| RILP | RILP\_HUMAN | Rab-interacting lysosomal protein | AESS[79.9663]EDETSSPAPSK | S | 355.0 | 64323.55 | 64323.55 | 0.0 | 0.0 |
| ACTB | ACTB\_HUMAN | Actin, cytoplasmic 1 | EITALAPS[79.9663]TM[15.9949]K | S |  | 64212.047 | 0.0 | 64212.047 | 0.0 |
| NECTIN1 | NECT1\_HUMAN | Nectin-1 | AGPLGGS[79.9663]SYEEEEEEEEGGGGGER | S | 434.0 | 63967.58 | 0.0 | 63967.58 | 0.0 |
| DMTN | DEMA\_HUMAN | Isoform 4 of Dematin | S[79.9663]SSLPAYGRTTLSR | S | 287.0 | 63135.523 | 0.0 | 63135.523 | 0.0 |
| GYPA | GLPA\_HUMAN | Isoform 2 of Glycophorin-A | PLPSPDTDVPLSS[79.9663]VEIENPETSDQ | S | 139.0 | 63022.979 | 0.0 | 63022.979 | 0.0 |
| BAG6 | BAG6\_HUMAN | Isoform 2 of Large proline-rich protein BAG6 | PLTS[79.9663]PESLSR | S | 1081.0 | 62913.9 | 0.0 | 62913.9 | 0.0 |
| SPTB | SPTB1\_HUMAN | Spectrin beta chain, erythrocytic | n[42.0106]TSATEFENVGNQPPYS[79.9663]R | S | 17.0 | 62308.729 | 0.0 | 54208.93 | 8099.799 |
| SPTB | SPTB1\_HUMAN | Spectrin beta chain, erythrocytic | LLS[79.9663]GEDVGQDEGATR | S | 767.0 | 62007.983 | 0.0 | 41085.508 | 20922.475 |
| EPB41 | EPB41\_HUMAN | Isoform 5 of Protein 4.1 | S[79.9663]QVSEEEGKEVESDK | S | 92.0 | 61995.8 | 0.0 | 61995.8 | 0.0 |
| ADD2 | ADDB\_HUMAN | Beta-adducin | GLSQMTTSADT[79.9663]DVDTSKDK | T | 675.0 | 61491.723 | 0.0 | 61491.723 | 0.0 |
| HSP90AA1 | HS90A\_HUMAN | Isoform 2 of Heat shock protein HSP 90-alpha | ESEDKPEIEDVGS[79.9663]DEEEEKK | S | 263.0 | 61302.187 | 0.0 | 61302.187 | 0.0 |
| KRT6A | K2C6A\_HUMAN | Keratin, type II cytoskeletal 6A | AIGGGLSSVGGGSS[79.9663]TIK | S | 547.0 | 60901.58 | 0.0 | 60901.58 | 0.0 |
| SLC4A1 | B3AT\_HUMAN | Band 3 anion transport protein | ATFDEEEGRDEY[79.9663]DEVAM[15.9949]PV | Y |  | 60623.127 | 0.0 | 0.0 | 60623.127 |
| ADD2 | ADDB\_HUMAN | Beta-adducin | SRSPST[79.9663]ES[79.9663]QLM[15.9949]SK | T |  | 60364.824 | 0.0 | 60364.824 | 0.0 |
| FLOT1 | FLOT1\_HUMAN | Flotillin-1 | ITLVSS[79.9663]GSGTM[15.9949]GAAK | S |  | 60330.195 | 0.0 | 0.0 | 60330.195 |
| SPTA1 | SPTA1\_HUMAN | Spectrin alpha chain, erythrocytic 1 | LTLS[79.9663]HPSDAPQIQEM[15.9949]K | S |  | 60136.78 | 0.0 | 60136.78 | 0.0 |
| KRT13 | K1C13\_HUMAN | Keratin, type I cytoskeletal 13 | S[79.9663]EMEC[57.0215]QNQEYK | S |  | 60108.676 | 0.0 | 0.0 | 60108.676 |
| ADD2 | ADDB\_HUMAN | Beta-adducin | GLSQM[15.9949]TT[79.9663]SADTDVDTSKDK | T | 671.0 | 60030.883 | 0.0 | 60030.883 | 0.0 |
| PI4K2A | P4K2A\_HUMAN | Phosphatidylinositol 4-kinase type 2-alpha | VAAAAGS[79.9663]GPSPPGSPGHDR | S | 44.0 | 59356.883 | 0.0 | 17647.227 | 41709.656 |
| PRKAR1A | KAP0\_HUMAN | cAMP-dependent protein kinase type I-alpha regulatory subunit | EDEIS[79.9663]PPPPNPVVK | S | 83.0 | 59073.09 | 0.0 | 59073.09 | 0.0 |
| UBAC1 | UBAC1\_HUMAN | Ubiquitin-associated domain-containing protein 1 | KPS[79.9663]PEELDK | S | 332.0 | 59002.613 | 0.0 | 59002.613 | 0.0 |
| SVIP | SVIP\_HUMAN | Small VCP/p97-interacting protein | GILDVQS[79.9663]VQEK | S | 46.0 | 58862.957 | 0.0 | 58862.957 | 0.0 |
| SPTB | SPTB1\_HUMAN | Spectrin beta chain, erythrocytic | QY[79.9663]QDHLNTR | Y | 928.0 | 58209.6 | 0.0 | 46448.99 | 11760.61 |
| SPTA1 | SPTA1\_HUMAN | Spectrin alpha chain, erythrocytic 1 | LQAT[79.9663]YWYHR | T | 369.0 | 57456.4 | 0.0 | 57456.4 | 0.0 |
| ADD2 | ADDB\_HUMAN | Beta-adducin | TESVTS[79.9663]GPM[15.9949]SPEGSPSK | S |  | 56290.94 | 0.0 | 39757.46 | 16533.48 |
| CYBRD1 | CYBR1\_HUMAN | Plasma membrane ascorbate-dependent reductase CYBRD1 | GSM[15.9949]PAY[79.9663]SGNNM[15.9949]DK | Y |  | 56224.836 | 0.0 | 0.0 | 56224.836 |
| CYBRD1 | CYBR1\_HUMAN | Plasma membrane ascorbate-dependent reductase CYBRD1 | SDSELNS[79.9663]EVAAR | S | 266.0 | 56028.545 | 0.0 | 22422.51 | 33606.035 |
| EIF4B | IF4B\_HUMAN | Eukaryotic translation initiation factor 4B | S[79.9663]QSSDTEQQSPTSGGGK | S | 495.0 | 55506.667 | 0.0 | 39457.562 | 16049.105 |
| DMTN | DEMA\_HUMAN | Isoform 4 of Dematin | Q[-17.0265]PLTSPGSVS[79.9663]PSR | S | 16.0 | 55383.95 | 0.0 | 0.0 | 55383.95 |
| SLC12A7 | S12A7\_HUMAN | Solute carrier family 12 member 7 | T[79.9663]LM[15.9949]M[15.9949]EQR | T |  | 55303.638 | 0.0 | 14037.828 | 41265.81 |
| KRT6A | K2C6A\_HUMAN | Keratin, type II cytoskeletal 6A | AIGGGLSSVGGGSST[79.9663]IK | T | 548.0 | 55206.59 | 0.0 | 55206.59 | 0.0 |
| ACTB | ACTB\_HUMAN | Actin, cytoplasmic 1 | n[42.0106]DDDIAALVVDNGS[79.9663]GMC[57.0215]K | S |  | 54852.6876 | 0.0 | 54852.6876 | 0.0 |
| PGM1 | PGM1\_HUMAN | Phosphoglucomutase-1 | AIGGIILTAS[79.9663]HNPGGPNGDFGIK | S | 117.0 | 54511.841 | 54511.841 | 0.0 | 0.0 |
| SPTA1 | SPTA1\_HUMAN | Spectrin alpha chain, erythrocytic 1 | FTM[15.9949]GHS[79.9663]AHEETK | S | 124.0 | 54441.938 | 0.0 | 54441.938 | 0.0 |
| RAB7A | RAB7A\_HUMAN | Ras-related protein Rab-7a | ATIGADFLT[79.9663]K | T | 47.0 | 54400.69 | 0.0 | 54400.69 | 0.0 |
| ADD2 | ADDB\_HUMAN | Beta-adducin | TESVTSGPM[15.9949]S[79.9663]PEGSPS[79.9663]KSPSK | S |  | 54366.0 | 0.0 | 54366.0 | 0.0 |
| ADD2 | ADDB\_HUMAN | Beta-adducin | TESVTSGPM[15.9949]SPEGS[79.9663]PSKS[79.9663]PSK | S |  | 54330.77 | 0.0 | 54330.77 | 0.0 |
| HRNR | HORN\_HUMAN | Hornerin | Q[-17.0265]SLGHGQHGSGSGQSPS[79.9663]PSR | S | 542.0 | 54268.348 | 0.0 | 32996.491 | 21271.857 |
| SPTA1 | SPTA1\_HUMAN | Spectrin alpha chain, erythrocytic 1 | ETVVES[79.9663]SGPK | S | 12.0 | 53982.188 | 0.0 | 53982.188 | 0.0 |
| SLC4A1 | B3AT\_HUMAN | Band 3 anion transport protein | RYQSSPAKPDSSFY[79.9663]K | Y | 359.0 | 53893.43 | 0.0 | 53893.43 | 0.0 |
| EPB41 | EPB41\_HUMAN | Isoform 5 of Protein 4.1 | AS[79.9663]RSLDGAAAVDSADR | S | 540.0 | 53778.582 | 0.0 | 53778.582 | 0.0 |
| STX7 | STX7\_HUMAN | Isoform 2 of Syntaxin-7 | n[42.0106]SY[79.9663]TPGVGGDPAQLAQR | Y | 3.0 | 53731.492 | 0.0 | 0.0 | 53731.492 |
| DMTN | DEMA\_HUMAN | Isoform 4 of Dematin | GAEEEEEEEDDDS[79.9663]GEEM[15.9949]KALR | S |  | 52995.406 | 0.0 | 52995.406 | 0.0 |
| NACA | NACAM\_HUMAN | Nascent polypeptide-associated complex subunit alpha, muscle-specific form | GAPTTPAATPPS[79.9663]PK | S | 766.0 | 52909.432 | 0.0 | 25917.547 | 26991.885 |
| ACTB | ACTB\_HUMAN | Actin, cytoplasmic 1 | Q[-17.0265]EY[79.9663]DESGPSIVHR | Y | 362.0 | 52861.8094 | 0.0 | 52861.8094 | 0.0 |
| ANK1 | ANK1\_HUMAN | Isoform Br21 of Ankyrin-1 | LVPLVQATFPENAVT[79.9663]KR | T | 1101.0 | 52764.56 | 0.0 | 52764.56 | 0.0 |
| KRT13 | K1C13\_HUMAN | Keratin, type I cytoskeletal 13 | MIGFPSSAGSVS[79.9663]PR | S | 427.0 | 52563.085 | 0.0 | 52563.085 | 0.0 |
| PGM1 | PGM1\_HUMAN | Phosphoglucomutase-1 | AIGGIILT[79.9663]ASHNPGGPNGDFGIK | T | 115.0 | 52204.71 | 52204.71 | 0.0 | 0.0 |
| SPTA1 | SPTA1\_HUMAN | Spectrin alpha chain, erythrocytic 1 | LTLSHPS[79.9663]DAPQIQEM[15.9949]K | S |  | 52150.484 | 0.0 | 52150.484 | 0.0 |
| CA2 | CAH2\_HUMAN | Carbonic anhydrase 2 | GERQS[79.9663]PVDIDTHTAK | S | 29.0 | 51811.75 | 0.0 | 51811.75 | 0.0 |
| IGHA1 | IGHA1\_HUMAN | Isoform 1 of Immunoglobulin heavy constant alpha 1 | TPLT[79.9663]ATLSK | T | 216.0 | 51393.277 | 0.0 | 51393.277 | 0.0 |
| SPTB | SPTB1\_HUMAN | Spectrin beta chain, erythrocytic | FAALEKPT[79.9663]TLELK | T | 2072.0 | 51160.27 | 0.0 | 51160.27 | 0.0 |
| SPTA1 | SPTA1\_HUMAN | Spectrin alpha chain, erythrocytic 1 | FSSDFDELS[79.9663]GWMNEK | S | 383.0 | 50879.62 | 0.0 | 50879.62 | 0.0 |
| DMTN | DEMA\_HUMAN | Isoform 4 of Dematin | VFAMS[79.9663]PEEFGK | S | 383.0 | 50764.598 | 0.0 | 50764.598 | 0.0 |
| ADD1 | ADDA\_HUMAN | Isoform 3 of Alpha-adducin | EKS[79.9663]PPDQPAVPHPPPSTPIK | S | 600.0 | 50047.21 | 0.0 | 50047.21 | 0.0 |
| KRT4 | K2C4\_HUMAN | Keratin, type II cytoskeletal 4 | GAFS[79.9663]SVSM[15.9949]SGGAGR | S |  | 48787.664 | 0.0 | 48787.664 | 0.0 |
| CTPS1 | PYRG1\_HUMAN | CTP synthase 1 | SGSS[79.9663]SPDSEITELK | S | 574.0 | 48750.683 | 0.0 | 33593.96 | 15156.723 |
| SPTA1 | SPTA1\_HUMAN | Spectrin alpha chain, erythrocytic 1 | LEDS[79.9663]YHLQVFK | S | 52.0 | 48647.65 | 0.0 | 48647.65 | 0.0 |
| C2CD2L | C2C2L\_HUMAN | Phospholipid transfer protein C2CD2L | EAGLSQS[79.9663]HDDLSNATATPSVR | S | 662.0 | 47949.81 | 0.0 | 0.0 | 47949.81 |
| ATP7A | ATP7A\_HUMAN | Isoform 1 of Copper-transporting ATPase 1 | YNASS[79.9663]VTPESLR | S | 325.0 | 47903.78 | 0.0 | 0.0 | 47903.78 |
| SMIM1 | SMIM1\_HUMAN | Small integral membrane protein 1 | MQPQES[79.9663]HVHYSR | S | 6.0 | 47884.07 | 0.0 | 47884.07 | 0.0 |
| NEDD4L | NED4L\_HUMAN | Isoform 2 of E3 ubiquitin-protein ligase NEDD4-like | S[79.9663]LSSPTVTLSAPLEGAK | S | 446.0 | 47792.035 | 0.0 | 37425.84 | 10366.195 |
| SPTA1 | SPTA1\_HUMAN | Spectrin alpha chain, erythrocytic 1 | Q[-17.0265]DTLDAS[79.9663]LQSFQQER | S | 1976.0 | 47743.498 | 0.0 | 47743.498 | 0.0 |
| C2CD2L | C2C2L\_HUMAN | Phospholipid transfer protein C2CD2L | NLGTPTSS[79.9663]TPRPSITPTK | S | 421.0 | 47742.887 | 0.0 | 47742.887 | 0.0 |
| GYPC | GLPC\_HUMAN | Isoform Glycophorin-D of Glycophorin-C | GTEFAES[79.9663]ADAALQGDPALQDAGDSSR | S | 104.0 | 46918.586 | 0.0 | 46918.586 | 0.0 |
| KRT13 | K1C13\_HUMAN | Keratin, type I cytoskeletal 13 | MIGFPSSAGS[79.9663]VSPR | S | 425.0 | 46613.53 | 0.0 | 46613.53 | 0.0 |
| SLC16A1 | MOT1\_HUMAN | Monocarboxylate transporter 1 | EEET[79.9663]SIDVAGKPNEVTK | T | 466.0 | 46386.97 | 0.0 | 0.0 | 46386.97 |
| HSPA8 | HSP7C\_HUMAN | Heat shock cognate 71 kDa protein | NQVAMNPT[79.9663]NTVFDAK | T | 64.0 | 46317.84 | 0.0 | 46317.84 | 0.0 |
| ANK1 | ANK1\_HUMAN | Isoform Br21 of Ankyrin-1 | ELGESEGLS[79.9663]DDEETISTR | S |  | 45807.477 | 0.0 | 45807.477 | 0.0 |
| STOM | STOM\_HUMAN | Stomatin | NLS[79.9663]QILSDR | S | 161.0 | 45603.594 | 0.0 | 45603.594 | 0.0 |
| ADD1 | ADDA\_HUMAN | Isoform 3 of Alpha-adducin | GS[79.9663]EENLDEAR | S | 586.0 | 45295.51 | 0.0 | 45295.51 | 0.0 |
| EPB41 | EPB41\_HUMAN | Isoform 5 of Protein 4.1 | DVPIVHT[79.9663]ETK | T | 760.0 | 45053.656 | 0.0 | 45053.656 | 0.0 |
| NRBP1 | NRBP\_HUMAN | Nuclear receptor-binding protein | TPT[79.9663]PEPAEVETR | T | 433.0 | 45035.047 | 0.0 | 0.0 | 45035.047 |
| ARHGDIA | GDIR1\_HUMAN | Rho GDP-dissociation inhibitor 1 | AEEY[79.9663]EFLTPVEEAPK | Y | 156.0 | 44989.295 | 44989.295 | 0.0 | 0.0 |
| ANK1 | ANK1\_HUMAN | Isoform Br21 of Ankyrin-1 | VVTDETS[79.9663]FVLVSDK | S | 794.0 | 43965.617 | 0.0 | 43965.617 | 0.0 |
| DSC2 | DSC2\_HUMAN | Isoform 2B of Desmocollin-2 | AINDT[79.9663]AAR | T | 631.0 | 43623.633 | 43623.633 | 0.0 | 0.0 |
| SPTA1 | SPTA1\_HUMAN | Spectrin alpha chain, erythrocytic 1 | FQS[79.9663]ADETGQDLVNANHEASDEVR | S | 428.0 | 43521.906 | 0.0 | 43521.906 | 0.0 |
| TGM3 | TGM3\_HUMAN | Protein-glutamine gamma-glutamyltransferase E | LKPNT[79.9663]PFAATSSM[15.9949]GLETEEQEPSIIGK | T |  | 43228.71 | 0.0 | 43228.71 | 0.0 |
| ANK1 | ANK1\_HUMAN | Isoform Br21 of Ankyrin-1 | NGASPNEVSSDGT[79.9663]TPLAIAK | T | 768.0 | 42861.453 | 0.0 | 42861.453 | 0.0 |
| KRT2 | K22E\_HUMAN | Keratin, type II cytoskeletal 2 epidermal | Y[79.9663]LDGLTAER | Y | 239.0 | 42620.3 | 0.0 | 0.0 | 42620.3 |
| EPB41 | EPB41\_HUMAN | Isoform 5 of Protein 4.1 | S[79.9663]M[15.9949]TPAQADLEFLENAK | S |  | 42269.629 | 0.0 | 42269.629 | 0.0 |
| DMTN | DEMA\_HUMAN | Isoform 4 of Dematin | LQS[79.9663]TEFSPSGSETGSPGLQIYPYEM[15.9949]LVVTNK | S |  | 42211.71 | 0.0 | 42211.71 | 0.0 |
| SLC4A1 | B3AT\_HUMAN | Band 3 anion transport protein | DEY[79.9663]DEVAM[15.9949]PV | Y |  | 42141.17 | 0.0 | 0.0 | 42141.17 |
| SPTB | SPTB1\_HUMAN | Spectrin beta chain, erythrocytic | M[15.9949]QLLAASY[79.9663]DLHR | Y | 1797.0 | 41774.67 | 0.0 | 41774.67 | 0.0 |
| YWHAE | 1433E\_HUMAN | 14-3-3 protein epsilon | Y[79.9663]LAEFATGNDR | Y | 131.0 | 41755.62 | 0.0 | 41755.62 | 0.0 |
| SPTA1 | SPTA1\_HUMAN | Spectrin alpha chain, erythrocytic 1 | VKS[79.9663]LNQNM[15.9949]ESLR | S |  | 41525.6 | 0.0 | 41525.6 | 0.0 |
| ANK1 | ANK1\_HUMAN | Isoform Br21 of Ankyrin-1 | Q[-17.0265]DDAT[79.9663]GAGQDSENEVSLVSGHQR | T | 1660.0 | 41495.875 | 0.0 | 0.0 | 41495.875 |
| EPS15L1 | EP15R\_HUMAN | Epidermal growth factor receptor substrate 15-like 1 | STPSHGSVSSLNSTGS[79.9663]LSPK | S | 253.0 | 41266.977 | 0.0 | 41266.977 | 0.0 |
| CSNK1A1 | KC1A\_HUMAN | Isoform 3 of Casein kinase I isoform alpha | AAQQAASSS[79.9663]GQGQQAQTPTGK | S | 313.0 | 41075.727 | 41075.727 | 0.0 | 0.0 |
| GYPA | GLPA\_HUMAN | Isoform 2 of Glycophorin-A | SPSDVKPLPSPDTDVPLSSVEIENPET[79.9663]SDQ | T | 147.0 | 40936.02 | 0.0 | 40936.02 | 0.0 |
| DMTN | DEMA\_HUMAN | Isoform 4 of Dematin | ESVGGS[79.9663]PQTK | S | 156.0 | 40653.23 | 0.0 | 40653.23 | 0.0 |
| ANK1 | ANK1\_HUMAN | Isoform Br21 of Ankyrin-1 | AEDS[79.9663]DAT[79.9663]GHEWK | S |  | 40550.76 | 0.0 | 40550.76 | 0.0 |
| SPTA1 | SPTA1\_HUMAN | Spectrin alpha chain, erythrocytic 1 | VEAADHQGIVPAVY[79.9663]VR | Y | 1030.0 | 40473.5775 | 0.0 | 40473.5775 | 0.0 |
| STOM | STOM\_HUMAN | Stomatin | LPDSFKDS[79.9663]PSK | S | 22.0 | 40405.105 | 0.0 | 40405.105 | 0.0 |
| ADD2 | ADDB\_HUMAN | Beta-adducin | EAETKS[79.9663]PLVSPSK | S | 613.0 | 40208.312 | 0.0 | 40208.312 | 0.0 |
| CCRL2 | CCRL2\_HUMAN | Isoform 2 of C-C chemokine receptor-like 2 | GQSAQGT[79.9663]SREEPDHSTEV | T | 333.0 | 40043.071 | 0.0 | 18077.88 | 21965.191 |
| ADD2 | ADDB\_HUMAN | Beta-adducin | FSEDDPEY[79.9663]M[15.9949]R | Y |  | 39907.383 | 0.0 | 39907.383 | 0.0 |
| CTTN | SRC8\_HUMAN | Src substrate cortactin | AKT[79.9663]QTPPVSPAPQPTEER | T | 399.0 | 39418.75 | 39418.75 | 0.0 | 0.0 |
| ANK1 | ANK1\_HUMAN | Isoform Br21 of Ankyrin-1 | EPGGS[79.9663]LSFLR | S | 1339.0 | 39017.46 | 0.0 | 39017.46 | 0.0 |
| CA1 | CAH1\_HUMAN | Carbonic anhydrase 1 | Y[79.9663]SSLAEAASK | Y | 129.0 | 38937.438 | 0.0 | 20783.45 | 18153.988 |
| ADD2 | ADDB\_HUMAN | Beta-adducin | DKTESVTSGPM[15.9949]SPEGSPSKSPS[79.9663]K | S | 162.0 | 38735.35 | 0.0 | 38735.35 | 0.0 |
| C2CD5 | C2CD5\_HUMAN | Isoform 3 of C2 domain-containing protein 5 | QS[79.9663]SSSDTDLSLTPK | S | 304.0 | 38426.223 | 0.0 | 38426.223 | 0.0 |
| EIF4B | IF4B\_HUMAN | Eukaryotic translation initiation factor 4B | SQSS[79.9663]DTEQQSPTSGGGK | S | 498.0 | 38341.77 | 0.0 | 38341.77 | 0.0 |
| LSM14A | LS14A\_HUMAN | Isoform 2 of Protein LSM14 homolog A | S[79.9663]PVSTRPLPSASQK | S | 216.0 | 38302.44 | 0.0 | 38302.44 | 0.0 |
| PI4K2A | P4K2A\_HUMAN | Phosphatidylinositol 4-kinase type 2-alpha | S[79.9663]SSESYTQSFQSR | S | 460.0 | 38283.26 | 0.0 | 11429.463 | 26853.797 |
| ERMAP | ERMAP\_HUMAN | Erythroid membrane-associated protein | VNS[79.9663]SLLPPK | S | 442.0 | 38279.875 | 0.0 | 0.0 | 38279.875 |
| METAP2 | MAP2\_HUMAN | Isoform 3 of Methionine aminopeptidase 2 | EEGAAS[79.9663]TAEEAAK | S | 29.0 | 38142.526 | 0.0 | 18975.932 | 19166.594 |
| VAMP3 | VAMP3\_HUMAN | Vesicle-associated membrane protein 3 | ADALQAGASQFET[79.9663]SAAK | T | 62.0 | 37584.512 | 22850.996 | 0.0 | 14733.516 |
| PSEN1 | PSN1\_HUMAN | Isoform 2 of Presenilin-1 | AAVQELSSS[79.9663]ILAGEDPEER | S | 367.0 | 37444.4245 | 0.0 | 7104.2295 | 30340.195 |
| TAB3 | TAB3\_HUMAN | Isoform 2 of TGF-beta-activated kinase 1 and MAP3K7-binding protein 3 | VIPNPT[79.9663]T[79.9663]VFK | T |  | 37387.0 | 0.0 | 37387.0 | 0.0 |
| EIF4B | IF4B\_HUMAN | Eukaryotic translation initiation factor 4B | SQS[79.9663]SDTEQQSPTSGGGK | S | 497.0 | 37348.504 | 0.0 | 37348.504 | 0.0 |
| ADD2 | ADDB\_HUMAN | Beta-adducin | VTM[15.9949]ILQS[79.9663]PSFR | S | 60.0 | 36864.367 | 0.0 | 36864.367 | 0.0 |
| EPB41 | EPB41\_HUMAN | Isoform 5 of Protein 4.1 | RASRS[79.9663]LDGAAAVDSADR | S | 542.0 | 36819.133 | 0.0 | 36819.133 | 0.0 |
| KRT6A | K2C6A\_HUMAN | Keratin, type II cytoskeletal 6A | AIGGGLSSVGGGS[79.9663]STIK | S | 546.0 | 36438.51 | 0.0 | 36438.51 | 0.0 |
| SLC2A1 | GTR1\_HUMAN | Solute carrier family 2, facilitated glucose transporter member 1 | LRGT[79.9663]ADVTHDLQEM[15.9949]K | T |  | 36337.71 | 0.0 | 0.0 | 36337.71 |
| PSMA5 | PSA5\_HUMAN | Proteasome subunit alpha type-5 | IT[79.9663]SPLMEPSSIEK | T | 55.0 | 35846.13 | 0.0 | 35846.13 | 0.0 |
| PIEZO1 | PIEZ1\_HUMAN | Piezo-type mechanosensitive ion channel component 1 | DPGLEPGPDS[79.9663]PGGSSPPR | S | 1391.0 | 35756.625 | 0.0 | 0.0 | 35756.625 |
| SPTB | SPTB1\_HUMAN | Spectrin beta chain, erythrocytic | VY[79.9663]TPHDGK | Y | 381.0 | 35331.89 | 0.0 | 0.0 | 35331.89 |
| DBNL | DBNL\_HUMAN | Isoform 2 of Drebrin-like protein | AM[15.9949]STTSISS[79.9663]PQPGK | S | 275.0 | 35093.85 | 35093.85 | 0.0 | 0.0 |
| USP24 | UBP24\_HUMAN | Ubiquitin carboxyl-terminal hydrolase 24 | VSDQNS[79.9663]PVLPK | S | 2047.0 | 34939.64 | 0.0 | 0.0 | 34939.64 |
| SPTA1 | SPTA1\_HUMAN | Spectrin alpha chain, erythrocytic 1 | ES[79.9663]LNEAQK | S | 1284.0 | 34867.95 | 0.0 | 34867.95 | 0.0 |
| SLC29A1 | S29A1\_HUMAN | Isoform 2 of Equilibrative nucleoside transporter 1 | EESGVSVSNS[79.9663]QPTNESHSIK | S | 273.0 | 34764.656 | 0.0 | 0.0 | 34764.656 |
| SLC43A1 | LAT3\_HUMAN | Large neutral amino acids transporter small subunit 3 | GT[79.9663]SENLPER | T | 284.0 | 34571.246 | 0.0 | 34571.246 | 0.0 |
| KRT6A | K2C6A\_HUMAN | Keratin, type II cytoskeletal 6A | SGFSSVS[79.9663]VSR | S | 37.0 | 34554.42 | 0.0 | 34554.42 | 0.0 |
| REEP4 | REEP4\_HUMAN | Receptor expression-enhancing protein 4 | S[79.9663]FSM[15.9949]QDLR | S |  | 34454.273 | 0.0 | 0.0 | 34454.273 |
| SERINC1 | SERC1\_HUMAN | Serine incorporator 1 | SDGS[79.9663]LEDGDDVHR | S | 364.0 | 34010.215 | 0.0 | 22105.729 | 11904.486 |
| ADD2 | ADDB\_HUMAN | Beta-adducin | EEEQTAEEILS[79.9663]K | S | 162.0 | 33924.94 | 0.0 | 33924.94 | 0.0 |
| HRNR | HORN\_HUMAN | Hornerin | YGQQGSGSGQS[79.9663]PSR | S | 542.0 | 33912.954 | 0.0 | 12637.858 | 21275.096 |
| BAG6 | BAG6\_HUMAN | Isoform 2 of Large proline-rich protein BAG6 | ENAS[79.9663]PAPGTTAEEAM[15.9949]SR | S |  | 33815.85 | 0.0 | 10160.358 | 23655.492 |
| DNAJC6 | AUXI\_HUMAN | Isoform 2 of Putative tyrosine-protein phosphatase auxilin | SATSTSAS[79.9663]PTLR | S | 570.0 | 33785.438 | 0.0 | 33785.438 | 0.0 |
| HBB | HBB\_HUMAN | Hemoglobin subunit beta | SAVT[79.9663]ALWGK | T | 13.0 | 33477.582 | 0.0 | 33477.582 | 0.0 |
| EPB41 | EPB41\_HUMAN | Isoform 5 of Protein 4.1 | LS[79.9663]M[15.9949]YGVDLHK | S |  | 33400.682 | 0.0 | 33400.682 | 0.0 |
| SPTB | SPTB1\_HUMAN | Spectrin beta chain, erythrocytic | AQEAS[79.9663]VLLR | S | 1267.0 | 33233.04 | 0.0 | 33233.04 | 0.0 |
| ANK1 | ANK1\_HUMAN | Isoform Br21 of Ankyrin-1 | YSILSESTPGS[79.9663]LSGTEQAEM[15.9949]K | S |  | 33062.55 | 0.0 | 33062.55 | 0.0 |
| YBX1 | YBOX1\_HUMAN | Y-box-binding protein 1 | AADPPAENSS[79.9663]APEAEQGGAE | S | 314.0 | 33014.36 | 33014.36 | 0.0 | 0.0 |
| CA1 | CAH1\_HUMAN | Carbonic anhydrase 1 | HDTSLKPIS[79.9663]VSYNPATAK | S | 49.0 | 32865.773 | 0.0 | 32865.773 | 0.0 |
| ANK1 | ANK1\_HUMAN | Isoform Br21 of Ankyrin-1 | ELQFS[79.9663]VEDINR | S | 1428.0 | 32776.942 | 0.0 | 8952.044 | 23824.898 |
| TRIM29 | TRI29\_HUMAN | Isoform Beta of Tripartite motif-containing protein 29 | TSYQPSS[79.9663]PGR | S | 489.0 | 32581.71 | 0.0 | 32581.71 | 0.0 |
| ANK1 | ANK1\_HUMAN | Isoform Br21 of Ankyrin-1 | ITHS[79.9663]PTVSQVT[79.9663]ER | S |  | 32549.104 | 0.0 | 32549.104 | 0.0 |
| CSNK1G1 | KC1G1\_HUMAN | Isoform 1S of Casein kinase I isoform gamma-1 | DRPS[79.9663]QQQPLR | S | 361.0 | 32420.305 | 0.0 | 17224.998 | 15195.307 |
| TMCC2 | TMCC2\_HUMAN | Isoform 2 of Transmembrane and coiled-coil domains protein 2 | ALSGSATLVS[79.9663]SPK | S | 469.0 | 31806.402 | 0.0 | 0.0 | 31806.402 |
| NEDD4L | NED4L\_HUMAN | Isoform 2 of E3 ubiquitin-protein ligase NEDD4-like | T[79.9663]SPQELSEELSR | T | 302.0 | 31627.814 | 0.0 | 31627.814 | 0.0 |
| SPTA1 | SPTA1\_HUMAN | Spectrin alpha chain, erythrocytic 1 | GVSEET[79.9663]LKEFSTIYK | T | 2272.0 | 31521.678 | 0.0 | 31521.678 | 0.0 |
| EPB41 | EPB41\_HUMAN | Isoform 5 of Protein 4.1 | PTSAPAITQGQVAEGGVLDAS[79.9663]AKK | S | 578.0 | 31512.951 | 0.0 | 31512.951 | 0.0 |
| MICALL2 | MILK2\_HUMAN | MICAL-like protein 2 | PAS[79.9663]PGPSLPAR | S | 649.0 | 31331.098 | 0.0 | 0.0 | 31331.098 |
| HBE1 | HBE\_HUMAN | Hemoglobin subunit epsilon | AAVTSLWS[79.9663]KMNVEEAGGEALGR | S | 17.0 | 31303.424 | 31303.424 | 0.0 | 0.0 |
| EPB41 | EPB41\_HUMAN | Isoform 5 of Protein 4.1 | EESPQS[79.9663]KAETELK | S | 191.0 | 30886.701 | 0.0 | 30886.701 | 0.0 |
| HSPB1 | HSPB1\_HUMAN | Heat shock protein beta-1 | Q[-17.0265]LS[79.9663]SGVSEIR | S | 82.0 | 30650.123 | 0.0 | 17994.988 | 12655.135 |
| ANK1 | ANK1\_HUMAN | Isoform Br21 of Ankyrin-1 | ITHSPTVSQVT[79.9663]ER | T | 1693.0 | 30572.576 | 0.0 | 30572.576 | 0.0 |
| DMTN | DEMA\_HUMAN | Isoform 4 of Dematin | QPLT[79.9663]SPGSVS[79.9663]PSR | T |  | 30473.645 | 0.0 | 30473.645 | 0.0 |
| SPTB | SPTB1\_HUMAN | Spectrin beta chain, erythrocytic | LSS[79.9663]SWESLQPEPSHPY | S | 2124.0 | 30468.316 | 0.0 | 30468.316 | 0.0 |
| HSP90AA1 | HS90A\_HUMAN | Isoform 2 of Heat shock protein HSP 90-alpha | DKEVS[79.9663]DDEAEEKEDK | S | 231.0 | 30440.196 | 0.0 | 17731.611 | 12708.585 |
| HSP90AB1 | HS90B\_HUMAN | Heat shock protein HSP 90-beta | EKEIS[79.9663]DDEAEEEK | S | 226.0 | 30209.5485 | 0.0 | 30209.5485 | 0.0 |
| ADD2 | ADDB\_HUMAN | Beta-adducin | SPS[79.9663]TESQLMSK | S | 532.0 | 30026.924 | 0.0 | 30026.924 | 0.0 |
| SPTB | SPTB1\_HUMAN | Spectrin beta chain, erythrocytic | ES[79.9663]QQLM[15.9949]DSHPEQK | S |  | 29785.313 | 0.0 | 29785.313 | 0.0 |
| YES1 | YES\_HUMAN | Tyrosine-protein kinase Yes | GAY[79.9663]SLSIR | Y | 194.0 | 29740.729 | 0.0 | 0.0 | 29740.729 |
| ACTBL2 | ACTBL\_HUMAN | Beta-actin-like protein 2 | EIIT[79.9663]LAPSTM[15.9949]K | T |  | 29709.572 | 0.0 | 29709.572 | 0.0 |
| SPTB | SPTB1\_HUMAN | Spectrin beta chain, erythrocytic | n[42.0106]TSAT[79.9663]EFENVGNQPPYSR | T | 5.0 | 29487.879 | 0.0 | 29487.879 | 0.0 |
| KRT5 | K2C5\_HUMAN | Keratin, type II cytoskeletal 5 | S[79.9663]FSTASAITPSVSR | S | 16.0 | 29384.873 | 0.0 | 29384.873 | 0.0 |
| VAMP3 | VAMP3\_HUMAN | Vesicle-associated membrane protein 3 | n[42.0106]ST[79.9663]GPTAATGSNR | T | 3.0 | 29251.082 | 0.0 | 0.0 | 29251.082 |
| HBB | HBB\_HUMAN | Hemoglobin subunit beta | S[79.9663]AVTALWGK | S | 10.0 | 29222.689 | 0.0 | 0.0 | 29222.689 |
| SPTB | SPTB1\_HUMAN | Spectrin beta chain, erythrocytic | M[15.9949]QLLAAS[79.9663]YDLHR | S | 1796.0 | 29185.752 | 0.0 | 29185.752 | 0.0 |
| ADD2 | ADDB\_HUMAN | Beta-adducin | GLSQMTTSADT[79.9663]DVDTSK | T | 675.0 | 29144.807 | 0.0 | 29144.807 | 0.0 |
| PRPSAP1 | KPRA\_HUMAN | Isoform 2 of Phosphoribosyl pyrophosphate synthase-associated protein 1 | AGLTHIIT[79.9663]M[15.9949]DLHQKEIQGFFSFPVDNLR | T |  | 29060.34 | 29060.34 | 0.0 | 0.0 |
| KRT5 | K2C5\_HUMAN | Keratin, type II cytoskeletal 5 | EY[79.9663]QELM[15.9949]NTK | Y |  | 29021.719 | 0.0 | 0.0 | 29021.719 |
| FLNA | FLNA\_HUMAN | Isoform 2 of Filamin-A | GT[79.9663]VEPQLEAR | T | 429.0 | 28979.414 | 0.0 | 0.0 | 28979.414 |
| ADD2 | ADDB\_HUMAN | Beta-adducin | ADEVEKS[79.9663]SSGM[15.9949]PIR | S |  | 28899.543 | 28899.543 | 0.0 | 0.0 |
| HBB | HBB\_HUMAN | Hemoglobin subunit beta | VVAGVANALAHKY[79.9663]H | Y | 146.0 | 28863.623 | 0.0 | 28863.623 | 0.0 |
| EPB41 | EPB41\_HUMAN | Isoform 5 of Protein 4.1 | HS[79.9663]NLM[15.9949]LEDLDK | S |  | 28834.674 | 0.0 | 28834.674 | 0.0 |
| STX7 | STX7\_HUMAN | Isoform 2 of Syntaxin-7 | EFGSLPT[79.9663]TPSEQR | T | 78.0 | 28777.102 | 0.0 | 28777.102 | 0.0 |
| SPTB | SPTB1\_HUMAN | Spectrin beta chain, erythrocytic | PAEETGPQEEEGET[79.9663]AGEAPVSHHAAT[79.9663]ER | T |  | 28665.031 | 0.0 | 28665.031 | 0.0 |
| ANK1 | ANK1\_HUMAN | Isoform Br21 of Ankyrin-1 | Q[-17.0265]DDATGAGQDS[79.9663]ENEVSLVSGHQR | S | 1666.0 | 28600.836 | 0.0 | 28600.836 | 0.0 |
| EPB41 | EPB41\_HUMAN | Isoform 5 of Protein 4.1 | TITYEAAQT[79.9663]VK | T | 591.0 | 28361.611 | 0.0 | 28361.611 | 0.0 |
| SLC14A1 | UT1\_HUMAN | Urea transporter 1 | n[42.0106]MEDS[79.9663]PTM[15.9949]VR | S |  | 28065.85 | 0.0 | 28065.85 | 0.0 |
| EPB41 | EPB41\_HUMAN | Isoform 5 of Protein 4.1 | DLDKS[79.9663]QEEIK | S | 674.0 | 28029.406 | 0.0 | 28029.406 | 0.0 |
| HSP90AB1 | HS90B\_HUMAN | Heat shock protein HSP 90-beta | IEDVGS[79.9663]DEEDDSGK | S | 255.0 | 27856.01 | 0.0 | 27856.01 | 0.0 |
| ABCC1 | MRP1\_HUMAN | Isoform 2 of Multidrug resistance-associated protein 1 | QLSSSSS[79.9663]YSGDISR | S | 919.0 | 27825.102 | 0.0 | 0.0 | 27825.102 |
| SNAP23 | SNP23\_HUMAN | Synaptosomal-associated protein 23 | M[15.9949]DNLS[79.9663]SEEIQQR | S | 5.0 | 27719.744 | 0.0 | 27719.744 | 0.0 |
| ADD2 | ADDB\_HUMAN | Beta-adducin | ADEVEKSS[79.9663]SGMPIR | S | 472.0 | 27589.309 | 27589.309 | 0.0 | 0.0 |
| YBX1 | YBOX1\_HUMAN | Y-box-binding protein 1 | AADPPAENS[79.9663]SAPEAEQGGAE | S | 313.0 | 27512.74 | 27512.74 | 0.0 | 0.0 |
| STOM | STOM\_HUMAN | Stomatin | VQNATLAVANITNADS[79.9663]ATR | S | 141.0 | 27508.442 | 0.0 | 14032.748 | 13475.694 |
| YWHAG | 1433G\_HUMAN | 14-3-3 protein gamma | YDDM[15.9949]AAAM[15.9949]KNVT[79.9663]ELNEPLSNEER | T | 31.0 | 27408.848 | 0.0 | 27408.848 | 0.0 |
| ANK1 | ANK1\_HUMAN | Isoform Br21 of Ankyrin-1 | ITHS[79.9663]PT[79.9663]VSQVTER | S |  | 27350.428 | 0.0 | 27350.428 | 0.0 |
| TMBIM1 | LFG3\_HUMAN | Protein lifeguard 3 | AVS[79.9663]DSFGPGEWDDR | S | 81.0 | 27194.848 | 0.0 | 27194.848 | 0.0 |
| ADD2 | ADDB\_HUMAN | Beta-adducin | SAGPQSQLLAS[79.9663]VIAEK | S | 522.0 | 27011.246 | 0.0 | 27011.246 | 0.0 |
| KRT2 | K22E\_HUMAN | Keratin, type II cytoskeletal 2 epidermal | GFSSGS[79.9663]AVVSGGSR | S | 26.0 | 27002.594 | 0.0 | 0.0 | 27002.594 |
| SPTA1 | SPTA1\_HUMAN | Spectrin alpha chain, erythrocytic 1 | ET[79.9663]GTLESQLEANK | T | 2189.0 | 26894.166 | 0.0 | 26894.166 | 0.0 |
| ABCC1 | MRP1\_HUMAN | Isoform 2 of Multidrug resistance-associated protein 1 | Q[-17.0265]LSSSSS[79.9663]YSGDISR | S | 919.0 | 26504.262 | 0.0 | 0.0 | 26504.262 |
| REPS1 | REPS1\_HUMAN | Isoform 2 of RalBP1-associated Eps domain-containing protein 1 | FVAS[79.9663]KNEQESR | S | 104.0 | 26490.883 | 0.0 | 26490.883 | 0.0 |
| SPTA1 | SPTA1\_HUMAN | Spectrin alpha chain, erythrocytic 1 | TGQEM[15.9949]IEGGHY[79.9663]ASDNVTTR | Y | 653.0 | 26401.027 | 0.0 | 26401.027 | 0.0 |
| SPTA1 | SPTA1\_HUMAN | Spectrin alpha chain, erythrocytic 1 | KY[79.9663]LKHQTFAHEVDGR | Y | 1538.0 | 26345.758 | 0.0 | 26345.758 | 0.0 |
| KRT13 | K1C13\_HUMAN | Keratin, type I cytoskeletal 13 | EVSTNT[79.9663]AM[15.9949]IQTSK | T |  | 26049.229 | 0.0 | 16373.407 | 9675.822 |
| EPB41 | EPB41\_HUMAN | Isoform 5 of Protein 4.1 | S[79.9663]EIPTKDVPIVHTETK | S | 748.0 | 25944.621 | 0.0 | 25944.621 | 0.0 |
| TSC22D4 | T22D4\_HUMAN | TSC22 domain family protein 4 | NGS[79.9663]PPPGAPSSR | S | 62.0 | 25467.297 | 0.0 | 0.0 | 25467.297 |
| DENND1A | DEN1A\_HUMAN | DENN domain-containing protein 1A | T[79.9663]GGTLSDPEVQR | T | 24.0 | 25192.102 | 0.0 | 25192.102 | 0.0 |
| SYAP1 | SYAP1\_HUMAN | Synapse-associated protein 1 | T[79.9663]PPVVIK | T | 248.0 | 24888.54 | 0.0 | 0.0 | 24888.54 |
| ACTBL2 | ACTBL\_HUMAN | Beta-actin-like protein 2 | M[15.9949]QKEIIT[79.9663]LAPSTM[15.9949]K | T |  | 24844.103 | 0.0 | 24844.103 | 0.0 |
| KRT10 | K1C10\_HUMAN | Keratin, type I cytoskeletal 10 | S[79.9663]QYEQLAEQNR | S | 323.0 | 24667.6123 | 0.0 | 13520.539 | 11147.0733 |
| HBA1 | HBA\_HUMAN | Hemoglobin subunit alpha | FLASVST[79.9663]VLTSK | T | 135.0 | 24524.46 | 0.0 | 0.0 | 24524.46 |
| EPN2 | EPN2\_HUMAN | Isoform 2 of Epsin-2 | AGGSPASYHGSTS[79.9663]PR | S | 258.0 | 24384.65 | 24384.65 | 0.0 | 0.0 |
| PSMA5 | PSA5\_HUMAN | Proteasome subunit alpha type-5 | ITS[79.9663]PLMEPSSIEK | S | 56.0 | 24307.719 | 0.0 | 24307.719 | 0.0 |
| PTGS1 | PGH1\_HUMAN | Isoform 2 of Prostaglandin G/H synthase 1 | LILIGET[79.9663]IK | T | 339.0 | 24187.945 | 0.0 | 24187.945 | 0.0 |
| CD44 | CD44\_HUMAN | Isoform 10 of CD44 antigen | S[79.9663]QEMVHLVNK | S | 706.0 | 24046.977 | 0.0 | 24046.977 | 0.0 |
| CTTN | SRC8\_HUMAN | Src substrate cortactin | T[79.9663]QTPPVSPAPQPTEER | T | 399.0 | 24030.8 | 0.0 | 24030.8 | 0.0 |
| DMTN | DEMA\_HUMAN | Isoform 4 of Dematin | T[79.9663]SLPHFHHPETSR | T | 123.0 | 23999.98 | 0.0 | 23999.98 | 0.0 |
| CLN6 | CLN6\_HUMAN | Isoform 2 of Ceroid-lipofuscinosis neuronal protein 6 | HGS[79.9663]VSADEAAR | S | 31.0 | 23904.506 | 0.0 | 0.0 | 23904.506 |
| ANK1 | ANK1\_HUMAN | Isoform Br21 of Ankyrin-1 | VVTDET[79.9663]SFVLVSDK | T | 793.0 | 23852.652 | 0.0 | 23852.652 | 0.0 |
| PIEZO1 | PIEZ1\_HUMAN | Piezo-type mechanosensitive ion channel component 1 | Q[-17.0265]DAVSGT[79.9663]PLLR | T | 734.0 | 23844.328 | 0.0 | 0.0 | 23844.328 |
| SPTB | SPTB1\_HUMAN | Spectrin beta chain, erythrocytic | LQT[79.9663]HADLNK | T | 1387.0 | 23803.607 | 0.0 | 23803.607 | 0.0 |
| SPTA1 | SPTA1\_HUMAN | Spectrin alpha chain, erythrocytic 1 | QDTLDASLQS[79.9663]FQQER | S | 1979.0 | 23727.336 | 0.0 | 23727.336 | 0.0 |
| YWHAZ | 1433Z\_HUMAN | 14-3-3 protein zeta/delta | Y[79.9663]LAEVAAGDDKK | Y | 128.0 | 23684.24 | 0.0 | 23684.24 | 0.0 |
| ABCC5 | MRP5\_HUMAN | ATP-binding cassette sub-family C member 5 | NATLAWDSSHSSIQNS[79.9663]PK | S | 509.0 | 23630.721 | 0.0 | 14671.268 | 8959.453 |
| SPTA1 | SPTA1\_HUMAN | Spectrin alpha chain, erythrocytic 1 | YQKHQS[79.9663]LEAEVQTK | S | 96.0 | 23618.125 | 0.0 | 23618.125 | 0.0 |
| CRNN | CRNN\_HUMAN | Cornulin | IS[79.9663]PQIQLSGQTEQTQK | S | 180.0 | 23609.36 | 0.0 | 0.0 | 23609.36 |
| KRT13 | K1C13\_HUMAN | Keratin, type I cytoskeletal 13 | M[15.9949]IGFPSS[79.9663]AGSVSPR | S | 422.0 | 23155.21 | 0.0 | 0.0 | 23155.21 |
| KRT33B | KT33B\_HUMAN | Keratin, type I cuticular Ha3-II | S[79.9663]DLEAQM[15.9949]ESLK | S |  | 23138.115 | 0.0 | 23138.115 | 0.0 |
| EHBP1L1 | EH1L1\_HUMAN | EH domain-binding protein 1-like protein 1 | ANEAGGQVGPEAPRPPET[79.9663]SPEM[15.9949]R | T |  | 23107.459 | 23107.459 | 0.0 | 0.0 |
| TMEM63B | CSCL2\_HUMAN | CSC1-like protein 2 | LTSVSS[79.9663]SVDFDQR | S | 114.0 | 23100.504 | 0.0 | 0.0 | 23100.504 |
| PNPLA6 | PLPL6\_HUMAN | Isoform 3 of Patatin-like phospholipase domain-containing protein 6 | VSQSTS[79.9663]SLVDTSVSATSR | S | 120.0 | 22897.135 | 0.0 | 0.0 | 22897.135 |
| PSMD2 | PSMD2\_HUMAN | 26S proteasome non-ATPase regulatory subunit 2 | APVQPQQS[79.9663]PAAAPGGTDEKPSGK | S | 16.0 | 22699.207 | 22699.207 | 0.0 | 0.0 |
| ANK1 | ANK1\_HUMAN | Isoform Br21 of Ankyrin-1 | ELVNY[79.9663]GANVNAQSQK | Y | 100.0 | 22494.755 | 0.0 | 12492.485 | 10002.27 |
| DMTN | DEMA\_HUMAN | Isoform 4 of Dematin | QRES[79.9663]VGGS[79.9663]PQTK | S |  | 22417.54 | 0.0 | 22417.54 | 0.0 |
| SPTA1 | SPTA1\_HUMAN | Spectrin alpha chain, erythrocytic 1 | SDDKSSLDS[79.9663]LEALM[15.9949]K | S |  | 22275.092 | 0.0 | 22275.092 | 0.0 |
| HSP90AA1 | HS90A\_HUMAN | Isoform 2 of Heat shock protein HSP 90-alpha | ESEDKPEIEDVGS[79.9663]DEEEEK | S | 263.0 | 22258.5565 | 0.0 | 22258.5565 | 0.0 |
| RAPGEF2 | RPGF2\_HUMAN | Rap guanine nucleotide exchange factor 2 | SETS[79.9663]PVAPR | S | 1022.0 | 22148.041 | 0.0 | 22148.041 | 0.0 |
| EPB42 | EPB42\_HUMAN | Isoform Long of Protein 4.2 | VLPT[79.9663]PQTQATQEGALLNK | T | 230.0 | 22027.129 | 0.0 | 22027.129 | 0.0 |
| MINK1 | MINK1\_HUMAN | Isoform 1 of Misshapen-like kinase 1 | VGVSSKPDS[79.9663]SPVLSPGNK | S | 777.0 | 21281.422 | 0.0 | 21281.422 | 0.0 |
| SPTB | SPTB1\_HUMAN | Spectrin beta chain, erythrocytic | LQT[79.9663]AYAGEK | T | 1864.0 | 20894.795 | 0.0 | 0.0 | 20894.795 |
| YWHAB | 1433B\_HUMAN | 14-3-3 protein beta/alpha | Y[79.9663]LSEVASGDNK | Y | 130.0 | 20783.02 | 0.0 | 20783.02 | 0.0 |
| FKBP4 | FKBP4\_HUMAN | Peptidyl-prolyl cis-trans isomerase FKBP4 | AEASSGDHPT[79.9663]DTEM[15.9949]KEEQK | T |  | 20767.068 | 20767.068 | 0.0 | 0.0 |
| ADD1 | ADDA\_HUMAN | Isoform 3 of Alpha-adducin | S[79.9663]LVQGELVTASK | S | 532.0 | 20733.848 | 0.0 | 20733.848 | 0.0 |
| SPTA1 | SPTA1\_HUMAN | Spectrin alpha chain, erythrocytic 1 | NLAVMS[79.9663]DKVK | S | 317.0 | 20651.203 | 0.0 | 20651.203 | 0.0 |
| HBA1 | HBA\_HUMAN | Hemoglobin subunit alpha | AAWGKVGAHAGEY[79.9663]GAEALER | Y | 25.0 | 20603.723 | 20603.723 | 0.0 | 0.0 |
| SPTB | SPTB1\_HUMAN | Spectrin beta chain, erythrocytic | S[79.9663]TASWAER | S | 2057.0 | 20532.031 | 0.0 | 20532.031 | 0.0 |
| ITSN1 | ITSN1\_HUMAN | Isoform 10 of Intersectin-1 | SAFTPATATGSS[79.9663]PSPVLGQGEK | S | 902.0 | 20439.303 | 0.0 | 20439.303 | 0.0 |
| SPTB | SPTB1\_HUMAN | Spectrin beta chain, erythrocytic | WQAFQT[79.9663]LVSER | T | 941.0 | 20357.713 | 0.0 | 20357.713 | 0.0 |
| VAMP3 | VAMP3\_HUMAN | Vesicle-associated membrane protein 3 | ADALQAGASQFETS[79.9663]AAK | S | 63.0 | 20229.736 | 0.0 | 0.0 | 20229.736 |
| SMIM1 | SMIM1\_HUMAN | Small integral membrane protein 1 | DGVSLGAVS[79.9663]S[79.9663]TEEASR | S |  | 20115.52 | 0.0 | 0.0 | 20115.52 |
| SMAP2 | SMAP2\_HUMAN | Isoform 2 of Stromal membrane-associated protein 2 | LYEAYLPET[79.9663]FRR | T | 100.0 | 20067.441 | 0.0 | 20067.441 | 0.0 |
| KRT5 | K2C5\_HUMAN | Keratin, type II cytoskeletal 5 | SFSTASAIT[79.9663]PSVSR | T | 24.0 | 19739.725 | 0.0 | 0.0 | 19739.725 |
| SPTBN4 | SPTN4\_HUMAN | Isoform 4 of Spectrin beta chain, non-erythrocytic 4 | ES[79.9663]WLNENQR | S | 443.0 | 19724.645 | 0.0 | 19724.645 | 0.0 |
| SPTA1 | SPTA1\_HUMAN | Spectrin alpha chain, erythrocytic 1 | VADDLLFEGLLT[79.9663]PEGAQIR | T | 1151.0 | 19669.896 | 0.0 | 19669.896 | 0.0 |
| KRT1 | K2C1\_HUMAN | Keratin, type II cytoskeletal 1 | SKAEAESLYQS[79.9663]K | S | 73.0 | 19655.09 | 0.0 | 0.0 | 19655.09 |
| KRT9 | K1C9\_HUMAN | Keratin, type I cytoskeletal 9 | S[79.9663]GGGGGGGLGSGGSIR | S | 14.0 | 19599.961 | 0.0 | 8602.018 | 10997.943 |
| MINPP1 | MINP1\_HUMAN | Multiple inositol polyphosphate phosphatase 1 | DPVASS[79.9663]LSPYFGTK | S | 42.0 | 19551.889 | 0.0 | 19551.889 | 0.0 |
| EPS15L1 | EP15R\_HUMAN | Epidermal growth factor receptor substrate 15-like 1 | STPSHGSVSSLNSTGSLS[79.9663]PK | S | 255.0 | 19501.133 | 0.0 | 19501.133 | 0.0 |
| ITSN2 | ITSN2\_HUMAN | Isoform 2 of Intersectin-2 | AQS[79.9663]LIDLGSSSSTSSTASLSGNSPK | S | 210.0 | 19480.049 | 0.0 | 19480.049 | 0.0 |
| SPTA1 | SPTA1\_HUMAN | Spectrin alpha chain, erythrocytic 1 | LQATY[79.9663]WYHR | Y | 370.0 | 19479.059 | 0.0 | 19479.059 | 0.0 |
| MICAL3 | MICA3\_HUMAN | [F-actin]-monooxygenase MICAL3 | GKS[79.9663]EEELEASK | S | 977.0 | 19439.945 | 0.0 | 0.0 | 19439.945 |
| SPTA1 | SPTA1\_HUMAN | Spectrin alpha chain, erythrocytic 1 | LSES[79.9663]HPDATEDLQR | S | 1252.0 | 19303.29 | 0.0 | 19303.29 | 0.0 |
| EPB41 | EPB41\_HUMAN | Isoform 5 of Protein 4.1 | EDEPPEQAEPEPT[79.9663]EAWK | T | 611.0 | 19153.088 | 0.0 | 19153.088 | 0.0 |
| PGK1 | PGK1\_HUMAN | Phosphoglycerate kinase 1 | AC[57.0215]ANPAAGS[79.9663]VILLENLR | S | 115.0 | 19143.594 | 19143.594 | 0.0 | 0.0 |
| YWHAH | 1433F\_HUMAN | 14-3-3 protein eta | YDDM[15.9949]AS[79.9663]AMK | S | 25.0 | 18640.256 | 0.0 | 0.0 | 18640.256 |
| ST13 | F10A1\_HUMAN | Hsc70-interacting protein | ADEPS[79.9663]SEESDLEIDK | S | 75.0 | 18233.979 | 0.0 | 18233.979 | 0.0 |
| DMTN | DEMA\_HUMAN | Isoform 4 of Dematin | ES[79.9663]VGGSPQTK | S | 152.0 | 18225.776 | 0.0 | 8946.084 | 9279.692 |
| KLC1 | KLC1\_HUMAN | Isoform J of Kinesin light chain 1 | AS[79.9663]SLNVLNVGGK | S |  | 18127.756 | 0.0 | 18127.756 | 0.0 |
| KEL | KELL\_HUMAN | Kell blood group glycoprotein | SQAGGM[15.9949]GTLWSQES[79.9663]TPEER | S | 28.0 | 18061.54 | 0.0 | 18061.54 | 0.0 |
| USP15 | UBP15\_HUMAN | Ubiquitin carboxyl-terminal hydrolase 15 | S[79.9663]PGASNFSTLPK | S | 229.0 | 18057.826 | 0.0 | 0.0 | 18057.826 |
| SPTB | SPTB1\_HUMAN | Spectrin beta chain, erythrocytic | LS[79.9663]SSWES[79.9663]LQPEPSHPY | S |  | 18036.793 | 0.0 | 18036.793 | 0.0 |
| YES1 | YES\_HUMAN | Tyrosine-protein kinase Yes | LIEDNEY[79.9663]TAR | Y | 426.0 | 18000.426 | 0.0 | 0.0 | 18000.426 |
| PNPLA6 | PLPL6\_HUMAN | Isoform 3 of Patatin-like phospholipase domain-containing protein 6 | VS[79.9663]QSTSSLVDTSVSATSR | S | 116.0 | 17866.406 | 0.0 | 0.0 | 17866.406 |
| FRMD4A | FRM4A\_HUMAN | FERM domain-containing protein 4A | AAGALGSASSGS[79.9663]M[15.9949]PNLAAR | S |  | 17824.115 | 0.0 | 17824.115 | 0.0 |
| MARCHF8 | MARH8\_HUMAN | Isoform 2 of E3 ubiquitin-protein ligase MARCHF8 | AGSPPSASAPAPVS[79.9663]SFSR | S | 65.0 | 17679.61 | 0.0 | 0.0 | 17679.61 |
| YBX3 | YBOX3\_HUMAN | Y-box-binding protein 3 | AGEAPTENPAPPT[79.9663]QQSSAE | T | 366.0 | 17626.402 | 17626.402 | 0.0 | 0.0 |
| PSMG1 | PSMG1\_HUMAN | Isoform 2 of Proteasome assembly chaperone 1 | AGT[79.9663]EDEEEEEEGRR | T | 18.0 | 17603.059 | 17603.059 | 0.0 | 0.0 |
| EPB41 | EPB41\_HUMAN | Isoform 5 of Protein 4.1 | KLSM[15.9949]Y[79.9663]GVDLHK | Y | 396.0 | 17581.262 | 0.0 | 17581.262 | 0.0 |
| KRT9 | K1C9\_HUMAN | Keratin, type I cytoskeletal 9 | FSSSSGY[79.9663]GGGSSR | Y | 53.0 | 17479.883 | 0.0 | 0.0 | 17479.883 |
| CYBRD1 | CYBR1\_HUMAN | Plasma membrane ascorbate-dependent reductase CYBRD1 | SDS[79.9663]ELNSEVAAR | S | 262.0 | 17417.39 | 0.0 | 0.0 | 17417.39 |
| ZDHHC5 | ZDHC5\_HUMAN | Isoform 2 of Palmitoyltransferase ZDHHC5 | S[79.9663]LGSASPGPGQPPLSSPTR | S | 679.0 | 17383.035 | 0.0 | 0.0 | 17383.035 |
| YBX3 | YBOX3\_HUMAN | Y-box-binding protein 3 | AGEAPTENPAPPTQQS[79.9663]SAE | S | 369.0 | 17319.537 | 17319.537 | 0.0 | 0.0 |
| GNG5 | GBG5\_HUMAN | Guanine nucleotide-binding protein G(I)/G(S)/G(O) subunit gamma-5 | n[42.0106]SGS[79.9663]SSVAAM[15.9949]KK | S |  | 17013.947 | 0.0 | 0.0 | 17013.947 |
| TNS1 | TENS1\_HUMAN | Tensin-1 | AASDGQYENQSPEAT[79.9663]SPRSPGVR | T | 1036.0 | 16985.934 | 16985.934 | 0.0 | 0.0 |
| ARHGAP23 | RHG23\_HUMAN | Rho GTPase-activating protein 23 | S[79.9663]ASQDRLEEVAAPR | S | 314.0 | 16946.475 | 0.0 | 16946.475 | 0.0 |
| AAK1 | AAK1\_HUMAN | AP2-associated protein kinase 1 | AGQTQPNPGILPIQPALT[79.9663]PR | T | 389.0 | 16794.912 | 16794.912 | 0.0 | 0.0 |
| KRT1 | K2C1\_HUMAN | Keratin, type II cytoskeletal 1 | AQY[79.9663]EDIAQK | Y | 358.0 | 16635.992 | 0.0 | 0.0 | 16635.992 |
| ACTB | ACTB\_HUMAN | Actin, cytoplasmic 1 | DLYANTVLSGGTTMY[79.9663]PGIADR | Y | 306.0 | 16622.422 | 0.0 | 16622.422 | 0.0 |
| SPTA1 | SPTA1\_HUMAN | Spectrin alpha chain, erythrocytic 1 | ETGTLES[79.9663]QLEANKR | S | 2194.0 | 16515.637 | 0.0 | 16515.637 | 0.0 |
| EPB41 | EPB41\_HUMAN | Isoform 5 of Protein 4.1 | KLS[79.9663]M[15.9949]YGVDLHK | S |  | 16494.168 | 0.0 | 16494.168 | 0.0 |
| SLC4A1 | B3AT\_HUMAN | Band 3 anion transport protein | S[79.9663]VTHANALTVM[15.9949]GK | S |  | 16442.84 | 0.0 | 0.0 | 16442.84 |
| CD44 | CD44\_HUMAN | Isoform 10 of CD44 antigen | ES[79.9663]SETPDQFM[15.9949]TADETR | S |  | 16392.646 | 0.0 | 0.0 | 16392.646 |
| PRPSAP1 | KPRA\_HUMAN | Isoform 2 of Phosphoribosyl pyrophosphate synthase-associated protein 1 | AGLT[79.9663]HIITM[15.9949]DLHQKEIQGFFSFPVDNLR | T |  | 16323.291 | 16323.291 | 0.0 | 0.0 |
| ACTB | ACTB\_HUMAN | Actin, cytoplasmic 1 | DSYVGDEAQS[79.9663]KR | S | 60.0 | 16214.265 | 0.0 | 16214.265 | 0.0 |
| ANK1 | ANK1\_HUMAN | Isoform Br21 of Ankyrin-1 | NGASPNEVS[79.9663]SDGTTPLAIAK | S | 764.0 | 16210.32 | 0.0 | 16210.32 | 0.0 |
| MAP4K4 | M4K4\_HUMAN | Isoform 6 of Mitogen-activated protein kinase kinase kinase kinase 4 | AAS[79.9663]SLNLSNGETESVK | S | 800.0 | 16168.198 | 16168.198 | 0.0 | 0.0 |
| RABEP1 | RABE1\_HUMAN | Isoform 2 of Rab GTPase-binding effector protein 1 | AQSTDS[79.9663]LGTSGSLQSK | S | 410.0 | 16166.094 | 16166.094 | 0.0 | 0.0 |
| STX7 | STX7\_HUMAN | Isoform 2 of Syntaxin-7 | EFGS[79.9663]LPTTPSEQR | S | 75.0 | 15937.097 | 0.0 | 0.0 | 15937.097 |
| RPS3 | RS3\_HUMAN | Isoform 2 of Small ribosomal subunit protein uS3 | DEILPT[79.9663]TPISEQK | T | 220.0 | 15933.131 | 0.0 | 15933.131 | 0.0 |
| ITSN1 | ITSN1\_HUMAN | Isoform 10 of Intersectin-1 | SAFTPATATGS[79.9663]SPSPVLGQGEK | S | 901.0 | 15916.252 | 0.0 | 15916.252 | 0.0 |
| GAPVD1 | GAPD1\_HUMAN | Isoform 2 of GTPase-activating protein and VPS9 domain-containing protein 1 | S[79.9663]SDIVSSVR | S | 902.0 | 15701.593 | 0.0 | 0.0 | 15701.593 |
| PIEZO1 | PIEZ1\_HUMAN | Piezo-type mechanosensitive ion channel component 1 | DPGLEPGPDSPGGS[79.9663]SPPR | S | 1395.0 | 15598.6875 | 0.0 | 0.0 | 15598.6875 |
| DMTN | DEMA\_HUMAN | Isoform 4 of Dematin | RGAEEEEEEEDDDS[79.9663]GEEMK | S | 226.0 | 15514.3097 | 0.0 | 15514.3097 | 0.0 |
| CTPS1 | PYRG1\_HUMAN | CTP synthase 1 | SGSSS[79.9663]PDSEITELK | S | 575.0 | 15343.927 | 0.0 | 0.0 | 15343.927 |
| PIEZO1 | PIEZ1\_HUMAN | Piezo-type mechanosensitive ion channel component 1 | SGSEEAVT[79.9663]DPGER | T | 1626.0 | 15249.543 | 0.0 | 0.0 | 15249.543 |
| KRT2 | K22E\_HUMAN | Keratin, type II cytoskeletal 2 epidermal | GFS[79.9663]SGSAVVSGGSR | S | 23.0 | 15086.0303 | 0.0 | 6927.9717 | 8158.0586 |
| CTPS1 | PYRG1\_HUMAN | CTP synthase 1 | S[79.9663]GSSSPDSEITELK | S | 571.0 | 14938.559 | 0.0 | 0.0 | 14938.559 |
| KCNN4 | KCNN4\_HUMAN | Intermediate conductance calcium-activated potassium channel protein 4 | EQVNS[79.9663]M[15.9949]VDISK | S |  | 14744.428 | 0.0 | 0.0 | 14744.428 |
| RTN4 | RTN4\_HUMAN | Reticulon-4 | TGVVFGASLFLLLS[79.9663]LTVFSIVSVTAYIALALLSVTIS[79.9663]FR | S |  | 14629.862 | 0.0 | 14629.862 | 0.0 |
| C2orf88 | SMAKA\_HUMAN | Small membrane A-kinase anchor protein | M[15.9949]AS[79.9663]PVNVKEEVK | S | 40.0 | 14608.293 | 0.0 | 14608.293 | 0.0 |
| CSNK1A1 | KC1A\_HUMAN | Isoform 3 of Casein kinase I isoform alpha | AAQQAASSSGQGQQAQT[79.9663]PTGF | T |  | 14598.561 | 14598.561 | 0.0 | 0.0 |
| KRT10 | K1C10\_HUMAN | Keratin, type I cytoskeletal 10 | SQY[79.9663]EQLAEQNR | Y | 325.0 | 14366.875 | 0.0 | 9238.817 | 5128.058 |
| CLCN5 | CLCN5\_HUMAN | H(+)/Cl(-) exchange transporter 5 | S[79.9663]YNGGGIGSSNR | S | 58.0 | 14314.143 | 0.0 | 0.0 | 14314.143 |
| CA2 | CAH2\_HUMAN | Carbonic anhydrase 2 | QS[79.9663]PVDIDTHTAK | S | 29.0 | 14306.531 | 0.0 | 14306.531 | 0.0 |
| HBA1 | HBA\_HUMAN | Hemoglobin subunit alpha | LLS[79.9663]HC[57.0215]LLVTLAAHLPAEFTPAVHASLDK | S |  | 14281.985 | 0.0 | 14281.985 | 0.0 |
| SLC2A4 | GLUT4\_HUMAN | Solute carrier family 2, facilitated glucose transporter member 4 | TPS[79.9663]LLEQEVK | S | 488.0 | 14174.658 | 0.0 | 0.0 | 14174.658 |
| TMEM63B | CSCL2\_HUMAN | CSC1-like protein 2 | LTSVS[79.9663]SSVDFDQR | S | 113.0 | 14164.016 | 0.0 | 0.0 | 14164.016 |
| DMTN | DEMA\_HUMAN | Isoform 4 of Dematin | GAEEEEEEEDDDS[79.9663]GEEMK | S | 226.0 | 14120.888 | 0.0 | 14120.888 | 0.0 |
| TNS1 | TENS1\_HUMAN | Tensin-1 | QGS[79.9663]PTPALPEKR | S | 1485.0 | 14112.258 | 0.0 | 14112.258 | 0.0 |
| GPS1 | CSN1\_HUMAN | Isoform 4 of COP9 signalosome complex subunit 1 | EGSQGELT[79.9663]PANSQSR | T | 479.0 | 13906.868 | 0.0 | 13906.868 | 0.0 |
| TNS1 | TENS1\_HUMAN | Tensin-1 | AASDGQYENQS[79.9663]PEATSPR | S | 1032.0 | 13799.512 | 13799.512 | 0.0 | 0.0 |
| SPTA1 | SPTA1\_HUMAN | Spectrin alpha chain, erythrocytic 1 | FT[79.9663]M[15.9949]GHSAHEETK | T |  | 13793.099 | 0.0 | 13793.099 | 0.0 |
| KRT6A | K2C6A\_HUMAN | Keratin, type II cytoskeletal 6A | AIGGGLSS[79.9663]VGGGSSTIK | S | 541.0 | 13399.236 | 0.0 | 0.0 | 13399.236 |
| KRT4 | K2C4\_HUMAN | Keratin, type II cytoskeletal 4 | S[79.9663]ISM[15.9949]SVAGSR | S |  | 13265.652 | 0.0 | 0.0 | 13265.652 |
| KRT14 | K1C14\_HUMAN | Keratin, type I cytoskeletal 14 | APST[79.9663]YGGGLSVSSSR | T | 45.0 | 13257.113 | 0.0 | 13257.113 | 0.0 |
| OXSR1 | OXSR1\_HUMAN | Serine/threonine-protein kinase OSR1 | TAQALSSGS[79.9663]GSQETK | S | 425.0 | 13040.043 | 0.0 | 13040.043 | 0.0 |
| FLCN | FLCN\_HUMAN | Isoform 2 of Folliculin | AHS[79.9663]PAEGASVESSSPGPK | S | 62.0 | 12970.675 | 12970.675 | 0.0 | 0.0 |
| AGER | RAGE\_HUMAN | Isoform 6 of Advanced glycosylation end product-specific receptor | AELNQSEEPEAGES[79.9663]STGGP | S | 399.0 | 12944.342 | 0.0 | 12944.342 | 0.0 |
| STOM | STOM\_HUMAN | Stomatin | VQNATLAVANIT[79.9663]NADSATR | T | 137.0 | 12759.211 | 0.0 | 0.0 | 12759.211 |
| HSP90AB1 | HS90B\_HUMAN | Heat shock protein HSP 90-beta | EKEIS[79.9663]DDEAEEEKGEK | S | 226.0 | 12738.033 | 0.0 | 12738.033 | 0.0 |
| ADD2 | ADDB\_HUMAN | Beta-adducin | GLSQM[15.9949]T[79.9663]TSADTDVDTSKDK | T | 670.0 | 12715.05 | 0.0 | 0.0 | 12715.05 |
| CANX | CALX\_HUMAN | Isoform 2 of Calnexin | QKS[79.9663]DAEEDGGTVSQEEEDR | S | 554.0 | 12644.244 | 0.0 | 0.0 | 12644.244 |
| ZNF185 | ZN185\_HUMAN | Isoform 3 of Zinc finger protein 185 | S[79.9663]TGSPTQETQAPFIAK | S |  | 12448.374 | 0.0 | 12448.374 | 0.0 |
| TMX1 | TMX1\_HUMAN | Thioredoxin-related transmembrane protein 1 | KVEEEQEADEEDVS[79.9663]EEEAESK | S | 247.0 | 12402.158 | 0.0 | 12402.158 | 0.0 |
| DENND1A | DEN1A\_HUMAN | DENN domain-containing protein 1A | TLRES[79.9663]DSAEGDEAESPEQQVR | S | 536.0 | 12367.331 | 0.0 | 12367.331 | 0.0 |
| KEL | KELL\_HUMAN | Kell blood group glycoprotein | S[79.9663]QAGGM[15.9949]GTLWSQESTPEER | S |  | 12340.848 | 0.0 | 12340.848 | 0.0 |
| KRT13 | K1C13\_HUMAN | Keratin, type I cytoskeletal 13 | EVSTNT[79.9663]AMIQTSK | T | 311.0 | 12301.186 | 0.0 | 12301.186 | 0.0 |
| EPB41 | EPB41\_HUMAN | Isoform 5 of Protein 4.1 | KEDEPPEQAEPEPT[79.9663]EAWKK | T |  | 12247.6045 | 0.0 | 12247.6045 | 0.0 |
| ITSN1 | ITSN1\_HUMAN | Isoform 10 of Intersectin-1 | STSM[15.9949]DSGSSES[79.9663]PASLK | S | 986.0 | 12210.791 | 0.0 | 12210.791 | 0.0 |
| TNS1 | TENS1\_HUMAN | Tensin-1 | SGSLGQPS[79.9663]PSAQR | S | 1228.0 | 12209.238 | 0.0 | 0.0 | 12209.238 |
| SPTB | SPTB1\_HUMAN | Spectrin beta chain, erythrocytic | HEAIETDTAAY[79.9663]EER | Y | 474.0 | 12168.19 | 0.0 | 12168.19 | 0.0 |
| HBB | HBB\_HUMAN | Hemoglobin subunit beta | LLVVYPWT[79.9663]QR | T | 39.0 | 12140.693 | 0.0 | 0.0 | 12140.693 |
| SPTB | SPTB1\_HUMAN | Spectrin beta chain, erythrocytic | ES[79.9663]QQLMDSHPEQK | S | 1018.0 | 12087.286 | 0.0 | 12087.286 | 0.0 |
| SPTB | SPTB1\_HUMAN | Spectrin beta chain, erythrocytic | ELY[79.9663]QQVVAQADLR | Y | 830.0 | 11953.998 | 0.0 | 11953.998 | 0.0 |
| PI4K2A | P4K2A\_HUMAN | Phosphatidylinositol 4-kinase type 2-alpha | SSS[79.9663]ESYTQSFQSR | S | 462.0 | 11820.8 | 0.0 | 11820.8 | 0.0 |
| C2orf88 | SMAKA\_HUMAN | Small membrane A-kinase anchor protein | QHES[79.9663]EEPFM[15.9949]PEER | S |  | 11753.588 | 0.0 | 11753.588 | 0.0 |
| HBB | HBB\_HUMAN | Hemoglobin subunit beta | VLGAFS[79.9663]DGLAHLDNLK | S | 73.0 | 11637.781 | 0.0 | 11637.781 | 0.0 |
| EPB41 | EPB41\_HUMAN | Isoform 5 of Protein 4.1 | LSM[15.9949]Y[79.9663]GVDLHK | Y | 396.0 | 11554.066 | 0.0 | 11554.066 | 0.0 |
| SPTA1 | SPTA1\_HUMAN | Spectrin alpha chain, erythrocytic 1 | GY[79.9663]VSLEDYTAFLIDK | Y | 2333.0 | 11535.143 | 0.0 | 11535.143 | 0.0 |
| RETREG2 | RETR2\_HUMAN | Reticulophagy regulator 2 | QALDS[79.9663]EEEEEDVAAK | S | 385.0 | 11361.458 | 0.0 | 0.0 | 11361.458 |
| RALBP1 | RBP1\_HUMAN | RalA-binding protein 1 | AGKEPAKPS[79.9663]PSR | S | 645.0 | 11299.227 | 11299.227 | 0.0 | 0.0 |
| UBR4 | UBR4\_HUMAN | Isoform 2 of E3 ubiquitin-protein ligase UBR4 | ELASPVS[79.9663]PELR | S | 181.0 | 11166.161 | 0.0 | 0.0 | 11166.161 |
| SPTA1 | SPTA1\_HUMAN | Spectrin alpha chain, erythrocytic 1 | GDC[57.0215]GDT[79.9663]LAATQSLLM[15.9949]K | T |  | 10975.114 | 0.0 | 10975.114 | 0.0 |
| TTN | TITIN\_HUMAN | Isoform 11 of Titin | QEQIQVT[79.9663]HGK |  |  | 10968.281 | 0.0 | 10968.281 | 0.0 |
| SPTA1 | SPTA1\_HUMAN |  | DLVAS[79.9663]EGLFHSHK | S | 299.0 | 10898.268 | 0.0 | 10898.268 | 0.0 |
| KRT2 | K22E\_HUMAN |  | DY[79.9663]QELM[15.9949]NVK | Y |  | 10836.935 | 0.0 | 0.0 | 10836.935 |
| ACTB | ACTB\_HUMAN |  | Q[-17.0265]EYDES[79.9663]GPSIVHR | S | 365.0 | 10721.951 | 0.0 | 10721.951 | 0.0 |
| PI4K2A | P4K2A\_HUMAN |  | VAAAAGSGPSPPGS[79.9663]PGHDR | S | 51.0 | 10451.304 | 0.0 | 0.0 | 10451.304 |
| ZDHHC2 | ZDHC2\_HUMAN |  | AGM[15.9949]S[79.9663]NPALTM[15.9949]ENET | S |  | 10318.062 | 0.0 | 0.0 | 10318.062 |
| EHBP1L1 | EH1L1\_HUMAN |  | AS[79.9663]SPEKAEEDRR | S | 1016.0 | 10290.576 | 0.0 | 10290.576 | 0.0 |
| CAT | CATA\_HUMAN |  | FS[79.9663]TVAGESGSADTVR | S | 114.0 | 10205.688 | 0.0 | 0.0 | 10205.688 |
| MINK1 | MINK1\_HUMAN |  | KGS[79.9663]VVNVNPTNTR | S | 993.0 | 10195.631 | 0.0 | 10195.631 | 0.0 |
| KRT1 | K2C1\_HUMAN |  | SGGGFS[79.9663]SGSAGIINYQR | S | 18.0 | 10087.029 | 0.0 | 0.0 | 10087.029 |
| ADD2 | ADDB\_HUMAN |  | T[79.9663]ESVT[79.9663]SGPM[15.9949]SPEGSPSK | T |  | 10074.091 | 0.0 | 10074.091 | 0.0 |
| ABCB6 | ABCB6\_HUMAN |  | APGIILLDEAT[79.9663]SALDTSNER | T | 754.0 | 9938.149 | 0.0 | 9938.149 | 0.0 |
| ZDHHC5 | ZDHC5\_HUMAN |  | SLGSASPGPGQPPLS[79.9663]SPTR | S | 693.0 | 9827.821 | 0.0 | 0.0 | 9827.821 |
| EPB41 | EPB41\_HUMAN |  | KLS[79.9663]MYGVDLHK | S | 394.0 | 9625.344 | 0.0 | 9625.344 | 0.0 |
| EPB41 | EPB41\_HUMAN |  | SM[15.9949]T[79.9663]PAQADLEFLENAK | T | 378.0 | 9617.631 | 0.0 | 9617.631 | 0.0 |
| HBD | HBD\_HUMAN |  | EFTPQM[15.9949]QAAY[79.9663]QK | Y | 131.0 | 9581.226 | 0.0 | 0.0 | 9581.226 |
| DMTN | DEMA\_HUMAN |  | VT[79.9663]SNLGKM[15.9949]ILK | T |  | 9429.336 | 0.0 | 9429.336 | 0.0 |
| DENND1A | DEN1A\_HUMAN |  | ES[79.9663]DSAEGDEAESPEQQVR | S | 536.0 | 9400.539 | 0.0 | 9400.539 | 0.0 |
| SKP1 | SKP1\_HUMAN |  | NDFT[79.9663]EEEEAQVR | T | 146.0 | 9387.485 | 0.0 | 9387.485 | 0.0 |
| GYPC | GLPC\_HUMAN |  | GT[79.9663]EFAESADAALQGDPALQDAGDSSR | T | 99.0 | 9346.153 | 0.0 | 9346.153 | 0.0 |
| STX7 | STX7\_HUMAN |  | TLNQLGTPQDS[79.9663]PELR | S | 45.0 | 9341.342 | 0.0 | 0.0 | 9341.342 |
| DMTN | DEMA\_HUMAN |  | TRS[79.9663]LPDR | S | 269.0 | 9289.287 | 0.0 | 0.0 | 9289.287 |
| LMNA | LMNA\_HUMAN |  | SGAQASSTPLS[79.9663]PTR | S | 22.0 | 9165.875 | 0.0 | 9165.875 | 0.0 |
| ABCC1 | MRP1\_HUMAN |  | QLSSSSSYS[79.9663]GDISR | S | 921.0 | 9046.215 | 0.0 | 0.0 | 9046.215 |
| HBA1 | HBA\_HUMAN |  | M[15.9949]FLSFPT[79.9663]TK | T | 39.0 | 8972.119 | 0.0 | 8972.119 | 0.0 |
| STX4 | STX4\_HUMAN |  | AIEPQKEEADENYNS[79.9663]VNTR | S | 117.0 | 8920.01 | 0.0 | 0.0 | 8920.01 |
| EFNB1 | EFNB1\_HUMAN |  | AAALS[79.9663]LSTLASPK | S | 281.0 | 8790.131 | 0.0 | 8790.131 | 0.0 |
| SGTA | SGTA\_HUMAN |  | S[79.9663]RTPSASNDDQQE | S | 301.0 | 8781.816 | 0.0 | 8781.816 | 0.0 |
| KRT13 | K1C13\_HUMAN |  | EVST[79.9663]NTAM[15.9949]IQTSK | T |  | 8778.927 | 0.0 | 0.0 | 8778.927 |
| ANK1 | ANK1\_HUMAN |  | KADAATS[79.9663]FLR | S | 15.0 | 8742.867 | 0.0 | 8742.867 | 0.0 |
| SLC2A1 | GTR1\_HUMAN |  | S[79.9663]FEM[15.9949]LILGR | S |  | 8680.132 | 0.0 | 8680.132 | 0.0 |
| EPB42 | EPB42\_HUMAN |  | Y[79.9663]PEGSLQEK | Y | 437.0 | 8579.043 | 0.0 | 0.0 | 8579.043 |
| ATP7A | ATP7A\_HUMAN |  | VSITSEVESTSNSPS[79.9663]SSSLQK | S | 359.0 | 8454.02 | 0.0 | 0.0 | 8454.02 |
| STOM | STOM\_HUMAN |  | VQNAT[79.9663]LAVANITNADSATR | T | 130.0 | 8121.2974 | 0.0 | 8121.2974 | 0.0 |
| ATP1A1 | AT1A1\_HUMAN |  | VDNSSLT[79.9663]GESEPQTR | T | 219.0 | 8111.884 | 0.0 | 0.0 | 8111.884 |
| ACTB | ACTB\_HUMAN |  | DS[79.9663]YVGDEAQSK | S | 52.0 | 8106.3467 | 0.0 | 8106.3467 | 0.0 |
| STOM | STOM\_HUMAN |  | EASM[15.9949]VITES[79.9663]PAALQLR | S | 244.0 | 8055.1265 | 0.0 | 0.0 | 8055.1265 |
| VAPB | VAPB\_HUMAN |  | T[79.9663]VQSNSPISALAPTGK | T | 201.0 | 8011.1445 | 0.0 | 8011.1445 | 0.0 |
| KEL | KELL\_HUMAN |  | SQAGGMGT[79.9663]LWSQESTPEER | T | 22.0 | 7917.831 | 0.0 | 7917.831 | 0.0 |
| KRT1 | K2C1\_HUMAN |  | Q[-17.0265]IS[79.9663]NLQQSISDAEQR | S | 420.0 | 7868.3857 | 0.0 | 0.0 | 7868.3857 |
| VPS4B | VPS4B\_HUMAN |  | LLEPVVS[79.9663]MSDM[15.9949]LR | S |  | 7832.5264 | 0.0 | 0.0 | 7832.5264 |
| PGK2 | PGK2\_HUMAN |  | AC[57.0215]ANPAPGS[79.9663]VILLENLR | S | 115.0 | 7733.62 | 7733.62 | 0.0 | 0.0 |
| ANK1 | ANK1\_HUMAN |  | LC[57.0215]QDY[79.9663]DTIGPEGGSLK | Y | 1073.0 | 7522.7626 | 0.0 | 7522.7626 | 0.0 |
| SPTA1 | SPTA1\_HUMAN |  | VVEVNQY[79.9663]ANEC[57.0215]AEENHPDLPLIQSK | Y |  | 7349.793 | 0.0 | 7349.793 | 0.0 |
| SLC43A1 | LAT3\_HUMAN |  | APSLEDGS[79.9663]DAFM[15.9949]SPQDVR | S |  | 7246.3667 | 0.0 | 0.0 | 7246.3667 |
| UBR4 | UBR4\_HUMAN |  | SNT[79.9663]PM[15.9949]GDKDDDDDDDADEK | T |  | 7008.3477 | 0.0 | 0.0 | 7008.3477 |
| SMIM1 | SMIM1\_HUMAN |  | DGVSLGAVS[79.9663]ST[79.9663]EEASR | S |  | 6919.9116 | 0.0 | 0.0 | 6919.9116 |
| DMTN | DEMA\_HUMAN |  | LQS[79.9663]TEFSPSGSETGSPGLQNGEGQR | S | 303.0 | 6734.373 | 0.0 | 6734.373 | 0.0 |
| KLC4 | KLC4\_HUMAN |  | AAS[79.9663]LNYLNQPSAAPLQVSR | S | 590.0 | 6711.754 | 6711.754 | 0.0 | 0.0 |
| EPB41 | EPB41\_HUMAN |  | S[79.9663]MT[79.9663]PAQADLEFLENAK | S |  | 6678.2188 | 0.0 | 6678.2188 | 0.0 |
| HBA1 | HBA\_HUMAN |  | n[42.0106]VLS[79.9663]PADKTNVK | S | 4.0 | 6640.5327 | 0.0 | 0.0 | 6640.5327 |
| PPP1R7 | PP1R7\_HUMAN |  | RVES[79.9663]EES[79.9663]GDEEGKK | S |  | 6449.858 | 0.0 | 6449.858 | 0.0 |
| HBA1 | HBA\_HUMAN |  | LLSHC[57.0215]LLVT[79.9663]LAAHLPAEFTPAVHASLDKFLASVSTVLTSK | T | 109.0 | 6033.9175 | 0.0 | 6033.9175 | 0.0 |
| TNS1 | TENS1\_HUMAN |  | EAFEEM[15.9949]EGTSPS[79.9663]SPPPSGVR | S | 1087.0 | 6028.6685 | 0.0 | 0.0 | 6028.6685 |
| CAT | CATA\_HUMAN |  | FNT[79.9663]ANDDNVTQVR | T | 434.0 | 6022.8096 | 0.0 | 0.0 | 6022.8096 |
| ABCB6 | ABCB6\_HUMAN |  | APGIILLDEATS[79.9663]ALDTSNER | S | 755.0 | 5848.844 | 0.0 | 5848.844 | 0.0 |
| SPTB | SPTB1\_HUMAN |  | GY[79.9663]QPC[57.0215]DPQVIQDR | Y |  | 5767.488 | 0.0 | 5767.488 | 0.0 |
| EPB42 | EPB42\_HUMAN |  | FQFT[79.9663]PTHVGLQR | T | 655.0 | 5659.7207 | 0.0 | 5659.7207 | 0.0 |
| CTTN | SRC8\_HUMAN |  | LPS[79.9663]SPVYEDAASFK | S | 417.0 | 5516.882 | 0.0 | 0.0 | 5516.882 |
| ABCA7 | ABCA7\_HUMAN |  | S[79.9663]LPLLGEEDEDVAR | S | 1765.0 | 5093.9053 | 0.0 | 0.0 | 5093.9053 |
| GPX1 | GPX1\_HUMAN |  | VLLIENVAS[79.9663]LUGTTVR | S | 47.0 | 5076.6 | 0.0 | 5076.6 | 0.0 |
| SPTA1 | SPTA1\_HUMAN |  | Q[-17.0265]EAFLENEDLGNSLGS[79.9663]AEALLQK | S | 510.0 | 4941.1177 | 0.0 | 4941.1177 | 0.0 |
| CAMP | CAMP\_HUMAN |  | WS[79.9663]LVLLLLGLVMPLAIIAQVLS[79.9663]Y[79.9663]KEAVLRAIDGINQR | S |  | 4766.8535 | 0.0 | 0.0 | 4766.8535 |
| ERMAP | ERMAP\_HUMAN |  | GNEYEALT[79.9663]SPQTSFR | T | 343.0 | 4495.1123 | 0.0 | 0.0 | 4495.1123 |
| DLG1 | DLG1\_HUMAN |  | EQM[15.9949]M[15.9949]NSSISSGSGS[79.9663]LR | S | 575.0 | 4486.9224 | 0.0 | 4486.9224 | 0.0 |
| ADD2 | ADDB\_HUMAN |  | GVS[79.9663]C[57.0215]SEVTASSLIK | S |  | 4414.4615 | 0.0 | 4414.4615 | 0.0 |
| ERMAP | ERMAP\_HUMAN |  | GNEYEALTS[79.9663]PQTSFR | S | 344.0 | 3927.3164 | 0.0 | 0.0 | 3927.3164 |
| HBA1 | HBA\_HUMAN |  | LLS[79.9663]HC[57.0215]LLVTLAAHLPAEFTPAVHASLDKFLASVSTVLTSK | S |  | 3667.7655 | 0.0 | 3351.1415 | 316.624 |
| RAB6A | RAB6A\_HUMAN |  | S[79.9663]TGGDFGNPLR | S | 2.0 | 3337.8066 | 0.0 | 3337.8066 | 0.0 |
| KRT9 | K1C9\_HUMAN |  | FSS[79.9663]SSGYGGGSSR | S | 49.0 | 3019.872 | 0.0 | 0.0 | 3019.872 |
| ADD2 | ADDB\_HUMAN |  | T[79.9663]ESVTS[79.9663]GPM[15.9949]SPEGSPSK | T |  | 2841.4177 | 0.0 | 2841.4177 | 0.0 |
| EPB41 | EPB41\_HUMAN |  | T[79.9663]QT[79.9663]VTISDNANAVK | T |  | 1275.9131 | 0.0 | 0.0 | 1275.9131 |
| KRT74 | K2C74\_HUMAN |  | LDS[79.9663]ELRS[79.9663]M[15.9949]RDLVEDYK | S |  | 0.0 | 0.0 | 0.0 | 0.0 |
| RTN3 | RTN3\_HUMAN |  | TGFVFGTTLIMLLS[79.9663]LAAFSVISVVS[79.9663]YLILALLSVTISFR | S |  | 0.0 | 0.0 | 0.0 | 0.0 |
| ADCY7 | ADCY7\_HUMAN |  | LT[79.9663]LAVLTIGSLLT[79.9663]VAIINLPLM[15.9949]PFQVPELPVGNETGLLAASSK | T |  | 0.0 | 0.0 | 0.0 | 0.0 |
| ACTBL2 | ACTBL\_HUMAN |  | MQKEIIT[79.9663]LAPSTM[15.9949]K | T |  | 0.0 | 0.0 | 0.0 | 0.0 |
| HBD | HBD\_HUMAN |  | VLGAFSDGLAHLDNLKGT[79.9663]FS[79.9663]QLSELHC[57.0215]DK | T |  | 0.0 | 0.0 | 0.0 | 0.0 |
| USP6NL | US6NL\_HUMAN |  | ALDAEDGKRGS[79.9663]TASQYDNVPGPELDSGAS[79.9663]VEEALER | S |  | 0.0 | 0.0 | 0.0 | 0.0 |
| UBA1 | UBA1\_HUMAN |  | DNPGVVT[79.9663]C[57.0215]LDEAR | T |  | 0.0 | 0.0 | 0.0 | 0.0 |
| FARSB | SYFB\_HUMAN |  | T[79.9663]YTIANQFPLNKLT[79.9663]ELLRHDMAAAGFT[79.9663]EALTFALC[57.0215]SQEDIADK | T |  | 0.0 | 0.0 | 0.0 | 0.0 |
| SPTB | SPTB1\_HUMAN |  | EQIY[79.9663]S[79.9663]S[79.9663]LDYGKDLTSVLILQR | Y |  | 0.0 | 0.0 | 0.0 | 0.0 |
| KRT6B | K2C6B\_HUMAN |  | EY[79.9663]QELM[15.9949]NVK | Y |  | 0.0 | 0.0 | 0.0 | 0.0 |
| KRT10 | K1C10\_HUMAN |  | AET[79.9663]EC[57.0215]QNTEYQQLLDIK | T |  | 0.0 | 0.0 | 0.0 | 0.0 |
| CAST | ICAL\_HUMAN |  | AAAPAPVSEAVC[57.0215]RT[79.9663]SM[15.9949]C[57.0215]SIQSAPPEPATLK | T |  | 0.0 | 0.0 | 0.0 | 0.0 |
| GYPA | GLPA\_HUMAN |  | DTYAATPRAHEVS[79.9663]EIS[79.9663]VRTVYPPEEETGER | S |  | 0.0 | 0.0 | 0.0 | 0.0 |
| GYPA | GLPA\_HUMAN |  | KSPSDVKPLPS[79.9663]PDT[79.9663]DVPLSSVEIENPETSDQ | S |  | 0.0 | 0.0 | 0.0 | 0.0 |
| GYPA | GLPA\_HUMAN |  | KS[79.9663]PS[79.9663]DVKPLPSPDTDVPLSSVEIENPETSDQ | S |  | 0.0 | 0.0 | 0.0 | 0.0 |
| GYPA | GLPA\_HUMAN |  | SPSDVKPLPS[79.9663]PDT[79.9663]DVPLSSVEIENPETSDQ | S |  | 0.0 | 0.0 | 0.0 | 0.0 |
| FKBP4 | FKBP4\_HUMAN |  | AEASSGDHPTDT[79.9663]EM[15.9949]KEEQK | T |  | 0.0 | 0.0 | 0.0 | 0.0 |
| EPB41 | EPB41\_HUMAN |  | VSLLDDTVY[79.9663]EC[57.0215]VVEK | Y |  | 0.0 | 0.0 | 0.0 | 0.0 |
| DMTN | DEMA\_HUMAN |  | Q[-17.0265]PLT[79.9663]S[79.9663]PGSVSPSR | T |  | 0.0 | 0.0 | 0.0 | 0.0 |
| CYBRD1 | CYBR1\_HUMAN |  | GSM[15.9949]PAYS[79.9663]GNNM[15.9949]DKSDSELNSEVAAR | S |  | 0.0 | 0.0 | 0.0 | 0.0 |
| CD44 | CD44\_HUMAN |  | KPSGLNGEASKS[79.9663]QEM[15.9949]VHLVNK | S |  | 0.0 | 0.0 | 0.0 | 0.0 |
| CD44 | CD44\_HUMAN |  | KPSGLNGEAS[79.9663]KSQEM[15.9949]VHLVNK | S |  | 0.0 | 0.0 | 0.0 | 0.0 |
| CAT | CATA\_HUMAN |  | AAQKADVLT[79.9663]T[79.9663]GAGNPVGDKLNVITVGPR | T |  | 0.0 | 0.0 | 0.0 | 0.0 |
| SLC4A1 | B3AT\_HUMAN |  | SVT[79.9663]HANALTVM[15.9949]GK | T |  | 0.0 | 0.0 | 0.0 | 0.0 |
| ADD2 | ADDB\_HUMAN |  | SPS[79.9663]T[79.9663]ESQLM[15.9949]SK | S |  | 0.0 | 0.0 | 0.0 | 0.0 |
| ACTB | ACTB\_HUMAN |  | DLYANTVLSGGTT[79.9663]M[15.9949]YPGIADR | T |  | 0.0 | 0.0 | 0.0 | 0.0 |
| ACTB | ACTB\_HUMAN |  | DLYANTVLSGGT[79.9663]TM[15.9949]YPGIADR | T |  | 0.0 | 0.0 | 0.0 | 0.0 |
| ACTB | ACTB\_HUMAN |  | DLYANTVLS[79.9663]GGTTM[15.9949]YPGIADR | S |  | 0.0 | 0.0 | 0.0 | 0.0 |
| TXN | THIO\_HUMAN |  | QIES[79.9663]KTAFQEALDAAGDK | S | 7.0 | 0.0 | 0.0 | 0.0 | 0.0 |
| ACTBL2 | ACTBL\_HUMAN |  | HQGVM[15.9949]VGMGQKDC[57.0215]Y[79.9663]VGDEAQSK | Y | 54.0 | 0.0 | 0.0 | 0.0 | 0.0 |
| EPB41 | EPB41\_HUMAN |  | LKAS[79.9663]NGDTPTHEDLTK | S | 56.0 | 0.0 | 0.0 | 0.0 | 0.0 |
| EPB41 | EPB41\_HUMAN |  | ASNGDT[79.9663]PTHEDLTK | T | 60.0 | 0.0 | 0.0 | 0.0 | 0.0 |
| SMIM5 | SMIM5\_HUMAN |  | KVQVQPT[79.9663]PP | T | 75.0 | 0.0 | 0.0 | 0.0 | 0.0 |
| EPB42 | EPB42\_HUMAN |  | INRT[79.9663]QATFPISSLGDR | T | 75.0 | 0.0 | 0.0 | 0.0 | 0.0 |
| EPB42 | EPB42\_HUMAN |  | TQAT[79.9663]FPISSLGDR | T | 78.0 | 0.0 | 0.0 | 0.0 | 0.0 |
| ADD2 | ADDB\_HUMAN |  | GNNS[79.9663]SNIWALR | S | 81.0 | 0.0 | 0.0 | 0.0 | 0.0 |
| LIN7A | LIN7A\_HUMAN |  | ARAT[79.9663]AKATVAAFAASEGHSHPR | T | 89.0 | 0.0 | 0.0 | 0.0 | 0.0 |
| HBA1 | HBA\_HUMAN |  | LLSHC[57.0215]LLVT[79.9663]LAAHLPAEFTPAVHASLDK | T | 109.0 | 0.0 | 0.0 | 0.0 | 0.0 |
| TXNL1 | TXNL1\_HUMAN |  | QHLENDPGS[79.9663]NEDTDIPK | S | 113.0 | 0.0 | 0.0 | 0.0 | 0.0 |
| TXNL1 | TXNL1\_HUMAN |  | Q[-17.0265]HLENDPGS[79.9663]NEDTDIPK | S | 113.0 | 0.0 | 0.0 | 0.0 | 0.0 |
| HBA1 | HBA\_HUMAN |  | LLSHC[57.0215]LLVTLAAHLPAEFT[79.9663]PAVHASLDKFLASVSTVLTSK | T | 119.0 | 0.0 | 0.0 | 0.0 | 0.0 |
| HTT | HD\_HUMAN |  | ALM[15.9949]DS[79.9663]NLPR | S | 161.0 | 0.0 | 0.0 | 0.0 | 0.0 |
| TSC22D4 | T22D4\_HUMAN |  | SFT[79.9663]GGLGQLVVPSK | T | 167.0 | 0.0 | 0.0 | 0.0 | 0.0 |
| ALAD | HEM2\_HUMAN |  | AGC[57.0215]QVVAPS[79.9663]DMMDGRVEAIKEALMAHGLGNR | S | 168.0 | 0.0 | 0.0 | 0.0 | 0.0 |
| KRT10 | K1C10\_HUMAN |  | Y[79.9663]ENEVALR | Y | 238.0 | 0.0 | 0.0 | 0.0 | 0.0 |
| CYBRD1 | CYBR1\_HUMAN |  | GSM[15.9949]PAYSGNNM[15.9949]DKS[79.9663]DSELNSEVAAR | S | 260.0 | 0.0 | 0.0 | 0.0 | 0.0 |
| SLC29A1 | S29A1\_HUMAN |  | EESGVSVS[79.9663]NSQPTNESHSIK | S | 271.0 | 0.0 | 0.0 | 0.0 | 0.0 |
| DMTN | DEMA\_HUMAN |  | SS[79.9663]SLPAYGR | S | 288.0 | 0.0 | 0.0 | 0.0 | 0.0 |
| SPAG9 | JIP4\_HUMAN |  | AGPSAQEPGSQT[79.9663]PLK | T | 292.0 | 0.0 | 0.0 | 0.0 | 0.0 |
| ACTBL2 | ACTBL\_HUMAN |  | DLYANTVLSGGS[79.9663]TMYPGIADR | S | 304.0 | 0.0 | 0.0 | 0.0 | 0.0 |
| ACTB | ACTB\_HUMAN |  | DLYANTVLSGGTTM[15.9949]Y[79.9663]PGIADR | Y | 306.0 | 0.0 | 0.0 | 0.0 | 0.0 |
| EIF4A1 | IF4A1\_HUMAN |  | S[79.9663]GSSRVLITTDLLAR | S | 320.0 | 0.0 | 0.0 | 0.0 | 0.0 |
| KRT5 | K2C5\_HUMAN |  | Y[79.9663]EELQQTAGR | Y | 365.0 | 0.0 | 0.0 | 0.0 | 0.0 |
| KRT71 | K2C71\_HUMAN |  | QAS[79.9663]NLETAIADAEQR | S | 370.0 | 0.0 | 0.0 | 0.0 | 0.0 |
| KRT73 | K2C73\_HUMAN |  | QC[57.0215]ANLET[79.9663]AIADAEQR | T | 376.0 | 0.0 | 0.0 | 0.0 | 0.0 |
| KIF1B | KIF1B\_HUMAN |  | Y[79.9663]LLASENQR | Y | 419.0 | 0.0 | 0.0 | 0.0 | 0.0 |
| ADD1 | ADDA\_HUMAN |  | GDEASEEGQNGS[79.9663]SPK | S | 464.0 | 0.0 | 0.0 | 0.0 | 0.0 |
| ADD1 | ADDA\_HUMAN |  | GDEASEEGQNGSS[79.9663]PK | S | 465.0 | 0.0 | 0.0 | 0.0 | 0.0 |
| KRT3 | K2C3\_HUMAN |  | YGVSGGGFSSASNRGGS[79.9663]IK | S | 613.0 | 0.0 | 0.0 | 0.0 | 0.0 |
| ATG9A | ATG9A\_HUMAN |  | AHSTM[15.9949]TGS[79.9663]GVDAR | S | 675.0 | 0.0 | 0.0 | 0.0 | 0.0 |
| ADD2 | ADDB\_HUMAN |  | TES[79.9663]VTSGPMSPEGSPSK | S | 686.0 | 0.0 | 0.0 | 0.0 | 0.0 |
| BMP2K | BMP2K\_HUMAN |  | AEHSSINQENGTANPIKNGKTS[79.9663]PASK | S | 715.0 | 0.0 | 0.0 | 0.0 | 0.0 |
| EPB41 | EPB41\_HUMAN |  | TLNINGQIPT[79.9663]GEGPPLVK | T | 725.0 | 0.0 | 0.0 | 0.0 | 0.0 |
| SLC4A1 | B3AT\_HUMAN |  | S[79.9663]VTHANALTVMGKASTPGAAAQIQEVK | S | 731.0 | 0.0 | 0.0 | 0.0 | 0.0 |
| SLC4A1 | B3AT\_HUMAN |  | SVTHANALTVMGKAS[79.9663]TPGAAAQIQEVK | S | 745.0 | 0.0 | 0.0 | 0.0 | 0.0 |
| ANK1 | ANK1\_HUMAN |  | NGASPNEVSSDGTT[79.9663]PLAIAK | T | 769.0 | 0.0 | 0.0 | 0.0 | 0.0 |
| SPTB | SPTB1\_HUMAN |  | DS[79.9663]PDVTHRLQALRELYQQVVAQADLR | S | 816.0 | 0.0 | 0.0 | 0.0 | 0.0 |
| SPTA1 | SPTA1\_HUMAN |  | VKS[79.9663]LNQNMESLR | S | 876.0 | 0.0 | 0.0 | 0.0 | 0.0 |
| NUP210 | PO210\_HUMAN |  | AASPIIT[79.9663]LVALDEALDNYTITFLIRGVAIGQTSLTASVTNK | T | 1029.0 | 0.0 | 0.0 | 0.0 | 0.0 |
| ANK1 | ANK1\_HUMAN |  | QS[79.9663]RNLKPDR | S | 1490.0 | 0.0 | 0.0 | 0.0 | 0.0 |
| SPTA1 | SPTA1\_HUMAN |  | HQT[79.9663]FAHEVDGR | T | 1543.0 | 0.0 | 0.0 | 0.0 | 0.0 |
| ANK1 | ANK1\_HUMAN |  | QDDAT[79.9663]GAGQDSENEVSLVSGHQR | T | 1660.0 | 0.0 | 0.0 | 0.0 | 0.0 |
| PRRC2A | PRC2A\_HUMAN |  | LS[79.9663]SNLGGPGSSR | S | 2065.0 | 0.0 | 0.0 | 0.0 | 0.0 |
| SPTA1 | SPTA1\_HUMAN |  | HLS[79.9663]DIIEER | S | 2135.0 | 0.0 | 0.0 | 0.0 | 0.0 |
