## Supplementary Resource - Deep Red for "Deep Red Blood Cell Proteome Defines the Band 3 N-Terminus Interactome as a Regulator of Hypoxic Adaptation via BLVRB-Dependent *S*-Nitroso Transfer": RBC Proteome 2025.html

Interactive Data Table


| Protein | Protein ID | Entry Name Link | Gene | Protein Description | Length | MW | Copy number/cell | Total Intensity (merged) | Intensity GeLC MS | Intensity Cellular | Intensity Soluble | Intensity Insoluble | Coverage GeLC MS | Coverage Cellular | Coverage Soluble | Coverage Insoluble |
| --- | --- | --- | --- | --- | --- | --- | --- | --- | --- | --- | --- | --- | --- | --- | --- | --- |
| sp\|P69905\|HBA\_HUMAN | P69905 | HBA\_HUMAN | HBA1 | Hemoglobin subunit alpha | 142 | 15.258 | 1.342200e+08 | 2.572954e+09 | 273960770.0 | 1586239740.0 | 491191200.0 | 221562368.0 | 99.3 | 100.0 | 100.0 | 97.2 |
| sp\|P68871\|HBB\_HUMAN | P68871 | HBB\_HUMAN | HBB | Hemoglobin subunit beta | 147 | 15.998 | 1.329834e+08 | 2.672886e+09 | 236245020.0 | 1678448640.0 | 441395808.0 | 316796544.0 | 100.0 | 100.0 | 97.3 | 95.9 |
| sp\|P02042\|HBD\_HUMAN | P02042 | HBD\_HUMAN | HBD | Hemoglobin subunit delta | 147 | 16.055 | 8.854062e+07 | 1.785953e+09 | 205425070.0 | 993851520.0 | 318668416.0 | 268007968.0 | 100.0 | 100.0 | 95.9 | 95.9 |
| sp\|P04040\|CATA\_HUMAN | P04040 | CATA\_HUMAN | CAT | Catalase | 527 | 59.756 | 8.170915e+06 | 6.134361e+08 | 192083660.0 | 381453152.0 | 21421516.0 | 18477810.0 | 85.77 | 97.5 | 60.5 | 40.0 |
| sp\|P30043\|BLVRB\_HUMAN | P30043 | BLVRB\_HUMAN | BLVRB | Flavin reductase (NADPH) | 206 | 22.119 | 1.309983e+07 | 3.640392e+08 | 187298910.0 | 133605856.0 | 22126030.0 | 21008442.0 | 97.57 | 98.1 | 86.9 | 63.6 |
| sp\|P02730\|B3AT\_HUMAN | P02730 | B3AT\_HUMAN | SLC4A1 | Band 3 anion transport protein | 911 | 101.792 | 6.459721e+06 | 8.261227e+08 | 147633730.0 | 78345408.0 | 223065056.0 | 377078528.0 | 50.05 | 53.0 | 46.1 | 36.3 |
| sp\|P00915\|CAH1\_HUMAN | P00915 | CAH1\_HUMAN | CA1 | Carbonic anhydrase 1 | 261 | 28.870 | 2.092553e+07 | 7.589976e+08 | 145683780.0 | 503337216.0 | 58389548.0 | 51587028.0 | 81.23 | 97.3 | 78.5 | 59.0 |
| sp\|P32119\|PRDX2\_HUMAN | P32119 | PRDX2\_HUMAN | PRDX2 | Peroxiredoxin-2 | 198 | 21.892 | 1.636504e+07 | 4.501111e+08 | 140192380.0 | 185795776.0 | 68263568.0 | 55859344.0 | 100.0 | 97.5 | 77.8 | 52.0 |
| sp\|P02549\|SPTA1\_HUMAN | P02549 | SPTA1\_HUMAN | SPTA1 | Spectrin alpha chain, erythrocytic 1 | 2419 | 280.014 | 1.841715e+06 | 6.479171e+08 | 95438340.0 | 60812440.0 | 364767360.0 | 126898928.0 | 89.05 | 81.3 | 82.6 | 57.5 |
| sp\|P11277\|SPTB1\_HUMAN | P11277 | SPTB1\_HUMAN | SPTB | Spectrin beta chain, erythrocytic | 2137 | 246.468 | 2.036585e+06 | 6.306380e+08 | 88724930.0 | 47162640.0 | 338950176.0 | 155800256.0 | 89.14 | 81.8 | 84.3 | 64.9 |
| sp\|P11171\|EPB41\_HUMAN | P11171 | EPB41\_HUMAN | EPB41 | Protein 4.1 | 864 | 97.017 | 4.012858e+06 | 4.891238e+08 | 83983500.0 | 35831644.0 | 314710720.0 | 54597968.0 | 81.25 | 73.5 | 78.2 | 46.9 |
| sp\|P16157\|ANK1\_HUMAN | P16157 | ANK1\_HUMAN | ANK1 | Ankyrin-1 | 1881 | 206.265 | 1.627559e+06 | 4.217737e+08 | 79516010.0 | 139440768.0 | 144406208.0 | 58410732.0 | 84.64 | 81.2 | 66.6 | 42.3 |
| sp\|P04406\|G3P\_HUMAN | P04406 | G3P\_HUMAN | GAPDH | Glyceraldehyde-3-phosphate dehydrogenase | 335 | 36.053 | 5.645676e+06 | 2.557258e+08 | 76904110.0 | 159309440.0 | 14283182.0 | 5229045.0 | 90.45 | 100.0 | 78.5 | 58.5 |
| sp\|P69891\|HBG1\_HUMAN | P69891 | HBG1\_HUMAN | HBG1 | Hemoglobin subunit gamma-1 | 147 | 16.140 | 3.261871e+07 | 6.614355e+08 | 71276744.0 | 378870688.0 | 168163968.0 | 43124112.0 | 92.52 | 100.0 | 95.2 | 90.5 |
| sp\|P69892\|HBG2\_HUMAN | P69892 | HBG2\_HUMAN | HBG2 | Hemoglobin subunit gamma-2 | 147 | 16.126 | 3.264703e+07 | 6.614355e+08 | 71276744.0 | 378870688.0 | 168163968.0 | 43124112.0 | 91.16 | 100.0 | 95.2 | 90.5 |
| sp\|P68133\|ACTS\_HUMAN | P68133 | ACTS\_HUMAN | ACTA1 | Actin, alpha skeletal muscle | 377 | 42.051 | 5.045322e+06 | 2.665522e+08 | 66157436.0 | 28298836.0 | 172095968.0 |  | 57.03 | 75.6 | 54.6 |  |
| sp\|P30041\|PRDX6\_HUMAN | P30041 | PRDX6\_HUMAN | PRDX6 | Peroxiredoxin-6 | 224 | 25.035 | 5.963165e+06 | 1.875606e+08 | 64981410.0 | 104999968.0 | 13881664.0 | 3697551.0 | 95.98 | 100.0 | 81.7 | 72.3 |
| sp\|P60709\|ACTB\_HUMAN | P60709 | ACTB\_HUMAN | ACTB | Actin, cytoplasmic 1 | 375 | 41.737 | 1.481314e+07 | 7.767577e+08 | 64081504.0 | 30324504.0 | 607397630.0 | 74954032.0 | 90.4 | 88.5 | 73.9 | 60.3 |
| sp\|P63261\|ACTG\_HUMAN | P63261 | ACTG\_HUMAN | ACTG1 | Actin, cytoplasmic 2 | 375 | 41.793 | 1.312755e+07 | 6.892937e+08 | 64081504.0 | 30324504.0 | 594887680.0 |  | 90.4 | 88.3 | 73.9 |  |
| sp\|P13716\|HEM2\_HUMAN | P13716 | HEM2\_HUMAN | ALAD | Delta-aminolevulinic acid dehydratase | 330 | 36.295 | 3.459988e+06 | 1.577751e+08 | 62011750.0 | 89737216.0 | 4592234.5 | 1433921.2 | 75.76 | 86.7 | 64.8 | 15.2 |
| sp\|Q6S8J3\|POTEE\_HUMAN | Q6S8J3 | POTEE\_HUMAN | POTEE | POTE ankyrin domain family member E | 1075 | 121.363 | 3.895923e+06 | 5.940374e+08 | 60554100.0 | 22822794.0 | 510660512.0 |  | 18.6 | 39.3 | 0.0 |  |
| sp\|P63267\|ACTH\_HUMAN | P63267 | ACTH\_HUMAN | ACTG2 | Actin, gamma-enteric smooth muscle | 376 | 41.360 | 1.150512e+06 | 5.978452e+07 | 59784524.0 |  |  |  | 52.66 |  |  |  |
| sp\|Q13228\|SBP1\_HUMAN | Q13228 | SBP1\_HUMAN | SELENBP1 | Methanethiol oxidase | 472 | 52.391 | 1.927831e+06 | 1.268945e+08 | 59096250.0 | 56327320.0 | 7672410.5 | 3798521.0 | 95.13 | 89.2 | 61.2 | 43.0 |
| sp\|P55072\|TERA\_HUMAN | P55072 | TERA\_HUMAN | VCP | Transitional endoplasmic reticulum ATPase | 806 | 89.322 | 1.338294e+06 | 1.501852e+08 | 55859484.0 | 80219960.0 | 9117976.0 | 4987768.5 | 84.12 | 89.2 | 58.1 | 39.0 |
| sp\|P11142\|HSP7C\_HUMAN | P11142 | HSP7C\_HUMAN | HSPA8 | Heat shock cognate 71 kDa protein | 646 | 70.898 | 1.705858e+06 | 1.519476e+08 | 52876680.0 | 30408126.0 | 58717768.0 | 9945055.0 | 72.76 | 83.9 | 65.0 | 54.6 |
| sp\|P04264\|K2C1\_HUMAN | P04264 | K2C1\_HUMAN | KRT1 | Keratin, type II cytoskeletal 1 | 644 | 66.039 | 4.084533e+06 | 3.388910e+08 | 52762664.0 | 30899744.0 | 65567064.0 | 189661504.0 | 75.31 | 81.1 | 78.7 | 71.6 |
| sp\|P00918\|CAH2\_HUMAN | P00918 | CAH2\_HUMAN | CA2 | Carbonic anhydrase 2 | 260 | 29.246 | 7.231903e+06 | 2.657273e+08 | 52714050.0 | 171261328.0 | 20647688.0 | 21104256.0 | 98.85 | 88.1 | 61.9 | 37.3 |
| sp\|P16452\|EPB42\_HUMAN | P16452 | EPB42\_HUMAN | EPB42 | Protein 4.2 | 691 | 77.009 | 2.438529e+06 | 2.359318e+08 | 51765590.0 | 57637120.0 | 76364616.0 | 50164488.0 | 72.36 | 73.5 | 58.8 | 44.0 |
| sp\|P0CG39\|POTEJ\_HUMAN | P0CG39 | POTEJ\_HUMAN | POTEJ | POTE ankyrin domain family member J | 1038 | 114.180 | 3.369742e+05 | 4.833968e+07 | 48339684.0 |  |  |  | 16.76 |  |  |  |
| sp\|P50395\|GDIB\_HUMAN | P50395 | GDIB\_HUMAN | GDI2 | Rab GDP dissociation inhibitor beta | 445 | 50.663 | 1.821411e+06 | 1.159354e+08 | 46087480.0 | 61781488.0 | 4622173.0 | 3444245.2 | 83.37 | 98.9 | 52.4 | 36.0 |
| sp\|P13798\|ACPH\_HUMAN | P13798 | ACPH\_HUMAN | APEH | Acylamino-acid-releasing enzyme | 732 | 81.225 | 6.341224e+05 | 6.471128e+07 | 43938310.0 | 17473492.0 | 2293292.5 | 1006184.7 | 61.75 | 77.7 | 31.3 | 22.3 |
| sp\|P00558\|PGK1\_HUMAN | P00558 | PGK1\_HUMAN | PGK1 | Phosphoglycerate kinase 1 | 417 | 44.615 | 1.891976e+06 | 1.060507e+08 | 43785210.0 | 57632740.0 | 3048808.0 | 1583960.4 | 95.68 | 99.3 | 69.8 | 32.9 |
| sp\|P35908\|K22E\_HUMAN | P35908 | K22E\_HUMAN | KRT2 | Keratin, type II cytoskeletal 2 epidermal | 639 | 65.433 | 3.294380e+06 | 2.708244e+08 | 43648344.0 | 18179246.0 | 64283584.0 | 144713184.0 | 92.49 | 90.6 | 89.0 | 75.4 |
| sp\|P07195\|LDHB\_HUMAN | P07195 | LDHB\_HUMAN | LDHB | L-lactate dehydrogenase B chain | 334 | 36.638 | 2.262768e+06 | 1.041571e+08 | 42523308.0 | 47593220.0 | 10444754.0 | 3595841.8 | 69.76 | 74.6 | 52.1 | 46.1 |
| sp\|P54652\|HSP72\_HUMAN | P54652 | HSP72\_HUMAN | HSPA2 | Heat shock-related 70 kDa protein 2 | 639 | 70.021 | 1.373827e+06 | 1.208586e+08 | 41342576.0 | 25358876.0 | 44407816.0 | 9749338.0 | 56.18 | 70.3 | 51.5 | 34.7 |
| sp\|Q01082\|SPTB2\_HUMAN | Q01082 | SPTB2\_HUMAN | SPTBN1 | Spectrin beta chain, non-erythrocytic 1 | 2364 | 274.609 | 8.219895e+05 | 2.835948e+08 | 40514516.0 | 7797625.0 | 220196368.0 | 15086253.0 | 7.91 | 29.2 | 44.6 | 20.6 |
| sp\|P34931\|HS71L\_HUMAN | P34931 | HS71L\_HUMAN | HSPA1L | Heat shock 70 kDa protein 1-like | 641 | 70.375 | 1.138826e+06 | 1.006915e+08 | 39633910.0 | 25428544.0 | 35629040.0 |  | 38.22 | 54.0 | 45.4 |  |
| sp\|P00352\|AL1A1\_HUMAN | P00352 | AL1A1\_HUMAN | ALDH1A1 | Aldehyde dehydrogenase 1A1 | 501 | 54.862 | 1.368398e+06 | 9.431943e+07 | 39252936.0 | 42611060.0 | 9215184.0 | 3240246.8 | 76.85 | 90.0 | 47.3 | 25.3 |
| sp\|Q562R1\|ACTBL\_HUMAN | Q562R1 | ACTBL\_HUMAN | ACTBL2 | Beta-actin-like protein 2 | 376 | 42.003 | 3.576690e+06 | 1.887465e+08 | 39155250.0 | 19011048.0 | 105798064.0 | 24782102.0 | 54.52 | 59.3 | 48.7 | 26.3 |
| sp\|P07738\|PMGE\_HUMAN | P07738 | PMGE\_HUMAN | BPGM | Bisphosphoglycerate mutase | 259 | 30.005 | 2.621385e+06 | 9.881925e+07 | 38859560.0 | 49874752.0 | 7371545.0 | 2713396.0 | 81.85 | 85.3 | 45.6 | 34.4 |
| sp\|P00491\|PNPH\_HUMAN | P00491 | PNPH\_HUMAN | PNP | Purine nucleoside phosphorylase | 289 | 32.118 | 2.103471e+06 | 8.487937e+07 | 37649264.0 | 38718916.0 | 5890223.5 | 2620962.2 | 94.46 | 98.3 | 73.7 | 32.9 |
| sp\|Q7Z794\|K2C1B\_HUMAN | Q7Z794 | K2C1B\_HUMAN | KRT77 | Keratin, type II cytoskeletal 1b | 578 | 61.901 | 2.779823e+06 | 2.161882e+08 | 36703916.0 | 11686512.0 | 57168020.0 | 110629776.0 | 36.51 | 44.8 | 46.0 | 38.6 |
| sp\|P0DMV8\|HS71A\_HUMAN | P0DMV8 | HS71A\_HUMAN | HSPA1A | Heat shock 70 kDa protein 1A | 641 | 70.052 | 1.247625e+06 | 1.098050e+08 | 35889960.0 | 28134880.0 | 36847716.0 | 8932396.0 | 68.8 | 87.7 | 47.4 | 30.1 |
| sp\|P17066\|HSP76\_HUMAN | P17066 | HSP76\_HUMAN | HSPA6 | Heat shock 70 kDa protein 6 | 643 | 71.028 | 6.879450e+05 | 6.139040e+07 | 35889960.0 | 25500440.0 |  |  | 20.37 | 56.9 |  |  |
| sp\|Q86VP6\|CAND1\_HUMAN | Q86VP6 | CAND1\_HUMAN | CAND1 | Cullin-associated NEDD8-dissociated protein 1 | 1230 | 136.376 | 5.025010e+05 | 8.609779e+07 | 35483940.0 | 43139060.0 | 2782912.5 | 4691875.0 | 62.28 | 72.7 | 38.5 | 22.4 |
| sp\|P20810\|ICAL\_HUMAN | P20810 | ICAL\_HUMAN | CAST | Calpastatin | 708 | 76.573 | 7.858458e+05 | 7.560145e+07 | 34652780.0 | 27494118.0 | 7143726.0 | 6310829.5 | 72.74 | 80.8 | 41.7 | 30.6 |
| sp\|Q08495\|DEMA\_HUMAN | Q08495 | DEMA\_HUMAN | DMTN | Dematin | 405 | 45.514 | 5.457743e+06 | 3.120867e+08 | 34448864.0 | 11698244.0 | 225101200.0 | 40838364.0 | 81.23 | 81.0 | 80.5 | 67.4 |
| sp\|P35611\|ADDA\_HUMAN | P35611 | ADDA\_HUMAN | ADD1 | Alpha-adducin | 737 | 80.955 | 1.535078e+06 | 1.561318e+08 | 33172998.0 | 35454300.0 | 75387736.0 | 12116722.0 | 68.25 | 64.7 | 47.2 | 33.1 |
| sp\|Q06830\|PRDX1\_HUMAN | Q06830 | PRDX1\_HUMAN | PRDX1 | Peroxiredoxin-1 | 199 | 22.110 | 4.171648e+06 | 1.158814e+08 | 32770900.0 | 33084102.0 | 30922870.0 | 19103486.0 | 80.4 | 97.5 | 50.3 | 49.2 |
| sp\|P04075\|ALDOA\_HUMAN | P04075 | ALDOA\_HUMAN | ALDOA | Fructose-bisphosphate aldolase A | 364 | 39.420 | 1.918413e+06 | 9.501144e+07 | 32689226.0 | 45189516.0 | 7924999.5 | 9207696.0 | 97.25 | 97.3 | 82.7 | 33.2 |
| sp\|P13645\|K1C10\_HUMAN | P13645 | K1C10\_HUMAN | KRT10 | Keratin, type I cytoskeletal 10 | 584 | 58.827 | 3.532021e+06 | 2.610460e+08 | 32406592.0 | 31329234.0 | 65021864.0 | 132288336.0 | 66.61 | 64.9 | 67.0 | 61.8 |
| sp\|Q12955\|ANK3\_HUMAN | Q12955 | ANK3\_HUMAN | ANK3 | Ankyrin-3 | 4377 | 480.410 | 2.187189e+05 | 1.320126e+08 | 32230902.0 | 28517192.0 | 49845000.0 | 21419520.0 | 3.72 | 36.4 | 22.4 | 13.2 |
| sp\|P22314\|UBA1\_HUMAN | P22314 | UBA1\_HUMAN | UBA1 | Ubiquitin-like modifier-activating enzyme 1 | 1058 | 117.849 | 4.790241e+05 | 7.092518e+07 | 31673768.0 | 35132776.0 | 2890614.5 | 1228018.5 | 68.24 | 75.8 | 26.2 | 21.3 |
| sp\|P07384\|CAN1\_HUMAN | P07384 | CAN1\_HUMAN | CAPN1 | Calpain-1 catalytic subunit | 714 | 81.890 | 8.610325e+05 | 8.858650e+07 | 31158822.0 | 31739716.0 | 14155091.0 | 11532876.0 | 63.03 | 70.4 | 41.6 | 29.7 |
| sp\|P07205\|PGK2\_HUMAN | P07205 | PGK2\_HUMAN | PGK2 | Phosphoglycerate kinase 2 | 417 | 44.796 | 6.764266e+05 | 3.806949e+07 | 30604340.0 | 6359238.0 | 1105909.9 |  | 33.33 | 60.4 | 24.9 |  |
| sp\|P52209\|6PGD\_HUMAN | P52209 | 6PGD\_HUMAN | PGD | 6-phosphogluconate dehydrogenase, decarboxylating | 483 | 53.140 | 7.381044e+05 | 4.927839e+07 | 29938460.0 | 13625312.0 | 3481745.0 | 2232871.8 | 54.24 | 71.6 | 42.7 | 25.1 |
| sp\|P11166\|GTR1\_HUMAN | P11166 | GTR1\_HUMAN | SLC2A1 | Solute carrier family 2, facilitated glucose transporter member 1 | 492 | 54.084 | 4.765785e+06 | 3.238325e+08 | 29639584.0 | 32739144.0 | 129708592.0 | 131745160.0 | 24.8 | 41.5 | 25.0 | 20.5 |
| sp\|P35612\|ADDB\_HUMAN | P35612 | ADDB\_HUMAN | ADD2 | Beta-adducin | 726 | 80.854 | 1.594012e+06 | 1.619236e+08 | 28970334.0 | 21164806.0 | 76499104.0 | 35289348.0 | 82.37 | 82.2 | 62.1 | 42.6 |
| sp\|P60174\|TPIS\_HUMAN | P60174 | TPIS\_HUMAN | TPI1 | Triosephosphate isomerase | 249 | 26.669 | 2.592762e+06 | 8.687334e+07 | 28954282.0 | 49952212.0 | 4260094.0 | 3706753.5 | 95.58 | 96.4 | 72.3 | 53.4 |
| sp\|P21980\|TGM2\_HUMAN | P21980 | TGM2\_HUMAN | TGM2 | Protein-glutamine gamma-glutamyltransferase 2 | 687 | 77.329 | 9.241315e+05 | 8.978283e+07 | 28250184.0 | 56986592.0 | 3226557.0 | 1319499.0 | 63.76 | 57.4 | 27.7 | 16.0 |
| sp\|P31939\|PUR9\_HUMAN | P31939 | PUR9\_HUMAN | ATIC | Bifunctional purine biosynthesis protein ATIC | 592 | 64.616 | 7.980077e+05 | 6.478348e+07 | 28043700.0 | 34823900.0 | 1494692.2 | 421190.8 | 85.98 | 84.8 | 61.1 | 25.8 |
| sp\|P48506\|GSH1\_HUMAN | P48506 | GSH1\_HUMAN | GCLC | Glutamate--cysteine ligase catalytic subunit | 637 | 72.766 | 5.651714e+05 | 5.166850e+07 | 27056306.0 | 19076728.0 | 3298562.0 | 2236904.5 | 70.02 | 82.4 | 35.3 | 16.5 |
| sp\|Q5TDH0\|DDI2\_HUMAN | Q5TDH0 | DDI2\_HUMAN | DDI2 | Protein DDI1 homolog 2 | 399 | 44.523 | 1.504517e+06 | 8.415864e+07 | 27053696.0 | 51405100.0 | 3028057.5 | 2671783.2 | 67.17 | 78.4 | 50.4 | 19.8 |
| sp\|P02724\|GLPA\_HUMAN | P02724 | GLPA\_HUMAN | GYPA | Glycophorin-A | 150 | 16.331 | 5.838069e+06 | 1.197841e+08 | 26838234.0 | 46698.812 | 83034608.0 | 9864561.0 | 37.33 | 20.7 | 49.3 | 41.3 |
| sp\|P23526\|SAHH\_HUMAN | P23526 | SAHH\_HUMAN | AHCY | Adenosylhomocysteinase | 432 | 47.716 | 1.452803e+06 | 8.709393e+07 | 26139122.0 | 58186968.0 | 1969298.9 | 798545.44 | 57.87 | 75.5 | 49.8 | 27.8 |
| sp\|P29401\|TKT\_HUMAN | P29401 | TKT\_HUMAN | TKT | Transketolase | 623 | 67.878 | 4.789697e+05 | 4.084644e+07 | 26118652.0 | 12193531.0 | 1715836.2 | 818420.9 | 61.64 | 74.0 | 52.2 | 21.7 |
| sp\|P35527\|K1C9\_HUMAN | P35527 | K1C9\_HUMAN | KRT9 | Keratin, type I cytoskeletal 9 | 623 | 62.064 | 2.201108e+06 | 1.716320e+08 | 25992004.0 | 23845086.0 | 29220224.0 | 92574688.0 | 73.68 | 77.8 | 62.0 | 61.3 |
| sp\|P53396\|ACLY\_HUMAN | P53396 | ACLY\_HUMAN | ACLY | ATP-citrate synthase | 1101 | 120.839 | 3.288770e+05 | 4.992956e+07 | 25609316.0 | 19100130.0 | 3258075.0 | 1962043.0 | 67.12 | 69.9 | 40.7 | 31.9 |
| sp\|O15020\|SPTN2\_HUMAN | O15020 | SPTN2\_HUMAN | SPTBN2 | Spectrin beta chain, non-erythrocytic 2 | 2390 | 271.325 | 4.208556e+05 | 1.434630e+08 | 25558820.0 | 7058091.0 | 101012144.0 | 9833991.0 | 7.36 | 34.0 | 22.0 | 14.5 |
| sp\|P00568\|KAD1\_HUMAN | P00568 | KAD1\_HUMAN | AK1 | Adenylate kinase isoenzyme 1 | 194 | 21.635 | 2.738216e+06 | 7.442894e+07 | 25373494.0 | 30846264.0 | 13355904.0 | 4853274.0 | 90.72 | 91.8 | 50.5 | 42.3 |
| sp\|P04259\|K2C6B\_HUMAN | P04259 | K2C6B\_HUMAN | KRT6B | Keratin, type II cytoskeletal 6B | 564 | 60.067 | 2.351095e+06 | 1.774285e+08 | 23876468.0 | 17024916.0 | 43860688.0 | 92666400.0 | 64.72 | 69.1 | 69.1 | 57.6 |
| sp\|P40925\|MDHC\_HUMAN | P40925 | MDHC\_HUMAN | MDH1 | Malate dehydrogenase, cytoplasmic | 334 | 36.426 | 1.782160e+06 | 8.155965e+07 | 23011868.0 | 35623708.0 | 15672060.0 | 7252013.5 | 61.38 | 71.3 | 47.0 | 31.4 |
| sp\|P54578\|UBP14\_HUMAN | P54578 | UBP14\_HUMAN | USP14 | Ubiquitin carboxyl-terminal hydrolase 14 | 494 | 56.069 | 9.277445e+05 | 6.535343e+07 | 22968922.0 | 36316980.0 | 3692575.8 | 2374950.2 | 79.15 | 73.9 | 43.5 | 29.1 |
| sp\|P62826\|RAN\_HUMAN | P62826 | RAN\_HUMAN | RAN | GTP-binding nuclear protein Ran | 216 | 24.423 | 2.432722e+06 | 7.464635e+07 | 22429348.0 | 31754988.0 | 12349978.0 | 8112035.0 | 59.26 | 74.1 | 34.7 | 30.6 |
| sp\|P11021\|BIP\_HUMAN | P11021 | BIP\_HUMAN | HSPA5 | Endoplasmic reticulum chaperone BiP | 654 | 72.333 | 4.427690e+05 | 4.023749e+07 | 22388678.0 | 2217331.2 | 10809096.0 | 4822382.0 | 55.5 | 55.2 | 44.2 | 32.1 |
| sp\|Q00013\|EM55\_HUMAN | Q00013 | EM55\_HUMAN | MPP1 | 55 kDa erythrocyte membrane protein | 466 | 52.296 | 1.290131e+06 | 8.476554e+07 | 22244584.0 | 12725755.0 | 18516302.0 | 31278896.0 | 95.28 | 93.3 | 61.6 | 40.3 |
| sp\|P06733\|ENOA\_HUMAN | P06733 | ENOA\_HUMAN | ENO1 | Alpha-enolase | 434 | 47.169 | 1.234673e+06 | 7.316875e+07 | 22202060.0 | 46062328.0 | 3497192.5 | 1407172.5 | 79.03 | 93.8 | 52.5 | 22.6 |
| sp\|P07900\|HS90A\_HUMAN | P07900 | HS90A\_HUMAN | HSP90AA1 | Heat shock protein HSP 90-alpha | 732 | 84.660 | 5.466614e+05 | 5.814519e+07 | 21909972.0 | 25542186.0 | 7824506.5 | 2868525.5 | 77.73 | 71.2 | 45.8 | 40.6 |
| sp\|Q15257\|PTPA\_HUMAN | Q15257 | PTPA\_HUMAN | PTPA | Serine/threonine-protein phosphatase 2A activator | 358 | 40.668 | 1.096834e+06 | 5.604167e+07 | 21324696.0 | 32100948.0 | 2284851.8 | 331170.25 | 62.57 | 68.2 | 46.6 | 28.2 |
| sp\|P78417\|GSTO1\_HUMAN | P78417 | GSTO1\_HUMAN | GSTO1 | Glutathione S-transferase omega-1 | 241 | 27.566 | 1.816559e+06 | 6.291301e+07 | 21293684.0 | 34512992.0 | 4643408.0 | 2462926.8 | 71.37 | 79.3 | 55.6 | 37.3 |
| sp\|P49368\|TCPG\_HUMAN | P49368 | TCPG\_HUMAN | CCT3 | T-complex protein 1 subunit gamma | 545 | 60.534 | 6.077157e+05 | 4.621862e+07 | 21150326.0 | 18901096.0 | 3526864.5 | 2640336.0 | 73.39 | 79.1 | 46.8 | 33.6 |
| sp\|Q7Z3Y8\|K1C27\_HUMAN | Q7Z3Y8 | K1C27\_HUMAN | KRT27 | Keratin, type I cytoskeletal 27 | 459 | 49.822 | 1.369586e+06 | 8.572895e+07 | 21015902.0 | 16518441.0 | 25928958.0 | 22265652.0 | 27.23 | 35.7 | 35.3 | 21.4 |
| sp\|P45974\|UBP5\_HUMAN | P45974 | UBP5\_HUMAN | USP5 | Ubiquitin carboxyl-terminal hydrolase 5 | 858 | 95.786 | 4.573587e+05 | 5.503971e+07 | 20805732.0 | 26989826.0 | 4924885.5 | 2319266.2 | 63.17 | 61.1 | 32.9 | 24.4 |
| sp\|P02533\|K1C14\_HUMAN | P02533 | K1C14\_HUMAN | KRT14 | Keratin, type I cytoskeletal 14 | 472 | 51.561 | 2.247979e+06 | 1.456233e+08 | 20736410.0 | 9855676.0 | 26943652.0 | 88087536.0 | 72.03 | 80.3 | 69.3 | 66.9 |
| sp\|P08779\|K1C16\_HUMAN | P08779 | K1C16\_HUMAN | KRT16 | Keratin, type I cytoskeletal 16 | 473 | 51.268 | 2.397952e+06 | 1.544557e+08 | 20736410.0 | 9511294.0 | 36120492.0 | 88087536.0 | 74.63 | 66.2 | 77.6 | 70.8 |
| sp\|P50502\|F10A1\_HUMAN | P50502 | F10A1\_HUMAN | ST13 | Hsc70-interacting protein | 369 | 41.332 | 1.109952e+06 | 5.763785e+07 | 20643814.0 | 24903852.0 | 9204580.0 | 2885605.0 | 59.08 | 54.7 | 47.2 | 33.9 |
| sp\|P02538\|K2C6A\_HUMAN | P02538 | K2C6A\_HUMAN | KRT6A | Keratin, type II cytoskeletal 6A | 564 | 60.045 | 2.303038e+06 | 1.737381e+08 | 20186074.0 | 17024916.0 | 43860688.0 | 92666400.0 | 64.89 | 69.3 | 70.4 | 57.6 |
| sp\|P13647\|K2C5\_HUMAN | P13647 | K2C5\_HUMAN | KRT5 | Keratin, type II cytoskeletal 5 | 590 | 62.378 | 1.768334e+06 | 1.385840e+08 | 20186074.0 | 17716166.0 | 40438248.0 | 60243476.0 | 63.9 | 68.1 | 62.9 | 60.5 |
| sp\|P09972\|ALDOC\_HUMAN | P09972 | ALDOC\_HUMAN | ALDOC | Fructose-bisphosphate aldolase C | 364 | 39.456 | 6.205250e+05 | 3.076022e+07 | 19798800.0 | 7617337.5 | 673218.94 | 2670867.5 | 59.89 | 70.1 | 28.3 | 28.8 |
| sp\|Q5XPI4\|RN123\_HUMAN | Q5XPI4 | RN123\_HUMAN | RNF123 | E3 ubiquitin-protein ligase RNF123 | 1314 | 148.515 | 2.589846e+05 | 4.832384e+07 | 19126360.0 | 25635944.0 | 602426.06 | 2959106.2 | 49.47 | 56.7 | 25.6 | 16.2 |
| sp\|P00338\|LDHA\_HUMAN | P00338 | LDHA\_HUMAN | LDHA | L-lactate dehydrogenase A chain | 332 | 36.689 | 7.937889e+05 | 3.658963e+07 | 18781572.0 | 15504099.0 | 1641568.8 | 662391.2 | 74.4 | 90.7 | 44.6 | 35.2 |
| sp\|Q01484\|ANK2\_HUMAN | Q01484 | ANK2\_HUMAN | ANK2 | Ankyrin-2 | 3957 | 435.270 | 3.432616e+04 | 1.877159e+07 | 18771594.0 |  |  |  | 3.21 |  |  |  |
| sp\|Q9NY33\|DPP3\_HUMAN | Q9NY33 | DPP3\_HUMAN | DPP3 | Dipeptidyl peptidase 3 | 737 | 82.589 | 3.251032e+05 | 3.373344e+07 | 18673946.0 | 13151860.0 | 1475110.5 | 432524.25 | 71.64 | 66.2 | 30.1 | 15.2 |
| sp\|Q7Z3Z0\|K1C25\_HUMAN | Q7Z3Z0 | K1C25\_HUMAN | KRT25 | Keratin, type I cytoskeletal 25 | 450 | 49.500 | 3.002186e+05 | 1.867067e+07 | 18670672.0 |  |  |  | 29.33 |  |  |  |
| sp\|O75955\|FLOT1\_HUMAN | O75955 | FLOT1\_HUMAN | FLOT1 | Flotillin-1 | 427 | 47.355 | 1.990209e+06 | 1.184082e+08 | 18309406.0 | 3544660.5 | 44582900.0 | 51971188.0 | 86.65 | 82.9 | 80.8 | 70.3 |
| sp\|Q9BYC9\|RM20\_HUMAN | Q9BYC9 | RM20\_HUMAN | MRPL20 | Large ribosomal subunit protein bL20m | 149 | 16.390 | 8.800744e+05 | 1.812239e+07 | 18122390.0 |  |  |  | 5.37 |  |  |  |
| sp\|P13489\|RINI\_HUMAN | P13489 | RINI\_HUMAN | RNH1 | Ribonuclease inhibitor | 461 | 49.973 | 8.828989e+05 | 5.543238e+07 | 17704652.0 | 30228386.0 | 6230139.0 | 1269207.8 | 67.25 | 79.2 | 37.3 | 22.8 |
| sp\|Q8NFI4\|F10A5\_HUMAN | Q8NFI4 | F10A5\_HUMAN | ST13P5 | Putative protein FAM10A5 | 369 | 40.590 | 3.459467e+05 | 1.764190e+07 | 17641904.0 |  |  |  | 35.77 |  |  |  |
| sp\|P07451\|CAH3\_HUMAN | P07451 | CAH3\_HUMAN | CA3 | Carbonic anhydrase 3 | 260 | 29.557 | 1.984112e+06 | 7.367901e+07 | 17461156.0 | 51210696.0 | 3286175.8 | 1720978.1 | 81.92 | 87.3 | 38.8 | 24.2 |
| sp\|P12035\|K2C3\_HUMAN | P12035 | K2C3\_HUMAN | KRT3 | Keratin, type II cytoskeletal 3 | 628 | 64.417 | 1.691726e+06 | 1.369139e+08 | 17362222.0 | 15307033.0 | 39576164.0 | 64668492.0 | 22.61 | 48.6 | 51.1 | 37.3 |
| sp\|Q01546\|K22O\_HUMAN | Q01546 | K22O\_HUMAN | KRT76 | Keratin, type II cytoskeletal 2 oral | 638 | 65.841 | 1.933918e+06 | 1.599748e+08 | 17362222.0 |  | 43860688.0 | 98751904.0 | 24.14 |  | 41.7 | 25.5 |
| sp\|P53004\|BIEA\_HUMAN | P53004 | BIEA\_HUMAN | BLVRA | Biliverdin reductase A | 296 | 33.428 | 9.709167e+05 | 4.077646e+07 | 17331104.0 | 19713898.0 | 3003888.5 | 727571.3 | 55.74 | 74.7 | 48.3 | 18.9 |
| sp\|P17174\|AATC\_HUMAN | P17174 | AATC\_HUMAN | GOT1 | Aspartate aminotransferase, cytoplasmic | 413 | 46.248 | 7.026416e+05 | 4.082667e+07 | 17193124.0 | 20928996.0 | 1770563.4 | 933984.75 | 77.24 | 79.2 | 43.8 | 21.5 |
| sp\|P21108\|PRPS3\_HUMAN | P21108 | PRPS3\_HUMAN | PRPS1L1 | Ribose-phosphate pyrophosphokinase 3 | 318 | 34.980 | 3.904927e+05 | 1.716129e+07 | 17161288.0 |  |  |  | 47.8 |  |  |  |
| sp\|P60891\|PRPS1\_HUMAN | P60891 | PRPS1\_HUMAN | PRPS1 | Ribose-phosphate pyrophosphokinase 1 | 318 | 34.834 | 1.204099e+06 | 5.269660e+07 | 17161288.0 | 29701772.0 | 4596410.5 | 1237129.8 | 65.41 | 83.6 | 37.1 | 23.9 |
| sp\|P00492\|HPRT\_HUMAN | P00492 | HPRT\_HUMAN | HPRT1 | Hypoxanthine-guanine phosphoribosyltransferase | 218 | 24.579 | 1.753529e+06 | 5.414948e+07 | 17130204.0 | 35138264.0 | 1485013.2 | 396003.25 | 74.77 | 87.2 | 52.8 | 24.3 |
| sp\|P31948\|STIP1\_HUMAN | P31948 | STIP1\_HUMAN | STIP1 | Stress-induced-phosphoprotein 1 | 543 | 62.639 | 5.359942e+05 | 4.218150e+07 | 17113184.0 | 19004392.0 | 3543441.8 | 2520484.0 | 81.4 | 92.3 | 60.2 | 42.7 |
| sp\|Q14974\|IMB1\_HUMAN | Q14974 | IMB1\_HUMAN | KPNB1 | Importin subunit beta-1 | 876 | 97.170 | 2.575188e+05 | 3.143825e+07 | 17108180.0 | 11411128.0 | 1332120.9 | 1586821.0 | 48.86 | 55.6 | 32.1 | 17.6 |
| sp\|P37837\|TALDO\_HUMAN | P37837 | TALDO\_HUMAN | TALDO1 | Transaldolase | 337 | 37.540 | 1.243942e+06 | 5.866938e+07 | 16964248.0 | 39171888.0 | 1695785.6 | 837455.9 | 64.69 | 68.8 | 51.6 | 32.6 |
| sp\|P22061\|PIMT\_HUMAN | P22061 | PIMT\_HUMAN | PCMT1 | Protein-L-isoaspartate(D-aspartate) O-methyltransferase | 227 | 24.636 | 2.215190e+06 | 6.856434e+07 | 16892626.0 | 40637964.0 | 8242549.5 | 2791201.5 | 75.33 | 94.7 | 51.1 | 24.7 |
| sp\|O15067\|PUR4\_HUMAN | O15067 | PUR4\_HUMAN | PFAS | Phosphoribosylformylglycinamidine synthase | 1338 | 144.734 | 1.450518e+05 | 2.637611e+07 | 16702781.0 | 8711966.0 | 573339.25 | 388027.4 | 52.09 | 64.3 | 20.9 | 16.1 |
| sp\|Q9UIA9\|XPO7\_HUMAN | Q9UIA9 | XPO7\_HUMAN | XPO7 | Exportin-7 | 1087 | 123.907 | 2.463348e+05 | 3.834764e+07 | 16674690.0 | 17877698.0 | 774520.4 | 3020736.0 | 55.38 | 60.4 | 20.7 | 21.3 |
| sp\|P09960\|LKHA4\_HUMAN | P09960 | LKHA4\_HUMAN | LTA4H | Leukotriene A-4 hydrolase | 611 | 69.285 | 3.429229e+05 | 2.985058e+07 | 16657110.0 | 12095976.0 | 807631.25 | 289867.3 | 72.5 | 74.6 | 36.2 | 17.5 |
| sp\|P40227\|TCPZ\_HUMAN | P40227 | TCPZ\_HUMAN | CCT6A | T-complex protein 1 subunit zeta | 531 | 58.024 | 4.327237e+05 | 3.154536e+07 | 16513348.0 | 9957963.0 | 3854709.2 | 1219343.5 | 71.0 | 80.6 | 43.3 | 26.9 |
| sp\|P31150\|GDIA\_HUMAN | P31150 | GDIA\_HUMAN | GDI1 | Rab GDP dissociation inhibitor alpha | 447 | 50.583 | 4.388495e+05 | 2.788928e+07 | 16506334.0 | 7432280.0 | 2615070.0 | 1335599.9 | 70.47 | 85.7 | 44.3 | 15.2 |
| sp\|P48507\|GSH0\_HUMAN | P48507 | GSH0\_HUMAN | GCLM | Glutamate--cysteine ligase regulatory subunit | 274 | 30.727 | 1.249135e+06 | 4.822215e+07 | 16424603.0 | 27411514.0 | 3441069.5 | 944964.1 | 43.8 | 52.9 | 35.8 | 12.8 |
| sp\|O14980\|XPO1\_HUMAN | O14980 | XPO1\_HUMAN | XPO1 | Exportin-1 | 1071 | 123.386 | 1.802281e+05 | 2.793866e+07 | 16424182.0 | 8807680.0 | 678870.0 | 2027926.5 | 61.72 | 56.6 | 23.5 | 16.2 |
| sp\|P08238\|HS90B\_HUMAN | P08238 | HS90B\_HUMAN | HSP90AB1 | Heat shock protein HSP 90-beta | 724 | 83.264 | 4.339388e+05 | 4.539447e+07 | 16420667.0 | 18475144.0 | 7824506.5 | 2674150.2 | 75.28 | 63.3 | 39.1 | 36.9 |
| sp\|Q58FF7\|H90B3\_HUMAN | Q58FF7 | H90B3\_HUMAN | HSP90AB3P | Putative heat shock protein HSP 90-beta-3 | 597 | 65.670 | 1.990245e+05 | 1.642067e+07 | 16420667.0 |  |  |  | 44.56 |  |  |  |
| sp\|P17858\|PFKAL\_HUMAN | P17858 | PFKAL\_HUMAN | PFKL | ATP-dependent 6-phosphofructokinase, liver type | 780 | 85.018 | 2.951322e+05 | 3.152424e+07 | 16179349.0 | 12339362.0 | 1791261.1 | 1214265.4 | 62.31 | 66.5 | 22.8 | 17.6 |
| sp\|Q99497\|PARK7\_HUMAN | Q99497 | PARK7\_HUMAN | PARK7 | Parkinson disease protein 7 | 189 | 19.891 | 1.877133e+06 | 4.691035e+07 | 16006750.0 | 22313140.0 | 6243470.0 | 2346994.8 | 94.71 | 97.9 | 64.6 | 48.1 |
| sp\|P30086\|PEBP1\_HUMAN | P30086 | PEBP1\_HUMAN | PEBP1 | Phosphatidylethanolamine-binding protein 1 | 187 | 21.057 | 1.046959e+06 | 2.769766e+07 | 15960214.0 | 8856350.0 | 2401876.8 | 479223.44 | 80.75 | 88.2 | 58.8 | 20.9 |
| sp\|Q8IZP2\|ST134\_HUMAN | Q8IZP2 | ST134\_HUMAN | ST13P4 | Putative protein FAM10A4 | 240 | 26.400 | 4.785065e+05 | 1.587117e+07 | 15871169.0 |  |  |  | 62.5 |  |  |  |
| sp\|P07203\|GPX1\_HUMAN | P07203 | GPX1\_HUMAN | GPX1 | Glutathione peroxidase 1 | 203 | 22.088 | 1.489238e+06 | 4.132736e+07 | 15777866.0 | 20400080.0 | 4305080.5 | 844331.8 | 68.97 | 89.7 | 33.0 | 28.1 |
| sp\|Q96G03\|PGM2\_HUMAN | Q96G03 | PGM2\_HUMAN | PGM2 | Phosphopentomutase | 612 | 68.283 | 3.634340e+05 | 3.117851e+07 | 15662150.0 | 12809509.0 | 1476316.1 | 1230532.9 | 66.83 | 73.5 | 34.0 | 20.8 |
| sp\|P34932\|HSP74\_HUMAN | P34932 | HSP74\_HUMAN | HSPA4 | Heat shock 70 kDa protein 4 | 840 | 94.331 | 3.531762e+05 | 4.185651e+07 | 15316735.0 | 24678260.0 | 1331266.6 | 530244.6 | 75.83 | 90.2 | 50.4 | 36.7 |
| sp\|P10768\|ESTD\_HUMAN | P10768 | ESTD\_HUMAN | ESD | S-formylglutathione hydrolase | 282 | 31.463 | 8.435899e+05 | 3.334638e+07 | 15147372.0 | 15984861.0 | 2004006.1 | 210136.27 | 82.62 | 85.1 | 59.6 | 12.8 |
| sp\|P78371\|TCPB\_HUMAN | P78371 | TCPB\_HUMAN | CCT2 | T-complex protein 1 subunit beta | 535 | 57.488 | 8.044150e+05 | 5.809978e+07 | 15125757.0 | 37215692.0 | 4107629.0 | 1650704.2 | 92.15 | 92.3 | 37.9 | 30.3 |
| sp\|P11908\|PRPS2\_HUMAN | P11908 | PRPS2\_HUMAN | PRPS2 | Ribose-phosphate pyrophosphokinase 2 | 318 | 34.769 | 7.900625e+05 | 3.451205e+07 | 15039958.0 | 19472096.0 |  |  | 60.38 | 57.2 |  |  |
| sp\|P00390\|GSHR\_HUMAN | P00390 | GSHR\_HUMAN | GSR | Glutathione reductase, mitochondrial | 522 | 56.257 | 3.357023e+05 | 2.372729e+07 | 14865250.0 | 6555599.5 | 1839737.4 | 466700.03 | 69.35 | 86.0 | 40.6 | 22.4 |
| sp\|P0CG47\|UBB\_HUMAN | P0CG47 | UBB\_HUMAN | UBB | Polyubiquitin-B | 229 | 25.762 | 3.918772e+06 | 1.268372e+08 | 14858498.0 | 42725872.0 | 31756282.0 | 37496576.0 | 98.69 | 99.6 | 96.9 | 84.1 |
| sp\|Q13200\|PSMD2\_HUMAN | Q13200 | PSMD2\_HUMAN | PSMD2 | 26S proteasome non-ATPase regulatory subunit 2 | 908 | 100.200 | 2.558548e+05 | 3.220909e+07 | 14809378.0 | 13084415.0 | 3395022.5 | 920274.0 | 74.23 | 76.5 | 46.8 | 27.6 |
| sp\|Q9Y2J2\|E41L3\_HUMAN | Q9Y2J2 | E41L3\_HUMAN | EPB41L3 | Band 4.1-like protein 3 | 1087 | 120.678 | 3.054489e+05 | 4.631096e+07 | 14799449.0 | 5593638.0 | 25917870.0 |  | 11.87 | 34.6 | 23.6 |  |
| sp\|Q5T4S7\|UBR4\_HUMAN | Q5T4S7 | UBR4\_HUMAN | UBR4 | E3 ubiquitin-protein ligase UBR4 | 5183 | 573.841 | 3.673491e+04 | 2.648425e+07 | 14798792.0 | 9080100.0 | 1449045.1 | 1156315.9 | 43.04 | 52.6 | 22.1 | 16.8 |
| sp\|Q9UQ80\|PA2G4\_HUMAN | Q9UQ80 | PA2G4\_HUMAN | PA2G4 | Proliferation-associated protein 2G4 | 394 | 43.787 | 4.250582e+05 | 2.338357e+07 | 14666164.0 | 4927065.5 | 2694794.8 | 1095541.8 | 73.35 | 89.6 | 39.6 | 23.6 |
| sp\|P15311\|EZRI\_HUMAN | P15311 | EZRI\_HUMAN | EZR | Ezrin | 586 | 69.413 | 3.597417e+05 | 3.137247e+07 | 14641890.0 | 14742701.0 | 1987882.1 |  | 59.39 | 72.0 | 42.0 |  |
| sp\|P26038\|MOES\_HUMAN | P26038 | MOES\_HUMAN | MSN | Moesin | 577 | 67.820 | 4.354445e+05 | 3.710289e+07 | 14641890.0 | 19441912.0 | 1994925.8 | 1024165.6 | 76.08 | 89.9 | 42.3 | 32.9 |
| sp\|P35241\|RADI\_HUMAN | P35241 | RADI\_HUMAN | RDX | Radixin | 583 | 68.564 | 3.646372e+05 | 3.141046e+07 | 14641890.0 | 14742701.0 | 1527041.9 | 498826.03 | 78.73 | 70.8 | 41.9 | 28.0 |
| sp\|P63104\|1433Z\_HUMAN | P63104 | 1433Z\_HUMAN | YWHAZ | 14-3-3 protein zeta/delta | 245 | 27.745 | 2.782555e+06 | 9.699417e+07 | 14640914.0 | 30940104.0 | 45872624.0 | 5540532.0 | 81.63 | 82.9 | 81.6 | 44.1 |
| sp\|Q9C0C9\|UBE2O\_HUMAN | Q9C0C9 | UBE2O\_HUMAN | UBE2O | (E3-independent) E2 ubiquitin-conjugating enzyme | 1292 | 141.293 | 2.128810e+05 | 3.778980e+07 | 14534392.0 | 20415454.0 | 1941952.8 | 898003.9 | 59.75 | 55.0 | 32.8 | 16.3 |
| sp\|P49247\|RPIA\_HUMAN | P49247 | RPIA\_HUMAN | RPIA | Ribose-5-phosphate isomerase | 311 | 33.269 | 8.445938e+05 | 3.530244e+07 | 14520016.0 | 18779972.0 | 1654950.4 | 347506.16 | 56.91 | 80.4 | 45.0 | 29.6 |
| sp\|P50990\|TCPQ\_HUMAN | P50990 | TCPQ\_HUMAN | CCT8 | T-complex protein 1 subunit theta | 548 | 59.621 | 5.134289e+05 | 3.845889e+07 | 14361931.0 | 18602812.0 | 3198879.8 | 2295265.5 | 88.87 | 90.9 | 58.6 | 35.9 |
| sp\|P30153\|2AAA\_HUMAN | P30153 | 2AAA\_HUMAN | PPP2R1A | Serine/threonine-protein phosphatase 2A 65 kDa regulatory subunit A alpha isoform | 589 | 65.309 | 6.371053e+05 | 5.227588e+07 | 14319600.0 | 26598584.0 | 7937255.0 | 3420444.2 | 73.34 | 70.6 | 46.3 | 23.9 |
| sp\|P38606\|VATA\_HUMAN | P38606 | VATA\_HUMAN | ATP6V1A | V-type proton ATPase catalytic subunit A | 617 | 68.304 | 3.109190e+05 | 2.668152e+07 | 14159551.0 | 10781413.0 | 1300490.8 | 440060.7 | 60.29 | 73.9 | 43.9 | 20.3 |
| sp\|Q13630\|FCL\_HUMAN | Q13630 | FCL\_HUMAN | GFUS | GDP-L-fucose synthase | 321 | 35.893 | 6.764049e+05 | 3.050237e+07 | 14039206.0 | 13553287.0 | 2094927.8 | 814953.25 | 68.54 | 87.5 | 29.9 | 14.3 |
| sp\|P07954\|FUMH\_HUMAN | P07954 | FUMH\_HUMAN | FH | Fumarate hydratase, mitochondrial | 510 | 54.637 | 4.937998e+05 | 3.389651e+07 | 13940765.0 | 18190504.0 | 1364392.4 | 400845.75 | 64.12 | 71.0 | 31.0 | 20.8 |
| sp\|P30566\|PUR8\_HUMAN | P30566 | PUR8\_HUMAN | ADSL | Adenylosuccinate lyase | 484 | 54.889 | 8.521589e+05 | 5.876559e+07 | 13741652.0 | 41568392.0 | 2557880.0 | 897662.0 | 58.06 | 77.1 | 27.9 | 20.0 |
| sp\|P28289\|TMOD1\_HUMAN | P28289 | TMOD1\_HUMAN | TMOD1 | Tropomodulin-1 | 359 | 40.569 | 1.632036e+06 | 8.318427e+07 | 13691526.0 | 10836951.0 | 52309736.0 | 6346054.5 | 74.09 | 76.6 | 74.1 | 40.9 |
| sp\|Q92526\|TCPW\_HUMAN | Q92526 | TCPW\_HUMAN | CCT6B | T-complex protein 1 subunit zeta-2 | 530 | 58.300 | 1.845582e+05 | 1.351821e+07 | 13518207.0 |  |  |  | 18.11 |  |  |  |
| sp\|P48643\|TCPE\_HUMAN | P48643 | TCPE\_HUMAN | CCT5 | T-complex protein 1 subunit epsilon | 541 | 59.671 | 6.087568e+05 | 4.563776e+07 | 13479263.0 | 26064512.0 | 3654296.5 | 2439688.5 | 79.11 | 96.3 | 55.8 | 33.8 |
| sp\|Q6XQN6\|PNCB\_HUMAN | Q6XQN6 | PNCB\_HUMAN | NAPRT | Nicotinate phosphoribosyltransferase | 538 | 57.578 | 3.954345e+05 | 2.860542e+07 | 13138693.0 | 13939846.0 | 1113787.8 | 413090.38 | 62.83 | 84.9 | 19.1 | 11.5 |
| sp\|O43488\|ARK72\_HUMAN | O43488 | ARK72\_HUMAN | AKR7A2 | Aflatoxin B1 aldehyde reductase member 2 | 359 | 39.589 | 4.400858e+05 | 2.188916e+07 | 13060648.0 | 6707460.0 | 1171444.4 | 949603.25 | 51.53 | 65.2 | 30.1 | 22.0 |
| sp\|P06744\|G6PI\_HUMAN | P06744 | G6PI\_HUMAN | GPI | Glucose-6-phosphate isomerase | 558 | 63.147 | 3.616929e+05 | 2.869524e+07 | 12987442.0 | 14313587.0 | 754515.9 | 639692.0 | 65.59 | 71.0 | 25.8 | 19.7 |
| sp\|Q9UKV8\|AGO2\_HUMAN | Q9UKV8 | AGO2\_HUMAN | AGO2 | Protein argonaute-2 | 859 | 97.208 | 2.535114e+05 | 3.096112e+07 | 12970810.0 | 14011112.0 | 2485174.5 | 1494024.2 | 76.72 | 82.9 | 47.7 | 30.3 |
| sp\|Q99832\|TCPH\_HUMAN | Q99832 | TCPH\_HUMAN | CCT7 | T-complex protein 1 subunit eta | 543 | 59.367 | 4.735937e+05 | 3.532386e+07 | 12970544.0 | 14891492.0 | 5693786.5 | 1768040.8 | 80.85 | 82.5 | 41.1 | 25.8 |
| sp\|P30613\|KPYR\_HUMAN | P30613 | KPYR\_HUMAN | PKLR | Pyruvate kinase PKLR | 574 | 61.830 | 4.875832e+05 | 3.787609e+07 | 12958199.0 | 17503836.0 | 4441820.0 | 2972231.0 | 77.53 | 83.6 | 51.2 | 26.8 |
| sp\|P20073\|ANXA7\_HUMAN | P20073 | ANXA7\_HUMAN | ANXA7 | Annexin A7 | 488 | 52.739 | 8.407000e+05 | 5.570447e+07 | 12886233.0 | 24856408.0 | 15416262.0 | 2545571.8 | 43.44 | 46.1 | 37.3 | 22.7 |
| sp\|P54725\|RD23A\_HUMAN | P54725 | RD23A\_HUMAN | RAD23A | UV excision repair protein RAD23 homolog A | 363 | 39.609 | 6.403282e+05 | 3.186498e+07 | 12831938.0 | 8197535.0 | 6994608.0 | 3840898.8 | 71.35 | 61.2 | 27.8 | 25.1 |
| sp\|P27105\|STOM\_HUMAN | P27105 | STOM\_HUMAN | STOM | Stomatin | 288 | 31.731 | 1.285234e+07 | 5.123694e+08 | 12777166.0 | 15212854.0 | 101701928.0 | 382677440.0 | 73.26 | 67.4 | 74.0 | 69.4 |
| sp\|Q9NTK5\|OLA1\_HUMAN | Q9NTK5 | OLA1\_HUMAN | OLA1 | Obg-like ATPase 1 | 396 | 44.744 | 7.389980e+05 | 4.154275e+07 | 12643438.0 | 23719130.0 | 3741073.2 | 1439107.1 | 84.85 | 90.4 | 45.7 | 24.0 |
| sp\|Q14254\|FLOT2\_HUMAN | Q14254 | FLOT2\_HUMAN | FLOT2 | Flotillin-2 | 428 | 47.064 | 3.886782e+06 | 2.298244e+08 | 12561971.0 | 5122124.0 | 31424904.0 | 180715392.0 | 76.4 | 68.9 | 85.5 | 61.4 |
| sp\|P28066\|PSA5\_HUMAN | P28066 | PSA5\_HUMAN | PSMA5 | Proteasome subunit alpha type-5 | 241 | 26.411 | 8.585930e+05 | 2.848980e+07 | 12500761.0 | 11505440.0 | 3604079.5 | 879515.6 | 61.41 | 76.8 | 43.2 | 12.9 |
| sp\|P31946\|1433B\_HUMAN | P31946 | 1433B\_HUMAN | YWHAB | 14-3-3 protein beta/alpha | 246 | 28.082 | 2.524503e+06 | 8.906786e+07 | 12448090.0 | 33641560.0 | 35978472.0 | 6999737.5 | 74.8 | 81.7 | 74.8 | 46.7 |
| sp\|P29144\|TPP2\_HUMAN | P29144 | TPP2\_HUMAN | TPP2 | Tripeptidyl-peptidase 2 | 1249 | 138.350 | 1.891386e+05 | 3.287581e+07 | 12301903.0 | 18877836.0 | 919397.8 | 776670.7 | 69.58 | 67.6 | 32.4 | 22.2 |
| sp\|P50991\|TCPD\_HUMAN | P50991 | TCPD\_HUMAN | CCT4 | T-complex protein 1 subunit delta | 539 | 57.924 | 3.887157e+05 | 2.828836e+07 | 12078802.0 | 11167455.0 | 4090922.0 | 951182.94 | 81.45 | 86.5 | 50.8 | 33.6 |
| sp\|Q6PCE3\|PGM2L\_HUMAN | Q6PCE3 | PGM2L\_HUMAN | PGM2L1 | Glucose 1,6-bisphosphate synthase | 622 | 70.442 | 2.826534e+05 | 2.501514e+07 | 12059154.0 | 10526695.0 | 1409857.0 | 1019437.75 | 61.74 | 61.9 | 35.0 | 20.6 |
| sp\|Q5XKE5\|K2C79\_HUMAN | Q5XKE5 | K2C79\_HUMAN | KRT79 | Keratin, type II cytoskeletal 79 | 535 | 57.836 | 1.558550e+06 | 1.132494e+08 | 11995080.0 | 7747810.5 | 29278980.0 | 64227568.0 | 17.94 | 45.6 | 35.5 | 40.7 |
| sp\|O75155\|CAND2\_HUMAN | O75155 | CAND2\_HUMAN | CAND2 | Cullin-associated NEDD8-dissociated protein 2 | 1236 | 135.256 | 1.273758e+05 | 2.164515e+07 | 11990026.0 | 9655120.0 |  |  | 8.01 | 39.3 |  |  |
| sp\|O95747\|OXSR1\_HUMAN | O95747 | OXSR1\_HUMAN | OXSR1 | Serine/threonine-protein kinase | 527 | 58.022 | 5.084435e+05 | 3.706402e+07 | 11871533.0 | 16657904.0 | 4459764.0 | 4074822.5 | 59.58 | 57.7 | 38.5 | 20.5 |
| sp\|Q9UNZ2\|NSF1C\_HUMAN | Q9UNZ2 | NSF1C\_HUMAN | NSFL1C | NSFL1 cofactor p47 | 370 | 40.573 | 6.241611e+05 | 3.181639e+07 | 11658888.0 | 15726646.0 | 3011114.5 | 1419744.6 | 67.84 | 83.2 | 62.2 | 53.2 |
| sp\|P17987\|TCPA\_HUMAN | P17987 | TCPA\_HUMAN | TCP1 | T-complex protein 1 subunit alpha | 556 | 60.344 | 5.526739e+05 | 4.190060e+07 | 11645105.0 | 24075260.0 | 4946920.0 | 1233310.0 | 86.15 | 86.2 | 63.7 | 24.8 |
| sp\|P04921\|GLPC\_HUMAN | P04921 | GLPC\_HUMAN | GYPC | Glycophorin-C | 128 | 13.811 | 9.113955e+05 | 1.581427e+07 | 11615196.0 | 652442.06 | 233774.05 | 3312858.2 | 29.69 | 20.3 | 27.3 | 27.3 |
| sp\|Q8WUM4\|PDC6I\_HUMAN | Q8WUM4 | PDC6I\_HUMAN | PDCD6IP | Programmed cell death 6-interacting protein | 868 | 96.023 | 2.441829e+05 | 2.945830e+07 | 11612823.0 | 10860865.0 | 4967902.0 | 2016707.5 | 66.47 | 68.7 | 45.2 | 31.6 |
| sp\|P48426\|PI42A\_HUMAN | P48426 | PI42A\_HUMAN | PIP4K2A | Phosphatidylinositol 5-phosphate 4-kinase type-2 alpha | 406 | 46.225 | 7.119190e+05 | 4.134516e+07 | 11571266.0 | 24632328.0 | 3323602.2 | 1817963.0 | 42.61 | 59.9 | 34.7 | 27.3 |
| sp\|P62258\|1433E\_HUMAN | P62258 | 1433E\_HUMAN | YWHAE | 14-3-3 protein epsilon | 255 | 29.174 | 3.319649e+06 | 1.216761e+08 | 11430686.0 | 38354692.0 | 61895352.0 | 9995369.0 | 90.98 | 87.1 | 77.3 | 63.1 |
| sp\|Q00610\|CLH1\_HUMAN | Q00610 | CLH1\_HUMAN | CLTC | Clathrin heavy chain 1 | 1675 | 191.615 | 1.596283e+05 | 3.842878e+07 | 11306893.0 | 24053186.0 | 1115553.8 | 1953144.8 | 73.13 | 66.3 | 29.8 | 12.7 |
| sp\|P49189\|AL9A1\_HUMAN | P49189 | AL9A1\_HUMAN | ALDH9A1 | 4-trimethylaminobutyraldehyde dehydrogenase | 494 | 53.802 | 4.266250e+05 | 2.883779e+07 | 11160042.0 | 15106418.0 | 1388893.4 | 1182432.1 | 50.4 | 67.8 | 34.0 | 22.1 |
| sp\|Q99460\|PSMD1\_HUMAN | Q99460 | PSMD1\_HUMAN | PSMD1 | 26S proteasome non-ATPase regulatory subunit 1 | 953 | 105.836 | 1.639435e+05 | 2.179941e+07 | 10995966.0 | 9529000.0 | 757392.25 | 517052.34 | 64.74 | 63.8 | 32.2 | 22.6 |
| sp\|P19013\|K2C4\_HUMAN | P19013 | K2C4\_HUMAN | KRT4 | Keratin, type II cytoskeletal 4 | 520 | 56.144 | 1.968291e+06 | 1.388385e+08 | 10789362.0 | 2750871.2 | 40085784.0 | 85212472.0 | 51.15 | 49.8 | 53.8 | 40.8 |
| sp\|P08397\|HEM3\_HUMAN | P08397 | HEM3\_HUMAN | HMBS | Porphobilinogen deaminase | 361 | 39.330 | 6.499353e+05 | 3.211525e+07 | 10759160.0 | 14115370.0 | 5570144.5 | 1670570.8 | 63.71 | 72.0 | 38.5 | 22.2 |
| sp\|P23528\|COF1\_HUMAN | P23528 | COF1\_HUMAN | CFL1 | Cofilin-1 | 166 | 18.502 | 3.231742e+06 | 7.512292e+07 | 10716846.0 | 51986416.0 | 8632455.0 | 3787204.8 | 88.55 | 94.0 | 77.7 | 51.2 |
| sp\|P27348\|1433T\_HUMAN | P27348 | 1433T\_HUMAN | YWHAQ | 14-3-3 protein theta | 245 | 27.764 | 2.174329e+06 | 7.584455e+07 | 10642503.0 | 26033750.0 | 33203148.0 | 5965146.0 | 64.49 | 76.3 | 60.8 | 42.9 |
| sp\|P25325\|THTM\_HUMAN | P25325 | THTM\_HUMAN | MPST | 3-mercaptopyruvate sulfurtransferase | 297 | 33.178 | 8.229021e+05 | 3.430169e+07 | 10540394.0 | 20686828.0 | 1803592.2 | 1270876.5 | 71.38 | 72.7 | 25.3 | 19.9 |
| sp\|P62714\|PP2AB\_HUMAN | P62714 | PP2AB\_HUMAN | PPP2CB | Serine/threonine-protein phosphatase 2A catalytic subunit beta isoform | 309 | 35.575 | 3.496916e+05 | 1.562958e+07 | 10442413.0 | 4209064.5 | 978098.8 |  | 65.05 | 71.5 | 39.5 |  |
| sp\|P67775\|PP2AA\_HUMAN | P67775 | PP2AA\_HUMAN | PPP2CA | Serine/threonine-protein phosphatase 2A catalytic subunit alpha isoform | 309 | 35.594 | 3.185726e+05 | 1.424631e+07 | 10442413.0 | 3731665.0 |  | 72232.06 | 66.67 | 72.2 |  | 15.2 |
| sp\|P09104\|ENOG\_HUMAN | P09104 | ENOG\_HUMAN | ENO2 | Gamma-enolase | 434 | 47.269 | 7.209745e+05 | 4.281673e+07 | 10430608.0 | 31798440.0 | 436363.53 | 151314.66 | 70.74 | 81.8 | 28.1 | 18.4 |
| sp\|P55786\|PSA\_HUMAN | P55786 | PSA\_HUMAN | NPEPPS | Puromycin-sensitive aminopeptidase | 919 | 103.276 | 1.756374e+05 | 2.278944e+07 | 10418850.0 | 11545520.0 | 431286.0 | 393781.97 | 62.35 | 59.2 | 26.6 | 16.5 |
| sp\|Q9BWD1\|THIC\_HUMAN | Q9BWD1 | THIC\_HUMAN | ACAT2 | Acetyl-CoA acetyltransferase, cytosolic | 397 | 41.351 | 3.533883e+05 | 1.835927e+07 | 10401582.0 | 6588214.5 | 1140023.1 | 229449.11 | 73.55 | 89.7 | 29.2 | 22.9 |
| sp\|P35998\|PRS7\_HUMAN | P35998 | PRS7\_HUMAN | PSMC2 | 26S proteasome regulatory subunit 7 | 433 | 48.634 | 4.561472e+05 | 2.787162e+07 | 10367969.0 | 13342302.0 | 2882203.8 | 1279140.6 | 80.6 | 83.4 | 50.8 | 38.1 |
| sp\|P13929\|ENOB\_HUMAN | P13929 | ENOB\_HUMAN | ENO3 | Beta-enolase | 434 | 46.987 | 7.302556e+05 | 4.310918e+07 | 10359576.0 | 30803900.0 | 365909.6 | 1579792.8 | 26.73 | 55.1 | 40.1 | 22.1 |
| sp\|P20618\|PSB1\_HUMAN | P20618 | PSB1\_HUMAN | PSMB1 | Proteasome subunit beta type-1 | 241 | 26.489 | 8.406344e+05 | 2.797627e+07 | 10269384.0 | 15850116.0 | 980258.1 | 876515.25 | 64.32 | 74.3 | 52.7 | 30.7 |
| sp\|O14744\|ANM5\_HUMAN | O14744 | ANM5\_HUMAN | PRMT5 | Protein arginine N-methyltransferase 5 | 637 | 72.684 | 2.052055e+05 | 1.873894e+07 | 10193073.0 | 6008741.5 | 1779996.4 | 757130.44 | 55.89 | 55.7 | 26.8 | 22.8 |
| sp\|O75083\|WDR1\_HUMAN | O75083 | WDR1\_HUMAN | WDR1 | WD repeat-containing protein 1 | 606 | 66.194 | 1.786732e+05 | 1.485920e+07 | 10162261.0 | 4063295.0 | 458988.3 | 174650.78 | 75.25 | 75.9 | 32.8 | 12.5 |
| sp\|P06132\|DCUP\_HUMAN | P06132 | DCUP\_HUMAN | UROD | Uroporphyrinogen decarboxylase | 367 | 40.787 | 3.858871e+05 | 1.977421e+07 | 10090537.0 | 8980536.0 | 703136.94 |  | 56.13 | 72.8 | 44.7 |  |
| sp\|Q14CN4\|K2C72\_HUMAN | Q14CN4 | K2C72\_HUMAN | KRT72 | Keratin, type II cytoskeletal 72 | 511 | 55.877 | 1.297039e+06 | 9.105488e+07 | 10063231.0 |  | 28626170.0 | 52365480.0 | 24.07 |  | 29.9 | 29.4 |
| sp\|P07477\|TRY1\_HUMAN | P07477 | TRY1\_HUMAN | PRSS1 | Serine protease 1 | 247 | 26.558 | 3.028425e+05 | 1.010484e+07 | 10063122.0 |  | 41717.18 |  | 14.17 |  | 15.4 |  |
| sp\|P02008\|HBAZ\_HUMAN | P02008 | HBAZ\_HUMAN | HBZ | Hemoglobin subunit zeta | 142 | 15.637 | 4.331563e+06 | 8.509718e+07 | 10060629.0 | 29615096.0 | 14313132.0 | 31108326.0 | 31.69 | 84.5 | 73.9 | 56.3 |
| sp\|Q16531\|DDB1\_HUMAN | Q16531 | DDB1\_HUMAN | DDB1 | DNA damage-binding protein 1 | 1140 | 126.968 | 1.094433e+05 | 1.745824e+07 | 10038887.0 | 6124839.0 | 802815.06 | 491702.62 | 55.96 | 56.8 | 23.6 | 14.4 |
| sp\|Q9H479\|FN3K\_HUMAN | Q9H479 | FN3K\_HUMAN | FN3K | Fructosamine-3-kinase | 309 | 35.171 | 5.682860e+05 | 2.511128e+07 | 10029087.0 | 12545017.0 | 1614797.0 | 922379.44 | 47.9 | 51.8 | 36.9 | 29.8 |
| sp\|P60842\|IF4A1\_HUMAN | P60842 | IF4A1\_HUMAN | EIF4A1 | Eukaryotic initiation factor 4A-I | 406 | 46.154 | 4.270058e+05 | 2.476055e+07 | 9998650.0 | 10964517.0 | 2926101.8 | 871279.9 | 62.07 | 70.9 | 50.0 | 29.6 |
| sp\|Q7Z6Z7\|HUWE1\_HUMAN | Q7Z6Z7 | HUWE1\_HUMAN | HUWE1 | E3 ubiquitin-protein ligase HUWE1 | 4374 | 481.891 | 3.225837e+04 | 1.953028e+07 | 9967093.0 | 7619781.0 | 438242.3 | 1505161.4 | 46.8 | 49.9 | 19.8 | 16.7 |
| sp\|Q16851\|UGPA\_HUMAN | Q16851 | UGPA\_HUMAN | UGP2 | UTP--glucose-1-phosphate uridylyltransferase | 508 | 56.940 | 3.033099e+05 | 2.169807e+07 | 9958010.0 | 9437842.0 | 1529674.0 | 772546.75 | 69.69 | 76.2 | 39.0 | 35.0 |
| sp\|P09211\|GSTP1\_HUMAN | P09211 | GSTP1\_HUMAN | GSTP1 | Glutathione S-transferase P | 210 | 23.356 | 1.281548e+06 | 3.760541e+07 | 9935022.0 | 26981692.0 | 688697.0 |  | 69.52 | 75.2 | 50.5 |  |
| sp\|Q96P70\|IPO9\_HUMAN | Q96P70 | IPO9\_HUMAN | IPO9 | Importin-9 | 1041 | 115.963 | 1.823887e+05 | 2.657262e+07 | 9889237.0 | 14552780.0 | 341046.03 | 1789559.2 | 34.68 | 35.6 | 22.4 | 18.1 |
| sp\|O95373\|IPO7\_HUMAN | O95373 | IPO7\_HUMAN | IPO7 | Importin-7 | 1038 | 119.517 | 1.409448e+05 | 2.116391e+07 | 9872958.0 | 9477057.0 | 310099.9 | 1503791.5 | 42.29 | 49.5 | 20.3 | 13.0 |
| sp\|Q8IZ83\|A16A1\_HUMAN | Q8IZ83 | A16A1\_HUMAN | ALDH16A1 | Aldehyde dehydrogenase family 16 member A1 | 802 | 85.127 | 2.026497e+05 | 2.167357e+07 | 9827561.0 | 10267242.0 | 1146954.2 | 431810.3 | 45.51 | 43.6 | 20.3 | 11.7 |
| sp\|P04632\|CPNS1\_HUMAN | P04632 | CPNS1\_HUMAN | CAPNS1 | Calpain small subunit 1 | 268 | 28.316 | 1.775996e+06 | 6.318166e+07 | 9777763.0 | 24018734.0 | 18967788.0 | 10417373.0 | 51.12 | 66.0 | 34.0 | 18.3 |
| sp\|Q9BSL1\|UBAC1\_HUMAN | Q9BSL1 | UBAC1\_HUMAN | UBAC1 | Ubiquitin-associated domain-containing protein 1 | 405 | 45.338 | 4.917910e+05 | 2.801303e+07 | 9772331.0 | 13500473.0 | 3437994.2 | 1302233.5 | 53.83 | 65.7 | 41.5 | 37.5 |
| sp\|Q00169\|PIPNA\_HUMAN | Q00169 | PIPNA\_HUMAN | PITPNA | Phosphatidylinositol transfer protein alpha isoform | 270 | 31.806 | 4.889645e+05 | 1.953905e+07 | 9595351.0 | 8316475.0 | 1627228.8 |  | 84.07 | 82.2 | 44.1 |  |
| sp\|P54920\|SNAA\_HUMAN | P54920 | SNAA\_HUMAN | NAPA | Alpha-soluble NSF attachment protein | 295 | 33.233 | 1.022388e+06 | 4.268765e+07 | 9525021.0 | 29198508.0 | 2473957.8 | 1490165.9 | 88.81 | 91.2 | 49.5 | 32.5 |
| sp\|P62195\|PRS8\_HUMAN | P62195 | PRS8\_HUMAN | PSMC5 | 26S proteasome regulatory subunit 8 | 406 | 45.626 | 5.046951e+05 | 2.893068e+07 | 9520929.0 | 16943156.0 | 1643970.6 | 822626.94 | 76.6 | 78.1 | 47.3 | 39.9 |
| sp\|Q14240\|IF4A2\_HUMAN | Q14240 | IF4A2\_HUMAN | EIF4A2 | Eukaryotic initiation factor 4A-II | 407 | 46.402 | 3.181862e+05 | 1.854963e+07 | 9518714.0 | 5661363.5 | 2548177.2 | 821375.06 | 59.46 | 65.4 | 39.6 | 26.5 |
| sp\|P17980\|PRS6A\_HUMAN | P17980 | PRS6A\_HUMAN | PSMC3 | 26S proteasome regulatory subunit 6A | 439 | 49.204 | 7.697020e+05 | 4.758173e+07 | 9493285.0 | 34846700.0 | 2483391.5 | 758350.0 | 75.17 | 74.0 | 50.1 | 30.3 |
| sp\|P19012\|K1C15\_HUMAN | P19012 | K1C15\_HUMAN | KRT15 | Keratin, type I cytoskeletal 15 | 456 | 49.212 | 1.697983e+06 | 1.049836e+08 | 9490180.0 | 3765312.2 | 15613007.0 | 76115104.0 | 28.95 | 41.2 | 53.7 | 44.1 |
| sp\|Q04695\|K1C17\_HUMAN | Q04695 | K1C17\_HUMAN | KRT17 | Keratin, type I cytoskeletal 17 | 432 | 48.106 | 1.743646e+06 | 1.053840e+08 | 9490180.0 | 4534416.0 | 15244303.0 | 76115104.0 | 71.3 | 69.4 | 70.6 | 67.8 |
| sp\|P62191\|PRS4\_HUMAN | P62191 | PRS4\_HUMAN | PSMC1 | 26S proteasome regulatory subunit 4 | 440 | 49.185 | 5.208396e+05 | 3.218503e+07 | 9366512.0 | 18815402.0 | 2633997.2 | 1369114.2 | 72.27 | 79.5 | 48.9 | 36.6 |
| sp\|Q9NRV9\|HEBP1\_HUMAN | Q9NRV9 | HEBP1\_HUMAN | HEBP1 | Heme-binding protein 1 | 189 | 21.097 | 1.125667e+06 | 2.983650e+07 | 9301386.0 | 16404298.0 | 2572204.5 | 1558614.5 | 80.42 | 93.1 | 40.2 | 32.3 |
| sp\|O00410\|IPO5\_HUMAN | O00410 | IPO5\_HUMAN | IPO5 | Importin-5 | 1097 | 123.630 | 1.539127e+05 | 2.390647e+07 | 9256354.0 | 12668934.0 | 762517.0 | 1218660.1 | 52.42 | 63.6 | 21.3 | 12.7 |
| sp\|P13796\|PLSL\_HUMAN | P13796 | PLSL\_HUMAN | LCP1 | Plastin-2 | 627 | 70.288 | 2.599784e+05 | 2.295808e+07 | 9245579.0 | 12191763.0 | 1231428.1 | 289312.3 | 69.06 | 77.0 | 50.9 | 28.5 |
| sp\|P46063\|RECQ1\_HUMAN | P46063 | RECQ1\_HUMAN | RECQL | ATP-dependent DNA helicase Q1 | 649 | 73.457 | 9.927625e+04 | 9.162114e+06 | 9122842.0 | 39272.023 |  |  | 4.47 | 31.7 |  |  |
| sp\|P28072\|PSB6\_HUMAN | P28072 | PSB6\_HUMAN | PSMB6 | Proteasome subunit beta type-6 | 239 | 25.358 | 5.677250e+05 | 1.808715e+07 | 9084738.0 | 6280931.0 | 1690730.2 | 1030753.4 | 53.97 | 81.2 | 15.9 | 15.9 |
| sp\|Q16401\|PSMD5\_HUMAN | Q16401 | PSMD5\_HUMAN | PSMD5 | 26S proteasome non-ATPase regulatory subunit 5 | 504 | 56.196 | 4.436828e+05 | 3.132530e+07 | 9049226.0 | 16449311.0 | 4566034.0 | 1260724.5 | 58.33 | 64.1 | 31.5 | 21.2 |
| sp\|P24666\|PPAC\_HUMAN | P24666 | PPAC\_HUMAN | ACP1 | Low molecular weight phosphotyrosine protein phosphatase | 158 | 18.042 | 1.166692e+06 | 2.644587e+07 | 9044246.0 | 12297654.0 | 4420004.0 | 683970.1 | 77.22 | 75.9 | 51.3 | 38.0 |
| sp\|P08237\|PFKAM\_HUMAN | P08237 | PFKAM\_HUMAN | PFKM | ATP-dependent 6-phosphofructokinase, muscle type | 780 | 85.183 | 2.013075e+05 | 2.154419e+07 | 9041212.0 | 11517250.0 | 421672.62 | 564051.75 | 60.0 | 68.3 | 31.2 | 17.4 |
| sp\|P11413\|G6PD\_HUMAN | P11413 | G6PD\_HUMAN | G6PD | Glucose-6-phosphate 1-dehydrogenase | 515 | 59.257 | 3.447193e+05 | 2.566388e+07 | 9023663.0 | 13984627.0 | 1934595.4 | 720999.25 | 77.86 | 89.7 | 38.3 | 27.2 |
| sp\|O43681\|GET3\_HUMAN | O43681 | GET3\_HUMAN | GET3 | ATPase GET3 | 348 | 38.793 | 5.848229e+05 | 2.850328e+07 | 9015666.0 | 17833574.0 | 1449856.9 | 204185.47 | 47.99 | 66.7 | 30.7 | 6.6 |
| sp\|Q9BS40\|LXN\_HUMAN | Q9BS40 | LXN\_HUMAN | LXN | Latexin | 222 | 25.750 | 7.566402e+05 | 2.447844e+07 | 8972524.0 | 9284480.0 | 5263434.0 | 958005.8 | 28.83 | 53.2 | 31.5 | 19.4 |
| sp\|P61981\|1433G\_HUMAN | P61981 | 1433G\_HUMAN | YWHAG | 14-3-3 protein gamma | 247 | 28.303 | 2.201875e+06 | 7.829651e+07 | 8972164.0 | 28822734.0 | 33521892.0 | 6979715.5 | 63.97 | 76.1 | 75.3 | 38.5 |
| sp\|Q9Y3I1\|FBX7\_HUMAN | Q9Y3I1 | FBX7\_HUMAN | FBXO7 | F-box only protein 7 | 522 | 58.503 | 5.023947e+05 | 3.692668e+07 | 8842294.0 | 23205462.0 | 4713899.0 | 165028.06 | 22.99 | 42.9 | 16.1 | 9.4 |
| sp\|P52907\|CAZA1\_HUMAN | P52907 | CAZA1\_HUMAN | CAPZA1 | F-actin-capping protein subunit alpha-1 | 286 | 32.923 | 7.891709e+05 | 3.264282e+07 | 8825528.0 | 19055772.0 | 4205277.0 | 556240.25 | 80.42 | 90.2 | 24.1 | 22.7 |
| sp\|Q9HC38\|GLOD4\_HUMAN | Q9HC38 | GLOD4\_HUMAN | GLOD4 | Glyoxalase domain-containing protein 4 | 313 | 34.793 | 6.284527e+05 | 2.747145e+07 | 8726520.0 | 16406729.0 | 1369290.4 | 968913.7 | 70.29 | 86.9 | 47.0 | 29.1 |
| sp\|Q16775\|GLO2\_HUMAN | Q16775 | GLO2\_HUMAN | HAGH | Hydroxyacylglutathione hydrolase, mitochondrial | 308 | 33.806 | 6.810310e+05 | 2.892530e+07 | 8720882.0 | 13767982.0 | 3880365.5 | 2556067.5 | 60.06 | 76.0 | 39.6 | 18.2 |
| sp\|P22234\|PUR6\_HUMAN | P22234 | PUR6\_HUMAN | PAICS | Bifunctional phosphoribosylaminoimidazole carboxylase/phosphoribosylaminoimidazole succinocarboxamide synthetase | 425 | 47.079 | 4.552743e+05 | 2.692883e+07 | 8497330.0 | 14585311.0 | 2306109.8 | 1540080.8 | 62.12 | 74.8 | 36.0 | 24.2 |
| sp\|Q5VVQ6\|OTU1\_HUMAN | Q5VVQ6 | OTU1\_HUMAN | YOD1 | Ubiquitin thioesterase OTU1 | 348 | 38.322 | 5.429682e+05 | 2.614205e+07 | 8471189.0 | 12433209.0 | 3915710.5 | 1321945.2 | 45.11 | 67.8 | 30.7 | 23.9 |
| sp\|Q16658\|FSCN1\_HUMAN | Q16658 | FSCN1\_HUMAN | FSCN1 | Fascin | 493 | 54.530 | 3.507857e+05 | 2.403225e+07 | 8419960.0 | 12101373.0 | 3088047.0 | 422872.5 | 65.11 | 75.5 | 30.4 | 20.9 |
| sp\|O60610\|DIAP1\_HUMAN | O60610 | DIAP1\_HUMAN | DIAPH1 | Protein diaphanous homolog 1 | 1272 | 141.347 | 1.689992e+05 | 3.001155e+07 | 8385127.0 | 7651152.0 | 11309740.0 | 2665534.5 | 50.16 | 59.4 | 46.7 | 26.7 |
| sp\|Q13867\|BLMH\_HUMAN | Q13867 | BLMH\_HUMAN | BLMH | Bleomycin hydrolase | 455 | 52.562 | 5.273943e+05 | 3.482768e+07 | 8362482.0 | 25873454.0 | 306213.4 | 285526.5 | 51.87 | 65.5 | 26.8 | 10.8 |
| sp\|P13646\|K1C13\_HUMAN | P13646 | K1C13\_HUMAN | KRT13 | Keratin, type I cytoskeletal 13 | 458 | 49.588 | 1.435599e+06 | 8.943896e+07 | 8357445.5 | 17076808.0 | 16502900.0 | 47501808.0 | 48.91 | 56.6 | 59.0 | 47.8 |
| sp\|P18669\|PGAM1\_HUMAN | P18669 | PGAM1\_HUMAN | PGAM1 | Phosphoglycerate mutase 1 | 254 | 28.804 | 9.139184e+05 | 3.307329e+07 | 8317384.0 | 24010974.0 | 744930.4 |  | 72.05 | 79.1 | 56.3 |  |
| sp\|Q9NSB2\|KRT84\_HUMAN | Q9NSB2 | KRT84\_HUMAN | KRT84 | Keratin, type II cuticular Hb4 | 600 | 64.842 | 4.232721e+05 | 3.448206e+07 | 8239358.5 |  | 26242700.0 |  | 37.83 |  | 39.3 |  |
| sp\|P28074\|PSB5\_HUMAN | P28074 | PSB5\_HUMAN | PSMB5 | Proteasome subunit beta type-5 | 263 | 28.480 | 6.746418e+05 | 2.413962e+07 | 8210075.5 | 13689027.0 | 1736830.1 | 503686.62 | 65.4 | 80.6 | 57.4 | 37.6 |
| sp\|P50570\|DYN2\_HUMAN | P50570 | DYN2\_HUMAN | DNM2 | Dynamin-2 | 870 | 98.064 | 1.703184e+05 | 2.098401e+07 | 8209004.5 | 7724715.0 | 4019466.5 | 1030821.6 | 66.32 | 60.9 | 41.4 | 23.1 |
| sp\|P36959\|GMPR1\_HUMAN | P36959 | GMPR1\_HUMAN | GMPR | GMP reductase 1 | 345 | 37.419 | 3.750492e+05 | 1.763184e+07 | 8205551.0 | 6743781.0 | 1820789.1 | 861717.1 | 61.45 | 75.1 | 26.7 | 23.8 |
| sp\|P28070\|PSB4\_HUMAN | P28070 | PSB4\_HUMAN | PSMB4 | Proteasome subunit beta type-4 | 264 | 29.204 | 6.854682e+05 | 2.515051e+07 | 8130750.0 | 14469084.0 | 2027937.1 | 522739.25 | 53.03 | 72.3 | 21.6 | 18.6 |
| sp\|P25788\|PSA3\_HUMAN | P25788 | PSA3\_HUMAN | PSMA3 | Proteasome subunit alpha type-3 | 255 | 28.433 | 5.842657e+05 | 2.087134e+07 | 8118266.0 | 10755485.0 | 1554623.2 | 442961.62 | 63.92 | 69.8 | 48.6 | 38.8 |
| sp\|P29972\|AQP1\_HUMAN | P29972 | AQP1\_HUMAN | AQP1 | Aquaporin-1 | 269 | 28.526 | 7.851602e+05 | 2.813949e+07 | 8107059.5 | 964541.25 | 4571946.0 | 14495948.0 | 40.89 | 52.0 | 16.4 | 8.9 |
| sp\|P16930\|FAAA\_HUMAN | P16930 | FAAA\_HUMAN | FAH | Fumarylacetoacetase | 419 | 46.374 | 2.871523e+05 | 1.673031e+07 | 8050092.0 | 7585448.0 | 948665.1 | 146104.48 | 66.11 | 74.9 | 23.4 | 15.3 |
| sp\|P30154\|2AAB\_HUMAN | P30154 | 2AAB\_HUMAN | PPP2R1B | Serine/threonine-protein phosphatase 2A 65 kDa regulatory subunit A beta isoform | 601 | 66.214 | 4.108712e+05 | 3.418005e+07 | 8048953.5 | 24008230.0 | 2122869.2 |  | 69.55 | 66.7 | 33.4 |  |
| sp\|P14735\|IDE\_HUMAN | P14735 | IDE\_HUMAN | IDE | Insulin-degrading enzyme | 1019 | 117.968 | 1.187501e+05 | 1.760010e+07 | 8020294.5 | 8852301.0 | 407523.06 | 319983.75 | 63.89 | 55.8 | 25.2 | 17.9 |
| sp\|P21281\|VATB2\_HUMAN | P21281 | VATB2\_HUMAN | ATP6V1B2 | V-type proton ATPase subunit B, brain isoform | 511 | 56.501 | 2.563384e+05 | 1.819646e+07 | 7845537.0 | 9354208.0 | 558498.9 | 438218.88 | 67.71 | 75.1 | 32.9 | 16.8 |
| sp\|Q5QJ74\|TBCEL\_HUMAN | Q5QJ74 | TBCEL\_HUMAN | TBCEL | Tubulin-specific chaperone cofactor E-like protein | 424 | 48.195 | 3.653458e+05 | 2.212194e+07 | 7822681.5 | 12773782.0 | 993100.7 | 532376.3 | 54.72 | 65.3 | 33.3 | 20.5 |
| sp\|P46459\|NSF\_HUMAN | P46459 | NSF\_HUMAN | NSF | Vesicle-fusing ATPase | 744 | 82.594 | 3.231309e+05 | 3.353082e+07 | 7820845.0 | 22646076.0 | 1946706.9 | 1117193.8 | 68.28 | 67.7 | 47.6 | 29.3 |
| sp\|Q9Y281\|COF2\_HUMAN | Q9Y281 | COF2\_HUMAN | CFL2 | Cofilin-2 | 166 | 18.737 | 9.911409e+05 | 2.333204e+07 | 7802704.0 | 10725992.0 | 4019056.0 | 784283.3 | 44.58 | 83.1 | 52.4 | 13.9 |
| sp\|P20020\|AT2B1\_HUMAN | P20020 | AT2B1\_HUMAN | ATP2B1 | Plasma membrane calcium-transporting ATPase 1 | 1220 | 134.685 | 3.081221e+05 | 5.213858e+07 | 7768164.0 | 3110783.0 | 9166692.0 | 32092944.0 | 39.92 | 43.8 | 38.8 | 32.2 |
| sp\|P23634\|AT2B4\_HUMAN | P23634 | AT2B4\_HUMAN | ATP2B4 | Plasma membrane calcium-transporting ATPase 4 | 1241 | 137.920 | 2.633176e+05 | 4.562725e+07 | 7768164.0 | 3906483.5 | 10332389.0 | 23620218.0 | 46.74 | 51.5 | 37.1 | 30.4 |
| sp\|P18577\|RHCE\_HUMAN | P18577 | RHCE\_HUMAN | RHCE | Blood group Rh(CE) polypeptide | 417 | 45.560 | 4.869662e+05 | 2.787403e+07 | 7598603.0 | 1559491.0 | 14073177.0 | 4642755.0 | 22.78 | 26.6 | 18.2 | 16.5 |
| sp\|O43396\|TXNL1\_HUMAN | O43396 | TXNL1\_HUMAN | TXNL1 | Thioredoxin-like protein 1 | 289 | 32.251 | 7.219245e+05 | 2.925177e+07 | 7587695.0 | 19433682.0 | 1367233.4 | 863155.75 | 62.63 | 77.2 | 42.2 | 29.8 |
| sp\|Q13618\|CUL3\_HUMAN | Q13618 | CUL3\_HUMAN | CUL3 | Cullin-3 | 768 | 88.930 | 1.830161e+05 | 2.044818e+07 | 7549602.5 | 10263138.0 | 1603625.1 | 1031819.1 | 69.66 | 68.0 | 40.6 | 28.8 |
| sp\|O00231\|PSD11\_HUMAN | O00231 | PSD11\_HUMAN | PSMD11 | 26S proteasome non-ATPase regulatory subunit 11 | 422 | 47.464 | 3.045796e+05 | 1.816278e+07 | 7548485.0 | 9075170.0 | 1069380.2 | 469741.03 | 75.12 | 77.5 | 44.1 | 21.1 |
| sp\|Q02161\|RHD\_HUMAN | Q02161 | RHD\_HUMAN | RHD | Blood group Rh(D) polypeptide | 417 | 45.211 | 1.577745e+05 | 8.961860e+06 | 7528084.5 | 1433775.9 |  |  | 13.43 | 20.6 |  |  |
| sp\|Q01813\|PFKAP\_HUMAN | Q01813 | PFKAP\_HUMAN | PFKP | ATP-dependent 6-phosphofructokinase, platelet type | 784 | 85.596 | 1.077310e+05 | 1.158540e+07 | 7527313.0 | 3414149.2 | 643940.2 |  | 46.56 | 46.6 | 18.4 |  |
| sp\|Q14166\|TTL12\_HUMAN | Q14166 | TTL12\_HUMAN | TTLL12 | Tubulin--tyrosine ligase-like protein 12 | 644 | 74.404 | 2.868675e+05 | 2.681604e+07 | 7520855.5 | 17143830.0 | 2010333.1 | 141025.67 | 61.34 | 72.4 | 30.4 | 8.4 |
| sp\|P11586\|C1TC\_HUMAN | P11586 | C1TC\_HUMAN | MTHFD1 | C-1-tetrahydrofolate synthase, cytoplasmic | 935 | 101.531 | 1.221885e+05 | 1.558642e+07 | 7499359.0 | 6140862.5 | 980848.7 | 965345.0 | 68.88 | 72.6 | 38.3 | 22.5 |
| sp\|Q15102\|PA1B3\_HUMAN | Q15102 | PA1B3\_HUMAN | PAFAH1B3 | Platelet-activating factor acetylhydrolase IB subunit alpha1 | 231 | 25.734 | 7.928083e+05 | 2.563260e+07 | 7497756.5 | 15472766.0 | 1966308.8 | 695768.56 | 86.58 | 80.1 | 53.2 | 34.2 |
| sp\|P52565\|GDIR1\_HUMAN | P52565 | GDIR1\_HUMAN | ARHGDIA | Rho GDP-dissociation inhibitor 1 | 204 | 23.207 | 1.281736e+06 | 3.737099e+07 | 7486702.5 | 27921970.0 | 1572268.0 | 390047.94 | 57.35 | 97.5 | 50.0 | 26.5 |
| sp\|Q9Y4E8\|UBP15\_HUMAN | Q9Y4E8 | UBP15\_HUMAN | USP15 | Ubiquitin carboxyl-terminal hydrolase 15 | 981 | 112.419 | 1.536361e+05 | 2.169953e+07 | 7441666.5 | 8714331.0 | 4092786.2 | 1450741.5 | 67.58 | 66.5 | 42.1 | 24.2 |
| sp\|Q04917\|1433F\_HUMAN | Q04917 | 1433F\_HUMAN | YWHAH | 14-3-3 protein eta | 246 | 28.219 | 1.864303e+06 | 6.609600e+07 | 7418791.0 | 21915086.0 | 31305860.0 | 5456264.0 | 63.41 | 70.3 | 59.3 | 42.7 |
| sp\|Q99733\|NP1L4\_HUMAN | Q99733 | NP1L4\_HUMAN | NAP1L4 | Nucleosome assembly protein 1-like 4 | 375 | 42.823 | 3.989604e+05 | 2.146466e+07 | 7398976.5 | 11313926.0 | 2057438.5 | 694318.9 | 46.67 | 44.5 | 25.1 | 15.5 |
| sp\|O95757\|HS74L\_HUMAN | O95757 | HS74L\_HUMAN | HSPA4L | Heat shock 70 kDa protein 4L | 839 | 94.512 | 7.542072e+04 | 8.955602e+06 | 7344951.0 | 1610651.2 |  |  | 43.74 | 52.8 |  |  |
| sp\|P36969\|GPX4\_HUMAN | P36969 | GPX4\_HUMAN | GPX4 | Phospholipid hydroperoxide glutathione peroxidase | 197 | 22.175 | 4.592850e+05 | 1.279567e+07 | 7316096.0 | 3678619.0 | 1458454.5 | 342500.34 | 45.69 | 67.0 | 43.1 | 28.4 |
| sp\|Q3LXA3\|TKFC\_HUMAN | Q3LXA3 | TKFC\_HUMAN | TKFC | Triokinase/FMN cyclase | 575 | 58.947 | 2.879422e+05 | 2.132476e+07 | 7308967.0 | 13049043.0 | 741283.7 | 225470.36 | 69.22 | 63.5 | 28.2 | 19.5 |
| sp\|P48637\|GSHB\_HUMAN | P48637 | GSHB\_HUMAN | GSS | Glutathione synthetase | 474 | 52.385 | 3.006221e+05 | 1.978541e+07 | 7180746.0 | 10279053.0 | 1704435.2 | 621172.6 | 73.21 | 76.8 | 29.1 | 20.0 |
| sp\|Q9P2R3\|ANFY1\_HUMAN | Q9P2R3 | ANFY1\_HUMAN | ANKFY1 | Rabankyrin-5 | 1169 | 128.399 | 9.233870e+04 | 1.489576e+07 | 7177238.0 | 6989683.5 | 479842.88 | 248992.39 | 56.03 | 46.9 | 20.4 | 12.1 |
| sp\|Q15008\|PSMD6\_HUMAN | Q15008 | PSMD6\_HUMAN | PSMD6 | 26S proteasome non-ATPase regulatory subunit 6 | 389 | 45.531 | 3.707033e+05 | 2.120561e+07 | 7163977.5 | 12023732.0 | 1341639.5 | 676260.94 | 73.26 | 80.7 | 43.2 | 25.4 |
| sp\|P48739\|PIPNB\_HUMAN | P48739 | PIPNB\_HUMAN | PITPNB | Phosphatidylinositol transfer protein beta isoform | 271 | 31.540 | 3.886557e+05 | 1.540082e+07 | 7158962.0 | 6382644.0 | 1539036.0 | 320178.84 | 84.13 | 87.5 | 48.3 | 24.4 |
| sp\|P49720\|PSB3\_HUMAN | P49720 | PSB3\_HUMAN | PSMB3 | Proteasome subunit beta type-3 | 205 | 22.949 | 6.293309e+05 | 1.814513e+07 | 7107057.5 | 10178243.0 | 698576.75 | 161249.52 | 46.34 | 62.0 | 43.9 | 33.2 |
| sp\|Q9HA64\|KT3K\_HUMAN | Q9HA64 | KT3K\_HUMAN | FN3KRP | Ketosamine-3-kinase | 309 | 34.412 | 2.779437e+05 | 1.201666e+07 | 7093113.0 | 3401940.5 | 961761.0 | 559848.0 | 47.57 | 52.1 | 41.1 | 12.3 |
| sp\|P60900\|PSA6\_HUMAN | P60900 | PSA6\_HUMAN | PSMA6 | Proteasome subunit alpha type-6 | 246 | 27.399 | 4.953719e+05 | 1.705231e+07 | 7088469.5 | 7610080.0 | 1310540.1 | 1043219.75 | 55.69 | 64.2 | 56.9 | 28.5 |
| sp\|P47756\|CAPZB\_HUMAN | P47756 | CAPZB\_HUMAN | CAPZB | F-actin-capping protein subunit beta | 272 | 30.629 | 6.110657e+05 | 2.351461e+07 | 7081826.0 | 14285780.0 | 1651574.5 | 495433.75 | 61.03 | 65.0 | 48.0 | 24.5 |
| sp\|Q96Q15\|SMG1\_HUMAN | Q96Q15 | SMG1\_HUMAN | SMG1 | Serine/threonine-protein kinase SMG1 | 3661 | 410.501 | 1.388044e+04 | 7.158709e+06 | 7035418.0 | 123290.516 |  |  | 2.7 | 26.1 |  |  |
| sp\|P31947\|1433S\_HUMAN | P31947 | 1433S\_HUMAN | SFN | 14-3-3 protein sigma | 248 | 27.774 | 1.557689e+06 | 5.435460e+07 | 7009506.0 | 18947958.0 | 25197420.0 | 3199711.2 | 36.69 | 54.0 | 48.0 | 38.7 |
| sp\|Q96IU4\|ABHEB\_HUMAN | Q96IU4 | ABHEB\_HUMAN | ABHD14B | Putative protein-lysine deacylase ABHD14B | 210 | 22.346 | 5.581733e+05 | 1.567061e+07 | 6961952.0 | 6600565.0 | 1873804.4 | 234292.9 | 50.48 | 56.2 | 34.3 | 22.9 |
| sp\|P12955\|PEPD\_HUMAN | P12955 | PEPD\_HUMAN | PEPD | Xaa-Pro dipeptidase | 493 | 54.548 | 2.944027e+05 | 2.017613e+07 | 6956954.5 | 12715721.0 | 503450.06 |  | 43.81 | 56.8 | 26.8 |  |
| sp\|Q04760\|LGUL\_HUMAN | Q04760 | LGUL\_HUMAN | GLO1 | Lactoylglutathione lyase | 184 | 20.778 | 3.723793e+05 | 9.720899e+06 | 6910256.0 | 628405.8 | 1341245.8 | 840991.25 | 75.0 | 80.4 | 41.3 | 24.5 |
| sp\|Q9Y2V2\|CHSP1\_HUMAN | Q9Y2V2 | CHSP1\_HUMAN | CARHSP1 | Calcium-regulated heat-stable protein 1 | 147 | 15.892 | 1.434980e+06 | 2.865111e+07 | 6904055.0 | 15728537.0 | 4269495.5 | 1749022.1 | 79.59 | 96.6 | 36.1 | 30.6 |
| sp\|P05062\|ALDOB\_HUMAN | P05062 | ALDOB\_HUMAN | ALDOB | Fructose-bisphosphate aldolase B | 364 | 39.473 | 1.378477e+05 | 6.836233e+06 | 6809668.0 |  | 26565.21 |  | 19.23 |  | 12.4 |  |
| sp\|O43813\|LANC1\_HUMAN | O43813 | LANC1\_HUMAN | LANCL1 | Glutathione S-transferase LANCL1 | 399 | 45.283 | 2.243282e+05 | 1.276251e+07 | 6807710.5 | 4970843.0 | 414681.62 | 569279.25 | 44.61 | 66.9 | 17.5 | 14.8 |
| sp\|Q9Y4G8\|RPGF2\_HUMAN | Q9Y4G8 | RPGF2\_HUMAN | RAPGEF2 | Rap guanine nucleotide exchange factor 2 | 1499 | 167.417 | 5.162916e+04 | 1.085955e+07 | 6649092.5 | 3628059.0 | 372528.0 | 209868.52 | 47.77 | 50.2 | 23.0 | 6.3 |
| sp\|Q9UDX3\|S14L4\_HUMAN | Q9UDX3 | S14L4\_HUMAN | SEC14L4 | SEC14-like protein 4 | 406 | 46.644 | 3.066754e+05 | 1.797181e+07 | 6618446.0 | 7459926.0 | 3114700.2 | 778740.44 | 57.39 | 56.9 | 32.3 | 20.2 |
| sp\|P17612\|KAPCA\_HUMAN | P17612 | KAPCA\_HUMAN | PRKACA | cAMP-dependent protein kinase catalytic subunit alpha | 351 | 40.590 | 1.738172e+05 | 8.863982e+06 | 6615608.5 | 2248373.5 |  |  | 67.52 | 58.4 |  |  |
| sp\|P22694\|KAPCB\_HUMAN | P22694 | KAPCB\_HUMAN | PRKACB | cAMP-dependent protein kinase catalytic subunit beta | 351 | 40.623 | 1.986791e+05 | 1.014008e+07 | 6615608.5 | 2123869.0 | 761892.1 | 638708.1 | 42.74 | 49.3 | 24.8 | 16.8 |
| sp\|P43034\|LIS1\_HUMAN | P43034 | LIS1\_HUMAN | PAFAH1B1 | Platelet-activating factor acetylhydrolase IB subunit beta | 410 | 46.638 | 2.510465e+05 | 1.470995e+07 | 6543836.5 | 6741503.5 | 899135.25 | 525476.8 | 77.32 | 73.2 | 42.7 | 36.6 |
| sp\|P23276\|KELL\_HUMAN | P23276 | KELL\_HUMAN | KEL | Kell blood group glycoprotein | 732 | 82.824 | 2.863785e+05 | 2.979983e+07 | 6543615.0 | 5008143.5 | 6488292.5 | 11759780.0 | 46.17 | 52.6 | 28.7 | 30.7 |
| sp\|Q8N1N4\|K2C78\_HUMAN | Q8N1N4 | K2C78\_HUMAN | KRT78 | Keratin, type II cytoskeletal 78 | 520 | 56.866 | 4.515760e+05 | 3.226270e+07 | 6509388.5 | 941306.5 | 9321870.0 | 15490132.0 | 32.88 | 61.2 | 47.7 | 33.3 |
| sp\|Q9GZT8\|NIF3L\_HUMAN | Q9GZT8 | NIF3L\_HUMAN | NIF3L1 | NIF3-like protein 1 | 377 | 41.968 | 6.334619e+05 | 3.340073e+07 | 6491764.5 | 26467460.0 | 296276.44 | 145227.19 | 44.83 | 57.3 | 30.8 | 14.6 |
| sp\|P09543\|CN37\_HUMAN | P09543 | CN37\_HUMAN | CNP | 2',3'-cyclic-nucleotide 3'-phosphodiesterase | 421 | 47.579 | 2.419515e+05 | 1.446308e+07 | 6473185.5 | 5404557.5 | 1213951.6 | 1371384.5 | 52.02 | 60.1 | 42.0 | 28.0 |
| sp\|P35579\|MYH9\_HUMAN | P35579 | MYH9\_HUMAN | MYH9 | Myosin-9 | 1960 | 226.532 | 7.065788e+04 | 2.010978e+07 | 6436780.0 | 11281428.0 | 1004060.3 | 1387510.4 | 67.91 | 74.5 | 43.2 | 26.7 |
| sp\|Q00796\|DHSO\_HUMAN | Q00796 | DHSO\_HUMAN | SORD | Sorbitol dehydrogenase | 357 | 38.325 | 3.910798e+05 | 1.883062e+07 | 6415717.5 | 11139172.0 | 798930.56 | 476802.0 | 70.59 | 72.0 | 36.1 | 14.3 |
| sp\|P22392\|NDKB\_HUMAN | P22392 | NDKB\_HUMAN | NME2 | Nucleoside diphosphate kinase B | 152 | 17.298 | 2.140515e+06 | 4.651910e+07 | 6411249.0 | 38954004.0 |  | 1153842.4 | 74.34 | 96.1 |  | 60.5 |
| sp\|Q99436\|PSB7\_HUMAN | Q99436 | PSB7\_HUMAN | PSMB7 | Proteasome subunit beta type-7 | 277 | 29.965 | 5.283824e+05 | 1.989206e+07 | 6396659.0 | 11157399.0 | 1223723.9 | 1114279.4 | 48.01 | 66.4 | 27.8 | 17.3 |
| sp\|P05089\|ARGI1\_HUMAN | P05089 | ARGI1\_HUMAN | ARG1 | Arginase-1 | 322 | 34.735 | 5.239418e+05 | 2.286480e+07 | 6385879.0 | 11365061.0 | 1705869.4 | 3407994.5 | 69.57 | 89.8 | 33.9 | 29.5 |
| sp\|O60256\|KPRB\_HUMAN | O60256 | KPRB\_HUMAN | PRPSAP2 | Phosphoribosyl pyrophosphate synthase-associated protein 2 | 369 | 40.926 | 3.976135e+05 | 2.044455e+07 | 6381302.5 | 12426922.0 | 972662.1 | 663664.4 | 65.31 | 81.3 | 40.7 | 14.4 |
| sp\|P15374\|UCHL3\_HUMAN | P15374 | UCHL3\_HUMAN | UCHL3 | Ubiquitin carboxyl-terminal hydrolase isozyme L3 | 230 | 26.183 | 6.707987e+05 | 2.206626e+07 | 6365778.5 | 13980857.0 | 1315472.9 | 404154.0 | 73.48 | 71.7 | 44.3 | 25.2 |
| sp\|Q9Y265\|RUVB1\_HUMAN | Q9Y265 | RUVB1\_HUMAN | RUVBL1 | RuvB-like 1 | 456 | 50.228 | 4.602300e+05 | 2.904276e+07 | 6345535.5 | 19866670.0 | 1843868.9 | 986690.3 | 75.0 | 71.5 | 39.3 | 16.4 |
| sp\|Q9Y230\|RUVB2\_HUMAN | Q9Y230 | RUVB2\_HUMAN | RUVBL2 | RuvB-like 2 | 463 | 51.157 | 2.376721e+05 | 1.527567e+07 | 6309559.5 | 5910921.0 | 2411630.0 | 643563.44 | 71.71 | 77.1 | 44.1 | 36.1 |
| sp\|P52888\|THOP1\_HUMAN | P52888 | THOP1\_HUMAN | THOP1 | Thimet oligopeptidase | 689 | 78.840 | 1.156225e+05 | 1.145265e+07 | 6268833.0 | 4927396.0 | 256424.9 |  | 52.54 | 56.3 | 25.1 |  |
| sp\|O76003\|GLRX3\_HUMAN | O76003 | GLRX3\_HUMAN | GLRX3 | Glutaredoxin-3 | 335 | 37.432 | 4.791895e+05 | 2.253552e+07 | 6223186.0 | 11674584.0 | 3868285.2 | 769460.9 | 54.03 | 76.7 | 36.4 | 23.3 |
| sp\|P15531\|NDKA\_HUMAN | P15531 | NDKA\_HUMAN | NME1 | Nucleoside diphosphate kinase A | 152 | 17.149 | 3.491942e+06 | 7.523553e+07 | 6219795.0 | 63796808.0 | 3869310.0 | 1349619.8 | 67.76 | 80.3 | 57.9 | 50.7 |
| sp\|Q5VW32\|BROX\_HUMAN | Q5VW32 | BROX\_HUMAN | BROX | BRO1 domain-containing protein BROX | 411 | 46.476 | 5.593444e+05 | 3.266068e+07 | 6212629.0 | 25004204.0 | 891464.6 | 552377.5 | 57.18 | 69.3 | 26.8 | 14.1 |
| sp\|P62333\|PRS10\_HUMAN | P62333 | PRS10\_HUMAN | PSMC6 | 26S proteasome regulatory subunit 10B | 389 | 44.173 | 3.931663e+05 | 2.181978e+07 | 6168575.0 | 13427247.0 | 1654044.8 | 569913.06 | 78.15 | 76.6 | 40.9 | 25.4 |
| sp\|P14324\|FPPS\_HUMAN | P14324 | FPPS\_HUMAN | FDPS | Farnesyl pyrophosphate synthase | 419 | 48.275 | 1.621544e+05 | 9.834859e+06 | 6100327.0 | 3338225.5 | 396306.56 |  | 24.82 | 46.1 | 22.9 |  |
| sp\|Q02094\|RHAG\_HUMAN | Q02094 | RHAG\_HUMAN | RHAG | Ammonium transporter Rh type A | 409 | 44.198 | 1.465142e+06 | 8.135787e+07 | 6076198.5 | 196123.56 | 38354296.0 | 36731252.0 | 14.67 | 20.5 | 9.0 | 2.2 |
| sp\|Q9H9G7\|AGO3\_HUMAN | Q9H9G7 | AGO3\_HUMAN | AGO3 | Protein argonaute-3 | 860 | 97.360 | 1.105275e+05 | 1.351973e+07 | 6053703.0 | 7466024.0 |  |  | 40.93 | 55.5 |  |  |
| sp\|P60953\|CDC42\_HUMAN | P60953 | CDC42\_HUMAN | CDC42 | Cell division control protein 42 homolog | 191 | 21.259 | 1.267957e+06 | 3.386604e+07 | 6040966.5 | 25379596.0 | 1271849.8 | 1173629.0 | 50.26 | 72.8 | 33.5 | 15.7 |
| sp\|Q9UBQ7\|GRHPR\_HUMAN | Q9UBQ7 | GRHPR\_HUMAN | GRHPR | Glyoxylate reductase/hydroxypyruvate reductase | 328 | 35.668 | 3.508627e+05 | 1.572292e+07 | 6030012.0 | 8583723.0 | 1011273.7 | 97907.89 | 52.74 | 56.4 | 22.3 | 18.6 |
| sp\|Q01518\|CAP1\_HUMAN | Q01518 | CAP1\_HUMAN | CAP1 | Adenylyl cyclase-associated protein 1 | 475 | 51.901 | 2.894109e+05 | 1.887156e+07 | 6028008.5 | 9961658.0 | 2263582.5 | 618308.1 | 58.74 | 76.4 | 51.6 | 19.8 |
| sp\|Q9ULA0\|DNPEP\_HUMAN | Q9ULA0 | DNPEP\_HUMAN | DNPEP | Aspartyl aminopeptidase | 485 | 53.410 | 1.478393e+05 | 9.920411e+06 | 5993947.0 | 3443624.0 | 482840.4 |  | 56.49 | 57.9 | 17.7 |  |
| sp\|Q13404\|UB2V1\_HUMAN | Q13404 | UB2V1\_HUMAN | UBE2V1 | Ubiquitin-conjugating enzyme E2 variant 1 | 147 | 16.495 | 1.420970e+06 | 2.944791e+07 | 5958708.0 | 11447725.0 | 9312276.0 | 2729198.0 | 73.47 | 95.2 | 72.1 | 61.2 |
| sp\|P14868\|SYDC\_HUMAN | P14868 | SYDC\_HUMAN | DARS1 | Aspartate--tRNA ligase, cytoplasmic | 501 | 57.136 | 1.799846e+05 | 1.291999e+07 | 5913625.0 | 5973888.0 | 782305.4 | 250175.72 | 71.86 | 76.6 | 55.5 | 24.6 |
| sp\|P61088\|UBE2N\_HUMAN | P61088 | UBE2N\_HUMAN | UBE2N | Ubiquitin-conjugating enzyme E2 N | 152 | 17.138 | 2.379188e+06 | 5.122785e+07 | 5895382.5 | 35537024.0 | 8094311.0 | 1701133.9 | 70.39 | 92.1 | 53.3 | 24.3 |
| sp\|Q86X55\|CARM1\_HUMAN | Q86X55 | CARM1\_HUMAN | CARM1 | Histone-arginine methyltransferase CARM1 | 608 | 65.854 | 2.435294e+05 | 2.014887e+07 | 5878418.5 | 9748921.0 | 3645851.2 | 875676.4 | 32.07 | 51.8 | 22.7 | 18.8 |
| sp\|O75223\|GGCT\_HUMAN | O75223 | GGCT\_HUMAN | GGCT | Gamma-glutamylcyclotransferase | 188 | 21.008 | 6.234582e+05 | 1.645543e+07 | 5872863.5 | 6545471.5 | 2703791.0 | 1333301.9 | 57.98 | 64.9 | 47.3 | 39.4 |
| sp\|Q13617\|CUL2\_HUMAN | Q13617 | CUL2\_HUMAN | CUL2 | Cullin-2 | 745 | 86.983 | 2.219918e+05 | 2.425987e+07 | 5871331.0 | 16504822.0 | 1405284.0 | 478434.6 | 67.38 | 69.7 | 38.1 | 20.0 |
| sp\|Q14C86\|GAPD1\_HUMAN | Q14C86 | GAPD1\_HUMAN | GAPVD1 | GTPase-activating protein and VPS9 domain-containing protein 1 | 1478 | 164.980 | 1.004701e+05 | 2.082501e+07 | 5865538.0 | 10995370.0 | 1369036.5 | 2595067.8 | 50.47 | 51.4 | 25.7 | 18.7 |
| sp\|P04424\|ARLY\_HUMAN | P04424 | ARLY\_HUMAN | ASL | Argininosuccinate lyase | 464 | 51.658 | 1.971043e+05 | 1.279237e+07 | 5796844.0 | 5617809.0 | 1044264.6 | 333448.06 | 55.17 | 59.3 | 32.1 | 30.8 |
| sp\|O00233\|PSMD9\_HUMAN | O00233 | PSMD9\_HUMAN | PSMD9 | 26S proteasome non-ATPase regulatory subunit 9 | 223 | 24.682 | 1.204858e+06 | 3.736227e+07 | 5750579.0 | 24706468.0 | 3169730.8 | 3735497.0 | 52.02 | 62.3 | 51.1 | 30.5 |
| sp\|P61224\|RAP1B\_HUMAN | P61224 | RAP1B\_HUMAN | RAP1B | Ras-related protein Rap-1b | 184 | 20.825 | 6.285184e+05 | 1.644448e+07 | 5750500.5 | 7850069.5 | 1391083.2 | 1452827.2 | 54.89 | 70.1 | 52.7 | 34.2 |
| sp\|P62834\|RAP1A\_HUMAN | P62834 | RAP1A\_HUMAN | RAP1A | Ras-related protein Rap-1A | 184 | 20.987 | 6.252263e+05 | 1.648560e+07 | 5750500.5 | 7850069.5 | 1432203.0 | 1452827.2 | 54.89 | 70.1 | 46.7 | 29.9 |
| sp\|P54727\|RD23B\_HUMAN | P54727 | RD23B\_HUMAN | RAD23B | UV excision repair protein RAD23 homolog B | 409 | 43.171 | 3.438098e+05 | 1.864780e+07 | 5739285.0 | 10003508.0 | 756141.0 | 2148862.8 | 52.08 | 60.1 | 28.4 | 18.1 |
| sp\|P49721\|PSB2\_HUMAN | P49721 | PSB2\_HUMAN | PSMB2 | Proteasome subunit beta type-2 | 201 | 22.836 | 4.082063e+05 | 1.171162e+07 | 5724901.0 | 4425631.0 | 1306819.8 | 254267.06 | 65.17 | 95.5 | 35.3 | 12.4 |
| sp\|Q3SY84\|K2C71\_HUMAN | Q3SY84 | K2C71\_HUMAN | KRT71 | Keratin, type II cytoskeletal 71 | 523 | 57.292 | 8.175950e+05 | 5.885039e+07 | 5717723.0 | 3407582.0 |  | 49725088.0 | 35.18 | 41.7 |  | 34.4 |
| sp\|Q86Y46\|K2C73\_HUMAN | Q86Y46 | K2C73\_HUMAN | KRT73 | Keratin, type II cytoskeletal 73 | 540 | 58.923 | 1.181948e+06 | 8.749847e+07 | 5717723.0 |  | 32055656.0 | 49725088.0 | 23.52 |  | 24.3 | 24.3 |
| sp\|Q7RTS7\|K2C74\_HUMAN | Q7RTS7 | K2C74\_HUMAN | KRT74 | Keratin, type II cytoskeletal 74 | 529 | 58.190 | 7.820916e+04 | 5.717723e+06 | 5717723.0 |  |  |  | 15.5 |  |  |  |
| sp\|P25787\|PSA2\_HUMAN | P25787 | PSA2\_HUMAN | PSMA2 | Proteasome subunit alpha type-2 | 234 | 25.899 | 6.028507e+05 | 1.961598e+07 | 5704423.0 | 13023930.0 | 665956.5 | 221667.58 | 74.79 | 67.1 | 24.4 | 15.4 |
| sp\|P63010\|AP2B1\_HUMAN | P63010 | AP2B1\_HUMAN | AP2B1 | AP-2 complex subunit beta | 937 | 104.553 | 2.178354e+05 | 2.861423e+07 | 5683632.0 | 6839894.5 | 12051772.0 | 4038936.2 | 63.71 | 51.8 | 35.3 | 25.8 |
| sp\|Q8IUI8\|CRLF3\_HUMAN | Q8IUI8 | CRLF3\_HUMAN | CRLF3 | Cytokine receptor-like factor 3 | 442 | 49.766 | 3.229534e+05 | 2.019249e+07 | 5679939.0 | 13142169.0 | 914092.9 | 456285.56 | 38.46 | 57.7 | 26.5 | 11.8 |
| sp\|Q9UNS2\|CSN3\_HUMAN | Q9UNS2 | CSN3\_HUMAN | COPS3 | COP9 signalosome complex subunit 3 | 423 | 47.873 | 4.067214e+05 | 2.446272e+07 | 5674658.0 | 15630839.0 | 2198290.5 | 958935.94 | 57.92 | 75.4 | 40.0 | 17.7 |
| sp\|Q92905\|CSN5\_HUMAN | Q92905 | CSN5\_HUMAN | COPS5 | COP9 signalosome complex subunit 5 | 334 | 37.579 | 4.248777e+05 | 2.005979e+07 | 5666422.5 | 13473294.0 | 643906.6 | 276163.97 | 70.96 | 72.5 | 40.7 | 21.3 |
| sp\|O14818\|PSA7\_HUMAN | O14818 | PSA7\_HUMAN | PSMA7 | Proteasome subunit alpha type-7 | 248 | 27.887 | 7.543833e+05 | 2.643084e+07 | 5665756.5 | 17356520.0 | 2022953.8 | 1385613.5 | 76.21 | 85.9 | 44.8 | 37.5 |
| sp\|P38919\|IF4A3\_HUMAN | P38919 | IF4A3\_HUMAN | EIF4A3 | Eukaryotic initiation factor 4A-III | 411 | 46.871 | 1.941612e+05 | 1.143362e+07 | 5620333.5 | 4701963.0 | 900387.4 | 210938.3 | 46.47 | 56.2 | 34.8 | 29.0 |
| sp\|O95782\|AP2A1\_HUMAN | O95782 | AP2A1\_HUMAN | AP2A1 | AP-2 complex subunit alpha-1 | 977 | 107.546 | 1.996605e+05 | 2.697761e+07 | 5515581.0 | 7511882.0 | 6195453.5 | 7754697.0 | 64.48 | 63.9 | 46.1 | 32.4 |
| sp\|P30626\|SORCN\_HUMAN | P30626 | SORCN\_HUMAN | SRI | Sorcin | 198 | 21.676 | 1.295481e+06 | 3.527990e+07 | 5493805.0 | 2817421.2 | 22221760.0 | 4746914.0 | 63.13 | 70.2 | 39.9 | 39.9 |
| sp\|P23381\|SYWC\_HUMAN | P23381 | SYWC\_HUMAN | WARS1 | Tryptophan--tRNA ligase, cytoplasmic | 471 | 53.165 | 3.478915e+05 | 2.323736e+07 | 5487315.5 | 16683552.0 | 783895.25 | 282596.53 | 63.27 | 66.5 | 32.1 | 22.1 |
| sp\|Q96F85\|CNRP1\_HUMAN | Q96F85 | CNRP1\_HUMAN | CNRIP1 | CB1 cannabinoid receptor-interacting protein 1 | 164 | 18.648 | 6.098317e+05 | 1.428760e+07 | 5485139.0 | 5850934.0 | 2490360.0 | 461170.9 | 68.9 | 92.7 | 30.5 | 26.2 |
| sp\|P63208\|SKP1\_HUMAN | P63208 | SKP1\_HUMAN | SKP1 | S-phase kinase-associated protein 1 | 163 | 18.658 | 9.880752e+05 | 2.316180e+07 | 5471807.5 | 11448018.0 | 5712030.0 | 529940.94 | 88.34 | 81.6 | 71.8 | 20.2 |
| sp\|Q9GZP4\|PITH1\_HUMAN | Q9GZP4 | PITH1\_HUMAN | PITHD1 | PITH domain-containing protein 1 | 211 | 24.178 | 5.125487e+05 | 1.556943e+07 | 5467950.0 | 7671008.0 | 2144820.2 | 285649.78 | 85.31 | 96.2 | 81.5 | 45.0 |
| sp\|P63000\|RAC1\_HUMAN | P63000 | RAC1\_HUMAN | RAC1 | Ras-related C3 botulinum toxin substrate 1 | 192 | 21.450 | 7.120935e+05 | 1.919029e+07 | 5433205.5 | 10973018.0 | 982150.8 | 1801913.1 | 45.31 | 63.0 | 34.4 | 21.9 |
| sp\|P00441\|SODC\_HUMAN | P00441 | SODC\_HUMAN | SOD1 | Superoxide dismutase [Cu-Zn] | 154 | 15.936 | 7.054729e+05 | 1.412462e+07 | 5414202.0 | 2977457.5 | 4467134.0 | 1265828.5 | 76.62 | 92.9 | 48.1 | 16.9 |
| sp\|P78356\|PI42B\_HUMAN | P78356 | PI42B\_HUMAN | PIP4K2B | Phosphatidylinositol 5-phosphate 4-kinase type-2 beta | 416 | 47.378 | 2.021407e+05 | 1.203227e+07 | 5406707.5 | 5571382.5 | 1054180.0 |  | 30.29 | 37.7 | 24.8 |  |
| sp\|P14550\|AK1A1\_HUMAN | P14550 | AK1A1\_HUMAN | AKR1A1 | Aldo-keto reductase family 1 member A1 | 325 | 36.573 | 3.631738e+05 | 1.668754e+07 | 5397035.5 | 10755131.0 | 361456.12 | 173912.55 | 72.31 | 73.2 | 49.5 | 23.7 |
| sp\|O00232\|PSD12\_HUMAN | O00232 | PSD12\_HUMAN | PSMD12 | 26S proteasome non-ATPase regulatory subunit 12 | 456 | 52.904 | 1.886616e+05 | 1.253976e+07 | 5391549.0 | 6244232.5 | 733298.6 | 170677.1 | 61.4 | 80.7 | 35.5 | 27.2 |
| sp\|Q07812\|BAX\_HUMAN | Q07812 | BAX\_HUMAN | BAX | Apoptosis regulator BAX | 192 | 21.184 | 2.880732e+05 | 7.667045e+06 | 5371909.0 | 2059024.6 | 97088.89 | 139022.08 | 25.52 | 43.8 | 18.8 | 23.4 |
| sp\|E9PAV3\|NACAM\_HUMAN | E9PAV3 | NACAM\_HUMAN | NACA | Nascent polypeptide-associated complex subunit alpha, muscle-specific form | 2078 | 205.422 | 2.985953e+04 | 7.706322e+06 | 5365415.0 | 1425256.6 | 825070.5 | 90579.59 | 10.11 | 56.6 | 37.5 | 26.6 |
| sp\|Q13162\|PRDX4\_HUMAN | Q13162 | PRDX4\_HUMAN | PRDX4 | Peroxiredoxin-4 | 271 | 30.540 | 4.467204e+05 | 1.714044e+07 | 5320439.0 | 4522546.0 | 5614861.5 | 1682594.1 | 47.6 | 60.9 | 25.8 | 12.2 |
| sp\|Q9BT78\|CSN4\_HUMAN | Q9BT78 | CSN4\_HUMAN | COPS4 | COP9 signalosome complex subunit 4 | 406 | 46.269 | 2.927384e+05 | 1.701715e+07 | 5258103.0 | 9654690.0 | 1166264.6 | 938096.44 | 86.7 | 84.5 | 55.4 | 29.6 |
| sp\|Q13564\|ULA1\_HUMAN | Q13564 | ULA1\_HUMAN | NAE1 | NEDD8-activating enzyme E1 regulatory subunit | 534 | 60.246 | 1.297497e+05 | 9.820910e+06 | 5211668.0 | 3981316.5 | 358818.53 | 269107.25 | 53.0 | 59.7 | 31.5 | 11.2 |
| sp\|Q9UNM6\|PSD13\_HUMAN | Q9UNM6 | PSD13\_HUMAN | PSMD13 | 26S proteasome non-ATPase regulatory subunit 13 | 376 | 42.945 | 2.671647e+05 | 1.441481e+07 | 5204604.0 | 4131408.0 | 4648920.0 | 429875.16 | 76.86 | 73.4 | 40.7 | 18.1 |
| sp\|P62820\|RAB1A\_HUMAN | P62820 | RAB1A\_HUMAN | RAB1A | Ras-related protein Rab-1A | 205 | 22.678 | 7.247247e+05 | 2.064881e+07 | 5203992.0 | 9752550.0 | 2720904.8 | 2971360.0 | 76.59 | 92.2 | 61.5 | 62.4 |
| sp\|Q9H0U4\|RAB1B\_HUMAN | Q9H0U4 | RAB1B\_HUMAN | RAB1B | Ras-related protein Rab-1B | 201 | 22.171 | 6.761234e+05 | 1.883338e+07 | 5203992.0 | 7876969.0 | 2720904.8 | 3031519.2 | 73.13 | 89.1 | 68.7 | 48.8 |
| sp\|P61201\|CSN2\_HUMAN | P61201 | CSN2\_HUMAN | COPS2 | COP9 signalosome complex subunit 2 | 443 | 51.597 | 2.265238e+05 | 1.468437e+07 | 5197575.5 | 8341562.0 | 1145233.6 |  | 61.85 | 71.3 | 46.0 |  |
| sp\|P61163\|ACTZ\_HUMAN | P61163 | ACTZ\_HUMAN | ACTR1A | Alpha-centractin | 376 | 42.614 | 2.633053e+05 | 1.409708e+07 | 5195735.0 | 8057070.0 | 844271.2 |  | 57.71 | 73.4 | 21.3 |  |
| sp\|P25786\|PSA1\_HUMAN | P25786 | PSA1\_HUMAN | PSMA1 | Proteasome subunit alpha type-1 | 263 | 29.556 | 6.346227e+05 | 2.356560e+07 | 5195290.0 | 16092063.0 | 1114830.2 | 1163412.0 | 65.02 | 68.1 | 57.0 | 36.9 |
| sp\|O76011\|KRT34\_HUMAN | O76011 | KRT34\_HUMAN | KRT34 | Keratin, type I cuticular Ha4 | 394 | 44.720 | 2.759817e+05 | 1.550598e+07 | 5184456.5 |  |  | 10321522.0 | 52.54 |  |  | 28.9 |
| sp\|Q14525\|KT33B\_HUMAN | Q14525 | KT33B\_HUMAN | KRT33B | Keratin, type I cuticular Ha3-II | 404 | 46.214 | 8.376297e+05 | 4.863431e+07 | 5184456.5 | 67810.48 |  | 43382048.0 | 63.86 | 37.1 |  | 44.6 |
| sp\|P61006\|RAB8A\_HUMAN | P61006 | RAB8A\_HUMAN | RAB8A | Ras-related protein Rab-8A | 207 | 23.668 | 8.060915e+05 | 2.396973e+07 | 5171470.5 | 13779641.0 | 2264108.8 | 2754506.0 | 64.73 | 73.9 | 49.3 | 33.3 |
| sp\|P43686\|PRS6B\_HUMAN | P43686 | PRS6B\_HUMAN | PSMC4 | 26S proteasome regulatory subunit 6B | 418 | 47.366 | 3.565717e+05 | 2.121929e+07 | 5148730.0 | 13888917.0 | 1311229.1 | 870409.8 | 62.2 | 75.1 | 36.8 | 24.6 |
| sp\|Q93008\|USP9X\_HUMAN | Q93008 | USP9X\_HUMAN | USP9X | Probable ubiquitin carboxyl-terminal hydrolase FAF-X | 2554 | 290.463 | 5.073921e+04 | 1.851619e+07 | 5138449.0 | 11162909.0 | 396684.47 | 1818150.6 | 48.55 | 48.9 | 22.1 | 13.9 |
| sp\|P55036\|PSMD4\_HUMAN | P55036 | PSMD4\_HUMAN | PSMD4 | 26S proteasome non-ATPase regulatory subunit 4 | 377 | 40.737 | 2.653814e+05 | 1.358241e+07 | 5135884.0 | 6948234.0 | 1175703.0 | 322584.12 | 48.81 | 45.6 | 35.3 | 26.8 |
| sp\|Q06323\|PSME1\_HUMAN | Q06323 | PSME1\_HUMAN | PSME1 | Proteasome activator complex subunit 1 | 249 | 28.723 | 9.778347e+05 | 3.528681e+07 | 5131020.0 | 25568960.0 | 3744843.2 | 841985.7 | 71.49 | 79.5 | 71.5 | 50.2 |
| sp\|Q14738\|2A5D\_HUMAN | Q14738 | 2A5D\_HUMAN | PPP2R5D | Serine/threonine-protein phosphatase 2A 56 kDa regulatory subunit delta isoform | 602 | 69.992 | 9.017330e+04 | 7.929458e+06 | 5115138.5 | 1878540.8 | 630122.6 | 305656.34 | 50.17 | 54.5 | 32.7 | 13.8 |
| sp\|Q15435\|PP1R7\_HUMAN | Q15435 | PP1R7\_HUMAN | PPP1R7 | Protein phosphatase 1 regulatory subunit 7 | 360 | 41.564 | 4.639441e+05 | 2.422703e+07 | 5104099.0 | 12397001.0 | 6429344.0 | 296583.94 | 72.22 | 70.6 | 35.8 | 30.6 |
| sp\|O43790\|KRT86\_HUMAN | O43790 | KRT86\_HUMAN | KRT86 | Keratin, type II cuticular Hb6 | 486 | 53.501 | 4.472237e+05 | 3.006103e+07 | 5091105.5 | 747914.5 | 1119901.1 | 23102108.0 | 51.85 | 56.2 | 40.3 | 41.2 |
| sp\|Q14533\|KRT81\_HUMAN | Q14533 | KRT81\_HUMAN | KRT81 | Keratin, type II cuticular Hb1 | 505 | 55.550 | 7.294758e+04 | 5.091106e+06 | 5091105.5 |  |  |  | 39.8 |  |  |  |
| sp\|Q15323\|K1H1\_HUMAN | Q15323 | K1H1\_HUMAN | KRT31 | Keratin, type I cuticular Ha1 | 416 | 47.237 | 8.328752e+05 | 4.942872e+07 | 5085388.0 | 162763.7 | 798520.56 | 43382048.0 | 61.78 | 26.4 | 45.4 | 37.0 |
| sp\|Q9Y315\|DEOC\_HUMAN | Q9Y315 | DEOC\_HUMAN | DERA | Deoxyribose-phosphate aldolase | 318 | 35.231 | 3.598480e+05 | 1.592800e+07 | 5068687.0 | 10577742.0 | 281568.03 |  | 83.33 | 77.7 | 46.5 |  |
| sp\|Q13057\|COASY\_HUMAN | Q13057 | COASY\_HUMAN | COASY | Bifunctional coenzyme A synthase | 564 | 62.329 | 1.836057e+05 | 1.437783e+07 | 5051413.0 | 8439683.0 | 674603.6 | 212133.4 | 42.91 | 60.5 | 20.6 | 11.3 |
| sp\|P01116\|RASK\_HUMAN | P01116 | RASK\_HUMAN | KRAS | GTPase KRas | 189 | 21.656 | 2.776885e+05 | 7.555327e+06 | 4980617.5 | 699677.8 | 1249420.2 | 625611.6 | 52.91 | 75.7 | 39.7 | 33.3 |
| sp\|P09417\|DHPR\_HUMAN | P09417 | DHPR\_HUMAN | QDPR | Dihydropteridine reductase | 244 | 25.790 | 1.058987e+06 | 3.431305e+07 | 4968300.0 | 29011832.0 | 300129.56 | 32787.69 | 58.2 | 73.8 | 36.1 | 18.9 |
| sp\|A6NCN2\|KR87P\_HUMAN | A6NCN2 | KR87P\_HUMAN | KRT87P | Putative keratin-87 protein | 255 | 28.050 | 1.408335e+05 | 4.963135e+06 | 4963135.0 |  |  |  | 63.53 |  |  |  |
| sp\|Q92530\|PSMF1\_HUMAN | Q92530 | PSMF1\_HUMAN | PSMF1 | Proteasome inhibitor PI31 subunit | 271 | 29.817 | 4.133856e+05 | 1.548590e+07 | 4935958.0 | 3361514.0 | 5514069.5 | 1674356.6 | 50.55 | 52.0 | 37.3 | 26.2 |
| sp\|Q02790\|FKBP4\_HUMAN | Q02790 | FKBP4\_HUMAN | FKBP4 | Peptidyl-prolyl cis-trans isomerase FKBP4 | 459 | 51.805 | 2.420883e+05 | 1.575660e+07 | 4918896.5 | 9618730.0 | 761478.1 | 457497.56 | 78.43 | 79.7 | 63.6 | 35.3 |
| sp\|Q15181\|IPYR\_HUMAN | Q15181 | IPYR\_HUMAN | PPA1 | Inorganic pyrophosphatase | 289 | 32.660 | 2.893240e+05 | 1.187183e+07 | 4914315.5 | 6242413.0 | 572178.1 | 142925.66 | 77.16 | 76.5 | 41.2 | 23.9 |
| sp\|Q9UQ16\|DYN3\_HUMAN | Q9UQ16 | DYN3\_HUMAN | DNM3 | Dynamin-3 | 869 | 97.746 | 1.043088e+05 | 1.280965e+07 | 4910744.0 | 5223937.0 | 2674966.5 |  | 17.61 | 36.4 | 26.4 |  |
| sp\|O43765\|SGTA\_HUMAN | O43765 | SGTA\_HUMAN | SGTA | Small glutamine-rich tetratricopeptide repeat-containing protein alpha | 313 | 34.063 | 3.345157e+05 | 1.431583e+07 | 4835061.5 | 7778149.0 | 1323899.5 | 378721.94 | 49.2 | 44.1 | 29.4 | 19.8 |
| sp\|P61026\|RAB10\_HUMAN | P61026 | RAB10\_HUMAN | RAB10 | Ras-related protein Rab-10 | 200 | 22.541 | 6.660463e+05 | 1.886231e+07 | 4819102.0 | 9072416.0 | 2874205.0 | 2096582.4 | 61.5 | 72.0 | 55.5 | 25.5 |
| sp\|P62937\|PPIA\_HUMAN | P62937 | PPIA\_HUMAN | PPIA | Peptidyl-prolyl cis-trans isomerase A | 165 | 18.012 | 3.703665e+06 | 8.381289e+07 | 4814122.0 | 70968440.0 | 6344661.0 | 1685668.4 | 89.7 | 93.9 | 66.1 | 40.0 |
| sp\|Q14558\|KPRA\_HUMAN | Q14558 | KPRA\_HUMAN | PRPSAP1 | Phosphoribosyl pyrophosphate synthase-associated protein 1 | 356 | 39.394 | 3.193402e+05 | 1.580523e+07 | 4811827.0 | 10237970.0 | 619308.94 | 136126.3 | 59.83 | 67.4 | 39.6 | 7.3 |
| sp\|Q13098\|CSN1\_HUMAN | Q13098 | CSN1\_HUMAN | GPS1 | COP9 signalosome complex subunit 1 | 491 | 55.537 | 2.493596e+05 | 1.739906e+07 | 4803903.5 | 10161832.0 | 1597964.8 | 835357.44 | 54.79 | 59.7 | 35.4 | 26.3 |
| sp\|O75874\|IDHC\_HUMAN | O75874 | IDHC\_HUMAN | IDH1 | Isocitrate dehydrogenase [NADP] cytoplasmic | 414 | 46.659 | 2.551289e+05 | 1.495589e+07 | 4747612.5 | 9012291.0 | 966412.44 | 229571.34 | 65.46 | 62.1 | 44.7 | 35.7 |
| sp\|Q14847\|LASP1\_HUMAN | Q14847 | LASP1\_HUMAN | LASP1 | LIM and SH3 domain protein 1 | 261 | 29.717 | 6.545430e+05 | 2.443770e+07 | 4741734.5 | 16628432.0 | 1680083.2 | 1387447.4 | 62.45 | 81.2 | 56.3 | 43.7 |
| sp\|P35813\|PPM1A\_HUMAN | P35813 | PPM1A\_HUMAN | PPM1A | Protein phosphatase 1A | 382 | 42.448 | 2.986485e+05 | 1.592703e+07 | 4741118.0 | 10418277.0 | 362354.28 | 405276.38 | 69.63 | 79.8 | 42.9 | 14.9 |
| sp\|P05091\|ALDH2\_HUMAN | P05091 | ALDH2\_HUMAN | ALDH2 | Aldehyde dehydrogenase, mitochondrial | 517 | 56.381 | 1.239893e+05 | 8.782826e+06 | 4730328.5 |  | 2237093.2 | 1815404.8 | 18.96 |  | 11.4 | 16.1 |
| sp\|Q9UPN7\|PP6R1\_HUMAN | Q9UPN7 | PP6R1\_HUMAN | PPP6R1 | Serine/threonine-protein phosphatase 6 regulatory subunit 1 | 881 | 96.724 | 9.586931e+04 | 1.165014e+07 | 4708732.0 | 5305451.0 | 926427.75 | 709526.0 | 50.85 | 40.6 | 18.2 | 12.8 |
| sp\|O00507\|USP9Y\_HUMAN | O00507 | USP9Y\_HUMAN | USP9Y | Probable ubiquitin carboxyl-terminal hydrolase FAF-Y | 2555 | 281.050 | 1.331656e+04 | 4.702110e+06 | 4702110.5 |  |  |  | 22.39 |  |  |  |
| sp\|Q709C8\|VP13C\_HUMAN | Q709C8 | VP13C\_HUMAN | VPS13C | Intermembrane lipid transfer protein VPS13C | 3753 | 422.390 | 1.390390e+04 | 7.378489e+06 | 4690990.5 | 1528851.2 | 418802.2 | 739845.3 | 36.18 | 44.4 | 19.4 | 15.4 |
| sp\|P63241\|IF5A1\_HUMAN | P63241 | IF5A1\_HUMAN | EIF5A | Eukaryotic translation initiation factor 5A-1 | 154 | 16.832 | 1.589917e+06 | 3.362229e+07 | 4669757.5 | 20392956.0 | 5081185.0 | 3478387.2 | 72.08 | 76.0 | 61.7 | 37.0 |
| sp\|Q14203\|DCTN1\_HUMAN | Q14203 | DCTN1\_HUMAN | DCTN1 | Dynactin subunit 1 | 1278 | 141.695 | 7.076884e+04 | 1.259835e+07 | 4660896.0 | 6542280.5 | 978831.4 | 416343.84 | 60.33 | 63.1 | 40.9 | 16.8 |
| sp\|Q9NRY5\|F1142\_HUMAN | Q9NRY5 | F1142\_HUMAN | FAM114A2 | Protein FAM114A2 | 505 | 55.468 | 3.318101e+05 | 2.312327e+07 | 4656769.5 | 10491918.0 | 4764443.5 | 3210142.8 | 76.04 | 71.7 | 48.7 | 33.5 |
| sp\|Q9NS86\|LANC2\_HUMAN | Q9NS86 | LANC2\_HUMAN | LANCL2 | LanC-like protein 2 | 450 | 50.854 | 1.266107e+05 | 8.089334e+06 | 4654869.0 | 2406937.5 | 755256.5 | 272271.25 | 45.56 | 59.6 | 28.4 | 15.6 |
| sp\|Q08211\|DHX9\_HUMAN | Q08211 | DHX9\_HUMAN | DHX9 | ATP-dependent RNA helicase A | 1270 | 140.958 | 5.299544e+04 | 9.385240e+06 | 4633933.0 | 3880833.5 | 395007.3 | 475466.3 | 45.59 | 52.6 | 33.3 | 14.9 |
| sp\|P55060\|XPO2\_HUMAN | P55060 | XPO2\_HUMAN | CSE1L | Exportin-2 | 971 | 110.417 | 8.674329e+04 | 1.203342e+07 | 4633172.5 | 5739489.5 | 398130.62 | 1262625.4 | 49.74 | 49.3 | 29.8 | 19.9 |
| sp\|O95352\|ATG7\_HUMAN | O95352 | ATG7\_HUMAN | ATG7 | Ubiquitin-like modifier-activating enzyme ATG7 | 703 | 77.960 | 6.738927e+04 | 6.600544e+06 | 4629137.5 | 1383316.0 | 408375.34 | 179715.0 | 44.95 | 60.3 | 33.1 | 16.5 |
| sp\|P51148\|RAB5C\_HUMAN | P51148 | RAB5C\_HUMAN | RAB5C | Ras-related protein Rab-5C | 216 | 23.483 | 8.421711e+05 | 2.484683e+07 | 4580884.0 | 16513824.0 | 2182341.0 | 1569785.1 | 70.83 | 75.0 | 51.9 | 40.3 |
| sp\|Q16881\|TRXR1\_HUMAN | Q16881 | TRXR1\_HUMAN | TXNRD1 | Thioredoxin reductase 1, cytoplasmic | 649 | 70.906 | 2.215601e+05 | 1.973748e+07 | 4579227.0 | 14322973.0 | 627352.1 | 207929.44 | 46.69 | 64.6 | 20.8 | 11.2 |
| sp\|P00387\|NB5R3\_HUMAN | P00387 | NB5R3\_HUMAN | CYB5R3 | NADH-cytochrome b5 reductase 3 | 301 | 34.235 | 3.544667e+05 | 1.524625e+07 | 4572216.0 | 7608521.0 | 1346279.2 | 1719231.5 | 64.12 | 68.4 | 18.3 | 19.3 |
| sp\|P17812\|PYRG1\_HUMAN | P17812 | PYRG1\_HUMAN | CTPS1 | CTP synthase 1 | 591 | 66.690 | 1.897940e+05 | 1.590231e+07 | 4571202.0 | 4747388.5 | 6176085.0 | 407637.53 | 56.85 | 55.3 | 29.6 | 17.3 |
| sp\|P06753\|TPM3\_HUMAN | P06753 | TPM3\_HUMAN | TPM3 | Tropomyosin alpha-3 chain | 285 | 32.950 | 2.305664e+06 | 9.544838e+07 | 4563954.5 | 35914488.0 | 48930144.0 | 6039794.0 | 47.37 | 80.7 | 71.9 | 43.9 |
| sp\|Q7L5N1\|CSN6\_HUMAN | Q7L5N1 | CSN6\_HUMAN | COPS6 | COP9 signalosome complex subunit 6 | 327 | 36.163 | 5.118321e+05 | 2.325461e+07 | 4556730.0 | 17325982.0 | 1371901.9 |  | 62.39 | 59.9 | 18.7 |  |
| sp\|P23588\|IF4B\_HUMAN | P23588 | IF4B\_HUMAN | EIF4B | Eukaryotic translation initiation factor 4B | 611 | 69.151 | 1.881843e+05 | 1.634930e+07 | 4550397.5 | 10175576.0 | 1205698.0 | 417623.88 | 42.23 | 68.4 | 40.6 | 33.2 |
| sp\|Q6ICL3\|TNG2\_HUMAN | Q6ICL3 | TNG2\_HUMAN | TANGO2 | Transport and Golgi organization protein 2 homolog | 276 | 30.937 | 3.666061e+05 | 1.424935e+07 | 4544899.0 | 8583300.0 | 743902.1 | 377249.72 | 72.83 | 77.2 | 26.1 | 19.2 |
| sp\|P55209\|NP1L1\_HUMAN | P55209 | NP1L1\_HUMAN | NAP1L1 | Nucleosome assembly protein 1-like 1 | 391 | 45.374 | 3.420888e+05 | 1.950128e+07 | 4537257.0 | 14147118.0 | 816903.25 |  | 30.95 | 33.5 | 15.9 |  |
| sp\|Q92930\|RAB8B\_HUMAN | Q92930 | RAB8B\_HUMAN | RAB8B | Ras-related protein Rab-8B | 207 | 23.584 | 4.878208e+05 | 1.445423e+07 | 4536072.5 | 7203410.5 |  | 2714747.5 | 57.97 | 64.7 |  | 35.3 |
| sp\|O43242\|PSMD3\_HUMAN | O43242 | PSMD3\_HUMAN | PSMD3 | 26S proteasome non-ATPase regulatory subunit 3 | 534 | 60.978 | 1.770839e+05 | 1.356655e+07 | 4528838.5 | 7073000.0 | 1439580.5 | 525128.6 | 70.6 | 73.0 | 54.1 | 40.8 |
| sp\|Q96BJ3\|AIDA\_HUMAN | Q96BJ3 | AIDA\_HUMAN | AIDA | Axin interactor, dorsalization-associated protein | 306 | 35.023 | 1.856661e+05 | 8.169644e+06 | 4501851.0 | 3396687.8 | 271105.0 |  | 57.84 | 55.2 | 34.6 |  |
| sp\|P30740\|ILEU\_HUMAN | P30740 | ILEU\_HUMAN | SERPINB1 | Leukocyte elastase inhibitor | 379 | 42.742 | 2.346807e+05 | 1.260229e+07 | 4492038.0 | 6926491.0 | 866512.75 | 317245.22 | 68.6 | 63.1 | 36.9 | 24.5 |
| sp\|P21399\|ACOHC\_HUMAN | P21399 | ACOHC\_HUMAN | ACO1 | Cytoplasmic aconitate hydratase | 889 | 98.399 | 8.579545e+04 | 1.060650e+07 | 4471899.0 | 5729550.5 | 285892.34 | 119158.25 | 55.57 | 56.1 | 25.6 | 13.7 |
| sp\|O60664\|PLIN3\_HUMAN | O60664 | PLIN3\_HUMAN | PLIN3 | Perilipin-3 | 434 | 47.075 | 2.637087e+05 | 1.559667e+07 | 4449137.0 | 7705689.0 | 2392776.0 | 1049069.0 | 64.98 | 87.3 | 40.1 | 33.6 |
| sp\|P61020\|RAB5B\_HUMAN | P61020 | RAB5B\_HUMAN | RAB5B | Ras-related protein Rab-5B | 215 | 23.707 | 5.699800e+05 | 1.697670e+07 | 4447932.0 | 8762820.0 | 1886328.4 | 1879621.9 | 66.98 | 68.8 | 47.9 | 44.2 |
| sp\|P14618\|KPYM\_HUMAN | P14618 | KPYM\_HUMAN | PKM | Pyruvate kinase PKM | 531 | 57.937 | 1.664883e+05 | 1.211873e+07 | 4440535.0 | 728085.4 | 2624246.0 | 4325859.0 | 48.59 | 70.4 | 40.7 | 27.5 |
| sp\|Q10567\|AP1B1\_HUMAN | Q10567 | AP1B1\_HUMAN | AP1B1 | AP-1 complex subunit beta-1 | 949 | 104.607 | 1.669810e+05 | 2.194548e+07 | 4440249.0 | 5432602.5 | 9422034.0 | 2650593.5 | 57.64 | 51.1 | 34.1 | 19.1 |
| sp\|Q15286\|RAB35\_HUMAN | Q15286 | RAB35\_HUMAN | RAB35 | Ras-related protein Rab-35 | 201 | 23.025 | 4.598224e+05 | 1.330169e+07 | 4419701.5 | 6006817.5 | 2276127.5 | 599047.2 | 73.63 | 61.7 | 38.3 | 23.9 |
| sp\|Q14651\|PLSI\_HUMAN | Q14651 | PLSI\_HUMAN | PLS1 | Plastin-1 | 629 | 70.253 | 8.214628e+04 | 7.250533e+06 | 4404728.5 | 2270807.5 | 529484.25 | 45512.52 | 42.45 | 44.0 | 25.8 | 13.8 |
| sp\|Q96CW1\|AP2M1\_HUMAN | Q96CW1 | AP2M1\_HUMAN | AP2M1 | AP-2 complex subunit mu | 435 | 49.655 | 4.030717e+05 | 2.514562e+07 | 4375247.0 | 7335798.5 | 11292725.0 | 2141854.5 | 60.69 | 72.4 | 61.6 | 36.3 |
| sp\|P28482\|MK01\_HUMAN | P28482 | MK01\_HUMAN | MAPK1 | Mitogen-activated protein kinase 1 | 360 | 41.390 | 1.731591e+05 | 9.004463e+06 | 4340167.0 | 3303050.8 | 833834.44 | 527411.2 | 68.89 | 58.6 | 32.5 | 20.8 |
| sp\|P78386\|KRT85\_HUMAN | P78386 | KRT85\_HUMAN | KRT85 | Keratin, type II cuticular Hb5 | 507 | 55.802 | 3.454847e+05 | 2.422120e+07 | 4312369.0 | 747914.5 | 1088308.5 | 18072610.0 | 52.27 | 57.2 | 29.4 | 39.4 |
| sp\|Q9H2M9\|RBGPR\_HUMAN | Q9H2M9 | RBGPR\_HUMAN | RAB3GAP2 | Rab3 GTPase-activating protein non-catalytic subunit | 1393 | 155.985 | 4.223591e+04 | 8.277170e+06 | 4309825.5 | 1330480.2 | 1960636.4 | 676228.0 | 31.51 | 42.4 | 23.5 | 16.9 |
| sp\|O75832\|PSD10\_HUMAN | O75832 | PSD10\_HUMAN | PSMD10 | 26S proteasome non-ATPase regulatory subunit 10 | 226 | 24.428 | 2.601055e+05 | 7.982789e+06 | 4294664.5 | 2418398.0 | 666441.56 | 603285.4 | 69.91 | 81.4 | 23.9 | 14.2 |
| sp\|P37802\|TAGL2\_HUMAN | P37802 | TAGL2\_HUMAN | TAGLN2 | Transgelin-2 | 199 | 22.391 | 8.208736e+05 | 2.309229e+07 | 4281598.0 | 16018234.0 | 2022247.2 | 770206.06 | 79.9 | 87.4 | 63.3 | 51.8 |
| sp\|Q9H4A4\|AMPB\_HUMAN | Q9H4A4 | AMPB\_HUMAN | RNPEP | Aminopeptidase B | 650 | 72.596 | 1.054108e+05 | 9.614244e+06 | 4268115.0 | 5097407.5 | 196514.94 | 52206.76 | 51.08 | 50.0 | 17.5 | 12.3 |
| sp\|Q96PL5\|ERMAP\_HUMAN | Q96PL5 | ERMAP\_HUMAN | ERMAP | Erythroid membrane-associated protein | 475 | 52.605 | 8.277584e+05 | 5.470761e+07 | 4262127.5 | 2199394.0 | 15179467.0 | 33066624.0 | 42.74 | 53.1 | 36.0 | 31.6 |
| sp\|Q14532\|K1H2\_HUMAN | Q14532 | K1H2\_HUMAN | KRT32 | Keratin, type I cuticular Ha2 | 448 | 50.343 | 7.647913e+05 | 4.837257e+07 | 4261943.5 |  | 728581.44 | 43382048.0 | 16.07 |  | 34.6 | 23.7 |
| sp\|O14618\|CCS\_HUMAN | O14618 | CCS\_HUMAN | CCS | Copper chaperone for superoxide dismutase | 274 | 29.041 | 4.272733e+05 | 1.558958e+07 | 4225848.0 | 6642169.0 | 2978967.2 | 1742597.5 | 54.74 | 62.4 | 24.1 | 14.6 |
| sp\|Q9H115\|SNAB\_HUMAN | Q9H115 | SNAB\_HUMAN | NAPB | Beta-soluble NSF attachment protein | 298 | 33.557 | 2.475661e+05 | 1.043738e+07 | 4218752.0 | 5297312.5 | 921313.8 |  | 47.65 | 85.6 | 41.3 |  |
| sp\|Q07960\|RHG01\_HUMAN | Q07960 | RHG01\_HUMAN | ARHGAP1 | Rho GTPase-activating protein 1 | 439 | 50.436 | 1.780694e+05 | 1.128358e+07 | 4206707.0 | 5072134.5 | 1830213.0 | 174530.5 | 64.69 | 67.0 | 41.7 | 21.9 |
| sp\|O76015\|KRT38\_HUMAN | O76015 | KRT38\_HUMAN | KRT38 | Keratin, type I cuticular Ha8 | 456 | 50.160 | 6.647810e+04 | 4.189414e+06 | 4189414.0 |  |  |  | 23.68 |  |  |  |
| sp\|P55263\|ADK\_HUMAN | P55263 | ADK\_HUMAN | ADK | Adenosine kinase | 362 | 40.545 | 2.376512e+05 | 1.210583e+07 | 4184168.5 | 7246050.5 | 518537.06 | 157073.08 | 71.82 | 75.7 | 30.7 | 14.6 |
| sp\|P20339\|RAB5A\_HUMAN | P20339 | RAB5A\_HUMAN | RAB5A | Ras-related protein Rab-5A | 215 | 23.659 | 3.874480e+05 | 1.151667e+07 | 4172507.8 | 4298184.5 | 1881887.8 | 1164090.4 | 72.56 | 71.2 | 46.0 | 44.7 |
| sp\|Q96NA2\|RILP\_HUMAN | Q96NA2 | RILP\_HUMAN | RILP | Rab-interacting lysosomal protein | 401 | 44.200 | 2.911837e+05 | 1.616987e+07 | 4149220.5 | 9449541.0 | 2100865.0 | 470240.12 | 43.64 | 53.1 | 35.4 | 25.2 |
| sp\|O14964\|HGS\_HUMAN | O14964 | HGS\_HUMAN | HGS | Hepatocyte growth factor-regulated tyrosine kinase substrate | 777 | 86.192 | 1.048848e+05 | 1.135786e+07 | 4125592.8 | 5063873.0 | 1068417.5 | 1099978.0 | 38.61 | 38.5 | 26.8 | 15.3 |
| sp\|P67936\|TPM4\_HUMAN | P67936 | TPM4\_HUMAN | TPM4 | Tropomyosin alpha-4 chain | 248 | 28.522 | 1.779353e+06 | 6.376162e+07 | 4120050.0 | 31318746.0 | 28322820.0 |  | 25.81 | 51.6 | 55.6 |  |
| sp\|Q13616\|CUL1\_HUMAN | Q13616 | CUL1\_HUMAN | CUL1 | Cullin-1 | 776 | 89.679 | 1.223070e+05 | 1.378032e+07 | 4100039.5 | 7776694.5 | 1332267.0 | 571318.6 | 74.87 | 81.2 | 38.7 | 26.4 |
| sp\|O95336\|6PGL\_HUMAN | O95336 | 6PGL\_HUMAN | PGLS | 6-phosphogluconolactonase | 258 | 27.547 | 5.380422e+05 | 1.862121e+07 | 4089910.5 | 13567422.0 | 644607.75 | 319267.16 | 66.67 | 69.8 | 47.3 | 28.3 |
| sp\|Q86X76\|NIT1\_HUMAN | Q86X76 | NIT1\_HUMAN | NIT1 | Deaminated glutathione amidase | 327 | 35.896 | 1.983200e+05 | 8.943959e+06 | 4088688.5 | 4410342.0 | 354307.06 | 90621.37 | 48.32 | 53.5 | 37.0 | 15.3 |
| sp\|Q9NR45\|SIAS\_HUMAN | Q9NR45 | SIAS\_HUMAN | NANS | Sialic acid synthase | 359 | 40.308 | 1.798414e+05 | 9.107478e+06 | 4053094.0 | 4770099.5 | 224249.22 | 60035.56 | 57.38 | 63.2 | 35.4 | 17.5 |
| sp\|Q13126\|MTAP\_HUMAN | Q13126 | MTAP\_HUMAN | MTAP | S-methyl-5'-thioadenosine phosphorylase | 283 | 31.236 | 4.815346e+05 | 1.889731e+07 | 4045557.0 | 13753931.0 | 678200.9 | 419623.94 | 82.69 | 85.5 | 51.2 | 23.7 |
| sp\|Q9UJU6\|DBNL\_HUMAN | Q9UJU6 | DBNL\_HUMAN | DBNL | Drebrin-like protein | 430 | 48.207 | 2.305336e+05 | 1.396244e+07 | 4009681.2 | 8917329.0 | 655982.6 | 379451.38 | 63.95 | 62.3 | 37.9 | 30.5 |
| sp\|P15121\|ALDR\_HUMAN | P15121 | ALDR\_HUMAN | AKR1B1 | Aldo-keto reductase family 1 member B1 | 316 | 35.853 | 1.209938e+05 | 5.450115e+06 | 4005177.2 | 1337121.2 | 107816.16 |  | 56.96 | 68.7 | 18.4 |  |
| sp\|P47755\|CAZA2\_HUMAN | P47755 | CAZA2\_HUMAN | CAPZA2 | F-actin-capping protein subunit alpha-2 | 286 | 32.949 | 3.247661e+05 | 1.344405e+07 | 4000432.2 | 7243070.0 | 1888300.4 | 312245.16 | 77.27 | 86.7 | 16.1 | 18.9 |
| sp\|P30711\|GSTT1\_HUMAN | P30711 | GSTT1\_HUMAN | GSTT1 | Glutathione S-transferase theta-1 | 240 | 27.335 | 1.749804e+05 | 6.009325e+06 | 4000326.8 | 2008998.0 |  |  | 40.0 | 65.0 |  |  |
| sp\|Q8IXQ3\|CI040\_HUMAN | Q8IXQ3 | CI040\_HUMAN | C9orf40 | Uncharacterized protein C9orf40 | 194 | 21.063 | 1.998961e+05 | 5.289831e+06 | 3996947.0 | 439674.3 | 507529.12 | 345680.1 | 60.31 | 62.9 | 35.6 | 29.4 |
| sp\|Q99808\|S29A1\_HUMAN | Q99808 | S29A1\_HUMAN | SLC29A1 | Equilibrative nucleoside transporter 1 | 456 | 50.219 | 5.427680e+05 | 3.424518e+07 | 3994509.5 | 2029901.5 | 9231212.0 | 18989558.0 | 25.0 | 29.8 | 22.4 | 21.9 |
| sp\|P50148\|GNAQ\_HUMAN | P50148 | GNAQ\_HUMAN | GNAQ | Guanine nucleotide-binding protein G(q) subunit alpha | 359 | 42.142 | 1.189867e+05 | 6.299858e+06 | 3956907.8 | 655276.25 | 795580.8 | 892092.7 | 55.15 | 42.3 | 39.0 | 27.6 |
| sp\|Q93009\|UBP7\_HUMAN | Q93009 | UBP7\_HUMAN | USP7 | Ubiquitin carboxyl-terminal hydrolase 7 | 1102 | 128.302 | 6.581045e+04 | 1.060829e+07 | 3956160.0 | 5833948.5 | 541248.94 | 276934.4 | 66.79 | 64.6 | 27.8 | 22.6 |
| sp\|Q96FW1\|OTUB1\_HUMAN | Q96FW1 | OTUB1\_HUMAN | OTUB1 | Ubiquitin thioesterase OTUB1 | 271 | 31.284 | 2.369156e+05 | 9.311789e+06 | 3939952.8 | 4213379.0 | 1003541.3 | 154916.25 | 58.67 | 49.1 | 49.8 | 11.4 |
| sp\|Q9BQA1\|MEP50\_HUMAN | Q9BQA1 | MEP50\_HUMAN | WDR77 | Methylosome protein WDR77 | 342 | 36.724 | 1.754885e+05 | 8.096846e+06 | 3936699.5 | 3627951.5 | 413129.44 | 119065.89 | 57.89 | 57.6 | 18.1 | 12.0 |
| sp\|Q9NPQ8\|RIC8A\_HUMAN | Q9NPQ8 | RIC8A\_HUMAN | RIC8A | Synembryn-A | 531 | 59.710 | 2.422561e+05 | 1.817352e+07 | 3897538.0 | 12484645.0 | 1234711.1 | 556622.6 | 68.36 | 65.5 | 31.8 | 20.2 |
| sp\|O94973\|AP2A2\_HUMAN | O94973 | AP2A2\_HUMAN | AP2A2 | AP-2 complex subunit alpha-2 | 939 | 103.960 | 1.285195e+05 | 1.678620e+07 | 3884833.8 | 5039764.5 | 2992837.0 | 4868767.0 | 62.3 | 49.9 | 30.1 | 24.4 |
| sp\|O76054\|S14L2\_HUMAN | O76054 | S14L2\_HUMAN | SEC14L2 | SEC14-like protein 2 | 403 | 46.145 | 1.495984e+05 | 8.672986e+06 | 3868700.5 | 3643755.0 | 871099.0 | 289431.88 | 54.59 | 56.1 | 18.1 | 13.6 |
| sp\|Q86YZ3\|HORN\_HUMAN | Q86YZ3 | HORN\_HUMAN | HRNR | Hornerin | 2850 | 282.390 | 7.741965e+04 | 2.746741e+07 | 3852162.8 | 11848194.0 | 4842031.0 | 6925021.0 | 46.49 | 57.9 | 34.5 | 29.7 |
| sp\|Q7Z4W1\|DCXR\_HUMAN | Q7Z4W1 | DCXR\_HUMAN | DCXR | L-xylulose reductase | 244 | 25.913 | 4.339880e+05 | 1.412904e+07 | 3845283.0 | 9095597.0 | 850372.6 | 337785.44 | 69.67 | 68.0 | 47.1 | 27.0 |
| sp\|Q8WVM8\|SCFD1\_HUMAN | Q8WVM8 | SCFD1\_HUMAN | SCFD1 | Sec1 family domain-containing protein 1 | 642 | 72.380 | 1.302112e+05 | 1.184089e+07 | 3840403.5 | 4699416.5 | 2617275.0 | 683790.75 | 51.56 | 67.4 | 37.5 | 17.6 |
| sp\|P54577\|SYYC\_HUMAN | P54577 | SYYC\_HUMAN | YARS1 | Tyrosine--tRNA ligase, cytoplasmic | 528 | 59.143 | 1.167894e+05 | 8.678085e+06 | 3815563.8 | 4320673.0 | 382502.72 | 159345.44 | 71.97 | 74.1 | 39.0 | 32.8 |
| sp\|Q5VT06\|CE350\_HUMAN | Q5VT06 | CE350\_HUMAN | CEP350 | Centrosome-associated protein 350 | 3117 | 342.870 | 8.848036e+03 | 3.811479e+06 | 3811479.0 |  |  |  | 1.48 |  |  |  |
| sp\|P53680\|AP2S1\_HUMAN | P53680 | AP2S1\_HUMAN | AP2S1 | AP-2 complex subunit sigma | 142 | 17.018 | 3.317712e+05 | 7.093561e+06 | 3773474.8 | 1055719.0 | 1451617.0 | 812750.5 | 18.31 | 43.7 | 35.9 | 25.4 |
| sp\|Q9UHY1\|NRBP\_HUMAN | Q9UHY1 | NRBP\_HUMAN | NRBP1 | Nuclear receptor-binding protein | 535 | 59.845 | 8.071802e+04 | 6.068977e+06 | 3757922.2 | 1833409.0 | 395724.16 | 81922.06 | 27.85 | 44.7 | 17.0 | 12.3 |
| sp\|P53990\|IST1\_HUMAN | P53990 | IST1\_HUMAN | IST1 | IST1 homolog | 364 | 39.751 | 2.329029e+05 | 1.163162e+07 | 3752298.5 | 5296026.0 | 1743094.8 | 840196.25 | 44.23 | 39.0 | 25.5 | 19.0 |
| sp\|P20340\|RAB6A\_HUMAN | P20340 | RAB6A\_HUMAN | RAB6A | Ras-related protein Rab-6A | 208 | 23.593 | 3.774364e+05 | 1.118778e+07 | 3750183.2 | 5308671.0 | 1465939.1 | 662990.06 | 69.23 | 61.5 | 41.8 | 34.6 |
| sp\|Q9NT62\|ATG3\_HUMAN | Q9NT62 | ATG3\_HUMAN | ATG3 | Ubiquitin-like-conjugating enzyme ATG3 | 314 | 35.864 | 1.719069e+05 | 7.745851e+06 | 3748196.5 | 2840324.8 | 876251.8 | 281078.25 | 42.99 | 51.6 | 27.7 | 12.4 |
| sp\|Q9UNN5\|FAF1\_HUMAN | Q9UNN5 | FAF1\_HUMAN | FAF1 | FAS-associated factor 1 | 650 | 73.954 | 1.297984e+05 | 1.206003e+07 | 3726870.5 | 6791222.0 | 1073591.5 | 468342.6 | 38.0 | 46.9 | 19.8 | 9.5 |
| sp\|Q86UX7\|URP2\_HUMAN | Q86UX7 | URP2\_HUMAN | FERMT3 | Fermitin family homolog 3 | 667 | 75.953 | 1.380831e+05 | 1.317658e+07 | 3699597.5 | 7477306.0 | 1644190.4 | 355484.53 | 65.97 | 63.3 | 48.1 | 14.4 |
| sp\|P08670\|VIME\_HUMAN | P08670 | VIME\_HUMAN | VIM | Vimentin | 466 | 53.652 | 2.449432e+05 | 1.651081e+07 | 3696642.2 | 255391.6 | 871478.5 | 11687300.0 | 35.19 | 35.4 | 47.9 | 44.6 |
| sp\|P55010\|IF5\_HUMAN | P55010 | IF5\_HUMAN | EIF5 | Eukaryotic translation initiation factor 5 | 431 | 49.223 | 1.643735e+05 | 1.016523e+07 | 3687438.0 | 5419355.5 | 729656.5 | 328776.4 | 49.88 | 48.7 | 21.3 | 19.5 |
| sp\|Q6IA69\|NADE\_HUMAN | Q6IA69 | NADE\_HUMAN | NADSYN1 | Glutamine-dependent NAD(+) synthetase | 706 | 79.285 | 6.175446e+04 | 6.151436e+06 | 3686562.5 | 1908764.2 | 121001.45 | 435107.62 | 31.59 | 33.7 | 17.4 | 16.1 |
| sp\|Q9BV20\|MTNA\_HUMAN | Q9BV20 | MTNA\_HUMAN | MRI1 | Methylthioribose-1-phosphate isomerase | 369 | 39.150 | 2.180034e+05 | 1.072290e+07 | 3658618.5 | 5950882.5 | 891060.2 | 222339.1 | 51.49 | 77.5 | 25.2 | 16.8 |
| sp\|P09493\|TPM1\_HUMAN | P09493 | TPM1\_HUMAN | TPM1 | Tropomyosin alpha-1 chain | 284 | 32.709 | 1.272782e+06 | 5.230441e+07 | 3647544.0 | 24667706.0 | 21928876.0 | 2060288.8 | 40.14 | 78.5 | 66.2 | 43.3 |
| sp\|Q9NTJ4\|MA2C1\_HUMAN | Q9NTJ4 | MA2C1\_HUMAN | MAN2C1 | Alpha-mannosidase 2C1 | 1040 | 115.835 | 5.202062e+04 | 7.570638e+06 | 3642027.2 | 3745656.0 | 120433.14 | 62522.016 | 36.15 | 45.1 | 14.7 | 9.0 |
| sp\|P30085\|KCY\_HUMAN | P30085 | KCY\_HUMAN | CMPK1 | UMP-CMP kinase | 196 | 22.222 | 5.287727e+05 | 1.476282e+07 | 3640757.2 | 10006876.0 | 1065026.1 | 50161.56 | 68.37 | 78.1 | 37.8 | 22.4 |
| sp\|P30046\|DOPD\_HUMAN | P30046 | DOPD\_HUMAN | DDT | D-dopachrome decarboxylase | 118 | 12.712 | 1.493466e+06 | 2.385208e+07 | 3639705.2 | 18493536.0 | 1718838.4 |  | 50.0 | 100.0 | 60.2 |  |
| sp\|Q58FG1\|HS904\_HUMAN | Q58FG1 | HS904\_HUMAN | HSP90AA4P | Putative heat shock protein HSP 90-alpha A4 | 418 | 47.712 | 7.258068e+04 | 4.350767e+06 | 3630915.5 | 719851.1 |  |  | 21.29 | 29.4 |  |  |
| sp\|Q9NRW1\|RAB6B\_HUMAN | Q9NRW1 | RAB6B\_HUMAN | RAB6B | Ras-related protein Rab-6B | 208 | 23.462 | 3.258277e+05 | 9.604400e+06 | 3630391.8 | 4080370.5 | 1198299.2 | 695338.56 | 60.58 | 61.5 | 36.1 | 33.2 |
| sp\|P52566\|GDIR2\_HUMAN | P52566 | GDIR2\_HUMAN | ARHGDIB | Rho GDP-dissociation inhibitor 2 | 201 | 22.988 | 5.110826e+05 | 1.476078e+07 | 3627947.2 | 9207535.0 | 1452135.9 | 473165.06 | 67.16 | 91.5 | 59.7 | 35.3 |
| sp\|Q04446\|GLGB\_HUMAN | Q04446 | GLGB\_HUMAN | GBE1 | 1,4-alpha-glucan-branching enzyme | 702 | 80.474 | 7.802910e+04 | 7.889134e+06 | 3622105.0 | 3486345.8 | 531799.9 | 248883.67 | 44.3 | 40.9 | 19.8 | 13.5 |
| sp\|P61086\|UBE2K\_HUMAN | P61086 | UBE2K\_HUMAN | UBE2K | Ubiquitin-conjugating enzyme E2 K | 200 | 22.407 | 3.291220e+05 | 9.265265e+06 | 3621365.2 | 4332793.0 | 970880.25 | 340226.1 | 82.0 | 63.5 | 38.5 | 30.0 |
| sp\|Q15631\|TSN\_HUMAN | Q15631 | TSN\_HUMAN | TSN | Translin | 228 | 26.183 | 2.027279e+05 | 6.668836e+06 | 3600076.0 | 2541357.8 | 310443.7 | 216958.02 | 46.05 | 68.4 | 25.0 | 14.5 |
| sp\|Q96DG6\|CMBL\_HUMAN | Q96DG6 | CMBL\_HUMAN | CMBL | Carboxymethylenebutenolidase homolog | 245 | 28.048 | 3.056661e+05 | 1.077125e+07 | 3598859.5 | 6023697.0 | 610432.1 | 538265.9 | 51.43 | 70.2 | 38.8 | 28.6 |
| sp\|Q04323\|UBXN1\_HUMAN | Q04323 | UBXN1\_HUMAN | UBXN1 | UBX domain-containing protein 1 | 297 | 33.325 | 1.936221e+05 | 8.106664e+06 | 3597023.0 | 3466947.8 | 847611.94 | 195081.22 | 53.54 | 72.1 | 36.4 | 33.0 |
| sp\|P49327\|FAS\_HUMAN | P49327 | FAS\_HUMAN | FASN | Fatty acid synthase | 2511 | 273.427 | 2.319783e+04 | 7.969039e+06 | 3594166.0 | 3273005.5 | 642922.0 | 458945.3 | 58.26 | 54.7 | 27.1 | 17.3 |
| sp\|P48147\|PPCE\_HUMAN | P48147 | PPCE\_HUMAN | PREP | Prolyl endopeptidase | 710 | 80.700 | 9.137838e+04 | 9.264759e+06 | 3555376.0 | 5579750.0 | 79206.79 | 50426.418 | 66.9 | 64.9 | 14.5 | 8.6 |
| sp\|Q9Y570\|PPME1\_HUMAN | Q9Y570 | PPME1\_HUMAN | PPME1 | Protein phosphatase methylesterase 1 | 386 | 42.315 | 3.492001e+05 | 1.856461e+07 | 3544070.5 | 13030798.0 | 1369444.5 | 620298.5 | 50.0 | 60.4 | 35.0 | 20.5 |
| sp\|Q13336\|UT1\_HUMAN | Q13336 | UT1\_HUMAN | SLC14A1 | Urea transporter 1 | 389 | 42.528 | 1.351151e+06 | 7.219314e+07 | 3525818.2 | 1525443.0 | 18502256.0 | 48639620.0 | 19.54 | 14.4 | 16.7 | 20.3 |
| sp\|Q93034\|CUL5\_HUMAN | Q93034 | CUL5\_HUMAN | CUL5 | Cullin-5 | 780 | 90.955 | 6.834777e+04 | 7.810306e+06 | 3502916.5 | 3189596.8 | 890029.0 | 227764.03 | 56.54 | 57.8 | 33.6 | 22.4 |
| sp\|Q15149\|PLEC\_HUMAN | Q15149 | PLEC\_HUMAN | PLEC | Plectin | 4684 | 531.791 | 1.257672e+04 | 8.402832e+06 | 3489630.8 | 4017359.0 | 424504.47 | 471337.53 | 3.97 | 32.4 | 22.7 | 18.1 |
| sp\|Q9H8M7\|MINY3\_HUMAN | Q9H8M7 | MINY3\_HUMAN | MINDY3 | Ubiquitin carboxyl-terminal hydrolase MINDY-3 | 445 | 49.725 | 8.610910e+04 | 5.379488e+06 | 3484596.2 | 1569687.9 | 247342.27 | 77862.07 | 27.19 | 44.7 | 13.3 | 4.9 |
| sp\|P35606\|COPB2\_HUMAN | P35606 | COPB2\_HUMAN | COPB2 | Coatomer subunit beta' | 906 | 102.487 | 3.110172e+04 | 4.004704e+06 | 3483319.5 | 347339.16 | 125063.5 | 48981.47 | 19.21 | 31.7 | 17.3 | 8.4 |
| sp\|Q92539\|LPIN2\_HUMAN | Q92539 | LPIN2\_HUMAN | LPIN2 | Phosphatidate phosphatase LPIN2 | 896 | 99.399 | 1.189882e+05 | 1.485946e+07 | 3474347.8 | 9696372.0 | 819868.0 | 868875.75 | 48.66 | 43.8 | 25.9 | 21.7 |
| sp\|P61106\|RAB14\_HUMAN | P61106 | RAB14\_HUMAN | RAB14 | Ras-related protein Rab-14 | 215 | 23.897 | 4.040111e+05 | 1.212980e+07 | 3468107.0 | 5277042.0 | 2463011.8 | 921643.4 | 73.02 | 92.6 | 54.9 | 30.7 |
| sp\|Q92973\|TNPO1\_HUMAN | Q92973 | TNPO1\_HUMAN | TNPO1 | Transportin-1 | 898 | 102.355 | 6.074658e+04 | 7.811744e+06 | 3436105.0 | 4102338.5 | 273300.72 |  | 45.99 | 38.3 | 13.6 |  |
| sp\|P52294\|IMA5\_HUMAN | P52294 | IMA5\_HUMAN | KPNA1 | Importin subunit alpha-5 | 538 | 60.222 | 1.617644e+05 | 1.223926e+07 | 3416660.0 | 7339322.0 | 1128352.5 | 354927.1 | 30.86 | 36.8 | 22.1 | 18.0 |
| sp\|P43353\|AL3B1\_HUMAN | P43353 | AL3B1\_HUMAN | ALDH3B1 | Aldehyde dehydrogenase family 3 member B1 | 468 | 51.840 | 5.354201e+04 | 3.487199e+06 | 3415147.2 | 72052.29 |  |  | 13.68 | 34.4 |  |  |
| sp\|Q9P2T1\|GMPR2\_HUMAN | Q9P2T1 | GMPR2\_HUMAN | GMPR2 | GMP reductase 2 | 348 | 37.874 | 1.628458e+05 | 7.748808e+06 | 3390381.2 | 3672735.0 | 194064.78 | 491626.66 | 59.48 | 74.7 | 33.9 | 19.3 |
| sp\|P46060\|RAGP1\_HUMAN | P46060 | RAGP1\_HUMAN | RANGAP1 | Ran GTPase-activating protein 1 | 587 | 63.542 | 1.072200e+05 | 8.559610e+06 | 3375937.0 | 3093378.8 | 1576175.1 | 514118.84 | 54.17 | 68.7 | 38.3 | 14.5 |
| sp\|Q99627\|CSN8\_HUMAN | Q99627 | CSN8\_HUMAN | COPS8 | COP9 signalosome complex subunit 8 | 209 | 23.226 | 1.776310e+05 | 5.183348e+06 | 3375658.0 | 1469894.2 | 337796.03 |  | 51.67 | 93.3 | 37.3 |  |
| sp\|Q00577\|PURA\_HUMAN | Q00577 | PURA\_HUMAN | PURA | Transcriptional activator protein Pur-alpha | 322 | 34.911 | 3.297409e+05 | 1.446280e+07 | 3368513.0 | 8463077.0 | 1999875.6 | 631330.9 | 55.28 | 50.6 | 35.1 | 13.4 |
| sp\|Q9BRF8\|CPPED\_HUMAN | Q9BRF8 | CPPED\_HUMAN | CPPED1 | Serine/threonine-protein phosphatase CPPED1 | 314 | 35.548 | 1.737857e+05 | 7.761514e+06 | 3356002.2 | 3483303.2 | 686849.56 | 235358.95 | 55.1 | 65.3 | 24.8 | 21.3 |
| sp\|P05198\|IF2A\_HUMAN | P05198 | IF2A\_HUMAN | EIF2S1 | Eukaryotic translation initiation factor 2 subunit 1 | 315 | 36.112 | 2.286486e+05 | 1.037379e+07 | 3349217.5 | 6023959.0 | 816573.25 | 184037.34 | 73.33 | 81.6 | 56.2 | 31.4 |
| sp\|O00487\|PSDE\_HUMAN | O00487 | PSDE\_HUMAN | PSMD14 | 26S proteasome non-ATPase regulatory subunit 14 | 310 | 34.577 | 5.068186e+05 | 2.201694e+07 | 3338990.5 | 16260020.0 | 1834665.8 | 583267.1 | 59.03 | 91.0 | 35.2 | 29.0 |
| sp\|Q06124\|PTN11\_HUMAN | Q06124 | PTN11\_HUMAN | PTPN11 | Tyrosine-protein phosphatase non-receptor type 11 | 593 | 68.011 | 1.608396e+05 | 1.374325e+07 | 3328132.0 | 9413926.0 | 679019.2 | 322168.47 | 70.83 | 69.5 | 36.1 | 19.1 |
| sp\|O00743\|PPP6\_HUMAN | O00743 | PPP6\_HUMAN | PPP6C | Serine/threonine-protein phosphatase 6 catalytic subunit | 305 | 35.144 | 2.028418e+05 | 8.956241e+06 | 3326862.5 | 4668479.0 | 960899.3 |  | 48.85 | 60.0 | 20.0 |  |
| sp\|P25685\|DNJB1\_HUMAN | P25685 | DNJB1\_HUMAN | DNAJB1 | DnaJ homolog subfamily B member 1 | 340 | 38.044 | 3.452148e+05 | 1.650034e+07 | 3318865.0 | 11263286.0 | 1279222.8 | 638961.44 | 56.47 | 62.1 | 50.6 | 38.8 |
| sp\|P61225\|RAP2B\_HUMAN | P61225 | RAP2B\_HUMAN | RAP2B | Ras-related protein Rap-2b | 183 | 20.504 | 1.680962e+06 | 4.330257e+07 | 3317378.0 | 6044563.0 | 15419078.0 | 18521550.0 | 74.32 | 70.5 | 62.8 | 31.1 |
| sp\|O60684\|IMA7\_HUMAN | O60684 | IMA7\_HUMAN | KPNA6 | Importin subunit alpha-7 | 536 | 60.030 | 1.541117e+05 | 1.162308e+07 | 3279422.0 | 6842773.5 | 1292084.8 | 208796.9 | 41.79 | 37.1 | 26.7 | 10.1 |
| sp\|Q9GZZ9\|UBA5\_HUMAN | Q9GZZ9 | UBA5\_HUMAN | UBA5 | Ubiquitin-like modifier-activating enzyme 5 | 404 | 44.863 | 1.542577e+05 | 8.694657e+06 | 3253164.8 | 4328361.0 | 704767.9 | 408363.28 | 51.49 | 53.7 | 15.1 | 11.1 |
| sp\|P51665\|PSMD7\_HUMAN | P51665 | PSMD7\_HUMAN | PSMD7 | 26S proteasome non-ATPase regulatory subunit 7 | 324 | 37.025 | 2.000028e+05 | 9.303539e+06 | 3252009.0 | 4822186.0 | 1034928.0 | 194416.47 | 58.95 | 56.2 | 22.5 | 11.4 |
| sp\|P14625\|ENPL\_HUMAN | P14625 | ENPL\_HUMAN | HSP90B1 | Endoplasmin | 803 | 92.469 | 1.520822e+05 | 1.766818e+07 | 3249335.2 | 3810320.8 | 9784986.0 | 823535.5 | 52.3 | 49.6 | 43.8 | 30.6 |
| sp\|P68402\|PA1B2\_HUMAN | P68402 | PA1B2\_HUMAN | PAFAH1B2 | Platelet-activating factor acetylhydrolase IB subunit alpha2 | 229 | 25.569 | 4.618028e+05 | 1.483500e+07 | 3240904.5 | 10843464.0 | 750629.5 |  | 34.5 | 46.3 | 19.2 |  |
| sp\|P50895\|BCAM\_HUMAN | P50895 | BCAM\_HUMAN | BCAM | Basal cell adhesion molecule | 628 | 67.405 | 3.615589e+05 | 3.061881e+07 | 3231858.8 | 2240410.2 | 5126592.0 | 20019948.0 | 34.71 | 32.6 | 41.9 | 42.4 |
| sp\|Q9UL46\|PSME2\_HUMAN | Q9UL46 | PSME2\_HUMAN | PSME2 | Proteasome activator complex subunit 2 | 239 | 27.402 | 4.655766e+05 | 1.602841e+07 | 3228895.8 | 9786574.0 | 2317559.0 | 695384.4 | 80.75 | 97.5 | 55.2 | 33.1 |
| sp\|Q5T6V5\|QNG1\_HUMAN | Q5T6V5 | QNG1\_HUMAN | QNG1 | Queuosine 5'-phosphate N-glycosylase/hydrolase | 341 | 39.029 | 1.589764e+05 | 7.795380e+06 | 3208277.8 | 4376637.0 | 210464.83 |  | 56.3 | 53.7 | 29.0 |  |
| sp\|P22303\|ACES\_HUMAN | P22303 | ACES\_HUMAN | ACHE | Acetylcholinesterase | 614 | 67.796 | 2.268815e+05 | 1.932504e+07 | 3199781.0 | 633553.6 | 4388211.0 | 11103492.0 | 41.86 | 45.6 | 30.0 | 20.2 |
| sp\|P36507\|MP2K2\_HUMAN | P36507 | MP2K2\_HUMAN | MAP2K2 | Dual specificity mitogen-activated protein kinase kinase 2 | 400 | 44.424 | 2.642720e+05 | 1.474979e+07 | 3193359.5 | 8566060.0 | 2339817.0 | 650557.8 | 53.25 | 59.8 | 36.0 | 27.5 |
| sp\|O00764\|PDXK\_HUMAN | O00764 | PDXK\_HUMAN | PDXK | Pyridoxal kinase | 312 | 35.102 | 1.617570e+05 | 7.133655e+06 | 3188508.8 | 3662915.8 | 282230.3 |  | 43.91 | 66.0 | 16.7 |  |
| sp\|P35613\|BASI\_HUMAN | P35613 | BASI\_HUMAN | BSG | Basigin | 385 | 42.200 | 5.789569e+05 | 3.069558e+07 | 3183778.5 | 4094104.2 | 16560617.0 | 6857078.0 | 21.56 | 34.8 | 29.1 | 14.0 |
| sp\|P11766\|ADHX\_HUMAN | P11766 | ADHX\_HUMAN | ADH5 | Alcohol dehydrogenase class-3 | 374 | 39.724 | 1.556146e+05 | 7.766415e+06 | 3150567.0 | 4615848.0 |  |  | 42.51 | 58.3 |  |  |
| sp\|P68036\|UB2L3\_HUMAN | P68036 | UB2L3\_HUMAN | UBE2L3 | Ubiquitin-conjugating enzyme E2 L3 | 154 | 17.862 | 1.622033e+06 | 3.640047e+07 | 3139027.0 | 29755326.0 | 2314456.8 | 1191664.9 | 76.62 | 83.1 | 27.9 | 16.2 |
| sp\|Q14247\|SRC8\_HUMAN | Q14247 | SRC8\_HUMAN | CTTN | Src substrate cortactin | 550 | 61.586 | 1.208590e+05 | 9.351432e+06 | 3136379.2 | 4973098.5 | 586242.25 | 655712.1 | 52.0 | 58.9 | 44.5 | 26.5 |
| sp\|P62873\|GBB1\_HUMAN | P62873 | GBB1\_HUMAN | GNB1 | Guanine nucleotide-binding protein G(I)/G(S)/G(T) subunit beta-1 | 340 | 37.377 | 2.295019e+05 | 1.077725e+07 | 3125723.0 | 1244659.0 | 5709014.5 | 697850.0 | 46.18 | 39.4 | 31.2 | 17.9 |
| sp\|P62879\|GBB2\_HUMAN | P62879 | GBB2\_HUMAN | GNB2 | Guanine nucleotide-binding protein G(I)/G(S)/G(T) subunit beta-2 | 340 | 37.331 | 2.149056e+05 | 1.007940e+07 | 3125723.0 | 1244659.0 | 5709014.5 |  | 35.88 | 58.8 | 26.2 |  |
| sp\|Q9HAV0\|GBB4\_HUMAN | Q9HAV0 | GBB4\_HUMAN | GNB4 | Guanine nucleotide-binding protein subunit beta-4 | 340 | 37.567 | 8.511158e+04 | 4.017097e+06 | 3125723.0 | 891373.9 |  |  | 21.76 | 40.9 |  |  |
| sp\|Q9BRG1\|VPS25\_HUMAN | Q9BRG1 | VPS25\_HUMAN | VPS25 | Vacuolar protein-sorting-associated protein 25 | 176 | 20.748 | 2.431638e+05 | 6.338585e+06 | 3110006.5 | 2995565.5 | 233013.19 |  | 57.39 | 63.1 | 34.7 |  |
| sp\|P68371\|TBB4B\_HUMAN | P68371 | TBB4B\_HUMAN | TUBB4B | Tubulin beta-4B chain | 445 | 49.831 | 2.484451e+05 | 1.555418e+07 | 3084202.5 | 8331895.0 | 3591840.2 | 546245.6 | 60.45 | 57.8 | 46.7 | 38.0 |
| sp\|P23743\|DGKA\_HUMAN | P23743 | DGKA\_HUMAN | DGKA | Diacylglycerol kinase alpha | 735 | 82.630 | 4.630867e+04 | 4.807477e+06 | 3072486.8 | 1488362.4 | 207809.75 | 38817.754 | 43.67 | 52.0 | 25.2 | 12.2 |
| sp\|P49589\|SYCC\_HUMAN | P49589 | SYCC\_HUMAN | CARS1 | Cysteine--tRNA ligase, cytoplasmic | 748 | 85.473 | 7.419527e+04 | 7.967504e+06 | 3061472.0 | 4593680.5 | 312351.78 |  | 58.56 | 60.3 | 30.5 |  |
| sp\|P48729\|KC1A\_HUMAN | P48729 | KC1A\_HUMAN | CSNK1A1 | Casein kinase I isoform alpha | 337 | 38.915 | 2.249533e+05 | 1.099832e+07 | 3059169.8 | 6894431.5 | 677546.0 | 367177.12 | 55.79 | 58.5 | 24.9 | 30.0 |
| sp\|Q13619\|CUL4A\_HUMAN | Q13619 | CUL4A\_HUMAN | CUL4A | Cullin-4A | 759 | 87.680 | 7.899510e+04 | 8.701975e+06 | 3056582.5 | 4957409.0 | 497779.62 | 190204.3 | 65.88 | 73.5 | 40.3 | 18.2 |
| sp\|Q5H9R7\|PP6R3\_HUMAN | Q5H9R7 | PP6R3\_HUMAN | PPP6R3 | Serine/threonine-protein phosphatase 6 regulatory subunit 3 | 873 | 97.669 | 7.112287e+04 | 8.727366e+06 | 3054814.8 | 5111739.0 | 194579.5 | 366232.22 | 58.08 | 50.3 | 18.7 | 19.6 |
| sp\|Q08722\|CD47\_HUMAN | Q08722 | CD47\_HUMAN | CD47 | Leukocyte surface antigen CD47 | 323 | 35.214 | 3.721840e+06 | 1.646608e+08 | 3051152.2 |  | 43727440.0 | 117882184.0 | 12.69 |  | 13.9 | 15.5 |
| sp\|O75116\|ROCK2\_HUMAN | O75116 | ROCK2\_HUMAN | ROCK2 | Rho-associated protein kinase 2 | 1388 | 160.900 | 4.754277e+04 | 9.610760e+06 | 3047164.0 | 2387892.5 | 3899311.5 | 276391.53 | 50.0 | 53.6 | 32.1 | 20.5 |
| sp\|P01112\|RASH\_HUMAN | P01112 | RASH\_HUMAN | HRAS | GTPase HRas | 189 | 21.298 | 1.857616e+05 | 4.970635e+06 | 3046304.0 | 377564.56 | 741281.2 | 805485.2 | 46.56 | 67.2 | 36.5 | 34.4 |
| sp\|Q9NWV4\|CZIB\_HUMAN | Q9NWV4 | CZIB\_HUMAN | CZIB | CXXC motif containing zinc binding protein | 160 | 18.048 | 3.741383e+05 | 8.483565e+06 | 3045142.8 | 5438422.0 |  |  | 76.88 | 73.1 |  |  |
| sp\|P07686\|HEXB\_HUMAN | P07686 | HEXB\_HUMAN | HEXB | Beta-hexosaminidase subunit beta | 556 | 61.160 | 3.959667e+04 | 3.042589e+06 | 3042589.0 |  |  |  | 3.6 |  |  |  |
| sp\|O76014\|KRT37\_HUMAN | O76014 | KRT37\_HUMAN | KRT37 | Keratin, type I cuticular Ha7 | 449 | 49.390 | 4.880538e+04 | 3.028474e+06 | 3028474.0 |  |  |  | 17.15 |  |  |  |
| sp\|O75131\|CPNE3\_HUMAN | O75131 | CPNE3\_HUMAN | CPNE3 | Copine-3 | 537 | 60.131 | 5.905058e+04 | 4.461077e+06 | 3022133.0 | 760573.56 | 432427.34 | 245943.23 | 37.06 | 35.0 | 25.0 | 14.0 |
| sp\|P61586\|RHOA\_HUMAN | P61586 | RHOA\_HUMAN | RHOA | Transforming protein RhoA | 193 | 21.768 | 5.318795e+05 | 1.454618e+07 | 2997589.2 | 10989452.0 | 197313.61 | 361825.7 | 43.52 | 54.4 | 31.1 | 10.4 |
| sp\|P04792\|HSPB1\_HUMAN | P04792 | HSPB1\_HUMAN | HSPB1 | Heat shock protein beta-1 | 205 | 22.783 | 4.977743e+05 | 1.424822e+07 | 2979117.2 | 7346938.0 | 2368668.0 | 1553494.0 | 68.29 | 75.6 | 62.0 | 26.3 |
| sp\|Q8WYN0\|ATG4A\_HUMAN | Q8WYN0 | ATG4A\_HUMAN | ATG4A | Cysteine protease ATG4A | 398 | 45.378 | 9.528052e+04 | 5.432086e+06 | 2973285.0 | 2243161.8 | 215638.97 |  | 30.9 | 34.4 | 12.6 |  |
| sp\|P0DPH7\|TBA3C\_HUMAN | P0DPH7 | TBA3C\_HUMAN | TUBA3C | Tubulin alpha-3C chain | 450 | 49.500 | 4.773455e+04 | 2.968624e+06 | 2968624.2 |  |  |  | 53.56 |  |  |  |
| sp\|P68366\|TBA4A\_HUMAN | P68366 | TBA4A\_HUMAN | TUBA4A | Tubulin alpha-4A chain | 448 | 49.924 | 1.303149e+05 | 8.173734e+06 | 2968624.2 | 4698355.0 | 434063.53 | 72691.52 | 50.45 | 48.2 | 30.6 | 22.3 |
| sp\|Q9BQE3\|TBA1C\_HUMAN | Q9BQE3 | TBA1C\_HUMAN | TUBA1C | Tubulin alpha-1C chain | 449 | 49.390 | 4.784087e+04 | 2.968624e+06 | 2968624.2 |  |  |  | 54.34 |  |  |  |
| sp\|P68363\|TBA1B\_HUMAN | P68363 | TBA1B\_HUMAN | TUBA1B | Tubulin alpha-1B chain | 451 | 50.152 | 1.452689e+05 | 9.153308e+06 | 2968624.2 | 5473186.0 | 571535.7 | 139961.97 | 63.64 | 0.0 | 37.3 | 22.7 |
| sp\|P17655\|CAN2\_HUMAN | P17655 | CAN2\_HUMAN | CAPN2 | Calpain-2 catalytic subunit | 700 | 79.995 | 4.482981e+04 | 4.505541e+06 | 2968521.5 | 1005533.9 | 308972.75 | 222512.78 | 37.71 | 32.7 | 16.6 | 11.1 |
| sp\|Q16543\|CDC37\_HUMAN | Q16543 | CDC37\_HUMAN | CDC37 | Hsp90 co-chaperone Cdc37 | 378 | 44.468 | 2.656623e+05 | 1.484207e+07 | 2952374.0 | 10332254.0 | 931579.5 | 625866.75 | 50.53 | 65.3 | 39.7 | 31.2 |
| sp\|O00442\|RTCA\_HUMAN | O00442 | RTCA\_HUMAN | RTCA | RNA 3'-terminal phosphate cyclase | 366 | 39.337 | 1.259756e+05 | 6.225940e+06 | 2951732.5 | 2888386.5 | 379446.53 | 6374.1064 | 54.92 | 65.3 | 22.4 | 9.3 |
| sp\|Q9C0E2\|XPO4\_HUMAN | Q9C0E2 | XPO4\_HUMAN | XPO4 | Exportin-4 | 1151 | 130.139 | 3.453429e+04 | 5.646445e+06 | 2935229.5 | 2264720.0 | 79437.51 | 367057.88 | 41.53 | 38.9 | 17.6 | 13.0 |
| sp\|Q8TEB1\|DCA11\_HUMAN | Q8TEB1 | DCA11\_HUMAN | DCAF11 | DDB1- and CUL4-associated factor 11 | 546 | 61.670 | 8.258188e+04 | 6.398466e+06 | 2933168.0 | 3174733.5 | 235496.58 | 55068.168 | 50.37 | 45.6 | 23.4 | 13.0 |
| sp\|Q9Y3Q8\|T22D4\_HUMAN | Q9Y3Q8 | T22D4\_HUMAN | TSC22D4 | TSC22 domain family protein 4 | 395 | 41.026 | 1.675477e+05 | 8.636045e+06 | 2921887.8 | 4503631.5 | 1062146.8 | 148379.2 | 53.42 | 53.4 | 40.0 | 26.8 |
| sp\|Q14344\|GNA13\_HUMAN | Q14344 | GNA13\_HUMAN | GNA13 | Guanine nucleotide-binding protein subunit alpha-13 | 377 | 44.050 | 8.606820e+04 | 4.763277e+06 | 2912888.0 | 986249.0 | 175745.23 | 688394.44 | 48.01 | 69.5 | 28.9 | 22.0 |
| sp\|Q53EL6\|PDCD4\_HUMAN | Q53EL6 | PDCD4\_HUMAN | PDCD4 | Programmed cell death protein 4 | 469 | 51.590 | 4.460187e+04 | 2.890918e+06 | 2890917.8 |  |  |  | 3.41 |  |  |  |
| sp\|O76013\|KRT36\_HUMAN | O76013 | KRT36\_HUMAN | KRT36 | Keratin, type I cuticular Ha6 | 467 | 52.247 | 6.572125e+05 | 4.314042e+07 | 2884696.2 |  | 385678.7 | 39870044.0 | 30.41 |  | 32.1 | 23.6 |
| sp\|Q99598\|TSNAX\_HUMAN | Q99598 | TSNAX\_HUMAN | TSNAX | Translin-associated protein X | 290 | 33.112 | 1.038111e+05 | 4.318634e+06 | 2877464.2 | 1314953.9 | 126215.8 |  | 62.07 | 79.7 | 38.3 |  |
| sp\|P43487\|RANG\_HUMAN | P43487 | RANG\_HUMAN | RANBP1 | Ran-specific GTPase-activating protein | 201 | 23.310 | 4.218399e+05 | 1.235399e+07 | 2875031.0 | 6641274.5 | 1958168.2 | 879511.4 | 71.64 | 78.6 | 38.8 | 16.9 |
| sp\|O00629\|IMA3\_HUMAN | O00629 | IMA3\_HUMAN | KPNA4 | Importin subunit alpha-3 | 521 | 57.887 | 1.380380e+05 | 1.003915e+07 | 2874737.8 | 6103218.0 | 744074.9 | 317118.62 | 42.8 | 51.1 | 11.3 | 7.3 |
| sp\|Q92508\|PIEZ1\_HUMAN | Q92508 | PIEZ1\_HUMAN | PIEZO1 | Piezo-type mechanosensitive ion channel component 1 | 2521 | 286.790 | 6.577984e+04 | 2.370140e+07 | 2861977.5 | 1851011.6 | 2765261.5 | 16223146.0 | 27.29 | 27.9 | 22.4 | 16.3 |
| sp\|Q9NUQ9\|CYRIB\_HUMAN | Q9NUQ9 | CYRIB\_HUMAN | CYRIB | CYFIP-related Rac1 interactor B | 324 | 36.748 | 1.516243e+05 | 7.000348e+06 | 2857380.5 | 3975735.2 | 93628.195 | 73604.12 | 74.38 | 69.4 | 21.3 | 17.6 |
| sp\|Q13177\|PAK2\_HUMAN | Q13177 | PAK2\_HUMAN | PAK2 | Serine/threonine-protein kinase PAK 2 | 524 | 58.043 | 1.110053e+05 | 8.094885e+06 | 2852028.0 | 3585908.5 | 1367392.0 | 289556.75 | 65.08 | 63.4 | 41.0 | 24.2 |
| sp\|Q96IJ6\|GMPPA\_HUMAN | Q96IJ6 | GMPPA\_HUMAN | GMPPA | Mannose-1-phosphate guanyltransferase alpha | 420 | 46.291 | 2.410861e+05 | 1.402122e+07 | 2845628.2 | 10756705.0 | 270458.0 | 148429.19 | 37.86 | 34.8 | 18.8 | 15.0 |
| sp\|Q9NRF8\|PYRG2\_HUMAN | Q9NRF8 | PYRG2\_HUMAN | CTPS2 | CTP synthase 2 | 586 | 65.678 | 1.091043e+05 | 9.002827e+06 | 2835258.2 | 5727359.0 | 440209.75 |  | 55.12 | 54.4 | 19.3 |  |
| sp\|Q96GG9\|DCNL1\_HUMAN | Q96GG9 | DCNL1\_HUMAN | DCUN1D1 | DCN1-like protein 1 | 259 | 30.124 | 3.890798e+05 | 1.472544e+07 | 2834486.0 | 11256572.0 | 571367.0 | 63019.992 | 50.97 | 52.1 | 37.8 | 27.0 |
| sp\|Q969R2\|OSBP2\_HUMAN | Q969R2 | OSBP2\_HUMAN | OSBP2 | Oxysterol-binding protein 2 | 916 | 101.266 | 4.932806e+04 | 6.275884e+06 | 2828549.5 | 2336280.2 | 646441.3 | 464612.5 | 56.88 | 65.5 | 30.0 | 18.1 |
| sp\|P49588\|SYAC\_HUMAN | P49588 | SYAC\_HUMAN | AARS1 | Alanine--tRNA ligase, cytoplasmic | 968 | 106.810 | 4.097696e+04 | 5.498812e+06 | 2825596.8 | 2393230.5 | 193405.6 | 86578.64 | 53.62 | 52.2 | 21.1 | 7.2 |
| sp\|Q562E7\|WDR81\_HUMAN | Q562E7 | WDR81\_HUMAN | WDR81 | WD repeat-containing protein 81 | 1941 | 211.697 | 2.701316e+04 | 7.184676e+06 | 2824975.0 | 3559581.0 | 256565.4 | 543554.94 | 45.75 | 45.6 | 11.7 | 11.3 |
| sp\|Q02750\|MP2K1\_HUMAN | Q02750 | MP2K1\_HUMAN | MAP2K1 | Dual specificity mitogen-activated protein kinase kinase 1 | 393 | 43.439 | 2.227865e+05 | 1.215866e+07 | 2821384.5 | 7062215.5 | 1854338.8 | 420724.53 | 41.48 | 61.8 | 35.1 | 26.7 |
| sp\|Q13509\|TBB3\_HUMAN | Q13509 | TBB3\_HUMAN | TUBB3 | Tubulin beta-3 chain | 450 | 50.433 | 7.559198e+04 | 4.789693e+06 | 2818340.2 | 1716821.1 |  | 254531.53 | 38.22 | 42.7 |  | 23.3 |
| sp\|P53041\|PPP5\_HUMAN | P53041 | PPP5\_HUMAN | PPP5C | Serine/threonine-protein phosphatase 5 | 499 | 56.879 | 1.282602e+05 | 9.165599e+06 | 2817096.0 | 5847322.0 | 339265.4 | 161915.9 | 47.9 | 63.5 | 24.6 | 12.2 |
| sp\|Q6L8Q7\|PDE12\_HUMAN | Q6L8Q7 | PDE12\_HUMAN | PDE12 | 2',5'-phosphodiesterase 12 | 609 | 67.352 | 5.226224e+04 | 4.422376e+06 | 2800171.8 | 1204611.0 | 230528.75 | 187064.08 | 51.4 | 58.3 | 25.3 | 13.8 |
| sp\|Q9UKX3\|MYH13\_HUMAN | Q9UKX3 | MYH13\_HUMAN | MYH13 | Myosin-13 | 1938 | 223.605 | 2.353247e+04 | 6.610984e+06 | 2796665.0 | 1561164.8 | 2253154.5 |  | 7.89 | 38.2 | 22.2 |  |
| sp\|O95837\|GNA14\_HUMAN | O95837 | GNA14\_HUMAN | GNA14 | Guanine nucleotide-binding protein subunit alpha-14 | 355 | 41.571 | 1.081261e+05 | 5.647266e+06 | 2793563.8 | 1194282.6 | 944174.56 | 715245.06 | 10.42 | 38.3 | 34.4 | 22.5 |
| sp\|Q969T9\|WBP2\_HUMAN | Q969T9 | WBP2\_HUMAN | WBP2 | WW domain-binding protein 2 | 261 | 28.087 | 9.036996e+05 | 3.188942e+07 | 2779084.5 | 18060424.0 | 9135884.0 | 1914027.2 | 37.55 | 63.6 | 28.0 | 19.5 |
| sp\|P25686\|DNJB2\_HUMAN | P25686 | DNJB2\_HUMAN | DNAJB2 | DnaJ homolog subfamily B member 2 | 324 | 35.580 | 3.621318e+05 | 1.618787e+07 | 2772162.0 | 7670427.0 | 3733887.0 | 2011394.1 | 39.51 | 58.6 | 41.4 | 34.9 |
| sp\|Q99829\|CPNE1\_HUMAN | Q99829 | CPNE1\_HUMAN | CPNE1 | Copine-1 | 537 | 59.059 | 9.548752e+04 | 7.085163e+06 | 2771771.5 | 3745852.2 | 426548.1 | 140991.53 | 24.02 | 47.5 | 21.6 | 7.3 |
| sp\|O94919\|ENDD1\_HUMAN | O94919 | ENDD1\_HUMAN | ENDOD1 | Endonuclease domain-containing 1 protein | 500 | 55.017 | 2.187927e+05 | 1.512331e+07 | 2759408.0 | 7999706.0 | 1701546.4 | 2662652.2 | 31.2 | 70.0 | 32.2 | 21.0 |
| sp\|Q01432\|AMPD3\_HUMAN | Q01432 | AMPD3\_HUMAN | AMPD3 | AMP deaminase 3 | 767 | 88.812 | 9.324253e+04 | 1.040406e+07 | 2757741.2 | 6702584.5 | 690580.1 | 253154.2 | 55.8 | 59.6 | 28.3 | 22.4 |
| sp\|Q9UI12\|VATH\_HUMAN | Q9UI12 | VATH\_HUMAN | ATP6V1H | V-type proton ATPase subunit H | 483 | 55.883 | 1.245745e+05 | 8.746336e+06 | 2739430.2 | 5414240.5 | 308148.47 | 284516.78 | 58.8 | 60.7 | 33.1 | 22.6 |
| sp\|P24941\|CDK2\_HUMAN | P24941 | CDK2\_HUMAN | CDK2 | Cyclin-dependent kinase 2 | 298 | 33.930 | 4.528152e+05 | 1.930288e+07 | 2734119.2 | 4139155.0 | 12039869.0 | 389731.88 | 68.46 | 77.5 | 35.6 | 25.8 |
| sp\|O60488\|ACSL4\_HUMAN | O60488 | ACSL4\_HUMAN | ACSL4 | Long-chain-fatty-acid--CoA ligase 4 | 711 | 79.188 | 7.781890e+04 | 7.742150e+06 | 2720013.5 | 4700193.5 | 167771.62 | 154171.56 | 63.57 | 56.1 | 14.5 | 18.3 |
| sp\|O15400\|STX7\_HUMAN | O15400 | STX7\_HUMAN | STX7 | Syntaxin-7 | 261 | 29.816 | 9.776314e+05 | 3.662197e+07 | 2713462.8 | 2439968.0 | 8396724.0 | 23071814.0 | 47.51 | 51.0 | 45.2 | 34.5 |
| sp\|Q9NXR1\|NDE1\_HUMAN | Q9NXR1 | NDE1\_HUMAN | NDE1 | Nuclear distribution protein nudE homolog 1 | 335 | 37.721 | 1.032894e+05 | 4.895040e+06 | 2706972.2 | 1732935.8 | 337173.66 | 117958.42 | 55.22 | 70.1 | 65.7 | 30.1 |
| sp\|Q9ULH1\|ASAP1\_HUMAN | Q9ULH1 | ASAP1\_HUMAN | ASAP1 | Arf-GAP with SH3 domain, ANK repeat and PH domain-containing protein 1 | 1129 | 125.498 | 8.351572e+04 | 1.316807e+07 | 2703396.5 | 8165486.5 | 1303641.1 | 995547.06 | 51.55 | 50.8 | 35.8 | 20.3 |
| sp\|P42330\|AK1C3\_HUMAN | P42330 | AK1C3\_HUMAN | AKR1C3 | Aldo-keto reductase family 1 member C3 | 323 | 36.853 | 1.155748e+05 | 5.351223e+06 | 2695540.2 | 1421497.5 | 1077532.1 | 156653.48 | 59.75 | 77.7 | 21.4 | 8.4 |
| sp\|Q53FA7\|QORX\_HUMAN | Q53FA7 | QORX\_HUMAN | TP53I3 | Quinone oxidoreductase PIG3 | 332 | 35.536 | 1.366372e+05 | 6.100346e+06 | 2678775.5 | 2975789.0 | 445781.88 |  | 54.52 | 69.9 | 18.4 |  |
| sp\|P41091\|IF2G\_HUMAN | P41091 | IF2G\_HUMAN | EIF2S3 | Eukaryotic translation initiation factor 2 subunit 3 | 472 | 51.109 | 1.314102e+05 | 8.438080e+06 | 2675231.8 | 4975154.0 | 436109.75 | 351584.62 | 53.18 | 56.6 | 37.3 | 22.0 |
| sp\|Q9NP79\|VTA1\_HUMAN | Q9NP79 | VTA1\_HUMAN | VTA1 | Vacuolar protein sorting-associated protein VTA1 homolog | 307 | 33.879 | 1.840009e+05 | 7.831911e+06 | 2656979.0 | 3429499.5 | 1318842.0 | 426590.12 | 37.13 | 43.6 | 22.5 | 14.3 |
| sp\|Q9NR46\|SHLB2\_HUMAN | Q9NR46 | SHLB2\_HUMAN | SH3GLB2 | Endophilin-B2 | 395 | 43.974 | 1.907188e+05 | 1.053675e+07 | 2625992.5 | 5337661.5 | 1565052.8 | 1008042.4 | 55.7 | 47.3 | 31.9 | 17.2 |
| sp\|Q8TBC4\|UBA3\_HUMAN | Q8TBC4 | UBA3\_HUMAN | UBA3 | NEDD8-activating enzyme E1 catalytic subunit | 463 | 51.852 | 8.073188e+04 | 5.259297e+06 | 2625088.0 | 2335253.8 | 298955.44 |  | 65.01 | 76.0 | 28.5 |  |
| sp\|P49915\|GUAA\_HUMAN | P49915 | GUAA\_HUMAN | GMPS | GMP synthase [glutamine-hydrolyzing] | 693 | 76.715 | 9.426971e+04 | 9.085934e+06 | 2622437.5 | 5876588.0 | 453754.12 | 133154.8 | 62.34 | 75.0 | 25.0 | 21.4 |
| sp\|Q6P2E9\|EDC4\_HUMAN | Q6P2E9 | EDC4\_HUMAN | EDC4 | Enhancer of mRNA-decapping protein 4 | 1401 | 151.661 | 3.641756e+04 | 6.939081e+06 | 2615853.0 | 383367.44 | 3864959.0 | 74901.43 | 16.49 | 33.8 | 15.1 | 8.9 |
| sp\|P27361\|MK03\_HUMAN | P27361 | MK03\_HUMAN | MAPK3 | Mitogen-activated protein kinase 3 | 379 | 43.136 | 1.018291e+05 | 5.518600e+06 | 2614238.0 | 2904362.2 |  |  | 40.11 | 55.1 |  |  |
| sp\|Q3ZCM7\|TBB8\_HUMAN | Q3ZCM7 | TBB8\_HUMAN | TUBB8 | Tubulin beta-8 chain | 444 | 48.840 | 4.260399e+04 | 2.614226e+06 | 2614226.0 |  |  |  | 19.37 |  |  |  |
| sp\|Q5VYK3\|ECM29\_HUMAN | Q5VYK3 | ECM29\_HUMAN | ECPAS | Proteasome adapter and scaffold protein ECM29 | 1845 | 204.291 | 3.121876e+04 | 8.012757e+06 | 2614172.8 | 4987644.5 | 127260.7 | 283679.16 | 54.63 | 49.5 | 23.4 | 11.0 |
| sp\|Q9NP58\|ABCB6\_HUMAN | Q9NP58 | ABCB6\_HUMAN | ABCB6 | ATP-binding cassette sub-family B member 6 | 842 | 93.886 | 2.373285e+05 | 2.799419e+07 | 2602092.0 | 3299437.2 | 5791999.5 | 16300663.0 | 43.82 | 34.7 | 32.7 | 29.7 |
| sp\|Q9NQW7\|XPP1\_HUMAN | Q9NQW7 | XPP1\_HUMAN | XPNPEP1 | Xaa-Pro aminopeptidase 1 | 623 | 69.918 | 7.985527e+04 | 7.014710e+06 | 2595812.2 | 4297568.0 | 75173.37 | 46156.438 | 52.97 | 55.1 | 22.2 | 10.3 |
| sp\|P16070\|CD44\_HUMAN | P16070 | CD44\_HUMAN | CD44 | CD44 antigen | 742 | 81.538 | 3.793183e+05 | 3.885805e+07 | 2587035.0 | 997313.1 | 8292969.5 | 26980736.0 | 14.56 | 31.0 | 20.1 | 19.4 |
| sp\|Q9Y3B8\|ORN\_HUMAN | Q9Y3B8 | ORN\_HUMAN | REXO2 | Oligoribonuclease, mitochondrial | 237 | 26.833 | 1.754198e+05 | 5.913779e+06 | 2585613.5 | 2548601.5 | 93096.44 | 686467.9 | 49.37 | 62.0 | 48.1 | 22.8 |
| sp\|Q96KP4\|CNDP2\_HUMAN | Q96KP4 | CNDP2\_HUMAN | CNDP2 | Cytosolic non-specific dipeptidase | 475 | 52.878 | 1.306494e+05 | 8.679594e+06 | 2564482.8 | 5563722.5 | 405221.22 | 146167.88 | 60.21 | 73.9 | 24.4 | 11.4 |
| sp\|A6NDG6\|PGP\_HUMAN | A6NDG6 | PGP\_HUMAN | PGP | Glycerol-3-phosphate phosphatase | 321 | 34.006 | 3.718061e+05 | 1.588507e+07 | 2552385.5 | 11580530.0 | 1288395.5 | 463763.6 | 46.42 | 69.2 | 37.4 | 24.9 |
| sp\|P46379\|BAG6\_HUMAN | P46379 | BAG6\_HUMAN | BAG6 | Large proline-rich protein BAG6 | 1132 | 119.409 | 5.283004e+04 | 7.925654e+06 | 2548391.2 | 3429872.8 | 891729.7 | 1055660.6 | 39.4 | 37.9 | 24.7 | 11.1 |
| sp\|O95376\|ARI2\_HUMAN | O95376 | ARI2\_HUMAN | ARIH2 | E3 ubiquitin-protein ligase ARIH2 | 493 | 57.819 | 1.237893e+05 | 8.992304e+06 | 2541932.5 | 6219984.0 | 175294.2 | 55092.918 | 42.8 | 40.2 | 20.9 | 16.2 |
| sp\|Q7L9L4\|MOB1B\_HUMAN | Q7L9L4 | MOB1B\_HUMAN | MOB1B | MOB kinase activator 1B | 216 | 25.091 | 4.075055e+05 | 1.284602e+07 | 2540602.5 | 9142775.0 | 1011480.2 | 151160.97 | 27.78 | 42.1 | 32.9 | 8.8 |
| sp\|P07741\|APT\_HUMAN | P07741 | APT\_HUMAN | APRT | Adenine phosphoribosyltransferase | 180 | 19.608 | 7.935937e+05 | 1.955009e+07 | 2532694.5 | 14897785.0 | 1496362.8 | 623242.75 | 69.44 | 98.3 | 38.9 | 33.3 |
| sp\|P29992\|GNA11\_HUMAN | P29992 | GNA11\_HUMAN | GNA11 | Guanine nucleotide-binding protein subunit alpha-11 | 359 | 42.123 | 8.678876e+04 | 4.593037e+06 | 2527255.2 | 597532.7 | 743222.94 | 725026.2 | 33.15 | 41.5 | 31.5 | 23.4 |
| sp\|O00299\|CLIC1\_HUMAN | O00299 | CLIC1\_HUMAN | CLIC1 | Chloride intracellular channel protein 1 | 241 | 26.923 | 5.029115e+05 | 1.701109e+07 | 2519582.2 | 9487979.0 | 4703587.0 | 299942.97 | 78.01 | 90.9 | 42.7 | 29.5 |
| sp\|Q92995\|UBP13\_HUMAN | Q92995 | UBP13\_HUMAN | USP13 | Ubiquitin carboxyl-terminal hydrolase 13 | 863 | 94.930 | 2.110125e+04 | 2.516684e+06 | 2516684.5 |  |  |  | 5.79 |  |  |  |
| sp\|P61758\|PFD3\_HUMAN | P61758 | PFD3\_HUMAN | VBP1 | Prefoldin subunit 3 | 197 | 22.626 | 1.899498e+05 | 5.399628e+06 | 2500966.2 | 2338917.5 | 439043.9 | 120700.51 | 53.81 | 74.1 | 44.2 | 16.8 |
| sp\|Q5T619\|ZN648\_HUMAN | Q5T619 | ZN648\_HUMAN | ZNF648 | Zinc finger protein 648 | 568 | 62.480 | 3.149954e+04 | 2.472648e+06 | 2472648.5 |  |  |  | 2.11 |  |  |  |
| sp\|P06703\|S10A6\_HUMAN | P06703 | S10A6\_HUMAN | S100A6 | Protein S100-A6 | 90 | 10.180 | 4.970421e+06 | 6.357086e+07 | 2464689.8 | 28407724.0 | 13198229.0 | 19500214.0 | 54.44 | 71.1 | 65.6 | 45.6 |
| sp\|P07902\|GALT\_HUMAN | P07902 | GALT\_HUMAN | GALT | Galactose-1-phosphate uridylyltransferase | 379 | 43.363 | 8.784058e+04 | 4.785548e+06 | 2463937.0 | 1581698.6 | 617594.3 | 122318.266 | 33.25 | 35.4 | 15.6 | 10.6 |
| sp\|P63092\|GNAS2\_HUMAN | P63092 | GNAS2\_HUMAN | GNAS | Guanine nucleotide-binding protein G(s) subunit alpha isoforms short | 394 | 45.665 | 3.309008e+05 | 1.898447e+07 | 2462631.0 | 1406121.8 | 6712468.0 | 8403245.0 | 41.62 | 45.2 | 31.2 | 26.4 |
| sp\|P78318\|IGBP1\_HUMAN | P78318 | IGBP1\_HUMAN | IGBP1 | Immunoglobulin-binding protein 1 | 339 | 39.222 | 2.004060e+05 | 9.875466e+06 | 2462461.5 | 6451989.0 | 741719.9 | 219296.08 | 45.13 | 89.1 | 44.8 | 23.6 |
| sp\|P10644\|KAP0\_HUMAN | P10644 | KAP0\_HUMAN | PRKAR1A | cAMP-dependent protein kinase type I-alpha regulatory subunit | 381 | 42.982 | 5.325622e+05 | 2.875902e+07 | 2461495.2 | 4697509.0 | 20895790.0 | 704230.0 | 53.54 | 55.6 | 46.2 | 22.3 |
| sp\|P31153\|METK2\_HUMAN | P31153 | METK2\_HUMAN | MAT2A | S-adenosylmethionine synthase isoform type-2 | 395 | 43.661 | 9.234068e+04 | 5.065285e+06 | 2458623.2 | 2581702.5 | 24959.273 |  | 47.59 | 69.1 | 15.2 |  |
| sp\|Q01469\|FABP5\_HUMAN | Q01469 | FABP5\_HUMAN | FABP5 | Fatty acid-binding protein 5 | 135 | 15.164 | 4.187214e+05 | 7.977302e+06 | 2454578.0 | 2992507.0 | 716921.2 | 1813296.2 | 53.33 | 86.7 | 34.1 | 47.4 |
| sp\|P23919\|KTHY\_HUMAN | P23919 | KTHY\_HUMAN | DTYMK | Thymidylate kinase | 212 | 23.819 | 2.243286e+05 | 6.713135e+06 | 2445921.0 | 3771925.2 | 495288.44 |  | 73.58 | 78.8 | 43.9 |  |
| sp\|Q2VIR3\|IF2GL\_HUMAN | Q2VIR3 | IF2GL\_HUMAN | EIF2S3B | Eukaryotic translation initiation factor 2 subunit 3B | 472 | 51.920 | 3.732809e+04 | 2.434936e+06 | 2434936.5 |  |  |  | 47.25 |  |  |  |
| sp\|Q9H4A3\|WNK1\_HUMAN | Q9H4A3 | WNK1\_HUMAN | WNK1 | Serine/threonine-protein kinase WNK1 | 2382 | 250.794 | 4.657972e+04 | 1.467679e+07 | 2434061.8 | 8388526.5 | 2540863.8 | 1313340.9 | 29.97 | 30.5 | 17.8 | 15.0 |
| sp\|P35580\|MYH10\_HUMAN | P35580 | MYH10\_HUMAN | MYH10 | Myosin-10 | 1976 | 228.999 | 2.339846e+04 | 6.731906e+06 | 2418476.5 | 3064010.2 | 304981.47 | 944438.3 | 49.39 | 58.7 | 24.3 | 14.4 |
| sp\|Q8TAC1\|RFESD\_HUMAN | Q8TAC1 | RFESD\_HUMAN | RFESD | Rieske domain-containing protein | 157 | 17.762 | 3.488406e+05 | 7.784594e+06 | 2407895.0 | 5241268.5 | 86397.64 | 49033.117 | 52.87 | 47.1 | 42.0 | 18.5 |
| sp\|Q15418\|KS6A1\_HUMAN | Q15418 | KS6A1\_HUMAN | RPS6KA1 | Ribosomal protein S6 kinase alpha-1 | 735 | 82.723 | 5.062589e+04 | 5.261579e+06 | 2389020.2 | 2507793.5 | 364765.28 |  | 51.7 | 52.4 | 28.0 |  |
| sp\|P16152\|CBR1\_HUMAN | P16152 | CBR1\_HUMAN | CBR1 | Carbonyl reductase [NADPH] 1 | 277 | 30.375 | 2.243001e+05 | 8.559784e+06 | 2384557.8 | 5579905.0 | 505789.25 | 89532.44 | 75.45 | 81.9 | 49.5 | 22.0 |
| sp\|Q15173\|2A5B\_HUMAN | Q15173 | 2A5B\_HUMAN | PPP2R5B | Serine/threonine-protein phosphatase 2A 56 kDa regulatory subunit beta isoform | 497 | 57.393 | 5.027236e+04 | 3.624978e+06 | 2373571.8 | 1046488.8 | 65372.668 | 139544.4 | 35.81 | 36.4 | 13.7 | 6.0 |
| sp\|Q7L5Y1\|ENOF1\_HUMAN | Q7L5Y1 | ENOF1\_HUMAN | ENOSF1 | Mitochondrial enolase superfamily member 1 | 443 | 49.786 | 1.061891e+05 | 6.642086e+06 | 2372464.0 | 3647307.5 | 622314.4 |  | 57.56 | 82.8 | 27.1 |  |
| sp\|P07437\|TBB5\_HUMAN | P07437 | TBB5\_HUMAN | TUBB | Tubulin beta chain | 444 | 49.671 | 2.220045e+05 | 1.385421e+07 | 2371234.8 | 8331895.0 | 2774187.0 | 376893.84 | 53.6 | 47.5 | 40.3 | 41.4 |
| sp\|Q13885\|TBB2A\_HUMAN | Q13885 | TBB2A\_HUMAN | TUBB2A | Tubulin beta-2A chain | 445 | 49.907 | 2.149437e+05 | 1.347732e+07 | 2371234.8 | 8331895.0 | 2774187.0 |  | 46.07 | 43.6 | 44.7 |  |
| sp\|Q9GZN8\|ADSSP\_HUMAN | Q9GZN8 | ADSSP\_HUMAN | ADISSP | Adipose-secreted signaling protein | 174 | 19.291 | 3.734343e+05 | 9.050782e+06 | 2341161.0 | 5758497.0 | 951124.44 |  | 67.24 | 63.8 | 33.9 |  |
| sp\|Q8IY47\|KBTB2\_HUMAN | Q8IY47 | KBTB2\_HUMAN | KBTBD2 | Kelch repeat and BTB domain-containing protein 2 | 623 | 68.530 | 2.706839e+04 | 2.330560e+06 | 2330559.8 |  |  |  | 7.87 |  |  |  |
| sp\|O15084\|ANR28\_HUMAN | O15084 | ANR28\_HUMAN | ANKRD28 | Serine/threonine-protein phosphatase 6 regulatory ankyrin repeat subunit A | 1053 | 112.966 | 4.900620e+04 | 6.955301e+06 | 2320821.2 | 4254786.0 | 250859.95 | 128834.12 | 44.54 | 53.2 | 14.2 | 7.0 |
| sp\|Q8NCW5\|NNRE\_HUMAN | Q8NCW5 | NNRE\_HUMAN | NAXE | NAD(P)H-hydrate epimerase | 288 | 31.675 | 3.605523e+05 | 1.434835e+07 | 2315129.5 | 11797356.0 | 235867.88 |  | 48.26 | 60.1 | 30.2 |  |
| sp\|A4D1P6\|WDR91\_HUMAN | A4D1P6 | WDR91\_HUMAN | WDR91 | WD repeat-containing protein 91 | 747 | 83.344 | 7.997601e+04 | 8.374353e+06 | 2312838.2 | 4637027.5 | 1089307.2 | 335179.75 | 45.92 | 54.1 | 41.1 | 21.7 |
| sp\|O00154\|BACH\_HUMAN | O00154 | BACH\_HUMAN | ACOT7 | Cytosolic acyl coenzyme A thioester hydrolase | 380 | 41.796 | 1.297536e+05 | 6.813514e+06 | 2310423.8 | 4111527.2 | 356387.44 | 35175.09 | 45.79 | 71.8 | 29.5 | 20.0 |
| sp\|Q9ULP9\|TBC24\_HUMAN | Q9ULP9 | TBC24\_HUMAN | TBC1D24 | TBC1 domain family member 24 | 559 | 62.919 | 9.761814e+04 | 7.716662e+06 | 2304528.2 | 4014559.5 | 345667.72 | 1051906.8 | 44.19 | 43.3 | 34.2 | 20.8 |
| sp\|P56537\|IF6\_HUMAN | P56537 | IF6\_HUMAN | EIF6 | Eukaryotic translation initiation factor 6 | 245 | 26.599 | 3.106684e+05 | 1.038197e+07 | 2301406.5 | 6619190.0 | 1208155.4 | 253213.17 | 56.73 | 57.6 | 17.6 | 18.4 |
| sp\|O43598\|DNPH1\_HUMAN | O43598 | DNPH1\_HUMAN | DNPH1 | 2'-deoxynucleoside 5'-phosphate N-hydrolase 1 | 174 | 19.108 | 2.624790e+05 | 6.301253e+06 | 2286655.5 | 2671252.2 | 569864.3 | 773481.0 | 68.39 | 54.6 | 28.7 | 23.0 |
| sp\|P57737\|CORO7\_HUMAN | P57737 | CORO7\_HUMAN | CORO7 | Coronin-7 | 925 | 100.605 | 6.645110e+04 | 8.399218e+06 | 2285823.5 | 5532589.5 | 363787.0 | 217018.25 | 43.14 | 42.1 | 20.8 | 9.5 |
| sp\|P13639\|EF2\_HUMAN | P13639 | EF2\_HUMAN | EEF2 | Elongation factor 2 | 858 | 95.338 | 5.706514e+04 | 6.835245e+06 | 2284282.5 | 3199201.0 | 783400.8 | 568360.6 | 55.71 | 56.6 | 35.5 | 19.8 |
| sp\|Q16774\|KGUA\_HUMAN | Q16774 | KGUA\_HUMAN | GUK1 | Guanylate kinase | 197 | 21.726 | 4.277262e+05 | 1.167516e+07 | 2278727.2 | 7835778.0 | 1303620.8 | 257033.66 | 68.02 | 91.9 | 39.6 | 34.5 |
| sp\|P53602\|MVD1\_HUMAN | P53602 | MVD1\_HUMAN | MVD | Diphosphomevalonate decarboxylase | 400 | 43.405 | 7.985705e+04 | 4.354820e+06 | 2277075.5 | 1353238.2 | 500959.84 | 223546.28 | 54.5 | 64.0 | 37.5 | 23.2 |
| sp\|Q15365\|PCBP1\_HUMAN | Q15365 | PCBP1\_HUMAN | PCBP1 | Poly(rC)-binding protein 1 | 356 | 37.498 | 2.600330e+05 | 1.225050e+07 | 2276385.2 | 5354503.0 | 2997572.0 | 1622037.1 | 58.99 | 70.5 | 41.0 | 18.8 |
| sp\|P43490\|NAMPT\_HUMAN | P43490 | NAMPT\_HUMAN | NAMPT | Nicotinamide phosphoribosyltransferase | 491 | 55.521 | 5.714325e+04 | 3.986019e+06 | 2276196.5 | 1332683.4 | 351149.7 | 25989.14 | 61.71 | 59.7 | 24.0 | 10.0 |
| sp\|P61978\|HNRPK\_HUMAN | P61978 | HNRPK\_HUMAN | HNRNPK | Heterogeneous nuclear ribonucleoprotein K | 463 | 50.976 | 2.060186e+05 | 1.319439e+07 | 2262442.5 | 8771745.0 | 1520302.1 | 639903.9 | 45.36 | 71.9 | 49.0 | 38.7 |
| sp\|Q9NX46\|ADPRS\_HUMAN | Q9NX46 | ADPRS\_HUMAN | ADPRS | ADP-ribosylhydrolase ARH3 | 363 | 38.947 | 1.230608e+05 | 6.021586e+06 | 2260162.5 | 3396175.0 | 283158.4 | 82089.96 | 40.5 | 49.6 | 32.2 | 15.7 |
| sp\|Q9H2U2\|IPYR2\_HUMAN | Q9H2U2 | IPYR2\_HUMAN | PPA2 | Inorganic pyrophosphatase 2, mitochondrial | 334 | 37.920 | 5.319270e+04 | 2.534180e+06 | 2251518.8 | 169425.94 | 70325.64 | 42910.04 | 36.23 | 45.2 | 29.6 | 19.2 |
| sp\|P27708\|PYR1\_HUMAN | P27708 | PYR1\_HUMAN | CAD | CAD protein | 2225 | 242.984 | 2.624891e+04 | 8.013202e+06 | 2249844.8 | 2655995.2 | 2935625.8 | 171736.62 | 32.4 | 41.5 | 15.9 | 10.8 |
| sp\|P23921\|RIR1\_HUMAN | P23921 | RIR1\_HUMAN | RRM1 | Ribonucleoside-diphosphate reductase large subunit | 792 | 90.070 | 6.629847e+04 | 7.502411e+06 | 2249050.2 | 4691426.5 | 428090.94 | 133843.38 | 58.46 | 58.1 | 23.2 | 7.3 |
| sp\|Q9UHJ6\|SHPK\_HUMAN | Q9UHJ6 | SHPK\_HUMAN | SHPK | Sedoheptulokinase | 478 | 51.505 | 6.792234e+04 | 4.395206e+06 | 2246547.0 | 2132073.5 |  | 16585.271 | 46.23 | 57.1 |  | 10.7 |
| sp\|Q96GD0\|PLPP\_HUMAN | Q96GD0 | PLPP\_HUMAN | PDXP | Chronophin | 296 | 31.698 | 2.222156e+05 | 8.849597e+06 | 2245446.5 | 4846667.5 | 1340356.2 | 417126.8 | 59.8 | 71.6 | 37.8 | 17.9 |
| sp\|Q96K76\|UBP47\_HUMAN | Q96K76 | UBP47\_HUMAN | USP47 | Ubiquitin carboxyl-terminal hydrolase 47 | 1375 | 157.311 | 3.095726e+04 | 6.118412e+06 | 2241586.8 | 3203720.2 | 444027.16 | 229077.75 | 51.56 | 51.6 | 28.9 | 13.5 |
| sp\|Q13557\|KCC2D\_HUMAN | Q13557 | KCC2D\_HUMAN | CAMK2D | Calcium/calmodulin-dependent protein kinase type II subunit delta | 499 | 56.369 | 3.470423e+04 | 2.457762e+06 | 2234881.5 | 27675.875 | 131923.06 | 63281.906 | 6.81 | 43.5 | 17.6 | 13.2 |
| sp\|Q9H4G4\|GAPR1\_HUMAN | Q9H4G4 | GAPR1\_HUMAN | GLIPR2 | Golgi-associated plant pathogenesis-related protein 1 | 154 | 17.218 | 5.207865e+05 | 1.126574e+07 | 2232401.5 | 3995362.5 | 1986029.4 | 3051943.5 | 68.83 | 85.7 | 39.0 | 26.6 |
| sp\|Q92783\|STAM1\_HUMAN | Q92783 | STAM1\_HUMAN | STAM | Signal transducing adapter molecule 1 | 540 | 59.180 | 1.009280e+05 | 7.504187e+06 | 2224825.5 | 3996749.2 | 962452.94 | 320159.0 | 51.11 | 49.8 | 31.9 | 15.9 |
| sp\|P28838\|AMPL\_HUMAN | P28838 | AMPL\_HUMAN | LAP3 | Cytosol aminopeptidase | 519 | 56.166 | 9.045935e+04 | 6.383283e+06 | 2224486.8 | 3792091.2 | 283545.16 | 83159.87 | 69.94 | 69.2 | 30.4 | 15.8 |
| sp\|P10599\|THIO\_HUMAN | P10599 | THIO\_HUMAN | TXN | Thioredoxin | 105 | 11.737 | 2.509263e+06 | 3.700160e+07 | 2198915.0 | 25282748.0 | 7514765.5 | 2005166.8 | 60.0 | 76.2 | 75.2 | 54.3 |
| sp\|Q13620\|CUL4B\_HUMAN | Q13620 | CUL4B\_HUMAN | CUL4B | Cullin-4B | 913 | 103.982 | 5.193122e+04 | 6.784281e+06 | 2197974.8 | 4288469.0 | 289854.25 | 7982.8716 | 19.72 | 51.0 | 24.4 | 8.4 |
| sp\|Q9Y5P6\|GMPPB\_HUMAN | Q9Y5P6 | GMPPB\_HUMAN | GMPPB | Mannose-1-phosphate guanyltransferase beta | 360 | 39.834 | 1.027766e+05 | 5.143580e+06 | 2197622.5 | 2564741.0 | 381216.38 |  | 36.67 | 48.9 | 33.6 |  |
| sp\|P51812\|KS6A3\_HUMAN | P51812 | KS6A3\_HUMAN | RPS6KA3 | Ribosomal protein S6 kinase alpha-3 | 740 | 83.736 | 5.630118e+04 | 5.923070e+06 | 2193169.8 | 2982078.0 | 505721.44 | 242100.7 | 63.51 | 63.8 | 32.6 | 19.7 |
| sp\|Q8N0W3\|FCSK\_HUMAN | Q8N0W3 | FCSK\_HUMAN | FCSK | L-fucose kinase | 1084 | 117.623 | 2.901858e+04 | 4.288305e+06 | 2192284.8 | 2051336.8 | 44683.055 |  | 42.71 | 38.3 | 11.2 |  |
| sp\|P09525\|ANXA4\_HUMAN | P09525 | ANXA4\_HUMAN | ANXA4 | Annexin A4 | 319 | 35.883 | 1.269574e+05 | 5.723530e+06 | 2189527.0 | 3070004.5 | 295066.25 | 168932.19 | 59.25 | 68.0 | 34.2 | 34.2 |
| sp\|P18206\|VINC\_HUMAN | P18206 | VINC\_HUMAN | VCL | Vinculin | 1134 | 123.799 | 3.042272e+04 | 4.731865e+06 | 2189454.0 | 1845372.9 | 408566.2 | 288472.28 | 60.41 | 52.9 | 40.4 | 22.5 |
| sp\|Q02413\|DSG1\_HUMAN | Q02413 | DSG1\_HUMAN | DSG1 | Desmoglein-1 | 1049 | 113.748 | 5.947044e+04 | 8.498887e+06 | 2164670.0 | 747837.4 | 2223689.2 | 3362690.8 | 29.93 | 33.7 | 24.6 | 17.1 |
| sp\|Q9H008\|LHPP\_HUMAN | Q9H008 | LHPP\_HUMAN | LHPP | Phospholysine phosphohistidine inorganic pyrophosphate phosphatase | 270 | 29.165 | 3.176395e+05 | 1.163895e+07 | 2158462.8 | 9204403.0 | 181512.3 | 94568.33 | 43.7 | 56.7 | 20.7 | 13.0 |
| sp\|O15439\|MRP4\_HUMAN | O15439 | MRP4\_HUMAN | ABCC4 | ATP-binding cassette sub-family C member 4 | 1325 | 149.527 | 1.023204e+05 | 1.922202e+07 | 2158235.2 | 3415525.5 | 2695738.5 | 10952516.0 | 44.0 | 43.5 | 29.8 | 28.0 |
| sp\|P25789\|PSA4\_HUMAN | P25789 | PSA4\_HUMAN | PSMA4 | Proteasome subunit alpha type-4 | 261 | 29.484 | 2.714810e+05 | 1.005641e+07 | 2156176.2 | 5535902.0 | 1561335.1 | 802998.56 | 73.56 | 80.8 | 32.2 | 20.3 |
| sp\|P61018\|RAB4B\_HUMAN | P61018 | RAB4B\_HUMAN | RAB4B | Ras-related protein Rab-4B | 213 | 23.587 | 1.469226e+05 | 4.353901e+06 | 2146220.0 | 478980.47 | 1728700.8 |  | 33.33 | 57.7 | 37.6 |  |
| sp\|Q58WW2\|DCAF6\_HUMAN | Q58WW2 | DCAF6\_HUMAN | DCAF6 | DDB1- and CUL4-associated factor 6 | 860 | 96.292 | 5.559907e+04 | 6.726279e+06 | 2143188.0 | 3709862.0 | 418357.28 | 454871.47 | 44.65 | 52.1 | 28.0 | 15.9 |
| sp\|P20338\|RAB4A\_HUMAN | P20338 | RAB4A\_HUMAN | RAB4A | Ras-related protein Rab-4A | 218 | 24.390 | 8.257866e+04 | 2.530444e+06 | 2139705.8 | 390738.56 |  |  | 32.57 | 28.9 |  |  |
| sp\|Q13287\|NMI\_HUMAN | Q13287 | NMI\_HUMAN | NMI | N-myc-interactor | 307 | 35.057 | 7.016104e+04 | 3.090210e+06 | 2139022.5 | 803456.6 | 147730.52 |  | 38.44 | 46.6 | 25.1 |  |
| sp\|Q96FV2\|SCRN2\_HUMAN | Q96FV2 | SCRN2\_HUMAN | SCRN2 | Secernin-2 | 425 | 46.597 | 5.705574e+04 | 3.340215e+06 | 2137729.5 | 1065269.2 | 137216.55 |  | 36.24 | 48.7 | 12.0 |  |
| sp\|Q9Y613\|FHOD1\_HUMAN | Q9Y613 | FHOD1\_HUMAN | FHOD1 | FH1/FH2 domain-containing protein 1 | 1164 | 126.551 | 2.693306e+04 | 4.282214e+06 | 2135531.0 | 1648424.9 | 223631.47 | 274626.97 | 43.04 | 56.4 | 24.0 | 17.1 |
| sp\|O75326\|SEM7A\_HUMAN | O75326 | SEM7A\_HUMAN | SEMA7A | Semaphorin-7A | 666 | 74.824 | 2.769694e+05 | 2.603694e+07 | 2112497.0 | 1184158.5 | 8648468.0 | 14091814.0 | 34.53 | 47.4 | 39.9 | 32.1 |
| sp\|Q9UEW8\|STK39\_HUMAN | Q9UEW8 | STK39\_HUMAN | STK39 | STE20/SPS1-related proline-alanine-rich protein kinase | 545 | 59.474 | 4.357099e+04 | 3.255680e+06 | 2108179.5 | 682287.8 | 465212.62 |  | 18.9 | 33.2 | 14.9 |  |
| sp\|P46109\|CRKL\_HUMAN | P46109 | CRKL\_HUMAN | CRKL | Crk-like protein | 303 | 33.777 | 2.253734e+05 | 9.564028e+06 | 2096196.4 | 6665475.5 | 802355.94 |  | 66.34 | 91.4 | 35.6 |  |
| sp\|Q9Y5L0\|TNPO3\_HUMAN | Q9Y5L0 | TNPO3\_HUMAN | TNPO3 | Transportin-3 | 923 | 104.203 | 3.318129e+04 | 4.344008e+06 | 2089808.9 | 2021161.4 | 54506.895 | 178530.53 | 42.9 | 49.6 | 17.0 | 12.4 |
| sp\|Q92696\|PGTA\_HUMAN | Q92696 | PGTA\_HUMAN | RABGGTA | Geranylgeranyl transferase type-2 subunit alpha | 567 | 65.072 | 4.212576e+04 | 3.443967e+06 | 2089750.8 | 1054008.5 | 197999.88 | 102208.09 | 57.32 | 41.8 | 24.2 | 17.3 |
| sp\|P15924\|DESP\_HUMAN | P15924 | DESP\_HUMAN | DSP | Desmoplakin | 2871 | 331.774 | 2.896579e+04 | 1.207382e+07 | 2089625.4 | 1614479.9 | 3166658.2 | 5203059.5 | 41.38 | 48.5 | 44.0 | 35.9 |
| sp\|P36915\|GNL1\_HUMAN | P36915 | GNL1\_HUMAN | GNL1 | Guanine nucleotide-binding protein-like 1 | 607 | 68.661 | 7.450409e+04 | 6.426987e+06 | 2087768.2 | 3407326.0 | 437641.75 | 494251.03 | 47.78 | 57.5 | 18.1 | 10.5 |
| sp\|Q96CV9\|OPTN\_HUMAN | Q96CV9 | OPTN\_HUMAN | OPTN | Optineurin | 577 | 65.922 | 8.906535e+04 | 7.376601e+06 | 2079887.6 | 4361183.0 | 446137.1 | 489393.53 | 58.93 | 73.3 | 49.6 | 37.6 |
| sp\|P60510\|PP4C\_HUMAN | P60510 | PP4C\_HUMAN | PPP4C | Serine/threonine-protein phosphatase 4 catalytic subunit | 307 | 35.080 | 6.315520e+04 | 2.783466e+06 | 2074056.5 | 709409.06 |  |  | 17.26 | 26.1 |  |  |
| sp\|O43665\|RGS10\_HUMAN | O43665 | RGS10\_HUMAN | RGS10 | Regulator of G-protein signaling 10 | 181 | 21.210 | 8.576661e+05 | 2.285473e+07 | 2073335.5 | 20019402.0 | 546158.56 | 215833.23 | 57.46 | 72.4 | 32.6 | 27.6 |
| sp\|O95834\|EMAL2\_HUMAN | O95834 | EMAL2\_HUMAN | EML2 | Echinoderm microtubule-associated protein-like 2 | 649 | 70.679 | 4.633604e+04 | 4.114589e+06 | 2056465.2 | 1544590.6 | 513533.25 |  | 36.52 | 47.8 | 21.9 |  |
| sp\|Q6UWP2\|DHR11\_HUMAN | Q6UWP2 | DHR11\_HUMAN | DHRS11 | Dehydrogenase/reductase SDR family member 11 | 260 | 28.308 | 1.025905e+05 | 3.648660e+06 | 2054204.9 | 1504372.1 | 90083.19 |  | 51.92 | 63.5 | 32.7 |  |
| sp\|P55957\|BID\_HUMAN | P55957 | BID\_HUMAN | BID | BH3-interacting domain death agonist | 195 | 21.995 | 3.924433e+05 | 1.084471e+07 | 2051785.8 | 7003429.5 | 1287670.0 | 501827.4 | 61.03 | 68.2 | 23.1 | 21.5 |
| sp\|Q9UBS0\|KS6B2\_HUMAN | Q9UBS0 | KS6B2\_HUMAN | RPS6KB2 | Ribosomal protein S6 kinase beta-2 | 482 | 53.455 | 3.806209e+04 | 2.556220e+06 | 2051167.4 | 276313.53 | 148550.27 | 80188.33 | 18.88 | 28.2 | 15.1 | 5.4 |
| sp\|P16083\|NQO2\_HUMAN | P16083 | NQO2\_HUMAN | NQO2 | Ribosyldihydronicotinamide dehydrogenase [quinone] | 231 | 25.919 | 4.533731e+05 | 1.476356e+07 | 2051076.8 | 12477736.0 | 234749.22 |  | 51.95 | 79.2 | 34.6 |  |
| sp\|P45984\|MK09\_HUMAN | P45984 | MK09\_HUMAN | MAPK9 | Mitogen-activated protein kinase 9 | 424 | 48.139 | 3.900944e+04 | 2.359304e+06 | 2047739.0 | 261420.8 | 50144.43 |  | 24.29 | 36.1 | 18.4 |  |
| sp\|Q9UHD1\|CHRD1\_HUMAN | Q9UHD1 | CHRD1\_HUMAN | CHORDC1 | Cysteine and histidine-rich domain-containing protein 1 | 332 | 37.490 | 1.777169e+05 | 8.370689e+06 | 2024797.0 | 6099665.0 | 246226.95 |  | 54.82 | 76.5 | 23.5 |  |
| sp\|P02462\|CO4A1\_HUMAN | P02462 | CO4A1\_HUMAN | COL4A1 | Collagen alpha-1(IV) chain | 1669 | 183.590 | 8.744116e+03 | 2.016889e+06 | 2016889.4 |  |  |  | 3.71 |  |  |  |
| sp\|P42025\|ACTY\_HUMAN | P42025 | ACTY\_HUMAN | ACTR1B | Beta-centractin | 376 | 42.293 | 8.671981e+04 | 4.607910e+06 | 2014397.5 | 1672136.5 | 696463.44 | 224912.27 | 45.21 | 48.1 | 19.7 | 10.4 |
| sp\|Q96PU5\|NED4L\_HUMAN | Q96PU5 | NED4L\_HUMAN | NEDD4L | E3 ubiquitin-protein ligase NEDD4-like | 975 | 111.932 | 2.467738e+04 | 3.470327e+06 | 2010661.8 | 1170507.8 | 257840.25 | 31317.396 | 39.28 | 45.1 | 21.4 | 13.6 |
| sp\|Q6GYQ0\|RGPA1\_HUMAN | Q6GYQ0 | RGPA1\_HUMAN | RALGAPA1 | Ral GTPase-activating protein subunit alpha-1 | 2036 | 229.832 | 7.469900e+03 | 2.156962e+06 | 2008800.5 | 148161.25 |  |  | 3.83 | 18.8 |  |  |
| sp\|Q7Z6P3\|RAB44\_HUMAN | Q7Z6P3 | RAB44\_HUMAN | RAB44 | Ras-related protein Rab-44 | 1021 | 112.310 | 1.421747e+04 | 2.006125e+06 | 2006125.2 |  |  |  | 2.35 |  |  |  |
| sp\|Q15046\|SYK\_HUMAN | Q15046 | SYK\_HUMAN | KARS1 | Lysine--tRNA ligase | 597 | 68.048 | 2.371056e+05 | 2.027096e+07 | 1992862.0 | 1375279.4 |  | 16902816.0 | 38.53 | 44.7 |  | 6.2 |
| sp\|P48556\|PSMD8\_HUMAN | P48556 | PSMD8\_HUMAN | PSMD8 | 26S proteasome non-ATPase regulatory subunit 8 | 350 | 39.612 | 1.361109e+05 | 6.773868e+06 | 1992567.8 | 3687937.0 | 805436.06 | 287927.44 | 46.0 | 41.1 | 20.0 | 20.0 |
| sp\|O75822\|EIF3J\_HUMAN | O75822 | EIF3J\_HUMAN | EIF3J | Eukaryotic translation initiation factor 3 subunit J | 258 | 29.062 | 1.842888e+05 | 6.728862e+06 | 1990466.1 | 3449110.5 | 690035.75 | 599249.25 | 36.82 | 58.5 | 38.4 | 29.1 |
| sp\|P11233\|RALA\_HUMAN | P11233 | RALA\_HUMAN | RALA | Ras-related protein Ral-A | 206 | 23.567 | 1.401055e+05 | 4.148363e+06 | 1986477.8 | 1619773.5 | 339612.03 | 202499.39 | 33.01 | 40.8 | 18.9 | 21.4 |
| sp\|Q15366\|PCBP2\_HUMAN | Q15366 | PCBP2\_HUMAN | PCBP2 | Poly(rC)-binding protein 2 | 365 | 38.580 | 1.015094e+05 | 4.920232e+06 | 1986003.5 | 953973.94 | 1070078.9 | 910175.8 | 43.56 | 65.8 | 35.1 | 18.1 |
| sp\|P68400\|CSK21\_HUMAN | P68400 | CSK21\_HUMAN | CSNK2A1 | Casein kinase II subunit alpha | 391 | 45.144 | 1.036993e+05 | 5.881564e+06 | 1957906.5 | 3817132.8 | 106524.95 |  | 53.2 | 53.2 | 23.3 |  |
| sp\|Q92621\|NU205\_HUMAN | Q92621 | NU205\_HUMAN | NUP205 | Nuclear pore complex protein Nup205 | 2012 | 227.922 | 7.831951e+03 | 2.242711e+06 | 1955677.9 | 269053.8 |  | 17979.65 | 5.47 | 23.8 |  | 3.4 |
| sp\|Q13107\|UBP4\_HUMAN | Q13107 | UBP4\_HUMAN | USP4 | Ubiquitin carboxyl-terminal hydrolase 4 | 963 | 108.565 | 1.870370e+04 | 2.551141e+06 | 1950803.6 | 387295.62 | 213041.69 |  | 32.5 | 39.3 | 16.8 |  |
| sp\|Q9BY43\|CHM4A\_HUMAN | Q9BY43 | CHM4A\_HUMAN | CHMP4A | Charged multivesicular body protein 4a | 222 | 25.098 | 3.561900e+05 | 1.123151e+07 | 1947959.0 | 8158054.0 | 650379.25 | 475112.94 | 45.05 | 56.3 | 44.6 | 39.2 |
| sp\|Q96AT9\|RPE\_HUMAN | Q96AT9 | RPE\_HUMAN | RPE | Ribulose-phosphate 3-epimerase | 228 | 24.928 | 1.312577e+05 | 4.110827e+06 | 1946407.1 | 2020929.5 | 143490.55 |  | 35.53 | 52.6 | 18.0 |  |
| sp\|P61204\|ARF3\_HUMAN | P61204 | ARF3\_HUMAN | ARF3 | ADP-ribosylation factor 3 | 181 | 20.601 | 4.085674e+05 | 1.057473e+07 | 1946289.8 |  | 5627639.0 | 3000797.2 | 59.67 |  | 49.2 | 33.7 |
| sp\|P30622\|CLIP1\_HUMAN | P30622 | CLIP1\_HUMAN | CLIP1 | CAP-Gly domain-containing linker protein 1 | 1438 | 162.246 | 1.693620e+04 | 3.452289e+06 | 1942353.2 | 98564.484 | 1411371.8 |  | 4.87 | 31.7 | 20.9 |  |
| sp\|Q6ICG6\|K0930\_HUMAN | Q6ICG6 | K0930\_HUMAN | KIAA0930 | Uncharacterized protein KIAA0930 | 404 | 45.794 | 4.539941e+04 | 2.612016e+06 | 1941008.1 | 378662.38 | 280439.44 | 11906.496 | 20.54 | 48.8 | 33.9 | 10.1 |
| sp\|Q9UMX0\|UBQL1\_HUMAN | Q9UMX0 | UBQL1\_HUMAN | UBQLN1 | Ubiquilin-1 | 589 | 62.519 | 4.644640e+04 | 3.648222e+06 | 1938073.0 | 1710149.0 |  |  | 28.18 | 33.3 |  |  |
| sp\|Q12765\|SCRN1\_HUMAN | Q12765 | SCRN1\_HUMAN | SCRN1 | Secernin-1 | 414 | 46.382 | 1.302731e+05 | 7.591390e+06 | 1935993.5 | 5469631.5 | 185765.11 |  | 40.1 | 45.4 | 21.7 |  |
| sp\|O95861\|BPNT1\_HUMAN | O95861 | BPNT1\_HUMAN | BPNT1 | 3'(2'),5'-bisphosphate nucleotidase 1 | 308 | 33.392 | 1.107731e+05 | 4.647227e+06 | 1935930.6 | 2246363.8 | 373009.66 | 91923.35 | 65.91 | 72.1 | 43.2 | 22.4 |
| sp\|Q13561\|DCTN2\_HUMAN | Q13561 | DCTN2\_HUMAN | DCTN2 | Dynactin subunit 2 | 401 | 44.231 | 1.538601e+05 | 8.550073e+06 | 1930128.5 | 5462301.0 | 831679.94 | 325963.7 | 60.85 | 75.8 | 32.7 | 19.2 |
| sp\|P26599\|PTBP1\_HUMAN | P26599 | PTBP1\_HUMAN | PTBP1 | Polypyrimidine tract-binding protein 1 | 557 | 59.633 | 8.861458e+04 | 6.639097e+06 | 1929873.2 | 3397303.0 | 1122952.9 | 188967.56 | 56.91 | 59.5 | 20.0 | 9.4 |
| sp\|Q8WW22\|DNJA4\_HUMAN | Q8WW22 | DNJA4\_HUMAN | DNAJA4 | DnaJ homolog subfamily A member 4 | 397 | 44.798 | 1.977191e+05 | 1.112819e+07 | 1922626.5 | 6549670.0 | 1951732.6 | 704156.8 | 43.83 | 67.8 | 42.3 | 30.0 |
| sp\|P15170\|ERF3A\_HUMAN | P15170 | ERF3A\_HUMAN | GSPT1 | Eukaryotic peptide chain release factor GTP-binding subunit ERF3A | 499 | 55.756 | 4.926882e+04 | 3.451286e+06 | 1916165.6 | 1141773.2 | 393346.94 |  | 51.3 | 55.5 | 28.1 |  |
| sp\|Q15404\|RSU1\_HUMAN | Q15404 | RSU1\_HUMAN | RSU1 | Ras suppressor protein 1 | 277 | 31.540 | 1.900379e+05 | 7.530417e+06 | 1907729.0 | 4583976.0 | 1038711.9 |  | 42.6 | 72.6 | 27.1 |  |
| sp\|Q2TAA2\|IAH1\_HUMAN | Q2TAA2 | IAH1\_HUMAN | IAH1 | Isoamyl acetate-hydrolyzing esterase 1 homolog | 248 | 27.599 | 1.359761e+05 | 4.714906e+06 | 1907408.5 | 2570256.5 | 237241.22 |  | 35.48 | 62.9 | 19.8 |  |
| sp\|Q8NBF2\|NHLC2\_HUMAN | Q8NBF2 | NHLC2\_HUMAN | NHLRC2 | NHL repeat-containing protein 2 | 726 | 79.444 | 4.009775e+04 | 4.002195e+06 | 1905395.2 | 1703949.0 | 268811.38 | 124039.234 | 47.25 | 48.5 | 15.8 | 7.6 |
| sp\|P46934\|NEDD4\_HUMAN | P46934 | NEDD4\_HUMAN | NEDD4 | E3 ubiquitin-protein ligase NEDD4 | 1319 | 149.114 | 2.271880e+04 | 4.256189e+06 | 1882448.8 | 2025081.8 | 298081.28 | 50577.15 | 27.9 | 32.2 | 23.1 | 10.2 |
| sp\|Q9NSK0\|KLC4\_HUMAN | Q9NSK0 | KLC4\_HUMAN | KLC4 | Kinesin light chain 4 | 619 | 68.640 | 2.718106e+04 | 2.344018e+06 | 1873941.4 | 403897.7 | 66178.51 |  | 41.03 | 50.1 | 18.3 |  |
| sp\|P61081\|UBC12\_HUMAN | P61081 | UBC12\_HUMAN | UBE2M | NEDD8-conjugating enzyme Ubc12 | 183 | 20.900 | 3.444295e+05 | 9.044068e+06 | 1873875.5 | 6009215.5 | 905193.0 | 255784.48 | 43.72 | 51.9 | 45.4 | 26.8 |
| sp\|P46734\|MP2K3\_HUMAN | P46734 | MP2K3\_HUMAN | MAP2K3 | Dual specificity mitogen-activated protein kinase kinase 3 | 347 | 39.318 | 2.616711e+05 | 1.292600e+07 | 1857110.2 | 3877184.8 | 6885284.0 | 306424.03 | 65.13 | 62.8 | 38.6 | 28.8 |
| sp\|P08754\|GNAI3\_HUMAN | P08754 | GNAI3\_HUMAN | GNAI3 | Guanine nucleotide-binding protein G(i) subunit alpha-3 | 354 | 40.532 | 3.943940e+05 | 2.008378e+07 | 1836294.4 | 2200037.2 | 6577047.5 | 9470403.0 | 52.54 | 64.1 | 41.2 | 33.1 |
| sp\|P51149\|RAB7A\_HUMAN | P51149 | RAB7A\_HUMAN | RAB7A | Ras-related protein Rab-7a | 207 | 23.490 | 1.610094e+05 | 4.751725e+06 | 1834103.5 | 1590396.5 | 885613.0 | 441612.4 | 67.15 | 75.8 | 57.5 | 25.1 |
| sp\|P34913\|HYES\_HUMAN | P34913 | HYES\_HUMAN | EPHX2 | Bifunctional epoxide hydrolase 2 | 555 | 62.616 | 2.530905e+04 | 1.991032e+06 | 1820412.9 | 170619.53 |  |  | 16.76 | 35.0 |  |  |
| sp\|Q15691\|MARE1\_HUMAN | Q15691 | MARE1\_HUMAN | MAPRE1 | Microtubule-associated protein RP/EB family member 1 | 268 | 29.999 | 3.148178e+05 | 1.186542e+07 | 1812074.4 | 8718233.0 | 1010436.3 | 324677.84 | 67.91 | 75.4 | 44.0 | 25.7 |
| sp\|Q9UI26\|IPO11\_HUMAN | Q9UI26 | IPO11\_HUMAN | IPO11 | Importin-11 | 975 | 112.535 | 3.837911e+04 | 5.426248e+06 | 1810665.8 | 2892400.0 | 181915.34 | 541266.6 | 38.15 | 41.6 | 17.5 | 10.1 |
| sp\|P07237\|PDIA1\_HUMAN | P07237 | PDIA1\_HUMAN | P4HB | Protein disulfide-isomerase | 508 | 57.116 | 1.400930e+05 | 1.005290e+07 | 1807959.0 | 4513085.5 | 3074585.8 | 657267.9 | 54.33 | 65.2 | 48.8 | 28.5 |
| sp\|P04899\|GNAI2\_HUMAN | P04899 | GNAI2\_HUMAN | GNAI2 | Guanine nucleotide-binding protein G(i) subunit alpha-2 | 355 | 40.451 | 3.631543e+05 | 1.845600e+07 | 1804396.8 | 2487433.5 | 5799272.5 | 8364898.5 | 66.48 | 66.2 | 43.9 | 28.5 |
| sp\|P68104\|EF1A1\_HUMAN | P68104 | EF1A1\_HUMAN | EEF1A1 | Elongation factor 1-alpha 1 | 462 | 50.141 | 9.350540e+04 | 5.890428e+06 | 1803659.8 | 2352154.5 | 1051019.2 | 683594.1 | 32.47 | 36.8 | 31.8 | 28.6 |
| sp\|P04083\|ANXA1\_HUMAN | P04083 | ANXA1\_HUMAN | ANXA1 | Annexin A1 | 346 | 38.714 | 1.761791e+05 | 8.569188e+06 | 1800873.4 | 1882600.0 | 3815564.8 | 1070150.2 | 61.56 | 79.2 | 57.2 | 22.3 |
| sp\|P42566\|EPS15\_HUMAN | P42566 | EPS15\_HUMAN | EPS15 | Epidermal growth factor receptor substrate 15 | 896 | 98.656 | 6.690022e+04 | 8.292170e+06 | 1799198.8 | 1465134.2 | 4451702.5 | 576135.0 | 58.04 | 61.2 | 37.3 | 20.0 |
| sp\|Q9BVL4\|SELO\_HUMAN | Q9BVL4 | SELO\_HUMAN | SELENOO | Protein adenylyltransferase SelO, mitochondrial | 669 | 73.489 | 5.139919e+04 | 4.745651e+06 | 1798785.2 | 2582765.5 | 185237.12 | 178862.69 | 28.4 | 47.4 | 28.8 | 18.4 |
| sp\|O15484\|CAN5\_HUMAN | O15484 | CAN5\_HUMAN | CAPN5 | Calpain-5 | 640 | 73.169 | 9.836074e+04 | 9.042032e+06 | 1797365.2 | 2268762.0 | 1542895.8 | 3433009.5 | 48.59 | 46.7 | 29.1 | 22.3 |
| sp\|P0DP23\|CALM1\_HUMAN | P0DP23 | CALM1\_HUMAN | CALM1 | Calmodulin-1 | 149 | 16.838 | 6.068293e+05 | 1.283732e+07 | 1794094.8 | 1010186.1 | 8874783.0 | 1158251.8 | 44.97 | 52.3 | 65.1 | 42.3 |
| sp\|Q9H4M9\|EHD1\_HUMAN | Q9H4M9 | EHD1\_HUMAN | EHD1 | EH domain-containing protein 1 | 534 | 60.627 | 1.167660e+05 | 8.894055e+06 | 1789355.5 | 5696560.5 | 1107159.8 | 300979.2 | 69.29 | 71.0 | 31.8 | 15.4 |
| sp\|P81605\|DCD\_HUMAN | P81605 | DCD\_HUMAN | DCD | Dermcidin | 110 | 11.284 | 7.582236e+05 | 1.074923e+07 | 1783117.6 | 411883.6 | 1887521.9 | 6666711.0 | 47.27 | 47.3 | 31.8 | 31.8 |
| sp\|Q09666\|AHNK\_HUMAN | Q09666 | AHNK\_HUMAN | AHNAK | Neuroblast differentiation-associated protein AHNAK | 5890 | 629.101 | 3.965262e+03 | 3.134075e+06 | 1781256.5 | 134078.84 | 713289.6 | 505450.5 | 3.07 | 64.7 | 43.9 | 34.0 |
| sp\|Q15907\|RB11B\_HUMAN | Q15907 | RB11B\_HUMAN | RAB11B | Ras-related protein Rab-11B | 218 | 24.489 | 2.470979e+05 | 7.602515e+06 | 1780250.6 | 4953260.0 | 869004.56 |  | 61.01 | 51.4 | 33.9 |  |
| sp\|P37840\|SYUA\_HUMAN | P37840 | SYUA\_HUMAN | SNCA | Alpha-synuclein | 140 | 14.460 | 5.335367e+06 | 9.692811e+07 | 1774936.8 | 32609990.0 | 50819396.0 | 11723785.0 | 61.43 | 72.9 | 89.3 | 60.0 |
| sp\|Q8WZA9\|IRGQ\_HUMAN | Q8WZA9 | IRGQ\_HUMAN | IRGQ | Immunity-related GTPase family Q protein | 623 | 62.717 | 6.441582e+04 | 5.075688e+06 | 1774906.6 | 2352335.0 | 815070.2 | 133376.44 | 43.66 | 37.7 | 25.4 | 19.9 |
| sp\|Q96RS6\|NUDC1\_HUMAN | Q96RS6 | NUDC1\_HUMAN | NUDCD1 | NudC domain-containing protein 1 | 583 | 66.756 | 4.989078e+04 | 4.184348e+06 | 1763110.0 | 2421237.5 |  |  | 49.06 | 58.8 |  |  |
| sp\|Q9Y5K5\|UCHL5\_HUMAN | Q9Y5K5 | UCHL5\_HUMAN | UCHL5 | Ubiquitin carboxyl-terminal hydrolase isozyme L5 | 329 | 37.607 | 7.616591e+04 | 3.598707e+06 | 1754329.2 | 1187892.6 | 542379.1 | 114105.83 | 57.45 | 57.1 | 40.1 | 21.9 |
| sp\|Q07866\|KLC1\_HUMAN | Q07866 | KLC1\_HUMAN | KLC1 | Kinesin light chain 1 | 573 | 65.310 | 3.228484e+04 | 2.649083e+06 | 1752488.2 | 666201.94 | 144822.64 | 85569.74 | 30.37 | 44.0 | 25.5 | 21.1 |
| sp\|O95573\|ACSL3\_HUMAN | O95573 | ACSL3\_HUMAN | ACSL3 | Fatty acid CoA ligase Acsl3 | 720 | 80.420 | 1.313102e+05 | 1.326721e+07 | 1741696.0 | 1419390.5 | 170633.6 | 9935491.0 | 47.5 | 48.8 | 23.6 | 13.2 |
| sp\|O95347\|SMC2\_HUMAN | O95347 | SMC2\_HUMAN | SMC2 | Structural maintenance of chromosomes protein 2 | 1197 | 135.656 | 2.118270e+04 | 3.610252e+06 | 1739836.6 | 1553758.1 | 207207.39 | 109449.65 | 45.11 | 46.9 | 21.9 | 11.4 |
| sp\|P26639\|SYTC\_HUMAN | P26639 | SYTC\_HUMAN | TARS1 | Threonine--tRNA ligase 1, cytoplasmic | 723 | 83.435 | 4.737874e+04 | 4.966483e+06 | 1737823.6 | 3064595.8 | 164063.17 |  | 54.08 | 56.4 | 24.6 |  |
| sp\|P54136\|SYRC\_HUMAN | P54136 | SYRC\_HUMAN | RARS1 | Arginine--tRNA ligase, cytoplasmic | 660 | 75.379 | 3.601613e+04 | 3.410866e+06 | 1737447.5 | 1673418.1 |  |  | 55.76 | 60.2 |  |  |
| sp\|Q08554\|DSC1\_HUMAN | Q08554 | DSC1\_HUMAN | DSC1 | Desmocollin-1 | 894 | 99.987 | 5.693012e+04 | 7.151593e+06 | 1732766.1 | 1760173.1 | 1506065.0 | 2152588.5 | 17.34 | 34.2 | 19.9 | 20.5 |
| sp\|Q9Y4P8\|WIPI2\_HUMAN | Q9Y4P8 | WIPI2\_HUMAN | WIPI2 | WD repeat domain phosphoinositide-interacting protein 2 | 454 | 49.408 | 6.800759e+04 | 4.221549e+06 | 1726874.8 | 1165815.4 | 1049230.4 | 279628.06 | 38.99 | 60.4 | 23.8 | 6.2 |
| sp\|O75688\|PPM1B\_HUMAN | O75688 | PPM1B\_HUMAN | PPM1B | Protein phosphatase 1B | 479 | 52.643 | 7.902733e+04 | 5.226790e+06 | 1713460.5 | 3177202.2 | 189215.89 | 146911.9 | 39.25 | 61.8 | 32.2 | 27.3 |
| sp\|P05455\|LA\_HUMAN | P05455 | LA\_HUMAN | SSB | Lupus La protein | 408 | 46.837 | 9.026981e+04 | 5.311886e+06 | 1713124.9 | 3311055.5 | 130858.984 | 156846.64 | 56.37 | 61.3 | 24.3 | 19.1 |
| sp\|O60749\|SNX2\_HUMAN | O60749 | SNX2\_HUMAN | SNX2 | Sorting nexin-2 | 519 | 58.471 | 6.415674e+04 | 4.713027e+06 | 1710204.8 | 2148723.5 | 585594.5 | 268504.25 | 45.66 | 48.9 | 38.3 | 14.1 |
| sp\|Q8N3D4\|EH1L1\_HUMAN | Q8N3D4 | EH1L1\_HUMAN | EHBP1L1 | EH domain-binding protein 1-like protein 1 | 1523 | 161.854 | 3.654612e+04 | 7.431593e+06 | 1693237.1 | 4048293.0 | 1026837.75 | 663224.75 | 54.37 | 55.8 | 37.9 | 23.2 |
| sp\|Q9UKU0\|ACSL6\_HUMAN | Q9UKU0 | ACSL6\_HUMAN | ACSL6 | Long-chain-fatty-acid--CoA ligase 6 | 697 | 77.752 | 4.527764e+04 | 4.422955e+06 | 1689451.2 | 1206839.2 | 669792.56 | 856871.8 | 45.34 | 54.5 | 32.1 | 14.8 |
| sp\|P61221\|ABCE1\_HUMAN | P61221 | ABCE1\_HUMAN | ABCE1 | ATP-binding cassette sub-family E member 1 | 599 | 67.314 | 5.595323e+04 | 4.732032e+06 | 1687697.5 | 2528626.2 | 412568.8 | 103139.72 | 44.24 | 53.9 | 21.4 | 16.4 |
| sp\|Q9UN37\|VPS4A\_HUMAN | Q9UN37 | VPS4A\_HUMAN | VPS4A | Vacuolar protein sorting-associated protein 4A | 437 | 48.898 | 7.555822e+04 | 4.641838e+06 | 1685886.9 | 1983412.8 | 817244.8 | 155293.61 | 56.52 | 63.2 | 28.8 | 19.9 |
| sp\|P51811\|XK\_HUMAN | P51811 | XK\_HUMAN | XK | Endoplasmic reticulum membrane adapter protein XK | 444 | 50.902 | 2.815023e+05 | 1.800255e+07 | 1685608.6 | 515832.6 | 2802707.0 | 12998401.0 | 23.2 | 25.7 | 23.0 | 18.9 |
| sp\|Q9Y4D1\|DAAM1\_HUMAN | Q9Y4D1 | DAAM1\_HUMAN | DAAM1 | Disheveled-associated activator of morphogenesis 1 | 1078 | 123.473 | 6.200650e+04 | 9.618921e+06 | 1678216.9 | 1390613.1 | 1404454.4 | 5145636.5 | 49.63 | 49.0 | 39.1 | 32.2 |
| sp\|Q9H2K8\|TAOK3\_HUMAN | Q9H2K8 | TAOK3\_HUMAN | TAOK3 | Serine/threonine-protein kinase TAO3 | 898 | 105.406 | 2.729410e+04 | 3.614525e+06 | 1670779.2 | 1536199.6 | 308599.84 | 98946.08 | 40.98 | 42.0 | 31.5 | 20.3 |
| sp\|O43314\|VIP2\_HUMAN | O43314 | VIP2\_HUMAN | PPIP5K2 | Inositol hexakisphosphate and diphosphoinositol-pentakisphosphate kinase 2 | 1243 | 140.407 | 1.810246e+04 | 3.193328e+06 | 1665525.8 | 1389918.0 | 137883.9 |  | 39.5 | 49.1 | 21.8 |  |
| sp\|Q14693\|LPIN1\_HUMAN | Q14693 | LPIN1\_HUMAN | LPIN1 | Phosphatidate phosphatase LPIN1 | 890 | 97.900 | 1.353824e+04 | 1.665183e+06 | 1665183.1 |  |  |  | 3.82 |  |  |  |
| sp\|P52594\|AGFG1\_HUMAN | P52594 | AGFG1\_HUMAN | AGFG1 | Arf-GAP domain and FG repeat-containing protein 1 | 562 | 58.260 | 7.290564e+04 | 5.336405e+06 | 1659834.0 | 1745371.2 | 1672639.5 | 258559.88 | 35.59 | 40.2 | 24.9 | 10.1 |
| sp\|Q05639\|EF1A2\_HUMAN | Q05639 | EF1A2\_HUMAN | EEF1A2 | Elongation factor 1-alpha 2 | 463 | 50.930 | 2.590987e+04 | 1.657892e+06 | 1657891.5 |  |  |  | 22.03 |  |  |  |
| sp\|Q86TP1\|PRUN1\_HUMAN | Q86TP1 | PRUN1\_HUMAN | PRUNE1 | Exopolyphosphatase PRUNE1 | 453 | 50.200 | 1.264149e+05 | 7.972952e+06 | 1651060.8 | 5943953.5 | 256839.19 | 121098.86 | 33.55 | 49.7 | 28.3 | 15.2 |
| sp\|P61960\|UFM1\_HUMAN | P61960 | UFM1\_HUMAN | UFM1 | Ubiquitin-fold modifier 1 | 85 | 9.118 | 1.066806e+06 | 1.222087e+07 | 1648087.5 | 10355448.0 | 217334.0 |  | 65.88 | 72.9 | 43.5 |  |
| sp\|O94829\|IPO13\_HUMAN | O94829 | IPO13\_HUMAN | IPO13 | Importin-13 | 963 | 108.195 | 1.508050e+04 | 2.049936e+06 | 1645519.4 | 281572.53 |  | 122843.72 | 30.74 | 36.8 |  | 7.8 |
| sp\|Q9H9Q2\|CSN7B\_HUMAN | Q9H9Q2 | CSN7B\_HUMAN | COPS7B | COP9 signalosome complex subunit 7b | 264 | 29.622 | 1.924052e+05 | 7.160581e+06 | 1643998.8 | 4575651.0 | 708539.9 | 232391.58 | 36.36 | 42.8 | 23.9 | 19.3 |
| sp\|P13693\|TCTP\_HUMAN | P13693 | TCTP\_HUMAN | TPT1 | Translationally-controlled tumor protein | 172 | 19.595 | 2.042828e+05 | 5.029146e+06 | 1643857.2 | 3190012.8 | 195276.5 |  | 33.14 | 93.0 | 39.5 |  |
| sp\|Q9UBC2\|EP15R\_HUMAN | Q9UBC2 | EP15R\_HUMAN | EPS15L1 | Epidermal growth factor receptor substrate 15-like 1 | 864 | 94.255 | 8.415840e+04 | 9.965958e+06 | 1632667.9 | 4005998.2 | 3649514.2 | 677777.3 | 60.88 | 56.5 | 56.5 | 33.7 |
| sp\|Q92598\|HS105\_HUMAN | Q92598 | HS105\_HUMAN | HSPH1 | Heat shock protein 105 kDa | 858 | 96.865 | 4.827457e+04 | 5.874927e+06 | 1632218.5 | 3985818.0 | 175308.12 | 81581.93 | 54.2 | 58.0 | 27.5 | 14.3 |
| sp\|P21266\|GSTM3\_HUMAN | P21266 | GSTM3\_HUMAN | GSTM3 | Glutathione S-transferase Mu 3 | 225 | 26.560 | 1.276180e+05 | 4.258504e+06 | 1627257.5 | 2396429.2 | 234817.42 |  | 63.11 | 82.2 | 20.9 |  |
| sp\|Q9NZL4\|HPBP1\_HUMAN | Q9NZL4 | HPBP1\_HUMAN | HSPBP1 | Hsp70-binding protein 1 | 359 | 39.303 | 1.087464e+05 | 5.369797e+06 | 1621594.4 | 3425982.0 | 322220.25 |  | 48.75 | 61.3 | 14.8 |  |
| sp\|Q9NZD4\|AHSP\_HUMAN | Q9NZD4 | AHSP\_HUMAN | AHSP | Alpha-hemoglobin-stabilizing protein | 102 | 11.840 | 6.085445e+05 | 9.052347e+06 | 1614304.5 | 1653763.4 | 3440758.0 | 2343521.5 | 36.27 | 48.0 | 42.2 | 50.0 |
| sp\|Q9ULC4\|MCTS1\_HUMAN | Q9ULC4 | MCTS1\_HUMAN | MCTS1 | Malignant T-cell-amplified sequence 1 | 181 | 20.555 | 1.706976e+05 | 4.408209e+06 | 1608090.8 | 2440997.0 | 359121.0 |  | 66.3 | 79.6 | 31.5 |  |
| sp\|O75695\|XRP2\_HUMAN | O75695 | XRP2\_HUMAN | RP2 | Protein XRP2 | 350 | 39.641 | 1.587430e+05 | 7.905992e+06 | 1601578.2 | 1616518.1 | 2679813.5 | 2008082.5 | 36.29 | 48.0 | 36.9 | 29.1 |
| sp\|O43149\|ZZEF1\_HUMAN | O43149 | ZZEF1\_HUMAN | ZZEF1 | Zinc finger ZZ-type and EF-hand domain-containing protein 1 | 2961 | 331.075 | 7.706568e+03 | 3.205565e+06 | 1598141.5 | 1348465.2 | 55985.355 | 202972.69 | 35.09 | 37.7 | 13.6 | 7.3 |
| sp\|P26447\|S10A4\_HUMAN | P26447 | S10A4\_HUMAN | S100A4 | Protein S100-A4 | 101 | 11.729 | 2.454309e+06 | 3.616657e+07 | 1583313.9 | 27517592.0 | 2640349.0 | 4425320.0 | 55.45 | 65.3 | 43.6 | 44.6 |
| sp\|P10114\|RAP2A\_HUMAN | P10114 | RAP2A\_HUMAN | RAP2A | Ras-related protein Rap-2a | 183 | 20.615 | 1.014643e+06 | 2.627930e+07 | 1569950.5 | 2095378.0 | 8376547.0 | 14237429.0 | 69.4 | 65.6 | 48.6 | 34.4 |
| sp\|P51784\|UBP11\_HUMAN | P51784 | UBP11\_HUMAN | USP11 | Ubiquitin carboxyl-terminal hydrolase 11 | 963 | 109.817 | 1.876115e+04 | 2.588488e+06 | 1566733.8 | 756140.6 | 256000.84 | 9612.422 | 39.88 | 36.6 | 22.9 | 8.3 |
| sp\|Q9HAB8\|PPCS\_HUMAN | Q9HAB8 | PPCS\_HUMAN | PPCS | Phosphopantothenate--cysteine ligase | 311 | 34.005 | 1.367309e+05 | 5.841532e+06 | 1556129.8 | 3505955.0 | 697501.25 | 81945.56 | 50.16 | 51.1 | 41.5 | 18.3 |
| sp\|Q9BZE9\|ASPC1\_HUMAN | Q9BZE9 | ASPC1\_HUMAN | ASPSCR1 | Tether containing UBX domain for GLUT4 | 553 | 60.183 | 6.934018e+04 | 5.242953e+06 | 1551618.8 | 3222152.8 | 336307.5 | 132873.95 | 52.98 | 59.1 | 46.5 | 28.2 |
| sp\|Q96L46\|CPNS2\_HUMAN | Q96L46 | CPNS2\_HUMAN | CAPNS2 | Calpain small subunit 2 | 248 | 27.660 | 2.866158e+05 | 9.960233e+06 | 1550785.9 |  | 3252153.8 | 5157293.0 | 15.73 |  | 29.4 | 24.6 |
| sp\|Q13464\|ROCK1\_HUMAN | Q13464 | ROCK1\_HUMAN | ROCK1 | Rho-associated protein kinase 1 | 1354 | 158.175 | 1.827509e+04 | 3.631739e+06 | 1538089.8 | 1347852.0 | 620992.9 | 124803.99 | 41.88 | 46.8 | 34.0 | 21.0 |
| sp\|P51692\|STA5B\_HUMAN | P51692 | STA5B\_HUMAN | STAT5B | Signal transducer and activator of transcription 5B | 787 | 89.866 | 6.138021e+04 | 6.930122e+06 | 1532592.2 | 3809079.2 | 1217898.8 | 370552.25 | 53.37 | 47.3 | 30.9 | 13.7 |
| sp\|Q9BXD5\|NPL\_HUMAN | Q9BXD5 | NPL\_HUMAN | NPL | N-acetylneuraminate lyase | 320 | 35.163 | 9.610951e+04 | 4.245897e+06 | 1528839.0 | 2293646.5 | 423411.2 |  | 58.44 | 70.0 | 27.8 |  |
| sp\|Q9NP59\|S40A1\_HUMAN | Q9NP59 | S40A1\_HUMAN | SLC40A1 | Solute carrier family 40 member 1 | 571 | 62.542 | 3.261418e+05 | 2.562686e+07 | 1525521.0 | 1978993.8 | 6462757.5 | 15659584.0 | 13.84 | 22.4 | 18.0 | 16.6 |
| sp\|P0CF74\|IGLC6\_HUMAN | P0CF74 | IGLC6\_HUMAN | IGLC6 | Immunoglobulin lambda constant 6 | 106 | 11.660 | 1.040347e+05 | 1.524031e+06 | 1524031.0 |  |  |  | 47.17 |  |  |  |
| sp\|Q9HA65\|TBC17\_HUMAN | Q9HA65 | TBC17\_HUMAN | TBC1D17 | TBC1 domain family member 17 | 648 | 72.670 | 2.995217e+04 | 2.734643e+06 | 1506551.0 | 860363.56 | 171243.19 | 196485.23 | 24.85 | 38.4 | 16.8 | 8.6 |
| sp\|O14787\|TNPO2\_HUMAN | O14787 | TNPO2\_HUMAN | TNPO2 | Transportin-2 | 897 | 101.388 | 2.749534e+04 | 3.502376e+06 | 1506431.0 | 1995945.2 |  |  | 32.0 | 33.2 |  |  |
| sp\|Q9NRS6\|SNX15\_HUMAN | Q9NRS6 | SNX15\_HUMAN | SNX15 | Sorting nexin-15 | 342 | 38.291 | 9.501910e+04 | 4.571143e+06 | 1504984.1 | 2266808.2 | 605416.44 | 193933.97 | 61.11 | 52.9 | 27.5 | 29.8 |
| sp\|P21333\|FLNA\_HUMAN | P21333 | FLNA\_HUMAN | FLNA | Filamin-A | 2647 | 280.739 | 9.594653e+03 | 3.384147e+06 | 1498729.8 | 1506565.6 | 190918.55 | 187932.58 | 52.81 | 53.2 | 29.9 | 19.9 |
| sp\|P50552\|VASP\_HUMAN | P50552 | VASP\_HUMAN | VASP | Vasodilator-stimulated phosphoprotein | 380 | 39.830 | 1.154749e+05 | 5.778498e+06 | 1496052.4 | 3464900.5 | 474432.1 | 343113.22 | 37.89 | 49.2 | 33.4 | 24.7 |
| sp\|Q86TI2\|DPP9\_HUMAN | Q86TI2 | DPP9\_HUMAN | DPP9 | Dipeptidyl peptidase 9 | 863 | 98.263 | 4.394048e+04 | 5.424654e+06 | 1492423.1 | 3824473.5 | 107756.914 |  | 46.93 | 34.2 | 16.6 |  |
| sp\|P35240\|MERL\_HUMAN | P35240 | MERL\_HUMAN | NF2 | Merlin | 595 | 69.690 | 2.350081e+04 | 2.057645e+06 | 1491044.8 | 566600.1 |  |  | 31.09 | 50.6 |  |  |
| sp\|Q9Y6I3\|EPN1\_HUMAN | Q9Y6I3 | EPN1\_HUMAN | EPN1 | Epsin-1 | 576 | 60.293 | 6.983009e+04 | 5.289646e+06 | 1490784.5 | 2065615.4 | 1297715.5 | 435530.78 | 24.83 | 31.1 | 17.5 | 10.1 |
| sp\|P27797\|CALR\_HUMAN | P27797 | CALR\_HUMAN | CALR | Calreticulin | 417 | 48.142 | 3.348528e+05 | 2.025327e+07 | 1487544.9 | 3576421.8 | 12906818.0 | 2282485.5 | 47.72 | 43.4 | 42.9 | 20.6 |
| sp\|Q9Y4D8\|HECD4\_HUMAN | Q9Y4D8 | HECD4\_HUMAN | HECTD4 | Probable E3 ubiquitin-protein ligase HECTD4 | 3996 | 439.344 | 3.594541e+03 | 1.984108e+06 | 1480838.1 | 421293.16 |  | 81976.67 | 16.97 | 27.1 |  | 10.1 |
| sp\|Q9UHD9\|UBQL2\_HUMAN | Q9UHD9 | UBQL2\_HUMAN | UBQLN2 | Ubiquilin-2 | 624 | 65.696 | 4.054279e+04 | 3.346337e+06 | 1479166.9 | 1787111.5 | 63747.766 | 16310.852 | 24.36 | 23.1 | 12.5 | 5.0 |
| sp\|Q9HBL0\|TENS1\_HUMAN | Q9HBL0 | TENS1\_HUMAN | TNS1 | Tensin-1 | 1839 | 197.023 | 2.862574e+04 | 7.085831e+06 | 1477294.5 | 2748957.5 | 2094669.4 | 764909.4 | 25.18 | 44.1 | 27.9 | 17.9 |
| sp\|Q9BZ23\|PANK2\_HUMAN | Q9BZ23 | PANK2\_HUMAN | PANK2 | Pantothenate kinase 2, mitochondrial | 570 | 62.681 | 5.961216e+04 | 4.694484e+06 | 1473812.8 | 3060922.2 | 159749.5 |  | 35.44 | 63.0 | 26.5 |  |
| sp\|Q6P1N9\|TATD1\_HUMAN | Q6P1N9 | TATD1\_HUMAN | TATDN1 | Deoxyribonuclease TATDN1 | 297 | 33.602 | 8.624697e+04 | 3.641046e+06 | 1471261.5 | 1925403.2 | 166874.97 | 77505.94 | 59.26 | 78.1 | 33.0 | 21.9 |
| sp\|Q08AM6\|VAC14\_HUMAN | Q08AM6 | VAC14\_HUMAN | VAC14 | Protein VAC14 homolog | 782 | 87.973 | 6.219352e+04 | 6.874035e+06 | 1456652.2 | 4238141.5 | 549186.6 | 630054.25 | 47.95 | 63.9 | 17.9 | 14.8 |
| sp\|P19367\|HXK1\_HUMAN | P19367 | HXK1\_HUMAN | HK1 | Hexokinase-1 | 917 | 102.486 | 3.137532e+04 | 4.039893e+06 | 1451057.4 | 2326426.5 | 262408.88 |  | 39.59 | 40.2 | 17.8 |  |
| sp\|P20042\|IF2B\_HUMAN | P20042 | IF2B\_HUMAN | EIF2S2 | Eukaryotic translation initiation factor 2 subunit 2 | 333 | 38.388 | 1.158880e+05 | 5.589217e+06 | 1450263.6 | 3560792.8 | 273789.9 | 304370.84 | 58.86 | 67.6 | 48.0 | 36.3 |
| sp\|P08758\|ANXA5\_HUMAN | P08758 | ANXA5\_HUMAN | ANXA5 | Annexin A5 | 320 | 35.937 | 1.240365e+05 | 5.600264e+06 | 1448689.4 | 3145086.5 | 1006488.4 |  | 66.56 | 59.1 | 29.1 |  |
| sp\|O75387\|LAT3\_HUMAN | O75387 | LAT3\_HUMAN | SLC43A1 | Large neutral amino acids transporter small subunit 3 | 559 | 61.477 | 1.042364e+05 | 8.050991e+06 | 1436339.4 | 135536.02 | 1398896.5 | 5080219.0 | 13.77 | 12.9 | 21.8 | 13.6 |
| sp\|Q9Y3C8\|UFC1\_HUMAN | Q9Y3C8 | UFC1\_HUMAN | UFC1 | Ubiquitin-fold modifier-conjugating enzyme 1 | 167 | 19.458 | 1.578269e+05 | 3.858302e+06 | 1432904.1 | 1431846.6 | 809935.1 | 183616.44 | 37.13 | 65.3 | 46.1 | 5.4 |
| sp\|O95456\|PSMG1\_HUMAN | O95456 | PSMG1\_HUMAN | PSMG1 | Proteasome assembly chaperone 1 | 288 | 32.854 | 1.733969e+05 | 7.157258e+06 | 1426983.8 | 5235507.0 | 381875.5 | 112891.73 | 44.79 | 54.2 | 31.2 | 20.1 |
| sp\|P28161\|GSTM2\_HUMAN | P28161 | GSTM2\_HUMAN | GSTM2 | Glutathione S-transferase Mu 2 | 218 | 25.745 | 1.081211e+05 | 3.497200e+06 | 1424824.0 | 1850552.2 | 221823.5 |  | 73.85 | 88.1 | 44.5 |  |
| sp\|P16989\|YBOX3\_HUMAN | P16989 | YBOX3\_HUMAN | YBX3 | Y-box-binding protein 3 | 372 | 40.090 | 8.799790e+04 | 4.432263e+06 | 1415431.2 | 1352387.6 | 1415792.5 | 248651.52 | 46.24 | 55.1 | 52.2 | 24.5 |
| sp\|P49593\|PPM1F\_HUMAN | P49593 | PPM1F\_HUMAN | PPM1F | Protein phosphatase 1F | 454 | 49.831 | 7.780365e+04 | 4.870984e+06 | 1409745.2 | 3269676.5 | 191562.66 |  | 46.92 | 67.2 | 21.4 |  |
| sp\|P42858\|HD\_HUMAN | P42858 | HD\_HUMAN | HTT | Huntingtin | 3142 | 347.603 | 9.544834e+03 | 4.168397e+06 | 1405102.5 | 1738743.4 | 818674.4 | 205876.67 | 33.0 | 33.7 | 11.0 | 11.5 |
| sp\|A0AVT1\|UBA6\_HUMAN | A0AVT1 | UBA6\_HUMAN | UBA6 | Ubiquitin-like modifier-activating enzyme 6 | 1052 | 117.970 | 3.266640e+04 | 4.841612e+06 | 1400508.4 | 3227332.8 | 141651.48 | 72119.66 | 53.61 | 47.3 | 19.0 | 9.0 |
| sp\|P09488\|GSTM1\_HUMAN | P09488 | GSTM1\_HUMAN | GSTM1 | Glutathione S-transferase Mu 1 | 218 | 23.980 | 4.631633e+04 | 1.395406e+06 | 1395405.5 |  |  |  | 41.74 |  |  |  |
| sp\|Q03013\|GSTM4\_HUMAN | Q03013 | GSTM4\_HUMAN | GSTM4 | Glutathione S-transferase Mu 4 | 218 | 25.561 | 8.498343e+04 | 2.729162e+06 | 1395405.5 | 1333756.1 |  |  | 28.9 | 55.5 |  |  |
| sp\|Q9H2G2\|SLK\_HUMAN | Q9H2G2 | SLK\_HUMAN | SLK | STE20-like serine/threonine-protein kinase | 1235 | 142.695 | 2.325098e+04 | 4.168378e+06 | 1394916.5 | 2206560.0 | 321477.1 | 245424.9 | 50.77 | 54.6 | 24.5 | 20.6 |
| sp\|Q66K74\|MAP1S\_HUMAN | Q66K74 | MAP1S\_HUMAN | MAP1S | Microtubule-associated protein 1S | 1059 | 112.211 | 1.340163e+04 | 1.889341e+06 | 1394027.8 | 278272.8 | 217039.92 |  | 15.77 | 50.0 | 12.3 |  |
| sp\|P06730\|IF4E\_HUMAN | P06730 | IF4E\_HUMAN | EIF4E | Eukaryotic translation initiation factor 4E | 217 | 25.097 | 1.720398e+05 | 5.424602e+06 | 1389673.2 | 3214160.2 | 820768.6 |  | 35.02 | 50.2 | 42.4 |  |
| sp\|Q8TCD5\|NT5C\_HUMAN | Q8TCD5 | NT5C\_HUMAN | NT5C | 5'(3')-deoxyribonucleotidase, cytosolic type | 201 | 23.383 | 1.553711e+05 | 4.564442e+06 | 1386213.9 | 3154783.0 | 23444.717 |  | 43.28 | 82.1 | 20.4 |  |
| sp\|P27824\|CALX\_HUMAN | P27824 | CALX\_HUMAN | CANX | Calnexin | 592 | 67.568 | 2.131970e+05 | 1.809836e+07 | 1381768.2 | 2203038.8 | 8008587.0 | 6504968.5 | 33.78 | 39.4 | 40.7 | 34.0 |
| sp\|Q8NB90\|AFG2A\_HUMAN | Q8NB90 | AFG2A\_HUMAN | AFG2A | ATPase family gene 2 protein homolog A | 893 | 97.904 | 1.480593e+04 | 1.821181e+06 | 1381623.9 | 439557.34 |  |  | 9.74 | 38.7 |  |  |
| sp\|O15498\|YKT6\_HUMAN | O15498 | YKT6\_HUMAN | YKT6 | Synaptobrevin homolog YKT6 | 198 | 22.418 | 1.620747e+05 | 4.564879e+06 | 1381385.1 | 1553113.2 | 956347.1 | 674034.06 | 70.71 | 67.2 | 46.5 | 25.8 |
| sp\|Q9H8H3\|TMT1A\_HUMAN | Q9H8H3 | TMT1A\_HUMAN | TMT1A | N6-adenosine-methyltransferase TMT1A | 244 | 28.319 | 9.668606e+04 | 3.440004e+06 | 1379529.6 | 1557525.5 | 172437.12 | 330511.62 | 22.54 | 49.6 | 24.2 | 18.9 |
| sp\|Q9BS26\|ERP44\_HUMAN | Q9BS26 | ERP44\_HUMAN | ERP44 | Endoplasmic reticulum resident protein 44 | 406 | 46.971 | 1.741072e+05 | 1.027457e+07 | 1379102.0 | 640827.6 | 6624156.5 | 1630484.5 | 26.85 | 29.6 | 35.2 | 8.4 |
| sp\|P17927\|CR1\_HUMAN | P17927 | CR1\_HUMAN | CR1 | Complement receptor type 1 | 2039 | 223.663 | 1.907540e+04 | 5.360248e+06 | 1368624.2 | 592603.7 | 868965.56 | 2530055.0 | 17.95 | 36.2 | 31.7 | 18.1 |
| sp\|O95630\|STABP\_HUMAN | O95630 | STABP\_HUMAN | STAMBP | STAM-binding protein | 424 | 48.077 | 6.459800e+04 | 3.901877e+06 | 1368507.0 | 2144746.0 | 230875.0 | 157748.84 | 45.99 | 55.9 | 27.8 | 17.2 |
| sp\|Q96RL7\|VP13A\_HUMAN | Q96RL7 | VP13A\_HUMAN | VPS13A | Intermembrane lipid transfer protein VPS13A | 3174 | 360.276 | 1.425466e+04 | 6.452223e+06 | 1365212.5 | 2724298.2 | 1318734.1 | 1043978.4 | 34.66 | 40.1 | 23.9 | 17.5 |
| sp\|P05023\|AT1A1\_HUMAN | P05023 | AT1A1\_HUMAN | ATP1A1 | Sodium/potassium-transporting ATPase subunit alpha-1 | 1023 | 112.896 | 7.306512e+04 | 1.036348e+07 | 1361428.0 | 1007435.75 | 2236129.5 | 5758491.0 | 40.76 | 43.0 | 33.9 | 34.0 |
| sp\|P46108\|CRK\_HUMAN | P46108 | CRK\_HUMAN | CRK | Adapter molecule crk | 304 | 33.831 | 8.132882e+04 | 3.456818e+06 | 1361120.2 | 1904272.2 | 191425.34 |  | 65.13 | 64.8 | 28.9 |  |
| sp\|Q5R3I4\|TTC38\_HUMAN | Q5R3I4 | TTC38\_HUMAN | TTC38 | Tetratricopeptide repeat protein 38 | 469 | 52.787 | 6.668975e+04 | 4.422860e+06 | 1355049.9 | 3051669.0 |  | 16141.362 | 49.25 | 60.3 |  | 6.2 |
| sp\|P14317\|HCLS1\_HUMAN | P14317 | HCLS1\_HUMAN | HCLS1 | Hematopoietic lineage cell-specific protein | 486 | 54.014 | 4.352866e+04 | 2.953920e+06 | 1352357.8 | 1452259.2 | 149303.1 |  | 38.89 | 55.1 | 24.9 |  |
| sp\|P00813\|ADA\_HUMAN | P00813 | ADA\_HUMAN | ADA | Adenosine deaminase | 363 | 40.764 | 9.740800e+04 | 4.988713e+06 | 1347809.5 | 3269904.0 | 348799.44 | 22200.541 | 58.13 | 76.3 | 35.0 | 18.5 |
| sp\|P42229\|STA5A\_HUMAN | P42229 | STA5A\_HUMAN | STAT5A | Signal transducer and activator of transcription 5A | 794 | 90.647 | 3.612337e+04 | 4.113948e+06 | 1343749.8 | 2770198.2 |  |  | 48.11 | 47.7 |  |  |
| sp\|Q12846\|STX4\_HUMAN | Q12846 | STX4\_HUMAN | STX4 | Syntaxin-4 | 297 | 34.180 | 2.308798e+05 | 9.914598e+06 | 1343236.1 | 1586037.5 | 2028302.2 | 4957022.5 | 39.39 | 58.9 | 38.0 | 35.4 |
| sp\|Q9H0W9\|CK054\_HUMAN | Q9H0W9 | CK054\_HUMAN | C11orf54 | Ester hydrolase C11orf54 | 315 | 35.117 | 1.520065e+05 | 6.706513e+06 | 1337378.1 | 5303370.5 | 65763.94 |  | 38.73 | 49.5 | 16.5 |  |
| sp\|Q8WZA0\|LZIC\_HUMAN | Q8WZA0 | LZIC\_HUMAN | LZIC | Protein LZIC | 190 | 21.495 | 2.144275e+05 | 5.790755e+06 | 1337075.0 | 3415467.0 | 710182.6 | 328030.3 | 58.42 | 82.6 | 42.1 | 27.4 |
| sp\|Q7L8J4\|3BP5L\_HUMAN | Q7L8J4 | 3BP5L\_HUMAN | SH3BP5L | SH3 domain-binding protein 5-like | 393 | 43.499 | 2.554762e+04 | 1.396197e+06 | 1330820.0 | 65377.406 |  |  | 6.62 | 46.3 |  |  |
| sp\|O14562\|UBFD1\_HUMAN | O14562 | UBFD1\_HUMAN | UBFD1 | Ubiquitin domain-containing protein UBFD1 | 309 | 33.382 | 1.156349e+05 | 4.849741e+06 | 1328384.4 | 2930581.0 | 311015.34 | 279759.94 | 40.78 | 44.7 | 24.3 | 19.1 |
| sp\|Q9Y6M5\|ZNT1\_HUMAN | Q9Y6M5 | ZNT1\_HUMAN | SLC30A1 | Proton-coupled zinc antiporter SLC30A1 | 507 | 55.300 | 3.074310e+05 | 2.135945e+07 | 1326099.8 | 733885.5 | 5507476.5 | 13791985.0 | 30.57 | 34.5 | 24.3 | 21.5 |
| sp\|Q9H0P0\|5NT3A\_HUMAN | Q9H0P0 | 5NT3A\_HUMAN | NT5C3A | Cytosolic 5'-nucleotidase 3A | 336 | 37.948 | 7.223521e+04 | 3.443935e+06 | 1322623.0 | 1919180.5 | 202131.95 |  | 54.17 | 69.6 | 27.7 |  |
| sp\|P54252\|ATX3\_HUMAN | P54252 | ATX3\_HUMAN | ATXN3 | Ataxin-3 | 361 | 41.250 | 1.244869e+05 | 6.451561e+06 | 1319765.9 | 4164282.8 | 814605.56 | 152906.48 | 32.41 | 38.0 | 30.7 | 22.4 |
| sp\|P17931\|LEG3\_HUMAN | P17931 | LEG3\_HUMAN | LGALS3 | Galectin-3 | 250 | 26.152 | 8.937247e+05 | 2.936472e+07 | 1319633.9 | 1151317.2 | 25269308.0 | 1624457.0 | 25.6 | 33.6 | 38.4 | 22.4 |
| sp\|Q9Y376\|CAB39\_HUMAN | Q9Y376 | CAB39\_HUMAN | CAB39 | Calcium-binding protein 39 | 341 | 39.869 | 1.353419e+05 | 6.779301e+06 | 1317557.5 | 4557843.5 | 857103.6 | 46796.15 | 62.17 | 68.0 | 30.5 | 12.3 |
| sp\|P06396\|GELS\_HUMAN | P06396 | GELS\_HUMAN | GSN | Gelsolin | 782 | 85.698 | 4.719816e+04 | 5.081745e+06 | 1312059.6 | 2824267.2 | 421428.3 | 523990.03 | 37.47 | 53.2 | 29.7 | 16.6 |
| sp\|P33176\|KINH\_HUMAN | P33176 | KINH\_HUMAN | KIF5B | Kinesin-1 heavy chain | 963 | 109.685 | 2.900411e+04 | 3.996907e+06 | 1311777.8 | 2187660.5 | 308859.25 | 188609.08 | 58.36 | 69.3 | 38.2 | 26.2 |
| sp\|Q96A08\|H2B1A\_HUMAN | Q96A08 | H2B1A\_HUMAN | H2BC1 | Histone H2B type 1-A | 127 | 13.970 | 7.468058e+04 | 1.310754e+06 | 1310754.2 |  |  |  | 51.18 |  |  |  |
| sp\|Q9UIC8\|LCMT1\_HUMAN | Q9UIC8 | LCMT1\_HUMAN | LCMT1 | Leucine carboxyl methyltransferase 1 | 334 | 38.379 | 4.139504e+04 | 1.995993e+06 | 1304885.2 | 427873.6 | 206765.16 | 56469.305 | 40.72 | 58.7 | 24.3 | 13.2 |
| sp\|Q5JSH3\|WDR44\_HUMAN | Q5JSH3 | WDR44\_HUMAN | WDR44 | WD repeat-containing protein 44 | 913 | 101.366 | 2.858237e+04 | 3.640052e+06 | 1304198.0 | 1999557.6 | 256588.66 | 79708.14 | 69.0 | 54.3 | 31.3 | 19.9 |
| sp\|Q13153\|PAK1\_HUMAN | Q13153 | PAK1\_HUMAN | PAK1 | Serine/threonine-protein kinase PAK 1 | 545 | 60.647 | 4.784873e+04 | 3.645834e+06 | 1295132.8 | 1180199.4 | 1170502.0 |  | 46.42 | 42.9 | 25.0 |  |
| sp\|Q9NTJ3\|SMC4\_HUMAN | Q9NTJ3 | SMC4\_HUMAN | SMC4 | Structural maintenance of chromosomes protein 4 | 1288 | 147.182 | 1.558850e+04 | 2.882546e+06 | 1291072.8 | 1250967.8 | 231150.75 | 109354.51 | 45.19 | 51.3 | 27.8 | 20.6 |
| sp\|Q6DD88\|ATLA3\_HUMAN | Q6DD88 | ATLA3\_HUMAN | ATL3 | Atlastin-3 | 541 | 60.542 | 1.016955e+05 | 7.735270e+06 | 1289215.6 | 4182774.5 | 996875.1 | 1266405.0 | 39.0 | 35.7 | 43.1 | 21.4 |
| sp\|P61160\|ARP2\_HUMAN | P61160 | ARP2\_HUMAN | ACTR2 | Actin-related protein 2 | 394 | 44.761 | 4.697249e+04 | 2.641560e+06 | 1287695.0 | 991875.75 | 337112.25 | 24877.031 | 39.34 | 49.2 | 22.1 | 11.2 |
| sp\|O75396\|SC22B\_HUMAN | O75396 | SC22B\_HUMAN | SEC22B | Vesicle-trafficking protein SEC22b | 215 | 24.741 | 1.937388e+05 | 6.022142e+06 | 1285088.2 | 316539.06 | 911363.6 | 3509150.8 | 47.44 | 60.0 | 49.3 | 46.0 |
| sp\|P02768\|ALBU\_HUMAN | P02768 | ALBU\_HUMAN | ALB | Albumin | 609 | 69.367 | 1.595109e+05 | 1.390146e+07 | 1284554.8 | 3648235.0 | 5732798.0 | 3235869.5 | 37.11 | 69.8 | 62.6 | 54.8 |
| sp\|Q9NZN3\|EHD3\_HUMAN | Q9NZN3 | EHD3\_HUMAN | EHD3 | EH domain-containing protein 3 | 535 | 60.887 | 4.873017e+04 | 3.727689e+06 | 1284408.5 | 1684884.5 | 656423.56 | 101972.63 | 40.19 | 61.3 | 26.9 | 14.0 |
| sp\|Q5T447\|HECD3\_HUMAN | Q5T447 | HECD3\_HUMAN | HECTD3 | E3 ubiquitin-protein ligase HECTD3 | 861 | 97.113 | 2.598274e+04 | 3.170147e+06 | 1280337.6 | 1611764.8 | 216569.67 | 61474.93 | 48.43 | 50.5 | 21.7 | 19.9 |
| sp\|P54105\|ICLN\_HUMAN | P54105 | ICLN\_HUMAN | CLNS1A | Methylosome subunit pICln | 237 | 26.215 | 6.805356e+04 | 2.241392e+06 | 1279041.4 | 764694.94 | 197655.98 |  | 34.6 | 46.4 | 16.0 |  |
| sp\|P12081\|HARS1\_HUMAN | P12081 | HARS1\_HUMAN | HARS1 | Histidine--tRNA ligase, cytoplasmic | 509 | 57.411 | 3.768485e+04 | 2.718185e+06 | 1277036.2 | 1068908.8 | 239900.08 | 132339.94 | 36.74 | 42.6 | 24.4 | 21.8 |
| sp\|Q9NZD2\|GLTP\_HUMAN | Q9NZD2 | GLTP\_HUMAN | GLTP | Glycolipid transfer protein | 209 | 23.850 | 5.396341e+04 | 1.616981e+06 | 1274895.9 | 342085.0 |  |  | 10.53 | 44.0 |  |  |
| sp\|Q16186\|ADRM1\_HUMAN | Q16186 | ADRM1\_HUMAN | ADRM1 | Proteasomal ubiquitin receptor ADRM1 | 407 | 42.153 | 1.222705e+05 | 6.475413e+06 | 1271693.6 | 4795061.5 | 332555.28 | 76102.87 | 19.41 | 30.0 | 21.6 | 11.1 |
| sp\|Q92614\|MY18A\_HUMAN | Q92614 | MY18A\_HUMAN | MYO18A | Unconventional myosin-XVIIIa | 2054 | 233.115 | 1.533686e+04 | 4.491834e+06 | 1270082.0 | 2101455.5 | 871489.5 | 248807.14 | 39.68 | 52.0 | 22.0 | 15.1 |
| sp\|Q53TN4\|CYBR1\_HUMAN | Q53TN4 | CYBR1\_HUMAN | CYBRD1 | Plasma membrane ascorbate-dependent reductase CYBRD1 | 286 | 31.641 | 7.094991e+05 | 2.820455e+07 | 1265015.2 | 97487.64 | 5441309.0 | 21400738.0 | 13.99 | 24.8 | 22.7 | 22.7 |
| sp\|Q9UL25\|RAB21\_HUMAN | Q9UL25 | RAB21\_HUMAN | RAB21 | Ras-related protein Rab-21 | 225 | 24.348 | 2.362928e+05 | 7.228214e+06 | 1263870.2 | 3638753.0 | 1235856.0 | 1089734.5 | 45.78 | 70.2 | 51.1 | 25.3 |
| sp\|O75352\|MPU1\_HUMAN | O75352 | MPU1\_HUMAN | MPDU1 | Mannose-P-dolichol utilization defect 1 protein | 247 | 26.638 | 2.108939e+05 | 7.058017e+06 | 1257883.8 | 1316827.5 | 917540.0 | 3565765.8 | 13.36 | 26.7 | 8.1 | 8.5 |
| sp\|Q969X1\|LFG3\_HUMAN | Q969X1 | LFG3\_HUMAN | TMBIM1 | Protein lifeguard 3 | 311 | 34.607 | 5.147587e+05 | 2.238127e+07 | 1255645.5 | 1571618.8 | 7166747.5 | 12387262.0 | 4.82 | 28.6 | 9.0 | 9.0 |
| sp\|P50579\|MAP2\_HUMAN | P50579 | MAP2\_HUMAN | METAP2 | Methionine aminopeptidase 2 | 478 | 52.892 | 6.163864e+04 | 4.096002e+06 | 1254361.6 | 2423000.5 | 313805.12 | 104834.74 | 51.88 | 62.3 | 27.0 | 9.2 |
| sp\|Q9NRR5\|UBQL4\_HUMAN | Q9NRR5 | UBQL4\_HUMAN | UBQLN4 | Ubiquilin-4 | 601 | 63.853 | 2.302727e+04 | 1.847315e+06 | 1253127.5 | 497759.28 | 96428.37 |  | 32.95 | 28.5 | 13.1 |  |
| sp\|Q8WUD1\|RAB2B\_HUMAN | Q8WUD1 | RAB2B\_HUMAN | RAB2B | Ras-related protein Rab-2B | 216 | 24.214 | 1.058281e+05 | 3.219473e+06 | 1246296.5 | 1645110.9 | 215900.48 | 112165.6 | 73.61 | 69.9 | 63.4 | 32.9 |
| sp\|Q53GL7\|PAR10\_HUMAN | Q53GL7 | PAR10\_HUMAN | PARP10 | Protein mono-ADP-ribosyltransferase PARP10 | 1025 | 109.998 | 1.667324e+04 | 2.304210e+06 | 1242169.6 | 773069.25 | 230199.19 | 58771.75 | 40.0 | 48.1 | 18.6 | 11.8 |
| sp\|Q14376\|GALE\_HUMAN | Q14376 | GALE\_HUMAN | GALE | UDP-glucose 4-epimerase | 348 | 38.282 | 3.767819e+04 | 1.812182e+06 | 1236227.0 | 575955.2 |  |  | 41.09 | 55.2 |  |  |
| sp\|Q9UNP9\|PPIE\_HUMAN | Q9UNP9 | PPIE\_HUMAN | PPIE | Peptidyl-prolyl cis-trans isomerase E | 301 | 33.431 | 4.392727e+04 | 1.845019e+06 | 1226037.1 | 618981.56 |  |  | 19.27 | 50.5 |  |  |
| sp\|O60763\|USO1\_HUMAN | O60763 | USO1\_HUMAN | USO1 | General vesicular transport factor p115 | 962 | 107.895 | 2.817401e+04 | 3.819153e+06 | 1225449.0 | 1998620.6 | 165021.72 | 430061.88 | 43.45 | 34.9 | 18.3 | 15.3 |
| sp\|O94889\|KLH18\_HUMAN | O94889 | KLH18\_HUMAN | KLHL18 | Kelch-like protein 18 | 574 | 63.638 | 2.905443e+04 | 2.322983e+06 | 1220235.0 | 1017994.8 | 84753.055 |  | 23.69 | 34.0 | 8.9 |  |
| sp\|P12829\|MYL4\_HUMAN | P12829 | MYL4\_HUMAN | MYL4 | Myosin light chain 4 | 197 | 21.565 | 2.948903e+05 | 7.989639e+06 | 1210819.6 | 2498728.2 | 659817.25 | 3620273.8 | 68.53 | 79.2 | 44.2 | 32.5 |
| sp\|Q9HD40\|SPCS\_HUMAN | Q9HD40 | SPCS\_HUMAN | SEPSECS | O-phosphoseryl-tRNA(Sec) selenium transferase | 501 | 55.726 | 2.832681e+04 | 1.983228e+06 | 1209067.5 | 707277.2 | 57420.406 | 9463.328 | 34.33 | 60.1 | 12.8 | 17.2 |
| sp\|Q14697\|GANAB\_HUMAN | Q14697 | GANAB\_HUMAN | GANAB | Neutral alpha-glucosidase AB | 944 | 106.874 | 1.074376e+05 | 1.442598e+07 | 1204368.6 | 2499495.0 | 8474356.0 | 2247761.5 | 43.86 | 41.0 | 27.1 | 19.9 |
| sp\|P46926\|GNPI1\_HUMAN | P46926 | GNPI1\_HUMAN | GNPDA1 | Glucosamine-6-phosphate isomerase 1 | 289 | 32.669 | 1.527709e+05 | 6.270375e+06 | 1200892.8 | 3625911.8 | 1350534.5 | 93035.875 | 84.08 | 73.4 | 31.8 | 6.6 |
| sp\|Q15018\|ABRX2\_HUMAN | Q15018 | ABRX2\_HUMAN | ABRAXAS2 | BRISC complex subunit Abraxas 2 | 415 | 46.901 | 6.704041e+04 | 3.950353e+06 | 1200777.1 | 2485025.5 | 126243.0 | 138307.52 | 39.52 | 61.9 | 31.3 | 21.9 |
| sp\|P22059\|OSBP1\_HUMAN | P22059 | OSBP1\_HUMAN | OSBP | Oxysterol-binding protein 1 | 807 | 89.421 | 1.643369e+04 | 1.846256e+06 | 1199110.8 | 470761.6 | 176383.14 |  | 12.52 | 43.5 | 24.9 |  |
| sp\|Q8NB49\|AT11C\_HUMAN | Q8NB49 | AT11C\_HUMAN | ATP11C | Phospholipid-transporting ATPase IG | 1132 | 129.477 | 2.881050e+04 | 4.686628e+06 | 1198266.2 | 38905.758 | 598216.9 | 2851239.5 | 21.91 | 19.9 | 18.8 | 15.9 |
| sp\|O43707\|ACTN4\_HUMAN | O43707 | ACTN4\_HUMAN | ACTN4 | Alpha-actinin-4 | 911 | 104.854 | 2.039509e+04 | 2.686753e+06 | 1192194.5 | 1195104.9 | 209704.42 | 89749.66 | 58.62 | 60.0 | 29.0 | 17.0 |
| sp\|Q9UNH7\|SNX6\_HUMAN | Q9UNH7 | SNX6\_HUMAN | SNX6 | Sorting nexin-6 | 406 | 46.649 | 3.955212e+04 | 2.318085e+06 | 1191929.1 | 785489.7 | 203580.05 | 137085.83 | 40.15 | 37.2 | 18.5 | 21.4 |
| sp\|O95433\|AHSA1\_HUMAN | O95433 | AHSA1\_HUMAN | AHSA1 | Activator of 90 kDa heat shock protein ATPase homolog 1 | 338 | 38.274 | 1.275472e+05 | 6.133266e+06 | 1190585.6 | 2430751.5 | 2511928.5 |  | 53.25 | 65.4 | 43.2 |  |
| sp\|Q9UDY4\|DNJB4\_HUMAN | Q9UDY4 | DNJB4\_HUMAN | DNAJB4 | DnaJ homolog subfamily B member 4 | 337 | 37.807 | 1.152004e+05 | 5.471965e+06 | 1179275.4 | 3345372.0 | 661518.75 | 285799.22 | 40.06 | 61.7 | 30.9 | 23.1 |
| sp\|O95865\|DDAH2\_HUMAN | O95865 | DDAH2\_HUMAN | DDAH2 | N(G),N(G)-dimethylarginine dimethylaminohydrolase 2 | 285 | 29.644 | 1.026967e+05 | 3.824814e+06 | 1174125.5 | 2407248.5 | 158836.6 | 84603.36 | 28.07 | 57.9 | 23.9 | 24.2 |
| sp\|P84085\|ARF5\_HUMAN | P84085 | ARF5\_HUMAN | ARF5 | ADP-ribosylation factor 5 | 180 | 20.530 | 7.850087e+05 | 2.024793e+07 | 1171026.6 | 16283402.0 | 941337.44 | 1852161.9 | 41.11 | 75.6 | 56.7 | 33.9 |
| sp\|Q07889\|SOS1\_HUMAN | Q07889 | SOS1\_HUMAN | SOS1 | Son of sevenless homolog 1 | 1333 | 152.464 | 6.581250e+03 | 1.260645e+06 | 1170928.1 | 89717.09 |  |  | 4.58 | 34.4 |  |  |
| sp\|P08123\|CO1A2\_HUMAN | P08123 | CO1A2\_HUMAN | COL1A2 | Collagen alpha-2(I) chain | 1366 | 129.314 | 4.411177e+04 | 7.166666e+06 | 1170007.6 | 282019.56 | 3281573.0 | 2433066.0 | 13.76 | 76.6 | 54.5 | 39.2 |
| sp\|Q9HCM2\|PLXA4\_HUMAN | Q9HCM2 | PLXA4\_HUMAN | PLXNA4 | Plexin-A4 | 1894 | 208.340 | 4.465613e+03 | 1.168882e+06 | 1168882.5 |  |  |  | 4.12 |  |  |  |
| sp\|P61158\|ARP3\_HUMAN | P61158 | ARP3\_HUMAN | ACTR3 | Actin-related protein 3 | 418 | 47.371 | 7.521462e+04 | 4.476432e+06 | 1166010.6 | 2829341.0 | 311209.22 | 169870.97 | 50.0 | 60.3 | 28.7 | 14.4 |
| sp\|O43847\|NRDC\_HUMAN | O43847 | NRDC\_HUMAN | NRDC | Nardilysin | 1151 | 131.701 | 1.485516e+04 | 2.458010e+06 | 1163235.2 | 1274220.6 |  | 20554.102 | 35.45 | 38.7 |  | 6.3 |
| sp\|O43491\|E41L2\_HUMAN | O43491 | E41L2\_HUMAN | EPB41L2 | Band 4.1-like protein 2 | 1005 | 112.588 | 3.810270e+04 | 5.389705e+06 | 1158413.5 | 1040565.94 | 2495908.8 | 694816.7 | 49.55 | 59.4 | 45.5 | 25.4 |
| sp\|O60506\|HNRPQ\_HUMAN | O60506 | HNRPQ\_HUMAN | SYNCRIP | Heterogeneous nuclear ribonucleoprotein Q | 623 | 69.603 | 4.668057e+04 | 4.082077e+06 | 1158138.6 | 2286850.8 | 397945.84 | 239142.05 | 40.13 | 54.1 | 29.4 | 20.5 |
| sp\|B2RUZ4\|SMIM1\_HUMAN | B2RUZ4 | SMIM1\_HUMAN | SMIM1 | Small integral membrane protein 1 | 78 | 8.749 | 4.259469e+06 | 4.681996e+07 | 1155753.1 | 9536817.0 | 14059888.0 | 22067506.0 | 43.59 | 78.2 | 35.9 | 35.9 |
| sp\|Q92878\|RAD50\_HUMAN | Q92878 | RAD50\_HUMAN | RAD50 | DNA repair protein RAD50 | 1312 | 153.892 | 1.141480e+04 | 2.206995e+06 | 1153123.0 | 596795.44 | 457076.4 |  | 33.92 | 39.3 | 23.7 |  |
| sp\|Q6P597\|KLC3\_HUMAN | Q6P597 | KLC3\_HUMAN | KLC3 | Kinesin light chain 3 | 504 | 55.364 | 2.499889e+04 | 1.738863e+06 | 1152825.8 | 493574.0 | 92462.9 |  | 32.74 | 50.8 | 29.6 |  |
| sp\|Q9BUH6\|PAXX\_HUMAN | Q9BUH6 | PAXX\_HUMAN | PAXX | Protein PAXX | 204 | 21.640 | 1.192248e+05 | 3.241464e+06 | 1148858.6 | 1478644.0 | 333076.38 | 280884.53 | 38.24 | 72.5 | 52.9 | 24.5 |
| sp\|O00178\|GTPB1\_HUMAN | O00178 | GTPB1\_HUMAN | GTPBP1 | GTP-binding protein 1 | 669 | 72.454 | 2.953366e+04 | 2.688418e+06 | 1148830.1 | 1017887.2 | 358108.75 | 163591.52 | 42.15 | 35.9 | 25.0 | 17.6 |
| sp\|P22102\|PUR2\_HUMAN | P22102 | PUR2\_HUMAN | GART | Trifunctional purine biosynthetic protein adenosine-3 | 1010 | 107.767 | 2.496681e+04 | 3.380384e+06 | 1145805.8 | 2161930.8 | 72647.195 |  | 22.87 | 42.3 | 20.0 |  |
| sp\|Q52LW3\|RHG29\_HUMAN | Q52LW3 | RHG29\_HUMAN | ARHGAP29 | Rho GTPase-activating protein 29 | 1261 | 138.710 | 6.570814e+03 | 1.145102e+06 | 1145101.8 |  |  |  | 1.43 |  |  |  |
| sp\|Q9NZL9\|MAT2B\_HUMAN | Q9NZL9 | MAT2B\_HUMAN | MAT2B | Methionine adenosyltransferase 2 subunit beta | 334 | 37.552 | 6.387855e+04 | 3.013737e+06 | 1143357.1 | 1738785.6 | 112507.414 | 19086.443 | 54.19 | 68.3 | 32.3 | 8.7 |
| sp\|Q6GMV2\|SMYD5\_HUMAN | Q6GMV2 | SMYD5\_HUMAN | SMYD5 | Histone-lysine N-trimethyltransferase SMYD5 | 418 | 47.341 | 7.710006e+04 | 4.585738e+06 | 1140672.2 | 3247687.0 | 197379.28 |  | 22.73 | 38.5 | 10.5 |  |
| sp\|P26641\|EF1G\_HUMAN | P26641 | EF1G\_HUMAN | EEF1G | Elongation factor 1-gamma | 437 | 50.119 | 3.581350e+04 | 2.255103e+06 | 1139466.8 | 693216.94 | 297233.8 | 125185.18 | 33.64 | 44.4 | 25.2 | 13.3 |
| sp\|P62140\|PP1B\_HUMAN | P62140 | PP1B\_HUMAN | PPP1CB | Serine/threonine-protein phosphatase PP1-beta catalytic subunit | 327 | 37.187 | 5.153550e+04 | 2.407769e+06 | 1137866.9 | 1202824.9 | 36927.06 | 30150.08 | 37.31 | 66.7 | 24.5 | 3.1 |
| sp\|Q9Y263\|PLAP\_HUMAN | Q9Y263 | PLAP\_HUMAN | PLAA | Phospholipase A-2-activating protein | 795 | 87.157 | 3.291874e+04 | 3.604646e+06 | 1120822.8 | 2070879.9 | 202971.39 | 209971.67 | 51.57 | 51.6 | 25.2 | 21.1 |
| sp\|P45985\|MP2K4\_HUMAN | P45985 | MP2K4\_HUMAN | MAP2K4 | Dual specificity mitogen-activated protein kinase kinase 4 | 399 | 44.288 | 4.584429e+04 | 2.550871e+06 | 1118927.5 | 1323954.9 | 107988.734 |  | 31.83 | 61.2 | 36.8 |  |
| sp\|O00505\|IMA4\_HUMAN | O00505 | IMA4\_HUMAN | KPNA3 | Importin subunit alpha-4 | 521 | 57.811 | 5.336593e+04 | 3.876071e+06 | 1113829.8 | 2068034.1 | 694207.25 |  | 31.29 | 31.9 | 14.6 |  |
| sp\|Q15751\|HERC1\_HUMAN | Q15751 | HERC1\_HUMAN | HERC1 | Probable E3 ubiquitin-protein ligase HERC1 | 4861 | 532.228 | 1.773708e+03 | 1.186034e+06 | 1110356.6 | 75677.17 |  |  | 1.83 | 26.4 |  |  |
| sp\|Q09161\|NCBP1\_HUMAN | Q09161 | NCBP1\_HUMAN | NCBP1 | Nuclear cap-binding protein subunit 1 | 790 | 91.839 | 1.582218e+04 | 1.825621e+06 | 1106660.2 | 718960.6 |  |  | 41.27 | 51.1 |  |  |
| sp\|O76071\|CIAO1\_HUMAN | O76071 | CIAO1\_HUMAN | CIAO1 | Probable cytosolic iron-sulfur protein assembly protein CIAO1 | 339 | 37.840 | 6.091333e+04 | 2.895881e+06 | 1105771.0 | 1759848.8 |  | 30260.814 | 44.84 | 58.4 |  | 7.4 |
| sp\|Q9UNQ0\|ABCG2\_HUMAN | Q9UNQ0 | ABCG2\_HUMAN | ABCG2 | Broad substrate specificity ATP-binding cassette transporter ABCG2 | 655 | 72.314 | 1.813440e+05 | 1.647565e+07 | 1105089.8 | 228157.34 | 4049251.0 | 11093156.0 | 28.24 | 20.6 | 24.6 | 20.6 |
| sp\|O75608\|LYPA1\_HUMAN | O75608 | LYPA1\_HUMAN | LYPLA1 | Acyl-protein thioesterase 1 | 230 | 24.670 | 6.368155e+04 | 1.973785e+06 | 1103776.1 | 77236.16 | 488836.78 | 303936.38 | 42.17 | 47.4 | 13.9 | 13.9 |
| sp\|P53611\|PGTB2\_HUMAN | P53611 | PGTB2\_HUMAN | RABGGTB | Geranylgeranyl transferase type-2 subunit beta | 331 | 36.924 | 9.674133e+04 | 4.487846e+06 | 1096192.5 | 2931217.2 | 230357.16 | 230079.19 | 25.08 | 29.0 | 17.8 | 16.0 |
| sp\|Q15811\|ITSN1\_HUMAN | Q15811 | ITSN1\_HUMAN | ITSN1 | Intersectin-1 | 1721 | 195.422 | 1.199205e+04 | 2.944312e+06 | 1093571.4 | 840587.6 | 778682.8 | 231470.64 | 24.64 | 35.7 | 28.9 | 13.8 |
| sp\|P35237\|SPB6\_HUMAN | P35237 | SPB6\_HUMAN | SERPINB6 | Serpin B6 | 376 | 42.622 | 6.220668e+04 | 3.331102e+06 | 1091767.1 | 1889418.0 | 252619.61 | 97297.66 | 55.85 | 72.9 | 17.3 | 17.0 |
| sp\|Q53S08\|RAB6D\_HUMAN | Q53S08 | RAB6D\_HUMAN | RAB6D | Ras-related protein Rab-6D | 254 | 28.242 | 6.919458e+04 | 2.455188e+06 | 1090565.5 | 1364622.2 |  |  | 24.02 | 27.2 |  |  |
| sp\|P31943\|HNRH1\_HUMAN | P31943 | HNRH1\_HUMAN | HNRNPH1 | Heterogeneous nuclear ribonucleoprotein H | 449 | 49.229 | 4.315284e+04 | 2.668994e+06 | 1089715.0 | 1120214.9 | 257835.62 | 201228.34 | 37.64 | 47.7 | 38.5 | 19.8 |
| sp\|O95394\|AGM1\_HUMAN | O95394 | AGM1\_HUMAN | PGM3 | Phosphoacetylglucosamine mutase | 542 | 59.852 | 2.741421e+04 | 2.061444e+06 | 1088668.9 | 783120.25 | 135733.78 | 53921.125 | 37.64 | 46.1 | 22.1 | 7.7 |
| sp\|P57772\|SELB\_HUMAN | P57772 | SELB\_HUMAN | EEFSEC | Selenocysteine-specific elongation factor | 596 | 65.305 | 2.484733e+04 | 2.038653e+06 | 1086101.8 | 818994.5 | 87194.36 | 46362.01 | 32.21 | 43.5 | 21.1 | 11.9 |
| sp\|P42574\|CASP3\_HUMAN | P42574 | CASP3\_HUMAN | CASP3 | Caspase-3 | 277 | 31.608 | 7.552385e+04 | 2.999151e+06 | 1084159.0 | 1557723.5 | 240711.9 | 116556.484 | 23.83 | 55.6 | 22.7 | 19.9 |
| sp\|O15554\|KCNN4\_HUMAN | O15554 | KCNN4\_HUMAN | KCNN4 | Intermediate conductance calcium-activated potassium channel protein 4 | 427 | 47.696 | 2.858734e+05 | 1.713061e+07 | 1080984.6 | 2853527.5 | 2991992.0 | 10204110.0 | 24.12 | 35.4 | 27.4 | 25.1 |
| sp\|Q15477\|SKI2\_HUMAN | Q15477 | SKI2\_HUMAN | SKIC2 | Superkiller complex protein 2 | 1246 | 137.755 | 1.529545e+04 | 2.647201e+06 | 1080733.2 | 998090.4 | 521684.6 | 46692.59 | 42.3 | 47.2 | 15.0 | 11.1 |
| sp\|Q15833\|STXB2\_HUMAN | Q15833 | STXB2\_HUMAN | STXBP2 | Syntaxin-binding protein 2 | 593 | 66.453 | 6.204922e+04 | 5.180457e+06 | 1070966.5 | 1484875.6 | 2342760.2 | 281855.03 | 44.52 | 63.6 | 39.0 | 19.1 |
| sp\|Q15813\|TBCE\_HUMAN | Q15813 | TBCE\_HUMAN | TBCE | Tubulin-specific chaperone E | 527 | 59.346 | 1.904940e+04 | 1.420332e+06 | 1065926.0 | 293008.8 | 61397.633 |  | 25.05 | 43.3 | 22.2 |  |
| sp\|Q8NFH3\|NUP43\_HUMAN | Q8NFH3 | NUP43\_HUMAN | NUP43 | Nucleoporin Nup43 | 380 | 42.151 | 3.004747e+04 | 1.591230e+06 | 1061879.2 | 385714.72 | 143636.12 |  | 32.11 | 31.6 | 12.4 |  |
| sp\|Q9H444\|CHM4B\_HUMAN | Q9H444 | CHM4B\_HUMAN | CHMP4B | Charged multivesicular body protein 4b | 224 | 24.950 | 9.627640e+04 | 3.017919e+06 | 1061491.2 | 1288177.1 | 476729.28 | 191521.05 | 45.98 | 46.0 | 33.5 | 37.1 |
| sp\|Q9NYU2\|UGGG1\_HUMAN | Q9NYU2 | UGGG1\_HUMAN | UGGT1 | UDP-glucose:glycoprotein glucosyltransferase 1 | 1555 | 177.190 | 5.401876e+04 | 1.202544e+07 | 1059194.0 | 1355778.5 | 7804463.0 | 1806004.1 | 30.23 | 41.0 | 31.9 | 17.7 |
| sp\|Q16555\|DPYL2\_HUMAN | Q16555 | DPYL2\_HUMAN | DPYSL2 | Dihydropyrimidinase-related protein 2 | 572 | 62.294 | 2.155693e+04 | 1.687137e+06 | 1058529.8 | 478319.44 | 128539.125 | 21748.234 | 57.87 | 60.0 | 30.1 | 9.6 |
| sp\|Q15843\|NEDD8\_HUMAN | Q15843 | NEDD8\_HUMAN | NEDD8 | NEDD8 | 81 | 9.072 | 4.514088e+05 | 5.145057e+06 | 1058047.2 | 3671455.5 | 300440.0 | 115114.75 | 79.01 | 81.5 | 34.6 | 24.7 |
| sp\|P63096\|GNAI1\_HUMAN | P63096 | GNAI1\_HUMAN | GNAI1 | Guanine nucleotide-binding protein G(i) subunit alpha-1 | 354 | 38.940 | 2.160492e+04 | 1.056977e+06 | 1056977.4 |  |  |  | 40.4 |  |  |  |
| sp\|P31321\|KAP1\_HUMAN | P31321 | KAP1\_HUMAN | PRKAR1B | cAMP-dependent protein kinase type I-beta regulatory subunit | 381 | 43.073 | 1.602580e+05 | 8.672452e+06 | 1052972.4 | 476390.5 | 7143089.5 |  | 28.35 | 34.4 | 35.2 |  |
| sp\|P08174\|DAF\_HUMAN | P08174 | DAF\_HUMAN | CD55 | Complement decay-accelerating factor | 381 | 41.400 | 1.757894e+05 | 9.143450e+06 | 1052370.5 | 236331.86 | 5827964.5 | 2026783.1 | 7.35 | 50.9 | 38.6 | 28.6 |
| sp\|Q9BTU6\|P4K2A\_HUMAN | Q9BTU6 | P4K2A\_HUMAN | PI4K2A | Phosphatidylinositol 4-kinase type 2-alpha | 479 | 54.022 | 1.510192e+05 | 1.024991e+07 | 1049523.1 | 976514.4 | 1856539.0 | 6367335.0 | 65.97 | 49.1 | 40.7 | 41.1 |
| sp\|Q7Z6L1\|TCPR1\_HUMAN | Q7Z6L1 | TCPR1\_HUMAN | TECPR1 | Tectonin beta-propeller repeat-containing protein 1 | 1165 | 129.696 | 1.102373e+04 | 1.796273e+06 | 1048246.75 | 70611.984 | 271932.66 | 405481.62 | 15.28 | 25.3 | 16.7 | 10.1 |
| sp\|Q9BRP4\|PAAF1\_HUMAN | Q9BRP4 | PAAF1\_HUMAN | PAAF1 | Proteasomal ATPase-associated factor 1 | 392 | 42.190 | 8.024016e+04 | 4.253226e+06 | 1047249.6 | 3057221.0 | 121923.31 | 26832.025 | 30.61 | 41.3 | 22.7 | 15.3 |
| sp\|O95248\|MTMR5\_HUMAN | O95248 | MTMR5\_HUMAN | SBF1 | Myotubularin-related protein 5 | 1868 | 208.443 | 8.504881e+03 | 2.227269e+06 | 1044383.44 | 934409.9 | 103024.18 | 145451.72 | 22.22 | 42.6 | 18.1 | 12.3 |
| sp\|P51610\|HCFC1\_HUMAN | P51610 | HCFC1\_HUMAN | HCFC1 | Host cell factor 1 | 2035 | 208.732 | 6.711479e+03 | 1.760048e+06 | 1043522.1 | 624798.8 | 91726.84 |  | 6.0 | 30.7 | 17.2 |  |
| sp\|P49770\|EI2BB\_HUMAN | P49770 | EI2BB\_HUMAN | EIF2B2 | Translation initiation factor eIF-2B subunit beta | 351 | 38.990 | 5.969988e+04 | 2.924448e+06 | 1040877.25 | 1576988.9 | 272705.6 | 33876.1 | 33.9 | 48.7 | 44.4 | 18.5 |
| sp\|O95197\|RTN3\_HUMAN | O95197 | RTN3\_HUMAN | RTN3 | Reticulon-3 | 1032 | 112.611 | 5.536614e+04 | 7.833254e+06 | 1034889.06 | 3473661.2 | 1098808.2 | 2225895.8 | 2.62 | 15.8 | 16.3 | 5.5 |
| sp\|Q04637\|IF4G1\_HUMAN | Q04637 | IF4G1\_HUMAN | EIF4G1 | Eukaryotic translation initiation factor 4 gamma 1 | 1599 | 175.491 | 2.115251e+04 | 4.663735e+06 | 1034152.3 | 2306862.0 | 824199.75 | 498520.97 | 32.9 | 36.5 | 21.6 | 14.4 |
| sp\|Q6P1N0\|C2D1A\_HUMAN | Q6P1N0 | C2D1A\_HUMAN | CC2D1A | Coiled-coil and C2 domain-containing protein 1A | 951 | 104.062 | 1.868733e+04 | 2.443187e+06 | 1030728.1 | 817536.0 | 271281.44 | 323641.03 | 39.43 | 48.9 | 46.3 | 21.8 |
| sp\|P62495\|ERF1\_HUMAN | P62495 | ERF1\_HUMAN | ETF1 | Eukaryotic peptide chain release factor subunit 1 | 437 | 49.031 | 4.865063e+04 | 2.996929e+06 | 1029991.25 | 1711403.9 | 169806.92 | 85726.555 | 37.99 | 64.5 | 19.0 | 9.4 |
| sp\|Q14773\|ICAM4\_HUMAN | Q14773 | ICAM4\_HUMAN | ICAM4 | Intercellular adhesion molecule 4 | 271 | 29.265 | 6.657457e+05 | 2.447789e+07 | 1027664.0 | 130418.43 | 3749470.5 | 19570340.0 | 26.57 | 24.7 | 39.1 | 21.4 |
| sp\|Q9Y2Z0\|SGT1\_HUMAN | Q9Y2Z0 | SGT1\_HUMAN | SUGT1 | Protein SGT1 homolog | 365 | 41.024 | 8.530115e+04 | 4.396530e+06 | 1027634.9 | 1972194.5 | 683729.56 | 712970.9 | 57.81 | 68.5 | 44.1 | 20.3 |
| sp\|P43405\|KSYK\_HUMAN | P43405 | KSYK\_HUMAN | SYK | Tyrosine-protein kinase SYK | 635 | 72.066 | 2.627605e+04 | 2.379072e+06 | 1027514.0 | 1107332.5 | 163507.62 | 80717.82 | 40.94 | 54.3 | 23.8 | 15.7 |
| sp\|O95260\|ATE1\_HUMAN | O95260 | ATE1\_HUMAN | ATE1 | Arginyl-tRNA--protein transferase 1 | 518 | 59.090 | 4.718514e+04 | 3.502970e+06 | 1022817.06 | 2226667.5 | 187060.47 | 66424.58 | 21.04 | 47.7 | 25.9 | 13.5 |
| sp\|Q8IXJ6\|SIR2\_HUMAN | Q8IXJ6 | SIR2\_HUMAN | SIRT2 | NAD-dependent protein deacetylase sirtuin-2 | 389 | 43.182 | 9.000858e+04 | 4.883192e+06 | 1021160.4 | 2682832.8 | 1179199.2 |  | 44.22 | 53.5 | 43.2 |  |
| sp\|P61019\|RAB2A\_HUMAN | P61019 | RAB2A\_HUMAN | RAB2A | Ras-related protein Rab-2A | 212 | 23.546 | 1.161026e+05 | 3.434601e+06 | 1020836.9 | 1943211.1 | 338276.62 | 132276.1 | 66.51 | 71.2 | 53.8 | 33.0 |
| sp\|Q9UBW8\|CSN7A\_HUMAN | Q9UBW8 | CSN7A\_HUMAN | COPS7A | COP9 signalosome complex subunit 7a | 275 | 30.277 | 1.595584e+05 | 6.069455e+06 | 1014569.25 | 4178302.0 | 734806.4 | 141777.39 | 57.09 | 51.3 | 35.3 | 29.8 |
| sp\|O00186\|STXB3\_HUMAN | O00186 | STXB3\_HUMAN | STXBP3 | Syntaxin-binding protein 3 | 592 | 67.764 | 3.589797e+04 | 3.056230e+06 | 1011116.25 | 1654403.0 | 257241.05 | 133469.86 | 34.12 | 37.5 | 19.3 | 16.7 |
| sp\|Q5D862\|FILA2\_HUMAN | Q5D862 | FILA2\_HUMAN | FLG2 | Filaggrin-2 | 2391 | 248.073 | 2.128118e+04 | 6.632729e+06 | 1011014.2 | 1567704.1 | 1420305.2 | 2633705.8 | 23.3 | 32.6 | 21.9 | 18.7 |
| sp\|O75351\|VPS4B\_HUMAN | O75351 | VPS4B\_HUMAN | VPS4B | Vacuolar protein sorting-associated protein 4B | 444 | 49.302 | 3.442903e+04 | 2.132586e+06 | 1010491.2 | 419868.1 | 702226.5 |  | 31.31 | 46.2 | 28.2 |  |
| sp\|Q9C0K7\|STRAB\_HUMAN | Q9C0K7 | STRAB\_HUMAN | STRADB | STE20-related kinase adapter protein beta | 418 | 47.026 | 3.569367e+04 | 2.108854e+06 | 1009982.06 | 341955.97 | 203829.22 | 553086.44 | 27.51 | 35.9 | 15.6 | 13.2 |
| sp\|P46439\|GSTM5\_HUMAN | P46439 | GSTM5\_HUMAN | GSTM5 | Glutathione S-transferase Mu 5 | 218 | 23.980 | 3.346408e+04 | 1.008197e+06 | 1008196.6 |  |  |  | 31.19 |  |  |  |
| sp\|Q8IUR7\|ARMC8\_HUMAN | Q8IUR7 | ARMC8\_HUMAN | ARMC8 | Armadillo repeat-containing protein 8 | 673 | 75.509 | 2.220574e+04 | 2.106595e+06 | 1008089.75 | 852300.06 | 77904.64 | 168300.95 | 37.0 | 52.3 | 32.8 | 21.5 |
| sp\|O43592\|XPOT\_HUMAN | O43592 | XPOT\_HUMAN | XPOT | Exportin-T | 962 | 109.964 | 1.045874e+05 | 1.444931e+07 | 1007999.06 | 13441315.0 |  |  | 22.56 | 33.1 |  |  |
| sp\|P50995\|ANX11\_HUMAN | P50995 | ANX11\_HUMAN | ANXA11 | Annexin A11 | 505 | 54.390 | 8.080056e+04 | 5.521418e+06 | 1007014.75 | 1528175.5 | 2776595.5 | 209632.03 | 38.02 | 37.4 | 36.2 | 23.4 |
| sp\|O15144\|ARPC2\_HUMAN | O15144 | ARPC2\_HUMAN | ARPC2 | Actin-related protein 2/3 complex subunit 2 | 300 | 34.333 | 8.745070e+04 | 3.772178e+06 | 1006224.8 | 2451677.5 | 266876.2 | 47399.645 | 44.67 | 47.3 | 33.7 | 6.3 |
| sp\|Q9UPU5\|UBP24\_HUMAN | Q9UPU5 | UBP24\_HUMAN | USP24 | Ubiquitin carboxyl-terminal hydrolase 24 | 2620 | 294.365 | 1.141261e+04 | 4.220738e+06 | 1001199.44 | 2705565.5 | 126202.836 | 387769.84 | 37.18 | 46.9 | 16.5 | 12.8 |
| sp\|Q15819\|UB2V2\_HUMAN | Q15819 | UB2V2\_HUMAN | UBE2V2 | Ubiquitin-conjugating enzyme E2 variant 2 | 145 | 16.363 | 1.084797e+06 | 2.230123e+07 | 1000480.1 | 9182024.0 | 9195484.0 | 2923237.5 | 63.45 | 89.7 | 71.7 | 54.5 |
| sp\|P06702\|S10A9\_HUMAN | P06702 | S10A9\_HUMAN | S100A9 | Protein S100-A9 | 114 | 13.242 | 4.003748e+05 | 6.660970e+06 | 999036.94 | 923846.8 | 3640939.2 | 1097147.2 | 63.16 | 90.4 | 72.8 | 50.0 |
| sp\|Q9NR09\|BIRC6\_HUMAN | Q9NR09 | BIRC6\_HUMAN | BIRC6 | Baculoviral IAP repeat-containing protein 6 | 4857 | 530.269 | 1.761524e+03 | 1.173551e+06 | 998753.0 | 174797.95 |  |  | 6.05 | 23.9 |  |  |
| sp\|Q8TAT6\|NPL4\_HUMAN | Q8TAT6 | NPL4\_HUMAN | NPLOC4 | Nuclear protein localization protein 4 homolog | 608 | 68.120 | 2.561500e+04 | 2.192231e+06 | 998620.8 | 1025395.0 | 123271.01 | 44943.703 | 43.26 | 52.6 | 16.6 | 12.8 |
| sp\|Q96CD2\|COAC\_HUMAN | Q96CD2 | COAC\_HUMAN | PPCDC | Phosphopantothenoylcysteine decarboxylase | 204 | 22.395 | 9.663800e+04 | 2.719043e+06 | 990307.56 | 1322724.9 | 406011.0 |  | 53.43 | 76.5 | 30.4 |  |
| sp\|O15440\|MRP5\_HUMAN | O15440 | MRP5\_HUMAN | ABCC5 | ATP-binding cassette sub-family C member 5 | 1437 | 160.660 | 1.626266e+04 | 3.282589e+06 | 989609.3 | 307247.78 | 304249.28 | 1681483.0 | 21.36 | 28.0 | 23.0 | 16.7 |
| sp\|P50453\|SPB9\_HUMAN | P50453 | SPB9\_HUMAN | SERPINB9 | Serpin B9 | 376 | 42.404 | 5.537720e+04 | 2.950224e+06 | 988150.5 | 1806030.5 | 156042.75 |  | 56.12 | 60.6 | 23.1 |  |
| sp\|P43250\|GRK6\_HUMAN | P43250 | GRK6\_HUMAN | GRK6 | G protein-coupled receptor kinase 6 | 576 | 65.991 | 3.220433e+04 | 2.670030e+06 | 988011.6 | 1258171.6 | 143773.89 | 280072.5 | 27.26 | 49.1 | 24.8 | 16.1 |
| sp\|Q8NE71\|ABCF1\_HUMAN | Q8NE71 | ABCF1\_HUMAN | ABCF1 | ATP-binding cassette sub-family F member 1 | 845 | 95.926 | 7.564420e+04 | 9.116521e+06 | 986633.6 | 1841387.4 | 273278.8 | 6015221.5 | 33.61 | 45.9 | 21.7 | 20.0 |
| sp\|P30101\|PDIA3\_HUMAN | P30101 | PDIA3\_HUMAN | PDIA3 | Protein disulfide-isomerase A3 | 505 | 56.782 | 3.663338e+05 | 2.613393e+07 | 984833.1 | 2565554.8 | 20203826.0 | 2379720.2 | 50.69 | 65.7 | 48.9 | 28.9 |
| sp\|P62136\|PP1A\_HUMAN | P62136 | PP1A\_HUMAN | PPP1CA | Serine/threonine-protein phosphatase PP1-alpha catalytic subunit | 330 | 37.512 | 5.021588e+04 | 2.366620e+06 | 984674.2 | 1321038.6 | 60906.72 |  | 38.79 | 67.6 | 17.6 |  |
| sp\|Q9NVS9\|PNPO\_HUMAN | Q9NVS9 | PNPO\_HUMAN | PNPO | Pyridoxine-5'-phosphate oxidase | 261 | 29.988 | 7.379618e+04 | 2.780344e+06 | 983388.75 | 1672311.0 | 124644.46 |  | 40.61 | 58.6 | 23.0 |  |
| sp\|Q16181\|SEPT7\_HUMAN | Q16181 | SEPT7\_HUMAN | SEPTIN7 | Septin-7 | 437 | 50.680 | 5.003642e+04 | 3.185958e+06 | 982305.25 | 2019273.0 | 184379.34 |  | 35.47 | 65.9 | 27.5 |  |
| sp\|Q9UH65\|SWP70\_HUMAN | Q9UH65 | SWP70\_HUMAN | SWAP70 | Switch-associated protein 70 | 585 | 68.998 | 2.506230e+04 | 2.172574e+06 | 982082.06 | 1037820.6 | 69521.79 | 83149.695 | 42.91 | 55.4 | 26.3 | 11.8 |
| sp\|Q08623\|HDHD1\_HUMAN | Q08623 | HDHD1\_HUMAN | PUDP | Pseudouridine-5'-phosphatase | 228 | 25.249 | 5.059768e+04 | 1.605062e+06 | 981558.8 | 623503.7 | 0.0 |  | 28.07 | 42.1 | 22.8 |  |
| sp\|Q5R372\|RBG1L\_HUMAN | Q5R372 | RBG1L\_HUMAN | RABGAP1L | Rab GTPase-activating protein 1-like | 815 | 92.513 | 2.834241e+04 | 3.294251e+06 | 975990.4 | 2039480.6 | 64732.414 | 214047.33 | 31.78 | 38.4 | 18.5 | 16.2 |
| sp\|Q9NZ32\|ARP10\_HUMAN | Q9NZ32 | ARP10\_HUMAN | ACTR10 | Actin-related protein 10 | 417 | 46.307 | 4.427499e+04 | 2.575860e+06 | 972153.5 | 1586647.0 | 17059.84 |  | 49.88 | 47.5 | 27.3 |  |
| sp\|O75165\|DJC13\_HUMAN | O75165 | DJC13\_HUMAN | DNAJC13 | DnaJ homolog subfamily C member 13 | 2243 | 254.415 | 1.089985e+04 | 3.484017e+06 | 970933.2 | 2163532.2 | 59202.355 | 290348.75 | 37.27 | 46.3 | 14.8 | 16.8 |
| sp\|O95155\|UBE4B\_HUMAN | O95155 | UBE4B\_HUMAN | UBE4B | Ubiquitin conjugation factor E4 B | 1302 | 146.185 | 9.718649e+03 | 1.784949e+06 | 970605.0 | 814344.3 |  |  | 36.87 | 43.1 |  |  |
| sp\|P11940\|PABP1\_HUMAN | P11940 | PABP1\_HUMAN | PABPC1 | Polyadenylate-binding protein 1 | 636 | 70.671 | 2.984674e+04 | 2.650057e+06 | 969591.6 | 1107460.2 | 330455.88 | 242549.62 | 37.58 | 51.4 | 34.9 | 21.1 |
| sp\|Q96KG9\|SCYL1\_HUMAN | Q96KG9 | SCYL1\_HUMAN | SCYL1 | N-terminal kinase-like protein | 808 | 89.631 | 2.626590e+04 | 2.957792e+06 | 969009.5 | 1274312.1 | 525930.94 | 188539.81 | 36.51 | 44.8 | 23.8 | 17.8 |
| sp\|Q9NP72\|RAB18\_HUMAN | Q9NP72 | RAB18\_HUMAN | RAB18 | Ras-related protein Rab-18 | 206 | 22.977 | 8.000851e+04 | 2.309652e+06 | 966853.7 | 579892.1 | 504008.62 | 258898.05 | 48.54 | 71.4 | 54.4 | 25.2 |
| sp\|Q53H96\|P5CR3\_HUMAN | Q53H96 | P5CR3\_HUMAN | PYCR3 | Pyrroline-5-carboxylate reductase 3 | 274 | 28.663 | 2.388296e+05 | 8.600563e+06 | 966115.75 | 5492708.5 | 1782237.2 | 359501.22 | 49.64 | 58.4 | 28.8 | 28.1 |
| sp\|P51858\|HDGF\_HUMAN | P51858 | HDGF\_HUMAN | HDGF | Hepatoma-derived growth factor | 240 | 26.788 | 2.407395e+05 | 8.102233e+06 | 963764.8 | 6263032.0 | 589882.4 | 285554.2 | 70.83 | 78.8 | 55.0 | 33.3 |
| sp\|Q9Y5Z4\|HEBP2\_HUMAN | Q9Y5Z4 | HEBP2\_HUMAN | HEBP2 | Heme-binding protein 2 | 205 | 22.875 | 7.193011e+04 | 2.067231e+06 | 962823.94 | 908020.94 | 165619.88 | 30766.166 | 32.68 | 43.4 | 31.7 | 16.6 |
| sp\|Q9UJW0\|DCTN4\_HUMAN | Q9UJW0 | DCTN4\_HUMAN | DCTN4 | Dynactin subunit 4 | 460 | 52.337 | 7.266321e+04 | 4.777938e+06 | 959478.2 | 3147765.5 | 604925.94 | 65768.68 | 45.87 | 44.6 | 28.7 | 13.9 |
| sp\|O60884\|DNJA2\_HUMAN | O60884 | DNJA2\_HUMAN | DNAJA2 | DnaJ homolog subfamily A member 2 | 412 | 45.746 | 4.992323e+04 | 2.869280e+06 | 954238.6 | 1480912.2 | 434129.34 |  | 29.13 | 52.7 | 34.7 |  |
| sp\|Q00534\|CDK6\_HUMAN | Q00534 | CDK6\_HUMAN | CDK6 | Cyclin-dependent kinase 6 | 326 | 36.938 | 2.894455e+05 | 1.343252e+07 | 953554.25 | 731465.0 | 11747498.0 |  | 48.77 | 44.2 | 43.6 |  |
| sp\|O15162\|PLS1\_HUMAN | O15162 | PLS1\_HUMAN | PLSCR1 | Phospholipid scramblase 1 | 318 | 35.049 | 1.875600e+05 | 8.259106e+06 | 953021.94 | 109920.55 | 2293047.8 | 4903115.5 | 19.5 | 31.8 | 7.5 | 7.9 |
| sp\|Q8NEG4\|FA83F\_HUMAN | Q8NEG4 | FA83F\_HUMAN | FAM83F | Protein FAM83F | 500 | 55.486 | 1.371970e+04 | 9.564125e+05 | 945754.6 | 10657.918 |  |  | 5.8 | 39.4 |  |  |
| sp\|Q6PGP7\|SKI3\_HUMAN | Q6PGP7 | SKI3\_HUMAN | SKIC3 | Superkiller complex protein 3 | 1564 | 175.486 | 8.691368e+03 | 1.916231e+06 | 941236.56 | 802679.7 | 101136.33 | 71177.94 | 36.0 | 35.5 | 14.3 | 7.9 |
| sp\|O75157\|T22D2\_HUMAN | O75157 | T22D2\_HUMAN | TSC22D2 | TSC22 domain family protein 2 | 780 | 79.228 | 2.709565e+04 | 2.697090e+06 | 938835.75 | 935940.2 | 759527.5 | 62786.215 | 5.51 | 11.7 | 7.9 | 5.9 |
| sp\|Q9NSY1\|BMP2K\_HUMAN | Q9NSY1 | BMP2K\_HUMAN | BMP2K | BMP-2-inducible protein kinase | 1161 | 129.172 | 2.129454e+04 | 3.455841e+06 | 937221.56 | 896229.2 | 1326895.5 | 295494.6 | 35.14 | 48.5 | 30.3 | 12.4 |
| sp\|P35236\|PTN7\_HUMAN | P35236 | PTN7\_HUMAN | PTPN7 | Tyrosine-protein phosphatase non-receptor type 7 | 360 | 40.529 | 7.382560e+04 | 3.759153e+06 | 934343.25 | 2234946.2 | 589863.9 |  | 51.94 | 71.4 | 29.4 |  |
| sp\|O43390\|HNRPR\_HUMAN | O43390 | HNRPR\_HUMAN | HNRNPR | Heterogeneous nuclear ribonucleoprotein R | 633 | 70.943 | 4.623980e+04 | 4.121380e+06 | 932938.75 | 2879810.8 | 308630.8 |  | 14.53 | 40.8 | 20.4 |  |
| sp\|Q12840\|KIF5A\_HUMAN | Q12840 | KIF5A\_HUMAN | KIF5A | Kinesin heavy chain isoform 5A | 1032 | 113.520 | 6.533315e+03 | 9.318009e+05 | 931800.9 |  |  |  | 13.86 |  |  |  |
| sp\|Q14CX7\|NAA25\_HUMAN | Q14CX7 | NAA25\_HUMAN | NAA25 | N-alpha-acetyltransferase 25, NatB auxiliary subunit | 972 | 112.292 | 1.312393e+04 | 1.851526e+06 | 931022.4 | 920503.56 |  |  | 33.54 | 32.8 |  |  |
| sp\|Q86YS7\|C2CD5\_HUMAN | Q86YS7 | C2CD5\_HUMAN | C2CD5 | C2 domain-containing protein 5 | 1000 | 110.447 | 2.662652e+04 | 3.694754e+06 | 929127.2 | 1936576.4 | 541484.25 | 287566.28 | 32.6 | 30.9 | 30.2 | 17.1 |
| sp\|Q96AZ6\|ISG20\_HUMAN | Q96AZ6 | ISG20\_HUMAN | ISG20 | Interferon-stimulated gene 20 kDa protein | 181 | 20.363 | 1.258760e+05 | 3.220342e+06 | 928662.75 | 2038414.9 | 130166.93 | 123096.95 | 59.12 | 75.7 | 74.0 | 26.0 |
| sp\|P58546\|MTPN\_HUMAN | P58546 | MTPN\_HUMAN | MTPN | Myotrophin | 118 | 12.895 | 3.475462e+05 | 5.630552e+06 | 928222.4 | 3492650.0 | 826012.56 | 383667.5 | 57.63 | 64.4 | 39.0 | 22.0 |
| sp\|Q6P1Q0\|LTMD1\_HUMAN | Q6P1Q0 | LTMD1\_HUMAN | LETMD1 | LETM1 domain-containing protein 1 | 360 | 39.600 | 1.862570e+04 | 9.266697e+05 | 926669.7 |  |  |  | 3.06 |  |  |  |
| sp\|P36543\|VATE1\_HUMAN | P36543 | VATE1\_HUMAN | ATP6V1E1 | V-type proton ATPase subunit E 1 | 226 | 26.145 | 2.288721e+05 | 7.517935e+06 | 926454.94 | 5737281.0 | 629156.25 | 225043.27 | 33.19 | 58.8 | 35.4 | 31.4 |
| sp\|Q7L7X3\|TAOK1\_HUMAN | Q7L7X3 | TAOK1\_HUMAN | TAOK1 | Serine/threonine-protein kinase TAO1 | 1001 | 116.070 | 7.695971e+03 | 1.122278e+06 | 924857.94 | 197420.34 |  |  | 6.39 | 26.5 |  |  |
| sp\|P36873\|PP1G\_HUMAN | P36873 | PP1G\_HUMAN | PPP1CC | Serine/threonine-protein phosphatase PP1-gamma catalytic subunit | 323 | 36.984 | 4.514460e+04 | 2.097668e+06 | 923958.9 | 1173709.4 |  |  | 33.44 | 68.7 |  |  |
| sp\|O60333\|KIF1B\_HUMAN | O60333 | KIF1B\_HUMAN | KIF1B | Kinesin-like protein KIF1B | 1816 | 204.476 | 1.169053e+04 | 3.003266e+06 | 920375.6 | 1918189.2 | 164701.25 |  | 10.19 | 24.7 | 14.5 |  |
| sp\|P14314\|GLU2B\_HUMAN | P14314 | GLU2B\_HUMAN | PRKCSH | Glucosidase 2 subunit beta | 528 | 59.425 | 1.884335e+05 | 1.406839e+07 | 918036.0 | 776994.6 | 9895896.0 | 2477462.2 | 23.86 | 27.1 | 30.3 | 19.3 |
| sp\|Q969G6\|RIFK\_HUMAN | Q969G6 | RIFK\_HUMAN | RFK | Riboflavin kinase | 155 | 17.623 | 6.236130e+04 | 1.380741e+06 | 914593.0 | 466147.75 |  |  | 12.9 | 41.3 |  |  |
| sp\|O60814\|H2B1K\_HUMAN | O60814 | H2B1K\_HUMAN | H2BC12 | Histone H2B type 1-K | 126 | 13.890 | 1.570336e+05 | 2.740387e+06 | 912991.7 | 1827395.2 |  |  | 61.9 | 81.0 |  |  |
| sp\|Q9NQR4\|NIT2\_HUMAN | Q9NQR4 | NIT2\_HUMAN | NIT2 | Omega-amidase NIT2 | 276 | 30.608 | 6.406703e+04 | 2.463693e+06 | 910412.9 | 1386664.4 | 123713.5 | 42902.64 | 71.38 | 59.8 | 30.1 | 20.3 |
| sp\|P19474\|RO52\_HUMAN | P19474 | RO52\_HUMAN | TRIM21 | E3 ubiquitin-protein ligase TRIM21 | 475 | 54.170 | 6.843711e+04 | 4.657659e+06 | 909535.75 | 3382864.5 | 341889.8 | 23368.787 | 42.11 | 40.4 | 28.6 | 17.3 |
| sp\|Q5VWZ2\|LYPL1\_HUMAN | Q5VWZ2 | LYPL1\_HUMAN | LYPLAL1 | Lysophospholipase-like protein 1 | 237 | 26.316 | 8.253142e+04 | 2.728704e+06 | 906080.94 | 1730120.5 | 92502.06 |  | 30.38 | 40.9 | 26.6 |  |
| sp\|Q9UEY8\|ADDG\_HUMAN | Q9UEY8 | ADDG\_HUMAN | ADD3 | Gamma-adducin | 706 | 79.155 | 1.322136e+05 | 1.314836e+07 | 904139.4 | 1692987.0 | 9566050.0 | 985181.44 | 37.11 | 46.9 | 39.7 | 19.5 |
| sp\|P12814\|ACTN1\_HUMAN | P12814 | ACTN1\_HUMAN | ACTN1 | Alpha-actinin-1 | 892 | 103.058 | 1.308066e+04 | 1.693669e+06 | 903837.1 | 637260.0 | 135610.81 | 16960.676 | 19.73 | 39.5 | 28.4 | 8.1 |
| sp\|Q99536\|VAT1\_HUMAN | Q99536 | VAT1\_HUMAN | VAT1 | Synaptic vesicle membrane protein VAT-1 homolog | 393 | 41.920 | 5.571073e+04 | 2.934116e+06 | 903736.56 | 1546034.9 | 345547.22 | 138797.2 | 39.69 | 72.0 | 17.8 | 15.0 |
| sp\|P52597\|HNRPF\_HUMAN | P52597 | HNRPF\_HUMAN | HNRNPF | Heterogeneous nuclear ribonucleoprotein F | 415 | 45.672 | 7.170402e+04 | 4.114440e+06 | 898009.75 | 2423289.2 | 460272.0 | 332868.9 | 37.11 | 54.0 | 31.1 | 22.9 |
| sp\|Q8TDQ7\|GNPI2\_HUMAN | Q8TDQ7 | GNPI2\_HUMAN | GNPDA2 | Glucosamine-6-phosphate isomerase 2 | 276 | 31.085 | 1.294463e+05 | 5.055424e+06 | 897637.1 | 4082106.0 | 75681.3 |  | 46.01 | 72.8 | 27.9 |  |
| sp\|Q9UBS4\|DJB11\_HUMAN | Q9UBS4 | DJB11\_HUMAN | DNAJB11 | DnaJ homolog subfamily B member 11 | 358 | 40.514 | 4.187481e+04 | 2.131450e+06 | 893558.3 | 161042.16 | 899298.6 | 177551.02 | 9.22 | 47.5 | 30.7 | 19.6 |
| sp\|P27448\|MARK3\_HUMAN | P27448 | MARK3\_HUMAN | MARK3 | MAP/microtubule affinity-regulating kinase 3 | 753 | 84.429 | 5.486966e+04 | 5.820242e+06 | 887841.5 | 2229619.2 | 902443.6 | 1800337.8 | 52.06 | 66.0 | 45.8 | 37.8 |
| sp\|Q96QK1\|VPS35\_HUMAN | Q96QK1 | VPS35\_HUMAN | VPS35 | Vacuolar protein sorting-associated protein 35 | 796 | 91.707 | 2.583370e+04 | 2.976502e+06 | 887378.2 | 1960858.8 | 73784.914 | 54480.344 | 39.45 | 42.0 | 19.5 | 7.0 |
| sp\|Q9H993\|ARMT1\_HUMAN | Q9H993 | ARMT1\_HUMAN | ARMT1 | Damage-control phosphatase ARMT1 | 441 | 51.172 | 2.095548e+04 | 1.347247e+06 | 886568.5 | 460678.47 |  |  | 24.72 | 40.6 |  |  |
| sp\|P55795\|HNRH2\_HUMAN | P55795 | HNRH2\_HUMAN | HNRNPH2 | Heterogeneous nuclear ribonucleoprotein H2 | 449 | 49.264 | 2.237933e+04 | 1.385141e+06 | 881366.5 | 503774.28 |  |  | 16.93 | 42.8 |  |  |
| sp\|O94966\|UBP19\_HUMAN | O94966 | UBP19\_HUMAN | USP19 | Ubiquitin carboxyl-terminal hydrolase 19 | 1318 | 145.651 | 1.523937e+04 | 2.788672e+06 | 881347.7 | 1765648.9 | 141675.86 |  | 32.4 | 36.0 | 21.9 |  |
| sp\|O75663\|TIPRL\_HUMAN | O75663 | TIPRL\_HUMAN | TIPRL | TIP41-like protein | 272 | 31.444 | 1.213323e+05 | 4.793263e+06 | 879220.94 | 3236297.5 | 526335.06 | 151409.45 | 59.19 | 55.5 | 26.5 | 18.0 |
| sp\|O60547\|GMDS\_HUMAN | O60547 | GMDS\_HUMAN | GMDS | GDP-mannose 4,6 dehydratase | 372 | 41.950 | 4.916186e+04 | 2.591060e+06 | 874507.56 | 1494461.0 | 222090.97 |  | 54.03 | 67.7 | 27.7 |  |
| sp\|P14923\|PLAK\_HUMAN | P14923 | PLAK\_HUMAN | JUP | Junction plakoglobin | 745 | 81.745 | 6.864380e+04 | 7.049846e+06 | 869735.9 | 1562986.4 | 1821366.2 | 2795758.0 | 36.51 | 49.0 | 36.6 | 31.5 |
| sp\|Q13283\|G3BP1\_HUMAN | Q13283 | G3BP1\_HUMAN | G3BP1 | Ras GTPase-activating protein-binding protein 1 | 466 | 52.164 | 3.776462e+04 | 2.474989e+06 | 868953.56 | 1357883.9 | 201143.12 | 47008.016 | 32.19 | 54.7 | 35.6 | 14.8 |
| sp\|Q9Y5S9\|RBM8A\_HUMAN | Q9Y5S9 | RBM8A\_HUMAN | RBM8A | RNA-binding protein 8A | 174 | 19.889 | 1.214635e+05 | 3.035119e+06 | 868888.44 | 1991408.1 | 174822.44 |  | 31.61 | 38.5 | 26.4 |  |
| sp\|Q13596\|SNX1\_HUMAN | Q13596 | SNX1\_HUMAN | SNX1 | Sorting nexin-1 | 522 | 59.070 | 1.746623e+04 | 1.296234e+06 | 868297.7 | 427936.12 |  |  | 17.62 | 34.5 |  |  |
| sp\|Q9Y371\|SHLB1\_HUMAN | Q9Y371 | SHLB1\_HUMAN | SH3GLB1 | Endophilin-B1 | 365 | 40.796 | 6.673431e+04 | 3.420455e+06 | 864046.7 | 951126.7 | 959263.75 | 646017.9 | 26.03 | 33.7 | 17.8 | 13.2 |
| sp\|P35609\|ACTN2\_HUMAN | P35609 | ACTN2\_HUMAN | ACTN2 | Alpha-actinin-2 | 894 | 98.340 | 6.981866e+03 | 8.626187e+05 | 862618.7 |  |  |  | 14.54 |  |  |  |
| sp\|O75935\|DCTN3\_HUMAN | O75935 | DCTN3\_HUMAN | DCTN3 | Dynactin subunit 3 | 186 | 21.119 | 5.888618e+04 | 1.562442e+06 | 860543.0 | 62218.977 | 529327.1 | 110352.94 | 24.19 | 41.9 | 62.4 | 31.7 |
| sp\|Q9H9S4\|CB39L\_HUMAN | Q9H9S4 | CB39L\_HUMAN | CAB39L | Calcium-binding protein 39-like | 337 | 39.088 | 2.936496e+04 | 1.442082e+06 | 859165.1 | 582916.9 |  |  | 25.82 | 37.1 |  |  |
| sp\|P31944\|CASPE\_HUMAN | P31944 | CASPE\_HUMAN | CASP14 | Caspase-14 | 242 | 27.680 | 2.034909e+05 | 7.076660e+06 | 858486.94 | 834835.4 | 2883999.0 | 2499338.2 | 40.08 | 57.9 | 42.6 | 29.8 |
| sp\|Q9UKL6\|PPCT\_HUMAN | Q9UKL6 | PPCT\_HUMAN | PCTP | Phosphatidylcholine transfer protein | 214 | 24.843 | 9.881479e+04 | 3.084204e+06 | 857588.4 | 1764973.9 | 280502.06 | 181140.03 | 24.3 | 21.0 | 24.3 | 13.1 |
| sp\|P13667\|PDIA4\_HUMAN | P13667 | PDIA4\_HUMAN | PDIA4 | Protein disulfide-isomerase A4 | 645 | 72.932 | 4.355579e+04 | 3.990995e+06 | 856197.9 | 722616.3 | 2158034.0 | 254146.88 | 41.09 | 49.6 | 46.8 | 22.6 |
| sp\|Q8TBZ2\|MYBPP\_HUMAN | Q8TBZ2 | MYBPP\_HUMAN | MYCBPAP | MYCBP-associated protein | 947 | 104.170 | 6.525789e+03 | 8.540688e+05 | 854068.75 |  |  |  | 1.9 |  |  |  |
| sp\|O14523\|C2C2L\_HUMAN | O14523 | C2C2L\_HUMAN | C2CD2L | Phospholipid transfer protein C2CD2L | 706 | 76.181 | 1.405284e+05 | 1.345017e+07 | 853937.44 | 423610.44 | 4107296.8 | 8065329.0 | 38.24 | 35.7 | 30.6 | 28.0 |
| sp\|Q7Z478\|DHX29\_HUMAN | Q7Z478 | DHX29\_HUMAN | DHX29 | ATP-dependent RNA helicase DHX29 | 1369 | 155.236 | 1.604734e+04 | 3.129772e+06 | 853444.25 | 94047.35 | 1444100.0 | 738180.4 | 11.83 | 31.8 | 18.8 | 10.0 |
| sp\|P53618\|COPB\_HUMAN | P53618 | COPB\_HUMAN | COPB1 | Coatomer subunit beta | 953 | 107.142 | 1.222536e+04 | 1.645654e+06 | 853412.3 | 691702.4 |  | 100539.57 | 24.87 | 35.4 |  | 6.9 |
| sp\|Q9Y5X3\|SNX5\_HUMAN | Q9Y5X3 | SNX5\_HUMAN | SNX5 | Sorting nexin-5 | 404 | 46.816 | 4.798862e+04 | 2.822602e+06 | 851658.8 | 1172229.1 | 673021.25 | 125693.09 | 38.37 | 48.3 | 28.0 | 20.0 |
| sp\|Q7Z4H3\|HDDC2\_HUMAN | Q7Z4H3 | HDDC2\_HUMAN | HDDC2 | 5'-deoxynucleotidase HDDC2 | 204 | 23.390 | 1.632203e+05 | 4.796468e+06 | 850846.0 | 3740281.5 | 141796.44 | 63544.297 | 49.51 | 62.7 | 20.1 | 33.8 |
| sp\|Q9Y5X1\|SNX9\_HUMAN | Q9Y5X1 | SNX9\_HUMAN | SNX9 | Sorting nexin-9 | 595 | 66.592 | 3.427432e+04 | 2.867531e+06 | 849129.5 | 1786352.6 | 232048.42 |  | 43.7 | 50.3 | 16.8 |  |
| sp\|Q9P2X3\|IMPCT\_HUMAN | Q9P2X3 | IMPCT\_HUMAN | IMPACT | Protein IMPACT | 320 | 36.476 | 7.251489e+04 | 3.323162e+06 | 843984.94 | 2270337.2 | 149666.06 | 59173.93 | 43.12 | 45.0 | 24.4 | 13.1 |
| sp\|P13987\|CD59\_HUMAN | P13987 | CD59\_HUMAN | CD59 | CD59 glycoprotein | 128 | 14.177 | 8.440340e+05 | 1.503355e+07 | 842516.1 |  | 6346833.0 | 7844197.5 | 17.97 |  | 28.1 | 28.1 |
| sp\|P52788\|SPSY\_HUMAN | P52788 | SPSY\_HUMAN | SMS | Spermine synthase | 366 | 41.268 | 2.505957e+04 | 1.299285e+06 | 841345.25 | 457939.3 |  |  | 45.9 | 30.6 |  |  |
| sp\|Q96BN8\|OTUL\_HUMAN | Q96BN8 | OTUL\_HUMAN | OTULIN | Ubiquitin thioesterase otulin | 352 | 40.263 | 9.671542e+04 | 4.892367e+06 | 840364.6 | 3088791.0 | 787297.06 | 175914.0 | 46.88 | 49.4 | 50.0 | 8.2 |
| sp\|P48444\|COPD\_HUMAN | P48444 | COPD\_HUMAN | ARCN1 | Coatomer subunit delta | 511 | 57.210 | 2.671201e+04 | 1.919975e+06 | 840337.2 | 819267.6 | 187698.98 | 72671.66 | 37.18 | 62.8 | 31.3 | 15.7 |
| sp\|P09651\|ROA1\_HUMAN | P09651 | ROA1\_HUMAN | HNRNPA1 | Heterogeneous nuclear ribonucleoprotein A1 | 372 | 38.747 | 1.082226e+05 | 5.268333e+06 | 840297.4 | 3749768.0 | 595586.9 | 82680.47 | 34.95 | 53.2 | 35.2 | 30.4 |
| sp\|Q9BTW9\|TBCD\_HUMAN | Q9BTW9 | TBCD\_HUMAN | TBCD | Tubulin-specific chaperone D | 1192 | 132.600 | 8.371192e+03 | 1.394595e+06 | 840234.1 | 554360.5 |  |  | 23.99 | 36.1 |  |  |
| sp\|Q15057\|ACAP2\_HUMAN | Q15057 | ACAP2\_HUMAN | ACAP2 | Arf-GAP with coiled-coil, ANK repeat and PH domain-containing protein 2 | 778 | 88.029 | 2.898996e+04 | 3.206199e+06 | 838671.25 | 2077838.8 | 149691.75 | 139997.52 | 40.36 | 55.3 | 31.6 | 17.9 |
| sp\|Q99447\|PCY2\_HUMAN | Q99447 | PCY2\_HUMAN | PCYT2 | Ethanolamine-phosphate cytidylyltransferase | 389 | 43.835 | 5.046531e+04 | 2.779273e+06 | 837972.6 | 1772479.5 | 90143.766 | 78676.73 | 51.93 | 66.8 | 43.7 | 15.7 |
| sp\|O94903\|PLPHP\_HUMAN | O94903 | PLPHP\_HUMAN | PLPBP | Pyridoxal phosphate homeostasis protein | 275 | 30.344 | 1.379203e+05 | 5.257972e+06 | 832487.7 | 4147839.2 | 173372.52 | 104272.164 | 40.0 | 70.5 | 24.7 | 17.5 |
| sp\|Q92542\|NICA\_HUMAN | Q92542 | NICA\_HUMAN | NCSTN | Nicastrin | 709 | 78.411 | 1.144735e+05 | 1.127714e+07 | 832192.25 | 71161.836 | 3913730.0 | 6460059.5 | 16.5 | 21.6 | 22.6 | 15.1 |
| sp\|Q8WXX5\|DNJC9\_HUMAN | Q8WXX5 | DNJC9\_HUMAN | DNAJC9 | DnaJ homolog subfamily C member 9 | 260 | 29.910 | 1.147788e+05 | 4.313156e+06 | 831962.25 | 2991586.0 | 310789.12 | 178818.22 | 51.92 | 69.2 | 50.0 | 18.8 |
| sp\|O75170\|PP6R2\_HUMAN | O75170 | PP6R2\_HUMAN | PPP6R2 | Serine/threonine-protein phosphatase 6 regulatory subunit 2 | 966 | 104.942 | 2.114317e+04 | 2.787639e+06 | 828502.3 | 1812423.6 | 55683.64 | 91029.58 | 29.92 | 50.3 | 25.3 | 13.4 |
| sp\|P78527\|PRKDC\_HUMAN | P78527 | PRKDC\_HUMAN | PRKDC | DNA-dependent protein kinase catalytic subunit | 4128 | 469.089 | 2.569769e+03 | 1.514490e+06 | 824889.75 | 556646.6 |  | 132954.08 | 16.72 | 34.3 |  | 11.3 |
| sp\|Q15021\|CND1\_HUMAN | Q15021 | CND1\_HUMAN | NCAPD2 | Condensin complex subunit 1 | 1401 | 157.182 | 9.457530e+03 | 1.867660e+06 | 824522.8 | 869851.25 |  | 173285.5 | 39.83 | 45.4 |  | 12.0 |
| sp\|P23193\|TCEA1\_HUMAN | P23193 | TCEA1\_HUMAN | TCEA1 | Transcription elongation factor A protein 1 | 301 | 33.970 | 1.172652e+05 | 5.004742e+06 | 822559.9 | 3900033.8 | 142611.81 | 139536.25 | 40.2 | 67.4 | 23.3 | 30.6 |
| sp\|P60981\|DEST\_HUMAN | P60981 | DEST\_HUMAN | DSTN | Destrin | 165 | 18.506 | 8.190439e+04 | 1.904307e+06 | 822511.0 | 947969.6 | 133826.0 |  | 36.97 | 67.3 | 33.9 |  |
| sp\|O14617\|AP3D1\_HUMAN | O14617 | AP3D1\_HUMAN | AP3D1 | AP-3 complex subunit delta-1 | 1153 | 130.158 | 7.580515e+03 | 1.239615e+06 | 820489.1 | 419125.62 |  |  | 10.06 | 30.3 |  |  |
| sp\|Q99816\|TS101\_HUMAN | Q99816 | TS101\_HUMAN | TSG101 | Tumor susceptibility gene 101 protein | 390 | 43.944 | 2.283217e+05 | 1.260561e+07 | 818658.5 | 2306007.8 | 3457148.5 | 6023796.5 | 35.38 | 46.9 | 38.5 | 24.1 |
| sp\|P33527\|MRP1\_HUMAN | P33527 | MRP1\_HUMAN | ABCC1 | Multidrug resistance-associated protein 1 | 1531 | 171.591 | 3.723683e+04 | 8.027574e+06 | 817904.0 | 794939.5 | 1008091.25 | 5406639.0 | 31.22 | 33.2 | 25.3 | 29.1 |
| sp\|Q8TEX9\|IPO4\_HUMAN | Q8TEX9 | IPO4\_HUMAN | IPO4 | Importin-4 | 1081 | 118.715 | 9.783715e+03 | 1.459239e+06 | 817693.0 | 641546.4 |  |  | 26.46 | 42.6 |  |  |
| sp\|Q63HN8\|RN213\_HUMAN | Q63HN8 | RN213\_HUMAN | RNF213 | E3 ubiquitin-protein ligase RNF213 | 5207 | 591.407 | 2.002430e+03 | 1.487856e+06 | 817106.3 | 366708.6 |  | 304041.16 | 10.12 | 31.9 |  | 9.5 |
| sp\|Q6VN20\|RBP10\_HUMAN | Q6VN20 | RBP10\_HUMAN | RANBP10 | Ran-binding protein 10 | 620 | 67.257 | 2.332387e+04 | 1.970858e+06 | 816287.56 | 683239.4 | 388535.12 | 82795.84 | 33.23 | 47.7 | 22.9 | 12.3 |
| sp\|P24043\|LAMA2\_HUMAN | P24043 | LAMA2\_HUMAN | LAMA2 | Laminin subunit alpha-2 | 3122 | 343.420 | 1.883361e+03 | 8.125991e+05 | 812599.06 |  |  |  | 0.74 |  |  |  |
| sp\|P46019\|KPB2\_HUMAN | P46019 | KPB2\_HUMAN | PHKA2 | Phosphorylase b kinase regulatory subunit alpha, liver isoform | 1235 | 138.408 | 8.040371e+03 | 1.398152e+06 | 810852.1 | 587300.0 |  |  | 37.09 | 44.6 |  |  |
| sp\|P57764\|GSDMD\_HUMAN | P57764 | GSDMD\_HUMAN | GSDMD | Gasdermin-D | 484 | 52.801 | 1.856732e+04 | 1.231710e+06 | 809599.6 | 306711.53 | 115398.91 |  | 16.53 | 34.1 | 13.0 |  |
| sp\|Q96T76\|MMS19\_HUMAN | Q96T76 | MMS19\_HUMAN | MMS19 | MMS19 nucleotide excision repair protein homolog | 1030 | 113.290 | 1.150940e+04 | 1.638179e+06 | 808346.25 | 811599.75 |  | 18232.559 | 24.47 | 45.0 |  | 11.6 |
| sp\|P05388\|RLA0\_HUMAN | P05388 | RLA0\_HUMAN | RPLP0 | Large ribosomal subunit protein uL10 | 317 | 34.274 | 7.962491e+04 | 3.428711e+06 | 807317.1 | 2166491.5 | 380467.12 | 74435.234 | 48.58 | 54.6 | 30.3 | 5.7 |
| sp\|Q13576\|IQGA2\_HUMAN | Q13576 | IQGA2\_HUMAN | IQGAP2 | Ras GTPase-activating-like protein IQGAP2 | 1575 | 180.578 | 1.612152e+04 | 3.657530e+06 | 806692.44 | 1464675.4 | 1316623.0 | 69539.56 | 43.11 | 46.5 | 32.5 | 10.8 |
| sp\|Q9Y624\|JAM1\_HUMAN | Q9Y624 | JAM1\_HUMAN | F11R | Junctional adhesion molecule A | 299 | 32.583 | 4.344759e+05 | 1.778582e+07 | 803357.2 | 3896624.0 | 2400969.5 | 10684869.0 | 34.78 | 47.5 | 33.1 | 34.4 |
| sp\|Q8IY17\|PLPL6\_HUMAN | Q8IY17 | PLPL6\_HUMAN | PNPLA6 | Patatin-like phospholipase domain-containing protein 6 | 1375 | 150.954 | 2.627857e+04 | 4.983833e+06 | 800689.6 | 1359363.0 | 1445690.8 | 1378089.2 | 35.49 | 41.8 | 27.6 | 17.3 |
| sp\|Q9H0R4\|HDHD2\_HUMAN | Q9H0R4 | HDHD2\_HUMAN | HDHD2 | Haloacid dehalogenase-like hydrolase domain-containing protein 2 | 259 | 28.536 | 4.622544e+05 | 1.657263e+07 | 800491.6 | 15772134.0 |  |  | 45.17 | 82.6 |  |  |
| sp\|Q7Z2Z2\|EFL1\_HUMAN | Q7Z2Z2 | EFL1\_HUMAN | EFL1 | Elongation factor-like GTPase 1 | 1120 | 125.430 | 9.853791e+03 | 1.552823e+06 | 797792.0 | 755030.94 |  |  | 11.79 | 37.9 |  |  |
| sp\|Q6KB66\|K2C80\_HUMAN | Q6KB66 | K2C80\_HUMAN | KRT80 | Keratin, type II cytoskeletal 80 | 452 | 50.525 | 8.888109e+04 | 5.641997e+06 | 796257.4 | 483837.9 | 1491051.4 | 2870850.0 | 44.25 | 38.9 | 47.1 | 55.3 |
| sp\|P19256\|LFA3\_HUMAN | P19256 | LFA3\_HUMAN | CD58 | Lymphocyte function-associated antigen 3 | 250 | 28.147 | 4.052968e+05 | 1.433252e+07 | 795466.44 | 1777242.0 | 5713519.5 | 6046290.0 | 14.0 | 19.6 | 11.2 | 18.4 |
| sp\|Q15386\|UBE3C\_HUMAN | Q15386 | UBE3C\_HUMAN | UBE3C | Ubiquitin-protein ligase E3C | 1083 | 123.923 | 1.256704e+04 | 1.956599e+06 | 795178.0 | 1076459.4 |  | 84961.98 | 37.67 | 33.1 |  | 9.3 |
| sp\|Q15185\|TEBP\_HUMAN | Q15185 | TEBP\_HUMAN | PTGES3 | Prostaglandin E synthase 3 | 160 | 18.697 | 2.482809e+05 | 5.832199e+06 | 795053.4 | 4564794.0 | 259978.62 | 212373.0 | 38.75 | 53.1 | 49.4 | 16.9 |
| sp\|P01040\|CYTA\_HUMAN | P01040 | CYTA\_HUMAN | CSTA | Cystatin-A | 98 | 11.006 | 1.909956e+05 | 2.641010e+06 | 791016.25 |  | 685126.9 | 1164867.0 | 76.53 |  | 83.7 | 36.7 |
| sp\|Q13610\|PWP1\_HUMAN | Q13610 | PWP1\_HUMAN | PWP1 | Periodic tryptophan protein 1 homolog | 501 | 55.828 | 2.448293e+04 | 1.717246e+06 | 786955.2 | 867896.1 | 55289.75 | 7105.061 | 33.13 | 30.5 | 25.1 | 11.4 |
| sp\|Q9UNF0\|PACN2\_HUMAN | Q9UNF0 | PACN2\_HUMAN | PACSIN2 | Protein kinase C and casein kinase substrate in neurons protein 2 | 486 | 55.739 | 2.463969e+04 | 1.725487e+06 | 786232.4 | 756656.6 | 131786.89 | 50810.812 | 24.9 | 34.6 | 17.1 | 11.5 |
| sp\|Q04656\|ATP7A\_HUMAN | Q04656 | ATP7A\_HUMAN | ATP7A | Copper-transporting ATPase 1 | 1500 | 163.373 | 6.366839e+04 | 1.306837e+07 | 786135.7 | 854342.75 | 2334662.2 | 9093225.0 | 26.53 | 28.6 | 26.5 | 22.1 |
| sp\|P47897\|SYQ\_HUMAN | P47897 | SYQ\_HUMAN | QARS1 | Glutamine--tRNA ligase | 775 | 87.799 | 2.480189e+04 | 2.735845e+06 | 783197.94 | 1332479.8 | 620167.25 |  | 26.58 | 49.7 | 21.4 |  |
| sp\|P34896\|GLYC\_HUMAN | P34896 | GLYC\_HUMAN | SHMT1 | Serine hydroxymethyltransferase, cytosolic | 483 | 53.083 | 3.775620e+04 | 2.518030e+06 | 782094.2 | 1083490.8 | 652445.06 |  | 47.41 | 63.4 | 32.3 |  |
| sp\|Q9BYT8\|NEUL\_HUMAN | Q9BYT8 | NEUL\_HUMAN | NLN | Neurolysin, mitochondrial | 704 | 80.652 | 1.001422e+04 | 1.014728e+06 | 777308.8 | 237419.0 |  |  | 27.41 | 33.2 |  |  |
| sp\|Q13131\|AAPK1\_HUMAN | Q13131 | AAPK1\_HUMAN | PRKAA1 | 5'-AMP-activated protein kinase catalytic subunit alpha-1 | 559 | 64.009 | 3.701310e+04 | 2.976553e+06 | 776524.8 | 755003.06 | 1071998.4 | 373027.1 | 33.99 | 39.5 | 16.1 | 15.2 |
| sp\|P13861\|KAP2\_HUMAN | P13861 | KAP2\_HUMAN | PRKAR2A | cAMP-dependent protein kinase type II-alpha regulatory subunit | 404 | 45.518 | 2.948664e+04 | 1.686264e+06 | 775782.6 | 730967.75 | 179513.62 |  | 42.33 | 47.0 | 25.2 |  |
| sp\|Q9UI10\|EI2BD\_HUMAN | Q9UI10 | EI2BD\_HUMAN | EIF2B4 | Translation initiation factor eIF-2B subunit delta | 523 | 57.557 | 3.702692e+04 | 2.677521e+06 | 772794.2 | 1646591.2 | 220610.08 | 37525.773 | 43.02 | 60.4 | 43.4 | 17.2 |
| sp\|Q12959\|DLG1\_HUMAN | Q12959 | DLG1\_HUMAN | DLG1 | Disks large homolog 1 | 904 | 100.455 | 1.703746e+04 | 2.150273e+06 | 771173.75 | 592745.8 | 700062.94 | 86290.4 | 39.49 | 35.7 | 22.8 | 12.3 |
| sp\|Q15276\|RABE1\_HUMAN | Q15276 | RABE1\_HUMAN | RABEP1 | Rab GTPase-binding effector protein 1 | 862 | 99.290 | 1.384842e+04 | 1.727519e+06 | 768026.4 | 583417.25 | 197093.23 | 178982.44 | 44.9 | 44.1 | 28.8 | 23.9 |
| sp\|Q92990\|GLMN\_HUMAN | Q92990 | GLMN\_HUMAN | GLMN | Glomulin | 594 | 68.208 | 2.584426e+04 | 2.214709e+06 | 763542.4 | 1425886.0 | 25280.271 |  | 35.86 | 37.2 | 8.6 |  |
| sp\|Q96HE9\|PRR11\_HUMAN | Q96HE9 | PRR11\_HUMAN | PRR11 | Proline-rich protein 11 | 360 | 39.600 | 1.532751e+04 | 7.625772e+05 | 762577.25 |  |  |  | 3.33 |  |  |  |
| sp\|Q9P0J7\|KCMF1\_HUMAN | Q9P0J7 | KCMF1\_HUMAN | KCMF1 | E3 ubiquitin-protein ligase KCMF1 | 381 | 41.945 | 7.777277e+04 | 4.098500e+06 | 759684.25 | 2822994.5 | 515821.06 |  | 44.09 | 43.6 | 16.5 |  |
| sp\|Q9Y3C6\|PPIL1\_HUMAN | Q9Y3C6 | PPIL1\_HUMAN | PPIL1 | Peptidyl-prolyl cis-trans isomerase-like 1 | 166 | 18.237 | 3.205841e+05 | 7.345350e+06 | 759485.6 | 6343508.5 | 200681.12 | 41674.383 | 31.93 | 74.7 | 30.1 | 13.3 |
| sp\|Q96EP5\|DAZP1\_HUMAN | Q96EP5 | DAZP1\_HUMAN | DAZAP1 | DAZ-associated protein 1 | 407 | 43.383 | 3.237893e+04 | 1.764815e+06 | 758703.6 | 860077.75 | 119099.35 | 26934.236 | 21.62 | 45.7 | 11.5 | 6.1 |
| sp\|P50416\|CPT1A\_HUMAN | P50416 | CPT1A\_HUMAN | CPT1A | Carnitine O-palmitoyltransferase 1, liver isoform | 773 | 88.368 | 4.613329e+04 | 5.121847e+06 | 756071.1 | 958373.56 | 1546152.0 | 1861250.6 | 27.17 | 39.6 | 31.7 | 23.8 |
| sp\|O43143\|DHX15\_HUMAN | O43143 | DHX15\_HUMAN | DHX15 | ATP-dependent RNA helicase DHX15 | 795 | 90.933 | 2.035694e+04 | 2.325686e+06 | 755792.8 | 1303855.0 | 223747.83 | 42290.594 | 37.11 | 41.4 | 28.2 | 11.4 |
| sp\|Q8TF17\|S3TC2\_HUMAN | Q8TF17 | S3TC2\_HUMAN | SH3TC2 | SH3 domain and tetratricopeptide repeat-containing protein 2 | 1288 | 144.777 | 4.552063e+03 | 8.279898e+05 | 755324.0 |  |  | 72665.805 | 1.55 |  |  | 13.6 |
| sp\|Q92538\|GBF1\_HUMAN | Q92538 | GBF1\_HUMAN | GBF1 | Golgi-specific brefeldin A-resistance guanine nucleotide exchange factor 1 | 1859 | 206.446 | 5.843696e+03 | 1.515693e+06 | 752410.06 | 517707.0 | 77992.6 | 167583.45 | 19.53 | 37.5 | 19.6 | 11.7 |
| sp\|Q9NWV8\|BABA1\_HUMAN | Q9NWV8 | BABA1\_HUMAN | BABAM1 | BRISC and BRCA1-A complex member 1 | 329 | 36.560 | 5.204480e+04 | 2.390565e+06 | 749186.56 | 1350264.4 | 233405.67 | 57708.844 | 23.1 | 52.9 | 24.9 | 16.7 |
| sp\|Q9H7D7\|WDR26\_HUMAN | Q9H7D7 | WDR26\_HUMAN | WDR26 | WD repeat-containing protein 26 | 661 | 72.124 | 2.173456e+04 | 1.969463e+06 | 749000.6 | 972956.8 | 172473.77 | 75031.9 | 23.75 | 23.0 | 15.0 | 6.7 |
| sp\|Q12882\|DPYD\_HUMAN | Q12882 | DPYD\_HUMAN | DPYD | Dihydropyrimidine dehydrogenase [NADP(+)] | 1025 | 111.401 | 8.146163e+03 | 1.140143e+06 | 748801.3 | 391341.8 |  |  | 32.29 | 40.8 |  |  |
| sp\|Q14116\|IL18\_HUMAN | Q14116 | IL18\_HUMAN | IL18 | Interleukin-18 | 193 | 22.326 | 1.025591e+05 | 2.876751e+06 | 747637.6 | 2129113.8 |  |  | 32.12 | 60.1 |  |  |
| sp\|O43583\|DENR\_HUMAN | O43583 | DENR\_HUMAN | DENR | Density-regulated protein | 198 | 22.092 | 3.779559e+05 | 1.049043e+07 | 747509.56 | 7019236.5 | 2457373.5 | 266311.84 | 22.73 | 58.6 | 41.4 | 31.3 |
| sp\|Q9C0H9\|SRCN1\_HUMAN | Q9C0H9 | SRCN1\_HUMAN | SRCIN1 | SRC kinase signaling inhibitor 1 | 1183 | 130.130 | 4.565843e+03 | 7.464754e+05 | 746475.44 |  |  |  | 1.94 |  |  |  |
| sp\|Q8WVJ2\|NUDC2\_HUMAN | Q8WVJ2 | NUDC2\_HUMAN | NUDCD2 | NudC domain-containing protein 2 | 157 | 17.676 | 9.805466e+04 | 2.177556e+06 | 746380.06 | 382635.1 | 723962.2 | 324579.0 | 35.03 | 47.8 | 30.6 | 17.2 |
| sp\|Q9UJ70\|NAGK\_HUMAN | Q9UJ70 | NAGK\_HUMAN | NAGK | N-acetyl-D-glucosamine kinase | 344 | 37.376 | 6.404424e+04 | 3.007392e+06 | 744998.5 | 1825574.9 | 313112.97 | 123705.53 | 52.33 | 44.8 | 27.0 | 22.1 |
| sp\|Q86VN1\|VPS36\_HUMAN | Q86VN1 | VPS36\_HUMAN | VPS36 | Vacuolar protein-sorting-associated protein 36 | 386 | 43.817 | 3.626526e+04 | 1.996414e+06 | 740609.6 | 852041.94 | 336648.3 | 67114.3 | 63.99 | 60.9 | 39.9 | 16.6 |
| sp\|P16615\|AT2A2\_HUMAN | P16615 | AT2A2\_HUMAN | ATP2A2 | Sarcoplasmic/endoplasmic reticulum calcium ATPase 2 | 1042 | 114.757 | 2.501182e+04 | 3.606133e+06 | 737047.44 | 1456267.8 | 687782.56 | 725034.75 | 14.11 | 34.1 | 25.2 | 23.2 |
| sp\|Q9BTY7\|HGH1\_HUMAN | Q9BTY7 | HGH1\_HUMAN | HGH1 | Protein HGH1 homolog | 390 | 42.129 | 4.085593e+04 | 2.162487e+06 | 734763.6 | 721903.56 | 598463.7 | 107355.73 | 55.13 | 53.3 | 38.7 | 12.6 |
| sp\|Q9NTJ5\|SAC1\_HUMAN | Q9NTJ5 | SAC1\_HUMAN | SACM1L | Phosphatidylinositol-3-phosphatase SAC1 | 587 | 66.967 | 7.991581e+04 | 6.723736e+06 | 734196.3 | 2569042.5 | 879134.1 | 2541363.5 | 52.64 | 52.3 | 33.2 | 25.4 |
| sp\|Q9Y5K6\|CD2AP\_HUMAN | Q9Y5K6 | CD2AP\_HUMAN | CD2AP | CD2-associated protein | 639 | 71.451 | 6.639985e+04 | 5.960636e+06 | 730748.8 | 4513589.0 | 528808.5 | 187489.52 | 59.62 | 72.6 | 32.2 | 18.5 |
| sp\|Q9NTX5\|ECHD1\_HUMAN | Q9NTX5 | ECHD1\_HUMAN | ECHDC1 | Ethylmalonyl-CoA decarboxylase | 307 | 33.698 | 7.831696e+04 | 3.315714e+06 | 723394.9 | 2223395.2 | 261572.84 | 107351.34 | 48.86 | 54.7 | 46.3 | 9.1 |
| sp\|Q9UK41\|VPS28\_HUMAN | Q9UK41 | VPS28\_HUMAN | VPS28 | Vacuolar protein sorting-associated protein 28 homolog | 221 | 25.425 | 1.058390e+05 | 3.380833e+06 | 722631.25 | 1536010.6 | 943824.06 | 178367.08 | 43.44 | 76.0 | 56.1 | 19.9 |
| sp\|Q92734\|TFG\_HUMAN | Q92734 | TFG\_HUMAN | TFG | Protein TFG | 400 | 43.448 | 1.389874e+05 | 7.586866e+06 | 720242.6 | 1236657.5 | 4996265.0 | 633700.94 | 26.25 | 37.0 | 25.8 | 9.0 |
| sp\|P24298\|ALAT1\_HUMAN | P24298 | ALAT1\_HUMAN | GPT | Alanine aminotransferase 1 | 496 | 54.637 | 1.674334e+04 | 1.149334e+06 | 719783.1 | 331745.16 | 75723.23 | 22082.135 | 49.8 | 66.3 | 23.2 | 5.8 |
| sp\|O43708\|MAAI\_HUMAN | O43708 | MAAI\_HUMAN | GSTZ1 | Maleylacetoacetate isomerase | 216 | 24.212 | 9.703590e+04 | 2.951755e+06 | 718164.8 | 2233590.0 |  |  | 32.41 | 44.4 |  |  |
| sp\|Q9Y4G6\|TLN2\_HUMAN | Q9Y4G6 | TLN2\_HUMAN | TLN2 | Talin-2 | 2542 | 271.613 | 8.222640e+03 | 2.805944e+06 | 713652.4 | 1745711.4 | 240109.86 | 106470.266 | 35.05 | 50.5 | 25.9 | 12.8 |
| sp\|Q8IWB7\|WDFY1\_HUMAN | Q8IWB7 | WDFY1\_HUMAN | WDFY1 | WD repeat and FYVE domain-containing protein 1 | 410 | 46.324 | 4.052103e+04 | 2.358325e+06 | 712039.2 | 1499284.5 | 100802.0 | 46199.344 | 44.63 | 50.7 | 29.5 | 14.4 |
| sp\|P19784\|CSK22\_HUMAN | P19784 | CSK22\_HUMAN | CSNK2A2 | Casein kinase II subunit alpha' | 350 | 41.213 | 2.340759e+05 | 1.212015e+07 | 708928.9 | 1232260.5 | 10178962.0 |  | 58.29 | 60.6 | 13.1 |  |
| sp\|Q13526\|PIN1\_HUMAN | Q13526 | PIN1\_HUMAN | PIN1 | Peptidyl-prolyl cis-trans isomerase NIMA-interacting 1 | 163 | 18.243 | 2.824452e+05 | 6.473626e+06 | 706214.3 | 5669481.5 | 97930.36 |  | 46.63 | 89.6 | 35.6 |  |
| sp\|O00159\|MYO1C\_HUMAN | O00159 | MYO1C\_HUMAN | MYO1C | Unconventional myosin-Ic | 1063 | 121.682 | 1.453405e+04 | 2.221929e+06 | 702027.25 | 1410804.6 | 109096.68 |  | 37.44 | 43.6 | 15.4 |  |
| sp\|Q14653\|IRF3\_HUMAN | Q14653 | IRF3\_HUMAN | IRF3 | Interferon regulatory factor 3 | 427 | 47.219 | 1.911302e+04 | 1.133870e+06 | 699021.7 | 434848.0 |  |  | 30.21 | 27.4 |  |  |
| sp\|P01023\|A2MG\_HUMAN | P01023 | A2MG\_HUMAN | A2M | Alpha-2-macroglobulin | 1474 | 163.291 | 9.871259e+03 | 2.025126e+06 | 696694.4 | 727382.25 | 351314.56 | 249734.4 | 5.77 | 25.6 | 28.1 | 10.9 |
| sp\|Q9NQ48\|LZTL1\_HUMAN | Q9NQ48 | LZTL1\_HUMAN | LZTFL1 | Leucine zipper transcription factor-like protein 1 | 299 | 34.592 | 2.433928e+04 | 1.057793e+06 | 696531.1 | 361261.75 |  |  | 33.78 | 40.1 |  |  |
| sp\|P01834\|IGKC\_HUMAN | P01834 | IGKC\_HUMAN | IGKC | Immunoglobulin kappa constant | 107 | 11.765 | 3.821865e+05 | 5.649167e+06 | 696475.8 | 2173156.5 | 2479357.2 | 300177.8 | 48.6 | 72.0 | 55.1 | 18.7 |
| sp\|P62805\|H4\_HUMAN | P62805 | H4\_HUMAN | H4C1 | Histone H4 | 103 | 11.367 | 2.204962e+05 | 3.148939e+06 | 695449.0 | 220540.2 | 1754586.6 | 478362.9 | 50.49 | 66.0 | 64.1 | 60.2 |
| sp\|P0DTE7\|AMY1B\_HUMAN | P0DTE7 | AMY1B\_HUMAN | AMY1B | Alpha-amylase 1B | 511 | 57.768 | 2.820215e+04 | 2.046853e+06 | 694833.9 |  | 1352019.5 |  | 21.14 |  | 29.9 |  |
| sp\|Q9H2H8\|PPIL3\_HUMAN | Q9H2H8 | PPIL3\_HUMAN | PPIL3 | Peptidyl-prolyl cis-trans isomerase-like 3 | 161 | 18.155 | 2.555698e+05 | 5.829387e+06 | 693735.44 | 4848968.0 | 230086.94 | 56596.203 | 62.11 | 72.0 | 49.1 | 29.2 |
| sp\|Q9NZM3\|ITSN2\_HUMAN | Q9NZM3 | ITSN2\_HUMAN | ITSN2 | Intersectin-2 | 1697 | 193.461 | 1.337526e+04 | 3.250969e+06 | 689968.44 | 155707.31 | 2308879.5 | 96413.94 | 9.37 | 27.6 | 24.1 | 5.1 |
| sp\|Q92841\|DDX17\_HUMAN | Q92841 | DDX17\_HUMAN | DDX17 | Probable ATP-dependent RNA helicase DDX17 | 729 | 80.272 | 2.192335e+04 | 2.210997e+06 | 688967.6 | 849347.9 | 524450.2 | 148231.4 | 38.41 | 55.3 | 37.9 | 22.4 |
| sp\|Q9Y2H1\|ST38L\_HUMAN | Q9Y2H1 | ST38L\_HUMAN | STK38L | Serine/threonine-protein kinase 38-like | 464 | 54.003 | 2.726215e+04 | 1.849674e+06 | 688633.4 | 1081782.8 | 79257.305 |  | 35.13 | 61.4 | 25.0 |  |
| sp\|Q7Z2W4\|ZCCHV\_HUMAN | Q7Z2W4 | ZCCHV\_HUMAN | ZC3HAV1 | Zinc finger CCCH-type antiviral protein 1 | 902 | 101.431 | 4.424470e+04 | 5.638312e+06 | 685065.9 | 3407514.0 | 905217.5 | 640514.2 | 40.69 | 67.6 | 30.0 | 17.4 |
| sp\|Q92835\|SHIP1\_HUMAN | Q92835 | SHIP1\_HUMAN | INPP5D | Phosphatidylinositol 3,4,5-trisphosphate 5-phosphatase 1 | 1189 | 133.292 | 2.248275e+04 | 3.765049e+06 | 682289.25 | 1949706.9 | 860631.75 | 272421.5 | 45.92 | 56.9 | 30.1 | 22.0 |
| sp\|Q99961\|SH3G1\_HUMAN | Q99961 | SH3G1\_HUMAN | SH3GL1 | Endophilin-A2 | 368 | 41.490 | 2.988194e+04 | 1.557648e+06 | 681662.25 | 632279.06 | 169337.78 | 74369.19 | 35.05 | 51.1 | 31.0 | 15.5 |
| sp\|Q8N122\|RPTOR\_HUMAN | Q8N122 | RPTOR\_HUMAN | RPTOR | Regulatory-associated protein of mTOR | 1335 | 149.038 | 9.020470e+03 | 1.689053e+06 | 680970.06 | 909994.44 |  | 98088.766 | 32.13 | 41.8 |  | 10.8 |
| sp\|P41226\|UBA7\_HUMAN | P41226 | UBA7\_HUMAN | UBA7 | Ubiquitin-like modifier-activating enzyme 7 | 1012 | 111.694 | 1.048725e+04 | 1.471664e+06 | 677357.56 | 794306.56 |  |  | 24.01 | 36.6 |  |  |
| sp\|Q99426\|TBCB\_HUMAN | Q99426 | TBCB\_HUMAN | TBCB | Tubulin-folding cofactor B | 244 | 27.326 | 1.041532e+05 | 3.575739e+06 | 677106.4 | 2471420.2 | 320508.2 | 106704.67 | 43.03 | 81.1 | 34.8 | 13.9 |
| sp\|Q14232\|EI2BA\_HUMAN | Q14232 | EI2BA\_HUMAN | EIF2B1 | Translation initiation factor eIF-2B subunit alpha | 305 | 33.712 | 7.366259e+04 | 3.119958e+06 | 677057.1 | 2293930.8 | 148969.6 |  | 44.92 | 55.1 | 30.8 |  |
| sp\|Q4G0N4\|NAKD2\_HUMAN | Q4G0N4 | NAKD2\_HUMAN | NADK2 | NAD kinase 2, mitochondrial | 442 | 49.433 | 2.621495e+04 | 1.628108e+06 | 671773.25 | 824501.2 | 131833.36 |  | 45.02 | 36.0 | 25.6 |  |
| sp\|Q9BX66\|SRBS1\_HUMAN | Q9BX66 | SRBS1\_HUMAN | SORBS1 | Sorbin and SH3 domain-containing protein 1 | 1292 | 142.513 | 6.793258e+03 | 1.216325e+06 | 671313.4 | 256619.27 | 181103.05 | 107289.64 | 6.42 | 43.3 | 22.0 | 7.3 |
| sp\|Q9C075\|K1C23\_HUMAN | Q9C075 | K1C23\_HUMAN | KRT23 | Keratin, type I cytoskeletal 23 | 422 | 48.131 | 7.697715e+04 | 4.654830e+06 | 669093.75 | 173419.16 | 275691.56 | 3536626.0 | 19.67 | 55.0 | 37.2 | 30.6 |
| sp\|Q9NR50\|EI2BG\_HUMAN | Q9NR50 | EI2BG\_HUMAN | EIF2B3 | Translation initiation factor eIF-2B subunit gamma | 452 | 50.240 | 2.031893e+04 | 1.282530e+06 | 668506.5 | 586706.8 | 27316.941 |  | 35.84 | 56.2 | 19.9 |  |
| sp\|Q8N9N7\|LRC57\_HUMAN | Q8N9N7 | LRC57\_HUMAN | LRRC57 | Leucine-rich repeat-containing protein 57 | 239 | 26.754 | 8.636260e+04 | 2.902897e+06 | 668469.3 | 1318262.8 | 497601.34 | 418563.2 | 51.88 | 43.5 | 21.3 | 24.3 |
| sp\|P12882\|MYH1\_HUMAN | P12882 | MYH1\_HUMAN | MYH1 | Myosin-1 | 1939 | 223.145 | 1.889218e+04 | 5.296468e+06 | 667129.6 | 1561164.8 | 2253154.5 | 815019.6 | 16.81 | 50.5 | 39.6 | 20.9 |
| sp\|Q96TA1\|NIBA2\_HUMAN | Q96TA1 | NIBA2\_HUMAN | NIBAN2 | Protein Niban 2 | 746 | 84.138 | 3.693891e+04 | 3.904752e+06 | 666521.06 | 1454484.6 | 140538.16 | 1643208.2 | 45.44 | 39.3 | 17.3 | 17.2 |
| sp\|P51991\|ROA3\_HUMAN | P51991 | ROA3\_HUMAN | HNRNPA3 | Heterogeneous nuclear ribonucleoprotein A3 | 378 | 39.595 | 5.588378e+04 | 2.779990e+06 | 666498.2 | 1570714.1 | 349008.78 | 193769.08 | 35.19 | 59.3 | 39.4 | 23.0 |
| sp\|Q9BU02\|THTPA\_HUMAN | Q9BU02 | THTPA\_HUMAN | THTPA | Thiamine-triphosphatase | 230 | 25.566 | 3.367991e+04 | 1.081810e+06 | 663390.75 | 278873.94 | 139545.0 |  | 18.26 | 49.6 | 49.1 |  |
| sp\|Q9Y623\|MYH4\_HUMAN | Q9Y623 | MYH4\_HUMAN | MYH4 | Myosin-4 | 1939 | 223.071 | 2.166115e+04 | 6.070741e+06 | 661407.7 | 1599662.4 | 2994651.2 | 815019.6 | 15.42 | 53.7 | 36.2 | 22.6 |
| sp\|P46736\|BRCC3\_HUMAN | P46736 | BRCC3\_HUMAN | BRCC3 | Lys-63-specific deubiquitinase BRCC36 | 316 | 36.072 | 5.362948e+04 | 2.430474e+06 | 660827.0 | 1699932.2 | 69714.48 |  | 48.1 | 61.1 | 25.6 |  |
| sp\|Q8TCA0\|LRC20\_HUMAN | Q8TCA0 | LRC20\_HUMAN | LRRC20 | Leucine-rich repeat-containing protein 20 | 184 | 20.509 | 4.980485e+04 | 1.283315e+06 | 660461.4 | 390457.2 | 191168.22 | 41228.234 | 61.96 | 60.9 | 25.5 | 14.7 |
| sp\|P29508\|SPB3\_HUMAN | P29508 | SPB3\_HUMAN | SERPINB3 | Serpin B3 | 390 | 44.565 | 3.921032e+04 | 2.195389e+06 | 659983.94 | 294423.22 | 960101.4 | 280880.3 | 31.28 | 36.4 | 41.0 | 17.4 |
| sp\|P02461\|CO3A1\_HUMAN | P02461 | CO3A1\_HUMAN | COL3A1 | Collagen alpha-1(III) chain | 1466 | 138.564 | 7.601372e+03 | 1.323304e+06 | 657675.25 | 276401.2 |  | 389227.38 | 12.89 | 67.3 |  | 30.5 |
| sp\|Q9UBV8\|PEF1\_HUMAN | Q9UBV8 | PEF1\_HUMAN | PEF1 | Peflin | 284 | 30.381 | 2.826673e+05 | 1.078934e+07 | 657620.1 | 2045506.8 | 1473026.8 | 6613185.0 | 26.41 | 26.4 | 16.2 | 8.8 |
| sp\|Q8IW45\|NNRD\_HUMAN | Q8IW45 | NNRD\_HUMAN | NAXD | ATP-dependent (S)-NAD(P)H-hydrate dehydratase | 347 | 36.576 | 3.863882e+04 | 1.775568e+06 | 656467.94 | 996165.9 | 99613.15 | 23320.803 | 51.3 | 58.8 | 16.7 | 8.6 |
| sp\|P31323\|KAP3\_HUMAN | P31323 | KAP3\_HUMAN | PRKAR2B | cAMP-dependent protein kinase type II-beta regulatory subunit | 418 | 46.302 | 2.118316e+04 | 1.232275e+06 | 655787.44 | 486793.47 | 89694.03 |  | 46.41 | 42.6 | 27.5 |  |
| sp\|Q9UKK9\|NUDT5\_HUMAN | Q9UKK9 | NUDT5\_HUMAN | NUDT5 | ADP-sugar pyrophosphatase | 219 | 24.328 | 3.190853e+05 | 9.752823e+06 | 653098.0 | 8257454.5 | 608947.94 | 233322.98 | 41.55 | 68.0 | 31.1 | 27.4 |
| sp\|P53985\|MOT1\_HUMAN | P53985 | MOT1\_HUMAN | SLC16A1 | Monocarboxylate transporter 1 | 500 | 53.944 | 1.992389e+05 | 1.350313e+07 | 651516.75 | 1101570.1 | 3203615.8 | 8546426.0 | 9.4 | 18.2 | 20.8 | 14.0 |
| sp\|Q00613\|HSF1\_HUMAN | Q00613 | HSF1\_HUMAN | HSF1 | Heat shock factor protein 1 | 529 | 58.190 | 8.899680e+03 | 6.506388e+05 | 650638.75 |  |  |  | 10.4 |  |  |  |
| sp\|P35219\|CAH8\_HUMAN | P35219 | CAH8\_HUMAN | CA8 | Carbonic anhydrase-related protein | 290 | 32.973 | 3.799836e+04 | 1.574130e+06 | 650410.5 | 923719.0 |  |  | 34.14 | 41.0 |  |  |
| sp\|Q99615\|DNJC7\_HUMAN | Q99615 | DNJC7\_HUMAN | DNAJC7 | DnaJ homolog subfamily C member 7 | 494 | 56.441 | 1.496393e+04 | 1.061103e+06 | 645820.1 | 408890.3 | 6392.269 |  | 9.11 | 51.6 | 22.7 |  |
| sp\|Q9Y4X5\|ARI1\_HUMAN | Q9Y4X5 | ARI1\_HUMAN | ARIH1 | E3 ubiquitin-protein ligase ARIH1 | 557 | 64.118 | 1.713902e+04 | 1.380648e+06 | 644859.5 | 564065.56 | 132064.25 | 39659.055 | 17.41 | 17.6 | 19.2 | 4.5 |
| sp\|Q15438\|CYH1\_HUMAN | Q15438 | CYH1\_HUMAN | CYTH1 | Cytohesin-1 | 398 | 46.413 | 2.071589e+04 | 1.207982e+06 | 644531.1 | 466497.12 | 96953.67 |  | 32.66 | 40.2 | 23.4 |  |
| sp\|Q6P9A3\|ZN549\_HUMAN | Q6P9A3 | ZN549\_HUMAN | ZNF549 | Zinc finger protein 549 | 640 | 74.439 | 3.520598e+04 | 3.292563e+06 | 642732.1 | 1013881.3 | 821436.75 | 814513.2 | 3.44 | 22.2 | 15.8 | 6.1 |
| sp\|Q13144\|EI2BE\_HUMAN | Q13144 | EI2BE\_HUMAN | EIF2B5 | Translation initiation factor eIF-2B subunit epsilon | 721 | 80.380 | 1.254411e+04 | 1.266791e+06 | 642184.6 | 550258.4 | 74348.05 |  | 29.4 | 35.9 | 14.8 |  |
| sp\|Q8N8A2\|ANR44\_HUMAN | Q8N8A2 | ANR44\_HUMAN | ANKRD44 | Serine/threonine-protein phosphatase 6 regulatory ankyrin repeat subunit B | 993 | 107.603 | 1.232806e+04 | 1.666618e+06 | 640650.4 | 959832.3 | 66135.76 |  | 14.1 | 36.6 | 12.6 |  |
| sp\|O75340\|PDCD6\_HUMAN | O75340 | PDCD6\_HUMAN | PDCD6 | Programmed cell death protein 6 | 191 | 21.868 | 2.461762e+05 | 6.763512e+06 | 635543.75 | 3304794.0 | 2468378.8 | 354795.72 | 42.41 | 76.4 | 45.0 | 28.3 |
| sp\|P10155\|RO60\_HUMAN | P10155 | RO60\_HUMAN | RO60 | RNA-binding protein RO60 | 538 | 60.671 | 2.437478e+04 | 1.857972e+06 | 634095.7 | 1168574.8 | 55301.21 |  | 29.74 | 41.3 | 14.5 |  |
| sp\|Q9NWZ3\|IRAK4\_HUMAN | Q9NWZ3 | IRAK4\_HUMAN | IRAK4 | Interleukin-1 receptor-associated kinase 4 | 460 | 51.530 | 2.340837e+04 | 1.515474e+06 | 633222.0 | 669808.0 | 212443.8 |  | 34.57 | 49.6 | 43.0 |  |
| sp\|Q8NBS9\|TXND5\_HUMAN | Q8NBS9 | TXND5\_HUMAN | TXNDC5 | Thioredoxin domain-containing protein 5 | 432 | 47.629 | 5.958373e+04 | 3.565466e+06 | 633220.44 | 699617.8 | 1544476.0 | 688151.5 | 21.76 | 37.7 | 35.4 | 19.9 |
| sp\|P16455\|MGMT\_HUMAN | P16455 | MGMT\_HUMAN | MGMT | Methylated-DNA--protein-cysteine methyltransferase | 207 | 22.770 | 2.209861e+04 | 6.321864e+05 | 632186.44 |  |  |  | 13.04 |  |  |  |
| sp\|P35573\|GDE\_HUMAN | P35573 | GDE\_HUMAN | AGL | Glycogen debranching enzyme | 1532 | 174.764 | 6.876468e+03 | 1.509853e+06 | 629532.56 | 795628.3 | 54444.562 | 30247.273 | 37.14 | 36.0 | 10.7 | 7.4 |
| sp\|Q08188\|TGM3\_HUMAN | Q08188 | TGM3\_HUMAN | TGM3 | Protein-glutamine gamma-glutamyltransferase E | 693 | 76.632 | 5.986989e+04 | 5.764156e+06 | 626696.8 | 779925.9 | 2382925.5 | 1974608.1 | 34.78 | 42.3 | 35.6 | 23.4 |
| sp\|Q13393\|PLD1\_HUMAN | Q13393 | PLD1\_HUMAN | PLD1 | Phospholipase D1 | 1074 | 124.184 | 6.984397e+03 | 1.089712e+06 | 626632.44 | 104273.88 | 117935.25 | 240870.31 | 16.85 | 33.1 | 17.7 | 7.7 |
| sp\|Q9Y673\|ALG5\_HUMAN | Q9Y673 | ALG5\_HUMAN | ALG5 | Dolichyl-phosphate beta-glucosyltransferase | 324 | 36.946 | 1.974570e+04 | 9.165522e+05 | 626364.25 | 99785.55 | 65624.27 | 124778.086 | 21.91 | 35.5 | 21.9 | 16.0 |
| sp\|O43427\|FIBP\_HUMAN | O43427 | FIBP\_HUMAN | FIBP | Acidic fibroblast growth factor intracellular-binding protein | 364 | 41.878 | 1.933708e+04 | 1.017405e+06 | 625737.5 | 321980.97 | 69686.54 |  | 39.84 | 35.7 | 27.7 |  |
| sp\|Q5EBM0\|CMPK2\_HUMAN | Q5EBM0 | CMPK2\_HUMAN | CMPK2 | UMP-CMP kinase 2, mitochondrial | 449 | 49.448 | 2.607200e+04 | 1.619721e+06 | 623943.2 | 884541.5 | 73726.63 | 37509.43 | 37.19 | 50.8 | 23.8 | 11.8 |
| sp\|Q8WU79\|SMAP2\_HUMAN | Q8WU79 | SMAP2\_HUMAN | SMAP2 | Stromal membrane-associated protein 2 | 429 | 46.786 | 1.585682e+04 | 9.320715e+05 | 622045.0 | 189989.55 | 120036.92 |  | 18.88 | 30.1 | 12.1 |  |
| sp\|Q8IZ07\|AN13A\_HUMAN | Q8IZ07 | AN13A\_HUMAN | ANKRD13A | Ankyrin repeat domain-containing protein 13A | 590 | 67.619 | 4.843274e+04 | 4.114575e+06 | 620607.8 | 2897835.0 | 560264.06 | 35867.94 | 49.15 | 55.6 | 19.8 | 8.8 |
| sp\|Q01415\|GALK2\_HUMAN | Q01415 | GALK2\_HUMAN | GALK2 | N-acetylgalactosamine kinase | 458 | 50.378 | 3.020653e+04 | 1.911873e+06 | 620430.1 | 1291442.6 |  |  | 22.27 | 39.1 |  |  |
| sp\|Q93088\|BHMT1\_HUMAN | Q93088 | BHMT1\_HUMAN | BHMT | Betaine--homocysteine S-methyltransferase 1 | 406 | 44.998 | 1.242405e+04 | 7.023824e+05 | 618621.2 |  | 83761.25 |  | 8.37 |  | 22.2 |  |
| sp\|O60271\|JIP4\_HUMAN | O60271 | JIP4\_HUMAN | SPAG9 | C-Jun-amino-terminal kinase-interacting protein 4 | 1321 | 146.205 | 2.573648e+04 | 4.727467e+06 | 618406.6 | 3918563.0 | 116111.02 | 74386.61 | 42.54 | 45.6 | 15.5 | 16.3 |
| sp\|Q9Y4L1\|HYOU1\_HUMAN | Q9Y4L1 | HYOU1\_HUMAN | HYOU1 | Hypoxia up-regulated protein 1 | 999 | 111.335 | 3.443682e+04 | 4.816948e+06 | 615879.7 | 400399.53 | 3459782.0 | 340886.5 | 34.63 | 45.9 | 43.3 | 21.0 |
| sp\|Q969U7\|PSMG2\_HUMAN | Q969U7 | PSMG2\_HUMAN | PSMG2 | Proteasome assembly chaperone 2 | 264 | 29.396 | 8.902564e+04 | 3.287914e+06 | 613893.94 | 2513426.5 | 160593.95 |  | 21.97 | 37.9 | 22.7 |  |
| sp\|Q9NPH2\|INO1\_HUMAN | Q9NPH2 | INO1\_HUMAN | ISYNA1 | Inositol-3-phosphate synthase 1 | 558 | 61.068 | 1.359110e+04 | 1.042763e+06 | 613841.56 | 428921.1 |  |  | 21.68 | 38.5 |  |  |
| sp\|Q9NVA2\|SEP11\_HUMAN | Q9NVA2 | SEP11\_HUMAN | SEPTIN11 | Septin-11 | 429 | 49.398 | 3.717161e+05 | 2.306948e+07 | 613265.6 | 780713.75 | 21675504.0 |  | 32.63 | 38.5 | 26.3 |  |
| sp\|P50452\|SPB8\_HUMAN | P50452 | SPB8\_HUMAN | SERPINB8 | Serpin B8 | 374 | 42.767 | 2.846012e+04 | 1.529194e+06 | 612866.44 | 530478.56 | 195679.77 | 190169.52 | 12.3 | 33.2 | 34.0 | 26.2 |
| sp\|P31942\|HNRH3\_HUMAN | P31942 | HNRH3\_HUMAN | HNRNPH3 | Heterogeneous nuclear ribonucleoprotein H3 | 346 | 36.926 | 5.929260e+04 | 2.750743e+06 | 612529.4 | 1940536.9 | 164902.75 | 32773.52 | 32.66 | 50.0 | 36.7 | 17.9 |
| sp\|Q15042\|RB3GP\_HUMAN | Q15042 | RB3GP\_HUMAN | RAB3GAP1 | Rab3 GTPase-activating protein catalytic subunit | 981 | 110.524 | 2.402648e+04 | 3.336290e+06 | 612309.94 | 1556052.9 | 913610.06 | 254317.39 | 42.3 | 51.7 | 37.1 | 21.6 |
| sp\|Q9BTV6\|DPH7\_HUMAN | Q9BTV6 | DPH7\_HUMAN | DPH7 | Diphthine methyltransferase | 452 | 50.575 | 1.397576e+04 | 8.880318e+05 | 610321.2 | 277710.6 |  |  | 12.17 | 32.3 |  |  |
| sp\|P35052\|GPC1\_HUMAN | P35052 | GPC1\_HUMAN | GPC1 | Glypican-1 | 558 | 61.380 | 7.907141e+03 | 6.097664e+05 | 609766.44 |  |  |  | 5.56 |  |  |  |
| sp\|Q13148\|TADBP\_HUMAN | Q13148 | TADBP\_HUMAN | TARDBP | TAR DNA-binding protein 43 | 414 | 44.740 | 4.128274e+04 | 2.320500e+06 | 609171.8 | 1372613.2 | 159994.4 | 178720.81 | 25.6 | 29.7 | 12.3 | 10.6 |
| sp\|Q9BXB4\|OSB11\_HUMAN | Q9BXB4 | OSB11\_HUMAN | OSBPL11 | Oxysterol-binding protein-related protein 11 | 747 | 83.643 | 7.632479e+03 | 8.020702e+05 | 608850.3 | 193219.9 |  |  | 8.7 | 34.1 |  |  |
| sp\|Q99755\|PI51A\_HUMAN | Q99755 | PI51A\_HUMAN | PIP5K1A | Phosphatidylinositol 4-phosphate 5-kinase type-1 alpha | 562 | 62.633 | 2.012809e+04 | 1.583883e+06 | 608665.94 | 777434.4 | 197782.45 |  | 24.73 | 40.2 | 40.7 |  |
| sp\|P46940\|IQGA1\_HUMAN | P46940 | IQGA1\_HUMAN | IQGAP1 | Ras GTPase-activating-like protein IQGAP1 | 1657 | 189.252 | 6.741952e+03 | 1.603036e+06 | 608383.94 | 780732.8 | 116279.67 | 97639.69 | 39.11 | 35.9 | 22.5 | 10.3 |
| sp\|Q96JH7\|VCIP1\_HUMAN | Q96JH7 | VCIP1\_HUMAN | VCPIP1 | Deubiquitinating protein VCPIP1 | 1222 | 134.321 | 1.505558e+04 | 2.540730e+06 | 607988.2 | 1543518.0 | 281098.3 | 108125.62 | 36.99 | 52.5 | 29.3 | 21.6 |
| sp\|O75191\|XYLB\_HUMAN | O75191 | XYLB\_HUMAN | XYLB | Xylulose kinase | 536 | 58.382 | 1.715130e+04 | 1.258036e+06 | 607461.2 | 650575.0 |  |  | 24.63 | 51.9 |  |  |
| sp\|Q12906\|ILF3\_HUMAN | Q12906 | ILF3\_HUMAN | ILF3 | Interleukin enhancer-binding factor 3 | 894 | 95.338 | 1.609902e+04 | 1.928336e+06 | 605074.0 | 1151394.5 | 171867.2 |  | 33.67 | 47.2 | 27.7 |  |
| sp\|Q9HCE7\|SMUF1\_HUMAN | Q9HCE7 | SMUF1\_HUMAN | SMURF1 | E3 ubiquitin-protein ligase SMURF1 | 757 | 83.270 | 5.779256e+03 | 6.046132e+05 | 604613.25 |  |  |  | 3.43 |  |  |  |
| sp\|Q9H814\|PHAX\_HUMAN | Q9H814 | PHAX\_HUMAN | PHAX | Phosphorylated adapter RNA export protein | 394 | 44.403 | 2.472790e+04 | 1.379484e+06 | 603464.6 | 776019.44 |  |  | 32.99 | 39.3 |  |  |
| sp\|Q12905\|ILF2\_HUMAN | Q12905 | ILF2\_HUMAN | ILF2 | Interleukin enhancer-binding factor 2 | 390 | 43.062 | 2.145485e+04 | 1.160745e+06 | 602538.06 | 489332.22 | 68874.86 |  | 32.31 | 46.7 | 19.2 |  |
| sp\|P07099\|HYEP\_HUMAN | P07099 | HYEP\_HUMAN | EPHX1 | Epoxide hydrolase 1 | 455 | 52.949 | 3.316157e+04 | 2.206023e+06 | 601819.0 | 732610.9 | 431429.5 | 440163.2 | 23.74 | 31.9 | 23.1 | 21.1 |
| sp\|P36871\|PGM1\_HUMAN | P36871 | PGM1\_HUMAN | PGM1 | Phosphoglucomutase-1 | 562 | 61.449 | 5.333205e+04 | 4.117374e+06 | 601075.6 | 1916919.6 | 89530.61 | 1509847.9 | 50.18 | 50.2 | 23.1 | 16.2 |
| sp\|Q9NQX3\|GEPH\_HUMAN | Q9NQX3 | GEPH\_HUMAN | GPHN | Gephyrin | 736 | 79.748 | 3.701936e+04 | 3.709077e+06 | 600890.3 | 2766775.2 | 230091.95 | 111319.73 | 48.37 | 76.9 | 36.8 | 8.4 |
| sp\|Q9UL15\|BAG5\_HUMAN | Q9UL15 | BAG5\_HUMAN | BAG5 | BAG family molecular chaperone regulator 5 | 447 | 51.200 | 2.813595e+04 | 1.809875e+06 | 599412.25 | 1086292.6 | 124170.42 |  | 41.83 | 52.8 | 15.4 |  |
| sp\|Q9BXJ9\|NAA15\_HUMAN | Q9BXJ9 | NAA15\_HUMAN | NAA15 | N-alpha-acetyltransferase 15, NatA auxiliary subunit | 866 | 101.272 | 1.551406e+04 | 1.973931e+06 | 596141.9 | 1200930.8 | 142261.38 | 34596.53 | 42.15 | 42.8 | 23.8 | 16.1 |
| sp\|Q14980\|NUMA1\_HUMAN | Q14980 | NUMA1\_HUMAN | NUMA1 | Nuclear mitotic apparatus protein 1 | 2115 | 238.260 | 8.839971e+03 | 2.646178e+06 | 595776.5 | 585757.5 | 1349191.9 | 115452.43 | 20.95 | 51.1 | 33.4 | 18.2 |
| sp\|P62942\|FKB1A\_HUMAN | P62942 | FKB1A\_HUMAN | FKBP1A | Peptidyl-prolyl cis-trans isomerase FKBP1A | 108 | 11.951 | 5.164275e+05 | 7.754089e+06 | 594115.8 | 4829126.0 | 1019666.6 | 1311180.8 | 41.67 | 61.1 | 38.9 | 13.0 |
| sp\|Q8TD19\|NEK9\_HUMAN | Q8TD19 | NEK9\_HUMAN | NEK9 | Serine/threonine-protein kinase Nek9 | 979 | 107.168 | 7.281229e+03 | 9.803631e+05 | 594089.5 | 309549.97 | 68272.555 | 8451.092 | 13.38 | 30.7 | 17.7 | 5.6 |
| sp\|Q9NVG8\|TBC13\_HUMAN | Q9NVG8 | TBC13\_HUMAN | TBC1D13 | TBC1 domain family member 13 | 400 | 46.554 | 4.157543e+04 | 2.431705e+06 | 593859.06 | 1837846.2 |  |  | 43.0 | 48.5 |  |  |
| sp\|Q15291\|RBBP5\_HUMAN | Q15291 | RBBP5\_HUMAN | RBBP5 | Retinoblastoma-binding protein 5 | 538 | 59.153 | 8.912968e+03 | 6.623938e+05 | 593850.4 | 68543.44 |  |  | 7.25 | 27.3 |  |  |
| sp\|P04908\|H2A1B\_HUMAN | P04908 | H2A1B\_HUMAN | H2AC4 | Histone H2A type 1-B/E | 130 | 14.135 | 7.396308e+04 | 1.313494e+06 | 593241.9 |  |  | 720251.6 | 35.38 |  |  | 0.0 |
| sp\|P16104\|H2AX\_HUMAN | P16104 | H2AX\_HUMAN | H2AX | Histone H2AX | 143 | 15.145 | 1.665552e+05 | 3.169163e+06 | 593241.9 | 273274.4 | 1665131.9 | 637514.4 | 27.27 | 69.9 | 59.4 | 47.6 |
| sp\|P23284\|PPIB\_HUMAN | P23284 | PPIB\_HUMAN | PPIB | Peptidyl-prolyl cis-trans isomerase B | 216 | 23.743 | 1.864240e+05 | 5.561020e+06 | 592861.9 | 1958736.2 | 2255137.5 | 754284.75 | 43.98 | 72.7 | 48.6 | 28.2 |
| sp\|Q9NWX6\|THG1\_HUMAN | Q9NWX6 | THG1\_HUMAN | THG1L | Probable tRNA(His) guanylyltransferase | 298 | 34.831 | 5.401386e+04 | 2.363678e+06 | 590258.25 | 947858.25 | 654500.44 | 171061.02 | 46.98 | 49.0 | 31.9 | 16.4 |
| sp\|Q8TF66\|LRC15\_HUMAN | Q8TF66 | LRC15\_HUMAN | LRRC15 | Leucine-rich repeat-containing protein 15 | 581 | 64.366 | 9.281440e+03 | 7.505663e+05 | 588877.7 |  | 9277.2295 | 152411.34 | 27.19 |  | 7.1 | 6.2 |
| sp\|Q96CN7\|ISOC1\_HUMAN | Q96CN7 | ISOC1\_HUMAN | ISOC1 | Isochorismatase domain-containing protein 1 | 298 | 32.237 | 5.797637e+04 | 2.348134e+06 | 588284.7 | 1608730.5 | 151118.31 |  | 40.27 | 44.3 | 16.8 |  |
| sp\|O43747\|AP1G1\_HUMAN | O43747 | AP1G1\_HUMAN | AP1G1 | AP-1 complex subunit gamma-1 | 822 | 91.351 | 1.211029e+04 | 1.389905e+06 | 588234.56 | 569045.25 | 232624.86 |  | 26.03 | 42.5 | 18.0 |  |
| sp\|Q9UHP3\|UBP25\_HUMAN | Q9UHP3 | UBP25\_HUMAN | USP25 | Ubiquitin carboxyl-terminal hydrolase 25 | 1055 | 122.218 | 1.006879e+04 | 1.546072e+06 | 587018.25 | 799430.3 | 116553.15 | 43070.273 | 40.57 | 39.0 | 21.9 | 8.3 |
| sp\|P02452\|CO1A1\_HUMAN | P02452 | CO1A1\_HUMAN | COL1A1 | Collagen alpha-1(I) chain | 1464 | 138.911 | 2.497801e+04 | 4.359248e+06 | 583907.44 | 1310947.6 | 554697.5 | 1909695.4 | 24.18 | 68.6 | 50.3 | 46.9 |
| sp\|Q6FI81\|CPIN1\_HUMAN | Q6FI81 | CPIN1\_HUMAN | CIAPIN1 | Anamorsin | 312 | 33.582 | 8.735785e+04 | 3.685748e+06 | 583564.94 | 2136256.0 | 375028.72 | 590898.6 | 29.81 | 37.5 | 41.3 | 39.1 |
| sp\|Q9Y696\|CLIC4\_HUMAN | Q9Y696 | CLIC4\_HUMAN | CLIC4 | Chloride intracellular channel protein 4 | 253 | 28.772 | 1.113305e+05 | 4.024402e+06 | 583027.6 | 2894658.2 | 546716.1 |  | 37.15 | 83.0 | 40.7 |  |
| sp\|Q15019\|SEPT2\_HUMAN | Q15019 | SEPT2\_HUMAN | SEPTIN2 | Septin-2 | 361 | 41.487 | 2.376333e+04 | 1.238616e+06 | 582902.8 | 570637.25 | 85075.53 |  | 53.46 | 60.9 | 31.9 |  |
| sp\|P01876\|IGHA1\_HUMAN | P01876 | IGHA1\_HUMAN | IGHA1 | Immunoglobulin heavy constant alpha 1 | 398 | 42.849 | 6.979009e+04 | 3.757090e+06 | 580470.7 |  | 1881348.9 | 1295270.2 | 16.83 |  | 28.3 | 19.5 |
| sp\|Q5VZ03\|NXNL2\_HUMAN | Q5VZ03 | NXNL2\_HUMAN | NXNL2 | Nucleoredoxin-like protein 2 | 156 | 17.614 | 2.026209e+05 | 4.483935e+06 | 580437.2 |  | 3408720.5 | 494777.25 | 8.33 |  | 12.8 | 8.3 |
| sp\|P06737\|PYGL\_HUMAN | P06737 | PYGL\_HUMAN | PYGL | Glycogen phosphorylase, liver form | 847 | 97.149 | 1.305000e+04 | 1.592817e+06 | 580381.94 | 1012435.1 |  |  | 29.63 | 51.9 |  |  |
| sp\|P11216\|PYGB\_HUMAN | P11216 | PYGB\_HUMAN | PYGB | Glycogen phosphorylase, brain form | 843 | 96.696 | 1.073355e+04 | 1.303974e+06 | 580381.94 | 624078.7 | 99513.39 |  | 36.89 | 57.8 | 27.5 |  |
| sp\|P11217\|PYGM\_HUMAN | P11217 | PYGM\_HUMAN | PYGM | Glycogen phosphorylase, muscle form | 842 | 97.092 | 1.273076e+04 | 1.552941e+06 | 580381.94 | 208921.94 | 317196.44 | 446440.94 | 12.95 | 33.8 | 19.1 | 12.8 |
| sp\|Q9UJY5\|GGA1\_HUMAN | Q9UJY5 | GGA1\_HUMAN | GGA1 | ADP-ribosylation factor-binding protein GGA1 | 639 | 70.384 | 1.404495e+04 | 1.241970e+06 | 579168.06 | 403527.0 | 259275.38 |  | 21.91 | 32.6 | 25.0 |  |
| sp\|Q96GK7\|FAH2A\_HUMAN | Q96GK7 | FAH2A\_HUMAN | FAHD2A | Fumarylacetoacetate hydrolase domain-containing protein 2A | 314 | 34.596 | 2.235969e+04 | 9.718713e+05 | 578969.0 | 392902.28 |  |  | 48.41 | 37.6 |  |  |
| sp\|Q8IVD9\|NUDC3\_HUMAN | Q8IVD9 | NUDC3\_HUMAN | NUDCD3 | NudC domain-containing protein 3 | 361 | 40.822 | 2.146214e+04 | 1.100739e+06 | 578207.1 | 489840.16 | 32691.748 |  | 46.54 | 65.4 | 20.2 |  |
| sp\|Q9Y6E0\|STK24\_HUMAN | Q9Y6E0 | STK24\_HUMAN | STK24 | Serine/threonine-protein kinase 24 | 443 | 49.308 | 3.986341e+04 | 2.469500e+06 | 577672.6 | 1500988.8 | 291548.8 | 99289.8 | 40.86 | 46.0 | 36.3 | 7.7 |
| sp\|Q8IYU2\|HACE1\_HUMAN | Q8IYU2 | HACE1\_HUMAN | HACE1 | E3 ubiquitin-protein ligase HACE1 | 909 | 102.342 | 9.328617e+03 | 1.199467e+06 | 577492.56 | 559721.4 | 62253.074 |  | 27.06 | 27.2 | 17.5 |  |
| sp\|Q9BTE6\|AASD1\_HUMAN | Q9BTE6 | AASD1\_HUMAN | AARSD1 | Alanyl-tRNA editing protein Aarsd1 | 412 | 45.480 | 3.143329e+04 | 1.796087e+06 | 570948.1 | 1181033.5 | 44105.484 |  | 42.48 | 56.3 | 16.5 |  |
| sp\|O14976\|GAK\_HUMAN | O14976 | GAK\_HUMAN | GAK | Cyclin-G-associated kinase | 1311 | 143.191 | 1.628996e+04 | 2.930575e+06 | 570215.0 | 689569.56 | 1463808.1 | 206982.31 | 31.2 | 35.2 | 21.6 | 7.9 |
| sp\|Q9UHC7\|MKRN1\_HUMAN | Q9UHC7 | MKRN1\_HUMAN | MKRN1 | E3 ubiquitin-protein ligase makorin-1 | 482 | 53.349 | 1.525024e+04 | 1.022163e+06 | 570158.7 | 345153.4 | 54686.363 | 52164.29 | 6.85 | 32.8 | 22.6 | 12.4 |
| sp\|Q9NVZ3\|NECP2\_HUMAN | Q9NVZ3 | NECP2\_HUMAN | NECAP2 | Adaptin ear-binding coat-associated protein 2 | 263 | 28.339 | 4.341392e+04 | 1.545720e+06 | 568788.0 | 890440.3 | 86491.25 |  | 39.16 | 66.9 | 38.8 |  |
| sp\|Q9BVA0\|KTNB1\_HUMAN | Q9BVA0 | KTNB1\_HUMAN | KATNB1 | Katanin p80 WD40 repeat-containing subunit B1 | 655 | 72.334 | 9.065505e+03 | 8.238567e+05 | 567697.44 | 224443.61 | 31715.604 |  | 16.03 | 47.6 | 24.0 |  |
| sp\|Q9Y666\|S12A7\_HUMAN | Q9Y666 | S12A7\_HUMAN | SLC12A7 | Solute carrier family 12 member 7 | 1083 | 119.106 | 3.383750e+04 | 5.063480e+06 | 567243.1 | 352194.12 | 546536.44 | 3597506.0 | 20.87 | 29.9 | 22.6 | 19.5 |
| sp\|O94915\|FRYL\_HUMAN | O94915 | FRYL\_HUMAN | FRYL | Protein furry homolog-like | 3013 | 339.598 | 1.835317e+03 | 7.830569e+05 | 566623.75 | 216433.17 |  |  | 11.18 | 30.0 |  |  |
| sp\|P67809\|YBOX1\_HUMAN | P67809 | YBOX1\_HUMAN | YBX1 | Y-box-binding protein 1 | 324 | 35.924 | 5.325370e+04 | 2.403542e+06 | 566480.75 | 1265887.6 | 441086.38 | 130086.945 | 45.06 | 55.2 | 43.2 | 31.2 |
| sp\|Q3YEC7\|RABL6\_HUMAN | Q3YEC7 | RABL6\_HUMAN | RABL6 | Rab-like protein 6 | 729 | 79.549 | 1.724797e+04 | 1.723812e+06 | 565999.5 | 594574.06 | 231824.75 | 331413.78 | 24.14 | 31.8 | 19.1 | 14.4 |
| sp\|Q7KZI7\|MARK2\_HUMAN | Q7KZI7 | MARK2\_HUMAN | MARK2 | Serine/threonine-protein kinase MARK2 | 788 | 87.911 | 2.133397e+04 | 2.356308e+06 | 564844.06 | 1383981.2 | 407482.38 |  | 27.28 | 57.7 | 40.0 |  |
| sp\|P23258\|TBG1\_HUMAN | P23258 | TBG1\_HUMAN | TUBG1 | Tubulin gamma-1 chain | 451 | 51.170 | 1.100348e+04 | 7.073958e+05 | 564256.44 | 143139.31 |  |  | 37.92 | 25.3 |  |  |
| sp\|P41214\|EIF2D\_HUMAN | P41214 | EIF2D\_HUMAN | EIF2D | Eukaryotic translation initiation factor 2D | 584 | 64.706 | 1.937850e+04 | 1.575367e+06 | 563640.06 | 859043.6 | 128061.51 | 24621.984 | 44.01 | 40.1 | 18.3 | 7.9 |
| sp\|P67870\|CSK2B\_HUMAN | P67870 | CSK2B\_HUMAN | CSNK2B | Casein kinase II subunit beta | 215 | 24.942 | 6.475577e+04 | 2.029210e+06 | 563614.44 | 1465595.1 |  |  | 31.16 | 40.5 |  |  |
| sp\|O14950\|ML12B\_HUMAN | O14950 | ML12B\_HUMAN | MYL12B | Myosin regulatory light chain 12B | 172 | 19.779 | 7.706031e+04 | 1.914927e+06 | 561816.75 | 927123.1 | 425987.25 |  | 54.65 | 65.5 | 24.0 |  |
| sp\|Q9NZW5\|PALS2\_HUMAN | Q9NZW5 | PALS2\_HUMAN | PALS2 | Protein PALS2 | 540 | 61.117 | 3.811546e+04 | 2.926715e+06 | 560294.56 | 476392.12 | 950050.9 | 939977.2 | 47.96 | 44.1 | 49.1 | 35.6 |
| sp\|O00625\|PIR\_HUMAN | O00625 | PIR\_HUMAN | PIR | Pirin | 290 | 32.113 | 2.666172e+04 | 1.075688e+06 | 560139.9 | 515547.75 |  |  | 32.07 | 35.2 |  |  |
| sp\|Q96A49\|SYAP1\_HUMAN | Q96A49 | SYAP1\_HUMAN | SYAP1 | Synapse-associated protein 1 | 352 | 39.933 | 3.661129e+04 | 1.836809e+06 | 559006.25 | 728979.25 | 478409.06 | 70414.875 | 35.23 | 46.9 | 42.6 | 6.5 |
| sp\|O94804\|STK10\_HUMAN | O94804 | STK10\_HUMAN | STK10 | Serine/threonine-protein kinase 10 | 968 | 112.135 | 1.120650e+04 | 1.578805e+06 | 557849.3 | 796170.75 | 205496.97 | 19288.07 | 22.52 | 38.7 | 27.3 | 14.6 |
| sp\|Q99729\|ROAA\_HUMAN | Q99729 | ROAA\_HUMAN | HNRNPAB | Heterogeneous nuclear ribonucleoprotein A/B | 332 | 36.225 | 4.122505e+04 | 1.876233e+06 | 555130.75 | 1321101.9 |  |  | 29.22 | 36.4 |  |  |
| sp\|Q8N4P3\|MESH1\_HUMAN | Q8N4P3 | MESH1\_HUMAN | HDDC3 | Guanosine-3',5'-bis(diphosphate) 3'-pyrophosphohydrolase MESH1 | 179 | 20.329 | 1.465750e+05 | 3.743632e+06 | 553452.4 | 3117596.0 | 72584.0 |  | 39.11 | 60.9 | 33.5 |  |
| sp\|P05109\|S10A8\_HUMAN | P05109 | S10A8\_HUMAN | S100A8 | Protein S100-A8 | 93 | 10.835 | 4.809289e+05 | 6.546768e+06 | 551752.0 | 407689.56 | 5587326.5 |  | 39.78 | 68.8 | 47.3 |  |
| sp\|Q6NYC1\|JMJD6\_HUMAN | Q6NYC1 | JMJD6\_HUMAN | JMJD6 | Bifunctional arginine demethylase and lysyl-hydroxylase JMJD6 | 403 | 46.462 | 1.709353e+04 | 9.978076e+05 | 551676.5 | 446131.12 |  |  | 26.05 | 43.4 |  |  |
| sp\|Q8IYB7\|DI3L2\_HUMAN | Q8IYB7 | DI3L2\_HUMAN | DIS3L2 | DIS3-like exonuclease 2 | 885 | 99.279 | 1.410483e+04 | 1.759310e+06 | 549644.6 | 1107459.8 | 52130.1 | 50075.453 | 42.03 | 45.9 | 21.4 | 12.1 |
| sp\|Q9UJ41\|RABX5\_HUMAN | Q9UJ41 | RABX5\_HUMAN | RABGEF1 | Rab5 GDP/GTP exchange factor | 491 | 56.891 | 1.154005e+04 | 8.248376e+05 | 548723.75 | 276113.88 |  |  | 25.87 | 45.4 |  |  |
| sp\|Q6ZUJ8\|BCAP\_HUMAN | Q6ZUJ8 | BCAP\_HUMAN | PIK3AP1 | Phosphoinositide 3-kinase adapter protein 1 | 805 | 90.398 | 1.412668e+04 | 1.604412e+06 | 547375.4 | 890855.1 | 139082.5 | 27099.406 | 31.55 | 42.5 | 28.9 | 9.6 |
| sp\|Q01433\|AMPD2\_HUMAN | Q01433 | AMPD2\_HUMAN | AMPD2 | AMP deaminase 2 | 879 | 94.890 | 1.682089e+04 | 2.005333e+06 | 546067.4 | 1459266.0 |  |  | 22.07 | 42.8 |  |  |
| sp\|Q9H223\|EHD4\_HUMAN | Q9H223 | EHD4\_HUMAN | EHD4 | EH domain-containing protein 4 | 541 | 61.175 | 1.020066e+04 | 7.840057e+05 | 545735.6 | 172599.98 | 65670.11 |  | 25.51 | 33.6 | 20.7 |  |
| sp\|Q6PIU2\|NCEH1\_HUMAN | Q6PIU2 | NCEH1\_HUMAN | NCEH1 | Neutral cholesterol ester hydrolase 1 | 408 | 45.808 | 6.592299e+04 | 3.793983e+06 | 543871.2 | 1136584.4 | 916227.4 | 1197299.8 | 36.52 | 49.3 | 29.7 | 14.2 |
| sp\|P55212\|CASP6\_HUMAN | P55212 | CASP6\_HUMAN | CASP6 | Caspase-6 | 293 | 33.310 | 3.263010e+04 | 1.365558e+06 | 542808.5 | 822749.75 |  |  | 50.17 | 61.1 |  |  |
| sp\|Q14103\|HNRPD\_HUMAN | Q14103 | HNRPD\_HUMAN | HNRNPD | Heterogeneous nuclear ribonucleoprotein D0 | 355 | 38.434 | 5.453301e+04 | 2.633251e+06 | 541417.3 | 1995761.5 | 81816.83 | 14254.91 | 27.61 | 43.7 | 21.7 | 16.9 |
| sp\|Q9H3S7\|PTN23\_HUMAN | Q9H3S7 | PTN23\_HUMAN | PTPN23 | Tyrosine-protein phosphatase non-receptor type 23 | 1636 | 178.974 | 9.497101e+03 | 2.135493e+06 | 541366.25 | 884310.5 | 504371.2 | 205445.0 | 19.74 | 32.8 | 19.4 | 14.4 |
| sp\|Q27J81\|INF2\_HUMAN | Q27J81 | INF2\_HUMAN | INF2 | Inverted formin-2 | 1249 | 135.624 | 1.244528e+04 | 2.120598e+06 | 541189.3 | 843595.56 | 307770.94 | 428042.34 | 30.58 | 37.4 | 20.1 | 11.8 |
| sp\|Q15084\|PDIA6\_HUMAN | Q15084 | PDIA6\_HUMAN | PDIA6 | Protein disulfide-isomerase A6 | 440 | 48.121 | 3.474831e+04 | 2.100804e+06 | 541133.6 | 509204.66 | 906290.2 | 144175.16 | 30.23 | 35.7 | 25.0 | 14.3 |
| sp\|Q8IWB9\|TEX2\_HUMAN | Q8IWB9 | TEX2\_HUMAN | TEX2 | Testis-expressed protein 2 | 1127 | 125.303 | 5.507650e+03 | 8.670516e+05 | 540765.5 | 152414.38 | 74072.945 | 99798.766 | 4.08 | 33.8 | 16.9 | 12.0 |
| sp\|Q9BSJ8\|ESYT1\_HUMAN | Q9BSJ8 | ESYT1\_HUMAN | ESYT1 | Extended synaptotagmin-1 | 1104 | 122.856 | 1.389470e+04 | 2.144681e+06 | 539991.6 | 570119.1 | 572196.8 | 462373.25 | 37.05 | 35.7 | 17.0 | 8.2 |
| sp\|O43447\|PPIH\_HUMAN | O43447 | PPIH\_HUMAN | PPIH | Peptidyl-prolyl cis-trans isomerase H | 177 | 19.208 | 1.042176e+05 | 2.515013e+06 | 539966.2 | 1886686.4 | 88360.22 |  | 42.37 | 88.7 | 46.9 |  |
| sp\|Q5T3F8\|CSCL2\_HUMAN | Q5T3F8 | CSCL2\_HUMAN | TMEM63B | CSC1-like protein 2 | 832 | 94.958 | 6.555341e+04 | 7.820671e+06 | 539645.2 | 746992.0 | 2435573.5 | 4098460.0 | 13.1 | 17.9 | 19.4 | 16.3 |
| sp\|Q96S19\|MTL26\_HUMAN | Q96S19 | MTL26\_HUMAN | METTL26 | Methyltransferase-like 26 | 204 | 22.578 | 5.938946e+04 | 1.684659e+06 | 539179.75 | 1145479.1 |  |  | 43.14 | 49.0 |  |  |
| sp\|Q5JSL3\|DOC11\_HUMAN | Q5JSL3 | DOC11\_HUMAN | DOCK11 | Dedicator of cytokinesis protein 11 | 2073 | 237.671 | 2.116055e+03 | 6.318592e+05 | 537424.56 | 89940.91 |  | 4493.7573 | 1.83 | 36.1 |  | 8.8 |
| sp\|O15173\|PGRC2\_HUMAN | O15173 | PGRC2\_HUMAN | PGRMC2 | Membrane-associated progesterone receptor component 2 | 223 | 23.818 | 8.677148e+04 | 2.596567e+06 | 536456.4 | 993877.3 | 739245.8 | 326987.1 | 34.08 | 51.1 | 39.5 | 29.1 |
| sp\|Q92882\|OSTF1\_HUMAN | Q92882 | OSTF1\_HUMAN | OSTF1 | Osteoclast-stimulating factor 1 | 214 | 23.787 | 4.886660e+04 | 1.460390e+06 | 533659.25 | 554829.3 | 371901.9 |  | 36.92 | 64.5 | 30.4 |  |
| sp\|P42345\|MTOR\_HUMAN | P42345 | MTOR\_HUMAN | MTOR | Serine/threonine-protein kinase mTOR | 2549 | 288.892 | 8.815516e+03 | 3.199635e+06 | 533073.3 | 2365296.8 | 57590.42 | 243674.42 | 31.11 | 42.7 | 14.6 | 10.8 |
| sp\|Q96K21\|ANCHR\_HUMAN | Q96K21 | ANCHR\_HUMAN | ZFYVE19 | Abscission/NoCut checkpoint regulator | 471 | 51.546 | 3.042162e+04 | 1.970128e+06 | 532455.1 | 1052180.5 | 272570.62 | 112921.89 | 31.63 | 64.3 | 43.3 | 14.9 |
| sp\|O95273\|CCDB1\_HUMAN | O95273 | CCDB1\_HUMAN | CCNDBP1 | Cyclin-D1-binding protein 1 | 360 | 40.262 | 3.994903e+04 | 2.020779e+06 | 531258.4 | 1266184.0 | 186254.45 | 37081.83 | 23.06 | 30.8 | 21.7 | 10.0 |
| sp\|P49591\|SYSC\_HUMAN | P49591 | SYSC\_HUMAN | SARS1 | Serine--tRNA ligase, cytoplasmic | 514 | 58.777 | 1.623123e+04 | 1.198605e+06 | 530184.1 | 633245.7 | 35175.094 |  | 28.02 | 57.6 | 22.2 |  |
| sp\|Q14677\|EPN4\_HUMAN | Q14677 | EPN4\_HUMAN | CLINT1 | Clathrin interactor 1 | 625 | 68.259 | 4.497408e+04 | 3.856909e+06 | 529609.1 | 1141215.2 | 1676549.4 | 509534.94 | 28.48 | 29.9 | 32.6 | 19.7 |
| sp\|Q13591\|SEM5A\_HUMAN | Q13591 | SEM5A\_HUMAN | SEMA5A | Semaphorin-5A | 1074 | 118.140 | 3.565250e+03 | 5.291809e+05 | 529180.9 |  |  |  | 0.84 |  |  |  |
| sp\|Q14139\|UBE4A\_HUMAN | Q14139 | UBE4A\_HUMAN | UBE4A | Ubiquitin conjugation factor E4 A | 1066 | 122.561 | 9.006530e+03 | 1.386842e+06 | 528313.2 | 830403.56 | 7920.56 | 20204.62 | 18.86 | 34.2 | 16.2 | 6.5 |
| sp\|Q9BXS5\|AP1M1\_HUMAN | Q9BXS5 | AP1M1\_HUMAN | AP1M1 | AP-1 complex subunit mu-1 | 423 | 48.587 | 2.791557e+04 | 1.704056e+06 | 526099.1 | 633104.4 | 544852.25 |  | 41.37 | 58.4 | 42.3 |  |
| sp\|Q96GX9\|MTNB\_HUMAN | Q96GX9 | MTNB\_HUMAN | APIP | Methylthioribulose-1-phosphate dehydratase | 242 | 27.125 | 6.645729e+04 | 2.264798e+06 | 524222.97 | 1740575.4 |  |  | 60.74 | 68.2 |  |  |
| sp\|O75190\|DNJB6\_HUMAN | O75190 | DNJB6\_HUMAN | DNAJB6 | DnaJ homolog subfamily B member 6 | 326 | 36.087 | 2.708988e+04 | 1.228217e+06 | 521849.88 | 214861.53 | 491505.22 |  | 15.03 | 43.6 | 33.7 |  |
| sp\|Q53H82\|LACB2\_HUMAN | Q53H82 | LACB2\_HUMAN | LACTB2 | Endoribonuclease LACTB2 | 288 | 32.806 | 6.956940e+04 | 2.867403e+06 | 519351.44 | 2348051.2 |  |  | 43.75 | 55.2 |  |  |
| sp\|Q9NXR7\|BABA2\_HUMAN | Q9NXR7 | BABA2\_HUMAN | BABAM2 | BRISC and BRCA1-A complex member 2 | 383 | 43.552 | 2.511571e+04 | 1.374266e+06 | 519292.88 | 854973.0 |  |  | 28.2 | 48.3 |  |  |
| sp\|O94979\|SC31A\_HUMAN | O94979 | SC31A\_HUMAN | SEC31A | Protein transport protein Sec31A | 1220 | 133.015 | 1.254368e+04 | 2.096248e+06 | 519206.1 | 1250705.6 | 298529.3 | 27806.555 | 27.95 | 35.2 | 15.1 | 6.6 |
| sp\|Q9BZV1\|UBXN6\_HUMAN | Q9BZV1 | UBXN6\_HUMAN | UBXN6 | UBX domain-containing protein 6 | 441 | 49.754 | 2.866622e+04 | 1.791908e+06 | 517042.47 | 1168959.2 | 105905.91 |  | 35.15 | 62.6 | 12.9 |  |
| sp\|Q99747\|SNAG\_HUMAN | Q99747 | SNAG\_HUMAN | NAPG | Gamma-soluble NSF attachment protein | 312 | 34.746 | 3.501595e+04 | 1.528579e+06 | 513581.75 | 614155.25 | 156478.67 | 244363.44 | 45.19 | 48.7 | 30.1 | 28.8 |
| sp\|P14174\|MIF\_HUMAN | P14174 | MIF\_HUMAN | MIF | Macrophage migration inhibitory factor | 115 | 12.476 | 2.933867e+05 | 4.598678e+06 | 512145.6 | 4025868.2 | 60664.062 |  | 17.39 | 52.2 | 10.4 |  |
| sp\|Q6PHR2\|ULK3\_HUMAN | Q6PHR2 | ULK3\_HUMAN | ULK3 | Serine/threonine-protein kinase ULK3 | 472 | 53.444 | 1.205166e+04 | 8.092134e+05 | 512124.06 | 188065.75 | 109023.59 |  | 30.3 | 35.0 | 21.4 |  |
| sp\|O00182\|LEG9\_HUMAN | O00182 | LEG9\_HUMAN | LGALS9 | Galectin-9 | 355 | 39.518 | 1.072754e+05 | 5.326136e+06 | 511594.56 | 473949.53 | 3579549.0 | 761043.4 | 25.63 | 31.8 | 28.7 | 11.8 |
| sp\|Q05086\|UBE3A\_HUMAN | Q05086 | UBE3A\_HUMAN | UBE3A | Ubiquitin-protein ligase E3A | 875 | 100.688 | 2.530157e+04 | 3.200680e+06 | 511065.34 | 2616392.5 | 73222.586 |  | 34.86 | 48.2 | 13.6 |  |
| sp\|Q8IY33\|MILK2\_HUMAN | Q8IY33 | MILK2\_HUMAN | MICALL2 | MICAL-like protein 2 | 904 | 97.502 | 1.196655e+04 | 1.465884e+06 | 508394.88 | 333028.62 | 500035.25 | 124425.6 | 29.31 | 64.6 | 28.4 | 21.6 |
| sp\|Q16570\|ACKR1\_HUMAN | Q16570 | ACKR1\_HUMAN | ACKR1 | Atypical chemokine receptor 1 | 336 | 36.960 | 1.091902e+04 | 5.070288e+05 | 507028.8 |  |  |  | 6.85 |  |  |  |
| sp\|Q68D91\|MBLC2\_HUMAN | Q68D91 | MBLC2\_HUMAN | MBLAC2 | Acyl-coenzyme A thioesterase MBLAC2 | 279 | 31.372 | 4.772781e+04 | 1.881182e+06 | 506673.8 | 1146092.5 | 118540.67 | 109875.17 | 41.58 | 54.1 | 40.9 | 28.7 |
| sp\|P80217\|IN35\_HUMAN | P80217 | IN35\_HUMAN | IFI35 | Interferon-induced 35 kDa protein | 286 | 31.546 | 3.125386e+04 | 1.238697e+06 | 506549.78 | 700223.6 | 31923.67 |  | 25.52 | 60.1 | 29.0 |  |
| sp\|P10746\|HEM4\_HUMAN | P10746 | HEM4\_HUMAN | UROS | Uroporphyrinogen-III synthase | 265 | 28.628 | 1.187786e+05 | 4.272149e+06 | 502349.97 | 3490022.2 | 204883.81 | 74892.7 | 52.45 | 53.2 | 27.5 | 3.8 |
| sp\|P35754\|GLRX1\_HUMAN | P35754 | GLRX1\_HUMAN | GLRX | Glutaredoxin-1 | 106 | 11.776 | 4.344218e+05 | 6.427271e+06 | 502247.84 | 3901571.5 | 1858246.1 | 165205.47 | 23.58 | 43.4 | 24.5 | 26.4 |
| sp\|Q08257\|QOR\_HUMAN | Q08257 | QOR\_HUMAN | CRYZ | Quinone oxidoreductase | 329 | 35.207 | 5.601280e+04 | 2.477613e+06 | 501886.44 | 1956347.6 |  | 19379.05 | 46.81 | 54.4 |  | 10.0 |
| sp\|P55160\|NCKPL\_HUMAN | P55160 | NCKPL\_HUMAN | NCKAP1L | Nck-associated protein 1-like | 1127 | 128.153 | 5.549892e+03 | 8.935739e+05 | 501738.7 | 391835.2 |  |  | 10.38 | 26.3 |  |  |
| sp\|Q9H299\|SH3L3\_HUMAN | Q9H299 | SH3L3\_HUMAN | SH3BGRL3 | SH3 domain-binding glutamic acid-rich-like protein 3 | 93 | 10.438 | 2.431820e+05 | 3.189083e+06 | 499946.9 | 2158689.0 | 312485.7 | 217961.52 | 43.01 | 49.5 | 10.8 | 22.6 |
| sp\|Q01970\|PLCB3\_HUMAN | Q01970 | PLCB3\_HUMAN | PLCB3 | 1-phosphatidylinositol 4,5-bisphosphate phosphodiesterase beta-3 | 1234 | 138.799 | 5.345940e+03 | 9.322397e+05 | 499355.7 | 381270.75 | 51613.23 |  | 29.5 | 36.5 | 20.2 |  |
| sp\|P07814\|SYEP\_HUMAN | P07814 | SYEP\_HUMAN | EPRS1 | Bifunctional glutamate/proline--tRNA ligase | 1512 | 170.591 | 9.409077e+03 | 2.016602e+06 | 499112.62 | 1376929.8 | 46897.383 | 93662.59 | 13.62 | 34.1 | 24.0 | 14.0 |
| sp\|Q96H20\|SNF8\_HUMAN | Q96H20 | SNF8\_HUMAN | SNF8 | Vacuolar-sorting protein SNF8 | 258 | 28.864 | 3.895737e+04 | 1.412743e+06 | 497548.28 | 520199.28 | 313392.62 | 81603.28 | 30.23 | 27.9 | 20.9 | 8.9 |
| sp\|P29590\|PML\_HUMAN | P29590 | PML\_HUMAN | PML | Protein PML | 882 | 97.551 | 9.006957e+03 | 1.103893e+06 | 497057.22 | 496492.06 | 23914.729 | 86428.914 | 15.31 | 33.8 | 29.5 | 13.0 |
| sp\|Q9P265\|DIP2B\_HUMAN | Q9P265 | DIP2B\_HUMAN | DIP2B | Disco-interacting protein 2 homolog B | 1576 | 171.492 | 6.382367e+03 | 1.375127e+06 | 496445.53 | 751707.1 | 35582.605 | 91391.7 | 25.25 | 49.0 | 18.3 | 13.2 |
| sp\|Q13185\|CBX3\_HUMAN | Q13185 | CBX3\_HUMAN | CBX3 | Chromobox protein homolog 3 | 183 | 20.811 | 5.432697e+04 | 1.420449e+06 | 496179.94 | 733402.7 | 173715.78 | 17150.285 | 30.05 | 57.4 | 20.8 | 19.1 |
| sp\|A5D8V6\|VP37C\_HUMAN | A5D8V6 | VP37C\_HUMAN | VPS37C | Vacuolar protein sorting-associated protein 37C | 355 | 38.659 | 6.792205e+04 | 3.298971e+06 | 495223.56 | 1243220.2 | 1407275.2 | 153252.5 | 12.11 | 29.6 | 14.9 | 3.1 |
| sp\|Q4G0F5\|VP26B\_HUMAN | Q4G0F5 | VP26B\_HUMAN | VPS26B | Vacuolar protein sorting-associated protein 26B | 336 | 39.155 | 5.047463e+04 | 2.483005e+06 | 494306.4 | 1805811.5 | 182886.9 |  | 32.14 | 40.2 | 28.3 |  |
| sp\|Q93070\|NAR4\_HUMAN | Q93070 | NAR4\_HUMAN | ART4 | Ecto-ADP-ribosyltransferase 4 | 314 | 35.878 | 1.056337e+05 | 4.761543e+06 | 493775.6 | 592744.75 | 1819628.6 | 1855394.0 | 19.43 | 50.6 | 49.7 | 15.6 |
| sp\|Q9BPX3\|CND3\_HUMAN | Q9BPX3 | CND3\_HUMAN | NCAPG | Condensin complex subunit 3 | 1015 | 114.334 | 7.524642e+03 | 1.080882e+06 | 493213.8 | 587668.4 |  |  | 30.54 | 37.9 |  |  |
| sp\|Q9Y490\|TLN1\_HUMAN | Q9Y490 | TLN1\_HUMAN | TLN1 | Talin-1 | 2541 | 269.767 | 1.021267e+04 | 3.461350e+06 | 491589.6 | 1558611.9 | 1300612.5 | 110535.59 | 27.47 | 59.4 | 26.2 | 14.5 |
| sp\|P46976\|GLYG\_HUMAN | P46976 | GLYG\_HUMAN | GYG1 | Glycogenin-1 | 350 | 39.384 | 2.441546e+04 | 1.208097e+06 | 491446.5 | 412742.97 | 303907.9 |  | 18.0 | 51.1 | 14.0 |  |
| sp\|Q9Y316\|MEMO1\_HUMAN | Q9Y316 | MEMO1\_HUMAN | MEMO1 | Protein MEMO1 | 297 | 33.733 | 3.013822e+04 | 1.277290e+06 | 490667.88 | 786622.56 |  |  | 36.03 | 47.5 |  |  |
| sp\|Q13325\|IFIT5\_HUMAN | Q13325 | IFIT5\_HUMAN | IFIT5 | Interferon-induced protein with tetratricopeptide repeats 5 | 482 | 55.847 | 2.849906e+04 | 1.999620e+06 | 489905.22 | 1293562.5 | 216152.61 |  | 24.9 | 41.9 | 16.4 |  |
| sp\|O95372\|LYPA2\_HUMAN | O95372 | LYPA2\_HUMAN | LYPLA2 | Acyl-protein thioesterase 2 | 231 | 24.737 | 3.097942e+04 | 9.628031e+05 | 488106.66 | 334916.34 |  | 139780.12 | 30.3 | 58.4 |  | 26.4 |
| sp\|Q9H9A6\|LRC40\_HUMAN | Q9H9A6 | LRC40\_HUMAN | LRRC40 | Leucine-rich repeat-containing protein 40 | 602 | 68.250 | 2.535245e+04 | 2.173901e+06 | 488050.56 | 1297089.9 | 254212.0 | 134548.34 | 49.17 | 27.6 | 26.9 | 13.8 |
| sp\|Q8TEH3\|DEN1A\_HUMAN | Q8TEH3 | DEN1A\_HUMAN | DENND1A | DENN domain-containing protein 1A | 1009 | 110.577 | 5.129338e+04 | 7.125958e+06 | 487695.34 | 1000877.9 | 1174715.5 | 4462669.0 | 21.01 | 36.8 | 28.7 | 14.5 |
| sp\|Q8WVV4\|POF1B\_HUMAN | Q8WVV4 | POF1B\_HUMAN | POF1B | Protein POF1B | 589 | 68.065 | 1.335557e+04 | 1.142098e+06 | 486982.6 | 117928.305 | 255381.3 | 281806.12 | 11.88 | 33.3 | 22.4 | 15.8 |
| sp\|P51570\|GALK1\_HUMAN | P51570 | GALK1\_HUMAN | GALK1 | Galactokinase | 392 | 42.272 | 2.107195e+04 | 1.119115e+06 | 484687.28 | 493685.53 | 122386.04 | 18356.043 | 35.46 | 40.6 | 24.0 | 11.0 |
| sp\|Q12986\|NFX1\_HUMAN | Q12986 | NFX1\_HUMAN | NFX1 | Transcriptional repressor NF-X1 | 1120 | 124.395 | 5.333702e+03 | 8.335829e+05 | 484557.2 | 324673.56 | 8619.193 | 15732.946 | 11.16 | 45.3 | 27.2 | 18.9 |
| sp\|Q96T66\|NMNA3\_HUMAN | Q96T66 | NMNA3\_HUMAN | NMNAT3 | Nicotinamide/nicotinic acid mononucleotide adenylyltransferase 3 | 252 | 28.322 | 3.377524e+04 | 1.201820e+06 | 483601.78 | 619880.5 | 98337.9 |  | 44.44 | 65.1 | 26.2 |  |
| sp\|P47929\|LEG7\_HUMAN | P47929 | LEG7\_HUMAN | LGALS7 | Galectin-7 | 136 | 15.075 | 1.405280e+05 | 2.661566e+06 | 482930.78 | 153895.73 | 1010349.8 | 1014389.5 | 44.85 | 69.9 | 68.4 | 20.6 |
| sp\|A4D126\|ISPD\_HUMAN | A4D126 | ISPD\_HUMAN | CRPPA | D-ribitol-5-phosphate cytidylyltransferase | 451 | 49.873 | 3.805142e+04 | 2.384260e+06 | 482602.0 | 1587489.9 | 314167.62 |  | 29.49 | 61.2 | 24.6 |  |
| sp\|O94832\|MYO1D\_HUMAN | O94832 | MYO1D\_HUMAN | MYO1D | Unconventional myosin-Id | 1006 | 116.202 | 4.617297e+03 | 6.740911e+05 | 482269.5 | 191821.6 |  |  | 21.37 | 34.4 |  |  |
| sp\|Q92599\|SEPT8\_HUMAN | Q92599 | SEPT8\_HUMAN | SEPTIN8 | Septin-8 | 483 | 55.756 | 3.293267e+05 | 2.306937e+07 | 481791.56 | 901574.6 | 21686004.0 |  | 22.98 | 38.3 | 17.2 |  |
| sp\|Q7L5D6\|GET4\_HUMAN | Q7L5D6 | GET4\_HUMAN | GET4 | Golgi to ER traffic protein 4 homolog | 327 | 36.504 | 4.032897e+04 | 1.849587e+06 | 480702.9 | 927508.6 | 355708.25 | 85667.25 | 31.5 | 35.5 | 27.8 | 14.1 |
| sp\|Q9H3K6\|BOLA2\_HUMAN | Q9H3K6 | BOLA2\_HUMAN | BOLA2 | BolA-like protein 2 | 86 | 10.117 | 2.832875e+05 | 3.600778e+06 | 480316.3 | 2181284.5 | 869098.5 | 70078.37 | 61.63 | 77.9 | 79.1 | 38.4 |
| sp\|Q9NRW7\|VPS45\_HUMAN | Q9NRW7 | VPS45\_HUMAN | VPS45 | Vacuolar protein sorting-associated protein 45 | 570 | 65.077 | 2.570005e+04 | 2.101255e+06 | 478982.0 | 879423.3 | 742849.5 |  | 34.39 | 40.5 | 27.7 |  |
| sp\|Q93050\|VPP1\_HUMAN | Q93050 | VPP1\_HUMAN | ATP6V0A1 | V-type proton ATPase 116 kDa subunit a 1 | 837 | 96.413 | 2.840530e+04 | 3.440742e+06 | 475005.2 | 606622.75 | 832423.1 | 1526690.9 | 21.27 | 28.1 | 21.9 | 16.8 |
| sp\|Q9BVM2\|DPCD\_HUMAN | Q9BVM2 | DPCD\_HUMAN | DPCD | Protein DPCD | 203 | 23.240 | 7.784799e+04 | 2.273007e+06 | 474935.16 | 1396565.5 | 178567.53 | 222938.36 | 35.47 | 61.1 | 29.6 | 17.7 |
| sp\|P07355\|ANXA2\_HUMAN | P07355 | ANXA2\_HUMAN | ANXA2 | Annexin A2 | 339 | 38.604 | 1.135597e+05 | 5.507746e+06 | 473097.75 | 559286.3 | 2218829.5 | 2256532.0 | 44.84 | 54.6 | 47.5 | 45.1 |
| sp\|Q92747\|ARC1A\_HUMAN | Q92747 | ARC1A\_HUMAN | ARPC1A | Actin-related protein 2/3 complex subunit 1A | 370 | 41.569 | 2.033696e+04 | 1.062118e+06 | 472960.1 | 589157.9 |  |  | 30.54 | 48.9 |  |  |
| sp\|P0DPA2\|VSIG8\_HUMAN | P0DPA2 | VSIG8\_HUMAN | VSIG8 | V-set and immunoglobulin domain-containing protein 8 | 414 | 43.891 | 9.763432e+03 | 5.383877e+05 | 471653.2 |  |  | 66734.51 | 12.08 |  |  | 9.7 |
| sp\|P49407\|ARRB1\_HUMAN | P49407 | ARRB1\_HUMAN | ARRB1 | Beta-arrestin-1 | 418 | 47.066 | 2.252355e+04 | 1.331868e+06 | 471250.84 | 806706.75 | 53910.645 |  | 39.23 | 53.1 | 16.3 |  |
| sp\|Q00688\|FKBP3\_HUMAN | Q00688 | FKBP3\_HUMAN | FKBP3 | Peptidyl-prolyl cis-trans isomerase FKBP3 | 224 | 25.177 | 1.012602e+05 | 3.203022e+06 | 470990.75 | 2336061.8 | 242870.44 | 153099.0 | 41.52 | 52.7 | 19.6 | 14.3 |
| sp\|Q96M27\|PRRC1\_HUMAN | Q96M27 | PRRC1\_HUMAN | PRRC1 | Protein PRRC1 | 445 | 46.701 | 2.641896e+04 | 1.550098e+06 | 470195.88 | 461301.28 | 618600.5 |  | 24.04 | 25.4 | 19.1 |  |
| sp\|Q9UBN7\|HDAC6\_HUMAN | Q9UBN7 | HDAC6\_HUMAN | HDAC6 | Histone deacetylase 6 | 1215 | 131.419 | 1.121247e+04 | 1.851299e+06 | 469977.2 | 1289756.1 | 91565.45 |  | 8.07 | 15.8 | 3.6 |  |
| sp\|P61077\|UB2D3\_HUMAN | P61077 | UB2D3\_HUMAN | UBE2D3 | Ubiquitin-conjugating enzyme E2 D3 | 147 | 16.687 | 4.457423e+04 | 9.344998e+05 | 468823.28 | 465676.56 |  |  | 42.86 | 26.5 |  |  |
| sp\|Q15517\|CDSN\_HUMAN | Q15517 | CDSN\_HUMAN | CDSN | Corneodesmosin | 529 | 51.522 | 1.621209e+04 | 1.049419e+06 | 467470.3 |  | 224442.06 | 357506.38 | 13.99 |  | 30.4 | 18.3 |
| sp\|Q12907\|LMAN2\_HUMAN | Q12907 | LMAN2\_HUMAN | LMAN2 | Vesicular integral-membrane protein VIP36 | 356 | 40.229 | 2.578209e+05 | 1.303090e+07 | 466407.88 | 522575.94 | 6055418.5 | 5986501.0 | 32.3 | 45.5 | 24.7 | 20.8 |
| sp\|P78330\|SERB\_HUMAN | P78330 | SERB\_HUMAN | PSPH | Phosphoserine phosphatase | 225 | 25.008 | 3.427867e+04 | 1.077011e+06 | 466247.72 | 610763.0 |  |  | 38.22 | 40.0 |  |  |
| sp\|Q8NHH9\|ATLA2\_HUMAN | Q8NHH9 | ATLA2\_HUMAN | ATL2 | Atlastin-2 | 583 | 66.229 | 2.032513e+04 | 1.691215e+06 | 463499.97 | 760097.2 | 109085.08 | 358532.5 | 26.76 | 20.8 | 13.0 | 13.0 |
| sp\|P07339\|CATD\_HUMAN | P07339 | CATD\_HUMAN | CTSD | Cathepsin D | 412 | 44.552 | 4.246507e+04 | 2.376929e+06 | 461914.84 | 33822.133 | 1047917.06 | 833274.75 | 17.48 | 18.2 | 34.7 | 17.0 |
| sp\|Q9NSD9\|SYFB\_HUMAN | Q9NSD9 | SYFB\_HUMAN | FARSB | Phenylalanine--tRNA ligase beta subunit | 589 | 66.116 | 8.230064e+03 | 6.836391e+05 | 460301.38 | 109631.86 | 113705.89 |  | 20.2 | 47.0 | 18.8 |  |
| sp\|Q96L94\|SNX22\_HUMAN | Q96L94 | SNX22\_HUMAN | SNX22 | Sorting nexin-22 | 193 | 22.068 | 4.510237e+04 | 1.250488e+06 | 459919.38 | 529391.94 | 261176.5 |  | 31.61 | 46.6 | 51.8 |  |
| sp\|Q9UBQ0\|VPS29\_HUMAN | Q9UBQ0 | VPS29\_HUMAN | VPS29 | Vacuolar protein sorting-associated protein 29 | 182 | 20.506 | 8.338247e+04 | 2.148191e+06 | 459335.94 | 1583249.9 | 105605.02 |  | 29.67 | 60.4 | 20.3 |  |
| sp\|Q8NFF5\|FAD1\_HUMAN | Q8NFF5 | FAD1\_HUMAN | FLAD1 | FAD synthase | 587 | 65.266 | 1.848086e+04 | 1.515397e+06 | 456246.2 | 1059150.8 |  |  | 36.12 | 47.4 |  |  |
| sp\|P07947\|YES\_HUMAN | P07947 | YES\_HUMAN | YES1 | Tyrosine-protein kinase Yes | 543 | 60.801 | 1.738494e+04 | 1.328009e+06 | 452551.28 | 129528.42 |  | 745929.2 | 8.66 | 48.3 |  | 11.0 |
| sp\|Q8N6M0\|OTU6B\_HUMAN | Q8N6M0 | OTU6B\_HUMAN | OTUD6B | Deubiquitinase OTUD6B | 293 | 33.813 | 4.784605e+04 | 2.032577e+06 | 451802.38 | 1498675.2 | 82099.195 |  | 18.77 | 53.9 | 24.2 |  |
| sp\|Q8TDB8\|GTR14\_HUMAN | Q8TDB8 | GTR14\_HUMAN | SLC2A14 | Solute carrier family 2, facilitated glucose transporter member 14 | 520 | 56.320 | 1.096647e+05 | 7.759733e+06 | 451312.88 | 1656393.2 | 754846.75 | 4897180.5 | 12.12 | 18.1 | 21.0 | 14.6 |
| sp\|O75369\|FLNB\_HUMAN | O75369 | FLNB\_HUMAN | FLNB | Filamin-B | 2602 | 278.164 | 2.614158e+03 | 9.135871e+05 | 451088.12 | 310118.28 | 67083.21 | 85297.47 | 4.38 | 42.9 | 20.3 | 12.6 |
| sp\|P05976\|MYL1\_HUMAN | P05976 | MYL1\_HUMAN | MYL1 | Myosin light chain 1/3, skeletal muscle isoform | 194 | 21.145 | 5.131775e+04 | 1.363303e+06 | 450775.0 | 427900.7 | 361035.4 | 123592.03 | 14.43 | 28.9 | 31.4 | 24.7 |
| sp\|Q9BUK6\|MSTO1\_HUMAN | Q9BUK6 | MSTO1\_HUMAN | MSTO1 | Protein misato homolog 1 | 570 | 61.835 | 1.724867e+04 | 1.340007e+06 | 450596.53 | 744206.06 | 145204.48 |  | 14.21 | 35.1 | 17.4 |  |
| sp\|Q9UKT5\|FBX4\_HUMAN | Q9UKT5 | FBX4\_HUMAN | FBXO4 | F-box only protein 4 | 387 | 44.136 | 1.249492e+04 | 6.928569e+05 | 449698.7 | 157978.73 | 77454.56 | 7724.9272 | 23.26 | 41.9 | 16.3 | 9.3 |
| sp\|Q9GZZ1\|NAA50\_HUMAN | Q9GZZ1 | NAA50\_HUMAN | NAA50 | N-alpha-acetyltransferase 50 | 169 | 19.398 | 9.357878e+04 | 2.280612e+06 | 447659.62 | 1601852.6 | 231100.12 |  | 43.2 | 72.8 | 53.3 |  |
| sp\|Q7Z3C6\|ATG9A\_HUMAN | Q7Z3C6 | ATG9A\_HUMAN | ATG9A | Autophagy-related protein 9A | 839 | 94.447 | 8.093965e+03 | 9.604320e+05 | 447135.53 | 192746.95 |  | 320549.56 | 14.42 | 33.0 |  | 13.9 |
| sp\|P29692\|EF1D\_HUMAN | P29692 | EF1D\_HUMAN | EEF1D | Elongation factor 1-delta | 281 | 31.122 | 5.126834e+04 | 2.004629e+06 | 446867.12 | 1335197.5 | 176651.33 | 45912.64 | 38.79 | 31.3 | 37.0 | 12.1 |
| sp\|Q9Y2B0\|CNPY2\_HUMAN | Q9Y2B0 | CNPY2\_HUMAN | CNPY2 | Protein canopy homolog 2 | 182 | 20.652 | 9.278528e+04 | 2.407456e+06 | 446591.25 | 95680.305 | 1548038.4 | 317145.9 | 21.43 | 35.2 | 41.8 | 33.5 |
| sp\|O43776\|SYNC\_HUMAN | O43776 | SYNC\_HUMAN | NARS1 | Asparagine--tRNA ligase, cytoplasmic | 548 | 62.943 | 1.122019e+04 | 8.872882e+05 | 446317.88 | 430014.9 | 10955.425 |  | 25.91 | 26.8 | 19.2 |  |
| sp\|O95819\|M4K4\_HUMAN | O95819 | M4K4\_HUMAN | MAP4K4 | Mitogen-activated protein kinase kinase kinase kinase 4 | 1239 | 142.101 | 6.722942e+03 | 1.200255e+06 | 445820.22 | 573360.1 | 181074.95 |  | 18.0 | 31.7 | 19.0 |  |
| sp\|P29350\|PTN6\_HUMAN | P29350 | PTN6\_HUMAN | PTPN6 | Tyrosine-protein phosphatase non-receptor type 6 | 595 | 67.561 | 8.669220e+03 | 7.358567e+05 | 445078.25 | 266937.8 | 23840.627 |  | 22.86 | 47.2 | 20.0 |  |
| sp\|Q16204\|CCDC6\_HUMAN | Q16204 | CCDC6\_HUMAN | CCDC6 | Coiled-coil domain-containing protein 6 | 474 | 53.291 | 2.748585e+04 | 1.840265e+06 | 444655.94 | 1007967.5 | 251434.12 | 136206.98 | 37.34 | 47.9 | 24.9 | 19.0 |
| sp\|O75390\|CISY\_HUMAN | O75390 | CISY\_HUMAN | CS | Citrate synthase, mitochondrial | 466 | 51.712 | 8.653205e+03 | 5.621931e+05 | 444514.94 | 61751.977 | 15957.215 | 39968.953 | 5.36 | 26.4 | 21.5 | 8.6 |
| sp\|Q9BQS8\|FYCO1\_HUMAN | Q9BQS8 | FYCO1\_HUMAN | FYCO1 | FYVE and coiled-coil domain-containing protein 1 | 1478 | 166.983 | 2.888655e+03 | 6.060174e+05 | 444330.38 | 161687.02 |  |  | 6.9 | 35.9 |  |  |
| sp\|P49354\|FNTA\_HUMAN | P49354 | FNTA\_HUMAN | FNTA | Protein farnesyltransferase/geranylgeranyltransferase type-1 subunit alpha | 379 | 44.409 | 2.638143e+04 | 1.471927e+06 | 443207.94 | 1028719.5 |  |  | 21.9 | 24.0 |  |  |
| sp\|Q96KB5\|TOPK\_HUMAN | Q96KB5 | TOPK\_HUMAN | PBK | Lymphokine-activated killer T-cell-originated protein kinase | 322 | 36.085 | 4.415755e+04 | 2.001930e+06 | 442217.06 | 501470.78 | 1058241.8 |  | 34.47 | 37.6 | 21.7 |  |
| sp\|Q5T749\|KPRP\_HUMAN | Q5T749 | KPRP\_HUMAN | KPRP | Keratinocyte proline-rich protein | 579 | 64.136 | 3.117192e+04 | 2.511786e+06 | 441875.94 |  | 973688.5 | 1096221.2 | 9.67 |  | 21.9 | 11.9 |
| sp\|Q14008\|CKAP5\_HUMAN | Q14008 | CKAP5\_HUMAN | CKAP5 | Cytoskeleton-associated protein 5 | 2032 | 225.495 | 7.130505e+03 | 2.020107e+06 | 440436.8 | 1307456.4 | 211179.44 | 61034.43 | 17.57 | 44.8 | 18.1 | 8.4 |
| sp\|Q5TFE4\|NT5D1\_HUMAN | Q5TFE4 | NT5D1\_HUMAN | NT5DC1 | 5'-nucleotidase domain-containing protein 1 | 455 | 51.845 | 2.074563e+04 | 1.351297e+06 | 440150.34 | 911146.5 |  |  | 15.82 | 60.9 |  |  |
| sp\|Q9UKE5\|TNIK\_HUMAN | Q9UKE5 | TNIK\_HUMAN | TNIK | TRAF2 and NCK-interacting protein kinase | 1360 | 154.943 | 5.186006e+03 | 1.009537e+06 | 438960.94 | 406118.4 | 164457.31 |  | 12.65 | 35.1 | 21.0 |  |
| sp\|Q9UNW1\|MINP1\_HUMAN | Q9UNW1 | MINP1\_HUMAN | MINPP1 | Multiple inositol polyphosphate phosphatase 1 | 487 | 55.051 | 5.815374e+04 | 4.022166e+06 | 438228.1 | 389724.56 | 2463761.8 | 730452.0 | 31.62 | 44.6 | 37.0 | 16.6 |
| sp\|Q93100\|KPBB\_HUMAN | Q93100 | KPBB\_HUMAN | PHKB | Phosphorylase b kinase regulatory subunit beta | 1093 | 124.884 | 6.502510e+03 | 1.020246e+06 | 438139.56 | 582106.56 |  |  | 29.55 | 32.0 |  |  |
| sp\|Q9H8Y8\|GORS2\_HUMAN | Q9H8Y8 | GORS2\_HUMAN | GORASP2 | Golgi reassembly-stacking protein 2 | 452 | 47.145 | 1.694890e+04 | 1.003909e+06 | 436411.06 | 202212.6 | 365285.16 |  | 18.58 | 24.3 | 21.2 |  |
| sp\|Q8NBK3\|SUMF1\_HUMAN | Q8NBK3 | SUMF1\_HUMAN | SUMF1 | Formylglycine-generating enzyme | 374 | 40.556 | 1.972026e+04 | 1.004812e+06 | 435964.34 |  | 513344.88 | 55503.168 | 7.22 |  | 27.5 | 21.1 |
| sp\|Q2TAZ0\|ATG2A\_HUMAN | Q2TAZ0 | ATG2A\_HUMAN | ATG2A | Autophagy-related protein 2 homolog A | 1938 | 212.860 | 9.897256e+03 | 2.646830e+06 | 435582.78 | 851637.9 | 365688.8 | 993920.4 | 26.78 | 35.8 | 14.2 | 10.3 |
| sp\|Q6IBS0\|TWF2\_HUMAN | Q6IBS0 | TWF2\_HUMAN | TWF2 | Twinfilin-2 | 349 | 39.548 | 3.191272e+04 | 1.585643e+06 | 435216.44 | 1085922.5 | 64504.418 |  | 42.98 | 64.5 | 22.1 |  |
| sp\|Q9UIQ6\|LCAP\_HUMAN | Q9UIQ6 | LCAP\_HUMAN | LNPEP | Leucyl-cystinyl aminopeptidase | 1025 | 117.349 | 9.772448e+03 | 1.440787e+06 | 434894.06 | 143825.72 | 215797.05 | 646270.6 | 20.0 | 22.3 | 21.1 | 15.0 |
| sp\|Q96EM0\|T3HPD\_HUMAN | Q96EM0 | T3HPD\_HUMAN | L3HYPDH | Trans-3-hydroxy-L-proline dehydratase | 354 | 38.138 | 1.279653e+04 | 6.131507e+05 | 434706.53 | 178444.12 |  |  | 23.73 | 54.2 |  |  |
| sp\|Q8TC07\|TBC15\_HUMAN | Q8TC07 | TBC15\_HUMAN | TBC1D15 | TBC1 domain family member 15 | 691 | 79.491 | 1.216674e+04 | 1.215093e+06 | 434632.1 | 725234.7 | 55225.992 |  | 24.17 | 34.9 | 15.5 |  |
| sp\|Q9UMY4\|SNX12\_HUMAN | Q9UMY4 | SNX12\_HUMAN | SNX12 | Sorting nexin-12 | 162 | 18.885 | 9.945405e+04 | 2.359699e+06 | 433932.5 | 1363909.5 | 376450.9 | 185406.12 | 40.12 | 58.0 | 35.8 | 26.5 |
| sp\|Q8IWT0\|ARCH\_HUMAN | Q8IWT0 | ARCH\_HUMAN | ZBTB8OS | Protein archease | 167 | 19.491 | 2.624684e+04 | 6.427295e+05 | 433444.66 | 209284.83 |  |  | 20.36 | 19.2 |  |  |
| sp\|Q9NZJ9\|NUDT4\_HUMAN | Q9NZJ9 | NUDT4\_HUMAN | NUDT4 | Diphosphoinositol polyphosphate phosphohydrolase 2 | 180 | 20.306 | 6.431587e+04 | 1.640815e+06 | 432718.44 | 1208096.9 |  |  | 26.67 | 51.7 |  |  |
| sp\|Q9NRQ2\|PLS4\_HUMAN | Q9NRQ2 | PLS4\_HUMAN | PLSCR4 | Phospholipid scramblase 4 | 329 | 37.005 | 1.148194e+05 | 5.338178e+06 | 432193.28 | 83037.85 | 800598.94 | 4022347.5 | 17.93 | 28.9 | 19.8 | 14.6 |
| sp\|P28062\|PSB8\_HUMAN | P28062 | PSB8\_HUMAN | PSMB8 | Proteasome subunit beta type-8 | 276 | 30.354 | 2.349869e+04 | 8.961418e+05 | 432078.8 | 420851.62 | 43211.43 |  | 26.45 | 45.7 | 35.5 |  |
| sp\|Q12857\|NFIA\_HUMAN | Q12857 | NFIA\_HUMAN | NFIA | Nuclear factor 1 A-type | 509 | 55.944 | 2.624515e+04 | 1.844674e+06 | 430397.44 | 1047586.94 | 267310.6 | 99379.32 | 22.99 | 27.5 | 20.4 | 7.7 |
| sp\|Q9NQC3\|RTN4\_HUMAN | Q9NQC3 | RTN4\_HUMAN | RTN4 | Reticulon-4 | 1192 | 129.931 | 5.844840e+03 | 9.541191e+05 | 429870.25 | 214267.75 | 73409.23 | 236571.88 | 7.47 | 29.4 | 9.0 | 6.5 |
| sp\|O60784\|TOM1\_HUMAN | O60784 | TOM1\_HUMAN | TOM1 | Target of Myb1 membrane trafficking protein | 492 | 53.818 | 2.230067e+04 | 1.507866e+06 | 428720.0 | 536360.3 | 398232.75 | 144552.48 | 28.86 | 50.8 | 23.0 | 6.9 |
| sp\|Q7KZF4\|SND1\_HUMAN | Q7KZF4 | SND1\_HUMAN | SND1 | Staphylococcal nuclease domain-containing protein 1 | 910 | 101.997 | 7.230565e+03 | 9.265669e+05 | 428719.06 | 378129.7 |  | 119718.164 | 32.86 | 40.9 |  | 11.6 |
| sp\|P31327\|CPSM\_HUMAN | P31327 | CPSM\_HUMAN | CPS1 | Carbamoyl-phosphate synthase [ammonia], mitochondrial | 1500 | 164.939 | 3.912750e+03 | 8.108166e+05 | 428683.06 | 57091.113 | 185343.02 | 139699.44 | 6.13 | 29.9 | 15.9 | 11.6 |
| sp\|Q0JRZ9\|FCHO2\_HUMAN | Q0JRZ9 | FCHO2\_HUMAN | FCHO2 | F-BAR domain only protein 2 | 810 | 88.924 | 1.162320e+04 | 1.298559e+06 | 425247.97 | 631163.44 | 242147.67 |  | 32.96 | 44.9 | 20.5 |  |
| sp\|Q16512\|PKN1\_HUMAN | Q16512 | PKN1\_HUMAN | PKN1 | Serine/threonine-protein kinase N1 | 942 | 103.932 | 1.371011e+04 | 1.790223e+06 | 424728.8 | 1260702.9 | 104791.734 |  | 25.69 | 28.8 | 22.8 |  |
| sp\|Q9Y4P1\|ATG4B\_HUMAN | Q9Y4P1 | ATG4B\_HUMAN | ATG4B | Cysteine protease ATG4B | 393 | 44.294 | 2.773430e+04 | 1.543403e+06 | 424024.06 | 1119378.5 |  |  | 38.68 | 53.2 |  |  |
| sp\|O75044\|SRGP2\_HUMAN | O75044 | SRGP2\_HUMAN | SRGAP2 | SLIT-ROBO Rho GTPase-activating protein 2 | 1071 | 120.871 | 6.208142e+03 | 9.427594e+05 | 423952.12 | 321689.03 | 125815.86 | 71302.44 | 19.05 | 29.4 | 13.4 | 6.8 |
| sp\|P05161\|ISG15\_HUMAN | P05161 | ISG15\_HUMAN | ISG15 | Ubiquitin-like protein ISG15 | 165 | 17.888 | 9.196192e+04 | 2.066745e+06 | 423298.5 | 950780.25 | 526625.56 | 166041.1 | 36.97 | 52.1 | 41.2 | 18.2 |
| sp\|Q9UGI8\|TES\_HUMAN | Q9UGI8 | TES\_HUMAN | TES | Testin | 421 | 47.996 | 1.640161e+04 | 9.890282e+05 | 422967.62 | 438760.88 | 127299.71 |  | 31.59 | 43.2 | 35.6 |  |
| sp\|Q96C11\|FGGY\_HUMAN | Q96C11 | FGGY\_HUMAN | FGGY | FGGY carbohydrate kinase domain-containing protein | 551 | 59.993 | 1.014194e+04 | 7.644316e+05 | 420606.62 | 343825.0 |  |  | 22.87 | 33.4 |  |  |
| sp\|Q9BVC4\|LST8\_HUMAN | Q9BVC4 | LST8\_HUMAN | MLST8 | Target of rapamycin complex subunit LST8 | 326 | 35.876 | 1.699999e+04 | 7.662490e+05 | 419994.2 | 213387.19 | 132867.61 | 0.0 | 19.02 | 26.1 | 22.4 | 10.7 |
| sp\|P15104\|GLNA\_HUMAN | P15104 | GLNA\_HUMAN | GLUL | Glutamine synthetase | 373 | 42.064 | 4.926430e+04 | 2.603515e+06 | 419315.06 | 1576767.1 | 364103.84 | 243328.72 | 27.61 | 37.3 | 26.8 | 17.7 |
| sp\|P61626\|LYSC\_HUMAN | P61626 | LYSC\_HUMAN | LYZ | Lysozyme C | 148 | 16.537 | 1.524165e+05 | 3.166693e+06 | 417880.62 |  | 1958768.5 | 790043.9 | 24.32 |  | 37.2 | 30.4 |
| sp\|Q9P0L0\|VAPA\_HUMAN | Q9P0L0 | VAPA\_HUMAN | VAPA | Vesicle-associated membrane protein-associated protein A | 249 | 27.893 | 1.824904e+05 | 6.395173e+06 | 417721.0 | 1422611.0 | 2392799.8 | 2162041.2 | 24.9 | 58.6 | 49.8 | 36.1 |
| sp\|Q9NRN7\|ADPPT\_HUMAN | Q9NRN7 | ADPPT\_HUMAN | AASDHPPT | L-aminoadipate-semialdehyde dehydrogenase-phosphopantetheinyl transferase | 309 | 35.776 | 7.141200e+04 | 3.209816e+06 | 416524.97 | 2689445.5 | 103845.26 |  | 20.71 | 44.3 | 16.5 |  |
| sp\|Q9Y5K8\|VATD\_HUMAN | Q9Y5K8 | VATD\_HUMAN | ATP6V1D | V-type proton ATPase subunit D | 247 | 28.263 | 8.085761e+04 | 2.871153e+06 | 414694.75 | 1920649.8 | 206426.47 | 329381.75 | 26.72 | 57.5 | 28.7 | 6.9 |
| sp\|P13807\|GYS1\_HUMAN | P13807 | GYS1\_HUMAN | GYS1 | Glycogen [starch] synthase, muscle | 737 | 83.786 | 1.147360e+04 | 1.207781e+06 | 412859.7 | 717224.1 | 65319.86 | 12377.661 | 34.19 | 40.7 | 12.5 | 7.1 |
| sp\|O43264\|ZW10\_HUMAN | O43264 | ZW10\_HUMAN | ZW10 | Centromere/kinetochore protein zw10 homolog | 779 | 88.829 | 2.429950e+04 | 2.711873e+06 | 412091.8 | 1474743.5 | 668703.3 | 156334.22 | 34.53 | 40.9 | 26.3 | 14.8 |
| sp\|P39687\|AN32A\_HUMAN | P39687 | AN32A\_HUMAN | ANP32A | Acidic leucine-rich nuclear phosphoprotein 32 family member A | 249 | 28.585 | 3.971873e+04 | 1.426431e+06 | 411395.75 | 946925.94 | 68109.016 |  | 42.97 | 41.0 | 25.3 |  |
| sp\|O75368\|SH3L1\_HUMAN | O75368 | SH3L1\_HUMAN | SH3BGRL | Adapter SH3BGRL | 114 | 12.774 | 5.799916e+05 | 9.308202e+06 | 410618.12 | 7231265.5 | 1298057.6 | 368260.44 | 25.44 | 85.1 | 61.4 | 28.9 |
| sp\|Q8IU68\|TMC8\_HUMAN | Q8IU68 | TMC8\_HUMAN | TMC8 | Transmembrane channel-like protein 8 | 726 | 81.641 | 3.563135e+04 | 3.654751e+06 | 410565.78 |  | 340707.88 | 2903477.2 | 10.33 |  | 19.6 | 16.3 |
| sp\|P29353\|SHC1\_HUMAN | P29353 | SHC1\_HUMAN | SHC1 | SHC-transforming protein 1 | 583 | 62.822 | 1.432554e+04 | 1.130681e+06 | 409560.53 | 427898.47 | 293221.7 |  | 21.27 | 55.2 | 25.2 |  |
| sp\|O94888\|UBXN7\_HUMAN | O94888 | UBXN7\_HUMAN | UBXN7 | UBX domain-containing protein 7 | 489 | 54.862 | 1.247143e+04 | 8.596173e+05 | 409289.97 | 401405.84 | 48921.47 |  | 18.4 | 26.2 | 10.8 |  |
| sp\|Q8NDH3\|PEPL1\_HUMAN | Q8NDH3 | PEPL1\_HUMAN | NPEPL1 | Probable aminopeptidase NPEPL1 | 523 | 55.861 | 1.808166e+04 | 1.269008e+06 | 406159.0 | 829574.56 | 21403.477 | 11870.565 | 15.49 | 27.9 | 10.1 | 4.0 |
| sp\|Q14151\|SAFB2\_HUMAN | Q14151 | SAFB2\_HUMAN | SAFB2 | Scaffold attachment factor B2 | 953 | 104.830 | 3.076808e+03 | 4.052315e+05 | 405231.47 |  |  |  | 6.82 |  |  |  |
| sp\|P50748\|KNTC1\_HUMAN | P50748 | KNTC1\_HUMAN | KNTC1 | Kinetochore-associated protein 1 | 2209 | 250.749 | 2.324055e+03 | 7.321544e+05 | 404180.3 | 327974.1 |  |  | 7.97 | 26.8 |  |  |
| sp\|P28676\|GRAN\_HUMAN | P28676 | GRAN\_HUMAN | GCA | Grancalcin | 217 | 24.010 | 5.725194e+04 | 1.727028e+06 | 403088.44 | 1323940.0 |  |  | 25.35 | 52.1 |  |  |
| sp\|Q86TX2\|ACOT1\_HUMAN | Q86TX2 | ACOT1\_HUMAN | ACOT1 | Acyl-coenzyme A thioesterase 1 | 421 | 46.277 | 9.192565e+03 | 5.344648e+05 | 401808.6 | 132656.16 |  |  | 21.38 | 67.5 |  |  |
| sp\|Q5T1H1\|EYS\_HUMAN | Q5T1H1 | EYS\_HUMAN | EYS | Protein eyes shut homolog | 3165 | 348.150 | 9.166912e+02 | 4.009652e+05 | 400965.16 |  |  |  | 1.74 |  |  |  |
| sp\|P09105\|HBAT\_HUMAN | P09105 | HBAT\_HUMAN | HBQ1 | Hemoglobin subunit theta-1 | 142 | 15.508 | 6.329606e+05 | 1.233246e+07 | 400533.8 | 8974711.0 | 2854322.0 | 102890.28 | 70.42 | 100.0 | 35.9 | 22.5 |
| sp\|P16949\|STMN1\_HUMAN | P16949 | STMN1\_HUMAN | STMN1 | Stathmin | 149 | 17.303 | 5.917699e+04 | 1.286446e+06 | 398927.44 | 679166.0 | 208352.1 |  | 30.87 | 54.4 | 44.3 |  |
| sp\|Q14790\|CASP8\_HUMAN | Q14790 | CASP8\_HUMAN | CASP8 | Caspase-8 | 479 | 55.391 | 2.278827e+04 | 1.585870e+06 | 396249.38 | 1149189.1 | 33089.344 | 7342.3384 | 38.41 | 41.5 | 11.3 | 8.4 |
| sp\|Q9Y678\|COPG1\_HUMAN | Q9Y678 | COPG1\_HUMAN | COPG1 | Coatomer subunit gamma-1 | 874 | 97.718 | 5.128009e+03 | 6.295649e+05 | 395979.78 | 214385.06 | 19200.023 |  | 17.28 | 40.3 | 16.4 |  |
| sp\|Q96BW5\|PTER\_HUMAN | Q96BW5 | PTER\_HUMAN | PTER | Phosphotriesterase-related protein | 349 | 39.018 | 2.062734e+04 | 1.011173e+06 | 395095.84 | 616076.9 |  |  | 40.4 | 48.7 |  |  |
| sp\|Q6DN90\|IQEC1\_HUMAN | Q6DN90 | IQEC1\_HUMAN | IQSEC1 | IQ motif and SEC7 domain-containing protein 1 | 963 | 108.314 | 1.714013e+04 | 2.332469e+06 | 394921.03 | 1565359.5 | 136027.5 | 236160.94 | 26.58 | 51.0 | 36.4 | 14.3 |
| sp\|Q96IV0\|NGLY1\_HUMAN | Q96IV0 | NGLY1\_HUMAN | NGLY1 | Peptide-N(4)-(N-acetyl-beta-glucosaminyl)asparagine amidase | 654 | 74.390 | 1.681499e+04 | 1.571551e+06 | 394042.72 | 1177507.8 |  |  | 28.44 | 43.0 |  |  |
| sp\|P20930\|FILA\_HUMAN | P20930 | FILA\_HUMAN | FLG | Filaggrin | 4061 | 435.170 | 8.690256e+03 | 4.751259e+06 | 392436.88 | 1296585.6 | 823857.6 | 2238378.5 | 16.92 | 56.4 | 40.9 | 34.4 |
| sp\|Q15208\|STK38\_HUMAN | Q15208 | STK38\_HUMAN | STK38 | Serine/threonine-protein kinase 38 | 465 | 54.190 | 1.411502e+04 | 9.609876e+05 | 391144.88 | 569842.75 |  |  | 28.39 | 58.1 |  |  |
| sp\|P49959\|MRE11\_HUMAN | P49959 | MRE11\_HUMAN | MRE11 | Double-strand break repair protein MRE11 | 708 | 80.593 | 1.582813e+04 | 1.602670e+06 | 389818.22 | 555271.6 | 559993.44 | 97586.25 | 24.86 | 45.2 | 23.4 | 6.2 |
| sp\|Q15796\|SMAD2\_HUMAN | Q15796 | SMAD2\_HUMAN | SMAD2 | Mothers against decapentaplegic homolog 2 | 467 | 52.306 | 2.058614e+04 | 1.352831e+06 | 389781.66 | 919641.2 | 43408.07 |  | 18.63 | 39.0 | 10.1 |  |
| sp\|Q9Y5A7\|NUB1\_HUMAN | Q9Y5A7 | NUB1\_HUMAN | NUB1 | NEDD8 ultimate buster 1 | 615 | 70.538 | 6.474094e+03 | 5.737455e+05 | 389093.94 | 184651.58 |  |  | 13.17 | 23.7 |  |  |
| sp\|Q86YT6\|MIB1\_HUMAN | Q86YT6 | MIB1\_HUMAN | MIB1 | E3 ubiquitin-protein ligase MIB1 | 1006 | 110.136 | 1.529448e+04 | 2.116319e+06 | 387795.97 | 1334123.6 | 297827.1 | 96572.33 | 29.52 | 46.9 | 28.4 | 22.4 |
| sp\|P31941\|ABC3A\_HUMAN | P31941 | ABC3A\_HUMAN | APOBEC3A | DNA dC->dU-editing enzyme APOBEC-3A | 199 | 21.890 | 1.406919e+04 | 3.869297e+05 | 386929.66 |  |  |  | 22.61 |  |  |  |
| sp\|Q9H477\|RBSK\_HUMAN | Q9H477 | RBSK\_HUMAN | RBKS | Ribokinase | 322 | 34.143 | 2.454110e+04 | 1.052720e+06 | 386680.88 | 666039.06 |  |  | 26.09 | 37.9 |  |  |
| sp\|O75317\|UBP12\_HUMAN | O75317 | UBP12\_HUMAN | USP12 | Ubiquitin carboxyl-terminal hydrolase 12 | 370 | 42.858 | 1.793348e+04 | 9.656364e+05 | 385411.6 | 530950.0 | 21490.604 | 27784.166 | 11.89 | 42.2 | 17.0 | 9.7 |
| sp\|Q9Y2S2\|CRYL1\_HUMAN | Q9Y2S2 | CRYL1\_HUMAN | CRYL1 | Lambda-crystallin homolog | 319 | 35.419 | 3.656532e+04 | 1.627132e+06 | 385165.56 | 1241966.5 |  |  | 23.82 | 52.7 |  |  |
| sp\|Q8N4C8\|MINK1\_HUMAN | Q8N4C8 | MINK1\_HUMAN | MINK1 | Misshapen-like kinase 1 | 1332 | 149.822 | 6.318244e+03 | 1.189294e+06 | 384606.94 | 525328.5 | 242699.22 | 36659.08 | 18.09 | 38.7 | 21.2 | 11.3 |
| sp\|P15090\|FABP4\_HUMAN | P15090 | FABP4\_HUMAN | FABP4 | Fatty acid-binding protein, adipocyte | 132 | 14.719 | 4.536535e+04 | 8.389183e+05 | 383993.56 |  | 17938.57 | 436986.2 | 27.27 |  | 15.2 | 15.9 |
| sp\|Q9BUL8\|PDC10\_HUMAN | Q9BUL8 | PDC10\_HUMAN | PDCD10 | Programmed cell death protein 10 | 212 | 24.702 | 6.086483e+04 | 1.888929e+06 | 383594.1 | 1280948.5 | 224386.58 |  | 19.81 | 22.2 | 30.2 |  |
| sp\|A6NDU8\|RIMC1\_HUMAN | A6NDU8 | RIMC1\_HUMAN | RIMOC1 | RAB7A-interacting MON1-CCZ1 complex subunit 1 | 294 | 33.620 | 2.726237e+04 | 1.151539e+06 | 383357.72 | 531938.1 | 172956.05 | 63286.656 | 27.89 | 47.6 | 10.9 | 14.6 |
| sp\|Q13492\|PICAL\_HUMAN | Q13492 | PICAL\_HUMAN | PICALM | Phosphatidylinositol-binding clathrin assembly protein | 652 | 70.755 | 3.056590e+04 | 2.717137e+06 | 382410.84 | 1985992.4 | 226731.44 | 122002.37 | 24.39 | 39.3 | 23.6 | 12.9 |
| sp\|P78560\|CRADD\_HUMAN | P78560 | CRADD\_HUMAN | CRADD | Death domain-containing protein CRADD | 199 | 22.745 | 5.534484e+04 | 1.581540e+06 | 382018.56 | 994685.3 | 159674.84 | 45161.52 | 54.27 | 69.3 | 27.1 | 7.0 |
| sp\|Q8IWV7\|UBR1\_HUMAN | Q8IWV7 | UBR1\_HUMAN | UBR1 | E3 ubiquitin-protein ligase UBR1 | 1749 | 200.211 | 2.573123e+03 | 6.472404e+05 | 380743.44 | 266496.94 |  |  | 14.81 | 38.1 |  |  |
| sp\|Q5W0V3\|FHI2A\_HUMAN | Q5W0V3 | FHI2A\_HUMAN | FHIP2A | FHF complex subunit HOOK interacting protein 2A | 765 | 86.558 | 7.063006e+03 | 7.680932e+05 | 380497.34 | 257798.58 | 92198.375 | 37598.926 | 9.8 | 42.5 | 16.3 | 12.0 |
| sp\|P42226\|STAT6\_HUMAN | P42226 | STAT6\_HUMAN | STAT6 | Signal transducer and activator of transcription 6 | 847 | 94.135 | 6.858035e+03 | 8.110880e+05 | 380296.06 | 430791.94 |  |  | 23.02 | 31.5 |  |  |
| sp\|Q8TBN0\|R3GEF\_HUMAN | Q8TBN0 | R3GEF\_HUMAN | RAB3IL1 | Guanine nucleotide exchange factor for Rab-3A | 382 | 42.637 | 1.924485e+04 | 1.030904e+06 | 379879.56 | 388855.12 | 207462.67 | 54707.004 | 37.43 | 59.7 | 36.9 | 22.8 |
| sp\|Q15363\|TMED2\_HUMAN | Q15363 | TMED2\_HUMAN | TMED2 | Transmembrane emp24 domain-containing protein 2 | 201 | 22.761 | 2.064398e+05 | 5.903395e+06 | 378388.3 | 1289481.8 | 3393341.8 | 842183.25 | 16.42 | 55.2 | 31.3 | 18.9 |
| sp\|P49257\|LMAN1\_HUMAN | P49257 | LMAN1\_HUMAN | LMAN1 | Protein ERGIC-53 | 510 | 57.549 | 7.571235e+04 | 5.474212e+06 | 378158.16 | 192818.0 | 1928311.8 | 2974924.5 | 21.37 | 27.3 | 21.8 | 21.4 |
| sp\|P02794\|FRIH\_HUMAN | P02794 | FRIH\_HUMAN | FTH1 | Ferritin heavy chain | 183 | 21.226 | 1.327032e+05 | 3.538888e+06 | 377156.97 | 2844175.0 | 317555.94 |  | 32.24 | 63.4 | 41.5 |  |
| sp\|P55039\|DRG2\_HUMAN | P55039 | DRG2\_HUMAN | DRG2 | Developmentally-regulated GTP-binding protein 2 | 364 | 40.746 | 1.837771e+04 | 9.407917e+05 | 376545.38 | 513284.2 | 39967.85 | 10994.265 | 34.62 | 42.6 | 23.6 | 8.2 |
| sp\|Q13838\|DX39B\_HUMAN | Q13838 | DX39B\_HUMAN | DDX39B | Spliceosome RNA helicase DDX39B | 428 | 48.991 | 1.199128e+04 | 7.380725e+05 | 376517.03 | 361555.44 |  |  | 23.36 | 43.7 |  |  |
| sp\|P15735\|PHKG2\_HUMAN | P15735 | PHKG2\_HUMAN | PHKG2 | Phosphorylase b kinase gamma catalytic chain, liver/testis isoform | 406 | 46.442 | 1.179965e+04 | 6.884893e+05 | 374913.9 | 260752.88 | 52822.57 |  | 19.95 | 39.7 | 20.2 |  |
| sp\|Q12933\|TRAF2\_HUMAN | Q12933 | TRAF2\_HUMAN | TRAF2 | TNF receptor-associated factor 2 | 501 | 55.859 | 1.913269e+04 | 1.342723e+06 | 374769.12 | 821088.44 | 113309.266 | 33556.113 | 21.56 | 49.9 | 38.5 | 11.4 |
| sp\|P25311\|ZA2G\_HUMAN | P25311 | ZA2G\_HUMAN | AZGP1 | Zinc-alpha-2-glycoprotein | 298 | 34.259 | 9.677507e+04 | 4.165387e+06 | 373516.84 | 367068.8 | 1924700.0 | 1500100.9 | 32.21 | 39.3 | 43.6 | 23.8 |
| sp\|P61970\|NTF2\_HUMAN | P61970 | NTF2\_HUMAN | NUTF2 | Nuclear transport factor 2 | 127 | 14.478 | 5.688415e+05 | 1.034706e+07 | 372745.06 | 9974316.0 |  |  | 61.42 | 69.3 |  |  |
| sp\|Q9Y450\|HBS1L\_HUMAN | Q9Y450 | HBS1L\_HUMAN | HBS1L | HBS1-like protein | 684 | 75.473 | 2.564406e+04 | 2.431619e+06 | 372168.38 | 1764471.9 | 178692.08 | 116287.086 | 37.28 | 56.4 | 23.1 | 13.5 |
| sp\|P01871\|IGHM\_HUMAN | P01871 | IGHM\_HUMAN | IGHM | Immunoglobulin heavy constant mu | 474 | 51.924 | 9.950555e+04 | 6.491314e+06 | 372093.78 | 425246.44 | 4439046.0 | 1254927.5 | 15.82 | 40.0 | 34.2 | 27.8 |
| sp\|Q9UHW9\|S12A6\_HUMAN | Q9UHW9 | S12A6\_HUMAN | SLC12A6 | Solute carrier family 12 member 6 | 1150 | 127.617 | 1.763402e+04 | 2.827334e+06 | 371878.7 | 317486.56 | 504353.06 | 1633615.8 | 19.74 | 32.1 | 28.5 | 20.1 |
| sp\|Q8N573\|OXR1\_HUMAN | Q8N573 | OXR1\_HUMAN | OXR1 | Oxidation resistance protein 1 | 874 | 97.970 | 5.082880e+03 | 6.256337e+05 | 371812.7 | 253821.02 |  |  | 8.92 | 34.2 |  |  |
| sp\|Q16610\|ECM1\_HUMAN | Q16610 | ECM1\_HUMAN | ECM1 | Extracellular matrix protein 1 | 540 | 60.674 | 2.196704e+04 | 1.674524e+06 | 371677.75 | 156355.16 | 471578.7 | 674912.4 | 16.67 | 44.1 | 25.6 | 19.8 |
| sp\|Q7Z403\|TMC6\_HUMAN | Q7Z403 | TMC6\_HUMAN | TMC6 | Transmembrane channel-like protein 6 | 805 | 90.045 | 2.841024e+04 | 3.214043e+06 | 370818.3 |  | 523280.34 | 2319944.0 | 12.8 |  | 16.4 | 15.5 |
| sp\|Q13164\|MK07\_HUMAN | Q13164 | MK07\_HUMAN | MAPK7 | Mitogen-activated protein kinase 7 | 816 | 88.386 | 7.992846e+03 | 8.875689e+05 | 369233.78 | 417502.44 | 89538.47 | 11294.234 | 22.06 | 25.0 | 17.3 | 4.7 |
| sp\|Q5T2T1\|MPP7\_HUMAN | Q5T2T1 | MPP7\_HUMAN | MPP7 | MAGUK p55 subfamily member 7 | 576 | 65.524 | 1.646406e+04 | 1.355359e+06 | 367116.3 | 271076.47 | 614152.75 | 103013.625 | 40.62 | 40.3 | 32.8 | 16.8 |
| sp\|Q15831\|STK11\_HUMAN | Q15831 | STK11\_HUMAN | STK11 | Serine/threonine-protein kinase STK11 | 433 | 48.636 | 1.955091e+04 | 1.194653e+06 | 366763.94 | 559160.4 | 216225.0 | 52503.973 | 35.1 | 33.5 | 19.6 | 6.9 |
| sp\|P08708\|RS17\_HUMAN | P08708 | RS17\_HUMAN | RPS17 | Small ribosomal subunit protein eS17 | 135 | 15.550 | 1.346778e+05 | 2.631138e+06 | 365386.3 | 2072144.2 | 193607.7 |  | 47.41 | 77.8 | 29.6 |  |
| sp\|Q9NPB8\|GPCP1\_HUMAN | Q9NPB8 | GPCP1\_HUMAN | GPCPD1 | Glycerophosphocholine phosphodiesterase GPCPD1 | 672 | 76.035 | 1.691897e+04 | 1.616235e+06 | 363342.88 | 1233231.0 | 19661.174 |  | 26.79 | 39.3 | 17.0 |  |
| sp\|O15294\|OGT1\_HUMAN | O15294 | OGT1\_HUMAN | OGT | UDP-N-acetylglucosamine--peptide N-acetylglucosaminyltransferase 110 kDa subunit | 1046 | 116.925 | 4.757943e+03 | 6.989463e+05 | 363080.0 | 288656.97 | 47209.312 |  | 22.28 | 37.9 | 15.4 |  |
| sp\|P49840\|GSK3A\_HUMAN | P49840 | GSK3A\_HUMAN | GSK3A | Glycogen synthase kinase-3 alpha | 483 | 50.981 | 1.755723e+04 | 1.124557e+06 | 362146.62 | 631232.6 | 92715.516 | 38462.594 | 34.58 | 40.2 | 19.0 | 8.9 |
| sp\|Q9Y6M4\|KC1G3\_HUMAN | Q9Y6M4 | KC1G3\_HUMAN | CSNK1G3 | Casein kinase I isoform gamma-3 | 447 | 51.389 | 2.909465e+04 | 1.878454e+06 | 360946.47 | 465559.66 | 374854.2 | 677093.5 | 27.52 | 59.7 | 32.2 | 17.0 |
| sp\|Q9P1Z2\|CACO1\_HUMAN | Q9P1Z2 | CACO1\_HUMAN | CALCOCO1 | Calcium-binding and coiled-coil domain-containing protein 1 | 691 | 77.336 | 1.659656e+04 | 1.612564e+06 | 360536.56 | 993776.8 | 176568.72 | 81681.79 | 30.97 | 37.2 | 24.9 | 16.9 |
| sp\|Q8NG06\|TRI58\_HUMAN | Q8NG06 | TRI58\_HUMAN | TRIM58 | E3 ubiquitin-protein ligase TRIM58 | 486 | 54.766 | 1.737911e+04 | 1.195792e+06 | 360221.66 | 626936.8 | 208633.78 |  | 31.28 | 52.3 | 22.4 |  |
| sp\|O14979\|HNRDL\_HUMAN | O14979 | HNRDL\_HUMAN | HNRNPDL | Heterogeneous nuclear ribonucleoprotein D-like | 420 | 46.200 | 6.201739e+03 | 3.599752e+05 | 359975.2 |  |  |  | 8.1 |  |  |  |
| sp\|Q9BY32\|ITPA\_HUMAN | Q9BY32 | ITPA\_HUMAN | ITPA | Inosine triphosphate pyrophosphatase | 194 | 21.446 | 3.082306e+04 | 8.304993e+05 | 359056.5 | 453273.28 |  | 18169.484 | 28.35 | 64.4 |  | 9.3 |
| sp\|Q9H425\|CA198\_HUMAN | Q9H425 | CA198\_HUMAN | C1orf198 | Uncharacterized protein C1orf198 | 327 | 36.346 | 1.231177e+05 | 5.622044e+06 | 358945.44 | 703952.3 | 4260648.0 | 298498.44 | 31.5 | 72.2 | 52.0 | 48.3 |
| sp\|Q9NYL2\|M3K20\_HUMAN | Q9NYL2 | M3K20\_HUMAN | MAP3K20 | Mitogen-activated protein kinase kinase kinase 20 | 800 | 91.155 | 6.027894e+03 | 6.903403e+05 | 358531.56 | 269207.22 | 62601.53 |  | 7.25 | 36.2 | 10.0 |  |
| sp\|Q9UK59\|DBR1\_HUMAN | Q9UK59 | DBR1\_HUMAN | DBR1 | Lariat debranching enzyme | 544 | 61.555 | 9.694016e+03 | 7.496943e+05 | 358233.66 | 312673.2 | 52094.418 | 26693.031 | 19.85 | 23.9 | 14.3 | 8.6 |
| sp\|Q8NBX0\|SCPDL\_HUMAN | Q8NBX0 | SCPDL\_HUMAN | SCCPDH | Saccharopine dehydrogenase-like oxidoreductase | 429 | 47.151 | 1.296566e+04 | 7.680732e+05 | 357575.3 | 167478.94 | 243018.97 |  | 14.69 | 45.0 | 14.9 |  |
| sp\|Q9Y484\|WIPI4\_HUMAN | Q9Y484 | WIPI4\_HUMAN | WDR45 | WD repeat domain phosphoinositide-interacting protein 4 | 360 | 39.868 | 2.103213e+04 | 1.053477e+06 | 356690.34 | 346624.9 | 350161.38 |  | 22.78 | 33.9 | 25.8 |  |
| sp\|Q8NFY9\|KBTB8\_HUMAN | Q8NFY9 | KBTB8\_HUMAN | KBTBD8 | Kelch repeat and BTB domain-containing protein 8 | 601 | 68.823 | 1.317721e+04 | 1.139395e+06 | 356092.0 | 551469.75 | 231832.88 |  | 22.8 | 47.9 | 19.6 |  |
| sp\|Q9UHJ3\|SMBT1\_HUMAN | Q9UHJ3 | SMBT1\_HUMAN | SFMBT1 | Scm-like with four MBT domains protein 1 | 866 | 95.260 | 2.963085e+03 | 3.546270e+05 | 354627.03 |  |  |  | 3.23 |  |  |  |
| sp\|Q3V6T2\|GRDN\_HUMAN | Q3V6T2 | GRDN\_HUMAN | CCDC88A | Girdin | 1871 | 216.042 | 2.101391e+03 | 5.703773e+05 | 354400.75 | 215976.55 |  |  | 14.75 | 28.2 |  |  |
| sp\|P01857\|IGHG1\_HUMAN | P01857 | IGHG1\_HUMAN | IGHG1 | Immunoglobulin heavy constant gamma 1 | 399 | 43.912 | 1.322134e+05 | 7.294169e+06 | 353869.0 | 1037389.1 | 3709764.2 | 2193147.0 | 23.06 | 60.0 | 34.2 | 22.4 |
| sp\|Q92688\|AN32B\_HUMAN | Q92688 | AN32B\_HUMAN | ANP32B | Acidic leucine-rich nuclear phosphoprotein 32 family member B | 251 | 28.788 | 2.810539e+04 | 1.016525e+06 | 351290.0 | 665235.4 |  |  | 34.26 | 35.5 |  |  |
| sp\|P52564\|MP2K6\_HUMAN | P52564 | MP2K6\_HUMAN | MAP2K6 | Dual specificity mitogen-activated protein kinase kinase 6 | 334 | 37.492 | 1.889617e+04 | 8.900812e+05 | 350870.12 | 539211.06 |  |  | 26.95 | 67.4 |  |  |
| sp\|Q0VDG4\|SCRN3\_HUMAN | Q0VDG4 | SCRN3\_HUMAN | SCRN3 | Secernin-3 | 424 | 48.544 | 2.527425e+04 | 1.541456e+06 | 350456.53 | 1105394.4 | 85604.8 |  | 42.22 | 41.5 | 19.6 |  |
| sp\|Q8N6T3\|ARFG1\_HUMAN | Q8N6T3 | ARFG1\_HUMAN | ARFGAP1 | ADP-ribosylation factor GTPase-activating protein 1 | 406 | 44.668 | 8.297356e+04 | 4.656433e+06 | 349437.38 | 968780.9 | 244942.27 | 3093272.8 | 28.57 | 40.1 | 37.2 | 25.1 |
| sp\|Q15223\|NECT1\_HUMAN | Q15223 | NECT1\_HUMAN | NECTIN1 | Nectin-1 | 517 | 57.158 | 2.265024e+04 | 1.626548e+06 | 349425.16 | 206649.31 | 1042263.5 | 28210.453 | 12.38 | 36.9 | 27.7 | 6.4 |
| sp\|Q15750\|TAB1\_HUMAN | Q15750 | TAB1\_HUMAN | TAB1 | TGF-beta-activated kinase 1 and MAP3K7-binding protein 1 | 504 | 54.644 | 2.053475e+04 | 1.409773e+06 | 349412.56 | 908837.25 | 136871.05 | 14651.685 | 52.18 | 50.2 | 22.8 | 6.5 |
| sp\|P49321\|NASP\_HUMAN | P49321 | NASP\_HUMAN | NASP | Nuclear autoantigenic sperm protein | 788 | 85.238 | 1.263900e+04 | 1.353516e+06 | 349018.0 | 920486.75 | 48355.914 | 35654.938 | 38.32 | 47.7 | 20.2 | 17.1 |
| sp\|Q8WW12\|PCNP\_HUMAN | Q8WW12 | PCNP\_HUMAN | PCNP | PEST proteolytic signal-containing nuclear protein | 178 | 18.925 | 1.895527e+05 | 4.506953e+06 | 348541.56 | 3806370.5 | 312699.56 | 39341.61 | 35.96 | 68.5 | 53.9 | 34.8 |
| sp\|P07948\|LYN\_HUMAN | P07948 | LYN\_HUMAN | LYN | Tyrosine-protein kinase Lyn | 512 | 58.574 | 1.563718e+04 | 1.150749e+06 | 346585.97 | 250825.0 | 413729.5 | 139608.1 | 8.59 | 39.5 | 47.9 | 23.2 |
| sp\|O43426\|SYNJ1\_HUMAN | O43426 | SYNJ1\_HUMAN | SYNJ1 | Synaptojanin-1 | 1573 | 173.103 | 4.303904e+03 | 9.360183e+05 | 345308.44 | 329469.62 | 217869.36 | 43370.875 | 30.26 | 50.4 | 19.6 | 11.1 |
| sp\|Q7Z7H5\|TMED4\_HUMAN | Q7Z7H5 | TMED4\_HUMAN | TMED4 | Transmembrane emp24 domain-containing protein 4 | 227 | 25.943 | 8.574999e+04 | 2.794933e+06 | 344343.7 |  | 2450589.5 |  | 4.85 |  | 54.6 |  |
| sp\|Q9H6Z4\|RANB3\_HUMAN | Q9H6Z4 | RANB3\_HUMAN | RANBP3 | Ran-binding protein 3 | 567 | 60.210 | 1.908094e+04 | 1.443396e+06 | 343085.72 | 1025606.94 | 52877.12 | 21826.395 | 21.16 | 49.6 | 19.0 | 15.3 |
| sp\|Q92625\|ANS1A\_HUMAN | Q92625 | ANS1A\_HUMAN | ANKS1A | Ankyrin repeat and SAM domain-containing protein 1A | 1134 | 123.108 | 6.586905e+03 | 1.018790e+06 | 342866.06 | 610226.25 | 65698.0 |  | 15.87 | 40.3 | 16.2 |  |
| sp\|Q13835\|PKP1\_HUMAN | Q13835 | PKP1\_HUMAN | PKP1 | Plakophilin-1 | 747 | 82.861 | 1.285364e+05 | 1.338116e+07 | 342797.78 | 136826.88 | 769628.8 | 12131902.0 | 19.81 | 42.0 | 44.3 | 30.7 |
| sp\|Q9UN86\|G3BP2\_HUMAN | Q9UN86 | G3BP2\_HUMAN | G3BP2 | Ras GTPase-activating protein-binding protein 2 | 482 | 54.121 | 1.532341e+04 | 1.041930e+06 | 341753.16 | 648640.3 | 51536.324 |  | 19.09 | 52.7 | 36.3 |  |
| sp\|O75069\|TMCC2\_HUMAN | O75069 | TMCC2\_HUMAN | TMCC2 | Transmembrane and coiled-coil domains protein 2 | 709 | 77.449 | 2.351592e+05 | 2.288205e+07 | 341197.28 | 451271.5 | 20326756.0 | 1762821.8 | 20.73 | 42.9 | 26.8 | 20.2 |
| sp\|P49902\|5NTC\_HUMAN | P49902 | 5NTC\_HUMAN | NT5C2 | Cytosolic purine 5'-nucleotidase | 561 | 64.970 | 1.892241e+04 | 1.544566e+06 | 340264.8 | 1125957.6 | 78343.55 |  | 18.89 | 23.5 | 15.9 |  |
| sp\|Q92890\|UFD1\_HUMAN | Q92890 | UFD1\_HUMAN | UFD1 | Ubiquitin recognition factor in ER-associated degradation protein 1 | 307 | 34.500 | 3.570224e+04 | 1.547504e+06 | 340185.97 | 1207317.9 |  |  | 17.92 | 35.2 |  |  |
| sp\|Q9UBS8\|RNF14\_HUMAN | Q9UBS8 | RNF14\_HUMAN | RNF14 | E3 ubiquitin-protein ligase RNF14 | 474 | 53.837 | 1.779073e+04 | 1.203350e+06 | 339788.88 | 776159.1 | 87401.555 |  | 25.95 | 34.0 | 27.4 |  |
| sp\|Q9BZI7\|REN3B\_HUMAN | Q9BZI7 | REN3B\_HUMAN | UPF3B | Regulator of nonsense transcripts 3B | 483 | 57.762 | 1.595289e+04 | 1.157707e+06 | 338912.53 | 445932.94 | 273626.1 | 99235.77 | 32.51 | 34.0 | 37.9 | 22.4 |
| sp\|O95786\|RIGI\_HUMAN | O95786 | RIGI\_HUMAN | RIGI | Antiviral innate immune response receptor RIG-I | 925 | 106.600 | 5.282376e+03 | 7.074629e+05 | 338545.84 | 323503.9 | 45413.133 |  | 31.78 | 46.5 | 18.2 |  |
| sp\|P49903\|SPS1\_HUMAN | P49903 | SPS1\_HUMAN | SEPHS1 | Selenide, water dikinase 1 | 392 | 43.120 | 6.242989e+03 | 3.382116e+05 | 338211.56 |  |  |  | 7.4 |  |  |  |
| sp\|P07910\|HNRPC\_HUMAN | P07910 | HNRPC\_HUMAN | HNRNPC | Heterogeneous nuclear ribonucleoproteins C1/C2 | 306 | 33.670 | 2.514348e+04 | 1.063618e+06 | 338072.97 | 640008.0 | 85537.14 |  | 26.47 | 51.0 | 45.1 |  |
| sp\|Q9UKY7\|CDV3\_HUMAN | Q9UKY7 | CDV3\_HUMAN | CDV3 | Protein CDV3 homolog | 258 | 27.335 | 4.626993e+04 | 1.589041e+06 | 337880.2 | 1204457.4 | 46703.152 |  | 56.59 | 80.6 | 23.3 |  |
| sp\|O95816\|BAG2\_HUMAN | O95816 | BAG2\_HUMAN | BAG2 | BAG family molecular chaperone regulator 2 | 211 | 23.772 | 1.202182e+05 | 3.590485e+06 | 337427.7 | 426649.8 | 2542295.5 | 284111.66 | 37.44 | 59.7 | 44.1 | 26.1 |
| sp\|Q03154\|ACY1\_HUMAN | Q03154 | ACY1\_HUMAN | ACY1 | Aminoacylase-1 | 408 | 45.885 | 1.059477e+04 | 6.107725e+05 | 336491.12 | 274281.38 |  |  | 34.31 | 47.1 |  |  |
| sp\|Q9H0X4\|F234A\_HUMAN | Q9H0X4 | F234A\_HUMAN | FAM234A | Protein FAM234A | 552 | 59.660 | 4.004781e+04 | 3.001782e+06 | 335768.88 | 510327.25 | 921746.0 | 1233940.1 | 24.09 | 39.7 | 15.6 | 12.7 |
| sp\|Q14258\|TRI25\_HUMAN | Q14258 | TRI25\_HUMAN | TRIM25 | E3 ubiquitin/ISG15 ligase TRIM25 | 630 | 70.973 | 1.373067e+04 | 1.224340e+06 | 334956.47 | 629456.8 | 259927.1 |  | 30.32 | 54.3 | 31.6 |  |
| sp\|Q9H074\|PAIP1\_HUMAN | Q9H074 | PAIP1\_HUMAN | PAIP1 | Polyadenylate-binding protein-interacting protein 1 | 479 | 53.525 | 1.308878e+04 | 8.801830e+05 | 334029.6 | 429937.8 | 116215.625 |  | 27.77 | 45.7 | 22.3 |  |
| sp\|Q14141\|SEPT6\_HUMAN | Q14141 | SEPT6\_HUMAN | SEPTIN6 | Septin-6 | 434 | 49.717 | 7.810548e+04 | 4.878694e+06 | 333990.4 | 4544703.5 |  |  | 26.27 | 44.9 |  |  |
| sp\|Q5T750\|KPLCE\_HUMAN | Q5T750 | KPLCE\_HUMAN | KPLCE | Protein KPLCE | 250 | 26.238 | 1.182639e+05 | 3.898522e+06 | 332962.03 | 48904.188 | 1712304.2 | 1804351.2 | 9.6 | 39.2 | 30.4 | 30.8 |
| sp\|Q9BTX7\|TTPAL\_HUMAN | Q9BTX7 | TTPAL\_HUMAN | TTPAL | Alpha-tocopherol transfer protein-like | 342 | 38.515 | 1.989287e+04 | 9.625972e+05 | 331374.38 | 395739.0 | 235483.8 |  | 30.7 | 52.3 | 10.8 |  |
| sp\|Q08378\|GOGA3\_HUMAN | Q08378 | GOGA3\_HUMAN | GOLGA3 | Golgin subfamily A member 3 | 1498 | 167.355 | 3.160901e+03 | 6.646097e+05 | 331323.4 | 235826.17 | 42683.31 | 54776.84 | 12.08 | 34.2 | 20.6 | 10.4 |
| sp\|Q7Z4S6\|KI21A\_HUMAN | Q7Z4S6 | KI21A\_HUMAN | KIF21A | Kinesin-like protein KIF21A | 1674 | 187.179 | 1.619379e+03 | 3.808226e+05 | 330965.16 | 43899.08 | 5958.317 |  | 5.02 | 26.2 | 12.4 |  |
| sp\|Q96A32\|MYL11\_HUMAN | Q96A32 | MYL11\_HUMAN | MYL11 | Myosin regulatory light chain 11 | 169 | 19.015 | 3.737791e+04 | 8.929528e+05 | 330569.16 |  | 197669.56 | 364714.12 | 24.26 |  | 60.9 | 40.2 |
| sp\|P11234\|RALB\_HUMAN | P11234 | RALB\_HUMAN | RALB | Ras-related protein Ral-B | 206 | 23.409 | 8.107437e+04 | 2.384425e+06 | 329729.7 | 1785808.6 | 268886.38 |  | 28.64 | 35.4 | 23.3 |  |
| sp\|P49755\|TMEDA\_HUMAN | P49755 | TMEDA\_HUMAN | TMED10 | Transmembrane emp24 domain-containing protein 10 | 219 | 24.976 | 1.496291e+05 | 4.695222e+06 | 329576.84 | 281793.75 | 3631792.0 | 452059.62 | 32.88 | 60.3 | 40.6 | 25.1 |
| sp\|Q9Y266\|NUDC\_HUMAN | Q9Y266 | NUDC\_HUMAN | NUDC | Nuclear migration protein nudC | 331 | 38.243 | 1.889839e+04 | 9.080170e+05 | 328585.44 | 523018.06 | 56413.523 |  | 40.18 | 45.6 | 32.0 |  |
| sp\|Q13813\|SPTN1\_HUMAN | Q13813 | SPTN1\_HUMAN | SPTAN1 | Spectrin alpha chain, non-erythrocytic 1 | 2472 | 284.539 | 9.833499e+04 | 3.515337e+07 | 327963.66 | 1549305.1 | 23224804.0 | 10051301.0 | 2.18 | 29.7 | 33.8 | 11.4 |
| sp\|Q8WUA2\|PPIL4\_HUMAN | Q8WUA2 | PPIL4\_HUMAN | PPIL4 | Peptidyl-prolyl cis-trans isomerase-like 4 | 492 | 57.225 | 1.645931e+04 | 1.183354e+06 | 327417.0 | 732125.8 | 75518.805 | 48292.438 | 31.91 | 37.4 | 12.6 | 12.8 |
| sp\|P22626\|ROA2\_HUMAN | P22626 | ROA2\_HUMAN | HNRNPA2B1 | Heterogeneous nuclear ribonucleoproteins A2/B1 | 353 | 37.430 | 3.536081e+04 | 1.662873e+06 | 326984.28 | 1155167.8 | 61528.812 | 119192.375 | 40.51 | 49.6 | 32.0 | 22.7 |
| sp\|Q99570\|PI3R4\_HUMAN | Q99570 | PI3R4\_HUMAN | PIK3R4 | Phosphoinositide 3-kinase regulatory subunit 4 | 1358 | 153.103 | 4.331948e+03 | 8.332669e+05 | 326658.75 | 506608.12 |  |  | 20.62 | 26.1 |  |  |
| sp\|A0FGR8\|ESYT2\_HUMAN | A0FGR8 | ESYT2\_HUMAN | ESYT2 | Extended synaptotagmin-2 | 921 | 102.357 | 1.494748e+04 | 1.922218e+06 | 326436.97 | 613808.8 | 323262.3 | 658709.6 | 25.52 | 49.1 | 24.2 | 16.8 |
| sp\|P60520\|GBRL2\_HUMAN | P60520 | GBRL2\_HUMAN | GABARAPL2 | Gamma-aminobutyric acid receptor-associated protein-like 2 | 117 | 13.667 | 1.303175e+05 | 2.237655e+06 | 322536.3 | 638427.56 | 413884.12 | 862806.75 | 29.06 | 60.7 | 45.3 | 35.0 |
| sp\|Q6VY07\|PACS1\_HUMAN | Q6VY07 | PACS1\_HUMAN | PACS1 | Phosphofurin acidic cluster sorting protein 1 | 963 | 104.898 | 5.417715e+03 | 7.140038e+05 | 321965.0 | 374482.0 |  | 17556.816 | 26.27 | 31.6 |  | 7.9 |
| sp\|P02787\|TRFE\_HUMAN | P02787 | TRFE\_HUMAN | TF | Serotransferrin | 698 | 77.050 | 1.822020e+04 | 1.763774e+06 | 320872.7 | 352391.16 | 621023.3 | 469487.16 | 12.03 | 62.2 | 35.8 | 19.3 |
| sp\|O00161\|SNP23\_HUMAN | O00161 | SNP23\_HUMAN | SNAP23 | Synaptosomal-associated protein 23 | 211 | 23.354 | 2.633523e+05 | 7.727082e+06 | 319428.03 | 688357.5 | 1678046.1 | 5041250.0 | 38.39 | 56.4 | 84.8 | 53.6 |
| sp\|P42357\|HUTH\_HUMAN | P42357 | HUTH\_HUMAN | HAL | Histidine ammonia-lyase | 657 | 72.698 | 1.077099e+04 | 9.837741e+05 | 319385.5 | 44914.992 | 217665.75 | 401807.84 | 17.5 | 27.9 | 21.5 | 17.0 |
| sp\|Q9UHY7\|ENOPH\_HUMAN | Q9UHY7 | ENOPH\_HUMAN | ENOPH1 | Enolase-phosphatase E1 | 261 | 28.933 | 3.836508e+04 | 1.394590e+06 | 317622.34 | 1076968.1 |  |  | 44.83 | 59.4 |  |  |
| sp\|Q8TBF2\|PXL2B\_HUMAN | Q8TBF2 | PXL2B\_HUMAN | PRXL2B | Prostamide/prostaglandin F synthase | 198 | 21.223 | 2.972503e+04 | 7.925858e+05 | 317433.0 | 311238.25 | 163914.56 |  | 28.28 | 50.0 | 23.2 |  |
| sp\|P14616\|INSRR\_HUMAN | P14616 | INSRR\_HUMAN | INSRR | Insulin receptor-related protein | 1297 | 142.670 | 1.765098e+03 | 3.163871e+05 | 316387.06 |  |  |  | 3.16 |  |  |  |
| sp\|Q8TBE9\|NANP\_HUMAN | Q8TBE9 | NANP\_HUMAN | NANP | N-acylneuraminate-9-phosphatase | 248 | 27.813 | 3.285406e+04 | 1.148032e+06 | 315168.06 | 692630.9 | 140233.06 |  | 16.53 | 46.4 | 29.4 |  |
| sp\|Q15075\|EEA1\_HUMAN | Q15075 | EEA1\_HUMAN | EEA1 | Early endosome antigen 1 | 1411 | 162.466 | 6.329353e+03 | 1.291930e+06 | 314930.16 | 833909.1 | 66010.01 | 77080.75 | 44.22 | 47.9 | 26.2 | 15.3 |
| sp\|Q9P2W9\|STX18\_HUMAN | Q9P2W9 | STX18\_HUMAN | STX18 | Syntaxin-18 | 335 | 38.674 | 1.440100e+04 | 6.997272e+05 | 314629.6 | 220032.9 | 99581.53 | 65483.203 | 13.43 | 38.2 | 15.8 | 17.9 |
| sp\|P23610\|HAP40\_HUMAN | P23610 | HAP40\_HUMAN | F8A1 | 40-kDa huntingtin-associated protein | 371 | 39.103 | 1.078867e+04 | 5.300234e+05 | 313891.12 | 154822.0 | 61310.305 |  | 25.61 | 29.9 | 14.3 |  |
| sp\|O95989\|NUDT3\_HUMAN | O95989 | NUDT3\_HUMAN | NUDT3 | Diphosphoinositol polyphosphate phosphohydrolase 1 | 172 | 19.471 | 5.348290e+04 | 1.308339e+06 | 313466.44 | 989490.7 | 5382.1406 |  | 35.47 | 39.5 | 12.8 |  |
| sp\|O95825\|QORL1\_HUMAN | O95825 | QORL1\_HUMAN | CRYZL1 | Quinone oxidoreductase-like protein 1 | 349 | 38.697 | 1.714397e+04 | 8.335003e+05 | 312458.8 | 480049.34 | 40992.117 |  | 34.67 | 51.0 | 12.0 |  |
| sp\|Q8NBF6\|AVL9\_HUMAN | Q8NBF6 | AVL9\_HUMAN | AVL9 | Late secretory pathway protein AVL9 homolog | 648 | 71.947 | 4.891107e+03 | 4.421168e+05 | 311971.38 | 101425.68 | 28719.762 |  | 15.9 | 22.8 | 14.7 |  |
| sp\|Q9NVM6\|DJC17\_HUMAN | Q9NVM6 | DJC17\_HUMAN | DNAJC17 | DnaJ homolog subfamily C member 17 | 304 | 34.687 | 1.456853e+04 | 6.348916e+05 | 311432.47 | 314071.56 | 9387.575 |  | 26.32 | 58.9 | 21.4 |  |
| sp\|Q9BZK3\|NACP4\_HUMAN | Q9BZK3 | NACP4\_HUMAN | NACA4P | Putative nascent polypeptide-associated complex subunit alpha-like protein | 213 | 23.430 | 1.057628e+04 | 3.113311e+05 | 311331.06 |  |  |  | 14.08 |  |  |  |
| sp\|O95292\|VAPB\_HUMAN | O95292 | VAPB\_HUMAN | VAPB | Vesicle-associated membrane protein-associated protein B/C | 243 | 27.228 | 1.310777e+05 | 4.483959e+06 | 310401.62 | 1224454.1 | 928038.5 | 2021065.2 | 30.45 | 65.0 | 44.0 | 39.1 |
| sp\|Q68EM7\|RHG17\_HUMAN | Q68EM7 | RHG17\_HUMAN | ARHGAP17 | Rho GTPase-activating protein 17 | 881 | 95.437 | 1.119900e+04 | 1.342806e+06 | 308727.6 | 915526.3 | 83232.94 | 35318.9 | 26.79 | 40.5 | 14.1 | 6.8 |
| sp\|Q96P63\|SPB12\_HUMAN | Q96P63 | SPB12\_HUMAN | SERPINB12 | Serpin B12 | 405 | 46.276 | 5.562577e+04 | 3.234067e+06 | 308138.75 | 140455.05 | 731845.3 | 2053628.0 | 27.9 | 47.9 | 18.3 | 21.0 |
| sp\|P98171\|RHG04\_HUMAN | P98171 | RHG04\_HUMAN | ARHGAP4 | Rho GTPase-activating protein 4 | 946 | 105.026 | 8.362397e+03 | 1.103430e+06 | 307588.88 | 718549.8 | 77291.24 |  | 23.47 | 43.2 | 17.9 |  |
| sp\|Q9NYB0\|TE2IP\_HUMAN | Q9NYB0 | TE2IP\_HUMAN | TERF2IP | Telomeric repeat-binding factor 2-interacting protein 1 | 399 | 44.260 | 1.336247e+04 | 7.430453e+05 | 307176.75 | 425699.75 | 10168.754 |  | 13.78 | 44.9 | 14.0 |  |
| sp\|Q9Y5X2\|SNX8\_HUMAN | Q9Y5X2 | SNX8\_HUMAN | SNX8 | Sorting nexin-8 | 465 | 52.569 | 1.631742e+04 | 1.077701e+06 | 306415.7 | 481045.78 | 290239.56 |  | 23.66 | 50.1 | 36.1 |  |
| sp\|Q6UWP8\|SBSN\_HUMAN | Q6UWP8 | SBSN\_HUMAN | SBSN | Suprabasin | 590 | 60.541 | 2.868100e+04 | 2.181529e+06 | 304688.12 | 1106010.0 | 477720.62 | 293110.28 | 44.92 | 74.6 | 40.3 | 27.8 |
| sp\|Q9Y3X0\|CCDC9\_HUMAN | Q9Y3X0 | CCDC9\_HUMAN | CCDC9 | Coiled-coil domain-containing protein 9 | 531 | 59.703 | 1.933502e+04 | 1.450300e+06 | 304661.56 | 962555.0 | 113557.12 | 69526.7 | 26.93 | 39.9 | 29.8 | 26.0 |
| sp\|P08133\|ANXA6\_HUMAN | P08133 | ANXA6\_HUMAN | ANXA6 | Annexin A6 | 673 | 75.873 | 9.417255e+03 | 8.976949e+05 | 303867.3 | 593827.56 |  |  | 35.66 | 50.5 |  |  |
| sp\|P63279\|UBC9\_HUMAN | P63279 | UBC9\_HUMAN | UBE2I | SUMO-conjugating enzyme UBC9 | 158 | 18.007 | 3.464743e+04 | 7.838438e+05 | 303633.75 | 480210.1 |  |  | 20.25 | 62.0 |  |  |
| sp\|Q9H0R3\|TM222\_HUMAN | Q9H0R3 | TM222\_HUMAN | TMEM222 | Transmembrane protein 222 | 208 | 23.230 | 1.754022e+05 | 5.119192e+06 | 303584.5 | 839737.0 | 701812.9 | 3274058.0 | 26.44 | 23.6 | 38.5 | 18.3 |
| sp\|Q8TF05\|PP4R1\_HUMAN | Q8TF05 | PP4R1\_HUMAN | PPP4R1 | Serine/threonine-protein phosphatase 4 regulatory subunit 1 | 950 | 107.004 | 7.190113e+03 | 9.666135e+05 | 303518.34 | 542680.6 | 120414.59 |  | 12.11 | 34.6 | 15.8 |  |
| sp\|Q96SZ5\|AEDO\_HUMAN | Q96SZ5 | AEDO\_HUMAN | ADO | 2-aminoethanethiol dioxygenase | 270 | 29.751 | 1.427836e+04 | 5.336998e+05 | 303046.8 | 230653.05 |  |  | 31.85 | 64.1 |  |  |
| sp\|Q9ULC5\|ACSL5\_HUMAN | Q9ULC5 | ACSL5\_HUMAN | ACSL5 | Long-chain-fatty-acid--CoA ligase 5 | 683 | 75.991 | 1.556665e+04 | 1.486191e+06 | 302790.03 | 410169.7 | 380311.2 | 392919.94 | 23.87 | 27.5 | 14.6 | 17.0 |
| sp\|Q9Y3F4\|STRAP\_HUMAN | Q9Y3F4 | STRAP\_HUMAN | STRAP | Serine-threonine kinase receptor-associated protein | 350 | 38.438 | 1.934347e+04 | 9.341409e+05 | 301280.78 | 565096.75 |  | 67763.36 | 54.29 | 50.9 |  | 11.7 |
| sp\|O43633\|CHM2A\_HUMAN | O43633 | CHM2A\_HUMAN | CHMP2A | Charged multivesicular body protein 2a | 222 | 25.104 | 8.909020e+04 | 2.809895e+06 | 300347.84 | 1303261.5 | 1206285.2 |  | 23.42 | 49.1 | 40.5 |  |
| sp\|Q9Y5B8\|NDK7\_HUMAN | Q9Y5B8 | NDK7\_HUMAN | NME7 | Nucleoside diphosphate kinase 7 | 376 | 42.492 | 1.459469e+04 | 7.791467e+05 | 299823.4 | 479323.28 |  |  | 31.65 | 41.5 |  |  |
| sp\|P14209\|CD99\_HUMAN | P14209 | CD99\_HUMAN | CD99 | CD99 antigen | 185 | 18.848 | 4.417025e+05 | 1.045953e+07 | 297814.94 | 675466.3 | 5193967.0 | 4292283.5 | 12.97 | 13.0 | 13.0 | 13.0 |
| sp\|Q6ZVD8\|PHLP2\_HUMAN | Q6ZVD8 | PHLP2\_HUMAN | PHLPP2 | PH domain leucine-rich repeat-containing protein phosphatase 2 | 1323 | 146.751 | 1.767396e+03 | 3.258607e+05 | 295644.28 | 30216.459 |  |  | 2.27 | 20.8 |  |  |
| sp\|Q9BXP5\|SRRT\_HUMAN | Q9BXP5 | SRRT\_HUMAN | SRRT | Serrate RNA effector molecule homolog | 876 | 100.666 | 6.821920e+03 | 8.627930e+05 | 295219.7 | 549208.0 |  | 18365.297 | 22.03 | 41.6 |  | 13.5 |
| sp\|Q53T59\|H1BP3\_HUMAN | Q53T59 | H1BP3\_HUMAN | HS1BP3 | HCLS1-binding protein 3 | 392 | 42.780 | 1.220712e+04 | 6.561016e+05 | 295089.22 | 331264.56 | 29747.803 |  | 27.55 | 34.7 | 22.2 |  |
| sp\|Q9P227\|RHG23\_HUMAN | Q9P227 | RHG23\_HUMAN | ARHGAP23 | Rho GTPase-activating protein 23 | 1491 | 162.192 | 8.364993e+03 | 1.704559e+06 | 293930.6 | 592062.5 | 739919.7 | 78646.664 | 12.88 | 51.4 | 34.5 | 18.4 |
| sp\|Q04206\|TF65\_HUMAN | Q04206 | TF65\_HUMAN | RELA | Transcription factor p65 | 551 | 60.219 | 7.639039e+03 | 5.779489e+05 | 293840.75 | 225101.33 | 59006.797 |  | 9.8 | 46.8 | 18.1 |  |
| sp\|Q8ND76\|CCNY\_HUMAN | Q8ND76 | CCNY\_HUMAN | CCNY | Cyclin-Y | 341 | 39.337 | 5.418879e+04 | 2.678106e+06 | 292501.2 | 141619.28 | 788526.4 | 1455459.4 | 19.65 | 46.0 | 26.1 | 15.2 |
| sp\|Q9H3Z4\|DNJC5\_HUMAN | Q9H3Z4 | DNJC5\_HUMAN | DNAJC5 | DnaJ homolog subfamily C member 5 | 198 | 22.149 | 9.661898e+04 | 2.688647e+06 | 291212.4 | 1243818.2 | 711069.44 | 442546.56 | 24.24 | 18.2 | 19.7 | 15.7 |
| sp\|Q9NR19\|ACSA\_HUMAN | Q9NR19 | ACSA\_HUMAN | ACSS2 | Acetyl-coenzyme A synthetase, cytoplasmic | 701 | 78.580 | 5.281643e+03 | 5.214326e+05 | 290859.53 | 230573.1 |  |  | 16.12 | 32.4 |  |  |
| sp\|Q15003\|CND2\_HUMAN | Q15003 | CND2\_HUMAN | NCAPH | Condensin complex subunit 2 | 741 | 82.563 | 8.327748e+03 | 8.638338e+05 | 289127.38 | 316747.44 | 160525.34 | 97433.62 | 20.65 | 33.9 | 29.6 | 9.7 |
| sp\|P53384\|NUBP1\_HUMAN | P53384 | NUBP1\_HUMAN | NUBP1 | Cytosolic Fe-S cluster assembly factor NUBP1 | 320 | 34.534 | 1.471593e+04 | 6.384867e+05 | 288442.0 | 291581.5 | 58463.24 |  | 8.44 | 33.4 | 22.8 |  |
| sp\|Q9NZZ3\|CHMP5\_HUMAN | Q9NZZ3 | CHMP5\_HUMAN | CHMP5 | Charged multivesicular body protein 5 | 219 | 24.571 | 5.224444e+04 | 1.612798e+06 | 287607.62 | 836965.5 | 171204.27 | 317020.94 | 25.11 | 35.6 | 30.1 | 26.0 |
| sp\|P32929\|CGL\_HUMAN | P32929 | CGL\_HUMAN | CTH | Cystathionine gamma-lyase | 405 | 44.508 | 2.660186e+04 | 1.487535e+06 | 286533.0 | 1139273.8 | 61728.207 |  | 22.96 | 62.5 | 23.0 |  |
| sp\|Q8N392\|RHG18\_HUMAN | Q8N392 | RHG18\_HUMAN | ARHGAP18 | Rho GTPase-activating protein 18 | 663 | 74.977 | 8.072624e+03 | 7.604312e+05 | 285871.12 | 425930.56 | 41334.516 | 7295.0073 | 28.21 | 54.1 | 15.4 | 13.3 |
| sp\|P61421\|VA0D1\_HUMAN | P61421 | VA0D1\_HUMAN | ATP6V0D1 | V-type proton ATPase subunit d 1 | 351 | 40.329 | 5.425644e+04 | 2.749071e+06 | 285350.8 |  | 2270559.0 | 193161.02 | 15.95 |  | 25.1 | 18.5 |
| sp\|P04843\|RPN1\_HUMAN | P04843 | RPN1\_HUMAN | RPN1 | Dolichyl-diphosphooligosaccharide--protein glycosyltransferase subunit 1 | 607 | 68.569 | 3.041687e+04 | 2.620351e+06 | 284665.53 | 243395.31 | 1214643.1 | 877647.3 | 22.9 | 37.9 | 32.9 | 17.0 |
| sp\|Q6ZSR9\|YJ005\_HUMAN | Q6ZSR9 | YJ005\_HUMAN |  | Uncharacterized protein FLJ45252 | 355 | 39.050 | 1.188471e+04 | 5.830779e+05 | 283402.4 | 222277.22 | 77398.33 |  | 28.17 | 50.1 | 18.9 |  |
| sp\|P48200\|IREB2\_HUMAN | P48200 | IREB2\_HUMAN | IREB2 | Iron-responsive element-binding protein 2 | 963 | 105.059 | 6.350957e+03 | 8.382810e+05 | 283162.47 | 555118.5 |  |  | 20.56 | 41.3 |  |  |
| sp\|P54619\|AAKG1\_HUMAN | P54619 | AAKG1\_HUMAN | PRKAG1 | 5'-AMP-activated protein kinase subunit gamma-1 | 331 | 37.579 | 1.722091e+04 | 8.130522e+05 | 281299.0 | 326510.47 | 153623.75 | 51618.95 | 24.77 | 31.7 | 17.8 | 8.8 |
| sp\|Q96T51\|RUFY1\_HUMAN | Q96T51 | RUFY1\_HUMAN | RUFY1 | RUN and FYVE domain-containing protein 1 | 708 | 79.818 | 1.618018e+04 | 1.622562e+06 | 281041.97 | 1066982.2 | 171838.86 | 102698.63 | 18.79 | 45.8 | 26.3 | 24.3 |
| sp\|Q9BV68\|RN126\_HUMAN | Q9BV68 | RN126\_HUMAN | RNF126 | E3 ubiquitin-protein ligase RNF126 | 311 | 33.861 | 5.717408e+04 | 2.432294e+06 | 280967.88 | 1685461.6 | 302089.03 | 163775.69 | 7.07 | 16.1 | 15.1 | 10.3 |
| sp\|Q8TEA8\|DTD1\_HUMAN | Q8TEA8 | DTD1\_HUMAN | DTD1 | D-aminoacyl-tRNA deacylase 1 | 209 | 23.424 | 3.407566e+04 | 1.002819e+06 | 280807.9 | 669877.75 | 44507.31 | 7625.889 | 22.01 | 47.8 | 22.5 | 7.7 |
| sp\|O43318\|M3K7\_HUMAN | O43318 | M3K7\_HUMAN | MAP3K7 | Mitogen-activated protein kinase kinase kinase 7 | 606 | 67.196 | 9.232825e+03 | 7.794624e+05 | 280654.0 | 411179.0 | 87629.41 |  | 13.37 | 48.5 | 28.2 |  |
| sp\|P49773\|HINT1\_HUMAN | P49773 | HINT1\_HUMAN | HINT1 | Adenosine 5'-monophosphoramidase HINT1 | 126 | 13.802 | 7.075678e+05 | 1.226951e+07 | 279426.78 | 11439731.0 | 550352.5 |  | 53.97 | 92.1 | 30.2 |  |
| sp\|P78368\|KC1G2\_HUMAN | P78368 | KC1G2\_HUMAN | CSNK1G2 | Casein kinase I isoform gamma-2 | 415 | 47.457 | 1.998411e+04 | 1.191522e+06 | 278169.66 | 236259.06 |  | 677093.5 | 23.37 | 57.3 |  | 12.5 |
| sp\|Q8IUZ5\|AT2L2\_HUMAN | Q8IUZ5 | AT2L2\_HUMAN | PHYKPL | 5-phosphohydroxy-L-lysine phospho-lyase | 450 | 49.711 | 5.311607e+03 | 3.317383e+05 | 277609.75 | 54128.52 |  |  | 13.11 | 32.2 |  |  |
| sp\|P31749\|AKT1\_HUMAN | P31749 | AKT1\_HUMAN | AKT1 | RAC-alpha serine/threonine-protein kinase | 480 | 55.686 | 5.863797e+03 | 4.102439e+05 | 277603.94 | 132639.94 |  |  | 6.04 | 32.1 |  |  |
| sp\|A2RRP1\|NBAS\_HUMAN | A2RRP1 | NBAS\_HUMAN | NBAS | NBAS subunit of NRZ tethering complex | 2371 | 268.571 | 5.225010e+03 | 1.763045e+06 | 277433.75 | 786399.2 | 209185.25 | 490026.8 | 17.04 | 34.5 | 17.6 | 15.9 |
| sp\|Q96LD8\|SENP8\_HUMAN | Q96LD8 | SENP8\_HUMAN | SENP8 | Sentrin-specific protease 8 | 212 | 24.107 | 1.759904e+04 | 5.330271e+05 | 277220.4 | 255806.72 |  |  | 29.25 | 36.3 |  |  |
| sp\|Q969M7\|UBE2F\_HUMAN | Q969M7 | UBE2F\_HUMAN | UBE2F | NEDD8-conjugating enzyme UBE2F | 185 | 21.077 | 1.894523e+04 | 5.016788e+05 | 277069.25 | 224609.56 |  |  | 24.86 | 40.0 |  |  |
| sp\|Q9BT09\|CNPY3\_HUMAN | Q9BT09 | CNPY3\_HUMAN | CNPY3 | Protein canopy homolog 3 | 278 | 30.748 | 3.595698e+04 | 1.389048e+06 | 276277.97 |  | 1036857.0 | 75912.94 | 13.31 |  | 29.1 | 20.1 |
| sp\|Q7RTP6\|MICA3\_HUMAN | Q7RTP6 | MICA3\_HUMAN | MICAL3 | [F-actin]-monooxygenase MICAL3 | 2002 | 224.295 | 6.610093e+03 | 1.862706e+06 | 275855.0 | 1201933.8 | 205062.9 | 179854.19 | 16.43 | 36.0 | 24.8 | 15.9 |
| sp\|P18440\|ARY1\_HUMAN | P18440 | ARY1\_HUMAN | NAT1 | Arylamine N-acetyltransferase 1 | 290 | 33.899 | 1.356079e+04 | 5.775493e+05 | 275072.7 | 302476.62 | 0.0 |  | 24.83 | 34.8 | 6.6 |  |
| sp\|P32019\|I5P2\_HUMAN | P32019 | I5P2\_HUMAN | INPP5B | Type II inositol 1,4,5-trisphosphate 5-phosphatase | 993 | 112.852 | 4.349578e+03 | 6.166994e+05 | 274683.88 | 248331.56 | 93683.914 |  | 18.53 | 33.2 | 22.3 |  |
| sp\|Q9BWS9\|CHID1\_HUMAN | Q9BWS9 | CHID1\_HUMAN | CHID1 | Chitinase domain-containing protein 1 | 393 | 44.941 | 5.575076e+04 | 3.147826e+06 | 274360.7 | 471420.75 | 1803838.6 | 598205.5 | 20.1 | 33.6 | 31.3 | 12.5 |
| sp\|Q8N370\|LAT4\_HUMAN | Q8N370 | LAT4\_HUMAN | SLC43A2 | Large neutral amino acids transporter small subunit 4 | 569 | 62.747 | 1.304664e+05 | 1.028510e+07 | 274111.78 | 869969.1 | 1022131.8 | 8118891.5 | 10.9 | 16.5 | 14.2 | 16.3 |
| sp\|Q16773\|KAT1\_HUMAN | Q16773 | KAT1\_HUMAN | KYAT1 | Kynurenine--oxoglutarate transaminase 1 | 422 | 47.875 | 1.202203e+04 | 7.231092e+05 | 273946.38 | 449162.8 |  |  | 19.43 | 34.1 |  |  |
| sp\|Q15139\|KPCD1\_HUMAN | Q15139 | KPCD1\_HUMAN | PRKD1 | Serine/threonine-protein kinase D1 | 912 | 100.320 | 2.169896e+03 | 2.734914e+05 | 273491.4 |  |  |  | 2.3 |  |  |  |
| sp\|Q92681\|RSCA1\_HUMAN | Q92681 | RSCA1\_HUMAN | RSC1A1 | Regulatory solute carrier protein family 1 member 1 | 617 | 66.790 | 4.200034e+03 | 3.524370e+05 | 273174.75 | 79262.2 |  |  | 14.1 | 32.4 |  |  |
| sp\|Q8TBZ3\|WDR20\_HUMAN | Q8TBZ3 | WDR20\_HUMAN | WDR20 | WD repeat-containing protein 20 | 569 | 62.893 | 6.012913e+04 | 4.751212e+06 | 273026.66 | 308685.78 | 1366952.1 | 2802547.2 | 13.71 | 42.2 | 23.0 | 16.0 |
| sp\|Q9NZT1\|CALL5\_HUMAN | Q9NZT1 | CALL5\_HUMAN | CALML5 | Calmodulin-like protein 5 | 146 | 15.893 | 2.852698e+05 | 5.696117e+06 | 271946.66 | 303616.53 | 2002495.4 | 3118058.5 | 36.99 | 38.4 | 48.6 | 21.2 |
| sp\|P18433\|PTPRA\_HUMAN | P18433 | PTPRA\_HUMAN | PTPRA | Receptor-type tyrosine-protein phosphatase alpha | 802 | 90.719 | 9.804694e+03 | 1.117505e+06 | 271151.75 | 90647.64 | 402550.78 | 353154.72 | 15.21 | 31.3 | 17.8 | 11.0 |
| sp\|Q96RT1\|ERBIN\_HUMAN | Q96RT1 | ERBIN\_HUMAN | ERBIN | Erbin | 1412 | 158.298 | 5.718248e+03 | 1.137249e+06 | 270970.94 | 476378.22 | 161520.11 | 228379.81 | 14.02 | 31.5 | 24.2 | 14.0 |
| sp\|Q8WVY7\|UBCP1\_HUMAN | Q8WVY7 | UBCP1\_HUMAN | UBLCP1 | Ubiquitin-like domain-containing CTD phosphatase 1 | 318 | 36.805 | 1.382401e+04 | 6.392314e+05 | 270242.4 | 282975.78 | 86013.2 |  | 16.67 | 46.9 | 18.9 |  |
| sp\|Q6UWE0\|LRSM1\_HUMAN | Q6UWE0 | LRSM1\_HUMAN | LRSAM1 | E3 ubiquitin-protein ligase LRSAM1 | 723 | 83.594 | 6.900619e+03 | 7.247368e+05 | 270110.0 | 328992.03 | 125634.805 |  | 21.02 | 40.7 | 24.5 |  |
| sp\|Q7L576\|CYFP1\_HUMAN | Q7L576 | CYFP1\_HUMAN | CYFIP1 | Cytoplasmic FMR1-interacting protein 1 | 1253 | 145.182 | 5.895491e+03 | 1.075350e+06 | 269416.4 | 518241.22 | 44865.723 | 242826.75 | 22.91 | 34.1 | 14.9 | 13.2 |
| sp\|P04080\|CYTB\_HUMAN | P04080 | CYTB\_HUMAN | CSTB | Cystatin-B | 98 | 11.140 | 4.828791e+05 | 6.758351e+06 | 269073.0 | 5297843.5 | 1059652.2 | 131782.19 | 45.92 | 93.9 | 55.1 | 21.4 |
| sp\|Q5T0N5\|FBP1L\_HUMAN | Q5T0N5 | FBP1L\_HUMAN | FNBP1L | Formin-binding protein 1-like | 605 | 70.065 | 1.167222e+04 | 1.027476e+06 | 268932.44 | 651196.44 | 107347.0 |  | 17.36 | 42.5 | 29.6 |  |
| sp\|P50454\|SERPH\_HUMAN | P50454 | SERPH\_HUMAN | SERPINH1 | Serpin H1 | 418 | 46.441 | 7.982056e+04 | 4.657292e+06 | 268898.3 | 208543.44 | 4023520.8 | 156329.94 | 21.05 | 43.8 | 34.2 | 18.2 |
| sp\|P11172\|UMPS\_HUMAN | P11172 | UMPS\_HUMAN | UMPS | Uridine 5'-monophosphate synthase | 480 | 52.222 | 1.423115e+04 | 9.337073e+05 | 268717.25 | 497996.66 | 166993.4 | 0.0 | 23.12 | 42.1 | 18.1 | 5.8 |
| sp\|Q9NWU2\|GID8\_HUMAN | Q9NWU2 | GID8\_HUMAN | GID8 | Glucose-induced degradation protein 8 homolog | 228 | 26.749 | 1.558815e+04 | 5.238648e+05 | 268464.88 | 213936.62 | 41463.25 |  | 32.46 | 45.6 | 21.5 |  |
| sp\|Q9Y6Y8\|S23IP\_HUMAN | Q9Y6Y8 | S23IP\_HUMAN | SEC23IP | SEC23-interacting protein | 1000 | 111.076 | 8.313381e+03 | 1.160152e+06 | 266927.4 | 803270.75 | 89954.32 |  | 20.5 | 29.5 | 15.9 |  |
| sp\|Q9H9C1\|SPE39\_HUMAN | Q9H9C1 | SPE39\_HUMAN | VIPAS39 | Spermatogenesis-defective protein 39 homolog | 493 | 57.005 | 2.089335e+04 | 1.496367e+06 | 266525.84 | 1229841.1 |  |  | 7.71 | 53.1 |  |  |
| sp\|Q6NXT1\|ANR54\_HUMAN | Q6NXT1 | ANR54\_HUMAN | ANKRD54 | Ankyrin repeat domain-containing protein 54 | 300 | 32.505 | 3.086101e+04 | 1.260310e+06 | 266052.84 | 683267.25 | 130951.836 | 180038.11 | 10.0 | 55.0 | 18.7 | 12.3 |
| sp\|O14602\|IF1AY\_HUMAN | O14602 | IF1AY\_HUMAN | EIF1AY | Eukaryotic translation initiation factor 1A, Y-chromosomal | 144 | 16.442 | 3.640823e+04 | 7.520928e+05 | 265837.78 |  | 486255.0 |  | 24.31 |  | 19.4 |  |
| sp\|Q99683\|M3K5\_HUMAN | Q99683 | M3K5\_HUMAN | MAP3K5 | Mitogen-activated protein kinase kinase kinase 5 | 1374 | 154.537 | 3.356206e+03 | 6.516257e+05 | 265755.5 | 385870.22 |  |  | 9.17 | 35.9 |  |  |
| sp\|Q8TE77\|SSH3\_HUMAN | Q8TE77 | SSH3\_HUMAN | SSH3 | Protein phosphatase Slingshot homolog 3 | 659 | 72.996 | 1.261601e+04 | 1.157013e+06 | 265264.8 | 847804.5 | 43943.895 |  | 24.89 | 44.2 | 20.3 |  |
| sp\|Q13451\|FKBP5\_HUMAN | Q13451 | FKBP5\_HUMAN | FKBP5 | Peptidyl-prolyl cis-trans isomerase FKBP5 | 457 | 51.212 | 1.774405e+04 | 1.141673e+06 | 264378.7 | 848951.4 | 28342.848 |  | 36.32 | 59.1 | 21.0 |  |
| sp\|P32121\|ARRB2\_HUMAN | P32121 | ARRB2\_HUMAN | ARRB2 | Beta-arrestin-2 | 409 | 46.106 | 1.821953e+04 | 1.055387e+06 | 264225.12 | 738359.6 | 52802.625 |  | 23.96 | 53.3 | 24.7 |  |
| sp\|Q9H3N1\|TMX1\_HUMAN | Q9H3N1 | TMX1\_HUMAN | TMX1 | Thioredoxin-related transmembrane protein 1 | 280 | 31.791 | 1.101940e+05 | 4.401282e+06 | 263963.62 | 585269.25 | 870483.56 | 2681566.0 | 20.0 | 36.8 | 30.0 | 27.1 |
| sp\|Q96C23\|GALM\_HUMAN | Q96C23 | GALM\_HUMAN | GALM | Galactose mutarotase | 342 | 37.766 | 9.735101e+03 | 4.619113e+05 | 263636.6 | 198274.72 |  |  | 24.27 | 26.3 |  |  |
| sp\|O95236\|APOL3\_HUMAN | O95236 | APOL3\_HUMAN | APOL3 | Apolipoprotein L3 | 402 | 44.278 | 3.186453e+04 | 1.772608e+06 | 263570.44 | 319461.1 | 889780.7 | 299795.53 | 11.19 | 32.1 | 30.1 | 18.7 |
| sp\|O00267\|SPT5H\_HUMAN | O00267 | SPT5H\_HUMAN | SUPT5H | Transcription elongation factor SPT5 | 1087 | 121.000 | 2.939600e+03 | 4.468799e+05 | 263433.5 | 183446.36 |  |  | 24.56 | 31.6 |  |  |
| sp\|Q9NVE7\|PANK4\_HUMAN | Q9NVE7 | PANK4\_HUMAN | PANK4 | 4'-phosphopantetheine phosphatase | 773 | 85.991 | 7.207694e+03 | 7.786934e+05 | 263099.0 | 515594.44 |  |  | 20.31 | 36.7 |  |  |
| sp\|P37235\|HPCL1\_HUMAN | P37235 | HPCL1\_HUMAN | HPCAL1 | Hippocalcin-like protein 1 | 193 | 22.313 | 7.938094e+04 | 2.225314e+06 | 262862.22 | 1962452.2 |  |  | 52.33 | 65.3 |  |  |
| sp\|Q2NKX8\|ERC6L\_HUMAN | Q2NKX8 | ERC6L\_HUMAN | ERCC6L | DNA excision repair protein ERCC-6-like | 1250 | 141.103 | 3.533508e+03 | 6.264111e+05 | 262562.97 | 271457.47 | 73627.125 | 18763.5 | 18.32 | 37.7 | 18.7 | 8.2 |
| sp\|Q9Y6W3\|CAN7\_HUMAN | Q9Y6W3 | CAN7\_HUMAN | CAPN7 | Calpain-7 | 813 | 92.652 | 7.630706e+03 | 8.882529e+05 | 262136.02 | 553012.7 | 73104.21 |  | 26.94 | 48.0 | 21.2 |  |
| sp\|Q13409\|DC1I2\_HUMAN | Q13409 | DC1I2\_HUMAN | DYNC1I2 | Cytoplasmic dynein 1 intermediate chain 2 | 638 | 71.457 | 1.554079e+04 | 1.395196e+06 | 261950.56 | 206623.27 | 524740.1 | 401881.6 | 7.68 | 17.7 | 11.3 | 6.7 |
| sp\|Q5T1M5\|FKB15\_HUMAN | Q5T1M5 | FKB15\_HUMAN | FKBP15 | FK506-binding protein 15 | 1219 | 133.630 | 8.524922e+03 | 1.431237e+06 | 261402.97 | 997431.5 | 172402.53 |  | 21.49 | 30.9 | 16.9 |  |
| sp\|Q13042\|CDC16\_HUMAN | Q13042 | CDC16\_HUMAN | CDC16 | Cell division cycle protein 16 homolog | 620 | 71.656 | 4.785606e+03 | 4.308307e+05 | 261011.4 | 84941.695 | 84877.625 |  | 16.94 | 22.6 | 14.2 |  |
| sp\|Q9BVK6\|TMED9\_HUMAN | Q9BVK6 | TMED9\_HUMAN | TMED9 | Transmembrane emp24 domain-containing protein 9 | 235 | 27.277 | 7.785673e+04 | 2.668148e+06 | 260957.72 | 372701.1 | 1337657.8 | 696831.5 | 8.51 | 38.3 | 31.1 | 24.7 |
| sp\|Q9NV96\|CC50A\_HUMAN | Q9NV96 | CC50A\_HUMAN | TMEM30A | Cell cycle control protein 50A | 361 | 40.684 | 1.205566e+05 | 6.162145e+06 | 260570.84 | 139738.34 | 2553284.8 | 3208551.2 | 11.08 | 23.0 | 38.0 | 21.3 |
| sp\|Q9C0I1\|MTMRC\_HUMAN | Q9C0I1 | MTMRC\_HUMAN | MTMR12 | Myotubularin-related protein 12 | 747 | 86.148 | 7.578183e+03 | 8.202145e+05 | 260083.62 | 515316.9 | 44813.953 |  | 28.51 | 51.8 | 16.9 |  |
| sp\|P10909\|CLUS\_HUMAN | P10909 | CLUS\_HUMAN | CLU | Clusterin | 449 | 52.495 | 8.667901e+04 | 5.716748e+06 | 259185.69 | 480781.5 | 4497249.0 | 479531.94 | 22.05 | 29.8 | 31.8 | 10.7 |
| sp\|Q96CT7\|CC124\_HUMAN | Q96CT7 | CC124\_HUMAN | CCDC124 | Coiled-coil domain-containing protein 124 | 223 | 25.835 | 3.347044e+04 | 1.086393e+06 | 259083.0 | 634547.56 | 148172.72 | 44590.01 | 22.42 | 64.1 | 35.0 | 27.4 |
| sp\|Q8NEB9\|PK3C3\_HUMAN | Q8NEB9 | PK3C3\_HUMAN | PIK3C3 | Phosphatidylinositol 3-kinase catalytic subunit type 3 | 887 | 101.549 | 8.313807e+03 | 1.060700e+06 | 254956.89 | 701644.44 | 104099.12 |  | 26.49 | 42.7 | 18.4 |  |
| sp\|Q8WUY1\|THEM6\_HUMAN | Q8WUY1 | THEM6\_HUMAN | THEM6 | Protein THEM6 | 208 | 23.865 | 9.982560e+04 | 2.993095e+06 | 254441.16 | 992133.6 | 505119.6 | 1241400.8 | 28.37 | 51.0 | 39.9 | 30.8 |
| sp\|P13533\|MYH6\_HUMAN | P13533 | MYH6\_HUMAN | MYH6 | Myosin-6 | 1939 | 223.735 | 6.448471e+03 | 1.812624e+06 | 253521.95 | 1559102.5 |  |  | 13.87 | 31.7 |  |  |
| sp\|P86790\|CCZ1B\_HUMAN | P86790 | CCZ1B\_HUMAN | CCZ1B | Vacuolar fusion protein CCZ1 homolog B | 482 | 55.866 | 7.405323e+03 | 5.197670e+05 | 253404.4 | 266362.62 |  |  | 11.41 | 56.8 |  |  |
| sp\|Q14764\|MVP\_HUMAN | Q14764 | MVP\_HUMAN | MVP | Major vault protein | 893 | 99.327 | 6.348935e+03 | 7.922922e+05 | 252919.45 | 539372.75 |  |  | 27.88 | 40.3 |  |  |
| sp\|Q70J99\|UN13D\_HUMAN | Q70J99 | UN13D\_HUMAN | UNC13D | Protein unc-13 homolog D | 1090 | 123.282 | 4.983094e+03 | 7.718199e+05 | 252815.19 | 501802.12 | 17202.545 |  | 14.4 | 20.5 | 6.7 |  |
| sp\|P10398\|ARAF\_HUMAN | P10398 | ARAF\_HUMAN | ARAF | Serine/threonine-protein kinase A-Raf | 606 | 67.585 | 7.436343e+03 | 6.314324e+05 | 252307.58 | 294517.75 | 78209.67 | 6397.4395 | 17.16 | 28.1 | 28.1 | 13.4 |
| sp\|P35670\|ATP7B\_HUMAN | P35670 | ATP7B\_HUMAN | ATP7B | Copper-transporting ATPase 2 | 1465 | 157.263 | 2.299982e+04 | 4.544312e+06 | 250695.19 | 125625.234 | 697567.94 | 3470423.8 | 7.99 | 25.4 | 22.3 | 14.9 |
| sp\|P22735\|TGM1\_HUMAN | P22735 | TGM1\_HUMAN | TGM1 | Protein-glutamine gamma-glutamyltransferase K | 817 | 89.787 | 2.416405e+04 | 2.725840e+06 | 249338.36 | 423628.5 | 827838.2 | 1225034.8 | 18.6 | 25.1 | 24.5 | 20.1 |
| sp\|Q13496\|MTM1\_HUMAN | Q13496 | MTM1\_HUMAN | MTM1 | Myotubularin | 603 | 69.932 | 1.039863e+04 | 9.136274e+05 | 246845.11 | 666782.25 |  |  | 22.89 | 63.8 |  |  |
| sp\|A1IGU5\|ARH37\_HUMAN | A1IGU5 | ARH37\_HUMAN | ARHGEF37 | Rho guanine nucleotide exchange factor 37 | 675 | 76.278 | 8.150850e+03 | 7.811228e+05 | 246455.22 | 351169.1 | 183498.52 |  | 18.67 | 51.6 | 28.6 |  |
| sp\|O95081\|AGFG2\_HUMAN | O95081 | AGFG2\_HUMAN | AGFG2 | Arf-GAP domain and FG repeat-containing protein 2 | 481 | 48.963 | 7.947203e+03 | 4.888769e+05 | 246328.3 | 242548.56 |  |  | 8.73 | 34.7 |  |  |
| sp\|Q96B36\|AKTS1\_HUMAN | Q96B36 | AKTS1\_HUMAN | AKT1S1 | Proline-rich AKT1 substrate 1 | 256 | 27.383 | 2.359501e+04 | 8.117427e+05 | 245957.23 | 565785.5 |  |  | 14.84 | 40.2 |  |  |
| sp\|P15882\|CHIN\_HUMAN | P15882 | CHIN\_HUMAN | CHN1 | N-chimaerin | 459 | 50.490 | 3.875433e+03 | 2.458344e+05 | 245834.44 |  |  |  | 2.61 |  |  |  |
| sp\|P11279\|LAMP1\_HUMAN | P11279 | LAMP1\_HUMAN | LAMP1 | Lysosome-associated membrane glycoprotein 1 | 417 | 44.882 | 1.491942e+04 | 8.412815e+05 | 245564.33 | 67885.62 | 208972.89 | 318858.62 | 6.47 | 44.4 | 17.0 | 6.5 |
| sp\|Q9P035\|HACD3\_HUMAN | Q9P035 | HACD3\_HUMAN | HACD3 | Very-long-chain (3R)-3-hydroxyacyl-CoA dehydratase 3 | 362 | 43.160 | 4.019201e+04 | 2.179407e+06 | 244509.4 | 619681.25 | 367143.78 | 948072.6 | 15.47 | 34.8 | 21.3 | 8.8 |
| sp\|Q7L5Y9\|MAEA\_HUMAN | Q7L5Y9 | MAEA\_HUMAN | MAEA | E3 ubiquitin-protein transferase MAEA | 396 | 45.287 | 1.292589e+04 | 7.354468e+05 | 241989.14 | 393563.06 | 99894.63 |  | 17.93 | 63.6 | 34.3 |  |
| sp\|Q9HB07\|MYG1\_HUMAN | Q9HB07 | MYG1\_HUMAN | MYG1 | MYG1 exonuclease | 376 | 42.477 | 1.708235e+04 | 9.116299e+05 | 241829.1 | 669800.8 |  |  | 25.0 | 48.7 |  |  |
| sp\|Q2M2I8\|AAK1\_HUMAN | Q2M2I8 | AAK1\_HUMAN | AAK1 | AP2-associated protein kinase 1 | 961 | 103.885 | 8.515424e+03 | 1.111415e+06 | 241332.27 | 707590.06 | 142554.8 | 19937.889 | 22.58 | 38.5 | 28.5 | 11.6 |
| sp\|Q00765\|REEP5\_HUMAN | Q00765 | REEP5\_HUMAN | REEP5 | Receptor expression-enhancing protein 5 | 189 | 21.493 | 9.496818e+04 | 2.564439e+06 | 241212.9 | 1246042.9 | 17804.781 | 1059378.1 | 16.93 | 41.3 | 10.1 | 11.6 |
| sp\|Q9BYN0\|SRXN1\_HUMAN | Q9BYN0 | SRXN1\_HUMAN | SRXN1 | Sulfiredoxin-1 | 137 | 14.259 | 1.770998e+05 | 3.172665e+06 | 240833.34 | 2348039.0 | 506895.12 | 76897.92 | 40.15 | 88.3 | 62.0 | 21.9 |
| sp\|Q92581\|SL9A6\_HUMAN | Q92581 | SL9A6\_HUMAN | SLC9A6 | Sodium/hydrogen exchanger 6 | 701 | 77.917 | 5.704887e+03 | 5.584656e+05 | 240463.94 |  | 74210.68 | 243791.0 | 8.56 |  | 5.5 | 7.0 |
| sp\|P43378\|PTN9\_HUMAN | P43378 | PTN9\_HUMAN | PTPN9 | Tyrosine-protein phosphatase non-receptor type 9 | 593 | 68.020 | 7.222059e+03 | 6.171842e+05 | 239876.31 | 333590.62 | 43717.293 |  | 15.01 | 35.8 | 38.1 |  |
| sp\|Q6PJ69\|TRI65\_HUMAN | Q6PJ69 | TRI65\_HUMAN | TRIM65 | E3 ubiquitin-protein ligase TRIM65 | 517 | 57.353 | 1.301728e+04 | 9.379800e+05 | 239116.19 | 567223.1 | 131640.67 |  | 16.63 | 53.4 | 32.9 |  |
| sp\|O94830\|DDHD2\_HUMAN | O94830 | DDHD2\_HUMAN | DDHD2 | Phospholipase DDHD2 | 711 | 81.032 | 4.604443e+03 | 4.687603e+05 | 238547.75 | 230212.55 |  |  | 5.06 | 38.3 |  |  |
| sp\|O95870\|ABHGA\_HUMAN | O95870 | ABHGA\_HUMAN | ABHD16A | Phosphatidylserine lipase ABHD16A | 558 | 63.243 | 1.406934e+04 | 1.117901e+06 | 238291.0 | 207084.88 | 131015.44 | 541509.44 | 21.15 | 39.1 | 19.9 | 16.3 |
| sp\|Q92562\|FIG4\_HUMAN | Q92562 | FIG4\_HUMAN | FIG4 | Polyphosphoinositide phosphatase | 907 | 103.635 | 5.495390e+03 | 7.155206e+05 | 237669.81 | 386389.12 | 91461.7 |  | 10.25 | 33.1 | 25.5 |  |
| sp\|Q9UKB1\|FBW1B\_HUMAN | Q9UKB1 | FBW1B\_HUMAN | FBXW11 | F-box/WD repeat-containing protein 11 | 542 | 59.620 | 3.170530e+03 | 2.374876e+05 | 237487.62 |  |  |  | 5.9 |  |  |  |
| sp\|Q6PKG0\|LARP1\_HUMAN | Q6PKG0 | LARP1\_HUMAN | LARP1 | La-related protein 1 | 1096 | 123.510 | 3.834358e+03 | 5.949931e+05 | 236438.34 | 233804.6 | 124750.164 |  | 11.77 | 51.2 | 24.6 |  |
| sp\|Q9HAV4\|XPO5\_HUMAN | Q9HAV4 | XPO5\_HUMAN | XPO5 | Exportin-5 | 1204 | 136.311 | 1.826901e+03 | 3.128694e+05 | 235969.67 | 76899.75 |  |  | 18.52 | 30.8 |  |  |
| sp\|Q8NB37\|GALD1\_HUMAN | Q8NB37 | GALD1\_HUMAN | GATD1 | Glutamine amidotransferase-like class 1 domain-containing protein 1 | 220 | 23.298 | 1.481401e+04 | 4.336193e+05 | 235790.25 | 113782.94 | 84046.07 |  | 32.73 | 24.5 | 14.1 |  |
| sp\|Q9Y617\|SERC\_HUMAN | Q9Y617 | SERC\_HUMAN | PSAT1 | Phosphoserine aminotransferase | 370 | 40.423 | 6.080090e+03 | 3.087847e+05 | 234417.1 | 74367.55 |  |  | 29.73 | 29.2 |  |  |
| sp\|Q8NCG7\|DGLB\_HUMAN | Q8NCG7 | DGLB\_HUMAN | DAGLB | Diacylglycerol lipase-beta | 672 | 73.732 | 1.231578e+04 | 1.140867e+06 | 234413.34 | 78594.555 |  | 827859.6 | 14.88 | 19.3 |  | 12.2 |
| sp\|O75347\|TBCA\_HUMAN | O75347 | TBCA\_HUMAN | TBCA | Tubulin-specific chaperone A | 108 | 12.855 | 6.479378e+05 | 1.046460e+07 | 234289.73 | 7661693.5 | 1682779.4 | 885835.75 | 37.04 | 77.8 | 67.6 | 54.6 |
| sp\|P51580\|TPMT\_HUMAN | P51580 | TPMT\_HUMAN | TPMT | Thiopurine S-methyltransferase | 245 | 28.180 | 6.842736e+04 | 2.422634e+06 | 234039.25 | 2042078.8 | 146516.45 |  | 10.2 | 68.6 | 29.0 |  |
| sp\|Q9H910\|JUPI2\_HUMAN | Q9H910 | JUPI2\_HUMAN | JPT2 | Jupiter microtubule associated homolog 2 | 190 | 20.063 | 2.215550e+04 | 5.584633e+05 | 233943.38 | 86723.984 | 160122.88 | 77673.01 | 35.79 | 75.3 | 54.2 | 21.1 |
| sp\|Q96H79\|ZCCHL\_HUMAN | Q96H79 | ZCCHL\_HUMAN | ZC3HAV1L | Zinc finger CCCH-type antiviral protein 1-like | 300 | 32.962 | 2.241940e+04 | 9.284417e+05 | 233896.05 | 567605.1 | 126940.55 |  | 21.33 | 52.7 | 19.3 |  |
| sp\|Q9H2P0\|ADNP\_HUMAN | Q9H2P0 | ADNP\_HUMAN | ADNP | Activity-dependent neuroprotector homeobox protein | 1102 | 121.220 | 1.533891e+03 | 2.336070e+05 | 233607.03 |  |  |  | 2.72 |  |  |  |
| sp\|Q2LD37\|BLTP1\_HUMAN | Q2LD37 | BLTP1\_HUMAN | BLTP1 | Bridge-like lipid transfer protein family member 1 | 5005 | 555.482 | 8.349841e+03 | 5.827272e+06 | 233296.47 | 351701.88 | 1127380.9 | 4114893.2 | 1.06 | 27.5 | 20.1 | 11.5 |
| sp\|P53609\|PGTB1\_HUMAN | P53609 | PGTB1\_HUMAN | PGGT1B | Geranylgeranyl transferase type-1 subunit beta | 377 | 42.368 | 7.213280e+03 | 3.839617e+05 | 233082.52 | 150879.2 |  |  | 18.3 | 22.5 |  |  |
| sp\|Q9P0M2\|AKA7G\_HUMAN | Q9P0M2 | AKA7G\_HUMAN | AKAP7 | A-kinase anchor protein 7 isoform gamma | 348 | 39.518 | 1.379411e+04 | 6.848663e+05 | 233014.25 | 351607.8 | 100244.234 |  | 18.97 | 51.4 | 22.7 |  |
| sp\|O75061\|AUXI\_HUMAN | O75061 | AUXI\_HUMAN | DNAJC6 | Putative tyrosine-protein phosphatase auxilin | 913 | 99.997 | 9.258039e+03 | 1.163116e+06 | 232984.31 | 743529.9 | 109373.914 | 77228.16 | 9.09 | 32.4 | 15.3 | 7.7 |
| sp\|Q9UKX2\|MYH2\_HUMAN | Q9UKX2 | MYH2\_HUMAN | MYH2 | Myosin-2 | 1941 | 223.044 | 1.781707e+04 | 4.992799e+06 | 232970.39 | 2020358.1 | 1924450.6 | 815019.6 | 16.43 | 44.1 | 35.8 | 16.9 |
| sp\|Q9NRH2\|SNRK\_HUMAN | Q9NRH2 | SNRK\_HUMAN | SNRK | SNF-related serine/threonine-protein kinase | 765 | 84.276 | 2.735209e+03 | 2.896087e+05 | 232710.75 | 56897.984 |  |  | 3.53 | 23.9 |  |  |
| sp\|Q5MNZ9\|WIPI1\_HUMAN | Q5MNZ9 | WIPI1\_HUMAN | WIPI1 | WD repeat domain phosphoinositide-interacting protein 1 | 446 | 48.673 | 1.175454e+04 | 7.188048e+05 | 231476.9 | 487327.88 |  |  | 17.94 | 40.8 |  |  |
| sp\|P29218\|IMPA1\_HUMAN | P29218 | IMPA1\_HUMAN | IMPA1 | Inositol monophosphatase 1 | 277 | 30.189 | 4.058106e+04 | 1.539179e+06 | 230630.62 | 836617.25 | 326614.56 | 145316.92 | 16.25 | 47.3 | 36.1 | 22.7 |
| sp\|Q92769\|HDAC2\_HUMAN | Q92769 | HDAC2\_HUMAN | HDAC2 | Histone deacetylase 2 | 488 | 55.364 | 3.486433e+04 | 2.425079e+06 | 230553.23 | 2194525.8 |  |  | 19.26 | 46.7 |  |  |
| sp\|Q9UJX2\|CDC23\_HUMAN | Q9UJX2 | CDC23\_HUMAN | CDC23 | Cell division cycle protein 23 homolog | 597 | 68.834 | 9.408297e+03 | 8.136379e+05 | 230290.38 | 540388.7 | 42958.812 |  | 20.27 | 40.4 | 18.6 |  |
| sp\|Q2KHT3\|CL16A\_HUMAN | Q2KHT3 | CL16A\_HUMAN | CLEC16A | Protein CLEC16A | 1053 | 117.715 | 2.805618e+03 | 4.149326e+05 | 229916.0 | 185016.61 |  |  | 9.88 | 27.7 |  |  |
| sp\|P30520\|PURA2\_HUMAN | P30520 | PURA2\_HUMAN | ADSS2 | Adenylosuccinate synthetase isozyme 2 | 456 | 50.097 | 6.592015e+03 | 4.149035e+05 | 229047.89 | 185855.58 |  |  | 17.32 | 36.4 |  |  |
| sp\|Q96S97\|MYADM\_HUMAN | Q96S97 | MYADM\_HUMAN | MYADM | Myeloid-associated differentiation marker | 322 | 35.274 | 2.279095e+04 | 1.010030e+06 | 228456.31 |  | 506734.75 | 274839.03 | 10.25 |  | 15.8 | 12.4 |
| sp\|P27986\|P85A\_HUMAN | P27986 | P85A\_HUMAN | PIK3R1 | Phosphatidylinositol 3-kinase regulatory subunit alpha | 724 | 83.598 | 3.975510e+03 | 4.175475e+05 | 227976.47 | 189571.02 |  |  | 21.41 | 15.2 |  |  |
| sp\|P50402\|EMD\_HUMAN | P50402 | EMD\_HUMAN | EMD | Emerin | 254 | 28.994 | 9.112042e+04 | 3.319258e+06 | 227563.81 | 568202.8 | 501075.94 | 2022415.4 | 40.16 | 40.2 | 20.1 | 23.2 |
| sp\|P21283\|VATC1\_HUMAN | P21283 | VATC1\_HUMAN | ATP6V1C1 | V-type proton ATPase subunit C 1 | 382 | 43.942 | 1.326064e+04 | 7.320851e+05 | 227196.06 | 421152.3 | 83736.77 |  | 22.51 | 44.8 | 23.6 |  |
| sp\|P49356\|FNTB\_HUMAN | P49356 | FNTB\_HUMAN | FNTB | Protein farnesyltransferase subunit beta | 437 | 48.774 | 1.174380e+04 | 7.196379e+05 | 226854.88 | 492783.03 |  |  | 9.15 | 15.3 |  |  |
| sp\|P36551\|HEM6\_HUMAN | P36551 | HEM6\_HUMAN | CPOX | Oxygen-dependent coproporphyrinogen-III oxidase, mitochondrial | 454 | 50.152 | 2.022154e+04 | 1.274147e+06 | 226417.86 | 857901.6 | 189827.55 |  | 32.38 | 52.0 | 45.8 |  |
| sp\|P40763\|STAT3\_HUMAN | P40763 | STAT3\_HUMAN | STAT3 | Signal transducer and activator of transcription 3 | 770 | 88.068 | 3.289247e+03 | 3.639417e+05 | 225776.39 | 138165.31 |  |  | 20.78 | 38.2 |  |  |
| sp\|O95163\|ELP1\_HUMAN | O95163 | ELP1\_HUMAN | ELP1 | Elongator complex protein 1 | 1332 | 150.254 | 2.694126e+03 | 5.085822e+05 | 224555.03 | 284027.16 |  |  | 22.15 | 29.1 |  |  |
| sp\|P61962\|DCAF7\_HUMAN | P61962 | DCAF7\_HUMAN | DCAF7 | DDB1- and CUL4-associated factor 7 | 342 | 38.926 | 1.198743e+04 | 5.862504e+05 | 224358.5 | 202196.61 | 159695.25 |  | 20.76 | 41.2 | 18.4 |  |
| sp\|Q9NPF4\|OSGEP\_HUMAN | Q9NPF4 | OSGEP\_HUMAN | OSGEP | tRNA N6-adenosine threonylcarbamoyltransferase | 335 | 36.427 | 8.143253e+03 | 3.726821e+05 | 224118.06 | 148564.02 |  |  | 16.72 | 38.2 |  |  |
| sp\|Q14689\|DIP2A\_HUMAN | Q14689 | DIP2A\_HUMAN | DIP2A | Disco-interacting protein 2 homolog A | 1571 | 170.369 | 1.648446e+03 | 3.528438e+05 | 224113.1 | 128730.734 |  |  | 13.18 | 33.7 |  |  |
| sp\|P01859\|IGHG2\_HUMAN | P01859 | IGHG2\_HUMAN | IGHG2 | Immunoglobulin heavy constant gamma 2 | 395 | 43.806 | 4.298522e+04 | 2.365756e+06 | 223802.4 | 413267.4 | 1728686.2 |  | 17.72 | 34.7 | 23.0 |  |
| sp\|Q16576\|RBBP7\_HUMAN | Q16576 | RBBP7\_HUMAN | RBBP7 | Histone-binding protein RBBP7 | 425 | 47.820 | 7.522024e+03 | 4.519199e+05 | 223755.16 | 228164.73 |  |  | 16.24 | 26.1 |  |  |
| sp\|Q9Y5P4\|CERT\_HUMAN | Q9Y5P4 | CERT\_HUMAN | CERT1 | Ceramide transfer protein | 624 | 70.835 | 6.220765e+03 | 5.536163e+05 | 223714.17 | 307634.1 | 22268.0 |  | 25.0 | 38.0 | 5.6 |  |
| sp\|Q13442\|HAP28\_HUMAN | Q13442 | HAP28\_HUMAN | PDAP1 | 28 kDa heat- and acid-stable phosphoprotein | 181 | 20.630 | 4.104476e+04 | 1.063835e+06 | 223586.1 | 653976.0 | 125705.45 | 60566.957 | 37.02 | 53.6 | 37.0 | 34.8 |
| sp\|P42338\|PK3CB\_HUMAN | P42338 | PK3CB\_HUMAN | PIK3CB | Phosphatidylinositol 4,5-bisphosphate 3-kinase catalytic subunit beta isoform | 1070 | 122.762 | 3.598743e+03 | 5.550498e+05 | 221725.31 | 263674.5 | 21273.258 | 48376.758 | 15.7 | 39.6 | 9.5 | 9.0 |
| sp\|P12273\|PIP\_HUMAN | P12273 | PIP\_HUMAN | PIP | Prolactin-inducible protein | 146 | 16.572 | 2.400836e+04 | 4.998670e+05 | 221395.05 | 54788.188 | 223683.78 |  | 24.66 | 28.8 | 22.6 |  |
| sp\|Q8WZ82\|OVCA2\_HUMAN | Q8WZ82 | OVCA2\_HUMAN | OVCA2 | Esterase OVCA2 | 227 | 24.418 | 2.895759e+04 | 8.883612e+05 | 221337.83 | 667023.4 |  |  | 40.97 | 65.6 |  |  |
| sp\|Q9Y2A7\|NCKP1\_HUMAN | Q9Y2A7 | NCKP1\_HUMAN | NCKAP1 | Nck-associated protein 1 | 1128 | 128.790 | 1.499285e+03 | 2.425959e+05 | 220849.4 | 21746.465 |  |  | 3.1 | 32.0 |  |  |
| sp\|Q9UKV3\|ACINU\_HUMAN | Q9UKV3 | ACINU\_HUMAN | ACIN1 | Apoptotic chromatin condensation inducer in the nucleus | 1341 | 151.862 | 1.191181e+04 | 2.272710e+06 | 220285.39 | 1705210.8 | 347214.2 |  | 7.01 | 39.6 | 22.5 |  |
| sp\|P11055\|MYH3\_HUMAN | P11055 | MYH3\_HUMAN | MYH3 | Myosin-3 | 1940 | 213.400 | 8.199092e+02 | 2.198251e+05 | 219825.12 |  |  |  | 10.21 |  |  |  |
| sp\|Q96Q42\|ALS2\_HUMAN | Q96Q42 | ALS2\_HUMAN | ALS2 | Alsin | 1657 | 183.634 | 1.608456e+03 | 3.710902e+05 | 219201.84 | 151888.36 |  |  | 6.94 | 21.4 |  |  |
| sp\|Q9H5X1\|CIA2A\_HUMAN | Q9H5X1 | CIA2A\_HUMAN | CIAO2A | Cytosolic iron-sulfur assembly component 2A | 160 | 18.355 | 7.658679e+04 | 1.766141e+06 | 218519.0 | 1411153.8 | 78152.42 | 58315.902 | 23.75 | 44.4 | 13.8 | 6.9 |
| sp\|P30040\|ERP29\_HUMAN | P30040 | ERP29\_HUMAN | ERP29 | Endoplasmic reticulum resident protein 29 | 261 | 28.993 | 6.650475e+04 | 2.422495e+06 | 217862.94 | 714957.1 | 1344139.5 | 145535.8 | 18.01 | 47.1 | 42.9 | 22.2 |
| sp\|Q8N766\|EMC1\_HUMAN | Q8N766 | EMC1\_HUMAN | EMC1 | ER membrane protein complex subunit 1 | 993 | 111.759 | 5.269147e+03 | 7.398437e+05 | 217645.5 |  | 420488.4 | 101709.83 | 4.63 |  | 17.8 | 15.4 |
| sp\|P36405\|ARL3\_HUMAN | P36405 | ARL3\_HUMAN | ARL3 | ADP-ribosylation factor-like protein 3 | 182 | 20.456 | 2.345022e+04 | 6.026771e+05 | 217524.05 | 385153.06 |  |  | 43.96 | 56.0 |  |  |
| sp\|Q6PL24\|TMED8\_HUMAN | Q6PL24 | TMED8\_HUMAN | TMED8 | Protein TMED8 | 325 | 35.740 | 7.030458e+03 | 3.156860e+05 | 216913.66 | 70439.04 | 28333.305 |  | 27.69 | 37.2 | 12.3 |  |
| sp\|Q8N1G4\|LRC47\_HUMAN | Q8N1G4 | LRC47\_HUMAN | LRRC47 | Leucine-rich repeat-containing protein 47 | 583 | 63.473 | 8.174319e+03 | 6.518652e+05 | 215869.62 | 366037.47 | 60972.57 | 8985.518 | 29.33 | 51.1 | 18.2 | 7.5 |
| sp\|O15145\|ARPC3\_HUMAN | O15145 | ARPC3\_HUMAN | ARPC3 | Actin-related protein 2/3 complex subunit 3 | 178 | 20.547 | 3.943922e+04 | 1.018108e+06 | 215573.84 | 620896.4 | 181637.88 |  | 23.6 | 51.1 | 32.0 |  |
| sp\|P40926\|MDHM\_HUMAN | P40926 | MDHM\_HUMAN | MDH2 | Malate dehydrogenase, mitochondrial | 338 | 35.503 | 4.456960e+04 | 1.988021e+06 | 215234.12 | 548436.4 | 902070.6 | 322279.97 | 16.57 | 60.4 | 40.8 | 37.0 |
| sp\|O00139\|KIF2A\_HUMAN | O00139 | KIF2A\_HUMAN | KIF2A | Kinesin-like protein KIF2A | 706 | 79.955 | 5.181931e+04 | 5.205403e+06 | 214863.69 | 4981073.5 | 9466.137 |  | 5.67 | 31.4 | 16.1 |  |
| sp\|P0C0L5\|CO4B\_HUMAN | P0C0L5 | CO4B\_HUMAN | C4B | Complement C4-B | 1744 | 192.751 | 1.690680e+04 | 4.094257e+06 | 214229.88 |  | 3095308.0 | 784719.44 | 5.05 |  | 23.1 | 13.8 |
| sp\|Q6B0K9\|HBM\_HUMAN | Q6B0K9 | HBM\_HUMAN | HBM | Hemoglobin subunit mu | 141 | 15.618 | 1.503650e+05 | 2.950456e+06 | 213474.86 | 2010323.8 | 568440.2 | 158217.47 | 21.99 | 65.2 | 31.2 | 6.4 |
| sp\|Q969X5\|ERGI1\_HUMAN | Q969X5 | ERGI1\_HUMAN | ERGIC1 | Endoplasmic reticulum-Golgi intermediate compartment protein 1 | 290 | 32.592 | 4.028908e+04 | 1.649740e+06 | 212650.6 | 266213.5 | 596810.9 | 574064.94 | 24.83 | 29.7 | 39.3 | 20.0 |
| sp\|O15247\|CLIC2\_HUMAN | O15247 | CLIC2\_HUMAN | CLIC2 | Chloride intracellular channel protein 2 | 247 | 28.356 | 7.479524e+04 | 2.664625e+06 | 212641.28 | 2372536.0 | 79447.72 |  | 47.37 | 49.8 | 33.2 |  |
| sp\|P62256\|UBE2H\_HUMAN | P62256 | UBE2H\_HUMAN | UBE2H | Ubiquitin-conjugating enzyme E2 H | 183 | 20.655 | 2.469539e+04 | 6.408527e+05 | 212241.06 | 428611.66 |  |  | 25.68 | 36.1 |  |  |
| sp\|Q9H0J9\|PAR12\_HUMAN | Q9H0J9 | PAR12\_HUMAN | PARP12 | Protein mono-ADP-ribosyltransferase PARP12 | 701 | 79.064 | 2.980268e+03 | 2.960406e+05 | 212204.08 | 63452.91 | 20383.562 |  | 13.27 | 39.2 | 13.6 |  |
| sp\|Q9Y4B6\|DCAF1\_HUMAN | Q9Y4B6 | DCAF1\_HUMAN | DCAF1 | DDB1- and CUL4-associated factor 1 | 1507 | 169.007 | 2.648726e+03 | 5.624176e+05 | 211776.58 | 350641.0 |  |  | 13.14 | 29.9 |  |  |
| sp\|Q9H8W4\|PKHF2\_HUMAN | Q9H8W4 | PKHF2\_HUMAN | PLEKHF2 | Pleckstrin homology domain-containing family F member 2 | 249 | 27.798 | 7.419957e+04 | 2.591386e+06 | 211734.7 | 2144806.5 | 234844.86 |  | 23.29 | 61.4 | 32.9 |  |
| sp\|Q5SRE7\|PHYD1\_HUMAN | Q5SRE7 | PHYD1\_HUMAN | PHYHD1 | Phytanoyl-CoA dioxygenase domain-containing protein 1 | 291 | 32.411 | 9.522190e+03 | 3.877452e+05 | 211395.44 | 125201.39 | 51148.355 |  | 19.59 | 45.4 | 31.6 |  |
| sp\|Q9UK76\|JUPI1\_HUMAN | Q9UK76 | JUPI1\_HUMAN | JPT1 | Jupiter microtubule associated homolog 1 | 154 | 16.940 | 9.895894e+03 | 2.106132e+05 | 210613.2 |  |  |  | 33.77 |  |  |  |
| sp\|Q16222\|UAP1\_HUMAN | Q16222 | UAP1\_HUMAN | UAP1 | UDP-N-acetylhexosamine pyrophosphorylase | 522 | 58.769 | 5.970011e+03 | 4.407990e+05 | 210282.53 | 230516.45 |  |  | 22.41 | 51.5 |  |  |
| sp\|P53992\|SC24C\_HUMAN | P53992 | SC24C\_HUMAN | SEC24C | Protein transport protein Sec24C | 1094 | 118.325 | 3.212811e+03 | 4.776160e+05 | 210124.02 | 138660.1 | 97202.44 | 31629.465 | 6.76 | 31.1 | 16.7 | 11.9 |
| sp\|P33121\|ACSL1\_HUMAN | P33121 | ACSL1\_HUMAN | ACSL1 | Long-chain-fatty-acid--CoA ligase 1 | 698 | 76.780 | 2.167948e+03 | 2.091290e+05 | 209128.95 |  |  |  | 6.59 |  |  |  |
| sp\|O60486\|PLXC1\_HUMAN | O60486 | PLXC1\_HUMAN | PLXNC1 | Plexin-C1 | 1568 | 172.480 | 9.650316e+02 | 2.091209e+05 | 209120.9 |  |  |  | 1.59 |  |  |  |
| sp\|Q9Y4K4\|M4K5\_HUMAN | Q9Y4K4 | M4K5\_HUMAN | MAP4K5 | Mitogen-activated protein kinase kinase kinase kinase 5 | 846 | 95.024 | 3.764952e+03 | 4.494796e+05 | 209020.66 | 183466.8 | 56992.1 |  | 16.31 | 37.5 | 17.0 |  |
| sp\|O15357\|SHIP2\_HUMAN | O15357 | SHIP2\_HUMAN | INPPL1 | Phosphatidylinositol 3,4,5-trisphosphate 5-phosphatase 2 | 1258 | 138.599 | 8.507773e+03 | 1.481471e+06 | 208425.98 | 1147746.8 | 125298.26 |  | 18.28 | 28.6 | 27.3 |  |
| sp\|Q96SW2\|CRBN\_HUMAN | Q96SW2 | CRBN\_HUMAN | CRBN | Protein cereblon | 442 | 50.546 | 3.413647e+03 | 2.167816e+05 | 208412.62 |  | 8369.024 |  | 19.46 |  | 12.7 |  |
| sp\|P06576\|ATPB\_HUMAN | P06576 | ATPB\_HUMAN | ATP5F1B | ATP synthase subunit beta, mitochondrial | 529 | 56.560 | 1.567615e+04 | 1.113951e+06 | 208185.86 | 484056.1 | 288087.56 | 133621.39 | 31.0 | 62.2 | 28.9 | 19.8 |
| sp\|Q6IQ20\|NAPEP\_HUMAN | Q6IQ20 | NAPEP\_HUMAN | NAPEPLD | N-acyl-phosphatidylethanolamine-hydrolyzing phospholipase D | 393 | 45.596 | 8.618793e+03 | 4.937309e+05 | 208135.72 | 285595.22 |  |  | 17.05 | 58.3 |  |  |
| sp\|O14745\|NHRF1\_HUMAN | O14745 | NHRF1\_HUMAN | NHERF1 | Na(+)/H(+) exchange regulatory cofactor NHE-RF1 | 358 | 38.868 | 1.856303e+04 | 9.064802e+05 | 207474.56 | 583188.8 | 98615.21 | 17201.596 | 23.18 | 69.3 | 34.4 | 10.3 |
| sp\|Q9GZL7\|WDR12\_HUMAN | Q9GZL7 | WDR12\_HUMAN | WDR12 | Ribosome biogenesis protein WDR12 | 423 | 47.708 | 4.421924e+03 | 2.650450e+05 | 207197.02 | 57847.97 |  |  | 13.24 | 30.0 |  |  |
| sp\|Q15436\|SC23A\_HUMAN | Q15436 | SC23A\_HUMAN | SEC23A | Protein transport protein Sec23A | 765 | 86.161 | 1.006610e+04 | 1.089655e+06 | 206718.5 | 488518.16 | 394418.16 |  | 16.73 | 41.4 | 22.1 |  |
| sp\|Q96Q45\|TM237\_HUMAN | Q96Q45 | TM237\_HUMAN | TMEM237 | Transmembrane protein 237 | 408 | 45.526 | 6.512412e+04 | 3.724934e+06 | 206675.2 |  | 1154478.6 | 2363779.8 | 22.79 |  | 30.4 | 16.7 |
| sp\|Q53GQ0\|DHB12\_HUMAN | Q53GQ0 | DHB12\_HUMAN | HSD17B12 | Very-long-chain 3-oxoacyl-CoA reductase | 312 | 34.324 | 5.989811e+04 | 2.583022e+06 | 206591.84 | 1102899.1 | 911848.1 | 361683.28 | 38.78 | 52.2 | 27.6 | 15.4 |
| sp\|Q9UBW5\|BIN2\_HUMAN | Q9UBW5 | BIN2\_HUMAN | BIN2 | Bridging integrator 2 | 565 | 61.874 | 4.809495e+03 | 3.738736e+05 | 205263.84 | 140666.22 | 27943.574 |  | 11.33 | 33.3 | 10.4 |  |
| sp\|O60234\|GMFG\_HUMAN | O60234 | GMFG\_HUMAN | GMFG | Glia maturation factor gamma | 142 | 16.801 | 1.356337e+05 | 2.862990e+06 | 204857.95 | 2159055.8 | 258575.58 | 240500.97 | 28.17 | 57.7 | 40.1 | 20.4 |
| sp\|A5LHX3\|PSB11\_HUMAN | A5LHX3 | PSB11\_HUMAN | PSMB11 | Proteasome subunit beta type-11 | 300 | 33.000 | 4.937316e+03 | 2.047020e+05 | 204701.98 |  |  |  | 8.67 |  |  |  |
| sp\|B7ZAP0\|RBG10\_HUMAN | B7ZAP0 | RBG10\_HUMAN | RABGAP1L | Rab GTPase-activating protein 1-like, isoform 10 | 253 | 29.038 | 1.145464e+04 | 4.178932e+05 | 204423.92 | 213469.33 |  |  | 19.37 | 49.0 |  |  |
| sp\|P35558\|PCKGC\_HUMAN | P35558 | PCKGC\_HUMAN | PCK1 | Phosphoenolpyruvate carboxykinase, cytosolic [GTP] | 622 | 68.420 | 2.367479e+03 | 2.035102e+05 | 203510.25 |  |  |  | 5.14 |  |  |  |
| sp\|Q96S82\|UBL7\_HUMAN | Q96S82 | UBL7\_HUMAN | UBL7 | Ubiquitin-like protein 7 | 380 | 40.510 | 8.964028e+03 | 4.562287e+05 | 202135.27 | 194833.1 | 59260.34 |  | 18.95 | 19.7 | 8.9 |  |
| sp\|Q9Y6D5\|BIG2\_HUMAN | Q9Y6D5 | BIG2\_HUMAN | ARFGEF2 | Brefeldin A-inhibited guanine nucleotide-exchange protein 2 | 1785 | 202.038 | 3.583807e+03 | 9.096929e+05 | 202033.81 | 467657.56 |  | 240001.56 | 12.94 | 41.5 |  | 9.1 |
| sp\|Q96C86\|DCPS\_HUMAN | Q96C86 | DCPS\_HUMAN | DCPS | m7GpppX diphosphatase | 337 | 38.609 | 5.733068e+03 | 2.780948e+05 | 201936.94 | 76157.84 |  |  | 22.85 | 23.1 |  |  |
| sp\|Q96ER3\|SAAL1\_HUMAN | Q96ER3 | SAAL1\_HUMAN | SAAL1 | Protein SAAL1 | 474 | 53.558 | 1.015698e+04 | 6.834488e+05 | 200816.25 | 482632.5 |  |  | 14.14 | 16.7 |  |  |
| sp\|Q9H1A4\|APC1\_HUMAN | Q9H1A4 | APC1\_HUMAN | ANAPC1 | Anaphase-promoting complex subunit 1 | 1944 | 216.500 | 1.659065e+03 | 4.512720e+05 | 200799.98 | 250472.06 |  |  | 10.49 | 32.2 |  |  |
| sp\|O00221\|IKBE\_HUMAN | O00221 | IKBE\_HUMAN | NFKBIE | NF-kappa-B inhibitor epsilon | 500 | 55.000 | 2.905590e+03 | 2.007771e+05 | 200777.08 |  |  |  | 2.2 |  |  |  |
| sp\|O75558\|STX11\_HUMAN | O75558 | STX11\_HUMAN | STX11 | Syntaxin-11 | 287 | 33.196 | 4.234753e+04 | 1.766164e+06 | 200677.34 | 488027.1 | 656713.4 | 420746.0 | 33.8 | 51.9 | 48.8 | 31.7 |
| sp\|Q9NZ52\|GGA3\_HUMAN | Q9NZ52 | GGA3\_HUMAN | GGA3 | ADP-ribosylation factor-binding protein GGA3 | 723 | 78.315 | 9.774635e+03 | 9.617510e+05 | 200575.31 | 472959.25 | 241604.77 | 46611.688 | 14.8 | 28.8 | 22.8 | 7.2 |
| sp\|Q9C0B1\|FTO\_HUMAN | Q9C0B1 | FTO\_HUMAN | FTO | Alpha-ketoglutarate-dependent dioxygenase FTO | 505 | 58.282 | 6.075881e+03 | 4.448984e+05 | 200489.97 | 244408.45 |  |  | 24.55 | 40.2 |  |  |
| sp\|Q86V88\|MGDP1\_HUMAN | Q86V88 | MGDP1\_HUMAN | MDP1 | Magnesium-dependent phosphatase 1 | 176 | 20.109 | 4.242536e+04 | 1.071848e+06 | 199700.06 | 372228.4 | 475557.94 | 24361.564 | 13.07 | 71.0 | 19.3 | 7.4 |
| sp\|Q9Y6D6\|BIG1\_HUMAN | Q9Y6D6 | BIG1\_HUMAN | ARFGEF1 | Brefeldin A-inhibited guanine nucleotide-exchange protein 1 | 1849 | 208.767 | 3.345792e+03 | 8.775622e+05 | 199693.16 | 605034.0 |  | 72835.0 | 16.12 | 31.1 |  | 9.1 |
| sp\|Q9H2J4\|PDCL3\_HUMAN | Q9H2J4 | PDCL3\_HUMAN | PDCL3 | Phosducin-like protein 3 | 239 | 27.614 | 3.020559e+04 | 1.047934e+06 | 199443.31 | 815889.8 | 32600.725 |  | 24.69 | 42.7 | 17.6 |  |
| sp\|Q4KMP7\|TB10B\_HUMAN | Q4KMP7 | TB10B\_HUMAN | TBC1D10B | TBC1 domain family member 10B | 808 | 87.199 | 8.127721e+03 | 8.904252e+05 | 199059.88 | 411982.7 | 181896.67 | 97485.99 | 15.22 | 35.3 | 20.9 | 11.8 |
| sp\|P15056\|BRAF\_HUMAN | P15056 | BRAF\_HUMAN | BRAF | Serine/threonine-protein kinase B-raf | 766 | 84.437 | 5.699687e+03 | 6.046457e+05 | 198972.81 | 358696.8 | 46976.105 |  | 11.23 | 23.0 | 15.7 |  |
| sp\|P32856\|STX2\_HUMAN | P32856 | STX2\_HUMAN | STX2 | Syntaxin-2 | 288 | 33.341 | 5.097870e+04 | 2.135426e+06 | 198915.61 | 336891.38 | 574390.5 | 1025228.6 | 26.04 | 27.4 | 34.7 | 29.9 |
| sp\|A6NHY2\|AKD1B\_HUMAN | A6NHY2 | AKD1B\_HUMAN | ANKDD1B | Ankyrin repeat and death domain-containing protein 1B | 528 | 58.080 | 2.725200e+03 | 1.988576e+05 | 198857.61 |  |  |  | 2.08 |  |  |  |
| sp\|Q13724\|MOGS\_HUMAN | Q13724 | MOGS\_HUMAN | MOGS | Mannosyl-oligosaccharide glucosidase | 837 | 91.918 | 1.762415e+04 | 2.035288e+06 | 198135.69 | 244768.19 | 1335471.8 | 256911.94 | 11.11 | 34.9 | 31.8 | 17.7 |
| sp\|P60983\|GMFB\_HUMAN | P60983 | GMFB\_HUMAN | GMFB | Glia maturation factor beta | 142 | 16.713 | 1.147663e+05 | 2.409827e+06 | 197938.89 | 1818431.6 | 393456.3 |  | 26.06 | 64.1 | 44.4 |  |
| sp\|P25705\|ATPA\_HUMAN | P25705 | ATPA\_HUMAN | ATP5F1A | ATP synthase subunit alpha, mitochondrial | 553 | 59.751 | 1.995893e+04 | 1.498303e+06 | 196785.95 | 510621.4 | 432676.06 | 358219.66 | 15.91 | 46.5 | 34.2 | 21.7 |
| sp\|Q8NC96\|NECP1\_HUMAN | Q8NC96 | NECP1\_HUMAN | NECAP1 | Adaptin ear-binding coat-associated protein 1 | 275 | 29.737 | 1.079919e+04 | 4.034646e+05 | 196773.19 | 166348.03 | 40343.383 |  | 22.55 | 36.7 | 20.0 |  |
| sp\|Q9NXU5\|ARL15\_HUMAN | Q9NXU5 | ARL15\_HUMAN | ARL15 | ADP-ribosylation factor-like protein 15 | 204 | 22.876 | 3.410827e+04 | 9.802954e+05 | 196373.19 | 222112.11 | 191438.6 | 370371.53 | 21.08 | 36.3 | 21.6 | 10.8 |
| sp\|P38646\|GRP75\_HUMAN | P38646 | GRP75\_HUMAN | HSPA9 | Stress-70 protein, mitochondrial | 679 | 73.680 | 1.477249e+04 | 1.367478e+06 | 195807.39 |  | 814719.06 | 356951.66 | 7.66 |  | 33.1 | 15.8 |
| sp\|O75475\|PSIP1\_HUMAN | O75475 | PSIP1\_HUMAN | PSIP1 | PC4 and SFRS1-interacting protein | 530 | 58.300 | 2.673223e+03 | 1.958037e+05 | 195803.72 |  |  |  | 3.96 |  |  |  |
| sp\|Q9H4A6\|GOLP3\_HUMAN | Q9H4A6 | GOLP3\_HUMAN | GOLPH3 | Golgi phosphoprotein 3 | 298 | 33.811 | 1.413794e+04 | 6.005669e+05 | 195712.36 | 397648.12 | 7206.3887 |  | 17.79 | 38.6 | 29.5 |  |
| sp\|Q99933\|BAG1\_HUMAN | Q99933 | BAG1\_HUMAN | BAG1 | BAG family molecular chaperone regulator 1 | 345 | 38.779 | 8.466551e+04 | 4.124965e+06 | 195109.38 | 3745792.8 | 132322.95 | 51740.02 | 20.87 | 38.3 | 25.5 | 14.2 |
| sp\|Q9BXL5\|HEMGN\_HUMAN | Q9BXL5 | HEMGN\_HUMAN | HEMGN | Hemogen | 484 | 55.341 | 6.303430e+03 | 4.382693e+05 | 194240.48 | 244028.83 |  |  | 8.68 | 34.9 |  |  |
| sp\|P04920\|B3A2\_HUMAN | P04920 | B3A2\_HUMAN | SLC4A2 | Anion exchange protein 2 | 1241 | 137.009 | 9.661226e+03 | 1.663024e+06 | 193700.12 | 50788.504 | 355277.44 | 1063257.9 | 12.49 | 29.9 | 22.4 | 21.0 |
| sp\|Q9NUU7\|DD19A\_HUMAN | Q9NUU7 | DD19A\_HUMAN | DDX19A | ATP-dependent RNA helicase DDX19A | 478 | 53.975 | 6.419906e+03 | 4.353499e+05 | 192786.95 | 152301.38 | 90261.58 |  | 14.64 | 42.9 | 19.0 |  |
| sp\|Q00839\|HNRPU\_HUMAN | Q00839 | HNRPU\_HUMAN | HNRNPU | Heterogeneous nuclear ribonucleoprotein U | 825 | 90.584 | 3.503421e+03 | 3.987135e+05 | 192420.31 | 166852.44 | 13297.349 | 26143.434 | 8.0 | 38.8 | 30.3 | 14.4 |
| sp\|Q12792\|TWF1\_HUMAN | Q12792 | TWF1\_HUMAN | TWF1 | Twinfilin-1 | 350 | 40.283 | 1.560203e+04 | 7.896237e+05 | 191390.97 | 517888.78 | 80343.94 |  | 9.43 | 42.0 | 32.0 |  |
| sp\|P42765\|THIM\_HUMAN | P42765 | THIM\_HUMAN | ACAA2 | 3-ketoacyl-CoA thiolase, mitochondrial | 397 | 41.924 | 5.993500e+03 | 3.156897e+05 | 191285.78 | 83571.91 | 40831.996 |  | 11.84 | 45.8 | 14.6 |  |
| sp\|Q9BZF1\|OSBL8\_HUMAN | Q9BZF1 | OSBL8\_HUMAN | OSBPL8 | Oxysterol-binding protein-related protein 8 | 889 | 101.196 | 8.839794e+03 | 1.123887e+06 | 190547.78 | 265847.22 | 138476.64 | 529015.3 | 12.37 | 35.8 | 17.0 | 15.5 |
| sp\|Q12981\|SEC20\_HUMAN | Q12981 | SEC20\_HUMAN | BNIP1 | Vesicle transport protein SEC20 | 228 | 26.132 | 9.600291e+04 | 3.151913e+06 | 190441.89 | 590227.8 | 583963.06 | 1787280.2 | 34.65 | 50.0 | 40.4 | 28.1 |
| sp\|Q9ULZ3\|ASC\_HUMAN | Q9ULZ3 | ASC\_HUMAN | PYCARD | Apoptosis-associated speck-like protein containing a CARD | 195 | 21.627 | 9.749210e+03 | 2.649006e+05 | 190393.75 | 34159.117 | 40347.688 |  | 18.97 | 32.3 | 19.0 |  |
| sp\|P19174\|PLCG1\_HUMAN | P19174 | PLCG1\_HUMAN | PLCG1 | 1-phosphatidylinositol 4,5-bisphosphate phosphodiesterase gamma-1 | 1290 | 148.532 | 1.452841e+03 | 2.711161e+05 | 189897.4 | 81218.7 |  |  | 15.66 | 25.8 |  |  |
| sp\|Q8N0X7\|SPART\_HUMAN | Q8N0X7 | SPART\_HUMAN | SPART | Spartin | 666 | 72.833 | 1.217148e+04 | 1.113752e+06 | 189777.19 | 720727.44 | 191348.38 | 11899.397 | 15.47 | 26.9 | 24.9 | 9.9 |
| sp\|P23443\|KS6B1\_HUMAN | P23443 | KS6B1\_HUMAN | RPS6KB1 | Ribosomal protein S6 kinase beta-1 | 525 | 59.140 | 9.269081e+03 | 6.887080e+05 | 189528.03 | 472293.2 | 26886.787 |  | 8.95 | 31.4 | 12.0 |  |
| sp\|O00214\|LEG8\_HUMAN | O00214 | LEG8\_HUMAN | LGALS8 | Galectin-8 | 317 | 35.808 | 7.109077e+04 | 3.198235e+06 | 189486.6 |  | 2797135.2 | 211613.47 | 22.08 |  | 36.3 | 15.1 |
| sp\|P53621\|COPA\_HUMAN | P53621 | COPA\_HUMAN | COPA | Coatomer subunit alpha | 1224 | 138.346 | 4.942252e+03 | 8.590306e+05 | 189420.42 | 619958.8 | 49651.355 |  | 16.99 | 41.9 | 12.2 |  |
| sp\|P59998\|ARPC4\_HUMAN | P59998 | ARPC4\_HUMAN | ARPC4 | Actin-related protein 2/3 complex subunit 4 | 168 | 19.667 | 5.437708e+04 | 1.343604e+06 | 189350.08 | 783247.9 | 371005.62 |  | 42.86 | 60.7 | 29.2 |  |
| sp\|Q6N063\|OGFD2\_HUMAN | Q6N063 | OGFD2\_HUMAN | OGFOD2 | 2-oxoglutarate and iron-dependent oxygenase domain-containing protein 2 | 350 | 38.996 | 1.101142e+04 | 5.394865e+05 | 189047.84 | 350438.62 |  |  | 7.14 | 29.4 |  |  |
| sp\|Q9NUP9\|LIN7C\_HUMAN | Q9NUP9 | LIN7C\_HUMAN | LIN7C | Protein lin-7 homolog C | 197 | 21.834 | 8.150870e+04 | 2.235910e+06 | 188356.39 | 661363.9 | 1386190.2 |  | 25.38 | 52.3 | 43.7 |  |
| sp\|Q6DHV7\|ADAL\_HUMAN | Q6DHV7 | ADAL\_HUMAN | ADAL | Adenosine deaminase-like protein | 355 | 40.264 | 1.240489e+04 | 6.275191e+05 | 188085.84 | 439433.22 |  |  | 26.76 | 39.7 |  |  |
| sp\|P52735\|VAV2\_HUMAN | P52735 | VAV2\_HUMAN | VAV2 | Guanine nucleotide exchange factor VAV2 | 878 | 101.289 | 3.062918e+03 | 3.897758e+05 | 187619.14 | 202156.61 |  |  | 13.1 | 42.1 |  |  |
| sp\|Q14997\|PSME4\_HUMAN | Q14997 | PSME4\_HUMAN | PSME4 | Proteasome activator complex subunit 4 | 1843 | 211.334 | 2.006870e+04 | 5.328511e+06 | 186718.19 | 5141792.5 |  |  | 8.41 | 30.5 |  |  |
| sp\|P49427\|UB2R1\_HUMAN | P49427 | UB2R1\_HUMAN | CDC34 | Ubiquitin-conjugating enzyme E2 R1 | 236 | 26.737 | 2.083191e+04 | 6.997759e+05 | 186507.22 | 362913.34 | 150355.33 |  | 19.07 | 32.2 | 25.4 |  |
| sp\|Q9H2U1\|DHX36\_HUMAN | Q9H2U1 | DHX36\_HUMAN | DHX36 | ATP-dependent DNA/RNA helicase DHX36 | 1008 | 114.760 | 1.420690e+03 | 2.048364e+05 | 186387.34 | 18449.012 |  |  | 4.46 | 26.0 |  |  |
| sp\|P10809\|CH60\_HUMAN | P10809 | CH60\_HUMAN | HSPD1 | 60 kDa heat shock protein, mitochondrial | 573 | 61.055 | 1.504606e+04 | 1.154147e+06 | 186144.48 | 138744.78 | 384158.34 | 445099.88 | 15.53 | 42.6 | 27.4 | 24.3 |
| sp\|Q53FT3\|HIKES\_HUMAN | Q53FT3 | HIKES\_HUMAN | HIKESHI | Protein Hikeshi | 197 | 21.628 | 2.040968e+04 | 5.545871e+05 | 186120.75 | 225216.25 | 92416.336 | 50833.73 | 19.29 | 19.3 | 10.7 | 6.6 |
| sp\|Q14914\|PTGR1\_HUMAN | Q14914 | PTGR1\_HUMAN | PTGR1 | Prostaglandin reductase 1 | 329 | 35.870 | 2.034745e+04 | 9.169772e+05 | 186112.38 | 662955.94 | 67908.875 |  | 36.47 | 26.7 | 13.7 |  |
| sp\|Q8WYP5\|ELYS\_HUMAN | Q8WYP5 | ELYS\_HUMAN | AHCTF1 | Protein ELYS | 2266 | 249.260 | 5.931579e+02 | 1.857548e+05 | 185754.81 |  |  |  | 1.32 |  |  |  |
| sp\|Q9UI42\|CBPA4\_HUMAN | Q9UI42 | CBPA4\_HUMAN | CPA4 | Carboxypeptidase A4 | 421 | 47.351 | 1.207353e+04 | 7.182579e+05 | 185336.22 | 73581.78 | 241261.1 | 218078.8 | 15.68 | 26.4 | 16.4 | 6.9 |
| sp\|Q14894\|CRYM\_HUMAN | Q14894 | CRYM\_HUMAN | CRYM | Ketimine reductase mu-crystallin | 314 | 33.776 | 1.307823e+04 | 5.549762e+05 | 185297.4 | 369678.78 |  |  | 25.16 | 20.4 |  |  |
| sp\|Q8NBI5\|S43A3\_HUMAN | Q8NBI5 | S43A3\_HUMAN | SLC43A3 | Equilibrative nucleobase transporter 1 | 491 | 54.529 | 3.226967e+04 | 2.210748e+06 | 185149.11 |  | 721856.8 | 1303741.6 | 8.55 |  | 12.6 | 14.1 |
| sp\|Q8NEZ2\|VP37A\_HUMAN | Q8NEZ2 | VP37A\_HUMAN | VPS37A | Vacuolar protein sorting-associated protein 37A | 397 | 44.314 | 2.039499e+04 | 1.135485e+06 | 184638.12 | 476829.06 | 474018.22 |  | 17.63 | 21.9 | 22.4 |  |
| sp\|Q7Z4H8\|PLGT3\_HUMAN | Q7Z4H8 | PLGT3\_HUMAN | POGLUT3 | Protein O-glucosyltransferase 3 | 507 | 58.572 | 1.840443e+04 | 1.354346e+06 | 183632.47 | 236568.88 | 638897.0 | 295247.7 | 5.13 | 31.8 | 8.7 | 9.5 |
| sp\|P02788\|TRFL\_HUMAN | P02788 | TRFL\_HUMAN | LTF | Lactotransferrin | 710 | 78.182 | 1.861603e+04 | 1.828568e+06 | 183344.16 | 809613.25 | 416033.22 | 419576.88 | 15.07 | 45.6 | 38.9 | 28.3 |
| sp\|Q96A65\|EXOC4\_HUMAN | Q96A65 | EXOC4\_HUMAN | EXOC4 | Exocyst complex component 4 | 974 | 110.498 | 9.252415e+03 | 1.284478e+06 | 182973.4 | 1060775.9 | 27296.867 | 13431.892 | 9.65 | 39.0 | 18.5 | 8.1 |
| sp\|Q8IZY2\|ABCA7\_HUMAN | Q8IZY2 | ABCA7\_HUMAN | ABCA7 | Phospholipid-transporting ATPase ABCA7 | 2146 | 234.350 | 6.378607e+03 | 1.878053e+06 | 182768.47 | 146025.12 | 459510.5 | 1089749.4 | 13.7 | 34.9 | 22.2 | 19.9 |
| sp\|Q8N9V3\|WSDU1\_HUMAN | Q8N9V3 | WSDU1\_HUMAN | WDSUB1 | WD repeat, SAM and U-box domain-containing protein 1 | 476 | 52.833 | 4.804306e+03 | 3.188990e+05 | 182714.53 | 136184.44 |  |  | 8.82 | 27.9 |  |  |
| sp\|Q9Y6P5\|SESN1\_HUMAN | Q9Y6P5 | SESN1\_HUMAN | SESN1 | Sestrin-1 | 492 | 56.557 | 5.653932e+03 | 4.017483e+05 | 182370.92 | 130021.01 | 89356.39 |  | 20.33 | 22.8 | 25.6 |  |
| sp\|Q9NXA8\|SIR5\_HUMAN | Q9NXA8 | SIR5\_HUMAN | SIRT5 | NAD-dependent protein deacylase sirtuin-5, mitochondrial | 310 | 33.881 | 1.748762e+04 | 7.443960e+05 | 182270.66 | 483842.62 | 78282.695 |  | 15.16 | 61.6 | 24.8 |  |
| sp\|O00203\|AP3B1\_HUMAN | O00203 | AP3B1\_HUMAN | AP3B1 | AP-3 complex subunit beta-1 | 1094 | 121.320 | 3.039842e+03 | 4.633408e+05 | 182150.06 | 281190.7 |  |  | 18.37 | 24.8 |  |  |
| sp\|Q96EK7\|F120B\_HUMAN | Q96EK7 | F120B\_HUMAN | FAM120B | Constitutive coactivator of peroxisome proliferator-activated receptor gamma | 910 | 103.783 | 3.762451e+03 | 4.905850e+05 | 181964.47 | 308620.5 |  |  | 12.2 | 43.7 |  |  |
| sp\|O94905\|ERLN2\_HUMAN | O94905 | ERLN2\_HUMAN | ERLIN2 | Erlin-2 | 339 | 37.840 | 6.779625e+04 | 3.223101e+06 | 181774.81 | 88394.445 | 2440250.5 | 512681.5 | 23.3 | 37.2 | 45.1 | 37.2 |
| sp\|O60331\|PI51C\_HUMAN | O60331 | PI51C\_HUMAN | PIP5K1C | Phosphatidylinositol 4-phosphate 5-kinase type-1 gamma | 668 | 73.260 | 4.519042e+03 | 4.159398e+05 | 181462.33 | 234477.47 |  |  | 5.24 | 18.3 |  |  |
| sp\|Q15437\|SC23B\_HUMAN | Q15437 | SC23B\_HUMAN | SEC23B | Protein transport protein Sec23B | 767 | 86.479 | 3.803759e+03 | 4.132766e+05 | 181441.56 | 145650.31 | 86184.766 |  | 13.04 | 37.8 | 12.3 |  |
| sp\|P19623\|SPEE\_HUMAN | P19623 | SPEE\_HUMAN | SRM | Spermidine synthase | 302 | 33.825 | 4.919509e+03 | 2.090628e+05 | 180723.69 |  | 28339.105 |  | 14.57 |  | 19.2 |  |
| sp\|Q9UHV9\|PFD2\_HUMAN | Q9UHV9 | PFD2\_HUMAN | PFDN2 | Prefoldin subunit 2 | 154 | 16.648 | 8.403064e+04 | 1.757587e+06 | 180663.19 | 1485697.1 | 91226.875 |  | 14.94 | 70.1 | 26.0 |  |
| sp\|Q9NZT2\|OGFR\_HUMAN | Q9NZT2 | OGFR\_HUMAN | OGFR | Opioid growth factor receptor | 677 | 73.325 | 7.203606e+03 | 6.636196e+05 | 180453.34 | 473780.9 | 9385.312 |  | 36.04 | 46.1 | 21.0 |  |
| sp\|Q6ZVX7\|FBX50\_HUMAN | Q6ZVX7 | FBX50\_HUMAN | NCCRP1 | F-box only protein 50 | 275 | 30.847 | 3.229982e+04 | 1.251786e+06 | 180272.67 | 201895.55 | 553801.75 | 315816.25 | 26.91 | 35.3 | 16.4 | 17.1 |
| sp\|Q5VW36\|FOCAD\_HUMAN | Q5VW36 | FOCAD\_HUMAN | FOCAD | Focadhesin | 1801 | 200.058 | 3.845694e+03 | 9.666023e+05 | 180214.52 | 661881.0 |  | 124506.8 | 9.16 | 29.0 |  | 6.1 |
| sp\|Q8TAF3\|WDR48\_HUMAN | Q8TAF3 | WDR48\_HUMAN | WDR48 | WD repeat-containing protein 48 | 677 | 76.210 | 8.253267e+03 | 7.902327e+05 | 179924.28 | 425391.9 | 184916.52 |  | 20.53 | 35.7 | 15.7 |  |
| sp\|Q9Y285\|SYFA\_HUMAN | Q9Y285 | SYFA\_HUMAN | FARSA | Phenylalanine--tRNA ligase alpha subunit | 508 | 57.564 | 1.026132e+04 | 7.421149e+05 | 179655.58 | 464420.72 | 98038.625 |  | 18.11 | 36.8 | 32.5 |  |
| sp\|Q9BR76\|COR1B\_HUMAN | Q9BR76 | COR1B\_HUMAN | CORO1B | Coronin-1B | 489 | 54.235 | 5.794357e+03 | 3.948227e+05 | 179624.95 | 174436.97 | 40760.73 |  | 13.09 | 20.2 | 18.6 |  |
| sp\|Q13363\|CTBP1\_HUMAN | Q13363 | CTBP1\_HUMAN | CTBP1 | C-terminal-binding protein 1 | 440 | 47.535 | 5.655725e+03 | 3.377683e+05 | 179532.84 | 97287.28 | 60948.22 |  | 17.5 | 36.8 | 16.6 |  |
| sp\|P26640\|SYVC\_HUMAN | P26640 | SYVC\_HUMAN | VARS1 | Valine--tRNA ligase | 1264 | 140.476 | 4.501885e+03 | 7.945362e+05 | 179205.03 | 593482.56 | 21848.64 |  | 17.48 | 39.4 | 21.0 |  |
| sp\|O00151\|PDLI1\_HUMAN | O00151 | PDLI1\_HUMAN | PDLIM1 | PDZ and LIM domain protein 1 | 329 | 36.072 | 1.237266e+05 | 5.607255e+06 | 178878.86 | 253688.44 | 764845.4 | 4409842.0 | 29.79 | 53.5 | 21.9 | 16.4 |
| sp\|Q7Z392\|TPC11\_HUMAN | Q7Z392 | TPC11\_HUMAN | TRAPPC11 | Trafficking protein particle complex subunit 11 | 1133 | 128.881 | 2.506638e+03 | 4.058800e+05 | 178569.16 | 227310.89 |  |  | 9.09 | 26.8 |  |  |
| sp\|O15397\|IPO8\_HUMAN | O15397 | IPO8\_HUMAN | IPO8 | Importin-8 | 1037 | 114.070 | 1.243332e+03 | 1.781869e+05 | 178186.88 |  |  |  | 2.12 |  |  |  |
| sp\|Q5JRK9\|GGEE3\_HUMAN | Q5JRK9 | GGEE3\_HUMAN | PAGE2B | Putative G antigen family E member 3 | 111 | 12.041 | 2.533921e+04 | 3.833300e+05 | 178053.14 |  | 155204.67 | 50072.2 | 30.63 |  | 48.6 | 20.7 |
| sp\|P42356\|PI4KA\_HUMAN | P42356 | PI4KA\_HUMAN | PI4KA | Phosphatidylinositol 4-kinase alpha | 2102 | 236.830 | 1.479874e+03 | 4.403304e+05 | 178047.95 | 98639.25 | 19479.926 | 144163.28 | 4.38 | 33.6 | 21.4 | 16.0 |
| sp\|Q9H832\|UBE2Z\_HUMAN | Q9H832 | UBE2Z\_HUMAN | UBE2Z | Ubiquitin-conjugating enzyme E2 Z | 354 | 38.210 | 9.745820e+03 | 4.678564e+05 | 177558.95 | 290297.44 |  |  | 13.28 | 29.4 |  |  |
| sp\|Q9UBF2\|COPG2\_HUMAN | Q9UBF2 | COPG2\_HUMAN | COPG2 | Coatomer subunit gamma-2 | 871 | 97.622 | 3.619623e+03 | 4.439441e+05 | 177100.86 | 220292.64 | 46550.56 |  | 10.68 | 47.2 | 9.1 |  |
| sp\|Q8NAT1\|PMGT2\_HUMAN | Q8NAT1 | PMGT2\_HUMAN | POMGNT2 | Protein O-linked-mannose beta-1,4-N-acetylglucosaminyltransferase 2 | 580 | 66.615 | 5.871410e+03 | 4.913960e+05 | 176808.23 |  | 175269.84 | 139317.94 | 8.28 |  | 26.4 | 3.4 |
| sp\|P53582\|MAP11\_HUMAN | P53582 | MAP11\_HUMAN | METAP1 | Methionine aminopeptidase 1 | 386 | 43.215 | 5.532004e+03 | 3.003545e+05 | 176348.47 | 87230.914 | 20651.352 | 16123.769 | 11.4 | 40.9 | 6.7 | 11.7 |
| sp\|Q15650\|TRIP4\_HUMAN | Q15650 | TRIP4\_HUMAN | TRIP4 | Activating signal cointegrator 1 | 581 | 66.146 | 4.404567e+03 | 3.660361e+05 | 175263.89 | 190772.22 |  |  | 17.73 | 39.4 |  |  |
| sp\|Q86VP3\|PACS2\_HUMAN | Q86VP3 | PACS2\_HUMAN | PACS2 | Phosphofurin acidic cluster sorting protein 2 | 889 | 97.702 | 2.100897e+03 | 2.578845e+05 | 175008.44 | 38730.156 | 44145.938 |  | 18.9 | 33.4 | 19.5 |  |
| sp\|Q9BTM9\|URM1\_HUMAN | Q9BTM9 | URM1\_HUMAN | URM1 | Ubiquitin-related modifier 1 | 101 | 11.380 | 1.874057e+04 | 2.679428e+05 | 174761.23 | 93181.6 | 0.0 |  | 12.87 | 19.8 | 12.9 |  |
| sp\|Q9UQE7\|SMC3\_HUMAN | Q9UQE7 | SMC3\_HUMAN | SMC3 | Structural maintenance of chromosomes protein 3 | 1217 | 141.542 | 2.597501e+03 | 4.619109e+05 | 173982.53 | 254400.52 | 15668.136 | 17859.738 | 20.38 | 30.7 | 17.1 | 15.1 |
| sp\|P0DOY2\|IGLC2\_HUMAN | P0DOY2 | IGLC2\_HUMAN | IGLC2 | Immunoglobulin lambda constant 2 | 106 | 11.294 | 1.277924e+05 | 1.813301e+06 | 173654.48 | 310122.06 | 811322.4 | 518202.28 | 41.51 | 85.8 | 23.6 | 9.4 |
| sp\|P62330\|ARF6\_HUMAN | P62330 | ARF6\_HUMAN | ARF6 | ADP-ribosylation factor 6 | 175 | 20.082 | 1.718135e+05 | 4.334923e+06 | 173580.08 | 4087202.8 | 39866.883 | 34272.844 | 26.86 | 71.4 | 12.6 | 21.1 |
| sp\|O95429\|BAG4\_HUMAN | O95429 | BAG4\_HUMAN | BAG4 | BAG family molecular chaperone regulator 4 | 457 | 49.594 | 3.439021e+03 | 2.142798e+05 | 173447.89 | 40831.87 |  |  | 3.28 | 30.2 |  |  |
| sp\|Q9C0B5\|ZDHC5\_HUMAN | Q9C0B5 | ZDHC5\_HUMAN | ZDHHC5 | Palmitoyltransferase ZDHHC5 | 715 | 77.545 | 2.284921e+04 | 2.226087e+06 | 173387.02 | 59123.93 | 316476.1 | 1677100.1 | 11.47 | 42.7 | 31.3 | 28.1 |
| sp\|Q8N4S9\|MALD2\_HUMAN | Q8N4S9 | MALD2\_HUMAN | MARVELD2 | MARVEL domain-containing protein 2 | 558 | 64.168 | 5.718471e+04 | 4.610156e+06 | 172764.28 | 12754.344 | 2854348.0 | 1570289.1 | 12.9 | 23.8 | 33.3 | 36.6 |
| sp\|P50542\|PEX5\_HUMAN | P50542 | PEX5\_HUMAN | PEX5 | Peroxisomal targeting signal 1 receptor | 639 | 70.865 | 3.275857e+03 | 2.916580e+05 | 172171.44 | 119486.53 |  |  | 21.13 | 25.2 |  |  |
| sp\|Q8TF65\|GIPC2\_HUMAN | Q8TF65 | GIPC2\_HUMAN | GIPC2 | PDZ domain-containing protein GIPC2 | 315 | 34.354 | 1.716542e+04 | 7.408815e+05 | 172145.64 | 568735.9 |  |  | 26.67 | 35.6 |  |  |
| sp\|Q9UIW2\|PLXA1\_HUMAN | Q9UIW2 | PLXA1\_HUMAN | PLXNA1 | Plexin-A1 | 1896 | 211.067 | 4.766297e+03 | 1.263917e+06 | 171735.95 | 61123.605 | 340998.7 | 690058.75 | 6.38 | 38.9 | 22.5 | 16.0 |
| sp\|P41743\|KPCI\_HUMAN | P41743 | KPCI\_HUMAN | PRKCI | Protein kinase C iota type | 596 | 68.262 | 4.030977e+03 | 3.457056e+05 | 171315.11 | 174390.53 |  |  | 10.91 | 32.4 |  |  |
| sp\|Q6ZWT7\|MBOA2\_HUMAN | Q6ZWT7 | MBOA2\_HUMAN | MBOAT2 | Lysophospholipid acyltransferase 2 | 520 | 59.527 | 3.850274e+04 | 2.879538e+06 | 171201.1 | 804101.06 | 400823.9 | 1503411.6 | 13.85 | 27.3 | 16.9 | 10.6 |
| sp\|Q9GZS3\|SKI8\_HUMAN | Q9GZS3 | SKI8\_HUMAN | SKIC8 | Superkiller complex protein 8 | 305 | 33.581 | 1.387677e+04 | 5.854628e+05 | 170546.25 | 338526.6 | 76389.97 |  | 33.11 | 56.4 | 16.7 |  |
| sp\|Q8TDY2\|RBCC1\_HUMAN | Q8TDY2 | RBCC1\_HUMAN | RB1CC1 | RB1-inducible coiled-coil protein 1 | 1594 | 183.091 | 1.208067e+03 | 2.778915e+05 | 170124.88 | 107766.59 |  |  | 5.83 | 30.9 |  |  |
| sp\|Q9Y3E0\|GOT1B\_HUMAN | Q9Y3E0 | GOT1B\_HUMAN | GOLT1B | Vesicle transport protein GOT1B | 138 | 15.426 | 1.035354e+04 | 2.006594e+05 | 169847.81 |  | 30811.545 |  | 10.87 |  | 16.7 |  |
| sp\|O00429\|DNM1L\_HUMAN | O00429 | DNM1L\_HUMAN | DNM1L | Dynamin-1-like protein | 736 | 81.877 | 6.988566e+03 | 7.188978e+05 | 169511.69 | 435597.47 | 113788.664 |  | 30.57 | 43.6 | 21.7 |  |
| sp\|Q8IUW5\|RELL1\_HUMAN | Q8IUW5 | RELL1\_HUMAN | RELL1 | RELT-like protein 1 | 271 | 29.340 | 9.876803e+04 | 3.640774e+06 | 169000.88 | 385450.22 | 613748.25 | 2472574.2 | 32.84 | 60.5 | 53.1 | 27.3 |
| sp\|P17252\|KPCA\_HUMAN | P17252 | KPCA\_HUMAN | PRKCA | Protein kinase C alpha type | 672 | 76.750 | 3.071728e+03 | 2.961954e+05 | 168656.5 | 127538.86 |  |  | 11.16 | 34.2 |  |  |
| sp\|Q6P3W7\|SCYL2\_HUMAN | Q6P3W7 | SCYL2\_HUMAN | SCYL2 | SCY1-like protein 2 | 929 | 103.709 | 7.110025e+03 | 9.264132e+05 | 168559.61 | 514238.56 | 215593.12 | 28021.865 | 10.12 | 36.5 | 21.4 | 4.6 |
| sp\|Q9UJ68\|MSRA\_HUMAN | Q9UJ68 | MSRA\_HUMAN | MSRA | Mitochondrial peptide methionine sulfoxide reductase | 235 | 26.132 | 1.587187e+04 | 5.210960e+05 | 168320.25 | 352775.8 |  |  | 31.06 | 56.2 |  |  |
| sp\|O75342\|LX12B\_HUMAN | O75342 | LX12B\_HUMAN | ALOX12B | Arachidonate 12-lipoxygenase, 12R-type | 701 | 80.356 | 1.056363e+04 | 1.066470e+06 | 168319.48 | 85491.086 | 354191.53 | 458468.4 | 6.42 | 36.2 | 23.8 | 18.8 |
| sp\|Q14683\|SMC1A\_HUMAN | Q14683 | SMC1A\_HUMAN | SMC1A | Structural maintenance of chromosomes protein 1A | 1233 | 143.233 | 1.389499e+04 | 2.500452e+06 | 168177.95 | 2332274.2 |  |  | 16.79 | 40.6 |  |  |
| sp\|Q7Z3U7\|MON2\_HUMAN | Q7Z3U7 | MON2\_HUMAN | MON2 | Protein MON2 homolog | 1717 | 190.359 | 3.671917e+03 | 8.781798e+05 | 168133.02 | 599834.56 |  | 110212.195 | 11.01 | 21.8 |  | 10.9 |
| sp\|P41252\|SYIC\_HUMAN | P41252 | SYIC\_HUMAN | IARS1 | Isoleucine--tRNA ligase, cytoplasmic | 1262 | 144.498 | 1.085426e+03 | 1.970513e+05 | 167800.28 | 29251.055 |  |  | 8.72 | 20.7 |  |  |
| sp\|Q9H6U6\|BCAS3\_HUMAN | Q9H6U6 | BCAS3\_HUMAN | BCAS3 | BCAS3 microtubule associated cell migration factor | 928 | 101.237 | 6.486607e+03 | 8.250381e+05 | 167790.5 | 559539.25 | 97708.375 |  | 3.99 | 41.3 | 19.7 |  |
| sp\|O75064\|DEN4B\_HUMAN | O75064 | DEN4B\_HUMAN | DENND4B | DENN domain-containing protein 4B | 1496 | 163.847 | 1.533889e+03 | 3.157545e+05 | 167497.03 | 148257.42 |  |  | 10.23 | 25.3 |  |  |
| sp\|O43826\|G6PT1\_HUMAN | O43826 | G6PT1\_HUMAN | SLC37A4 | Glucose-6-phosphate exchanger SLC37A4 | 429 | 46.360 | 4.603884e+04 | 2.681544e+06 | 167155.1 | 607804.8 | 967926.94 | 938657.2 | 3.03 | 26.8 | 9.8 | 12.4 |
| sp\|O00421\|CCRL2\_HUMAN | O00421 | CCRL2\_HUMAN | CCRL2 | C-C chemokine receptor-like 2 | 344 | 39.513 | 1.168223e+05 | 5.799399e+06 | 167057.81 |  | 1125622.4 | 4506719.0 | 5.52 |  | 11.6 | 13.7 |
| sp\|Q9Y303\|NAGA\_HUMAN | Q9Y303 | NAGA\_HUMAN | AMDHD2 | N-acetylglucosamine-6-phosphate deacetylase | 409 | 43.748 | 5.079217e+03 | 2.791722e+05 | 166689.6 | 112482.59 |  |  | 15.16 | 35.5 |  |  |
| sp\|Q9NVX2\|NLE1\_HUMAN | Q9NVX2 | NLE1\_HUMAN | NLE1 | Notchless protein homolog 1 | 485 | 53.320 | 4.602072e+03 | 3.082909e+05 | 166355.12 | 141935.75 |  |  | 17.11 | 42.3 |  |  |
| sp\|P20936\|RASA1\_HUMAN | P20936 | RASA1\_HUMAN | RASA1 | Ras GTPase-activating protein 1 | 1047 | 116.403 | 2.101627e+03 | 3.073527e+05 | 166351.6 | 141001.05 |  |  | 9.55 | 23.7 |  |  |
| sp\|Q96P48\|ARAP1\_HUMAN | Q96P48 | ARAP1\_HUMAN | ARAP1 | Arf-GAP with Rho-GAP domain, ANK repeat and PH domain-containing protein 1 | 1450 | 162.192 | 2.740316e+03 | 5.584023e+05 | 166282.77 | 392119.5 |  |  | 10.9 | 38.1 |  |  |
| sp\|Q9ULC3\|RAB23\_HUMAN | Q9ULC3 | RAB23\_HUMAN | RAB23 | Ras-related protein Rab-23 | 237 | 26.659 | 2.508222e+04 | 8.400922e+05 | 165795.16 | 599310.25 | 74986.79 |  | 28.69 | 39.7 | 33.8 |  |
| sp\|Q9Y2X7\|GIT1\_HUMAN | Q9Y2X7 | GIT1\_HUMAN | GIT1 | ARF GTPase-activating protein GIT1 | 761 | 84.341 | 8.449682e+03 | 8.953571e+05 | 165616.05 | 507259.5 | 222481.55 |  | 11.96 | 51.4 | 18.4 |  |
| sp\|Q96D71\|REPS1\_HUMAN | Q96D71 | REPS1\_HUMAN | REPS1 | RalBP1-associated Eps domain-containing protein 1 | 796 | 86.662 | 5.985215e+03 | 6.516668e+05 | 165501.34 | 218303.81 | 232189.69 | 35671.984 | 10.3 | 29.8 | 22.2 | 7.0 |
| sp\|Q9BQG2\|NUD12\_HUMAN | Q9BQG2 | NUD12\_HUMAN | NUDT12 | NAD-capped RNA hydrolase NUDT12 | 462 | 52.076 | 7.482007e+03 | 4.895227e+05 | 164800.4 | 324722.34 |  |  | 16.45 | 50.9 |  |  |
| sp\|G2XKQ0\|SUMO5\_HUMAN | G2XKQ0 | SUMO5\_HUMAN | SUMO1P1 | Small ubiquitin-related modifier 5 | 101 | 11.110 | 1.179975e+04 | 1.647039e+05 | 164703.9 |  |  |  | 18.81 |  |  |  |
| sp\|O00399\|DCTN6\_HUMAN | O00399 | DCTN6\_HUMAN | DCTN6 | Dynactin subunit 6 | 190 | 20.747 | 4.864583e+04 | 1.267997e+06 | 164636.25 | 991995.06 | 111365.31 |  | 26.84 | 32.6 | 11.1 |  |
| sp\|Q9NPC3\|CIP1\_HUMAN | Q9NPC3 | CIP1\_HUMAN | CCNB1IP1 | E3 ubiquitin-protein ligase CCNB1IP1 | 277 | 30.470 | 4.290709e+03 | 1.642550e+05 | 164255.03 |  |  |  | 5.42 |  |  |  |
| sp\|O15523\|DDX3Y\_HUMAN | O15523 | DDX3Y\_HUMAN | DDX3Y | ATP-dependent RNA helicase DDX3Y | 660 | 73.154 | 5.361981e+03 | 4.928112e+05 | 163295.88 | 329515.28 |  |  | 13.48 | 38.5 |  |  |
| sp\|P49768\|PSN1\_HUMAN | P49768 | PSN1\_HUMAN | PSEN1 | Presenilin-1 | 467 | 52.668 | 4.974529e+04 | 3.291668e+06 | 162827.2 |  | 1397530.6 | 1731309.8 | 9.42 |  | 13.7 | 23.8 |
| sp\|Q8N2U9\|S66A2\_HUMAN | Q8N2U9 | S66A2\_HUMAN | SLC66A2 | Solute carrier family 66 member 2 | 271 | 30.478 | 2.482897e+04 | 9.507411e+05 | 161470.08 | 91063.53 | 167380.83 | 530826.7 | 17.34 | 37.3 | 14.8 | 18.5 |
| sp\|Q8NC44\|RETR2\_HUMAN | Q8NC44 | RETR2\_HUMAN | RETREG2 | Reticulophagy regulator 2 | 543 | 57.830 | 4.476449e+03 | 3.252400e+05 | 161381.8 | 59246.78 | 30566.08 | 74045.33 | 13.81 | 10.3 | 6.8 | 4.4 |
| sp\|Q7Z5R6\|AB1IP\_HUMAN | Q7Z5R6 | AB1IP\_HUMAN | APBB1IP | Amyloid beta A4 precursor protein-binding family B member 1-interacting protein | 666 | 73.183 | 3.263591e+03 | 3.000704e+05 | 161211.45 | 119423.0 |  | 19435.943 | 11.71 | 26.9 |  | 4.8 |
| sp\|Q13190\|STX5\_HUMAN | Q13190 | STX5\_HUMAN | STX5 | Syntaxin-5 | 355 | 39.673 | 2.675705e+04 | 1.333676e+06 | 160730.97 | 415549.7 | 194850.52 | 562545.25 | 26.76 | 44.8 | 31.8 | 25.1 |
| sp\|P08069\|IGF1R\_HUMAN | P08069 | IGF1R\_HUMAN | IGF1R | Insulin-like growth factor 1 receptor | 1367 | 154.793 | 8.448307e+03 | 1.643002e+06 | 160635.6 | 1094441.8 | 180297.7 | 207627.17 | 4.32 | 32.3 | 15.9 | 15.4 |
| sp\|P05141\|ADT2\_HUMAN | P05141 | ADT2\_HUMAN | SLC25A5 | ADP/ATP translocase 2 | 298 | 32.852 | 2.262054e+04 | 9.336454e+05 | 159746.06 | 578171.75 | 195727.6 |  | 6.71 | 55.4 | 29.2 |  |
| sp\|P38159\|RBMX\_HUMAN | P38159 | RBMX\_HUMAN | RBMX | RNA-binding motif protein, X chromosome | 391 | 42.332 | 2.861177e+04 | 1.521705e+06 | 159639.19 | 1173009.5 | 189056.75 |  | 13.81 | 55.2 | 42.5 |  |
| sp\|Q6ZS17\|RIPR1\_HUMAN | Q6ZS17 | RIPR1\_HUMAN | RIPOR1 | Rho family-interacting cell polarization regulator 1 | 1223 | 132.308 | 1.992413e+03 | 3.311942e+05 | 159520.19 | 98694.03 | 72979.97 |  | 6.05 | 30.7 | 20.4 |  |
| sp\|Q9C0C2\|TB182\_HUMAN | Q9C0C2 | TB182\_HUMAN | TNKS1BP1 | 182 kDa tankyrase-1-binding protein | 1729 | 181.796 | 1.242631e+03 | 2.838205e+05 | 158935.5 | 124885.04 |  |  | 3.41 | 37.0 |  |  |
| sp\|Q9C0K0\|BC11B\_HUMAN | Q9C0K0 | BC11B\_HUMAN | BCL11B | B-cell lymphoma/leukemia 11B | 894 | 98.340 | 1.281919e+03 | 1.583828e+05 | 158382.83 |  |  |  | 5.26 |  |  |  |
| sp\|Q7LBR1\|CHM1B\_HUMAN | Q7LBR1 | CHM1B\_HUMAN | CHMP1B | Charged multivesicular body protein 1b | 199 | 22.109 | 4.575064e+04 | 1.270818e+06 | 158201.77 | 604469.56 | 508146.88 |  | 10.05 | 47.2 | 30.7 |  |
| sp\|P35658\|NU214\_HUMAN | P35658 | NU214\_HUMAN | NUP214 | Nuclear pore complex protein Nup214 | 2090 | 213.620 | 2.558329e+03 | 6.866185e+05 | 158196.53 | 438868.06 | 78307.93 | 11245.987 | 2.73 | 27.0 | 13.8 | 5.5 |
| sp\|Q96NT3\|GUCD1\_HUMAN | Q96NT3 | GUCD1\_HUMAN | GUCD1 | Protein GUCD1 | 240 | 27.207 | 6.402002e+04 | 2.188334e+06 | 158130.44 | 104309.09 | 404981.3 | 1520913.2 | 10.0 | 22.5 | 16.7 | 16.2 |
| sp\|Q9Y2I1\|NISCH\_HUMAN | Q9Y2I1 | NISCH\_HUMAN | NISCH | Nischarin | 1504 | 166.629 | 5.503424e+03 | 1.152128e+06 | 158120.78 | 994007.1 |  |  | 5.05 | 24.0 |  |  |
| sp\|Q96G74\|OTUD5\_HUMAN | Q96G74 | OTUD5\_HUMAN | OTUD5 | OTU domain-containing protein 5 | 571 | 60.626 | 7.200138e+03 | 5.484246e+05 | 157921.25 | 262148.94 | 100783.75 | 27570.61 | 17.86 | 39.9 | 15.6 | 6.0 |
| sp\|Q92738\|US6NL\_HUMAN | Q92738 | US6NL\_HUMAN | USP6NL | USP6 N-terminal-like protein | 828 | 94.104 | 4.178696e+03 | 4.940445e+05 | 157760.97 | 336283.5 |  |  | 10.14 | 42.4 |  |  |
| sp\|Q86W56\|PARG\_HUMAN | Q86W56 | PARG\_HUMAN | PARG | Poly(ADP-ribose) glycohydrolase | 976 | 111.110 | 1.705235e+03 | 2.380425e+05 | 157684.6 | 80357.94 |  |  | 13.22 | 39.2 |  |  |
| sp\|Q6NUQ1\|RINT1\_HUMAN | Q6NUQ1 | RINT1\_HUMAN | RINT1 | RAD50-interacting protein 1 | 792 | 90.632 | 6.128537e+03 | 6.978395e+05 | 157485.06 | 134047.16 | 358579.97 | 47727.273 | 9.34 | 25.4 | 13.1 | 9.3 |
| sp\|P17813\|EGLN\_HUMAN | P17813 | EGLN\_HUMAN | ENG | Endoglin | 658 | 70.578 | 7.632983e+03 | 6.768319e+05 | 157374.97 | 89151.06 | 37150.62 | 393155.25 | 6.53 | 23.3 | 21.6 | 7.6 |
| sp\|Q9UPM8\|AP4E1\_HUMAN | Q9UPM8 | AP4E1\_HUMAN | AP4E1 | AP-4 complex subunit epsilon-1 | 1137 | 127.287 | 3.368003e+03 | 5.386091e+05 | 156827.66 | 198012.8 | 75040.88 | 108727.73 | 9.59 | 28.2 | 12.5 | 8.5 |
| sp\|Q9HDC9\|APMAP\_HUMAN | Q9HDC9 | APMAP\_HUMAN | APMAP | Adipocyte plasma membrane-associated protein | 416 | 46.480 | 7.947339e+04 | 4.640930e+06 | 156646.5 | 315764.16 | 3852442.5 | 316076.97 | 14.66 | 58.2 | 31.0 | 15.6 |
| sp\|Q9NZ08\|ERAP1\_HUMAN | Q9NZ08 | ERAP1\_HUMAN | ERAP1 | Endoplasmic reticulum aminopeptidase 1 | 941 | 107.235 | 2.549225e+04 | 3.434488e+06 | 156124.44 | 237620.12 | 2768420.0 | 272323.0 | 13.18 | 35.1 | 24.2 | 12.2 |
| sp\|Q15027\|ACAP1\_HUMAN | Q15027 | ACAP1\_HUMAN | ACAP1 | Arf-GAP with coiled-coil, ANK repeat and PH domain-containing protein 1 | 740 | 81.536 | 2.947688e+04 | 3.019590e+06 | 155996.36 | 2863593.8 |  |  | 9.59 | 42.0 |  |  |
| sp\|Q96AB6\|NTAN1\_HUMAN | Q96AB6 | NTAN1\_HUMAN | NTAN1 | Protein N-terminal asparagine amidohydrolase | 310 | 34.677 | 6.450976e+03 | 2.810503e+05 | 155724.64 | 125325.67 |  |  | 12.58 | 48.4 |  |  |
| sp\|Q96J01\|THOC3\_HUMAN | Q96J01 | THOC3\_HUMAN | THOC3 | THO complex subunit 3 | 351 | 38.610 | 3.205217e+03 | 1.554800e+05 | 155479.95 |  |  |  | 2.85 |  |  |  |
| sp\|P40306\|PSB10\_HUMAN | P40306 | PSB10\_HUMAN | PSMB10 | Proteasome subunit beta type-10 | 273 | 28.936 | 1.329726e+04 | 4.834124e+05 | 155469.5 | 327942.94 |  |  | 11.36 | 28.2 |  |  |
| sp\|Q8TEQ6\|GEMI5\_HUMAN | Q8TEQ6 | GEMI5\_HUMAN | GEMIN5 | Gem-associated protein 5 | 1508 | 168.589 | 2.445879e+03 | 5.180616e+05 | 155334.64 | 362727.0 |  |  | 9.42 | 32.3 |  |  |
| sp\|Q15334\|L2GL1\_HUMAN | Q15334 | L2GL1\_HUMAN | LLGL1 | Lethal(2) giant larvae protein homolog 1 | 1064 | 115.418 | 6.127466e+03 | 8.885290e+05 | 155019.6 | 733509.4 |  |  | 16.54 | 35.7 |  |  |
| sp\|Q0VDF9\|HSP7E\_HUMAN | Q0VDF9 | HSP7E\_HUMAN | HSPA14 | Heat shock 70 kDa protein 14 | 509 | 54.794 | 6.855503e+03 | 4.719430e+05 | 154901.9 | 317041.06 |  |  | 21.22 | 48.3 |  |  |
| sp\|P62993\|GRB2\_HUMAN | P62993 | GRB2\_HUMAN | GRB2 | Growth factor receptor-bound protein 2 | 217 | 25.206 | 1.583913e+04 | 5.015942e+05 | 154873.64 | 202169.67 | 144550.84 |  | 30.88 | 44.7 | 23.5 |  |
| sp\|Q96FS4\|SIPA1\_HUMAN | Q96FS4 | SIPA1\_HUMAN | SIPA1 | Signal-induced proliferation-associated protein 1 | 1042 | 112.149 | 2.578410e+03 | 3.632993e+05 | 153261.5 | 188213.9 | 21823.912 |  | 14.2 | 39.0 | 13.2 |  |
| sp\|Q86VX9\|MON1A\_HUMAN | Q86VX9 | MON1A\_HUMAN | MON1A | Vacuolar fusion protein MON1 homolog A | 652 | 72.895 | 2.782645e+03 | 2.548430e+05 | 152673.38 | 102169.62 |  |  | 17.94 | 31.4 |  |  |
| sp\|Q99873\|ANM1\_HUMAN | Q99873 | ANM1\_HUMAN | PRMT1 | Protein arginine N-methyltransferase 1 | 371 | 42.462 | 1.030902e+04 | 5.499648e+05 | 152471.3 | 371404.56 | 26088.975 |  | 13.21 | 51.2 | 19.7 |  |
| sp\|Q9Y3P9\|RBGP1\_HUMAN | Q9Y3P9 | RBGP1\_HUMAN | RABGAP1 | Rab GTPase-activating protein 1 | 1069 | 121.737 | 2.550018e+03 | 3.900166e+05 | 152262.14 | 237754.44 |  |  | 3.46 | 39.3 |  |  |
| sp\|Q9C0E8\|LNP\_HUMAN | Q9C0E8 | LNP\_HUMAN | LNPK | Endoplasmic reticulum junction formation protein lunapark | 428 | 47.740 | 1.493168e+04 | 8.955877e+05 | 151816.98 | 232360.05 |  | 511410.7 | 9.58 | 33.4 |  | 21.5 |
| sp\|P42224\|STAT1\_HUMAN | P42224 | STAT1\_HUMAN | STAT1 | Signal transducer and activator of transcription 1-alpha/beta | 750 | 87.335 | 4.187986e+03 | 4.595266e+05 | 151565.44 | 307961.2 |  |  | 24.0 | 46.5 |  |  |
| sp\|A6NHX0\|CAST2\_HUMAN | A6NHX0 | CAST2\_HUMAN | CASTOR2 | Cytosolic arginine sensor for mTORC1 subunit 2 | 329 | 36.056 | 7.319832e+03 | 3.315857e+05 | 151198.89 | 180386.83 |  |  | 24.32 | 22.5 |  |  |
| sp\|Q9NZN5\|ARHGC\_HUMAN | Q9NZN5 | ARHGC\_HUMAN | ARHGEF12 | Rho guanine nucleotide exchange factor 12 | 1544 | 173.232 | 2.178304e+03 | 4.740932e+05 | 150954.44 | 260948.16 | 62190.64 |  | 10.82 | 29.5 | 16.4 |  |
| sp\|P51617\|IRAK1\_HUMAN | P51617 | IRAK1\_HUMAN | IRAK1 | Interleukin-1 receptor-associated kinase 1 | 712 | 76.537 | 4.646130e+03 | 4.467659e+05 | 149767.42 | 270650.56 | 26347.889 |  | 12.36 | 28.2 | 11.1 |  |
| sp\|Q9HB90\|RRAGC\_HUMAN | Q9HB90 | RRAGC\_HUMAN | RRAGC | Ras-related GTP-binding protein C | 399 | 44.224 | 9.344521e+03 | 5.191971e+05 | 149288.17 | 315062.94 | 54845.96 |  | 12.28 | 18.0 | 21.1 |  |
| sp\|Q9Y2L5\|TPPC8\_HUMAN | Q9Y2L5 | TPPC8\_HUMAN | TRAPPC8 | Trafficking protein particle complex subunit 8 | 1435 | 160.997 | 2.117990e+03 | 4.284092e+05 | 149221.73 | 279187.44 |  |  | 14.56 | 27.6 |  |  |
| sp\|Q9P2J5\|SYLC\_HUMAN | Q9P2J5 | SYLC\_HUMAN | LARS1 | Leucine--tRNA ligase, cytoplasmic | 1176 | 134.466 | 9.806970e+02 | 1.656779e+05 | 149180.64 | 16497.227 |  |  | 10.37 | 32.6 |  |  |
| sp\|P14672\|GLUT4\_HUMAN | P14672 | GLUT4\_HUMAN | SLC2A4 | Solute carrier family 2, facilitated glucose transporter member 4 | 509 | 54.787 | 7.178270e+03 | 4.940996e+05 | 149096.53 | 231119.4 | 37643.234 | 76240.41 | 7.27 | 8.4 | 8.4 | 3.5 |
| sp\|Q969V3\|NCLN\_HUMAN | Q969V3 | NCLN\_HUMAN | NCLN | B | 563 | 62.974 | 1.679510e+04 | 1.328805e+06 | 148618.97 | 59357.664 | 748178.4 | 372649.66 | 12.79 | 18.3 | 30.6 | 15.8 |
| sp\|O43169\|CYB5B\_HUMAN | O43169 | CYB5B\_HUMAN | CYB5B | Cytochrome b5 type B | 150 | 16.695 | 3.738829e+04 | 7.842222e+05 | 148308.38 | 354175.62 | 97136.25 | 184601.97 | 43.33 | 26.7 | 10.7 | 18.7 |
| sp\|O15260\|SURF4\_HUMAN | O15260 | SURF4\_HUMAN | SURF4 | Surfeit locus protein 4 | 269 | 30.394 | 3.022641e+04 | 1.154228e+06 | 148084.89 | 270085.7 |  | 736057.3 | 8.55 | 30.5 |  | 11.5 |
| sp\|Q96QA5\|GSDMA\_HUMAN | Q96QA5 | GSDMA\_HUMAN | GSDMA | Gasdermin-A | 445 | 49.365 | 3.045039e+04 | 1.888553e+06 | 147891.3 |  | 739982.3 | 1000679.5 | 13.71 |  | 11.5 | 18.4 |
| sp\|Q9NW68\|BSDC1\_HUMAN | Q9NW68 | BSDC1\_HUMAN | BSDC1 | BSD domain-containing protein 1 | 430 | 47.163 | 5.942502e+03 | 3.521177e+05 | 147433.34 | 204684.4 |  |  | 12.79 | 31.9 |  |  |
| sp\|Q9P260\|RELCH\_HUMAN | Q9P260 | RELCH\_HUMAN | RELCH | RAB11-binding protein RELCH | 1216 | 134.630 | 2.103407e+03 | 3.557807e+05 | 146925.73 | 173959.62 |  | 34895.344 | 10.28 | 38.2 |  | 5.3 |
| sp\|P54760\|EPHB4\_HUMAN | P54760 | EPHB4\_HUMAN | EPHB4 | Ephrin type-B receptor 4 | 987 | 108.270 | 1.008678e+04 | 1.372075e+06 | 146853.58 | 78504.49 | 638119.9 | 508597.12 | 11.96 | 43.3 | 20.2 | 20.7 |
| sp\|P49006\|MRP\_HUMAN | P49006 | MRP\_HUMAN | MARCKSL1 | MARCKS-related protein | 195 | 19.529 | 3.121541e+04 | 7.658895e+05 | 146744.72 | 352307.38 | 131384.6 | 135452.83 | 33.85 | 24.6 | 24.6 | 23.6 |
| sp\|P27816\|MAP4\_HUMAN | P27816 | MAP4\_HUMAN | MAP4 | Microtubule-associated protein 4 | 1152 | 121.005 | 2.282177e+03 | 3.469523e+05 | 146252.06 | 55754.574 | 80007.39 | 64938.23 | 9.03 | 44.9 | 29.9 | 15.7 |
| sp\|Q9Y5B0\|CTDP1\_HUMAN | Q9Y5B0 | CTDP1\_HUMAN | CTDP1 | RNA polymerase II subunit A C-terminal domain phosphatase | 961 | 104.399 | 2.831144e+03 | 3.713432e+05 | 146154.31 | 203592.69 | 21596.217 |  | 10.82 | 49.5 | 27.1 |  |
| sp\|P02647\|APOA1\_HUMAN | P02647 | APOA1\_HUMAN | APOA1 | Apolipoprotein A-I | 267 | 30.778 | 3.332515e+04 | 1.288634e+06 | 144300.45 | 387155.5 | 388508.97 | 368669.28 | 36.33 | 62.2 | 61.4 | 25.8 |
| sp\|Q8WU76\|SCFD2\_HUMAN | Q8WU76 | SCFD2\_HUMAN | SCFD2 | Sec1 family domain-containing protein 2 | 684 | 75.127 | 8.925768e+03 | 8.424784e+05 | 144178.5 | 350993.12 | 263547.75 | 83759.05 | 8.48 | 41.2 | 9.8 | 6.9 |
| sp\|P48449\|LSS\_HUMAN | P48449 | LSS\_HUMAN | LSS | Lanosterol synthase | 732 | 83.309 | 2.914805e+03 | 3.050834e+05 | 143820.08 | 161263.34 |  |  | 17.62 | 30.2 |  |  |
| sp\|P16333\|NCK1\_HUMAN | P16333 | NCK1\_HUMAN | NCK1 | Cytoplasmic protein NCK1 | 377 | 42.864 | 3.911444e+03 | 2.106430e+05 | 143326.84 | 59351.285 |  | 7964.849 | 9.28 | 30.8 |  | 10.3 |
| sp\|P25445\|TNR6\_HUMAN | P25445 | TNR6\_HUMAN | FAS | Tumor necrosis factor receptor superfamily member 6 | 335 | 37.732 | 4.849246e+04 | 2.298800e+06 | 143245.31 | 369679.62 | 674749.7 | 1111125.5 | 14.93 | 35.8 | 29.3 | 23.0 |
| sp\|Q96Q05\|TPPC9\_HUMAN | Q96Q05 | TPPC9\_HUMAN | TRAPPC9 | Trafficking protein particle complex subunit 9 | 1148 | 126.280 | 9.012022e+02 | 1.429796e+05 | 142979.56 |  |  |  | 8.8 |  |  |  |
| sp\|Q9UIM3\|FKBPL\_HUMAN | Q9UIM3 | FKBPL\_HUMAN | FKBPL | FK506-binding protein-like | 349 | 38.176 | 7.802338e+03 | 3.742246e+05 | 142046.27 | 208471.88 |  | 23706.46 | 18.62 | 60.7 |  | 13.2 |
| sp\|Q86Y82\|STX12\_HUMAN | Q86Y82 | STX12\_HUMAN | STX12 | Syntaxin-12 | 276 | 31.642 | 2.413885e+04 | 9.596164e+05 | 141839.94 | 63308.36 | 119522.84 | 634945.3 | 23.55 | 23.2 | 26.8 | 30.1 |
| sp\|P14415\|AT1B2\_HUMAN | P14415 | AT1B2\_HUMAN | ATP1B2 | Sodium/potassium-transporting ATPase subunit beta-2 | 290 | 33.367 | 5.318720e+04 | 2.229674e+06 | 141359.02 | 136138.73 | 861307.44 | 1090869.1 | 4.83 | 33.8 | 20.0 | 14.5 |
| sp\|O60927\|PP1RB\_HUMAN | O60927 | PP1RB\_HUMAN | PPP1R11 | E3 ubiquitin-protein ligase PPP1R11 | 126 | 13.953 | 1.255079e+04 | 2.200167e+05 | 140823.16 | 79193.58 |  |  | 33.33 | 34.1 |  |  |
| sp\|Q9Y3D8\|KAD6\_HUMAN | Q9Y3D8 | KAD6\_HUMAN | AK6 | Adenylate kinase isoenzyme 6 | 172 | 20.061 | 6.889135e+03 | 1.736339e+05 | 139970.5 |  |  | 33663.367 | 11.05 |  |  | 11.0 |
| sp\|Q96SU4\|OSBL9\_HUMAN | Q96SU4 | OSBL9\_HUMAN | OSBPL9 | Oxysterol-binding protein-related protein 9 | 736 | 83.185 | 5.177864e+03 | 5.411439e+05 | 139920.03 | 401223.9 |  |  | 16.17 | 21.5 |  |  |
| sp\|Q9UKX7\|NUP50\_HUMAN | Q9UKX7 | NUP50\_HUMAN | NUP50 | Nuclear pore complex protein Nup50 | 468 | 50.144 | 7.382630e+03 | 4.651009e+05 | 139743.67 | 257106.44 | 49562.895 | 18687.943 | 14.74 | 65.0 | 24.6 | 15.2 |
| sp\|Q9UNE7\|CHIP\_HUMAN | Q9UNE7 | CHIP\_HUMAN | STUB1 | E3 ubiquitin-protein ligase CHIP | 303 | 34.856 | 1.743130e+04 | 7.633513e+05 | 139436.02 | 345760.94 | 278154.34 |  | 14.19 | 44.6 | 48.2 |  |
| sp\|Q13045\|FLII\_HUMAN | Q13045 | FLII\_HUMAN | FLII | Protein flightless-1 homolog | 1269 | 144.751 | 1.410549e+03 | 2.565233e+05 | 139290.75 | 117232.51 |  |  | 9.61 | 31.3 |  |  |
| sp\|Q92482\|AQP3\_HUMAN | Q92482 | AQP3\_HUMAN | AQP3 | Aquaporin-3 | 292 | 31.544 | 4.710849e+04 | 1.866952e+06 | 139203.03 | 58868.73 | 1104344.2 | 564535.94 | 2.74 | 13.0 | 4.5 | 5.1 |
| sp\|Q13485\|SMAD4\_HUMAN | Q13485 | SMAD4\_HUMAN | SMAD4 | Mothers against decapentaplegic homolog 4 | 552 | 60.439 | 4.733340e+03 | 3.594199e+05 | 139086.98 | 220332.9 |  |  | 19.93 | 29.2 |  |  |
| sp\|Q9Y6B6\|SAR1B\_HUMAN | Q9Y6B6 | SAR1B\_HUMAN | SAR1B | GTP-binding protein SAR1b | 198 | 22.410 | 7.506852e+04 | 2.113571e+06 | 138968.53 | 1546404.5 | 262583.4 | 165614.78 | 32.83 | 42.4 | 38.9 | 19.7 |
| sp\|Q8NHG7\|SVIP\_HUMAN | Q8NHG7 | SVIP\_HUMAN | SVIP | Small VCP/p97-interacting protein | 77 | 8.443 | 2.251988e+06 | 2.388801e+07 | 138896.03 | 63147.188 | 11902256.0 | 11783709.0 | 33.77 | 70.1 | 59.7 | 71.4 |
| sp\|P12268\|IMDH2\_HUMAN | P12268 | IMDH2\_HUMAN | IMPDH2 | Inosine-5'-monophosphate dehydrogenase 2 | 514 | 55.805 | 3.979920e+04 | 2.790388e+06 | 138369.84 | 117601.92 | 1592822.1 | 941593.9 | 9.73 | 18.1 | 15.6 | 10.9 |
| sp\|Q01105\|SET\_HUMAN | Q01105 | SET\_HUMAN | SET | Protein SET | 290 | 33.489 | 4.832067e+03 | 2.033070e+05 | 137085.48 |  | 31354.938 | 34866.547 | 17.93 |  | 19.3 | 12.1 |
| sp\|P41250\|GARS\_HUMAN | P41250 | GARS\_HUMAN | GARS1 | Glycine--tRNA ligase | 739 | 83.166 | 1.718955e+03 | 1.796088e+05 | 136826.6 | 42782.18 |  |  | 6.09 | 35.5 |  |  |
| sp\|P49790\|NU153\_HUMAN | P49790 | NU153\_HUMAN | NUP153 | Nuclear pore complex protein Nup153 | 1475 | 153.938 | 6.732163e+03 | 1.302020e+06 | 135993.77 | 418286.22 | 535418.06 | 212321.8 | 12.68 | 33.9 | 35.5 | 20.5 |
| sp\|Q9NXN4\|GDAP2\_HUMAN | Q9NXN4 | GDAP2\_HUMAN | GDAP2 | Ganglioside-induced differentiation-associated protein 2 | 497 | 56.225 | 5.222885e+03 | 3.689411e+05 | 135870.8 | 233070.34 |  |  | 11.07 | 36.0 |  |  |
| sp\|P31146\|COR1A\_HUMAN | P31146 | COR1A\_HUMAN | CORO1A | Coronin-1A | 461 | 51.026 | 8.676546e+03 | 5.562315e+05 | 134602.92 | 275370.66 | 88576.31 | 57681.6 | 11.28 | 48.2 | 34.1 | 19.1 |
| sp\|Q6ZV70\|LANC3\_HUMAN | Q6ZV70 | LANC3\_HUMAN | LANCL3 | LanC-like protein 3 | 420 | 46.319 | 3.058299e+03 | 1.779739e+05 | 134478.34 | 43495.523 |  |  | 11.9 | 24.0 |  |  |
| sp\|Q9P2Q2\|FRM4A\_HUMAN | Q9P2Q2 | FRM4A\_HUMAN | FRMD4A | FERM domain-containing protein 4A | 1039 | 115.458 | 4.016078e+03 | 5.825636e+05 | 134395.05 | 296780.78 | 121005.25 | 30382.477 | 9.43 | 41.0 | 19.6 | 13.0 |
| sp\|Q9ULT8\|HECD1\_HUMAN | Q9ULT8 | HECD1\_HUMAN | HECTD1 | E3 ubiquitin-protein ligase HECTD1 | 2610 | 289.368 | 1.343709e+03 | 4.885095e+05 | 134318.64 | 271559.9 | 53408.457 | 29222.516 | 5.25 | 31.1 | 15.2 | 7.5 |
| sp\|Q14457\|BECN1\_HUMAN | Q14457 | BECN1\_HUMAN | BECN1 | Beclin-1 | 450 | 51.896 | 9.081626e+03 | 5.921267e+05 | 133761.86 | 340947.7 | 117417.125 |  | 11.11 | 32.2 | 18.4 |  |
| sp\|Q9H871\|RMD5A\_HUMAN | Q9H871 | RMD5A\_HUMAN | RMND5A | E3 ubiquitin-protein transferase RMND5A | 391 | 43.993 | 1.211129e+04 | 6.694084e+05 | 133760.25 | 535648.2 |  |  | 15.86 | 34.8 |  |  |
| sp\|Q676U5\|A16L1\_HUMAN | Q676U5 | A16L1\_HUMAN | ATG16L1 | Autophagy-related protein 16-1 | 607 | 68.265 | 4.442426e+03 | 3.810092e+05 | 133699.55 | 222890.03 | 24419.576 |  | 6.26 | 26.0 | 19.1 |  |
| sp\|P14649\|MYL6B\_HUMAN | P14649 | MYL6B\_HUMAN | MYL6B | Myosin light chain 6B | 208 | 22.880 | 4.609079e+03 | 1.324913e+05 | 132491.28 |  |  |  | 6.25 |  |  |  |
| sp\|O95487\|SC24B\_HUMAN | O95487 | SC24B\_HUMAN | SEC24B | Protein transport protein Sec24B | 1268 | 137.418 | 5.655226e+03 | 9.763618e+05 | 132460.89 | 307622.5 | 442116.47 | 94161.9 | 9.54 | 20.3 | 14.2 | 4.2 |
| sp\|Q13459\|MYO9B\_HUMAN | Q13459 | MYO9B\_HUMAN | MYO9B | Unconventional myosin-IXb | 2157 | 243.401 | 7.562895e+03 | 2.312744e+06 | 131828.5 | 1957494.0 | 168631.7 | 54789.91 | 8.3 | 42.5 | 19.7 | 9.2 |
| sp\|P34949\|MPI\_HUMAN | P34949 | MPI\_HUMAN | MPI | Mannose-6-phosphate isomerase | 423 | 46.656 | 5.033981e+03 | 2.950776e+05 | 131708.88 | 163368.72 |  |  | 16.78 | 31.0 |  |  |
| sp\|O15143\|ARC1B\_HUMAN | O15143 | ARC1B\_HUMAN | ARPC1B | Actin-related protein 2/3 complex subunit 1B | 372 | 40.950 | 8.545220e+03 | 4.396370e+05 | 131707.31 | 237999.53 | 69930.21 |  | 15.59 | 33.9 | 12.4 |  |
| sp\|Q9H5N1\|RABE2\_HUMAN | Q9H5N1 | RABE2\_HUMAN | RABEP2 | Rab GTPase-binding effector protein 2 | 569 | 63.543 | 4.373379e+03 | 3.491419e+05 | 131549.0 | 206252.22 |  | 11340.675 | 18.98 | 43.8 |  | 15.5 |
| sp\|Q8N1F7\|NUP93\_HUMAN | Q8N1F7 | NUP93\_HUMAN | NUP93 | Nuclear pore complex protein Nup93 | 819 | 93.488 | 1.978607e+03 | 2.323981e+05 | 130804.83 | 75138.95 |  | 26454.287 | 14.41 | 33.2 |  | 9.4 |
| sp\|P0C870\|JMJD7\_HUMAN | P0C870 | JMJD7\_HUMAN | JMJD7 | Bifunctional peptidase and (3S)-lysyl hydroxylase JMJD7 | 316 | 35.932 | 3.641430e+03 | 1.643882e+05 | 130297.54 | 34090.625 |  |  | 18.67 | 11.1 |  |  |
| sp\|Q9BXW6\|OSBL1\_HUMAN | Q9BXW6 | OSBL1\_HUMAN | OSBPL1A | Oxysterol-binding protein-related protein 1 | 950 | 108.470 | 2.029038e+03 | 2.765140e+05 | 130222.47 | 146291.5 |  |  | 10.63 | 37.3 |  |  |
| sp\|Q7KYR7\|BT2A1\_HUMAN | Q7KYR7 | BT2A1\_HUMAN | BTN2A1 | Butyrophilin subfamily 2 member A1 | 527 | 59.633 | 7.318170e+03 | 5.482849e+05 | 130137.97 |  | 264492.34 | 153654.61 | 6.45 |  | 17.3 | 13.1 |
| sp\|Q9NZ43\|USE1\_HUMAN | Q9NZ43 | USE1\_HUMAN | USE1 | Vesicle transport protein USE1 | 259 | 29.371 | 2.777271e+04 | 1.024835e+06 | 128443.89 | 422794.6 | 136220.75 | 337376.25 | 16.6 | 40.2 | 34.4 | 30.1 |
| sp\|Q8NEZ5\|FBX22\_HUMAN | Q8NEZ5 | FBX22\_HUMAN | FBXO22 | F-box only protein 22 | 403 | 44.508 | 5.550874e+03 | 3.103964e+05 | 128388.29 | 182008.08 |  |  | 12.16 | 28.8 |  |  |
| sp\|Q9Y639\|NPTN\_HUMAN | Q9Y639 | NPTN\_HUMAN | NPTN | Neuroplastin | 398 | 44.387 | 5.231032e+04 | 2.917160e+06 | 128045.8 | 345696.38 | 2174350.0 | 269068.25 | 10.05 | 19.6 | 23.6 | 26.1 |
| sp\|Q15283\|RASA2\_HUMAN | Q15283 | RASA2\_HUMAN | RASA2 | Ras GTPase-activating protein 2 | 850 | 96.614 | 3.504006e+03 | 4.253261e+05 | 127403.27 | 297922.8 |  |  | 14.82 | 26.9 |  |  |
| sp\|Q02224\|CENPE\_HUMAN | Q02224 | CENPE\_HUMAN | CENPE | Centromere-associated protein E | 2701 | 297.110 | 3.405758e+02 | 1.271301e+05 | 127130.05 |  |  |  | 0.7 |  |  |  |
| sp\|Q15370\|ELOB\_HUMAN | Q15370 | ELOB\_HUMAN | ELOB | Elongin-B | 118 | 13.133 | 1.187175e+05 | 1.958826e+06 | 126811.39 | 1189851.0 | 483173.47 | 158990.38 | 10.17 | 39.0 | 35.6 | 28.0 |
| sp\|Q9UH62\|ARMX3\_HUMAN | Q9UH62 | ARMX3\_HUMAN | ARMCX3 | Armadillo repeat-containing X-linked protein 3 | 379 | 42.501 | 1.238972e+04 | 6.615729e+05 | 126541.336 | 535031.6 |  |  | 30.87 | 31.9 |  |  |
| sp\|Q96S38\|KS6C1\_HUMAN | Q96S38 | KS6C1\_HUMAN | RPS6KC1 | Ribosomal protein S6 kinase delta-1 | 1066 | 117.260 | 8.560323e+02 | 1.261122e+05 | 126112.22 |  |  |  | 6.19 |  |  |  |
| sp\|Q96Q11\|TRNT1\_HUMAN | Q96Q11 | TRNT1\_HUMAN | TRNT1 | CCA tRNA nucleotidyltransferase 1, mitochondrial | 434 | 50.128 | 5.291705e+03 | 3.332677e+05 | 126037.81 | 207229.86 |  |  | 19.59 | 21.9 |  |  |
| sp\|P69849\|NOMO3\_HUMAN | P69849 | NOMO3\_HUMAN | NOMO3 | B | 1222 | 134.134 | 5.350882e+03 | 9.017401e+05 | 125871.69 |  | 660635.1 | 115233.31 | 3.36 |  | 26.4 | 8.8 |
| sp\|Q6ZMI0\|PPR21\_HUMAN | Q6ZMI0 | PPR21\_HUMAN | PPP1R21 | Protein phosphatase 1 regulatory subunit 21 | 780 | 88.314 | 4.627986e+03 | 5.134980e+05 | 125584.83 | 371323.25 | 16589.945 |  | 13.85 | 45.6 | 18.2 |  |
| sp\|P01024\|CO3\_HUMAN | P01024 | CO3\_HUMAN | C3 | Complement C3 | 1663 | 187.148 | 1.555612e+04 | 3.657662e+06 | 125299.31 | 2251493.8 | 845401.4 | 435467.4 | 4.63 | 41.9 | 34.2 | 20.4 |
| sp\|P63151\|2ABA\_HUMAN | P63151 | 2ABA\_HUMAN | PPP2R2A | Serine/threonine-protein phosphatase 2A 55 kDa regulatory subunit B alpha isoform | 447 | 51.692 | 2.739561e+03 | 1.779186e+05 | 124868.89 |  | 40340.375 | 12709.375 | 14.99 |  | 16.8 | 18.6 |
| sp\|P55854\|SUMO3\_HUMAN | P55854 | SUMO3\_HUMAN | SUMO3 | Small ubiquitin-related modifier 3 | 103 | 11.637 | 2.249609e+04 | 3.289010e+05 | 124602.266 |  |  | 204298.73 | 11.65 |  |  | 22.3 |
| sp\|O60493\|SNX3\_HUMAN | O60493 | SNX3\_HUMAN | SNX3 | Sorting nexin-3 | 162 | 18.762 | 1.992309e+04 | 4.696270e+05 | 124539.5 | 345087.53 |  |  | 11.11 | 35.8 |  |  |
| sp\|Q8WWI5\|CTL1\_HUMAN | Q8WWI5 | CTL1\_HUMAN | SLC44A1 | Choline transporter-like protein 1 | 657 | 73.302 | 2.195070e+04 | 2.021536e+06 | 123944.07 | 100357.05 | 495826.12 | 1301408.4 | 6.24 | 35.0 | 28.0 | 21.6 |
| sp\|P16885\|PLCG2\_HUMAN | P16885 | PLCG2\_HUMAN | PLCG2 | 1-phosphatidylinositol 4,5-bisphosphate phosphodiesterase gamma-2 | 1265 | 139.150 | 7.070261e+02 | 1.236049e+05 | 123604.94 |  |  |  | 4.19 |  |  |  |
| sp\|Q9H3F6\|BACD3\_HUMAN | Q9H3F6 | BACD3\_HUMAN | KCTD10 | BTB/POZ domain-containing adapter for CUL3-mediated RhoA degradation protein 3 | 313 | 35.432 | 6.752126e+03 | 3.005754e+05 | 123308.09 | 38563.4 | 138703.88 |  | 6.39 | 31.3 | 26.2 |  |
| sp\|Q9H6Y2\|WDR55\_HUMAN | Q9H6Y2 | WDR55\_HUMAN | WDR55 | WD repeat-containing protein 55 | 383 | 42.070 | 3.713980e+03 | 1.963040e+05 | 123063.21 | 73240.84 |  |  | 14.36 | 30.0 |  |  |
| sp\|Q99943\|PLCA\_HUMAN | Q99943 | PLCA\_HUMAN | AGPAT1 | 1-acyl-sn-glycerol-3-phosphate acyltransferase alpha | 283 | 31.130 | 3.146442e+03 | 1.230597e+05 | 123059.734 |  |  |  | 17.67 |  |  |  |
| sp\|P40121\|CAPG\_HUMAN | P40121 | CAPG\_HUMAN | CAPG | Macrophage-capping protein | 348 | 38.499 | 1.377119e+04 | 6.660978e+05 | 122741.55 | 363108.75 | 118282.8 | 61964.72 | 10.63 | 36.8 | 15.2 | 9.8 |
| sp\|Q15172\|2A5A\_HUMAN | Q15172 | 2A5A\_HUMAN | PPP2R5A | Serine/threonine-protein phosphatase 2A 56 kDa regulatory subunit alpha isoform | 486 | 56.194 | 2.840741e+03 | 2.005574e+05 | 122713.47 | 77843.92 |  |  | 18.72 | 13.0 |  |  |
| sp\|Q96GA7\|SDSL\_HUMAN | Q96GA7 | SDSL\_HUMAN | SDSL | Serine dehydratase-like | 329 | 34.674 | 4.952550e+03 | 2.157496e+05 | 122233.41 | 93516.18 |  |  | 20.06 | 38.6 |  |  |
| sp\|Q9NZC3\|GDE1\_HUMAN | Q9NZC3 | GDE1\_HUMAN | GDE1 | Glycerophosphodiester phosphodiesterase 1 | 331 | 37.718 | 1.891040e+04 | 8.961206e+05 | 121938.03 | 285975.16 | 195769.7 | 292437.7 | 14.2 | 39.9 | 21.1 | 19.3 |
| sp\|Q9Y2I8\|WDR37\_HUMAN | Q9Y2I8 | WDR37\_HUMAN | WDR37 | WD repeat-containing protein 37 | 494 | 54.665 | 4.461655e+03 | 3.064238e+05 | 121925.37 | 101168.08 | 83330.33 |  | 9.72 | 27.1 | 27.7 |  |
| sp\|Q96MV1\|TLCD4\_HUMAN | Q96MV1 | TLCD4\_HUMAN | TLCD4 | TLC domain-containing protein 4 | 263 | 30.041 | 3.350865e+04 | 1.264703e+06 | 120942.625 | 879114.6 | 147527.84 | 117117.71 | 11.79 | 27.0 | 17.5 | 6.8 |
| sp\|Q9Y5Y2\|NUBP2\_HUMAN | Q9Y5Y2 | NUBP2\_HUMAN | NUBP2 | Cytosolic Fe-S cluster assembly factor NUBP2 | 271 | 28.825 | 8.936309e+03 | 3.236269e+05 | 120896.03 | 202730.89 |  |  | 13.28 | 30.3 |  |  |
| sp\|O00560\|SDCB1\_HUMAN | O00560 | SDCB1\_HUMAN | SDCBP | Syntenin-1 | 298 | 32.444 | 2.018585e+04 | 8.228083e+05 | 120778.03 | 631312.5 | 70717.74 |  | 21.14 | 52.3 | 23.2 |  |
| sp\|Q9H2C0\|GAN\_HUMAN | Q9H2C0 | GAN\_HUMAN | GAN | Gigaxonin | 597 | 67.638 | 6.162068e+03 | 5.236419e+05 | 120299.94 | 403341.94 |  |  | 7.37 | 47.6 |  |  |
| sp\|P80303\|NUCB2\_HUMAN | P80303 | NUCB2\_HUMAN | NUCB2 | Nucleobindin-2 | 420 | 50.223 | 3.540223e+04 | 2.233832e+06 | 120158.69 | 602397.25 | 1390368.8 | 120907.02 | 14.05 | 48.1 | 47.6 | 20.7 |
| sp\|P28749\|RBL1\_HUMAN | P28749 | RBL1\_HUMAN | RBL1 | Retinoblastoma-like protein 1 | 1068 | 117.480 | 8.138009e+02 | 1.201156e+05 | 120115.56 |  |  |  | 1.97 |  |  |  |
| sp\|Q14145\|KEAP1\_HUMAN | Q14145 | KEAP1\_HUMAN | KEAP1 | Kelch-like ECH-associated protein 1 | 624 | 69.666 | 2.283077e+03 | 1.998290e+05 | 120025.3 | 79803.7 |  |  | 16.19 | 52.2 |  |  |
| sp\|P78310\|CXAR\_HUMAN | P78310 | CXAR\_HUMAN | CXADR | Coxsackievirus and adenovirus receptor | 365 | 40.030 | 2.876754e+04 | 1.446790e+06 | 119465.92 | 171847.27 | 338344.75 | 817132.2 | 9.32 | 60.0 | 40.3 | 28.8 |
| sp\|O14841\|OPLA\_HUMAN | O14841 | OPLA\_HUMAN | OPLAH | 5-oxoprolinase | 1288 | 137.457 | 1.226998e+03 | 2.118984e+05 | 119267.92 | 92630.51 |  |  | 14.83 | 41.4 |  |  |
| sp\|P02545\|LMNA\_HUMAN | P02545 | LMNA\_HUMAN | LMNA | Prelamin-A/C | 664 | 74.139 | 6.815343e+04 | 6.348215e+06 | 118674.36 | 186907.53 | 574896.4 | 5467736.5 | 16.42 | 41.3 | 46.2 | 36.1 |
| sp\|Q8NEY1\|NAV1\_HUMAN | Q8NEY1 | NAV1\_HUMAN | NAV1 | Neuron navigator 1 | 1877 | 202.472 | 1.268643e+03 | 3.227167e+05 | 118280.05 | 204436.61 |  |  | 1.28 | 40.3 |  |  |
| sp\|Q9NQG5\|RPR1B\_HUMAN | Q9NQG5 | RPR1B\_HUMAN | RPRD1B | Regulation of nuclear pre-mRNA domain-containing protein 1B | 326 | 36.900 | 9.632463e+03 | 4.465611e+05 | 118070.1 | 328491.0 |  |  | 23.62 | 53.4 |  |  |
| sp\|P45880\|VDAC2\_HUMAN | P45880 | VDAC2\_HUMAN | VDAC2 | Voltage-dependent anion-selective channel protein 2 | 294 | 32.340 | 2.903403e+03 | 1.179681e+05 | 117968.06 |  |  |  | 17.01 |  |  |  |
| sp\|Q9ULV4\|COR1C\_HUMAN | Q9ULV4 | COR1C\_HUMAN | CORO1C | Coronin-1C | 474 | 53.249 | 7.571786e+03 | 5.065554e+05 | 117629.75 | 327022.62 | 61903.0 |  | 16.46 | 30.0 | 20.0 |  |
| sp\|P27701\|CD82\_HUMAN | P27701 | CD82\_HUMAN | CD82 | CD82 antigen | 267 | 29.626 | 2.394370e+04 | 8.912129e+05 | 117594.125 |  | 249346.19 | 524272.56 | 10.11 |  | 8.2 | 8.2 |
| sp\|O00273\|DFFA\_HUMAN | O00273 | DFFA\_HUMAN | DFFA | DNA fragmentation factor subunit alpha | 331 | 36.522 | 4.403382e+04 | 2.020496e+06 | 117427.3 | 1903069.0 |  |  | 21.15 | 53.8 |  |  |
| sp\|Q8N584\|TT39C\_HUMAN | Q8N584 | TT39C\_HUMAN | TTC39C | Tetratricopeptide repeat protein 39C | 583 | 65.870 | 3.068401e+03 | 2.539317e+05 | 117425.68 | 136506.06 |  |  | 6.69 | 25.9 |  |  |
| sp\|Q96RG2\|PASK\_HUMAN | Q96RG2 | PASK\_HUMAN | PASK | PAS domain-containing serine/threonine-protein kinase | 1323 | 142.929 | 1.992841e+03 | 3.578575e+05 | 117407.36 | 240450.17 |  |  | 6.12 | 32.7 |  |  |
| sp\|O15269\|SPTC1\_HUMAN | O15269 | SPTC1\_HUMAN | SPTLC1 | Serine palmitoyltransferase 1 | 473 | 52.744 | 3.506044e+03 | 2.323312e+05 | 117195.21 |  | 99489.53 | 15646.497 | 2.96 |  | 10.8 | 5.7 |
| sp\|P98172\|EFNB1\_HUMAN | P98172 | EFNB1\_HUMAN | EFNB1 | Ephrin-B1 | 346 | 38.007 | 2.757331e+04 | 1.316648e+06 | 116933.39 | 125828.72 | 638859.56 | 435026.2 | 17.92 | 50.9 | 36.7 | 23.1 |
| sp\|Q14155\|ARHG7\_HUMAN | Q14155 | ARHG7\_HUMAN | ARHGEF7 | Rho guanine nucleotide exchange factor 7 | 803 | 90.012 | 1.086467e+04 | 1.228667e+06 | 116898.33 | 543224.3 | 568544.25 |  | 7.72 | 30.8 | 25.5 |  |
| sp\|Q6NSW5\|DE10B\_HUMAN | Q6NSW5 | DE10B\_HUMAN | DENND10P1 | Putative DENN domain-containing protein 10 B | 357 | 40.514 | 7.241449e+03 | 3.685936e+05 | 116687.71 | 251905.88 |  |  | 12.32 | 14.3 |  |  |
| sp\|Q32MZ4\|LRRF1\_HUMAN | Q32MZ4 | LRRF1\_HUMAN | LRRFIP1 | Leucine-rich repeat flightless-interacting protein 1 | 808 | 89.253 | 2.063564e+04 | 2.313971e+06 | 116017.29 | 313633.84 | 22518.062 | 1861801.5 | 10.27 | 51.6 | 25.1 | 16.1 |
| sp\|Q5VZM2\|RRAGB\_HUMAN | Q5VZM2 | RRAGB\_HUMAN | RRAGB | Ras-related GTP-binding protein B | 374 | 43.250 | 3.566591e+03 | 1.938012e+05 | 115830.49 |  | 77970.75 |  | 7.75 |  | 6.4 |  |
| sp\|Q8IWV8\|UBR2\_HUMAN | Q8IWV8 | UBR2\_HUMAN | UBR2 | E3 ubiquitin-protein ligase UBR2 | 1755 | 200.538 | 1.085635e+03 | 2.735254e+05 | 115829.54 | 157695.9 |  |  | 5.58 | 33.7 |  |  |
| sp\|Q6R327\|RICTR\_HUMAN | Q6R327 | RICTR\_HUMAN | RICTOR | Rapamycin-insensitive companion of mTOR | 1708 | 192.218 | 3.045963e+03 | 7.355901e+05 | 115331.33 | 201215.28 | 266979.72 | 152063.75 | 7.14 | 28.6 | 8.8 | 10.1 |
| sp\|Q9UBP0\|SPAST\_HUMAN | Q9UBP0 | SPAST\_HUMAN | SPAST | Spastin | 616 | 67.197 | 2.955337e+03 | 2.495020e+05 | 115233.266 | 134268.73 |  |  | 8.28 | 41.4 |  |  |
| sp\|Q13951\|PEBB\_HUMAN | Q13951 | PEBB\_HUMAN | CBFB | Core-binding factor subunit beta | 182 | 21.508 | 1.240449e+04 | 3.351939e+05 | 115182.83 | 220011.1 |  |  | 14.29 | 53.8 |  |  |
| sp\|O95810\|CAVN2\_HUMAN | O95810 | CAVN2\_HUMAN | CAVIN2 | Caveolae-associated protein 2 | 425 | 47.173 | 1.061688e+04 | 6.292275e+05 | 114717.984 | 340043.72 | 174465.83 |  | 19.53 | 42.1 | 27.5 |  |
| sp\|Q8NBJ5\|GT251\_HUMAN | Q8NBJ5 | GT251\_HUMAN | COLGALT1 | Procollagen galactosyltransferase 1 | 622 | 71.636 | 2.200299e+04 | 1.980296e+06 | 114704.375 | 66276.69 | 1643444.1 | 155870.81 | 8.36 | 37.1 | 31.2 | 16.2 |
| sp\|Q86X10\|RLGPB\_HUMAN | Q86X10 | RLGPB\_HUMAN | RALGAPB | Ral GTPase-activating protein subunit beta | 1494 | 166.799 | 1.713538e+03 | 3.590908e+05 | 114335.38 | 244755.45 |  |  | 7.43 | 33.5 |  |  |
| sp\|P00167\|CYB5\_HUMAN | P00167 | CYB5\_HUMAN | CYB5A | Cytochrome b5 | 134 | 15.330 | 5.237715e+04 | 1.008791e+06 | 114282.39 | 152178.47 | 588952.25 | 153377.8 | 35.82 | 49.3 | 32.1 | 29.1 |
| sp\|P16383\|GCFC2\_HUMAN | P16383 | GCFC2\_HUMAN | GCFC2 | Intron Large complex component GCFC2 | 781 | 89.385 | 3.815911e+03 | 4.285288e+05 | 114204.58 | 314324.22 |  |  | 11.4 | 32.4 |  |  |
| sp\|O95396\|MOCS3\_HUMAN | O95396 | MOCS3\_HUMAN | MOCS3 | Adenylyltransferase and sulfurtransferase MOCS3 | 460 | 49.669 | 3.645871e+03 | 2.275118e+05 | 114106.875 | 113404.88 |  |  | 20.65 | 56.5 |  |  |
| sp\|P49184\|DNSL1\_HUMAN | P49184 | DNSL1\_HUMAN | DNASE1L1 | Deoxyribonuclease-1-like 1 | 302 | 33.893 | 2.243350e+04 | 9.552657e+05 | 114105.19 |  | 288899.12 | 552261.4 | 26.49 |  | 11.6 | 8.6 |
| sp\|P62633\|CNBP\_HUMAN | P62633 | CNBP\_HUMAN | CNBP | CCHC-type zinc finger nucleic acid binding protein | 177 | 19.470 | 4.659619e+03 | 1.139813e+05 | 113981.266 |  |  |  | 18.64 |  |  |  |
| sp\|Q8TB03\|CX038\_HUMAN | Q8TB03 | CX038\_HUMAN | CXorf38 | Uncharacterized protein CXorf38 | 319 | 35.090 | 2.579133e+03 | 1.137036e+05 | 113703.59 |  |  |  | 9.72 |  |  |  |
| sp\|Q5VSP4\|LC1L1\_HUMAN | Q5VSP4 | LC1L1\_HUMAN | LCN1P1 | Putative lipocalin 1-like protein 1 | 162 | 17.820 | 5.077002e+03 | 1.136664e+05 | 113666.43 |  |  |  | 13.58 |  |  |  |
| sp\|Q9H3H3\|CK068\_HUMAN | Q9H3H3 | CK068\_HUMAN | C11orf68 | UPF0696 protein C11orf68 | 292 | 31.430 | 2.561995e+04 | 1.011672e+06 | 113468.055 | 898204.0 |  |  | 18.15 | 51.7 |  |  |
| sp\|P52306\|GDS1\_HUMAN | P52306 | GDS1\_HUMAN | RAP1GDS1 | Rap1 GTPase-GDP dissociation stimulator 1 | 607 | 66.317 | 2.701211e+03 | 2.250612e+05 | 113372.19 | 111688.984 |  |  | 14.99 | 45.0 |  |  |
| sp\|O60568\|PLOD3\_HUMAN | O60568 | PLOD3\_HUMAN | PLOD3 | Multifunctional procollagen lysine hydroxylase and glycosyltransferase LH3 | 738 | 84.785 | 9.155358e+03 | 9.752400e+05 | 112996.06 | 28188.97 | 735386.4 | 98668.61 | 5.15 | 32.8 | 25.1 | 15.7 |
| sp\|Q8WVT3\|TPC12\_HUMAN | Q8WVT3 | TPC12\_HUMAN | TRAPPC12 | Trafficking protein particle complex subunit 12 | 735 | 79.375 | 2.547110e+03 | 2.540087e+05 | 112663.91 | 141344.81 |  |  | 17.28 | 36.7 |  |  |
| sp\|Q92499\|DDX1\_HUMAN | Q92499 | DDX1\_HUMAN | DDX1 | ATP-dependent RNA helicase DDX1 | 740 | 82.432 | 3.229860e+03 | 3.345005e+05 | 112400.14 | 222100.34 |  |  | 15.14 | 46.4 |  |  |
| sp\|Q9BUJ2\|HNRL1\_HUMAN | Q9BUJ2 | HNRL1\_HUMAN | HNRNPUL1 | Heterogeneous nuclear ribonucleoprotein U-like protein 1 | 856 | 95.739 | 2.341086e+03 | 2.815940e+05 | 112354.03 | 169240.02 |  |  | 7.48 | 28.5 |  |  |
| sp\|O14975\|S27A2\_HUMAN | O14975 | S27A2\_HUMAN | SLC27A2 | Long-chain fatty acid transport protein 2 | 620 | 70.312 | 4.429213e+03 | 3.912670e+05 | 112053.2 |  | 75244.99 | 203968.77 | 9.52 |  | 22.9 | 8.4 |
| sp\|Q8N8R5\|CB069\_HUMAN | Q8N8R5 | CB069\_HUMAN | C2orf69 | Mitochondrial protein C2orf69 | 385 | 43.448 | 1.127823e+04 | 6.156412e+05 | 112025.016 | 503616.22 |  |  | 9.35 | 41.6 |  |  |
| sp\|Q9UKW4\|VAV3\_HUMAN | Q9UKW4 | VAV3\_HUMAN | VAV3 | Guanine nucleotide exchange factor VAV3 | 847 | 97.776 | 1.713361e+03 | 2.104740e+05 | 111698.625 | 98775.35 |  |  | 5.67 | 28.9 |  |  |
| sp\|P31151\|S10A7\_HUMAN | P31151 | S10A7\_HUMAN | S100A7 | Protein S100-A7 | 101 | 11.471 | 2.560948e+04 | 3.690789e+05 | 111372.625 | 44991.508 | 212714.72 |  | 21.78 | 76.2 | 62.4 |  |
| sp\|Q8IWF6\|DEN6A\_HUMAN | Q8IWF6 | DEN6A\_HUMAN | DENND6A | Protein DENND6A | 608 | 69.575 | 3.431341e+03 | 2.999399e+05 | 110950.47 | 188989.42 |  |  | 12.34 | 34.0 |  |  |
| sp\|Q5T6F0\|DCA12\_HUMAN | Q5T6F0 | DCA12\_HUMAN | DCAF12 | DDB1- and CUL4-associated factor 12 | 453 | 49.830 | 1.769708e+03 | 1.107923e+05 | 110792.336 |  |  |  | 5.3 |  |  |  |
| sp\|Q06203\|PUR1\_HUMAN | Q06203 | PUR1\_HUMAN | PPAT | Amidophosphoribosyltransferase | 517 | 57.399 | 3.682103e+03 | 2.655323e+05 | 110241.75 | 155290.56 |  |  | 13.93 | 31.5 |  |  |
| sp\|Q8IZ81\|ELMD2\_HUMAN | Q8IZ81 | ELMD2\_HUMAN | ELMOD2 | ELMO domain-containing protein 2 | 293 | 34.961 | 1.272044e+04 | 5.587316e+05 | 110231.95 | 103384.67 | 74405.7 | 270709.25 | 12.63 | 36.2 | 24.9 | 24.2 |
| sp\|Q9UDY6\|TRI10\_HUMAN | Q9UDY6 | TRI10\_HUMAN | TRIM10 | Tripartite motif-containing protein 10 | 481 | 55.037 | 6.019043e+03 | 4.161974e+05 | 110050.625 | 197243.27 | 108903.5 |  | 11.02 | 48.6 | 21.2 |  |
| sp\|O95721\|SNP29\_HUMAN | O95721 | SNP29\_HUMAN | SNAP29 | Synaptosomal-associated protein 29 | 258 | 28.970 | 3.267460e+04 | 1.189258e+06 | 110045.234 | 807233.5 | 185760.75 | 86218.016 | 17.83 | 55.4 | 25.2 | 25.6 |
| sp\|Q12767\|TMM94\_HUMAN | Q12767 | TMM94\_HUMAN | TMEM94 | Transmembrane protein 94 | 1356 | 151.151 | 1.781134e+03 | 3.382398e+05 | 109787.19 |  | 27306.56 | 201146.05 | 2.36 |  | 6.8 | 9.3 |
| sp\|P98196\|AT11A\_HUMAN | P98196 | AT11A\_HUMAN | ATP11A | Phospholipid-transporting ATPase IH | 1134 | 129.756 | 8.473992e+03 | 1.381442e+06 | 109529.65 |  | 342141.28 | 929771.0 | 4.67 |  | 27.2 | 20.2 |
| sp\|O14662\|STX16\_HUMAN | O14662 | STX16\_HUMAN | STX16 | Syntaxin-16 | 325 | 37.031 | 6.286164e+03 | 2.924612e+05 | 109454.914 | 109420.22 | 73586.08 |  | 15.08 | 35.1 | 19.4 |  |
| sp\|Q9UJX5\|APC4\_HUMAN | Q9UJX5 | APC4\_HUMAN | ANAPC4 | Anaphase-promoting complex subunit 4 | 808 | 92.116 | 2.838168e+03 | 3.284659e+05 | 109278.47 | 219187.44 |  |  | 7.43 | 34.5 |  |  |
| sp\|Q5W0Z9\|ZDH20\_HUMAN | Q5W0Z9 | ZDH20\_HUMAN | ZDHHC20 | Palmitoyltransferase ZDHHC20 | 365 | 42.278 | 3.693968e+04 | 1.962116e+06 | 108636.18 |  | 279399.3 | 1574080.5 | 10.41 |  | 29.0 | 20.5 |
| sp\|Q8N8N7\|PTGR2\_HUMAN | Q8N8N7 | PTGR2\_HUMAN | PTGR2 | Prostaglandin reductase 2 | 351 | 38.499 | 4.049704e+03 | 1.958799e+05 | 108492.46 | 87387.45 |  |  | 12.25 | 26.2 |  |  |
| sp\|A1A5C7\|S22AN\_HUMAN | A1A5C7 | S22AN\_HUMAN | SLC22A23 | Solute carrier family 22 member 23 | 686 | 73.748 | 3.800231e+04 | 3.521092e+06 | 108368.24 | 37951.902 | 1823382.0 | 1551390.0 | 8.31 | 31.8 | 22.7 | 22.4 |
| sp\|Q16563\|SYPL1\_HUMAN | Q16563 | SYPL1\_HUMAN | SYPL1 | Synaptophysin-like protein 1 | 259 | 28.565 | 1.186332e+05 | 4.257528e+06 | 108025.42 | 3379506.5 | 224715.6 | 545280.1 | 13.9 | 43.6 | 13.5 | 16.6 |
| sp\|P0C7P3\|SLN14\_HUMAN | P0C7P3 | SLN14\_HUMAN | SLFN14 | Protein SLFN14 | 912 | 103.907 | 2.106889e+03 | 2.750450e+05 | 107863.66 | 167181.3 |  |  | 7.46 | 47.5 |  |  |
| sp\|Q9Y6B7\|AP4B1\_HUMAN | Q9Y6B7 | AP4B1\_HUMAN | AP4B1 | AP-4 complex subunit beta-1 | 739 | 83.260 | 2.751642e+03 | 2.878362e+05 | 107846.4 | 127189.125 | 52800.688 |  | 4.33 | 28.0 | 21.7 |  |
| sp\|Q9UJX6\|ANC2\_HUMAN | Q9UJX6 | ANC2\_HUMAN | ANAPC2 | Anaphase-promoting complex subunit 2 | 822 | 93.828 | 3.936581e+03 | 4.640542e+05 | 107513.086 | 356541.16 |  |  | 7.54 | 33.0 |  |  |
| sp\|Q9Y6W5\|WASF2\_HUMAN | Q9Y6W5 | WASF2\_HUMAN | WASF2 | Actin-binding protein WASF2 | 498 | 54.284 | 5.263188e+04 | 3.589532e+06 | 107442.47 | 3399964.8 | 82124.875 |  | 6.22 | 27.7 | 14.9 |  |
| sp\|P30419\|NMT1\_HUMAN | P30419 | NMT1\_HUMAN | NMT1 | Glycylpeptide N-tetradecanoyltransferase 1 | 496 | 56.806 | 3.851901e+03 | 2.749075e+05 | 107350.86 | 167556.6 |  |  | 7.46 | 36.3 |  |  |
| sp\|O00303\|EIF3F\_HUMAN | O00303 | EIF3F\_HUMAN | EIF3F | Eukaryotic translation initiation factor 3 subunit F | 357 | 37.564 | 2.819951e+03 | 1.330855e+05 | 107201.64 |  | 25883.82 |  | 17.93 |  | 17.6 |  |
| sp\|Q8NHP6\|MSPD2\_HUMAN | Q8NHP6 | MSPD2\_HUMAN | MOSPD2 | Motile sperm domain-containing protein 2 | 518 | 59.746 | 6.462906e+03 | 4.851252e+05 | 106899.59 | 42683.83 | 89635.37 | 245906.38 | 3.67 | 31.3 | 27.2 | 12.9 |
| sp\|Q8N4A0\|GALT4\_HUMAN | Q8N4A0 | GALT4\_HUMAN | GALNT4 | Polypeptide N-acetylgalactosaminyltransferase 4 | 578 | 66.666 | 9.805559e+03 | 8.212850e+05 | 106658.58 |  | 498752.47 | 215873.98 | 4.67 |  | 21.3 | 14.2 |
| sp\|Q6P587\|FAHD1\_HUMAN | Q6P587 | FAHD1\_HUMAN | FAHD1 | Acylpyruvase FAHD1, mitochondrial | 224 | 24.843 | 9.616663e+03 | 3.001550e+05 | 106636.836 | 193518.16 |  |  | 30.36 | 63.8 |  |  |
| sp\|Q92945\|FUBP2\_HUMAN | Q92945 | FUBP2\_HUMAN | KHSRP | Far upstream element-binding protein 2 | 711 | 73.115 | 7.500161e+03 | 6.889604e+05 | 106620.81 | 72301.89 | 365922.06 | 144115.61 | 5.06 | 54.4 | 24.1 | 17.9 |
| sp\|P06748\|NPM\_HUMAN | P06748 | NPM\_HUMAN | NPM1 | Nucleophosmin | 294 | 32.575 | 1.127651e+04 | 4.615050e+05 | 106537.23 | 177258.1 | 144988.62 | 32721.09 | 11.56 | 30.6 | 25.5 | 20.4 |
| sp\|Q8N5C8\|TAB3\_HUMAN | Q8N5C8 | TAB3\_HUMAN | TAB3 | TGF-beta-activated kinase 1 and MAP3K7-binding protein 3 | 712 | 78.653 | 6.825643e+03 | 6.744908e+05 | 106195.61 | 554476.9 | 13818.3125 |  | 10.11 | 39.6 | 13.9 |  |
| sp\|Q9BW62\|KATL1\_HUMAN | Q9BW62 | KATL1\_HUMAN | KATNAL1 | Katanin p60 ATPase-containing subunit A-like 1 | 490 | 55.392 | 9.219156e+03 | 6.415868e+05 | 105400.19 | 536186.56 |  |  | 20.2 | 70.0 |  |  |
| sp\|P21291\|CSRP1\_HUMAN | P21291 | CSRP1\_HUMAN | CSRP1 | Cysteine and glycine-rich protein 1 | 193 | 20.567 | 1.223713e+04 | 3.162042e+05 | 105196.91 | 211007.34 |  |  | 24.35 | 75.6 |  |  |
| sp\|P41227\|NAA10\_HUMAN | P41227 | NAA10\_HUMAN | NAA10 | N-alpha-acetyltransferase 10 | 235 | 26.459 | 2.133651e+04 | 7.092740e+05 | 105119.016 | 582986.7 | 21168.275 | 0.0 | 31.06 | 48.9 | 22.1 | 14.5 |
| sp\|B5ME19\|EIFCL\_HUMAN | B5ME19 | EIFCL\_HUMAN | EIF3CL | Eukaryotic translation initiation factor 3 subunit C-like protein | 914 | 105.473 | 9.504869e+02 | 1.259519e+05 | 104873.016 | 21078.844 |  |  | 9.74 | 26.3 |  |  |
| sp\|Q9H1K0\|RBNS5\_HUMAN | Q9H1K0 | RBNS5\_HUMAN | RBSN | Rabenosyn-5 | 784 | 88.870 | 3.815957e+03 | 4.260649e+05 | 104312.76 | 250011.6 | 55522.906 | 16217.626 | 8.55 | 43.4 | 16.8 | 7.7 |
| sp\|Q70IA6\|MOB2\_HUMAN | Q70IA6 | MOB2\_HUMAN | MOB2 | MOB kinase activator 2 | 237 | 26.927 | 1.826340e+04 | 6.178551e+05 | 103424.73 | 376478.25 | 100941.4 | 37010.72 | 20.25 | 27.4 | 21.1 | 21.5 |
| sp\|Q8NEN9\|PDZD8\_HUMAN | Q8NEN9 | PDZD8\_HUMAN | PDZD8 | PDZ domain-containing protein 8 | 1154 | 128.563 | 5.506991e+03 | 8.895032e+05 | 103083.25 | 184856.4 | 296112.03 | 305451.5 | 10.49 | 25.7 | 27.3 | 20.0 |
| sp\|Q9Y608\|LRRF2\_HUMAN | Q9Y608 | LRRF2\_HUMAN | LRRFIP2 | Leucine-rich repeat flightless-interacting protein 2 | 721 | 82.171 | 2.480346e+03 | 2.560637e+05 | 102811.85 | 153251.86 |  |  | 6.66 | 29.5 |  |  |
| sp\|P50583\|AP4A\_HUMAN | P50583 | AP4A\_HUMAN | NUDT2 | Bis(5'-nucleosyl)-tetraphosphatase [asymmetrical] | 147 | 16.829 | 6.988282e+04 | 1.477563e+06 | 102742.57 | 1374820.0 |  |  | 10.2 | 81.6 |  |  |
| sp\|P07311\|ACYP1\_HUMAN | P07311 | ACYP1\_HUMAN | ACYP1 | Acylphosphatase-1 | 99 | 11.261 | 2.318356e+05 | 3.280003e+06 | 102354.02 | 2526478.0 | 585374.06 | 65797.016 | 22.22 | 98.0 | 49.5 | 20.2 |
| sp\|P43307\|SSRA\_HUMAN | P43307 | SSRA\_HUMAN | SSR1 | Translocon-associated protein subunit alpha | 286 | 32.235 | 4.231821e+04 | 1.713847e+06 | 102347.766 |  | 1462118.6 | 149380.9 | 11.89 |  | 6.6 | 6.6 |
| sp\|Q14694\|UBP10\_HUMAN | Q14694 | UBP10\_HUMAN | USP10 | Ubiquitin carboxyl-terminal hydrolase 10 | 798 | 87.134 | 2.558523e+03 | 2.800877e+05 | 101802.27 | 178285.44 |  |  | 11.9 | 29.2 |  |  |
| sp\|A1L0T0\|HACL2\_HUMAN | A1L0T0 | HACL2\_HUMAN | ILVBL | 2-hydroxyacyl-CoA lyase 2 | 632 | 67.868 | 2.025407e+03 | 1.727009e+05 | 101497.125 | 71203.76 |  |  | 9.18 | 41.9 |  |  |
| sp\|P54868\|HMCS2\_HUMAN | P54868 | HMCS2\_HUMAN | HMGCS2 | Hydroxymethylglutaryl-CoA synthase, mitochondrial | 508 | 55.880 | 1.444768e+03 | 1.014312e+05 | 101431.2 |  |  |  | 8.07 |  |  |  |
| sp\|Q8IZP0\|ABI1\_HUMAN | Q8IZP0 | ABI1\_HUMAN | ABI1 | Abl interactor 1 | 508 | 55.081 | 3.908135e+03 | 2.704509e+05 | 100470.734 | 169980.2 |  |  | 8.27 | 30.9 |  |  |
| sp\|Q7Z3B4\|NUP54\_HUMAN | Q7Z3B4 | NUP54\_HUMAN | NUP54 | Nucleoporin p54 | 507 | 55.435 | 8.233001e+03 | 5.734022e+05 | 100122.06 | 254441.9 | 218838.27 |  | 8.48 | 32.3 | 16.8 |  |
| sp\|Q9BRX2\|PELO\_HUMAN | Q9BRX2 | PELO\_HUMAN | PELO | Protein pelota homolog | 385 | 43.359 | 9.041979e+03 | 4.925608e+05 | 99724.3 | 378164.4 | 14672.145 |  | 15.84 | 41.3 | 17.1 |  |
| sp\|Q92620\|PRP16\_HUMAN | Q92620 | PRP16\_HUMAN | DHX38 | Pre-mRNA-splicing factor ATP-dependent RNA helicase PRP16 | 1227 | 140.503 | 2.114175e+03 | 3.732018e+05 | 99645.61 | 273556.2 |  |  | 8.72 | 28.0 |  |  |
| sp\|Q9BRZ2\|TRI56\_HUMAN | Q9BRZ2 | TRI56\_HUMAN | TRIM56 | E3 ubiquitin-protein ligase TRIM56 | 755 | 81.488 | 5.254015e+03 | 5.379008e+05 | 99302.28 | 345688.88 | 92909.59 |  | 10.07 | 63.0 | 23.8 |  |
| sp\|P06454\|PTMA\_HUMAN | P06454 | PTMA\_HUMAN | PTMA | Prothymosin alpha | 111 | 12.203 | 8.719077e+04 | 1.336763e+06 | 99299.33 | 1112002.5 | 89098.05 | 36362.633 | 21.62 | 41.4 | 26.1 | 25.2 |
| sp\|Q9C0C7\|AMRA1\_HUMAN | Q9C0C7 | AMRA1\_HUMAN | AMBRA1 | Activating molecule in BECN1-regulated autophagy protein 1 | 1298 | 142.507 | 1.514131e+03 | 2.710922e+05 | 99269.734 | 134168.8 | 37653.617 |  | 14.33 | 32.1 | 21.3 |  |
| sp\|Q9NQ89\|CL004\_HUMAN | Q9NQ89 | CL004\_HUMAN | C12orf4 | Protein C12orf4 | 552 | 63.801 | 3.287895e+03 | 2.635497e+05 | 99163.14 | 164386.6 |  |  | 8.51 | 33.7 |  |  |
| sp\|Q9H330\|TM245\_HUMAN | Q9H330 | TM245\_HUMAN | TMEM245 | Transmembrane protein 245 | 879 | 97.357 | 8.990464e+03 | 1.099680e+06 | 98751.63 | 234321.4 | 150675.56 | 615931.75 | 4.21 | 16.5 | 11.1 | 5.6 |
| sp\|Q9H1E5\|TMX4\_HUMAN | Q9H1E5 | TMX4\_HUMAN | TMX4 | Thioredoxin-related transmembrane protein 4 | 349 | 38.952 | 4.546785e+04 | 2.225109e+06 | 98290.05 | 139469.3 | 987889.8 | 999460.1 | 16.05 | 41.3 | 19.5 | 14.0 |
| sp\|Q8N3R9\|PALS1\_HUMAN | Q8N3R9 | PALS1\_HUMAN | PALS1 | Protein PALS1 | 675 | 74.250 | 1.053371e+03 | 9.826414e+04 | 98264.14 |  |  |  | 1.63 |  |  |  |
| sp\|Q8IZ21\|PHAR4\_HUMAN | Q8IZ21 | PHAR4\_HUMAN | PHACTR4 | Phosphatase and actin regulator 4 | 702 | 78.211 | 1.876800e+03 | 1.844178e+05 | 98061.234 | 86356.61 |  |  | 12.11 | 39.2 |  |  |
| sp\|O14578\|CTRO\_HUMAN | O14578 | CTRO\_HUMAN | CIT | Citron Rho-interacting kinase | 2027 | 231.431 | 5.928837e+02 | 1.723885e+05 | 97442.39 | 74946.086 |  |  | 3.85 | 39.1 |  |  |
| sp\|P02675\|FIBB\_HUMAN | P02675 | FIBB\_HUMAN | FGB | Fibrinogen beta chain | 491 | 55.928 | 1.678502e+05 | 1.179420e+07 | 96683.85 | 5729465.5 | 4617274.0 | 1350773.6 | 7.54 | 65.0 | 49.9 | 26.1 |
| sp\|Q5T0F9\|C2D1B\_HUMAN | Q5T0F9 | C2D1B\_HUMAN | CC2D1B | Coiled-coil and C2 domain-containing protein 1B | 858 | 94.224 | 8.762679e+02 | 1.037327e+05 | 96153.17 | 7579.513 |  |  | 5.59 | 35.8 |  |  |
| sp\|Q96DM3\|RMC1\_HUMAN | Q96DM3 | RMC1\_HUMAN | RMC1 | Regulator of MON1-CCZ1 complex | 657 | 74.975 | 2.648377e+03 | 2.494672e+05 | 95907.07 | 153560.14 |  |  | 11.11 | 37.1 |  |  |
| sp\|Q9BVL2\|NUP58\_HUMAN | Q9BVL2 | NUP58\_HUMAN | NUP58 | Nucleoporin p58/p45 | 599 | 60.897 | 3.525764e+03 | 2.697530e+05 | 95864.98 | 173888.06 |  |  | 5.18 | 19.4 |  |  |
| sp\|P61769\|B2MG\_HUMAN | P61769 | B2MG\_HUMAN | B2M | Beta-2-microglobulin | 119 | 13.715 | 1.160851e+05 | 2.000273e+06 | 95661.766 |  | 1840078.2 | 64533.43 | 16.81 |  | 22.7 | 16.8 |
| sp\|Q8NCE2\|MTMRE\_HUMAN | Q8NCE2 | MTMRE\_HUMAN | MTMR14 | Myotubularin-related protein 14 | 650 | 72.203 | 2.475885e+03 | 2.245964e+05 | 95151.836 | 129444.56 |  |  | 5.08 | 21.7 |  |  |
| sp\|P48553\|TPC10\_HUMAN | P48553 | TPC10\_HUMAN | TRAPPC10 | Trafficking protein particle complex subunit 10 | 1259 | 142.189 | 1.231377e+03 | 2.199755e+05 | 94919.39 | 125056.125 |  |  | 7.78 | 25.9 |  |  |
| sp\|Q8TBX8\|PI42C\_HUMAN | Q8TBX8 | PI42C\_HUMAN | PIP4K2C | Phosphatidylinositol 5-phosphate 4-kinase type-2 gamma | 421 | 47.300 | 1.071797e+04 | 6.369288e+05 | 94268.22 | 524946.0 | 17714.605 |  | 10.21 | 31.8 | 12.4 |  |
| sp\|Q96FN9\|DTD2\_HUMAN | Q96FN9 | DTD2\_HUMAN | DTD2 | D-aminoacyl-tRNA deacylase 2 | 168 | 18.660 | 3.908697e+04 | 9.163488e+05 | 94244.54 | 806545.2 | 15559.033 |  | 6.55 | 67.3 | 25.0 |  |
| sp\|P11441\|UBL4A\_HUMAN | P11441 | UBL4A\_HUMAN | UBL4A | Ubiquitin-like protein 4A | 157 | 17.777 | 1.209250e+05 | 2.700795e+06 | 94038.24 | 756979.8 | 1692667.0 | 157110.1 | 13.38 | 61.1 | 45.2 | 29.9 |
| sp\|Q6ZVM7\|TM1L2\_HUMAN | Q6ZVM7 | TM1L2\_HUMAN | TOM1L2 | TOM1-like protein 2 | 507 | 55.556 | 1.912471e+03 | 1.334883e+05 | 93817.44 |  | 39670.824 |  | 7.5 |  | 13.6 |  |
| sp\|O60885\|BRD4\_HUMAN | O60885 | BRD4\_HUMAN | BRD4 | Bromodomain-containing protein 4 | 1362 | 152.219 | 3.376054e+03 | 6.456473e+05 | 93772.664 | 356281.28 | 127200.83 | 68392.56 | 3.23 | 26.8 | 14.0 | 10.4 |
| sp\|P08575\|PTPRC\_HUMAN | P08575 | PTPRC\_HUMAN | PTPRC | Receptor-type tyrosine-protein phosphatase C | 1306 | 147.486 | 1.677238e+03 | 3.107868e+05 | 93728.65 |  | 101335.92 | 115722.266 | 3.68 |  | 15.2 | 9.3 |
| sp\|Q9H267\|VP33B\_HUMAN | Q9H267 | VP33B\_HUMAN | VPS33B | Vacuolar protein sorting-associated protein 33B | 617 | 70.585 | 5.814010e+03 | 5.155910e+05 | 93519.836 | 422071.16 |  |  | 18.48 | 41.8 |  |  |
| sp\|O14920\|IKKB\_HUMAN | O14920 | IKKB\_HUMAN | IKBKB | Inhibitor of nuclear factor kappa-B kinase subunit beta | 756 | 86.564 | 2.499605e+03 | 2.718478e+05 | 93434.03 | 178413.77 |  |  | 8.07 | 47.0 |  |  |
| sp\|Q8IWD4\|CC117\_HUMAN | Q8IWD4 | CC117\_HUMAN | CCDC117 | Coiled-coil domain-containing protein 117 | 279 | 30.541 | 1.927495e+04 | 7.395946e+05 | 93377.375 | 581217.44 | 27690.688 | 37309.066 | 9.68 | 37.6 | 16.1 | 15.8 |
| sp\|Q05469\|LIPS\_HUMAN | Q05469 | LIPS\_HUMAN | LIPE | Hormone-sensitive lipase | 1076 | 116.598 | 1.847787e+03 | 2.706825e+05 | 93058.31 | 177624.19 |  |  | 3.53 | 37.5 |  |  |
| sp\|Q15424\|SAFB1\_HUMAN | Q15424 | SAFB1\_HUMAN | SAFB | Scaffold attachment factor B1 | 915 | 102.642 | 2.434582e+03 | 3.139544e+05 | 92815.17 | 206460.36 | 14678.903 |  | 10.38 | 36.8 | 25.5 |  |
| sp\|Q9BTE1\|DCTN5\_HUMAN | Q9BTE1 | DCTN5\_HUMAN | DCTN5 | Dynactin subunit 5 | 182 | 20.127 | 4.103992e+04 | 1.037774e+06 | 92676.25 | 818279.5 | 126817.984 |  | 6.04 | 30.8 | 10.4 |  |
| sp\|P21964\|COMT\_HUMAN | P21964 | COMT\_HUMAN | COMT | Catechol O-methyltransferase | 271 | 30.037 | 1.988096e+04 | 7.502587e+05 | 91983.58 | 540126.0 | 118149.14 |  | 16.61 | 47.6 | 15.1 |  |
| sp\|A6NFX1\|MFS2B\_HUMAN | A6NFX1 | MFS2B\_HUMAN | MFSD2B | Sphingosine-1-phosphate transporter MFSD2B | 504 | 53.743 | 1.807360e+04 | 1.220348e+06 | 90684.83 | 121357.266 | 317489.5 | 690816.44 | 6.75 | 16.9 | 10.7 | 10.5 |
| sp\|Q99501\|GA2L1\_HUMAN | Q99501 | GA2L1\_HUMAN | GAS2L1 | GAS2-like protein 1 | 681 | 72.717 | 2.748080e+03 | 2.510629e+05 | 90336.93 | 160725.94 |  |  | 6.61 | 48.9 |  |  |
| sp\|Q92934\|BAD\_HUMAN | Q92934 | BAD\_HUMAN | BAD | Bcl2-associated agonist of cell death | 168 | 18.392 | 3.422950e+04 | 7.909456e+05 | 90306.05 | 185620.84 | 515018.72 |  | 14.29 | 50.0 | 45.2 |  |
| sp\|O15127\|SCAM2\_HUMAN | O15127 | SCAM2\_HUMAN | SCAMP2 | Secretory carrier-associated membrane protein 2 | 329 | 36.649 | 1.138600e+04 | 5.242646e+05 | 90148.41 | 114093.125 | 24673.775 | 295349.25 | 12.16 | 14.0 | 15.5 | 6.4 |
| sp\|P07320\|CRGD\_HUMAN | P07320 | CRGD\_HUMAN | CRYGD | Gamma-crystallin D | 174 | 20.738 | 1.213267e+04 | 3.161116e+05 | 90113.11 | 225998.53 |  |  | 11.49 | 40.8 |  |  |
| sp\|Q9NSA3\|CNBP1\_HUMAN | Q9NSA3 | CNBP1\_HUMAN | CTNNBIP1 | Beta-catenin-interacting protein 1 | 81 | 8.910 | 8.043104e+03 | 9.003649e+04 | 90036.49 |  |  |  | 18.52 |  |  |  |
| sp\|P06213\|INSR\_HUMAN | P06213 | INSR\_HUMAN | INSR | Insulin receptor | 1382 | 156.333 | 3.941293e+03 | 7.741169e+05 | 89529.516 |  | 404217.7 | 280369.7 | 1.59 |  | 22.4 | 17.9 |
| sp\|Q05397\|FAK1\_HUMAN | Q05397 | FAK1\_HUMAN | PTK2 | Focal adhesion kinase 1 | 1052 | 119.233 | 1.622549e+03 | 2.430588e+05 | 89077.18 | 153981.66 |  |  | 6.56 | 22.9 |  |  |
| sp\|Q5TEC6\|H37\_HUMAN | Q5TEC6 | H37\_HUMAN | H3-7 | Histone H3-7 | 136 | 14.960 | 4.719724e+03 | 8.870853e+04 | 88708.53 |  |  |  | 14.71 |  |  |  |
| sp\|Q9H900\|ZWILC\_HUMAN | Q9H900 | ZWILC\_HUMAN | ZWILCH | Protein zwilch homolog | 591 | 67.214 | 3.899871e+03 | 3.293268e+05 | 88378.77 | 240948.06 |  |  | 7.61 | 27.2 |  |  |
| sp\|P05814\|CASB\_HUMAN | P05814 | CASB\_HUMAN | CSN2 | Beta-casein | 226 | 24.860 | 2.826968e+03 | 8.829563e+04 | 88295.63 |  |  |  | 11.95 |  |  |  |
| sp\|P06732\|KCRM\_HUMAN | P06732 | KCRM\_HUMAN | CKM | Creatine kinase M-type | 381 | 43.101 | 3.803920e+04 | 2.059851e+06 | 88165.61 | 227918.89 | 941516.44 | 802250.25 | 2.36 | 46.7 | 31.2 | 24.1 |
| sp\|Q8NB15\|ZN511\_HUMAN | Q8NB15 | ZN511\_HUMAN | ZNF511 | Zinc finger protein 511 | 252 | 28.265 | 7.876397e+03 | 2.797008e+05 | 88090.22 | 154213.11 | 37397.484 |  | 15.48 | 38.1 | 55.2 |  |
| sp\|Q9Y262\|EIF3L\_HUMAN | Q9Y262 | EIF3L\_HUMAN | EIF3L | Eukaryotic translation initiation factor 3 subunit L | 564 | 66.727 | 1.504431e+03 | 1.261220e+05 | 87883.28 |  |  | 38238.74 | 9.22 |  |  | 6.9 |
| sp\|P36639\|8ODP\_HUMAN | P36639 | 8ODP\_HUMAN | NUDT1 | Oxidized purine nucleoside triphosphate hydrolase | 156 | 17.952 | 8.254502e+04 | 1.861748e+06 | 87354.96 | 1774393.0 |  |  | 17.31 | 37.8 |  |  |
| sp\|Q14574\|DSC3\_HUMAN | Q14574 | DSC3\_HUMAN | DSC3 | Desmocollin-3 | 896 | 99.969 | 3.513522e+03 | 4.412911e+05 | 87128.64 | 19067.637 | 98564.84 | 236530.0 | 7.7 | 22.2 | 10.0 | 7.7 |
| sp\|O43399\|TPD54\_HUMAN | O43399 | TPD54\_HUMAN | TPD52L2 | Tumor protein D54 | 206 | 22.238 | 2.066341e+04 | 5.773178e+05 | 87047.83 | 435728.8 | 0.0 | 54541.19 | 11.65 | 61.2 | 26.7 | 14.1 |
| sp\|Q9H974\|QTRT2\_HUMAN | Q9H974 | QTRT2\_HUMAN | QTRT2 | Queuine tRNA-ribosyltransferase accessory subunit 2 | 415 | 46.713 | 2.441771e+03 | 1.433045e+05 | 86970.555 | 56333.938 |  |  | 11.08 | 28.9 |  |  |
| sp\|P00738\|HPT\_HUMAN | P00738 | HPT\_HUMAN | HP | Haptoglobin | 406 | 45.205 | 2.559023e+04 | 1.453375e+06 | 86507.375 | 654088.8 | 528876.7 | 183902.4 | 5.67 | 40.9 | 34.2 | 25.9 |
| sp\|P08865\|RSSA\_HUMAN | P08865 | RSSA\_HUMAN | RPSA | Small ribosomal subunit protein uS2 | 295 | 32.854 | 9.188239e+03 | 3.792606e+05 | 86481.32 | 99315.78 | 114452.234 | 79011.24 | 15.93 | 41.0 | 29.8 | 17.3 |
| sp\|P00505\|AATM\_HUMAN | P00505 | AATM\_HUMAN | GOT2 | Aspartate aminotransferase, mitochondrial | 430 | 47.518 | 5.104501e+03 | 3.047394e+05 | 86327.34 | 104861.07 | 113550.97 |  | 8.37 | 45.1 | 24.0 |  |
| sp\|O75122\|CLAP2\_HUMAN | O75122 | CLAP2\_HUMAN | CLASP2 | CLIP-associating protein 2 | 1294 | 141.064 | 4.445853e+03 | 7.879315e+05 | 86321.37 | 280567.94 | 139339.05 | 281703.16 | 8.73 | 49.5 | 25.7 | 20.5 |
| sp\|Q6NW29\|RWDD4\_HUMAN | Q6NW29 | RWDD4\_HUMAN | RWDD4 | RWD domain-containing protein 4 | 188 | 21.251 | 1.003325e+04 | 2.678788e+05 | 86188.23 | 181690.55 |  |  | 10.11 | 17.6 |  |  |
| sp\|Q9P287\|BCCIP\_HUMAN | Q9P287 | BCCIP\_HUMAN | BCCIP | BRCA2 and CDKN1A-interacting protein | 314 | 35.979 | 4.815251e+03 | 2.176633e+05 | 86161.77 | 131501.5 |  |  | 20.06 | 42.0 |  |  |
| sp\|P35222\|CTNB1\_HUMAN | P35222 | CTNB1\_HUMAN | CTNNB1 | Catenin beta-1 | 781 | 85.497 | 2.322307e+03 | 2.494524e+05 | 86122.85 | 19118.951 | 144210.58 |  | 5.63 | 29.8 | 12.8 |  |
| sp\|Q8IZ73\|RUSD2\_HUMAN | Q8IZ73 | RUSD2\_HUMAN | RPUSD2 | Pseudouridylate synthase RPUSD2 | 545 | 59.950 | 1.136102e+03 | 8.557039e+04 | 85570.39 |  |  |  | 7.16 |  |  |  |
| sp\|Q9BVQ7\|AFG2B\_HUMAN | Q9BVQ7 | AFG2B\_HUMAN | AFG2B | ATPase family gene 2 protein homolog B | 753 | 82.830 | 8.203176e+02 | 8.536638e+04 | 85366.375 |  |  |  | 1.59 |  |  |  |
| sp\|Q9BRT3\|MIEN1\_HUMAN | Q9BRT3 | MIEN1\_HUMAN | MIEN1 | Migration and invasion enhancer 1 | 115 | 12.403 | 1.796385e+05 | 2.799260e+06 | 85290.7 | 2236919.5 | 303636.44 | 173413.31 | 18.26 | 63.5 | 47.8 | 44.3 |
| sp\|Q13469\|NFAC2\_HUMAN | Q13469 | NFAC2\_HUMAN | NFATC2 | Nuclear factor of activated T-cells, cytoplasmic 2 | 925 | 101.750 | 6.609580e+02 | 8.449391e+04 | 84493.91 |  |  |  | 8.76 |  |  |  |
| sp\|Q9H7C9\|AAMDC\_HUMAN | Q9H7C9 | AAMDC\_HUMAN | AAMDC | Mth938 domain-containing protein | 122 | 13.332 | 9.197122e+04 | 1.540510e+06 | 84351.55 | 1240238.6 | 150951.56 | 64967.867 | 13.11 | 68.9 | 58.2 | 39.3 |
| sp\|Q9BRR6\|ADPGK\_HUMAN | Q9BRR6 | ADPGK\_HUMAN | ADPGK | ADP-dependent glucokinase | 497 | 54.089 | 7.529355e+03 | 5.116629e+05 | 84041.53 | 151546.2 | 249827.53 | 26247.594 | 5.84 | 28.8 | 13.5 | 5.0 |
| sp\|Q9Y277\|VDAC3\_HUMAN | Q9Y277 | VDAC3\_HUMAN | VDAC3 | Voltage-dependent anion-selective channel protein 3 | 283 | 30.659 | 1.005034e+04 | 3.871294e+05 | 83911.69 | 44774.668 | 258443.0 |  | 19.08 | 16.6 | 13.1 |  |
| sp\|Q9NQT8\|KI13B\_HUMAN | Q9NQT8 | KI13B\_HUMAN | KIF13B | Kinesin-like protein KIF13B | 1826 | 202.789 | 2.064146e+03 | 5.258984e+05 | 83332.79 | 322734.06 | 119831.6 |  | 3.72 | 35.1 | 13.4 |  |
| sp\|Q9NRX4\|PHP14\_HUMAN | Q9NRX4 | PHP14\_HUMAN | PHPT1 | 14 kDa phosphohistidine phosphatase | 125 | 13.833 | 2.102590e+05 | 3.654165e+06 | 83052.48 | 3290575.0 | 280537.3 |  | 19.2 | 88.8 | 46.4 |  |
| sp\|Q13613\|MTMR1\_HUMAN | Q13613 | MTMR1\_HUMAN | MTMR1 | Myotubularin-related protein 1 | 665 | 74.678 | 2.698394e+03 | 2.531717e+05 | 82905.766 | 170265.92 |  |  | 14.74 | 42.7 |  |  |
| sp\|O75436\|VP26A\_HUMAN | O75436 | VP26A\_HUMAN | VPS26A | Vacuolar protein sorting-associated protein 26A | 327 | 38.170 | 2.368427e+03 | 1.135793e+05 | 82581.58 |  | 30997.758 |  | 9.17 |  | 27.5 |  |
| sp\|Q86SJ6\|DSG4\_HUMAN | Q86SJ6 | DSG4\_HUMAN | DSG4 | Desmoglein-4 | 1040 | 114.400 | 5.675781e+02 | 8.157720e+04 | 81577.2 |  |  |  | 5.29 |  |  |  |
| sp\|Q9H1E1\|RNAS7\_HUMAN | Q9H1E1 | RNAS7\_HUMAN | RNASE7 | Ribonuclease 7 | 156 | 17.419 | 5.517299e+03 | 1.207444e+05 | 81465.78 | 39278.598 |  |  | 10.26 | 53.2 |  |  |
| sp\|Q8N3C0\|ASCC3\_HUMAN | Q8N3C0 | ASCC3\_HUMAN | ASCC3 | Activating signal cointegrator 1 complex subunit 3 | 2202 | 251.460 | 9.257635e+02 | 2.924732e+05 | 81417.22 | 211056.0 |  |  | 1.45 | 32.2 |  |  |
| sp\|Q9Y4K3\|TRAF6\_HUMAN | Q9Y4K3 | TRAF6\_HUMAN | TRAF6 | TNF receptor-associated factor 6 | 522 | 57.420 | 1.127407e+03 | 8.133191e+04 | 81331.914 |  |  |  | 6.51 |  |  |  |
| sp\|Q7L2H7\|EIF3M\_HUMAN | Q7L2H7 | EIF3M\_HUMAN | EIF3M | Eukaryotic translation initiation factor 3 subunit M | 374 | 42.503 | 1.664331e+04 | 8.887433e+05 | 81294.19 | 799796.1 |  | 7652.9746 | 8.82 | 42.8 |  | 8.0 |
| sp\|O95671\|ASML\_HUMAN | O95671 | ASML\_HUMAN | ASMTL | Probable bifunctional dTTP/UTP pyrophosphatase/methyltransferase protein | 621 | 68.857 | 3.779424e+03 | 3.269572e+05 | 81258.59 | 245698.6 |  |  | 15.94 | 43.8 |  |  |
| sp\|O43852\|CALU\_HUMAN | O43852 | CALU\_HUMAN | CALU | Calumenin | 315 | 37.107 | 5.744602e+04 | 2.678138e+06 | 81084.91 | 357186.06 | 2098674.2 | 141192.81 | 10.48 | 22.2 | 29.2 | 20.0 |
| sp\|P27695\|APEX1\_HUMAN | P27695 | APEX1\_HUMAN | APEX1 | DNA-(apurinic or apyrimidinic site) endonuclease | 318 | 35.555 | 3.164529e+03 | 1.413601e+05 | 80945.484 | 60414.637 |  |  | 16.98 | 37.7 |  |  |
| sp\|Q6ZNW5\|GDPP1\_HUMAN | Q6ZNW5 | GDPP1\_HUMAN | GDPGP1 | GDP-D-glucose phosphorylase 1 | 385 | 42.362 | 2.402497e+03 | 1.278664e+05 | 80887.37 | 46979.043 |  |  | 7.79 | 49.4 |  |  |
| sp\|O15234\|CASC3\_HUMAN | O15234 | CASC3\_HUMAN | CASC3 | Protein CASC3 | 703 | 76.278 | 2.776214e+03 | 2.660538e+05 | 80801.19 | 185252.58 |  |  | 8.11 | 28.4 |  |  |
| sp\|Q9H920\|RN121\_HUMAN | Q9H920 | RN121\_HUMAN | RNF121 | E3 ubiquitin ligase RNF121 | 327 | 37.882 | 3.912695e+03 | 1.862199e+05 | 80106.45 | 16061.7 | 23585.037 | 66466.7 | 7.95 | 22.0 | 13.8 | 13.5 |
| sp\|Q9Y2G3\|AT11B\_HUMAN | Q9Y2G3 | AT11B\_HUMAN | ATP11B | Phospholipid-transporting ATPase IF | 1177 | 134.190 | 1.327894e+04 | 2.238725e+06 | 79967.74 | 86120.88 | 802850.25 | 1269786.2 | 8.07 | 18.8 | 20.2 | 16.1 |
| sp\|Q9UDY2\|ZO2\_HUMAN | Q9UDY2 | ZO2\_HUMAN | TJP2 | Tight junction protein ZO-2 | 1190 | 133.958 | 1.540336e+03 | 2.592395e+05 | 79934.125 | 125854.01 | 53451.395 |  | 4.2 | 39.4 | 22.1 |  |
| sp\|O60826\|CCD22\_HUMAN | O60826 | CCD22\_HUMAN | CCDC22 | Coiled-coil domain-containing protein 22 | 627 | 70.756 | 2.239130e+03 | 1.990489e+05 | 79798.46 | 119250.42 |  |  | 8.61 | 41.0 |  |  |
| sp\|Q6BDS2\|BLT3A\_HUMAN | Q6BDS2 | BLT3A\_HUMAN | BLTP3A | Bridge-like lipid transfer protein family member 3A | 1440 | 159.485 | 1.791906e+03 | 3.590478e+05 | 79476.09 | 169314.3 | 93928.58 | 16328.861 | 6.04 | 27.3 | 16.1 | 10.3 |
| sp\|P53367\|ARFP1\_HUMAN | P53367 | ARFP1\_HUMAN | ARFIP1 | Arfaptin-1 | 373 | 41.738 | 2.866066e+04 | 1.502917e+06 | 79472.56 | 90149.805 | 1333294.5 |  | 10.46 | 45.3 | 20.6 |  |
| sp\|Q07817\|B2CL1\_HUMAN | Q07817 | B2CL1\_HUMAN | BCL2L1 | Bcl-2-like protein 1 | 233 | 26.049 | 2.942106e+04 | 9.628676e+05 | 79225.836 |  | 585571.4 | 298070.38 | 10.3 |  | 21.9 | 11.2 |
| sp\|Q9NSY0\|NRBP2\_HUMAN | Q9NSY0 | NRBP2\_HUMAN | NRBP2 | Nuclear receptor-binding protein 2 | 501 | 57.803 | 2.993808e+03 | 2.174160e+05 | 79163.98 | 138252.0 |  |  | 7.58 | 24.6 |  |  |
| sp\|Q96C19\|EFHD2\_HUMAN | Q96C19 | EFHD2\_HUMAN | EFHD2 | EF-hand domain-containing protein D2 | 240 | 26.697 | 1.899437e+04 | 6.370955e+05 | 79070.516 | 414659.0 | 83048.01 | 60318.0 | 15.0 | 79.6 | 18.8 | 22.5 |
| sp\|Q14156\|EFR3A\_HUMAN | Q14156 | EFR3A\_HUMAN | EFR3A | Protein EFR3 homolog A | 821 | 92.924 | 3.828656e+03 | 4.469834e+05 | 79063.586 | 39654.703 | 106049.36 | 222215.77 | 7.8 | 38.7 | 25.1 | 16.1 |
| sp\|Q9H773\|DCTP1\_HUMAN | Q9H773 | DCTP1\_HUMAN | DCTPP1 | dCTP pyrophosphatase 1 | 170 | 18.681 | 8.673229e+03 | 2.035626e+05 | 78704.6 | 124858.04 |  |  | 13.53 | 56.5 |  |  |
| sp\|Q6IAN0\|DRS7B\_HUMAN | Q6IAN0 | DRS7B\_HUMAN | DHRS7B | Dehydrogenase/reductase SDR family member 7B | 325 | 35.119 | 2.026203e+04 | 8.940098e+05 | 78333.53 | 406309.0 | 241890.69 | 167476.62 | 7.69 | 41.5 | 33.2 | 28.3 |
| sp\|Q7L4E1\|MIGA2\_HUMAN | Q7L4E1 | MIGA2\_HUMAN | MIGA2 | Mitoguardin 2 | 593 | 65.531 | 2.672592e+03 | 2.200375e+05 | 78109.09 | 58535.715 | 83392.69 |  | 4.72 | 17.9 | 21.4 |  |
| sp\|Q9UHX1\|PUF60\_HUMAN | Q9UHX1 | PUF60\_HUMAN | PUF60 | Poly(U)-binding-splicing factor PUF60 | 559 | 59.875 | 2.038923e+03 | 1.533781e+05 | 77574.17 | 75803.95 |  |  | 5.55 | 31.8 |  |  |
| sp\|Q9UGK8\|SRGEF\_HUMAN | Q9UGK8 | SRGEF\_HUMAN | SERGEF | Secretion-regulating guanine nucleotide exchange factor | 458 | 48.981 | 2.744323e+03 | 1.688807e+05 | 77425.766 | 91454.91 |  |  | 8.95 | 40.0 |  |  |
| sp\|P04844\|RPN2\_HUMAN | P04844 | RPN2\_HUMAN | RPN2 | Dolichyl-diphosphooligosaccharide--protein glycosyltransferase subunit 2 | 631 | 69.284 | 1.918950e+04 | 1.670374e+06 | 77291.06 | 166809.56 | 1133035.8 | 293237.4 | 12.52 | 26.1 | 15.1 | 9.4 |
| sp\|Q96FJ2\|DYL2\_HUMAN | Q96FJ2 | DYL2\_HUMAN | DYNLL2 | Dynein light chain 2, cytoplasmic | 89 | 10.350 | 7.577813e+04 | 9.853746e+05 | 76814.445 | 856512.8 |  | 52047.348 | 25.84 | 74.2 |  | 20.2 |
| sp\|Q9Y478\|AAKB1\_HUMAN | Q9Y478 | AAKB1\_HUMAN | PRKAB1 | 5'-AMP-activated protein kinase subunit beta-1 | 270 | 30.382 | 1.124805e+04 | 4.293493e+05 | 76432.13 | 290987.5 | 61929.633 |  | 16.3 | 33.7 | 17.4 |  |
| sp\|P35712\|SOX6\_HUMAN | P35712 | SOX6\_HUMAN | SOX6 | Transcription factor SOX-6 | 828 | 91.921 | 1.884188e+03 | 2.175986e+05 | 76301.98 | 141296.67 |  |  | 6.52 | 39.6 |  |  |
| sp\|Q9UJX4\|APC5\_HUMAN | Q9UJX4 | APC5\_HUMAN | ANAPC5 | Anaphase-promoting complex subunit 5 | 755 | 85.077 | 2.218082e+04 | 2.370866e+06 | 76228.35 | 2294637.5 |  |  | 11.13 | 28.2 |  |  |
| sp\|A8K2U0\|A2ML1\_HUMAN | A8K2U0 | A2ML1\_HUMAN | A2ML1 | Alpha-2-macroglobulin-like protein 1 | 1454 | 161.107 | 4.416125e+03 | 8.938669e+05 | 75715.5 | 44724.816 | 506115.62 | 267311.0 | 2.75 | 23.3 | 17.7 | 15.3 |
| sp\|Q96EC8\|YIPF6\_HUMAN | Q96EC8 | YIPF6\_HUMAN | YIPF6 | Protein YIPF6 | 236 | 26.256 | 1.457767e+04 | 4.808767e+05 | 75485.516 | 217578.97 | 61798.145 | 126014.04 | 8.47 | 8.9 | 22.9 | 5.1 |
| sp\|P55884\|EIF3B\_HUMAN | P55884 | EIF3B\_HUMAN | EIF3B | Eukaryotic translation initiation factor 3 subunit B | 814 | 92.482 | 1.891058e+03 | 2.197249e+05 | 74886.04 | 126448.586 | 12851.839 | 5538.4585 | 6.14 | 30.8 | 12.4 | 9.5 |
| sp\|P39656\|OST48\_HUMAN | P39656 | OST48\_HUMAN | DDOST | Dolichyl-diphosphooligosaccharide--protein glycosyltransferase 48 kDa subunit | 456 | 50.801 | 1.824758e+04 | 1.164648e+06 | 74519.67 |  | 951016.6 | 139111.92 | 6.8 |  | 29.6 | 7.7 |
| sp\|Q13546\|RIPK1\_HUMAN | Q13546 | RIPK1\_HUMAN | RIPK1 | Receptor-interacting serine/threonine-protein kinase 1 | 671 | 75.931 | 5.032463e+03 | 4.800836e+05 | 74427.86 | 405655.72 |  |  | 7.9 | 36.4 |  |  |
| sp\|O00189\|AP4M1\_HUMAN | O00189 | AP4M1\_HUMAN | AP4M1 | AP-4 complex subunit mu-1 | 453 | 49.977 | 2.991350e+03 | 1.878255e+05 | 74420.5 | 113405.03 |  |  | 6.4 | 48.1 |  |  |
| sp\|P02679\|FIBG\_HUMAN | P02679 | FIBG\_HUMAN | FGG | Fibrinogen gamma chain | 453 | 51.512 | 3.291483e+05 | 2.130184e+07 | 74401.99 | 17474846.0 | 3219505.8 | 533087.3 | 11.7 | 58.5 | 46.6 | 21.6 |
| sp\|Q99828\|CIB1\_HUMAN | Q99828 | CIB1\_HUMAN | CIB1 | Calcium and integrin-binding protein 1 | 191 | 21.703 | 9.089802e+04 | 2.478514e+06 | 73516.45 | 573537.3 | 1000465.8 | 830994.4 | 14.66 | 50.8 | 39.3 | 26.7 |
| sp\|Q15828\|CYTM\_HUMAN | Q15828 | CYTM\_HUMAN | CST6 | Cystatin-M | 149 | 16.511 | 2.516837e+04 | 5.220903e+05 | 73467.78 |  | 187294.0 | 261328.48 | 27.52 |  | 22.8 | 14.1 |
| sp\|Q96LB3\|IFT74\_HUMAN | Q96LB3 | IFT74\_HUMAN | IFT74 | Intraflagellar transport protein 74 homolog | 600 | 66.000 | 8.853422e+02 | 7.341288e+04 | 73412.88 |  |  |  | 1.33 |  |  |  |
| sp\|O14531\|DPYL4\_HUMAN | O14531 | DPYL4\_HUMAN | DPYSL4 | Dihydropyrimidinase-related protein 4 | 572 | 62.920 | 9.283302e+02 | 7.338518e+04 | 73385.18 |  |  |  | 2.8 |  |  |  |
| sp\|Q92851\|CASPA\_HUMAN | Q92851 | CASPA\_HUMAN | CASP10 | Caspase-10 | 521 | 58.951 | 1.773921e+03 | 1.313841e+05 | 73059.45 | 58324.61 |  |  | 11.71 | 44.7 |  |  |
| sp\|P05387\|RLA2\_HUMAN | P05387 | RLA2\_HUMAN | RPLP2 | Large ribosomal subunit protein P2 | 115 | 11.665 | 5.258861e+04 | 7.707145e+05 | 73055.266 | 567651.0 | 130008.266 |  | 46.09 | 64.3 | 79.1 |  |
| sp\|P51790\|CLCN3\_HUMAN | P51790 | CLCN3\_HUMAN | CLCN3 | H(+)/Cl(-) exchange transporter 3 | 818 | 90.966 | 1.168595e+04 | 1.335550e+06 | 73048.23 | 169173.6 | 210678.5 | 882649.9 | 4.03 | 17.7 | 13.3 | 17.5 |
| sp\|Q96FZ7\|CHMP6\_HUMAN | Q96FZ7 | CHMP6\_HUMAN | CHMP6 | Charged multivesicular body protein 6 | 201 | 23.485 | 1.221272e+04 | 3.603464e+05 | 72719.38 | 287627.0 |  |  | 6.47 | 41.8 |  |  |
| sp\|P01009\|A1AT\_HUMAN | P01009 | A1AT\_HUMAN | SERPINA1 | Alpha-1-antitrypsin | 418 | 46.737 | 3.077612e+04 | 1.807140e+06 | 72709.36 | 906701.44 | 827729.7 |  | 17.22 | 58.1 | 43.1 |  |
| sp\|Q8N9B5\|JMY\_HUMAN | Q8N9B5 | JMY\_HUMAN | JMY | Junction-mediating and -regulatory protein | 988 | 108.680 | 5.321273e+02 | 7.265781e+04 | 72657.81 |  |  |  | 2.73 |  |  |  |
| sp\|P08236\|BGLR\_HUMAN | P08236 | BGLR\_HUMAN | GUSB | Beta-glucuronidase | 651 | 74.732 | 1.244324e+03 | 1.168308e+05 | 72388.16 | 44442.605 |  |  | 5.07 | 22.3 |  |  |
| sp\|Q92609\|TBCD5\_HUMAN | Q92609 | TBCD5\_HUMAN | TBC1D5 | TBC1 domain family member 5 | 795 | 89.004 | 5.247252e+03 | 5.867574e+05 | 71815.45 | 514942.0 |  |  | 8.43 | 23.8 |  |  |
| sp\|P61764\|STXB1\_HUMAN | P61764 | STXB1\_HUMAN | STXBP1 | Syntaxin-binding protein 1 | 594 | 67.569 | 2.905440e+03 | 2.466474e+05 | 71280.39 | 175367.05 | 0.0 |  | 6.4 | 37.4 | 23.2 |  |
| sp\|O60502\|OGA\_HUMAN | O60502 | OGA\_HUMAN | OGA | Protein O-GlcNAcase | 916 | 100.760 | 5.629153e+02 | 7.126041e+04 | 71260.414 |  |  |  | 2.73 |  |  |  |
| sp\|Q96IG2\|FXL20\_HUMAN | Q96IG2 | FXL20\_HUMAN | FBXL20 | F-box/LRR-repeat protein 20 | 436 | 48.423 | 4.720205e+03 | 2.871638e+05 | 71175.28 | 215988.53 |  |  | 7.57 | 22.7 |  |  |
| sp\|Q8IYA8\|IHO1\_HUMAN | Q8IYA8 | IHO1\_HUMAN | IHO1 | Interactor of HORMAD1 protein 1 | 594 | 65.340 | 8.654031e+02 | 7.104193e+04 | 71041.93 |  |  |  | 2.53 |  |  |  |
| sp\|P35637\|FUS\_HUMAN | P35637 | FUS\_HUMAN | FUS | RNA-binding protein FUS | 526 | 53.426 | 4.150707e+03 | 2.786069e+05 | 70877.31 | 188368.23 |  | 19361.396 | 4.37 | 37.3 |  | 23.4 |
| sp\|Q66LE6\|2ABD\_HUMAN | Q66LE6 | 2ABD\_HUMAN | PPP2R2D | Serine/threonine-protein phosphatase 2A 55 kDa regulatory subunit B delta isoform | 453 | 52.042 | 3.092588e+03 | 2.022056e+05 | 70665.63 | 131540.0 |  |  | 7.95 | 26.3 |  |  |
| sp\|P30044\|PRDX5\_HUMAN | P30044 | PRDX5\_HUMAN | PRDX5 | Peroxiredoxin-5, mitochondrial | 214 | 22.086 | 1.107390e+05 | 3.072804e+06 | 70636.22 | 2065662.5 | 838475.0 | 98029.86 | 14.49 | 86.9 | 40.7 | 19.2 |
| sp\|Q9ULQ1\|TPC1\_HUMAN | Q9ULQ1 | TPC1\_HUMAN | TPCN1 | Two pore channel protein 1 | 816 | 94.147 | 1.991797e+03 | 2.355965e+05 | 70302.3 | 28371.834 | 40177.297 | 96745.06 | 2.08 | 9.7 | 9.9 | 9.6 |
| sp\|P41440\|S19A1\_HUMAN | P41440 | S19A1\_HUMAN | SLC19A1 | Reduced folate transporter | 591 | 64.868 | 5.462115e+04 | 4.451522e+06 | 70216.26 |  | 788603.4 | 3592702.2 | 8.46 |  | 12.0 | 12.4 |
| sp\|B1AK53\|ESPN\_HUMAN | B1AK53 | ESPN\_HUMAN | ESPN | Espin | 854 | 91.733 | 6.535563e+02 | 7.532267e+04 | 70116.53 |  | 5206.1436 |  | 4.45 |  | 22.1 |  |
| sp\|Q9Y2H2\|SAC2\_HUMAN | Q9Y2H2 | SAC2\_HUMAN | INPP5F | Phosphatidylinositide phosphatase SAC2 | 1132 | 128.407 | 5.707384e+02 | 9.207526e+04 | 69993.984 | 22081.281 |  |  | 5.48 | 40.6 |  |  |
| sp\|Q7L9B9\|EEPD1\_HUMAN | Q7L9B9 | EEPD1\_HUMAN | EEPD1 | Endonuclease/exonuclease/phosphatase family domain-containing protein 1 | 569 | 62.403 | 9.578836e+03 | 7.509921e+05 | 69936.16 | 142337.84 | 136821.28 | 401896.84 | 11.6 | 38.0 | 33.0 | 14.4 |
| sp\|Q99836\|MYD88\_HUMAN | Q99836 | MYD88\_HUMAN | MYD88 | Myeloid differentiation primary response protein MyD88 | 296 | 33.233 | 1.028772e+04 | 4.295424e+05 | 69660.234 | 252497.39 | 84438.67 | 22946.06 | 8.11 | 44.3 | 22.6 | 8.1 |
| sp\|Q5W111\|SPRY7\_HUMAN | Q5W111 | SPRY7\_HUMAN | SPRYD7 | SPRY domain-containing protein 7 | 196 | 21.666 | 2.447854e+04 | 6.663178e+05 | 69426.586 | 235955.75 | 192467.02 | 168468.4 | 11.22 | 54.6 | 54.6 | 31.1 |
| sp\|P61923\|COPZ1\_HUMAN | P61923 | COPZ1\_HUMAN | COPZ1 | Coatomer subunit zeta-1 | 177 | 20.198 | 2.935455e+04 | 7.449052e+05 | 69164.2 | 387826.28 | 263823.97 | 24090.756 | 10.73 | 48.6 | 27.1 | 6.2 |
| sp\|P85299\|PRR5\_HUMAN | P85299 | PRR5\_HUMAN | PRR5 | Proline-rich protein 5 | 388 | 42.753 | 3.515504e+03 | 1.888301e+05 | 69057.29 | 99964.7 | 19808.16 |  | 12.37 | 41.5 | 17.8 |  |
| sp\|Q05823\|RN5A\_HUMAN | Q05823 | RN5A\_HUMAN | RNASEL | 2-5A-dependent ribonuclease | 741 | 83.533 | 2.779047e+03 | 2.916562e+05 | 68835.3 | 217462.62 | 5358.2563 |  | 7.42 | 34.0 | 13.8 |  |
| sp\|P48163\|MAOX\_HUMAN | P48163 | MAOX\_HUMAN | ME1 | NADP-dependent malic enzyme | 572 | 64.150 | 3.427462e+03 | 2.762399e+05 | 68582.69 |  |  | 207657.22 | 8.74 |  |  | 10.1 |
| sp\|O95208\|EPN2\_HUMAN | O95208 | EPN2\_HUMAN | EPN2 | Epsin-2 | 641 | 68.482 | 7.283047e+03 | 6.266236e+05 | 68266.99 | 429949.0 | 128407.58 |  | 9.83 | 44.1 | 20.0 |  |
| sp\|O60437\|PEPL\_HUMAN | O60437 | PEPL\_HUMAN | PPL | Periplakin | 1756 | 204.747 | 1.599843e+04 | 4.115400e+06 | 68117.34 |  | 3711852.2 | 335430.94 | 1.14 |  | 31.8 | 17.3 |
| sp\|Q92834\|RPGR\_HUMAN | Q92834 | RPGR\_HUMAN | RPGR | X-linked retinitis pigmentosa GTPase regulator | 1020 | 113.387 | 1.313064e+03 | 1.870537e+05 | 68062.516 | 118991.21 |  |  | 3.92 | 20.7 |  |  |
| sp\|Q96QR8\|PURB\_HUMAN | Q96QR8 | PURB\_HUMAN | PURB | Transcriptional activator protein Pur-beta | 312 | 33.241 | 1.877644e+04 | 7.841598e+05 | 67952.04 | 548255.6 | 97101.375 | 70850.805 | 4.17 | 52.6 | 19.6 | 23.7 |
| sp\|P55735\|SEC13\_HUMAN | P55735 | SEC13\_HUMAN | SEC13 | Protein SEC13 homolog | 322 | 35.541 | 8.414746e+03 | 3.757403e+05 | 67355.64 | 253454.19 | 54930.504 |  | 12.42 | 23.3 | 15.2 |  |
| sp\|Q6E0U4\|DMKN\_HUMAN | Q6E0U4 | DMKN\_HUMAN | DMKN | Dermokine | 476 | 47.082 | 2.490711e+03 | 1.473314e+05 | 67090.375 |  | 80241.06 |  | 3.57 |  | 21.2 |  |
| sp\|Q9Y3R4\|NEUR2\_HUMAN | Q9Y3R4 | NEUR2\_HUMAN | NEU2 | Sialidase-2 | 380 | 42.254 | 2.898931e+03 | 1.538944e+05 | 66941.516 |  | 20814.4 | 66138.52 | 8.16 |  | 15.5 | 11.1 |
| sp\|Q9UM00\|TMCO1\_HUMAN | Q9UM00 | TMCO1\_HUMAN | TMCO1 | Calcium load-activated calcium channel | 239 | 27.079 | 1.462843e+04 | 4.976769e+05 | 66769.766 |  | 102909.66 | 327997.5 | 10.04 |  | 36.0 | 31.0 |
| sp\|P57081\|WDR4\_HUMAN | P57081 | WDR4\_HUMAN | WDR4 | tRNA (guanine-N(7)-)-methyltransferase non-catalytic subunit WDR4 | 412 | 45.490 | 2.400319e+03 | 1.371835e+05 | 66759.97 | 70423.56 |  |  | 7.77 | 30.8 |  |  |
| sp\|P19838\|NFKB1\_HUMAN | P19838 | NFKB1\_HUMAN | NFKB1 | Nuclear factor NF-kappa-B p105 subunit | 968 | 105.356 | 2.785557e+03 | 3.687131e+05 | 66554.14 | 296089.0 | 6069.9395 |  | 3.31 | 30.3 | 12.9 |  |
| sp\|Q9UHD2\|TBK1\_HUMAN | Q9UHD2 | TBK1\_HUMAN | TBK1 | Serine/threonine-protein kinase TBK1 | 729 | 83.642 | 1.287695e+03 | 1.353177e+05 | 66452.59 | 68865.125 |  |  | 6.04 | 24.7 |  |  |
| sp\|Q96FQ6\|S10AG\_HUMAN | Q96FQ6 | S10AG\_HUMAN | S100A16 | Protein S100-A16 | 103 | 11.801 | 5.432204e+04 | 8.054008e+05 | 66118.234 | 75573.8 | 257731.6 | 405977.12 | 11.65 | 43.7 | 35.0 | 35.0 |
| sp\|Q8IW75\|SPA12\_HUMAN | Q8IW75 | SPA12\_HUMAN | SERPINA12 | Serpin A12 | 414 | 47.175 | 1.119433e+04 | 6.634791e+05 | 66061.7 | 127429.11 | 236807.52 | 233180.77 | 9.9 | 48.1 | 31.4 | 20.0 |
| sp\|Q9BRX5\|PSF3\_HUMAN | Q9BRX5 | PSF3\_HUMAN | GINS3 | DNA replication complex GINS protein PSF3 | 216 | 24.535 | 1.018653e+04 | 3.139998e+05 | 65933.34 | 230365.4 | 17701.098 |  | 23.15 | 51.9 | 20.8 |  |
| sp\|Q9NYG2\|ZDHC3\_HUMAN | Q9NYG2 | ZDHC3\_HUMAN | ZDHHC3 | Palmitoyltransferase ZDHHC3 | 299 | 34.170 | 2.535901e+04 | 1.088665e+06 | 65757.21 | 9968.094 | 225527.98 | 787412.2 | 9.03 | 40.1 | 24.7 | 13.0 |
| sp\|Q15628\|TRADD\_HUMAN | Q15628 | TRADD\_HUMAN | TRADD | Tumor necrosis factor receptor type 1-associated DEATH domain protein | 312 | 34.247 | 6.855069e+03 | 2.949521e+05 | 65639.96 | 158778.16 | 70534.016 |  | 15.06 | 42.3 | 28.8 |  |
| sp\|Q8WYR4\|RSPH1\_HUMAN | Q8WYR4 | RSPH1\_HUMAN | RSPH1 | Radial spoke head 1 homolog | 309 | 33.990 | 1.533537e+03 | 6.548812e+04 | 65488.125 |  |  |  | 9.39 |  |  |  |
| sp\|Q8TCF1\|ZFAN1\_HUMAN | Q8TCF1 | ZFAN1\_HUMAN | ZFAND1 | AN1-type zinc finger protein 1 | 268 | 30.787 | 1.844300e+04 | 7.133720e+05 | 64587.81 | 648784.2 |  |  | 7.84 | 46.3 |  |  |
| sp\|O95562\|SFT2B\_HUMAN | O95562 | SFT2B\_HUMAN | SFT2D2 | Vesicle transport protein SFT2B | 160 | 17.779 | 1.332029e+05 | 2.975350e+06 | 64548.582 | 700906.0 | 705033.25 | 1504861.9 | 6.25 | 24.4 | 11.9 | 6.9 |
| sp\|Q15005\|SPCS2\_HUMAN | Q15005 | SPCS2\_HUMAN | SPCS2 | Signal peptidase complex subunit 2 | 226 | 25.003 | 4.643713e+03 | 1.458729e+05 | 64522.62 |  | 81350.305 |  | 8.41 |  | 35.4 |  |
| sp\|P14866\|HNRPL\_HUMAN | P14866 | HNRPL\_HUMAN | HNRNPL | Heterogeneous nuclear ribonucleoprotein L | 589 | 64.133 | 4.459029e+03 | 3.592849e+05 | 64465.03 | 290072.72 | 4747.1685 |  | 4.92 | 35.3 | 34.6 |  |
| sp\|P05026\|AT1B1\_HUMAN | P05026 | AT1B1\_HUMAN | ATP1B1 | Sodium/potassium-transporting ATPase subunit beta-1 | 303 | 35.061 | 1.020228e+04 | 4.494059e+05 | 64356.484 |  | 176981.9 | 208067.52 | 8.25 |  | 16.5 | 13.2 |
| sp\|Q96AE4\|FUBP1\_HUMAN | Q96AE4 | FUBP1\_HUMAN | FUBP1 | Far upstream element-binding protein 1 | 644 | 67.560 | 2.853919e+03 | 2.422414e+05 | 64222.055 | 178019.38 |  |  | 11.34 | 38.8 |  |  |
| sp\|P62253\|UB2G1\_HUMAN | P62253 | UB2G1\_HUMAN | UBE2G1 | Ubiquitin-conjugating enzyme E2 G1 | 170 | 19.509 | 8.878764e+03 | 2.176229e+05 | 64170.645 | 153452.3 |  |  | 15.29 | 45.9 |  |  |
| sp\|Q92485\|ASM3B\_HUMAN | Q92485 | ASM3B\_HUMAN | SMPDL3B | Acid sphingomyelinase-like phosphodiesterase 3b | 455 | 50.814 | 1.259693e+04 | 8.042024e+05 | 64086.273 | 19992.082 | 380021.75 | 340102.25 | 5.71 | 13.4 | 16.9 | 9.9 |
| sp\|P07476\|INVO\_HUMAN | P07476 | INVO\_HUMAN | IVL | Involucrin | 585 | 68.479 | 1.484692e+04 | 1.277353e+06 | 64029.523 | 92809.67 | 816263.9 | 304249.56 | 5.13 | 31.5 | 40.2 | 29.9 |
| sp\|Q8TC12\|RDH11\_HUMAN | Q8TC12 | RDH11\_HUMAN | RDH11 | Retinol dehydrogenase 11 | 318 | 35.386 | 1.803296e+04 | 8.017068e+05 | 63875.72 | 408552.44 | 58885.516 | 270393.16 | 13.84 | 29.6 | 20.8 | 14.2 |
| sp\|P62906\|RL10A\_HUMAN | P62906 | RL10A\_HUMAN | RPL10A | Large ribosomal subunit protein uL1 | 217 | 24.831 | 6.992635e+03 | 2.181485e+05 | 63531.406 | 154617.1 |  |  | 12.44 | 26.7 |  |  |
| sp\|Q9Y2T2\|AP3M1\_HUMAN | Q9Y2T2 | AP3M1\_HUMAN | AP3M1 | AP-3 complex subunit mu-1 | 418 | 46.939 | 6.771212e+03 | 3.993166e+05 | 63358.766 | 335957.88 |  |  | 7.66 | 56.9 |  |  |
| sp\|Q9H4L5\|OSBL3\_HUMAN | Q9H4L5 | OSBL3\_HUMAN | OSBPL3 | Oxysterol-binding protein-related protein 3 | 887 | 101.224 | 3.131372e+03 | 3.982312e+05 | 62592.637 | 335638.6 |  |  | 5.75 | 35.4 |  |  |
| sp\|Q99698\|LYST\_HUMAN | Q99698 | LYST\_HUMAN | LYST | Lysosomal-trafficking regulator | 3801 | 429.139 | 2.490080e+02 | 1.342544e+05 | 62540.06 | 71714.33 |  |  | 1.89 | 23.0 |  |  |
| sp\|Q6ZNJ1\|NBEL2\_HUMAN | Q6ZNJ1 | NBEL2\_HUMAN | NBEAL2 | Neurobeachin-like protein 2 | 2754 | 302.517 | 6.753197e+02 | 2.566707e+05 | 62435.586 | 194235.16 |  |  | 2.51 | 27.7 |  |  |
| sp\|Q8IYB3\|SRRM1\_HUMAN | Q8IYB3 | SRRM1\_HUMAN | SRRM1 | Serine/arginine repetitive matrix protein 1 | 904 | 99.440 | 4.946436e+02 | 6.179746e+04 | 61797.46 |  |  |  | 4.87 |  |  |  |
| sp\|P57088\|TMM33\_HUMAN | P57088 | TMM33\_HUMAN | TMEM33 | Transmembrane protein 33 | 247 | 27.978 | 1.956480e+04 | 6.877162e+05 | 61115.844 | 308949.53 | 82504.555 | 235146.27 | 15.79 | 41.7 | 26.3 | 23.1 |
| sp\|Q8IZH2\|XRN1\_HUMAN | Q8IZH2 | XRN1\_HUMAN | XRN1 | 5'-3' exoribonuclease 1 | 1706 | 194.107 | 6.389465e+02 | 1.558199e+05 | 61019.883 | 94799.99 |  |  | 1.64 | 23.9 |  |  |
| sp\|P49914\|MTHFS\_HUMAN | P49914 | MTHFS\_HUMAN | MTHFS | 5-formyltetrahydrofolate cyclo-ligase | 203 | 23.256 | 8.053580e+03 | 2.353104e+05 | 60328.168 | 174982.25 |  |  | 5.91 | 58.1 |  |  |
| sp\|P23396\|RS3\_HUMAN | P23396 | RS3\_HUMAN | RPS3 | Small ribosomal subunit protein uS3 | 243 | 26.688 | 6.790872e+03 | 2.276978e+05 | 59924.27 |  | 124810.92 | 42962.57 | 9.05 |  | 34.2 | 25.9 |
| sp\|Q99798\|ACON\_HUMAN | Q99798 | ACON\_HUMAN | ACO2 | Aconitate hydratase, mitochondrial | 780 | 85.425 | 1.212731e+03 | 1.301567e+05 | 59821.562 |  | 70335.125 |  | 3.59 |  | 17.6 |  |
| sp\|Q9BU89\|DOHH\_HUMAN | Q9BU89 | DOHH\_HUMAN | DOHH | Deoxyhypusine hydroxylase | 302 | 33.220 | 1.427385e+03 | 5.957417e+04 | 59574.168 |  |  |  | 8.28 |  |  |  |
| sp\|Q96AJ9\|VTI1A\_HUMAN | Q96AJ9 | VTI1A\_HUMAN | VTI1A | Vesicle transport through interaction with t-SNAREs homolog 1A | 217 | 25.218 | 5.697550e+03 | 1.805161e+05 | 59573.695 |  | 50430.523 | 70511.875 | 11.98 |  | 21.2 | 30.9 |
| sp\|Q9H0E2\|TOLIP\_HUMAN | Q9H0E2 | TOLIP\_HUMAN | TOLLIP | Toll-interacting protein | 274 | 30.282 | 4.399878e+03 | 1.673950e+05 | 59232.797 | 99368.56 | 8793.603 |  | 13.14 | 35.4 | 23.0 |  |
| sp\|Q15369\|ELOC\_HUMAN | Q15369 | ELOC\_HUMAN | ELOC | Elongin-C | 112 | 12.473 | 5.290060e+04 | 8.289889e+05 | 59067.844 | 159098.64 | 464228.12 | 146594.25 | 37.5 | 77.7 | 58.9 | 21.4 |
| sp\|Q9BRA2\|TXD17\_HUMAN | Q9BRA2 | TXD17\_HUMAN | TXNDC17 | Thioredoxin domain-containing protein 17 | 123 | 13.941 | 2.272305e+05 | 3.979950e+06 | 58681.832 | 3654009.8 | 43804.043 | 223454.58 | 11.38 | 96.7 | 13.8 | 13.0 |
| sp\|Q5GLZ8\|HERC4\_HUMAN | Q5GLZ8 | HERC4\_HUMAN | HERC4 | Probable E3 ubiquitin-protein ligase HERC4 | 1057 | 116.270 | 3.982855e+02 | 5.818076e+04 | 58180.758 |  |  |  | 2.27 |  |  |  |
| sp\|Q9HBF4\|ZFYV1\_HUMAN | Q9HBF4 | ZFYV1\_HUMAN | ZFYVE1 | Zinc finger FYVE domain-containing protein 1 | 777 | 87.176 | 3.962019e+03 | 4.339410e+05 | 57364.906 | 321079.34 | 55496.742 |  | 6.18 | 36.7 | 18.1 |  |
| sp\|P51809\|VAMP7\_HUMAN | P51809 | VAMP7\_HUMAN | VAMP7 | Vesicle-associated membrane protein 7 | 220 | 24.935 | 2.796013e+04 | 8.759226e+05 | 57319.53 |  | 171728.8 | 646874.25 | 9.55 |  | 25.5 | 30.0 |
| sp\|Q13158\|FADD\_HUMAN | Q13158 | FADD\_HUMAN | FADD | FAS-associated death domain protein | 208 | 23.279 | 1.462114e+04 | 4.276246e+05 | 56910.766 | 370713.88 |  |  | 15.38 | 43.3 |  |  |
| sp\|A0A1W2PR19\|GSTT4\_HUMAN | A0A1W2PR19 | GSTT4\_HUMAN | GSTT4 | Glutathione S-transferase theta-4 | 241 | 26.510 | 1.706116e+03 | 5.682448e+04 | 56824.477 |  |  |  | 12.45 |  |  |  |
| sp\|Q8IWA5\|CTL2\_HUMAN | Q8IWA5 | CTL2\_HUMAN | SLC44A2 | Choline transporter-like protein 2 | 706 | 80.124 | 2.007911e+04 | 2.021270e+06 | 56609.9 | 55800.76 | 457402.22 | 1451457.0 | 7.37 | 18.3 | 24.6 | 19.0 |
| sp\|Q13541\|4EBP1\_HUMAN | Q13541 | 4EBP1\_HUMAN | EIF4EBP1 | Eukaryotic translation initiation factor 4E-binding protein 1 | 118 | 12.580 | 8.880022e+03 | 1.403498e+05 | 56567.35 |  | 58411.53 | 25370.93 | 22.03 |  | 30.5 | 29.7 |
| sp\|P15498\|VAV\_HUMAN | P15498 | VAV\_HUMAN | VAV1 | Proto-oncogene vav | 845 | 98.314 | 2.980324e+03 | 3.681256e+05 | 56291.18 | 311834.44 |  |  | 6.27 | 29.3 |  |  |
| sp\|Q9NQE9\|HINT3\_HUMAN | Q9NQE9 | HINT3\_HUMAN | HINT3 | Adenosine 5'-monophosphoramidase HINT3 | 182 | 20.361 | 1.233762e+04 | 3.156078e+05 | 55969.97 | 221886.48 | 37751.383 |  | 13.74 | 43.4 | 24.2 |  |
| sp\|Q13501\|SQSTM\_HUMAN | Q13501 | SQSTM\_HUMAN | SQSTM1 | Sequestosome-1 | 440 | 47.687 | 4.513877e+03 | 2.704375e+05 | 55814.086 | 214623.38 |  |  | 6.59 | 21.1 |  |  |
| sp\|P20962\|PTMS\_HUMAN | P20962 | PTMS\_HUMAN | PTMS | Parathymosin | 102 | 11.530 | 1.941616e+04 | 2.812612e+05 | 55617.887 | 211799.4 | 13843.89 |  | 11.76 | 27.5 | 11.8 |  |
| sp\|Q9BRP8\|PYM1\_HUMAN | Q9BRP8 | PYM1\_HUMAN | PYM1 | Partner of Y14 and mago | 204 | 22.656 | 9.397254e+03 | 2.674862e+05 | 55349.348 | 151109.31 | 25098.084 | 35929.438 | 8.82 | 49.5 | 23.5 | 21.1 |
| sp\|Q07666\|KHDR1\_HUMAN | Q07666 | KHDR1\_HUMAN | KHDRBS1 | KH domain-containing, RNA-binding, signal transduction-associated protein 1 | 443 | 48.227 | 1.105820e+03 | 6.700266e+04 | 55174.99 | 11827.667 |  |  | 9.03 | 23.3 |  |  |
| sp\|O75348\|VATG1\_HUMAN | O75348 | VATG1\_HUMAN | ATP6V1G1 | V-type proton ATPase subunit G 1 | 118 | 13.758 | 2.063109e+05 | 3.566109e+06 | 54071.953 | 3218705.5 | 243239.0 | 50092.53 | 9.32 | 52.5 | 31.4 | 16.9 |
| sp\|Q8WZ73\|RFFL\_HUMAN | Q8WZ73 | RFFL\_HUMAN | RFFL | E3 ubiquitin-protein ligase rififylin | 363 | 40.514 | 1.676098e+03 | 8.531426e+04 | 54045.617 | 31268.64 |  |  | 11.29 | 31.7 |  |  |
| sp\|Q9UM54\|MYO6\_HUMAN | Q9UM54 | MYO6\_HUMAN | MYO6 | Unconventional myosin-VI | 1294 | 142.340 | 2.993309e+02 | 5.352980e+04 | 53529.8 |  |  |  | 2.78 |  |  |  |
| sp\|P84103\|SRSF3\_HUMAN | P84103 | SRSF3\_HUMAN | SRSF3 | Serine/arginine-rich splicing factor 3 | 164 | 18.040 | 2.359722e+03 | 5.348284e+04 | 53482.844 |  |  |  | 15.85 |  |  |  |
| sp\|Q13033\|STRN3\_HUMAN | Q13033 | STRN3\_HUMAN | STRN3 | Striatin-3 | 797 | 87.209 | 1.612578e+03 | 1.766848e+05 | 53204.53 | 123480.27 |  |  | 4.77 | 33.6 |  |  |
| sp\|Q13574\|DGKZ\_HUMAN | Q13574 | DGKZ\_HUMAN | DGKZ | Diacylglycerol kinase zeta | 928 | 103.981 | 2.070672e+03 | 2.705095e+05 | 53139.05 | 217370.47 |  |  | 5.5 | 42.6 |  |  |
| sp\|Q14152\|EIF3A\_HUMAN | Q14152 | EIF3A\_HUMAN | EIF3A | Eukaryotic translation initiation factor 3 subunit A | 1382 | 166.569 | 9.419376e+02 | 1.971213e+05 | 52738.895 | 127204.5 | 17177.867 |  | 4.27 | 33.8 | 22.5 |  |
| sp\|Q15814\|TBCC\_HUMAN | Q15814 | TBCC\_HUMAN | TBCC | Tubulin-specific chaperone C | 346 | 39.248 | 1.579769e+03 | 7.789834e+04 | 52385.312 | 25513.025 |  |  | 9.83 | 36.1 |  |  |
| sp\|O00559\|RCAS1\_HUMAN | O00559 | RCAS1\_HUMAN | EBAG9 | Receptor-binding cancer antigen expressed on SiSo cells | 213 | 24.377 | 4.425279e+03 | 1.355308e+05 | 52028.055 | 83502.77 |  |  | 16.9 | 28.2 |  |  |
| sp\|P23219\|PGH1\_HUMAN | P23219 | PGH1\_HUMAN | PTGS1 | Prostaglandin G/H synthase 1 | 599 | 68.686 | 1.390152e+03 | 1.199631e+05 | 51906.336 | 23583.725 | 44473.047 |  | 6.18 | 30.4 | 18.7 |  |
| sp\|A0A075B6P5\|KV228\_HUMAN | A0A075B6P5 | KV228\_HUMAN | IGKV2-28 | Immunoglobulin kappa variable 2-28 | 120 | 12.957 | 3.838134e+04 | 6.248010e+05 | 51418.523 |  | 573382.44 |  | 10.83 |  | 25.8 |  |
| sp\|O95070\|YIF1A\_HUMAN | O95070 | YIF1A\_HUMAN | YIF1A | Protein YIF1A | 293 | 32.011 | 1.854154e+04 | 7.456966e+05 | 51390.1 | 182742.84 | 111476.16 | 400087.5 | 5.12 | 17.7 | 37.2 | 11.6 |
| sp\|O75592\|MYCB2\_HUMAN | O75592 | MYCB2\_HUMAN | MYCBP2 | E3 ubiquitin-protein ligase MYCBP2 | 4678 | 513.636 | 4.059698e+02 | 2.619789e+05 | 51268.316 | 210710.62 |  |  | 1.45 | 31.0 |  |  |
| sp\|Q5VWP2\|TET5C\_HUMAN | Q5VWP2 | TET5C\_HUMAN | TENT5C | Terminal nucleotidyltransferase 5C | 391 | 44.944 | 1.096063e+03 | 6.189054e+04 | 50936.75 | 10953.792 |  |  | 8.18 | 19.7 |  |  |
| sp\|Q9BT73\|PSMG3\_HUMAN | Q9BT73 | PSMG3\_HUMAN | PSMG3 | Proteasome assembly chaperone 3 | 122 | 13.104 | 2.644787e+05 | 4.354234e+06 | 50931.9 | 4181880.2 | 121421.72 |  | 12.3 | 73.8 | 45.1 |  |
| sp\|Q9H9Q4\|NHEJ1\_HUMAN | Q9H9Q4 | NHEJ1\_HUMAN | NHEJ1 | Non-homologous end-joining factor 1 | 299 | 33.337 | 3.358857e+03 | 1.406809e+05 | 50708.883 | 66651.6 | 23320.428 |  | 8.36 | 40.8 | 8.4 |  |
| sp\|O14929\|HAT1\_HUMAN | O14929 | HAT1\_HUMAN | HAT1 | Histone acetyltransferase type B catalytic subunit | 419 | 49.541 | 1.987570e+03 | 1.237099e+05 | 50701.703 | 73008.16 |  |  | 7.4 | 21.5 |  |  |
| sp\|Q14320\|FA50A\_HUMAN | Q14320 | FA50A\_HUMAN | FAM50A | Protein FAM50A | 339 | 40.242 | 8.448696e+03 | 4.271559e+05 | 50629.453 | 376526.44 |  |  | 12.68 | 42.2 |  |  |
| sp\|Q8N5K1\|CISD2\_HUMAN | Q8N5K1 | CISD2\_HUMAN | CISD2 | CDGSH iron-sulfur domain-containing protein 2 | 135 | 15.278 | 1.696394e+05 | 3.256195e+06 | 50314.652 | 546220.06 | 1701414.5 | 958245.6 | 8.89 | 57.0 | 43.0 | 27.4 |
| sp\|Q14691\|PSF1\_HUMAN | Q14691 | PSF1\_HUMAN | GINS1 | DNA replication complex GINS protein PSF1 | 196 | 22.988 | 5.749193e+03 | 1.660448e+05 | 50289.062 | 115755.73 |  |  | 14.8 | 51.0 |  |  |
| sp\|O94760\|DDAH1\_HUMAN | O94760 | DDAH1\_HUMAN | DDAH1 | N(G),N(G)-dimethylarginine dimethylaminohydrolase 1 | 285 | 31.122 | 6.662206e+03 | 2.604970e+05 | 49886.895 | 210610.1 |  |  | 7.02 | 57.2 |  |  |
| sp\|Q6P1M0\|S27A4\_HUMAN | Q6P1M0 | S27A4\_HUMAN | SLC27A4 | Long-chain fatty acid transport protein 4 | 643 | 72.064 | 7.032978e+02 | 6.367586e+04 | 49757.168 | 13918.6875 |  |  | 3.89 | 30.5 |  |  |
| sp\|Q9UPT5\|EXOC7\_HUMAN | Q9UPT5 | EXOC7\_HUMAN | EXOC7 | Exocyst complex component 7 | 735 | 83.382 | 2.261406e+03 | 2.369016e+05 | 49507.727 | 187393.83 |  |  | 7.48 | 33.2 |  |  |
| sp\|Q99543\|DNJC2\_HUMAN | Q99543 | DNJC2\_HUMAN | DNAJC2 | DnaJ homolog subfamily C member 2 | 621 | 71.996 | 1.910089e+03 | 1.727742e+05 | 49197.51 | 123576.74 |  |  | 7.25 | 32.5 |  |  |
| sp\|Q9Y6A5\|TACC3\_HUMAN | Q9Y6A5 | TACC3\_HUMAN | TACC3 | Transforming acidic coiled-coil-containing protein 3 | 838 | 90.360 | 2.514531e+03 | 2.854633e+05 | 49187.617 | 236275.69 |  |  | 2.74 | 29.0 |  |  |
| sp\|O75175\|CNOT3\_HUMAN | O75175 | CNOT3\_HUMAN | CNOT3 | CCR4-NOT transcription complex subunit 3 | 753 | 81.872 | 1.937921e+03 | 1.993373e+05 | 49180.47 | 150156.86 |  |  | 2.79 | 18.7 |  |  |
| sp\|Q8TF71\|MOT10\_HUMAN | Q8TF71 | MOT10\_HUMAN | SLC16A10 | Monocarboxylate transporter 10 | 515 | 55.493 | 1.583640e+04 | 1.104109e+06 | 49094.79 | 46709.832 | 242591.81 | 765712.5 | 8.54 | 27.4 | 18.3 | 22.9 |
| sp\|Q9H1Y0\|ATG5\_HUMAN | Q9H1Y0 | ATG5\_HUMAN | ATG5 | Autophagy protein 5 | 275 | 32.447 | 2.280845e+03 | 9.297955e+04 | 48894.293 | 44085.26 |  |  | 8.0 | 43.6 |  |  |
| sp\|B0I1T2\|MYO1G\_HUMAN | B0I1T2 | MYO1G\_HUMAN | MYO1G | Unconventional myosin-Ig | 1018 | 116.442 | 5.118545e+02 | 7.488129e+04 | 48887.414 | 25993.873 |  |  | 2.16 | 35.3 |  |  |
| sp\|Q9NWM0\|SMOX\_HUMAN | Q9NWM0 | SMOX\_HUMAN | SMOX | Spermine oxidase | 555 | 61.819 | 2.348843e+03 | 1.824287e+05 | 47781.81 | 134646.89 |  |  | 5.41 | 33.3 |  |  |
| sp\|Q15125\|EBP\_HUMAN | Q15125 | EBP\_HUMAN | EBP | 3-beta-hydroxysteroid-Delta(8),Delta(7)-isomerase | 230 | 26.353 | 1.678985e+04 | 5.558966e+05 | 47652.207 | 146115.33 | 54414.645 | 307714.4 | 5.22 | 20.9 | 12.6 | 5.2 |
| sp\|P24390\|ERD21\_HUMAN | P24390 | ERD21\_HUMAN | KDELR1 | ER lumen protein-retaining receptor 1 | 212 | 24.542 | 3.526217e+04 | 1.087267e+06 | 47586.36 | 89273.96 | 264580.16 | 685826.5 | 5.19 | 5.2 | 9.0 | 5.2 |
| sp\|C9J7I0\|UMAD1\_HUMAN | C9J7I0 | UMAD1\_HUMAN | UMAD1 | UBAP1-MVB12-associated (UMA)-domain containing protein 1 | 137 | 15.200 | 8.994246e+04 | 1.717614e+06 | 47457.67 | 337093.8 | 1235434.0 | 97628.28 | 21.9 | 46.7 | 38.7 | 21.9 |
| sp\|Q9NUY8\|TBC23\_HUMAN | Q9NUY8 | TBC23\_HUMAN | TBC1D23 | TBC1 domain family member 23 | 699 | 78.322 | 7.667100e+02 | 7.544528e+04 | 47412.28 | 28032.998 |  |  | 5.01 | 24.7 |  |  |
| sp\|Q92551\|IP6K1\_HUMAN | Q92551 | IP6K1\_HUMAN | IP6K1 | Inositol hexakisphosphate kinase 1 | 441 | 50.236 | 2.058518e+03 | 1.299233e+05 | 47376.83 | 82546.45 |  |  | 5.9 | 55.1 |  |  |
| sp\|O75264\|SIM24\_HUMAN | O75264 | SIM24\_HUMAN | SMIM24 | Small integral membrane protein 24 | 130 | 14.964 | 3.359662e+04 | 6.316267e+05 | 46606.24 | 326596.6 | 122333.336 | 136090.48 | 18.46 | 35.4 | 28.5 | 28.5 |
| sp\|Q9H400\|LIME1\_HUMAN | Q9H400 | LIME1\_HUMAN | LIME1 | Lck-interacting transmembrane adapter 1 | 295 | 31.288 | 6.008436e+04 | 2.361872e+06 | 46443.81 | 65783.85 | 1208020.8 | 1041623.8 | 7.8 | 57.3 | 39.0 | 22.7 |
| sp\|P10515\|ODP2\_HUMAN | P10515 | ODP2\_HUMAN | DLAT | Dihydrolipoyllysine-residue acetyltransferase component of pyruvate dehydrogenase complex, mitochondrial | 647 | 68.997 | 1.173342e+03 | 1.017119e+05 | 46439.938 |  | 55271.977 |  | 6.65 |  | 28.6 |  |
| sp\|P47712\|PA24A\_HUMAN | P47712 | PA24A\_HUMAN | PLA2G4A | Cytosolic phospholipase A2 | 749 | 85.239 | 4.085640e+03 | 4.375379e+05 | 45340.465 | 392197.4 |  |  | 3.34 | 32.7 |  |  |
| sp\|Q00325\|S25A3\_HUMAN | Q00325 | S25A3\_HUMAN | SLC25A3 | Solute carrier family 25 member 3 | 362 | 40.095 | 2.598004e+03 | 1.308721e+05 | 44764.094 | 35862.945 | 50245.11 |  | 3.31 | 43.4 | 19.1 |  |
| sp\|Q9BW91\|NUDT9\_HUMAN | Q9BW91 | NUDT9\_HUMAN | NUDT9 | ADP-ribose pyrophosphatase, mitochondrial | 350 | 39.125 | 3.271614e+03 | 1.608176e+05 | 44502.023 | 116315.56 |  |  | 4.0 | 31.1 |  |  |
| sp\|P54687\|BCAT1\_HUMAN | P54687 | BCAT1\_HUMAN | BCAT1 | Branched-chain-amino-acid aminotransferase, cytosolic | 386 | 42.460 | 8.232584e+02 | 4.391707e+04 | 43917.066 |  |  |  | 7.25 |  |  |  |
| sp\|O43719\|HTSF1\_HUMAN | O43719 | HTSF1\_HUMAN | HTATSF1 | HIV Tat-specific factor 1 | 755 | 85.853 | 1.947501e+03 | 2.100633e+05 | 43878.047 | 166185.28 |  |  | 4.11 | 24.8 |  |  |
| sp\|Q6ZVK8\|NUD18\_HUMAN | Q6ZVK8 | NUD18\_HUMAN | NUDT18 | 8-oxo-dGDP phosphatase NUDT18 | 323 | 35.530 | 9.803109e+02 | 4.375989e+04 | 43759.887 |  |  |  | 7.43 |  |  |  |
| sp\|Q13263\|TIF1B\_HUMAN | Q13263 | TIF1B\_HUMAN | TRIM28 | Transcription intermediary factor 1-beta | 835 | 88.550 | 1.265923e+03 | 1.408358e+05 | 43719.645 | 97116.164 |  |  | 3.83 | 24.1 |  |  |
| sp\|Q96DA0\|ZG16B\_HUMAN | Q96DA0 | ZG16B\_HUMAN | ZG16B | Zymogen granule protein 16 homolog B | 208 | 22.739 | 2.439440e+04 | 6.969131e+05 | 43665.375 | 9820.392 | 643427.3 |  | 22.12 | 34.6 | 33.7 |  |
| sp\|O14772\|FPGT\_HUMAN | O14772 | FPGT\_HUMAN | FPGT | Fucose-1-phosphate guanylyltransferase | 607 | 68.010 | 7.496374e+02 | 6.405325e+04 | 43657.96 | 20395.293 |  |  | 4.45 | 33.8 |  |  |
| sp\|Q8WUX2\|CHAC2\_HUMAN | Q8WUX2 | CHAC2\_HUMAN | CHAC2 | Glutathione-specific gamma-glutamylcyclotransferase 2 | 184 | 20.875 | 2.090322e+04 | 5.482224e+05 | 43636.367 | 504586.06 |  |  | 14.13 | 33.7 |  |  |
| sp\|O94992\|HEXI1\_HUMAN | O94992 | HEXI1\_HUMAN | HEXIM1 | Protein HEXIM1 | 359 | 40.623 | 1.918647e+03 | 9.792290e+04 | 43514.08 | 54408.82 |  |  | 6.96 | 24.5 |  |  |
| sp\|Q5T5C0\|STXB5\_HUMAN | Q5T5C0 | STXB5\_HUMAN | STXBP5 | Syntaxin-binding protein 5 | 1151 | 127.573 | 1.727439e+03 | 2.768718e+05 | 43365.484 | 233506.33 |  |  | 3.13 | 32.1 |  |  |
| sp\|Q8TEA7\|TBCK\_HUMAN | Q8TEA7 | TBCK\_HUMAN | TBCK | TBC domain-containing protein kinase-like protein | 893 | 100.679 | 1.178152e+03 | 1.490244e+05 | 43323.62 | 105700.78 |  |  | 4.82 | 39.3 |  |  |
| sp\|Q6WCQ1\|MPRIP\_HUMAN | Q6WCQ1 | MPRIP\_HUMAN | MPRIP | Myosin phosphatase Rho-interacting protein | 1025 | 116.533 | 7.169871e+02 | 1.049730e+05 | 43004.914 | 61968.04 |  |  | 3.32 | 35.3 |  |  |
| sp\|Q6RW13\|ATRAP\_HUMAN | Q6RW13 | ATRAP\_HUMAN | AGTRAP | Type-1 angiotensin II receptor-associated protein | 159 | 17.419 | 1.581713e+04 | 3.461530e+05 | 42886.297 | 39138.027 |  | 264128.7 | 13.84 | 30.2 |  | 10.7 |
| sp\|Q5TAQ9\|DCAF8\_HUMAN | Q5TAQ9 | DCAF8\_HUMAN | DCAF8 | DDB1- and CUL4-associated factor 8 | 597 | 66.852 | 1.501584e+03 | 1.261192e+05 | 42461.336 | 83657.84 |  |  | 3.35 | 30.5 |  |  |
| sp\|P61916\|NPC2\_HUMAN | P61916 | NPC2\_HUMAN | NPC2 | NPC intracellular cholesterol transporter 2 | 151 | 16.610 | 2.029249e+03 | 4.234694e+04 | 42346.945 |  |  |  | 10.6 |  |  |  |
| sp\|Q08209\|PP2BA\_HUMAN | Q08209 | PP2BA\_HUMAN | PPP3CA | Protein phosphatase 3 catalytic subunit alpha | 521 | 58.688 | 3.850724e+03 | 2.839284e+05 | 42193.137 | 241735.31 |  |  | 6.33 | 35.1 |  |  |
| sp\|Q96CW6\|S7A6O\_HUMAN | Q96CW6 | S7A6O\_HUMAN | SLC7A6OS | Probable RNA polymerase II nuclear localization protein SLC7A6 | 309 | 35.028 | 2.147390e+03 | 9.450252e+04 | 42043.75 | 52458.77 |  |  | 3.56 | 17.8 |  |  |
| sp\|Q86VR2\|RETR3\_HUMAN | Q86VR2 | RETR3\_HUMAN | RETREG3 | Reticulophagy regulator 3 | 466 | 51.396 | 3.114118e+04 | 2.010859e+06 | 41966.754 | 78356.03 | 1879235.1 | 11300.716 | 3.22 | 17.6 | 17.8 | 8.8 |
| sp\|P41240\|CSK\_HUMAN | P41240 | CSK\_HUMAN | CSK | Tyrosine-protein kinase CSK | 450 | 50.704 | 2.040166e+03 | 1.299645e+05 | 41869.676 | 65688.43 |  | 22406.414 | 5.78 | 38.0 |  | 9.1 |
| sp\|Q96EP0\|RNF31\_HUMAN | Q96EP0 | RNF31\_HUMAN | RNF31 | E3 ubiquitin-protein ligase RNF31 | 1072 | 117.920 | 2.817331e+02 | 4.173905e+04 | 41739.05 |  |  |  | 3.08 |  |  |  |
| sp\|Q969T4\|UB2E3\_HUMAN | Q969T4 | UB2E3\_HUMAN | UBE2E3 | Ubiquitin-conjugating enzyme E2 E3 | 207 | 22.913 | 5.837057e+03 | 1.680324e+05 | 41242.984 | 98056.83 | 28732.58 |  | 9.66 | 47.3 | 38.6 |  |
| sp\|Q12968\|NFAC3\_HUMAN | Q12968 | NFAC3\_HUMAN | NFATC3 | Nuclear factor of activated T-cells, cytoplasmic 3 | 1075 | 115.594 | 2.394892e+04 | 3.478071e+06 | 41070.49 | 3437000.5 |  |  | 2.79 | 17.7 |  |  |
| sp\|P12277\|KCRB\_HUMAN | P12277 | KCRB\_HUMAN | CKB | Creatine kinase B-type | 381 | 42.644 | 1.008180e+03 | 5.401485e+04 | 40883.492 | 13131.362 |  |  | 8.92 | 44.4 |  |  |
| sp\|O75915\|PRAF3\_HUMAN | O75915 | PRAF3\_HUMAN | ARL6IP5 | PRA1 family protein 3 | 188 | 20.680 | 1.571089e+03 | 4.081957e+04 | 40819.566 |  |  |  | 5.85 |  |  |  |
| sp\|Q9BV57\|MTND\_HUMAN | Q9BV57 | MTND\_HUMAN | ADI1 | Acireductone dioxygenase | 179 | 21.498 | 1.391329e+04 | 3.757898e+05 | 40515.92 | 335273.88 |  |  | 17.32 | 59.2 |  |  |
| sp\|P61254\|RL26\_HUMAN | P61254 | RL26\_HUMAN | RPL26 | Large ribosomal subunit protein uL24 | 145 | 15.950 | 2.020856e+03 | 4.049611e+04 | 40496.105 |  |  |  | 6.21 |  |  |  |
| sp\|Q9BUB5\|MKNK1\_HUMAN | Q9BUB5 | MKNK1\_HUMAN | MKNK1 | MAP kinase-interacting serine/threonine-protein kinase 1 | 465 | 51.342 | 1.429517e+03 | 9.221025e+04 | 40215.266 | 51994.984 |  |  | 5.38 | 34.8 |  |  |
| sp\|P30047\|GFRP\_HUMAN | P30047 | GFRP\_HUMAN | GCHFR | GTP cyclohydrolase 1 feedback regulatory protein | 84 | 9.698 | 3.261418e+04 | 3.973798e+05 | 39809.25 | 357570.5 |  |  | 19.05 | 82.1 |  |  |
| sp\|P02649\|APOE\_HUMAN | P02649 | APOE\_HUMAN | APOE | Apolipoprotein E | 317 | 36.154 | 2.472623e+04 | 1.123134e+06 | 39510.07 |  | 1068872.9 | 14750.748 | 10.09 |  | 37.9 | 27.4 |
| sp\|O15258\|RER1\_HUMAN | O15258 | RER1\_HUMAN | RER1 | Protein RER1 | 196 | 22.958 | 1.563093e+04 | 4.508542e+05 | 39117.68 | 187028.0 | 93138.25 | 131570.28 | 14.8 | 58.7 | 24.5 | 17.3 |
| sp\|Q969E2\|SCAM4\_HUMAN | Q969E2 | SCAM4\_HUMAN | SCAMP4 | Secretory carrier-associated membrane protein 4 | 229 | 25.728 | 1.664205e+04 | 5.379354e+05 | 38775.934 | 30138.508 | 348482.94 | 120538.02 | 4.8 | 9.6 | 10.0 | 10.0 |
| sp\|P30050\|RL12\_HUMAN | P30050 | RL12\_HUMAN | RPL12 | Large ribosomal subunit protein uL11 | 165 | 17.819 | 3.775289e+03 | 8.451829e+04 | 38062.508 |  | 28053.164 | 18402.615 | 15.15 |  | 32.7 | 17.6 |
| sp\|Q9Y3T6\|R3HC1\_HUMAN | Q9Y3T6 | R3HC1\_HUMAN | R3HCC1 | R3H and coiled-coil domain-containing protein 1 | 440 | 49.092 | 2.881028e+03 | 1.776950e+05 | 37684.996 | 140010.05 |  |  | 8.41 | 46.1 |  |  |
| sp\|Q01844\|EWS\_HUMAN | Q01844 | EWS\_HUMAN | EWSR1 | RNA-binding protein EWS | 656 | 68.478 | 6.863092e+02 | 5.904566e+04 | 37365.566 | 21680.098 |  |  | 2.13 | 31.9 |  |  |
| sp\|Q13445\|TMED1\_HUMAN | Q13445 | TMED1\_HUMAN | TMED1 | Transmembrane emp24 domain-containing protein 1 | 227 | 25.206 | 5.009876e+04 | 1.586529e+06 | 37195.68 | 314809.38 | 719577.8 | 514946.25 | 7.49 | 36.6 | 26.4 | 22.0 |
| sp\|P24557\|THAS\_HUMAN | P24557 | THAS\_HUMAN | TBXAS1 | Thromboxane-A synthase | 533 | 58.630 | 5.045230e+02 | 3.716362e+04 | 37163.62 |  |  |  | 2.25 |  |  |  |
| sp\|Q9NWX5\|ASB6\_HUMAN | Q9NWX5 | ASB6\_HUMAN | ASB6 | Ankyrin repeat and SOCS box protein 6 | 421 | 47.136 | 3.019438e+03 | 1.788117e+05 | 36516.72 | 142295.0 |  |  | 6.18 | 28.0 |  |  |
| sp\|Q6UUV9\|CRTC1\_HUMAN | Q6UUV9 | CRTC1\_HUMAN | CRTC1 | CREB-regulated transcription coactivator 1 | 634 | 69.740 | 4.150504e+02 | 3.636637e+04 | 36366.37 |  |  |  | 2.84 |  |  |  |
| sp\|Q6ZNE5\|BAKOR\_HUMAN | Q6ZNE5 | BAKOR\_HUMAN | ATG14 | Beclin 1-associated autophagy-related key regulator | 492 | 55.309 | 3.456784e+03 | 2.402068e+05 | 36271.53 | 203935.25 |  |  | 7.11 | 17.9 |  |  |
| sp\|Q8NAA5\|LR75A\_HUMAN | Q8NAA5 | LR75A\_HUMAN | LRRC75A | Leucine-rich repeat-containing protein 75A | 344 | 37.780 | 2.857439e+03 | 1.356301e+05 | 36119.08 | 99511.0 |  |  | 11.05 | 47.1 |  |  |
| sp\|Q99735\|MGST2\_HUMAN | Q99735 | MGST2\_HUMAN | MGST2 | Microsomal glutathione S-transferase 2 | 147 | 16.621 | 2.212601e+04 | 4.620376e+05 | 36068.156 |  | 193124.39 | 232845.05 | 9.52 |  | 10.9 | 9.5 |
| sp\|O95674\|CDS2\_HUMAN | O95674 | CDS2\_HUMAN | CDS2 | Phosphatidate cytidylyltransferase 2 | 445 | 51.418 | 2.402287e+03 | 1.551877e+05 | 36032.14 | 46024.527 |  | 73131.03 | 4.27 | 28.3 |  | 14.4 |
| sp\|Q96JJ7\|TMX3\_HUMAN | Q96JJ7 | TMX3\_HUMAN | TMX3 | Protein disulfide-isomerase TMX3 | 454 | 51.872 | 1.448960e+04 | 9.442926e+05 | 35931.336 | 89145.64 | 394309.3 | 424906.34 | 9.91 | 33.3 | 28.0 | 29.7 |
| sp\|Q9BXI6\|TB10A\_HUMAN | Q9BXI6 | TB10A\_HUMAN | TBC1D10A | TBC1 domain family member 10A | 508 | 55.880 | 5.081840e+02 | 3.567751e+04 | 35677.508 |  |  |  | 2.36 |  |  |  |
| sp\|Q15427\|SF3B4\_HUMAN | Q15427 | SF3B4\_HUMAN | SF3B4 | Splicing factor 3B subunit 4 | 424 | 44.386 | 4.516289e+03 | 2.518517e+05 | 35478.31 | 170264.36 | 46109.05 |  | 3.3 | 30.0 | 12.7 |  |
| sp\|Q15311\|RBP1\_HUMAN | Q15311 | RBP1\_HUMAN | RALBP1 | RalA-binding protein 1 | 655 | 76.063 | 2.199292e+04 | 2.101713e+06 | 35190.33 | 138359.1 | 1928164.0 |  | 4.73 | 33.3 | 23.7 |  |
| sp\|Q14749\|GNMT\_HUMAN | Q14749 | GNMT\_HUMAN | GNMT | Glycine N-methyltransferase | 295 | 32.450 | 8.508322e+02 | 3.468772e+04 | 34687.723 |  |  |  | 6.78 |  |  |  |
| sp\|Q8ND04\|SMG8\_HUMAN | Q8ND04 | SMG8\_HUMAN | SMG8 | Nonsense-mediated mRNA decay factor SMG8 | 991 | 109.684 | 1.154330e+03 | 1.590708e+05 | 34216.746 | 124854.016 |  |  | 3.63 | 44.7 |  |  |
| sp\|Q12965\|MYO1E\_HUMAN | Q12965 | MYO1E\_HUMAN | MYO1E | Unconventional myosin-Ie | 1108 | 121.880 | 2.201474e+02 | 3.371034e+04 | 33710.336 |  |  |  | 3.7 |  |  |  |
| sp\|Q9NWS6\|F118A\_HUMAN | Q9NWS6 | F118A\_HUMAN | FAM118A | Protein FAM118A | 357 | 40.269 | 5.043882e+03 | 2.551837e+05 | 33567.28 | 221616.42 |  |  | 8.96 | 49.6 |  |  |
| sp\|Q9C0D3\|ZY11B\_HUMAN | Q9C0D3 | ZY11B\_HUMAN | ZYG11B | Protein zyg-11 homolog B | 744 | 83.921 | 7.141980e+02 | 7.530199e+04 | 33097.89 | 42204.1 |  |  | 2.96 | 23.1 |  |  |
| sp\|Q5MNZ6\|WIPI3\_HUMAN | Q5MNZ6 | WIPI3\_HUMAN | WDR45B | WD repeat domain phosphoinositide-interacting protein 3 | 344 | 38.122 | 3.860557e+03 | 1.849025e+05 | 32982.766 | 151919.75 |  |  | 4.07 | 44.5 |  |  |
| sp\|P63027\|VAMP2\_HUMAN | P63027 | VAMP2\_HUMAN | VAMP2 | Vesicle-associated membrane protein 2 | 116 | 12.663 | 1.725964e+05 | 2.745905e+06 | 32313.723 |  | 2713590.8 |  | 14.66 |  | 34.5 |  |
| sp\|Q9Y2G2\|CARD8\_HUMAN | Q9Y2G2 | CARD8\_HUMAN | CARD8 | Caspase recruitment domain-containing protein 8 | 537 | 60.652 | 1.064568e+04 | 8.112146e+05 | 32092.66 | 779121.9 |  |  | 1.86 | 26.6 |  |  |
| sp\|P81133\|SIM1\_HUMAN | P81133 | SIM1\_HUMAN | SIM1 | Single-minded homolog 1 | 766 | 84.260 | 3.027946e+02 | 3.205433e+04 | 32054.332 |  |  |  | 1.31 |  |  |  |
| sp\|Q9Y5S1\|TRPV2\_HUMAN | Q9Y5S1 | TRPV2\_HUMAN | TRPV2 | Transient receptor potential cation channel subfamily V member 2 | 764 | 85.981 | 1.073867e+04 | 1.160032e+06 | 32053.797 |  | 489128.56 | 638849.75 | 5.37 |  | 26.0 | 16.9 |
| sp\|P08195\|4F2\_HUMAN | P08195 | 4F2\_HUMAN | SLC3A2 | 4F2 cell-surface antigen heavy chain | 630 | 67.994 | 7.948492e+03 | 6.790043e+05 | 32036.545 | 23727.463 | 468171.5 | 155068.77 | 8.73 | 18.7 | 26.7 | 22.4 |
| sp\|Q9NR12\|PDLI7\_HUMAN | Q9NR12 | PDLI7\_HUMAN | PDLIM7 | PDZ and LIM domain protein 7 | 457 | 49.845 | 1.639881e+03 | 1.026954e+05 | 32026.338 | 70669.11 |  |  | 5.47 | 46.6 |  |  |
| sp\|Q92643\|GPI8\_HUMAN | Q92643 | GPI8\_HUMAN | PIGK | GPI-anchor transamidase | 395 | 45.252 | 2.302857e+03 | 1.309248e+05 | 31796.059 |  | 62143.355 | 36985.4 | 5.06 |  | 7.6 | 13.9 |
| sp\|Q5JPI9\|EFMT2\_HUMAN | Q5JPI9 | EFMT2\_HUMAN | EEF1AKMT2 | EEF1A lysine methyltransferase 2 | 291 | 31.830 | 3.598523e+03 | 1.439057e+05 | 31774.488 | 96680.59 | 15450.652 |  | 5.5 | 22.0 | 12.0 |  |
| sp\|Q07955\|SRSF1\_HUMAN | Q07955 | SRSF1\_HUMAN | SRSF1 | Serine/arginine-rich splicing factor 1 | 248 | 27.280 | 9.156138e+02 | 3.138151e+04 | 31381.51 |  |  |  | 4.03 |  |  |  |
| sp\|Q9Y3C5\|RNF11\_HUMAN | Q9Y3C5 | RNF11\_HUMAN | RNF11 | RING finger protein 11 | 154 | 17.444 | 1.420011e+03 | 3.112110e+04 | 31121.098 | 0.0 |  |  | 7.79 | 57.1 |  |  |
| sp\|Q96S53\|TESK2\_HUMAN | Q96S53 | TESK2\_HUMAN | TESK2 | Dual specificity testis-specific protein kinase 2 | 571 | 63.639 | 1.696866e+03 | 1.356714e+05 | 31062.574 | 104608.78 |  |  | 3.33 | 42.9 |  |  |
| sp\|P05386\|RLA1\_HUMAN | P05386 | RLA1\_HUMAN | RPLP1 | Large ribosomal subunit protein P1 | 114 | 11.514 | 9.686603e+03 | 1.401248e+05 | 31062.113 | 87193.82 | 9834.834 | 12033.995 | 14.04 | 14.0 | 14.0 | 14.0 |
| sp\|Q9HCY8\|S10AE\_HUMAN | Q9HCY8 | S10AE\_HUMAN | S100A14 | Protein S100-A14 | 104 | 11.662 | 1.137670e+05 | 1.666888e+06 | 30910.82 | 21463.996 | 824014.5 | 790498.8 | 14.42 | 42.3 | 40.4 | 44.2 |
| sp\|Q9UJC5\|SH3L2\_HUMAN | Q9UJC5 | SH3L2\_HUMAN | SH3BGRL2 | SH3 domain-binding glutamic acid-rich-like protein 2 | 107 | 12.326 | 5.058391e+05 | 7.833425e+06 | 30854.562 | 7784694.0 | 17876.7 |  | 13.08 | 72.9 | 42.1 |  |
| sp\|P80188\|NGAL\_HUMAN | P80188 | NGAL\_HUMAN | LCN2 | Neutrophil gelatinase-associated lipocalin | 198 | 22.588 | 2.941282e+03 | 8.347024e+04 | 30724.145 | 45111.625 | 7634.465 |  | 7.58 | 25.8 | 11.6 |  |
| sp\|P19447\|ERCC3\_HUMAN | P19447 | ERCC3\_HUMAN | ERCC3 | General transcription and DNA repair factor IIH helicase subunit XPB | 782 | 86.020 | 2.832592e+02 | 3.061262e+04 | 30612.623 |  |  |  | 1.15 |  |  |  |
| sp\|Q5T0T0\|MARH8\_HUMAN | Q5T0T0 | MARH8\_HUMAN | MARCHF8 | E3 ubiquitin-protein ligase MARCHF8 | 291 | 32.965 | 3.255147e+04 | 1.348158e+06 | 30538.37 | 98200.96 | 243862.02 | 975556.6 | 6.19 | 27.1 | 26.1 | 23.4 |
| sp\|Q92820\|GGH\_HUMAN | Q92820 | GGH\_HUMAN | GGH | Gamma-glutamyl hydrolase | 318 | 35.964 | 8.003654e+03 | 3.616375e+05 | 30422.865 |  | 265048.56 | 66166.08 | 11.64 |  | 28.0 | 33.0 |
| sp\|Q9UHQ4\|BAP29\_HUMAN | Q9UHQ4 | BAP29\_HUMAN | BCAP29 | B-cell receptor-associated protein 29 | 241 | 28.320 | 2.487428e+04 | 8.850358e+05 | 30291.69 | 457946.56 | 78542.39 | 318255.2 | 9.96 | 37.8 | 13.7 | 21.6 |
| sp\|P55211\|CASP9\_HUMAN | P55211 | CASP9\_HUMAN | CASP9 | Caspase-9 | 416 | 46.281 | 1.919665e+03 | 1.116209e+05 | 30070.5 | 81550.36 |  |  | 14.9 | 17.1 |  |  |
| sp\|O75629\|CREG1\_HUMAN | O75629 | CREG1\_HUMAN | CREG1 | Protein CREG1 | 220 | 24.075 | 2.206211e+03 | 6.673144e+04 | 29899.758 | 36831.684 |  |  | 9.55 | 30.5 |  |  |
| sp\|Q96DE0\|NUD16\_HUMAN | Q96DE0 | NUD16\_HUMAN | NUDT16 | U8 snoRNA-decapping enzyme | 195 | 21.273 | 1.149421e+04 | 3.072026e+05 | 29802.514 | 277400.1 |  |  | 16.41 | 34.4 |  |  |
| sp\|Q08174\|PCDH1\_HUMAN | Q08174 | PCDH1\_HUMAN | PCDH1 | Protocadherin-1 | 1060 | 114.743 | 4.308010e+03 | 6.210407e+05 | 29761.543 | 165441.89 | 228487.0 | 197350.27 | 1.13 | 25.7 | 17.0 | 13.6 |
| sp\|O15243\|OBRG\_HUMAN | O15243 | OBRG\_HUMAN | LEPROT | Leptin receptor gene-related protein | 131 | 14.254 | 3.361350e+04 | 6.019599e+05 | 28806.174 |  | 550048.4 | 23105.365 | 9.92 |  | 10.7 | 9.9 |
| sp\|Q6YP21\|KAT3\_HUMAN | Q6YP21 | KAT3\_HUMAN | KYAT3 | Kynurenine--oxoglutarate transaminase 3 | 454 | 51.400 | 2.782033e+03 | 1.796564e+05 | 28538.58 | 151117.78 |  |  | 2.64 | 25.6 |  |  |
| sp\|O00519\|FAAH1\_HUMAN | O00519 | FAAH1\_HUMAN | FAAH | Fatty-acid amide hydrolase 1 | 579 | 63.066 | 7.675175e+02 | 6.081360e+04 | 28229.338 | 15560.155 | 17024.11 |  | 1.9 | 27.8 | 6.0 |  |
| sp\|Q9UGM3\|DMBT1\_HUMAN | Q9UGM3 | DMBT1\_HUMAN | DMBT1 | Deleted in malignant brain tumors 1 protein | 2413 | 260.735 | 4.342521e+02 | 1.422520e+05 | 28201.104 |  | 114050.914 |  | 6.96 |  | 14.1 |  |
| sp\|P01034\|CYTC\_HUMAN | P01034 | CYTC\_HUMAN | CST3 | Cystatin-C | 146 | 15.799 | 4.249294e+03 | 8.434581e+04 | 28018.145 |  | 56327.664 |  | 10.96 |  | 18.5 |  |
| sp\|O43768\|ENSA\_HUMAN | O43768 | ENSA\_HUMAN | ENSA | Alpha-endosulfine | 121 | 13.389 | 1.255597e+04 | 2.112106e+05 | 27765.39 | 35429.492 | 98263.03 | 49752.67 | 11.57 | 57.9 | 57.9 | 41.3 |
| sp\|O15305\|PMM2\_HUMAN | O15305 | PMM2\_HUMAN | PMM2 | Phosphomannomutase 2 | 246 | 28.082 | 2.056912e+04 | 7.257062e+05 | 27747.277 | 595202.8 | 102756.16 |  | 10.57 | 44.7 | 40.2 |  |
| sp\|Q8NB16\|MLKL\_HUMAN | Q8NB16 | MLKL\_HUMAN | MLKL | Mixed lineage kinase domain-like protein | 471 | 54.479 | 9.504904e+02 | 6.505700e+04 | 27727.523 | 37329.473 |  |  | 4.88 | 41.6 |  |  |
| sp\|Q9UKJ5\|CHIC2\_HUMAN | Q9UKJ5 | CHIC2\_HUMAN | CHIC2 | Cysteine-rich hydrophobic domain-containing protein 2 | 165 | 19.254 | 1.338693e+05 | 3.238314e+06 | 27576.145 |  | 493033.94 | 2717704.0 | 7.88 |  | 33.3 | 38.8 |
| sp\|Q9H939\|PPIP2\_HUMAN | Q9H939 | PPIP2\_HUMAN | PSTPIP2 | Proline-serine-threonine phosphatase-interacting protein 2 | 334 | 38.858 | 1.377160e+03 | 6.723294e+04 | 27520.46 | 39712.48 |  |  | 3.29 | 30.2 |  |  |
| sp\|O14737\|PDCD5\_HUMAN | O14737 | PDCD5\_HUMAN | PDCD5 | Programmed cell death protein 5 | 125 | 14.285 | 1.604317e+05 | 2.879305e+06 | 27203.18 | 2683516.0 | 137133.14 | 31452.666 | 10.4 | 85.6 | 57.6 | 23.2 |
| sp\|P12109\|CO6A1\_HUMAN | P12109 | CO6A1\_HUMAN | COL6A1 | Collagen alpha-1(VI) chain | 1028 | 108.529 | 3.666940e+02 | 4.999963e+04 | 27159.457 | 22840.174 |  |  | 2.43 | 43.6 |  |  |
| sp\|P40818\|UBP8\_HUMAN | P40818 | UBP8\_HUMAN | USP8 | Ubiquitin carboxyl-terminal hydrolase 8 | 1118 | 127.523 | 7.384798e+02 | 1.183162e+05 | 27056.383 | 91259.84 |  |  | 1.79 | 37.9 |  |  |
| sp\|P01591\|IGJ\_HUMAN | P01591 | IGJ\_HUMAN | JCHAIN | Immunoglobulin J chain | 159 | 18.099 | 7.024349e+04 | 1.597268e+06 | 27005.516 | 115780.72 | 1057873.2 | 396608.56 | 7.55 | 28.9 | 49.1 | 32.7 |
| sp\|P62857\|RS28\_HUMAN | P62857 | RS28\_HUMAN | RPS28 | Small ribosomal subunit protein eS28 | 69 | 7.590 | 2.828472e+03 | 2.697185e+04 | 26971.852 |  |  |  | 17.39 |  |  |  |
| sp\|P18031\|PTN1\_HUMAN | P18031 | PTN1\_HUMAN | PTPN1 | Tyrosine-protein phosphatase non-receptor type 1 | 435 | 49.967 | 5.087536e+03 | 3.193801e+05 | 26868.781 |  | 261518.08 | 30993.262 | 2.99 |  | 29.2 | 17.7 |
| sp\|Q9NXG2\|THUM1\_HUMAN | Q9NXG2 | THUM1\_HUMAN | THUMPD1 | THUMP domain-containing protein 1 | 353 | 39.315 | 9.384170e+03 | 4.635230e+05 | 26807.258 | 410094.25 | 26621.516 |  | 4.25 | 30.6 | 13.3 |  |
| sp\|Q8N0Z6\|TTC5\_HUMAN | Q8N0Z6 | TTC5\_HUMAN | TTC5 | Tetratricopeptide repeat protein 5 | 440 | 48.928 | 5.537700e+03 | 3.404113e+05 | 26539.613 | 313871.72 |  |  | 5.91 | 33.4 |  |  |
| sp\|O15371\|EIF3D\_HUMAN | O15371 | EIF3D\_HUMAN | EIF3D | Eukaryotic translation initiation factor 3 subunit D | 548 | 60.280 | 3.482234e+02 | 2.637232e+04 | 26372.32 |  |  |  | 2.55 |  |  |  |
| sp\|Q15274\|NADC\_HUMAN | Q15274 | NADC\_HUMAN | QPRT | Nicotinate-nucleotide pyrophosphorylase [carboxylating] | 297 | 30.846 | 1.821006e+03 | 7.057118e+04 | 26253.785 | 44317.4 |  |  | 8.08 | 19.9 |  |  |
| sp\|O75312\|ZPR1\_HUMAN | O75312 | ZPR1\_HUMAN | ZPR1 | Zinc finger protein ZPR1 | 459 | 50.925 | 1.059570e+03 | 6.779188e+04 | 26045.215 | 41746.67 |  |  | 6.32 | 38.1 |  |  |
| sp\|P04279\|SEMG1\_HUMAN | P04279 | SEMG1\_HUMAN | SEMG1 | Semenogelin-1 | 462 | 52.131 | 4.612660e+03 | 3.021097e+05 | 25840.809 | 54914.395 | 144812.22 | 76542.24 | 3.46 | 43.5 | 42.0 | 24.5 |
| sp\|Q9Y365\|STA10\_HUMAN | Q9Y365 | STA10\_HUMAN | STARD10 | START domain-containing protein 10 | 291 | 33.049 | 9.300065e+02 | 3.861548e+04 | 25783.578 |  | 12831.905 |  | 4.12 |  | 11.3 |  |
| sp\|Q04864\|REL\_HUMAN | Q04864 | REL\_HUMAN | REL | Proto-oncogene c-Rel | 619 | 68.520 | 5.067854e+03 | 4.362733e+05 | 25701.781 | 410571.47 |  |  | 4.2 | 24.6 |  |  |
| sp\|P10619\|PPGB\_HUMAN | P10619 | PPGB\_HUMAN | CTSA | Lysosomal protective protein | 480 | 54.466 | 9.987850e+02 | 6.834624e+04 | 25177.584 | 0.0 | 43168.656 |  | 2.71 | 17.7 | 12.9 |  |
| sp\|Q92597\|NDRG1\_HUMAN | Q92597 | NDRG1\_HUMAN | NDRG1 | Protein NDRG1 | 394 | 42.835 | 2.833525e+03 | 1.524906e+05 | 24917.691 | 48840.574 | 55322.984 | 23409.303 | 3.55 | 31.7 | 20.3 | 7.1 |
| sp\|P21796\|VDAC1\_HUMAN | P21796 | VDAC1\_HUMAN | VDAC1 | Voltage-dependent anion-selective channel protein 1 | 283 | 30.773 | 5.244989e+03 | 2.027830e+05 | 24727.082 |  | 140177.77 | 37878.17 | 10.25 |  | 32.9 | 17.7 |
| sp\|O60762\|DPM1\_HUMAN | O60762 | DPM1\_HUMAN | DPM1 | Dolichol-phosphate mannosyltransferase subunit 1 | 260 | 29.634 | 2.544705e+03 | 9.474251e+04 | 24335.826 | 20280.68 | 50126.0 |  | 6.15 | 49.2 | 10.8 |  |
| sp\|Q96BI3\|APH1A\_HUMAN | Q96BI3 | APH1A\_HUMAN | APH1A | Gamma-secretase subunit APH-1A | 265 | 28.996 | 5.366102e+04 | 1.954853e+06 | 24101.234 | 36300.61 | 429888.38 | 1464563.1 | 4.91 | 5.3 | 5.3 | 8.3 |
| sp\|Q99504\|EYA3\_HUMAN | Q99504 | EYA3\_HUMAN | EYA3 | Eyes absent homolog 3 | 573 | 62.663 | 1.559957e+03 | 1.228120e+05 | 24092.48 | 98719.54 |  |  | 3.32 | 9.8 |  |  |
| sp\|Q8IWE2\|NXP20\_HUMAN | Q8IWE2 | NXP20\_HUMAN | FAM114A1 | Protein NOXP20 | 563 | 60.742 | 1.216743e+03 | 9.285494e+04 | 23889.34 | 48360.234 | 20605.37 |  | 5.51 | 22.2 | 15.1 |  |
| sp\|A2RU67\|F234B\_HUMAN | A2RU67 | F234B\_HUMAN | FAM234B | Protein FAM234B | 622 | 67.039 | 4.955356e+03 | 4.173684e+05 | 23696.031 | 57167.105 | 181529.69 | 154975.58 | 2.09 | 13.7 | 19.1 | 12.1 |
| sp\|P02671\|FIBA\_HUMAN | P02671 | FIBA\_HUMAN | FGA | Fibrinogen alpha chain | 866 | 94.973 | 8.638082e+04 | 1.030706e+07 | 23623.422 | 7243572.0 | 2398926.5 | 640934.5 | 3.58 | 50.1 | 42.6 | 26.8 |
| sp\|Q9Y606\|PUS1\_HUMAN | Q9Y606 | PUS1\_HUMAN | PUS1 | Pseudouridylate synthase 1 homolog | 427 | 47.470 | 1.820457e+03 | 1.085718e+05 | 23502.225 | 85069.54 |  |  | 4.92 | 33.5 |  |  |
| sp\|Q15653\|IKBB\_HUMAN | Q15653 | IKBB\_HUMAN | NFKBIB | NF-kappa-B inhibitor beta | 356 | 37.771 | 2.182570e+03 | 1.035723e+05 | 23140.715 | 80431.62 |  |  | 3.65 | 21.1 |  |  |
| sp\|Q9NRK6\|ABCBA\_HUMAN | Q9NRK6 | ABCBA\_HUMAN | ABCB10 | ATP-binding cassette sub-family B member 10, mitochondrial | 738 | 79.148 | 4.423632e+02 | 4.398819e+04 | 23039.615 |  | 20948.578 |  | 2.17 |  | 23.2 |  |
| sp\|Q7Z5U6\|WDR53\_HUMAN | Q7Z5U6 | WDR53\_HUMAN | WDR53 | WD repeat-containing protein 53 | 358 | 38.989 | 1.565621e+03 | 7.669127e+04 | 22727.092 | 53964.176 |  |  | 4.19 | 19.0 |  |  |
| sp\|P01833\|PIGR\_HUMAN | P01833 | PIGR\_HUMAN | PIGR | Polymeric immunoglobulin receptor | 764 | 83.284 | 7.353925e+03 | 7.694811e+05 | 22414.98 | 103394.13 | 491775.28 | 151896.66 | 2.09 | 36.4 | 30.4 | 18.3 |
| sp\|P54709\|AT1B3\_HUMAN | P54709 | AT1B3\_HUMAN | ATP1B3 | Sodium/potassium-transporting ATPase subunit beta-3 | 279 | 31.513 | 2.281522e+03 | 9.032992e+04 | 22410.256 | 67919.66 |  |  | 5.38 | 37.6 |  |  |
| sp\|Q9HD47\|MOG1\_HUMAN | Q9HD47 | MOG1\_HUMAN | RANGRF | Ran guanine nucleotide release factor | 186 | 20.460 | 8.677341e+02 | 2.230537e+04 | 22305.371 |  |  |  | 15.05 |  |  |  |
| sp\|Q7Z422\|SZRD1\_HUMAN | Q7Z422 | SZRD1\_HUMAN | SZRD1 | SUZ domain-containing protein 1 | 152 | 16.720 | 1.036032e+03 | 2.176338e+04 | 21763.383 |  |  |  | 13.82 |  |  |  |
| sp\|O75533\|SF3B1\_HUMAN | O75533 | SF3B1\_HUMAN | SF3B1 | Splicing factor 3B subunit 1 | 1304 | 143.440 | 1.187740e+02 | 2.140469e+04 | 21404.688 |  |  |  | 1.92 |  |  |  |
| sp\|O00264\|PGRC1\_HUMAN | O00264 | PGRC1\_HUMAN | PGRMC1 | Membrane-associated progesterone receptor component 1 | 195 | 21.671 | 2.070835e+04 | 5.638214e+05 | 21283.52 | 116798.7 | 233338.16 | 192401.03 | 10.26 | 39.0 | 34.9 | 22.6 |
| sp\|Q9Y530\|OARD1\_HUMAN | Q9Y530 | OARD1\_HUMAN | OARD1 | ADP-ribose glycohydrolase OARD1 | 152 | 17.025 | 2.983005e+04 | 6.380552e+05 | 21236.734 | 552852.94 | 63965.516 |  | 7.89 | 61.8 | 48.0 |  |
| sp\|Q7L5L3\|GDPD3\_HUMAN | Q7L5L3 | GDPD3\_HUMAN | GDPD3 | Lysophospholipase D GDPD3 | 318 | 36.596 | 7.608116e+02 | 3.498065e+04 | 21216.75 |  | 13763.902 |  | 4.4 |  | 15.1 |  |
| sp\|Q86V21\|AACS\_HUMAN | Q86V21 | AACS\_HUMAN | AACS | Acetoacetyl-CoA synthetase | 672 | 73.920 | 2.252987e+02 | 2.092367e+04 | 20923.668 |  |  |  | 2.68 |  |  |  |
| sp\|P37198\|NUP62\_HUMAN | P37198 | NUP62\_HUMAN | NUP62 | Nuclear pore glycoprotein p62 | 522 | 53.255 | 4.263136e+03 | 2.852376e+05 | 20780.781 | 234674.9 |  | 29781.873 | 2.87 | 18.2 |  | 5.9 |
| sp\|Q5F1R6\|DJC21\_HUMAN | Q5F1R6 | DJC21\_HUMAN | DNAJC21 | DnaJ homolog subfamily C member 21 | 531 | 62.028 | 8.346718e+02 | 6.504601e+04 | 20598.795 | 44447.22 |  |  | 4.52 | 26.7 |  |  |
| sp\|P0DP57\|SLUR2\_HUMAN | P0DP57 | SLUR2\_HUMAN | SLURP2 | Secreted Ly-6/uPAR domain-containing protein 2 | 97 | 10.160 | 3.673486e+03 | 4.689098e+04 | 20546.46 |  | 26344.516 |  | 13.4 |  | 27.8 |  |
| sp\|Q9P003\|CNIH4\_HUMAN | Q9P003 | CNIH4\_HUMAN | CNIH4 | Protein cornichon homolog 4 | 139 | 15.290 | 1.068231e+03 | 2.052058e+04 | 20520.584 |  |  |  | 14.39 |  |  |  |
| sp\|Q96S52\|PIGS\_HUMAN | Q96S52 | PIGS\_HUMAN | PIGS | GPI transamidase component PIG-S | 555 | 61.656 | 3.527122e+03 | 2.732203e+05 | 20244.617 | 39444.953 | 174439.0 | 39091.72 | 3.06 | 26.3 | 15.7 | 10.8 |
| sp\|O43175\|SERA\_HUMAN | O43175 | SERA\_HUMAN | PHGDH | D-3-phosphoglycerate dehydrogenase | 533 | 58.630 | 2.731967e+02 | 2.012392e+04 | 20123.918 |  |  |  | 2.81 |  |  |  |
| sp\|Q9BY41\|HDAC8\_HUMAN | Q9BY41 | HDAC8\_HUMAN | HDAC8 | Histone deacetylase 8 | 377 | 41.470 | 3.845524e+02 | 2.003580e+04 | 20035.799 |  |  |  | 4.24 |  |  |  |
| sp\|Q6P9B6\|MEAK7\_HUMAN | Q6P9B6 | MEAK7\_HUMAN | MEAK7 | MTOR-associated protein MEAK7 | 456 | 50.994 | 4.948755e+03 | 3.170533e+05 | 20024.164 | 297029.1 |  |  | 3.51 | 43.6 |  |  |
| sp\|P31483\|TIA1\_HUMAN | P31483 | TIA1\_HUMAN | TIA1 | Cytotoxic granule associated RNA binding protein TIA1 | 386 | 42.963 | 3.812575e+03 | 2.057928e+05 | 19628.836 | 164152.94 | 22010.994 |  | 3.11 | 39.1 | 11.7 |  |
| sp\|Q8WUA7\|TB22A\_HUMAN | Q8WUA7 | TB22A\_HUMAN | TBC1D22A | TBC1 domain family member 22A | 517 | 59.121 | 7.577384e+02 | 5.628312e+04 | 19579.295 | 36703.83 |  |  | 1.93 | 28.2 |  |  |
| sp\|Q06187\|BTK\_HUMAN | Q06187 | BTK\_HUMAN | BTK | Tyrosine-protein kinase BTK | 659 | 76.281 | 8.160091e+02 | 7.820392e+04 | 19324.352 | 58879.57 |  |  | 3.79 | 42.2 |  |  |
| sp\|Q96AD5\|PLPL2\_HUMAN | Q96AD5 | PLPL2\_HUMAN | PNPLA2 | Patatin-like phospholipase domain-containing protein 2 | 504 | 55.316 | 9.518136e+02 | 6.614847e+04 | 19267.314 | 46881.16 |  |  | 2.58 | 28.4 |  |  |
| sp\|Q658Y4\|F91A1\_HUMAN | Q658Y4 | F91A1\_HUMAN | FAM91A1 | Protein FAM91A1 | 838 | 93.909 | 1.094853e+03 | 1.291755e+05 | 18875.426 | 110300.06 |  |  | 5.13 | 37.4 |  |  |
| sp\|Q15493\|RGN\_HUMAN | Q15493 | RGN\_HUMAN | RGN | Regucalcin | 299 | 33.253 | 9.373997e+02 | 3.916272e+04 | 18483.469 |  | 20679.248 |  | 5.69 |  | 35.5 |  |
| sp\|O43524\|FOXO3\_HUMAN | O43524 | FOXO3\_HUMAN | FOXO3 | Forkhead box protein O3 | 673 | 71.277 | 1.829957e+03 | 1.638730e+05 | 18345.422 | 145527.56 |  |  | 2.82 | 16.8 |  |  |
| sp\|P46977\|STT3A\_HUMAN | P46977 | STT3A\_HUMAN | STT3A | Dolichyl-diphosphooligosaccharide--protein glycosyltransferase subunit STT3A | 705 | 80.530 | 3.881710e+03 | 3.927335e+05 | 18227.719 |  | 113552.07 | 260953.7 | 1.56 |  | 12.1 | 9.6 |
| sp\|Q13308\|PTK7\_HUMAN | Q13308 | PTK7\_HUMAN | PTK7 | Inactive tyrosine-protein kinase 7 | 1070 | 118.392 | 5.686741e+03 | 8.458687e+05 | 18220.004 |  | 145864.12 | 681784.56 | 1.12 |  | 14.0 | 19.2 |
| sp\|Q86UX6\|ST32C\_HUMAN | Q86UX6 | ST32C\_HUMAN | STK32C | Serine/threonine-protein kinase 32C | 486 | 53.460 | 2.705365e+02 | 1.817071e+04 | 18170.713 |  |  |  | 2.47 |  |  |  |
| sp\|P27482\|CALL3\_HUMAN | P27482 | CALL3\_HUMAN | CALML3 | Calmodulin-like protein 3 | 149 | 16.891 | 1.393703e+05 | 2.957623e+06 | 18068.148 |  | 2463636.5 | 475918.62 | 11.41 |  | 66.4 | 59.1 |
| sp\|P06028\|GLPB\_HUMAN | P06028 | GLPB\_HUMAN | GYPB | Glycophorin-B | 91 | 9.782 | 5.397773e+03 | 6.633756e+04 | 17900.363 |  | 23763.973 | 24673.219 | 18.68 |  | 18.7 | 18.7 |
| sp\|Q7Z3D6\|GLUCM\_HUMAN | Q7Z3D6 | GLUCM\_HUMAN | DGLUCY | D-glutamate cyclase, mitochondrial | 616 | 67.760 | 2.089521e+02 | 1.778842e+04 | 17788.42 |  |  |  | 1.95 |  |  |  |
| sp\|Q9H9H4\|VP37B\_HUMAN | Q9H9H4 | VP37B\_HUMAN | VPS37B | Vacuolar protein sorting-associated protein 37B | 285 | 31.307 | 1.012418e+04 | 3.982159e+05 | 17677.293 | 320096.4 | 60442.176 |  | 9.82 | 49.8 | 35.8 |  |
| sp\|P35268\|RL22\_HUMAN | P35268 | RL22\_HUMAN | RPL22 | Large ribosomal subunit protein eL22 | 128 | 14.787 | 4.591717e+03 | 8.530458e+04 | 17202.586 |  | 53050.105 | 15051.893 | 11.72 |  | 10.2 | 27.3 |
| sp\|Q96RF0\|SNX18\_HUMAN | Q96RF0 | SNX18\_HUMAN | SNX18 | Sorting nexin-18 | 628 | 68.894 | 1.046611e+03 | 9.059076e+04 | 17157.854 | 73432.91 |  |  | 2.71 | 40.4 |  |  |
| sp\|Q96BS2\|CHP3\_HUMAN | Q96BS2 | CHP3\_HUMAN | TESC | Calcineurin B homologous protein 3 | 214 | 24.750 | 1.448823e+04 | 4.505134e+05 | 17027.281 | 241245.84 | 75346.0 | 116894.31 | 6.07 | 43.9 | 21.5 | 9.8 |
| sp\|P52434\|RPAB3\_HUMAN | P52434 | RPAB3\_HUMAN | POLR2H | DNA-directed RNA polymerases I, II, and III subunit RPABC3 | 150 | 17.143 | 1.602523e+04 | 3.451504e+05 | 17001.016 | 307361.56 | 15018.578 | 5769.2314 | 8.67 | 14.7 | 29.3 | 8.7 |
| sp\|P22695\|QCR2\_HUMAN | P22695 | QCR2\_HUMAN | UQCRC2 | Cytochrome b-c1 complex subunit 2, mitochondrial | 453 | 48.443 | 1.651706e+03 | 1.005266e+05 | 16920.13 |  | 83606.43 |  | 3.53 |  | 16.8 |  |
| sp\|Q9H0W8\|SMG9\_HUMAN | Q9H0W8 | SMG9\_HUMAN | SMG9 | Nonsense-mediated mRNA decay factor SMG9 | 520 | 57.651 | 5.999141e+04 | 4.345233e+06 | 16771.666 | 4328461.0 |  |  | 3.08 | 58.5 |  |  |
| sp\|Q9NY35\|CLDN1\_HUMAN | Q9NY35 | CLDN1\_HUMAN | CLDND1 | Claudin domain-containing protein 1 | 253 | 28.603 | 4.784614e+04 | 1.719395e+06 | 16637.543 |  | 397617.53 | 1305139.9 | 5.93 |  | 27.3 | 14.2 |
| sp\|Q8NC51\|SERB1\_HUMAN | Q8NC51 | SERB1\_HUMAN | SERBP1 | SERPINE1 mRNA-binding protein 1 | 408 | 44.965 | 2.399058e+03 | 1.355291e+05 | 16515.004 | 68345.48 | 33964.312 | 16704.285 | 3.92 | 30.9 | 31.6 | 19.1 |
| sp\|P40855\|PEX19\_HUMAN | P40855 | PEX19\_HUMAN | PEX19 | Peroxisomal biogenesis factor 19 | 299 | 32.807 | 3.216284e+03 | 1.325678e+05 | 16431.238 | 116136.55 |  |  | 12.71 | 49.5 |  |  |
| sp\|O76094\|SRP72\_HUMAN | O76094 | SRP72\_HUMAN | SRP72 | Signal recognition particle subunit SRP72 | 671 | 74.606 | 6.857534e+02 | 6.427749e+04 | 16307.632 | 47969.855 |  |  | 2.53 | 43.1 |  |  |
| sp\|P60866\|RS20\_HUMAN | P60866 | RS20\_HUMAN | RPS20 | Small ribosomal subunit protein uS10 | 119 | 13.373 | 9.748266e+03 | 1.637847e+05 | 16235.831 | 16446.242 | 131102.64 |  | 9.24 | 62.2 | 9.2 |  |
| sp\|P31751\|AKT2\_HUMAN | P31751 | AKT2\_HUMAN | AKT2 | RAC-beta serine/threonine-protein kinase | 481 | 55.769 | 2.045156e+03 | 1.432968e+05 | 16147.061 | 127149.72 |  |  | 2.29 | 37.0 |  |  |
| sp\|P23246\|SFPQ\_HUMAN | P23246 | SFPQ\_HUMAN | SFPQ | Splicing factor, proline- and glutamine-rich | 707 | 76.149 | 2.048445e+02 | 1.959773e+04 | 16029.815 |  |  | 3567.9128 | 2.12 |  |  | 19.2 |
| sp\|P62899\|RL31\_HUMAN | P62899 | RL31\_HUMAN | RPL31 | Large ribosomal subunit protein eL31 | 125 | 14.463 | 1.153234e+03 | 2.095526e+04 | 15853.477 |  |  | 5101.78 | 11.2 |  |  | 28.0 |
| sp\|P20645\|MPRD\_HUMAN | P20645 | MPRD\_HUMAN | M6PR | Cation-dependent mannose-6-phosphate receptor | 277 | 30.470 | 4.050743e+02 | 1.550688e+04 | 15506.877 |  |  |  | 5.05 |  |  |  |
| sp\|Q6ZNC8\|MBOA1\_HUMAN | Q6ZNC8 | MBOA1\_HUMAN | MBOAT1 | Lysophospholipid acyltransferase 1 | 495 | 56.557 | 4.183460e+03 | 2.972618e+05 | 15265.207 | 240918.34 |  | 41078.27 | 2.22 | 17.4 |  | 6.5 |
| sp\|Q7Z7N9\|T179B\_HUMAN | Q7Z7N9 | T179B\_HUMAN | TMEM179B | Transmembrane protein 179B | 219 | 24.090 | 4.998631e+02 | 1.512882e+04 | 15128.82 |  |  |  | 7.31 |  |  |  |
| sp\|O15194\|CTDSL\_HUMAN | O15194 | CTDSL\_HUMAN | CTDSPL | CTD small phosphatase-like protein | 276 | 31.129 | 1.221301e+03 | 4.776446e+04 | 14872.554 | 32891.91 |  |  | 3.62 | 8.4 |  |  |
| sp\|Q8NBQ5\|DHB11\_HUMAN | Q8NBQ5 | DHB11\_HUMAN | HSD17B11 | Estradiol 17-beta-dehydrogenase 11 | 300 | 32.964 | 7.873978e+03 | 3.261003e+05 | 14734.75 | 230172.42 | 81193.164 |  | 6.0 | 53.7 | 14.0 |  |
| sp\|O15511\|ARPC5\_HUMAN | O15511 | ARPC5\_HUMAN | ARPC5 | Actin-related protein 2/3 complex subunit 5 | 151 | 16.320 | 1.911813e+04 | 3.919969e+05 | 14728.323 | 245894.73 | 131373.88 |  | 7.28 | 46.4 | 20.5 |  |
| sp\|P49411\|EFTU\_HUMAN | P49411 | EFTU\_HUMAN | TUFM | Elongation factor Tu, mitochondrial | 455 | 49.875 | 1.922693e+03 | 1.204786e+05 | 14561.334 |  | 52330.29 | 53586.992 | 2.42 |  | 31.4 | 13.3 |
| sp\|Q6UW02\|CP20A\_HUMAN | Q6UW02 | CP20A\_HUMAN | CYP20A1 | Cytochrome P450 20A1 | 462 | 52.432 | 3.714355e+03 | 2.446791e+05 | 14302.204 | 230376.94 |  |  | 3.03 | 40.3 |  |  |
| sp\|Q5T013\|HYI\_HUMAN | Q5T013 | HYI\_HUMAN | HYI | Putative hydroxypyruvate isomerase | 277 | 30.406 | 5.101131e+03 | 1.948691e+05 | 14299.462 | 180569.61 |  |  | 7.22 | 44.8 |  |  |
| sp\|P07737\|PROF1\_HUMAN | P07737 | PROF1\_HUMAN | PFN1 | Profilin-1 | 140 | 15.054 | 2.676235e+05 | 5.061665e+06 | 14228.931 | 4333517.0 | 531585.1 | 182333.6 | 11.43 | 80.7 | 72.9 | 42.9 |
| sp\|P82979\|SARNP\_HUMAN | P82979 | SARNP\_HUMAN | SARNP | SAP domain-containing ribonucleoprotein | 210 | 23.100 | 4.841891e+02 | 1.405219e+04 | 14052.195 |  |  |  | 9.05 |  |  |  |
| sp\|P01019\|ANGT\_HUMAN | P01019 | ANGT\_HUMAN | AGT | Angiotensinogen | 485 | 53.154 | 2.479272e+03 | 1.655684e+05 | 13861.591 |  | 151706.77 |  | 4.95 |  | 12.2 |  |
| sp\|Q9BYM8\|HOIL1\_HUMAN | Q9BYM8 | HOIL1\_HUMAN | RBCK1 | RanBP-type and C3HC4-type zinc finger-containing protein 1 | 510 | 57.572 | 8.024184e+02 | 5.804026e+04 | 13706.672 | 44333.586 |  |  | 2.16 | 40.0 |  |  |
| sp\|Q96E40\|SACA9\_HUMAN | Q96E40 | SACA9\_HUMAN | SPACA9 | Sperm acrosome-associated protein 9 | 222 | 24.420 | 4.424058e+02 | 1.357324e+04 | 13573.243 |  |  |  | 5.86 |  |  |  |
| sp\|O43379\|WDR62\_HUMAN | O43379 | WDR62\_HUMAN | WDR62 | WD repeat-containing protein 62 | 1518 | 165.954 | 5.429395e+02 | 1.132026e+05 | 13342.851 | 99859.73 |  |  | 1.05 | 36.9 |  |  |
| sp\|Q9NZ09\|UBAP1\_HUMAN | Q9NZ09 | UBAP1\_HUMAN | UBAP1 | Ubiquitin-associated protein 1 | 502 | 55.084 | 7.955406e+02 | 5.505604e+04 | 13124.912 | 41931.125 |  |  | 3.59 | 16.7 |  |  |
| sp\|P18510\|IL1RA\_HUMAN | P18510 | IL1RA\_HUMAN | IL1RN | Interleukin-1 receptor antagonist protein | 177 | 19.470 | 5.261870e+02 | 1.287132e+04 | 12871.324 |  |  |  | 10.73 |  |  |  |
| sp\|Q9BQI3\|E2AK1\_HUMAN | Q9BQI3 | E2AK1\_HUMAN | EIF2AK1 | Eukaryotic translation initiation factor 2-alpha kinase 1 | 630 | 71.106 | 1.541786e+03 | 1.377361e+05 | 12425.162 | 125310.89 |  |  | 1.9 | 26.2 |  |  |
| sp\|Q01081\|U2AF1\_HUMAN | Q01081 | U2AF1\_HUMAN | U2AF1 | Splicing factor U2AF 35 kDa subunit | 240 | 26.400 | 3.656980e+02 | 1.212952e+04 | 12129.523 |  |  |  | 5.42 |  |  |  |
| sp\|Q9Y383\|LC7L2\_HUMAN | Q9Y383 | LC7L2\_HUMAN | LUC7L2 | Putative RNA-binding protein Luc7-like 2 | 392 | 46.514 | 2.750928e+03 | 1.607608e+05 | 11556.507 | 149204.27 |  |  | 3.83 | 32.7 |  |  |
| sp\|Q9Y314\|NOSIP\_HUMAN | Q9Y314 | NOSIP\_HUMAN | NOSIP | Nitric oxide synthase-interacting protein | 301 | 33.110 | 2.760548e+02 | 1.148343e+04 | 11483.432 |  |  |  | 6.98 |  |  |  |
| sp\|O75907\|DGAT1\_HUMAN | O75907 | DGAT1\_HUMAN | DGAT1 | Diacylglycerol O-acyltransferase 1 | 488 | 55.278 | 9.729144e+02 | 6.756847e+04 | 11455.169 | 22115.35 | 8745.43 | 25252.525 | 4.3 | 28.7 | 12.1 | 24.0 |
| sp\|Q05209\|PTN12\_HUMAN | Q05209 | PTN12\_HUMAN | PTPN12 | Tyrosine-protein phosphatase non-receptor type 12 | 780 | 88.106 | 9.661451e+02 | 1.069461e+05 | 11186.275 | 95759.84 |  |  | 2.05 | 32.2 |  |  |
| sp\|Q5JS54\|PSMG4\_HUMAN | Q5JS54 | PSMG4\_HUMAN | PSMG4 | Proteasome assembly chaperone 4 | 123 | 13.775 | 9.260438e+04 | 1.602656e+06 | 10925.111 | 1591730.8 |  |  | 20.33 | 94.3 |  |  |
| sp\|Q7L775\|EPMIP\_HUMAN | Q7L775 | EPMIP\_HUMAN | EPM2AIP1 | EPM2A-interacting protein 1 | 607 | 66.770 | 1.276290e+02 | 1.070651e+04 | 10706.506 |  |  |  | 2.97 |  |  |  |
| sp\|P33993\|MCM7\_HUMAN | P33993 | MCM7\_HUMAN | MCM7 | DNA replication licensing factor MCM7 | 719 | 81.308 | 3.485549e+02 | 3.560587e+04 | 10597.796 | 25008.078 |  |  | 1.81 | 34.1 |  |  |
| sp\|Q9BXK5\|B2L13\_HUMAN | Q9BXK5 | B2L13\_HUMAN | BCL2L13 | Bcl-2-like protein 13 | 485 | 52.723 | 2.906670e+03 | 1.925365e+05 | 10477.6045 | 110871.51 | 71187.36 |  | 1.86 | 29.9 | 20.2 |  |
| sp\|P53597\|SUCA\_HUMAN | P53597 | SUCA\_HUMAN | SUCLG1 | Succinate--CoA ligase [ADP/GDP-forming] subunit alpha, mitochondrial | 346 | 36.250 | 3.484977e+02 | 1.587176e+04 | 10127.734 |  | 5744.0254 |  | 4.34 |  | 21.7 |  |
| sp\|P49862\|KLK7\_HUMAN | P49862 | KLK7\_HUMAN | KLK7 | Kallikrein-7 | 253 | 27.525 | 1.726387e+03 | 5.970114e+04 | 10071.754 | 22701.225 | 26928.162 |  | 6.32 | 65.2 | 25.7 |  |
| sp\|Q14696\|MESD\_HUMAN | Q14696 | MESD\_HUMAN | MESD | LRP chaperone MESD | 234 | 26.077 | 7.244794e+04 | 2.373563e+06 | 9684.8545 | 961616.1 | 1379408.4 | 22853.844 | 5.56 | 55.6 | 41.5 | 24.8 |
| sp\|P19320\|VCAM1\_HUMAN | P19320 | VCAM1\_HUMAN | VCAM1 | Vascular cell adhesion protein 1 | 739 | 81.290 | 9.475013e+01 | 9.676852e+03 | 9676.852 |  |  |  | 2.03 |  |  |  |
| sp\|Q86WQ0\|NR2CA\_HUMAN | Q86WQ0 | NR2CA\_HUMAN | NR2C2AP | Nuclear receptor 2C2-associated protein | 139 | 15.876 | 8.792282e+04 | 1.753718e+06 | 9362.918 | 1744355.4 |  |  | 10.07 | 46.0 |  |  |
| sp\|Q5K4L6\|S27A3\_HUMAN | Q5K4L6 | S27A3\_HUMAN | SLC27A3 | Long-chain fatty acid transport protein 3 | 683 | 75.130 | 9.255496e+01 | 8.736354e+03 | 8736.3545 |  |  |  | 3.51 |  |  |  |
| sp\|P35080\|PROF2\_HUMAN | P35080 | PROF2\_HUMAN | PFN2 | Profilin-2 | 140 | 15.046 | 1.252639e+03 | 2.367905e+04 | 8428.034 |  | 15251.012 |  | 10.0 |  | 10.0 |  |
| sp\|Q9NRD1\|FBX6\_HUMAN | Q9NRD1 | FBX6\_HUMAN | FBXO6 | F-box only protein 6 | 293 | 33.933 | 3.704036e+03 | 1.579118e+05 | 8346.142 | 124275.71 | 25289.977 |  | 4.1 | 22.2 | 22.9 |  |
| sp\|Q16643\|DREB\_HUMAN | Q16643 | DREB\_HUMAN | DBN1 | Drebrin | 649 | 71.429 | 7.123097e+02 | 6.392350e+04 | 8164.295 | 55759.207 |  |  | 2.77 | 28.8 |  |  |
| sp\|P51178\|PLCD1\_HUMAN | P51178 | PLCD1\_HUMAN | PLCD1 | 1-phosphatidylinositol 4,5-bisphosphate phosphodiesterase delta-1 | 756 | 83.160 | 7.680177e+01 | 8.024221e+03 | 8024.2207 |  |  |  | 1.98 |  |  |  |
| sp\|P13010\|XRCC5\_HUMAN | P13010 | XRCC5\_HUMAN | XRCC5 | X-ray repair cross-complementing protein 5 | 732 | 82.705 | 9.831131e+02 | 1.021533e+05 | 7864.7236 | 65617.875 | 16284.114 | 12386.591 | 1.91 | 32.5 | 9.4 | 4.2 |
| sp\|Q5JTD0\|TJAP1\_HUMAN | Q5JTD0 | TJAP1\_HUMAN | TJAP1 | Tight junction-associated protein 1 | 557 | 61.270 | 1.004890e+02 | 7.735416e+03 | 7735.416 |  |  |  | 2.51 |  |  |  |
| sp\|Q86Y37\|CACL1\_HUMAN | Q86Y37 | CACL1\_HUMAN | CACUL1 | CDK2-associated and cullin domain-containing protein 1 | 369 | 41.064 | 1.260409e+03 | 6.502644e+04 | 7655.5146 | 57370.93 |  |  | 5.15 | 40.4 |  |  |
| sp\|Q9BZH6\|WDR11\_HUMAN | Q9BZH6 | WDR11\_HUMAN | WDR11 | WD repeat-containing protein 11 | 1224 | 136.685 | 2.808518e+03 | 4.822977e+05 | 6816.2974 | 475481.44 |  |  | 2.21 | 34.6 |  |  |
| sp\|P31040\|SDHA\_HUMAN | P31040 | SDHA\_HUMAN | SDHA | Succinate dehydrogenase [ubiquinone] flavoprotein subunit, mitochondrial | 664 | 72.692 | 3.376171e+02 | 3.083388e+04 | 6789.6074 | 24044.275 |  |  | 1.96 | 19.6 |  |  |
| sp\|Q9UBI9\|HDC\_HUMAN | Q9UBI9 | HDC\_HUMAN | HECA | Headcase protein homolog | 543 | 59.730 | 8.315197e+01 | 6.239966e+03 | 6239.966 |  |  |  | 5.71 |  |  |  |
| sp\|O15372\|EIF3H\_HUMAN | O15372 | EIF3H\_HUMAN | EIF3H | Eukaryotic translation initiation factor 3 subunit H | 352 | 39.930 | 2.602770e+03 | 1.305726e+05 | 5519.38 | 104303.98 | 17206.541 | 3542.739 | 5.4 | 33.0 | 15.1 | 13.9 |
| sp\|P49069\|CAMLG\_HUMAN | P49069 | CAMLG\_HUMAN | CAMLG | Guided entry of tail-anchored proteins factor CAMLG | 296 | 32.560 | 1.217778e+02 | 4.981610e+03 | 4981.6104 |  |  |  | 7.43 |  |  |  |
| sp\|Q9P2E9\|RRBP1\_HUMAN | Q9P2E9 | RRBP1\_HUMAN | RRBP1 | Ribosome-binding protein 1 | 1410 | 152.456 | 2.922815e+02 | 5.598388e+04 | 4487.5293 | 51496.35 |  |  | 1.28 | 45.3 |  |  |
| sp\|O14981\|BTAF1\_HUMAN | O14981 | BTAF1\_HUMAN | BTAF1 | TATA-binding protein-associated factor 172 | 1849 | 206.887 | 7.732420e+01 | 2.009860e+04 | 3817.4717 | 16281.128 |  |  | 0.92 | 31.9 |  |  |
| sp\|Q6ZTZ1\|MSD1\_HUMAN | Q6ZTZ1 | MSD1\_HUMAN | MSANTD1 | Myb/SANT-like DNA-binding domain-containing protein 1 | 278 | 30.580 | 0.000000e+00 | 0.000000e+00 | 0.0 |  |  |  | 14.03 |  |  |  |
| sp\|Q15398\|DLGP5\_HUMAN | Q15398 | DLGP5\_HUMAN | DLGAP5 | Disks large-associated protein 5 | 846 | 93.060 | 0.000000e+00 | 0.000000e+00 | 0.0 |  |  |  | 2.25 |  |  |  |
| sp\|P36957\|ODO2\_HUMAN | P36957 | ODO2\_HUMAN | DLST | Dihydrolipoyllysine-residue succinyltransferase component of 2-oxoglutarate dehydrogenase complex, mitochondrial | 453 | 48.755 | 1.537946e+03 | 9.420572e+04 | 0.0 | 34097.277 | 60108.445 |  | 3.09 | 29.6 | 17.2 |  |
| sp\|P51452\|DUS3\_HUMAN | P51452 | DUS3\_HUMAN | DUSP3 | Dual specificity protein phosphatase 3 | 185 | 20.478 | 1.963179e+03 | 5.050852e+04 | 0.0 | 50508.516 |  |  | 7.57 | 33.0 |  |  |
| sp\|O43684\|BUB3\_HUMAN | O43684 | BUB3\_HUMAN | BUB3 | Mitotic checkpoint protein BUB3 | 328 | 37.155 | 7.641859e+02 | 3.567249e+04 | 0.0 | 35672.492 |  |  | 4.27 | 29.6 |  |  |
| sp\|Q96JG6\|VPS50\_HUMAN | Q96JG6 | VPS50\_HUMAN | VPS50 | Syndetin | 964 | 111.174 | 1.681938e+02 | 2.349256e+04 | 0.0 | 23492.564 |  |  | 1.35 | 24.7 |  |  |
| sp\|P61599\|NAA20\_HUMAN | P61599 | NAA20\_HUMAN | NAA20 | N-alpha-acetyltransferase 20 | 178 | 20.368 | 3.849859e+03 | 9.851682e+04 | 0.0 | 98516.82 |  |  | 11.8 | 18.0 |  |  |
| sp\|A0A0B4J2D5\|GAL3B\_HUMAN | A0A0B4J2D5 | GAL3B\_HUMAN | GATD3B | Putative glutamine amidotransferase-like class 1 domain-containing protein 3B, mitochondrial | 268 | 28.142 | 5.167129e+03 | 1.826928e+05 |  | 182692.81 |  |  |  | 58.2 |  |  |
| sp\|A6NED2\|RCCD1\_HUMAN | A6NED2 | RCCD1\_HUMAN | RCCD1 | RCC1 domain-containing protein 1 | 376 | 40.079 | 2.747628e+03 | 1.383541e+05 |  | 138354.08 |  |  |  | 19.1 |  |  |
| sp\|A7E2V4\|ZSWM8\_HUMAN | A7E2V4 | ZSWM8\_HUMAN | ZSWIM8 | Zinc finger SWIM domain-containing protein 8 | 1837 | 197.297 | 1.721605e+02 | 4.267476e+04 |  | 42674.76 |  |  |  | 30.8 |  |  |
| sp\|O00308\|WWP2\_HUMAN | O00308 | WWP2\_HUMAN | WWP2 | NEDD4-like E3 ubiquitin-protein ligase WWP2 | 870 | 98.912 | 5.100511e+03 | 6.338402e+05 |  | 355423.28 | 261399.66 | 17017.287 |  | 28.3 | 24.7 | 11.5 |
| sp\|O00471\|EXOC5\_HUMAN | O00471 | EXOC5\_HUMAN | EXOC5 | Exocyst complex component 5 | 708 | 81.853 | 2.564497e+03 | 2.637266e+05 |  | 263726.6 |  |  |  | 44.6 |  |  |
| sp\|O00534\|VMA5A\_HUMAN | O00534 | VMA5A\_HUMAN | VWA5A | von Willebrand factor A domain-containing protein 5A | 786 | 86.489 | 1.336818e+03 | 1.452614e+05 |  | 145261.44 |  |  |  | 42.1 |  |  |
| sp\|O00571\|DDX3X\_HUMAN | O00571 | DDX3X\_HUMAN | DDX3X | ATP-dependent RNA helicase DDX3X | 662 | 73.243 | 7.456300e+03 | 6.861304e+05 |  | 329515.28 | 239240.69 | 117374.42 |  | 34.3 | 23.4 | 13.0 |
| sp\|O00746\|NDKM\_HUMAN | O00746 | NDKM\_HUMAN | NME4 | Nucleoside diphosphate kinase, mitochondrial | 187 | 20.659 | 4.498907e+03 | 1.167706e+05 |  | 116770.586 |  |  |  | 59.9 |  |  |
| sp\|O14530\|TXND9\_HUMAN | O14530 | TXND9\_HUMAN | TXNDC9 | Thioredoxin domain-containing protein 9 | 226 | 26.534 | 4.687492e+03 | 1.562645e+05 |  | 156264.52 |  |  |  | 43.4 |  |  |
| sp\|O14933\|UB2L6\_HUMAN | O14933 | UB2L6\_HUMAN | UBE2L6 | Ubiquitin/ISG15-conjugating enzyme E2 L6 | 153 | 17.769 | 2.115268e+04 | 4.722213e+05 |  | 472221.3 |  |  |  | 47.7 |  |  |
| sp\|O14983\|AT2A1\_HUMAN | O14983 | AT2A1\_HUMAN | ATP2A1 | Sarcoplasmic/endoplasmic reticulum calcium ATPase 1 | 1001 | 110.252 | 1.407741e+04 | 1.949964e+06 |  | 579887.7 | 800394.7 | 569681.1 |  | 34.9 | 21.3 | 17.8 |
| sp\|O15027\|SC16A\_HUMAN | O15027 | SC16A\_HUMAN | SEC16A | Protein transport protein Sec16A | 2357 | 251.894 | 1.132196e+02 | 3.583080e+04 |  | 35830.797 |  |  |  | 29.5 |  |  |
| sp\|O15116\|LSM1\_HUMAN | O15116 | LSM1\_HUMAN | LSM1 | U6 snRNA-associated Sm-like protein LSm1 | 133 | 15.179 | 3.440521e+04 | 6.561219e+05 |  | 656121.9 |  |  |  | 54.1 |  |  |
| sp\|O15212\|PFD6\_HUMAN | O15212 | PFD6\_HUMAN | PFDN6 | Prefoldin subunit 6 | 129 | 14.583 | 5.851691e+04 | 1.072125e+06 |  | 984777.9 | 87347.13 |  |  | 82.2 | 34.1 |  |
| sp\|O15231\|ZN185\_HUMAN | O15231 | ZN185\_HUMAN | ZNF185 | Zinc finger protein 185 | 689 | 73.525 | 1.294486e+03 | 1.195775e+05 |  | 51744.258 | 52818.895 | 15014.326 |  | 40.9 | 35.8 | 19.6 |
| sp\|O15270\|SPTC2\_HUMAN | O15270 | SPTC2\_HUMAN | SPTLC2 | Serine palmitoyltransferase 2 | 562 | 62.924 | 4.740369e+03 | 3.747534e+05 |  | 285477.28 | 89276.13 |  |  | 36.5 | 24.2 |  |
| sp\|O15335\|CHAD\_HUMAN | O15335 | CHAD\_HUMAN | CHAD | Chondroadherin | 359 | 40.476 | 8.663380e+02 | 4.405570e+04 |  | 31978.543 | 12077.157 |  |  | 38.7 | 29.2 |  |
| sp\|O15355\|PPM1G\_HUMAN | O15355 | PPM1G\_HUMAN | PPM1G | Protein phosphatase 1G | 546 | 59.272 | 8.482470e+02 | 6.316683e+04 |  | 63166.83 |  |  |  | 34.4 |  |  |
| sp\|O43414\|ERI3\_HUMAN | O43414 | ERI3\_HUMAN | ERI3 | ERI1 exoribonuclease 3 | 337 | 37.238 | 9.466853e+03 | 4.429035e+05 |  | 260124.38 | 115403.28 | 67375.84 |  | 50.1 | 22.8 | 9.8 |
| sp\|O43617\|TPPC3\_HUMAN | O43617 | TPPC3\_HUMAN | TRAPPC3 | Trafficking protein particle complex subunit 3 | 180 | 20.274 | 1.667107e+04 | 4.246393e+05 |  | 424639.3 |  |  |  | 30.0 |  |  |
| sp\|O43815\|STRN\_HUMAN | O43815 | STRN\_HUMAN | STRN | Striatin | 780 | 86.132 | 1.067539e+03 | 1.155221e+05 |  | 115522.15 |  |  |  | 34.1 |  |  |
| sp\|O60361\|NDK8\_HUMAN | O60361 | NDK8\_HUMAN | NME2P1 | Putative nucleoside diphosphate kinase | 137 | 15.529 | 1.839043e+06 | 3.588002e+07 |  | 33331278.0 | 2548737.2 |  |  | 93.4 | 52.6 |  |
| sp\|O60733\|PLPL9\_HUMAN | O60733 | PLPL9\_HUMAN | PLA2G6 | 85/88 kDa calcium-independent phospholipase A2 | 806 | 89.903 | 3.448276e+03 | 3.894874e+05 |  | 155171.12 | 234316.28 |  |  | 29.0 | 19.9 |  |
| sp\|O60841\|IF2P\_HUMAN | O60841 | IF2P\_HUMAN | EIF5B | Eukaryotic translation initiation factor 5B | 1220 | 138.827 | 4.253578e+02 | 7.419002e+04 |  | 74190.02 |  |  |  | 23.4 |  |  |
| sp\|O60888\|CUTA\_HUMAN | O60888 | CUTA\_HUMAN | CUTA | Protein CutA | 179 | 19.116 | 1.220493e+05 | 2.931227e+06 |  | 2722343.2 | 199342.67 | 9541.089 |  | 48.6 | 23.5 | 29.6 |
| sp\|O60925\|PFD1\_HUMAN | O60925 | PFD1\_HUMAN | PFDN1 | Prefoldin subunit 1 | 122 | 14.210 | 2.172285e+04 | 3.878181e+05 |  | 245753.11 | 142065.03 |  |  | 47.5 | 43.4 |  |
| sp\|O75391\|SPAG7\_HUMAN | O75391 | SPAG7\_HUMAN | SPAG7 | Sperm-associated antigen 7 | 227 | 26.034 | 2.071067e+03 | 6.774109e+04 |  | 67741.09 |  |  |  | 59.5 |  |  |
| sp\|O75531\|BAF\_HUMAN | O75531 | BAF\_HUMAN | BANF1 | Barrier-to-autointegration factor | 89 | 10.059 | 6.004701e+04 | 7.588629e+05 |  | 758862.94 |  |  |  | 55.1 |  |  |
| sp\|O75717\|WDHD1\_HUMAN | O75717 | WDHD1\_HUMAN | WDHD1 | WD repeat and HMG-box DNA-binding protein 1 | 1129 | 125.967 | 8.952671e+02 | 1.416859e+05 |  | 141685.88 |  |  |  | 21.7 |  |  |
| sp\|O75821\|EIF3G\_HUMAN | O75821 | EIF3G\_HUMAN | EIF3G | Eukaryotic translation initiation factor 3 subunit G | 320 | 35.611 | 2.591318e+03 | 1.159370e+05 |  | 45091.84 | 46586.37 | 24258.768 |  | 26.2 | 15.9 | 10.6 |
| sp\|O94851\|MICA2\_HUMAN | O94851 | MICA2\_HUMAN | MICAL2 | [F-actin]-monooxygenase MICAL2 | 1957 | 219.061 | 5.554671e+02 | 1.528764e+05 |  | 152876.44 |  |  |  | 33.8 |  |  |
| sp\|O95256\|I18RA\_HUMAN | O95256 | I18RA\_HUMAN | IL18RAP | Interleukin-18 receptor accessory protein | 599 | 68.310 | 1.410860e+04 | 1.210836e+06 |  | 1210836.0 |  |  |  | 36.6 |  |  |
| sp\|O95777\|LSM8\_HUMAN | O95777 | LSM8\_HUMAN | LSM8 | U6 snRNA-associated Sm-like protein LSm8 | 96 | 10.403 | 2.218870e+04 | 2.900064e+05 |  | 290006.4 |  |  |  | 64.6 |  |  |
| sp\|O95881\|TXD12\_HUMAN | O95881 | TXD12\_HUMAN | TXNDC12 | Thioredoxin domain-containing protein 12 | 172 | 19.206 | 1.370998e+04 | 3.308194e+05 |  | 227966.33 | 102853.09 |  |  | 49.4 | 30.2 |  |
| sp\|P00450\|CERU\_HUMAN | P00450 | CERU\_HUMAN | CP | Ceruloplasmin | 1065 | 122.205 | 7.835853e+02 | 1.203074e+05 |  | 66948.336 | 53359.086 |  |  | 25.6 | 11.8 |  |
| sp\|P00488\|F13A\_HUMAN | P00488 | F13A\_HUMAN | F13A1 | Coagulation factor XIII A chain | 732 | 83.268 | 2.663885e+03 | 2.786832e+05 |  | 278683.2 |  |  |  | 37.8 |  |  |
| sp\|P01008\|ANT3\_HUMAN | P01008 | ANT3\_HUMAN | SERPINC1 | Antithrombin-III | 464 | 52.602 | 5.725215e+03 | 3.783652e+05 |  | 157825.6 | 220539.6 |  |  | 39.4 | 28.4 |  |
| sp\|P02100\|HBE\_HUMAN | P02100 | HBE\_HUMAN | HBE1 | Hemoglobin subunit epsilon | 147 | 16.203 | 2.906687e+07 | 5.917126e+08 |  | 423136512.0 | 140009136.0 | 28566998.0 |  | 100.0 | 72.1 | 48.3 |
| sp\|P02747\|C1QC\_HUMAN | P02747 | C1QC\_HUMAN | C1QC | Complement C1q subcomponent subunit C | 245 | 25.774 | 4.741012e+04 | 1.535218e+06 |  | 976062.75 | 141200.23 | 417954.8 |  | 46.5 | 39.2 | 35.5 |
| sp\|P02751\|FINC\_HUMAN | P02751 | FINC\_HUMAN | FN1 | Fibronectin | 2477 | 272.320 | 4.897334e+03 | 1.675546e+06 |  | 954305.4 | 654872.9 | 66367.945 |  | 28.4 | 22.2 | 6.7 |
| sp\|P02766\|TTHY\_HUMAN | P02766 | TTHY\_HUMAN | TTR | Transthyretin | 147 | 15.887 | 1.541592e+04 | 3.077008e+05 |  | 206945.94 | 100754.81 |  |  | 54.4 | 8.8 |  |
| sp\|P02771\|FETA\_HUMAN | P02771 | FETA\_HUMAN | AFP | Alpha-fetoprotein | 609 | 68.678 | 8.604344e+03 | 7.424250e+05 |  | 724079.0 | 18345.965 |  |  | 26.9 | 16.6 |  |
| sp\|P02790\|HEMO\_HUMAN | P02790 | HEMO\_HUMAN | HPX | Hemopexin | 462 | 51.676 | 1.592326e+04 | 1.033804e+06 |  | 834075.6 | 199727.92 |  |  | 39.4 | 29.7 |  |
| sp\|P02792\|FRIL\_HUMAN | P02792 | FRIL\_HUMAN | FTL | Ferritin light chain | 175 | 20.020 | 2.986221e+04 | 7.511092e+05 |  | 751109.25 |  |  |  | 45.7 |  |  |
| sp\|P04003\|C4BPA\_HUMAN | P04003 | C4BPA\_HUMAN | C4BPA | C4b-binding protein alpha chain | 597 | 67.033 | 4.988476e+02 | 4.201203e+04 |  | 42012.03 |  |  |  | 57.3 |  |  |
| sp\|P04114\|APOB\_HUMAN | P04114 | APOB\_HUMAN | APOB | Apolipoprotein B-100 | 4563 | 515.605 | 2.314410e+03 | 1.499252e+06 |  | 1294167.5 | 160651.1 | 44433.273 |  | 31.9 | 18.3 | 7.2 |
| sp\|P04196\|HRG\_HUMAN | P04196 | HRG\_HUMAN | HRG | Histidine-rich glycoprotein | 525 | 59.578 | 2.612186e+03 | 1.955272e+05 |  | 195527.17 |  |  |  | 29.1 |  |  |
| sp\|P04275\|VWF\_HUMAN | P04275 | VWF\_HUMAN | VWF | von Willebrand factor | 2813 | 309.265 | 2.106612e+04 | 8.185260e+06 |  | 0.0 | 8185260.5 |  |  | 29.2 | 14.3 |  |
| sp\|P05164\|PERM\_HUMAN | P05164 | PERM\_HUMAN | MPO | Myeloperoxidase | 745 | 83.869 | 6.242214e+03 | 6.577445e+05 |  | 423378.25 | 175624.47 | 58741.805 |  | 49.3 | 35.3 | 19.2 |
| sp\|P05546\|HEP2\_HUMAN | P05546 | HEP2\_HUMAN | SERPIND1 | Heparin cofactor 2 | 499 | 57.071 | 7.650461e+02 | 5.485551e+04 |  | 54855.51 |  |  |  | 35.5 |  |  |
| sp\|P07864\|LDHC\_HUMAN | P07864 | LDHC\_HUMAN | LDHC | L-lactate dehydrogenase C chain | 332 | 36.311 | 7.023068e+03 | 3.203924e+05 |  | 320392.4 |  |  |  | 31.3 |  |  |
| sp\|P07951\|TPM2\_HUMAN | P07951 | TPM2\_HUMAN | TPM2 | Tropomyosin beta chain | 284 | 32.851 | 1.411078e+06 | 5.823940e+07 |  | 25898348.0 | 28322820.0 | 4018235.2 |  | 58.8 | 63.0 | 42.6 |
| sp\|P08246\|ELNE\_HUMAN | P08246 | ELNE\_HUMAN | ELANE | Neutrophil elastase | 267 | 28.518 | 3.963386e+03 | 1.420046e+05 |  | 96834.86 | 45169.78 |  |  | 50.2 | 24.0 |  |
| sp\|P09497\|CLCB\_HUMAN | P09497 | CLCB\_HUMAN | CLTB | Clathrin light chain B | 229 | 25.190 | 3.577188e+04 | 1.132106e+06 |  | 509727.94 | 416583.7 | 205794.44 |  | 22.3 | 27.1 | 24.5 |
| sp\|P09874\|PARP1\_HUMAN | P09874 | PARP1\_HUMAN | PARP1 | Poly [ADP-ribose] polymerase 1 | 1014 | 113.084 | 1.465143e+03 | 2.081605e+05 |  | 208160.55 |  |  |  | 38.0 |  |  |
| sp\|P0C024\|NUDT7\_HUMAN | P0C024 | NUDT7\_HUMAN | NUDT7 | Peroxisomal coenzyme A diphosphatase NUDT7 | 238 | 26.942 | 7.898591e+03 | 2.673601e+05 |  | 267360.12 |  |  |  | 58.0 |  |  |
| sp\|P0C0L4\|CO4A\_HUMAN | P0C0L4 | CO4A\_HUMAN | C4A | Complement C4-A | 1744 | 192.785 | 1.545330e+04 | 3.742930e+06 |  | 647622.06 | 3095308.0 |  |  | 28.4 | 20.4 |  |
| sp\|P0CAP2\|GRL1A\_HUMAN | P0CAP2 | GRL1A\_HUMAN | POLR2M | DNA-directed RNA polymerase II subunit GRINL1A | 368 | 41.740 | 2.625272e+03 | 1.376715e+05 |  | 137671.48 |  |  |  | 51.4 |  |  |
| sp\|P11215\|ITAM\_HUMAN | P11215 | ITAM\_HUMAN | ITGAM | Integrin alpha-M | 1152 | 127.179 | 1.107325e+03 | 1.769325e+05 |  | 161057.06 | 15875.449 |  |  | 31.3 | 20.1 |  |
| sp\|P11678\|PERE\_HUMAN | P11678 | PERE\_HUMAN | EPX | Eosinophil peroxidase | 715 | 81.040 | 1.306520e+03 | 1.330248e+05 |  | 122247.45 | 10777.318 |  |  | 47.0 | 17.1 |  |
| sp\|P12111\|CO6A3\_HUMAN | P12111 | CO6A3\_HUMAN | COL6A3 | Collagen alpha-3(VI) chain | 3177 | 343.669 | 2.341601e+03 | 1.011045e+06 |  | 1008780.06 | 2264.614 |  |  | 32.4 | 18.0 |  |
| sp\|P12270\|TPR\_HUMAN | P12270 | TPR\_HUMAN | TPR | Nucleoprotein TPR | 2363 | 267.293 | 3.867687e+02 | 1.298841e+05 |  | 129884.12 |  |  |  | 23.9 |  |  |
| sp\|P14621\|ACYP2\_HUMAN | P14621 | ACYP2\_HUMAN | ACYP2 | Acylphosphatase-2 | 99 | 11.140 | 4.626900e+04 | 6.475785e+05 |  | 624975.44 | 14704.591 | 7898.4863 |  | 85.9 | 28.3 | 13.1 |
| sp\|P15259\|PGAM2\_HUMAN | P15259 | PGAM2\_HUMAN | PGAM2 | Phosphoglycerate mutase 2 | 253 | 28.766 | 6.518279e+05 | 2.355752e+07 |  | 22710078.0 | 589749.6 | 257691.8 |  | 58.5 | 35.6 | 13.8 |
| sp\|P16401\|H15\_HUMAN | P16401 | H15\_HUMAN | H1-5 | Histone H1.5 | 226 | 22.580 | 9.624906e+03 | 2.730471e+05 |  | 152437.12 | 120610.0 |  |  | 39.4 | 34.1 |  |
| sp\|P18621\|RL17\_HUMAN | P18621 | RL17\_HUMAN | RPL17 | 60S ribosomal protein L17 | 184 | 21.397 | 8.970959e+03 | 2.411620e+05 |  | 211619.95 | 29542.068 |  |  | 51.1 | 41.8 |  |
| sp\|P19338\|NUCL\_HUMAN | P19338 | NUCL\_HUMAN | NCL | Nucleolin | 710 | 76.614 | 1.001109e+03 | 9.636219e+04 |  | 96362.19 |  |  |  | 35.9 |  |  |
| sp\|P19652\|A1AG2\_HUMAN | P19652 | A1AG2\_HUMAN | ORM2 | Alpha-1-acid glycoprotein 2 | 201 | 23.603 | 1.303634e+04 | 3.865805e+05 |  | 219336.03 | 167244.48 |  |  | 39.8 | 21.4 |  |
| sp\|P19823\|ITIH2\_HUMAN | P19823 | ITIH2\_HUMAN | ITIH2 | Inter-alpha-trypsin inhibitor heavy chain H2 | 946 | 106.463 | 9.317991e+02 | 1.246345e+05 |  | 124634.46 |  |  |  | 29.8 |  |  |
| sp\|P23497\|SP100\_HUMAN | P23497 | SP100\_HUMAN | SP100 | Nuclear autoantigen Sp-100 | 879 | 100.417 | 2.864682e+03 | 3.614105e+05 |  | 361410.5 |  |  |  | 35.6 |  |  |
| sp\|P25398\|RS12\_HUMAN | P25398 | RS12\_HUMAN | RPS12 | 40S ribosomal protein S12 | 132 | 14.515 | 1.138087e+04 | 2.075437e+05 |  | 71250.55 | 136293.19 |  |  | 61.4 | 28.8 |  |
| sp\|P27449\|VATL\_HUMAN | P27449 | VATL\_HUMAN | ATP6V0C | V-type proton ATPase 16 kDa proteolipid subunit c | 155 | 15.736 | 2.955300e+04 | 5.842693e+05 |  | 182256.4 | 207870.62 | 194142.3 |  | 11.6 | 11.6 | 11.6 |
| sp\|P27694\|RFA1\_HUMAN | P27694 | RFA1\_HUMAN | RPA1 | Replication protein A 70 kDa DNA-binding subunit | 616 | 68.138 | 2.084009e+03 | 1.784046e+05 |  | 178404.64 |  |  |  | 41.7 |  |  |
| sp\|P29074\|PTN4\_HUMAN | P29074 | PTN4\_HUMAN | PTPN4 | Tyrosine-protein phosphatase non-receptor type 4 | 926 | 105.911 | 1.269799e+03 | 1.689636e+05 |  | 168963.62 |  |  |  | 24.5 |  |  |
| sp\|P30048\|PRDX3\_HUMAN | P30048 | PRDX3\_HUMAN | PRDX3 | Thioredoxin-dependent peroxide reductase, mitochondrial | 256 | 27.693 | 6.571899e+03 | 2.286536e+05 |  | 228653.6 |  |  |  | 41.8 |  |  |
| sp\|P31689\|DNJA1\_HUMAN | P31689 | DNJA1\_HUMAN | DNAJA1 | DnaJ homolog subfamily A member 1 | 397 | 44.868 | 2.001507e+03 | 1.128265e+05 |  | 112826.47 |  |  |  | 36.0 |  |  |
| sp\|P35749\|MYH11\_HUMAN | P35749 | MYH11\_HUMAN | MYH11 | Myosin-11 | 1972 | 227.339 | 3.388298e+04 | 9.677713e+06 |  | 9188798.0 | 488914.62 |  |  | 36.9 | 20.7 |  |
| sp\|P42336\|PK3CA\_HUMAN | P42336 | PK3CA\_HUMAN | PIK3CA | Phosphatidylinositol 4,5-bisphosphate 3-kinase catalytic subunit alpha isoform | 1068 | 124.284 | 3.100866e+02 | 4.841894e+04 |  | 48418.945 |  |  |  | 40.5 |  |  |
| sp\|P43652\|AFAM\_HUMAN | P43652 | AFAM\_HUMAN | AFM | Afamin | 599 | 69.069 | 2.429337e+03 | 2.108085e+05 |  | 210808.47 |  |  |  | 35.2 |  |  |
| sp\|P46013\|KI67\_HUMAN | P46013 | KI67\_HUMAN | MKI67 | Proliferation marker protein Ki-67 | 3256 | 358.694 | 3.302949e+02 | 1.488480e+05 |  | 148848.05 |  |  |  | 51.8 |  |  |
| sp\|P46527\|CDN1B\_HUMAN | P46527 | CDN1B\_HUMAN | CDKN1B | Cyclin-dependent kinase inhibitor 1B | 198 | 22.073 | 2.721004e+03 | 7.545842e+04 |  | 75458.42 |  |  |  | 44.4 |  |  |
| sp\|P47813\|IF1AX\_HUMAN | P47813 | IF1AX\_HUMAN | EIF1AX | Eukaryotic translation initiation factor 1A, X-chromosomal | 144 | 16.460 | 3.047399e+05 | 6.301969e+06 |  | 6301969.0 |  |  |  | 59.0 |  |  |
| sp\|P48735\|IDHP\_HUMAN | P48735 | IDHP\_HUMAN | IDH2 | Isocitrate dehydrogenase [NADP], mitochondrial | 452 | 50.909 | 1.577277e+04 | 1.008834e+06 |  | 357351.03 | 493944.34 | 157538.33 |  | 39.8 | 36.9 | 25.7 |
| sp\|P49750\|YLPM1\_HUMAN | P49750 | YLPM1\_HUMAN | YLPM1 | YLP motif-containing protein 1 | 2146 | 241.645 | 1.576506e+03 | 4.786198e+05 |  | 457162.47 | 21457.285 |  |  | 27.0 | 19.1 |  |
| sp\|P49789\|FHIT\_HUMAN | P49789 | FHIT\_HUMAN | FHIT | Bis(5'-adenosyl)-triphosphatase | 147 | 16.858 | 3.880176e+04 | 8.218161e+05 |  | 745006.9 | 76809.22 |  |  | 87.1 | 15.6 |  |
| sp\|P49796\|RGS3\_HUMAN | P49796 | RGS3\_HUMAN | RGS3 | Regulator of G-protein signaling 3 | 1198 | 132.336 | 2.560254e+02 | 4.256751e+04 |  | 42567.51 |  |  |  | 39.1 |  |  |
| sp\|P52272\|HNRPM\_HUMAN | P52272 | HNRPM\_HUMAN | HNRNPM | Heterogeneous nuclear ribonucleoprotein M | 730 | 77.516 | 1.016809e+03 | 9.902571e+04 |  | 96328.3 |  | 2697.4072 |  | 58.9 |  | 26.3 |
| sp\|P53675\|CLH2\_HUMAN | P53675 | CLH2\_HUMAN | CLTCL1 | Clathrin heavy chain 2 | 1640 | 187.030 | 4.166648e+04 | 9.790733e+06 |  | 9790733.0 |  |  |  | 36.2 |  |  |
| sp\|P54819\|KAD2\_HUMAN | P54819 | KAD2\_HUMAN | AK2 | Adenylate kinase 2, mitochondrial | 239 | 26.478 | 3.359898e+03 | 1.117708e+05 |  | 111770.81 |  |  |  | 74.1 |  |  |
| sp\|P55196\|AFAD\_HUMAN | P55196 | AFAD\_HUMAN | AFDN | Afadin | 1824 | 206.804 | 1.210350e+02 | 3.144756e+04 |  | 31447.555 |  |  |  | 38.2 |  |  |
| sp\|P60660\|MYL6\_HUMAN | P60660 | MYL6\_HUMAN | MYL6 | Myosin light polypeptide 6 | 151 | 16.930 | 1.164479e+05 | 2.476885e+06 |  | 2272034.8 | 134754.52 | 70095.72 |  | 41.7 | 30.5 | 29.8 |
| sp\|P61247\|RS3A\_HUMAN | P61247 | RS3A\_HUMAN | RPS3A | 40S ribosomal protein S3a | 264 | 29.945 | 4.646581e+03 | 1.748135e+05 |  | 143390.95 | 14505.574 | 16916.996 |  | 54.2 | 19.3 | 20.8 |
| sp\|P61326\|MGN\_HUMAN | P61326 | MGN\_HUMAN | MAGOH | Protein mago nashi homolog | 146 | 17.164 | 4.833182e+04 | 1.042242e+06 |  | 973091.0 | 69151.48 |  |  | 67.8 | 19.9 |  |
| sp\|P61457\|PHS\_HUMAN | P61457 | PHS\_HUMAN | PCBD1 | Pterin-4-alpha-carbinolamine dehydratase | 104 | 12.000 | 9.499960e+03 | 1.432255e+05 |  | 143225.45 |  |  |  | 86.5 |  |  |
| sp\|P61956\|SUMO2\_HUMAN | P61956 | SUMO2\_HUMAN | SUMO2 | Small ubiquitin-related modifier 2 | 95 | 10.871 | 4.972725e+04 | 6.791741e+05 |  | 404030.38 | 275143.72 |  |  | 43.2 | 32.6 |  |
| sp\|P61966\|AP1S1\_HUMAN | P61966 | AP1S1\_HUMAN | AP1S1 | AP-1 complex subunit sigma-1A | 158 | 18.733 | 1.318014e+04 | 3.102020e+05 |  | 194600.52 | 115601.53 |  |  | 50.0 | 43.7 |  |
| sp\|P62341\|SELT\_HUMAN | P62341 | SELT\_HUMAN | SELENOT | Thioredoxin reductase-like selenoprotein T | 195 | 22.324 | 2.495223e+03 | 6.998395e+04 |  | 69983.95 |  |  |  | 42.1 |  |  |
| sp\|P62877\|RBX1\_HUMAN | P62877 | RBX1\_HUMAN | RBX1 | E3 ubiquitin-protein ligase RBX1 | 108 | 12.274 | 1.398204e+05 | 2.156125e+06 |  | 737077.5 | 1345633.0 | 73414.13 |  | 31.5 | 27.8 | 18.5 |
| sp\|P63165\|SUMO1\_HUMAN | P63165 | SUMO1\_HUMAN | SUMO1 | Small ubiquitin-related modifier 1 | 101 | 11.557 | 4.625474e+04 | 6.716121e+05 |  | 574062.8 | 97549.26 |  |  | 57.4 | 51.5 |  |
| sp\|P82970\|HMGN5\_HUMAN | P82970 | HMGN5\_HUMAN | HMGN5 | High mobility group nucleosome-binding domain-containing protein 5 | 282 | 31.525 | 3.513142e+03 | 1.391451e+05 |  | 139145.1 |  |  |  | 40.8 |  |  |
| sp\|P84095\|RHOG\_HUMAN | P84095 | RHOG\_HUMAN | RHOG | Rho-related GTP-binding protein RhoG | 191 | 21.309 | 5.009772e+03 | 1.341214e+05 |  | 134121.44 |  |  |  | 46.1 |  |  |
| sp\|P98160\|PGBM\_HUMAN | P98160 | PGBM\_HUMAN | HSPG2 | Basement membrane-specific heparan sulfate proteoglycan core protein | 4391 | 468.830 | 6.519489e+01 | 3.840132e+04 |  | 38401.316 |  |  |  | 26.3 |  |  |
| sp\|P98175\|RBM10\_HUMAN | P98175 | RBM10\_HUMAN | RBM10 | RNA-binding protein 10 | 930 | 103.533 | 3.667125e+02 | 4.770036e+04 |  | 47700.363 |  |  |  | 39.0 |  |  |
| sp\|P99999\|CYC\_HUMAN | P99999 | CYC\_HUMAN | CYCS | Cytochrome c | 105 | 11.749 | 1.263742e+04 | 1.865419e+05 |  | 186541.89 |  |  |  | 64.8 |  |  |
| sp\|Q01804\|OTUD4\_HUMAN | Q01804 | OTUD4\_HUMAN | OTUD4 | OTU domain-containing protein 4 | 1114 | 124.045 | 1.128122e+03 | 1.758136e+05 |  | 175813.6 |  |  |  | 32.0 |  |  |
| sp\|Q02241\|KIF23\_HUMAN | Q02241 | KIF23\_HUMAN | KIF23 | Kinesin-like protein KIF23 | 960 | 110.059 | 9.418557e+02 | 1.302348e+05 |  | 98139.42 | 32095.395 |  |  | 33.4 | 22.0 |  |
| sp\|Q02487\|DSC2\_HUMAN | Q02487 | DSC2\_HUMAN | DSC2 | Desmocollin-2 | 901 | 99.962 | 2.522489e+04 | 3.167972e+06 |  | 32574.902 | 1402416.9 | 1732979.9 |  | 26.5 | 20.6 | 5.4 |
| sp\|Q03393\|PTPS\_HUMAN | Q03393 | PTPS\_HUMAN | PTS | 6-pyruvoyl tetrahydrobiopterin synthase | 145 | 16.386 | 4.261152e+03 | 8.772373e+04 |  | 87723.734 |  |  |  | 62.1 |  |  |
| sp\|Q05655\|KPCD\_HUMAN | Q05655 | KPCD\_HUMAN | PRKCD | Protein kinase C delta type | 676 | 77.505 | 6.854790e+02 | 6.674843e+04 |  | 66748.43 |  |  |  | 31.7 |  |  |
| sp\|Q08J23\|NSUN2\_HUMAN | Q08J23 | NSUN2\_HUMAN | NSUN2 | RNA cytosine C(5)-methyltransferase NSUN2 | 767 | 86.471 | 2.124445e+03 | 2.307986e+05 |  | 230798.6 |  |  |  | 34.9 |  |  |
| sp\|Q12800\|TFCP2\_HUMAN | Q12800 | TFCP2\_HUMAN | TFCP2 | Alpha-globin transcription factor CP2 | 502 | 57.256 | 1.829383e+03 | 1.315961e+05 |  | 131596.06 |  |  |  | 34.3 |  |  |
| sp\|Q12830\|BPTF\_HUMAN | Q12830 | BPTF\_HUMAN | BPTF | Nucleosome-remodeling factor subunit BPTF | 3046 | 338.262 | 2.738016e+03 | 1.163607e+06 |  | 1163607.1 |  |  |  | 25.0 |  |  |
| sp\|Q13105\|ZBT17\_HUMAN | Q13105 | ZBT17\_HUMAN | ZBTB17 | Zinc finger and BTB domain-containing protein 17 | 803 | 87.928 | 3.995627e+02 | 4.413969e+04 |  | 44139.688 |  |  |  | 45.5 |  |  |
| sp\|Q13113\|PDZ1I\_HUMAN | Q13113 | PDZ1I\_HUMAN | PDZK1IP1 | PDZK1-interacting protein 1 | 114 | 12.227 | 1.365609e+05 | 2.097796e+06 |  | 192904.69 | 363149.0 | 1541742.8 |  | 34.2 | 32.5 | 32.5 |
| sp\|Q13303\|KCAB2\_HUMAN | Q13303 | KCAB2\_HUMAN | KCNAB2 | Voltage-gated potassium channel subunit beta-2 | 367 | 41.000 | 2.374817e+03 | 1.223295e+05 |  | 122329.47 |  |  |  | 43.1 |  |  |
| sp\|Q13310\|PABP4\_HUMAN | Q13310 | PABP4\_HUMAN | PABPC4 | Polyadenylate-binding protein 4 | 644 | 70.783 | 3.403753e+03 | 3.026942e+05 |  | 302694.22 |  |  |  | 39.6 |  |  |
| sp\|Q13347\|EIF3I\_HUMAN | Q13347 | EIF3I\_HUMAN | EIF3I | Eukaryotic translation initiation factor 3 subunit I | 325 | 36.502 | 1.874648e+03 | 8.597131e+04 |  | 85971.31 |  |  |  | 45.2 |  |  |
| sp\|Q13478\|IL18R\_HUMAN | Q13478 | IL18R\_HUMAN | IL18R1 | Interleukin-18 receptor 1 | 541 | 62.304 | 2.818566e+03 | 2.206283e+05 |  | 220628.31 |  |  |  | 28.3 |  |  |
| sp\|Q13614\|MTMR2\_HUMAN | Q13614 | MTMR2\_HUMAN | MTMR2 | Myotubularin-related protein 2 | 643 | 73.381 | 9.319010e+02 | 8.591531e+04 |  | 85915.31 |  |  |  | 33.6 |  |  |
| sp\|Q13627\|DYR1A\_HUMAN | Q13627 | DYR1A\_HUMAN | DYRK1A | Dual specificity tyrosine-phosphorylation-regulated kinase 1A | 763 | 85.584 | 2.598726e+03 | 2.794282e+05 |  | 173178.05 | 106250.11 |  |  | 26.9 | 10.6 |  |
| sp\|Q14011\|CIRBP\_HUMAN | Q14011 | CIRBP\_HUMAN | CIRBP | Cold-inducible RNA-binding protein | 172 | 18.648 | 9.064379e+03 | 2.123672e+05 |  | 187691.28 | 24675.932 |  |  | 48.8 | 18.0 |  |
| sp\|Q14019\|COTL1\_HUMAN | Q14019 | COTL1\_HUMAN | COTL1 | Coactosin-like protein | 142 | 15.945 | 1.708149e+04 | 3.421900e+05 |  | 342190.03 |  |  |  | 56.3 |  |  |
| sp\|Q14157\|UBP2L\_HUMAN | Q14157 | UBP2L\_HUMAN | UBAP2L | Ubiquitin-associated protein 2-like | 1087 | 114.535 | 2.128292e+03 | 3.062574e+05 |  | 253714.69 | 52542.734 |  |  | 20.6 | 14.7 |  |
| sp\|Q14324\|MYPC2\_HUMAN | Q14324 | MYPC2\_HUMAN | MYBPC2 | Myosin-binding protein C, fast-type | 1141 | 128.072 | 3.163976e+04 | 5.091016e+06 |  | 223628.84 | 4867387.5 |  |  | 44.4 | 18.7 |  |
| sp\|Q14353\|GAMT\_HUMAN | Q14353 | GAMT\_HUMAN | GAMT | Guanidinoacetate N-methyltransferase | 236 | 26.318 | 4.840536e+03 | 1.600529e+05 |  | 160052.88 |  |  |  | 64.4 |  |  |
| sp\|Q14624\|ITIH4\_HUMAN | Q14624 | ITIH4\_HUMAN | ITIH4 | Inter-alpha-trypsin inhibitor heavy chain H4 | 930 | 103.357 | 2.620929e+03 | 3.403395e+05 |  | 120085.79 | 196753.47 | 23500.236 |  | 30.3 | 23.7 | 7.8 |
| sp\|Q14667\|K0100\_HUMAN | Q14667 | K0100\_HUMAN | KIAA0100 | Protein KIAA0100 | 2235 | 253.700 | 5.266996e+02 | 1.678806e+05 |  | 39469.89 | 63605.164 | 64805.59 |  | 20.9 | 11.6 | 12.1 |
| sp\|Q14919\|NC2A\_HUMAN | Q14919 | NC2A\_HUMAN | DRAP1 | Dr1-associated corepressor | 205 | 22.350 | 7.980638e+03 | 2.240951e+05 |  | 206826.55 | 17268.527 |  |  | 21.5 | 21.5 |  |
| sp\|Q15048\|LRC14\_HUMAN | Q15048 | LRC14\_HUMAN | LRRC14 | Leucine-rich repeat-containing protein 14 | 493 | 54.513 | 3.660204e+03 | 2.506816e+05 |  | 250681.62 |  |  |  | 41.0 |  |  |
| sp\|Q15637\|SF01\_HUMAN | Q15637 | SF01\_HUMAN | SF1 | Splicing factor 1 | 639 | 68.330 | 3.277666e+03 | 2.813800e+05 |  | 267538.66 | 13841.352 |  |  | 33.3 | 14.1 |  |
| sp\|Q15642\|CIP4\_HUMAN | Q15642 | CIP4\_HUMAN | TRIP10 | Cdc42-interacting protein 4 | 601 | 68.352 | 1.868623e+03 | 1.604686e+05 |  | 160468.62 |  |  |  | 51.1 |  |  |
| sp\|Q15836\|VAMP3\_HUMAN | Q15836 | VAMP3\_HUMAN | VAMP3 | Vesicle-associated membrane protein 3 | 100 | 11.309 | 1.421721e+06 | 2.020020e+07 |  | 1335315.1 | 3703231.0 | 15161654.0 |  | 54.0 | 54.0 | 66.0 |
| sp\|Q16864\|VATF\_HUMAN | Q16864 | VATF\_HUMAN | ATP6V1F | V-type proton ATPase subunit F | 119 | 13.370 | 1.548527e+04 | 2.601161e+05 |  | 260116.11 |  |  |  | 51.3 |  |  |
| sp\|Q17R31\|TATD3\_HUMAN | Q17R31 | TATD3\_HUMAN | TATDN3 | Putative deoxyribonuclease TATDN3 | 274 | 30.333 | 6.745531e+03 | 2.570684e+05 |  | 257068.39 |  |  |  | 62.4 |  |  |
| sp\|Q567U6\|CCD93\_HUMAN | Q567U6 | CCD93\_HUMAN | CCDC93 | Coiled-coil domain-containing protein 93 | 631 | 73.198 | 2.135380e+03 | 1.963774e+05 |  | 196377.4 |  |  |  | 28.1 |  |  |
| sp\|Q5T0D9\|TPRGL\_HUMAN | Q5T0D9 | TPRGL\_HUMAN | TPRG1L | Tumor protein p63-regulated gene 1-like protein | 272 | 30.212 | 2.043384e+03 | 7.756156e+04 |  | 77561.56 |  |  |  | 19.5 |  |  |
| sp\|Q5T2E6\|ARMD3\_HUMAN | Q5T2E6 | ARMD3\_HUMAN | ARMH3 | Armadillo-like helical domain-containing protein 3 | 689 | 78.710 | 2.103726e+03 | 2.080349e+05 |  | 208034.89 |  |  |  | 26.0 |  |  |
| sp\|Q5T6J7\|GNTK\_HUMAN | Q5T6J7 | GNTK\_HUMAN | IDNK | Probable gluconokinase | 187 | 20.578 | 1.047380e+04 | 2.707850e+05 |  | 270785.0 |  |  |  | 50.8 |  |  |
| sp\|Q5TA50\|CPTP\_HUMAN | Q5TA50 | CPTP\_HUMAN | CPTP | Ceramide-1-phosphate transfer protein | 214 | 24.365 | 9.154200e+03 | 2.802231e+05 |  | 280223.12 |  |  |  | 36.9 |  |  |
| sp\|Q5TAX3\|TUT4\_HUMAN | Q5TAX3 | TUT4\_HUMAN | TUT4 | Terminal uridylyltransferase 4 | 1644 | 185.166 | 1.478313e+05 | 3.439100e+07 |  | 34390996.0 |  |  |  | 27.1 |  |  |
| sp\|Q5VZ89\|DEN4C\_HUMAN | Q5VZ89 | DEN4C\_HUMAN | DENND4C | DENN domain-containing protein 4C | 1909 | 212.711 | 5.770706e+02 | 1.542184e+05 |  | 154218.36 |  |  |  | 28.1 |  |  |
| sp\|Q6GMV3\|PTRD1\_HUMAN | Q6GMV3 | PTRD1\_HUMAN | PTRHD1 | Putative peptidyl-tRNA hydrolase PTRHD1 | 140 | 15.805 | 7.349915e+04 | 1.459466e+06 |  | 448239.5 | 451468.47 | 559758.0 |  | 68.6 | 32.9 | 21.4 |
| sp\|Q6IQ22\|RAB12\_HUMAN | Q6IQ22 | RAB12\_HUMAN | RAB12 | Ras-related protein Rab-12 | 244 | 27.248 | 6.341650e+04 | 2.170971e+06 |  | 457566.25 | 1713404.8 |  |  | 47.1 | 30.3 |  |
| sp\|Q6IQ49\|SDE2\_HUMAN | Q6IQ49 | SDE2\_HUMAN | SDE2 | Replication stress response regulator SDE2 | 451 | 49.742 | 6.553393e+02 | 4.095497e+04 |  | 40954.97 |  |  |  | 52.1 |  |  |
| sp\|Q6NXE6\|ARMC6\_HUMAN | Q6NXE6 | ARMC6\_HUMAN | ARMC6 | Armadillo repeat-containing protein 6 | 501 | 54.142 | 2.572383e+03 | 1.749795e+05 |  | 174979.47 |  |  |  | 37.7 |  |  |
| sp\|Q6P1A2\|MBOA5\_HUMAN | Q6P1A2 | MBOA5\_HUMAN | LPCAT3 | Lysophospholipid acyltransferase 5 | 487 | 56.035 | 3.001418e+04 | 2.113017e+06 |  | 529654.25 | 388399.03 | 1194964.0 |  | 14.0 | 12.9 | 10.7 |
| sp\|Q6UX04\|CWC27\_HUMAN | Q6UX04 | CWC27\_HUMAN | CWC27 | Spliceosome-associated protein CWC27 homolog | 472 | 53.847 | 4.420512e+03 | 2.990551e+05 |  | 299055.1 |  |  |  | 30.1 |  |  |
| sp\|Q6ZT21\|TMPPE\_HUMAN | Q6ZT21 | TMPPE\_HUMAN | TMPPE | Transmembrane protein with metallophosphoesterase domain | 453 | 49.453 | 1.127262e+03 | 7.003815e+04 |  | 55466.355 | 14571.797 |  |  | 7.1 | 6.8 |  |
| sp\|Q6ZW49\|PAXI1\_HUMAN | Q6ZW49 | PAXI1\_HUMAN | PAXIP1 | PAX-interacting protein 1 | 1069 | 121.341 | 4.412307e+03 | 6.726521e+05 |  | 672652.06 |  |  |  | 27.5 |  |  |
| sp\|Q712K3\|UB2R2\_HUMAN | Q712K3 | UB2R2\_HUMAN | UBE2R2 | Ubiquitin-conjugating enzyme E2 R2 | 238 | 27.166 | 1.479605e+04 | 5.049970e+05 |  | 395006.12 | 109990.86 |  |  | 36.6 | 10.9 |  |
| sp\|Q7L266\|ASGL1\_HUMAN | Q7L266 | ASGL1\_HUMAN | ASRGL1 | Isoaspartyl peptidase/L-asparaginase | 308 | 32.055 | 1.095871e+04 | 4.413389e+05 |  | 441338.9 |  |  |  | 47.4 |  |  |
| sp\|Q7Z406\|MYH14\_HUMAN | Q7Z406 | MYH14\_HUMAN | MYH14 | Myosin-14 | 1995 | 227.871 | 2.442817e+04 | 6.993543e+06 |  | 5560906.0 | 395352.7 | 1037284.0 |  | 41.4 | 23.7 | 16.0 |
| sp\|Q7Z434\|MAVS\_HUMAN | Q7Z434 | MAVS\_HUMAN | MAVS | Mitochondrial antiviral-signaling protein | 540 | 56.528 | 1.659757e+03 | 1.178760e+05 |  | 85681.375 | 32194.617 |  |  | 42.4 | 6.5 |  |
| sp\|Q7Z460\|CLAP1\_HUMAN | Q7Z460 | CLAP1\_HUMAN | CLASP1 | CLIP-associating protein 1 | 1538 | 169.451 | 6.475772e+02 | 1.378646e+05 |  | 137864.62 |  |  |  | 39.3 |  |  |
| sp\|Q7Z7E8\|UB2Q1\_HUMAN | Q7Z7E8 | UB2Q1\_HUMAN | UBE2Q1 | Ubiquitin-conjugating enzyme E2 Q1 | 422 | 46.127 | 3.982015e+02 | 2.307678e+04 |  | 23076.781 |  |  |  | 31.0 |  |  |
| sp\|Q8IWY9\|CDAN1\_HUMAN | Q8IWY9 | CDAN1\_HUMAN | CDAN1 | Codanin-1 | 1227 | 134.120 | 2.509102e+02 | 4.227942e+04 |  | 42279.42 |  |  |  | 21.9 |  |  |
| sp\|Q8IYB5\|SMAP1\_HUMAN | Q8IYB5 | SMAP1\_HUMAN | SMAP1 | Stromal membrane-associated protein 1 | 467 | 50.386 | 2.958891e+03 | 1.873079e+05 |  | 187307.86 |  |  |  | 22.5 |  |  |
| sp\|Q8IYI6\|EXOC8\_HUMAN | Q8IYI6 | EXOC8\_HUMAN | EXOC8 | Exocyst complex component 8 | 725 | 81.799 | 1.429739e+03 | 1.469339e+05 |  | 146933.88 |  |  |  | 30.8 |  |  |
| sp\|Q8N0Y7\|PGAM4\_HUMAN | Q8N0Y7 | PGAM4\_HUMAN | PGAM4 | Probable phosphoglycerate mutase 4 | 254 | 28.777 | 6.284392e+05 | 2.272092e+07 |  | 22720922.0 |  |  |  | 57.5 |  |  |
| sp\|Q8N163\|CCAR2\_HUMAN | Q8N163 | CCAR2\_HUMAN | CCAR2 | Cell cycle and apoptosis regulator protein 2 | 923 | 102.902 | 2.206397e+02 | 2.852493e+04 |  | 28524.93 |  |  |  | 38.4 |  |  |
| sp\|Q8N3F0\|MTURN\_HUMAN | Q8N3F0 | MTURN\_HUMAN | MTURN | Maturin | 131 | 14.925 | 4.743046e+04 | 8.893829e+05 |  | 889382.94 |  |  |  | 29.8 |  |  |
| sp\|Q8N6H7\|ARFG2\_HUMAN | Q8N6H7 | ARFG2\_HUMAN | ARFGAP2 | ADP-ribosylation factor GTPase-activating protein 2 | 521 | 56.720 | 5.528608e+03 | 3.939754e+05 |  | 393975.44 |  |  |  | 47.0 |  |  |
| sp\|Q8N7R7\|CCYL1\_HUMAN | Q8N7R7 | CCYL1\_HUMAN | CCNYL1 | Cyclin-Y-like protein 1 | 359 | 40.705 | 2.154082e+04 | 1.101608e+06 |  | 85724.34 |  | 1015883.94 |  | 32.0 |  | 14.8 |
| sp\|Q8ND56\|LS14A\_HUMAN | Q8ND56 | LS14A\_HUMAN | LSM14A | Protein LSM14 homolog A | 463 | 50.530 | 8.712571e+03 | 5.531116e+05 |  | 461181.62 | 91930.01 |  |  | 32.6 | 33.0 |  |
| sp\|Q8NFH8\|REPS2\_HUMAN | Q8NFH8 | REPS2\_HUMAN | REPS2 | RalBP1-associated Eps domain-containing protein 2 | 660 | 71.534 | 9.094056e+02 | 8.173109e+04 |  | 81731.09 |  |  |  | 53.0 |  |  |
| sp\|Q8NFU3\|TSTD1\_HUMAN | Q8NFU3 | TSTD1\_HUMAN | TSTD1 | Thiosulfate:glutathione sulfurtransferase | 115 | 12.530 | 1.854484e+04 | 2.919385e+05 |  | 82323.805 | 209614.66 |  |  | 81.7 | 26.1 |  |
| sp\|Q8NI08\|NCOA7\_HUMAN | Q8NI08 | NCOA7\_HUMAN | NCOA7 | Nuclear receptor coactivator 7 | 942 | 106.162 | 3.646170e+02 | 4.863212e+04 |  | 48632.12 |  |  |  | 29.5 |  |  |
| sp\|Q8TAG9\|EXOC6\_HUMAN | Q8TAG9 | EXOC6\_HUMAN | EXOC6 | Exocyst complex component 6 | 804 | 93.722 | 1.477973e+03 | 1.740304e+05 |  | 174030.38 |  |  |  | 23.9 |  |  |
| sp\|Q8TCT1\|PHOP1\_HUMAN | Q8TCT1 | PHOP1\_HUMAN | PHOSPHO1 | Phosphoethanolamine/phosphocholine phosphatase | 267 | 29.713 | 1.175204e+04 | 4.387093e+05 |  | 378726.25 | 33534.555 | 26448.508 |  | 44.9 | 13.5 | 18.7 |
| sp\|Q8TEU7\|RPGF6\_HUMAN | Q8TEU7 | RPGF6\_HUMAN | RAPGEF6 | Rap guanine nucleotide exchange factor 6 | 1601 | 179.423 | 5.886832e+03 | 1.327018e+06 |  | 1327018.2 |  |  |  | 33.7 |  |  |
| sp\|Q8WU90\|ZC3HF\_HUMAN | Q8WU90 | ZC3HF\_HUMAN | ZC3H15 | Zinc finger CCCH domain-containing protein 15 | 426 | 48.602 | 1.883223e+03 | 1.149934e+05 |  | 114993.45 |  |  |  | 29.8 |  |  |
| sp\|Q8WVL7\|ANR49\_HUMAN | Q8WVL7 | ANR49\_HUMAN | ANKRD49 | Ankyrin repeat domain-containing protein 49 | 239 | 27.290 | 6.619070e+03 | 2.269434e+05 |  | 226943.44 |  |  |  | 15.9 |  |  |
| sp\|Q8WXB1\|MT21A\_HUMAN | Q8WXB1 | MT21A\_HUMAN | METTL21A | Protein N-lysine methyltransferase METTL21A | 218 | 24.600 | 6.020077e+03 | 1.860605e+05 |  | 178809.52 | 7251.021 |  |  | 29.4 | 18.8 |  |
| sp\|Q8WZ42\|TITIN\_HUMAN | Q8WZ42 | TITIN\_HUMAN | TTN | Titin | 34350 | 3816.030 | 1.020155e+04 | 4.890972e+07 |  | 17180152.0 | 22018874.0 | 9710691.0 |  | 33.7 | 16.7 | 8.5 |
| sp\|Q92817\|EVPL\_HUMAN | Q92817 | EVPL\_HUMAN | EVPL | Envoplakin | 2033 | 231.604 | 2.007295e+03 | 5.840828e+05 |  | 205761.56 | 246556.55 | 131764.67 |  | 30.9 | 30.8 | 15.0 |
| sp\|Q92974\|ARHG2\_HUMAN | Q92974 | ARHG2\_HUMAN | ARHGEF2 | Rho guanine nucleotide exchange factor 2 | 986 | 111.543 | 4.387514e+02 | 6.148625e+04 |  | 61486.25 |  |  |  | 36.1 |  |  |
| sp\|Q969H8\|MYDGF\_HUMAN | Q969H8 | MYDGF\_HUMAN | MYDGF | Myeloid-derived growth factor | 173 | 18.795 | 6.567409e+04 | 1.550792e+06 |  | 322704.06 | 1228087.9 |  |  | 29.5 | 29.5 |  |
| sp\|Q96A33\|CCD47\_HUMAN | Q96A33 | CCD47\_HUMAN | CCDC47 | PAT complex subunit CCDC47 | 483 | 55.874 | 4.969885e+03 | 3.488778e+05 |  | 146543.6 | 202334.16 |  |  | 24.6 | 26.1 |  |
| sp\|Q96B23\|CR025\_HUMAN | Q96B23 | CR025\_HUMAN | C18orf25 | Uncharacterized protein C18orf25 | 404 | 43.395 | 8.186028e+03 | 4.463033e+05 |  | 424630.75 |  | 21672.52 |  | 40.3 |  | 5.7 |
| sp\|Q96BM9\|ARL8A\_HUMAN | Q96BM9 | ARL8A\_HUMAN | ARL8A | ADP-ribosylation factor-like protein 8A | 186 | 21.416 | 9.944298e+03 | 2.675652e+05 |  | 267565.2 |  |  |  | 40.3 |  |  |
| sp\|Q96BY7\|ATG2B\_HUMAN | Q96BY7 | ATG2B\_HUMAN | ATG2B | Autophagy-related protein 2 homolog B | 2078 | 232.763 | 1.366986e+03 | 3.997560e+05 |  | 237283.56 | 162472.47 |  |  | 31.5 | 15.0 |  |
| sp\|Q96CN4\|EVI5L\_HUMAN | Q96CN4 | EVI5L\_HUMAN | EVI5L | EVI5-like protein | 794 | 91.376 | 8.156014e+02 | 9.363264e+04 |  | 93632.64 |  |  |  | 19.4 |  |  |
| sp\|Q96DX4\|RSPRY\_HUMAN | Q96DX4 | RSPRY\_HUMAN | RSPRY1 | RING finger and SPRY domain-containing protein 1 | 576 | 64.180 | 6.858723e+03 | 5.530445e+05 |  | 191478.5 | 238469.9 | 123096.14 |  | 24.0 | 8.3 | 13.7 |
| sp\|Q96E09\|PBIR1\_HUMAN | Q96E09 | PBIR1\_HUMAN | PABIR1 | PPP2R1A-PPP2R2A-interacting phosphatase regulator 1 | 287 | 30.529 | 8.733912e+03 | 3.349952e+05 |  | 334995.16 |  |  |  | 46.0 |  |  |
| sp\|Q96F24\|NRBF2\_HUMAN | Q96F24 | NRBF2\_HUMAN | NRBF2 | Nuclear receptor-binding factor 2 | 287 | 32.378 | 2.471880e+03 | 1.005529e+05 |  | 51183.97 | 49368.926 |  |  | 22.6 | 18.8 |  |
| sp\|Q96GS4\|BORC6\_HUMAN | Q96GS4 | BORC6\_HUMAN | BORCS6 | BLOC-1-related complex subunit 6 | 357 | 37.226 | 9.961734e+02 | 4.659062e+04 |  | 46590.617 |  |  |  | 50.7 |  |  |
| sp\|Q96HA7\|TONSL\_HUMAN | Q96HA7 | TONSL\_HUMAN | TONSL | Tonsoku-like protein | 1378 | 150.929 | 2.972449e+04 | 5.636431e+06 |  | 5636431.0 |  |  |  | 30.9 |  |  |
| sp\|Q96JB1\|DYH8\_HUMAN | Q96JB1 | DYH8\_HUMAN | DNAH8 | Dynein axonemal heavy chain 8 | 4490 | 514.664 | 1.944035e+02 | 1.257029e+05 |  | 125702.85 |  |  |  | 28.4 |  |  |
| sp\|Q96K17\|BT3L4\_HUMAN | Q96K17 | BT3L4\_HUMAN | BTF3L4 | Transcription factor BTF3 homolog 4 | 158 | 17.271 | 9.575481e+03 | 2.077759e+05 |  | 207775.92 |  |  |  | 61.4 |  |  |
| sp\|Q96MW1\|CCD43\_HUMAN | Q96MW1 | CCD43\_HUMAN | CCDC43 | Coiled-coil domain-containing protein 43 | 224 | 25.248 | 1.654707e+03 | 5.248863e+04 |  | 52488.633 |  |  |  | 38.4 |  |  |
| sp\|Q96N06\|SPT33\_HUMAN | Q96N06 | SPT33\_HUMAN | SPATA33 | Spermatogenesis-associated protein 33 | 139 | 15.461 | 1.612590e+04 | 3.132411e+05 |  | 271265.62 | 24265.443 | 17710.025 |  | 69.8 | 38.1 | 36.7 |
| sp\|Q96NY7\|CLIC6\_HUMAN | Q96NY7 | CLIC6\_HUMAN | CLIC6 | Chloride intracellular channel protein 6 | 704 | 73.012 | 9.493655e+02 | 8.708530e+04 |  | 87085.3 |  |  |  | 26.3 |  |  |
| sp\|Q96PE3\|INP4A\_HUMAN | Q96PE3 | INP4A\_HUMAN | INPP4A | Inositol polyphosphate-4-phosphatase type I A | 977 | 109.956 | 4.704789e+02 | 6.499445e+04 |  | 64994.45 |  |  |  | 23.2 |  |  |
| sp\|Q96RL1\|UIMC1\_HUMAN | Q96RL1 | UIMC1\_HUMAN | UIMC1 | BRCA1-A complex subunit RAP80 | 719 | 79.727 | 6.964136e+02 | 6.975733e+04 |  | 69757.33 |  |  |  | 29.6 |  |  |
| sp\|Q96S66\|CLCC1\_HUMAN | Q96S66 | CLCC1\_HUMAN | CLCC1 | Chloride channel CLIC-like protein 1 | 551 | 62.023 | 2.694363e+03 | 2.099549e+05 |  | 60536.613 |  | 149418.33 |  | 39.2 |  | 29.6 |
| sp\|Q96TC7\|RMD3\_HUMAN | Q96TC7 | RMD3\_HUMAN | RMDN3 | Regulator of microtubule dynamics protein 3 | 470 | 52.118 | 5.334335e+03 | 3.492892e+05 |  | 136947.31 | 137201.81 | 75140.12 |  | 36.0 | 19.6 | 10.2 |
| sp\|Q99442\|SEC62\_HUMAN | Q99442 | SEC62\_HUMAN | SEC62 | Translocation protein SEC62 | 399 | 45.862 | 1.356946e+04 | 7.818664e+05 |  | 310762.06 | 120020.31 | 351084.06 |  | 20.6 | 23.6 | 23.1 |
| sp\|Q99471\|PFD5\_HUMAN | Q99471 | PFD5\_HUMAN | PFDN5 | Prefoldin subunit 5 | 154 | 17.328 | 3.394376e+04 | 7.389678e+05 |  | 544052.2 | 87790.84 | 107124.75 |  | 72.1 | 35.7 | 26.0 |
| sp\|Q99614\|TTC1\_HUMAN | Q99614 | TTC1\_HUMAN | TTC1 | Tetratricopeptide repeat protein 1 | 292 | 33.526 | 1.303026e+04 | 5.488478e+05 |  | 548847.75 |  |  |  | 47.9 |  |  |
| sp\|Q9BPZ7\|SIN1\_HUMAN | Q9BPZ7 | SIN1\_HUMAN | MAPKAP1 | Target of rapamycin complex 2 subunit MAPKAP1 | 522 | 59.123 | 3.005859e+03 | 2.232761e+05 |  | 144429.34 | 78846.77 |  |  | 40.8 | 9.8 |  |
| sp\|Q9BQ69\|MACD1\_HUMAN | Q9BQ69 | MACD1\_HUMAN | MACROD1 | ADP-ribose glycohydrolase MACROD1 | 325 | 35.505 | 2.330211e+03 | 1.039446e+05 |  | 103944.61 |  |  |  | 36.0 |  |  |
| sp\|Q9BR61\|ACBD6\_HUMAN | Q9BR61 | ACBD6\_HUMAN | ACBD6 | Acyl-CoA-binding domain-containing protein 6 | 282 | 31.151 | 2.210165e+03 | 8.649954e+04 |  | 86499.54 |  |  |  | 32.6 |  |  |
| sp\|Q9BSU1\|PHAF1\_HUMAN | Q9BSU1 | PHAF1\_HUMAN | PHAF1 | Phagosome assembly factor 1 | 422 | 47.524 | 9.287945e+03 | 5.545616e+05 |  | 554561.6 |  |  |  | 47.2 |  |  |
| sp\|Q9BUP3\|HTAI2\_HUMAN | Q9BUP3 | HTAI2\_HUMAN | HTATIP2 | Oxidoreductase HTATIP2 | 242 | 27.049 | 1.918353e+04 | 6.519240e+05 |  | 571299.9 | 80624.05 |  |  | 28.9 | 28.5 |  |
| sp\|Q9BV40\|VAMP8\_HUMAN | Q9BV40 | VAMP8\_HUMAN | VAMP8 | Vesicle-associated membrane protein 8 | 100 | 11.438 | 1.011803e+05 | 1.453996e+06 |  | 203690.56 | 449797.16 | 800508.6 |  | 40.0 | 53.0 | 55.0 |
| sp\|Q9BVJ7\|DUS23\_HUMAN | Q9BVJ7 | DUS23\_HUMAN | DUSP23 | Dual specificity protein phosphatase 23 | 150 | 16.588 | 4.920241e+04 | 1.025410e+06 |  | 878170.0 | 147240.05 |  |  | 84.0 | 37.3 |  |
| sp\|Q9BVM4\|GGACT\_HUMAN | Q9BVM4 | GGACT\_HUMAN | GGACT | Gamma-glutamylaminecyclotransferase | 153 | 17.329 | 8.275760e+04 | 1.801767e+06 |  | 1801766.6 |  |  |  | 63.4 |  |  |
| sp\|Q9BW85\|YJU2\_HUMAN | Q9BW85 | YJU2\_HUMAN | YJU2 | Splicing factor YJU2 | 323 | 37.086 | 2.888036e+03 | 1.345643e+05 |  | 134564.27 |  |  |  | 39.6 |  |  |
| sp\|Q9BWU0\|NADAP\_HUMAN | Q9BWU0 | NADAP\_HUMAN | SLC4A1AP | Kanadaptin | 796 | 88.814 | 1.056172e+03 | 1.178510e+05 |  | 117851.04 |  |  |  | 21.7 |  |  |
| sp\|Q9C0B0\|UNK\_HUMAN | Q9C0B0 | UNK\_HUMAN | UNK | RING finger protein unkempt homolog | 810 | 88.084 | 3.924816e+03 | 4.343436e+05 |  | 111380.76 | 75409.086 | 247553.8 |  | 41.5 | 18.8 | 11.6 |
| sp\|Q9H0N5\|PHS2\_HUMAN | Q9H0N5 | PHS2\_HUMAN | PCBD2 | Pterin-4-alpha-carbinolamine dehydratase 2 | 130 | 14.365 | 4.140821e+04 | 7.473246e+05 |  | 520315.3 | 227009.25 |  |  | 83.8 | 25.4 |  |
| sp\|Q9H0R8\|GBRL1\_HUMAN | Q9H0R8 | GBRL1\_HUMAN | GABARAPL1 | Gamma-aminobutyric acid receptor-associated protein-like 1 | 117 | 14.044 | 1.858089e+04 | 3.278494e+05 |  | 327849.44 |  |  |  | 22.2 |  |  |
| sp\|Q9H1H9\|KI13A\_HUMAN | Q9H1H9 | KI13A\_HUMAN | KIF13A | Kinesin-like protein KIF13A | 1805 | 202.308 | 9.197105e+02 | 2.337660e+05 |  | 226489.8 |  | 7276.207 |  | 27.3 |  | 6.7 |
| sp\|Q9H254\|SPTN4\_HUMAN | Q9H254 | SPTN4\_HUMAN | SPTBN4 | Spectrin beta chain, non-erythrocytic 4 | 2564 | 288.985 | 1.046132e+05 | 3.798208e+07 |  | 2280758.8 | 34325976.0 | 1375346.0 |  | 29.0 | 22.0 | 11.8 |
| sp\|Q9H446\|RWDD1\_HUMAN | Q9H446 | RWDD1\_HUMAN | RWDD1 | RWD domain-containing protein 1 | 243 | 27.940 | 5.339831e+03 | 1.874438e+05 |  | 187443.8 |  |  |  | 29.6 |  |  |
| sp\|Q9H4E7\|DEFI6\_HUMAN | Q9H4E7 | DEFI6\_HUMAN | DEF6 | Differentially expressed in FDCP 6 homolog | 631 | 73.910 | 7.321654e+03 | 6.798758e+05 |  | 679875.75 |  |  |  | 37.9 |  |  |
| sp\|Q9HB71\|CYBP\_HUMAN | Q9HB71 | CYBP\_HUMAN | CACYBP | Calcyclin-binding protein | 228 | 26.210 | 1.275120e+04 | 4.198897e+05 |  | 419889.66 |  |  |  | 34.2 |  |  |
| sp\|Q9HCD6\|TANC2\_HUMAN | Q9HCD6 | TANC2\_HUMAN | TANC2 | Protein TANC2 | 1990 | 219.650 | 8.726321e+03 | 2.408128e+06 |  | 2370561.8 | 14303.661 | 23262.48 |  | 27.5 | 13.6 | 7.1 |
| sp\|Q9HCP0\|KC1G1\_HUMAN | Q9HCP0 | KC1G1\_HUMAN | CSNK1G1 | Casein kinase I isoform gamma-1 | 422 | 48.511 | 1.125456e+04 | 6.859396e+05 |  | 305683.12 | 380256.44 |  |  | 43.8 | 21.6 |  |
| sp\|Q9HD42\|CHM1A\_HUMAN | Q9HD42 | CHM1A\_HUMAN | CHMP1A | Charged multivesicular body protein 1a | 196 | 21.703 | 1.473145e+04 | 4.016821e+05 |  | 401682.06 |  |  |  | 24.5 |  |  |
| sp\|Q9HD45\|TM9S3\_HUMAN | Q9HD45 | TM9S3\_HUMAN | TM9SF3 | Transmembrane 9 superfamily member 3 | 589 | 67.888 | 1.267064e+04 | 1.080709e+06 |  | 187388.86 | 351196.7 | 542123.25 |  | 22.2 | 13.1 | 10.2 |
| sp\|Q9NP61\|ARFG3\_HUMAN | Q9NP61 | ARFG3\_HUMAN | ARFGAP3 | ADP-ribosylation factor GTPase-activating protein 3 | 516 | 56.928 | 2.404607e+03 | 1.719836e+05 |  | 171983.64 |  |  |  | 35.7 |  |  |
| sp\|Q9NQP4\|PFD4\_HUMAN | Q9NQP4 | PFD4\_HUMAN | PFDN4 | Prefoldin subunit 4 | 134 | 15.314 | 6.189368e+04 | 1.190836e+06 |  | 1178838.9 | 7697.143 | 4300.3535 |  | 41.8 | 11.2 | 11.2 |
| sp\|Q9NQS1\|AVEN\_HUMAN | Q9NQS1 | AVEN\_HUMAN | AVEN | Cell death regulator Aven | 362 | 38.506 | 7.594580e+03 | 3.674086e+05 |  | 367408.62 |  |  |  | 49.7 |  |  |
| sp\|Q9NQY0\|BIN3\_HUMAN | Q9NQY0 | BIN3\_HUMAN | BIN3 | Bridging integrator 3 | 253 | 29.665 | 2.063457e+04 | 7.690541e+05 |  | 655333.25 | 113720.81 |  |  | 47.8 | 33.2 |  |
| sp\|Q9NRL2\|BAZ1A\_HUMAN | Q9NRL2 | BAZ1A\_HUMAN | BAZ1A | Bromodomain adjacent to zinc finger domain protein 1A | 1556 | 178.702 | 4.452170e+04 | 9.995817e+06 |  | 9995817.0 |  |  |  | 19.6 |  |  |
| sp\|Q9NTZ6\|RBM12\_HUMAN | Q9NTZ6 | RBM12\_HUMAN | RBM12 | RNA-binding protein 12 | 932 | 97.395 | 6.805668e+03 | 8.327691e+05 |  | 390305.8 | 442463.3 |  |  | 25.6 | 23.6 |  |
| sp\|Q9NUJ3\|T11L1\_HUMAN | Q9NUJ3 | T11L1\_HUMAN | TCP11L1 | T-complex protein 11-like protein 1 | 509 | 57.035 | 1.273878e+03 | 9.128227e+04 |  | 91282.266 |  |  |  | 44.4 |  |  |
| sp\|Q9NV35\|NUD15\_HUMAN | Q9NV35 | NUD15\_HUMAN | NUDT15 | Nucleotide triphosphate diphosphatase NUDT15 | 164 | 18.609 | 5.683087e+03 | 1.328692e+05 |  | 132869.25 |  |  |  | 37.8 |  |  |
| sp\|Q9NV70\|EXOC1\_HUMAN | Q9NV70 | EXOC1\_HUMAN | EXOC1 | Exocyst complex component 1 | 894 | 101.982 | 8.825154e+02 | 1.130741e+05 |  | 113074.05 |  |  |  | 30.8 |  |  |
| sp\|Q9NYT0\|PLEK2\_HUMAN | Q9NYT0 | PLEK2\_HUMAN | PLEK2 | Pleckstrin-2 | 353 | 39.971 | 1.045585e+03 | 5.250750e+04 |  | 52507.5 |  |  |  | 21.5 |  |  |
| sp\|Q9P0G3\|KLK14\_HUMAN | Q9P0G3 | KLK14\_HUMAN | KLK14 | Kallikrein-14 | 267 | 29.122 | 6.447866e+03 | 2.359143e+05 |  | 235914.34 |  |  |  | 57.7 |  |  |
| sp\|Q9P0S9\|TM14C\_HUMAN | Q9P0S9 | TM14C\_HUMAN | TMEM14C | Transmembrane protein 14C | 112 | 11.565 | 3.289466e+04 | 4.779562e+05 |  | 387731.5 | 90224.73 |  |  | 67.0 | 8.9 |  |
| sp\|Q9P2D3\|HTR5B\_HUMAN | Q9P2D3 | HTR5B\_HUMAN | HEATR5B | HEAT repeat-containing protein 5B | 2071 | 224.302 | 2.138236e+02 | 6.025678e+04 |  | 60256.78 |  |  |  | 31.7 |  |  |
| sp\|Q9UBG3\|CRNN\_HUMAN | Q9UBG3 | CRNN\_HUMAN | CRNN | Cornulin | 495 | 53.533 | 1.361956e+04 | 9.160135e+05 |  | 60504.082 | 774552.44 | 80956.98 |  | 42.0 | 52.3 | 15.4 |
| sp\|Q9UHG3\|PCYOX\_HUMAN | Q9UHG3 | PCYOX\_HUMAN | PCYOX1 | Prenylcysteine oxidase 1 | 505 | 56.640 | 6.300014e+03 | 4.483136e+05 |  | 172293.88 | 276019.75 |  |  | 37.6 | 20.0 |  |
| sp\|Q9UI08\|EVL\_HUMAN | Q9UI08 | EVL\_HUMAN | EVL | Ena/VASP-like protein | 416 | 44.620 | 7.654595e+02 | 4.291103e+04 |  | 42911.027 |  |  |  | 55.3 |  |  |
| sp\|Q9UKT8\|FBXW2\_HUMAN | Q9UKT8 | FBXW2\_HUMAN | FBXW2 | F-box/WD repeat-containing protein 2 | 454 | 51.512 | 4.768392e+03 | 3.086011e+05 |  | 308601.12 |  |  |  | 33.0 |  |  |
| sp\|Q9UL26\|RB22A\_HUMAN | Q9UL26 | RB22A\_HUMAN | RAB22A | Ras-related protein Rab-22A | 194 | 21.855 | 9.518377e+03 | 2.613550e+05 |  | 261355.02 |  |  |  | 73.7 |  |  |
| sp\|Q9UNF1\|MAGD2\_HUMAN | Q9UNF1 | MAGD2\_HUMAN | MAGED2 | Melanoma-associated antigen D2 | 606 | 64.954 | 1.615733e+03 | 1.318538e+05 |  | 84115.305 | 47738.527 |  |  | 37.5 | 31.4 |  |
| sp\|Q9UPN3\|MACF1\_HUMAN | Q9UPN3 | MACF1\_HUMAN | MACF1 | Microtubule-actin cross-linking factor 1, isoforms 1/2/3/5 | 7388 | 838.308 | 1.111227e+02 | 1.170371e+05 |  | 117037.125 |  |  |  | 32.2 |  |  |
| sp\|Q9UPT8\|ZC3H4\_HUMAN | Q9UPT8 | ZC3H4\_HUMAN | ZC3H4 | Zinc finger CCCH domain-containing protein 4 | 1303 | 140.257 | 4.442082e+02 | 7.827593e+04 |  | 78275.93 |  |  |  | 34.2 |  |  |
| sp\|Q9Y237\|PIN4\_HUMAN | Q9Y237 | PIN4\_HUMAN | PIN4 | Peptidyl-prolyl cis-trans isomerase NIMA-interacting 4 | 131 | 13.810 | 3.368803e+04 | 5.845026e+05 |  | 557947.6 |  | 26554.953 |  | 64.9 |  | 29.0 |
| sp\|Q9Y2T7\|YBOX2\_HUMAN | Q9Y2T7 | YBOX2\_HUMAN | YBX2 | Y-box-binding protein 2 | 364 | 38.518 | 2.792680e+04 | 1.351456e+06 |  | 910369.94 | 441086.38 |  |  | 64.6 | 28.6 |  |
| sp\|Q9Y3A5\|SBDS\_HUMAN | Q9Y3A5 | SBDS\_HUMAN | SBDS | Ribosome maturation protein SBDS | 250 | 28.764 | 1.375584e+04 | 4.971113e+05 |  | 497111.34 |  |  |  | 39.6 |  |  |
| sp\|Q9Y3S2\|ZN330\_HUMAN | Q9Y3S2 | ZN330\_HUMAN | ZNF330 | Zinc finger protein 330 | 320 | 36.201 | 2.957382e+03 | 1.345071e+05 |  | 134507.06 |  |  |  | 18.8 |  |  |
| sp\|Q9Y485\|DMXL1\_HUMAN | Q9Y485 | DMXL1\_HUMAN | DMXL1 | DmX-like protein 1 | 3027 | 337.839 | 1.417800e+02 | 6.017859e+04 |  | 60178.594 |  |  |  | 22.9 |  |  |
| sp\|Q9Y4Y9\|LSM5\_HUMAN | Q9Y4Y9 | LSM5\_HUMAN | LSM5 | U6 snRNA-associated Sm-like protein LSm5 | 91 | 9.937 | 6.048375e+03 | 7.551116e+04 |  | 75511.164 |  |  |  | 61.5 |  |  |
| sp\|Q9Y4Z0\|LSM4\_HUMAN | Q9Y4Z0 | LSM4\_HUMAN | LSM4 | U6 snRNA-associated Sm-like protein LSm4 | 139 | 15.350 | 2.550920e+04 | 4.919516e+05 |  | 491951.56 |  |  |  | 53.2 |  |  |
| sp\|Q9Y587\|AP4S1\_HUMAN | Q9Y587 | AP4S1\_HUMAN | AP4S1 | AP-4 complex subunit sigma-1 | 144 | 17.005 | 6.301858e+04 | 1.346364e+06 |  | 1234080.8 | 99181.57 | 13101.426 |  | 58.3 | 33.3 | 13.2 |
| sp\|Q9Y5R8\|TPPC1\_HUMAN | Q9Y5R8 | TPPC1\_HUMAN | TRAPPC1 | Trafficking protein particle complex subunit 1 | 145 | 16.832 | 4.773908e+03 | 1.009548e+05 |  | 100954.78 |  |  |  | 49.0 |  |  |
| sp\|Q9Y6R4\|M3K4\_HUMAN | Q9Y6R4 | M3K4\_HUMAN | MAP3K4 | Mitogen-activated protein kinase kinase kinase 4 | 1608 | 181.685 | 1.035006e+02 | 2.362539e+04 |  | 23625.387 |  |  |  | 28.9 |  |  |
| sp\|Q9Y6U3\|SCIN\_HUMAN | Q9Y6U3 | SCIN\_HUMAN | SCIN | Scinderin | 715 | 80.489 | 6.551431e+02 | 6.625061e+04 |  | 66250.61 |  |  |  | 30.6 |  |  |
| sp\|P78385\|KRT83\_HUMAN | P78385 | KRT83\_HUMAN | KRT83 | Keratin, type II cuticular Hb3 | 493 | 54.195 | 3.667253e+05 | 2.496992e+07 |  | 747914.5 | 1119901.1 | 23102108.0 |  | 45.8 | 35.1 | 35.5 |
| sp\|Q13136\|LIPA1\_HUMAN | Q13136 | LIPA1\_HUMAN | PPFIA1 | Liprin-alpha-1 | 1202 | 135.779 | 9.095768e+01 | 1.551633e+04 |  | 15516.334 |  |  |  | 34.4 |  |  |
| sp\|O95271\|TNKS1\_HUMAN | O95271 | TNKS1\_HUMAN | TNKS | Poly [ADP-ribose] polymerase tankyrase-1 | 1327 | 142.039 | 7.425606e+02 | 1.325124e+05 |  | 132512.44 | 0.0 |  |  | 29.3 | 11.2 |  |
| sp\|O95714\|HERC2\_HUMAN | O95714 | HERC2\_HUMAN | HERC2 | E3 ubiquitin-protein ligase HERC2 | 4834 | 527.228 | 7.873379e+02 | 5.215270e+05 |  | 521527.0 |  |  |  | 27.0 |  |  |
| sp\|P00747\|PLMN\_HUMAN | P00747 | PLMN\_HUMAN | PLG | Plasminogen | 810 | 90.569 | 2.125176e+04 | 2.418197e+06 |  | 2223816.8 | 94143.8 | 100236.22 |  | 41.6 | 25.8 | 20.7 |
| sp\|P04439\|HLAA\_HUMAN | P04439 | HLAA\_HUMAN | HLA-A | HLA class I histocompatibility antigen, A alpha chain | 365 | 40.841 | 7.020584e+03 | 3.602357e+05 |  | 74885.83 | 241485.56 | 43864.3 |  | 34.0 | 38.6 | 9.0 |
| sp\|Q58FF6\|H90B4\_HUMAN | Q58FF6 | H90B4\_HUMAN | HSP90AB4P | Putative heat shock protein HSP 90-beta 4 | 505 | 58.264 | 2.086744e+05 | 1.527519e+07 |  | 10266690.0 | 5008501.0 |  |  | 32.5 | 19.0 |  |
| sp\|P13535\|MYH8\_HUMAN | P13535 | MYH8\_HUMAN | MYH8 | Myosin-8 | 1937 | 222.763 | 1.496367e+04 | 4.187918e+06 |  | 1599662.4 | 1773236.4 | 815019.6 |  | 39.3 | 26.5 | 18.2 |
| sp\|P12883\|MYH7\_HUMAN | P12883 | MYH7\_HUMAN | MYH7 | Myosin-7 | 1935 | 223.097 | 1.149062e+04 | 3.220732e+06 |  | 1559102.5 | 1661629.1 |  |  | 35.2 | 22.4 |  |
| sp\|P10412\|H14\_HUMAN | P10412 | H14\_HUMAN | H1-4 | Histone H1.4 | 219 | 21.865 | 2.613189e+04 | 7.178563e+05 |  | 172526.1 | 359040.2 | 186290.03 |  | 46.1 | 23.5 | 26.5 |
| sp\|P16403\|H12\_HUMAN | P16403 | H12\_HUMAN | H1-2 | Histone H1.2 | 213 | 21.365 | 5.253042e+03 | 1.410038e+05 |  | 141003.83 |  |  |  | 62.9 |  |  |
| sp\|P41236\|IPP2\_HUMAN | P41236 | IPP2\_HUMAN | PPP1R2 | Protein phosphatase inhibitor 2 | 205 | 23.015 | 1.289698e+04 | 3.729205e+05 |  | 372920.53 |  |  |  | 29.8 |  |  |
| sp\|Q04828\|AK1C1\_HUMAN | Q04828 | AK1C1\_HUMAN | AKR1C1 | Aldo-keto reductase family 1 member C1 | 323 | 36.788 | 1.559521e+04 | 7.207994e+05 |  | 720799.44 |  |  |  | 57.9 |  |  |
| sp\|P49792\|RBP2\_HUMAN | P49792 | RBP2\_HUMAN | RANBP2 | E3 SUMO-protein ligase RanBP2 | 3224 | 358.199 | 1.567528e+02 | 7.054347e+04 |  | 70543.47 |  |  |  | 30.5 |  |  |
| sp\|P62328\|TYB4\_HUMAN | P62328 | TYB4\_HUMAN | TMSB4X | Thymosin beta-4 | 44 | 5.053 | 4.357180e+04 | 2.766126e+05 |  | 178257.1 | 74443.84 | 23911.627 |  | 77.3 | 27.3 | 56.8 |
| sp\|P62837\|UB2D2\_HUMAN | P62837 | UB2D2\_HUMAN | UBE2D2 | Ubiquitin-conjugating enzyme E2 D2 | 147 | 16.735 | 8.260961e+04 | 1.736894e+06 |  | 1054403.0 | 682491.5 |  |  | 26.5 | 12.2 |  |
| sp\|Q71U36\|TBA1A\_HUMAN | Q71U36 | TBA1A\_HUMAN | TUBA1A | Tubulin alpha-1A chain | 451 | 50.136 | 8.689072e+04 | 5.473186e+06 |  | 5473186.0 |  |  |  | 47.7 |  |  |
| sp\|P04350\|TBB4A\_HUMAN | P04350 | TBB4A\_HUMAN | TUBB4A | Tubulin beta-4A chain | 444 | 49.586 | 2.902404e+04 | 1.808148e+06 |  | 1808148.2 |  |  |  | 45.5 |  |  |
| sp\|P68431\|H31\_HUMAN | P68431 | H31\_HUMAN | H3C1 | Histone H3.1 | 136 | 15.404 | 1.358649e+05 | 2.629407e+06 |  | 726589.06 | 1097835.1 | 804982.94 |  | 66.9 | 76.5 | 23.5 |
| sp\|P18085\|ARF4\_HUMAN | P18085 | ARF4\_HUMAN | ARF4 | ADP-ribosylation factor 4 | 180 | 20.511 | 6.668410e+05 | 1.718408e+07 |  | 16283402.0 | 383538.2 | 517139.5 |  | 66.1 | 45.0 | 33.9 |
| sp\|P84077\|ARF1\_HUMAN | P84077 | ARF1\_HUMAN | ARF1 | ADP-ribosylation factor 1 | 181 | 20.697 | 1.028932e+06 | 2.675538e+07 |  | 26755378.0 |  |  |  | 81.2 |  |  |
| sp\|Q14571\|ITPR2\_HUMAN | Q14571 | ITPR2\_HUMAN | ITPR2 | Inositol 1,4,5-trisphosphate receptor type 2 | 2701 | 308.064 | 1.887700e+02 | 7.306191e+04 |  | 28146.293 |  | 44915.617 |  | 28.1 |  | 8.2 |
| sp\|Q93077\|H2A1C\_HUMAN | Q93077 | H2A1C\_HUMAN | H2AC6 | Histone H2A type 1-C | 130 | 14.105 | 8.077177e+04 | 1.431363e+06 |  | 1431363.2 |  |  |  | 70.0 |  |  |
| sp\|Q16777\|H2A2C\_HUMAN | Q16777 | H2A2C\_HUMAN | H2AC20 | Histone H2A type 2-C | 129 | 13.988 | 2.231910e+05 | 3.922378e+06 |  | 1472348.8 | 1729777.4 | 720251.6 |  | 65.1 | 60.5 | 55.8 |
| sp\|P0C0S8\|H2A1\_HUMAN | P0C0S8 | H2A1\_HUMAN | H2AC11 | Histone H2A type 1 | 130 | 14.091 | 8.316713e+04 | 1.472349e+06 |  | 1472348.8 |  |  |  | 0.0 |  |  |
| sp\|Q5QNW6\|H2B2F\_HUMAN | Q5QNW6 | H2B2F\_HUMAN | H2BC18 | Histone H2B type 2-F | 126 | 13.920 | 1.163973e+05 | 2.035632e+06 |  | 2035631.5 |  |  |  | 81.0 |  |  |
| sp\|Q5SW79\|CE170\_HUMAN | Q5SW79 | CE170\_HUMAN | CEP170 | Centrosomal protein of 170 kDa | 1584 | 175.293 | 5.843463e+02 | 1.286921e+05 |  | 128692.14 |  |  |  | 32.1 |  |  |
| sp\|Q7L523\|RRAGA\_HUMAN | Q7L523 | RRAGA\_HUMAN | RRAGA | Ras-related GTP-binding protein A | 313 | 36.566 | 9.280648e+03 | 4.263565e+05 |  | 426356.53 |  |  |  | 23.0 |  |  |
| sp\|Q7Z5G4\|GOGA7\_HUMAN | Q7Z5G4 | GOGA7\_HUMAN | GOLGA7 | Golgin subfamily A member 7 | 137 | 15.824 | 3.256152e+05 | 6.473485e+06 |  | 560248.1 | 1146854.2 | 4766382.5 |  | 30.7 | 42.3 | 44.5 |
| sp\|Q6ZT62\|BGIN\_HUMAN | Q6ZT62 | BGIN\_HUMAN | BARGIN | Bargin | 677 | 73.599 | 1.174700e+04 | 1.086216e+06 |  | 1086215.8 |  |  |  | 41.2 |  |  |
| sp\|Q9BZL6\|KPCD2\_HUMAN | Q9BZL6 | KPCD2\_HUMAN | PRKD2 | Serine/threonine-protein kinase D2 | 878 | 96.722 | 7.191288e+02 | 8.738746e+04 |  | 87387.46 |  |  |  | 28.8 |  |  |
| sp\|Q9Y297\|FBW1A\_HUMAN | Q9Y297 | FBW1A\_HUMAN | BTRC | F-box/WD repeat-containing protein 1A | 605 | 68.867 | 1.077680e+03 | 9.324338e+04 |  | 93243.38 |  |  |  | 54.2 |  |  |
| sp\|Q9UP95\|S12A4\_HUMAN | Q9UP95 | S12A4\_HUMAN | SLC12A4 | Solute carrier family 12 member 4 | 1085 | 120.650 | 7.840684e+03 | 1.188498e+06 |  | 249738.22 |  | 938759.7 |  | 18.8 |  | 10.5 |
| sp\|O00291\|HIP1\_HUMAN | O00291 | HIP1\_HUMAN | HIP1 | Huntingtin-interacting protein 1 | 1037 | 116.221 | 3.743320e+02 | 5.465862e+04 |  | 54658.62 |  |  |  | 36.8 |  |  |
| sp\|O14828\|SCAM3\_HUMAN | O14828 | SCAM3\_HUMAN | SCAMP3 | Secretory carrier-associated membrane protein 3 | 347 | 38.287 | 3.435428e+03 | 1.652530e+05 |  | 75986.18 | 16309.646 | 72957.164 |  | 30.0 | 17.9 | 12.7 |
| sp\|O15541\|R113A\_HUMAN | O15541 | R113A\_HUMAN | RNF113A | E3 ubiquitin-protein ligase RNF113A | 343 | 38.787 | 2.289376e+03 | 1.115631e+05 |  | 111563.05 |  |  |  | 39.1 |  |  |
| sp\|O43752\|STX6\_HUMAN | O43752 | STX6\_HUMAN | STX6 | Syntaxin-6 | 255 | 29.176 | 1.690958e+04 | 6.198343e+05 |  | 47746.957 | 221983.12 | 350104.22 |  | 47.1 | 29.4 | 22.4 |
| sp\|O75884\|RBBP9\_HUMAN | O75884 | RBBP9\_HUMAN | RBBP9 | Serine hydrolase RBBP9 | 186 | 21.000 | 8.670243e+03 | 2.287535e+05 |  | 228753.48 |  |  |  | 53.8 |  |  |
| sp\|O75891\|AL1L1\_HUMAN | O75891 | AL1L1\_HUMAN | ALDH1L1 | Cytosolic 10-formyltetrahydrofolate dehydrogenase | 902 | 98.829 | 9.759729e+02 | 1.211823e+05 |  | 121182.336 |  |  |  | 29.9 |  |  |
| sp\|O95166\|GBRAP\_HUMAN | O95166 | GBRAP\_HUMAN | GABARAP | Gamma-aminobutyric acid receptor-associated protein | 117 | 13.918 | 5.295095e+04 | 9.259078e+05 |  | 925907.8 |  |  |  | 21.4 |  |  |
| sp\|P01011\|AACT\_HUMAN | P01011 | AACT\_HUMAN | SERPINA3 | Alpha-1-antichymotrypsin | 423 | 47.651 | 7.399960e+03 | 4.430151e+05 |  | 74643.13 | 229943.11 | 138428.86 |  | 30.5 | 23.9 | 17.5 |
| sp\|P02763\|A1AG1\_HUMAN | P02763 | A1AG1\_HUMAN | ORM1 | Alpha-1-acid glycoprotein 1 | 201 | 23.540 | 1.013903e+04 | 2.998610e+05 |  | 182466.0 | 117395.05 |  |  | 31.3 | 20.4 |  |
| sp\|P08697\|A2AP\_HUMAN | P08697 | A2AP\_HUMAN | SERPINF2 | Alpha-2-antiplasmin | 491 | 54.566 | 3.348162e+03 | 2.295334e+05 |  | 46981.79 | 182551.56 |  |  | 36.5 | 25.9 |  |
| sp\|P20774\|MIME\_HUMAN | P20774 | MIME\_HUMAN | OGN | Mimecan | 298 | 33.922 | 8.957287e+02 | 3.817465e+04 |  | 33970.117 | 4204.5366 |  |  | 21.8 | 21.1 |  |
| sp\|P25205\|MCM3\_HUMAN | P25205 | MCM3\_HUMAN | MCM3 | DNA replication licensing factor MCM3 | 808 | 90.981 | 2.668410e+02 | 3.050144e+04 |  | 30501.443 |  |  |  | 29.0 |  |  |
| sp\|P40616\|ARL1\_HUMAN | P40616 | ARL1\_HUMAN | ARL1 | ADP-ribosylation factor-like protein 1 | 181 | 20.418 | 2.682880e+03 | 6.882269e+04 |  | 68822.69 |  |  |  | 39.8 |  |  |
| sp\|P49747\|COMP\_HUMAN | P49747 | COMP\_HUMAN | COMP | Cartilage oligomeric matrix protein | 757 | 82.860 | 4.107788e+02 | 4.276319e+04 |  | 42763.19 |  |  |  | 28.5 |  |  |
| sp\|P52758\|RIDA\_HUMAN | P52758 | RIDA\_HUMAN | RIDA | 2-iminobutanoate/2-iminopropanoate deaminase | 137 | 14.494 | 2.510160e+04 | 4.570953e+05 |  | 301049.56 | 156045.75 |  |  | 50.4 | 32.1 |  |
| sp\|Q14566\|MCM6\_HUMAN | Q14566 | MCM6\_HUMAN | MCM6 | DNA replication licensing factor MCM6 | 821 | 92.889 | 5.507749e+02 | 6.427699e+04 |  | 64276.992 |  |  |  | 36.5 |  |  |
| sp\|Q16595\|FRDA\_HUMAN | Q16595 | FRDA\_HUMAN | FXN | Frataxin, mitochondrial | 210 | 23.135 | 2.270186e+03 | 6.598545e+04 |  | 65985.45 |  |  |  | 47.1 |  |  |
| sp\|Q6IA17\|SIGIR\_HUMAN | Q6IA17 | SIGIR\_HUMAN | SIGIRR | Single Ig IL-1-related receptor | 410 | 45.679 | 3.070713e+03 | 1.762272e+05 |  | 176227.22 |  |  |  | 19.0 |  |  |
| sp\|Q8IVV7\|GID4\_HUMAN | Q8IVV7 | GID4\_HUMAN | GID4 | Glucose-induced degradation protein 4 homolog | 300 | 33.514 | 3.032203e+03 | 1.276738e+05 |  | 127673.78 |  |  |  | 32.7 |  |  |
| sp\|Q8IX07\|FOG1\_HUMAN | Q8IX07 | FOG1\_HUMAN | ZFPM1 | Zinc finger protein ZFPM1 | 1006 | 104.888 | 3.735460e+02 | 4.922514e+04 |  | 49225.145 |  |  |  | 48.3 |  |  |
| sp\|Q8IYD1\|ERF3B\_HUMAN | Q8IYD1 | ERF3B\_HUMAN | GSPT2 | Eukaryotic peptide chain release factor GTP-binding subunit ERF3B | 628 | 68.883 | 1.094595e+04 | 9.472898e+05 |  | 947289.75 |  |  |  | 43.2 |  |  |
| sp\|Q8NDA8\|MROH1\_HUMAN | Q8NDA8 | MROH1\_HUMAN | MROH1 | Maestro heat-like repeat-containing protein family member 1 | 1641 | 181.249 | 1.302010e+02 | 2.964880e+04 |  | 29648.797 |  |  |  | 28.6 |  |  |
| sp\|Q92503\|S14L1\_HUMAN | Q92503 | S14L1\_HUMAN | SEC14L1 | SEC14-like protein 1 | 715 | 81.250 | 2.488891e+02 | 2.540659e+04 |  | 25406.588 |  |  |  | 45.0 |  |  |
| sp\|Q96DC7\|TMCO6\_HUMAN | Q96DC7 | TMCO6\_HUMAN | TMCO6 | Transmembrane and coiled-coil domain-containing protein 6 | 493 | 54.442 | 2.270945e+03 | 1.553309e+05 |  | 155330.9 |  |  |  | 14.4 |  |  |
| sp\|Q99685\|MGLL\_HUMAN | Q99685 | MGLL\_HUMAN | MGLL | Monoglyceride lipase | 303 | 33.261 | 2.793781e+03 | 1.167468e+05 |  | 116746.766 |  |  |  | 32.0 |  |  |
| sp\|Q9BQK8\|LPIN3\_HUMAN | Q9BQK8 | LPIN3\_HUMAN | LPIN3 | Phosphatidate phosphatase LPIN3 | 851 | 93.614 | 5.404344e+02 | 6.356249e+04 |  | 63562.492 |  |  |  | 30.2 |  |  |
| sp\|Q9BYJ1\|LOXE3\_HUMAN | Q9BYJ1 | LOXE3\_HUMAN | ALOXE3 | Hydroperoxide isomerase ALOXE3 | 711 | 80.543 | 3.184930e+03 | 3.222885e+05 |  | 61713.92 |  | 260574.58 |  | 41.9 |  | 9.6 |
| sp\|Q9BZL4\|PP12C\_HUMAN | Q9BZL4 | PP12C\_HUMAN | PPP1R12C | Protein phosphatase 1 regulatory subunit 12C | 782 | 84.881 | 1.600338e+02 | 1.706630e+04 |  | 17066.3 |  |  |  | 41.3 |  |  |
| sp\|Q9C0D9\|EPT1\_HUMAN | Q9C0D9 | EPT1\_HUMAN | SELENOI | Ethanolaminephosphotransferase 1 | 397 | 45.229 | 3.642680e+03 | 2.069928e+05 |  | 206992.78 |  |  |  | 14.6 |  |  |
| sp\|Q9H0F6\|SHRPN\_HUMAN | Q9H0F6 | SHRPN\_HUMAN | SHARPIN | Sharpin | 387 | 39.949 | 1.723083e+03 | 8.648268e+04 |  | 86482.68 |  |  |  | 52.5 |  |  |
| sp\|Q9H3U1\|UN45A\_HUMAN | Q9H3U1 | UN45A\_HUMAN | UNC45A | Protein unc-45 homolog A | 944 | 103.077 | 1.405076e+02 | 1.819612e+04 |  | 18196.121 |  |  |  | 45.3 |  |  |
| sp\|Q9NWH9\|SLTM\_HUMAN | Q9NWH9 | SLTM\_HUMAN | SLTM | SAFB-like transcription modulator | 1034 | 117.148 | 9.903791e+02 | 1.457651e+05 |  | 127318.34 | 18446.748 |  |  | 30.9 | 27.7 |  |
| sp\|Q9P1F3\|ABRAL\_HUMAN | Q9P1F3 | ABRAL\_HUMAN | ABRACL | Costars family protein ABRACL | 81 | 9.056 | 1.576959e+04 | 1.794213e+05 |  | 179421.34 |  |  |  | 50.6 |  |  |
| sp\|Q9UL33\|TPC2L\_HUMAN | Q9UL33 | TPC2L\_HUMAN | TRAPPC2L | Trafficking protein particle complex subunit 2-like protein | 140 | 16.146 | 4.025946e+03 | 8.166765e+04 |  | 81667.65 |  |  |  | 76.4 |  |  |
| sp\|Q9UMR2\|DD19B\_HUMAN | Q9UMR2 | DD19B\_HUMAN | DDX19B | ATP-dependent RNA helicase DDX19B | 479 | 53.927 | 1.757473e+03 | 1.190726e+05 |  | 119072.65 |  |  |  | 40.5 |  |  |
| sp\|Q9Y217\|MTMR6\_HUMAN | Q9Y217 | MTMR6\_HUMAN | MTMR6 | Myotubularin-related protein 6 | 621 | 71.968 | 7.542352e+02 | 6.819670e+04 |  | 68196.7 |  |  |  | 22.4 |  |  |
| sp\|Q9Y333\|LSM2\_HUMAN | Q9Y333 | LSM2\_HUMAN | LSM2 | U6 snRNA-associated Sm-like protein LSm2 | 95 | 10.835 | 5.377170e+04 | 7.319810e+05 |  | 731981.0 |  |  |  | 74.7 |  |  |
| sp\|Q9Y394\|DHRS7\_HUMAN | Q9Y394 | DHRS7\_HUMAN | DHRS7 | Dehydrogenase/reductase SDR family member 7 | 339 | 38.299 | 3.322069e+03 | 1.598502e+05 |  | 115518.664 |  | 44331.543 |  | 23.9 |  | 9.7 |
| sp\|Q9Y3B3\|TMED7\_HUMAN | Q9Y3B3 | TMED7\_HUMAN | TMED7 | Transmembrane emp24 domain-containing protein 7 | 224 | 25.172 | 4.408016e+04 | 1.394049e+06 |  | 379854.78 | 1014194.06 |  |  | 51.3 | 25.4 |  |
| sp\|Q9Y3D6\|FIS1\_HUMAN | Q9Y3D6 | FIS1\_HUMAN | FIS1 | Mitochondrial fission 1 protein | 152 | 16.938 | 3.527525e+04 | 7.506705e+05 |  | 353391.84 | 231127.45 | 166151.22 |  | 26.3 | 15.8 | 8.6 |
| sp\|Q9Y5Q8\|TF3C5\_HUMAN | Q9Y5Q8 | TF3C5\_HUMAN | GTF3C5 | General transcription factor 3C polypeptide 5 | 519 | 59.571 | 4.440623e+02 | 3.323502e+04 |  | 33235.02 |  |  |  | 51.6 |  |  |
| sp\|A6NMB1\|SIG16\_HUMAN | A6NMB1 | SIG16\_HUMAN | SIGLEC16 | Sialic acid-binding Ig-like lectin 16 | 481 | 52.992 | 9.601587e+03 | 6.392496e+05 |  | 639249.6 |  |  |  | 49.9 |  |  |
| sp\|P07478\|TRY2\_HUMAN | P07478 | TRY2\_HUMAN | PRSS2 | Trypsin-2 | 247 | 26.488 | 2.567528e+04 | 8.544399e+05 |  | 854439.9 |  |  |  | 10.9 |  |  |
| sp\|P49757\|NUMB\_HUMAN | P49757 | NUMB\_HUMAN | NUMB | Protein numb homolog | 651 | 70.804 | 3.300240e+02 | 2.935760e+04 |  | 29357.598 |  |  |  | 23.2 |  |  |
| sp\|Q9NR56\|MBNL1\_HUMAN | Q9NR56 | MBNL1\_HUMAN | MBNL1 | Muscleblind-like protein 1 | 388 | 41.817 | 4.783300e+03 | 2.513030e+05 |  | 251302.98 |  |  |  | 24.5 |  |  |
| sp\|B7ZAQ6\|GPHRA\_HUMAN | B7ZAQ6 | GPHRA\_HUMAN | GPR89A | Golgi pH regulator A | 455 | 52.917 | 1.017300e+05 | 6.763342e+06 |  | 6657968.0 |  | 105373.734 |  | 17.4 |  | 12.5 |
| sp\|O14641\|DVL2\_HUMAN | O14641 | DVL2\_HUMAN | DVL2 | Segment polarity protein dishevelled homolog DVL-2 | 736 | 78.948 | 7.713641e+02 | 7.650991e+04 |  | 76509.914 |  |  |  | 31.5 |  |  |
| sp\|P01111\|RASN\_HUMAN | P01111 | RASN\_HUMAN | NRAS | GTPase NRas | 189 | 21.229 | 4.721572e+04 | 1.259312e+06 |  | 354449.62 | 904862.25 |  |  | 78.3 | 40.7 |  |
| sp\|P05155\|IC1\_HUMAN | P05155 | IC1\_HUMAN | SERPING1 | Plasma protease C1 inhibitor | 500 | 55.154 | 3.656068e+02 | 2.533427e+04 |  | 25334.273 |  |  |  | 32.4 |  |  |
| sp\|P22894\|MMP8\_HUMAN | P22894 | MMP8\_HUMAN | MMP8 | Neutrophil collagenase | 467 | 53.412 | 2.358328e+02 | 1.582560e+04 |  | 12089.981 | 3735.6174 |  |  | 44.1 | 21.4 |  |
| sp\|P28907\|CD38\_HUMAN | P28907 | CD38\_HUMAN | CD38 | ADP-ribosyl cyclase/cyclic ADP-ribose hydrolase 1 | 300 | 34.328 | 2.292623e+04 | 9.887770e+05 |  | 168968.45 | 326756.3 | 493052.25 |  | 39.7 | 45.7 | 31.7 |
| sp\|P47736\|RPGP1\_HUMAN | P47736 | RPGP1\_HUMAN | RAP1GAP | Rap1 GTPase-activating protein 1 | 663 | 73.361 | 4.910126e+02 | 4.525588e+04 |  | 45255.88 |  |  |  | 40.0 |  |  |
| sp\|P49736\|MCM2\_HUMAN | P49736 | MCM2\_HUMAN | MCM2 | DNA replication licensing factor MCM2 | 904 | 101.896 | 4.423653e+02 | 5.663114e+04 |  | 56631.14 |  |  |  | 35.0 |  |  |
| sp\|P51648\|AL3A2\_HUMAN | P51648 | AL3A2\_HUMAN | ALDH3A2 | Aldehyde dehydrogenase family 3 member A2 | 485 | 54.848 | 1.489570e+06 | 1.026452e+08 |  | 86409712.0 | 16175653.0 | 59857.617 |  | 40.4 | 21.9 | 2.5 |
| sp\|P55327\|TPD52\_HUMAN | P55327 | TPD52\_HUMAN | TPD52 | Tumor protein D52 | 224 | 24.327 | 3.164071e+03 | 9.670566e+04 |  | 96705.664 |  |  |  | 19.2 |  |  |
| sp\|Q14165\|MLEC\_HUMAN | Q14165 | MLEC\_HUMAN | MLEC | Malectin | 292 | 32.234 | 1.891122e+04 | 7.658627e+05 |  | 214561.28 | 252203.81 | 299097.6 |  | 16.1 | 47.9 | 31.5 |
| sp\|Q5JSP0\|FGD3\_HUMAN | Q5JSP0 | FGD3\_HUMAN | FGD3 | FYVE, RhoGEF and PH domain-containing protein 3 | 725 | 79.401 | 4.076518e+02 | 4.066610e+04 |  | 40666.1 |  |  |  | 33.7 |  |  |
| sp\|Q6NXT6\|TAPT1\_HUMAN | Q6NXT6 | TAPT1\_HUMAN | TAPT1 | Transmembrane anterior posterior transformation protein 1 homolog | 567 | 64.260 | 6.616768e+03 | 5.341999e+05 |  | 106572.89 | 99699.4 | 327927.62 |  | 23.6 | 16.2 | 18.5 |
| sp\|Q7Z4V5\|HDGR2\_HUMAN | Q7Z4V5 | HDGR2\_HUMAN | HDGFL2 | Hepatoma-derived growth factor-related protein 2 | 671 | 74.317 | 6.232229e+02 | 5.819005e+04 |  | 58190.047 |  |  |  | 32.2 |  |  |
| sp\|Q8IWW6\|RHG12\_HUMAN | Q8IWW6 | RHG12\_HUMAN | ARHGAP12 | Rho GTPase-activating protein 12 | 846 | 96.254 | 3.150013e+02 | 3.809327e+04 |  | 38093.27 |  |  |  | 21.3 |  |  |
| sp\|Q8IWX8\|CHERP\_HUMAN | Q8IWX8 | CHERP\_HUMAN | CHERP | Calcium homeostasis endoplasmic reticulum protein | 916 | 103.702 | 3.940204e+02 | 5.133612e+04 |  | 51336.12 |  |  |  | 30.5 |  |  |
| sp\|Q8N142\|PURA1\_HUMAN | Q8N142 | PURA1\_HUMAN | ADSS1 | Adenylosuccinate synthetase isozyme 1 | 457 | 50.208 | 3.242241e+03 | 2.045198e+05 |  | 89722.67 | 87962.49 | 26834.63 |  | 29.8 | 12.5 | 5.7 |
| sp\|Q8NBJ7\|SUMF2\_HUMAN | Q8NBJ7 | SUMF2\_HUMAN | SUMF2 | Inactive C-alpha-formylglycine-generating enzyme 2 | 301 | 33.843 | 2.216040e+04 | 9.422445e+05 |  | 53275.496 | 748850.9 | 140118.14 |  | 28.2 | 35.2 | 24.9 |
| sp\|Q8WVB3\|HEXD\_HUMAN | Q8WVB3 | HEXD\_HUMAN | HEXD | Hexosaminidase D | 486 | 53.790 | 9.039307e+02 | 6.108771e+04 |  | 61087.71 |  |  |  | 22.0 |  |  |
| sp\|Q96AA3\|RFT1\_HUMAN | Q96AA3 | RFT1\_HUMAN | RFT1 | Protein RFT1 homolog | 541 | 60.335 | 6.680187e+03 | 5.063783e+05 |  | 244762.72 |  | 261615.56 |  | 24.6 |  | 10.0 |
| sp\|Q9BY42\|RTF2\_HUMAN | Q9BY42 | RTF2\_HUMAN | RTF2 | Replication termination factor 2 | 306 | 33.887 | 2.118344e+03 | 9.018760e+04 |  | 90187.6 |  |  |  | 31.7 |  |  |
| sp\|Q9GZM8\|NDEL1\_HUMAN | Q9GZM8 | NDEL1\_HUMAN | NDEL1 | Nuclear distribution protein nudE-like 1 | 345 | 38.375 | 2.163724e+04 | 1.043200e+06 |  | 1043199.6 |  |  |  | 52.8 |  |  |
| sp\|Q9H6E5\|STPAP\_HUMAN | Q9H6E5 | STPAP\_HUMAN | TUT1 | Speckle targeted PIP5K1A-regulated poly(A) polymerase | 874 | 93.847 | 4.301122e+04 | 5.071300e+06 |  | 5071300.5 |  |  |  | 33.1 |  |  |
| sp\|Q9HAS0\|NJMU\_HUMAN | Q9HAS0 | NJMU\_HUMAN | C17orf75 | Protein Njmu-R1 | 396 | 44.622 | 8.050847e+02 | 4.513441e+04 |  | 45134.41 |  |  |  | 18.7 |  |  |
| sp\|Q9Y508\|RN114\_HUMAN | Q9Y508 | RN114\_HUMAN | RNF114 | E3 ubiquitin-protein ligase RNF114 | 228 | 25.694 | 2.520641e+03 | 8.136917e+04 |  | 81369.17 |  |  |  | 62.7 |  |  |
| sp\|Q9Y6K9\|NEMO\_HUMAN | Q9Y6K9 | NEMO\_HUMAN | IKBKG | NF-kappa-B essential modulator | 419 | 48.198 | 1.890481e+03 | 1.144771e+05 |  | 114477.06 |  |  |  | 27.9 |  |  |
| sp\|P01889\|HLAB\_HUMAN | P01889 | HLAB\_HUMAN | HLA-B | HLA class I histocompatibility antigen, B alpha chain | 362 | 40.460 | 3.392764e+03 | 1.724633e+05 |  | 49768.258 | 122695.04 |  |  | 19.6 | 27.9 |  |
| sp\|O00160\|MYO1F\_HUMAN | O00160 | MYO1F\_HUMAN | MYO1F | Unconventional myosin-If | 1098 | 124.844 | 1.198632e+02 | 1.880056e+04 |  | 18800.557 |  |  |  | 30.0 |  |  |
| sp\|O15126\|SCAM1\_HUMAN | O15126 | SCAM1\_HUMAN | SCAMP1 | Secretory carrier-associated membrane protein 1 | 338 | 37.920 | 2.720180e+03 | 1.295935e+05 |  | 40718.363 | 6621.9194 | 82253.19 |  | 24.9 | 14.2 | 6.5 |
| sp\|O43572\|AKA10\_HUMAN | O43572 | AKA10\_HUMAN | AKAP10 | A-kinase anchor protein 10, mitochondrial | 662 | 73.818 | 5.384434e+02 | 4.993666e+04 |  | 49936.656 |  |  |  | 47.0 |  |  |
| sp\|O60307\|MAST3\_HUMAN | O60307 | MAST3\_HUMAN | MAST3 | Microtubule-associated serine/threonine-protein kinase 3 | 1309 | 143.137 | 3.577998e+02 | 6.434416e+04 |  | 64344.156 |  |  |  | 29.6 |  |  |
| sp\|O95486\|SC24A\_HUMAN | O95486 | SC24A\_HUMAN | SEC24A | Protein transport protein Sec24A | 1093 | 119.749 | 4.412454e+02 | 6.638489e+04 |  | 66384.89 |  |  |  | 20.0 |  |  |
| sp\|P07858\|CATB\_HUMAN | P07858 | CATB\_HUMAN | CTSB | Cathepsin B | 339 | 37.822 | 2.156950e+04 | 1.024948e+06 |  | 84502.734 | 940444.8 |  |  | 33.3 | 30.7 |  |
| sp\|P0DME0\|SETLP\_HUMAN | P0DME0 | SETLP\_HUMAN | SETSIP | Protein SETSIP | 302 | 34.882 | 2.931725e+03 | 1.284818e+05 |  | 128481.84 |  |  |  | 11.7 |  |  |
| sp\|P11169\|GTR3\_HUMAN | P11169 | GTR3\_HUMAN | SLC2A3 | Solute carrier family 2, facilitated glucose transporter member 3 | 496 | 53.924 | 1.034124e+05 | 7.006027e+06 |  | 1656393.2 | 792415.9 | 4557218.0 |  | 27.2 | 21.4 | 17.1 |
| sp\|Q9BX46\|RBM24\_HUMAN | Q9BX46 | RBM24\_HUMAN | RBM24 | RNA-binding protein 24 | 236 | 24.776 | 7.985773e+03 | 2.485795e+05 |  | 248579.5 |  |  |  | 15.3 |  |  |
| sp\|Q9H3Q1\|BORG4\_HUMAN | Q9H3Q1 | BORG4\_HUMAN | CDC42EP4 | Cdc42 effector protein 4 | 356 | 37.980 | 1.091847e+03 | 5.209954e+04 |  | 52099.54 |  |  |  | 50.8 |  |  |
| sp\|Q9HBH5\|RDH14\_HUMAN | Q9HBH5 | RDH14\_HUMAN | RDH14 | Retinol dehydrogenase 14 | 336 | 36.865 | 6.042953e+02 | 2.798856e+04 |  | 27988.562 |  |  |  | 15.8 |  |  |
| sp\|Q9HD15\|SRA1\_HUMAN | Q9HD15 | SRA1\_HUMAN | SRA1 | Steroid receptor RNA activator 1 | 224 | 24.400 | 3.981075e+03 | 1.220414e+05 |  | 122041.445 |  |  |  | 58.0 |  |  |
| sp\|Q9NRX5\|SERC1\_HUMAN | Q9NRX5 | SERC1\_HUMAN | SERINC1 | Serine incorporator 1 | 453 | 50.495 | 7.930852e+03 | 5.031360e+05 |  | 55422.957 | 73385.055 | 374327.97 |  | 22.3 | 13.2 | 6.2 |
| sp\|Q9NRZ7\|PLCC\_HUMAN | Q9NRZ7 | PLCC\_HUMAN | AGPAT3 | 1-acyl-sn-glycerol-3-phosphate acyltransferase gamma | 376 | 43.381 | 1.220055e+04 | 6.649607e+05 |  | 181783.72 |  | 483177.0 |  | 27.4 |  | 24.2 |
| sp\|Q9UKG1\|DP13A\_HUMAN | Q9UKG1 | DP13A\_HUMAN | APPL1 | DCC-interacting protein 13-alpha | 709 | 79.663 | 1.318895e+03 | 1.320031e+05 |  | 132003.1 |  |  |  | 28.2 |  |  |
| sp\|A0MZ66\|SHOT1\_HUMAN | A0MZ66 | SHOT1\_HUMAN | SHTN1 | Shootin-1 | 631 | 71.640 | 7.128772e+02 | 6.416341e+04 |  | 64163.41 |  |  |  | 44.5 |  |  |
| sp\|P07305\|H10\_HUMAN | P07305 | H10\_HUMAN | H1-0 | Histone H1.0 | 194 | 20.863 | 4.466172e+03 | 1.170656e+05 |  | 30253.14 | 86812.47 |  |  | 7.2 | 26.3 |  |
| sp\|P09622\|DLDH\_HUMAN | P09622 | DLDH\_HUMAN | DLD | Dihydrolipoyl dehydrogenase, mitochondrial | 509 | 54.177 | 7.637808e+03 | 5.198773e+05 |  | 72870.94 | 399603.28 | 47403.094 |  | 44.6 | 24.0 | 15.5 |
| sp\|P10643\|CO7\_HUMAN | P10643 | CO7\_HUMAN | C7 | Complement component C7 | 843 | 93.518 | 2.258713e+03 | 2.653832e+05 |  | 48852.152 | 206012.53 | 10518.561 |  | 29.2 | 27.4 | 15.7 |
| sp\|P98179\|RBM3\_HUMAN | P98179 | RBM3\_HUMAN | RBM3 | RNA-binding protein 3 | 157 | 17.170 | 7.298900e+03 | 1.574508e+05 |  | 157450.8 |  |  |  | 52.9 |  |  |
| sp\|Q58FG0\|HS905\_HUMAN | Q58FG0 | HS905\_HUMAN | HSP90AA5P | Putative heat shock protein HSP 90-alpha A5 | 334 | 38.738 | 2.225554e+05 | 1.083160e+07 |  | 10831595.0 |  |  |  | 46.4 |  |  |
| sp\|Q659C4\|LAR1B\_HUMAN | Q659C4 | LAR1B\_HUMAN | LARP1B | La-related protein 1B | 914 | 105.322 | 3.669117e+02 | 4.855096e+04 |  | 48550.96 |  |  |  | 42.3 |  |  |
| sp\|Q86U90\|YRDC\_HUMAN | Q86U90 | YRDC\_HUMAN | YRDC | YrdC domain-containing protein, mitochondrial | 279 | 29.328 | 4.638033e+03 | 1.708966e+05 |  | 159285.67 | 11610.946 |  |  | 38.0 | 29.7 |  |
| sp\|Q86WN1\|FCSD1\_HUMAN | Q86WN1 | FCSD1\_HUMAN | FCHSD1 | F-BAR and double SH3 domains protein 1 | 690 | 76.942 | 8.597669e+02 | 8.311154e+04 |  | 83111.54 |  |  |  | 34.6 |  |  |
| sp\|Q8IUR0\|TPPC5\_HUMAN | Q8IUR0 | TPPC5\_HUMAN | TRAPPC5 | Trafficking protein particle complex subunit 5 | 188 | 20.783 | 5.517096e+03 | 1.440575e+05 |  | 144057.53 |  |  |  | 36.7 |  |  |
| sp\|Q8N5M9\|JAGN1\_HUMAN | Q8N5M9 | JAGN1\_HUMAN | JAGN1 | Protein jagunal homolog 1 | 183 | 21.125 | 2.435150e+04 | 6.463080e+05 |  | 278164.94 | 93444.96 | 274698.1 |  | 18.6 | 13.1 | 13.1 |
| sp\|Q8NBM8\|PCYXL\_HUMAN | Q8NBM8 | PCYXL\_HUMAN | PCYOX1L | Prenylcysteine oxidase-like | 494 | 54.646 | 3.230747e+03 | 2.218086e+05 |  | 13797.65 | 208010.97 |  |  | 11.3 | 19.4 |  |
| sp\|Q92636\|FAN\_HUMAN | Q92636 | FAN\_HUMAN | NSMAF | Protein FAN | 917 | 104.372 | 5.262540e+02 | 6.900755e+04 |  | 69007.55 |  |  |  | 21.4 |  |  |
| sp\|Q92733\|PRCC\_HUMAN | Q92733 | PRCC\_HUMAN | PRCC | Proline-rich protein PRCC | 491 | 52.418 | 7.793218e+02 | 5.132328e+04 |  | 51323.28 |  |  |  | 34.8 |  |  |
| sp\|Q99439\|CNN2\_HUMAN | Q99439 | CNN2\_HUMAN | CNN2 | Calponin-2 | 309 | 33.697 | 1.026760e+03 | 4.346877e+04 |  | 43468.773 | 0.0 |  |  | 46.0 | 28.5 |  |
| sp\|Q9BX10\|GTPB2\_HUMAN | Q9BX10 | GTPB2\_HUMAN | GTPBP2 | GTP-binding protein 2 | 602 | 65.768 | 1.969088e+03 | 1.627035e+05 |  | 162703.52 |  |  |  | 27.7 |  |  |
| sp\|Q9BZF3\|OSBL6\_HUMAN | Q9BZF3 | OSBL6\_HUMAN | OSBPL6 | Oxysterol-binding protein-related protein 6 | 934 | 106.306 | 2.457458e+02 | 3.282170e+04 |  | 32821.7 |  |  |  | 28.3 |  |  |
| sp\|Q9H6E4\|CC134\_HUMAN | Q9H6E4 | CC134\_HUMAN | CCDC134 | Coiled-coil domain-containing protein 134 | 229 | 26.561 | 1.368933e+04 | 4.568184e+05 |  | 87675.625 | 369142.8 |  |  | 50.2 | 32.3 |  |
| sp\|Q9P2E3\|ZNFX1\_HUMAN | Q9P2E3 | ZNFX1\_HUMAN | ZNFX1 | NFX1-type zinc finger-containing protein 1 | 1918 | 220.227 | 0.000000e+00 | 0.000000e+00 |  | 0.0 |  |  |  | 30.3 |  |  |
| sp\|Q9UJX3\|APC7\_HUMAN | Q9UJX3 | APC7\_HUMAN | ANAPC7 | Anaphase-promoting complex subunit 7 | 565 | 63.133 | 3.550411e+02 | 2.816127e+04 |  | 28161.266 |  |  |  | 41.9 |  |  |
| sp\|Q9UK45\|LSM7\_HUMAN | Q9UK45 | LSM7\_HUMAN | LSM7 | U6 snRNA-associated Sm-like protein LSm7 | 103 | 11.602 | 6.452412e+03 | 9.405288e+04 |  | 94052.875 |  |  |  | 37.9 |  |  |
| sp\|Q9UM07\|PADI4\_HUMAN | Q9UM07 | PADI4\_HUMAN | PADI4 | Protein-arginine deiminase type-4 | 663 | 74.079 | 2.668115e+03 | 2.483229e+05 |  | 238962.48 | 9360.448 |  |  | 34.2 | 17.3 |  |
| sp\|Q15032\|R3HD1\_HUMAN | Q15032 | R3HD1\_HUMAN | R3HDM1 | R3H domain-containing protein 1 | 1099 | 120.696 | 1.969826e+04 | 2.987019e+06 |  | 2987019.0 |  |  |  | 25.6 |  |  |
| sp\|Q66K14\|TBC9B\_HUMAN | Q66K14 | TBC9B\_HUMAN | TBC1D9B | TBC1 domain family member 9B | 1250 | 140.525 | 3.147766e+02 | 5.557420e+04 |  | 55574.203 |  |  |  | 27.6 |  |  |
| sp\|Q9BPX7\|CG025\_HUMAN | Q9BPX7 | CG025\_HUMAN | C7orf25 | UPF0415 protein C7orf25 | 421 | 46.451 | 7.296619e+02 | 4.258277e+04 |  | 42582.766 |  |  |  | 28.7 |  |  |
| sp\|Q9NRD5\|PICK1\_HUMAN | Q9NRD5 | PICK1\_HUMAN | PICK1 | PRKCA-binding protein | 415 | 46.600 | 8.860595e+02 | 5.187594e+04 |  | 51875.938 |  |  |  | 21.4 |  |  |
| sp\|Q9UBF6\|RBX2\_HUMAN | Q9UBF6 | RBX2\_HUMAN | RNF7 | RING-box protein 2 | 113 | 12.683 | 2.332467e+04 | 3.716676e+05 |  | 371667.56 |  |  |  | 43.4 |  |  |
| sp\|A2RUC4\|TYW5\_HUMAN | A2RUC4 | TYW5\_HUMAN | TYW5 | tRNA wybutosine-synthesizing protein 5 | 315 | 36.548 | 4.590024e+03 | 2.107636e+05 |  | 210763.64 |  |  |  | 33.7 |  |  |
| sp\|P02745\|C1QA\_HUMAN | P02745 | C1QA\_HUMAN | C1QA | Complement C1q subcomponent subunit A | 245 | 26.017 | 1.206940e+04 | 3.945119e+05 |  | 394511.88 |  |  |  | 49.8 |  |  |
| sp\|P62081\|RS7\_HUMAN | P62081 | RS7\_HUMAN | RPS7 | 40S ribosomal protein S7 | 194 | 22.127 | 4.504736e+03 | 1.252302e+05 |  | 125230.195 |  |  |  | 19.6 |  |  |
| sp\|Q53RE8\|ANR39\_HUMAN | Q53RE8 | ANR39\_HUMAN | ANKRD39 | Ankyrin repeat domain-containing protein 39 | 183 | 19.651 | 1.596679e+03 | 3.942025e+04 |  | 39420.246 |  |  |  | 45.4 |  |  |
| sp\|Q5T4F4\|ZFY27\_HUMAN | Q5T4F4 | ZFY27\_HUMAN | ZFYVE27 | Protrudin | 411 | 45.843 | 5.647407e+03 | 3.252664e+05 |  | 44260.598 | 84192.43 | 196813.42 |  | 25.1 | 19.5 | 16.5 |
| sp\|Q9Y3E5\|PTH2\_HUMAN | Q9Y3E5 | PTH2\_HUMAN | PTRH2 | Peptidyl-tRNA hydrolase 2, mitochondrial | 179 | 19.194 | 7.238266e+02 | 1.745489e+04 |  | 17454.893 |  |  |  | 55.9 |  |  |
| sp\|P02765\|FETUA\_HUMAN | P02765 | FETUA\_HUMAN | AHSG | Alpha-2-HS-glycoprotein | 367 | 39.341 | 5.388864e+03 | 2.663543e+05 |  | 17219.668 | 99570.266 | 149564.38 |  | 38.1 | 18.5 | 5.7 |
| sp\|P13727\|PRG2\_HUMAN | P13727 | PRG2\_HUMAN | PRG2 | Bone marrow proteoglycan | 222 | 25.206 | 8.571812e+02 | 2.714524e+04 |  | 27145.242 |  |  |  | 50.5 |  |  |
| sp\|Q00536\|CDK16\_HUMAN | Q00536 | CDK16\_HUMAN | CDK16 | Cyclin-dependent kinase 16 | 496 | 55.716 | 9.773729e+02 | 6.841595e+04 |  | 68415.95 |  |  |  | 28.2 |  |  |
| sp\|Q12802\|AKP13\_HUMAN | Q12802 | AKP13\_HUMAN | AKAP13 | A-kinase anchor protein 13 | 2813 | 307.550 | 2.457839e+01 | 9.496999e+03 |  | 9496.999 |  |  |  | 31.2 |  |  |
| sp\|Q14596\|NBR1\_HUMAN | Q14596 | NBR1\_HUMAN | NBR1 | Next to BRCA1 gene 1 protein | 966 | 107.413 | 2.352934e+02 | 3.175293e+04 |  | 31752.926 |  |  |  | 23.1 |  |  |
| sp\|Q8N138\|ORML3\_HUMAN | Q8N138 | ORML3\_HUMAN | ORMDL3 | ORM1-like protein 3 | 153 | 17.495 | 2.677901e+04 | 5.886073e+05 |  | 292950.88 |  | 295656.47 |  | 28.8 |  | 22.2 |
| sp\|Q8N2K0\|ABD12\_HUMAN | Q8N2K0 | ABD12\_HUMAN | ABHD12 | Lysophosphatidylserine lipase ABHD12 | 398 | 45.097 | 2.083348e+03 | 1.180393e+05 |  | 58430.555 |  | 59608.723 |  | 46.0 |  | 16.3 |
| sp\|Q96N21\|AP4AT\_HUMAN | Q96N21 | AP4AT\_HUMAN | TEPSIN | AP-4 complex accessory subunit Tepsin | 525 | 55.137 | 1.735774e+03 | 1.202413e+05 |  | 120241.26 |  |  |  | 35.4 |  |  |
| sp\|P19086\|GNAZ\_HUMAN | P19086 | GNAZ\_HUMAN | GNAZ | Guanine nucleotide-binding protein G(z) subunit alpha | 355 | 40.924 | 5.550078e+03 | 2.853608e+05 |  | 158407.42 |  | 126953.375 |  | 33.8 |  | 20.8 |
| sp\|Q8NFH4\|NUP37\_HUMAN | Q8NFH4 | NUP37\_HUMAN | NUP37 | Nucleoporin Nup37 | 326 | 36.708 | 7.808533e+02 | 3.601201e+04 |  | 36012.008 |  |  |  | 25.5 |  |  |
| sp\|Q9NZI7\|UBIP1\_HUMAN | Q9NZI7 | UBIP1\_HUMAN | UBP1 | Upstream-binding protein 1 | 540 | 60.491 | 1.416589e+03 | 1.076594e+05 |  | 107659.35 |  |  |  | 15.0 |  |  |
| sp\|O75477\|ERLN1\_HUMAN | O75477 | ERLN1\_HUMAN | ERLIN1 | Erlin-1 | 348 | 39.171 | 4.608980e+04 | 2.268227e+06 |  | 89658.375 | 1709635.4 | 468933.7 |  | 42.0 | 30.5 | 29.3 |
| sp\|P05114\|HMGN1\_HUMAN | P05114 | HMGN1\_HUMAN | HMGN1 | Non-histone chromosomal protein HMG-14 | 100 | 10.659 | 1.543205e+03 | 2.066604e+04 |  | 20666.043 |  |  |  | 47.0 |  |  |
| sp\|Q92504\|S39A7\_HUMAN | Q92504 | S39A7\_HUMAN | SLC39A7 | Zinc transporter SLC39A7 | 469 | 50.118 | 2.085490e+03 | 1.313164e+05 |  | 70410.64 | 24574.64 | 36331.16 |  | 13.4 | 4.7 | 5.3 |
| sp\|Q96KP1\|EXOC2\_HUMAN | Q96KP1 | EXOC2\_HUMAN | EXOC2 | Exocyst complex component 2 | 924 | 104.066 | 3.882508e+02 | 5.076195e+04 |  | 50761.953 |  |  |  | 26.4 |  |  |
| sp\|Q9H1P3\|OSBL2\_HUMAN | Q9H1P3 | OSBL2\_HUMAN | OSBPL2 | Oxysterol-binding protein-related protein 2 | 480 | 55.201 | 6.700365e+02 | 4.646892e+04 |  | 46468.918 |  |  |  | 39.4 |  |  |
| sp\|Q9NUM3\|S39A9\_HUMAN | Q9NUM3 | S39A9\_HUMAN | SLC39A9 | Zinc transporter ZIP9 | 307 | 32.251 | 9.591891e+02 | 3.886553e+04 |  | 38865.527 |  |  |  | 4.9 |  |  |
| sp\|Q9Y296\|TPPC4\_HUMAN | Q9Y296 | TPPC4\_HUMAN | TRAPPC4 | Trafficking protein particle complex subunit 4 | 219 | 24.340 | 3.659841e+03 | 1.119180e+05 |  | 111918.016 |  |  |  | 55.7 |  |  |
| sp\|O95985\|TOP3B\_HUMAN | O95985 | TOP3B\_HUMAN | TOP3B | DNA topoisomerase 3-beta-1 | 862 | 96.662 | 4.467375e+03 | 5.425320e+05 |  | 542532.0 |  |  |  | 43.6 |  |  |
| sp\|Q86UK7\|ZN598\_HUMAN | Q86UK7 | ZN598\_HUMAN | ZNF598 | E3 ubiquitin-protein ligase ZNF598 | 904 | 98.637 | 1.626820e+02 | 2.016027e+04 |  | 20160.273 |  |  |  | 39.5 |  |  |
| sp\|Q8NAG6\|ANKL1\_HUMAN | Q8NAG6 | ANKL1\_HUMAN | ANKLE1 | Ankyrin repeat and LEM domain-containing protein 1 | 615 | 66.890 | 1.917655e+03 | 1.611569e+05 |  | 161156.86 |  |  |  | 25.9 |  |  |
| sp\|Q96BH1\|RNF25\_HUMAN | Q96BH1 | RNF25\_HUMAN | RNF25 | E3 ubiquitin-protein ligase RNF25 | 459 | 51.219 | 5.719588e+02 | 3.680552e+04 |  | 36805.523 |  |  |  | 49.9 |  |  |
| sp\|Q96DZ1\|ERLEC\_HUMAN | Q96DZ1 | ERLEC\_HUMAN | ERLEC1 | Endoplasmic reticulum lectin 1 | 483 | 54.858 | 5.999338e+03 | 4.134857e+05 |  | 15083.77 | 398401.9 |  |  | 42.4 | 35.6 |  |
| sp\|O60825\|F262\_HUMAN | O60825 | F262\_HUMAN | PFKFB2 | 6-phosphofructo-2-kinase/fructose-2,6-bisphosphatase 2 | 505 | 58.477 | 5.062060e+02 | 3.719029e+04 |  | 37190.285 |  |  |  | 44.8 |  |  |
| sp\|P31949\|S10AB\_HUMAN | P31949 | S10AB\_HUMAN | S100A11 | Protein S100-A11 | 105 | 11.740 | 9.485248e+04 | 1.399052e+06 |  | 117698.05 | 719192.75 | 562161.5 |  | 61.9 | 44.8 | 15.2 |
| sp\|P33316\|DUT\_HUMAN | P33316 | DUT\_HUMAN | DUT | Deoxyuridine 5'-triphosphate nucleotidohydrolase, mitochondrial | 252 | 26.563 | 8.410919e+02 | 2.806970e+04 |  | 28069.697 |  |  |  | 69.0 |  |  |
| sp\|Q9BTT0\|AN32E\_HUMAN | Q9BTT0 | AN32E\_HUMAN | ANP32E | Acidic leucine-rich nuclear phosphoprotein 32 family member E | 268 | 30.692 | 1.552167e+03 | 5.985228e+04 |  | 59852.28 |  |  |  | 10.1 |  |  |
| sp\|Q86VQ1\|GLCI1\_HUMAN | Q86VQ1 | GLCI1\_HUMAN | GLCCI1 | Glucocorticoid-induced transcript 1 protein | 547 | 58.024 | 3.949073e+02 | 2.878857e+04 |  | 28788.566 |  |  |  | 50.1 |  |  |
| sp\|Q8NFD5\|ARI1B\_HUMAN | Q8NFD5 | ARI1B\_HUMAN | ARID1B | AT-rich interactive domain-containing protein 1B | 2236 | 243.943 | 1.441435e+02 | 4.417743e+04 |  | 44177.43 |  |  |  | 19.5 |  |  |
| sp\|O60284\|ST18\_HUMAN | O60284 | ST18\_HUMAN | ST18 | Suppression of tumorigenicity 18 protein | 1047 | 115.155 | 6.633431e+02 | 9.597059e+04 |  | 95970.586 |  |  |  | 33.4 |  |  |
| sp\|P01031\|CO5\_HUMAN | P01031 | CO5\_HUMAN | C5 | Complement C5 | 1676 | 188.305 | 9.486705e+02 | 2.244370e+05 |  | 42398.656 | 182038.31 |  |  | 22.5 | 13.5 |  |
| sp\|Q9UQN3\|CHM2B\_HUMAN | Q9UQN3 | CHM2B\_HUMAN | CHMP2B | Charged multivesicular body protein 2b | 213 | 23.907 | 5.641899e+03 | 1.694601e+05 |  | 169460.14 |  |  |  | 45.5 |  |  |
| sp\|P55084\|ECHB\_HUMAN | P55084 | ECHB\_HUMAN | HADHB | Trifunctional enzyme subunit beta, mitochondrial | 474 | 51.294 | 2.355830e+03 | 1.518195e+05 |  | 81098.586 | 70720.95 |  |  | 36.3 | 23.0 |  |
| sp\|Q7Z6E9\|RBBP6\_HUMAN | Q7Z6E9 | RBBP6\_HUMAN | RBBP6 | E3 ubiquitin-protein ligase RBBP6 | 1792 | 201.564 | 2.911643e+02 | 7.373407e+04 |  | 73734.07 |  |  |  | 33.7 |  |  |
| sp\|Q10570\|CPSF1\_HUMAN | Q10570 | CPSF1\_HUMAN | CPSF1 | Cleavage and polyadenylation specificity factor subunit 1 | 1443 | 160.884 | 2.292629e+04 | 4.634084e+06 |  | 4634083.5 |  |  |  | 29.0 |  |  |
| sp\|Q96HR9\|REEP6\_HUMAN | Q96HR9 | REEP6\_HUMAN | REEP6 | Receptor expression-enhancing protein 6 | 211 | 23.418 | 2.940787e+03 | 8.652280e+04 |  | 86522.805 |  |  |  | 42.7 |  |  |
| sp\|Q9Y679\|AUP1\_HUMAN | Q9Y679 | AUP1\_HUMAN | AUP1 | Lipid droplet-regulating VLDL assembly factor AUP1 | 410 | 45.787 | 1.820237e+03 | 1.047098e+05 |  | 104709.8 |  |  |  | 39.5 |  |  |
| sp\|P12429\|ANXA3\_HUMAN | P12429 | ANXA3\_HUMAN | ANXA3 | Annexin A3 | 323 | 36.375 | 5.959309e+04 | 2.723429e+06 |  | 2723429.2 |  |  |  | 39.3 |  |  |
| sp\|P24752\|THIL\_HUMAN | P24752 | THIL\_HUMAN | ACAT1 | Acetyl-CoA acetyltransferase, mitochondrial | 427 | 45.200 | 4.578251e+02 | 2.599891e+04 |  | 18732.604 | 7266.3105 |  |  | 32.6 | 14.3 |  |
| sp\|Q6UX06\|OLFM4\_HUMAN | Q6UX06 | OLFM4\_HUMAN | OLFM4 | Olfactomedin-4 | 510 | 57.280 | 2.026482e+03 | 1.458354e+05 |  | 29842.635 | 82778.21 | 33214.566 |  | 33.5 | 6.5 | 10.0 |
| sp\|Q8WVR3\|TPC14\_HUMAN | Q8WVR3 | TPC14\_HUMAN | TRAPPC14 | Trafficking protein particle complex subunit 14 | 580 | 62.597 | 4.147533e+02 | 3.261824e+04 |  | 23091.479 | 9526.763 |  |  | 41.0 | 18.8 |  |
| sp\|Q14669\|TRIPC\_HUMAN | Q14669 | TRIPC\_HUMAN | TRIP12 | E3 ubiquitin-protein ligase TRIP12 | 1992 | 220.434 | 1.444562e+02 | 4.000663e+04 |  | 17665.244 | 5667.6514 | 16673.738 |  | 26.6 | 17.3 | 11.4 |
| sp\|Q9UMZ2\|SYNRG\_HUMAN | Q9UMZ2 | SYNRG\_HUMAN | SYNRG | Synergin gamma | 1314 | 140.654 | 2.947914e+02 | 5.209355e+04 |  | 34065.816 | 18027.738 |  |  | 27.9 | 15.4 |  |
| sp\|Q9Y3E7\|CHMP3\_HUMAN | Q9Y3E7 | CHMP3\_HUMAN | CHMP3 | Charged multivesicular body protein 3 | 222 | 25.073 | 1.276523e+04 | 4.021168e+05 |  | 257975.75 | 144141.08 |  |  | 38.7 | 25.2 |  |
| sp\|P36404\|ARL2\_HUMAN | P36404 | ARL2\_HUMAN | ARL2 | ADP-ribosylation factor-like protein 2 | 184 | 20.878 | 1.208966e+04 | 3.171174e+05 |  | 317117.44 |  |  |  | 15.8 |  |  |
| sp\|Q96G28\|CFA36\_HUMAN | Q96G28 | CFA36\_HUMAN | CFAP36 | Cilia- and flagella-associated protein 36 | 342 | 39.447 | 4.304049e+02 | 2.133086e+04 |  | 21330.861 |  |  |  | 41.2 |  |  |
| sp\|Q9UHB9\|SRP68\_HUMAN | Q9UHB9 | SRP68\_HUMAN | SRP68 | Signal recognition particle subunit SRP68 | 627 | 70.730 | 3.645721e+02 | 3.239696e+04 |  | 32396.959 |  |  |  | 35.7 |  |  |
| sp\|Q6P179\|ERAP2\_HUMAN | Q6P179 | ERAP2\_HUMAN | ERAP2 | Endoplasmic reticulum aminopeptidase 2 | 960 | 110.462 | 1.237294e+03 | 1.717130e+05 |  | 27636.186 | 128056.83 | 16019.95 |  | 21.7 | 21.1 | 8.0 |
| sp\|O15042\|SR140\_HUMAN | O15042 | SR140\_HUMAN | U2SURP | U2 snRNP-associated SURP motif-containing protein | 1029 | 118.292 | 2.210063e+02 | 3.284560e+04 |  | 32845.598 |  |  |  | 30.6 |  |  |
| sp\|O75449\|KTNA1\_HUMAN | O75449 | KTNA1\_HUMAN | KATNA1 | Katanin p60 ATPase-containing subunit A1 | 491 | 55.965 | 1.479601e+03 | 1.040347e+05 |  | 104034.7 |  |  |  | 40.3 |  |  |
| sp\|Q9H9A5\|CNO10\_HUMAN | Q9H9A5 | CNO10\_HUMAN | CNOT10 | CCR4-NOT transcription complex subunit 10 | 744 | 82.310 | 3.332543e+02 | 3.446240e+04 |  | 34462.4 |  |  |  | 29.7 |  |  |
| sp\|O60343\|TBCD4\_HUMAN | O60343 | TBCD4\_HUMAN | TBC1D4 | TBC1 domain family member 4 | 1298 | 146.563 | 4.901559e+01 | 9.025593e+03 |  | 9025.593 |  |  |  | 25.8 |  |  |
| sp\|Q9H1Z4\|WDR13\_HUMAN | Q9H1Z4 | WDR13\_HUMAN | WDR13 | WD repeat-containing protein 13 | 485 | 53.715 | 8.066139e+02 | 5.443502e+04 |  | 54435.023 |  |  |  | 29.1 |  |  |
| sp\|P29374\|ARI4A\_HUMAN | P29374 | ARI4A\_HUMAN | ARID4A | AT-rich interactive domain-containing protein 4A | 1257 | 142.752 | 8.259969e+03 | 1.481418e+06 |  | 1481418.5 |  |  |  | 25.5 |  |  |
| sp\|Q05084\|ICA69\_HUMAN | Q05084 | ICA69\_HUMAN | ICA1 | Islet cell autoantigen 1 | 483 | 54.645 | 5.099971e+02 | 3.501348e+04 |  | 35013.48 |  |  |  | 34.0 |  |  |
| sp\|Q8NDI1\|EHBP1\_HUMAN | Q8NDI1 | EHBP1\_HUMAN | EHBP1 | EH domain-binding protein 1 | 1231 | 140.017 | 3.795484e+02 | 6.676750e+04 |  | 66767.5 |  |  |  | 27.3 |  |  |
| sp\|Q9NRR3\|C42S2\_HUMAN | Q9NRR3 | C42S2\_HUMAN | CDC42SE2 | CDC42 small effector protein 2 | 84 | 9.223 | 1.583432e+04 | 1.834801e+05 |  | 66567.3 |  | 116912.77 |  | 46.4 |  | 22.6 |
| sp\|Q96AX9\|MIB2\_HUMAN | Q96AX9 | MIB2\_HUMAN | MIB2 | E3 ubiquitin-protein ligase MIB2 | 955 | 103.658 | 6.695660e+01 | 8.719938e+03 |  | 8719.938 |  |  |  | 21.2 |  |  |
| sp\|P62263\|RS14\_HUMAN | P62263 | RS14\_HUMAN | RPS14 | 40S ribosomal protein S14 | 151 | 16.273 | 2.312183e+04 | 4.727234e+05 |  | 114100.14 | 143584.17 | 215039.05 |  | 44.4 | 23.8 | 16.6 |
| sp\|Q9H3S4\|TPK1\_HUMAN | Q9H3S4 | TPK1\_HUMAN | TPK1 | Thiamin pyrophosphokinase 1 | 243 | 27.265 | 1.496480e+04 | 5.126177e+05 |  | 512617.66 |  |  |  | 11.1 |  |  |
| sp\|Q9HAU5\|RENT2\_HUMAN | Q9HAU5 | RENT2\_HUMAN | UPF2 | Regulator of nonsense transcripts 2 | 1272 | 147.810 | 1.354887e+03 | 2.516077e+05 |  | 251607.7 |  |  |  | 22.0 |  |  |
| sp\|O60264\|SMCA5\_HUMAN | O60264 | SMCA5\_HUMAN | SMARCA5 | SWI/SNF-related matrix-associated actin-dependent regulator of chromatin subfamily A member 5 | 1052 | 121.905 | 1.922769e+02 | 2.944867e+04 |  | 29448.672 |  |  |  | 34.9 |  |  |
| sp\|O95159\|ZFPL1\_HUMAN | O95159 | ZFPL1\_HUMAN | ZFPL1 | Zinc finger protein-like 1 | 310 | 34.114 | 1.303955e+03 | 5.588720e+04 |  | 55887.203 |  |  |  | 31.0 |  |  |
| sp\|Q15121\|PEA15\_HUMAN | Q15121 | PEA15\_HUMAN | PEA15 | Astrocytic phosphoprotein PEA-15 | 130 | 15.040 | 2.249152e+04 | 4.249950e+05 |  | 277898.72 | 84607.5 | 62488.793 |  | 33.8 | 33.8 | 13.1 |
| sp\|Q9BRT9\|SLD5\_HUMAN | Q9BRT9 | SLD5\_HUMAN | GINS4 | DNA replication complex GINS protein SLD5 | 223 | 26.047 | 3.499331e+03 | 1.145144e+05 |  | 114514.35 |  |  |  | 24.7 |  |  |
| sp\|P02786\|TFR1\_HUMAN | P02786 | TFR1\_HUMAN | TFRC | Transferrin receptor protein 1 | 760 | 84.871 | 3.557157e+02 | 3.792970e+04 |  | 33720.387 |  | 4209.3174 |  | 22.8 |  | 12.1 |
| sp\|Q68DH5\|LMBD2\_HUMAN | Q68DH5 | LMBD2\_HUMAN | LMBRD2 | G-protein coupled receptor-associated protein LMBRD2 | 695 | 81.172 | 2.202799e+03 | 2.246458e+05 |  | 73889.414 | 24729.96 | 126026.45 |  | 19.9 | 10.2 | 7.8 |
| sp\|Q8TAA3\|PSMA8\_HUMAN | Q8TAA3 | PSMA8\_HUMAN | PSMA8 | Proteasome subunit alpha-type 8 | 256 | 28.530 | 2.183406e+05 | 7.826244e+06 |  | 7826244.5 |  |  |  | 48.8 |  |  |
| sp\|P04156\|PRIO\_HUMAN | P04156 | PRIO\_HUMAN | PRNP | Major prion protein | 253 | 27.661 | 4.831259e+04 | 1.678979e+06 |  | 218176.16 | 532516.1 | 928287.0 |  | 29.6 | 17.4 | 19.8 |
| sp\|P63218\|GBG5\_HUMAN | P63218 | GBG5\_HUMAN | GNG5 | Guanine nucleotide-binding protein G(I)/G(S)/G(O) subunit gamma-5 | 68 | 7.318 | 6.354441e+05 | 5.842341e+06 |  | 421753.6 | 2218463.2 | 3202124.2 |  | 89.7 | 41.2 | 41.2 |
| sp\|Q05315\|LEG10\_HUMAN | Q05315 | LEG10\_HUMAN | CLC | Galectin-10 | 142 | 16.453 | 9.922782e+02 | 2.051142e+04 |  | 20511.418 |  |  |  | 71.8 |  |  |
| sp\|Q96P50\|ACAP3\_HUMAN | Q96P50 | ACAP3\_HUMAN | ACAP3 | Arf-GAP with coiled-coil, ANK repeat and PH domain-containing protein 3 | 834 | 92.495 | 1.019642e+03 | 1.184904e+05 |  | 118490.39 |  |  |  | 35.4 |  |  |
| sp\|Q9NRZ5\|PLCD\_HUMAN | Q9NRZ5 | PLCD\_HUMAN | AGPAT4 | 1-acyl-sn-glycerol-3-phosphate acyltransferase delta | 378 | 44.021 | 4.593400e+03 | 2.540454e+05 |  | 72413.195 |  | 181632.2 |  | 34.9 |  | 19.3 |
| sp\|P62070\|RRAS2\_HUMAN | P62070 | RRAS2\_HUMAN | RRAS2 | Ras-related protein R-Ras2 | 204 | 23.400 | 1.057009e+04 | 3.107504e+05 |  | 200618.81 | 58240.992 | 51890.555 |  | 27.0 | 37.3 | 15.7 |
| sp\|Q16698\|DECR\_HUMAN | Q16698 | DECR\_HUMAN | DECR1 | 2,4-dienoyl-CoA reductase [(3E)-enoyl-CoA-producing], mitochondrial | 335 | 36.068 | 3.886220e+03 | 1.761029e+05 |  | 68552.484 | 72705.336 | 34845.11 |  | 61.2 | 29.9 | 20.9 |
| sp\|Q9H1I8\|ASCC2\_HUMAN | Q9H1I8 | ASCC2\_HUMAN | ASCC2 | Activating signal cointegrator 1 complex subunit 2 | 757 | 86.360 | 5.824308e+02 | 6.319375e+04 |  | 63193.746 |  |  |  | 19.3 |  |  |
| sp\|Q13542\|4EBP2\_HUMAN | Q13542 | 4EBP2\_HUMAN | EIF4EBP2 | Eukaryotic translation initiation factor 4E-binding protein 2 | 120 | 12.939 | 9.194347e+03 | 1.494647e+05 |  | 114468.94 | 19803.682 | 15192.124 |  | 23.3 | 12.5 | 12.5 |
| sp\|Q9Y4E1\|WAC2C\_HUMAN | Q9Y4E1 | WAC2C\_HUMAN | WASHC2C | WASH complex subunit 2C | 1320 | 147.154 | 3.164198e+02 | 5.849960e+04 |  | 58499.6 |  |  |  | 27.3 |  |  |
| sp\|O60613\|SEP15\_HUMAN | O60613 | SEP15\_HUMAN | SELENOF | Selenoprotein F | 165 | 18.092 | 2.749547e+03 | 6.249782e+04 |  | 62497.816 |  |  |  | 34.5 |  |  |
| sp\|Q5JXB2\|UE2NL\_HUMAN | Q5JXB2 | UE2NL\_HUMAN | UBE2NL | Putative ubiquitin-conjugating enzyme E2 N-like | 153 | 17.377 | 1.015928e+06 | 2.217967e+07 |  | 22179668.0 |  |  |  | 45.1 |  |  |
| sp\|P61619\|S61A1\_HUMAN | P61619 | S61A1\_HUMAN | SEC61A1 | Protein transport protein Sec61 subunit alpha isoform 1 | 476 | 52.265 | 1.293593e+04 | 8.494267e+05 |  | 74918.19 | 60656.98 | 713851.5 |  | 23.9 | 6.7 | 19.1 |
| sp\|Q8WXG6\|MADD\_HUMAN | Q8WXG6 | MADD\_HUMAN | MADD | MAP kinase-activating death domain protein | 1647 | 183.303 | 4.315294e+01 | 9.937956e+03 |  | 9937.956 |  |  |  | 25.9 |  |  |
| sp\|P02746\|C1QB\_HUMAN | P02746 | C1QB\_HUMAN | C1QB | Complement C1q subcomponent subunit B | 253 | 26.722 | 7.725990e+03 | 2.593822e+05 |  | 99533.07 | 159849.17 |  |  | 54.2 | 27.7 |  |
| sp\|P49419\|AL7A1\_HUMAN | P49419 | AL7A1\_HUMAN | ALDH7A1 | Alpha-aminoadipic semialdehyde dehydrogenase | 539 | 58.487 | 1.549957e+03 | 1.138928e+05 |  | 58817.977 | 13184.98 | 41889.816 |  | 33.6 | 18.2 | 10.9 |
| sp\|Q8NEU8\|DP13B\_HUMAN | Q8NEU8 | DP13B\_HUMAN | APPL2 | DCC-interacting protein 13-beta | 664 | 74.493 | 1.356195e+03 | 1.269272e+05 |  | 100440.41 | 26486.82 |  |  | 35.8 | 7.5 |  |
| sp\|Q9Y3A3\|PHOCN\_HUMAN | Q9Y3A3 | PHOCN\_HUMAN | MOB4 | MOB-like protein phocein | 225 | 26.032 | 1.764201e+03 | 5.769958e+04 |  | 57699.58 |  |  |  | 44.9 |  |  |
| sp\|P62312\|LSM6\_HUMAN | P62312 | LSM6\_HUMAN | LSM6 | U6 snRNA-associated Sm-like protein LSm6 | 80 | 9.128 | 1.512203e+04 | 1.734215e+05 |  | 42188.6 | 131232.9 |  |  | 27.5 | 42.5 |  |
| sp\|Q6ZTN6\|AN13D\_HUMAN | Q6ZTN6 | AN13D\_HUMAN | ANKRD13D | Ankyrin repeat domain-containing protein 13D | 518 | 68.172 | 2.780447e+04 | 2.381430e+06 |  | 2381429.8 |  |  |  | 21.4 |  |  |
| sp\|P06850\|CRF\_HUMAN | P06850 | CRF\_HUMAN | CRH | Corticoliberin | 196 | 21.422 | 5.676680e+04 | 1.527818e+06 |  | 1527817.8 |  |  |  | 44.4 |  |  |
| sp\|Q9NYL9\|TMOD3\_HUMAN | Q9NYL9 | TMOD3\_HUMAN | TMOD3 | Tropomodulin-3 | 352 | 39.595 | 3.772862e+04 | 1.876845e+06 |  | 52200.21 | 1824644.8 |  |  | 32.7 | 43.5 |  |
| sp\|P48454\|PP2BC\_HUMAN | P48454 | PP2BC\_HUMAN | PPP3CC | Serine/threonine-protein phosphatase 2B catalytic subunit gamma isoform | 512 | 58.129 | 4.809763e+02 | 3.512641e+04 |  | 35126.406 |  |  |  | 36.3 |  |  |
| sp\|Q8NHG8\|ZNRF2\_HUMAN | Q8NHG8 | ZNRF2\_HUMAN | ZNRF2 | E3 ubiquitin-protein ligase ZNRF2 | 242 | 24.115 | 2.478412e+04 | 7.508928e+05 |  | 750892.75 |  |  |  | 57.9 |  |  |
| sp\|Q2TAM9\|TUSC1\_HUMAN | Q2TAM9 | TUSC1\_HUMAN | TUSC1 | Tumor suppressor candidate gene 1 protein | 212 | 23.390 | 1.034161e+03 | 3.039035e+04 |  | 30390.352 |  |  |  | 53.3 |  |  |
| sp\|Q8N806\|UBR7\_HUMAN | Q8N806 | UBR7\_HUMAN | UBR7 | Putative E3 ubiquitin-protein ligase UBR7 | 425 | 47.999 | 5.222902e+02 | 3.149642e+04 |  | 31496.42 |  |  |  | 25.9 |  |  |
| sp\|Q9P289\|STK26\_HUMAN | Q9P289 | STK26\_HUMAN | STK26 | Serine/threonine-protein kinase 26 | 416 | 46.529 | 9.833300e+03 | 5.748310e+05 |  | 574831.0 |  |  |  | 30.3 |  |  |
| sp\|Q9Y3I0\|RTCB\_HUMAN | Q9Y3I0 | RTCB\_HUMAN | RTCB | RNA-splicing ligase RtcB homolog | 505 | 55.210 | 8.189562e+02 | 5.680618e+04 |  | 56806.176 |  |  |  | 36.8 |  |  |
| sp\|Q86XP3\|DDX42\_HUMAN | Q86XP3 | DDX42\_HUMAN | DDX42 | ATP-dependent RNA helicase DDX42 | 938 | 102.975 | 3.868384e+02 | 5.004705e+04 |  | 50047.05 |  |  |  | 31.3 |  |  |
| sp\|Q86TB9\|PATL1\_HUMAN | Q86TB9 | PATL1\_HUMAN | PATL1 | Protein PAT1 homolog 1 | 770 | 86.850 | 8.019879e+02 | 8.750941e+04 |  | 87509.414 |  |  |  | 33.0 |  |  |
| sp\|Q13188\|STK3\_HUMAN | Q13188 | STK3\_HUMAN | STK3 | Serine/threonine-protein kinase 3 | 491 | 56.301 | 1.287329e+03 | 9.105897e+04 |  | 91058.97 |  |  |  | 45.6 |  |  |
| sp\|Q02978\|M2OM\_HUMAN | Q02978 | M2OM\_HUMAN | SLC25A11 | Mitochondrial 2-oxoglutarate/malate carrier protein | 314 | 34.062 | 2.868783e+03 | 1.227679e+05 |  | 51437.516 | 71330.43 |  |  | 37.6 | 25.2 |  |
| sp\|Q9ULU4\|PKCB1\_HUMAN | Q9ULU4 | PKCB1\_HUMAN | ZMYND8 | Protein kinase C-binding protein 1 | 1186 | 131.692 | 5.057432e+01 | 8.367709e+03 |  | 8367.709 |  |  |  | 31.6 |  |  |
| sp\|Q12972\|PP1R8\_HUMAN | Q12972 | PP1R8\_HUMAN | PPP1R8 | Nuclear inhibitor of protein phosphatase 1 | 351 | 38.479 | 1.298151e+03 | 6.275755e+04 |  | 62757.55 |  |  |  | 29.6 |  |  |
| sp\|O75781\|PALM\_HUMAN | O75781 | PALM\_HUMAN | PALM | Paralemmin-1 | 387 | 42.076 | 2.017060e+03 | 1.066278e+05 |  | 44250.5 | 62377.312 |  |  | 55.3 | 35.7 |  |
| sp\|P17661\|DESM\_HUMAN | P17661 | DESM\_HUMAN | DES | Desmin | 470 | 53.536 | 5.202773e+03 | 3.499435e+05 |  | 113127.39 | 236816.1 |  |  | 38.7 | 35.1 |  |
| sp\|P07996\|TSP1\_HUMAN | P07996 | TSP1\_HUMAN | THBS1 | Thrombospondin-1 | 1170 | 129.383 | 2.702239e+02 | 4.392564e+04 |  | 43925.64 |  |  |  | 29.8 |  |  |
| sp\|Q7Z3J2\|VP35L\_HUMAN | Q7Z3J2 | VP35L\_HUMAN | VPS35L | VPS35 endosomal protein-sorting factor-like | 963 | 109.563 | 2.857352e+03 | 3.933189e+05 |  | 393318.88 |  |  |  | 22.7 |  |  |
| sp\|Q96ME1\|FXL18\_HUMAN | Q96ME1 | FXL18\_HUMAN | FBXL18 | F-box/LRR-repeat protein 18 | 805 | 78.919 | 3.880623e+02 | 3.847691e+04 |  | 38476.906 |  |  |  | 35.3 |  |  |
| sp\|O14974\|MYPT1\_HUMAN | O14974 | MYPT1\_HUMAN | PPP1R12A | Protein phosphatase 1 regulatory subunit 12A | 1030 | 115.281 | 2.023733e+01 | 2.931083e+03 |  | 2931.0833 |  |  |  | 25.5 |  |  |
| sp\|Q15041\|AR6P1\_HUMAN | Q15041 | AR6P1\_HUMAN | ARL6IP1 | ADP-ribosylation factor-like protein 6-interacting protein 1 | 203 | 23.363 | 1.879361e+04 | 5.516402e+05 |  | 551640.25 |  |  |  | 13.8 |  |  |
| sp\|Q96CS3\|FAF2\_HUMAN | Q96CS3 | FAF2\_HUMAN | FAF2 | FAS-associated factor 2 | 445 | 52.623 | 2.237406e+03 | 1.479236e+05 |  | 41712.71 | 106210.92 |  |  | 25.8 | 17.3 |  |
| sp\|P07585\|PGS2\_HUMAN | P07585 | PGS2\_HUMAN | DCN | Decorin | 359 | 39.747 | 3.519115e+02 | 1.757337e+04 |  | 13432.287 | 4141.079 |  |  | 56.0 | 29.0 |  |
| sp\|Q13615\|MTMR3\_HUMAN | Q13615 | MTMR3\_HUMAN | MTMR3 | Myotubularin-related protein 3 | 1198 | 133.619 | 1.008396e+02 | 1.692842e+04 |  | 16928.418 |  |  |  | 35.6 |  |  |
| sp\|Q9NZB2\|F120A\_HUMAN | Q9NZB2 | F120A\_HUMAN | FAM120A | Constitutive coactivator of PPAR-gamma-like protein 1 | 1118 | 121.888 | 9.547762e+01 | 1.462109e+04 |  | 9396.723 | 5224.365 |  |  | 27.9 | 18.2 |  |
| sp\|Q9UI15\|TAGL3\_HUMAN | Q9UI15 | TAGL3\_HUMAN | TAGLN3 | Transgelin-3 | 199 | 22.473 | 2.167193e+05 | 6.118934e+06 |  | 6118934.5 |  |  |  | 64.8 |  |  |
| sp\|Q9P253\|VPS18\_HUMAN | Q9P253 | VPS18\_HUMAN | VPS18 | Vacuolar protein sorting-associated protein 18 homolog | 973 | 110.186 | 1.732397e+02 | 2.398231e+04 |  | 23982.305 |  |  |  | 29.2 |  |  |
| sp\|O95831\|AIFM1\_HUMAN | O95831 | AIFM1\_HUMAN | AIFM1 | Apoptosis-inducing factor 1, mitochondrial | 613 | 66.901 | 3.235077e+02 | 2.719158e+04 |  | 27191.578 |  |  |  | 44.0 |  |  |
| sp\|P09132\|SRP19\_HUMAN | P09132 | SRP19\_HUMAN | SRP19 | Signal recognition particle 19 kDa protein | 144 | 16.156 | 1.660088e+03 | 3.369629e+04 |  | 33696.285 |  |  |  | 55.6 |  |  |
| sp\|Q13555\|KCC2G\_HUMAN | Q13555 | KCC2G\_HUMAN | CAMK2G | Calcium/calmodulin-dependent protein kinase type II subunit gamma | 558 | 62.607 | 6.014765e+02 | 4.731062e+04 |  | 47310.625 |  |  |  | 37.8 |  |  |
| sp\|Q8TE68\|ES8L1\_HUMAN | Q8TE68 | ES8L1\_HUMAN | EPS8L1 | Epidermal growth factor receptor kinase substrate 8-like protein 1 | 723 | 80.251 | 8.463726e+02 | 8.533540e+04 |  | 27820.984 | 57514.414 |  |  | 38.3 | 21.2 |  |
| sp\|Q8WVV9\|HNRLL\_HUMAN | Q8WVV9 | HNRLL\_HUMAN | HNRNPLL | Heterogeneous nuclear ribonucleoprotein L-like | 542 | 60.083 | 2.059657e+02 | 1.554761e+04 |  | 15547.613 |  |  |  | 36.7 |  |  |
| sp\|P47224\|MSS4\_HUMAN | P47224 | MSS4\_HUMAN | RABIF | Guanine nucleotide exchange factor MSS4 | 123 | 13.839 | 1.444598e+04 | 2.511707e+05 |  | 171100.58 | 80070.14 |  |  | 56.9 | 44.7 |  |
| sp\|Q96S06\|LMF1\_HUMAN | Q96S06 | LMF1\_HUMAN | LMF1 | Lipase maturation factor 1 | 567 | 64.873 | 1.953384e+02 | 1.592094e+04 |  | 15920.942 |  |  |  | 25.9 |  |  |
| sp\|Q8N1G2\|CMTR1\_HUMAN | Q8N1G2 | CMTR1\_HUMAN | CMTR1 | Cap-specific mRNA (nucleoside-2'-O-)-methyltransferase 1 | 835 | 95.321 | 1.692919e+02 | 2.027412e+04 |  | 20274.12 |  |  |  | 36.2 |  |  |
| sp\|Q9BYI3\|HYCCI\_HUMAN | Q9BYI3 | HYCCI\_HUMAN | FAM126A | Hyccin | 521 | 57.625 | 5.469454e+02 | 3.959789e+04 |  | 39597.895 |  |  |  | 33.6 |  |  |
| sp\|A8MPS7\|YDJC\_HUMAN | A8MPS7 | YDJC\_HUMAN | YDJC | Carbohydrate deacetylase | 323 | 34.466 | 1.615134e+03 | 6.993855e+04 |  | 69938.55 |  |  |  | 47.1 |  |  |
| sp\|P48047\|ATPO\_HUMAN | P48047 | ATPO\_HUMAN | ATP5PO | ATP synthase subunit O, mitochondrial | 213 | 23.277 | 2.647869e+03 | 7.743561e+04 |  | 10093.582 | 67342.03 |  |  | 60.1 | 30.5 |  |
| sp\|Q96GZ6\|S41A3\_HUMAN | Q96GZ6 | S41A3\_HUMAN | SLC41A3 | Solute carrier family 41 member 3 | 507 | 54.767 | 3.079608e+02 | 2.119003e+04 |  | 21190.031 |  |  |  | 20.1 |  |  |
| sp\|P57721\|PCBP3\_HUMAN | P57721 | PCBP3\_HUMAN | PCBP3 | Poly(rC)-binding protein 3 | 371 | 39.465 | 9.446106e+03 | 4.683625e+05 |  | 468362.47 |  |  |  | 44.5 |  |  |
| sp\|Q13480\|GAB1\_HUMAN | Q13480 | GAB1\_HUMAN | GAB1 | GRB2-associated-binding protein 1 | 694 | 76.616 | 1.818323e+03 | 1.750281e+05 |  | 21623.941 |  | 153404.12 |  | 26.9 |  | 12.8 |
| sp\|O60645\|EXOC3\_HUMAN | O60645 | EXOC3\_HUMAN | EXOC3 | Exocyst complex component 3 | 745 | 85.567 | 1.410741e+02 | 1.516599e+04 |  | 15165.989 |  |  |  | 27.2 |  |  |
| sp\|Q8IVT5\|KSR1\_HUMAN | Q8IVT5 | KSR1\_HUMAN | KSR1 | Kinase suppressor of Ras 1 | 923 | 102.160 | 2.663324e+02 | 3.418393e+04 |  | 34183.93 |  |  |  | 40.8 |  |  |
| sp\|Q9Y4U1\|MMAC\_HUMAN | Q9Y4U1 | MMAC\_HUMAN | MMACHC | Cyanocobalamin reductase / alkylcobalamin dealkylase | 282 | 31.728 | 1.081335e+03 | 4.310424e+04 |  | 43104.24 |  |  |  | 29.1 |  |  |
| sp\|Q08752\|PPID\_HUMAN | Q08752 | PPID\_HUMAN | PPID | Peptidyl-prolyl cis-trans isomerase D | 370 | 40.764 | 1.249249e+03 | 6.397983e+04 |  | 63979.83 |  |  |  | 56.5 |  |  |
| sp\|Q92804\|RBP56\_HUMAN | Q92804 | RBP56\_HUMAN | TAF15 | TATA-binding protein-associated factor 2N | 592 | 61.830 | 2.492112e+03 | 1.935904e+05 |  | 110119.71 | 83470.734 |  |  | 35.1 | 22.0 |  |
| sp\|P55289\|CAD12\_HUMAN | P55289 | CAD12\_HUMAN | CDH12 | Cadherin-12 | 794 | 88.332 | 7.171036e+04 | 7.958242e+06 |  | 7958241.5 |  |  |  | 23.4 |  |  |
| sp\|Q99805\|TM9S2\_HUMAN | Q99805 | TM9S2\_HUMAN | TM9SF2 | Transmembrane 9 superfamily member 2 | 663 | 75.776 | 1.411815e+04 | 1.344085e+06 |  | 410501.03 |  | 933584.0 |  | 28.4 |  | 10.3 |
| sp\|P08319\|ADH4\_HUMAN | P08319 | ADH4\_HUMAN | ADH4 | All-trans-retinol dehydrogenase [NAD(+)] ADH4 | 380 | 40.222 | 5.646217e+03 | 2.853240e+05 |  | 285324.03 |  |  |  | 64.5 |  |  |
| sp\|P04004\|VTNC\_HUMAN | P04004 | VTNC\_HUMAN | VTN | Vitronectin | 478 | 54.306 | 1.292685e+04 | 8.819777e+05 |  | 45063.27 | 762551.06 | 74363.336 |  | 31.6 | 16.3 | 11.3 |
| sp\|O75113\|N4BP1\_HUMAN | O75113 | N4BP1\_HUMAN | N4BP1 | NEDD4-binding protein 1 | 896 | 100.379 | 1.279576e+02 | 1.613712e+04 |  | 16137.121 |  |  |  | 29.0 |  |  |
| sp\|Q8N5D0\|WDTC1\_HUMAN | Q8N5D0 | WDTC1\_HUMAN | WDTC1 | WD and tetratricopeptide repeats protein 1 | 677 | 75.920 | 1.100775e+02 | 1.049958e+04 |  | 10499.576 |  |  |  | 43.6 |  |  |
| sp\|Q9BYV8\|CEP41\_HUMAN | Q9BYV8 | CEP41\_HUMAN | CEP41 | Centrosomal protein of 41 kDa | 373 | 41.368 | 3.662258e+02 | 1.903402e+04 |  | 15453.599 | 3580.4258 |  |  | 47.7 | 36.5 |  |
| sp\|Q9Y3L5\|RAP2C\_HUMAN | Q9Y3L5 | RAP2C\_HUMAN | RAP2C | Ras-related protein Rap-2c | 183 | 20.745 | 5.707258e+04 | 1.487504e+06 |  | 1487503.9 |  |  |  | 66.7 |  |  |
| sp\|Q99952\|PTN18\_HUMAN | Q99952 | PTN18\_HUMAN | PTPN18 | Tyrosine-protein phosphatase non-receptor type 18 | 460 | 50.482 | 3.208447e+02 | 2.034926e+04 |  | 20349.256 |  |  |  | 40.0 |  |  |
| sp\|Q9NSY2\|STAR5\_HUMAN | Q9NSY2 | STAR5\_HUMAN | STARD5 | StAR-related lipid transfer protein 5 | 213 | 23.794 | 4.053089e+02 | 1.211632e+04 |  | 12116.319 |  |  |  | 32.9 |  |  |
| sp\|O75808\|CAN15\_HUMAN | O75808 | CAN15\_HUMAN | CAPN15 | Calpain-15 | 1086 | 117.314 | 8.154851e+01 | 1.201941e+04 |  | 12019.406 |  |  |  | 36.9 |  |  |
| sp\|P02686\|MBP\_HUMAN | P02686 | MBP\_HUMAN | MBP | Myelin basic protein | 304 | 33.117 | 2.695172e+02 | 1.121385e+04 |  | 11213.847 |  |  |  | 28.3 |  |  |
| sp\|O14545\|TRAD1\_HUMAN | O14545 | TRAD1\_HUMAN | TRAFD1 | TRAF-type zinc finger domain-containing protein 1 | 582 | 64.841 | 2.049941e+02 | 1.669968e+04 |  | 16699.682 |  |  |  | 30.1 |  |  |
| sp\|O15211\|RGL2\_HUMAN | O15211 | RGL2\_HUMAN | RGL2 | Ral guanine nucleotide dissociation stimulator-like 2 | 777 | 83.549 | 0.000000e+00 | 0.000000e+00 |  | 0.0 |  |  |  | 35.1 |  |  |
| sp\|P04179\|SODM\_HUMAN | P04179 | SODM\_HUMAN | SOD2 | Superoxide dismutase [Mn], mitochondrial | 222 | 24.750 | 1.463528e+03 | 4.550858e+04 |  | 45508.58 |  |  |  | 14.9 |  |  |
| sp\|P24539\|AT5F1\_HUMAN | P24539 | AT5F1\_HUMAN | ATP5PB | ATP synthase F(0) complex subunit B1, mitochondrial | 256 | 28.909 | 2.035143e+03 | 7.391712e+04 |  | 24584.744 | 40261.293 | 9071.086 |  | 38.3 | 22.7 | 7.8 |
| sp\|Q01085\|TIAR\_HUMAN | Q01085 | TIAR\_HUMAN | TIAL1 | Nucleolysin TIAR | 375 | 41.591 | 2.793548e+03 | 1.459731e+05 |  | 118658.73 | 27314.33 |  |  | 24.8 | 13.1 |  |
| sp\|Q15654\|TRIP6\_HUMAN | Q15654 | TRIP6\_HUMAN | TRIP6 | Thyroid receptor-interacting protein 6 | 476 | 50.288 | 9.901887e+02 | 6.256039e+04 |  | 62560.395 |  |  |  | 58.0 |  |  |
| sp\|Q9NVJ2\|ARL8B\_HUMAN | Q9NVJ2 | ARL8B\_HUMAN | ARL8B | ADP-ribosylation factor-like protein 8B | 186 | 21.539 | 1.233530e+04 | 3.338047e+05 |  | 173583.38 |  | 160221.31 |  | 50.0 |  | 9.7 |
| sp\|Q9Y426\|C2CD2\_HUMAN | Q9Y426 | C2CD2\_HUMAN | C2CD2 | C2 domain-containing protein 2 | 696 | 75.533 | 1.736179e+03 | 1.647587e+05 |  | 13266.716 | 70987.49 | 80504.54 |  | 17.1 | 9.8 | 11.2 |
| sp\|Q13232\|NDK3\_HUMAN | Q13232 | NDK3\_HUMAN | NME3 | Nucleoside diphosphate kinase 3 | 169 | 19.015 | 9.362397e+02 | 2.236663e+04 |  | 22366.63 |  |  |  | 30.2 |  |  |
| sp\|Q9UPQ9\|TNR6B\_HUMAN | Q9UPQ9 | TNR6B\_HUMAN | TNRC6B | Trinucleotide repeat-containing gene 6B protein | 1833 | 194.002 | 2.678590e+01 | 6.528744e+03 |  | 6528.744 |  |  |  | 24.1 |  |  |
| sp\|Q9H777\|RNZ1\_HUMAN | Q9H777 | RNZ1\_HUMAN | ELAC1 | Zinc phosphodiesterase ELAC protein 1 | 363 | 40.019 | 2.379082e+02 | 1.196170e+04 |  | 11961.696 |  |  |  | 27.5 |  |  |
| sp\|Q8WV44\|TRI41\_HUMAN | Q8WV44 | TRI41\_HUMAN | TRIM41 | E3 ubiquitin-protein ligase TRIM41 | 630 | 71.670 | 1.409056e+02 | 1.268770e+04 |  | 12687.7 |  |  |  | 26.8 |  |  |
| sp\|Q5VTR2\|BRE1A\_HUMAN | Q5VTR2 | BRE1A\_HUMAN | RNF20 | E3 ubiquitin-protein ligase BRE1A | 975 | 113.662 | 5.986470e+02 | 8.548762e+04 |  | 28043.352 |  | 57444.27 |  | 26.8 |  | 13.0 |
| sp\|O14556\|G3PT\_HUMAN | O14556 | G3PT\_HUMAN | GAPDHS | Glyceraldehyde-3-phosphate dehydrogenase, testis-specific | 408 | 44.501 | 7.923212e+04 | 4.429842e+06 |  | 4429841.5 |  |  |  | 25.7 |  |  |
| sp\|Q12766\|HMGX3\_HUMAN | Q12766 | HMGX3\_HUMAN | HMGXB3 | HMG domain-containing protein 3 | 1292 | 141.106 | 2.071941e+02 | 3.673161e+04 |  | 36731.61 |  |  |  | 27.6 |  |  |
| sp\|Q96BZ8\|LENG1\_HUMAN | Q96BZ8 | LENG1\_HUMAN | LENG1 | Leukocyte receptor cluster member 1 | 264 | 30.528 | 3.741770e+02 | 1.435134e+04 |  | 14351.345 |  |  |  | 40.5 |  |  |
| sp\|Q9Y6M1\|IF2B2\_HUMAN | Q9Y6M1 | IF2B2\_HUMAN | IGF2BP2 | Insulin-like growth factor 2 mRNA-binding protein 2 | 599 | 66.121 | 3.210000e+02 | 2.666623e+04 |  | 26666.232 |  |  |  | 21.0 |  |  |
| sp\|Q96PU4\|UHRF2\_HUMAN | Q96PU4 | UHRF2\_HUMAN | UHRF2 | E3 ubiquitin-protein ligase UHRF2 | 802 | 89.985 | 7.161671e+03 | 8.096581e+05 |  | 809658.1 |  |  |  | 33.5 |  |  |
| sp\|A8MXQ7\|IQAK1\_HUMAN | A8MXQ7 | IQAK1\_HUMAN | IQANK1 | IQ motif and ankyrin repeat domain-containing protein 1 | 560 | 63.562 | 1.561249e+02 | 1.246772e+04 |  | 12467.715 |  |  |  | 40.4 |  |  |
| sp\|Q96AG4\|LRC59\_HUMAN | Q96AG4 | LRC59\_HUMAN | LRRC59 | Leucine-rich repeat-containing protein 59 | 307 | 34.930 | 1.034427e+04 | 4.539579e+05 |  | 23887.201 | 261108.28 | 168962.42 |  | 36.5 | 17.6 | 6.5 |
| sp\|Q9BW19\|KIFC1\_HUMAN | Q9BW19 | KIFC1\_HUMAN | KIFC1 | Kinesin-like protein KIFC1 | 673 | 73.748 | 9.495312e+01 | 8.797852e+03 |  | 8797.852 |  |  |  | 38.9 |  |  |
| sp\|Q6XPR3\|RPTN\_HUMAN | Q6XPR3 | RPTN\_HUMAN | RPTN | Repetin | 784 | 90.731 | 1.158356e+02 | 1.320428e+04 |  | 8468.824 | 4735.461 |  |  | 42.0 | 38.4 |  |
| sp\|Q9Y3Z3\|SAMH1\_HUMAN | Q9Y3Z3 | SAMH1\_HUMAN | SAMHD1 | Deoxynucleoside triphosphate triphosphohydrolase SAMHD1 | 626 | 72.201 | 6.401000e+02 | 5.806417e+04 |  | 58064.17 |  |  |  | 40.9 |  |  |
| sp\|Q86YN1\|DOPP1\_HUMAN | Q86YN1 | DOPP1\_HUMAN | DOLPP1 | Dolichyldiphosphatase 1 | 238 | 27.031 | 3.550792e+03 | 1.205881e+05 |  | 120588.1 |  |  |  | 5.5 |  |  |
| sp\|Q8TAQ2\|SMRC2\_HUMAN | Q8TAQ2 | SMRC2\_HUMAN | SMARCC2 | SWI/SNF complex subunit SMARCC2 | 1214 | 132.879 | 1.264364e+02 | 2.110793e+04 |  | 21107.934 |  |  |  | 20.4 |  |  |
| sp\|O43240\|KLK10\_HUMAN | O43240 | KLK10\_HUMAN | KLK10 | Kallikrein-10 | 276 | 30.170 | 2.207437e+02 | 8.367213e+03 |  | 8367.213 |  |  |  | 43.8 |  |  |
| sp\|P78406\|RAE1L\_HUMAN | P78406 | RAE1L\_HUMAN | RAE1 | mRNA export factor RAE1 | 368 | 40.968 | 9.902100e+02 | 5.096702e+04 |  | 50967.02 |  |  |  | 23.1 |  |  |
| sp\|Q09028\|RBBP4\_HUMAN | Q09028 | RBBP4\_HUMAN | RBBP4 | Histone-binding protein RBBP4 | 425 | 47.656 | 2.449144e+03 | 1.466388e+05 |  | 146638.84 |  |  |  | 17.9 |  |  |
| sp\|Q15126\|PMVK\_HUMAN | Q15126 | PMVK\_HUMAN | PMVK | Phosphomevalonate kinase | 192 | 21.995 | 3.476445e+03 | 9.606749e+04 |  | 37837.473 | 38512.44 | 19717.574 |  | 52.1 | 47.4 | 19.8 |
| sp\|Q96DT6\|ATG4C\_HUMAN | Q96DT6 | ATG4C\_HUMAN | ATG4C | Cysteine protease ATG4C | 458 | 52.497 | 8.823130e+02 | 5.819348e+04 |  | 58193.477 |  |  |  | 26.4 |  |  |
| sp\|Q9UK55\|ZPI\_HUMAN | Q9UK55 | ZPI\_HUMAN | SERPINA10 | Protein Z-dependent protease inhibitor | 444 | 50.707 | 5.696193e+02 | 3.628856e+04 |  | 36288.56 |  |  |  | 34.9 |  |  |
| sp\|Q9Y337\|KLK5\_HUMAN | Q9Y337 | KLK5\_HUMAN | KLK5 | Kallikrein-5 | 293 | 32.020 | 9.573737e+02 | 3.851412e+04 |  | 38514.12 |  |  |  | 23.2 |  |  |
| sp\|O60669\|MOT2\_HUMAN | O60669 | MOT2\_HUMAN | SLC16A7 | Monocarboxylate transporter 2 | 478 | 52.200 | 1.537359e+04 | 1.008238e+06 |  | 45270.965 | 265755.6 | 697211.5 |  | 21.3 | 7.3 | 5.6 |
| sp\|O75063\|XYLK\_HUMAN | O75063 | XYLK\_HUMAN | FAM20B | Glycosaminoglycan xylosylkinase | 409 | 46.432 | 1.026887e+04 | 5.990417e+05 |  | 38177.797 | 509632.78 | 51231.094 |  | 25.7 | 24.7 | 7.1 |
| sp\|P17540\|KCRS\_HUMAN | P17540 | KCRS\_HUMAN | CKMT2 | Creatine kinase S-type, mitochondrial | 419 | 47.504 | 1.468307e+03 | 8.763229e+04 |  | 32351.568 | 39142.492 | 16138.234 |  | 25.1 | 31.3 | 16.0 |
| sp\|P60468\|SC61B\_HUMAN | P60468 | SC61B\_HUMAN | SEC61B | Protein transport protein Sec61 subunit beta | 96 | 9.974 | 3.904382e+04 | 4.892590e+05 |  | 63263.37 | 171181.88 | 254813.75 |  | 78.1 | 55.2 | 42.7 |
| sp\|Q92879\|CELF1\_HUMAN | Q92879 | CELF1\_HUMAN | CELF1 | CUGBP Elav-like family member 1 | 486 | 52.063 | 5.230018e+02 | 3.420972e+04 |  | 34209.723 |  |  |  | 25.1 |  |  |
| sp\|Q9NX55\|HYPK\_HUMAN | Q9NX55 | HYPK\_HUMAN | HYPK | Huntingtin-interacting protein K | 121 | 13.651 | 1.653629e+03 | 2.836088e+04 |  | 28360.883 |  |  |  | 63.6 |  |  |
| sp\|O14815\|CAN9\_HUMAN | O14815 | CAN9\_HUMAN | CAPN9 | Calpain-9 | 690 | 79.097 | 0.000000e+00 | 0.000000e+00 |  | 0.0 |  |  |  | 23.2 |  |  |
| sp\|Q9H270\|VPS11\_HUMAN | Q9H270 | VPS11\_HUMAN | VPS11 | Vacuolar protein sorting-associated protein 11 homolog | 941 | 107.837 | 2.189187e+02 | 2.965978e+04 |  | 29659.775 |  |  |  | 29.6 |  |  |
| sp\|Q8N108\|MIER1\_HUMAN | Q8N108 | MIER1\_HUMAN | MIER1 | Mesoderm induction early response protein 1 | 512 | 57.983 | 2.568483e+02 | 1.871089e+04 |  | 18710.893 |  |  |  | 27.1 |  |  |
| sp\|O14791\|APOL1\_HUMAN | O14791 | APOL1\_HUMAN | APOL1 | Apolipoprotein L1 | 398 | 43.974 | 1.774028e+02 | 9.801074e+03 |  | 9801.074 |  |  |  | 25.6 |  |  |
| sp\|Q06033\|ITIH3\_HUMAN | Q06033 | ITIH3\_HUMAN | ITIH3 | Inter-alpha-trypsin inhibitor heavy chain H3 | 890 | 99.849 | 1.700952e+02 | 2.133796e+04 |  | 21337.957 |  |  |  | 27.6 |  |  |
| sp\|Q9BRR8\|GPTC1\_HUMAN | Q9BRR8 | GPTC1\_HUMAN | GPATCH1 | G patch domain-containing protein 1 | 931 | 103.345 | 7.418845e+01 | 9.632587e+03 |  | 9632.587 |  |  |  | 21.9 |  |  |
| sp\|P35030\|TRY3\_HUMAN | P35030 | TRY3\_HUMAN | PRSS3 | Trypsin-3 | 304 | 32.529 | 5.126652e+03 | 2.095182e+05 |  | 107548.31 | 55696.406 | 46273.48 |  | 38.8 | 22.0 | 9.2 |
| sp\|Q86WR0\|CCD25\_HUMAN | Q86WR0 | CCD25\_HUMAN | CCDC25 | Coiled-coil domain-containing protein 25 | 208 | 24.479 | 2.250887e+03 | 6.922524e+04 |  | 69225.24 |  |  |  | 44.7 |  |  |
| sp\|Q9Y312\|AAR2\_HUMAN | Q9Y312 | AAR2\_HUMAN | AAR2 | Protein AAR2 homolog | 384 | 43.472 | 4.594058e+02 | 2.509130e+04 |  | 25091.303 |  |  |  | 34.6 |  |  |
| sp\|Q92600\|CNOT9\_HUMAN | Q92600 | CNOT9\_HUMAN | CNOT9 | CCR4-NOT transcription complex subunit 9 | 299 | 33.631 | 6.597011e+02 | 2.787431e+04 |  | 27874.31 |  |  |  | 25.4 |  |  |
| sp\|Q1KMD3\|HNRL2\_HUMAN | Q1KMD3 | HNRL2\_HUMAN | HNRNPUL2 | Heterogeneous nuclear ribonucleoprotein U-like protein 2 | 747 | 85.105 | 2.648097e+02 | 2.831432e+04 |  | 28314.318 |  |  |  | 23.8 |  |  |
| sp\|Q6IA86\|ELP2\_HUMAN | Q6IA86 | ELP2\_HUMAN | ELP2 | Elongator complex protein 2 | 826 | 92.500 | 3.060958e+02 | 3.557266e+04 |  | 35572.656 |  |  |  | 21.2 |  |  |
| sp\|Q8NHS0\|DNJB8\_HUMAN | Q8NHS0 | DNJB8\_HUMAN | DNAJB8 | DnaJ homolog subfamily B member 8 | 232 | 25.686 | 2.795908e+03 | 9.022699e+04 |  | 90226.99 |  |  |  | 24.6 |  |  |
| sp\|P05186\|PPBT\_HUMAN | P05186 | PPBT\_HUMAN | ALPL | Alkaline phosphatase, tissue-nonspecific isozyme | 524 | 57.305 | 7.209175e+03 | 5.190333e+05 |  | 448150.62 | 29863.047 | 41019.62 |  | 26.9 | 21.9 | 14.7 |
| sp\|Q6IBW4\|CNDH2\_HUMAN | Q6IBW4 | CNDH2\_HUMAN | NCAPH2 | Condensin-2 complex subunit H2 | 605 | 68.227 | 3.480204e+02 | 2.983171e+04 |  | 29831.707 |  |  |  | 25.1 |  |  |
| sp\|P42331\|RHG25\_HUMAN | P42331 | RHG25\_HUMAN | ARHGAP25 | Rho GTPase-activating protein 25 | 645 | 73.435 | 3.704563e+02 | 3.417884e+04 |  | 34178.836 |  |  |  | 36.6 |  |  |
| sp\|O00124\|UBXN8\_HUMAN | O00124 | UBXN8\_HUMAN | UBXN8 | UBX domain-containing protein 8 | 270 | 30.541 | 4.211520e+02 | 1.615992e+04 |  | 16159.921 |  |  |  | 34.8 |  |  |
| sp\|O15151\|MDM4\_HUMAN | O15151 | MDM4\_HUMAN | MDM4 | Protein Mdm4 | 490 | 54.864 | 1.859522e+02 | 1.281758e+04 |  | 12817.575 |  |  |  | 18.6 |  |  |
| sp\|P04746\|AMYP\_HUMAN | P04746 | AMYP\_HUMAN | AMY2A | Pancreatic alpha-amylase | 511 | 57.707 | 2.269294e+03 | 1.645267e+05 |  | 164526.72 |  |  |  | 9.6 |  |  |
| sp\|P05413\|FABPH\_HUMAN | P05413 | FABPH\_HUMAN | FABP3 | Fatty acid-binding protein, heart | 133 | 14.858 | 6.907239e+02 | 1.289383e+04 |  | 12893.831 |  |  |  | 39.1 |  |  |
| sp\|P06729\|CD2\_HUMAN | P06729 | CD2\_HUMAN | CD2 | T-cell surface antigen CD2 | 351 | 39.448 | 6.768782e+02 | 3.354692e+04 |  | 33546.92 |  |  |  | 33.3 |  |  |
| sp\|P14780\|MMP9\_HUMAN | P14780 | MMP9\_HUMAN | MMP9 | Matrix metalloproteinase-9 | 707 | 78.458 | 9.411297e+02 | 9.276921e+04 |  | 61706.207 | 25513.541 | 5549.459 |  | 35.9 | 15.4 | 6.9 |
| sp\|P30519\|HMOX2\_HUMAN | P30519 | HMOX2\_HUMAN | HMOX2 | Heme oxygenase 2 | 316 | 36.033 | 8.503421e+02 | 3.849562e+04 |  | 20831.764 | 17663.852 |  |  | 38.9 | 19.3 |  |
| sp\|Q02878\|RL6\_HUMAN | Q02878 | RL6\_HUMAN | RPL6 | 60S ribosomal protein L6 | 288 | 32.728 | 2.511642e+03 | 1.032748e+05 |  | 14443.54 | 65251.27 | 23579.969 |  | 34.4 | 25.0 | 23.6 |
| sp\|Q04941\|PLP2\_HUMAN | Q04941 | PLP2\_HUMAN | PLP2 | Proteolipid protein 2 | 152 | 16.691 | 6.589150e+05 | 1.381748e+07 |  | 1025806.2 | 1258191.5 | 11533484.0 |  | 28.9 | 24.3 | 24.3 |
| sp\|Q13217\|DNJC3\_HUMAN | Q13217 | DNJC3\_HUMAN | DNAJC3 | DnaJ homolog subfamily C member 3 | 504 | 57.580 | 6.426049e+03 | 4.648714e+05 |  | 36363.113 | 428508.28 |  |  | 22.8 | 26.2 |  |
| sp\|Q14657\|LAGE3\_HUMAN | Q14657 | LAGE3\_HUMAN | LAGE3 | EKC/KEOPS complex subunit LAGE3 | 143 | 14.804 | 8.826685e+02 | 1.641700e+04 |  | 16417.002 |  |  |  | 45.5 |  |  |
| sp\|Q8WUF8\|F172A\_HUMAN | Q8WUF8 | F172A\_HUMAN | FAM172A | Cotranscriptional regulator FAM172A | 416 | 47.972 | 4.905618e+02 | 2.956642e+04 |  | 29566.416 |  |  |  | 26.7 |  |  |
| sp\|Q9BTD8\|RBM42\_HUMAN | Q9BTD8 | RBM42\_HUMAN | RBM42 | RNA-binding protein 42 | 480 | 50.414 | 4.055538e+02 | 2.568720e+04 |  | 25687.201 |  |  |  | 22.5 |  |  |
| sp\|Q9BVG9\|PTSS2\_HUMAN | Q9BVG9 | PTSS2\_HUMAN | PTDSS2 | Phosphatidylserine synthase 2 | 487 | 56.253 | 1.935949e+02 | 1.368223e+04 |  | 13682.226 |  |  |  | 25.7 |  |  |
| sp\|Q9H079\|KTBL1\_HUMAN | Q9H079 | KTBL1\_HUMAN | KATNBL1 | KATNB1-like protein 1 | 304 | 34.767 | 0.000000e+00 | 0.000000e+00 |  | 0.0 |  |  |  | 39.5 |  |  |
| sp\|Q9H496\|IFG15\_HUMAN | Q9H496 | IFG15\_HUMAN | TOR1AIP2 | Torsin-1A-interacting protein 2, isoform IFRG15 | 131 | 15.348 | 7.910528e+03 | 1.525367e+05 |  | 152536.72 |  |  |  | 20.6 |  |  |
| sp\|Q9NWW5\|CLN6\_HUMAN | Q9NWW5 | CLN6\_HUMAN | CLN6 | Ceroid-lipofuscinosis neuronal protein 6 | 311 | 35.919 | 2.126429e+03 | 9.596045e+04 |  | 25119.654 |  | 70840.8 |  | 7.1 |  | 3.5 |
| sp\|P05543\|THBG\_HUMAN | P05543 | THBG\_HUMAN | SERPINA7 | Thyroxine-binding globulin | 415 | 46.325 | 2.371577e+03 | 1.380288e+05 |  | 130840.0 | 7188.81 |  |  | 14.7 | 17.8 |  |
| sp\|Q9Y282\|ERGI3\_HUMAN | Q9Y282 | ERGI3\_HUMAN | ERGIC3 | Endoplasmic reticulum-Golgi intermediate compartment protein 3 | 383 | 43.222 | 3.092153e+04 | 1.679125e+06 |  | 80068.516 | 827268.94 | 771787.2 |  | 25.8 | 25.1 | 20.6 |
| sp\|O00391\|QSOX1\_HUMAN | O00391 | QSOX1\_HUMAN | QSOX1 | Sulfhydryl oxidase 1 | 747 | 82.578 | 4.572070e+02 | 4.743451e+04 |  | 17838.111 | 29596.395 |  |  | 38.3 | 17.4 |  |
| sp\|P80723\|BASP1\_HUMAN | P80723 | BASP1\_HUMAN | BASP1 | Brain acid soluble protein 1 | 227 | 22.693 | 1.392552e+03 | 3.970274e+04 |  | 25787.465 | 13915.278 |  |  | 80.6 | 47.6 |  |
| sp\|Q8TBA6\|GOGA5\_HUMAN | Q8TBA6 | GOGA5\_HUMAN | GOLGA5 | Golgin subfamily A member 5 | 731 | 83.024 | 7.563030e+02 | 7.888903e+04 |  | 26609.01 |  | 52280.02 |  | 25.4 |  | 13.5 |
| sp\|Q15417\|CNN3\_HUMAN | Q15417 | CNN3\_HUMAN | CNN3 | Calponin-3 | 329 | 36.414 | 0.000000e+00 | 0.000000e+00 |  | 0.0 |  |  |  | 37.7 |  |  |
| sp\|Q9NVK5\|FGOP2\_HUMAN | Q9NVK5 | FGOP2\_HUMAN | FGFR1OP2 | FGFR1 oncogene partner 2 | 253 | 29.426 | 1.340814e+03 | 4.956978e+04 |  | 49569.777 |  |  |  | 35.2 |  |  |
| sp\|P55210\|CASP7\_HUMAN | P55210 | CASP7\_HUMAN | CASP7 | Caspase-7 | 303 | 34.277 | 1.188027e+02 | 5.116187e+03 |  | 5116.187 |  |  |  | 33.0 |  |  |
| sp\|Q9UKG9\|OCTC\_HUMAN | Q9UKG9 | OCTC\_HUMAN | CROT | Peroxisomal carnitine O-octanoyltransferase | 612 | 70.178 | 1.620758e+02 | 1.429014e+04 |  | 14290.137 |  |  |  | 46.9 |  |  |
| sp\|P13804\|ETFA\_HUMAN | P13804 | ETFA\_HUMAN | ETFA | Electron transfer flavoprotein subunit alpha, mitochondrial | 333 | 35.080 | 3.263600e+03 | 1.438380e+05 |  | 27503.959 | 72097.16 | 44236.883 |  | 51.1 | 33.9 | 16.2 |
| sp\|Q32P28\|P3H1\_HUMAN | Q32P28 | P3H1\_HUMAN | P3H1 | Prolyl 3-hydroxylase 1 | 736 | 83.394 | 4.262239e+03 | 4.465702e+05 |  | 27312.984 | 353723.72 | 65533.53 |  | 25.4 | 21.2 | 16.7 |
| sp\|Q9UEU0\|VTI1B\_HUMAN | Q9UEU0 | VTI1B\_HUMAN | VTI1B | Vesicle transport through interaction with t-SNAREs homolog 1B | 232 | 26.688 | 4.275565e+04 | 1.433596e+06 |  | 48527.676 | 800782.06 | 584285.9 |  | 31.5 | 44.8 | 31.0 |
| sp\|Q16621\|NFE2\_HUMAN | Q16621 | NFE2\_HUMAN | NFE2 | Transcription factor NF-E2 45 kDa subunit | 373 | 41.473 | 1.255862e+03 | 6.543716e+04 |  | 65437.16 |  |  |  | 15.0 |  |  |
| sp\|O14880\|MGST3\_HUMAN | O14880 | MGST3\_HUMAN | MGST3 | Microsomal glutathione S-transferase 3 | 152 | 16.516 | 1.960240e+03 | 4.067535e+04 |  | 40675.35 |  |  |  | 34.9 |  |  |
| sp\|O60238\|BNI3L\_HUMAN | O60238 | BNI3L\_HUMAN | BNIP3L | BCL2/adenovirus E1B 19 kDa protein-interacting protein 3-like | 219 | 23.930 | 5.156876e+03 | 1.550410e+05 |  | 44022.105 | 111018.875 |  |  | 37.4 | 31.5 |  |
| sp\|P15151\|PVR\_HUMAN | P15151 | PVR\_HUMAN | PVR | Poliovirus receptor | 417 | 45.303 | 2.419543e+03 | 1.377138e+05 |  | 0.0 | 28963.098 | 108750.7 |  | 17.3 | 8.4 | 16.3 |
| sp\|P20290\|BTF3\_HUMAN | P20290 | BTF3\_HUMAN | BTF3 | Transcription factor BTF3 | 206 | 22.168 | 9.180823e+02 | 2.556968e+04 |  | 25569.68 |  |  |  | 37.9 |  |  |
| sp\|P30049\|ATPD\_HUMAN | P30049 | ATPD\_HUMAN | ATP5F1D | ATP synthase subunit delta, mitochondrial | 168 | 17.490 | 2.389012e+03 | 5.249588e+04 |  | 37783.832 | 14712.05 |  |  | 12.5 | 13.1 |  |
| sp\|P32969\|RL9\_HUMAN | P32969 | RL9\_HUMAN | RPL9 | 60S ribosomal protein L9 | 192 | 21.863 | 1.713558e+03 | 4.706799e+04 |  | 47067.99 |  |  |  | 39.6 |  |  |
| sp\|P49459\|UBE2A\_HUMAN | P49459 | UBE2A\_HUMAN | UBE2A | Ubiquitin-conjugating enzyme E2 A | 152 | 17.315 | 2.091057e+03 | 4.548892e+04 |  | 45488.918 |  |  |  | 28.9 |  |  |
| sp\|P56377\|AP1S2\_HUMAN | P56377 | AP1S2\_HUMAN | AP1S2 | AP-1 complex subunit sigma-2 | 157 | 18.615 | 5.097878e+03 | 1.192256e+05 |  | 119225.62 |  |  |  | 31.2 |  |  |
| sp\|P62244\|RS15A\_HUMAN | P62244 | RS15A\_HUMAN | RPS15A | 40S ribosomal protein S15a | 130 | 14.840 | 1.363584e+03 | 2.542336e+04 |  | 25423.36 |  |  |  | 31.5 |  |  |
| sp\|P62917\|RL8\_HUMAN | P62917 | RL8\_HUMAN | RPL8 | 60S ribosomal protein L8 | 257 | 28.025 | 3.222519e+03 | 1.134640e+05 |  | 15469.812 | 97994.234 |  |  | 30.4 | 24.1 |  |
| sp\|P63098\|CANB1\_HUMAN | P63098 | CANB1\_HUMAN | PPP3R1 | Calcineurin subunit B type 1 | 170 | 19.300 | 1.338515e+03 | 3.245619e+04 |  | 32456.195 |  |  |  | 17.6 |  |  |
| sp\|P63146\|UBE2B\_HUMAN | P63146 | UBE2B\_HUMAN | UBE2B | Ubiquitin-conjugating enzyme E2 B | 152 | 17.312 | 3.306473e+03 | 7.191663e+04 |  | 71916.63 |  |  |  | 54.6 |  |  |
| sp\|Q8N5M4\|TTC9C\_HUMAN | Q8N5M4 | TTC9C\_HUMAN | TTC9C | Tetratricopeptide repeat protein 9C | 171 | 20.013 | 8.882599e+02 | 2.233415e+04 |  | 22334.15 |  |  |  | 22.2 |  |  |
| sp\|Q8NDC0\|MISSL\_HUMAN | Q8NDC0 | MISSL\_HUMAN | MAPK1IP1L | MAPK-interacting and spindle-stabilizing protein-like | 245 | 24.269 | 1.425898e+05 | 4.347679e+06 |  | 32501.904 | 4253335.0 | 61842.125 |  | 6.9 | 6.9 | 6.9 |
| sp\|Q8WY22\|BRI3B\_HUMAN | Q8WY22 | BRI3B\_HUMAN | BRI3BP | BRI3-binding protein | 251 | 27.836 | 1.433453e+03 | 5.013112e+04 |  | 17227.547 | 12345.464 | 20558.111 |  | 14.7 | 6.8 | 9.6 |
| sp\|Q9H6U8\|ALG9\_HUMAN | Q9H6U8 | ALG9\_HUMAN | ALG9 | Alpha-1,2-mannosyltransferase ALG9 | 611 | 69.863 | 1.397847e+03 | 1.226942e+05 |  | 10148.549 | 40309.055 | 72236.61 |  | 20.9 | 17.0 | 10.3 |
| sp\|Q9H867\|MT21D\_HUMAN | Q9H867 | MT21D\_HUMAN | VCPKMT | Protein N-lysine methyltransferase METTL21D | 229 | 25.807 | 1.552523e+03 | 5.033761e+04 |  | 50337.613 |  |  |  | 21.4 |  |  |
| sp\|Q9HCN8\|SDF2L\_HUMAN | Q9HCN8 | SDF2L\_HUMAN | SDF2L1 | Stromal cell-derived factor 2-like protein 1 | 221 | 23.598 | 9.639508e+03 | 2.857901e+05 |  | 65879.89 | 219910.22 |  |  | 33.5 | 26.2 |  |
| sp\|Q9UNI6\|DUS12\_HUMAN | Q9UNI6 | DUS12\_HUMAN | DUSP12 | Dual specificity protein phosphatase 12 | 340 | 37.687 | 1.495248e+03 | 7.079818e+04 |  | 70798.18 |  |  |  | 36.5 |  |  |
| sp\|Q9UNK0\|STX8\_HUMAN | Q9UNK0 | STX8\_HUMAN | STX8 | Syntaxin-8 | 236 | 26.907 | 1.391402e+04 | 4.703652e+05 |  | 24908.281 | 89096.195 | 356360.75 |  | 23.7 | 24.6 | 23.7 |
| sp\|Q9Y2Q5\|LTOR2\_HUMAN | Q9Y2Q5 | LTOR2\_HUMAN | LAMTOR2 | Ragulator complex protein LAMTOR2 | 125 | 13.508 | 2.550062e+04 | 4.327719e+05 |  | 96652.914 | 336118.94 |  |  | 56.8 | 39.2 |  |
| sp\|Q9Y547\|IFT25\_HUMAN | Q9Y547 | IFT25\_HUMAN | HSPB11 | Intraflagellar transport protein 25 homolog | 144 | 16.297 | 1.981904e+04 | 4.057957e+05 |  | 405795.66 |  |  |  | 8.3 |  |  |
| sp\|O75379\|VAMP4\_HUMAN | O75379 | VAMP4\_HUMAN | VAMP4 | Vesicle-associated membrane protein 4 | 141 | 16.397 | 2.915317e+03 | 6.005752e+04 |  | 0.0 | 44374.047 | 15683.472 |  | 27.7 | 9.9 | 31.2 |
| sp\|Q13907\|IDI1\_HUMAN | Q13907 | IDI1\_HUMAN | IDI1 | Isopentenyl-diphosphate Delta-isomerase 1 | 227 | 26.319 | 1.455038e+03 | 4.811282e+04 |  | 48112.824 |  |  |  | 18.5 |  |  |
| sp\|Q15025\|TNIP1\_HUMAN | Q15025 | TNIP1\_HUMAN | TNIP1 | TNFAIP3-interacting protein 1 | 636 | 71.864 | 1.013667e+02 | 9.152161e+03 |  | 9152.161 |  |  |  | 36.6 |  |  |
| sp\|Q9NQ34\|TMM9B\_HUMAN | Q9NQ34 | TMM9B\_HUMAN | TMEM9B | Transmembrane protein 9B | 198 | 22.531 | 4.206044e+02 | 1.190615e+04 |  | 11906.153 |  |  |  | 25.3 |  |  |
| sp\|P01602\|KV105\_HUMAN | P01602 | KV105\_HUMAN | IGKV1-5 | Immunoglobulin kappa variable 1-5 | 117 | 12.782 | 6.314271e+03 | 1.014003e+05 |  | 7413.7305 |  | 93986.56 |  | 36.8 |  | 25.6 |
| sp\|P51965\|UB2E1\_HUMAN | P51965 | UB2E1\_HUMAN | UBE2E1 | Ubiquitin-conjugating enzyme E2 E1 | 193 | 21.404 | 4.575463e+02 | 1.230402e+04 |  | 12304.023 |  |  |  | 24.9 |  |  |
| sp\|Q969M3\|YIPF5\_HUMAN | Q969M3 | YIPF5\_HUMAN | YIPF5 | Protein YIPF5 | 257 | 27.989 | 3.195962e+03 | 1.123844e+05 |  | 83735.67 |  | 28648.744 |  | 16.7 |  | 4.7 |
| sp\|Q13114\|TRAF3\_HUMAN | Q13114 | TRAF3\_HUMAN | TRAF3 | TNF receptor-associated factor 3 | 568 | 64.490 | 5.349577e+02 | 4.334400e+04 |  | 43343.996 |  |  |  | 25.2 |  |  |
| sp\|Q66PJ3\|AR6P4\_HUMAN | Q66PJ3 | AR6P4\_HUMAN | ARL6IP4 | ADP-ribosylation factor-like protein 6-interacting protein 4 | 237 | 26.375 | 1.044400e+03 | 3.460801e+04 |  | 34608.008 |  |  |  | 40.5 |  |  |
| sp\|O95295\|SNAPN\_HUMAN | O95295 | SNAPN\_HUMAN | SNAPIN | SNARE-associated protein Snapin | 136 | 14.874 | 2.869269e+03 | 5.361869e+04 |  | 53618.69 |  |  |  | 50.7 |  |  |
| sp\|P46782\|RS5\_HUMAN | P46782 | RS5\_HUMAN | RPS5 | 40S ribosomal protein S5 | 204 | 22.876 | 2.276251e+03 | 6.542104e+04 |  | 13548.379 |  | 51872.66 |  | 21.1 |  | 17.2 |
| sp\|Q96NT1\|NP1L5\_HUMAN | Q96NT1 | NP1L5\_HUMAN | NAP1L5 | Nucleosome assembly protein 1-like 5 | 182 | 19.593 | 1.795720e+03 | 4.420350e+04 |  | 44203.5 |  |  |  | 18.1 |  |  |
| sp\|Q9BSD7\|NTPCR\_HUMAN | Q9BSD7 | NTPCR\_HUMAN | NTPCR | Cancer-related nucleoside-triphosphatase | 190 | 20.713 | 8.115151e+02 | 2.111819e+04 |  | 21118.193 |  |  |  | 43.7 |  |  |
| sp\|Q9H4A5\|GLP3L\_HUMAN | Q9H4A5 | GLP3L\_HUMAN | GOLPH3L | Golgi phosphoprotein 3-like | 285 | 32.767 | 1.165525e+03 | 4.798166e+04 |  | 47981.66 |  |  |  | 33.7 |  |  |
| sp\|P35270\|SPRE\_HUMAN | P35270 | SPRE\_HUMAN | SPR | Sepiapterin reductase | 261 | 28.048 | 1.779731e+03 | 6.271530e+04 |  | 62715.3 |  |  |  | 39.5 |  |  |
| sp\|Q6P3S1\|DEN1B\_HUMAN | Q6P3S1 | DEN1B\_HUMAN | DENND1B | DENN domain-containing protein 1B | 775 | 86.552 | 1.909240e+02 | 2.076132e+04 |  | 20761.316 |  |  |  | 23.5 |  |  |
| sp\|O14907\|TX1B3\_HUMAN | O14907 | TX1B3\_HUMAN | TAX1BP3 | Tax1-binding protein 3 | 124 | 13.735 | 2.785149e+03 | 4.806115e+04 |  | 39631.605 | 8429.549 |  |  | 44.4 | 13.7 |  |
| sp\|Q06828\|FMOD\_HUMAN | Q06828 | FMOD\_HUMAN | FMOD | Fibromodulin | 376 | 43.179 | 5.468008e+02 | 2.966326e+04 |  | 29663.258 |  |  |  | 11.7 |  |  |
| sp\|O00400\|ACATN\_HUMAN | O00400 | ACATN\_HUMAN | SLC33A1 | Acetyl-coenzyme A transporter 1 | 549 | 60.909 | 2.774835e+02 | 2.123419e+04 |  | 21234.191 |  |  |  | 19.1 |  |  |
| sp\|Q8NCL4\|GALT6\_HUMAN | Q8NCL4 | GALT6\_HUMAN | GALNT6 | Polypeptide N-acetylgalactosaminyltransferase 6 | 622 | 71.159 | 1.266546e+02 | 1.132317e+04 |  | 11323.165 |  |  |  | 28.6 |  |  |
| sp\|Q8N9F7\|GDPD1\_HUMAN | Q8N9F7 | GDPD1\_HUMAN | GDPD1 | Lysophospholipase D GDPD1 | 314 | 36.167 | 1.097555e+03 | 4.987188e+04 |  | 49871.88 |  |  |  | 34.4 |  |  |
| sp\|P15814\|IGLL1\_HUMAN | P15814 | IGLL1\_HUMAN | IGLL1 | Immunoglobulin lambda-like polypeptide 1 | 213 | 22.963 | 7.046368e+02 | 2.032877e+04 |  | 0.0 | 17818.639 | 2510.1316 |  | 41.8 | 35.2 | 28.2 |
| sp\|Q96RQ1\|ERGI2\_HUMAN | Q96RQ1 | ERGI2\_HUMAN | ERGIC2 | Endoplasmic reticulum-Golgi intermediate compartment protein 2 | 377 | 42.549 | 9.868158e+03 | 5.275244e+05 |  | 33190.918 | 286067.47 | 208266.06 |  | 24.7 | 15.1 | 7.4 |
| sp\|P24534\|EF1B\_HUMAN | P24534 | EF1B\_HUMAN | EEF1B2 | Elongation factor 1-beta | 225 | 24.764 | 1.350825e+03 | 4.202783e+04 |  | 10043.057 | 31984.773 |  |  | 9.8 | 15.1 |  |
| sp\|P50053\|KHK\_HUMAN | P50053 | KHK\_HUMAN | KHK | Ketohexokinase | 298 | 32.523 | 6.587578e+02 | 2.691742e+04 |  | 26917.424 |  |  |  | 57.0 |  |  |
| sp\|Q8WVE0\|EFMT1\_HUMAN | Q8WVE0 | EFMT1\_HUMAN | EEF1AKMT1 | EEF1A lysine methyltransferase 1 | 214 | 24.506 | 5.752622e+02 | 1.771150e+04 |  | 17711.502 |  |  |  | 48.6 |  |  |
| sp\|Q9BWQ6\|YIPF2\_HUMAN | Q9BWQ6 | YIPF2\_HUMAN | YIPF2 | Protein YIPF2 | 316 | 35.151 | 9.198338e+02 | 4.062227e+04 |  | 40622.273 |  |  |  | 7.9 |  |  |
| sp\|Q8WV19\|SFT2A\_HUMAN | Q8WV19 | SFT2A\_HUMAN | SFT2D1 | Vesicle transport protein SFT2A | 159 | 17.804 | 7.361690e+02 | 1.646692e+04 |  | 16466.916 |  |  |  | 17.0 |  |  |
| sp\|O95219\|SNX4\_HUMAN | O95219 | SNX4\_HUMAN | SNX4 | Sorting nexin-4 | 450 | 51.909 | 6.767872e+02 | 4.413793e+04 |  | 44137.926 |  |  |  | 25.6 |  |  |
| sp\|P17096\|HMGA1\_HUMAN | P17096 | HMGA1\_HUMAN | HMGA1 | High mobility group protein HMG-I/HMG-Y | 107 | 11.676 | 2.769074e+03 | 4.062055e+04 |  | 11710.062 | 28910.49 |  |  | 44.9 | 57.9 |  |
| sp\|P63244\|RACK1\_HUMAN | P63244 | RACK1\_HUMAN | RACK1 | Receptor of activated protein C kinase 1 | 317 | 35.077 | 1.131443e+03 | 4.986229e+04 |  | 7539.539 | 42322.746 |  |  | 54.9 | 12.0 |  |
| sp\|P07358\|CO8B\_HUMAN | P07358 | CO8B\_HUMAN | C8B | Complement component C8 beta chain | 591 | 66.948 | 1.654567e+03 | 1.391679e+05 |  | 56372.36 | 59058.332 | 23737.236 |  | 38.9 | 14.2 | 4.9 |
| sp\|Q63HM1\|KFA\_HUMAN | Q63HM1 | KFA\_HUMAN | AFMID | Kynurenine formamidase | 303 | 33.992 | 1.002370e+03 | 4.280769e+04 |  | 42807.695 |  |  |  | 43.2 |  |  |
| sp\|Q86V85\|GP180\_HUMAN | Q86V85 | GP180\_HUMAN | GPR180 | Integral membrane protein GPR180 | 440 | 49.395 | 2.521210e+03 | 1.564621e+05 |  | 44753.324 | 71347.44 | 40361.312 |  | 23.4 | 7.7 | 5.7 |
| sp\|Q9HCN4\|GPN1\_HUMAN | Q9HCN4 | GPN1\_HUMAN | GPN1 | GPN-loop GTPase 1 | 374 | 41.740 | 3.184253e+02 | 1.669849e+04 |  | 16698.486 |  |  |  | 19.0 |  |  |
| sp\|Q6UW68\|TM205\_HUMAN | Q6UW68 | TM205\_HUMAN | TMEM205 | Transmembrane protein 205 | 189 | 21.198 | 4.321871e+03 | 1.151023e+05 |  | 29336.494 |  | 85765.766 |  | 11.6 |  | 20.1 |
| sp\|Q9NRA0\|SPHK2\_HUMAN | Q9NRA0 | SPHK2\_HUMAN | SPHK2 | Sphingosine kinase 2 | 654 | 69.217 | 1.712114e+02 | 1.488890e+04 |  | 14888.901 |  |  |  | 28.6 |  |  |
| sp\|P51571\|SSRD\_HUMAN | P51571 | SSRD\_HUMAN | SSR4 | Translocon-associated protein subunit delta | 173 | 18.999 | 3.035138e+04 | 7.244798e+05 |  | 66258.6 | 524990.06 | 133231.16 |  | 14.5 | 24.9 | 13.3 |
| sp\|P51957\|NEK4\_HUMAN | P51957 | NEK4\_HUMAN | NEK4 | Serine/threonine-protein kinase Nek4 | 841 | 94.597 | 7.017473e+01 | 8.340177e+03 |  | 8340.177 |  |  |  | 26.9 |  |  |
| sp\|Q86TV6\|TTC7B\_HUMAN | Q86TV6 | TTC7B\_HUMAN | TTC7B | Tetratricopeptide repeat protein 7B | 843 | 94.179 | 1.625511e+02 | 1.923362e+04 |  | 19233.621 |  |  |  | 26.1 |  |  |
| sp\|Q8N9R8\|SCAI\_HUMAN | Q8N9R8 | SCAI\_HUMAN | SCAI | Protein SCAI | 606 | 70.399 | 2.098388e+02 | 1.855963e+04 |  | 18559.635 |  |  |  | 24.4 |  |  |
| sp\|Q99633\|PRP18\_HUMAN | Q99633 | PRP18\_HUMAN | PRPF18 | Pre-mRNA-splicing factor 18 | 342 | 39.860 | 4.161752e+02 | 2.084158e+04 |  | 20841.582 |  |  |  | 25.4 |  |  |
| sp\|Q9UK97\|FBX9\_HUMAN | Q9UK97 | FBX9\_HUMAN | FBXO9 | F-box only protein 9 | 447 | 52.329 | 3.736111e+02 | 2.456288e+04 |  | 24562.885 |  |  |  | 29.3 |  |  |
| sp\|O43548\|TGM5\_HUMAN | O43548 | TGM5\_HUMAN | TGM5 | Protein-glutamine gamma-glutamyltransferase 5 | 720 | 80.778 | 7.950206e+01 | 8.068422e+03 |  | 8068.422 |  |  |  | 20.8 |  |  |
| sp\|O75716\|STK16\_HUMAN | O75716 | STK16\_HUMAN | STK16 | Serine/threonine-protein kinase 16 | 305 | 34.656 | 5.117042e+02 | 2.227997e+04 |  | 22279.97 |  |  |  | 29.2 |  |  |
| sp\|Q9NPR2\|SEM4B\_HUMAN | Q9NPR2 | SEM4B\_HUMAN | SEMA4B | Semaphorin-4B | 837 | 92.766 | 2.086313e+01 | 2.431562e+03 |  | 2431.562 |  |  |  | 20.9 |  |  |
| sp\|Q9NZ45\|CISD1\_HUMAN | Q9NZ45 | CISD1\_HUMAN | CISD1 | CDGSH iron-sulfur domain-containing protein 1 | 108 | 12.199 | 2.821306e+03 | 4.324058e+04 |  | 9987.72 | 33252.863 |  |  | 52.8 | 75.9 |  |
| sp\|P16671\|CD36\_HUMAN | P16671 | CD36\_HUMAN | CD36 | Platelet glycoprotein 4 | 472 | 53.053 | 9.491693e+03 | 6.326606e+05 |  | 0.0 | 158603.31 | 474057.3 |  | 30.9 | 5.7 | 4.7 |
| sp\|P01619\|KV320\_HUMAN | P01619 | KV320\_HUMAN | IGKV3-20 | Immunoglobulin kappa variable 3-20 | 116 | 12.557 | 1.947401e+03 | 3.072264e+04 |  | 30722.639 |  |  |  | 25.0 |  |  |
| sp\|P00966\|ASSY\_HUMAN | P00966 | ASSY\_HUMAN | ASS1 | Argininosuccinate synthase | 412 | 46.530 | 1.298801e+02 | 7.592643e+03 |  | 7592.6426 |  |  |  | 22.3 |  |  |
| sp\|Q53FV1\|ORML2\_HUMAN | Q53FV1 | ORML2\_HUMAN | ORMDL2 | ORM1-like protein 2 | 153 | 17.363 | 6.840349e+02 | 1.492176e+04 |  | 14921.765 |  |  |  | 20.3 |  |  |
| sp\|O14569\|C56D2\_HUMAN | O14569 | C56D2\_HUMAN | CYB561D2 | Transmembrane reductase CYB561D2 | 222 | 23.974 | 4.004387e+03 | 1.206129e+05 |  | 71934.25 | 27985.484 | 20693.148 |  | 5.4 | 5.4 | 5.4 |
| sp\|P83916\|CBX1\_HUMAN | P83916 | CBX1\_HUMAN | CBX1 | Chromobox protein homolog 1 | 185 | 21.418 | 3.454869e+03 | 9.296676e+04 |  | 92966.76 |  |  |  | 29.7 |  |  |
| sp\|Q4G0W2\|DUS28\_HUMAN | Q4G0W2 | DUS28\_HUMAN | DUSP28 | Dual specificity phosphatase 28 | 176 | 18.324 | 4.884814e+02 | 1.124567e+04 |  | 11245.673 |  |  |  | 54.5 |  |  |
| sp\|P20160\|CAP7\_HUMAN | P20160 | CAP7\_HUMAN | AZU1 | Azurocidin | 251 | 26.886 | 4.890975e+02 | 1.652109e+04 |  | 16521.094 |  |  |  | 31.5 |  |  |
| sp\|O75635\|SPB7\_HUMAN | O75635 | SPB7\_HUMAN | SERPINB7 | Serpin B7 | 380 | 42.905 | 7.690292e+03 | 4.145414e+05 |  | 46330.887 | 167077.19 | 201133.28 |  | 40.5 | 40.3 | 26.8 |
| sp\|P51884\|LUM\_HUMAN | P51884 | LUM\_HUMAN | LUM | Lumican | 338 | 38.429 | 6.290762e+02 | 3.037243e+04 |  | 30372.428 |  |  |  | 35.8 |  |  |
| sp\|Q8TB24\|RIN3\_HUMAN | Q8TB24 | RIN3\_HUMAN | RIN3 | Ras and Rab interactor 3 | 985 | 107.854 | 1.198864e+02 | 1.624513e+04 |  | 16245.13 |  |  |  | 22.6 |  |  |
| sp\|Q99470\|SDF2\_HUMAN | Q99470 | SDF2\_HUMAN | SDF2 | Stromal cell-derived factor 2 | 211 | 23.026 | 5.098191e+03 | 1.474863e+05 |  | 107236.88 | 40249.445 |  |  | 24.6 | 18.5 |  |
| sp\|P05120\|PAI2\_HUMAN | P05120 | PAI2\_HUMAN | SERPINB2 | Plasminogen activator inhibitor 2 | 415 | 46.596 | 3.685833e+03 | 2.157752e+05 |  | 32590.943 | 121862.2 | 61322.04 |  | 29.2 | 27.7 | 11.1 |
| sp\|P02810\|PRPC\_HUMAN | P02810 | PRPC\_HUMAN | PRH1 | Salivary acidic proline-rich phosphoprotein 1/2 | 166 | 17.015 | 1.308898e+03 | 2.798047e+04 |  | 27980.467 |  |  |  | 26.5 |  |  |
| sp\|Q96JN8\|NEUL4\_HUMAN | Q96JN8 | NEUL4\_HUMAN | NEURL4 | Neuralized-like protein 4 | 1562 | 166.907 | 8.457067e+01 | 1.773419e+04 |  | 17734.195 |  |  |  | 18.5 |  |  |
| sp\|O75015\|FCG3B\_HUMAN | O75015 | FCG3B\_HUMAN | FCGR3B | Low affinity immunoglobulin gamma Fc region receptor III-B | 233 | 26.216 | 6.310262e+02 | 2.078409e+04 |  | 16227.026 | 4557.062 |  |  | 11.6 | 6.4 |  |
| sp\|P10301\|RRAS\_HUMAN | P10301 | RRAS\_HUMAN | RRAS | Ras-related protein R-Ras | 218 | 23.480 | 2.561569e+03 | 7.556511e+04 |  | 75565.11 |  |  |  | 46.8 |  |  |
| sp\|Q7L1V2\|MON1B\_HUMAN | Q7L1V2 | MON1B\_HUMAN | MON1B | Vacuolar fusion protein MON1 homolog B | 547 | 59.217 | 1.572791e+02 | 1.170131e+04 |  | 11701.31 |  |  |  | 30.5 |  |  |
| sp\|Q9Y5K3\|PCY1B\_HUMAN | Q9Y5K3 | PCY1B\_HUMAN | PCYT1B | Choline-phosphate cytidylyltransferase B | 369 | 41.940 | 1.283111e+03 | 6.760983e+04 |  | 19105.703 |  | 48504.13 |  | 48.0 |  | 8.4 |
| sp\|P42768\|WASP\_HUMAN | P42768 | WASP\_HUMAN | WAS | Wiskott-Aldrich syndrome protein | 502 | 52.913 | 9.965248e+02 | 6.624722e+04 |  | 66247.22 |  |  |  | 31.3 |  |  |
| sp\|Q8TBQ9\|KISHA\_HUMAN | Q8TBQ9 | KISHA\_HUMAN | TMEM167A | Protein kish-A | 72 | 8.060 | 6.211982e+04 | 6.290459e+05 |  | 156307.97 | 154574.25 | 318163.7 |  | 77.8 | 38.9 | 12.5 |
| sp\|Q8IWE4\|DCNL3\_HUMAN | Q8IWE4 | DCNL3\_HUMAN | DCUN1D3 | DCN1-like protein 3 | 304 | 34.291 | 6.071718e+02 | 2.615826e+04 |  | 26158.264 |  |  |  | 51.0 |  |  |
| sp\|Q8TCT9\|HM13\_HUMAN | Q8TCT9 | HM13\_HUMAN | HM13 | Minor histocompatibility antigen H13 | 377 | 41.488 | 2.610960e+03 | 1.360943e+05 |  | 61417.47 |  | 74676.82 |  | 15.9 |  | 16.4 |
| sp\|Q07065\|CKAP4\_HUMAN | Q07065 | CKAP4\_HUMAN | CKAP4 | Cytoskeleton-associated protein 4 | 602 | 66.022 | 1.797047e+03 | 1.490614e+05 |  | 14972.5 | 122086.84 | 12002.05 |  | 40.4 | 32.2 | 12.8 |
| sp\|Q13490\|BIRC2\_HUMAN | Q13490 | BIRC2\_HUMAN | BIRC2 | Baculoviral IAP repeat-containing protein 2 | 618 | 69.900 | 2.651837e+02 | 2.328848e+04 |  | 23288.484 |  |  |  | 48.1 |  |  |
| sp\|Q14739\|LBR\_HUMAN | Q14739 | LBR\_HUMAN | LBR | Delta(14)-sterol reductase LBR | 615 | 70.703 | 2.131555e+02 | 1.893440e+04 |  | 18934.398 |  |  |  | 26.3 |  |  |
| sp\|Q9H1C7\|CYTM1\_HUMAN | Q9H1C7 | CYTM1\_HUMAN | CYSTM1 | Cysteine-rich and transmembrane domain-containing protein 1 | 97 | 10.631 | 1.325817e+05 | 1.770822e+06 |  | 44034.938 | 94781.5 | 1632005.6 |  | 11.3 | 11.3 | 11.3 |
| sp\|A5PLN9\|TPC13\_HUMAN | A5PLN9 | TPC13\_HUMAN | TRAPPC13 | Trafficking protein particle complex subunit 13 | 417 | 46.524 | 3.224349e+02 | 1.884674e+04 |  | 18846.74 |  |  |  | 37.4 |  |  |
| sp\|P12956\|XRCC6\_HUMAN | P12956 | XRCC6\_HUMAN | XRCC6 | X-ray repair cross-complementing protein 6 | 609 | 69.843 | 5.286728e+02 | 4.639028e+04 |  | 46390.28 |  |  |  | 24.1 |  |  |
| sp\|P25815\|S100P\_HUMAN | P25815 | S100P\_HUMAN | S100P | Protein S100-P | 95 | 10.400 | 1.975796e+04 | 2.581622e+05 |  | 27794.148 | 101729.48 | 128638.58 |  | 53.7 | 13.7 | 24.2 |
| sp\|O43292\|GPAA1\_HUMAN | O43292 | GPAA1\_HUMAN | GPAA1 | Glycosylphosphatidylinositol anchor attachment 1 protein | 621 | 67.623 | 2.481873e+03 | 2.108585e+05 |  | 42621.26 |  | 168237.28 |  | 14.2 |  | 7.1 |
| sp\|O60831\|PRAF2\_HUMAN | O60831 | PRAF2\_HUMAN | PRAF2 | PRA1 family protein 2 | 178 | 19.258 | 2.450174e+03 | 5.928233e+04 |  | 48501.39 | 10780.945 |  |  | 24.7 | 21.3 |  |
| sp\|P62314\|SMD1\_HUMAN | P62314 | SMD1\_HUMAN | SNRPD1 | Small nuclear ribonucleoprotein Sm D1 | 119 | 13.282 | 9.882024e+02 | 1.649022e+04 |  | 16490.223 |  |  |  | 31.9 |  |  |
| sp\|Q5XUX1\|FBXW9\_HUMAN | Q5XUX1 | FBXW9\_HUMAN | FBXW9 | F-box/WD repeat-containing protein 9 | 458 | 50.760 | 2.465729e+02 | 1.572476e+04 |  | 15724.761 |  |  |  | 36.5 |  |  |
| sp\|Q8IXW5\|RPAP2\_HUMAN | Q8IXW5 | RPAP2\_HUMAN | RPAP2 | Putative RNA polymerase II subunit B1 CTD phosphatase RPAP2 | 612 | 69.509 | 1.765808e+02 | 1.542061e+04 |  | 15420.613 |  |  |  | 27.0 |  |  |
| sp\|P78314\|3BP2\_HUMAN | P78314 | 3BP2\_HUMAN | SH3BP2 | SH3 domain-binding protein 2 | 561 | 62.244 | 1.527421e+02 | 1.194465e+04 |  | 11944.6455 |  |  |  | 53.1 |  |  |
| sp\|Q8NEW0\|ZNT7\_HUMAN | Q8NEW0 | ZNT7\_HUMAN | SLC30A7 | Zinc transporter 7 | 376 | 41.626 | 1.286342e+04 | 6.727261e+05 |  | 42789.895 | 79846.945 | 550089.3 |  | 9.6 | 14.4 | 13.8 |
| sp\|O75901\|RASF9\_HUMAN | O75901 | RASF9\_HUMAN | RASSF9 | Ras association domain-containing protein 9 | 435 | 50.021 | 9.539996e+02 | 5.995394e+04 |  | 59953.94 |  |  |  | 16.1 |  |  |
| sp\|Q9NR31\|SAR1A\_HUMAN | Q9NR31 | SAR1A\_HUMAN | SAR1A | GTP-binding protein SAR1a | 198 | 22.367 | 2.261603e+04 | 6.355377e+05 |  | 443011.0 | 192526.72 |  |  | 22.7 | 21.2 |  |
| sp\|O43924\|PDE6D\_HUMAN | O43924 | PDE6D\_HUMAN | PDE6D | Retinal rod rhodopsin-sensitive cGMP 3',5'-cyclic phosphodiesterase subunit delta | 150 | 17.420 | 3.170320e+03 | 6.938545e+04 |  | 69385.45 |  |  |  | 38.0 |  |  |
| sp\|Q4G0A6\|MINY4\_HUMAN | Q4G0A6 | MINY4\_HUMAN | MINDY4 | Probable ubiquitin carboxyl-terminal hydrolase MINDY-4 | 757 | 84.372 | 1.014832e+02 | 1.075745e+04 |  | 10757.455 |  |  |  | 35.4 |  |  |
| sp\|Q8TDL5\|BPIB1\_HUMAN | Q8TDL5 | BPIB1\_HUMAN | BPIFB1 | BPI fold-containing family B member 1 | 484 | 52.442 | 1.161761e+04 | 7.654433e+05 |  | 765443.3 |  |  |  | 22.5 |  |  |
| sp\|P49913\|CAMP\_HUMAN | P49913 | CAMP\_HUMAN | CAMP | Cathelicidin antimicrobial peptide | 170 | 19.301 | 3.642428e+03 | 8.832587e+04 |  | 79120.39 | 9205.476 |  |  | 30.6 | 10.6 |  |
| sp\|Q9H7B4\|SMYD3\_HUMAN | Q9H7B4 | SMYD3\_HUMAN | SMYD3 | Histone-lysine N-methyltransferase SMYD3 | 428 | 49.097 | 2.594403e+02 | 1.600330e+04 |  | 16003.3 |  |  |  | 31.3 |  |  |
| sp\|O15530\|PDPK1\_HUMAN | O15530 | PDPK1\_HUMAN | PDPK1 | 3-phosphoinositide-dependent protein kinase 1 | 556 | 63.152 | 6.253488e+02 | 4.961655e+04 |  | 49616.55 |  |  |  | 39.4 |  |  |
| sp\|Q8IWP9\|CC28A\_HUMAN | Q8IWP9 | CC28A\_HUMAN | CCDC28A | Coiled-coil domain-containing protein 28A | 274 | 30.367 | 1.749974e+03 | 6.676526e+04 |  | 2280.695 | 64484.562 |  |  | 39.8 | 19.3 |  |
| sp\|Q6IC98\|GRAM4\_HUMAN | Q6IC98 | GRAM4\_HUMAN | GRAMD4 | GRAM domain-containing protein 4 | 578 | 66.408 | 1.532727e+02 | 1.278800e+04 |  | 12787.996 |  |  |  | 34.4 |  |  |
| sp\|Q96QF0\|RAB3I\_HUMAN | Q96QF0 | RAB3I\_HUMAN | RAB3IP | Rab-3A-interacting protein | 476 | 53.021 | 3.591068e+03 | 2.392152e+05 |  | 239215.19 |  |  |  | 23.7 |  |  |
| sp\|Q15077\|P2RY6\_HUMAN | Q15077 | P2RY6\_HUMAN | P2RY6 | P2Y purinoceptor 6 | 328 | 36.429 | 4.282396e+02 | 1.959978e+04 |  | 19599.78 |  |  |  | 29.9 |  |  |
| sp\|Q8N371\|KDM8\_HUMAN | Q8N371 | KDM8\_HUMAN | KDM8 | Bifunctional peptidase and arginyl-hydroxylase JMJD5 | 416 | 47.270 | 4.912218e+02 | 2.917296e+04 |  | 29172.955 |  |  |  | 33.9 |  |  |
| sp\|Q86YS6\|RAB43\_HUMAN | Q86YS6 | RAB43\_HUMAN | RAB43 | Ras-related protein Rab-43 | 212 | 23.339 | 8.585500e+03 | 2.517474e+05 |  | 251747.4 |  |  |  | 43.4 |  |  |
| sp\|Q9BVC6\|TM109\_HUMAN | Q9BVC6 | TM109\_HUMAN | TMEM109 | Transmembrane protein 109 | 243 | 26.210 | 1.878919e+04 | 6.187174e+05 |  | 51641.734 | 139336.56 | 427739.06 |  | 20.6 | 23.0 | 17.7 |
| sp\|Q9NRP0\|OSTC\_HUMAN | Q9NRP0 | OSTC\_HUMAN | OSTC | Oligosaccharyltransferase complex subunit | 149 | 16.829 | 1.976188e+03 | 4.178339e+04 |  | 41783.395 |  |  |  | 12.1 |  |  |
| sp\|Q9NXJ5\|PGPI\_HUMAN | Q9NXJ5 | PGPI\_HUMAN | PGPEP1 | Pyroglutamyl-peptidase 1 | 209 | 23.138 | 1.624005e+03 | 4.720961e+04 |  | 47209.61 |  |  |  | 18.7 |  |  |
| sp\|Q9HBI1\|PARVB\_HUMAN | Q9HBI1 | PARVB\_HUMAN | PARVB | Beta-parvin | 364 | 41.714 | 2.644354e+02 | 1.385858e+04 |  | 13858.576 |  |  |  | 34.6 |  |  |
| sp\|P62280\|RS11\_HUMAN | P62280 | RS11\_HUMAN | RPS11 | 40S ribosomal protein S11 | 158 | 18.431 | 8.781625e+02 | 2.033485e+04 |  | 20334.848 |  |  |  | 27.2 |  |  |
| sp\|O95619\|YETS4\_HUMAN | O95619 | YETS4\_HUMAN | YEATS4 | YEATS domain-containing protein 4 | 227 | 26.499 | 2.291789e+03 | 7.629944e+04 |  | 76299.44 |  |  |  | 15.4 |  |  |
| sp\|P62851\|RS25\_HUMAN | P62851 | RS25\_HUMAN | RPS25 | 40S ribosomal protein S25 | 125 | 13.742 | 1.932881e+03 | 3.337123e+04 |  | 33371.23 |  |  |  | 34.4 |  |  |
| sp\|Q8N335\|GPD1L\_HUMAN | Q8N335 | GPD1L\_HUMAN | GPD1L | Glycerol-3-phosphate dehydrogenase 1-like protein | 351 | 38.419 | 6.582393e+02 | 3.177218e+04 |  | 31772.18 |  |  |  | 33.0 |  |  |
| sp\|Q9NYZ2\|MFRN1\_HUMAN | Q9NYZ2 | MFRN1\_HUMAN | SLC25A37 | Mitoferrin-1 | 338 | 37.323 | 4.950981e+02 | 2.321587e+04 |  | 23215.87 |  |  |  | 38.5 |  |  |
| sp\|Q8NGD3\|OR4K5\_HUMAN | Q8NGD3 | OR4K5\_HUMAN | OR4K5 | Olfactory receptor 4K5 | 323 | 36.258 | 1.110567e+02 | 5.059015e+03 |  | 5059.0146 |  |  |  | 29.1 |  |  |
| sp\|Q3ZCW2\|LEGL\_HUMAN | Q3ZCW2 | LEGL\_HUMAN | LGALSL | Galectin-related protein | 172 | 18.986 | 8.014849e+02 | 1.911816e+04 |  | 19118.156 |  |  |  | 21.5 |  |  |
| sp\|Q9NWW6\|NRK1\_HUMAN | Q9NWW6 | NRK1\_HUMAN | NMRK1 | Nicotinamide riboside kinase 1 | 199 | 23.193 | 0.000000e+00 | 0.000000e+00 |  | 0.0 |  |  |  | 20.1 |  |  |
| sp\|Q9Y3A6\|TMED5\_HUMAN | Q9Y3A6 | TMED5\_HUMAN | TMED5 | Transmembrane emp24 domain-containing protein 5 | 229 | 26.005 | 2.575655e+04 | 8.415148e+05 |  | 66101.55 | 544628.0 | 230785.22 |  | 26.2 | 16.2 | 11.8 |
| sp\|O95858\|TSN15\_HUMAN | O95858 | TSN15\_HUMAN | TSPAN15 | Tetraspanin-15 | 294 | 33.165 | 3.843473e+03 | 1.601478e+05 |  | 160147.8 |  |  |  | 25.9 |  |  |
| sp\|P60484\|PTEN\_HUMAN | P60484 | PTEN\_HUMAN | PTEN | Phosphatidylinositol 3,4,5-trisphosphate 3-phosphatase and dual-specificity protein phosphatase PTEN | 403 | 47.166 | 6.276125e+02 | 3.719100e+04 |  | 37190.996 |  |  |  | 17.4 |  |  |
| sp\|Q9NZ01\|TECR\_HUMAN | Q9NZ01 | TECR\_HUMAN | TECR | Very-long-chain enoyl-CoA reductase | 308 | 36.034 | 4.716891e+03 | 2.135431e+05 |  | 36121.8 | 31884.133 | 145537.14 |  | 16.9 | 12.3 | 6.8 |
| sp\|Q9Y3Q3\|TMED3\_HUMAN | Q9Y3Q3 | TMED3\_HUMAN | TMED3 | Transmembrane emp24 domain-containing protein 3 | 217 | 24.777 | 9.749025e+02 | 3.034779e+04 |  | 30347.79 |  |  |  | 26.7 |  |  |
| sp\|P0DPK5\|H3Y2\_HUMAN | P0DPK5 | H3Y2\_HUMAN | H3Y2 | Histone H3.X | 147 | 16.466 | 3.006138e+02 | 6.218908e+03 |  | 6218.9077 |  |  |  | 41.5 |  |  |
| sp\|Q8N9U9\|SPOT1\_HUMAN | Q8N9U9 | SPOT1\_HUMAN | SPANXA2-OT1 | Putative uncharacterized protein SPANXA2-OT1 | 137 | 13.254 | 3.077939e+04 | 5.125358e+05 |  | 512535.75 |  |  |  | 75.2 |  |  |
| sp\|Q8NG68\|TTL\_HUMAN | Q8NG68 | TTL\_HUMAN | TTL | Tubulin--tyrosine ligase | 377 | 43.212 | 1.471528e+03 | 7.988955e+04 |  | 79889.555 |  |  |  | 29.4 |  |  |
| sp\|Q9Y276\|BCS1\_HUMAN | Q9Y276 | BCS1\_HUMAN | BCS1L | Mitochondrial chaperone BCS1 | 419 | 47.534 | 3.170543e+03 | 1.893455e+05 |  | 189345.55 |  |  |  | 22.7 |  |  |
| sp\|P59665\|DEF1\_HUMAN | P59665 | DEF1\_HUMAN | DEFA1 | Neutrophil defensin 1 | 94 | 10.201 | 0.000000e+00 | 0.000000e+00 |  | 0.0 |  |  |  | 43.6 |  |  |
| sp\|P62249\|RS16\_HUMAN | P62249 | RS16\_HUMAN | RPS16 | 40S ribosomal protein S16 | 146 | 16.445 | 1.724278e+03 | 3.562529e+04 |  | 35625.29 |  |  |  | 29.5 |  |  |
| sp\|Q9Y5U2\|TSSC4\_HUMAN | Q9Y5U2 | TSSC4\_HUMAN | TSSC4 | Protein TSSC4 | 329 | 34.326 | 0.000000e+00 | 0.000000e+00 |  | 0.0 |  |  |  | 25.8 |  |  |
| sp\|A6NI79\|CCD69\_HUMAN | A6NI79 | CCD69\_HUMAN | CCDC69 | Coiled-coil domain-containing protein 69 | 296 | 34.796 | 4.543568e+03 | 1.986294e+05 |  |  | 140840.98 | 57788.406 |  |  | 27.7 | 22.0 |
| sp\|O14656\|TOR1A\_HUMAN | O14656 | TOR1A\_HUMAN | TOR1A | Torsin-1A | 332 | 37.809 | 1.624986e+04 | 7.719018e+05 |  |  | 684230.6 | 87671.195 |  |  | 28.3 | 16.0 |
| sp\|O14657\|TOR1B\_HUMAN | O14657 | TOR1B\_HUMAN | TOR1B | Torsin-1B | 336 | 37.979 | 9.900922e+03 | 4.724287e+05 |  |  | 379268.28 | 93160.45 |  |  | 24.7 | 19.0 |
| sp\|O15078\|CE290\_HUMAN | O15078 | CE290\_HUMAN | CEP290 | Centrosomal protein of 290 kDa | 2479 | 290.386 | 3.942563e+02 | 1.438373e+05 |  |  | 143837.28 |  |  |  | 18.2 |  |
| sp\|O43504\|LTOR5\_HUMAN | O43504 | LTOR5\_HUMAN | LAMTOR5 | Ragulator complex protein LAMTOR5 | 91 | 9.614 | 8.195848e+03 | 9.899543e+04 |  |  | 98995.43 |  |  |  | 26.4 |  |
| sp\|O75718\|CRTAP\_HUMAN | O75718 | CRTAP\_HUMAN | CRTAP | Cartilage-associated protein | 401 | 46.562 | 3.754194e+03 | 2.196168e+05 |  |  | 219616.75 |  |  |  | 28.9 |  |
| sp\|O75787\|RENR\_HUMAN | O75787 | RENR\_HUMAN | ATP6AP2 | Renin receptor | 350 | 39.008 | 3.681677e+03 | 1.804332e+05 |  |  | 180433.22 |  |  |  | 15.4 |  |
| sp\|O94766\|B3GA3\_HUMAN | O94766 | B3GA3\_HUMAN | B3GAT3 | Galactosylgalactosylxylosylprotein 3-beta-glucuronosyltransferase 3 | 335 | 37.122 | 6.924699e+03 | 3.229605e+05 |  |  | 322960.5 |  |  |  | 31.3 |  |
| sp\|O95171\|SCEL\_HUMAN | O95171 | SCEL\_HUMAN | SCEL | Sciellin | 688 | 77.552 | 4.020405e+03 | 3.917238e+05 |  |  | 333114.38 | 58609.43 |  |  | 49.3 | 12.2 |
| sp\|O95196\|CSPG5\_HUMAN | O95196 | CSPG5\_HUMAN | CSPG5 | Chondroitin sulfate proteoglycan 5 | 566 | 60.016 | 1.746402e+05 | 1.316826e+07 |  |  | 13168263.0 |  |  |  | 12.9 |  |
| sp\|O95297\|MPZL1\_HUMAN | O95297 | MPZL1\_HUMAN | MPZL1 | Myelin protein zero-like protein 1 | 269 | 29.082 | 1.809048e+04 | 6.609849e+05 |  |  | 186056.33 | 474928.6 |  |  | 24.5 | 22.3 |
| sp\|O95479\|G6PE\_HUMAN | O95479 | G6PE\_HUMAN | H6PD | GDH/6PGL endoplasmic bifunctional protein | 791 | 88.893 | 1.084239e+03 | 1.210904e+05 |  |  | 121090.45 |  |  |  | 15.7 |  |
| sp\|O95807\|TM50A\_HUMAN | O95807 | TM50A\_HUMAN | TMEM50A | Transmembrane protein 50A | 157 | 17.400 | 6.081483e+04 | 1.329462e+06 |  |  | 413415.56 | 916046.3 |  |  | 14.6 | 5.1 |
| sp\|O95833\|CLIC3\_HUMAN | O95833 | CLIC3\_HUMAN | CLIC3 | Chloride intracellular channel protein 3 | 236 | 26.648 | 2.466732e+03 | 8.258550e+04 |  |  | 82585.5 |  |  |  | 14.0 |  |
| sp\|O96005\|CLPT1\_HUMAN | O96005 | CLPT1\_HUMAN | CLPTM1 | Cleft lip and palate transmembrane protein 1 | 669 | 76.097 | 4.567166e+02 | 4.366481e+04 |  |  | 43664.805 |  |  |  | 18.7 |  |
| sp\|P00751\|CFAB\_HUMAN | P00751 | CFAB\_HUMAN | CFB | Complement factor B | 764 | 85.533 | 3.436296e+03 | 3.692678e+05 |  |  | 193287.83 | 175979.98 |  |  | 24.1 | 12.8 |
| sp\|P01042\|KNG1\_HUMAN | P01042 | KNG1\_HUMAN | KNG1 | Kininogen-1 | 644 | 71.957 | 3.516860e+04 | 3.179401e+06 |  |  | 3179401.2 |  |  |  | 26.4 |  |
| sp\|P02743\|SAMP\_HUMAN | P02743 | SAMP\_HUMAN | APCS | Serum amyloid P-component | 223 | 25.387 | 3.567179e+03 | 1.137767e+05 |  |  | 113776.74 |  |  |  | 20.2 |  |
| sp\|P02748\|CO9\_HUMAN | P02748 | CO9\_HUMAN | C9 | Complement component C9 | 559 | 63.173 | 2.222225e+03 | 1.763749e+05 |  |  | 176374.86 |  |  |  | 16.3 |  |
| sp\|P02760\|AMBP\_HUMAN | P02760 | AMBP\_HUMAN | AMBP | Protein AMBP | 352 | 38.999 | 1.035585e+04 | 5.074071e+05 |  |  | 319490.0 | 187917.12 |  |  | 37.2 | 10.2 |
| sp\|P05090\|APOD\_HUMAN | P05090 | APOD\_HUMAN | APOD | Apolipoprotein D | 189 | 21.276 | 2.347528e+04 | 6.275060e+05 |  |  | 265196.53 | 362309.5 |  |  | 41.8 | 28.6 |
| sp\|P05556\|ITB1\_HUMAN | P05556 | ITB1\_HUMAN | ITGB1 | Integrin beta-1 | 798 | 88.415 | 8.112714e+03 | 9.011753e+05 |  |  | 362142.84 | 539032.44 |  |  | 20.6 | 18.2 |
| sp\|P06241\|FYN\_HUMAN | P06241 | FYN\_HUMAN | FYN | Tyrosine-protein kinase Fyn | 537 | 60.762 | 8.870398e+03 | 6.771616e+05 |  |  | 236299.02 | 440862.6 |  |  | 27.7 | 15.5 |
| sp\|P07602\|SAP\_HUMAN | P07602 | SAP\_HUMAN | PSAP | Prosaposin | 524 | 58.113 | 1.587320e+04 | 1.158924e+06 |  |  | 849858.06 | 309066.03 |  |  | 30.2 | 18.1 |
| sp\|P08727\|K1C19\_HUMAN | P08727 | K1C19\_HUMAN | KRT19 | Keratin, type I cytoskeletal 19 | 400 | 44.106 | 1.663611e+06 | 9.218637e+07 |  |  | 16071268.0 | 76115104.0 |  |  | 44.8 | 37.2 |
| sp\|P09496\|CLCA\_HUMAN | P09496 | CLCA\_HUMAN | CLTA | Clathrin light chain A | 248 | 27.077 | 2.639970e+04 | 8.980834e+05 |  |  | 670128.94 | 227954.5 |  |  | 16.9 | 23.4 |
| sp\|P10606\|COX5B\_HUMAN | P10606 | COX5B\_HUMAN | COX5B | Cytochrome c oxidase subunit 5B, mitochondrial | 129 | 13.696 | 7.857178e+03 | 1.352002e+05 |  |  | 135200.25 | 0.0 |  |  | 41.9 | 22.5 |
| sp\|P13073\|COX41\_HUMAN | P13073 | COX41\_HUMAN | COX4I1 | Cytochrome c oxidase subunit 4 isoform 1, mitochondrial | 169 | 19.577 | 7.250712e+03 | 1.783380e+05 |  |  | 178338.03 |  |  |  | 39.6 |  |
| sp\|P13671\|CO6\_HUMAN | P13671 | CO6\_HUMAN | C6 | Complement component C6 | 934 | 104.786 | 8.032248e+02 | 1.057444e+05 |  |  | 83620.07 | 22124.365 |  |  | 16.2 | 8.1 |
| sp\|P13797\|PLST\_HUMAN | P13797 | PLST\_HUMAN | PLS3 | Plastin-3 | 630 | 70.811 | 7.990066e+03 | 7.108341e+05 |  |  | 710834.06 |  |  |  | 30.3 |  |
| sp\|P18124\|RL7\_HUMAN | P18124 | RL7\_HUMAN | RPL7 | 60S ribosomal protein L7 | 248 | 29.226 | 1.037372e+03 | 3.809088e+04 |  |  | 38090.875 |  |  |  | 27.4 |  |
| sp\|P20929\|NEBU\_HUMAN | P20929 | NEBU\_HUMAN | NEB | Nebulin | 6669 | 772.914 | 6.653279e+01 | 6.460766e+04 |  |  | 64607.664 |  |  |  | 20.9 |  |
| sp\|P22830\|HEMH\_HUMAN | P22830 | HEMH\_HUMAN | FECH | Ferrochelatase, mitochondrial | 423 | 47.862 | 2.973800e+03 | 1.788215e+05 |  |  | 178821.48 |  |  |  | 23.6 |  |
| sp\|P23141\|EST1\_HUMAN | P23141 | EST1\_HUMAN | CES1 | Liver carboxylesterase 1 | 567 | 62.521 | 1.307768e+03 | 1.027244e+05 |  |  | 102724.42 |  |  |  | 30.3 |  |
| sp\|P26885\|FKBP2\_HUMAN | P26885 | FKBP2\_HUMAN | FKBP2 | Peptidyl-prolyl cis-trans isomerase FKBP2 | 142 | 15.649 | 2.855657e+04 | 5.614483e+05 |  |  | 561448.3 |  |  |  | 32.4 |  |
| sp\|P28331\|NDUS1\_HUMAN | P28331 | NDUS1\_HUMAN | NDUFS1 | NADH-ubiquinone oxidoreductase 75 kDa subunit, mitochondrial | 727 | 79.468 | 7.643036e+02 | 7.630893e+04 |  |  | 76308.93 |  |  |  | 20.9 |  |
| sp\|P30838\|AL3A1\_HUMAN | P30838 | AL3A1\_HUMAN | ALDH3A1 | Aldehyde dehydrogenase, dimeric NADP-preferring | 453 | 50.395 | 7.115982e+03 | 4.505463e+05 |  |  | 390414.56 | 60131.766 |  |  | 26.3 | 11.3 |
| sp\|P32926\|DSG3\_HUMAN | P32926 | DSG3\_HUMAN | DSG3 | Desmoglein-3 | 999 | 107.533 | 1.458106e+03 | 1.969917e+05 |  |  | 153557.98 | 43433.742 |  |  | 18.7 | 10.1 |
| sp\|P32942\|ICAM3\_HUMAN | P32942 | ICAM3\_HUMAN | ICAM3 | Intercellular adhesion molecule 3 | 547 | 59.541 | 2.438333e+03 | 1.824006e+05 |  |  | 182400.62 |  |  |  | 15.2 |  |
| sp\|P34897\|GLYM\_HUMAN | P34897 | GLYM\_HUMAN | SHMT2 | Serine hydroxymethyltransferase, mitochondrial | 504 | 55.993 | 2.301673e+03 | 1.619178e+05 |  |  | 161917.75 |  |  |  | 20.4 |  |
| sp\|P35232\|PHB1\_HUMAN | P35232 | PHB1\_HUMAN | PHB1 | Prohibitin 1 | 272 | 29.804 | 8.228676e+03 | 3.081213e+05 |  |  | 308121.28 |  |  |  | 36.4 |  |
| sp\|P36952\|SPB5\_HUMAN | P36952 | SPB5\_HUMAN | SERPINB5 | Serpin B5 | 375 | 42.100 | 6.745119e+03 | 3.567704e+05 |  |  | 284304.44 | 72466.0 |  |  | 32.0 | 19.5 |
| sp\|P39060\|COIA1\_HUMAN | P39060 | COIA1\_HUMAN | COL18A1 | Collagen alpha-1(XVIII) chain | 1754 | 178.188 | 8.900259e+02 | 1.992500e+05 |  |  | 199249.97 |  |  |  | 21.7 |  |
| sp\|P46781\|RS9\_HUMAN | P46781 | RS9\_HUMAN | RPS9 | 40S ribosomal protein S9 | 194 | 22.591 | 1.782987e+03 | 5.060587e+04 |  |  | 50605.867 |  |  |  | 40.2 |  |
| sp\|P48594\|SPB4\_HUMAN | P48594 | SPB4\_HUMAN | SERPINB4 | Serpin B4 | 390 | 44.854 | 1.329694e+04 | 7.493249e+05 |  |  | 749324.94 |  |  |  | 34.4 |  |
| sp\|P48723\|HSP13\_HUMAN | P48723 | HSP13\_HUMAN | HSPA13 | Heat shock 70 kDa protein 13 | 471 | 51.927 | 4.520808e+03 | 2.949351e+05 |  |  | 294935.12 |  |  |  | 20.2 |  |
| sp\|P51659\|DHB4\_HUMAN | P51659 | DHB4\_HUMAN | HSD17B4 | Peroxisomal multifunctional enzyme type 2 | 736 | 79.686 | 1.286648e+03 | 1.288128e+05 |  |  | 89488.195 | 39324.58 |  |  | 19.7 | 9.4 |
| sp\|P52179\|MYOM1\_HUMAN | P52179 | MYOM1\_HUMAN | MYOM1 | Myomesin-1 | 1685 | 187.627 | 4.777369e+02 | 1.126163e+05 |  |  | 112616.31 |  |  |  | 18.9 |  |
| sp\|P55145\|MANF\_HUMAN | P55145 | MANF\_HUMAN | MANF | Mesencephalic astrocyte-derived neurotrophic factor | 182 | 20.700 | 8.982116e+03 | 2.335964e+05 |  |  | 233596.4 |  |  |  | 16.5 |  |
| sp\|P56557\|TM50B\_HUMAN | P56557 | TM50B\_HUMAN | TMEM50B | Transmembrane protein 50B | 158 | 17.936 | 1.764494e+04 | 3.976153e+05 |  |  | 224165.66 | 173449.6 |  |  | 12.0 | 12.0 |
| sp\|P58107\|EPIPL\_HUMAN | P58107 | EPIPL\_HUMAN | EPPK1 | Epiplakin | 5088 | 555.658 | 3.650867e+02 | 2.548712e+05 |  |  | 254871.2 |  |  |  | 15.6 |  |
| sp\|P62269\|RS18\_HUMAN | P62269 | RS18\_HUMAN | RPS18 | 40S ribosomal protein S18 | 152 | 17.719 | 1.116120e+04 | 2.484663e+05 |  |  | 179280.84 | 69185.43 |  |  | 38.2 | 25.0 |
| sp\|P62753\|RS6\_HUMAN | P62753 | RS6\_HUMAN | RPS6 | 40S ribosomal protein S6 | 249 | 28.681 | 1.257567e+04 | 4.531506e+05 |  |  | 403489.38 | 49661.234 |  |  | 27.7 | 19.7 |
| sp\|P62913\|RL11\_HUMAN | P62913 | RL11\_HUMAN | RPL11 | 60S ribosomal protein L11 | 178 | 20.252 | 2.955822e+03 | 7.520788e+04 |  |  | 75207.875 |  |  |  | 30.9 |  |
| sp\|P80108\|PHLD\_HUMAN | P80108 | PHLD\_HUMAN | GPLD1 | Phosphatidylinositol-glycan-specific phospholipase D | 840 | 92.336 | 6.446629e+02 | 7.478610e+04 |  |  | 74786.1 |  |  |  | 19.9 |  |
| sp\|Q02218\|ODO1\_HUMAN | Q02218 | ODO1\_HUMAN | OGDH | 2-oxoglutarate dehydrogenase, mitochondrial | 1023 | 115.935 | 1.189830e+03 | 1.733072e+05 |  |  | 112108.6 | 61198.65 |  |  | 13.5 | 8.6 |
| sp\|Q02383\|SEMG2\_HUMAN | Q02383 | SEMG2\_HUMAN | SEMG2 | Semenogelin-2 | 582 | 65.444 | 1.272138e+03 | 1.045974e+05 |  |  | 104597.445 |  |  |  | 27.1 |  |
| sp\|Q02818\|NUCB1\_HUMAN | Q02818 | NUCB1\_HUMAN | NUCB1 | Nucleobindin-1 | 461 | 53.879 | 3.429131e+03 | 2.321244e+05 |  |  | 232124.38 |  |  |  | 33.4 |  |
| sp\|Q09327\|MGAT3\_HUMAN | Q09327 | MGAT3\_HUMAN | MGAT3 | Beta-1,4-mannosyl-glycoprotein 4-beta-N-acetylglucosaminyltransferase | 533 | 61.313 | 2.284276e+03 | 1.759618e+05 |  |  | 175961.77 |  |  |  | 27.4 |  |
| sp\|Q10471\|GALT2\_HUMAN | Q10471 | GALT2\_HUMAN | GALNT2 | Polypeptide N-acetylgalactosaminyltransferase 2 | 571 | 64.733 | 4.497874e+03 | 3.658054e+05 |  |  | 248249.62 | 117555.8 |  |  | 24.9 | 13.7 |
| sp\|Q11201\|SIA4A\_HUMAN | Q11201 | SIA4A\_HUMAN | ST3GAL1 | CMP-N-acetylneuraminate-beta-galactosamide-alpha-2,3-sialyltransferase 1 | 340 | 39.075 | 2.000378e+03 | 9.820377e+04 |  |  | 98203.766 |  |  |  | 25.0 |  |
| sp\|Q12913\|PTPRJ\_HUMAN | Q12913 | PTPRJ\_HUMAN | PTPRJ | Receptor-type tyrosine-protein phosphatase eta | 1337 | 145.941 | 2.128744e+03 | 3.903174e+05 |  |  | 286262.78 | 104054.6 |  |  | 17.7 | 8.5 |
| sp\|Q13261\|I15RA\_HUMAN | Q13261 | I15RA\_HUMAN | IL15RA | Interleukin-15 receptor subunit alpha | 267 | 28.233 | 1.289506e+02 | 4.574015e+03 |  |  | 4574.015 |  |  |  | 12.7 |  |
| sp\|Q13443\|ADAM9\_HUMAN | Q13443 | ADAM9\_HUMAN | ADAM9 | Disintegrin and metalloproteinase domain-containing protein 9 | 819 | 90.556 | 1.925820e+03 | 2.191038e+05 |  |  | 35729.934 | 183373.9 |  |  | 30.5 | 17.5 |
| sp\|Q13510\|ASAH1\_HUMAN | Q13510 | ASAH1\_HUMAN | ASAH1 | Acid ceramidase | 395 | 44.660 | 3.717022e+03 | 2.085600e+05 |  |  | 102521.81 | 106038.164 |  |  | 22.3 | 14.4 |
| sp\|Q13530\|SERC3\_HUMAN | Q13530 | SERC3\_HUMAN | SERINC3 | Serine incorporator 3 | 473 | 52.580 | 2.971525e+03 | 1.962986e+05 |  |  | 66701.61 | 129596.97 |  |  | 9.3 | 12.5 |
| sp\|Q13586\|STIM1\_HUMAN | Q13586 | STIM1\_HUMAN | STIM1 | Stromal interaction molecule 1 | 685 | 77.423 | 5.369606e+04 | 5.223115e+06 |  |  | 5223115.0 |  |  |  | 20.9 |  |
| sp\|Q14134\|TRI29\_HUMAN | Q14134 | TRI29\_HUMAN | TRIM29 | Tripartite motif-containing protein 29 | 588 | 65.835 | 3.390179e+03 | 2.804120e+05 |  |  | 156373.72 | 124038.3 |  |  | 29.1 | 12.9 |
| sp\|Q14160\|SCRIB\_HUMAN | Q14160 | SCRIB\_HUMAN | SCRIB | Protein scribble homolog | 1630 | 174.915 | 8.323764e+02 | 1.829212e+05 |  |  | 35524.28 | 147396.88 |  |  | 19.0 | 14.0 |
| sp\|Q14204\|DYHC1\_HUMAN | Q14204 | DYHC1\_HUMAN | DYNC1H1 | Cytoplasmic dynein 1 heavy chain 1 | 4646 | 532.408 | 3.406006e+01 | 2.278280e+04 |  |  | 22782.799 |  |  |  | 15.8 |  |
| sp\|Q14257\|RCN2\_HUMAN | Q14257 | RCN2\_HUMAN | RCN2 | Reticulocalbin-2 | 317 | 36.876 | 1.377842e+04 | 6.383523e+05 |  |  | 638352.3 |  |  |  | 19.2 |  |
| sp\|Q14315\|FLNC\_HUMAN | Q14315 | FLNC\_HUMAN | FLNC | Filamin-C | 2725 | 291.022 | 6.247410e+01 | 2.284247e+04 |  |  | 22842.467 |  |  |  | 18.3 |  |
| sp\|Q14554\|PDIA5\_HUMAN | Q14554 | PDIA5\_HUMAN | PDIA5 | Protein disulfide-isomerase A5 | 519 | 59.594 | 7.635563e+03 | 5.716902e+05 |  |  | 571690.2 |  |  |  | 31.8 |  |
| sp\|Q15006\|EMC2\_HUMAN | Q15006 | EMC2\_HUMAN | EMC2 | ER membrane protein complex subunit 2 | 297 | 34.834 | 4.157939e+03 | 1.819695e+05 |  |  | 181969.5 |  |  |  | 28.3 |  |
| sp\|Q15109\|RAGE\_HUMAN | Q15109 | RAGE\_HUMAN | AGER | Advanced glycosylation end product-specific receptor | 404 | 42.803 | 6.735533e+03 | 3.622124e+05 |  |  | 342357.0 | 19855.41 |  |  | 19.6 | 11.6 |
| sp\|Q15155\|NOMO1\_HUMAN | Q15155 | NOMO1\_HUMAN | NOMO1 | Nodal modulator 1 | 1222 | 134.324 | 3.914631e+03 | 6.606351e+05 |  |  | 660635.1 |  |  |  | 24.5 |  |
| sp\|Q15165\|PON2\_HUMAN | Q15165 | PON2\_HUMAN | PON2 | Serum paraoxonase/arylesterase 2 | 354 | 39.381 | 1.527980e+03 | 7.559995e+04 |  |  | 75599.95 |  |  |  | 20.1 |  |
| sp\|Q15293\|RCN1\_HUMAN | Q15293 | RCN1\_HUMAN | RCN1 | Reticulocalbin-1 | 331 | 38.890 | 4.032647e+04 | 1.970359e+06 |  |  | 1891608.8 | 78749.75 |  |  | 24.2 | 13.3 |
| sp\|Q15904\|VAS1\_HUMAN | Q15904 | VAS1\_HUMAN | ATP6AP1 | V-type proton ATPase subunit S1 | 470 | 52.026 | 7.530715e+03 | 4.922365e+05 |  |  | 492236.47 |  |  |  | 15.1 |  |
| sp\|Q16706\|MA2A1\_HUMAN | Q16706 | MA2A1\_HUMAN | MAN2A1 | Alpha-mannosidase 2 | 1144 | 131.141 | 9.976507e+03 | 1.643744e+06 |  |  | 1643743.9 |  |  |  | 14.2 |  |
| sp\|Q16891\|MIC60\_HUMAN | Q16891 | MIC60\_HUMAN | IMMT | MIC | 758 | 83.678 | 8.981274e+02 | 9.442052e+04 |  |  | 94420.52 |  |  |  | 16.2 |  |
| sp\|Q2M2I5\|K1C24\_HUMAN | Q2M2I5 | K1C24\_HUMAN | KRT24 | Keratin, type I cytoskeletal 24 | 525 | 55.087 | 4.268882e+04 | 2.954476e+06 |  |  | 2954475.8 |  |  |  | 23.0 |  |
| sp\|Q4G148\|GXLT1\_HUMAN | Q4G148 | GXLT1\_HUMAN | GXYLT1 | Glucoside xylosyltransferase 1 | 440 | 50.567 | 2.075399e+03 | 1.318518e+05 |  |  | 131851.77 |  |  |  | 19.5 |  |
| sp\|Q4KMQ2\|ANO6\_HUMAN | Q4KMQ2 | ANO6\_HUMAN | ANO6 | Anoctamin-6 | 910 | 106.165 | 4.576256e+03 | 6.103920e+05 |  |  | 192531.83 | 417860.16 |  |  | 17.1 | 20.9 |
| sp\|Q5VY43\|PEAR1\_HUMAN | Q5VY43 | PEAR1\_HUMAN | PEAR1 | Platelet endothelial aggregation receptor 1 | 1037 | 110.666 | 2.085001e+03 | 2.898929e+05 |  |  | 289892.94 |  |  |  | 29.5 |  |
| sp\|Q658P3\|STEA3\_HUMAN | Q658P3 | STEA3\_HUMAN | STEAP3 | Metalloreductase STEAP3 | 488 | 54.601 | 2.748517e+03 | 1.885455e+05 |  |  | 90474.51 | 98071.01 |  |  | 6.8 | 9.2 |
| sp\|Q6IAA8\|LTOR1\_HUMAN | Q6IAA8 | LTOR1\_HUMAN | LAMTOR1 | Ragulator complex protein LAMTOR1 | 161 | 17.745 | 3.051152e+04 | 6.802320e+05 |  |  | 456095.16 | 224136.83 |  |  | 34.2 | 6.8 |
| sp\|Q6UW63\|PLGT2\_HUMAN | Q6UW63 | PLGT2\_HUMAN | POGLUT2 | Protein O-glucosyltransferase 2 | 502 | 58.043 | 7.416634e+02 | 5.408462e+04 |  |  | 54084.625 |  |  |  | 15.5 |  |
| sp\|Q6ZN66\|GBP6\_HUMAN | Q6ZN66 | GBP6\_HUMAN | GBP6 | Guanylate-binding protein 6 | 633 | 72.427 | 1.814109e+03 | 1.650749e+05 |  |  | 165074.92 |  |  |  | 22.4 |  |
| sp\|Q6ZRP7\|QSOX2\_HUMAN | Q6ZRP7 | QSOX2\_HUMAN | QSOX2 | Sulfhydryl oxidase 2 | 698 | 77.529 | 2.580287e+03 | 2.513329e+05 |  |  | 78826.37 | 172506.55 |  |  | 22.3 | 17.2 |
| sp\|Q6ZV73\|FGD6\_HUMAN | Q6ZV73 | FGD6\_HUMAN | FGD6 | FYVE, RhoGEF and PH domain-containing protein 6 | 1430 | 160.816 | 5.568673e+03 | 1.125118e+06 |  |  | 828320.2 | 296797.84 |  |  | 24.9 | 13.6 |
| sp\|Q70UQ0\|IKIP\_HUMAN | Q70UQ0 | IKIP\_HUMAN | IKBIP | Inhibitor of nuclear factor kappa-B kinase-interacting protein | 350 | 39.309 | 2.925340e+03 | 1.444726e+05 |  |  | 97999.59 | 46473.023 |  |  | 27.1 | 19.4 |
| sp\|Q71RC9\|SMIM5\_HUMAN | Q71RC9 | SMIM5\_HUMAN | SMIM5 | Small integral membrane protein 5 | 77 | 8.540 | 9.958157e+05 | 1.068449e+07 |  |  | 1859187.6 | 8825307.0 |  |  | 26.0 | 24.7 |
| sp\|Q7Z2K6\|ERMP1\_HUMAN | Q7Z2K6 | ERMP1\_HUMAN | ERMP1 | Endoplasmic reticulum metallopeptidase 1 | 904 | 100.231 | 3.031939e+03 | 3.818033e+05 |  |  | 175309.78 | 206493.48 |  |  | 12.9 | 15.0 |
| sp\|Q7Z2K8\|GRIN1\_HUMAN | Q7Z2K8 | GRIN1\_HUMAN | GPRIN1 | G protein-regulated inducer of neurite outgrowth 1 | 1008 | 102.399 | 1.035480e+04 | 1.332155e+06 |  |  | 1332154.9 |  |  |  | 45.0 |  |
| sp\|Q7Z7M9\|GALT5\_HUMAN | Q7Z7M9 | GALT5\_HUMAN | GALNT5 | Polypeptide N-acetylgalactosaminyltransferase 5 | 940 | 106.266 | 5.729944e+02 | 7.650007e+04 |  |  | 76500.07 |  |  |  | 23.7 |  |
| sp\|Q86SF2\|GALT7\_HUMAN | Q86SF2 | GALT7\_HUMAN | GALNT7 | N-acetylgalactosaminyltransferase 7 | 657 | 75.389 | 7.473109e+02 | 7.078259e+04 |  |  | 70782.59 |  |  |  | 23.7 |  |
| sp\|Q86SR1\|GLT10\_HUMAN | Q86SR1 | GLT10\_HUMAN | GALNT10 | Polypeptide N-acetylgalactosaminyltransferase 10 | 603 | 68.992 | 4.716969e+03 | 4.088640e+05 |  |  | 321765.6 | 87098.43 |  |  | 28.7 | 10.0 |
| sp\|Q86UE4\|LYRIC\_HUMAN | Q86UE4 | LYRIC\_HUMAN | MTDH | Protein LYRIC | 582 | 63.837 | 6.017650e+02 | 4.826325e+04 |  |  | 48263.25 |  |  |  | 20.3 |  |
| sp\|Q8IXB1\|DJC10\_HUMAN | Q8IXB1 | DJC10\_HUMAN | DNAJC10 | DnaJ homolog subfamily C member 10 | 793 | 91.080 | 1.104139e+03 | 1.263467e+05 |  |  | 126346.71 |  |  |  | 8.3 |  |
| sp\|Q8IZN3\|ZDH14\_HUMAN | Q8IZN3 | ZDH14\_HUMAN | ZDHHC14 | Palmitoyltransferase ZDHHC14 | 488 | 53.388 | 1.219804e+04 | 8.181841e+05 |  |  | 89491.03 | 728693.06 |  |  | 14.1 | 20.1 |
| sp\|Q8N129\|CNPY4\_HUMAN | Q8N129 | CNPY4\_HUMAN | CNPY4 | Protein canopy homolog 4 | 248 | 28.310 | 2.314241e+04 | 8.231246e+05 |  |  | 823124.6 |  |  |  | 31.5 |  |
| sp\|Q8NBI6\|XXLT1\_HUMAN | Q8NBI6 | XXLT1\_HUMAN | XXYLT1 | Xyloside xylosyltransferase 1 | 393 | 43.807 | 3.255256e+03 | 1.791620e+05 |  |  | 179161.97 |  |  |  | 26.7 |  |
| sp\|Q8NBL1\|PGLT1\_HUMAN | Q8NBL1 | PGLT1\_HUMAN | POGLUT1 | Protein O-glucosyltransferase 1 | 392 | 46.189 | 4.713071e+03 | 2.735015e+05 |  |  | 217298.81 | 56202.67 |  |  | 30.9 | 9.4 |
| sp\|Q8NBP0\|TTC13\_HUMAN | Q8NBP0 | TTC13\_HUMAN | TTC13 | Tetratricopeptide repeat protein 13 | 860 | 96.813 | 4.666454e+03 | 5.675940e+05 |  |  | 567594.0 |  |  |  | 23.8 |  |
| sp\|Q8NEX9\|DR9C7\_HUMAN | Q8NEX9 | DR9C7\_HUMAN | SDR9C7 | Short-chain dehydrogenase/reductase family 9C member 7 | 313 | 35.263 | 9.633765e+03 | 4.268079e+05 |  |  | 130471.266 | 296336.66 |  |  | 38.7 | 14.4 |
| sp\|Q8NFQ8\|TOIP2\_HUMAN | Q8NFQ8 | TOIP2\_HUMAN | TOR1AIP2 | Torsin-1A-interacting protein 2 | 470 | 51.263 | 7.970814e+03 | 5.133622e+05 |  |  | 492640.44 | 20721.73 |  |  | 39.1 | 13.8 |
| sp\|Q8TAX7\|MUC7\_HUMAN | Q8TAX7 | MUC7\_HUMAN | MUC7 | Mucin-7 | 377 | 39.159 | 6.895548e+03 | 3.392482e+05 |  |  | 339248.16 |  |  |  | 17.8 |  |
| sp\|Q8TCJ2\|STT3B\_HUMAN | Q8TCJ2 | STT3B\_HUMAN | STT3B | Dolichyl-diphosphooligosaccharide--protein glycosyltransferase subunit STT3B | 826 | 93.674 | 4.687781e+03 | 5.517007e+05 |  |  | 133697.78 | 418002.88 |  |  | 18.2 | 12.3 |
| sp\|Q8TDN4\|CABL1\_HUMAN | Q8TDN4 | CABL1\_HUMAN | CABLES1 | CDK5 and ABL1 enzyme substrate 1 | 633 | 67.599 | 5.489729e+02 | 4.662387e+04 |  |  | 46623.867 |  |  |  | 11.4 |  |
| sp\|Q8TEM1\|PO210\_HUMAN | Q8TEM1 | PO210\_HUMAN | NUP210 | Nuclear pore membrane glycoprotein 210 | 1887 | 205.111 | 5.866196e+02 | 1.511690e+05 |  |  | 110442.73 | 40726.24 |  |  | 14.3 | 6.3 |
| sp\|Q92896\|GSLG1\_HUMAN | Q92896 | GSLG1\_HUMAN | GLG1 | Golgi apparatus protein 1 | 1179 | 134.552 | 2.007874e+03 | 3.394249e+05 |  |  | 222236.58 | 117188.336 |  |  | 27.7 | 14.8 |
| sp\|Q969N2\|PIGT\_HUMAN | Q969N2 | PIGT\_HUMAN | PIGT | GPI transamidase component PIG-T | 578 | 65.700 | 3.095578e+03 | 2.555196e+05 |  |  | 255519.6 |  |  |  | 21.1 |  |
| sp\|Q96AQ6\|PBIP1\_HUMAN | Q96AQ6 | PBIP1\_HUMAN | PBXIP1 | Pre-B-cell leukemia transcription factor-interacting protein 1 | 731 | 80.643 | 1.774175e+04 | 1.797547e+06 |  |  | 1797547.0 |  |  |  | 20.1 |  |
| sp\|Q96DR5\|BPIA2\_HUMAN | Q96DR5 | BPIA2\_HUMAN | BPIFA2 | BPI fold-containing family A member 2 | 249 | 27.011 | 3.226951e+03 | 1.095091e+05 |  |  | 109509.11 |  |  |  | 14.5 |  |
| sp\|Q96EU7\|C1GLC\_HUMAN | Q96EU7 | C1GLC\_HUMAN | C1GALT1C1 | C1GALT1-specific chaperone 1 | 318 | 36.382 | 2.723542e+03 | 1.244909e+05 |  |  | 124490.94 |  |  |  | 26.7 |  |
| sp\|Q96HE7\|ERO1A\_HUMAN | Q96HE7 | ERO1A\_HUMAN | ERO1A | ERO1-like protein alpha | 468 | 54.393 | 3.685828e+03 | 2.518809e+05 |  |  | 213064.17 | 38816.746 |  |  | 20.1 | 9.8 |
| sp\|Q99623\|PHB2\_HUMAN | Q99623 | PHB2\_HUMAN | PHB2 | Prohibitin-2 | 299 | 33.296 | 9.241645e+03 | 3.865970e+05 |  |  | 386597.0 |  |  |  | 57.9 |  |
| sp\|Q99714\|HCD2\_HUMAN | Q99714 | HCD2\_HUMAN | HSD17B10 | 3-hydroxyacyl-CoA dehydrogenase type-2 | 261 | 26.923 | 2.162996e+03 | 7.316381e+04 |  |  | 73163.81 |  |  |  | 31.4 |  |
| sp\|Q9BSF0\|SMAKA\_HUMAN | Q9BSF0 | SMAKA\_HUMAN | C2orf88 | Small membrane A-kinase anchor protein | 95 | 10.970 | 1.898018e+05 | 2.615917e+06 |  |  | 1333824.4 | 1282093.0 |  |  | 68.4 | 38.9 |
| sp\|Q9BTV4\|TMM43\_HUMAN | Q9BTV4 | TMM43\_HUMAN | TMEM43 | Transmembrane protein 43 | 400 | 44.876 | 3.138038e+03 | 1.769251e+05 |  |  | 53311.664 | 123613.43 |  |  | 13.5 | 12.0 |
| sp\|Q9H0V9\|LMA2L\_HUMAN | Q9H0V9 | LMA2L\_HUMAN | LMAN2L | VIP36-like protein | 348 | 39.711 | 2.021580e+04 | 1.008600e+06 |  |  | 333990.47 | 674609.44 |  |  | 17.5 | 15.2 |
| sp\|Q9H488\|OFUT1\_HUMAN | Q9H488 | OFUT1\_HUMAN | POFUT1 | GDP-fucose protein O-fucosyltransferase 1 | 388 | 43.956 | 3.518565e+03 | 1.943126e+05 |  |  | 194312.58 |  |  |  | 28.1 |  |
| sp\|Q9H9B4\|SFXN1\_HUMAN | Q9H9B4 | SFXN1\_HUMAN | SFXN1 | Sideroflexin-1 | 322 | 35.619 | 1.209731e+03 | 5.413621e+04 |  |  | 54136.21 |  |  |  | 27.3 |  |
| sp\|Q9HC84\|MUC5B\_HUMAN | Q9HC84 | MUC5B\_HUMAN | MUC5B | Mucin-5B | 5762 | 596.340 | 2.856989e+02 | 2.140522e+05 |  |  | 159289.34 | 54762.875 |  |  | 8.0 | 4.1 |
| sp\|Q9NS00\|C1GLT\_HUMAN | Q9NS00 | C1GLT\_HUMAN | C1GALT1 | Glycoprotein-N-acetylgalactosamine 3-beta-galactosyltransferase 1 | 363 | 42.203 | 2.114847e+03 | 1.121346e+05 |  |  | 112134.555 |  |  |  | 22.3 |  |
| sp\|Q9NUE0\|ZDH18\_HUMAN | Q9NUE0 | ZDH18\_HUMAN | ZDHHC18 | Palmitoyltransferase ZDHHC18 | 388 | 42.031 | 8.036344e+03 | 4.243707e+05 |  |  | 65240.758 | 359129.97 |  |  | 26.0 | 11.1 |
| sp\|Q9NZH8\|IL36G\_HUMAN | Q9NZH8 | IL36G\_HUMAN | IL36G | Interleukin-36 gamma | 169 | 18.721 | 1.472497e+04 | 3.463384e+05 |  |  | 170863.03 | 175475.4 |  |  | 44.4 | 37.3 |
| sp\|Q9UBC9\|SPRR3\_HUMAN | Q9UBC9 | SPRR3\_HUMAN | SPRR3 | Small proline-rich protein 3 | 169 | 18.154 | 1.687964e+05 | 3.849928e+06 |  |  | 2720714.2 | 1129214.0 |  |  | 89.9 | 66.9 |
| sp\|Q9UBS9\|SUCO\_HUMAN | Q9UBS9 | SUCO\_HUMAN | SUCO | SUN domain-containing ossification factor | 1254 | 139.430 | 6.020525e+02 | 1.054649e+05 |  |  | 105464.86 |  |  |  | 10.0 |  |
| sp\|Q9UIV8\|SPB13\_HUMAN | Q9UIV8 | SPB13\_HUMAN | SERPINB13 | Serpin B13 | 391 | 44.276 | 2.895569e+03 | 1.610718e+05 |  |  | 161071.78 |  |  |  | 43.2 |  |
| sp\|Q9UJZ1\|STML2\_HUMAN | Q9UJZ1 | STML2\_HUMAN | STOML2 | Stomatin-like protein 2, mitochondrial | 356 | 38.534 | 3.924901e+03 | 1.900159e+05 |  |  | 190015.9 |  |  |  | 31.5 |  |
| sp\|Q9ULI4\|KI26A\_HUMAN | Q9ULI4 | KI26A\_HUMAN | KIF26A | Kinesin-like protein KIF26A | 1882 | 194.590 | 3.664986e+02 | 8.960041e+04 |  |  | 89600.41 |  |  |  | 31.1 |  |
| sp\|Q9UMX5\|NENF\_HUMAN | Q9UMX5 | NENF\_HUMAN | NENF | Neudesin | 172 | 18.856 | 2.024750e+04 | 4.796650e+05 |  |  | 479665.03 |  |  |  | 53.5 |  |
| sp\|Q9UP52\|TFR2\_HUMAN | Q9UP52 | TFR2\_HUMAN | TFR2 | Transferrin receptor protein 2 | 801 | 88.755 | 9.246243e+02 | 1.031039e+05 |  |  | 103103.945 |  |  |  | 11.0 |  |
| sp\|Q9Y2G5\|OFUT2\_HUMAN | Q9Y2G5 | OFUT2\_HUMAN | POFUT2 | GDP-fucose protein O-fucosyltransferase 2 | 429 | 49.976 | 2.252336e+03 | 1.414203e+05 |  |  | 141420.31 |  |  |  | 17.5 |  |
| sp\|Q9Y4K1\|CRBG1\_HUMAN | Q9Y4K1 | CRBG1\_HUMAN | CRYBG1 | Beta/gamma crystallin domain-containing protein 1 | 1723 | 188.676 | 1.802356e+02 | 4.272425e+04 |  |  | 42724.254 |  |  |  | 20.5 |  |
| sp\|Q9Y5W9\|SNX11\_HUMAN | Q9Y5W9 | SNX11\_HUMAN | SNX11 | Sorting nexin-11 | 270 | 30.433 | 4.569886e+03 | 1.747299e+05 |  |  | 97953.19 | 76776.73 |  |  | 37.8 | 12.6 |
| sp\|A0A024RBG1\|NUD4B\_HUMAN | A0A024RBG1 | NUD4B\_HUMAN | NUDT4B | Diphosphoinositol polyphosphate phosphohydrolase NUDT4B | 181 | 20.434 | 4.721117e+03 | 1.212035e+05 |  |  | 121203.53 |  |  |  | 21.7 |  |
| sp\|O00148\|DX39A\_HUMAN | O00148 | DX39A\_HUMAN | DDX39A | ATP-dependent RNA helicase DDX39A | 427 | 49.130 | 1.978095e+03 | 1.220987e+05 |  |  | 122098.7 |  |  |  | 16.4 |  |
| sp\|O60911\|CATL2\_HUMAN | O60911 | CATL2\_HUMAN | CTSV | Cathepsin L2 | 334 | 37.329 | 4.530146e+02 | 2.124593e+04 |  |  | 21245.926 |  |  |  | 21.3 |  |
| sp\|P01780\|HV307\_HUMAN | P01780 | HV307\_HUMAN | IGHV3-7 | Immunoglobulin heavy variable 3-7 | 117 | 12.943 | 1.061831e+04 | 1.726662e+05 |  |  | 80572.79 | 92093.44 |  |  | 45.3 | 15.4 |
| sp\|P01861\|IGHG4\_HUMAN | P01861 | IGHG4\_HUMAN | IGHG4 | Immunoglobulin heavy constant gamma 4 | 327 | 43.832 | 8.148258e+03 | 4.487177e+05 |  |  | 448717.72 |  |  |  | 33.0 |  |
| sp\|P01860\|IGHG3\_HUMAN | P01860 | IGHG3\_HUMAN | IGHG3 | Immunoglobulin heavy constant gamma 3 | 377 | 49.093 | 2.802718e+04 | 1.728686e+06 |  |  | 1728686.2 |  |  |  | 44.6 |  |
| sp\|P48668\|K2C6C\_HUMAN | P48668 | K2C6C\_HUMAN | KRT6C | Keratin, type II cytoskeletal 6C | 564 | 60.025 | 1.810379e+06 | 1.365271e+08 |  |  | 43860688.0 | 92666400.0 |  |  | 66.7 | 57.6 |
| sp\|P09228\|CYTT\_HUMAN | P09228 | CYTT\_HUMAN | CST2 | Cystatin-SA | 141 | 16.445 | 1.452529e+04 | 3.001069e+05 |  |  | 300106.9 |  |  |  | 34.0 |  |
| sp\|P12236\|ADT3\_HUMAN | P12236 | ADT3\_HUMAN | SLC25A6 | ADP/ATP translocase 3 | 298 | 32.866 | 8.212003e+03 | 3.390885e+05 |  |  | 207934.48 | 131154.06 |  |  | 35.6 | 32.6 |
| sp\|P22105\|TENX\_HUMAN | P22105 | TENX\_HUMAN | TNXB | Tenascin-X | 4244 | 458.388 | 1.114597e+02 | 6.419016e+04 |  |  | 64190.156 |  |  |  | 17.6 |  |
| sp\|Q00535\|CDK5\_HUMAN | Q00535 | CDK5\_HUMAN | CDK5 | Cyclin-dependent-like kinase 5 | 292 | 33.304 | 2.817349e+05 | 1.178838e+07 |  |  | 11788382.0 |  |  |  | 29.1 |  |
| sp\|P51793\|CLCN4\_HUMAN | P51793 | CLCN4\_HUMAN | CLCN4 | H(+)/Cl(-) exchange transporter 4 | 760 | 84.917 | 6.110564e+03 | 6.519182e+05 |  |  | 82635.29 | 569282.94 |  |  | 13.9 | 12.1 |
| sp\|P0CG38\|POTEI\_HUMAN | P0CG38 | POTEI\_HUMAN | POTEI | POTE ankyrin domain family member I | 1075 | 121.282 | 3.257142e+06 | 4.963068e+08 |  |  | 496306816.0 |  |  |  | 28.7 |  |
| sp\|Q9BYX7\|ACTBM\_HUMAN | Q9BYX7 | ACTBM\_HUMAN | POTEKP | Putative beta-actin-like protein 3 | 375 | 42.016 | 9.252426e+06 | 4.884133e+08 |  |  | 488413280.0 |  |  |  | 45.3 |  |
| sp\|P68032\|ACTC\_HUMAN | P68032 | ACTC\_HUMAN | ACTC1 | Actin, alpha cardiac muscle 1 | 377 | 42.019 | 4.603463e+06 | 2.430231e+08 |  |  | 172095968.0 | 70927104.0 |  |  | 59.7 | 41.1 |
| sp\|P62807\|H2B1C\_HUMAN | P62807 | H2B1C\_HUMAN | H2BC4 | Histone H2B type 1-C/E/F/G/I | 126 | 13.906 | 7.944862e+04 | 1.388052e+06 |  |  | 1388052.0 |  |  |  | 69.0 |  |
| sp\|Q08043\|ACTN3\_HUMAN | Q08043 | ACTN3\_HUMAN | ACTN3 | Alpha-actinin-3 | 901 | 103.241 | 1.532415e+03 | 1.987677e+05 |  |  | 198767.69 |  |  |  | 23.0 |  |
| sp\|Q7Z3Y9\|K1C26\_HUMAN | Q7Z3Y9 | K1C26\_HUMAN | KRT26 | Keratin, type I cytoskeletal 26 | 468 | 51.911 | 2.391246e+05 | 1.559556e+07 |  |  | 15595556.0 |  |  |  | 12.0 |  |
| sp\|Q15714\|T22D1\_HUMAN | Q15714 | T22D1\_HUMAN | TSC22D1 | TSC22 domain family protein 1 | 1073 | 109.677 | 5.680698e+03 | 7.827704e+05 |  |  | 782770.4 |  |  |  | 6.2 |  |
| sp\|Q9Y4X0\|AMMR1\_HUMAN | Q9Y4X0 | AMMR1\_HUMAN | AMMECR1 | AMME syndrome candidate gene 1 protein | 333 | 35.463 | 1.402858e+04 | 6.250379e+05 |  |  | 209843.6 | 415194.34 |  |  | 17.1 | 21.0 |
| sp\|Q9Y5F7\|PCDGL\_HUMAN | Q9Y5F7 | PCDGL\_HUMAN | PCDHGC4 | Protocadherin gamma-C4 | 938 | 101.214 | 1.188608e+04 | 1.511459e+06 |  |  | 1419473.6 | 91985.586 |  |  | 10.8 | 8.1 |
| sp\|O14735\|CDIPT\_HUMAN | O14735 | CDIPT\_HUMAN | CDIPT | CDP-diacylglycerol--inositol 3-phosphatidyltransferase | 213 | 23.539 | 1.632529e+04 | 4.827987e+05 |  |  | 162652.97 | 320145.7 |  |  | 22.1 | 15.0 |
| sp\|O95214\|LERL1\_HUMAN | O95214 | LERL1\_HUMAN | LEPROTL1 | Leptin receptor overlapping transcript-like 1 | 131 | 14.428 | 1.430356e+04 | 2.592790e+05 |  |  | 259279.03 |  |  |  | 9.9 |  |
| sp\|P02511\|CRYAB\_HUMAN | P02511 | CRYAB\_HUMAN | CRYAB | Alpha-crystallin B chain | 175 | 20.159 | 6.000909e+03 | 1.519859e+05 |  |  | 84644.87 | 67340.98 |  |  | 29.1 | 29.7 |
| sp\|P04062\|GLCM\_HUMAN | P04062 | GLCM\_HUMAN | GBA | Lysosomal acid glucosylceramidase | 536 | 59.716 | 4.814198e+03 | 3.611868e+05 |  |  | 221934.4 | 139252.38 |  |  | 9.1 | 14.6 |
| sp\|P04217\|A1BG\_HUMAN | P04217 | A1BG\_HUMAN | A1BG | Alpha-1B-glycoprotein | 495 | 54.254 | 7.568527e+02 | 5.158938e+04 |  |  | 51589.38 |  |  |  | 18.6 |  |
| sp\|P06727\|APOA4\_HUMAN | P06727 | APOA4\_HUMAN | APOA4 | Apolipoprotein A-IV | 396 | 45.372 | 1.874764e+03 | 1.068690e+05 |  |  | 106868.98 |  |  |  | 26.0 |  |
| sp\|P07357\|CO8A\_HUMAN | P07357 | CO8A\_HUMAN | C8A | Complement component C8 alpha chain | 584 | 65.163 | 8.898549e+02 | 7.285132e+04 |  |  | 72851.32 |  |  |  | 17.3 |  |
| sp\|P15309\|PPAP\_HUMAN | P15309 | PPAP\_HUMAN | ACP3 | Prostatic acid phosphatase | 386 | 44.566 | 2.589704e+03 | 1.450010e+05 |  |  | 91057.57 | 53943.43 |  |  | 8.3 | 7.5 |
| sp\|P46779\|RL28\_HUMAN | P46779 | RL28\_HUMAN | RPL28 | 60S ribosomal protein L28 | 137 | 15.748 | 3.641910e+03 | 7.205627e+04 |  |  | 72056.266 |  |  |  | 21.9 |  |
| sp\|P55809\|SCOT1\_HUMAN | P55809 | SCOT1\_HUMAN | OXCT1 | Succinyl-CoA:3-ketoacid coenzyme A transferase 1, mitochondrial | 520 | 56.158 | 3.997533e+03 | 2.820466e+05 |  |  | 282046.56 |  |  |  | 19.4 |  |
| sp\|P59768\|GBG2\_HUMAN | P59768 | GBG2\_HUMAN | GNG2 | Guanine nucleotide-binding protein G(I)/G(S)/G(O) subunit gamma-2 | 71 | 7.850 | 6.242025e+04 | 6.156194e+05 |  |  | 443076.38 | 172543.05 |  |  | 69.0 | 46.5 |
| sp\|P67812\|SC11A\_HUMAN | P67812 | SC11A\_HUMAN | SEC11A | Signal peptidase complex catalytic subunit SEC11A | 179 | 20.625 | 3.339768e+04 | 8.654210e+05 |  |  | 430975.94 | 434445.06 |  |  | 47.5 | 13.4 |
| sp\|Q07020\|RL18\_HUMAN | Q07020 | RL18\_HUMAN | RPL18 | 60S ribosomal protein L18 | 188 | 21.634 | 2.896670e+03 | 7.873232e+04 |  |  | 62239.75 | 16492.568 |  |  | 28.7 | 12.2 |
| sp\|Q08357\|S20A2\_HUMAN | Q08357 | S20A2\_HUMAN | SLC20A2 | Sodium-dependent phosphate transporter 2 | 652 | 70.392 | 1.093371e+04 | 9.669589e+05 |  |  | 89195.31 | 877763.6 |  |  | 13.3 | 17.5 |
| sp\|Q13438\|OS9\_HUMAN | Q13438 | OS9\_HUMAN | OS9 | Protein | 667 | 75.562 | 2.311960e+02 | 2.194830e+04 |  |  | 21948.297 |  |  |  | 19.2 |  |
| sp\|Q96LJ7\|DHRS1\_HUMAN | Q96LJ7 | DHRS1\_HUMAN | DHRS1 | Dehydrogenase/reductase SDR family member 1 | 313 | 33.909 | 1.078807e+03 | 4.595955e+04 |  |  | 45959.55 |  |  |  | 41.5 |  |
| sp\|Q9NX62\|IMPA3\_HUMAN | Q9NX62 | IMPA3\_HUMAN | BPNT2 | Golgi-resident adenosine 3',5'-bisphosphate 3'-phosphatase | 359 | 38.681 | 8.470469e+03 | 4.116445e+05 |  |  | 299167.75 | 112476.74 |  |  | 11.7 | 12.8 |
| sp\|Q9ULT0\|TTC7A\_HUMAN | Q9ULT0 | TTC7A\_HUMAN | TTC7A | Tetratricopeptide repeat protein 7A | 858 | 96.185 | 5.611814e+02 | 6.781531e+04 |  |  | 33823.5 | 33991.812 |  |  | 21.1 | 14.7 |
| sp\|Q9Y6N5\|SQOR\_HUMAN | Q9Y6N5 | SQOR\_HUMAN | SQOR | Sulfide:quinone oxidoreductase, mitochondrial | 450 | 49.961 | 6.890482e+02 | 4.325117e+04 |  |  | 43251.17 |  |  |  | 22.2 |  |
| sp\|P0C7P4\|UCRIL\_HUMAN | P0C7P4 | UCRIL\_HUMAN | UQCRFS1P1 | Putative cytochrome b-c1 complex subunit Rieske-like protein 1 | 283 | 30.815 | 4.278404e+03 | 1.656384e+05 |  |  | 150970.56 | 14667.876 |  |  | 24.7 | 4.7 |
| sp\|O15431\|COPT1\_HUMAN | O15431 | COPT1\_HUMAN | SLC31A1 | High affinity copper uptake protein 1 | 190 | 21.091 | 1.350328e+03 | 3.578111e+04 |  |  | 35781.105 |  |  |  | 5.8 |  |
| sp\|O43760\|SNG2\_HUMAN | O43760 | SNG2\_HUMAN | SYNGR2 | Synaptogyrin-2 | 224 | 24.810 | 3.732648e+04 | 1.163485e+06 |  |  | 206067.55 | 957417.75 |  |  | 22.8 | 20.5 |
| sp\|P08559\|ODPA\_HUMAN | P08559 | ODPA\_HUMAN | PDHA1 | Pyruvate dehydrogenase E1 component subunit alpha, somatic form, mitochondrial | 390 | 43.296 | 1.198526e+03 | 6.519471e+04 |  |  | 65194.707 |  |  |  | 29.5 |  |
| sp\|P22307\|SCP2\_HUMAN | P22307 | SCP2\_HUMAN | SCP2 | Sterol carrier protein 2 | 547 | 58.994 | 1.589209e+03 | 1.177893e+05 |  |  | 117789.34 |  |  |  | 13.0 |  |
| sp\|P36578\|RL4\_HUMAN | P36578 | RL4\_HUMAN | RPL4 | 60S ribosomal protein L4 | 427 | 47.697 | 1.163651e+03 | 6.973180e+04 |  |  | 69731.805 |  |  |  | 32.3 |  |
| sp\|Q86UD5\|SL9B2\_HUMAN | Q86UD5 | SL9B2\_HUMAN | SLC9B2 | Sodium/hydrogen exchanger 9B2 | 537 | 57.564 | 2.815452e+03 | 2.036180e+05 |  |  | 36074.15 | 167543.86 |  |  | 21.4 | 6.7 |
| sp\|Q8WWI1\|LMO7\_HUMAN | Q8WWI1 | LMO7\_HUMAN | LMO7 | LIM domain only protein 7 | 1683 | 192.696 | 3.723925e+03 | 9.015520e+05 |  |  | 901552.0 |  |  |  | 17.2 |  |
| sp\|Q9HCJ1\|ANKH\_HUMAN | Q9HCJ1 | ANKH\_HUMAN | ANKH | Progressive ankylosis protein homolog | 492 | 54.241 | 1.025567e+04 | 6.988901e+05 |  |  | 41888.867 | 657001.25 |  |  | 9.6 | 17.7 |
| sp\|Q9Y5U9\|IR3IP\_HUMAN | Q9Y5U9 | IR3IP\_HUMAN | IER3IP1 | Immediate early response 3-interacting protein 1 | 82 | 8.969 | 1.120983e+04 | 1.263165e+05 |  |  | 126316.52 |  |  |  | 34.1 |  |
| sp\|A0A075B6I9\|LV746\_HUMAN | A0A075B6I9 | LV746\_HUMAN | IGLV7-46 | Immunoglobulin lambda variable 7-46 | 117 | 12.468 | 1.502141e+04 | 2.353015e+05 |  |  | 235301.47 |  |  |  | 21.4 |  |
| sp\|Q9GZU7\|CTDS1\_HUMAN | Q9GZU7 | CTDS1\_HUMAN | CTDSP1 | Carboxy-terminal domain RNA polymerase II polypeptide A small phosphatase 1 | 261 | 29.203 | 2.674997e+03 | 9.814496e+04 |  |  | 60256.42 | 37888.535 |  |  | 11.5 | 7.7 |
| sp\|O75356\|ENTP5\_HUMAN | O75356 | ENTP5\_HUMAN | ENTPD5 | Nucleoside diphosphate phosphatase ENTPD5 | 428 | 47.517 | 1.691641e+03 | 1.009891e+05 |  |  | 100989.05 |  |  |  | 17.8 |  |
| sp\|O75367\|H2AY\_HUMAN | O75367 | H2AY\_HUMAN | MACROH2A1 | Core histone macro-H2A.1 | 372 | 39.184 | 2.746620e+03 | 1.352149e+05 |  |  | 116110.766 | 19104.125 |  |  | 28.5 | 17.5 |
| sp\|P39023\|RL3\_HUMAN | P39023 | RL3\_HUMAN | RPL3 | 60S ribosomal protein L3 | 403 | 46.109 | 1.107611e+03 | 6.416381e+04 |  |  | 64163.812 |  |  |  | 22.3 |  |
| sp\|P46776\|RL27A\_HUMAN | P46776 | RL27A\_HUMAN | RPL27A | 60S ribosomal protein L27a | 148 | 16.561 | 3.881107e+03 | 8.075313e+04 |  |  | 70461.33 | 10291.798 |  |  | 27.0 | 15.5 |
| sp\|P49585\|PCY1A\_HUMAN | P49585 | PCY1A\_HUMAN | PCYT1A | Choline-phosphate cytidylyltransferase A | 367 | 41.731 | 1.021441e+03 | 5.355366e+04 |  |  | 53553.656 |  |  |  | 37.9 |  |
| sp\|P61604\|CH10\_HUMAN | P61604 | CH10\_HUMAN | HSPE1 | 10 kDa heat shock protein, mitochondrial | 102 | 10.932 | 8.752219e+03 | 1.202084e+05 |  |  | 120208.44 |  |  |  | 55.9 |  |
| sp\|Q5J8M3\|EMC4\_HUMAN | Q5J8M3 | EMC4\_HUMAN | EMC4 | ER membrane protein complex subunit 4 | 183 | 20.087 | 4.079757e+03 | 1.029595e+05 |  |  | 81712.39 | 21247.14 |  |  | 26.2 | 16.4 |
| sp\|Q92506\|DHB8\_HUMAN | Q92506 | DHB8\_HUMAN | HSD17B8 | (3R)-3-hydroxyacyl-CoA dehydrogenase | 261 | 26.974 | 1.586008e+03 | 5.374870e+04 |  |  | 53748.703 |  |  |  | 33.0 |  |
| sp\|Q99618\|CDCA3\_HUMAN | Q99618 | CDCA3\_HUMAN | CDCA3 | Cell division cycle-associated protein 3 | 268 | 28.998 | 1.052478e+03 | 3.834406e+04 |  |  | 38344.055 |  |  |  | 41.8 |  |
| sp\|O43759\|SNG1\_HUMAN | O43759 | SNG1\_HUMAN | SYNGR1 | Synaptogyrin-1 | 233 | 25.456 | 1.425959e+04 | 4.560521e+05 |  |  | 166573.66 | 289478.47 |  |  | 14.6 | 5.2 |
| sp\|P63167\|DYL1\_HUMAN | P63167 | DYL1\_HUMAN | DYNLL1 | Dynein light chain 1, cytoplasmic | 89 | 10.366 | 2.077633e+04 | 2.705809e+05 |  |  | 270580.94 |  |  |  | 24.7 |  |
| sp\|Q9BX97\|PLVAP\_HUMAN | Q9BX97 | PLVAP\_HUMAN | PLVAP | Plasmalemma vesicle-associated protein | 442 | 50.594 | 8.720747e+03 | 5.543319e+05 |  |  | 58512.79 | 495819.1 |  |  | 25.6 | 21.3 |
| sp\|Q9Y6A9\|SPCS1\_HUMAN | Q9Y6A9 | SPCS1\_HUMAN | SPCS1 | Signal peptidase complex subunit 1 | 169 | 18.298 | 2.662077e+03 | 6.119860e+04 |  |  | 28941.2 | 32257.395 |  |  | 27.2 | 7.1 |
| sp\|P42677\|RS27\_HUMAN | P42677 | RS27\_HUMAN | RPS27 | 40S ribosomal protein S27 | 84 | 9.461 | 4.885714e+03 | 5.807406e+04 |  |  | 58074.062 |  |  |  | 44.0 |  |
| sp\|P00367\|DHE3\_HUMAN | P00367 | DHE3\_HUMAN | GLUD1 | Glutamate dehydrogenase 1, mitochondrial | 558 | 61.398 | 3.360963e+02 | 2.592598e+04 |  |  | 25925.977 |  |  |  | 8.1 |  |
| sp\|P18077\|RL35A\_HUMAN | P18077 | RL35A\_HUMAN | RPL35A | 60S ribosomal protein L35a | 110 | 12.538 | 1.264343e+04 | 1.991638e+05 |  |  | 199163.8 |  |  |  | 34.5 |  |
| sp\|Q8NCT1\|ARRD4\_HUMAN | Q8NCT1 | ARRD4\_HUMAN | ARRDC4 | Arrestin domain-containing protein 4 | 418 | 45.479 | 2.987824e+03 | 1.707195e+05 |  |  | 95173.34 | 75546.14 |  |  | 17.7 | 14.8 |
| sp\|P16150\|LEUK\_HUMAN | P16150 | LEUK\_HUMAN | SPN | Leukosialin | 400 | 40.322 | 1.283032e+03 | 6.499752e+04 |  |  | 49397.895 | 15599.628 |  |  | 28.5 | 13.0 |
| sp\|P36955\|PEDF\_HUMAN | P36955 | PEDF\_HUMAN | SERPINF1 | Pigment epithelium-derived factor | 418 | 46.312 | 1.524102e+03 | 8.867979e+04 |  |  | 40341.04 | 48338.746 |  |  | 19.1 | 10.0 |
| sp\|P10253\|LYAG\_HUMAN | P10253 | LYAG\_HUMAN | GAA | Lysosomal alpha-glucosidase | 952 | 105.324 | 4.707128e+02 | 6.228745e+04 |  |  | 40047.566 | 22239.88 |  |  | 8.2 | 7.6 |
| sp\|P55273\|CDN2D\_HUMAN | P55273 | CDN2D\_HUMAN | CDKN2D | Cyclin-dependent kinase 4 inhibitor D | 166 | 17.700 | 8.328127e+02 | 1.851986e+04 |  |  | 18519.863 |  |  |  | 25.9 |  |
| sp\|P62310\|LSM3\_HUMAN | P62310 | LSM3\_HUMAN | LSM3 | U6 snRNA-associated Sm-like protein LSm3 | 102 | 11.845 | 4.401061e+03 | 6.549523e+04 |  |  | 49853.688 | 15641.539 |  |  | 19.6 | 11.8 |
| sp\|Q12797\|ASPH\_HUMAN | Q12797 | ASPH\_HUMAN | ASPH | Aspartyl/asparaginyl beta-hydroxylase | 758 | 85.863 | 3.608270e+02 | 3.892443e+04 |  |  | 38924.426 |  |  |  | 6.2 |  |
| sp\|P62750\|RL23A\_HUMAN | P62750 | RL23A\_HUMAN | RPL23A | 60S ribosomal protein L23a | 156 | 17.695 | 5.143555e+03 | 1.143487e+05 |  |  | 96393.8 | 17954.86 |  |  | 35.3 | 17.3 |
| sp\|P62701\|RS4X\_HUMAN | P62701 | RS4X\_HUMAN | RPS4X | 40S ribosomal protein S4, X isoform | 263 | 29.598 | 8.165805e+02 | 3.036537e+04 |  |  | 30365.367 |  |  |  | 16.0 |  |
| sp\|O15533\|TPSN\_HUMAN | O15533 | TPSN\_HUMAN | TAPBP | Tapasin | 448 | 47.571 | 3.447264e+03 | 2.060316e+05 |  |  | 206031.64 |  |  |  | 10.3 |  |
| sp\|P62424\|RL7A\_HUMAN | P62424 | RL7A\_HUMAN | RPL7A | 60S ribosomal protein L7a | 266 | 29.996 | 1.228704e+03 | 4.630497e+04 |  |  | 46304.97 |  |  |  | 25.6 |  |
| sp\|P31930\|QCR1\_HUMAN | P31930 | QCR1\_HUMAN | UQCRC1 | Cytochrome b-c1 complex subunit 1, mitochondrial | 480 | 52.646 | 1.514235e+03 | 1.001557e+05 |  |  | 100155.7 |  |  |  | 6.5 |  |
| sp\|P33908\|MA1A1\_HUMAN | P33908 | MA1A1\_HUMAN | MAN1A1 | Mannosyl-oligosaccharide 1,2-alpha-mannosidase IA | 653 | 72.969 | 9.429631e+02 | 8.644706e+04 |  |  | 86447.06 |  |  |  | 15.3 |  |
| sp\|Q0VGL1\|LTOR4\_HUMAN | Q0VGL1 | LTOR4\_HUMAN | LAMTOR4 | Ragulator complex protein LAMTOR4 | 99 | 10.741 | 3.794290e+04 | 5.120264e+05 |  |  | 512026.38 |  |  |  | 24.2 |  |
| sp\|P09958\|FURIN\_HUMAN | P09958 | FURIN\_HUMAN | FURIN | Furin | 794 | 86.678 | 2.125943e+02 | 2.315142e+04 |  |  | 23151.424 |  |  |  | 12.0 |  |
| sp\|P19971\|TYPH\_HUMAN | P19971 | TYPH\_HUMAN | TYMP | Thymidine phosphorylase | 482 | 49.955 | 3.834698e+03 | 2.406729e+05 |  |  | 44601.58 | 196071.36 |  |  | 38.4 | 15.8 |
| sp\|P42773\|CDN2C\_HUMAN | P42773 | CDN2C\_HUMAN | CDKN2C | Cyclin-dependent kinase 4 inhibitor C | 168 | 18.127 | 3.971732e+03 | 9.045300e+04 |  |  | 90453.0 |  |  |  | 35.1 |  |
| sp\|Q9HCM1\|RESF1\_HUMAN | Q9HCM1 | RESF1\_HUMAN | RESF1 | Retroelement silencing factor 1 | 1747 | 194.857 | 1.953720e+02 | 4.782946e+04 |  |  | 47829.457 |  |  |  | 24.8 |  |
| sp\|P00734\|THRB\_HUMAN | P00734 | THRB\_HUMAN | F2 | Prothrombin | 622 | 70.037 | 9.138012e+02 | 8.040747e+04 |  |  | 80407.47 |  |  |  | 9.3 |  |
| sp\|Q6ZVZ8\|ASB18\_HUMAN | Q6ZVZ8 | ASB18\_HUMAN | ASB18 | Ankyrin repeat and SOCS box protein 18 | 466 | 50.803 | 1.421694e+04 | 9.074292e+05 |  |  | 907429.2 |  |  |  | 19.5 |  |
| sp\|P16219\|ACADS\_HUMAN | P16219 | ACADS\_HUMAN | ACADS | Short-chain specific acyl-CoA dehydrogenase, mitochondrial | 412 | 44.297 | 8.885825e+02 | 4.945261e+04 |  |  | 49452.61 |  |  |  | 14.1 |  |
| sp\|P19404\|NDUV2\_HUMAN | P19404 | NDUV2\_HUMAN | NDUFV2 | NADH dehydrogenase [ubiquinone] flavoprotein 2, mitochondrial | 249 | 27.392 | 7.388850e+02 | 2.542832e+04 |  |  | 25428.324 |  |  |  | 49.8 |  |
| sp\|P05783\|K1C18\_HUMAN | P05783 | K1C18\_HUMAN | KRT18 | Keratin, type I cytoskeletal 18 | 430 | 48.058 | 1.948392e+04 | 1.176411e+06 |  |  | 1176411.2 |  |  |  | 20.7 |  |
| sp\|P30825\|SL7A1\_HUMAN | P30825 | SL7A1\_HUMAN | SLC7A1 | High affinity cationic amino acid transporter 1 | 629 | 67.638 | 1.108096e+03 | 9.416405e+04 |  |  | 94164.05 |  |  |  | 8.1 |  |
| sp\|P10321\|HLAC\_HUMAN | P10321 | HLAC\_HUMAN | HLA-C | HLA class I histocompatibility antigen, C alpha chain | 366 | 40.649 | 1.794085e+03 | 9.162417e+04 |  |  | 91624.17 |  |  |  | 24.0 |  |
| sp\|P49748\|ACADV\_HUMAN | P49748 | ACADV\_HUMAN | ACADVL | Very long-chain specific acyl-CoA dehydrogenase, mitochondrial | 655 | 70.390 | 6.655249e+02 | 5.885623e+04 |  |  | 31563.838 | 27292.389 |  |  | 24.6 | 15.1 |
| sp\|P08842\|STS\_HUMAN | P08842 | STS\_HUMAN | STS | Steryl-sulfatase | 583 | 65.492 | 7.858581e+02 | 6.466206e+04 |  |  | 64662.062 |  |  |  | 16.1 |  |
| sp\|P55899\|FCGRN\_HUMAN | P55899 | FCGRN\_HUMAN | FCGRT | IgG receptor FcRn large subunit p51 | 365 | 39.743 | 4.112373e+02 | 2.053385e+04 |  |  | 20533.846 |  |  |  | 7.4 |  |
| sp\|Q99674\|CGRE1\_HUMAN | Q99674 | CGRE1\_HUMAN | CGREF1 | Cell growth regulator with EF hand domain protein 1 | 318 | 33.456 | 8.779562e+02 | 3.690320e+04 |  |  | 36903.2 |  |  |  | 41.5 |  |
| sp\|P47900\|P2RY1\_HUMAN | P47900 | P2RY1\_HUMAN | P2RY1 | P2Y purinoceptor 1 | 373 | 42.072 | 9.049091e+02 | 4.783164e+04 |  |  | 47831.64 |  |  |  | 17.7 |  |
| sp\|Q8IWT6\|LRC8A\_HUMAN | Q8IWT6 | LRC8A\_HUMAN | LRRC8A | Volume-regulated anion channel subunit LRRC8A | 810 | 94.199 | 1.393397e+03 | 1.649067e+05 |  |  | 45834.97 | 119071.69 |  |  | 11.9 | 16.5 |
| sp\|Q99622\|C10\_HUMAN | Q99622 | C10\_HUMAN | C12orf57 | Protein C10 | 126 | 13.178 | 1.812879e+03 | 3.001480e+04 |  |  | 30014.803 |  |  |  | 13.5 |  |
| sp\|Q9Y2T3\|GUAD\_HUMAN | Q9Y2T3 | GUAD\_HUMAN | GDA | Guanine deaminase | 454 | 51.003 | 1.586676e+03 | 1.016719e+05 |  |  | 41192.117 | 60479.82 |  |  | 20.0 | 12.8 |
| sp\|O75112\|LDB3\_HUMAN | O75112 | LDB3\_HUMAN | LDB3 | LIM domain-binding protein 3 | 727 | 77.135 | 8.723680e+03 | 8.454119e+05 |  |  | 845411.9 |  |  |  | 22.6 |  |
| sp\|P43007\|SATT\_HUMAN | P43007 | SATT\_HUMAN | SLC1A4 | Neutral amino acid transporter A | 532 | 55.723 | 3.972620e+03 | 2.781177e+05 |  |  | 66589.1 | 211528.6 |  |  | 12.4 | 3.8 |
| sp\|P62318\|SMD3\_HUMAN | P62318 | SMD3\_HUMAN | SNRPD3 | Small nuclear ribonucleoprotein Sm D3 | 126 | 13.916 | 1.956853e+03 | 3.421288e+04 |  |  | 34212.883 |  |  |  | 35.7 |  |
| sp\|Q6Y288\|B3GLT\_HUMAN | Q6Y288 | B3GLT\_HUMAN | B3GLCT | Beta-1,3-glucosyltransferase | 498 | 56.564 | 6.994979e+02 | 4.970999e+04 |  |  | 49709.99 |  |  |  | 3.0 |  |
| sp\|P09601\|HMOX1\_HUMAN | P09601 | HMOX1\_HUMAN | HMOX1 | Heme oxygenase 1 | 288 | 32.819 | 1.034068e+03 | 4.263748e+04 |  |  | 42637.484 |  |  |  | 26.7 |  |
| sp\|O14967\|CLGN\_HUMAN | O14967 | CLGN\_HUMAN | CLGN | Calmegin | 610 | 70.039 | 3.209794e+02 | 2.824452e+04 |  |  | 28244.523 |  |  |  | 18.2 |  |
| sp\|O60218\|AK1BA\_HUMAN | O60218 | AK1BA\_HUMAN | AKR1B10 | Aldo-keto reductase family 1 member B10 | 316 | 36.020 | 4.743993e+02 | 2.146866e+04 |  |  | 21468.658 |  |  |  | 8.9 |  |
| sp\|Q16850\|CP51A\_HUMAN | Q16850 | CP51A\_HUMAN | CYP51A1 | Lanosterol 14-alpha demethylase | 509 | 57.278 | 2.574837e+02 | 1.852912e+04 |  |  | 18529.117 |  |  |  | 15.7 |  |
| sp\|P79483\|DRB3\_HUMAN | P79483 | DRB3\_HUMAN | HLA-DRB3 | HLA class II histocompatibility antigen, DR beta 3 chain | 266 | 29.962 | 9.549158e+02 | 3.594620e+04 |  |  | 35946.203 |  |  |  | 22.6 |  |
| sp\|Q9H361\|PABP3\_HUMAN | Q9H361 | PABP3\_HUMAN | PABPC3 | Polyadenylate-binding protein 3 | 631 | 70.031 | 6.207696e+03 | 5.461827e+05 |  |  | 303633.03 | 242549.62 |  |  | 24.7 | 20.8 |
| sp\|P08311\|CATG\_HUMAN | P08311 | CATG\_HUMAN | CTSG | Cathepsin G | 255 | 28.837 | 1.118857e+03 | 4.053607e+04 |  |  | 40536.074 |  |  |  | 23.1 |  |
| sp\|Q9Y2E5\|MA2B2\_HUMAN | Q9Y2E5 | MA2B2\_HUMAN | MAN2B2 | Epididymis-specific alpha-mannosidase | 1009 | 113.979 | 9.834397e+01 | 1.408282e+04 |  |  | 14082.823 |  |  |  | 6.7 |  |
| sp\|Q8TAV4\|STML3\_HUMAN | Q8TAV4 | STML3\_HUMAN | STOML3 | Stomatin-like protein 3 | 291 | 32.135 | 2.907676e+04 | 1.173928e+06 |  |  | 92555.32 | 1081372.5 |  |  | 17.5 | 14.1 |
| sp\|P51688\|SPHM\_HUMAN | P51688 | SPHM\_HUMAN | SGSH | N-sulphoglucosamine sulphohydrolase | 502 | 56.695 | 9.486156e+02 | 6.756973e+04 |  |  | 53936.605 | 13633.122 |  |  | 22.3 | 12.2 |
| sp\|Q07021\|C1QBP\_HUMAN | Q07021 | C1QBP\_HUMAN | C1QBP | Complement component 1 Q subcomponent-binding protein, mitochondrial | 282 | 31.362 | 5.449884e+02 | 2.147376e+04 |  |  | 21473.762 |  |  |  | 14.9 |  |
| sp\|Q96S99\|PKHF1\_HUMAN | Q96S99 | PKHF1\_HUMAN | PLEKHF1 | Pleckstrin homology domain-containing family F member 1 | 279 | 31.195 | 2.931502e+04 | 1.148927e+06 |  |  | 1058484.8 | 90442.02 |  |  | 20.1 | 12.2 |
| sp\|Q8NC56\|LEMD2\_HUMAN | Q8NC56 | LEMD2\_HUMAN | LEMD2 | LEM domain-containing protein 2 | 503 | 56.975 | 3.692216e+03 | 2.642948e+05 |  |  | 25311.756 | 238983.05 |  |  | 27.2 | 25.2 |
| sp\|P28325\|CYTD\_HUMAN | P28325 | CYTD\_HUMAN | CST5 | Cystatin-D | 142 | 16.080 | 3.754646e+03 | 7.585291e+04 |  |  | 75852.91 |  |  |  | 16.9 |  |
| sp\|Q8WUR7\|CO040\_HUMAN | Q8WUR7 | CO040\_HUMAN | C15orf40 | UPF0235 protein C15orf40 | 153 | 16.353 | 1.351175e+03 | 2.776044e+04 |  |  | 27760.438 |  |  |  | 13.7 |  |
| sp\|O00478\|BT3A3\_HUMAN | O00478 | BT3A3\_HUMAN | BTN3A3 | Butyrophilin subfamily 3 member A3 | 584 | 65.002 | 3.823980e+02 | 3.122910e+04 |  |  | 31229.104 |  |  |  | 20.9 |  |
| sp\|O15427\|MOT4\_HUMAN | O15427 | MOT4\_HUMAN | SLC16A3 | Monocarboxylate transporter 4 | 465 | 49.469 | 3.508541e+03 | 2.180604e+05 |  |  | 33203.477 | 184856.95 |  |  | 14.0 | 9.7 |
| sp\|Q96EY5\|MB12A\_HUMAN | Q96EY5 | MB12A\_HUMAN | MVB12A | Multivesicular body subunit 12A | 273 | 28.783 | 1.721549e+03 | 6.225477e+04 |  |  | 62254.77 |  |  |  | 31.5 |  |
| sp\|Q9BQT8\|ODC\_HUMAN | Q9BQT8 | ODC\_HUMAN | SLC25A21 | Mitochondrial 2-oxodicarboxylate carrier | 299 | 33.303 | 1.480167e+03 | 6.193143e+04 |  |  | 61931.43 |  |  |  | 27.8 |  |
| sp\|A0A075B6K5\|LV39\_HUMAN | A0A075B6K5 | LV39\_HUMAN | IGLV3-9 | Immunoglobulin lambda variable 3-9 | 115 | 12.332 | 6.095506e+02 | 9.444097e+03 |  |  | 9444.097 |  |  |  | 23.5 |  |
| sp\|Q13467\|FZD5\_HUMAN | Q13467 | FZD5\_HUMAN | FZD5 | Frizzled-5 | 585 | 64.507 | 8.311085e+02 | 6.735684e+04 |  |  | 9847.73 | 57509.11 |  |  | 22.7 | 11.8 |
| sp\|Q04837\|SSBP\_HUMAN | Q04837 | SSBP\_HUMAN | SSBP1 | Single-stranded DNA-binding protein, mitochondrial | 148 | 17.260 | 1.133365e+03 | 2.457695e+04 |  |  | 24576.947 |  |  |  | 32.4 |  |
| sp\|B2RPK0\|HGB1A\_HUMAN | B2RPK0 | HGB1A\_HUMAN | HMGB1P1 | Putative high mobility group protein B1-like 1 | 211 | 24.238 | 6.818902e+02 | 2.076483e+04 |  |  | 20764.83 |  |  |  | 23.2 |  |
| sp\|O43402\|EMC8\_HUMAN | O43402 | EMC8\_HUMAN | EMC8 | ER membrane protein complex subunit 8 | 210 | 23.773 | 1.478815e+03 | 4.416873e+04 |  |  | 44168.734 |  |  |  | 13.8 |  |
| sp\|P42167\|LAP2B\_HUMAN | P42167 | LAP2B\_HUMAN | TMPO | Lamina-associated polypeptide 2, isoforms beta/gamma | 454 | 50.670 | 2.126750e+02 | 1.353894e+04 |  |  | 13538.9375 |  |  |  | 27.1 |  |
| sp\|Q6ZT12\|UBR3\_HUMAN | Q6ZT12 | UBR3\_HUMAN | UBR3 | E3 ubiquitin-protein ligase UBR3 | 1888 | 212.433 | 1.228459e+03 | 3.278686e+05 |  |  | 327868.62 |  |  |  | 13.8 |  |
| sp\|P01877\|IGHA2\_HUMAN | P01877 | IGHA2\_HUMAN | IGHA2 | Immunoglobulin heavy constant alpha 2 | 340 | 42.334 | 2.194349e+04 | 1.167111e+06 |  |  | 1167110.8 |  |  |  | 25.6 |  |
| sp\|Q13642\|FHL1\_HUMAN | Q13642 | FHL1\_HUMAN | FHL1 | Four and a half LIM domains protein 1 | 323 | 36.263 | 1.108396e+03 | 5.049819e+04 |  |  | 33661.69 | 16836.504 |  |  | 18.6 | 25.7 |
| sp\|P15144\|AMPN\_HUMAN | P15144 | AMPN\_HUMAN | ANPEP | Aminopeptidase N | 967 | 109.540 | 1.344309e+02 | 1.850073e+04 |  |  | 7751.1973 | 10749.532 |  |  | 13.9 | 9.1 |
| sp\|P15880\|RS2\_HUMAN | P15880 | RS2\_HUMAN | RPS2 | 40S ribosomal protein S2 | 293 | 31.324 | 2.410244e+03 | 9.485392e+04 |  |  | 44545.58 | 50308.344 |  |  | 23.9 | 17.7 |
| sp\|Q92859\|NEO1\_HUMAN | Q92859 | NEO1\_HUMAN | NEO1 | Neogenin | 1461 | 160.017 | 3.656602e+02 | 7.351247e+04 |  |  | 24507.275 | 49005.195 |  |  | 10.7 | 13.5 |
| sp\|O60235\|TM11D\_HUMAN | O60235 | TM11D\_HUMAN | TMPRSS11D | Transmembrane protease serine 11D | 418 | 46.263 | 4.468539e+02 | 2.597266e+04 |  |  | 25972.662 |  |  |  | 8.6 |  |
| sp\|Q9BTV7\|CABL2\_HUMAN | Q9BTV7 | CABL2\_HUMAN | CABLES2 | CDK5 and ABL1 enzyme substrate 2 | 478 | 52.235 | 2.521256e+02 | 1.654610e+04 |  |  | 16546.102 |  |  |  | 29.7 |  |
| sp\|P19235\|EPOR\_HUMAN | P19235 | EPOR\_HUMAN | EPOR | Erythropoietin receptor | 508 | 55.065 | 1.935583e+05 | 1.339074e+07 |  |  | 13319961.0 | 70778.42 |  |  | 8.9 | 5.3 |
| sp\|O00258\|GET1\_HUMAN | O00258 | GET1\_HUMAN | GET1 | Guided entry of tail-anchored proteins factor 1 | 174 | 19.780 | 2.511176e+03 | 6.240517e+04 |  |  | 39336.184 | 23068.986 |  |  | 8.0 | 23.0 |
| sp\|P12830\|CADH1\_HUMAN | P12830 | CADH1\_HUMAN | CDH1 | Cadherin-1 | 882 | 97.456 | 1.715808e+02 | 2.100847e+04 |  |  | 21008.465 |  |  |  | 14.6 |  |
| sp\|Q5BJH7\|YIF1B\_HUMAN | Q5BJH7 | YIF1B\_HUMAN | YIF1B | Protein YIF1B | 314 | 34.435 | 3.459184e+03 | 1.496549e+05 |  |  | 54534.2 | 95120.68 |  |  | 36.6 | 12.7 |
| sp\|Q9H1A3\|METL9\_HUMAN | Q9H1A3 | METL9\_HUMAN | METTL9 | Protein-L-histidine N-pros-methyltransferase | 318 | 36.536 | 3.576841e+02 | 1.641866e+04 |  |  | 16418.664 |  |  |  | 15.7 |  |
| sp\|P84098\|RL19\_HUMAN | P84098 | RL19\_HUMAN | RPL19 | 60S ribosomal protein L19 | 196 | 23.466 | 1.094362e+03 | 3.226392e+04 |  |  | 32263.92 |  |  |  | 23.0 |  |
| sp\|Q9NX40\|OCAD1\_HUMAN | Q9NX40 | OCAD1\_HUMAN | OCIAD1 | OCIA domain-containing protein 1 | 245 | 27.626 | 8.009636e+02 | 2.780020e+04 |  |  | 27800.203 |  |  |  | 34.7 |  |
| sp\|Q9NZQ9\|TMOD4\_HUMAN | Q9NZQ9 | TMOD4\_HUMAN | TMOD4 | Tropomodulin-4 | 345 | 39.335 | 1.668185e+05 | 8.244046e+06 |  |  | 8244046.0 |  |  |  | 9.0 |  |
| sp\|P61011\|SRP54\_HUMAN | P61011 | SRP54\_HUMAN | SRP54 | Signal recognition particle 54 kDa protein | 504 | 55.705 | 1.393620e+02 | 9.753392e+03 |  |  | 9753.392 |  |  |  | 23.2 |  |
| sp\|Q6P1J6\|PLB1\_HUMAN | Q6P1J6 | PLB1\_HUMAN | PLB1 | Phospholipase B1, membrane-associated | 1458 | 163.081 | 1.238138e+02 | 2.536821e+04 |  |  | 25368.205 |  |  |  | 13.8 |  |
| sp\|P49821\|NDUV1\_HUMAN | P49821 | NDUV1\_HUMAN | NDUFV1 | NADH dehydrogenase [ubiquinone] flavoprotein 1, mitochondrial | 464 | 50.817 | 3.615658e+02 | 2.308413e+04 |  |  | 23084.13 |  |  |  | 11.6 |  |
| sp\|A0A0C4DH25\|KVD20\_HUMAN | A0A0C4DH25 | KVD20\_HUMAN | IGKV3D-20 | Immunoglobulin kappa variable 3D-20 | 116 | 12.515 | 2.136176e+03 | 3.358807e+04 |  |  | 33588.066 |  |  |  | 13.8 |  |
| sp\|Q99661\|KIF2C\_HUMAN | Q99661 | KIF2C\_HUMAN | KIF2C | Kinesin-like protein KIF2C | 725 | 81.313 | 8.285338e+01 | 8.464228e+03 |  |  | 8464.228 |  |  |  | 23.6 |  |
| sp\|Q9NUQ2\|PLCE\_HUMAN | Q9NUQ2 | PLCE\_HUMAN | AGPAT5 | 1-acyl-sn-glycerol-3-phosphate acyltransferase epsilon | 364 | 42.072 | 8.011418e+02 | 4.234671e+04 |  |  | 15877.345 | 26469.367 |  |  | 19.5 | 14.6 |
| sp\|P13674\|P4HA1\_HUMAN | P13674 | P4HA1\_HUMAN | P4HA1 | Prolyl 4-hydroxylase subunit alpha-1 | 534 | 61.049 | 1.434834e+02 | 1.100519e+04 |  |  | 11005.188 |  |  |  | 12.2 |  |
| sp\|P29622\|KAIN\_HUMAN | P29622 | KAIN\_HUMAN | SERPINA4 | Kallistatin | 427 | 48.542 | 0.000000e+00 | 0.000000e+00 |  |  | 0.0 |  |  |  | 7.0 |  |
| sp\|P28330\|ACADL\_HUMAN | P28330 | ACADL\_HUMAN | ACADL | Long-chain specific acyl-CoA dehydrogenase, mitochondrial | 430 | 47.656 | 1.249655e+02 | 7.482121e+03 |  |  | 7482.1206 |  |  |  | 18.8 |  |
| sp\|Q02809\|PLOD1\_HUMAN | Q02809 | PLOD1\_HUMAN | PLOD1 | Procollagen-lysine,2-oxoglutarate 5-dioxygenase 1 | 727 | 83.550 | 4.661020e+02 | 4.892654e+04 |  |  | 48926.543 |  |  |  | 20.1 |  |
| sp\|A0A075B6R9\|KVD24\_HUMAN | A0A075B6R9 | KVD24\_HUMAN | IGKV2D-24 | Probable non-functional immunoglobulin kappa variable 2D-24 | 120 | 13.079 | 3.124218e+03 | 5.133729e+04 |  |  | 51337.293 |  |  |  | 16.7 |  |
| sp\|P48067\|SC6A9\_HUMAN | P48067 | SC6A9\_HUMAN | SLC6A9 | Sodium- and chloride-dependent glycine transporter 1 | 706 | 78.260 | 1.818132e+03 | 1.787650e+05 |  |  | 64039.688 | 114725.3 |  |  | 3.0 | 4.8 |
| sp\|Q9H7F0\|AT133\_HUMAN | Q9H7F0 | AT133\_HUMAN | ATP13A3 | Polyamine-transporting ATPase 13A3 | 1226 | 138.043 | 4.994935e+02 | 8.662861e+04 |  |  | 12357.263 | 74271.35 |  |  | 13.8 | 11.0 |
| sp\|Q9P0I2\|EMC3\_HUMAN | Q9P0I2 | EMC3\_HUMAN | EMC3 | ER membrane protein complex subunit 3 | 261 | 29.952 | 9.773515e+02 | 3.677848e+04 |  |  | 36778.48 |  |  |  | 16.5 |  |
| sp\|Q9Y2W1\|TR150\_HUMAN | Q9Y2W1 | TR150\_HUMAN | THRAP3 | Thyroid hormone receptor-associated protein 3 | 955 | 108.666 | 8.152181e+01 | 1.112973e+04 |  |  | 11129.731 |  |  |  | 24.4 |  |
| sp\|P02585\|TNNC2\_HUMAN | P02585 | TNNC2\_HUMAN | TNNC2 | Troponin C, skeletal muscle | 160 | 18.122 | 3.624430e+04 | 8.252072e+05 |  |  | 825207.25 | 0.0 |  |  | 21.9 | 27.5 |
| sp\|Q9Y241\|HIG1A\_HUMAN | Q9Y241 | HIG1A\_HUMAN | HIGD1A | HIG1 domain family member 1A, mitochondrial | 93 | 10.143 | 1.782243e+03 | 2.271174e+04 |  |  | 22711.74 |  |  |  | 20.4 |  |
| sp\|P28065\|PSB9\_HUMAN | P28065 | PSB9\_HUMAN | PSMB9 | Proteasome subunit beta type-9 | 219 | 23.264 | 2.476464e+03 | 7.238249e+04 |  |  | 38482.69 | 33899.805 |  |  | 21.9 | 9.1 |
| sp\|Q5JTC6\|AMER1\_HUMAN | Q5JTC6 | AMER1\_HUMAN | AMER1 | APC membrane recruitment protein 1 | 1135 | 124.029 | 2.545108e+01 | 3.965944e+03 |  |  | 3965.944 |  |  |  | 18.8 |  |
| sp\|P43251\|BTD\_HUMAN | P43251 | BTD\_HUMAN | BTD | Biotinidase | 543 | 61.133 | 5.914037e+02 | 4.542311e+04 |  |  | 45423.11 |  |  |  | 8.7 |  |
| sp\|Q8WU20\|FRS2\_HUMAN | Q8WU20 | FRS2\_HUMAN | FRS2 | Fibroblast growth factor receptor substrate 2 | 508 | 57.029 | 6.428011e+02 | 4.605635e+04 |  |  | 30051.256 | 16005.093 |  |  | 23.2 | 14.2 |
| sp\|Q9BQR3\|PRS27\_HUMAN | Q9BQR3 | PRS27\_HUMAN | PRSS27 | Serine protease 27 | 290 | 31.940 | 3.491875e+02 | 1.401234e+04 |  |  | 14012.341 |  |  |  | 20.7 |  |
| sp\|Q2PZI1\|D19L1\_HUMAN | Q2PZI1 | D19L1\_HUMAN | DPY19L1 | Probable C-mannosyltransferase DPY19L1 | 675 | 77.319 | 1.484905e+02 | 1.442454e+04 |  |  | 14424.542 |  |  |  | 13.0 |  |
| sp\|Q6UWP7\|LCLT1\_HUMAN | Q6UWP7 | LCLT1\_HUMAN | LCLAT1 | Lysocardiolipin acyltransferase 1 | 414 | 48.920 | 8.950560e+02 | 5.501154e+04 |  |  | 27670.723 | 27340.82 |  |  | 11.1 | 8.0 |
| sp\|Q5HYA8\|MKS3\_HUMAN | Q5HYA8 | MKS3\_HUMAN | TMEM67 | Meckelin | 995 | 111.745 | 2.495075e+02 | 3.502909e+04 |  |  | 6803.7466 | 28225.348 |  |  | 8.8 | 6.5 |
| sp\|P20674\|COX5A\_HUMAN | P20674 | COX5A\_HUMAN | COX5A | Cytochrome c oxidase subunit 5A, mitochondrial | 150 | 16.762 | 7.372330e+03 | 1.552558e+05 |  |  | 155255.78 |  |  |  | 31.3 |  |
| sp\|O75386\|TULP3\_HUMAN | O75386 | TULP3\_HUMAN | TULP3 | Tubby-related protein 3 | 442 | 49.642 | 1.030187e+03 | 6.425140e+04 |  |  | 24988.734 | 39262.67 |  |  | 10.2 | 5.2 |
| sp\|O96000\|NDUBA\_HUMAN | O96000 | NDUBA\_HUMAN | NDUFB10 | NADH dehydrogenase [ubiquinone] 1 beta subcomplex subunit 10 | 172 | 20.777 | 9.742979e+02 | 2.543266e+04 |  |  | 25432.658 |  |  |  | 40.7 |  |
| sp\|P78382\|S35A1\_HUMAN | P78382 | S35A1\_HUMAN | SLC35A1 | CMP-sialic acid transporter | 337 | 36.779 | 1.012194e+03 | 4.677143e+04 |  |  | 12523.185 | 34248.25 |  |  | 6.2 | 17.8 |
| sp\|Q8TF09\|DLRB2\_HUMAN | Q8TF09 | DLRB2\_HUMAN | DYNLRB2 | Dynein light chain roadblock-type 2 | 96 | 10.855 | 2.236076e+03 | 3.049535e+04 |  |  | 30495.346 |  |  |  | 20.8 |  |
| sp\|Q9HB40\|RISC\_HUMAN | Q9HB40 | RISC\_HUMAN | SCPEP1 | Retinoid-inducible serine carboxypeptidase | 452 | 50.831 | 8.979266e+02 | 5.734382e+04 |  |  | 57343.824 |  |  |  | 7.3 |  |
| sp\|Q9NS69\|TOM22\_HUMAN | Q9NS69 | TOM22\_HUMAN | TOMM22 | Mitochondrial import receptor subunit TOM22 homolog | 142 | 15.522 | 3.462481e+02 | 6.752309e+03 |  |  | 6752.3086 |  |  |  | 16.2 |  |
| sp\|Q9UGL9\|CRCT1\_HUMAN | Q9UGL9 | CRCT1\_HUMAN | CRCT1 | Cysteine-rich C-terminal protein 1 | 99 | 9.736 | 2.829805e+03 | 3.461420e+04 |  |  | 34614.2 |  |  |  | 26.3 |  |
| sp\|Q9UHA4\|LTOR3\_HUMAN | Q9UHA4 | LTOR3\_HUMAN | LAMTOR3 | Ragulator complex protein LAMTOR3 | 124 | 13.623 | 3.945722e+03 | 6.753306e+04 |  |  | 67533.06 |  |  |  | 8.1 |  |
| sp\|Q9Y5M8\|SRPRB\_HUMAN | Q9Y5M8 | SRPRB\_HUMAN | SRPRB | Signal recognition particle receptor subunit beta | 271 | 29.702 | 1.031693e+03 | 3.849935e+04 |  |  | 13109.173 | 25390.18 |  |  | 11.8 | 23.2 |
| sp\|O75489\|NDUS3\_HUMAN | O75489 | NDUS3\_HUMAN | NDUFS3 | NADH dehydrogenase [ubiquinone] iron-sulfur protein 3, mitochondrial | 264 | 30.242 | 1.517258e+03 | 5.764836e+04 |  |  | 57648.363 |  |  |  | 17.4 |  |
| sp\|Q8WWA1\|TMM40\_HUMAN | Q8WWA1 | TMM40\_HUMAN | TMEM40 | Transmembrane protein 40 | 233 | 25.495 | 3.690223e+02 | 1.182020e+04 |  |  | 11820.2 |  |  |  | 18.9 |  |
| sp\|Q92963\|RIT1\_HUMAN | Q92963 | RIT1\_HUMAN | RIT1 | GTP-binding protein Rit1 | 219 | 25.145 | 1.655132e+03 | 5.228791e+04 |  |  | 23182.912 | 29105.0 |  |  | 22.8 | 24.2 |
| sp\|P31025\|LCN1\_HUMAN | P31025 | LCN1\_HUMAN | LCN1 | Lipocalin-1 | 176 | 19.250 | 5.031072e+03 | 1.216770e+05 |  |  | 42151.15 | 79525.84 |  |  | 6.8 | 10.8 |
| sp\|P01700\|LV147\_HUMAN | P01700 | LV147\_HUMAN | IGLV1-47 | Immunoglobulin lambda variable 1-47 | 117 | 12.284 | 2.251176e+04 | 3.474292e+05 |  |  | 347429.22 |  |  |  | 11.1 |  |
| sp\|P21810\|PGS1\_HUMAN | P21810 | PGS1\_HUMAN | BGN | Biglycan | 368 | 41.654 | 0.000000e+00 | 0.000000e+00 |  |  | 0.0 |  |  |  | 13.9 |  |
| sp\|P53634\|CATC\_HUMAN | P53634 | CATC\_HUMAN | CTSC | Dipeptidyl peptidase 1 | 463 | 51.854 | 2.369982e+02 | 1.543990e+04 |  |  | 15439.898 |  |  |  | 5.4 |  |
| sp\|O00748\|EST2\_HUMAN | O00748 | EST2\_HUMAN | CES2 | Cocaine esterase | 559 | 61.807 | 2.811089e+03 | 2.182878e+05 |  |  | 218287.81 |  |  |  | 15.9 |  |
| sp\|Q92520\|FAM3C\_HUMAN | Q92520 | FAM3C\_HUMAN | FAM3C | Protein FAM3C | 227 | 24.680 | 1.320261e+03 | 4.093758e+04 |  |  | 26964.725 | 13972.851 |  |  | 31.7 | 18.9 |
| sp\|O00459\|P85B\_HUMAN | O00459 | P85B\_HUMAN | PIK3R2 | Phosphatidylinositol 3-kinase regulatory subunit beta | 728 | 81.545 | 3.632905e+02 | 3.721933e+04 |  |  | 37219.33 |  |  |  | 2.4 |  |
| sp\|P12532\|KCRU\_HUMAN | P12532 | KCRU\_HUMAN | CKMT1A | Creatine kinase U-type, mitochondrial | 417 | 47.037 | 7.796239e+02 | 4.607251e+04 |  |  | 18065.805 | 28006.71 |  |  | 24.7 | 14.4 |
| sp\|Q3ZAQ7\|VMA21\_HUMAN | Q3ZAQ7 | VMA21\_HUMAN | VMA21 | Vacuolar ATPase assembly integral membrane protein VMA21 | 101 | 11.354 | 1.018651e+03 | 1.453087e+04 |  |  | 14530.87 |  |  |  | 34.7 |  |
| sp\|P42704\|LPPRC\_HUMAN | P42704 | LPPRC\_HUMAN | LRPPRC | Leucine-rich PPR motif-containing protein, mitochondrial | 1394 | 157.905 | 0.000000e+00 | 0.000000e+00 |  |  | 0.0 |  |  |  | 11.9 |  |
| sp\|A0A0C4DH41\|HV461\_HUMAN | A0A0C4DH41 | HV461\_HUMAN | IGHV4-61 | Immunoglobulin heavy variable 4-61 | 118 | 13.066 | 2.177581e+03 | 3.574655e+04 |  |  | 35746.547 |  |  |  | 7.8 |  |
| sp\|Q10589\|BST2\_HUMAN | Q10589 | BST2\_HUMAN | BST2 | Bone marrow stromal antigen 2 | 180 | 19.769 | 1.245823e+04 | 3.094270e+05 |  |  | 73623.27 | 235803.7 |  |  | 10.6 | 16.1 |
| sp\|Q9UNE0\|EDAR\_HUMAN | Q9UNE0 | EDAR\_HUMAN | EDAR | Tumor necrosis factor receptor superfamily member EDAR | 448 | 48.582 | 2.803668e+03 | 1.711272e+05 |  |  | 171127.25 |  |  |  | 14.5 |  |
| sp\|Q8N357\|S35F6\_HUMAN | Q8N357 | S35F6\_HUMAN | SLC35F6 | Solute carrier family 35 member F6 | 371 | 40.214 | 2.571400e+02 | 1.299165e+04 |  |  | 12991.645 |  |  |  | 7.0 |  |
| sp\|Q8IXS8\|F126B\_HUMAN | Q8IXS8 | F126B\_HUMAN | FAM126B | Protein FAM126B | 530 | 58.647 | 1.992729e+02 | 1.468288e+04 |  |  | 14682.878 |  |  |  | 9.8 |  |
| sp\|O95867\|LY66C\_HUMAN | O95867 | LY66C\_HUMAN | LY6G6C | Lymphocyte antigen 6 complex locus protein G6c | 125 | 13.821 | 7.918748e+02 | 1.375033e+04 |  |  | 13750.331 |  |  |  | 27.2 |  |
| sp\|Q92997\|DVL3\_HUMAN | Q92997 | DVL3\_HUMAN | DVL3 | Segment polarity protein dishevelled homolog DVL-3 | 716 | 78.055 | 9.996524e+01 | 9.803178e+03 |  |  | 9803.178 |  |  |  | 9.5 |  |
| sp\|O00115\|DNS2A\_HUMAN | O00115 | DNS2A\_HUMAN | DNASE2 | Deoxyribonuclease-2-alpha | 360 | 39.581 | 6.298598e+02 | 3.132188e+04 |  |  | 31321.88 |  |  |  | 10.0 |  |
| sp\|P53999\|TCP4\_HUMAN | P53999 | TCP4\_HUMAN | SUB1 | Activated RNA polymerase II transcriptional coactivator p15 | 127 | 14.395 | 1.527191e+03 | 2.761990e+04 |  |  | 27619.9 |  |  |  | 36.2 |  |
| sp\|P05156\|CFAI\_HUMAN | P05156 | CFAI\_HUMAN | CFI | Complement factor I | 583 | 65.750 | 2.017154e+02 | 1.666295e+04 |  |  | 16662.953 |  |  |  | 20.9 |  |
| sp\|Q6UXH1\|CREL2\_HUMAN | Q6UXH1 | CREL2\_HUMAN | CRELD2 | Protein disulfide isomerase CRELD2 | 353 | 38.192 | 1.078247e+03 | 5.173778e+04 |  |  | 51737.78 |  |  |  | 5.4 |  |
| sp\|O43670\|ZN207\_HUMAN | O43670 | ZN207\_HUMAN | ZNF207 | BUB3-interacting and GLEBS motif-containing protein ZNF207 | 478 | 50.751 | 3.045971e+02 | 1.942171e+04 |  |  | 19421.71 |  |  |  | 8.4 |  |
| sp\|P08493\|MGP\_HUMAN | P08493 | MGP\_HUMAN | MGP | Matrix Gla protein | 103 | 12.353 | 9.345252e+02 | 1.450376e+04 |  |  | 14503.761 |  |  |  | 27.2 |  |
| sp\|Q16842\|SIA4B\_HUMAN | Q16842 | SIA4B\_HUMAN | ST3GAL2 | CMP-N-acetylneuraminate-beta-galactosamide-alpha-2,3-sialyltransferase 2 | 350 | 40.173 | 2.301535e+02 | 1.161633e+04 |  |  | 11616.332 |  |  |  | 21.7 |  |
| sp\|P50336\|PPOX\_HUMAN | P50336 | PPOX\_HUMAN | PPOX | Protoporphyrinogen oxidase | 477 | 50.765 | 1.044322e+03 | 6.660642e+04 |  |  | 24329.057 | 42277.367 |  |  | 16.4 | 8.2 |
| sp\|Q9BSI4\|TINF2\_HUMAN | Q9BSI4 | TINF2\_HUMAN | TINF2 | TERF1-interacting nuclear factor 2 | 451 | 50.023 | 2.018299e+03 | 1.268447e+05 |  |  | 126844.71 |  |  |  | 18.6 |  |
| sp\|Q8IU60\|DCP2\_HUMAN | Q8IU60 | DCP2\_HUMAN | DCP2 | m7GpppN-mRNA hydrolase | 420 | 48.423 | 1.269117e+02 | 7.720947e+03 |  |  | 7720.947 |  |  |  | 16.7 |  |
| sp\|Q6P5S2\|LEG1H\_HUMAN | Q6P5S2 | LEG1H\_HUMAN | LEG1 | Protein LEG1 homolog | 330 | 37.926 | 7.892872e+02 | 3.760878e+04 |  |  | 37608.78 |  |  |  | 11.2 |  |
| sp\|Q9UBV7\|B4GT7\_HUMAN | Q9UBV7 | B4GT7\_HUMAN | B4GALT7 | Beta-1,4-galactosyltransferase 7 | 327 | 37.406 | 4.987321e+02 | 2.343828e+04 |  |  | 23438.28 |  |  |  | 12.2 |  |
| sp\|Q14108\|SCRB2\_HUMAN | Q14108 | SCRB2\_HUMAN | SCARB2 | Lysosome membrane protein 2 | 478 | 54.290 | 1.034111e+03 | 7.053492e+04 |  |  | 70534.92 |  |  |  | 7.5 |  |
| sp\|Q9BWL3\|CA043\_HUMAN | Q9BWL3 | CA043\_HUMAN | C1orf43 | Protein C1orf43 | 253 | 28.779 | 1.428405e+03 | 5.164689e+04 |  |  | 51646.895 |  |  |  | 19.4 |  |
| sp\|Q96GS6\|AB17A\_HUMAN | Q96GS6 | AB17A\_HUMAN | ABHD17A | Alpha/beta hydrolase domain-containing protein 17A | 310 | 33.990 | 4.707458e+02 | 2.010272e+04 |  |  | 11238.04 | 8864.678 |  |  | 13.2 | 5.8 |
| sp\|Q9NNX1\|TUFT1\_HUMAN | Q9NNX1 | TUFT1\_HUMAN | TUFT1 | Tuftelin | 390 | 44.264 | 3.339625e+02 | 1.857229e+04 |  |  | 18572.295 |  |  |  | 8.7 |  |
| sp\|P23467\|PTPRB\_HUMAN | P23467 | PTPRB\_HUMAN | PTPRB | Receptor-type tyrosine-protein phosphatase beta | 1997 | 224.301 | 2.421491e+02 | 6.823877e+04 |  |  | 68238.77 |  |  |  | 12.5 |  |
| sp\|A6NHG4\|DDTL\_HUMAN | A6NHG4 | DDTL\_HUMAN | DDTL | D-dopachrome decarboxylase-like protein | 134 | 14.195 | 1.427994e+05 | 2.546707e+06 |  |  |  | 2546706.8 |  |  |  | 23.7 |
| sp\|A7XYQ1\|SOBP\_HUMAN | A7XYQ1 | SOBP\_HUMAN | SOBP | Sine oculis-binding protein homolog | 873 | 92.658 | 1.485395e+04 | 1.729188e+06 |  |  |  | 1729187.8 |  |  |  | 6.3 |
| sp\|O14732\|IMPA2\_HUMAN | O14732 | IMPA2\_HUMAN | IMPA2 | Inositol monophosphatase 2 | 288 | 31.321 | 4.720242e+03 | 1.857450e+05 |  |  |  | 185744.97 |  |  |  | 17.0 |
| sp\|O75150\|BRE1B\_HUMAN | O75150 | BRE1B\_HUMAN | RNF40 | E3 ubiquitin-protein ligase BRE1B | 1001 | 113.678 | 4.583497e+02 | 6.546219e+04 |  |  |  | 65462.195 |  |  |  | 17.6 |
| sp\|O94986\|CE152\_HUMAN | O94986 | CE152\_HUMAN | CEP152 | Centrosomal protein of 152 kDa | 1710 | 195.626 | 2.489790e+03 | 6.119367e+05 |  |  |  | 611936.7 |  |  |  | 13.9 |
| sp\|O95183\|VAMP5\_HUMAN | O95183 | VAMP5\_HUMAN | VAMP5 | Vesicle-associated membrane protein 5 | 116 | 12.805 | 9.378540e+03 | 1.508801e+05 |  |  |  | 150880.11 |  |  |  | 52.6 |
| sp\|P00519\|ABL1\_HUMAN | P00519 | ABL1\_HUMAN | ABL1 | Tyrosine-protein kinase ABL1 | 1130 | 122.873 | 4.197091e+01 | 6.479209e+03 |  |  |  | 6479.2095 |  |  |  | 13.5 |
| sp\|P17900\|SAP3\_HUMAN | P17900 | SAP3\_HUMAN | GM2A | Ganglioside GM2 activator | 193 | 20.838 | 2.281086e+04 | 5.971931e+05 |  |  |  | 597193.06 |  |  |  | 13.0 |
| sp\|P20700\|LMNB1\_HUMAN | P20700 | LMNB1\_HUMAN | LMNB1 | Lamin-B1 | 586 | 66.408 | 7.989071e+02 | 6.665517e+04 |  |  |  | 66655.17 |  |  |  | 20.6 |
| sp\|P21439\|MDR3\_HUMAN | P21439 | MDR3\_HUMAN | ABCB4 | Phosphatidylcholine translocator ABCB4 | 1286 | 141.523 | 4.170370e+02 | 7.415130e+04 |  |  |  | 74151.305 |  |  |  | 9.6 |
| sp\|P37287\|PIGA\_HUMAN | P37287 | PIGA\_HUMAN | PIGA | Phosphatidylinositol N-acetylglucosaminyltransferase subunit A | 484 | 54.127 | 3.559642e+02 | 2.420681e+04 |  |  |  | 24206.805 |  |  |  | 13.0 |
| sp\|P48634\|PRC2A\_HUMAN | P48634 | PRC2A\_HUMAN | PRRC2A | Protein PRRC2A | 2157 | 228.863 | 1.538236e+02 | 4.422989e+04 |  |  |  | 44229.887 |  |  |  | 18.5 |
| sp\|P51798\|CLCN7\_HUMAN | P51798 | CLCN7\_HUMAN | CLCN7 | H(+)/Cl(-) exchange transporter 7 | 805 | 88.679 | 7.859357e+02 | 8.756387e+04 |  |  |  | 87563.87 |  |  |  | 6.8 |
| sp\|P55011\|S12A2\_HUMAN | P55011 | S12A2\_HUMAN | SLC12A2 | Solute carrier family 12 member 2 | 1212 | 131.447 | 8.451377e+02 | 1.395710e+05 |  |  |  | 139571.03 |  |  |  | 14.0 |
| sp\|P61803\|DAD1\_HUMAN | P61803 | DAD1\_HUMAN | DAD1 | Dolichyl-diphosphooligosaccharide--protein glycosyltransferase subunit DAD1 | 113 | 12.497 | 6.179919e+04 | 9.702992e+05 |  |  |  | 970299.2 |  |  |  | 35.4 |
| sp\|P98194\|AT2C1\_HUMAN | P98194 | AT2C1\_HUMAN | ATP2C1 | Calcium-transporting ATPase type 2C member 1 | 919 | 100.577 | 4.068770e+02 | 5.141371e+04 |  |  |  | 51413.71 |  |  |  | 11.4 |
| sp\|Q13423\|NNTM\_HUMAN | Q13423 | NNTM\_HUMAN | NNT | NAD(P) transhydrogenase, mitochondrial | 1086 | 113.896 | 6.139637e+02 | 8.785538e+04 |  |  |  | 87855.375 |  |  |  | 10.2 |
| sp\|Q5BKX6\|S45A4\_HUMAN | Q5BKX6 | S45A4\_HUMAN | SLC45A4 | Solute carrier family 45 member 4 | 768 | 83.878 | 2.243347e+03 | 2.364078e+05 |  |  |  | 236407.77 |  |  |  | 13.7 |
| sp\|Q6A163\|K1C39\_HUMAN | Q6A163 | K1C39\_HUMAN | KRT39 | Keratin, type I cytoskeletal 39 | 491 | 55.651 | 1.710214e+03 | 1.195751e+05 |  |  |  | 119575.06 |  |  |  | 27.9 |
| sp\|Q6NUJ1\|SAPL1\_HUMAN | Q6NUJ1 | SAPL1\_HUMAN | PSAPL1 | Proactivator polypeptide-like 1 | 521 | 56.627 | 6.290724e+02 | 4.475498e+04 |  |  |  | 44754.977 |  |  |  | 20.2 |
| sp\|Q6ZRV2\|FA83H\_HUMAN | Q6ZRV2 | FA83H\_HUMAN | FAM83H | Protein FAM83H | 1179 | 127.122 | 9.778302e+02 | 1.561713e+05 |  |  |  | 156171.33 |  |  |  | 19.0 |
| sp\|Q7Z4F1\|LRP10\_HUMAN | Q7Z4F1 | LRP10\_HUMAN | LRP10 | Low-density lipoprotein receptor-related protein 10 | 713 | 76.193 | 5.488740e+02 | 5.254179e+04 |  |  |  | 52541.79 |  |  |  | 9.4 |
| sp\|Q8IUK8\|CBLN2\_HUMAN | Q8IUK8 | CBLN2\_HUMAN | CBLN2 | Cerebellin-2 | 224 | 24.084 | 2.948852e+05 | 8.922751e+06 |  |  |  | 8922751.0 |  |  |  | 4.9 |
| sp\|Q8IWU2\|LMTK2\_HUMAN | Q8IWU2 | LMTK2\_HUMAN | LMTK2 | Serine/threonine-protein kinase LMTK2 | 1503 | 164.900 | 2.675460e+02 | 5.542890e+04 |  |  |  | 55428.9 |  |  |  | 9.2 |
| sp\|Q92544\|TM9S4\_HUMAN | Q92544 | TM9S4\_HUMAN | TM9SF4 | Transmembrane 9 superfamily member 4 | 642 | 74.519 | 2.023884e+03 | 1.894828e+05 |  |  |  | 189482.81 |  |  |  | 14.2 |
| sp\|Q9H0U3\|MAGT1\_HUMAN | Q9H0U3 | MAGT1\_HUMAN | MAGT1 | Magnesium transporter protein 1 | 335 | 38.037 | 6.522702e+03 | 3.117102e+05 |  |  |  | 311710.16 |  |  |  | 10.7 |
| sp\|Q9H6H4\|REEP4\_HUMAN | Q9H6H4 | REEP4\_HUMAN | REEP4 | Receptor expression-enhancing protein 4 | 257 | 29.395 | 3.156354e+03 | 1.165672e+05 |  |  |  | 116567.18 |  |  |  | 20.2 |
| sp\|Q9NVA4\|T184C\_HUMAN | Q9NVA4 | T184C\_HUMAN | TMEM184C | Transmembrane protein 184C | 438 | 50.142 | 1.893948e+03 | 1.193128e+05 |  |  |  | 119312.75 |  |  |  | 9.4 |
| sp\|Q9UHN6\|CEIP2\_HUMAN | Q9UHN6 | CEIP2\_HUMAN | CEMIP2 | Cell surface hyaluronidase | 1383 | 154.374 | 7.994624e+02 | 1.550563e+05 |  |  |  | 155056.27 |  |  |  | 12.8 |
| sp\|Q9UIJ5\|ZDHC2\_HUMAN | Q9UIJ5 | ZDHC2\_HUMAN | ZDHHC2 | Palmitoyltransferase ZDHHC2 | 367 | 42.022 | 9.886524e+03 | 5.219603e+05 |  |  |  | 521960.34 |  |  |  | 22.3 |
| sp\|Q9ULF5\|S39AA\_HUMAN | Q9ULF5 | S39AA\_HUMAN | SLC39A10 | Zinc transporter ZIP10 | 831 | 94.132 | 6.214964e+03 | 7.350097e+05 |  |  |  | 735009.7 |  |  |  | 8.9 |
| sp\|Q9Y446\|PKP3\_HUMAN | Q9Y446 | PKP3\_HUMAN | PKP3 | Plakophilin-3 | 797 | 87.082 | 9.210196e+02 | 1.007661e+05 |  |  |  | 100766.09 |  |  |  | 11.5 |
| sp\|A2A3N6\|PIPSL\_HUMAN | A2A3N6 | PIPSL\_HUMAN | PIPSL | Putative PIP5K1A and PSMD4-like protein | 862 | 95.048 | 6.240350e+02 | 7.451936e+04 |  |  |  | 74519.36 |  |  |  | 8.5 |
| sp\|O60812\|HNRC1\_HUMAN | O60812 | HNRC1\_HUMAN | HNRNPCL1 | Heterogeneous nuclear ribonucleoprotein C-like 1 | 293 | 32.142 | 1.259439e+03 | 5.085894e+04 |  |  |  | 50858.94 |  |  |  | 17.1 |
| sp\|P08729\|K2C7\_HUMAN | P08729 | K2C7\_HUMAN | KRT7 | Keratin, type II cytoskeletal 7 | 469 | 51.386 | 7.599318e+05 | 4.906102e+07 |  |  |  | 49061024.0 |  |  |  | 22.2 |
| sp\|P05787\|K2C8\_HUMAN | P05787 | K2C8\_HUMAN | KRT8 | Keratin, type II cytoskeletal 8 | 483 | 53.704 | 8.617210e+05 | 5.814206e+07 |  |  |  | 58142064.0 |  |  |  | 31.7 |
| sp\|P54707\|AT12A\_HUMAN | P54707 | AT12A\_HUMAN | ATP12A | Potassium-transporting ATPase alpha chain 2 | 1039 | 115.511 | 1.083684e+04 | 1.572690e+06 |  |  |  | 1572689.6 |  |  |  | 8.0 |
| sp\|Q13733\|AT1A4\_HUMAN | Q13733 | AT1A4\_HUMAN | ATP1A4 | Sodium/potassium-transporting ATPase subunit alpha-4 | 1029 | 114.166 | 1.758183e+04 | 2.521843e+06 |  |  |  | 2521843.0 |  |  |  | 12.4 |
| sp\|Q92764\|KRT35\_HUMAN | Q92764 | KRT35\_HUMAN | KRT35 | Keratin, type I cuticular Ha5 | 455 | 50.361 | 5.834851e+05 | 3.691826e+07 |  |  |  | 36918264.0 |  |  |  | 34.9 |
| sp\|Q7Z3Y7\|K1C28\_HUMAN | Q7Z3Y7 | K1C28\_HUMAN | KRT28 | Keratin, type I cytoskeletal 28 | 464 | 50.567 | 9.124400e+05 | 5.796805e+07 |  |  |  | 57968048.0 |  |  |  | 15.9 |
| sp\|P62491\|RB11A\_HUMAN | P62491 | RB11A\_HUMAN | RAB11A | Ras-related protein Rab-11A | 216 | 24.394 | 2.965081e+04 | 9.087339e+05 |  |  |  | 908733.9 |  |  |  | 31.9 |
| sp\|Q12756\|KIF1A\_HUMAN | Q12756 | KIF1A\_HUMAN | KIF1A | Kinesin-like protein KIF1A | 1690 | 191.064 | 6.928337e+02 | 1.663125e+05 |  |  |  | 166312.55 |  |  |  | 10.4 |
| sp\|Q96IW7\|SC22A\_HUMAN | Q96IW7 | SC22A\_HUMAN | SEC22A | Vesicle-trafficking protein SEC22a | 307 | 34.948 | 5.234749e+03 | 2.298452e+05 |  |  |  | 229845.16 |  |  |  | 12.4 |
| sp\|Q99879\|H2B1M\_HUMAN | Q99879 | H2B1M\_HUMAN | H2BC14 | Histone H2B type 1-M | 126 | 13.989 | 4.691387e+04 | 8.245274e+05 |  |  |  | 824527.4 |  |  |  | 65.1 |
| sp\|P19634\|SL9A1\_HUMAN | P19634 | SL9A1\_HUMAN | SLC9A1 | Sodium/hydrogen exchanger 1 | 815 | 90.763 | 2.385517e+02 | 2.720248e+04 |  |  |  | 27202.477 |  |  |  | 6.4 |
| sp\|Q9BUN8\|DERL1\_HUMAN | Q9BUN8 | DERL1\_HUMAN | DERL1 | Derlin-1 | 251 | 28.801 | 7.544540e+03 | 2.729968e+05 |  |  |  | 272996.75 |  |  |  | 19.5 |
| sp\|Q9H3U5\|MFSD1\_HUMAN | Q9H3U5 | MFSD1\_HUMAN | MFSD1 | Major facilitator superfamily domain-containing protein 1 | 465 | 51.209 | 2.817482e+03 | 1.812695e+05 |  |  |  | 181269.47 |  |  |  | 10.5 |
| sp\|P48029\|SC6A8\_HUMAN | P48029 | SC6A8\_HUMAN | SLC6A8 | Sodium- and chloride-dependent creatine transporter 1 | 635 | 70.523 | 1.702361e+02 | 1.508341e+04 |  |  |  | 15083.414 |  |  |  | 6.3 |
| sp\|Q15758\|AAAT\_HUMAN | Q15758 | AAAT\_HUMAN | SLC1A5 | Neutral amino acid transporter B(0) | 541 | 56.598 | 3.368064e+03 | 2.394962e+05 |  |  |  | 239496.16 |  |  |  | 6.1 |
| sp\|Q9H7F4\|T185B\_HUMAN | Q9H7F4 | T185B\_HUMAN | TMEM185B | Transmembrane protein 185B | 350 | 40.599 | 9.523364e+03 | 4.857612e+05 |  |  |  | 485761.25 |  |  |  | 9.1 |
| sp\|Q9UKR5\|ERG28\_HUMAN | Q9UKR5 | ERG28\_HUMAN | ERG28 | Ergosterol biosynthetic protein 28 homolog | 140 | 15.864 | 1.157969e+04 | 2.307951e+05 |  |  |  | 230795.14 |  |  |  | 45.0 |
| sp\|Q9Y5Y0\|FLVC1\_HUMAN | Q9Y5Y0 | FLVC1\_HUMAN | FLVCR1 | Feline leukemia virus subgroup C receptor-related protein 1 | 555 | 59.863 | 2.863801e+03 | 2.153865e+05 |  |  |  | 215386.53 |  |  |  | 3.8 |
| sp\|Q5VW38\|GP107\_HUMAN | Q5VW38 | GP107\_HUMAN | GPR107 | Protein GPR107 | 600 | 66.990 | 1.862853e+03 | 1.567854e+05 |  |  |  | 156785.4 |  |  |  | 7.7 |
| sp\|Q9P0T7\|TMEM9\_HUMAN | Q9P0T7 | TMEM9\_HUMAN | TMEM9 | Proton-transporting V-type ATPase complex assembly regulator TMEM9 | 183 | 20.574 | 1.474599e+04 | 3.811621e+05 |  |  |  | 381162.12 |  |  |  | 19.7 |
| sp\|O00461\|GOLI4\_HUMAN | O00461 | GOLI4\_HUMAN | GOLIM4 | Golgi integral membrane protein 4 | 696 | 81.880 | 9.850985e+01 | 1.013385e+04 |  |  |  | 10133.8545 |  |  |  | 7.3 |
| sp\|Q9BVT8\|TMUB1\_HUMAN | Q9BVT8 | TMUB1\_HUMAN | TMUB1 | Transmembrane and ubiquitin-like domain-containing protein 1 | 246 | 26.261 | 5.868976e+03 | 1.936381e+05 |  |  |  | 193638.06 |  |  |  | 8.9 |
| sp\|P48509\|CD151\_HUMAN | P48509 | CD151\_HUMAN | CD151 | CD151 antigen | 253 | 28.295 | 1.227789e+03 | 4.364660e+04 |  |  |  | 43646.6 |  |  |  | 9.1 |
| sp\|Q9NY59\|NSMA2\_HUMAN | Q9NY59 | NSMA2\_HUMAN | SMPD3 | Sphingomyelin phosphodiesterase 3 | 655 | 71.081 | 6.545884e+02 | 5.845733e+04 |  |  |  | 58457.33 |  |  |  | 12.7 |
| sp\|Q92902\|HPS1\_HUMAN | Q92902 | HPS1\_HUMAN | HPS1 | Hermansky-Pudlak syndrome 1 protein | 700 | 79.292 | 6.574385e+03 | 6.549402e+05 |  |  |  | 654940.25 |  |  |  | 6.7 |
| sp\|O95969\|SG1D2\_HUMAN | O95969 | SG1D2\_HUMAN | SCGB1D2 | Secretoglobin family 1D member 2 | 90 | 9.925 | 8.553106e+03 | 1.066526e+05 |  |  |  | 106652.62 |  |  |  | 22.2 |
| sp\|P09668\|CATH\_HUMAN | P09668 | CATH\_HUMAN | CTSH | Pro-cathepsin H | 335 | 37.394 | 1.562143e+03 | 7.339050e+04 |  |  |  | 73390.5 |  |  |  | 21.5 |
| sp\|Q01650\|LAT1\_HUMAN | Q01650 | LAT1\_HUMAN | SLC7A5 | Large neutral amino acids transporter small subunit 1 | 507 | 55.010 | 6.498570e+02 | 4.491347e+04 |  |  |  | 44913.47 |  |  |  | 13.0 |
| sp\|Q5FWE3\|PRRT3\_HUMAN | Q5FWE3 | PRRT3\_HUMAN | PRRT3 | Proline-rich transmembrane protein 3 | 981 | 102.197 | 1.679698e+02 | 2.156684e+04 |  |  |  | 21566.844 |  |  |  | 9.7 |
| sp\|Q03519\|TAP2\_HUMAN | Q03519 | TAP2\_HUMAN | TAP2 | Antigen peptide transporter 2 | 686 | 75.664 | 1.293764e+03 | 1.229876e+05 |  |  |  | 122987.64 |  |  |  | 3.8 |
| sp\|Q9UQ49\|NEUR3\_HUMAN | Q9UQ49 | NEUR3\_HUMAN | NEU3 | Sialidase-3 | 428 | 48.252 | 3.331787e+02 | 2.019806e+04 |  |  |  | 20198.062 |  |  |  | 10.7 |
| sp\|Q9HD20\|AT131\_HUMAN | Q9HD20 | AT131\_HUMAN | ATP13A1 | Endoplasmic reticulum transmembrane helix translocase | 1204 | 132.955 | 8.985547e+01 | 1.500950e+04 |  |  |  | 15009.505 |  |  |  | 10.0 |
| sp\|A0A075B6S2\|KVD29\_HUMAN | A0A075B6S2 | KVD29\_HUMAN | IGKV2D-29 | Immunoglobulin kappa variable 2D-29 | 120 | 13.143 | 6.557866e+03 | 1.082865e+05 |  |  |  | 108286.47 |  |  |  | 16.7 |
| sp\|P00403\|COX2\_HUMAN | P00403 | COX2\_HUMAN | MT-CO2 | Cytochrome c oxidase subunit 2 | 227 | 25.565 | 4.451137e+03 | 1.429664e+05 |  |  |  | 142966.39 |  |  |  | 11.9 |
| sp\|Q9BQ50\|TREX2\_HUMAN | Q9BQ50 | TREX2\_HUMAN | TREX2 | Three prime repair exonuclease 2 | 236 | 25.922 | 1.439944e+03 | 4.689550e+04 |  |  |  | 46895.5 |  |  |  | 15.7 |
| sp\|P51788\|CLCN2\_HUMAN | P51788 | CLCN2\_HUMAN | CLCN2 | Chloride channel protein 2 | 898 | 98.535 | 1.327336e+02 | 1.643193e+04 |  |  |  | 16431.928 |  |  |  | 8.8 |
| sp\|P23490\|LORI\_HUMAN | P23490 | LORI\_HUMAN | LORICRIN | Loricrin | 312 | 25.761 | 1.815144e+04 | 5.874772e+05 |  |  |  | 587477.2 |  |  |  | 6.7 |
| sp\|Q9H490\|PIGU\_HUMAN | Q9H490 | PIGU\_HUMAN | PIGU | Phosphatidylinositol glycan anchor biosynthesis class U protein | 435 | 50.052 | 1.165324e+03 | 7.327999e+04 |  |  |  | 73279.99 |  |  |  | 7.1 |
| sp\|Q9HC07\|TM165\_HUMAN | Q9HC07 | TM165\_HUMAN | TMEM165 | Transmembrane protein 165 | 324 | 34.906 | 0.000000e+00 | 0.000000e+00 |  |  |  | 0.0 |  |  |  | 17.6 |
| sp\|Q7Z3F1\|GP155\_HUMAN | Q7Z3F1 | GP155\_HUMAN | GPR155 | Integral membrane protein GPR155 | 870 | 96.919 | 1.764191e+02 | 2.148184e+04 |  |  |  | 21481.844 |  |  |  | 5.1 |
| sp\|Q71RG4\|TMUB2\_HUMAN | Q71RG4 | TMUB2\_HUMAN | TMUB2 | Transmembrane and ubiquitin-like domain-containing protein 2 | 321 | 33.788 | 1.219483e+03 | 5.176728e+04 |  |  |  | 51767.277 |  |  |  | 4.4 |
| sp\|P0C671\|BNIP5\_HUMAN | P0C671 | BNIP5\_HUMAN | BNIP5 | Protein BNIP5 | 652 | 71.930 | 7.268870e+02 | 6.568922e+04 |  |  |  | 65689.22 |  |  |  | 11.8 |
| sp\|P02652\|APOA2\_HUMAN | P02652 | APOA2\_HUMAN | APOA2 | Apolipoprotein A-II | 100 | 11.175 | 8.221087e+03 | 1.154234e+05 |  |  |  | 115423.414 |  |  |  | 17.0 |
| sp\|P22528\|SPR1B\_HUMAN | P22528 | SPR1B\_HUMAN | SPRR1B | Cornifin-B | 89 | 9.888 | 8.049152e+02 | 9.999441e+03 |  |  |  | 9999.441 |  |  |  | 61.8 |
| sp\|Q9NZH6\|IL37\_HUMAN | Q9NZH6 | IL37\_HUMAN | IL37 | Interleukin-37 | 218 | 24.126 | 2.635420e+03 | 7.988261e+04 |  |  |  | 79882.61 |  |  |  | 17.9 |
| sp\|O15118\|NPC1\_HUMAN | O15118 | NPC1\_HUMAN | NPC1 | NPC intracellular cholesterol transporter 1 | 1278 | 142.167 | 9.647718e+01 | 1.723219e+04 |  |  |  | 17232.193 |  |  |  | 5.3 |
| sp\|Q96NS1\|YPEL4\_HUMAN | Q96NS1 | YPEL4\_HUMAN | YPEL4 | Protein yippee-like 4 | 127 | 14.301 | 6.283463e+03 | 1.128971e+05 |  |  |  | 112897.06 |  |  |  | 37.0 |
| sp\|Q6ZMB5\|T184A\_HUMAN | Q6ZMB5 | T184A\_HUMAN | TMEM184A | Transmembrane protein 184A | 413 | 45.777 | 5.735635e+02 | 3.298724e+04 |  |  |  | 32987.24 |  |  |  | 8.2 |
| sp\|P51151\|RAB9A\_HUMAN | P51151 | RAB9A\_HUMAN | RAB9A | Ras-related protein Rab-9A | 201 | 22.838 | 5.485694e+02 | 1.574007e+04 |  |  |  | 15740.074 |  |  |  | 11.4 |
| sp\|P08582\|TRFM\_HUMAN | P08582 | TRFM\_HUMAN | MELTF | Melanotransferrin | 738 | 80.215 | 3.285945e+02 | 3.311563e+04 |  |  |  | 33115.633 |  |  |  | 8.0 |
| sp\|Q15124\|PGM5\_HUMAN | Q15124 | PGM5\_HUMAN | PGM5 | Phosphoglucomutase-like protein 5 | 567 | 62.225 | 0.000000e+00 | 0.000000e+00 |  |  |  | 0.0 |  |  |  | 10.6 |
| sp\|Q86Y34\|AGRG3\_HUMAN | Q86Y34 | AGRG3\_HUMAN | ADGRG3 | Adhesion G protein-coupled receptor G3 | 549 | 60.861 | 1.583144e+02 | 1.210533e+04 |  |  |  | 12105.331 |  |  |  | 4.4 |
| sp\|Q96HA4\|CA159\_HUMAN | Q96HA4 | CA159\_HUMAN | C1orf159 | Uncharacterized protein C1orf159 | 380 | 40.283 | 5.985987e+03 | 3.029526e+05 |  |  |  | 302952.62 |  |  |  | 17.1 |
| sp\|Q9BSR8\|YIPF4\_HUMAN | Q9BSR8 | YIPF4\_HUMAN | YIPF4 | Protein YIPF4 | 244 | 27.083 | 9.728034e+02 | 3.310084e+04 |  |  |  | 33100.84 |  |  |  | 18.4 |
| sp\|Q9HAT2\|SIAE\_HUMAN | Q9HAT2 | SIAE\_HUMAN | SIAE | Sialate O-acetylesterase | 523 | 58.315 | 4.857278e+02 | 3.558692e+04 |  |  |  | 35586.918 |  |  |  | 15.9 |
| sp\|Q8TEQ8\|PIGO\_HUMAN | Q8TEQ8 | PIGO\_HUMAN | PIGO | GPI ethanolamine phosphate transferase 3 | 1089 | 118.699 | 1.365916e+02 | 2.036987e+04 |  |  |  | 20369.871 |  |  |  | 5.0 |
| sp\|P60228\|EIF3E\_HUMAN | P60228 | EIF3E\_HUMAN | EIF3E | Eukaryotic translation initiation factor 3 subunit E | 445 | 52.221 | 1.510149e+02 | 9.907913e+03 |  |  |  | 9907.913 |  |  |  | 10.3 |
| sp\|Q14956\|GPNMB\_HUMAN | Q14956 | GPNMB\_HUMAN | GPNMB | Transmembrane glycoprotein NMB | 572 | 63.923 | 2.432334e+03 | 1.953428e+05 |  |  |  | 195342.83 |  |  |  | 14.3 |
| sp\|Q8IVJ1\|S41A1\_HUMAN | Q8IVJ1 | S41A1\_HUMAN | SLC41A1 | Solute carrier family 41 member 1 | 513 | 54.901 | 6.503957e+02 | 4.486163e+04 |  |  |  | 44861.63 |  |  |  | 3.5 |
| sp\|P25942\|TNR5\_HUMAN | P25942 | TNR5\_HUMAN | CD40 | Tumor necrosis factor receptor superfamily member 5 | 277 | 30.619 | 4.336772e+02 | 1.668302e+04 |  |  |  | 16683.021 |  |  |  | 14.4 |
| sp\|Q9Y5E5\|PCDB4\_HUMAN | Q9Y5E5 | PCDB4\_HUMAN | PCDHB4 | Protocadherin beta-4 | 795 | 87.270 | 5.143689e+02 | 5.639711e+04 |  |  |  | 56397.105 |  |  |  | 4.4 |
| sp\|P01920\|DQB1\_HUMAN | P01920 | DQB1\_HUMAN | HLA-DQB1 | HLA class II histocompatibility antigen, DQ beta 1 chain | 261 | 29.991 | 3.002563e+02 | 1.131358e+04 |  |  |  | 11313.583 |  |  |  | 10.3 |
| sp\|Q8IY95\|TM192\_HUMAN | Q8IY95 | TM192\_HUMAN | TMEM192 | Transmembrane protein 192 | 271 | 30.922 | 1.712155e+03 | 6.651625e+04 |  |  |  | 66516.25 |  |  |  | 3.3 |
| sp\|Q96K19\|RN170\_HUMAN | Q96K19 | RN170\_HUMAN | RNF170 | E3 ubiquitin-protein ligase RNF170 | 258 | 29.815 | 3.738498e+02 | 1.400390e+04 |  |  |  | 14003.902 |  |  |  | 8.9 |
| sp\|Q9NX94\|WBP1L\_HUMAN | Q9NX94 | WBP1L\_HUMAN | WBP1L | WW domain binding protein 1-like | 342 | 37.828 | 7.139888e+02 | 3.393297e+04 |  |  |  | 33932.973 |  |  |  | 11.7 |
| sp\|P33764\|S10A3\_HUMAN | P33764 | S10A3\_HUMAN | S100A3 | Protein S100-A3 | 101 | 11.713 | 5.504545e+02 | 8.100405e+03 |  |  |  | 8100.405 |  |  |  | 18.8 |
| sp\|Q6XR72\|ZNT10\_HUMAN | Q6XR72 | ZNT10\_HUMAN | SLC30A10 | Zinc transporter 10 | 485 | 52.684 | 5.668226e+02 | 3.751829e+04 |  |  |  | 37518.29 |  |  |  | 7.8 |
| sp\|P36406\|TRI23\_HUMAN | P36406 | TRI23\_HUMAN | TRIM23 | E3 ubiquitin-protein ligase TRIM23 | 574 | 64.067 | 1.005848e+02 | 8.096250e+03 |  |  |  | 8096.25 |  |  |  | 10.6 |
| sp\|Q96G23\|CERS2\_HUMAN | Q96G23 | CERS2\_HUMAN | CERS2 | Ceramide synthase 2 | 380 | 44.876 | 3.877199e+02 | 2.185996e+04 |  |  |  | 21859.963 |  |  |  | 2.4 |
| sp\|P04181\|OAT\_HUMAN | P04181 | OAT\_HUMAN | OAT | Ornithine aminotransferase, mitochondrial | 439 | 48.535 | 2.194632e+02 | 1.338239e+04 |  |  |  | 13382.394 |  |  |  | 4.3 |
| sp\|P16298\|PP2BB\_HUMAN | P16298 | PP2BB\_HUMAN | PPP3CB | Serine/threonine-protein phosphatase 2B catalytic subunit beta isoform | 524 | 59.024 | 2.200950e+02 | 1.632134e+04 |  |  |  | 16321.343 |  |  |  | 4.8 |
| sp\|Q96A83\|COQA1\_HUMAN | Q96A83 | COQA1\_HUMAN | COL26A1 | Collagen alpha-1(XXVI) chain | 441 | 45.381 | 1.061224e+04 | 6.050599e+05 |  |  |  | 605059.94 |  |  |  | 20.0 |
| sp\|P57054\|PIGP\_HUMAN | P57054 | PIGP\_HUMAN | PIGP | Phosphatidylinositol N-acetylglucosaminyltransferase subunit P | 158 | 18.089 | 1.951892e+03 | 4.435959e+04 |  |  |  | 44359.594 |  |  |  | 24.1 |
| sp\|Q8N697\|S15A4\_HUMAN | Q8N697 | S15A4\_HUMAN | SLC15A4 | Solute carrier family 15 member 4 | 577 | 62.034 | 0.000000e+00 | 0.000000e+00 |  |  |  | 0.0 |  |  |  | 9.5 |
| sp\|Q8WVP7\|LMBR1\_HUMAN | Q8WVP7 | LMBR1\_HUMAN | LMBR1 | Limb region 1 protein homolog | 490 | 55.098 | 2.138476e+02 | 1.480326e+04 |  |  |  | 14803.258 |  |  |  | 6.3 |
| sp\|Q9UHJ9\|PGAP2\_HUMAN | Q9UHJ9 | PGAP2\_HUMAN | PGAP2 | Post-GPI attachment to proteins factor 2 | 254 | 29.400 | 8.591477e+02 | 3.173455e+04 |  |  |  | 31734.547 |  |  |  | 11.8 |
| sp\|Q13433\|S39A6\_HUMAN | Q13433 | S39A6\_HUMAN | SLC39A6 | Zinc transporter ZIP6 | 755 | 85.047 | 7.609091e+01 | 8.130344e+03 |  |  |  | 8130.344 |  |  |  | 4.6 |
| sp\|O95476\|CNEP1\_HUMAN | O95476 | CNEP1\_HUMAN | CTDNEP1 | CTD nuclear envelope phosphatase 1 | 244 | 28.377 | 1.021264e+03 | 3.641009e+04 |  |  |  | 36410.09 |  |  |  | 7.8 |
| sp\|Q8NFB2\|T185A\_HUMAN | Q8NFB2 | T185A\_HUMAN | TMEM185A | Transmembrane protein 185A | 350 | 40.647 | 4.143118e+03 | 2.115792e+05 |  |  |  | 211579.19 |  |  |  | 5.1 |
| sp\|Q6ZUX7\|LHPL2\_HUMAN | Q6ZUX7 | LHPL2\_HUMAN | LHFPL2 | LHFPL tetraspan subfamily member 2 protein | 228 | 24.486 | 7.964795e+03 | 2.450245e+05 |  |  |  | 245024.55 |  |  |  | 7.5 |
| sp\|Q8WW43\|APH1B\_HUMAN | Q8WW43 | APH1B\_HUMAN | APH1B | Gamma-secretase subunit APH-1B | 257 | 28.460 | 1.860520e+03 | 6.652522e+04 |  |  |  | 66525.22 |  |  |  | 9.3 |
| sp\|Q8NBM4\|UBAC2\_HUMAN | Q8NBM4 | UBAC2\_HUMAN | UBAC2 | Ubiquitin-associated domain-containing protein 2 | 344 | 38.964 | 2.153015e+02 | 1.053969e+04 |  |  |  | 10539.686 |  |  |  | 18.0 |
| sp\|P01911\|DRB1\_HUMAN | P01911 | DRB1\_HUMAN | HLA-DRB1 | HLA class II histocompatibility antigen, DRB1 beta chain | 266 | 29.966 | 1.453010e+03 | 5.470341e+04 |  |  |  | 54703.41 |  |  |  | 3.5 |
| sp\|A0A0A0MRZ8\|KVD11\_HUMAN | A0A0A0MRZ8 | KVD11\_HUMAN | IGKV3D-11 | Immunoglobulin kappa variable 3D-11 | 115 | 12.625 | 1.035239e+04 | 1.642061e+05 |  |  |  | 164206.1 |  |  |  | 7.8 |
| sp\|Q14242\|SELPL\_HUMAN | Q14242 | SELPL\_HUMAN | SELPLG | P-selectin glycoprotein ligand 1 | 412 | 43.201 | 2.058401e+03 | 1.117226e+05 |  |  |  | 111722.6 |  |  |  | 3.4 |
| sp\|Q8N377\|YJ004\_HUMAN | Q8N377 | YJ004\_HUMAN |  | Putative uncharacterized protein LOC387726 | 158 | 17.380 | 7.759663e+03 | 1.694376e+05 |  |  |  | 169437.6 |  |  |  | 11.4 |
| sp\|Q8N0U8\|VKORL\_HUMAN | Q8N0U8 | VKORL\_HUMAN | VKORC1L1 | Vitamin K epoxide reductase complex subunit 1-like protein 1 | 176 | 19.836 | 4.929801e+03 | 1.228572e+05 |  |  |  | 122857.22 |  |  |  | 9.1 |
| sp\|Q9H1E3\|NUCKS\_HUMAN | Q9H1E3 | NUCKS\_HUMAN | NUCKS1 | Nuclear ubiquitous casein and cyclin-dependent kinase substrate 1 | 243 | 27.296 | 8.383837e+02 | 2.875140e+04 |  |  |  | 28751.4 |  |  |  | 14.8 |
| sp\|Q9UBH0\|I36RA\_HUMAN | Q9UBH0 | I36RA\_HUMAN | IL36RN | Interleukin-36 receptor antagonist protein | 155 | 16.962 | 1.491073e+03 | 3.177555e+04 |  |  |  | 31775.553 |  |  |  | 12.9 |
| sp\|Q9Y672\|ALG6\_HUMAN | Q9Y672 | ALG6\_HUMAN | ALG6 | Dolichyl pyrophosphate Man9GlcNAc2 alpha-1,3-glucosyltransferase | 507 | 58.121 | 7.155274e+02 | 5.224882e+04 |  |  |  | 52248.824 |  |  |  | 1.8 |
| sp\|Q8IXI1\|MIRO2\_HUMAN | Q8IXI1 | MIRO2\_HUMAN | RHOT2 | Mitochondrial Rho GTPase 2 | 618 | 68.118 | 1.211166e+03 | 1.036532e+05 |  |  |  | 103653.23 |  |  |  | 5.7 |
| sp\|Q92521\|PIGB\_HUMAN | Q92521 | PIGB\_HUMAN | PIGB | GPI mannosyltransferase 3 | 554 | 65.056 | 1.961082e+02 | 1.602877e+04 |  |  |  | 16028.769 |  |  |  | 5.4 |
| sp\|Q9H0V1\|TM168\_HUMAN | Q9H0V1 | TM168\_HUMAN | TMEM168 | Transmembrane protein 168 | 697 | 79.755 | 1.320295e+02 | 1.322958e+04 |  |  |  | 13229.582 |  |  |  | 5.9 |
| sp\|P12318\|FCG2A\_HUMAN | P12318 | FCG2A\_HUMAN | FCGR2A | Low affinity immunoglobulin gamma Fc region receptor II-a | 317 | 35.001 | 1.194376e+03 | 5.252169e+04 |  |  |  | 52521.69 |  |  |  | 6.0 |
| sp\|P55808\|XG\_HUMAN | P55808 | XG\_HUMAN | XG | Glycoprotein Xg | 180 | 19.723 | 6.578863e+03 | 1.630200e+05 |  |  |  | 163020.02 |  |  |  | 3.9 |
| sp\|Q9UN36\|NDRG2\_HUMAN | Q9UN36 | NDRG2\_HUMAN | NDRG2 | Protein NDRG2 | 371 | 40.798 | 3.115207e+02 | 1.596772e+04 |  |  |  | 15967.719 |  |  |  | 8.4 |
| sp\|Q99720\|SGMR1\_HUMAN | Q99720 | SGMR1\_HUMAN | SIGMAR1 | Sigma non-opioid intracellular receptor 1 | 223 | 25.128 | 7.921295e+03 | 2.500756e+05 |  |  |  | 250075.56 |  |  |  | 3.6 |
| sp\|P29034\|S10A2\_HUMAN | P29034 | S10A2\_HUMAN | S100A2 | Protein S100-A2 | 98 | 11.117 | 8.587672e+03 | 1.199445e+05 |  |  |  | 119944.46 |  |  |  | 9.2 |
| sp\|Q9NZU0\|FLRT3\_HUMAN | Q9NZU0 | FLRT3\_HUMAN | FLRT3 | Leucine-rich repeat transmembrane protein FLRT3 | 649 | 73.004 | 0.000000e+00 | 0.000000e+00 |  |  |  | 0.0 |  |  |  | 1.2 |
| sp\|O94886\|CSCL1\_HUMAN | O94886 | CSCL1\_HUMAN | TMEM63A | CSC1-like protein 1 | 807 | 92.126 | 1.047284e+02 | 1.212171e+04 |  |  |  | 12121.714 |  |  |  | 6.2 |
| sp\|Q7Z698\|SPRE2\_HUMAN | Q7Z698 | SPRE2\_HUMAN | SPRED2 | Sprouty-related, EVH1 domain-containing protein 2 | 418 | 47.558 | 5.406915e+02 | 3.230653e+04 |  |  |  | 32306.525 |  |  |  | 15.3 |
| sp\|Q99519\|NEUR1\_HUMAN | Q99519 | NEUR1\_HUMAN | NEU1 | Sialidase-1 | 415 | 45.467 | 6.199361e+03 | 3.541281e+05 |  |  |  | 354128.1 |  |  |  | 14.5 |
| sp\|P01903\|DRA\_HUMAN | P01903 | DRA\_HUMAN | HLA-DRA | HLA class II histocompatibility antigen, DR alpha chain | 254 | 28.621 | 1.144328e+03 | 4.114836e+04 |  |  |  | 41148.36 |  |  |  | 5.9 |
| sp\|P83881\|RL36A\_HUMAN | P83881 | RL36A\_HUMAN | RPL36A | 60S ribosomal protein L36a | 106 | 12.441 | 0.000000e+00 | 0.000000e+00 |  |  |  | 0.0 |  |  |  | 18.9 |
| sp\|P62304\|RUXE\_HUMAN | P62304 | RUXE\_HUMAN | SNRPE | Small nuclear ribonucleoprotein E | 92 | 10.804 | 1.090083e+03 | 1.479657e+04 |  |  |  | 14796.573 |  |  |  | 21.7 |
