## Supplementary figures and images for "Deep Red Blood Cell Proteome Defines the Band 3 N-Terminus Interactome as a Regulator of Hypoxic Adaptation via BLVRB-Dependent *S*-Nitroso Transfer"

### DALL·E 2024-02-16 08.37.18 - Create an image of a single red blood cell on a black background. The red blood cell should be detailed and realistic, capturing its distinctive bicon.webp

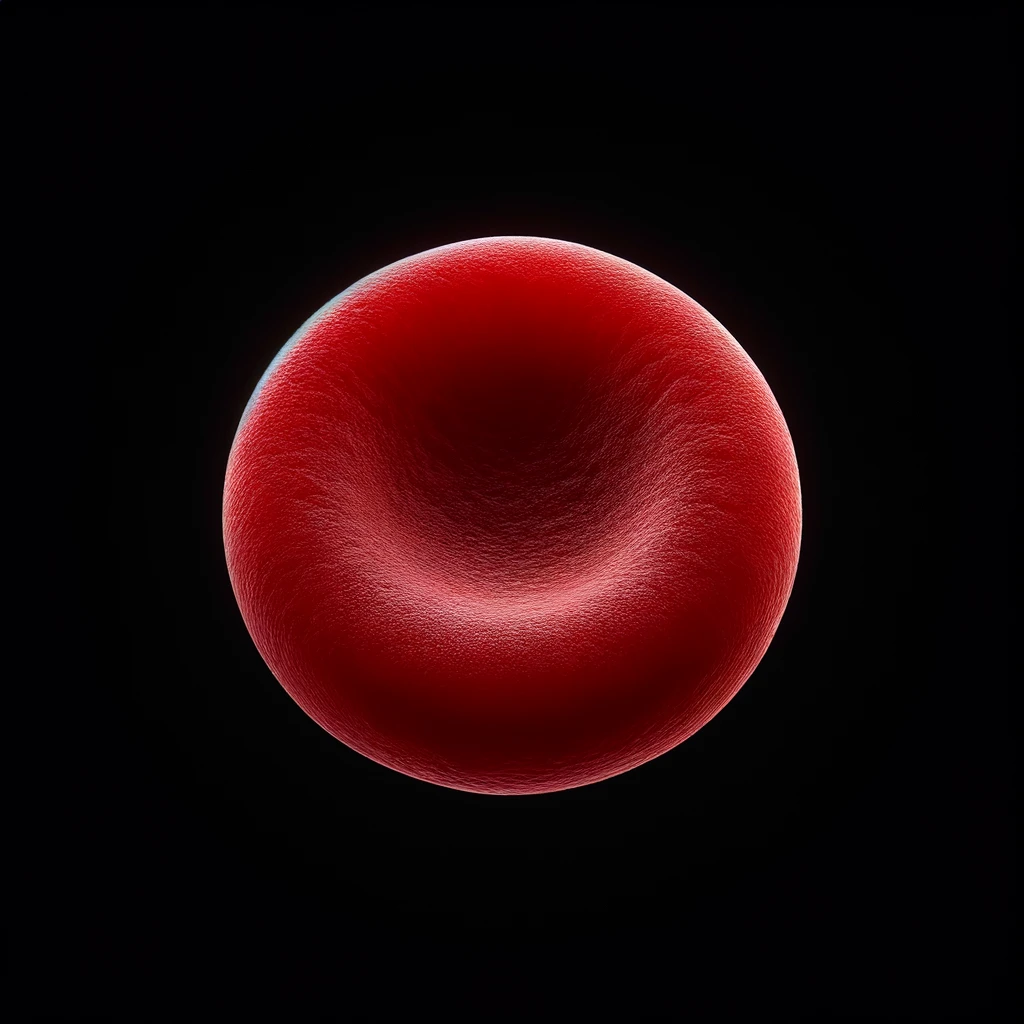
